# Evolutionary origin and functional diversification of plant GBF1-type ARF guanine-nucleotide exchange factors

**DOI:** 10.1101/2025.06.03.657657

**Authors:** Manoj K. Singh, Theresa Lauster, Kerstin Huhn, Sandra Richter, Marika Kientz, Richard A. Neher, Gerd Jürgens

## Abstract

Large ARF GTPase-activating guanine-nucleotide exchange factors (ARF-GEFs) are essential regulators of membrane trafficking across the eukaryotes. Although conserved, the GBF1-type ARF-GEFs underwent plant-specific evolution. Of the 3 paralogs in *Arabidopsis thaliana*, AtGNOM-LIKE1 (AtGNL1) performs the GBF1 task of retrograde Golgi-ER traffic whereas AtGNOM mediates polar recycling from endosomes to the basal plasma membrane. AtGNL2 is specific to pollen and is functionally equivalent to AtGNOM. To clarify their evolutionary origin and functional diversification in the plant lineage, we established a phylogenetic tree and ortho-synteny groups (OSGs), which enabled identification of AtGNOM and AtGNL1 orthologs. In addition, somatic paralogs from several species were functionally analysed in transgenic Arabidopsis plants and tested for GNOM-specific DCB-DCB domain interaction in yeast two-hybrid assays. Our results support the following scenario. The ancient plant-specific GBF1-type ARF-GEF was functionally equivalent to AtGNOM and mediated both polar recycling and secretory trafficking. GNL2 only arose in the early flowering plants and then evolved independently. Genome duplication in the basal eudicots – but neither in monocots nor in magnoliids – gave rise to the *AtGNL1* ortholog, distinguished by its unique OSG. The AtGNL1 ortholog lost the ability of DCB-DCB interaction and was free to evolve independently, eventually acquiring AtGNL1 function in the rosid clade of core eudicots. In eudicot evolution, the *AtGNOM* ortholog underwent repeated transposition to different OSGs whereas the *AtGNL1* ortholog was recurrently lost but eventually positively selected. This evolutionary process uncoupled polar recycling required for gravitropic response or lateral root initiation from secretory trafficking required for growth.

## INTRODUCTION

Membrane trafficking delivers roughly one-third of eukaryotic proteins to their sites of action on endomembrane compartments or the plasma membrane as well as to the vacuole or lysosome for degradation. The formation of cargo carriers such as membrane vesicles requires the action of small GTPases of the ARF family which need to be activated through GDP-GTP exchange by specific guanine-nucleotide exchange factors (ARF-GEFs). The family of large ARF-GEFs is conserved between plants and non-plant eukaryotes and comprises two classes named after the respective mammalian prototypes GBF1 and BIG1 (Pipaliya et al., 2019). GBF1 regulates the formation of COPI vesicles mediating retrograde traffic from the Golgi stack(s) to the endoplasmic reticulum (ER). This indirectly also affects the anterograde ER-Golgi traffic and thus the early section of the secretory pathway. By contrast, BIG1 regulates the late secretory pathway from the *trans*-Golgi network (TGN) to the plasma membrane (reviews: Casanova, 2007; Anders & Jürgens, 2008; Bui et al., 2009; Singh & Jürgens, 2018).

In the flowering plant *Arabidopsis thaliana*, there are three GBF1-related paralogs – GNOM, GNOM-LIKE1 (GNL1) and GNL2 – with different roles in membrane trafficking. GNL1 performs the eukaryotic ancestral function in COPI-dependent retrograde Golgi-ER traffic and is thus essentially the plant counterpart to GBF1 (Richter et al., 2007; Teh and Moore, 2007). GNOM, however, has a unique and plant-specific role, mediating polar recycling of proteins from endosomes to the basal plasma membrane (Geldner et al., 2003). One prominent cargo protein delivered along this pathway is the PIN1 protein which transports the signaling molecule auxin out of the cell (Geldner et al., 2003). GNOM can compensate for the lack of GNL1 whereas GNL1 cannot take over GNOM function (Richter et al., 2007). The third paralog – GNL2 – is functionally equivalent to GNOM but acts exclusively in pollen development (Richter et al., 2011).

Large ARF-GEFs share a conserved domain organization: an N-terminal dimerization (DCB) domain, a Homology Upstream of SEC7 (HUS) domain, the catalytic SEC7 domain and 3 or 4 Homology Downstream of SEC7 (HDS1-3/4) domains (Anders and Jürgens, 2008). The DCB domain of GNOM interacts with another copy of itself and with the complementary ΔDCB fragment of GNOM, and this latter interaction appears to mediate membrane association of GNOM (Grebe et al., 2000; Anders et al., 2008). GNL1 shares the DCB-ΔDCB interaction such that in yeast two-hybrid interaction assays, the two parts of one paralog each can also interact with the complementary parts of the other paralog (Brumm and Singh et al., 2022). Nonetheless, GNOM-GNL1 heterodimer formation does not occur in Arabidopsis plants, which appears to be prevented by the DCB-DCB interaction occurring between GNOM, but not GNL1, proteins nor between GNOM and GNL1 (Brumm and Singh et al., 2022).

Here, we analyse the evolution of GBF1-type ARF-GEFs in the plant lineage (Streptophyta), using phylogenetic sequence comparisons and synteny analyses as well as functional studies of GNOM-related ARF-GEFs in transgenic Arabidopsis plants and DCB-DCB interaction using yeast two-hybrid assays. Our results suggest that the ancient plant-specific GBF1-type ARF-GEF was functionally most closely related to Arabidopsis GNOM (AtGNOM) whereas GNL1 – although functionally more similar to non-plant GBF1 – appears to be derived. In addition, the evolution towards AtGNL1 of a GNOM duplicate required an initial step of disrupting DCB-DCB interaction. This appears to have occurred early in eudicot evolution whereas the two paralogs became functionally distinct later and only within the rosid clade of core eudicots.

## RESULTS

### A phylogenetic tree of GBF1-type ARF guanine-nucleotide exchange factors in Streptophyta

Genome sequences from >1,500 plant species are publicly available, which facilitated our phylogenetic analysis of GNOM-related ARF-GEFs (for overview see Marks et al., 2021; Kress et al., 2022; Pucker et al., 2022; Xie et al., 2024; monthly updated website; Plant Biotechnology Information (https://www.plabipd.de/index.ep /); for genome databases used, see Materials and Methods).

Our analysis included genomes from streptophytic algae, non-angiosperm land plants comprising mosses, lycopods, ferns and gymnosperms as well as basal angiosperms and the 5 major mesangiosperm clades of monocots, magnoliids and their sister clade Chloranthales, and eudicots and their sister clade Ceratophyllales (papers on angiosperm phylogeny include Yang et al., 2020; Zeng et al., 2017; Stull et al., 2023; Guo et al., 2021; OTPTI, 2019; Zhang et al., 2020a; Wang et al., 2022; Chanderbali et al., 2022; Xue et al., 2022; Li et al., 2021a; Wang et al., 2023; Zuntini et al., 2024; review: Guo et al., 2023). We initially chose 175 streptophytic species, for which genomic sequence information was available at the onset of this study, and analysed the phylogeny of GNOM-related ARF-GEFs with the Nextstrain program (Figure 1; Hadfield et al., 2018).

**Figure 1.**
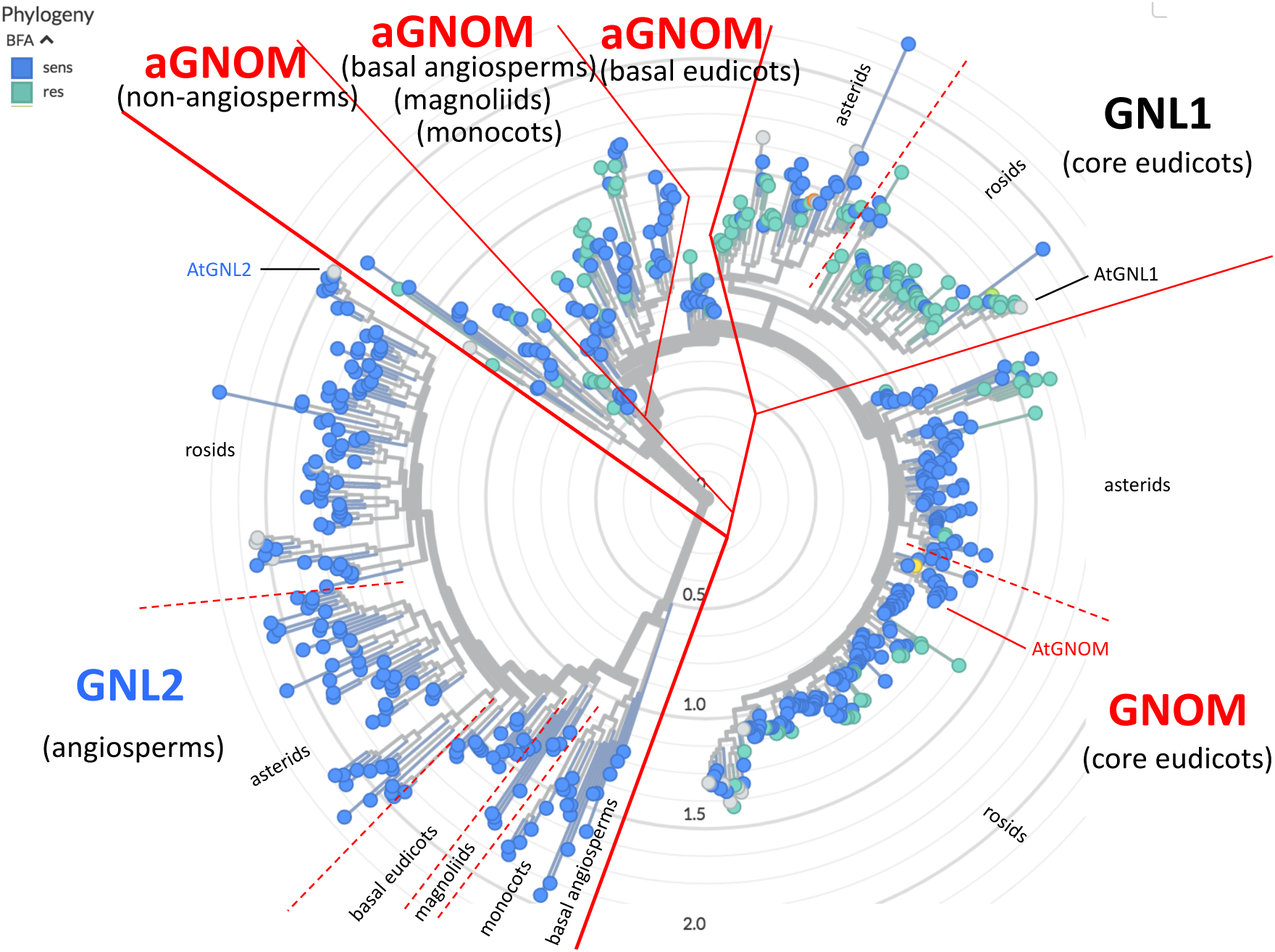
Phylogenetic analysis of GBF1-related ARF-GEFs from 175 plant species

Major clades are indicated (based on sequence similarity, using the Nextstrain program). ‘Sens’ (blue) and ‘res’ (aquamarine) indicate predicted sensitivity or resistance to brefeldin A (BFA) based on the analysis of *Arabidopsis thaliana* AtGNOM (sens), AtGNL1 (res) and AtGNL2 (sens) proteins. aGNOM, ancient GNOM. Gene sequences and list of species analysed in Suppl. Data file 1.

Pollen-specific GNL2 orthologs form a distinct clade comprising sequences from angiosperms only (Figure 1). Most species have a single gene encoding the AtGNL2 ortholog. Exceptions predominantly include basal angiosperms, basal monocots and magnoliids as well as polyploid species (Suppl. Table S1D-H). The GNL2 clade likely reflects the phylogenetic relationship of angiosperms and will not be considered further.

The other GBF1-related ARF-GEF sequences were grouped into 5 clades of which 3 represented “ancient GNOM” (aGNOM) in (i) non-angiosperm streptophytes (17 species), (ii) basal angiosperms, monocots, magnoliids, Chloranthales and Ceratophyllales (14 species), and (iii) basal eudicots (10 species) while 2 other clades were designated (iv) “GNL1” and (v) “GNOM”. The last two clades each comprise sequences from both asterids (49 species) and rosids (85 species) of the core eudicots (Figure 1). Thus, sequence diversity of the non-GNL2 paralogs only became detectable as AtGNOM-related and AtGNL1-related clades in the core eudicots with their split into the major branches of asterids and rosids.

Many non-angiosperm streptophytes – especially algae and mosses – have a single gene encoding a GBF1-related ARF-GEF which is always more closely related to AtGNOM than to AtGNL1 (Suppl. Table S1A-B). By contrast, gymnosperm species of different orders generally have two paralogs named GNOMa and GNOMb, which are >80% identical. These paralogs form two separate branches of a phylogenetic tree and may have originated from a whole-genome duplication in a common ancestor of gymnosperms (Suppl. Figure S1; Li et al., 2015; Stull et al., 2021). There is a trend of increasing sequence similarity to AtGNOM, from approx. 50% in algae to at least 60% in mosses and ferns, and more than 70% in gymnosperms (Suppl. Table S1A-C). When another *GNOM*-related gene copy is present in the genome of a non-angiosperm species, the protein encoded is also more closely related to AtGNOM than to AtGNL1. In general, however, the second paralog tends to be less similar to AtGNOM than the first one. In conclusion, the non-angiosperm species have AtGNOM-related ARF-GEFs. If duplicated, one of the two paralogs appears to be under less selection pressure than the other one.

In basal angiosperms (aka ANA clade) where the AtGNL2-related ARF-GEF makes its first appearance, several species have AtGNOM-related ARF-GEFs encoded by single-copy genes (Suppl. Table S1D). The proportion is very similar in magnoliids (7 of 11 species analysed) (Suppl. Table S1D). Several orders of monocots including the highly derived Poales also include species with a single-copy gene encoding a GNOM-related ARF-GEF (11 of 52 species analysed; Suppl. Table S1E). When another gene copy is present the protein encoded is generally less related by sequence to AtGNOM and even less to AtGNL1 than the first one. The GNOM-related paralogs of the monocot species analysed do not form two separate branches of GNOMa and GNOMb. Rather, each order has their own pair of GNOM paralogs (Suppl. Figure S2). The basal eudicots usually have two paralogs which both are more closely related to AtGNOM than to AtGNL1 (Suppl. Table S1F).

The core eudicots display the distinct clades of AtGNL1-related and AtGNOM-related ARF-GEFs (Figure 1). The former have reduced similarity to AtGNOM compared to the latter (Suppl. Table S1G-H). Nonetheless, both paralogs from the same species are more closely related to AtGNOM than AtGNL1, although this difference becomes less pronounced for the AtGNL1-related paralog. In the rosids but not the asterids, the AtGNL1-related paralog tends to show higher sequence similarity to AtGNL1 than does the AtGNOM-related paralog (Suppl. Table S1G-H). Markedly increased similarity to AtGNL1 as opposed to AtGNOM was only displayed among the AtGNL1-related ARF-GEFs in the species of Brassicales (Suppl. Table S1H). Taken together, these results suggest that one paralog is under selection for GNOM function whereas the additional one(s) appear(s) to drift, with the possible exception of those in rosid species. In line with this, many species which are scattered among all major clades of angiosperms have only a single copy of *GNOM*-related gene, including 44 in the monocots and core eudicots (Suppl. Table S1E, G-H). This suggests recurrent loss of the drifting paralog(s) (see below).

### Ortho-synteny groups identify *AtGNOM* and *AtGNL1* orthologs in eudicot evolution

One way to determine orthology among homologous genes is to assess their syntenic relationship, as also recently reported for Brassicaceae (Walden and Schranz, 2023). In the core eudicots, each of the two genes encoding GNOM-related paralogs is embedded in its own specific syntenic environment which appears to be largely conserved between different orders and which we therefore designate ‘ortho-synteny group’ (OSG) (Figure 2; see Suppl. Figure S3A). The standard GNOM-OSG1 is present in 11 of 17 orders of the rosid clade and in 6 of 15 orders of the asterid clade (Figure 2A-B). Likewise, 13 of 17 orders of the rosid clade and 10 of 15 orders of the asterid clade have the standard GNL1-OSG2 (Figure 2C-D). The non-standard OSGs will be presented in the next section.

**Figure 2.**
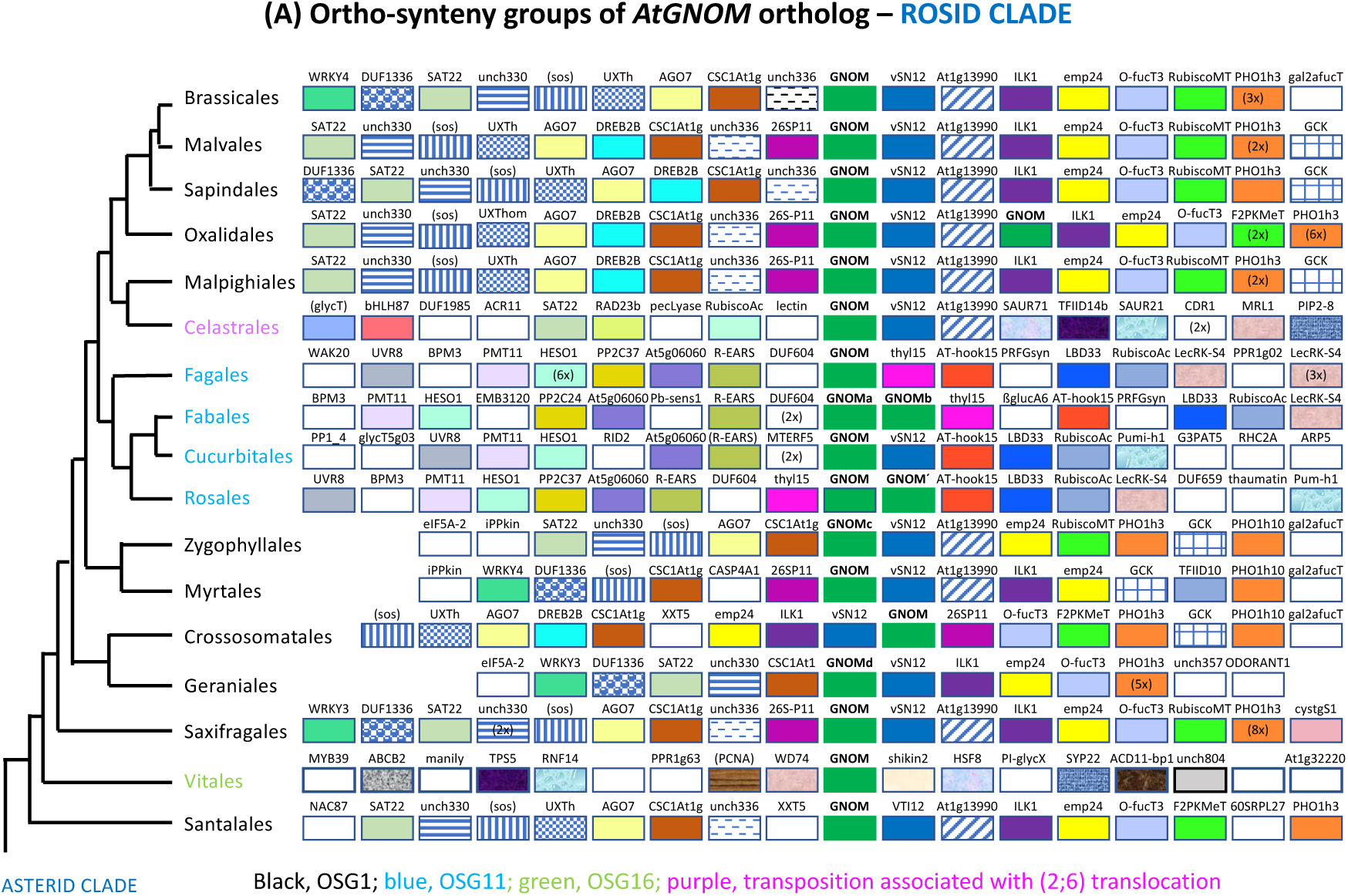
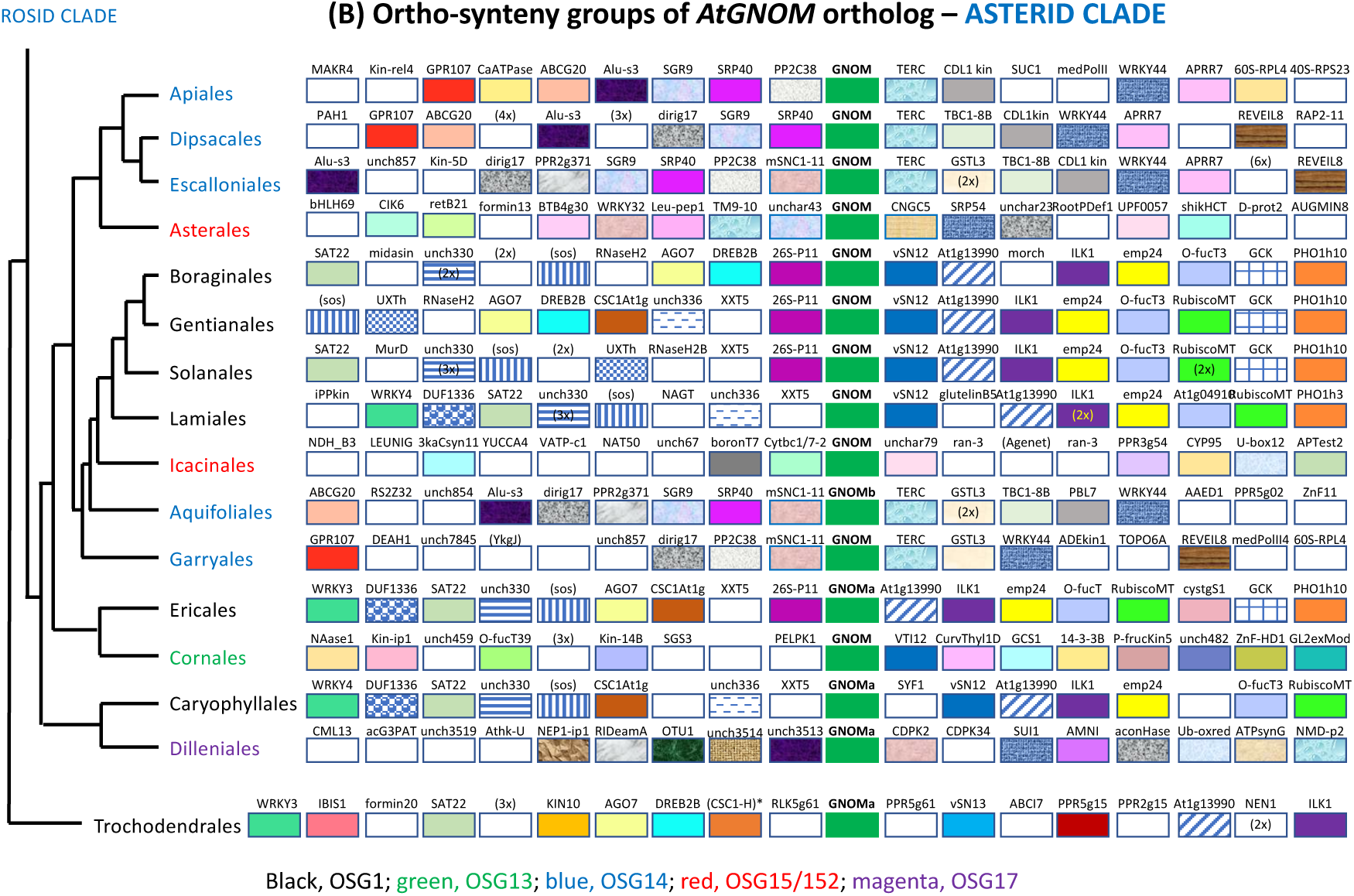
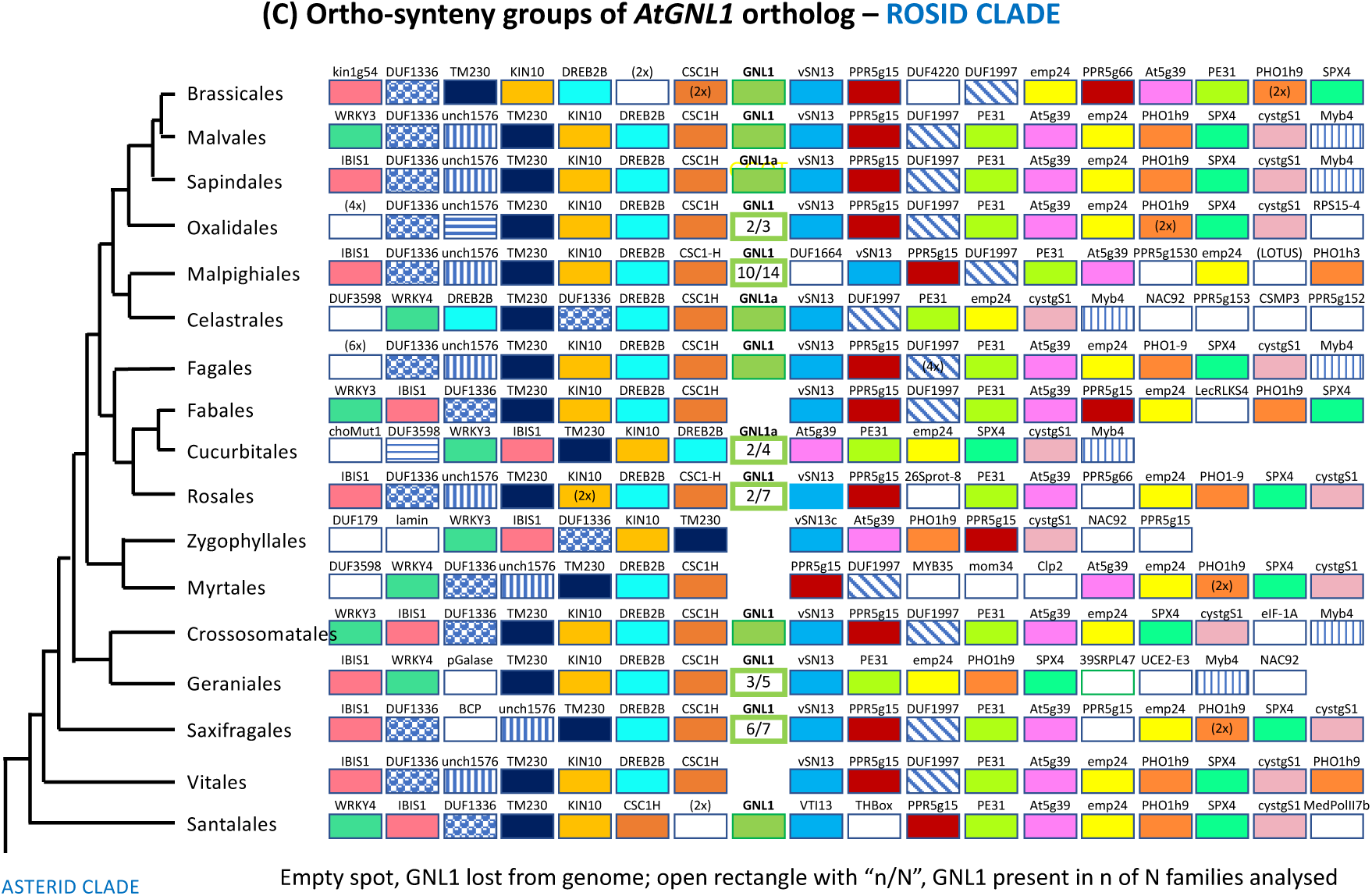
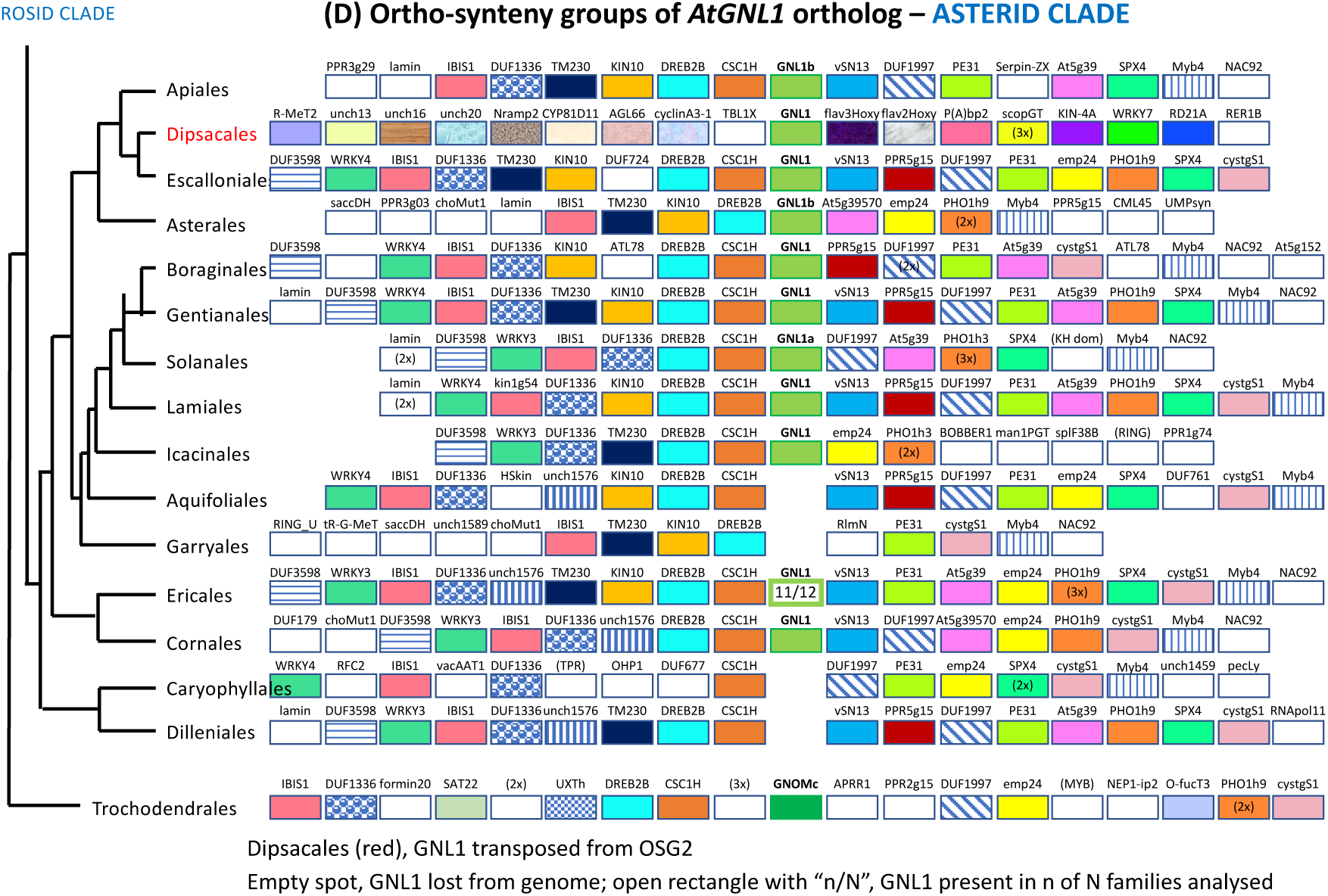
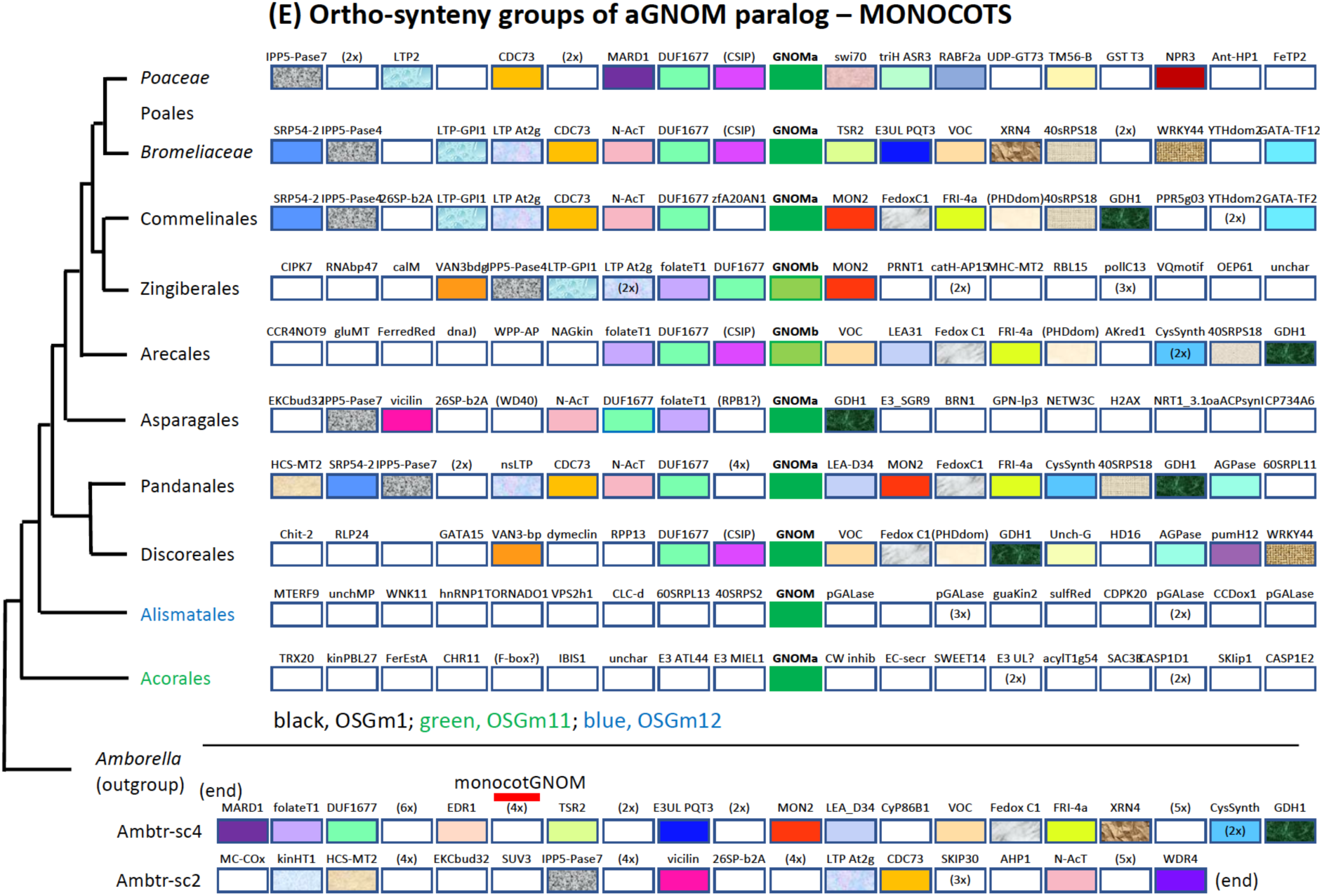
A,B. Ortho-synteny groups of *AtGNOM* orthologs in the core eudicot clades Rosidae **(A) and Asteridae (B).** The flanking genes are colour-coded or pattern-coded (see Suppl. Figure S3A). The positions of AtGNOM orthologs in different OSGs have been mapped onto the genome of *Theobroma cacao* (see Suppl. Figure S3B). The cladogram on the left is based on OTPTI (2019). Note OSG11 is shared by orders of the N_2_-fixing clade (A, light blue) whereas OSG14 is shared by several orders from among Lamiids and Campanulids (B, blue). For details within orders, see Suppl. Figures S4-S35. Figure 2C,D. Ortho-synteny groups of *AtGNL1* orthologs in the core eudicot clades Rosidae (C) and Asteridae (D). The flanking genes are colour-coded or pattern-coded (see Suppl. Figure S3A). Figure 2E. **Ortho-synteny groups of *aGNOM* in the monocot clade.** The recurrent flanking genes are colour-coded. The position of *aGNOM* is indicated in the relevant precursor OSGs (themselves lacking *aGNOM* which is located at sc2, Mb 9.6, at least 2 Mb away from monocotGNOM) in the basal angiosperm *Amborella trichopoda* (bottom). ‘GNOMa’ (dark green) and ‘GNOMb’ (light green) are distinguished by being slightly more or less closely related to AtGNOM, respectively. The cladogram on the left is based on OTPTI (2019). Note the different OSGs in the basal orders Acorales (OSGm11, green) and Alismatales (OSGm12, blue). For details within orders, see Suppl. Figure S36.

The cladogram on the left is based on OTPTI (2019). Note that the AtGNL1 ortholog was lost from OSG2 in several orders; empty spot, loss of *AtGNL1* ortholog in all families analysed; open rectangle, *AtGNL1* ortholog present in “n” of “N” families analysed. For details within orders, see Suppl. Figures S4-S35.

The paralogous gene most closely related to *AtGNOM* (designated *AtGNOM* ortholog from here on) is usually flanked by a conserved set of genes such as those encoding v-SNARE VTI12 (Uemura et al., 2004) and CSC1-like channel At1g69450 (Hou et al., 2014). The other paralogous gene, which is less closely related to *AtGNOM* (designated *AtGNL1* ortholog from here on), is flanked by a different conserved set of genes such as those encoding the paralogs v-SNARE VTI13 and CSC1-like channel HYP1. Other flanking genes also encode paralogs. These include phosphate transporter PHO1 homologs 3 or 10 and PHO1 homolog 9 (Wang et al., 2004), the functionally overlapping transcriptional regulators WRKY3 and WRKY4 (Liu et al., 2023a), DUF1336 domain proteins (Lv et al., 2023), AP2-related transcription factors involved in ethylene response named RAP2-11, ERN1, DREB or ERF119 (Feng et al., 2020) and ER-located emp24 domain proteins (Montesinos et al., 2013) (Figure 2; for detailed information on families within orders, see Suppl. Figures S4-S35). These observations suggest that the paralogous chromosomal segments arose by duplication as will be discussed in a later section (see below).

### Alternative OSGs by transposition of the *AtGNOM* ortholog

Although the unique OSGs of the two *GNOM*-related paralogs are largely conserved among the core eudicots, the *ARF-GEF* gene was lost from its standard OSG in several orders, with paralog-specific consequences (Figure 2). The *AtGNL1* ortholog generally disappeared from the respective genome altogether. This loss occurred in 4 of 17 rosid orders and in 4 of 15 asterid orders; in the asterid order Dipsacales, however, the *AtGNL1* ortholog was transposed (Figure 2C-D). By contrast, the *AtGNOM* ortholog remained in all genomes analysed but integrated into different genomic regions, establishing new GNOM-OSGs. This change of genomic location occurred in 6 of 17 rosid orders and in 9 of 15 asterid orders (Figure 2A-B). In several additional orders, both transposition of the *AtGNOM* ortholog and loss of the *AtGNL1* ortholog only affected one or more but not all families. Each transposition event was confined to a specific group of closely related orders or families within an order (colour-coded in Figure 2A-B). The insertion sites were mapped onto the *Theobroma cacao* (Malvales) genome as a reference (Suppl Figure S3B). In the following, we present specific cases such as the 4 orders of the N2-fixing fabid clade among the rosids and 6 orders among the asterids which purportedly do not form a monophyletic clade. All alternative OSGs as well as the loss of the *GNOM* paralogs from their respective standard OSGs are documented in detail in Suppl Figures S4-S35.

The fabid species analysed represent 3, 7, 4 and 5 families of the orders Fabales, Rosales, Cucurbitales and Fagales, respectively (Suppl. Figures S10-S13). In each case, the standard OSG1 lacks the *AtGNOM* ortholog which instead has been inserted into a different genomic environment designated OSG11 (see Figure 2A for overview; details in Suppl Figures S10-S13). This new OSG11 includes marker genes encoding the transcription factor AThook15 and the thylakoid luminal protein 15kDa (Thyl-15) (Figure 2A; Suppl Figures S10-S13). Essentially the same change occurred in all 4 orders of fabids, although there are differences in the position of the *AtGNOM* ortholog relative to the marker genes AThook15 and Thyl-15 (Figure 2A; Suppl Figures S10-S13). These and other differences as well as the occurrence of tandem duplications in Rosales and Fabales but not in Fagales or Cucurbitales suggest subsequent events of local reorganization. Regarding the loss of the *AtGNL1* ortholog from OSG2, this event happened in all 3 families of Fabales, in most families of Rosales and Cucurbitales but not at all in Fagales (Figure 2C; Suppl Figures S10-S13).

A complex case of *AtGNOM* ortholog transposition is presented in the asterids, involving orders from the two major clades, lamiids and campanulids (Figure 2B; Suppl. Figures S25-S29). The campanulid orders Apiales, Dipsacales and Escalloniales share the same transposition event to OSG14 with the non-core lamiids Aquifoliales and Garryales. These five orders are not considered members of a monophyletic clade, unlike the N2-fixing fabids discussed above (Figure 2; Stull et al., 2020; Zhang et al., 2020b). In the Araliaceae family of the Apiales, the *AtGNOM* ortholog underwent transposition to OSG141, which however might be secondary (Suppl. Figure S27). In Asterales, which are purportedly phylogenetically related to Apiales, Dipsacales and Escalloniales, the *AtGNOM* ortholog was transposed to different OSGs (OSG15 in Asteraceae and Goodeniaceae, OSG151 in Campanulaceae; these are just 8 Mb apart, suggesting consecutive events) (Suppl. Figure S30). Interestingly, the basal lamiid order Icacinales appears to have the *AtGNOM* ortholog transposed to nearly the same place as that in *Campanulaceae* (OSG152) such that the two *AtGNOM* orthologs are only about 10 genes apart (Suppl. Figure S35). By contrast, the core-lamiid orders of the asterids display the standard ortho-synteny groups OSG1 and OSG2 (Figure 2B, 2D; Suppl. Figures S21-S24).

In conclusion, the common syntenic relationships (“standard” GNOM-OSG1 and GNL1-OSG2) are detectable across the core eudicots, which suggests that they originated in the basal eudicots, i.e. before the split between rosids and asterids, and that the transposition of the *AtGNOM* ortholog to new OSGs as well as the loss of the *AtGNL1* ortholog were secondary.

In contrast to the core eudicots with their distinct GNOM-OSG1 and GNL1-OSG2, the monocots display a different pattern (Figure 2E). In 8 of 10 orders analysed, a set of largely conserved genes flanks the single-copy *AtGNOM*-paralogous gene or one of the two paralogs named GNOMa or GNOMb. This monocot-specific ortho-synteny group OSGm1 appears derived from 2 precursor segments present in the genome of the basal angiosperm *Amborella trichopoda* and 2 Mb away from AmbtrGNOM (Figure 2E, bottom). The other GNOM paralog might be embedded in the same OSGm1 (3 orders) or transposed to new genomic locations that differ between orders (Suppl. Figure S36). In 2 basal orders, none of the GNOM paralogs is present in the monocot standard OSGm1. In conclusion, the OSG analysis strongly suggests the GNOM paralogs lack divergent evolutionary trajectories in the monocot clade of angiosperms.

### Assembly of the standard OSGs 1 and 2 in the basal eudicots

The early-diverging eudicots with sequenced genomes include Ranunculales, Proteales, Buxales and Trochodendrales. Also, Gunnerales are commonly regarded as sister clade to the core eudicots (OTPTI, 2019; Zuntini et al., 2024). Additional orders such as Saxifragales, Santalales and Dilleniales have not uncontroversially been placed, although they were here assigned to the core eudicots (see Figure 2; OTPTI, 2019; Liu et al., 2023b; Zuntini et al., 2024; see Suppl. Figure S37). The last 3 orders display the standard ortho-synteny groups OSG1 and OSG2 as present in the core eudicots (Suppl Figures S18, S20, S34).

In the basal angiosperms such as Amborella or Nymphaea, the future gene sets flanking the *GNOM*-related paralogs in the core eudicots are still two separate synteny groups (Figure 3). In addition, they are located on different chromosomes (chr4, chr10) than AmbtrGNOM (chr3). These sets are first joined to form a contiguous synteny group in basal eudicots but not in monocots nor magnoliids (see Suppl. Figure S38). This formation of an incipient standard OSG was detected in all 12 families of basal eudicots analysed (7 in Ranunculales, 3 in Proteales, and 1 each in Buxales and Trochodendrales). In Ranunculales, Proteales and Buxales, these protoOSGs still lack the *GNOM*-related paralogs which are located at different positions). Trochodendrales is the only basal-eudicot order in which the GNOM-related paralogs are part of the incipient OSG1/2 (Figure 3; Suppl. Figures S38). In this regard, Trochodendrales are closer to the core eudicots than are Buxales, although the two orders have been proposed sister clades (Wang et al., 2022; Chanderbali et al., 2022; Sun et al., 2016; Zuntini et al., 2024).

**Figure 3.**
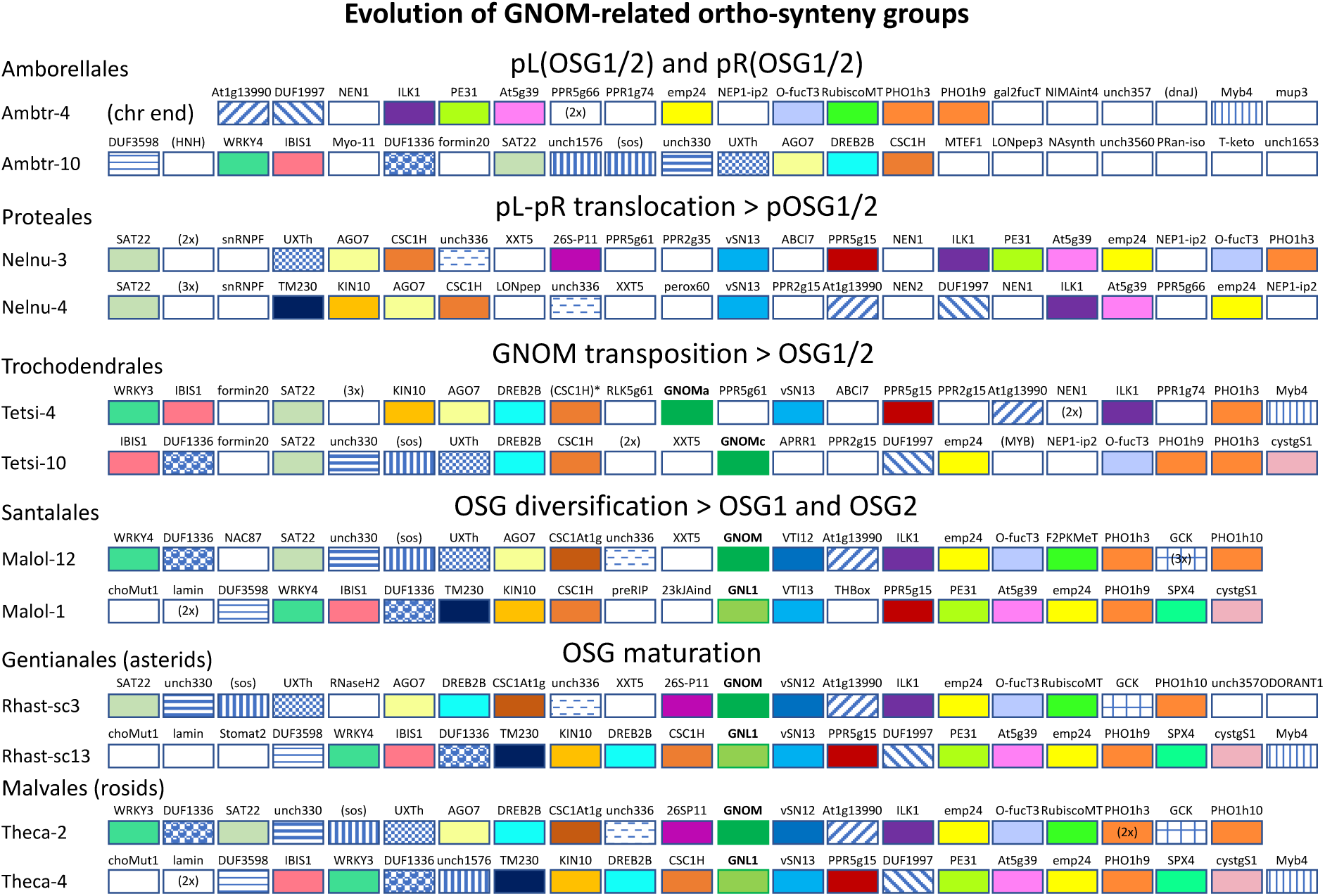
Evolution of the standard OSGs 1 and 2 in the basal eudicots.

Flanking genes are colour-coded or pattern-coded as in Figure 2. *Amborella trichopoda* (Ambtr) represents the basal angiosperms, *Nelumbo nucifera* (Nelnu) and *Tetracentron sinense* (Tetsi) the basal eudicots, *Malania oleifera* (Malol), *Rhazya stricta* (Rhast) and *Theobroma cacao* (Theca) the core eudicots. For explanation, see text.

The protoOSGs also include other genes that are not present in the mature OSGs of core eudicots. Furthermore, there is no clear distinction between (proto)OSG1 and (proto)OSG2. For example, OSG1 marker *SAT22* is linked to both OSG1 members *AGO7* or *O-fucT3* and to OSG2 members *IBIS1* or *CGS1*. Moreover, only VTI13 and CSC1-like HYP1 but not their paralogs VTI12 or CSC1-like At1g69450, respectively, are present in basal eudicots, unlike the situation in core eudicots (Figure 3). Nonetheless, the two standard OSGs appear to have originated in the basal eudicots. This is supported by the analysis of OSG1 and OSG2 in *Myrothamnus flabellifolia* which is the only species of the order Gunnerales whose genome has been assembled (phytozome 13; https://phytozome-next.jgi.doe.gov/). Gunnerales represent the sister clade to the core eudicots (OTPTI, 2019; Zuntini et al., 2024). Although *MyrflGNOM* and five flanking genes have undergone a transposition of to a different genomic location, superimposed on an inversion of the remaining OSG1, this OSG1 has the mature organization characteristic of those in core eudicots (Suppl. Figure S38). Similarly, OSG2 also resembles GNL1-OSG2 from core eudicots, except that *MyrflGNL1* has been lost from the genome.

### Functional analysis of GNOM-related paralogs in transgenic Arabidopsis

To assess whether GNOM-related ARF-GEFs from other species are functionally equivalent to AtGNOM or AtGNL1, we generated transgenic Arabidopsis plants that expressed the coding sequence (*CDS*) or, in some cases, the genomic sequence of *GNOM*-paralogous genes from the *AtGNOM cis*-regulatory sequences. This assay was based on the earlier observation that the *CDS* of *AtGNL1* expressed from the *AtGNOM cis*-regulatory sequences was unable to rescue *gnom* mutant plants but rescued *gnl1* mutant plants (Richter et al., 2007). As a positive control, we first tested putative GNOM and GNL1 orthologs from *Brassica napus*, which show 95% and 87% sequence identity to their *Arabidopsis thaliana* counterparts, respectively (Suppl. Table S1H). BranaGNOM and BranaGNL1 indeed behaved as expected in that transgenic *BranaGNOM* rescued the *gnom^sgt^* deletion mutant whereas *BranaGNL1* did not (Figure 4A-B; Suppl. Table S2). In addition, BranaGNOM did not interact with AtGNL1 and conversely, BranaGNL1 did not interact with AtGNOM in coimmunoprecipitation experiments whereas BranaGNOM and BranaGNL1 interacted with AtGNOM and AtGNL1, respectively (Figure 4C-D). Furthermore, their subcellular localization mirrored that of their respective *Arabidopsis thaliana* counterparts (Figure 5A-D). Thus, BranaGNOM and BranaGNL1 appear to be functionally true orthologs of AtGNOM and AtGNL1, respectively.

**Figure 4.**
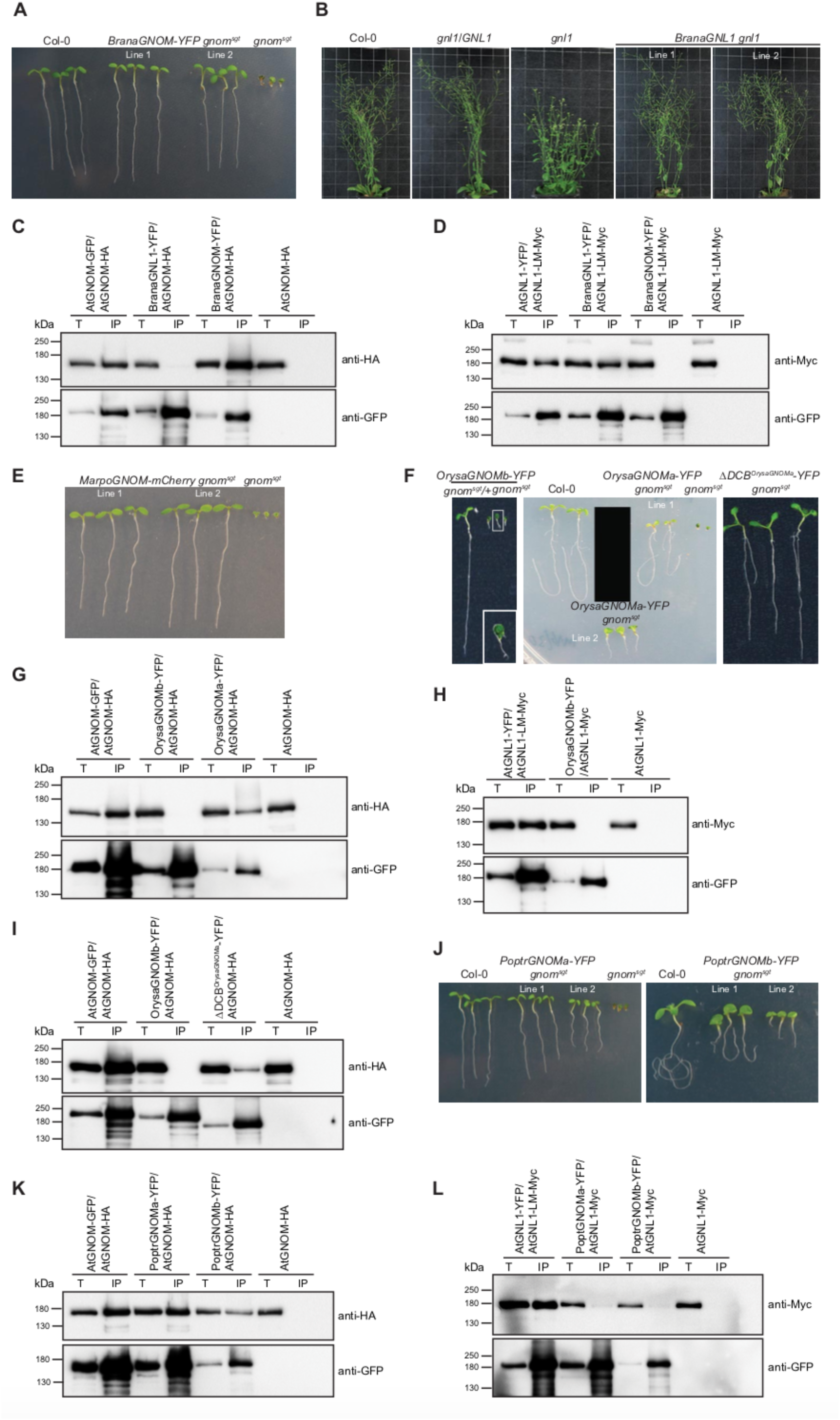
Functional analysis and interaction studies of *GNOM*-related paralogs in transgenic *Arabidopsis thaliana*. (**A-D**) *Brassica napus*. (**A-B**) Rescue of (A) *gnom^sgt^* by *BnGNOM*, (B) *gnl1* by *BnGNL1*; (**C-D**) co-immunoprecipitation interaction assays of BnGNOM-YFP and BnGNL1-YFP with (C) AtGNOM-HA, (D) AtGNL1-LM-Myc, using GFP-Trap-agarose followed by immunoblot analysis with anti-GFP and anti-Myc/anti-HA antibodies. (**E**) *Marchantia polymorpha*: rescue of *gnom^sgt^* by *MarpoGNOM*. (**F-I**) *Oryza sativa*. (**F**) Rescue of *gnom^sgt^*by *OsGNOMa*, *OsGNOMb* and *βDCB^OsGNOMa^*; (**G-I**) co-immunoprecipitation interaction assay of (G) OsGNOMa-YFP and OsGNOMb-YFP with AtGNOM-HA, (H) OsGNOMb-YFP with AtGNL1-LM-Myc, (I) OsGNOMb-YFP and βDCB^OsGNOMa^-YFP with AtGNOM-HA. (**J-L**) *Populus trichocarpa*. (**J**) Rescue of *gnom^sgt^* by *PtGNOMa* and *PtGNOMb*; (**K-L**) co-immunoprecipitation interaction assay of PtGNOMa-YFP and PtGNOMb-YFP with (K) AtGNOM, (L) AtGNL1-Myc. T, total; IP, immunoprecipitate; kDa, kilodalton. For quantitative rescue data, see Suppl. Table S2.

**Figure 5.**
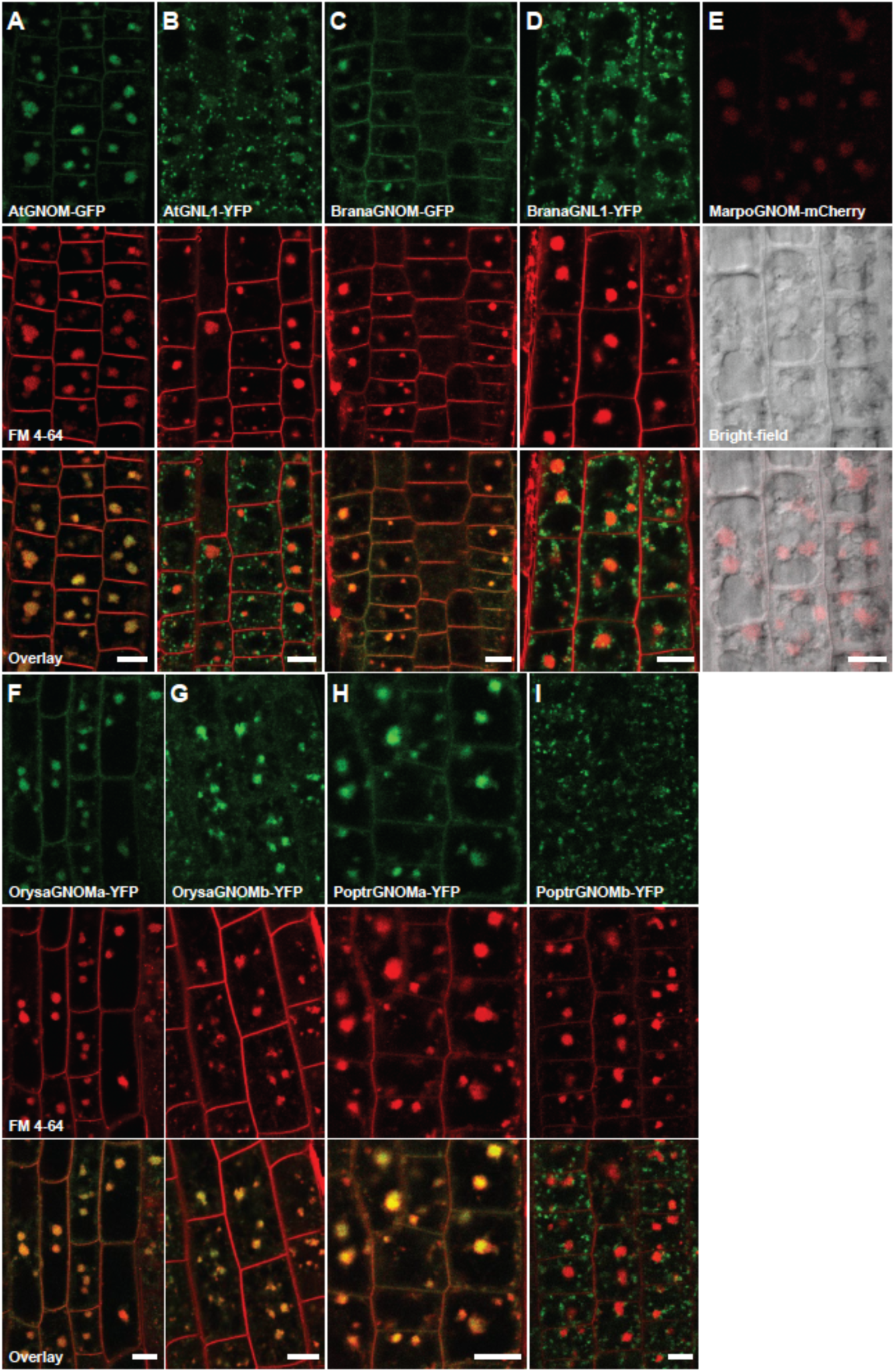
Subcellular accumulation of GNOM-related paralogs in transgenic *Arabidopsis thalana* Live-cell imaging of AtGNOM-GFP (**A**), AtGNL1-YFP (**B**), BranaGNOM-YFP(**C**), BranaGNL1-YFP (**D**), MarpoGNOM-mCherry (**E**), OrysaGNOMa-YFP (**F**), OrysaGNOMb-YFP (**G**), PoptrGNOMa-YFP (**H**) and PoptrGNOMb-YFP (**I**) after BFA treatment (50μM for 1h) of seedlings in the presence of SynaptoRed C2 (FM4-64 equivalent; A-D, F-I). MarpoGNOM-mCherry (E, red signals) was imaged in bright field. Scale bar, 10µm.

In order to identify the ancient GBF1-type ARF-GEF in plants, we tested the only GNOM-related ARF-GEF from the liverwort *Marchantia polymorpha* representing the earliest-diverging land plant clade. MarpoGNOM is 65% identical to AtGNOM but only 55% to AtGNL1. When expressed in transgenic *Arabidopsis thaliana* plants, MarpoGNOM tagged with mCherry rescued the *gnom^sgt^* deletion mutant (Figure 4E). This result supports the notion that the ancient GBF1-type plant ARF-GEF was functionally equivalent to AtGNOM, implying that the AtGNL1 ortholog originated later in plant evolution.

Next, we tested the two GNOM-related ARF-GEFs from the monocotyledonous angiosperm *Oryza sativa* representing the ‘ancient GNOM’ clade (see Figure 1). These proteins have comparatively low sequence identity values, sharing 70% or 62% identical amino acid residues with AtGNOM and 56% or 55% with AtGNL1, respectively (Suppl. Table S1E).

Nonetheless, both GNOM-related paralogs from rice were able to rescue the axis formation defect of the *gnom^sgt^* deletion mutant, thus surpassing AtGNL1 (Figure 4F; Richter et al., 2007). However, there was a quantitative difference in that the more closely related OrysaGNOMa performed much better that the more diverged OrysaGNOMb (Figure 4F; Suppl. Table S2). In addition, OrysaGNOMa but not OrysaGNOMb interacted with AtGNOM and OrysaGNOMb also did not interact with AtGNL1 in coimmunoprecipitation experiments (Figure 4G-I). Also, with its DCB domain deleted, OrysaGNOMa was still able to rescue the *gnom^sgt^*deletion mutant, thus behaving just like AtGNOM (Brumm and Singh et al., 2022). These results are consistent with the OSG analysis and the phylogenetic analysis of GNOM-related sequences which placed the monocots including *Oryza sativa* into the “ancient GNOM” clade (see Figure 1, Figure 2E).

Note BFA resistance of PoptrGNOMb prevents its accumulation in BFA compartments. We then analysed the two GNOM-related paralogs from *Populus trichocarpa* which is in the order of Malpighiales and thus in the rosid clade of core eudicots. Our analysis of syntenic relationships suggested that the *AtGNL1* ortholog has been lost from the *Populus trichocarpa* genome and the *AtGNOM* ortholog transposed to OSG12 and duplicated in tandem (see Suppl. Figure S8). This interpretation was functionally assessed in transgenic Arabidopsis plants (Figure 4J). Both *PoptrGNOM* transgenes were able to rescue the Arabidopsis *gnom^sgt^* deletion mutant, supporting their interpretation of being *AtGNOM* orthologs (Suppl. Table S2). In addition, both PoptrGNOM proteins interacted with AtGNOM but did not interact with AtGNL1 in co-immunoprecipitation experiments with extracts from transgenic Arabidopsis seedlings (Figure 4K-L). In conclusion, the functional analysis of *GNOM* paralogs from other plant species is consistent with their synteny-based classification of being either *AtGNOM* or *AtGNL1* orthologs.

We also tested the subcellular localization of the GNOM paralogs from other species in Arabidopsis seedling roots. We previously used brefeldin A (BFA) to visualize the localization of AtGNOM in endosomal BFA compartments (Geldner et al., 2003). All but 2 paralogs tested localized to BFA compartments (Figure 5). One exception was GNL1 from *Brassica napus* which behaves like AtGNL1 (Figure 5D, compare with Figure 5B). The other exception was PoptrGNOMb (Figure 5I). This result is consistent with its predicted BFA resistance since AtGNOM when rendered BFA-resistant did also not accumulate in BFA compartments (Geldner et al., 2003).

### DCB domain interaction in GNOM-related ARF-GEFs

In Arabidopsis, the DCB domain of GNOM interacts with both the complementary ΔDCB fragment and with the DCB domain of another GNOM protein (Anders et al., 2008). The DCB domain of GNL1 can only interact with the complementary ΔDCB fragment of either GNOM or GNL1 but not with another DCB domain of either GNOM or GNL1 (Brumm and Singh et al., 2022). Swapping experiments indicated that the N-terminal half of the DCB domain determines its interaction behaviour (Brumm and Singh et al., 2022). Thus, the DCB-DCB interaction is a distinguishing feature of GNOM.

We presumed that the difference in DCB-DCB interaction behaviour between AtGNOM and AtGNL1 arose during evolution of the eudicots, reflecting the different OSGs in which the GNOM paralogs are embedded. We thus tested the DCB domains of GNOM paralogs from species representing different clades for interaction with themselves, with the DCB of another GNOM paralog from the same species (if present), with the DCB domain of AtGNOM and with the complementary ΔDCB fragment of AtGNOM (Table 1).

**Table 1.** DCB-DCB interaction behaviour of GNOM-related paralogs in the yeast two-hybrid assay.

| Species | Order | Gene | DCB x self | x DCB(gene 2) | x DCB<sup>GNOM</sup> | x ΔDCB<sup>GNOM</sup> |
| --- | --- | --- | --- | --- | --- | --- |
| *core eudicots* |  |  |  |  |  |  |
| Arabidopsis thaliana | Brassicales | GNOM | Y | N | Y | Y |
|  |  | GNL1 | N | N | N | Y |
| Brassica napus | Brassicales | Brana1 | Y | N | Y | Y |
|  |  | Brana2 | N | N | N | Y |
| Capsella rubella | Brassicales | Capru1 | Y | N | Y | Y |
|  |  | Capru2 | N | N | N | Y |
| Pistacia vera | Sapindales | Pisve1 | Y | N | Y | Y |
|  |  | Pisve2 | N | N | N | Y |
| Populus trichocarpa | Malpighiales | Poptr1 | Y | Y | Y | Y |
|  |  | Poptr2 | Y | Y | Y | Y |
| Salix brachista | Malpighiales | Salbr1 | Y | Y | Y | Y |
|  |  | Salbr2 | Y | Y | Y | Y |
| Ricinus communis | Malpighiales | Ricco1 | Y | N | Y | Y |
|  |  | Ricco2 | N | N | N | Y |
| Amaranthus hypochondriacus | Caryophyllales | Amahy1 | Y | Y | Y | Y |
|  |  | Amahy2 | Y | Y | Y | Y |
| *basal eudicots* |  |  |  |  |  |  |
| Aquilegia coerulea | Ranunculales | Aquco1 | Y | Y | n.d. | Y |
|  |  | Aquco2 | Y | Y | n.d. | Y |
| *monocots* |  |  |  |  |  |  |
| Oryza sativa | Poales | Orysa1 | Y | Y | Y | Y |
|  |  | Orysa2 | Y | Y | Y | Y |
| *basal angiosperms* |  |  |  |  |  |  |
| Amborella trichopoda | Amborellales | Ambtr | Y | x | Y | Y |
| *streptophytic algae* |  |  |  |  |  |  |
| Klebsormidium nitens | Klebsormidiales | Kleni | Y | x | Y | Y |

The DCB domains of GNOM paralogs from the species indicated were tested for interaction with themselves (DCB x self), with the DCB domain of another paralog from the same species (x DCB(gene2)), with DCB of AtGNOM (x DCB^GNOM^) and the complementary ΔDCB fragment of AtGNOM (βDCB^GNOM^). Y, interaction; N, no interaction; n.d., not done; x, only 1 GNOM paralog gene in that species. (for gene numbers, see Suppl. Table S1A-H)

We first tested the DCB domain of GNOM-related ARF-GEFs encoded by single-copy genes in their respective species including the streptophytic alga *Klebsormidium nitens* and the basal angiosperm *Amborella trichopoda* (Table 1). The two DCB domains interacted with themselves, with the DCB domain of AtGNOM and with the ΔDCB fragment of AtGNOM, which we used as a positive control. Thus, DCB-DCB interaction appears to be an ancient feature of GNOM-related ARF-GEFs.

We then examined the DCB domain interaction behaviour of GNOM-related ARF-GEFs that are encoded by two genes in the same species and are both more closely related to AtGNOM than to AtGNL1. This group of species included the monocot *Oryza sativa* and the basal eudicot *Aquilegia coerulea* in the order of Ranunculales. The DCB domain of these four ARF-GEFs interacted with itself, with the DCB domain of the paralog from the same species and with the DCB domain and βDCB fragment of AtGNOM (Table 1). These results are consistent with the higher sequence similarity to AtGNOM than to AtGNL1. There was also no difference in DCB-DCB interaction behaviour between the two GNOM paralogs from *Amaranthus hypochondriacus* (Caryophyllales, asterids, core eudicots). For the rosid species *Salix brachista* and *Populus trichocarpa* (Malpighiales), the results were the same, both paralogs behaving like AtGNOM in the DCB-DCB interaction assay. These results are consistent with the analyses of sequence similarities and syntenic relationships, suggesting that the *At1GNL1* ortholog was lost whereas the *AtGNOM* ortholog was duplicated (see above).

Finally, we identified species in which the DCB domain of the paralogous GNOM-related ARF-GEFs behaved differently, one like AtGNOM, the other like AtGNL1. This species group included *Ricinus communis* (Malpighiales), *Pistacia vera* (Sapindales), *Capsella rubella* (Brassicales) and *Brassica napus* (Brassicales).

In conclusion, the DCB-DCB interaction behaviour is consistent with the sequence similarities and syntenic relationships of the GNOM paralogs, strongly suggesting that the evolutionary trajectories leading to AtGNOM and AtGNL1 were started early in eudicot evolution.

## DISCUSSION

### The original plant-specific GBF1-type ARF-GEF was functionally equivalent to AtGNOM

The only GNOM paralog from the moss *Marchantia polymorpha* was able to completely rescue transgenic Arabidopsis that lacked the endogenous *GNOM* gene, indicating its functional equivalence to AtGNOM. Several other species from streptophytic algae to basal angiosperms have a single-copy gene encoding a GNOM paralog which each are more closely related by sequence to AtGNOM than to AtGNL1. Moreover, the DCB domain of the single-copy paralog from the alga *Klebsormidium nitens* and the basal angiosperm *Amborella trichopoda* both displayed the DCB-DCB interaction that distinguishes AtGNOM from AtGNL1. Thus, a single-copy gene encoding an AtGNOM-orthologous ARF-GEF seems to be the evolutionary starting point for the duplication and subsequent diversification of GNOM-related paralogs in the angiosperm lineage.

### Duplication and functional divergence of GNOM paralogs in eudicot evolution

Being able to distinguish the GNOM-related paralogs by their OSG embedding (see Figure 2), we can sketch a likely scenario of how duplication and functional diversification of the somatic paralogs might have occurred in eudicot evolution (Figure 6).

**Figure 6.**
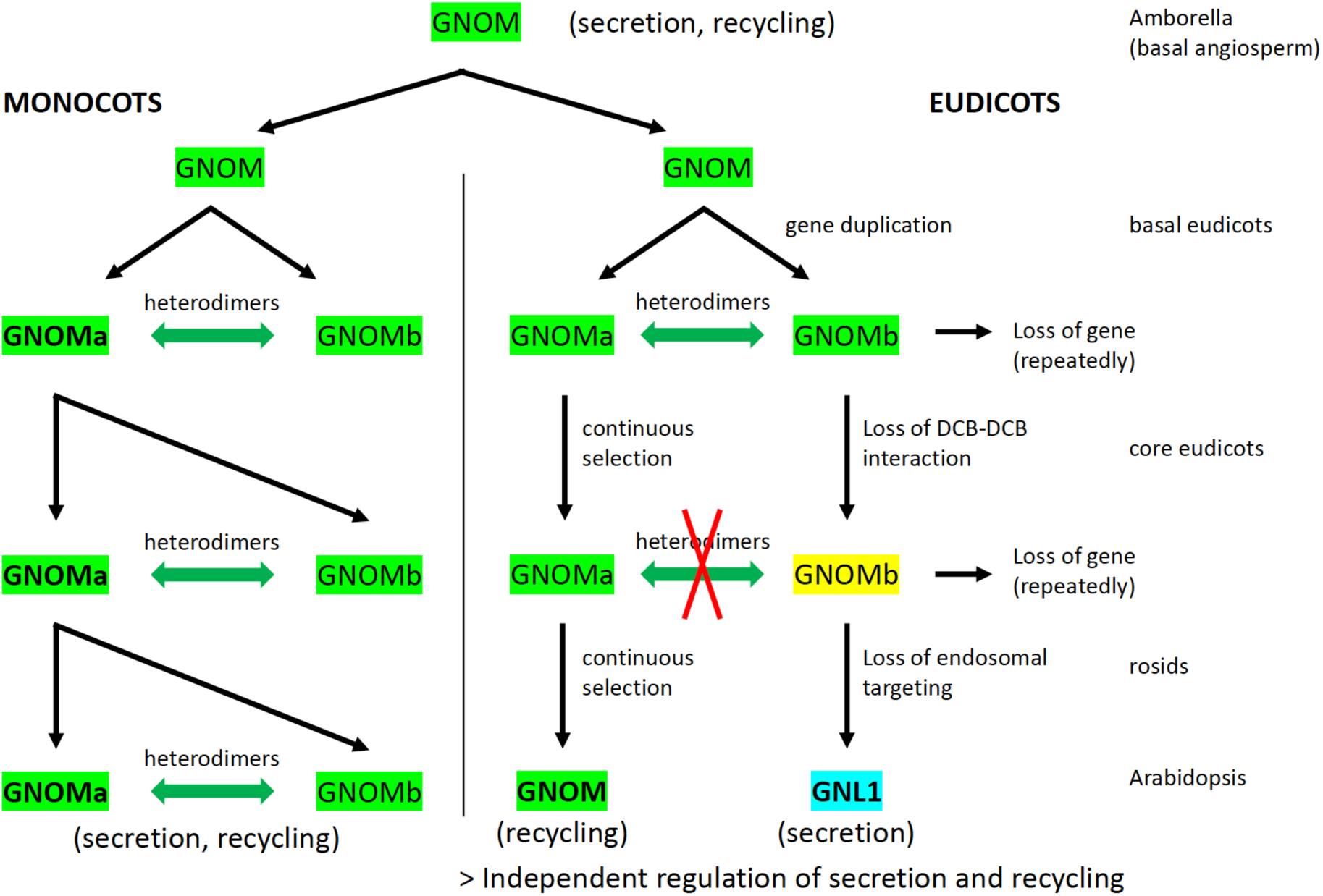
Evolutionary trajectories of GNOM paralogs in angiosperms (model). Following gene duplication, the formation of GNOMa-GNOMb heterodimers is eventually disrupted in eudicots but not in monocots. In monocots (*left*), GNOM-paralogous gene duplicates did not initiate divergent evolutionary trajectories. In eudicots (*right*), however, while paralog GNOMa (i.e. AtGNOM ortholog) is under strong selection, paralog GNOMb (i.e. AGNL1 ortholog) is free to evolve, repeatedly resulting in degeneration and eventual loss from the genome. In the rosid branch of core eudicots, paralog GNOMb not only experiences loss of endosomal targeting which is a unique feature of AtGNOM, but also seems to acquire a new (selectable) function, eventually giving rise to AtGNL1.

Gene duplication of the somatic GNOM-related paralogs occurred repeatedly, e.g. as part of the ancient whole-genome duplication (WGD) events (ψ, α, τ, ν) specific to each clade of mesangiosperms (approx. 120 mio years ago) or during later WGD events at the cretaceous-paleogene boundary (approx. 66 mio years ago) or even more recently (less than 20 mio years ago; Zhang et al., 2020b). The loss of DCB-DCB interaction as the result of mutation in one of the two GNOM paralogs can be envisaged as the next critical step, which occurred in eudicot evolution (Figure 6). As shown in Arabidopsis, the exclusive DCB-DCB interaction of GNOM essentially prevents the formation of GNOM-GNL1 heterodimers (Brumm and Singh et al., 2022). Following the loss of DCB-DCB interaction, one of the two paralogs (the *AtGNOM* ortholog) was then under continuous selection while the other paralog GNOMb (the *AtGNL1* ortholog) was free to drift, accumulating mutations over time. GNOMb was eventually lost from the genome in many species. However, an important change occurred in the rosid branch of the core eudicots when paralog GNOMb started to become more closely related to AtGNL1 and might have lost the ability of associating with endosomal membranes. As a consequence, there might have been selection for improving the ability of GNOMb to perform the ancient GBF1 function in retrograde trafficking from Golgi stacks to the ER. The advantage of this functional specialization would have been independent regulation of the secretory pathway from GNOM-dependent endosomal recycling to the plasma membrane. For example, the fast root gravitropic response only requires GNOM-dependent endosomal recycling but no de-novo secretion mediated by GNL1 (Richter et al., 2007).

When the functional change of the AtGNL1 ortholog occurred in rosid evolution cannot precisely be inferred from the available data. The increasing sequence similarity that all GNOM paralogs experience in Brassicales suggest that the change has occurred earlier. Additionally, the *AtGNL1* ortholog is no longer lost from the genome in Sapindales, Malvales and Brassicales, in contrast to the closely related orders of Oxalidales and Malpighiales (see Figure 2).

Whole-genome duplications occurred independently in the major clades of angiosperms such as monocots and eudicots (Zhang et al., 2020b). Unlike the eudicots, however, the monocots did not establish distinct OSGs of the two GNOM paralogs that would be maintained during monocot evolution. Moreover, the GNOM paralogs of *Oryza sativa*, which represents the highly diverged grass family in our analysis, can still form heterodimers by DCB-DCB interaction and displayed GNOM function in transgenic Arabidopsis plants, although to different degrees. Thus, no evolution towards an AtGNL1 equivalent appears to have occurred in the non-eudicots.

### Concluding remarks

The ancient plant-specific GBF1-type ARF-GEF was a bi-functional GNOM ortholog, mediating both endosomal recycling and retrograde Golgi-ER traffic. Although GNL1 looks ancient because of its functional similarity to mammalian GBF1, it is derived. This was most convincingly revealed by the analysis of the relevant ortho-synteny groups (OSGs), which helped to track the evolutionary trajectories of paralogs. This approach enabled the identification of orthologs among paralogous proteins and thus helped distinguish different evolutionary steps, from early separation of the dimerising orthologs to their later functional diversification.

## MATERIAL AND METHODS

### Plant growth and Media

Arabidopsis seedlings were initially grown on Agar plates (1/2MS media, 1% sucrose and 0.8% agar) for 6-7 days and then transferred to soil under long day light condition (16h) at 23°C.

The following previously published lines were used in this study: *gnom^sgt^* mutant (Brumm et al., 2020), *gnl1* mutant (Richter et al., 2007), *GNOM-HA (*Anders et al., 2008)*, GNOM-GFP (*Geldner *et al., 2003), GNL1-LM-Myc* (Richter et al., 2007), *GNL1-Myc* (Richter et al., 2007).

### Cloning in plant expression vectors, generation of transgenic plants and genotyping

The genomic DNA of *BranaGNL1*, *OrysaGNOMb*, *OrysaGNOMa* lacking first *271 amino acids* (*ΔDCB^OrysaGNOMa^*) and *PoptrGNOMb* were cloned in frame with c-terminal YFP in *pGreenII(Bar)* binary vector containing GNOM regulatory sequences using ApaI and NotI restriction sites.

Full length coding sequences of *BranaGNOM*, *PoptrGNOMa* and *OrysaGNOMa* were synthesized by the company BaseClear B.V. (Leiden, Netherlands) and cloned into *pGII(Bar)-pGNOM::YFP* vector (Brumm and Singh et al., 2022) using SalI and NotI (*BranaGNOM*), SalI and SmaI (*PoptrGNOMa*) or PspXI and SmaI (*OrysaGNOMa*) restriction enzymes.

To generate *pGII(Bar)-pGNOM::mCherry* plasmid, the *mCherry* coding sequence was PCR amplified from an existing template and cloned into *pGII(Bar)-pGNOM:: YFPv* using XmaI and XbaI enzymes, to replace *YFPv* with *mCherry*. *MarpoGNOM* CDS was synthesised by the company Genewiz Germany GmbH (Leipzig, Germany). *The pUC-GW:MarpoGNOM* plasmid was digested with HpaI and SmaI and *MarpoGNOM* CDS was ligated into PspXI and SmaI digested *pGII(Bar)-pGNOM::mCherry* vector.

Binary vector constructs were transformed into *gnom^sgt^/GNOM, or gnl1/GNL1* mutant plants by *Agrobacterium* mediated floral dip method and T1 transgenic plants were selected using BASTA (Bayer) or phosphinothricin. Genotyping of *gnom^sgt^* and *gnl1* was performed as described earlier (Brumm et al., 2020; Richter et al., 2007).

### BFA-treatment and confocal microscopy

5-days old *Arabidopsis* seedlings were incubated with 50μM Brefeldin A (Sigma) and 2μM SynaptoRed C2 (FM4-64 equivalent; Sigma) for 1h in liquid growth media (0.5MS and 1% sucrose) and live-cell imaging was performed using Leica confocal microscope (TCS-SP8). Images adjustments (brightness and contrast only) were performed using LAS X (Leica) software and Adobe Photoshop CS3.

### CoIP and western blot analysis

CoIP was performed as described earlier (Brumm and Singh et al., 2022). Briefly, *Arabidopsis* seedlings were ground in liquid nitrogen and solubilised in ice-cold lysis buffer (50mM Tris pH 7.5, 150mM NaCl, 2mM EDTA, 1% Triton X-100) with protease inhibitors (cOmplete EDTA-free®, Roche). Total cell lysate was cleared by centrifugation at 10,000 x g for 15 min at 4^0^C and filtering the supernatant through two layers of Miracloth® (Merck Millipore). Immunoprecipitation was achieved by adding ChromoTek GFP-Trap^®^ Agarose beads (proteintech) into supernatant and incubating the reaction tube in cold room for 3h with end-to-end rotation. The beads were washed 3-4 times with ice-cold wash buffer (50mM Tris pH 7.5, 150mM NaCl, 2mM EDTA, 0.1% Triton X-100) and boiled in 2x Laemmli buffer to elute the bound protein.

After separation on SDS-PAGE, proteins were transferred to PVDF membrane (Merck Millipore) and immunodetection was performed using following primary and secondary antibodies: mouse anti-GFP (Roche), 1:1000; mouse anti-c-Myc 9E10 (Santa Cruz Biotechnology), 1:1000; POD-conjugated anti-HA (Roche), 1:1500; POD-conjugated anti-mouse (Sigma), 1: 5000. Chemiluminescence signal detection was performed using BM-chemiluminescence blotting substrate (Roche) and FusionFx7 imaging system (PeqLab).

### Yeast two-hybrid assays

Arabidopsis DCB^GNOM^ and ΔDCB^GNOM^ constructs and yeast two-hybrid method were previously described (Grebe et al., 2000; Anders et al., 2008). DCB domains of different GNOM related paralogs were synthesized by the company BaseClear B.V. (Leiden, Netherlands) and cloned into standard/modified *pEG202* and *pJG4-5* yeast vectors using EcoRI and XhoI (*BranaGNOM*, *BranaGNL1, CapruGNOMa*, *CapruGNOMb*, *PoptrGNOMa*, *PoptrGNOMb, AmhyGNOMa, OrysaGNOMa*, *OrysaGNOMb*, *AmbtrGNOM, KleniGNOM)* or XhoI (*AmhyGNOMb, AqucoGNOMa, AqucoGNOMb*) or NotI (*RiccoGNOMa*, *RiccoGNOMb*, *PisveGNOMa*, *PisveGNOMb, SalbrGNOMa, SalbrGNOMb*) enzyme sites.

### Phylogenetic analysis of plant GBF1-related sequences

All sequences (amino acids) were individually aligned using MAFFT (Katoh and Stanley, 2013) to the references AtGNOM, AtGNL1, and AtGNL2. Parts not present in any of these three reference sequences were removed from each sequence. From these sequences, a multiple sequence alignment was constructed using MAFFT, a phylogenetic tree was inferred using IQ-Tree (Nguyen et al., 2015) with a JTT-R5 model. The tree was rooted at mid-point, ancestral sequences reconstructed using TreeTime (Sagulenko et al., 2018) and visualized using Nextstrain (Hadfield et al., 2018).

### Sequence analysis

Blast searches were done at NCBI. Initially, AtGNOM was used to search for GNOM-related sequences in the genomes of other plant species using TblastN. Genomic sequences including the candidate homologs were then searched against the non-redundant protein database, limiting the search to one or two closely related species (if that genome was annotated) and to *Theobroma cacao* (Malvales) as the reference genome. If the genome of interest had not been annotated the GNOM-related gene was annotated and the conceptual translation product was aligned with *Arabidopsis thaliana* sequences using blastP to determine the percent identity to AtGNOM and AtGNL1 or AtGNL2.

Genome databases from which genome assemblies were accessed included NCBI (https://www.ncbi.nlm.nih.gov/assembly/), CNCB-NGDC (https://ngdc.cncb.ac.cn), EnsemblPlants (http://plants.ensembl.org/index.html), Plant GARDEN (Genome and Resource Database Entry; https://plantgarden.jp/en/index<u>), Phytozome v13</u> (https://phytozome-next.jgi.doe.gov/<u>); CoGe Comparative Genomics</u> (https://genomevolution.org/coge/<u>). In addition, annotated genome data files were retrieved</u> <u>from public repositories like Dryad and Figshare.</u>

### Identification and assembly of ortho-synteny groups (OSGs)

Different categories of genome assemblies were analysed to identify ortho-synteny groups (OSGs): (i) Complete genome fully annotated including tentative assignment of protein function; (ii) complete genome fully annotated but no tentative assignment of protein function (genes “uncharacterized”); (iii) possibly complete coverage of genome but fragmented (scaffolds or contigs) and with genes annotated; (iv) possibly complete coverage of genome partitioned into (virtual) chromosomes, or linkage groups, scaffolds or contigs without any further annotation. In general, 10 genes flanking GNOM paralogs on each side in annotated genome assemblies (categories i-iii) were compared with the *Theobroma cacao* (Theca) reference genome to identify standard GNOM-OSG1 (Theca chr2), GNL1-OSG2 (Theca chr4); see Suppl. Figure S8A. New OSGs due to transposition of GNOM were identified in blastP searches against the translation products of the Theca genome. OSGs in category (iv) genomes were identified by blastX searches against the translation products of the Theca reference genome. Apparent discrepancies were resolved by re-annotation of the OSG-covering segments in the genome of interest.

## SUPPLEMENTARY INFORMATION

### Suppl. Data file S1.

ARF-GEFs analysed with the Nextstrain program; (1a) species list, (1b) ARF-GEF sequences

### Suppl. Figures

S1-S2 Phylogeny of GNOM-related paralogs in gymnosperms and monocots S3A OSG1 and OSG2 marker genes

S3B GNOM transpositions mapped onto Theobroma cacao genome

S4-S35 Detailed analysis of GNOM-OSGs and GNL1-OSGs within core eudicot orders S36 GNOM-OSGs in monocots

S37 angiosperm cladogram

S38 origin of OSGs in basal eudicots

### Suppl. Tables

S1A-H Information on GBF1-type ARF-GEFs

S2 Rescue of Arabidopsis gnom by GNOM-related transgenes from other species S3 List of primers

## REFERENCES

Anders N, Jürgens G (2008). Large ARF guanine nucleotide exchange factors in membrane trafficking. Cell Mol Life Sci 65, 3433–3445. doi: 10.1007/s00018-008-8227-7.

Anders N, Nielsen M, Keicher J, Stierhof YD, Furutani M, Tasaka M, Skriver K, Jürgens G (2008). Membrane association of the Arabidopsis ARF exchange factor GNOM involves interaction of conserved domains. Plant Cell 20, 142–151. doi: 10.1105/tpc.107.056515.

Brumm S, Singh MK, Nielsen ME, Richter S, Beckmann H, Stierhof YD, Fischer AM, Kumaran M, Sundaresan V, Jürgens G (2020). Coordinated activation of ARF1 GTPases by ARF-GEF GNOM dimers is essential for vesicle trafficking in Arabidopsis. Plant Cell. 2020 Aug;32(8):2491–2507. doi: 10.1105/tpc.20.00240.

Brumm S, Singh MK, Kriechbaum C, Richter S, Huhn K, Kucera T, Baumann S, Wolters H, Takada S, Jürgens G (2022). N-terminal domain of ARF-GEF GNOM prevents heterodimerization with functionally divergent GNL1 in Arabidopsis. Plant J 112, 772–785. doi: 10.1111/tpj.15979.

Bui QT, Golinelli-Cohen MP, Jackson CL (2009). Large Arf1 guanine nucleotide exchange factors: evolution, domain structure, and roles in membrane trafficking and human disease. Mol Genet Genomics 282, 329–350. doi: 10.1007/s00438-009-0473-3.

Cai L, Xi Z, Lemmon EM, Lemmon AR, Mast A, Buddenhagen CE, Liu L, Davis CC. (2021). The Perfect Storm: Gene Tree Estimation Error, Incomplete Lineage Sorting, and Ancient Gene Flow Explain the Most Recalcitrant Ancient Angiosperm Clade, Malpighiales. Syst Biol. 70, 491–507. doi: 10.1093/sysbio/syaa083.

Casanova JE (2007) Regulation of Arf activation: the Sec7 family of guanine nucleotide exchange factors. Traffic 8, 1476–1485. doi: 10.1111/j.1600-0854.2007.00634.x.

Chanderbali AS, Jin L, Xu Q, Zhang Y, Zhang J, Jian S, Carroll E, Sankoff D, Albert VA, Howarth DG, Soltis DE, Soltis PS (2022). Buxus and Tetracentron genomes help resolve eudicot genome history. Nat Commun 13, 643. doi: 10.1038/s41467-022-28312-w.

Clarkson JJ, Zuntini AR, Maurin O, Downie SR, Plunkett GM, Nicolas AN, Smith JF, Feist MAE, et al. (2021). A higher-level nuclear phylogenomic study of the carrot family (Apiaceae). Am J Bot. 108, 1252–1269. doi: 10.1002/ajb2.1701.

Feng K, Hou XL, Xing GM, Liu JX, Duan AQ, Xu ZS, Li MY, Zhuang J, Xiong AS (2020). Advances in AP2/ERF super-family transcription factors in plant, Crit Rev Biotech 40, 750–776. doi: 10.1080/07388551.2020.1768509.

Fonseca LHM. (2021). Combining molecular and geographical data to infer the phylogeny of Lamiales and its dispersal patterns in and out of the tropics. Mol Phylogenet Evol. 164, 107287. doi: 10.1016/j.ympev.2021.107287.

Geldner N, Anders N, Wolters H, Keicher J, Kornberger W, Muller P, Delbarre A, Ueda T, Nakano A, Jürgens G (2003). The Arabidopsis GNOM ARF-GEF mediates endosomal recycling, auxin transport, and auxin-dependent plant growth. Cell 112, 219–230. doi: 10.1016/s0092-8674(03)00003-5.

Gong X, Zhang H, Guo Y, Yu S, Tang M (2025). Chromosome-level genome assembly of *Iodes seguinii* and its metabonomic implications for rheumatoid arthritis treatment. Plant Genome 18, e20534. doi: 10.1002/tpg2.20534.

Grebe M, Gadea J, Steinmann T, Kientz M, Rahfeld JU, Salchert K, Koncz C, Jürgens G (2000). A conserved domain of the arabidopsis GNOM protein mediates subunit interaction and cyclophilin 5 binding. Plant Cell 12, 343–356. doi: 10.1105/tpc.12.3.343

Guo X, Fang D, Sahu SK, Yang S, Guang X, Folk R, Smith SA, Chanderbali AS, Chen S, Liu M, Yang T, Zhang S, Liu X, Xu X, Soltis PS, Soltis DE, Liu H (2021). Chloranthus genome provides insights into the early diversification of angiosperms. Nat Commun 12, 6930. doi: 10.1038/s41467-021-26922-4.

Guo C, Luo Y, Gao LM, Yi TS, Li HT, Yang JB, Li DZ (2023). Phylogenomics and the flowering plant tree of life. J Integr Plant Biol 65, 299–323. doi: 10.1111/jipb.13415.

Hadfield J, Megill C, Bell SM, Huddleston J, Potter B, Callender C, Sagulenko P, Bedford T, Neher RA (2018). Nextstrain: real-time tracking of pathogen evolution. Bioinformatics 34, 4121–4123. doi: 10.1093/bioinformatics/bty407.

Hou C, Tian W, Kleist T, He K, Garcia V, Bai F, Hao Y, Luan S, Li L (2014). DUF221 proteins are a family of osmosensitive calcium-permeable cation channels conserved across eukaryotes. Cell Res 24, 632–635. doi: 10.1038/cr.2014.14.

Joyce EM, Appelhans MS, Buerki S, Cheek M, de Vos JM, Pirani JR, Zuntini AR, et al. (2023). Phylogenomic analyses of Sapindales support new family relationships, rapid Mid-Cretaceous Hothouse diversification, and heterogeneous histories of gene duplication. Front Plant Sci. 14, 1063174. doi: 10.3389/fpls.2023.1063174.

Katoh K and Stanley DM (2013). MAFFT multiple sequence alignment software version 7: improvements in performance and usability. Mol Biol Evol 30, 772–780. doi: 10.1093/molbev/mst010.

Kress WJ, Soltis DE, Kersey PJ, Wegrzyn JL, Leebens-Mack JH, Gostel MR, Liu X, Soltis PS (2022). Green plant genomes: What we know in an era of rapidly expanding opportunities. Proc Natl Acad Sci USA 119, e2115640118.

Lee AK, Gilman IS, Srivastav M, Lerner AD, Donoghue MJ, Clement WL. (2021). Reconstructing Dipsacales phylogeny using Angiosperms353: issues and insights. Am J Bot. 108, 1122–1142. doi: 10.1002/ajb2.1695.

Li Z, Baniaga AE, Sessa EB, Scascitelli M, Graham SW, Rieseberg LH, Barker MS (2015). Early genome duplications in conifers and other seed plants. Sci Adv 1, e1501084. doi: 10.1126/sciadv.1501084.

Li, H.T., Luo, Y., Gan, L., Ma, P.F., Gao, L.M., Yang, J.B., Cai, J., Gitzendanner, M.A., Fritsch, P.W., Zhang, T., Jin, J.J., Zeng, C.X., Wang, H., Yu, W.B., Zhang, R., van der Bank, M., Olmstead, R.G., Hollingsworth, P.M., Chase, M.W., Soltis, D.E., Soltis, P.S., Yi, T.S., and Li, D.Z. (2021a). Plastid phylogenomic insights into relationships of all flowering plant families. BMC Biol 19, 232.

Liu L, Chen M, Folk RA, Wang M, Zhao T, Shang F, Soltis DE, Li P (2023b). Phylogenomic and syntenic data demonstrate complex evolutionary processes in early radiation of the rosids. Mol Ecol Resour 23, 1673–1688. doi: 10.1111/1755-0998.13833.

Liu Y, Zhang Y, Liu Y, Lin L, Xiong X, Zhang D, Li S, Yu X, Li Y (2023a). Genome-Wide Identification and Characterization of WRKY Transcription Factors and Their Expression Profile in *Loropetalum chinense* var. *rubrum*. Plants (Basel) 12, 2131. doi: 10.3390/plants12112131.

Lv P, Wan J, Zhang C, Hina A, Al Amin GM, Begum N, Zhao T (2023). Unraveling the Diverse Roles of Neglected Genes Containing Domains of Unknown Function (DUFs): Progress and Perspective. Int J Mol Sci. 2023 Feb 20;24(4):4187. doi: 10.3390/ijms24044187.

Marks RA, Hotaling S, Frandsen PB, VanBuren R (2021). Representation and participation across 20 years of plant genome sequencing. Nat Plants 7, 1571–1578. doi: 10.1038/s41477-021-01031-8.

Maurin O, Anest A, Bellot S, Biffin E, Brewer G, Charles-Dominique T, Cowan RS, et al (2021) A nuclear phylogenomic study of the angiosperm order Myrtales, exploring the potential and limitations of the universal Angiosperms353 probe set. Am J Bot 108, 1087–1111. doi: 10.1002/ajb2.1699.

Montesinos JC, Langhans M, Sturm S, Hillmer S, Aniento F, Robinson DG, Marcote MJ (2013). Putative p24 complexes in Arabidopsis contain members of the delta and beta subfamilies and cycle in the early secretory pathway. J Exp Bot. 2013 Aug;64(11):3147–67. doi: 10.1093/jxb/ert157.

Nguyen LT, Schmidt HA, von Haeseler A, Minh BQ (2015). IQ-TREE: a fast and effective stochastic algorithm for estimating maximum-likelihood phylogenies. Mol Biol Evol 32, 268–274. doi: 10.1093/molbev/msu300.

(OTPTI) One Thousand Plant Transcriptomes Initiative (2019). One thousand plant transcriptomes and the phylogenomics of green plants. Nature 574, 679–685.

Pipaliya SV, Schlacht A, Klinger CM, Kahn RA, Dacks J (2019). Ancient complement and lineage-specific evolution of the Sec7 ARF GEF proteins in eukaryotes. Mol Biol Cell 30, 1846–1863. doi: 10.1091/mbc.E19-01-0073.

Pucker B, Irisarri I, de Vries J, Xu B (2022). Plant genome sequence assembly in the era of long reads: Progress, challenges and future directions. Quant Plant Biol 3, e5. doi: 10.1017/qpb.2021.18.

RanOmics group; Becker A, Bachelier JB, Carrive L, Conde E, Silva N, Damerval C, Del Rio C, Deveaux Y, et al. (2024). A cornucopia of diversity-Ranunculales as a model lineage. J Exp Bot 75, 1800–1822. doi: 10.1093/jxb/erad492.

Richter S, Geldner N, Schrader J, Wolters H, Stierhof YD, Rios G, Koncz C, Robinson DG, Jürgens G (2007). Functional diversification of closely related ARF-GEFs in protein secretion and recycling. Nature 448, 488–492. doi: 10.1038/nature05967.

Richter S, Müller LM, Stierhof YD, Mayer U, Takada N, Kost B, Vieten A, Geldner N, Koncz C, Jürgens G (2011). Polarized cell growth in Arabidopsis requires endosomal recycling mediated by GBF1-related ARF exchange factors. Nat Cell Biol 14, 80–86. doi: 10.1038/ncb2389.

Sagulenko P, Puller V, Neher RA (2018). TreeTime: Maximum-likelihood phylodynamic analysis. Virus Evol. 2018 Jan 8; 4, vex042. doi: 10.1093/ve/vex042.

Singh MK, Jürgens G (2018). Specificity of plant membrane trafficking - ARFs, regulators and coat proteins. Semin Cell Dev Biol 80, 85–93. doi: 10.1016/j.semcdb.2017.10.005.

Stull GW, Soltis PS, Soltis DE, Gitzendanner MA, Smith SA (2020). Nuclear phylogenomic analyses of asterids conflict with plastome trees and support novel relationships among major lineages. Am J Bot 107, 790–805. doi: 10.1002/ajb2.1468.

Stull GW, Qu XJ, Parins-Fukuchi C, Yang YY, Yang JB, Yang ZY, Hu Y, Ma H, Soltis PS, Soltis DE, Li DZ, Smith SA, Yi TS (2021). Gene duplications and phylogenomic conflict underlie major pulses of phenotypic evolution in gymnosperms. Nat Plants 7, 1015–1025. doi: 10.1038/s41477-021-00964-4.

Stull GW, Pham KK, Soltis PS, Soltis DE (2023). Deep reticulation: the long legacy of hybridization in vascular plant evolution. Plant J 114, 743–766. doi: 10.1111/tpj.16142.

Sun Y, Moore MJ, Zhang S, Soltis PS, Soltis DE, Zhao T, Meng A, Li X, Li J, Wang H (2016). Phylogenomic and structural analyses of 18 complete plastomes across nearly all families of early-diverging eudicots, including an angiosperm-wide analysis of IR gene content evolution. Mol Phylogenet Evol 96, 93–101. doi: 10.1016/j.ympev.2015.12.006.

Teh O-K, Moore I (2007). An ARF-GEF acting at the Golgi and in selective endocytosis in polarized plant cells. Nature 448, 493–496. doi: 10.1038/nature06023.

Thomas SK, Liu X, Du ZY, Dong Y, Cummings A, Pokorny L, Xiang QJ, Leebens-Mack JH. (2021). Comprehending Cornales: phylogenetic reconstruction of the order using the Angiosperms353 probe set. Am J Bot 108, 1112–1121. doi: 10.1002/ajb2.1696.

Uemura T, Ueda T, Ohniwa RL, Nakano A, Takeyasu K, Sato MH (2004). Systematic analysis of SNARE molecules in Arabidopsis: dissection of the post-Golgi network in plant cells. Cell Struct Funct 29, 49–65. doi: 10.1247/csf.29.49.

Walden N, Schranz ME (2023). Synteny Identifies Reliable Orthologs for Phylogenomics and Comparative Genomics of the Brassicaceae. Genome Biol Evol 15, evad034. doi: 10.1093/gbe/evad034.

Wang Z, Li Y, Sun P, Zhu M, Wang D, Lu Z, Hu H, Xu R, Zhang J, Ma J, Liu J, Yang Y (2022). A high-quality Buxus austro-yunnanensis (Buxales) genome provides new insights into karyotype evolution in early eudicots. BMC Biol 20, 216. doi: 10.1186/s12915-022-01420-1.

Wang Y, Ribot C, Rezzonico E, Poirier Y (2004). Structure and expression profile of the Arabidopsis PHO1 gene family indicates a broad role in inorganic phosphate homeostasis. Plant Physiol 135, 400–411. doi: 10.1104/pp.103.037945.

Wang Z, Zhou J, Pan J, Cheng W, Fang J, Lv Q, Lin X, Cheng W, Zhang L, Cheng K (2023). Insights into the Superrosids phylogeny and flavonoid synthesis from the telomere-to-telomere gap-free genome assembly of *Penthorum chinense* Pursh. Hortic Res 11, uhad274. doi: 10.1093/hr/uhad274.

Weigend M, Luebert F, Gottschling M, Couvreur TLP, Hilger HH, Miller JS. (2014). From capsules to nutlets-phylogenetic relationships in the Boraginales. Cladistics. 30, 508–518. doi: 10.1111/cla.12061.

Xie L, Gong X, Yang K, Huang Y, Zhang S, Shen L, Sun Y, Wu D, Ye C, Zhu QH, Fan L (2024). Technology-enabled great leap in deciphering plant genomes. Nat Plants 10, 551–566. doi: 10.1038/s41477-024-01655-6.

Xu K, Lin C, Lee SY, Mao L, Meng K. (2022). Comparative analysis of complete Ilex (Aquifoliaceae) chloroplast genomes: insights into evolutionary dynamics and phylogenetic relationships. BMC Genomics 23, 203. doi: 10.1186/s12864-022-08397-9.

Xue, J.Y., Dong, S.S., Wang, M.Q., Song, T.Q., Zhou, G.C., Li, Z., Van de Peer, Y., Shao, Z.Q., et al. (2022). Mitochondrial genes from 18 angiosperms fill sampling gaps for phylogenomic inferences of the early diversification of flowering plants. J Syst. Evol. 60: 773– 788. doi: 10.1111/jse.12708.

Yang L, Su D, Chang X, Foster CSP, Sun L, Huang CH, Zhou X, Zeng L, Ma H, Zhong B (2020). Phylogenomic Insights into Deep Phylogeny of Angiosperms Based on Broad Nuclear Gene Sampling. Plant Commun 1, 100027. doi: 10.1016/j.xplc.2020.100027.

Zeng L, Zhang N, Zhang Q, Endress PK, Huang J, Ma H (2017). Resolution of deep eudicot phylogeny and their temporal diversification using nuclear genes from transcriptomic and genomic datasets. New Phyt 214, 1338–1354. doi: 10.1111/nph.14503.

Zhang C, Zhang T, Luebert F, Xiang Y, Huang CH, Hu Y, Rees M, Frohlich MW, Qi J, Weigend M, Ma H (2020c). Asterid Phylogenomics/Phylotranscriptomics Uncover Morphological Evolutionary Histories and Support Phylogenetic Placement for Numerous Whole-Genome Duplications. Mol Biol Evol 37, 3188–3210. doi: 10.1093/molbev/msaa160.

Zhang L, Chen F, Zhang X, Li Z, Zhao Y, Lohaus R, Chang X, Dong W, Ho SYW, Liu X, et al. (2020a). The water lily genome and the early evolution of flowering plants. Nature 577, 79–84. doi: 10.1038/s41586-019-1852-5.

Zhang L, Wu S, Chang X, Wang X, Zhao Y, Xia Y, Trigiano RN, Jiao Y, Chen F (2020b). The ancient wave of polyploidization events in flowering plants and their facilitated adaptation to environmental stress. Plant Cell Environ 43, 2847–2856. doi: 10.1111/pce.13898.

Zhang G, Ma H (2024) Nuclear phylogenomics of angiosperms and insights into their relationships and evolution. J Integr Plant Biol 66, 546. doi: 10.1111/jipb.13609.

Zhao SW, Guo JF, Kong L, Nie S, Yan XM, Shi TL, Tian XC, Ma HY, Bao YT, Li ZC, Chen ZY, Zhang RG, Ma YP, El-Kassaby YA, Porth I, Zhao W, Mao JF. (2023). Haplotype-resolved genome assembly of Coriaria nepalensis a non-legume nitrogen-fixing shrub. Sci Data 10, 259. doi: 10.1038/s41597-023-02171-6.

Zuntini AR, Carruthers T, Maurin O, Bailey PC, Leempoel K, Brewer GE, Epitawalage N, Françoso E, Gallego-Paramo B, et al. (2024). Phylogenomics and the rise of the angiosperms. Nature 629, 843–850. doi: 10.1038/s41586-024-07324-0.

