## Supplemental Information for "Evolutionary origin and functional diversification of plant GBF1-type ARF guanine-nucleotide exchange factors"

Submitted to BIORXIV 3 June 2025

#### SUPPLEMENTAL INFORMATION

| | | page |
|---|---|---|
| **Suppl. Data file S1. Nextstrain analysis** | | **3** |
| (S1A) | Species list | 3 |
| (S1B) | ARF-GEF sequences | 25 |
| **Suppl. Figures** | | **221** |
| **S1.** | **GNOM phylogeny / gymnosperms** | **221** |
| **S2** | **GNOM phylogeny / monocots** | **222** |
| **S3A** | **Marker genes in OSG1 and OSG2** | **223** |
| **S3B** | **GNOM transpositions mapped onto Theobroma cacao genome** | **224** |
| **S4-S35** | **Cladograms and OSGs of core eudicot orders** | **225** |
| S4 | Brassicales | 225 |
| S5 | Malvales | 229 |
| S6 | Sapindales | 232 |
| S7 | Oxalidales | 237 |
| S8 | Malpighiales | 239 |
| S9 | Celastrales | 247 |
| S10 | Fagales | 249 |
| S11 | Fabales | 252 |
| S12 | Cucurbitales | 257 |
| S13 | Rosales | 261 |
| S14 | Zygophyllales | 265 |
| S15 | Myrtales | 267 |
| S16 | Crossosomatales | 270 |
| S17 | Geraniales | 272 |
| S18 | Saxifragales | 275 |
| S19 | Vitales | 280 |
| S20 | Santalales | 282 |

S21 Lamiales 285
S22 Boraginales 291
S23 Gentianales 295
S24 Solanales 298
S25 Aquifoliales 300
S26 Garryales 303
S27 Apiales 304
S28 Dipsacales 312
S29 Escalloniales 316
S30 Asterales 317
S31 Ericales 323
S32 Cornales 330
S33 Caryophyllales 334
S34 Dilleniales 339
S35 Icacinales 342
**S36 GNOM-OSGs in monocots 344**
**S37 cladogram of angiosperms 348**
**S38 Origin of OSG1 and OSG2 in basal eudicots 349**

**Suppl. Tables 353**
**S1. Information on GBF1-type plant ARF-GEFs 353**
S1A. Charophyta 353
S1B. Bryophyta, Lycophyta, ferns 354
S1C. Gymnosperms 356
S1D. Basal angiosperms, magnoliids, chloranthales, ceratophyllales 358
S1E. Monocots 363
S1F. Basal eudicots 374
S1G. Core eudicots – superasterids 380
S1H. Core eudicots – superrosids 416
**S2. Rescue of *gnom*$^{sgt}$ by *GNOM*-related paralogs 452**
**S3. List of primers 454**

| Order | Family | Species | Gene name | BFA | Class | Division |
|---|---|---|---|---|---|---|
| Chlamydomonadales | | Chlamydomonas reinhardtii | Cre07.g334100 | sens | Chlorophyceae | CHLOROPHYTA |
| Chlamydomonadales | | Volvox carteri | Vocar.0006s0037 | sens | Chlorophyceae | CHLOROPHYTA |
| Trebouxiophyceae | | Coccomyxa subellipsoidea | Cocsu_102 | res | Trebouxiophyceae | CHLOROPHYTA |
| Mamiellales | | Ostreococcus lucimarinus | Ostlu_88403 | sens | Mamiellophyceae | CHLOROPHYTA |
| Mamiellales | | Micromonas sp. RCC299 | Mic_59868 | sens | Mamiellophyceae | CHLOROPHYTA |
| Mamiellales | | Micromonas pusilla | Micpu_211597 | sens | Mamiellophyceae | CHLOROPHYTA |
| Klebsormidiales | | Klebsormidium fl. (nitens) | kfl00216_0140 | sens | Klebsormidiophyceae | CHAROPHYTA |
| Charales | | Chara braunii | Chabr_g19322 | res | Charophyceae | CHAROPHYTA |
| Marchantiales | | Marchantia polymorpha | Mapoly0100s0023 | res | Marchantiopsida | MARCHANTIOPHYTA |
| Funariales | | Physcomitrella patens | Pp3c22_6150V3 | sens | Bryopsida | BRYOPHYTA |
| Funariales | | Physcomitrella patens | Pp3c15_11320V3 | sens | Bryopsida | BRYOPHYTA |
| Funariales | | Physcomitrella patens | Pp3c9_11830V3 | sens | Bryopsida | BRYOPHYTA |
| Sphagnales | | Sphagnum fallax | Sphfalx0165s0016 | sens | Sphagnopsida | BRYOPHYTA |
| Selaginellales | | Selaginella moellendorfii | Selmo_101122 | sens | Lycopodiopsida | LYCOPODIOPHYTA |
| Saviniales | | Salvinia_cucullata | Sacu_s0267.g026961 | res | Polypodiopsida | FERNS |
| Saviniales | | Azolla filiculoides | Azfi_s0087.g042331 | sens | Polypodiopsida | FERNS |
| Saviniales | | Azolla filiculoides | Azfi_s0094.g043397 | res | Polypodiopsida | FERNS |
| Pinales | | Picea abies | comp95351 | res | Pinopsida | PINOPHYTA |
| Pinales | | Picea abies | comp77326+79708 | sens | Pinopsida | PINOPHYTA |
| Pinales | | Taxus baccata | TBA00004200+4199 (combined) | sens | Pinopsida | PINOPHYTA |
| Gnetales | | Gnetum montanum | GMO00001745+1746 (combined) | sens | Gnetopsida | GNETOPHYTA |
| Gnetales | | Gnetum montanum | GMO000012041+1433 (combined) | ??? | Gnetopsida | GNETOPHYTA |
| Amborellales | | Amborella trichopoda | AmTr_sc02.551 | sens | (basal angiosperms) | ANGIOSPERMS |
| Amborellales | | Amborella trichopoda | AmTr_sc17.131+132+133+129 | sens | (basal angiosperms) | ANGIOSPERMS |
| Alismatales | | Spirodela polyrhiza | Spipo3G0098500 | sens | Monocots | ANGIOSPERMS |
| Alismatales | | Spirodela polyrhiza | Spipo21G0014600+14700 | sens | Monocots | ANGIOSPERMS |

| | | | | | | |
|---|---|---|---|---|---|---|
| Alismatales | | Zostera marina | Zosma108g00240 | sens | Monocots | ANGIOSPERMS |
| Alismatales | | Zostera marina | Zosma56g00900 | sens | Monocots | ANGIOSPERMS |
| Alismatales | | Zostera marina | Zosma123g00350.1 | sens | Monocots | ANGIOSPERMS |
| Alismatales | | Zostera marina | Zosma111g00160+00150 | sens | Monocots | ANGIOSPERMS |
| Dioscoreales | | Dioscorea rotundata | Dioro_chr8_5252329 | sens | Monocots | ANGIOSPERMS |
| Dioscoreales | | | Dioro_chr4_5930135 | sens | Monocots | ANGIOSPERMS |
| Asparagales | | Asparagus officinalis | Aspof_LOC109850386 | sens | Monocots | ANGIOSPERMS |
| Asparagales | | Asparagus officinalis | Aspof_LOC109851574 | sens | Monocots | ANGIOSPERMS |
| Arecales | | Phoenix dactylifera | Phoda_LOC103712126 | res | Monocots | ANGIOSPERMS |
| Arecales | | Phoenix dactylifera | Phoda_LOC103696334 | sens | Monocots | ANGIOSPERMS |
| Arecales | | Phoenix dactylifera | Phoda_LOC103695708 | sens, res | Monocots | ANGIOSPERMS |
| Zingiberales | | Musa acuminata | Musac_LOC103988820 | sens | Monocots | ANGIOSPERMS |
| Zingiberales | | Musa acuminata | Musac_LOC103986838 | sens | Monocots | ANGIOSPERMS |
| Zingiberales | | Musa acuminata | Musac_LOC103990130 | sens | Monocots | ANGIOSPERMS |
| Poales | | Oryza sativa | LOC_Os03g46330 | sens | Monocots | ANGIOSPERMS |
| Poales | | Oryza sativa | LOC_Os02g22090 | sens | Monocots | ANGIOSPERMS |
| Poales | | Oryza sativa | LOC_Os04g02690 | sens | Monocots | ANGIOSPERMS |
| Poales | | Sorghum bicolor | Sobic.001G143600 | sens | Monocots | ANGIOSPERMS |
| Poales | | Sorghum bicolor | Sobic.001G301200 | sens | Monocots | ANGIOSPERMS |
| Poales | | Sorghum bicolor | Sobic.008G100600 | sens | Monocots | ANGIOSPERMS |
| Poales | | Brachypodium distachyon | Bradi1g12920 | sens | Monocots | ANGIOSPERMS |
| Poales | | Brachypodium distachyon | Bradi1g74210 | sens | Monocots | ANGIOSPERMS |
| Poales | | Brachypodium distachyon | Bradi2g48310 | sens | Monocots | ANGIOSPERMS |
| Poales | | Oropetium thomaeum | Oropetium_20150105_08249A | sens | Monocots | ANGIOSPERMS |
| Poales | | Oropetium thomaeum | Oropetium_20150105_25567A | sens | Monocots | ANGIOSPERMS |
| Poales | | Ananas comosus | Aco005192 | sens | Monocots | ANGIOSPERMS |
| Poales | | Ananas comosus | Aco000228 | res | Monocots | ANGIOSPERMS |
| Poales | | Ananas comosus | Aco005934 | sens | Monocots | ANGIOSPERMS |

| | | | | | | |
|---|---|---|---|---|---|---|
| Laurales | | Cinnamomum micranthum | CKAN_00834500 | res | Magnoliids | ANGIOSPERMS |
| Laurales | | Cinnamomum micranthum | CKAN_01177000 | sens | Magnoliids | ANGIOSPERMS |
| Ceratophyl-lales | | Ceratophyllum demersum | Cerde_20156 | sens | Non-eudicot (sister to eudicots) | ANGIOSPERMS |
| Ceratophyl-lales | | Ceratophyllum demersum | Cerde_25991 | sens | Non-eudicot (sister to eudicots) | ANGIOSPERMS |
| Ceratophyl-lales | | Ceratophyllum demersum | Cerde_10541 | sens | Non-eudicot (sister to eudicots) | ANGIOSPERMS |
| Ranunculales | Ranunculaceae | Aquilegia coerulea | Aquca_014_00263 | sens | (basal eudicots) | ANGIOSPERMS |
| Ranunculales | Ranunculaceae | Aquilegia coerulea | Aquca_045_00153 | res | (basal eudicots) | ANGIOSPERMS |
| Ranunculales | Ranunculaceae | Aquilegia coerulea | Aquca_007_00074 | sens | (basal eudicots) | ANGIOSPERMS |
| Ranunculales | Ranunculaceae | Aquilegia coerulea | Aquca_020_00178 | sens | (basal eudicots) | ANGIOSPERMS |
| Ranunculales | Papaveraceae | Eschscholzia californica | Eca_sc010404_g0150 | sens | (basal eudicots) | ANGIOSPERMS |
| Ranunculales | Papaveraceae | Eschscholzia californica | Eca_sc000188_g0230 | res | (basal eudicots) | ANGIOSPERMS |
| Ranunculales | Papaveraceae | Eschscholzia californica | Eca_sc001331_g0490 | sens | (basal eudicots) | ANGIOSPERMS |
| Ranunculales | Papaveraceae | Papaver somniferum | Papso_LOC113307666 | sens | (basal eudicots) | ANGIOSPERMS |
| Ranunculales | Papaveraceae | Papaver somniferum | Papso_LOC113297613 | res | (basal eudicots) | ANGIOSPERMS |
| Ranunculales | Papaveraceae | Papaver somniferum | Papso_LOC113316139 | sens | (basal eudicots) | ANGIOSPERMS |
| Ranunculales | Papaveraceae | Macleaya cordata | Macco_1831g8 | sens | (basal eudicots) | ANGIOSPERMS |
| Ranunculales | Papaveraceae | Macleaya cordata | Macco_9095g21 | res | (basal eudicots) | ANGIOSPERMS |
| Ranunculales | Papaveraceae | Macleaya cordata | Macco_1789g18 | sens | (basal eudicots) | ANGIOSPERMS |
| Ranunculales | Circaeasteraceae | Kingdonia uniflora | Kinun_030821 | sens | (basal eudicots) | ANGIOSPERMS |
| Ranunculales | Circaeasteraceae | Kingdonia uniflora | Kinun_019872 | res | (basal eudicots) | ANGIOSPERMS |
| Ranunculales | Circaeasteraceae | Kingdonia uniflora | Kinun_020572 | sens | (basal eudicots) | ANGIOSPERMS |
| Ranunculales | Lardizabalaceae | Akebia trifoliata | AketrGNOMa_chr14 | sens | (basal eudicots) | ANGIOSPERMS |
| Ranunculales | Lardizabalaceae | Akebia trifoliata | AketrGNOMb_chr11 | res | (basal eudicots) | ANGIOSPERMS |
| Ranunculales | Lardizabalaceae | Akebia trifoliata | AketrGNL2_chr10 | sens | (basal eudicots) | ANGIOSPERMS |

| | | | | | | |
|---|---|---|---|---|---|---|
| Proteales | Proteaceae | Macadamia integrifolia | Mint05g10540 | sens | (basal eudicots) | ANGIOSPERMS |
| Proteales | Proteaceae | Macadamia integrifolia | MintSC289g00040 | sens | (basal eudicots) | ANGIOSPERMS |
| Proteales | Proteaceae | Macadamia integrifolia | Mint05g17010 | sens | (basal eudicots) | ANGIOSPERMS |
| Proteales | Nelumbonaceae | Nelumbo nucifera | Nelnu_LOC104605370 | sens | (basal eudicots) | ANGIOSPERMS |
| Proteales | Nelumbonaceae | Nelumbo nucifera | Nelnu_LOC104612314 | sens | (basal eudicots) | ANGIOSPERMS |
| Proteales | Nelumbonaceae | Nelumbo nucifera | Nelnu_LOC104585650 | sens | (basal eudicots) | ANGIOSPERMS |
| Trochondendrales | Trochodendraceae | Trochodendronaralioides | Troal_0.675 | sens | (basal eudicots) | ANGIOSPERMS |
| Trochondendrales | Trochodendraceae | Trochodendronaralioides | Troal_13.1500 | res | (basal eudicots) | ANGIOSPERMS |
| Trochondendrales | Trochodendraceae | Trochodendronaralioides | Troal_12.117 | sens | (basal eudicots) | ANGIOSPERMS |
| Trochondendrales | Trochodendraceae | Trochodendronaralioides | Troal_17.673 | sens | (basal eudicots) | ANGIOSPERMS |
| Trochondendrales | Trochodendraceae | Tetracentron sinense | Tetsi_007283 | sens | (basal eudicots) | ANGIOSPERMS |
| Trochondendrales | Trochodendraceae | Tetracentron sinense | Tetsi_000870 | res | (basal eudicots) | ANGIOSPERMS |
| Trochondendrales | Trochodendraceae | Tetracentron sinense | Tetsi_015395 | sens | (basal eudicots) | ANGIOSPERMS |
| Trochondendrales | Trochodendraceae | Tetracentron sinense | Tetsi_012649 | sens | (basal eudicots) | ANGIOSPERMS |
| Caryophyllales | Amaranthaceae | Amaranthus cruentus | Amacr03g15130 | sens | (core eudicots) | ANGIOSPERMS |
| Caryophyllales | Amaranthaceae | Amaranthus cruentus | Amacr12g11020 | res | (core eudicots) | ANGIOSPERMS |
| Caryophyllales | Amaranthaceae | Amaranthus cruentus | Amacr14g03290 | sens | (core eudicots) | ANGIOSPERMS |
| caryophyllales | Amaranthaceae | Spinacia oleracea | Spiol_LOC110785052 | sens | (core eudicots) | ANGIOSPERMS |
| Caryophyllales | Amaranthaceae | Spinacia oleracea | Spiol_LOC110804592 | sens | (core eudicots) | ANGIOSPERMS |
| Caryophyllales | Amaranthaceae | Spinacia oleracea | Spiol_LOC110776164 | res | (core eudicots) | ANGIOSPERMS |
| Caryophyllales | Amaranthaceae | Spinacia oleracea | Spiol_LOC110777245 | sens | (core eudicots) | ANGIOSPERMS |
| Caryophyllales | Amaranthaceae | Beta vulgaris | Betvu_LOC104897851 | sens | (core eudicots) | ANGIOSPERMS |
| Caryophyllales | Amaranthaceae | Beta vulgaris | Betvu_LOC104906035 | res | (core eudicots) | ANGIOSPERMS |
| Caryophyllales | Amaranthaceae | Beta vulgaris | Betvu_LOC104902176 | sens | (core eudicots) | ANGIOSPERMS |
| Caryophyllales | Amaranthaceae | Suaeda aralocaspica | Suaar_00002098 | sens | (core eudicots) | ANGIOSPERMS |

| | | | | | | |
|---|---|---|---|---|---|---|
| Caryophyllales | Amaranthaceae | Suaeda aralocaspica | Suaar_00025615 | res | (core eudicots) | ANGIOSPERMS |
| Caryophyllales | Amaranthaceae | Suaeda aralocaspica | Suaar_00007839 | sens | (core eudicots) | ANGIOSPERMS |
| Caryophyllales | Droseraceae | Drosera spatulata | Drosp_00009154 | sens | (core eudicots) | ANGIOSPERMS |
| Caryophyllales | Droseraceae | Drosera spatulata | Drosp_00004000 | sens | (core eudicots) | ANGIOSPERMS |
| Caryophyllales | Droseraceae | Drosera spatulata | Drosp_00017218 | sens | (core eudicots) | ANGIOSPERMS |
| Caryophyllales | Droseraceae | Drosera spatulata | Drosp_00008616 | sens | (core eudicots) | ANGIOSPERMS |
| Caryophyllales | Portulacaceae | Portulaca amilis | Poram_023444 | sens | (core eudicots) | ANGIOSPERMS |
| Caryophyllales | Portulacaceae | Portulaca amilis | Poram_037122 | res | (core eudicots) | ANGIOSPERMS |
| Caryophyllales | Portulacaceae | Portulaca amilis | Poram_026381 | sens | (core eudicots) | ANGIOSPERMS |
| Caryophyllales | Simondsiaceae | Simmondsia chinensis | Simch_030709 | sens | (core eudicots) | ANGIOSPERMS |
| Caryophyllales | Simondsiaceae | Simmondsia chinensis | Simch_029603 | res | (core eudicots) | ANGIOSPERMS |
| Caryophyllales | Simondsiaceae | Simmondsia chinensis | Simch_015694 | sens | (core eudicots) | ANGIOSPERMS |
| Caryophyllales | Polygonaceae | Fagopyrum tartaricum | FagtaGNOMa_chr5-18.6 | sens | (core eudicots) | ANGIOSPERMS |
| Caryophyllales | Polygonaceae | Fagopyrum tartaricum | FagtaGNOMb_chr5-43.3 | sens | (core eudicots) | ANGIOSPERMS |
| Caryophyllales | Polygonaceae | Fagopyrum tartaricum | FagtaGNOMc_chr6-36.2 | res | (core eudicots) | ANGIOSPERMS |
| Caryophyllales | Polygonaceae | Fagopyrum tartaricum | FagtaGNL2a_chr5-51.6 | sens | (core eudicots) | ANGIOSPERMS |
| Caryophyllales | Polygonaceae | Fagopyrum tartaricum | FagtaGNL2b_chr6-47.9 | sens | (core eudicots) | ANGIOSPERMS |
| Caryophyllales | Caryophyllaceae | Dianthus caryophyllus | Diaca_4337 | sens | (core eudicots) | ANGIOSPERMS |
| Caryophyllales | Caryophyllaceae | Dianthus caryophyllus | DiacaGNL1_sc4 | sens | (core eudicots) | ANGIOSPERMS |
| Caryophyllales | Caryophyllaceae | Dianthus caryophyllus | Diaca_14696 | sens | (core eudicots) | ANGIOSPERMS |
| Caryophyllales | Cactaceae | Carnegiea gigantea | CargiGNOMa_sc8504 | sens | (core eudicots) | ANGIOSPERMS |
| Caryophyllales | Cactaceae | Carnegiea gigantea | CargiGNOMb_sc1597 | sens | (core eudicots) | ANGIOSPERMS |
| Caryophyllales | Cactaceae | Carnegiea gigantea | CargiGNL2_sc1566 | sens | (core eudicots) | ANGIOSPERMS |
| Cornales | Nyssaceae | Camptotheca acuminata | Cac_g011358 | sens | (basal asterids) | ANGIOSPERMS |
| Cornales | Nyssaceae | Camptotheca acuminata | Cac_g006307 | res | (basal asterids) | ANGIOSPERMS |
| Cornales | Nyssaceae | Camptotheca acuminata | Cac_g011770 | sens | (basal asterids) | ANGIOSPERMS |
| Cornales | Nyssaceae | Nyssa sinensis | Nyssi_012948 | sens | (basal asterids) | ANGIOSPERMS |

| | | | | | | |
|---|---|---|---|---|---|---|
| Cornales | Nyssaceae | Nyssa sinensis | Nyssi_012781 | sens | (basal asterids) | ANGIOSPERMS |
| Cornales | Nyssaceae | Nyssa sinensis | Nyssi_000634 | res | (basal asterids) | ANGIOSPERMS |
| Cornales | Nyssaceae | Nyssa sinensis | NyssiGNL2 | sens | (basal asterids) | ANGIOSPERMS |
| Cornales | Hydrangeaceae | Hydrangea quercifolia | Hyque_10G111800 | sens | (basal asterids) | ANGIOSPERMS |
| Cornales | Hydrangeaceae | Hydrangea quercifolia | Hyque_13G093400 | res | (basal asterids) | ANGIOSPERMS |
| Cornales | Hydrangeaceae | Hydrangea quercifolia | Hyque_18G092600 | sens | (basal asterids) | ANGIOSPERMS |
| Ericales | Actinidiaceae | Actinidia chinensis | CEY00_Acc06361 | sens | (basal asterids) | ANGIOSPERMS |
| Ericales | Actinidiaceae | Actinidia chinensis | CEY00_Acc31499 | sens | (basal asterids) | ANGIOSPERMS |
| Ericales | Actinidiaceae | Actinidia chinensis | CEY00_Acc02326 | res | (basal asterids) | ANGIOSPERMS |
| Ericales | Actinidiaceae | Actinidia chinensis | CEY00_Acc09920 | sens | (basal asterids) | ANGIOSPERMS |
| Ericales | Theaceae | Camellia sinensis | Camsi_LOC114306371 | sens | (basal asterids) | ANGIOSPERMS |
| Ericales | Theaceae | Camellia sinensis | Camsi_LOC114271324 | res | (basal asterids) | ANGIOSPERMS |
| Ericales | Theaceae | Camellia sinensis | Camsi_LOC114282320 | sens | (basal asterids) | ANGIOSPERMS |
| Ericales | Ericaceae | Rhododendron simsii | Rhosi_10G0094900 | sens | (basal asterids) | ANGIOSPERMS |
| Ericales | Ericaceae | Rhododendron simsii | Rhosi_08G0072200 | res | (basal asterids) | ANGIOSPERMS |
| Ericales | Ericaceae | Rhododendron simsii | Rhosi_03G0244100 | sens | (basal asterids) | ANGIOSPERMS |
| Ericales | Ebenaceae | Diospyros oleifera | Diool_1472.179 | sens | (basal asterids) | ANGIOSPERMS |
| Ericales | Ebenaceae | Diospyros oleifera | Diool_124.634 | res | (basal asterids) | ANGIOSPERMS |
| Ericales | Ebenaceae | Diospyros oleifera | Diool_968.73 | sens | (basal asterids) | ANGIOSPERMS |
| Ericales | Ebenaceae | Diospyros oleifera | Diool_92.671 | sens | (basal asterids) | ANGIOSPERMS |
| Ericales | Roridulaceae | Roridula gorgonias | Rorgo_13293 | sens | (basal asterids) | ANGIOSPERMS |
| Ericales | Roridulaceae | Roridula gorgonias | Rorgo_03512 | sens | (basal asterids) | ANGIOSPERMS |
| Ericales | Roridulaceae | Roridula gorgonias | Rorgo_08858 | sens | (basal asterids) | ANGIOSPERMS |
| Ericales | Clethraceae | Clethra arborea | Clear_17208 | sens | (basal asterids) | ANGIOSPERMS |
| Ericales | Clethraceae | Clethra arborea | Clear_18756 | res | (basal asterids) | ANGIOSPERMS |
| Ericales | Clethraceae | Clethra arborea | Clear_06455 | sens | (basal asterids) | ANGIOSPERMS |

| | | | | | | |
|---|---|---|---|---|---|---|
| Ericales | Primulaceae | Aegiceras corniculatum | Aegco_05G 0197980 | sens | (basal asterids) | ANGIOSPERM S |
| Ericales | Primulaceae | Aegiceras corniculatum | Aegco_14G 0264380 | sens | (basal asterids) | ANGIOSPERM S |
| Ericales | Primulaceae | Aegiceras corniculatum | Aegco_02G 0321210 | sens | (basal asterids) | ANGIOSPERM S |
| Ericales | Primulaceae | Primula veris | Prive_BG88 09307 | sens | (basal asterids) | ANGIOSPERM S |
| Ericales | Primulaceae | Primula veris | Prive_BG88 16056 | sens | (basal asterids) | ANGIOSPERM S |
| Asterales | Campanula-ceae | Platycodon grandiflorus | PlagrGNOM _chr7 | sens | (campanulid asterids) | ANGIOSPERM S |
| Asterales | Campanula-ceae | Platycodon grandiflorus | PlagrGNL1_ chr1 | res | (campanulid asterids) | ANGIOSPERM S |
| Asterales | Campanula-ceae | Platycodon grandiflorus | PlagrGNL2_ chr9 | sens | (campanulid asterids) | ANGIOSPERM S |
| Asterales | Asteraceae | Cynara cardunculus | Cynca_LOC 112507087 | sens | (campanulid asterids) | ANGIOSPERM S |
| Asterales | Asteraceae | Cynara cardunculus | Cynca_LOC 112512756 | sens | (campanulid asterids) | ANGIOSPERM S |
| Asterales | Asteraceae | Cynara cardunculus | Cynca_LOC 112516397 | sens | (campanulid asterids) | ANGIOSPERM S |
| Asterales | Asteraceae | Lactuca sativa | Lacsa_LOC 111878063 | sens | (campanulid asterids) | ANGIOSPERM S |
| Asterales | Asteraceae | Lactuca sativa | Lacsa_LOC 111879745 | sens | (campanulid asterids) | ANGIOSPERM S |
| Asterales | Asteraceae | Lactuca sativa | Lacsa_ LOC111920 050 | res | (campanulid asterids) | ANGIOSPERM S |
| Asterales | Asteraceae | Lactuca sativa | Lacsa_LOC 111897921 | sens | (campanulid asterids) | ANGIOSPERM S |
| Apiales | | Daucus carota | Dauca_LOC 108218573 | sens | (campanulid asterids) | ANGIOSPERM S |
| Apiales | | Daucus carota | Dauca_LOC 108218683 | sens | (campanulid asterids) | ANGIOSPERM S |
| Apiales | | Daucus carota | Dauca_LOC 108214229 | res | (campanulid asterids) | ANGIOSPERM S |
| Apiales | | Daucus carota | Dauca_LOC 108212563 | res | (campanulid asterids) | ANGIOSPERM S |
| Apiales | | Daucus carota | Dauca_LOC 108199672 | sens | (campanulid asterids) | ANGIOSPERM S |
| Apiales | | Centella asiatica | Cenas_5.10 79 | sens | campanulid asterids) | ANGIOSPERM S |
| Apiales | | Centella asiatica | Cenas_1.46 25 | sens | campanulid asterids) | ANGIOSPERM S |
| Dipsacales | | Lonicera japonica | Lonja_GWH PAAZE0055 99 | sens | (campanulid asterids) | ANGIOSPERM S |
| Dipsacales | | Lonicera japonica | Lonja_GWH PAAZE0284 49 | sens | (campanulid asterids) | ANGIOSPERM S |
| Dipsacales | | Lonicera japonica | Lonja_GWH PAAZE0117 10 | res | (campanulid asterids) | ANGIOSPERM S |

| | | | | | | |
|---|---|---|---|---|---|---|
| Dipsacales | | Lonicera japonica | Lonja_GWHPAAZE011712 | res | (campanulid asterids) | ANGIOSPERMS |
| Dipsacales | | Lonicera japonica | Lonja_GWHPAAZE011900 | res | (campanulid asterids) | ANGIOSPERMS |
| Dipsacales | | Lonicera japonica | Lonja_GWHPAAZE01190 | res | (campanulid asterids) | ANGIOSPERMS |
| Dipsacales | | Lonicera japonica | Lonja_GWHPAAZE025641 | sens | (campanulid asterids) | ANGIOSPERMS |
| Gentianales | Apocynaceae | Catharanthus roseus | CRO_018704 | sens | (lamiid asterids) | ANGIOSPERMS |
| Gentianales | Apocynaceae | Catharanthus roseus | CRO_021518 | sens | (lamiid asterids) | ANGIOSPERMS |
| Gentianales | Apocynaceae | Catharanthus roseus | CRO_013169 | sens | (lamiid asterids) | ANGIOSPERMS |
| Gentianales | Apocynaceae | Calotropis gigantea | Calgi_g003986 | sens | (lamiid asterids) | ANGIOSPERMS |
| Gentianales | Apocynaceae | Calotropis gigantea | Calgi_g002145 | sens | (lamiid asterids) | ANGIOSPERMS |
| Gentianales | Apocynaceae | Calotropis gigantea | Calgi_g000739 | sens | (lamiid asterids) | ANGIOSPERMS |
| Gentianales | Rubiaceae | Coffea arabica | Cofar_LOC113716666 | sens | (lamiid asterids) | ANGIOSPERMS |
| Gentianales | Rubiaceae | Coffea arabica | Cofar_LOC113718983 | sens | (lamiid asterids) | ANGIOSPERMS |
| Gentianales | Rubiaceae | Coffea arabica | Cofar_LOC113692407 | sens | (lamiid asterids) | ANGIOSPERMS |
| Gentianales | Rubiaceae | Coffea arabica | Cofar_LOC113695788 | sens | (lamiid asterids) | ANGIOSPERMS |
| Gentianales | Rubiaceae | Coffea canephora | Cofca_T00032293001 | sens | (lamiid asterids) | ANGIOSPERMS |
| Gentianales | Rubiaceae | Coffea canephora | Cofca_T00039609001 | res(?) | (lamiid asterids) | ANGIOSPERMS |
| Gentianales | Rubiaceae | Coffea canephora | T00041903001 | sens | (lamiid asterids) | ANGIOSPERMS |
| Gentianales | Rubiaceae | Chiococca alba | Chial_g3955 | sens | (lamiid asterids) | ANGIOSPERMS |
| Gentianales | Rubiaceae | Chiococca alba | Chial_g16800 | res | (lamiid asterids) | ANGIOSPERMS |
| Gentianales | Rubiaceae | Chiococca alba | Chial_g8037 | sens | (lamiid asterids) | ANGIOSPERMS |
| Gentianales | Gelsemiaceae | Gelsemium sempervirens | GelseGNOM_sc8 | sens | (lamiid asterids) | ANGIOSPERMS |
| Gentianales | Gelsemiaceae | Gelsemium sempervirens | GelseGNL1_sc0 | res | (lamiid asterids) | ANGIOSPERMS |
| Gentianales | Gelsemiaceae | Gelsemium sempervirens | GelseGNL2_sc147 | sens | (lamiid asterids) | ANGIOSPERMS |

| | | | | | | |
|---|---|---|---|---|---|---|
| Lamiales | Lentibulariaceae | Genlisea aurea | Genau_EPS60000 | sens | (lamiid asterids) | ANGIOSPERMS |
| Lamiales | Lentibulariaceae | Genlisea aurea | Genau_EPS65079 | sens | (lamiid asterids) | ANGIOSPERMS |
| Lamiales | Lentibulariaceae | Genlisea aurea | Genau_EPS71757 | sens | (lamiid asterids) | ANGIOSPERMS |
| Lamiales | Lentibulariaceae | Genlisea aurea | Genau_EPS59939 | sens | (lamiid asterids) | ANGIOSPERMS |
| Lamiales | Lentibulariaceae | Utricularia gibba | UtrgiGNOM_ctg749 | sens | (lamiid asterids) | ANGIOSPERMS |
| Lamiales | Lentibulariaceae | Utricularia gibba | UtrgiGNL1_ctg 736 | sens | (lamiid asterids) | ANGIOSPERMS |
| Lamiales | Lentibulariaceae | Utricularia gibba | UtrgiGNL2_chr1 | sens | (lamiid asterids) | ANGIOSPERMS |
| Lamiales | Pedaliaceae | Sesamum indicum | Sesin_LOC105159998 | sens | (lamiid asterids) | ANGIOSPERMS |
| Lamiales | Pedaliaceae | Sesamum indicum | Sesin_LOC105177279 | res | (lamiid asterids) | ANGIOSPERMS |
| Lamiales | Pedaliaceae | Sesamum indicum | Sesin_LOC105158259 | sens | (lamiid asterids) | ANGIOSPERMS |
| Lamiales | Phrymaceae | Erythranthe guttatus | Erygu_LOC105967563 | sens | (lamiid asterids) | ANGIOSPERMS |
| Lamiales | Phrymaceae | Erythranthe guttatus | Erygu_LOC105970904 | res | (lamiid asterids) | ANGIOSPERMS |
| Lamiales | Phrymaceae | Erythranthe guttatus | Erygu_LOC105976035 | sens | (lamiid asterids) | ANGIOSPERMS |
| Lamiales | Orobanchaceae | Lindenbergia philippensis | Liphi_05G047100 | sens | (lamiid asterids) | ANGIOSPERMS |
| Lamiales | Orobanchaceae | Lindenbergia philippensis | Liphi_09G029200 | sens | (lamiid asterids) | ANGIOSPERMS |
| Lamiales | Orobanchaceae | Lindenbergia philippensis | Liphi_08G097200 | sens | (lamiid asterids) | ANGIOSPERMS |
| Lamiales | Lamiaceae | Callicarpa americana | Calam_14G038300 | sens | (lamiid asterids) | ANGIOSPERMS |
| Lamiales | Lamiaceae | Callicarpa americana | Calam_09G131100 | res | (lamiid asterids) | ANGIOSPERMS |
| Lamiales | Lamiaceae | Callicarpa americana | Calam_12G060000 | sens | (lamiid asterids) | ANGIOSPERMS |
| Lamiales | Oleaceae | Fraxinus excelsior | FraexGNOMa_chr7 | res | (lamiid asterids) | ANGIOSPERMS |
| Lamiales | Oleaceae | Fraxinus excelsior | FraexGNOMb_chr15 | res | (lamiid asterids) | ANGIOSPERMS |
| Lamiales | Oleaceae | Fraxinus excelsior | FraexGNOMc_chr8 | sens | (lamiid asterids) | ANGIOSPERMS |
| Lamiales | Oleaceae | Fraxinus excelsior | FraexGNL2_chr1 | sens | (lamiid asterids) | ANGIOSPERMS |
| Lamiales | Acanthaceae | Andrographis paniculata | AndpaGNOM_chr19 | sens | (lamiid asterids) | ANGIOSPERMS |
| Lamiales | Acanthaceae | Andrographis paniculata | AndpaGNL1_chr2 | sens | (lamiid asterids) | ANGIOSPERMS |

| | | | | | | |
|---|---|---|---|---|---|---|
| Lamiales | Acanthaceae | Andrographis paniculata | AndpaGNL2_chr10 | sens | (lamiid asterids) | ANGIOSPERMS |
| Lamiales | Plantaginaceae | Antirrhinum majus | Am03g32310 | sens | (lamiid asterids) | ANGIOSPERMS |
| Lamiales | Plantaginaceae | Antirrhinum majus | Am06g34280 | sens | (lamiid asterids) | ANGIOSPERMS |
| Lamiales | Plantaginaceae | Antirrhinum majus | Am06g18620 | sens | (lamiid asterids) | ANGIOSPERMS |
| Lamiales | Bignoniaceae | Handroanthus impetiginosus | Hanim_22191 | sens | (lamiid asterids) | ANGIOSPERMS |
| Lamiales | Bignoniaceae | Handroanthus impetiginosus | Hanim_24118 | res | (lamiid asterids) | ANGIOSPERMS |
| Lamiales | Bignoniaceae | Handroanthus impetiginosus | Hanim_17220 | sens | (lamiid asterids) | ANGIOSPERMS |
| Lamiales | Gesneriaceae | Primulina huaijiensis | PrihuGNOMa_chr14 | sens | (lamiid asterids) | ANGIOSPERMS |
| Lamiales | Gesneriaceae | Primulina huaijiensis | PrihuGNOMb_chr3-29 | sens | (lamiid asterids) | ANGIOSPERMS |
| Lamiales | Gesneriaceae | Primulina huaijiensis | PrihuGNOMc_chr3-28 | sens | (lamiid asterids) | ANGIOSPERMS |
| Lamiales | Gesneriaceae | Primulina huaijiensis | PrihuGNOMd_sc40239 | sens | (lamiid asterids) | ANGIOSPERMS |
| Lamiales | Gesneriaceae | Primulina huaijiensis | PrihuGNL2_chr3-24 | sens | (lamiid asterids) | ANGIOSPERMS |
| Solanales | Solanaceae | Solanum lycopersicum | Solyc05g013040 | sens | (lamiid asterids) | ANGIOSPERMS |
| Solanales | Solanaceae | Solanum lycopersicum | Solyc04g005560 | sens | (lamiid asterids) | ANGIOSPERMS |
| Solanales | Solanaceae | Solanum lycopersicum | Solyc02g067010 | sens | (lamiid asterids) | ANGIOSPERMS |
| Solanales | Solanaceae | Solanum lycopersicum | Solyc02g088290 | sens | (lamiid asterids) | ANGIOSPERMS |
| Solanales | Solanaceae | Solanum lycopersicum | Solyc03g070470 | sens | (lamiid asterids) | ANGIOSPERMS |
| Solanales | Convolvulaceae | Ipomoea nil | Iponi_LOC109166281 | sens | (lamiid asterids) | ANGIOSPERMS |
| Solanales | Convolvulaceae | Ipomoea nil | Iponi_LOC109153697 | res | (lamiid asterids) | ANGIOSPERMS |
| Solanales | Convolvulaceae | Ipomoea nil | Iponi_LOC109180388 | sens | (lamiid asterids) | ANGIOSPERMS |
| Solanales | Convolvulaceae | Ipomoea nil | Iponi_LOC109147747 | sens | (lamiid asterids) | ANGIOSPERMS |

| | | | | | | |
|---|---|---|---|---|---|---|
| Solanales | Convolvulaceae | Cuscuta australis | Cusau_000320 | sens | (lamiid asterids) | ANGIOSPERMS |
| Solanales | Convolvulaceae | Cuscuta australis | Cusau_005101 | sens | (lamiid asterids) | ANGIOSPERMS |
| Solanales | Convolvulaceae | Cuscuta australis | Cusau_001637 | sens | (lamiid asterids) | ANGIOSPERMS |
| Garryales | | Eucommia ulmoides | Eucul_013014 | sens | (lamiid asterids) | ANGIOSPERMS |
| Garryales | | Eucommia ulmoides | Eucul_026230 | sens | (lamiid asterids) | ANGIOSPERMS |
| Boraginales | | Echium plantagineum | EchplGNOMa_LG8 | sens | (lamiid asterids) | ANGIOSPERMS |
| Boraginales | | Echium plantagineum | EchplGNOMb_LG3-261 | sens | (lamiid asterids) | ANGIOSPERMS |
| Boraginales | | Echium plantagineum | EchplGNOMc_LG3-262 | sens | (lamiid asterids) | ANGIOSPERMS |
| Boraginales | | Echium plantagineum | EchplGNL2_LG4 | sens | (lamiid asterids) | ANGIOSPERMS |
| Saxifragales | Crassulaceae | Kalanchoe fedtschenkoi | Kalfe_61s0170 | sens | (core eudicots) | ANGIOSPERMS |
| Saxifragales | Crassulaceae | Kalanchoe fedtschenkoi | Kalfe_43s0155 | sens | (core eudicots) | ANGIOSPERMS |
| Saxifragales | Crassulaceae | Kalanchoe fedtschenkoi | Kalfe_1222s0075 | sens | (core eudicots) | ANGIOSPERMS |
| Saxifragales | Crassulaceae | Kalanchoe fedtschenkoi | Kalfe_96s0081 | res | (core eudicots) | ANGIOSPERMS |
| Saxifragales | Crassulaceae | Kalanchoe fedtschenkoi | Kalfe_31s0117 | sens | (core eudicots) | ANGIOSPERMS |
| Saxifragales | Crassulaceae | Sedum album | SedalGNOM_ctg13 | sens | (core eudicots) | ANGIOSPERMS |
| Saxifragales | Crassulaceae | Sedum album | SedalGNL1a_sc317 | sens | (core eudicots) | ANGIOSPERMS |
| Saxifragales | Crassulaceae | Sedum album | SedalGNL1b_sc25 | res | (core eudicots) | ANGIOSPERMS |
| Saxifragales | Crassulaceae | Sedum album | SedalGNL2_ctg425 | sens | (core eudicots) | ANGIOSPERMS |
| Saxifragales | Crassulaceae | Rhodiola crenulata | Rhocr_CCG027441 | sens | (core eudicots) | ANGIOSPERMS |
| Saxifragales | Crassulaceae | Rhodiola crenulata | Rhocr_CCG023694 | sens | (core eudicots) | ANGIOSPERMS |
| Saxifragales | Crassulaceae | Rhodiola crenulata | Rhocr_CCG028218 | sens | (core eudicots) | ANGIOSPERMS |
| Saxifragales | Hamamelida-ceae | Hamamelis virginiana | HamviGNOMa_ctg1436 | sens | (core eudicots) | ANGIOSPERMS |

| | | | | | | |
|---|---|---|---|---|---|---|
| Saxifragales | Hamamelida-ceae | Hamamelis virginiana | HamviGNOMb_ctg1436 | res | (core eudicots) | ANGIOSPERMS |
| Saxifragales | Hamamelida-ceae | Hamamelis virginiana | HamviGNL2_ctg1273 | sens | (core eudicots) | ANGIOSPERMS |
| Santalales | Santalaceae | Santalum album | Sanal_SaT02071 | sens | (core eudicots) | ANGIOSPERMS |
| Santalales | Santalaceae | Santalum album | Sanal_SaT03034 | res | (core eudicots) | ANGIOSPERMS |
| Santalales | Santalaceae | Santalum album | Sanal_SaT05437 | sens | (core eudicots) | ANGIOSPERMS |
| Santalales | Olacaceae | Malania oleifera | Maole_005233 | sens | (core eudicots) | ANGIOSPERMS |
| Santalales | Olacaceae | Malania oleifera | Maole_012409 | res | (core eudicots) | ANGIOSPERMS |
| Santalales | Olacaceae | Malania oleifera | Maole_013225 | sens | (core eudicots) | ANGIOSPERMS |
| Santalales | Olacaceae | Malania oleifera | Maole_014154 | sens | (core eudicots) | ANGIOSPERMS |
| Vitales | Vitaceae | Vitis vinifera | Vitvi_LOC100247152 | sens | (core eudicots) | ANGIOSPERMS |
| Vitales | Vitaceae | Vitis vinifera | Vitvi_LOC100255904 | sens | (core eudicots) | ANGIOSPERMS |
| Vitales | Vitaceae | Vitis riparia | Vitri_LOC117929271 | sens | (core eudicots) | ANGIOSPERMS |
| Vitales | vitaceae | Vitis riparia | Vitri_LOC117925203 | sens | (core eudicots) | ANGIOSPERMS |
| Vitales | Vitaceae | Muscadinia rotundifolia | VITMro_g023260 | sens | (core eudicots) | ANGIOSPERMS |
| Vitales | Vitaceae | Muscadinia rotundifolia | VITMRO_g058050 | sens | (core eudicots) | ANGIOSPERMS |
| Vitales | Vitaceae | Muscadinia rotundifolia | VITMro_g386820 | sens | (core eudicots) | ANGIOSPERMS |
| Vitales | Vitaceae | Muscadinia rotundifolia | VITMro_g138390 | sens | (core eudicots) | ANGIOSPERMS |
| Fabales | Fabaceae | Medicago trunculata | Medtr_LOC25489342 | sens | (fabid rosids) | ANGIOSPERMS |
| Fabales | Fabaceae | Medicago trunculata | Medtr_LOC25489344 | sens | (fabid rosids) | ANGIOSPERMS |
| Fabales | Fabaceae | Medicago trunculata | Medtr_LOC11438427 | sens | (fabid rosids) | ANGIOSPERMS |
| Fabales | Fabaceae | Medicago trunculata | Medtr_LOC11437178 | sens | (fabid rosids) | ANGIOSPERMS |
| Fabales | Fabaceae | Trifolium pratense | Tp_L195_g002385 | sens | (fabid rosids) | ANGIOSPERMS |
| Fabales | Fabaceae | Trifolium pratense | Tp57577_mRNA23676 | sens | (fabid rosids) | ANGIOSPERMS |
| Fabales | Fabaceae | Trifolium pratense | Tp57577_mRNA2954 | sens | (fabid rosids) | ANGIOSPERMS |
| Fabales | Fabaceae | Trifolium pratense | Tp57577_mRNA31612 | sens | (fabid rosids) | ANGIOSPERMS |
| Fabales | Fabaceae | Phaseolus vulgaris | Phvul.001G235300 | sens | (fabid rosids) | ANGIOSPERMS |
| Fabales | Fabaceae | Phaseolus vulgaris | Phvul.008G231400 | sens | (fabid rosids) | ANGIOSPERMS |

| | | | | | | |
|---|---|---|---|---|---|---|
| Fabales | Fabaceae | Phaseolus vulgaris | Phvul.001G 235400 | res | (fabid rosids) | ANGIOSPERM S |
| Fabales | Fabaceae | Phaseolus vulgaris | Phvul.004G 161900 | sens | (fabid rosids) | ANGIOSPERM S |
| Fabales | Fabaceae | Vigna angularis | Vigan_LOC1 08329051 | sens | (fabid rosids) | ANGIOSPERM S |
| Fabales | Fabaceae | Vigna angularis | Vigan_LOC1 08338084 | sens | (fabid rosids) | ANGIOSPERM S |
| Fabales | Fabaceae | Vigna angularis | Vigan_LOC1 08329296 | res | (fabid rosids) | ANGIOSPERM S |
| Fabales | Fabaceae | Vigna angularis | Vigan10g01 7100 | sens | (fabid rosids) | ANGIOSPERM S |
| Fabales | Fabaceae | Arachis hypogaea | Arahy_LOC1 12733642 | sens | (fabid rosids) | ANGIOSPERM S |
| Fabales | Fabaceae | Arachis hypogaea | Arahy_LOC1 12791591 | sens | (fabid rosids) | ANGIOSPERM S |
| Fabales | Fabaceae | Arachis hypogaea | Arahy_LOC1 12765119 | sens | (fabid rosids) | ANGIOSPERM S |
| Fabales | Fabaceae | Arachis hypogaea | Arahy_LOC1 12749204 | sens | (fabid rosids) | ANGIOSPERM S |
| Fabales | Fabaceae | Abrus precatorius | Abrpr_LOC1 13846820 | sens | (fabid rosids) | ANGIOSPERM S |
| Fabales | Fabaceae | Abrus precatorius | Abrpr_LOC1 13849046 | sens | (fabid rosids) | ANGIOSPERM S |
| Fabales | Fabaceae | Abrus precatorius | Abrpr_LOC1 13846103 | sens | (fabid rosids) | ANGIOSPERM S |
| Fabales | Fabaceae | Abrus precatorius | Abrpr_LOC1 13866795 | sens | (fabid rosids) | ANGIOSPERM S |
| Fabales | Fabaceae | Cajanus cajan | Cajca_LOC1 09801720 | sens | (fabid rosids) | ANGIOSPERM S |
| Fabales | Fabaceae | Cajanus cajan | Cajca_LOC1 09808001 | sens | (fabid rosids) | ANGIOSPERM S |
| Fabales | Fabaceae | Cajanus cajan | Cajca_LOC1 09801940 | res | (fabid rosids) | ANGIOSPERM S |
| Fabales | Fabaceae | Cajanus cajan | Cajca_LOC1 09813450 | sens | (fabid rosids) | ANGIOSPERM S |
| Fabales | Fabaceae | Prosopis alba | Proal_LOC1 14732932 | res | (fabid rosids) | ANGIOSPERM S |
| Fabales | Fabaceae | Prosopis alba | Proal_LOC1 14748431 | sens | (fabid rosids) | ANGIOSPERM S |
| Fabales | Fabaceae | Prosopis alba | Proal_LOC1 14759906 | sens | (fabid rosids) | ANGIOSPERM S |
| Fabales | Fabaceae | Cercis canadensis | CercaGNOM a_sc32 | sens | (fabid rosids) | ANGIOSPERM S |
| Fabales | Fabaceae | Cercis canadensis | CercaGNOM b_sc32 | res | (fabid rosids) | ANGIOSPERM S |
| Fabales | Fabaceae | Cercis canadensis | CercaGNL2_ sc515 | sens | (fabid rosids) | ANGIOSPERM S |
| Rosales | Rosaceae | Fragaria vesca | Frave_LOC1 01300393 | sens | (fabid rosids) | ANGIOSPERM S |
| Rosales | Rosaceae | Fragaria vesca | Frave_LOC1 01300670 | res | (fabid rosids) | ANGIOSPERM S |
| Rosales | Rosaceae | Fragaria vesca | Frave_LOC1 01305794 | sens | (fabid rosids) | ANGIOSPERM S |

| | | | | | | |
|---|---|---|---|---|---|---|
| Rosales | Rosaceae | Fragaria vesca | Frave_LOC101307629 | sens | (fabid rosids) | ANGIOSPERMS |
| Rosales | Rosaceae | Prunus persica | Prupe_LOC18785443 | sens | (fabid rosids) | ANGIOSPERMS |
| Rosales | Rosaceae | Prunus persica | Prupe_LOC18784935 | sens | (fabid rosids) | ANGIOSPERMS |
| Rosales | Rosaceae | Prunus persica | Prupe_LOC18770144 | sens | (fabid rosids) | ANGIOSPERMS |
| Rosales | Rosaceae | Malus domestica | MDP0000129069 | sens | (fabid rosids) | ANGIOSPERMS |
| Rosales | Rosaceae | Malus domestica | MDP0000427621 | sens | (fabid rosids) | ANGIOSPERMS |
| Rosales | Rosaceae | Malus domestica | MDP0000410984 | sens | (fabid rosids) | ANGIOSPERMS |
| Rosales | Rosaceae | Malus domestica | MDP0000802196 | sens | (fabid rosids) | ANGIOSPERMS |
| Rosales | Rosaceae | Malus domestica | MDP0000427619 | res | (fabid rosids) | ANGIOSPERMS |
| Rosales | Rosaceae | Malus domestica | MDP0000565847 | sens | (fabid rosids) | ANGIOSPERMS |
| Rosales | Cannabaceae | Trema orientale | Treor_270070 | sens | (fabid rosids) | ANGIOSPERMS |
| Rosales | Cannabaceae | Trema orientale | Treor_079340 | sens | (fabid rosids) | ANGIOSPERMS |
| Rosales | Cannabaceae | Trema orientale | Treor_095630 | sens | (fabid rosids) | ANGIOSPERMS |
| Rosales | Cannabaceae | Parasponia andersonii | Paran_256170 | sens | (fabid rosids) | ANGIOSPERMS |
| Rosales | Cannabaceae | Parasponia andersonii | Paran_321370 | sens | (fabid rosids) | ANGIOSPERMS |
| Rosales | Cannabaceae | Parasponia andersonii | Paran_058240 | sens | (fabid rosids) | ANGIOSPERMS |
| Rosales | Cannabaceae | Cannabis sativa | Cansa_LOC115707648 | sens | (fabid rosids) | ANGIOSPERMS |
| Rosales | Cannabaceae | Cannabis sativa | Cansa_LOC115704658 | res | (fabid rosids) | ANGIOSPERMS |
| Rosales | Cannabaceae | Cannabis sativa | Cansa_LOC115716528 | sens | (fabid rosids) | ANGIOSPERMS |
| Rosales | Moraceae | Morus notabilis | Morno_LOC21408061 | sens | (fabid rosids) | ANGIOSPERMS |
| Rosales | Moraceae | Morus notabilis | Morno_LOC21408060 | res | (fabid rosids) | ANGIOSPERMS |
| Rosales | Moraceae | Morus notabilis | Morno_LOC21403706 | sens | (fabid rosids) | ANGIOSPERMS |
| Rosales | Rhamnaceae | Ziziphus jujuba | Zizju_LOC107413409 | sens | (fabid rosids) | ANGIOSPERMS |
| Rosales | Rhamnaceae | Ziziphus jujuba | Zizju_LOC107413408 | res | (fabid rosids) | ANGIOSPERMS |
| Rosales | Rhamnaceae | Ziziphus jujuba | Zizju_LOC107431830 | sens | (fabid rosids) | ANGIOSPERMS |

| | | | | | | |
|---|---|---|---|---|---|---|
| Rosales | Rhamnaceae | Rhamnella rubrinervis | Rharu_RR24768 | sens | (fabid rosids) | ANGIOSPERMS |
| Rosales | Rhamnaceae | Rhamnella rubrinervis | Rharu_RR24766 | res | (fabid rosids) | ANGIOSPERMS |
| Rosales | Rhamnaceae | Rhamnella rubrinervis | RharuGNL2_LG7 | sens | (fabid rosids) | ANGIOSPERMS |
| Rosales | Rosaceae | Rosa chinensis | Rosch_LOC112182803 | sens | (fabid rosids) | ANGIOSPERMS |
| Rosales | Rosaceae | Rosa chinensis | Rosch_LOC112182802 | sens | (fabid rosids) | ANGIOSPERMS |
| Rosales | Rosaceae | Rosa chinensis | Rosch_LOC112190848 | sens | (fabid rosids) | ANGIOSPERMS |
| Rosales | Rosaceae | Pyrus x bretschneideri | Pyrxb_LOC103935740 | sens | (fabid rosids) | ANGIOSPERMS |
| Rosales | Rosaceae | Pyrus x bretschneideri | Pyrxb_LOC103942956 | sens | (fabid rosids) | ANGIOSPERMS |
| Rosales | Rosaceae | Pyrus x bretschneideri | Pyrxb_LOC103935729 | sens | (fabid rosids) | ANGIOSPERMS |
| Rosales | Rosaceae | Pyrus x bretschneideri | Pyrxb_LOC103942957 | res | (fabid rosids) | ANGIOSPERMS |
| Rosales | Rosaceae | Pyrus x bretschneideri | Pyrxb_LOC103929457 | sens | (fabid rosids) | ANGIOSPERMS |
| Fagales | Betulaceae | Betula pendula | Betpe_c0365.g0008 | sens | (fabid rosids) | ANGIOSPERMS |
| Fagales | Betulaceae | Betula pendula | Betpe_c0281.g0095 | res | (fabid rosids) | ANGIOSPERMS |
| Fagales | Betulaceae | Betula pendula | Betpe_c1159.g0023 | sens | (fabid rosids) | ANGIOSPERMS |
| Fagales | Betulaceae | Carpinus fangiana | Carfa_001430 | sens | (fabid rosids) | ANGIOSPERMS |
| Fagales | Betulaceae | Carpinus fangiana | Carfa_007592 | sens | (fabid rosids) | ANGIOSPERMS |
| Fagales | Betulaceae | Carpinus fangiana | Carfa_003276 | sens | (fabid rosids) | ANGIOSPERMS |
| Fagales | Juglandaceae | Juglans regia | Jugre_LOC108991852 | sens | (fabid rosids) | ANGIOSPERMS |
| Fagales | Juglandaceae | Juglans regia | Jugre_LOC109006351 | sens | (fabid rosids) | ANGIOSPERMS |
| Fagales | Juglandaceae | Juglans regia | Jugre_LOC108985951 | res | (fabid rosids) | ANGIOSPERMS |
| Fagales | Juglandaceae | Juglans regia | Jugre_LOC108992663 | sens | (fabid rosids) | ANGIOSPERMS |
| Fagales | Fagaceae | Quercus suber | Quesu_LOC112039782 | sens | (fabid rosids) | ANGIOSPERMS |
| Fagales | Fagaceae | Quercus suber | Quesu_LOC111985027 | sens | (fabid rosids) | ANGIOSPERMS |
| Fagales | Fagaceae | Quercus lobata | Quelo_LOC115954940 | sens | (fabid rosids) | ANGIOSPERMS |

| | | | | | | |
|---|---|---|---|---|---|---|
| Fagales | Fagaceae | Quercus lobata | Quelo_LOC 115976920 | res | (fabid rosids) | ANGIOSPERM S |
| Fagales | Fagaceae | Quercus lobata | Quelo_LOC 115981000 | sens | (fabid rosids) | ANGIOSPERM S |
| Fagales | Fagaceae | Fagus sylvatica | FagsyGNOM _chr9 | sens | (fabid rosids) | ANGIOSPERM S |
| Fagales | Fagaceae | Fagus sylvatica | FagsyGNL1 _chr8 | res | (fabid rosids) | ANGIOSPERM S |
| Fagales | Fagaceae | Fagus sylvatica | FagsyGNL2 _chr12 | sens | (fabid rosids) | ANGIOSPERM S |
| Fagales | Myricaceae | Morella rubra | Morru_MR3 G014762 | sens | (fabid rosids) | ANGIOSPERM S |
| Fagales | Myricaceae | Morella rubra | Morru_MR2 G025548 | res | (fabid rosids) | ANGIOSPERM S |
| Fagales | Myricaceae | Morella rubra | Morru_MR0 G007998 | sens | (fabid rosids) | ANGIOSPERM S |
| Fagales | Casuarinace ae | Casuarina equisetifolia | CaseqGNO M_sc309 | sens | (fabid rosids) | ANGIOSPERM S |
| Fagales | Casuarinace ae | Casuarina equisetifolia | CaseqGNL1 _sc190 | res | (fabid rosids) | ANGIOSPERM S |
| Fagales | Casuarinace ae | Casuarina equisetifolia | CaseqGNL2 _sc144 | sens | (fabid rosids) | ANGIOSPERM S |
| Cucurbitales | Cucurbitace ae | Cucumis sativus | Cucsa_LOC 101204638 | sens | (fabid rosids) | ANGIOSPERM S |
| Cucurbitales | Cucurbitace ae | Cucumis sativus | Cucsa_LOC 101207938 | res | (fabid rosids) | ANGIOSPERM S |
| Cucurbitales | Cucurbitace ae | Cucumis sativus | Cucsa_LOC 101210509 | sens | (fabid rosids) | ANGIOSPERM S |
| Cucurbitales | Cucurbitace ae | Cucurbita maxima | Cucma_LOC 111490452 | sens | (fabid rosids) | ANGIOSPERM S |
| Cucurbitales | Cucurbitace ae | Cucurbita maxima | Cucma_LOC 111476959 | res | (fabid rosids) | ANGIOSPERM S |
| Cucurbitales | Cucurbitace ae | Cucurbita maxima | Cucma_LOC 111465412 | sens | (fabid rosids) | ANGIOSPERM S |
| Cucurbitales | Cucurbitace ae | Momordica charantia | Momch_LOC 111010039 | sens | (fabid rosids) | ANGIOSPERM S |
| Cucurbitales | Cucurbitace ae | Momordica charantia | Momch_LOC 111025550 | sens | (fabid rosids) | ANGIOSPERM S |
| Cucurbitales | Cucurbitace ae | Momordica charantia | Momch_LOC 111024228 | sens | (fabid rosids) | ANGIOSPERM S |
| Cucurbitales | Cucurbitace ae | Benincasa hispida | Benhi_LOC1 20080467 | sens | (fabid rosids) | ANGIOSPERM S |
| Cucurbitales | Cucurbitace ae | Benincasa hispida | Benhi_LOC1 20079606 | sens | (fabid rosids) | ANGIOSPERM S |
| Cucurbitales | Cucurbitace ae | Citrullus lanatus | CitlaGNOM_ chr7 | sens | (fabid rosids) | ANGIOSPERM S |
| Cucurbitales | Cucurbitace ae | Citrullus lanatus | CitlaGNL2_c hr 11 | sens | (fabid rosids) | ANGIOSPERM S |

| | | | | | | |
|---|---|---|---|---|---|---|
| Cucurbitales | Datiscaceae | Datisca glomerata | Datgl_137S20405 | sens | (fabid rosids) | ANGIOSPERMS |
| Cucurbitales | Datiscaceae | Datisca glomerata | Datgl_891S35907 | res | (fabid rosids) | ANGIOSPERMS |
| Cucurbitales | Datiscaceae | Datisca glomerata | Datgl_574S11525 | sens | (fabid rosids) | ANGIOSPERMS |
| Cucurbitales | Begoniaceae | Begonia fuchsioides | Begfu_1501S08048 | sens | (fabid rosids) | ANGIOSPERMS |
| Cucurbitales | Begoniaceae | Begonia fuchsioides | Begfu_4464S35236 | res | (fabid rosids) | ANGIOSPERMS |
| Cucurbitales | Begoniaceae | Begonia fuchsioides | Begfu1692S08615 | sens | (fabid rosids) | ANGIOSPERMS |
| Malpighiales | Salicaceae | Populus trichocarpa | Poptr_LOC18107297 | sens | (COM clade rosids) | ANGIOSPERMS |
| Malpighiales | Salicaceae | Populus trichocarpa | Poptr_LOC18107296 | res | (COM clade rosids) | ANGIOSPERMS |
| Malpighiales | Salicaceae | Populus trichocarpa | Poptr_LOC7458753 | sens | (COM clade rosids) | ANGIOSPERMS |
| Malpighiales | Salicaceae | Salix brachista | Salbr_026992 | sens | (COM clade rosids) | ANGIOSPERMS |
| Malpighiales | Salicaceae | Salix brachista | Salbr_026991 | res | (COM clade rosids) | ANGIOSPERMS |
| Malpighiales | Salicaceae | Salix brachista | Salbr_028175 | sens | (COM clade rosids) | ANGIOSPERMS |
| Malpighiales | Rhizophoraceae | Kandelia obovata | Kanob_002465a | sens | (COM clade rosids) | ANGIOSPERMS |
| Malpighiales | Rhizophoraceae | Kandelia obovata | Kanob_002465b | res | (COM clade rosids) | ANGIOSPERMS |
| Malpighiales | Rhizophoraceae | Rhizophora apiculata | Rhiap_sc34.1 | sens | (COM clade rosids) | ANGIOSPERMS |
| Malpighiales | Rhizophoraceae | Rhizophora apiculata | Rhiap_sc34.2 | res | (COM clade rosids) | ANGIOSPERMS |
| Malpighiales | Rhizophoraceae | Rhizophora apiculata | Rhiap_sc6.10 | sens | (COM clade rosids) | ANGIOSPERMS |
| Malpighiales | Euphorbiaceae | Jatropha curcas | Jatcu_LOC105630489 | sens | (COM clade rosids) | ANGIOSPERMS |
| Malpighiales | Euphorbiaceae | Jatropha curcas | Jatcu_LOC105633077 | sens | (COM clade rosids) | ANGIOSPERMS |
| Malpighiales | Euphorbiaceae | Jatropha curcas | Jatcu_LOC105638327 | sens | (COM clade rosids) | ANGIOSPERMS |
| Malpighiales | Euphorbiaceae | Ricinus communis | Ricco_LOC8281853 | sens | (COM clade rosids) | ANGIOSPERMS |
| Malpighiales | Euphorbiaceae | Ricinus communis | Ricco_LOC8283443 | res | (COM clade rosids) | ANGIOSPERMS |
| Malpighiales | Euphorbiaceae | Ricinus communis | Ricco_LOC107261358 | sens | (COM clade rosids) | ANGIOSPERMS |
| Malpighiales | Linaceae | Linum usitatissimum | Lus10026627 | sens | (COM clade rosids) | ANGIOSPERMS |
| Malpighiales | Linaceae | Linum usitatissimum | Lus10030432 | sens | (COM clade rosids) | ANGIOSPERMS |

| | | | | | | |
|---|---|---|---|---|---|---|
| Malpighiales | Linaceae | Linum usitatissimu m | Lus1003714 0 | sens | (COM clade rosids) | ANGIOSPERM S |
| Malpighiales | Linaceae | Linum usitatissimu m | Lus1003678 9 | sens | (COM clade rosids) | ANGIOSPERM S |
| Malpighiales | Linaceae | Linum usitatissimu m | Lus1002177 1 | sens | (COM clade rosids) | ANGIOSPERM S |
| Malpighiales | Linaceae | Linum usitatissimu m | Lus1003461 5 | sens | (COM clade rosids) | ANGIOSPERM S |
| Malpighiales | Euphorbiace ae | Manihot esculenta | Manes_LOC 110625274 | sens | (COM clade rosids) | ANGIOSPERM S |
| Malpighiales | Euphorbiace ae | Manihot esculenta | Manes_LOC 110625243 | sens | (COM clade rosids) | ANGIOSPERM S |
| Malpighiales | Euphorbiace ae | Manihot esculenta | Manes_LOC 110609669 | res | (COM clade rosids) | ANGIOSPERM S |
| Malpighiales | Euphorbiace ae | Manihot esculenta | Manes_LOC 110610972 | sens | (COM clade rosids) | ANGIOSPERM S |
| Malpighiales | Podostemac eae | Cladopus chinensis | Clach_0019 36 | sens | (COM clade rosids) | ANGIOSPERM S |
| Malpighiales | Podostemac eae | Cladopus chinensis | Clach_0196 47 | sens | (COM clade rosids) | ANGIOSPERM S |
| Malpighiales | Podostemac eae | Cladopus chinensis | Clach_0036 99 | sens | (COM clade rosids) | ANGIOSPERM S |
| Malpighiales | Podostemac eae | Cladopus chinensis | Clach_0252 06 | sens | (COM clade rosids) | ANGIOSPERM S |
| Malpighiales | Euphorbiace ae | Hevea brasiliensis | Hevbr_LOC1 10645704 | sens | (COM clade rosids) | ANGIOSPERM S |
| Malpighiales | Euphorbiace ae | Hevea brasiliensis | Hevbr_LOC1 10663786 | res | (COM clade rosids) | ANGIOSPERM S |
| Malpighiales | Euphorbiace ae | Hevea brasiliensis | Hevbr_LOC1 10653973 | sens | (COM clade rosids) | ANGIOSPERM S |
| Malpighiales | Euphorbiace ae | Vernicia fordii | Verfo_0544 07 | sens | (COM clade rosids) | ANGIOSPERM S |
| Malpighiales | Euphorbiace ae | Vernicia fordii | Verfo_0532 60 | res | (COM clade rosids) | ANGIOSPERM S |
| Malpighiales | Euphorbiace ae | Vernicia fordii | Verfo_0357 62 | sens | (COM clade rosids) | ANGIOSPERM S |
| Celastrales | Celastracea e | Tripterygium wilfordii | Triwi_LOC11 9991987 | sens | (COM clade rosids) | ANGIOSPERM S |
| Celastrales | Celastracea e | Tripterygium wilfordii | Triwi_LOC12 0016528 | sens | (COM clade rosids) | ANGIOSPERM S |
| Celastrales | Celastracea e | Tripterygium wilfordii | Triwi_LOC11 9996657 | res | (COM clade rosids) | ANGIOSPERM S |
| Celastrales | Celastracea e | Tripterygium wilfordii | Triwi_LOC11 9996659 | res | (COM clade rosids) | ANGIOSPERM S |
| Celastrales | Celastracea e | Tripterygium wilfordii | Triwi_LOC11 9996549 | sens | (COM clade rosids) | ANGIOSPERM S |
| Oxalidales | Cephalotace ae | Cephalotus follicularis | Cepfo_2467 5 | sens | (COM clade rosids) | ANGIOSPERM S |
| Oxalidales | Cephalotace ae | Cephalotus follicularis | Cepfo_2105 5 | res | (COM clade rosids) | ANGIOSPERM S |

| | | | | | | |
|---|---|---|---|---|---|---|
| Oxalidales | Cephalotaceae | Cephalotus follicularis | Cepfo_22974 | sens | (COM clade rosids) | ANGIOSPERMS |
| Oxalidales | Oxalidaceae | Averrhoa carambola | Aveca_019952 | sens | (COM clade rosids) | ANGIOSPERMS |
| Oxalidales | Oxalidaceae | Averrhoa carambola | Aveca_019955 | sens | (COM clade rosids) | ANGIOSPERMS |
| Oxalidales | Oxalidaceae | Averrhoa carambola | Aveca_014651 | sens | (COM clade rosids) | ANGIOSPERMS |
| Myrtales | Myrtaceae | Eucalyptus grandis | Eucgr_LOC104433426 | sens | (malvid rosids) | ANGIOSPERMS |
| Myrtales | Myrtaceae | Eucalyptus grandis | Eucgr_LOC104440351 | sens | (malvid rosids) | ANGIOSPERMS |
| Myrtales | Myrtaceae | Syzygium oleosum | Syzol_LOC115671905 | sens | rosids | ANGIOSPERMS |
| Myrtales | Myrtaceae | Syzygium oleosum | Syzol_LOC115690464 | sens | rosids | ANGIOSPERMS |
| Myrtales | Myrtaceae | Corymbia citriodora | Corci_B2628 | sens | rosids | ANGIOSPERMS |
| Myrtales | Myrtaceae | Corymbia citriodora | Corci_C2224 | sens | rosids | ANGIOSPERMS |
| Myrtales | Myrtaceae | Rhodamnia argentea | Rhoar_LOC115755598 | sens | rosids | ANGIOSPERMS |
| Myrtales | Myrtaceae | Rhodamnia argentea | Rhoar_LOC115729910 | sens | rosids | ANGIOSPERMS |
| Myrtales | Myrtaceae | Rhodamnia argentea | Rhoar_LOC115729907 | sens | rosids | ANGIOSPERMS |
| Myrtales | Lythraceae | Punica granatum | Pungr_LOC116199793 | sens | rosids | ANGIOSPERMS |
| Myrtales | Lythraceae | Punica granatum | Pungr_LOC116209960 | res | rosids | ANGIOSPERMS |
| Myrtales | Lythraceae | Punica granatum | Pungr_LOC116200818 | sens | rosids | ANGIOSPERMS |
| Sapindales | Rutaceae | Citrus sinensis | Citsi_LOC102627469 | sens | (malvid rosids) | ANGIOSPERMS |
| Sapindales | Rutaceae | Citrus sinensis | Citsi_LOC102624619 | res | (malvid rosids) | ANGIOSPERMS |
| Sapindales | Rutaceae | Citrus sinensis | Citsi_LOC102625158 | sens | (malvid rosids) | ANGIOSPERMS |
| Sapindales | Anacardiaceae | Anacardium occidentale | Anaoc_0003s0363 | sens | (malvid rosids) | ANGIOSPERMS |
| Sapindales | Anacardiaceae | Anacardium occidentale | Anaoc_0018s0976 | sens | (malvid rosids) | ANGIOSPERMS |
| Sapindales | Anacardiaceae | Anacardium occidentale | Anaoc_0010s1629 | res | (malvid rosids) | ANGIOSPERMS |
| Sapindales | Anacardiaceae | Anacardium occidentale | Anaoc_0019s0415 | sens | (malvid rosids) | ANGIOSPERMS |
| Sapindales | Anacardiaceae | Pistacia vera | Pisve_LOC116120515 | sens | (malvid rosids) | ANGIOSPERMS |
| Sapindales | Anacardiaceae | Pistacia vera | Pisve_LOC116112061 | res | (malvid rosids) | ANGIOSPERMS |
| Sapindales | Anacardiaceae | Pistacia vera | Pisve_LOC116127888 | sens | (malvid rosids) | ANGIOSPERMS |
| Sapindales | Sapindaceae | Acer yangbiense | Aceya_EZV62_017872 | sens | (malvid rosids) | ANGIOSPERMS |

| | | | | | | |
|---|---|---|---|---|---|---|
| Sapindales | Sapindaceae | Acer yangbiense | Aceya_EZV62_015425 | res | (malvid rosids) | ANGIOSPERMS |
| Sapindales | Sapindaceae | Acer yangbiense | Aceya_EZV62_012732 | sens | (malvid rosids) | ANGIOSPERMS |
| Sapindales | Sapindaceae | Xanthoceras sorbifolium | XansoGNOMa | sens | (malvid rosids) | ANGIOSPERMS |
| Sapindales | Sapindaceae | Xanthoceras sorbifolium | XansoGNOMb | sens | (malvid rosids) | ANGIOSPERMS |
| Sapindales | Sapindaceae | Xanthoceras sorbifolium | Xanso_XS14G07389 | res | (malvid rosids) | ANGIOSPERMS |
| Sapindales | Sapindaceae | Xanthoceras sorbifolium | Xanso_XS04G12756 | sens | (malvid rosids) | ANGIOSPERMS |
| Sapindales | Sapindaceae | Dimocarpus longan | Dimlo_026160 | sens | (malvid rosids) | ANGIOSPERMS |
| Sapindales | Sapindaceae | Dimocarpus longan | Dimlo_022708 | res | (malvid rosids) | ANGIOSPERMS |
| Sapindales | Sapindaceae | Dimocarpus longan | Dimlo_019311 | sens | (malvid rosids) | ANGIOSPERMS |
| Sapindales | Meliaceae | Azadirachta indica | AzainGNOM_ctg18078 | sens | (malvid rosids) | ANGIOSPERMS |
| Sapindales | Meliaceae | Azadirachta indica | AzainGNL1_ctg9988 | sens | (malvid rosids) | ANGIOSPERMS |
| Sapindales | Meliaceae | Azadirachta indica | AzainGNL2_ctg12079 | sens | (malvid rosids) | ANGIOSPERMS |
| Malvales | Malvaceae | Theobroma cacao | Theca_LOC18610338 | sens | (malvid rosids) | ANGIOSPERMS |
| Malvales | Malvaceae | Theobroma cacao | Theca_LOC18601571 | res | (malvid rosids) | ANGIOSPERMS |
| Malvales | Malvaceae | Theobroma cacao | Theca_LOC18588627 | sens | (malvid rosids) | ANGIOSPERMS |
| Malvales | Malvaceae | Gossypium raimondii | Gosra_LOC105779596 | sens | (malvid rosids) | ANGIOSPERMS |
| Malvales | Malvaceae | Gossypium raimondii | Gosra_LOC105798157 | res | (malvid rosids) | ANGIOSPERMS |
| Malvales | Malvaceae | Gossypium raimondii | Gosra_LOC105780543 | sens | (malvid rosids) | ANGIOSPERMS |
| Malvales | Malvaceae | Gossypium raimondii | Gosra_LOC105769188 | sens | (malvid rosids) | ANGIOSPERMS |
| Malvales | Malvaceae | Durio zibethinus | Durzi_LOC111281800 | sens | (malvid rosids) | ANGIOSPERMS |
| Malvales | Malvaceae | Durio zibethinus | Durzi_LOC111286997 | sens | (malvid rosids) | ANGIOSPERMS |
| Malvales | Malvaceae | Durio zibethinus | Durzi_LOC111291690 | res | (malvid rosids) | ANGIOSPERMS |
| Malvales | Malvaceae | Durio zibethinus | Durzi_LOC111284518 | sens | (malvid rosids) | ANGIOSPERMS |
| Malvales | Malvaceae | Durio zibethinus | Durzi_LOC111303223 | sens | (malvid rosids) | ANGIOSPERMS |
| Malvales | Malvaceae | Corchorus capsularis | CCACVL1_06544 | sens | (malvid rosids) | ANGIOSPERMS |
| Malvales | Malvaceae | Corchorus capsularis | CCACVL1_13722 | sens | (malvid rosids) | ANGIOSPERMS |
| Malvales | Malvaceae | Corchorus capsularis | CCACVL1_09433 | sens | (malvid rosids) | ANGIOSPERMS |

| | | | | | | |
|---|---|---|---|---|---|---|
| Malvales | Malvaceae | Corchorus olitorius | COLO4_27976 | sens | (malvid rosids) | ANGIOSPERMS |
| Malvales | Malvaceae | Corchorus olitorius | COLO4_12228 | sens | (malvid rosids) | ANGIOSPERMS |
| Malvales | Malvaceae | Corchorus olitorius | COLO4_34896 | sens | (malvid rosids) | ANGIOSPERMS |
| Malvales | Thymelaeaceae | Aquilaria sinensis | Aqusi_92.109 | sens | (malvid rosids) | ANGIOSPERMS |
| Malvales | Thymelaeaceae | Aquilaria sinensis | Aqusi_3.680 | res | (malvid rosids) | ANGIOSPERMS |
| Malvales | Thymelaeaceae | Aquilaria sinensis | Aqusi_sc277.24 | sens | (malvid rosids) | ANGIOSPERMS |
| Brassicales | Caricaceae | Carica papaya | Carpa_LOC110823971 | ??? | (malvid rosids) | ANGIOSPERMS |
| Brassicales | Caricaceae | Carica papaya | Carpa_LOC110820219 | res | (malvid rosids) | ANGIOSPERMS |
| Brassicales | Caricaceae | Carica papaya | Carpa_LOC110818636 | sens | (malvid rosids) | ANGIOSPERMS |
| Brassicales | Moringaceae | Moringa oleifera | Morol36145S10705 | sens | (malvid rosids) | ANGIOSPERMS |
| Brassicales | Moringaceae | Moringa oleifera | Morol33726S06189 | res | (malvid rosids) | ANGIOSPERMS |
| Brassicales | Moringaceae | Moringa oleifera | Morol36117S09591 | sens | (malvid rosids) | ANGIOSPERMS |
| Brassicales | Cleomaceae | Tarenaya hassleriana | Tarha_LOC104824072 | sens | (malvid rosids) | ANGIOSPERMS |
| Brassicales | Cleomaceae | Tarenaya hassleriana | Tarha_LOC104805178 | sens | (malvid rosids) | ANGIOSPERMS |
| Brassicales | Cleomaceae | Tarenaya hassleriana | Tarha_LOC104805485 | sens | (malvid rosids) | ANGIOSPERMS |
| Brassicales | Cleomaceae | Tarenaya hassleriana | Tarha_LOC104823043 | ? (V) | (malvid rosids) | ANGIOSPERMS |
| Brassicales | Cleomaceae | Tarenaya hassleriana | Tarha_LOC104799071 | sens | (malvid rosids) | ANGIOSPERMS |
| Brassicales | Cleomaceae | Cleome violacea | Clevi_0004s1596 | sens | (malvid rosids) | ANGIOSPERMS |
| Brassicales | Cleomaceae | Cleome violacea | Clevi_0004s2072 | res | (malvid rosids) | ANGIOSPERMS |
| Brassicales | Cleomaceae | Cleome violacea | Clevi_0068s0067 | sens | (malvid rosids) | ANGIOSPERMS |
| Brassicales | Brassicaceae | Aethionema arabicum | AetarGNOMa_sc2513 | sens | (malvid rosids) | ANGIOSPERMS |
| Brassicales | Brassicaceae | Aethionema arabicum | AetarGNOMb_sc4296 | sens | (malvid rosids) | ANGIOSPERMS |
| Brassicales | Brassicaceae | Aethionema arabicum | AetarGNL1_sc5916 | res | (malvid rosids) | ANGIOSPERMS |
| Brassicales | Brassicaceae | Aethionema arabicum | AetarGNL2_sc580 | sens | (malvid rosids) | ANGIOSPERMS |
| Brassicales | Brassicaceae | Microthlaspi erraticum | MERR_LOCUS32759 | sens | (malvid rosids) | ANGIOSPERMS |
| Brassicales | Brassicaceae | Microthlaspi erraticum | MERR_LOCUS42656 | res | (malvid rosids) | ANGIOSPERMS |
| Brassicales | Brassicaceae | Microthlaspi erraticum | MERR_LOCUS31978 | sens | (malvid rosids) | ANGIOSPERMS |

| | | | | | | |
|---|---|---|---|---|---|---|
| Brassicales | Brassicaceae | Myagrum perfoliatum | Myper_0002s2159 | sens | (malvid rosids) | ANGIOSPERMS |
| Brassicales | Brassicaceae | Myagrum perfoliatum | Myper_0019s0546 | res | (malvid rosids) | ANGIOSPERMS |
| Brassicales | Brassicaceae | Myagrum perfoliatum | Myper_0004s2011 | sens | (malvid rosids) | ANGIOSPERMS |
| Brassicales | Brassicaceae | Capsella rubella | Capru_LOC17897608 | sens | (malvid rosids) | ANGIOSPERMS |
| Brassicales | Brassicaceae | Capsella rubella | Capru_LOC17879376 | res | (malvid rosids) | ANGIOSPERMS |
| Brassicales | Brassicaceae | Capsella rubella | Capru_LOC17881284 | sens | (malvid rosids) | ANGIOSPERMS |
| Brassicales | Brassicaceae | Arabidopsis lyrata | AL1G25820 | sens | (malvid rosids) | ANGIOSPERMS |
| Brassicales | Brassicaceae | Arabidopsis lyrata | AL7G50810 | res | (malvid rosids) | ANGIOSPERMS |
| Brassicales | Brassicaceae | Arabidopsis lyrata | AL6G31120 | sens | (malvid rosids) | ANGIOSPERMS |
| Brassicales | Brassicaceae | Arabidopsis thaliana | GNOM / At1g13980 | sens | (malvid rosids) | ANGIOSPERMS |
| Brassicales | Brassicaceae | Arabidopsis thaliana | GNL1 / AT5G39500 | res | (malvid rosids) | ANGIOSPERMS |
| Brassicales | Brassicaceae | Arabidopsis thaliana | GNL2 / AT5G19610 | sens | (malvid rosids) | ANGIOSPERMS |

**Plant GBF1-related ARF-GRFs    FINAL LIST         5 Jan 2022**

Suppl. Data 1. ARF-GEFs analysed with the Nextstrain program

**175 total number of species analysed** (1 May 2022) (see Suppl. Data 1a)
 6 Chlorophyta
 2 Charophyta
 3 Bryophyta
 1 Lycopodiophyta
 2 ferns
 3 gymnosperms
 1 basal angiosperms (ANA clade)
11 monocots
 1 magnoliids
 1 Ceratophyllales
10 basal eudicots
49 asterids
85 rosids

**>At1g13980**
MGRLKLHSGIKAIEEEPEDFECTDSSNTTTLACMIDTEIAAVLAVMRRNVRWGGRYMSGDDQLEHSLIQSLKALR
KQVFSWNQPWHTISPMLYLQPFLDVIRSDETGAPIISIALSSVYKILNLNVIDQNTANIEDAMHLVVDSVTSCRF
EVTDPASEEVVLMKILQVLLACMKNKASVMLSNQHVCTVVNTCFRVVHQAGMKGELLQRVARHTMHELVRCIFSH
LPDVERTETTLVNRAGSIKQEKAGVDSDYAIVSKPVEDGNANSEYDVENSMATFATGAQSLMDDGPVGPGSRKPA
SPYDLHIMTEPYGVPSMVEIFHFLCSLLNVVEHVGMGSRSNTIAFDEDVPLFALNLINSAIELGGSSIRHHPRLL
SLIQDELFRNLMQFGLSMSPLILSMVCSIVLNLYQHLRTELKLQLEAFFSCVILRLAQGKYGPSYQQQEVAMEAL
VNFCRQKSFMVEMYANLDCDITCSNVFEELSNLLSKSTFPVNCPLSAMHILALDGLIAVIQGMAERISNGLTGLD
LGPVHLDEYTPFWMVKCDNYSDPNHWVSFVRRRKYIKRRLMIGADHFNRDPKKGLEFLQGTHLLPDKLDPQSVAC
FFRYTAGLDKNLVGDFLGNHDEFCVQVLNEFAGTFDFQYMNLDTALRLFLETFRLPGESQKIQRVLEAFSERYYM
QSPEILANKDAALVLSYSIIMLNTDQHNVQVKKKMTEEDFIRNNRHINGGNDLPREFLSELFHSICNNEIRTTPE
QGAGFPEMTPSRWIDLMHKSKKTAPYILADSRAYLDHDMFAIMSGPTIAAISVVFDHAEHEDVYQTCIDGFLAIA
KISACHHLEDVLDDLVVSLCKFTTLLNPSSVDEPVLAFGDDGKARMATITIFTIANKYGDYIRTGWRNILDCILR
LHKLGLLPARVASDAADESEHSSEQGQGKPLANSLSSAHLQSMGTPRRSSGLMGRFSQLLSLDTEEPRSQPTEQQ
LAAHQRTLQTIQKCHIDSIFTESKFLQAESLLQLARALIWAAGRPQKGTSSPEDEDTAVFCLELLIAITLNNRDR
IVLLWQGVYEHIATIAQSTVMPCNLVDKAIFGLLRICQRLLPYKESLADELLRSLQLVLKLDARVADAYCEQIAI
EVSRLVKANANHIRSQAGWRTITSLLSITARHPEASESGFDAVSFVMSEGTHLYPANYVLCVDAARQFAESRVGQ
SERSIRALDLMGDSLEFLAKWALSAKENMGEEDFGKMSQDIGEMWLRLVQGLRKVCLDQREDVRNHALQSLQKCL
GGVDGINLAHSMWSQCFDKVIFTVLDDLLEIAAGSQKDYRNMEGTLLLAIKLLSKVFLQQLQELSQLSTFCKLWL
GVLTRMEKYMKVKVRGKKSDKLQESVPELLKNILLVMKTKGVLLQRSALGGDSLWELTWLHVNNIAPSMRLELFP
DQESSQLGDDETVSNGLSSPENTTGS*
**>At5g39500**
MGYQNHPSGSNSFHGEFKRCHSKPSKGAVASMINSEIGAVLAVMRRNVRWGVRYIADDDQLEHSLIHSLKELRKQ
IFSWQSNWQYVDPRLYIQPFLDVILSDETGAPITGVALSSVYKILTLEVFTLETVNVGEAMHIIVDAVKSCRFEV
TDPASEEVVLMKILQVLLACVKSKASNGLSNQDICTIVNTCLRVVHQSSSKSELLQRIARHTMHELIRCIFSQLP
FISPLANECELHVDNKVGTVDWDPNSGEKRVENGNIASISDTLGTDKDDPSSEMVIPETDLRNDEKKTEVSDDLN
AAANGENAMMAPYGIPCMVEIFHFLCTLLNVGENGEVNSRSNPIAFDEDVPLFALGLINSAIELGGPSFREHPKL
LTLIQDDLFCNLMQFGMSMSPLILSTVCSIVLNLYLNLRTELKVQLEAFFSYVLLRIAQSKHGSSYQQQEVAMEA
LVDLCRQHTFIAEVFANFDCDITCSNVFEDVSNLLSKNAFPVNGPLSAMHILALDGLISMVQGMAERVGEELPAS
DVPTHEERYEEFWTVRCENYGDPNFWVPFVRKVKHIKKKLMLGADRFNRDPNKGLQYLQGVHLLPEKLDPKSVAC
FFRYTCGLDKNVMGDFLGNHDQFCIQVLHEFAKTFDFQNMNLATALRLFVGTFKLSGEAQKIHRVLEAFSERYYE
QSPHILIDKDAAFVLAYSIILLNTDQHNAQVKTRMTEEDFIRNNRTINGGADLPREYLSEIYHSIRHSEIQMDED
KGTGFQLMTASRWISVIYKSKETSPYIQCDAASHLDRDMFYIVSGPTIAATSVVFEQAEQEDVLRRCIDGLLAIA
KLSAYYHLNSVLDDLVVSLCKFTPFFAPLSADEAVLVLGEDARARMATEAVFLIANKYGDYISAGWKNILECVLS
LNKLHILPDHIASDAADDPELSTSNLEQEKPSANPVPVVSQSQPSAMPRKSSSFIGRFLLSFDSEETKPLPSEEE
LAAYKHARGIVKDCHIDSIFSDSKFLQAESLQQLVNSLIRASGKDEASSVFCLELLIAVTLNNRDRILLIWPTVY
EHILGIVQLTLTPCTLVEKAVFGVLKICQRLLPYKENLTDELLKSLQLVLKLKAKVADAYCERIAQEVVRLVKAN
ASHVRSRTGWRTIISLLSITARHPEASEAGFEALRFIMSEGAHLLPSNYELCLDAASHFAESRVGEVDRSISAID
LMSNSVFCLARWSQEAKNSIGETDAMMKLSEDIGKMWLKLVKNLKKVCLDQRDEVRNHAISMLQRAIAGADGIML
PQPLWFQCFDSAVFILLDDVLTFSIENSRKTLKKTVEETLVLATKLMSKAFLQSLQDISQQPSFCRLWVGVLNRL
ETYMSTEFRGKRSEKVNELIPELLKNTLLVMKATGVLLPGDDIGSDSFWQLTWLHVNKISPSLQSEVFPQEELDQ
FQRRNAKPEDPPVPGNEV*
**>At5g19610**

MDRIAVRAKRKELGISCMLNTEVGAVLAVIRRPLSESYLSPQETDHCDSSVQQSLKSLRALIFNPQQDWRTIDPS
VYLSPFLEVIQSDEIPASATAVALSSILKILKIEIFDEKTPGAKDAMNSIVSGITSCRLEKTDLVSEDAVMMRIL
QVLTGIMKHPSSELLEDQAVCTIVNTCFQVVQQSTGRGDLLQRNGRYTMHELIQIIFSRLPDFEVRGDEGGEDSE
SDTDEIDMSGGYGIRCCIDIFHFLCSLLNVVEVVENLEGTNVHTADEDVQIFALVLINSAIELSGDAIGQHPKLL
RMVQDDLFHHLIHYGASSSPLVLSMICSCILNIYHFLRKFMRLQLEAFFSFVLLRVTAFTGFLPLQEVALEGLIN
FCRQPAFIVEAYVNYDCDPMCRNIFEETGKVLCRHTFPTSGPLTSIQIQAFEGLVILIHNIADNMDREEDEGNEE
DDNNSNVIKPSPVEIHEYIPFWIDKPKEDFETWVDHIRVRKAQKRKLAIAANHFNRDEKKGLEYLKYNYLVSDPL
DPMALASFFRFTPGLDKTMIGDYLGDPDELHLSVLRSFTHTFEFTGMNLDTALRTFLESFRLPGESQKIERMIEA
FSERFYDQQSSDIFASKDTVHILCYSLIMLNTDQHNPQVRRKMTEDEFIRNNRAINAGNDLPKEYLSELFQSIAT
NAFALSTHSGPVEMNPNRWIELMNRTKTTQPFSLCQFDRRIGRDMFATIAGPSIAAVSAFFEHSDDDEVLHECVD
AMISIARVAQYGLEDILDELIASFCKFTTLLNPYTTPEETLFAFSHDMKPRMATLAVFTLANTFGDSIRGGWRNI
VDCLLKLRKLQLLPQSVIEFEINEENGGSESDMNNVSSQDTKFNRRQGSSLMGRFSHFLALDNVEESVALGMSEF
EQNLKVIKQCRIGQIFSKSSVLPDVAVLNLGRSLIYAAAGKGQKFSTAIEEEETVKFCWDLIITIALSNVHRFNM
FWPSYHEYLLNVANFPLFSPIPFVEKGLPGLFRVCIKILASNLQDHLPEELIFRSLTIMWKIDKEIIETCYDTIT
EFVSKIIIDYSANLHTNIGWKSVLQLLSLCGRHPETKEQAVDALIGLMSFNASHLSQSSYAYCIDCAFSFVALRN
SSVEKNLKILDLMADSVTMLVKWYKTASTDTANSYSPASNTSSSSSMEENNLRGVNFVHHLFLKLSEAFRKTTLA
RREEIRNRAVTSLEKSFTMGHEDLGFTPSGCIYCIDHVIFPTIDDLHEKLLDYSRRENAEREMRSMEGTLKIAMK
VLMNVFLVYLEQIVESAEFRTFWLGVLRRMDTCMKADLGEYGDNKLQEVVPELLTTMIGTMKEKEILVQKEDDDL
WEITYIQIQWIAPALKDELFPDEEI*

**>AL1G25820**

MGRLKLHSGIKAIEEEPEDFESTDSSNTTTLACMIDTEIAAVLAVMRRNVRWGGRYMSGDDQLEHSLIQSLKALR
KQVFSWNQPWHTISPMLYLQPFLDVIRSDETGAPITSIALSSVYKILNLNVIDQNTANIEDAMHLVVDSVTSCRF
EVTDPASEEVVLMKILQVLLACMKNKASVMLSNQHVCTVVNTCFRVVHQAGMKGELLQRVARHTMHELVRCIFSH
LPDVDRTESTLVNRAGSIKQEKAGVDSDYAIVSKPVEDGNANSEYDNENSVATFATGAQSLMDDGLVGPGSRKPA
SPYDLHIMTEPYGVPSMVEIFHFLCSLLNVVEHVGMGSRSNTIAFDEDVPLFALNLINSAIELGGSSIRHHPRLL
SLIQDELFRNLMQFGLSMSPLILSMVCSIVLNLYQHLRTELKLQLEAFFSCVILRLAQGKYGPSYQQQEVAMEAL
VNFCRQKSFMVEMYANLDCDITCSNVFEELSNLLSKSTFPVNCPLSAMHILALDGLIAVIQGMAERISNGLTGLD
LGPVHLDEYTPFWMVKCDNYSDPNHWVSFVRRRKYIKRRLMIGADHFNRDPKKGLEFLQGTHLLPDKLDPQSVAC
FFRYTAGLDKNLVGDFLGNHDEFCVQVLHEFAGTFDFQYMNLDTALRLFLETFRLPGESQKIQRVLEAFSERYYM
QSPEILANKDAALVLSYSIIMLNTDQHNVQVKKKMTEEDFIRNNRHINGGNDLPREFLSELFHSICNNEIRTTPE
QGAGFPEMTPSRWIDLMHKSKKTAPYIMADSRAYLDHDMFAIMSGPTIAAISVVFDHAEHEDVYQTCVDGFLAIA
KISACHHLEDVLDDLVVSLCKFTTLLNPSSVDEPVLAFGDDAKARMATITIFTIANKYGDYIRTGWRNILDCILR
LHKLGLLPARVASDAADESELSSEQGQGKPLANSLSSAHLQSMGTPRRSSGLMGRFSQLLSLDTEEPRSQPTEQQ
LAAHQRTLQTIQKCHIDSIFTESKFLQAESLLQLARALIWAAGRPQKGTSSPEDEDTAVFCLELLIAITLNNRDR
IVLLWQGVYEHIATIAQSTVMPCNLVDKAIFGLLRICQRLLPYKESLADELLRSLQLVLKLDARVADAYCEQIAI
EVSRLVKANANHIRSQAGWRTITSLLSITARHPEASEAGFNAVSFVMSEGTHLYPANYVLCVDAARQFAESRVGQ
SERSIRALDLMGDSLEYLAKWALSAKENMGEEDFGKMSQDIGEMWLRLVQGLRKVCLDQREDVRNHALQALQKCL
GGVDGINLAHSMWSQCFDKVIFTVLDDLLEIAGGSQKDYRNMEGTLLLAIKLLSKVFLQQLQELSQLSTFCKLWL
GVLTRMEKYMKVKVRGKKSDKLQESVPELLKNILLVMKTKGVLLQRSALGGDSLWELTWLHVNNIAPSMRLELFP
DQESTQLGDDETVSNGLSSPEAGANTTGS*

**>AL7G50810**

MGYQNHPSGSNSFHGEFKRCHSKPSKGAVASMINSEIGAVLAVMRRNVRWGVRYIADDDQLEHSLIHSLKELRKQ
IFSWQSNWQYVDPRLYIQPFLDVILSDETGAPITGVALSSVYKILTLEIFTLETVNVGEAMHIIVDAVKSCRFEV
TDPASEEVVLMKILQVLLACVKSKASNGLSNQDICTIVNTCLRVVHQSSSKSELLQRIARHTMHELIRCIFSQLP
FISPLANESELHVHDKVGIVDWDQNSGEKRVENGNIASVSDTLGTDKDSPSSEMVIPETELRNDEKKTEVSDDLN
ADANGENAMMAPFGIPCMVEIFHFLCTLLNVGENGEVNSRSNPIAFDEDVPLFALGLINSAIELGGPFFRDHPKL
LTLIQDELFCNLMQFGMSMSPLILSTVCSIVLNLYLNLRTELKVQLEAFFSYVLLRIAQSKHGSSYQQQEVAMEA
LVDLCRQHTFIAEMFANFDCDITYSNVFEDVSNLLSKSAFPVNGPISAMHILALDGLISMVQGMAERVGEEFPAS
DVPTHEERYEEFWTVRCENYGDPNFWVPFVRKAKHIKKKLMLGADHFNRDPKKGLQYLQGMHLLPEELDPKSVAC
FFRYTCGLDKNLIGDFLGNHDQFCIQVLHEFAKTFDFQNMNLATALRLFVGTFRLPGEAQKIQRVLEAFSERYYE
QSPHILIDKDAAFVLAYSIILLNTDQHNAQVRTRMTEEDFIRNNRTINGGADLPREYLSEIYHSIRHSQIEIEMN
PDEGTGFQLMTASRWISVIYKSKETSPYIQCDTASHLDRDMFYIVSGPTIAATSVVFEQTEQEDVLQRCIDGLLA
IAKLSAYYHLNSVLDDLVVSLCKFTPFFAPLSADEAVLALGEDARARMATEAVFLIANKYGDYISSGWKNILECV
LSLNKLHILPDHIASDAADDPELSTSSLEQEKPSANPLPVISQSQPSATPRKSSSFIGRFLMSFDSEETKPLPTE
EELAAYKHARGIVKDCHIDSIFSDSKFLQAESLQQLVNSLIKASGKDEASSVFCLELLIAVTLNNRDRILLIWQT
VYEHILGIVQPTLTPCTLVEKAVFGVLKICQRLLPYKENLTDELLKSLQLVLKLKPRVADAYCERITQEVVHLVK
ANASHIRSHTGWRTIISLLSITARHPEASDAGFEALRFIMSEGAHLLPSNYELCLDAAKNFAISRVGEIDRSISA
IDLMSNSVFCLARWSQEAKNSIGETDAMMKLSEDIGEMWLALVNKLQIVCYDQRDQVRNHAILMLQRAIAGADGI
MLPQPIWFQCFDSAVFPLLDDKSLAFAIENSRKNFKKTVEETLVLATKLMSKAFLQSLQDISQQPSFCRLWVGVLN
RLETYMSTEFRGKRSEKVHELIPELLKNTLLVMKATGVLLPGDDIGSDSFWQLTWLHVKKISPSLQSEVFPQEEL
DQFQRRNAKPEDSPVPENEA*

**>AL6G31120**

MDRIAVRAKRKELGISCMLNTEVGAVLAVIRRPLSESYLSPQETDHCDSSVQQSLKSLRALIFNPQQDWRTIDPS
VYLSPFLEVIQSDEIPASATAVALSSILKILKIEIFDEKTPGAKDAMNSIVSGITSCRLEKTDLLSEDAVMMRIL
QVLIGIMKHPSSELLEDQAVCTIVNTCFQVVQQSTGRGDLLQRNGRYAMHELIQIIFSRLPDFEVRGDDGGEESE
SDTDEIDMSGGYGIRCCIDIFHFLCSLLNVVEVVENSEGTSVHTADEDVQIFALVLINSAIELSGDAIGQHPKLL
RMVQDDLFHHLIHYGASSSPLVLSMISSCILNIYHFLRKFMRLQLEAFFSFVLLRVTAFTGFLPLQEVALEGLIN
FCRQPAFIVEAYVNYDCDPMCRNIFEETGKVLCRHTFPTSGPLTSIQIQAFEGLVILIHNIADNMDREEEEGAEE
DDNSNAIKPSPVEIHEYIPFWIDKPKEDFETWVDHIRVRKAQKRKLAIAANHFNRDEKKGLEYLKYNYLVSDPLD
PMALASFFRFTPGLDKTMIGDYLGDPDELHLSVLKSFTHTFEFTGMNLDTALRTFLESFRLPGESQKIERMIEAF
SERFYDQQSSDIFASKDTVHILCYSLIMLNTDQHNPQVRKKMTEDEFIRNNRAINAGNDLPKEYLSELFQSIATN
AFALSTHSGPVEMNPNRWIELMNRTKTTQPFSLCQFDRRIGRDMFATIAGPSIAAVSAFFEHSDDDEVLHECVDA
MISIARVAQYGLEDILDELIASFCKFTTLLNPYTTPEETLFAFSHDMKPRMATLAVFTLANNFGDSIRGGWRNIV
DCLLKLRKLQLLPQSVIEFEITNEENNGGSESDMNIVSNQDTKFNRRQGSSLMGRFSHFLALDSVEESLALGMSE
FEQNLKVIKQCRIGQIFSKSSVLPDVAVLNLGRSLIYAAAGKGQKFSTAIEEEETVKFCWDLIITIALSNVHRFN
MFWPSYHEYLLNVANFPLFSPIPFVEKGLPGLFRVCIKILASNLQDHLPEELIFRSLTIMWKIDKEIIETCYDTI
TEFVSKIITVYSANLHTNIGWKSVLQLLSLCGRHPETKEQAVDALIGLMSINASHLSQSSYAYCIDCAFSFVALR
NSSVEKNLKILDLMADSVTMLVKWYKTASTDTANSYSPASNTSSSSSMEENSLRGVNFVHHLFLKLSEAFRKTTL
ARREEIRNRAVTSLEKSFTMGHEDLGFTPSGCIYCIDHVIFPTIDDLHEKLLDYSRRENAEREMRSMEGTLKIAM
KMLMNVFLVYLQQIVESAEFRTFWLGVLRRMDTCMKADLGEYGDNKLQEVVPELLTTMIGTMKEKKILVQKEDDD
LWEITYIQIQWIAPSLKDELFPDEEI*

**>Capru_LOC17897608**

MGRLKLHSGINSIEEEPEDFESTDSSNTTTLACMIDTEIAAVLAVMRRNVRWGGRYMAGDDQLEHSLIQSLKALR
KQVFSWNQPWHTISPILYLQPFLDVIRSDETGAPITSIALSSVYKILNLNVIDQHTANIEDAMHLVVDSVTSCRF
EVTDPASEEVVLMKILQVLLACMKNKASVMLSNQHVCTVVNTCFRVVHQAGMKGELLQRVARHTMHELVRCIFAH
LPDVDRTENTLVNRAGSIKQEKAGVDSDYAIVSKPVEDGNGNSEYDTENSVATFASGAQTLMDDGPVGPGSRKPA
SPYDLHIMTEPYGVPSMVEIFHFLCSLLNVVEHVGMGSRSNTIAFDEDVPLFALNLINSAIELGGSSIRHHPRLL
SLIQDELFRNLMQFGLSMSPLILSMVCSIVLNLYQHLRTELKLQLEAFFSCVILRLAQGKYGPSYQQQEVAMEAL
VNFCRQKSFMVEMYANLDCDITCSNVFEELSNLLSKSTFPVNCPLSAMHILALDGLIAVIQGMAERISNGLTGLD
LGPVHLDEYTPFWMVKCDNYSDPNHWVSFVRRRKYIKRRLMIGADHFNRDPKKGLEFLQGTHLLPDKLDPQSVAC
FFRYTAGLDKNLVGDFLGNHDEFCVQVLNEFAGTFDFQYMNLDTALRLFLETFRLPGESQKIQRVLEAFSERYYM
QSPEILANKDAALVLSYSIIMLNTDQHNVQVKKKMTEEDFIRNNRHINGGNDLPREFLSELYHSICNNEIRTTPE
QGAGFPEMTPSRWIDLMHKSKKTAPYIMADSRAYLDHDMFAIMSGPTIAAISVVFDHAEHEDVYQTCIDGFLAIA
KISACHHLEDVLDDLVVSLCKFTTLLNPSSVDEPVLAFGDDAKARMATITIFTIANKYGDYIRTGWRNILDCILR
LHKLGLLPARVASDAADESELSSEQGQGKPLANSLSSAHLQSMGTPRRSSGLMGRFSQLLSLDTEEPRSQPTEQQ
LAAHQRTLQTIQKCHIDSIFTESKFLQAESLLQLARALIWAAGRPQKGTSSPEDEDTAVFCLELLIAITLNNRDR
IVLLWQGVYEHIATIAQSTLMPCNLVDKAIFGLLRICQRLLPYKESLADELLRSLQLVLKLDARVADAYCEQIAV
EVSRLVKANANHIRSQAGWRTITSLLSITAKHPEASEAGFDAVSFVMSEGTHLYPANYVLCVDAARQFAESRVGQ
SERSIRALDLMEDSLKYLAKWVRTAKENMGEEDFGKMSQDIGEMWLWLVQGLRKVCLVQREDVRNHALQSLQKCL
GGVDGINLGHSMWSQCFDKVIFTLLDDLLELAAGSQKDYRNMEGTLLLAIKLLSKVFLQQLQELSQLSTFCKLWL
GVLTRMEKYMKVKVRGKKSDKLQESVPELLKNILLVMKTKGVLLQRSALGGDSLWELTWLHVNNIVPSMRLELFP
DQESTQLGDDETVANGLSSPEAGANTAGL*

**>Capru_LOC17879376**

MGYQNHPSGSNSFHGEFKSCHSKPSKGAVASMINSEIGAVLAVMRRNVRWGVRYIADDDQLEHSLIHSLKELRKR
IFSWQSNWQNVDPILYIRPFLDVILSDETGAPITGVALSSVYKILTLDIFTLETVNVGEAMHIIVDAVKSCRFEV
TDPASEEVVLMKILQVLLACIKSKASSGLSNQDICTIVNTCLRVVHQSSSKSELLQRIARHTMHELIRYIFSQLP
SISPLAIENELHVDDKVGTVDWDQNSGENKVGNGNIASISDTPGTENNSPSSETVIPETELKNDEKKTEVNDDLN
VPANGENAMMAPFGIPCMVEIFHFLCTLLNVGEHGEVNSRSNPIAFDEDVPLFALGLINSAIELGGTSFREYPKL
LTLIQDELFCNLMQFGVSMSPLILSTVCSIVLNLYLNLRTELKVQLEAFFSHVLLRIAQSKHGSSYQQQEVAMEA
LVDLCRQHTFMAEVFANFDCDITCSNVFEDVANLLSKSAFPVNGPLSAMHILALDGLISMVQGMAERVGEELPSS
DVHTHEEGYEEFWTARCENYGDPNFWVPFVRKAKRIKKKLMLGADNFNRDPKKGLQYLQGMHLLPEELDPKSVAC
FFRYTCGLDMDLIGDYLGNHDQFCIQVLYEFAKTFDFQNMNLATALRLFVGTFRLPGESQKIQRVLEAFSERYYE
QSPHILINKDAAFVLAYSIILLNTDQHHSQVKTRMTEEDFIRNNRGINGGADLPREYLSEIYHSICNIEIQMNAT
KGTSFQMMTASRWISVIYKSKETSPYILCDAASHLDRDMFCIVSGPTIAATSVVFEQAEQEDVFQRCVDGLLAIA
KLSAYYHLNSVVDDVVVSLCKFTPFSTPLSADEAVLALGEDARARMATEAVFLIANKYGDYISSAWKNILECVLS
LHKLQILPVHIASDAADDTEPSTSNLEQEKPSTNPLSVVSESQPSVTPRKTSSFIGRFLWSFDMEETKPLPTEEE
QVAYKHARGIVNDCHIDSIFSDSKFLQAESLQHLVNSLIKAAEKNEASAVFCLELLIAVTLNNRDRILLIWQTVY
EHISGIIQSTSTPCTLVEKAVFGVLKICQRLLPYKENLTDELLKSLQLVLKLNANVADAYCERITQEVGRLVKAN
ASHVRSHIGWRTIISLLSITARHPEASDAGFEALRFIMSEGAHLLPSNYILCLDAAWQFAGSRVNEIDRSISAID
LMSNSVFCLARWSQEAKNSVGEDDAMVKLSRDIGEMWLRLVHNLKRVCRDPRDQVRNHAILMTTRAIAGADGIML
PQPLWFQCFDSEVFPLLDETLADSILNSKKTFKKTVEETLVLATKLMSKAFLQSLQDISQQPSFCKLWLGVLNRF
ETYMSTKFRGKRSEKIHELIPELLKNTLLVMKTTGVLLPGDDIGSDSFWQLTWLHVKKISPSLQSEVFPEEELDQ
LQTRNAKPEDSQTSENDAGD*

**>Capru_LOC17881284**

MERIAVKAKRKELGISCMLNTEVGAVLAVIRRPLAESYLSPQETDHCDSSVQQSLKSLRALIFNPQQDWRTIDPS
VYLSPFLEVIQSDEIPASATAVALSSILKILKIEIFDEKTPGAKDAMNSIVSGITSCRLEKTDLVSEDAVMMRIL
QVLIGIMKHPSSDLLEDQAVCTIVNTCFQVVQQSSGRGDLLQRNGRYAMHELIQIIFSRLQDFEVRGEEGGEESE
SDTDEIDMSNGYGIRCCIDIFHFLCSLLNVVEVVENSEGTNVHTADEDVQIFALVLINCAIELSGDAIGQHPKLL
RMVQDDLFHHLIHYGASSSPLVLSMISSCILNIYHFLRKFMRLQLEAFFSFVLLKVTALTGFLQLQEVALEGLIN
FCRQPAFIVEAYVNYDCDPMCRNIFEETGKVLCRHTFPTSGPLTSLQIQAFEGLVILIHNIADNMDKEEDEGDEE
IDQSNAIKPSPVEIHEYIPFWVDKPKEDFETWVEHIRVRKAQKRKLAIAANHFNRDEKKGLEYLKYNYLVSDPPD
PMALASFFRFTPGLDKTMIGDYLGDPDEFHLSVLRSFTYTFEFTGMNLDTALRTFLESFRLPGESQKIERMIEAF
SERFYKQQSTDIFASKDTVHILCYSLIMLNTDQHNPQVKKKMTEDEFIRNNRAINANNDLPREYLSELFQSIATN
AFALSTHSGPVEMNPNRWIELMNRTKTTQPFSMCQFDRRIGRDMFATIAGPSIAAVAAFFEHSDDDEVLHECVDA
MISIARVAQYGLEDILDELIASFCKFTTLLNPYTTPEETLFAFSHDMKPRMATLAVFTLANNFGDSIRGGWRNIV
DCLLKLRKLQLLPQSVIEFETDDEAINNGGSEPDMNTVSNQDIKFNRRQGSSLMGRFSHFLALDSVEESVALGMS
EFEQNLKVIKQCRIGQIFSRSAVLPDVAVLNLGRSLIYAAAGKGQKFSTAIEEEETVKFCWDLIIAVALSNIHRF
NMFWPSYHEYLLNVANFPLFSPVPFVEKGLPGLFKVCLKILASNLQDHLPEELIFRSLTIMWKIDKEIIETCYDT
ITEFVSNIITDYSGNLHTNIGWKSVLQLLSLCGRHPETKDQAVDALIGLMSVNASNLSQSSYAYCIDCAFSFVAL
RNSSVEKNLNILDLMAESVTMLIKWYKTASTDSVNSNSPASNTSSSSSLEENSMRGVNFVHHLFLKLSEAFRKTT
LARREEIRNKAVTSLEKSFTMSHEDLGFTPSGCIYCIDHVIFPTIDDLHEKLLDYSRRENAEREMRSMEGTLKIA
MKMLMNVFLVYLEQIVESAEFRTFWLGVLRRMDTCMKADLGEYGDNKLQEVVPELLITMIGTMKEKEILVQKEDD
DLWEITYIQIQWIAPSLKDELFPDEEI*

**> Myper_0002s2159**

MGRLKLHPGIKAIEEEPEDFESTESSNTTTLACMIDTEIAAVLAVMRRNVRWGGRYMSGDDQLEHSLIQSLKALR
KQLFSWNQPWHTISPMLYLQPFLDVIRSDETGAPITSIALSSVYKILSLNVIDQNTANIEDAMHLVVDSVTSCRF
EVTDPASEEVVLMKILQVLLACMKNRASVMLSNQHVCTVVNTCFRVVHQAGMKGELLQRVARHTMHELVRCIFSH
LPDVDRTETTLVNRAGSINQEKAGVDNDYAIVSKPVEDGNANSEYDTENSVATFASGAQSLMDDGPAGPGSRKPA
SPYDLHIMTEPYGVPSMVEIFHFLCSLLNVVEHVGMGARSNTIAFDEDVPLFALNLINSAIELGGSSIRHHPRLL
SLIQDELFRNLMQFGLSMSPLILSMVCSIVLNLYQHLRTELKLQLEAFFSCVILRLAQGKYGPSYQQQEVAMEAL
VNFCRQKNFMVEMYANLDCDITCSNVFEELSNLLSKSTFPVNCPLSAMHILALDGLIAVIQGMAERISNGLTGLD
LGPVHLDEYTPFWMVKCDNYSDPNHWVSFVRRRKYIKRRLMIGADHFNRDPKKGLEFLQGTHLLPDKLDPQSVAC
FFRYTAGLDKNLVGDFLGNHDEFCVQVLNEFAGTFDFQYMNLDTALRLFLETFRLPGESQKIQRVLEAFSERYYM
QSPEILANKDAALVLSYSIIMLNTDQHNVQVKKKMTEEDFIRNNRHINGGNDLPREFLSELYHSICNNEIRTTPE
QGSGFPEMTPSRWIDLMHKSKKTAPYIMADSRAYLDHDMFAIMSGPTIAAISVVFDHAEHEDVYQTCIDGFLAIA
KISACHHLEDVLDDLVVSLCKFTTLLNPSSVDEPVLAFGDDAKARMATITIFTIANKYGDYIRTGWRNILDCILR
LHKLGLLPARVASDAADESELSSEQGQGKPLANSLSSAHLQSMGTPRRSSGLMGRFSQLLSLDTEEPRSQPTEQQ
LAAHQRTLQTIQKCHIDSIFTESKFLQAESLLQLARALIWAAGRPQKGNNSPEDEDTAVFCLELLIAITLNNRDR
IVLLWQGVYEHIATIAQSTVMPCNLVEKAIFGLLRICQRLLPYKESLADELLRSLQLVLKLDARVADAYCENIAL
EVSRLVKANANHIRSQAGWRTITSLLSITARHPEASEAGFNAVSFVMSEGTHLFPANYVLCVDAARQFAESRVGQ
SERSIRALDLMGDSLEYLAKWARSAKENMGEEDFARMSQDFGEMWLRLVQGLRKVCLDQREDVRNHALQSLQKCL
GGVDGINLAHSMWSQCFDKVIFTVLDDLLEIASGSQKDYRNMEGTLLLAIKLLSKVFLQLVQELSQLSTFCKLWL
GVLTRMEKYMKVKVRGKKSDKLQETVPELLKNILLVMKTKGVLLQRSALGGDSLWELTWLHVNNIAPSMRLELFP
DQESTQLGDDETVLNPSSNGLASPEAGANIITA*

**>Myper_0019s0546**

MGYQNHHHPSGSNSFHGGEFKSCHSNKPSKGAVASMINSEIGAVLAVMRRNVRWGVRYIADDDQLEHSLIHSLKE
LRKQIFSWQSNWQNVDPKLYIQPFLDVILSDETGAPITGVALSAVYKILTLDVFTLDTVNVGEAMHIIVDAVKSC
RFEVTDPASEEVVLMKILQVLLACIKSKAANGLSNQDICTIVNTCLRVVNQSSSKSELLQRIARHTMHELVRCIF
SQLPYISPLANESELQVGDKVGTVDWDQNSGENKVENGNIDSVSDNLGTDKDSPSSQMVIPETELQNDEKGTEVS
DDLNVAANGESAMLAPYGIPCMVEIFHFLCTLLNVGENGEVNSRSNPIAFDEDVPLFALGLINSVIELGGSSFRE
HPKLLSLIQDELFCNLMQFGVSMSPLILSTVCSIVLNLYLSLRTELKVQLEAFFSCVLLRIAQSKHGSSYQQQEV
AMEALVDLCRQHTFMAEVFANFDCDITCSNVFEDVSNLLSKSAFPVNGPLSAMHILALDGLISMVQGMAERVGEE
LPASDVSKHEEGYEAFWTVRCENYGDPNCWVPFVRKSKHIKKKLMVGADHFNSDPKKGLQQLQAMHLLPEELDPK
SVACFFRYTCGIDKNLIGDFLGNHDQFCVQVLNEFAKTFDFQNMNLDNALRLFVGTFRLPGESQKIQRVLEAFSE
RYYEQSPQILIDKDAALLLSYSIILLNTDHHNTQVKKKMTEEDFIRNNRCINGGADLPREYLSELYHSICGSEIQ
MIPDQGTGFQMMTASRWISVIYKSKETSPYIICDSASYLDRDMFHIVSGPTIAATSVVFEQAEQDDVIQRCVDGL
LAIAKLSAYYNLNSVLDDLVVSLCKFTPFFASLSVDEAVLALGEDARARMATEAVFLIANRYGDYINSGWKNILE
CVLSLHKLQILPAHVASDAADDQEPSTSNSEQERLLANPVSVVSQVQPSATPRKSSSFIGRFSQLLSFDMEETKP
LPTEEELAAYKHARGIVKDCHIDSIFSDSKFLQAESLQQLVNSLIKAAAKDEASAVFCLELLIAVTLNNRDRILL
IWQTVYEHISGIVQSTSTPCPLVEKAVFGVLKICQRLLPYKENLTDELLKSLQLVLKLDARVADAYCEPITQEVA
RLVKANASHVRSHIGWRTIISLLSITARHPEASDAGFEALRFIMSEGAHLLPSNYILCLDAARQFAESRVGEIDR
SISAIDLMSNSVFCLARWSQEAKNSVGEDEAMMKLSQDIGEMWLRLVNNLKRVCLDQRDQVRNHAILMLQRAIAG
ADGIMLPQPLWFQCFDSAIFPLLDELLAVSIDNSRKTFKKTVEETLVLATKLMSKAFLQSLQDISQQPSFCRLWL
GVLNRMETYMSTKFRGKRSEKVHELIPELLKNTLLVMKTTGVLLPGDDIGSDSFWQLTWLHVKKISPSLQSEVFP
QEELDQFQRRNAKPEDTPPETVVE*

**>Myper_0004s2011**

MERVAVKAKRKELGISCMLNTEVGAVLAVIRRPLAESSYLSPQETDHCDSSVQHSLKSLRALIFNPQQDWRTIDP

SVYLSPFLEVIQSDEIPASATAVALSSILKILKIEIFDEKTPGAKDAMNSIVSGITSCRLEKTDPISEDAVMMRI
LQVLTGIMKHPASELLEDQAVCTIVNTCFQVVQQSAGRGDLLQRNGRYTMHELIQIIFSRLPDFEVRGDEGGEDS
ESDTDEIDMSGGYGIRCCIDIFHFLCSLLNVVEVVENSEGTSVHTADEDVQIFALVLINSAIELSGDAIGQHPKL
LRMVQDDLFHHLIHYGASSSPLVLSMISSCILNIYHFLRKFVRLQLEAFFSFVLLKVTAFTGFLQLQEVALEGLI
NFCRQPAFIVEAYVNYDCDPMCRSVFEETGKVLCRHTFPTSGPLTSLQIQAFEGLVILIHNIADNMDREEDEGGD
EEESNAIKPSAVEIHEYTPFWVEKPKEDFETWVDHIRVRKAQKRKLAIAANHFNRDEKKGLEYLKYNHLVSDPPD
PMALASFFRFTPGLDKTMIGDYLGDPDELHLSVLKSFTHTFEFTGMNLDTALRTFLESFRLPGESQKIERMIEAF
SERFYEQQSSEIFATKDTVHILCYSLIMLNTDQHNPQVKKKMTEDEFIRNNRAINAGKDLPREYLSELFQSISTN
AFALSTHSGPVEMNPNRWIELMNRTRTTQPFSLCQFDRRIGRDMFATIAGPSIAAVSAFFEHSDDDEVLHECVDA
LISIARVAQYGLEDILDELIASFCKFTTLLNPYTTPEETLFAFSHDMKPRMATLAVFTLANNFGDSIRGGWRNIV
DCLLKLRKLQLLPQSAIEFETTDEETLNKSGSEPDLNILPSQETKSNRRQGSSLMGRFSHFLALDSVEESVALGM
SEFEQNLKVIKQCRIGQIFSKSSVLPDVAVLNLGRSLIYAAAGKGQKFSTATEEEETVKFCWDLIIAVALSNIHR
FSMFWPSYHEYLINVTTFPLFSPVPFVEKGIPGLFKVCIKILASQLQDQLPEELIFRSITLMWKIDKEIIDTCYD
TITEFVSKIITEHSANLHTQIGWKSVLQLVSLCGRHPETKEQAVDALIGLMSTNASHLSQPSYAYCIDCAFSFVA
LRNSAVEKNLRILDLMTDSVTMLVKWYKTASTETGNNFSVASNTSSSSSAEENSLRGVNFVHHLFLKLAEAFRKT
TLARREEIRNRAVTSLEKSFIMAHEDLGFTPSGCIYCIDHVVFPTIDDLHEKLLDYSRRENAEREMRSMEGTLKT
AMKMLQNVFLHYLEQIVGSAEFRTFWLGVLRRMDTCMKADLGEYGDNKLQEVVPELLTIMIGTMKEKEILVQKED
DDLWEITYIQIQWIAPSLKDELFPDEEI*

**>MERR_LOCUS32759**

MGRLKLHSGIKAIEEEPEDFESTDSSNTTTLACMIDTEVAAVLAVMRRNVRWEGRYMSGDDQLEHSLIQSLKALR
KQLFSWSQPWHTISPMLYLQPFLDVIQSDETGAPITSIALSSVYKILSLNVIDQNTANIEDAMHLVVVSVTSCRF
EVTDPASEEVVLMKILQVLLACMRNKASVMLSNQHVCTVVNTCFRVVHQAGMKGELLQRVARHTMHELVRCIFSH
LPDVDRTENTLVNRAGNIDQEKAGDDGEYAIVSKPVEDGNANSVYDTENSVATFAPGGQSLMDDGPVGPGSRKPA
SPYDLHIMTEPYGVPSMVEIFHFLSSLLNVVEHAGMGPRSNTIAFDEDLPLFALNLINSAIELGGSSIRHHPRLL
SLIQDELFRNLMQFGLSMSPLILSMVCSIVLNLYQHLRTELKLQLEAFFSCVILRLAQGKYGPSYQQQEVAMEAL
VNFCRQKNFMVEMYANLDCDITCSNVFEELSNLLSKSTFPVNCPLSAMHILALDGLIAVIQGMAERISNGLTGLD
LGPVHLDEYTPFWMVKCDNYSDPNHWVSFVRRRKYIKRRLMIGADHFNRDPKKGLEFLQGTHLLPDKLDPQSVAC
FFRYTAGLDKNLVGDFLGNHDEFCVQVLNEFAGTFDFQYMNLDTALRLFLETFRLPGESQKIQRVLEAFSERYYM
QSPEILANKDAALVLSYSIIMLNTDQHNVQVKKKMTEEDFIRNNRHINGGNDLPREFLSELYHSICNNEIRTTPE
QGSGFPEMTPSRWIDLMHKSKKTAPYIMADSRAYLDHDMFAIMSGPTIAAISVVFDHAEHEDVYQTCIDGFLAIA
KISACHHLEDVLDDLVVSLCKFTTLLNPSSVDEPVLAFGDDAKARMATITIFTIANKYGDYIRTGWRNILDCILR
LHKLGLLPARVASDAAEESELSSEQGQGKPLANSLSSAHLQSIGTPRRSSGLMGRFSQLLSLDTEEPRSQPTEQQ
LAAHQRTLQTIQKCHIDSIFTESKFLQAESLLQLARALIWAAGRPQKGNSSPEDEDTAVFCLELLIAITLNNRDR
IVLLWQGVYEHIATIAQSTVMPCNLVEKAIFGLLRICQRLLPYKESLADELLRSLQLVLKLDARVADAYCENIAL
EVSRLVKANANHIRSQAGWRTITSLLSVTARHPEASEAGFNAISFVMSEGSHLFPANYVLCVDAARQFAESRVGQ
SERSIRAVDLMGDSLEYLAKWARSAKENMGEEEFGKKSQDIGEMWLRLVQGLRKVCLDQREDVRNHALQSLQKCL
GGVNGINLAHSMWSQCFDKVIFTVLDDLLEIASGSQKDYRNMEGTLLLAIKLLSKVFLQLLQELSQLSTFCKLWL
GVLTRMEKYMKVKVRGKKSDKLQESVPELLKNILLVMKTKGVLLQRSALGGDSLWELTWLHVNNIAPSMRLELFP
DQDNTQIGDDETPSNGLSSPETGANTTG*

**>MERR_LOCUS42656**

MGYQNHPSGSNSFHGELKSCHSKPSKGALASMINSEIGAVLAVMRRNVRWGVRYIADDDQLEHSLIHSLKELRKQ
IFSWQCNWQNVDPKLYIQPFLDVILSDETGAPITGVALSSVYKILTLDIFTLDTVNVGEAMHIIVDAVKSCRFEV
TDPASEEVVLMKILQVLLACIKSKAANGLSNQDICTIVNTCLRVVHQSSSKSELLQRIARHTMHELVRCIFCQLP
YISPLANESELHVGDKVGTVDWDPNAGENKVENGDIGGVSDTLITEKDCSSSEVAIPETEPKNEEKETEVSDDVN
VAANGESVMMAPFGIPCMVEIFHFLCTLLNVGENGEVSSRSNPIAFDEDVPLFALGLINSAIELGGPSFREHPKL
LALIQDELFCNLMQFGVSMSPLILSTVCSIVLNLYLSLRSELKIQLEAFFSCVLLRIAQSKHGSSYQQQEVAMEA
LVDLCRQHTFMAEVFANFDCDITCSNVFEDVSNLLSKSAFPVNGPLSAMHILALDGLISMVQGMAERVGEEVSAS
DVSADEEGYEAFWTVRCENYGDPNFWVPFVRKAKHIKKKLMLGADNFNSDPKKGLKQLQTMHLLPEELDPKSVAC
FFRYTGGLDKNLIGDYLGNHDQFCVQVLHEFAKTFDFQNMNLDTALRLFVGTFRLPGESQKIQRVLEAFSERYYE
QSPQILINKDAALLLSYSIILLNTDHHNTQVKKKMTEEDFIRNNRLINGGADLPREYLSELYHSICDSEIQMIPD
QGTGFQMMTASRWISVVYKSKETSPYIICDSASHLDRDMFSIAFGPTIAATSVVFEQAEQEDVLQRCVDGLLAIA
KLSAYYHLNSVLDDLVVSLCKFTPFFAQISPDEAVLALGEDARARMATEAVFLIANRYGDYISSSWKNILECVLS
LQKLQILPAHIASDAADDPEPSTSNLEQEKPSANPVTVVAQVHPSATPRKSSSFIGRFSQLLYFDMEETKPLPTE
EDLAAYKHARGIVKDCHIDSIFSDSKFLQAESLQQLVNSLIKAAEKNEASAVFCLELLIAVTLNNRDRILLIWQT
VYEHISGIVQSTSTPCPLVEKAVFGVLKICQRLLPYKENLTDELLKSLQLVLKLDARVADAYCEPITQEVARLVK
ANASHVRSHVGWRTIISLLSITARHPEASDAGFEALRFIMSEGAHLLPSNYILCLDAARQFAESRVGEIDRSISA
IDLMSNSVLCLARWSQEAKNSVGEDEAMVKLSQDIGEMWLRLVNNLKRVCLDQRDQVRNHAILMLHRAIAGADGI
MLPQPLWFQCFDSAIFPLLDELLTFSVENSRKTFKKTVEETLVLATKLMSKAFLQSLQDISQQPSFCRLWLGVLD
RMETYMSTKFRGKRSEKVHELIPELLKNTLLVMKTTGVLLPGDDIGSDSFWQLTWLHVKKISPSLQSEVFPQEEL
DQFQRRNAKPEETPVPENEG*

**>MERR_LOCUS31978**

MDRITVKAKRKELGISCMLNTEVGAVLAVIRRPIAEGYLSPQETDHCDSSVQQSLKSLRALIFNPQQDWRTIDPS
VYLSPFLEVIQSDEIPASATAVALSSILKILKIEIFDEKTPGAKDSMITIVSGITSCRLEKTDSVSEDAVMMRIL

QVLVGIMKHPASELLEDQAICTIVNTCFQVVQQSSGRGDLLQRNGRYTMHELIQIIFSRLPDFEVRGDEGGEDSE
SDTDEIDMSGGYGIRCCIDIFHFLCSLLNVVEVVENSEGTSVHTADEDVQIFALVLINSAIELSGDAIGQHPKLL
RMVQDDLFHHLIHYGASSSPLVLSMISSCILNIYHFLRKFTRLQLEGFFSFVLLKVTAYTGFLQLQEVALEGLIN
FCRQPAFIVEAYVNYDCDPICKNVFEETGKVLCRHTFPTSGPLTSLQIQAFEGLVILIHNIADNMDREEDEGDEE
DSNVTKPSPVEIHEYIPFWIEKPKEDFETWVDHIRVRKAQKRKLAIAANHFNRDEKKGLEYLKYNHLVSDPPDPM
ALASFFRFTPGLDKTMIGDYLGDPDELHLSVLKSFTHTFEFTGMNLDTALRTFLESFRLPGESQKIERMIEAFSE
RFYEQQSSEIFASKDTVHILCYSLIMLNTDQHNPQVKKKMTEDEFIRNNRAINAGKDLPREYLSELFQSISSNAF
ALSTHSGPVEMNPNRWIELMNRTKTTQPFSLCQFDRRIGRDMFATIAGPSIAAVSAFFEHSDDDEVLHECVDALI
SIARVAQYGLEDILDELIASFCKFTTLLNPYTTPEETLFAFSHDMKPRMATLAVFTLANNFGDSIRGGWRNIVDC
LLKLRKLQLLPQSAIEFETTDEENLNNVGSETDLNNVSGPDTKSNRRQGSSLMGRFSHFLALDSVEESVALGMSE
FEQNLKVIKQCRIGQIFSKSSVLPDVAVLNLGRSLIYAAAGKGQKFSTAIEEEETVKFCWDLITAVALSNVHRFN
MFWPSYHEYLLNVANFPLFSPIPFVEKGIPGLFKVCIKILATNLQDNLPEELIFRSITIMWKIDKEIIDTCYDTI
TEFVSKIITEYSANLHTQIGWKSVLQLLSLCGRHPETKEQAVDALIGLMSTNASHLSQPSYAYCIDCAFSFVALR
NSAVEKNLRILDLMTDSVSMLIKWYKTASTETVSNYSVASNTSSSSSAEENSLRGVNFVHHLFLKLSEAFRKTTL
ARREEIRNRAVTSLEKSFIMGHVDLGFTPSGCIYCIDHVIFPTIDDLHEKLLDYSRRENAEREMRSMEGTLKIAM
KMLHNVFLHYLEQIVGSAEFRTFWLGVLRRMDTCMKADLGEYGDNKLQEVVPELLTTMIGTMKEKEILVQKEDDD
LWEITYIQIQWIAPALKDELFPDEEM*

**>AetarGNOMa_sc2513**

MGRLKLQSGIKAIEEEPEDLESIYSSKTTALACMIDTEIAAVLAVMRRSVRWGGRYMSGDDQLEHSLIQSLKALR
KQVFSWQQPWHTINPALYLQPFLDVIRSDETGAPPIMTIALSSVYKILSPDIIDESTVNIEESMHLVVDSVTSCR
FEVTDPASEEVVLMKILQVLLACMKNKASVVLSNQHVCTVVNTCFRIVHQAGMKSELLQRVARHTMHELVRCIFS
HLPDVDRTESAIVNRAGSIKQEIAGVDSDNAVVGKPTNNENANSENDTENSLVAFASGAPASADEGLVGPGGRKE
ASPYDLHLMTEPYGVPSMVEVFHFLCSLLNVVEHDGMGPRSNTIAFDEDMPMFALSMISSAIELGGSSIRHHPRL
LGLIQDELFRNLMQFGLSTSPLILSMVCSIVLSLYQHLRSELKLQLEAFFSCVILRLAQGKYGSSYQQQEVAMES
LVNFCRQKSFMVEMYANLDCDITCSNVFEDLSNLLSKSAFPVNCPLSSMHILALDGLIAVIQGMAERISNGLAGL
DLGRVHLDEYTPFWMVKCDNYSDPNHWVPFVRRRKYIKRRLMIGADHFNRDPKKGLEFLQGTHLLPDKLDPQSVA
CFFRYTAGLDKNLVGDFLGNHDEFCVQVLVEFAGTFDFQYMNLDTALRLFLETFRLPGESQKIQRVLEAFSERYY
EQSPEILANKDAALVLSYSIIMLNTDQHNVQVKKKMTEEDFIRNNRHINGGSDLPREFLSELFYSICNNEIRTTP
EQGAGFPEMNPSRWIDLMHKSKKTAPYIVADSRAYLDHDMFAIMSGPTIAAISVVFDHAEQEDVYQTCIEGFLAI
AKISACHHledvlddlvvSLCKFTTLLNPSSVDEPVLAFGDDAKARMATITIFTIANKYGDYIRSGWRNIVDCIL
RLHKLGLLPARVVSDAADEAELSSDPGQGKPVTNSLSSVHLPSMGTPRRSSGLMGRFSQLLSLETEEPRSQPTEQ
QLGAHQRTLQTIQKCHIDSIFTESKFLQAESLLQLARALIWAAGRPQKGNSSPEDEDTAVFCLELLIAITLNNRD
RIVLLWQGVYEHIANIVQSTVMPCALVEKAIFGLLRICQRLLPYKESLADELLRSLQLVLKLDARVADAYCEQVA
LEVSRLVKANASHIRSQAGWRTITSLLSVTARHPEASEAGFSAISFVMSDGAHLSPANYILCADAARQFAESRVG
QSERSICALDLIGGSVECLAKWARATRERMGEDDSGRMSQESGEMWLRLVQGLRKVCLDQREDVRNHALESLQQC
LVGVDGIQLAHSMWSQCFDKVIFTVLDDLLEIAQGSQKDYRNMEGTLLLAVKLLSKVFlqllqdlvqlSTFCKLW
LGVLTRmekymkvkvkgkksekLQEVVPGLLKNILLVMKTKEVLVQRSALGGDSLWELTWLHVNNITPSMRSEVF
PDHDPPEFTQLGGLAGDETISNGLASPKGESNTAG*

**>AetarGNOMb_sc4296**

MGRLKMHSGIKSIEEEPEDFESTDSSKTSTLACMIDTEISAVLAVMRRNVRWGGRYMSGDDQLEHSLIQSLKALR
KQIFSWNQPWHTISPTSYLRPFLDVIRSDETGAPITSIALSSVYKILSLNVIDQNTANIEDAMHLIVDSVTSCRF
EVTDPASEEVVLMKILQVLLACMKNKASVMLSNQHVCTVVNTCFRVVHQAGMKGELLQRVARHTMHELVRCIFSH
LPNVDRTENTLVNRAGSIKQEEADVDNDYAIVSKPDDDGNANSEYDAENTIVTFASGVQGLMNDGSVGLGSKKEV
SPYDLHLMTEPYGVPSMVEIFQFLCSLLNMVEHVGMGPRENTISFDENVPLFALKLINSAIELGGSSIRHHPRLL
SLIQDELFRNLMQFGLSMSPLILSIVCSIVLNLYQHLRTELKLQLEAFFSCVILRLAQGKYGASYQQQEVAMEAL
VNFCRKKSFMVEMYANLDCDITCSNVFEELSNLLSKSAFPVNCPLSAMHILAFDGLIAVIQGMAERVSNGVSGLD
LGPVHLDEYTPFWMVKCENYSDPSHWVQFVRRRKFIKRRLMIGADHFNRDPKKGLEFLQGTHLLPDKLDPQSVAC
FFRYTAGLDKNLVGDFLGNHDDFCVQVLNEFAGTFDFQYMNLDTALRLFLETFRLPGEAQKIQRVLEAFSERYYV
QSPEILANKDAALVLSYSIIMLNTDQHNVQVKKKMTEEDFIRNNRHINGGSDLPREFLSELYHSICNNEIRTIPE
KGAGFSEMNPSRWIDLMHKSKKTAPYIMANSRAYLDHDMFAIMSGPTIAAISVVFDHAEHEDVYQTCIDGFLAIA
KISACHHledvlddlvvSLCKFTTLLNPSSVDEPVLAFGDDakariatitiftiaNKYGDYIRMGWRNILDCILR
LHKLGLLPARVASEAASDESELPSEPGQGKPltsslssahlssMGTPRRSSGLMGRFSQLLSLDTEEPRSQPTEQ
QLAAHQRTLQTIQKCHIDSIFTESKFLQAESLLQLARALIWAAGRPQKGNNSPEDEDTAVFCLELLIAITLNNRD
RIVLLWQGVYEHIASIAQSTVIPCALVEKAVFGLLRICQRVLPYKESLADELLRSLQLVLKLDARVADAYCEQIA
LEVNRLVKANANHIRSQAGWRTITSLLSITARHPEASEAGFNALSFVMSDGTHLVPANYVLCADIARQFAESRVG
QSDRSIRSLDLMADSQECLAKWFHSAKEIMTEEEDFTKMLQDIGEMWLRLVQGLRKVCLDQREDIRNHALQSLQK
CLDRVDEIKLAQTVWSQCFDKVIFTVLDDLLEIASGTQKDYRNMEGTlllaikllsklflqllqdlskfsTFCKL
WLGVLTRMEKYMKVKIKGKKSEKLQESIPELLKNILLVMKTKEVLIQRSALGSDSLWELTWLHVNNIDPSLRLEL
FPDHEPVQPGLTSPEAGTTTAGL*

**>AetarGNL1_sc5916**

MRYQNLPSGSNSFHGEFKSCHSKSPKGSVACMINSEIGAVLAVMRRNVRWGVRYNADDEQLDHSLIHSLKELRKH
IFSWQSNWQNVDPRLYIQPFLDVILSDETGAPITGVALSSVYKILTLDIFTLDTVNVGEAMHIIVDAVKSCRFEV
TDPASEEVVLMKILQVLLACIKSNASNGLSNQDICTIVNTCLRVVHQSSSKSELLQRIARHTMHELVRCIFSQLP

YISPLANESELHVGMSVVTVDWDQNSGEKKVQNGDIGNTSDSLSADKGSLSPEMAILETKPTNDEEGTEVTDDLN
VAENGESTMMAPFGIPCMVEIFHFLCSLLNVGKNTEVNSRSNPIAFDEDVPLFALGLINSAIELGGPSFREHPKL
LILIQDELFCNLMQLGMSMSPLILSTVCSIVLNLYLDLRTELKVQLEAFFSCVLLRIAQSKHGSSYQQQEVAMEA
LVDLCRQHAFMAEVYANFDCDITCSNVFEDISNLLSKSAFPVNGPLSAMHILALDGLISMVQGMAKRVGEELPAS
DVPTQEEGYEAFWTVKCENYGDPNLWVPFVRKAKHLKKRLMLGADNFNEDPKKGLQYLQGMHLLPDELDPKSVAC
FFRYTCGLDKNLIGDFLGNHDQFCVQVLHEFAKTFDFQNMNLDTALRLFVGTFRLPGESQKIQRVLEAFSERYYE
QSPQILIDKDaalllSYSIILLNTDHHNMQVKKKMTEEDFIRNNRCINGGADLPREYLSELYHSICDSEIQMIPD
QGTGFQMMTASRWISVIHKSKEISPYIVCDTASHLDHDMFAILSGPTIAATSVVFEQAEQEDVLQRCVDGLLAIA
KISAYYHLNsvlddlvvslCKFTPLFAPLSIDEAVLALGDDARARMATEAVFLIANRYGDYIREGWRNVLDCVLR
LQKLQVLPAHIASDAADDLETSTSNPEQEKPAANPLSVVSHVQPSATPRKSSSLIGRFSQLLSFDMEETKPLPTE
EEVAAYKHAREIVKNCHIDSIFSDSKFLQAESLQQLVNSLITAAKKDDASAVFCLELLIAVTLNNRDRILLIWQT
VYDYISSIVQSISTPCTLVEKTVFGVLKICQRLLPYKENLTDEllkslqlvlklDARVADAYCEPITQEVGRLVK
ANASHVRSHAGWRTIISLLSITARHPEASEAGFEALRFIMAEGAHLLPSNYILCLDAARQFAESRVGETDRSISA
IDLMANSVFCLVRWSQEAKNSVGEDEAIKVSKDIGEMWLRLVNNLKRICLDQRDEVRNHAILMLQRSVAGADGIM
LPQSLWFQCFDSAIFPLLDELLEVSIENSRKNLTKTAEETLVHATKLMSKAFLQSLQDVSQQPSFCRLWLGVLNR
METYMATKFRGKRSEKVHELIPELLKNTLLVMKTTGILLPGDDIGSDSFWQLTWLHVKKISPSLQSEVFPQEEVD
QFQRRNAKQEDLPVPA*

**>AetarGNL2_sc580**

MEERVVAKARRKEVGISCMLNTEVGAVLAVIRRPIADAYIASQETDHCDSSVQQSLKSLRALIFNPQQEWRTIDP
SIYLSPFLEVVQSDEIPASATAVALSSILKMLKIEIFDEKTPGAKDGMNSIVSGITSCRLEKTDSMSEDAVMMRI
LQVLIGIMKHPASDLLEDQAVCTIVNTCFQVVQQSAGRGDLLQRNGRYTMHELIQIIFSRLPdfevsgdedgeds
esdtdeidmsGGYGIRCCIDIFHFLCSllnvvelvenleGTTVHTADEDVQIFALVLINSAIELSGDAIGQHPKL
LRMVQDDLFHHLIHYGAsssplvlsmissCILNIYHFLRKFVRLQLEAFFSFVLLRVTSFTGFFQLQEVALEGLI
NFCRQPAFIIEAYVNYDCDPMCRSIFEEMGKVLCRHTFPTSGPLTSLQIQAFEGLVILIHNIADYMdreeddgde
eeeeekeedKNHLIKPTPIEIHEYIPFWIDKINEDFETWVEHIRVRKAQKRKLAIAANHFNRDEKKGLEYLKYNY
LVSDPPDPKALASFFRFTPGLDKTMIGDYLGDPDELHLSVLRSFTHTFEFSGMNLDTALRTFLESFRLPGESQKI
ERMIEAFSERFYEQQSSEIFASKDTVHILCYSLIMLNTDQHNPQVKKKMTEDEFIRNNRAINAGKDLPREYLSEL
FQSISSNAFALSTNTNPVEMNPNRWIELMNKTKTTKSFSVCEFDRRIGRDMFATIAGPSIAAISAFFEHSDDDEV
LHECVDALISIARVAQYGLEDILDELVASFCKFTTLLNPYTTPEETLFAFSHDMKPRMATLAVFTLANNFGDSIR
GGWRNIVECllklkklqllPNSVVEFEAIDEKTTNNGGSEHDVQIAPTQDSKFNRRQGSSSMGKFSHFLALDSVE
ESIALGVSEFEQNLKVIKQCRIGQIFSKSSILPDVAVLNLGRSLMYAAAGKGQKFSTAIEEEETVKFCWDLIIAI
ALSNIHRFNLFWPSYHEYLLHVVNFPLFSPAPFVENGIPGLFKLCIKILASQLQDHLPEELIFRTITIMWKIDKE
MIDTCYEIITGFVSKIITEYSENLHTQIGWKSVLQLLSLCGRHPETKDQAVDALICLMSTNASHLSQSSYAYCID
CAFSFVALRNSSVEKNLKILDLMTDSVTLLIQWYTSATTEattnnysmasntsstssaeesLRGVNFVHHLFLKL
SEAFRKTTLARREDIRNRAVTSLEKSFIMGHEDLGFTPSGCIYCIDHIIFPTIDDLHEKLLDYSRRENAEREMRS
MEGTLRLGMIMLKNVFLTYLNQIVGSTEFRTFWLGVLRRMDTCMKADLGDYGDSKLREVVPELLTTMIGTMKEKE
ILVQKEDDDLWEITYIQIQWIAPSLKDELFPDEDI*

**>Clevi_0004s1596**

MGRLKLQSGIKAIEEEPEDFESTDSSNTVTLACMIDTEIAAVLAVMRRNVRWGGRYMSGDDQLEHSLIQSLKALR
KQIFSWQQPWHTVSPALYLQPFLDVIRSDETGAAITSVALSSVYKILTLDVIDQNTANIEDAMHLVVDSVTSCRF
EVTDPASEEVVLMKILQVLLACMKNKASVVLSNQHVCTVVNTCFRIVHQAGMKGELLQRVARHTMHQLVRCIFSH
LPDIDRTENALVNRAGSIKQETAGVDGDYAFGSKPTNNGNANSEYDTENSLAISGAPSLMDDGTVGSGSRKDASP
YDLHLMTEPYGVPSMVEIFNFLCSLLNVVEHVGMGPRSNTIAFDEDVPLFALSLINSAIELGGVSIRHHPRLLCL
IQDELFRNLMQFGLSMSPLILSMVCSIVLNLYHHLRTELKLQLEAFFSCVILRLAQGKYGASYQQQEVTMEALVN
FCRQKTFMVEMYANLDCDITCSNVFEDLSNLLSKSAFPVNCPLSAMHILALDGLIAVIQGMAERIGNGLAGLELG
PVHLDEYTPFWMVKCENYSDPNHWVSFVRRRKYIKRRLMIGADHFNRDPKKGLEFLQGTHLLPDKLDPQSVACFF
RYTAGLDKNLVGDFLGNHDEFCVQVLHEFAGTFDFQDMNLDTALRLFLETFRLPGESQKIQRVLEAFSDRYYEQS
PQIFANKDASLLLSYSLIMLNTDQHNVQVKKKMTEEDFIRNNRHINGGSDLPREYLCELYQAICSNEIRTSPEQG
AGYPEMTPSRWIDLMHKSKKTAPFIVADSRAYLDHDMFAIMSGPTIAAISVVFDHAEHEDVYQTCIDGFLAVAKI
SACHHLEDVLDDLVVSLCKFTTLLNPSSVDEPVLAFGDDTKARMATVTVFTIANKYGDYIRTGWRNILDCILRLH
KLGLLPARVASDAADESELSPDPGQGKPVTNSLSSVHLPSMGTPRRSSGLMGRFSQLLSLDTEEPRSQPTEQQLA
AHQRTLQTIQKCHIDSIFTESKFLQAESLLQLARALIWAAGRPQKGNSSPEDEDTAVFCLELLIAITLNNRDRIV
LLWQGVYEHIANIVQSTVMPCALVEKAIFGLLRICQRLLPYKESLADELLRSLQLVLKLDARVADAYCEQIALEV
SRLVKANASHIRSQAGWRTITSLLSITARHPEASEAGFDAISFVMSDGAHLLPANYILCTDAARQFAESRVGQSE
RSIRALDLMGGSVECLAKWARTTKESMGEEESGRMFQDIGEMWLRLVQGLRKVCLDQREDVRNHALQSLQKCLAG
VDGIQIAHNLWSQCFDKVIFTVLDDLLEIAQGSQKDYRNMDGTLLLAVKLLSKVFLQLLQDLAQLTTFCKLWLGV
LTRVEKYMKVKVRGKKSEKLQEAVPDLLKNILLVMKTKGVLVQRSALGGDSLWELTWLHVNNIAPTLRSELFPDH
ETAQYTQPGGLAVDETMPNGVVGTEGGTATG*

**>Clevi_0004s2072**

MTMGHLNLSSGSNSFHGELKYCHSKSLKGALASTINSEIGAVLAVMRRNVRWGVRYIPDDDQLEHSLIHSLKELR
KQIFSWQSNWQGVDPRLYIQPFLDVIRSDETGAPITGVALSSVYKILTLDILTLDTVNVGEAIHLIVDAVKSCRF
EVTDPASEEVVLMKILQVLLACVKSKASCRLSNLDVCTIVNTCLRVVHQSSSKSELLQRIARHTMHELVRCIFSH
VPYISPSEHSLTNGSRSHVGDAVGTVERDQNSLDKLIENGNIGSESDTLGALNNSSSPVAENPETELKNDERSTE

VNDSVNVAEHGENTMMEPFGIPCMIDIFNFLCSLLNVRENSDASSRSNPIAFDEDVPLFALGLINSAVELGGSSF
SKHPKLLRLIQDELFCNLMQFGLSMSPLILSTVCNIVLNLYLNLRAELKVQLEAFFSCVLLRIAQNKHGSSYQQQ
EVAMEALVDFCRQHAFMAEMYANFDCDITCTNVFEDVSNLLSKNAFPVNGPLSAMHILALDGLISMVQGMAERVG
EELATSDASTDQEGYEAFWTVRCENYGDPNFWVPFVRKAKQIKKRLMVGADYFNGDPKKGLQFLQGMQLLPENLE
PQSVAAFFRYTCGLDKNLIGDFLGNHDPFCVEVLHEFAKTFDFRNMNLDTALRLFVGTFRLPGESQKIQRVLEAF
SERYYEQSPQILINKDAALLLSYSLILLNTDHHNVQVKKKMTETDFVRNNRGINGGKDLPLEYLSELYHSICENE
IQMIPDQETGFPRMTASRWISVIHKSKETSPYIVCDTAAHLDRDMFSIVSGPTIAATSVVFDQAEQEDVLQRCVN
GFLAIAKLSAYYHLDSVLDDLVVSLCKFTPLLAPLSVDETVLALVEDARARMATSAVFSIANKYGDYIRSGWKNI
LECVLILEKLQLLPARMASDATEDMEPSSDNLEQERPATNHLPVSQAQSSATPRKSSSLIGRFSQLLSFDTEEIK
PQPTEEELAAHKNAREIVKDCHIDCIFSESKFLQAESLLQLVNSLITAAEKDEDSAVFCLELLITITLNNRDRIL
LIWQTVYEYIANIVQSTSTPCALVEKAVFGLLKICQRLLPYKENLTDELLKSLQLVLKLDARVADAYCEPITQEV
ARLVKANATHIRSHTGWRTIISLLSITARHPEASESGFEALMFIMSEGAHLLPSNYVLCLDAARQFAESRVGEID
RSISAIDLMANSVFCLGRWSREAKNSVGEDAAAKLSQDIGEMWIRVVNSLKRVCLDQRDEVRNHAILMLQRSVAG
ADGFPLPSALWFQCFDSVIFPLLDELLEVSQENARKNFRKAAEKTLVLATKLMSKAFLQSLQDLSQQPSFCRLWL
GVLNRLETYMSAKFRGKRSEKIHELIPELLKNTLLVMKTTGILLPGEDIGSDSFWQLTWLHVKKICPSLLSEVFP
QDELERLQRRNATAGEIPLPEKGAAAVQSSQTPA*
**>Clevi_0068s0067**
MDKAAARSRRKELGISCMLNTEVGAVLAVIRRPLADNYHSPQEADYSDSSVQQSLKSLRALIFNPQQEWRTIDPS
IYLSPFLEVVQSDEIPASATAVAISSILKILKIEIFDEKTPGAKDALNAIVSGITSCRLEKTDLVSEDAVMMRVL
QMLVAIMKHPASNLLEDQVVCTIVNTCFQVVQQSASRGDLLQRNGRYTMHELIQIIFSRLPDFEVRGDEGEDSES
DTDEVDVNGGYGIRCCIDIFHFLCSLLNVVEVVENSEGTTVHTADEDVQIFALVLINSAIELSGDAIGQHPKLLR
MIQDDLFHHLIHYGASSSPLVLSMISSCILNIYHFLRKFVRLQLEAFISFVLLKVTSLSGFLQLQEVALEGIINF
CRQPSFILEAYVNYDCDPICRSLFEEIGKVLCRNTFPTSGPLTSLQIQAFEGLVIVIHNIADNIDKYYDDGDGDG
NGDEDSSSYKPRPVEITEYRPFWIEQPKEDLDVWVDHIRVRKAQKRKLLIAANHFNRDEKKGLEYLKFNNLVSDP
PDPEAFSSFFRFTPGLDKTMIGDYLGDPEEFHIKVLRQFTHTFEFSGMTLDTALRTFLETFRLPGESQKIERIIE
AFSERFYEQQSSEIFATKDTVHILCYSLIMLNTDQHNPQVKKKMTEDEFIRNNRAINAGKDLPREYLSDLFQSIS
SNAIALSSNSFPVEMNPNKWIELINRTKTSQPFSLCEFDRRIGRDMFASIAGPSIAALSSFFEHAEDDEMLHECI
DALISIARVAQYGLEDILDELLASFCKFTTLLNPYATPEETLFTFSHDMKPRIATLAVFTLANNFGDSIRGGWRN
IVDCLLKLKKLKLLPRSIIEFEMADENTSGSVGSESGATPPNQDTKLNRRRSGGLMGRFSHFLSLDNIEESVALG
MSEFEQNLKVIKQCRIGQIFSNSSNLPDAAVLNLGRSLIFAAAGKGQRFSTPIEEEETVIFCWDLIIAITLANVH
RFNMFWPSYHDYLLTVANFPLFSPVPFVEKGIPGIFKVCVKILASQLEDRLPEELIFRSITLMWKMEKEIIDTCC
DVIAQYTSKIVTEYSGNLHTQIGWKSVLQLLSLCGRHPETYEEAVDALICLMSTNASHLSRSSYAYCIDCAFSFV
ALRNTPLEKNLKILDLMTDSVNLLIQWYKTATTEPVNNYSIASNTSTSSMEENLRGGNFAHFLFVKLCEAFKKTS
LARREEIRNRVVTSLQKSFIAAQDLEFTPTGCINCINHVIFPTVDDLHEKLLDYSRRDNAEKEMRSMEGTLKMAM
EMLTNVYLHYLKQISESPDFRTFWLGVLRRMDTCMKADLGDYGENKLQDVVPELLKNMISTMKEREILVKKEDDD
LWEITYIQIQWIAPTLKEELFPEEEV*
**>Tarha_LOC104824072**
MGRLKSQPAIKAIEEEPEDFESTDSSNTVTLACMIDTEIAAVLAVMRRNVRWGGRYMSGDDQLEHSLIQSLKALR
KQIFSWQQLWHTITPVLYLQPFLDVIRSDETGAPITSIALSSVYKILTLDVIDQNTANIEDAMHLVVDSVTSCRF
EVTDPASEEVVLMKILQVLLACMKNKASAVLSNQHVCTVVNTCFRIVHQAGMKGELLQRVARHTMHELVRCIFSH
LPDIDRTENALVNRAGSFDQESAGVDSDYTFGSKPTNIGNANSDYDTENSLATFASGVPGVMDDGTVGAGSRKDA
SPYDLHLMTEPYGVPSMVEIFHFLCSLLNVVEHVGMAPQSNTIAFDEDVPLFALSLINSAIELGGASIRHHPRLL
SLVQDELFRNLMQFGLSMSPLILSMVCSIVLNVYHHLRTELKLQLEAFFSCVILRLAQGKYGASYQQQEVAMEAL
VNFCRQKTFMVEMYANLDCDITCSNVFEDLSNLLSKSAFPVNCPLSAMHLLALDGLIAVIQGMAERIGNGLTGLD
LGPVHLDEYTPFWMVKCDNYSDPNHWVSFVRRRKYIKRRLIIGADHFNRDPKKGLEFLQGTHLLPDKLDPQSVAC
FFRYTAGLDKNLVGDFLGNHDEFCVQVLHEFAGTFDFQDMNLDTALRLFLETFRLPGESQKIQRVLEAFSERYYE
QSPHIFANKDAALLLSYSLIMLNTDQHNVQVKKKMTEEDFIRNNRHINGGSDLPREYLSELYHAICNNEIRTSPE
QCAGYPEMTPSRWIDLMHKSKTTAPYIVANSRAYLDHDMFAIMSGPTIAAISVVFDHVEHEDVYQTCIDGFLAVA
KISACHHLEDVLDDLVVSLCKFTTLLNPSSVDEPVLAFGDDTKARMATVTVFTIANKYGDYIRTGWRNILDCILR
LHKLGLLPARVASDAADESELSSDPGQGKPITNSLSSVHLPSMGTPRRSSGLMGRFSQLLSLDAEEPRSQPTEQQ
LAAHQRTIQTIQKCHIDSVFTESKFLQAESLLQLARALIWMAGRPQKGNGSPEDEDTAVFCLELLIAITLNNRDR
IVLLWQGVYEHIANIVQSTVMPSALVEKAVFGLLRICQRLLPYKESLADELLRSLQLVLKLDARVADAYCEQIAL
EVSRLVKANTSHIRSQAGWRTITSLLSITARHPEASEAGFDAISFVMSSGAHLLPANYILCIDAARQFAESRVGQ
SERSIHALNLMAGSVECLVKWAHTAKESMREDESGRMVQDIGEMWLRLVQGLRKVCLDQREDVRNHALQSLQKCL
AGIGIHGIQIGHSIWSQCFDKVIFTVLDDLLEIAQGSQKDYRNMEGTLLLAVKLLSRVFLLLLQELAQLTTFCKL
WLGILTRMEKYMKVKVRGKKSEKLQESVPDLLKNILLAMKTKGVLVQRSALGGDSLWELTWLHVNNIVPSLRSEL
FPDHDSGLAVDGTVANPSSGLAAS*
**>Tarha_LOC104805178**
MGRLKLQSGINAIEEEPEDFDSSDSSNTVTLACMIDTEIAAVLAVMRRNVRWGSRYMSGDDQLEHSLIQSLKSLR
KQIFSWQQPWHTISPALYLRPFLDVIRSDETGAPITSIALSAVYKILTLDVIDQNTANIEDAMHLVVDSVTSCRF
EVADPASEEVVLMKILQVLLACMKNKASVVLSNQHVCTVVNTCFRIVHQAGXQRVARHTMHELVRCIFSHLPDID
RTENALVNRAGSIKQENAGVDGDYAFASKPTNNGNANSECATENTLATFEYGAPGLMDDGRVGNGSRKDASPYDS
HLMTEPYGVPSMVEIFHFLCSLLNVVEHVGMSPQSNTIAFDEDVPLFALSLINSSIELGGVSIRHHPRLLSLIQD

ELFRNLMQFGLSMSPLILSMVCSIVLNLYHHLRTELKLHLEAFFSCVILRLAQGKYGASYQQQEVAMEALVNFCR
QKTFMVEMYANLDCDITCTNVFEELSNLLSKSAFPVNGPLSAMHILALDGLIAVIQGMAERIGNGLAGLDLSPVH
LDEYTPFWMVKCDNYGDPNHWVSFVRRRKYIKRRLMIGADHFNRDPKKGLEFLQGTYLLPDKLDPQSVACFFRYT
AGLDKNLVGDFLGNHDEFCVQVLHEFAGTFNFQDMNLDTALRLFLETFRLPGESQKIQRVLEAFSERYYELSPQI
FANKDAALLLSYSLIMLNTDQHNVQVKKKMTEEDFIRNNRHINGGSDLPREYLSELYHAICNNEIRTTPEQGAGY
PEMTPSRWIDLMHKSKKTAPYIVADSRAYLDHDMFGIMSGPTIAAISVVFDHTEHEDVYQTCIAGFLAVAKISAC
HNLEDVLDDLVVSLCKFTTLLNPSSVDEPVLAFGDDTKARMATVAVFTIANKYGDYIRTGWRNILDCILRLHKLG
LLPARVASDAADESELSSDPGQGKPITNSLSSVHLPSVGTPRRSSGLMGRFSQLLSLDTEEPRSQPTEQQLAAHQ
RTLQTIQKCHIDSIFTESKFLKAESLLQLARALIWAAGRPQRGNSSPEDEDTAVFCLELLIAITLNNRDRIVLLW
QGVYEHIANIVQSTVMPCALVEKAIFGLLRICQRLLPYKESLADELLRSLQLMLKLDARVADAYCEQIALEVSRL
VKANASHIRSQAGWRTITSLLSITARHPEASEAGFDAISFIMSDGSHLLLANYILCIDAARQFAESRVGQSERSI
CALDLMGGSVECLAKWARTARESMEGNESGRMFQDIGEMWLRLAQGLRKVCLDQREDVRSHALLSLQKCLVGVDG
IQLANNGMWAQCFDKVIFTVLDDLLEIAQGSPKDYWNMEGTMHLAIKLLSKVFLQLLRDLAQLTTFCKIWLGVLT
RMEKYMKVKVRGKKSEKLQEAVPDLLKNILQVMKAKGVLVQRNALGGDSLWELTWLHVNNISPSLRSELCPDHDP
AQLTQPGGGSAVDDAVANPPNGLAGSEGAAAATG*
**>Tarha_LOC104805485**
MGHLNFPSGRNSFKYCHSKSSKGALAFMINSEIGVVLAVMRRNVRWGVRYISDDDQLEHSLIHSLKELRKQIFSW
QNNWQSVDPRLYIQPFLDVIKSDETGAPITGVALSSVHKILTLDVLTLDTVNVGEAIHLIVDAVKSCRFEVTDPA
SEEVVLMKILQVLLACVKSKASCRLSNLDVCTIVNSCLSVVHQSSSKSELLQRIARHTMHELVRCIFSQLPYISP
SENAFANGSRLHVGDAVGTPDWDQTSRDKQVDNGNIGSESDALSTTENSSSPVMENPETEMKKDERSTEVNDPVA
GNGENTMMEPFGIPCMVEIFHFLCSLLNIRENSEISSRSNPIAFDEDVPLFALGLINSAIELGGPSFHEHPKLLR
LIQDELFSNLMQFGLSMSPLILSTVCSIVFNLYLNLRTVLKVQIEAFFSCVLLRIAQNKHGSSYQQQEVAMEALV
DLCRQHAFMAEVYANFDCDITCANVFEDITNLLSKSAFPVNGPLSAMHILALDGLVSMAQGMGERVGEESSAADT
STDQEAYEAFWTVRCENYEDPNFWVPFVRKTKHIKKRLMVGADHFNRDPKKGLLFLQGMHLLPENQDPESVAAFF
RYTCGLDKNLIGDFLGNHDPFCVQVLHEFAKTFDFRNMNLDMALRLFLGTFRLPGESQKIQRVLEAFSERYYEQS
PQILINKDAALLLSYSLIMLNTDQHNVQVKKKMTEADFIRNNRCINGGEDLPKEYLSELYHSICENEIQMVPDQE
TGFPAITASRWISVIHKSKETSPYIVCDTSAHLDHDMFSIVSGPTIAATSVVFDQAEQEDVLQRCVDGFLAIAKL
SACYHLDSVLDDLVASLCKFTPLLAPLSDDETVLALAEDVKARMATAAVFTIANKYGDYIRSGWKNVLECVLSLQ
KLQLLSAHMASDATEDMEPSFSNPELERPSTNLLQATQVQSSSTPRKSSSLIGRFSQLLYFDTEETKPQPTEEEL
AVHKHAREVVKDCRVDSIFSESKFLQAESLQQLVNSLITSAEKDEDSAVFCLELLITITLNNRDRILLIWQTVYE
YIANIVQSTSTPCKLVEKAVFGLLKICQRLLPYKENLTDELLKSLQLVLKLDARVADAYCEPITQEVARLVKGNA
AHIRSHSGWRTIISLLSITARHPEASESGFEALMFIMSEGAHLLPSNYILCLDAARQFAESRVGEISRSISAIDL
MANSVFCLGRWYREAKNSVGEDAAKKLSQDMGEMWIRLVNSLKRVCLDQRDEIRNHTISMLQRSVAGADGFPLPT
NLWFQCFDSVIFPLLDEILELSKENARKKNFKKASEETLVLATKLMSKAFLQSLQDLSKQPSFCRLWLGVLNHLE
TYMSAKFRGKQSEKVHELIPELLKNTLLVMKTTGILLPGEDIGSDSFWQLTWLHVNKICPSLLPEVFPQDELELS
QRQNAMTGEFSLPEKGAVA*
**>Tarha_LOC104823043**
MTMGHLSSPSGSNPFKYCHSKSSKGALASTINSEISAVLAVMRRNVRWGVRYIPDDDQLEHSLVHSLKELRKQIF
SWQSNWQSVDPKLYIQPFLDVIRSDETGAPITGVALSSVYKILTLDILTLDTVNVEEAIHLIVDAVKSCRFEVTD
PASEEVVLMKILQVLLACVKSKASFRLSNLDVCTIVNTCLRVVHQSSSKSELLQRIARHTMHELVRCIFSQLSYI
SPSEHSLANGSRSHVGDTVGTVDWDQNSLDKQVENGNIGSESDTPSATKNSSSAVSEIPETGLRNGERSTEVNDS
VNVDENGENAMMEPFGIPCMVEIFHFLCSLLNIRENNEVSSRSNPIAFDEDVPLFALGLINSAIELGGSSFREHP
KLLRLIQDELFCNLMQFGLSMSPLILSTVCSIFLNLYLNLRTELKVQLEAFFSYVLLRIAQNKYGSSYQQQEVAM
EALVDLCGQRAFMAEMYANFDCDITCINVFEDVTNLLSKNAFPVNGPLSAMHILALDGLISMVQGMAERVGEELL
PAVDASTYQEAYEAFWTVRCANYGDPNFWVPFVRKTKHVKKRLMVGADHFNRDPKKGLQFLQGMHLFPEILDPES
VAAFFRYTCGLDKNLIGDFLGNHDPFCVQVLHEFAKTFDFRHMNLDTALRLFLGTFRLPGESQKIQRVLEAFSER
YYEQGNDERGIVFGFVFDAVVLNTDQHNVQVKKKMTESDFIRNNRCINGGEDLPKEYLSELYHSICENEILMIPD
QETGFSIMTASQWISVIYKSRETSPYIICDASAHLDRDMFSILSGPTVAAASVVFDQAEQEGVLQRCVDGFLAIA
KLSAYYHLDSVLNDLVVSLSKFTPLLAPLSEDETVLALVDYARTRMATAAVFSIAHKYGDYIRSGWKSVLECVLS
LQKLKLLPAHMASDATEDTKPSSSNPEQERPSTMPLPVMQVHSSSTPRRSSSLIGRFSQLLYFDTEETKPQPTEE
ELAAHKQAREIVKDFHIDSIFSESKFLQAESLQQLVNPLITAAEKDGDSAVFCLELLITITLNNRDRILLIWQAV
YEYIANVVQSTSVPSTLVEKAVFGLLKICQRLLPYKENLTDELLKSLQLILKLNARVADAYCETITREVAHLVKA
NATHIRSHIGWRTIISLLSITARHPEASESGFEALVFIMSEGAHLLPSNYILCLDAARQFAESRVGEISRSISAI
DLMANSVFCLGKWSREAKNSVGEDAAEKLNQDIGEMWIRLVNSLKRVCLDQRDEVRNHSILMVQRSVAGADGFPL
PPDLWFRCFDSVIFPLLDELLEISREKAGKTFGKSAEETLVLATKLMSKAFLQSLEYLWKQPSFCKLWLGVLIRL
ETYMSAKFRGKRSEKIHELIPELLKNTLLVMKTSGILLPGEDVGSDSFWQLTWLHVKKTCPSLFSEIFPQDEIEH
LQRQNAKAGEVLLPENGTPA*
**>Tarha_LOC104799071**
MDKAAVKSRRKELGISCMLNTEVGAVLAVIRRPLMDNYLSPQDCDYSDSSVQQSLKSLRALIFNPQQEWRTIDPS
IYLSPFLEVVQSDEIPASATAVALSSILKILKIEIFDEKTPGAKDAMNSIVSGITSCRLEKTDPVSEDAVMMRIL
QVLVGIMKHPASNLLEDQAVCTIVNTCFQVVQQSARRGDLLQRNGRYTMHELIQIIFARLPDFEVRGDEGEDSES
DTDEIDVSGGYGIRCCIDIFHFLCSLLNVVEVVESAEGTTVHTADEDVQIFALVLINSAIELCGDAIGQHPKLLR
MIQDDLFHHLIHYGSSSSPLVLSMICSCILNIYHFLRKFVRLQLEAFFAFVLLRVTSMSGYIQLQEVALEGIINF

CRQPSFIVESYVNYDCDPICRSLFEEIGKVLCRHTFPTSGPLTSLQIQAFEGLVIVIHNIADNIDKEYDDGEGDC
DNDSSSLKPRPVEIKEYTPFWIEKANDDFDAWIDHIRVRKAQKRKLLIASNHFNRDEKKGLEYLKFNNLVSDPPD
PKAFASFFRFTPGLDKTIIGDYLGDPDDFHVKVLKDFTQTFEFTGMSLDTALRTFLETFRLPGESQKIERIIEAF
SERFFQQQSSEIFASKDTVHILCYSLIMLNTDQHNPQVKKKMTEDEFIRNNRAINAGKDLPREYLSELFQSISSN
AIVLSSSSFSVEMNPNKWIELINRTKMLQPFSLCEFDRRIGRDMFASIAGPSIAALSSFFEHADDDEMLHECVDA
LISIARVAQYGLEDILDELVASFCKFTTLLNPYSTPEETLFTFSHDMKPRIATLAVFTLANSFGDSIRGGWRNIV
DCLLKLKKLRLLPQSVVEFDTSGENVSGNGIGSESDGNPSSRDTKLNRRQNACLIGRFSHFLSVDSMEDSVALGM
SEFEQNLKVIKQCRIGQIFSNSSNLPDTAILNLGRSLIFAAAGRGQRFSTPTEEEETVMFCWDLIIAVALANVHR
FNMFWPSYHDYLLTVANFPLFSPVPFVEKGIPGILTVCTKIIASQFEDRLPEELIFRSITLMWKMEKEIIDTCCD
TITEYACKIIMEYSGNLQTQIGWKSVLQLLSLCGRHPETYEQSVDALISLMSTNASHLSRSSYAYCIDCAFSFVT
LRNSPLEKNLKILDLMTDSVSLLIQWYKTASAEPGNSYSVASSGSNSSMEENLRGGGNFAHFLFLKLCEAFRKTS
LARREEIRNRAVASLQKSFVTGEELEFTPTGCLTCINHVIFPTIDDLHEKLLDYSRRDNAEREMRAMEGTLKMAM
QMVTNVYMHYLKLIVESPDFRTFWLGVLRRMDTCMKADLGDYGENKLHEVVPELLKSMISTMKEREILVRKQDDD
LWEITYIQIQWIAPSLKEELFPDEEL*

**>Morol36145S10705**

MGRLKLQPGIKAIEEEPEDFDDTYSNKATLACMINTEVGAVLAVMRRNVRWGGRYMSGDDQLEHSLIQSLKALRK
QVFSWQHQWHTINPTLYLQPFLDVIRSDETGAPITGVALSSVYKILTLDVIDQNTVNVEDAMHLLVDAVTSCRFE
VTDPASEEVVLMKILQVLLACMKSKASVMLSNQHVCTLVNTCFRIVHQAGSKGELLQRVARHTMHELIRCIFSHL
CNMDNTEHALVNRAGAVKQEVGGLDNDYAFGSKPKENGNANSEYDGQHSSASFASNTSMDRGTAMMEDGKIGTGN
GQDAVPYDLHLMTEPYGVPCMVEIFHFLCSLLNVVEHMGMGPQSNTIAFDEDVPLFALGLINSAIELGGPSIRHH
PRLLSLIQDELFRNLMQFGLSMSPLILSMVCSIVLNLYHHLRTELKLQLEAFFACVILRLAQGRYGASYQQQEVA
MEALVDFCRQKTFMVEMYANLDCDITCSNVFEDLANLLSKSAFPVNCPLSAMHILALDGLIAVIQGMAERIGNGS
VSLEQAPVNLEEYTPFWMVKCDNYSDAHYWVPFVRRRKYIKRRLMIGADHFNRDPKKGLEFLQGTHLLPDKLDPQ
SVACFFRFTAGLDKNLVGDFLGNHDEFCVQVLHEFAGTFDFQDMNLDTALRLFLETFRLPGESQKIQRVLEAFSE
RYYEQSPQILANKDAALLLSYSLIMLNTDQHNVQVKKKMTEEDFIRNNRHINGGSDLPREFLSELYHSICKNEIR
TTPEQGAGFPEMTPSRWIDLMHKSKKTAPYIVANSRAYLDHDMFAIMSGPTIAAISVVFDHAEHEDVYQTCIDGF
LAVAKISACHHLEDVLDDLVVSLCKFTTLLNPSSVEEPVLAFGDDTKARMATVTVFTIANRYGDYIRTGWRNILD
CILRLHKLGLLPARVASDAADESELSAEPGHGKPITNSLSAAHMPSMGTPRRSSGLMGRFSQLLSLDTEEPRSQP
TEQQLAAHQRTLQTIQKCHIDSIFTESKFLQAESLLQLARALIWAAGRPQKGSSSPEDEDTAVFCLELLIAITLN
NRDRIVLLWQGVYEHIANIVQSTMMPCALVEKAVFGLLRICQRLLPYKENLADELLRSLQLVLKLDARVADAYCE
QITLEVSRLVKANASHIKSQMGWRTITSLLSITARHPEASEAGFDALLFIMSDGAHLLPANYILCADAARQFAES
RVGQAERSVRALDLMAGSVDCLERWARDTKEAMGEEEAAKMSQDIGEMWLRLVQGLRKVCLDQREDVRNHAVLSL
QKCLAGIDGIQLAHGLWLQCFDLVIFTMLDDLLEIAQGHSQKDYRNMEGSLILAVKLLSKVFLQLLNDLSQLTTF
CKLWLGVLSRMEKYMKVKVRGKKSEKLQETVPELLKGILLVMKTRGVLVQRSALGGDSLWELTWLHVNNIAPSLQ
VEVFPDQDLGHLQLKQAETGGGLVSDGTVSAPSNEAVASEGAGAAG*

**>Morol33726S06189**

MGHLNSASGSNSFHEELKHYPAKLSGGALACMVNSEIGAVLAVMRRNVRWGVRYMPDDDQLEHSLIHALKALRKQ
IFSWQNQWHHINPAVYLQPFLDVIQSDETGAPITGVALSSVYRILTLDVLNLDTVNVGDAMHLIVNAVTSCRFEV
TDPGSEEVVLMKILQVLLACMKSKAAVRLSNQHVCNIVNTCFRVVHQASSKGELLQRIARHTLHELVRCIFSHLP
DLENSEHTFSNKRSSYIRETVDAVDGDHQSSGNQQENGNVSVQYDVQPSTAYASSAVIGGLKAENEKDINNIVVI
DGMEATENGQRAMTEPFGIPCMVEIFHFLCSLLNVIEHIGIGPRSNPVAYDEDVPLFALGLINSAIELGGPSFSK
HPKLLGLIQDELFRNLMQFGLSMSPLILSTVCSIVLNLYHHLRTELKVQLEAFFSCVLCGLAQSKHGSSYQQQEV
AMEALVDFCRQQTFMAEMYANYDSDITCSNVFEDLANLLSKNAFPVNGPLSAMHILALDGLTSMIQGMAERIGSE
LSASDDTSLDGEYEAFWTLRCENYTNPSCWVPFVRKSKYIKRKLMVGADHFNRDPKKGLEFLQGMHLLPDKLDPQ
SVAAFFRYTSGLDKNLIGDFLGNHDEFCVQVLYEFARTFDFRDMNLDTALRLFLGTFRLPGESQKIQRVLEAFAE
RYYEHSPQILFNKDAALLLSYSLILLNTDQHNVQVKKKMTEEDFIRNNRHINGGKDLPREFLTELYHSICENEIQ
MIPDQGGGFSVMTSGRWINVMHKSKETTPYIVCDSKALLDRDMFAILSGPTIAAISVVFDQVEREDILQTCIDGL
LAIAKISAFYHFDDVLDDLVVSLCKFTTLLTPLSVEEALVTFGEDTRARIATTAVFTIANKYGDYIRSGWENILD
CVLSLHKLGFLPAYMASDAAEDMEHSSEVEQGKPNPLSTSKTHSVATPRKSSSLIGRFSQLLSFDMEETRLQPTE
EQLAAHQHTREIIQSCHIDSIFMESKFLHAESLLQLVRALILAAGRLGKVSGSTEGEDTAVLCLELLIAITLNNR
DRILLIWQGVYEHISNIVQSTAMPCALVEKAVFGLLRICERLLPYKENLTDELLKSLQLILKLDARVADAYCEPI
TQEVMRLVKANATHIRSHAGWRTIISLLSITARHPEASEAGFEALAFIMFEGAHLLPSNYILCVDAARQFAESRV
GEIDRSVSAIDMMASSVGCLVRWSSETKNAVEEEAAMKVRQDIGEMWLRLVQGLRKVCLDQREEVRNHAILMLQR
SFAGVDGIHLPNAIWFQSFDMVIFTLLDDLLEIAQGNSPKDYRKMEGTLVLATKFMSKVFLQSLQDLSGQPSFCK
LWLGVLNRMERLMKIKFRGKRSEKIHELIPELLKNTLLVMKTTGILMPSDEIGGDSFWQLTWLHVNKICPSLQSD
VFPQRELEQMQAKHIKTGGIPVSDGAVLVPSSETTA*

**>Morol36117S09591**

MKNPSDQNQGEESNCHNCTSCRQRKKDTAKSKRKEVGLSCMLNTEVGAVLAVIRRPQDPTTQYLSPQDDCCDSAV
QQSLKSLRALIFNPQQEWRTIDPSIYLSPFLDVTQSDDIPAVGTAVALSAILKIIKLEIFDEKTPGAKDAMNSIV
TGITGCRLEKTDPVSEDAVMMKILQVLTGIMRHPASVLLTDQAVCTIINTCFQVVQQSASRGDLLQRSARYTMHD
LIQIIFSRLPEFEVKEAESESDTDDADLDSGYGVRCAVDIFHFLCSLLNVVEIVETEGTASRTADEDVQLFALVL
INAAIELSGDDITKHSKLLRMIQDDLFHHLIHYGIGSSPLVLSMICSTVLNVYHFLRRFIRLQLEAFFSFVLLRY
VAFGSSVQLQEVGIEGMINFCRQPTFIFEVYVNYDCDPICRSLFEEIGKLLCKHSFPASGPLTSLQIQSFEGLLI

MIHNIADHIDREIDSGSSGPDLVEISESRPFWLEKPKDEAGTWVEYTREKKLQKRKILIAANHFNRDEKKGLEYL
KLSGLISEPPDPKAYAFFFRYTPSLDKAKIGDYLGDPDEFNIQVLQEFTQTFEFTGMTLDAAFRTYLETFRLPGE
SQKIHRVVQAFSERFYEQQSSEIFATKDALFIFCYSLIMLNTDQHNPQVKRKMTEEEFIRNNRGINGGRDFPREY
LSDIFHSIAANAFTMSGHSGPVEMNPNRWIELMNRSKIMQPFNLCDFDHRLGRDMFACIAGPSVAALSAFFDHAD
EDEILHECIEGLISVARIAQCGLEDTLDELIASFCKFTTLLNPYATPEETLFSFSHDMKPRMATLAVFTIANNFG
DSIRGGWRNIVDCLLKLKKLKLLPQTVIEFDAASTSTSDRPPHSRSESGVIFPTHDAKFGKRLNSGIVSRFSHFL
SLESMEDSVSLGINEFEQNLKVIKQCQVGNIFRNSSCLPDESFLNLGRSLIFAAGGKGQKFSTPVEEEDTIGFCW
DLIVSISLSNIQRFQSFWPAFHDYLLGVAQFPLFSTIPFTEKAILGLFKICLKLLASYQAERLPEELIFKSINLM
WKLDKEILDTCCESITQSVSKVLIEYPANLQSQLGWKSVLHLLSICGRHPETYDQAVETLIMLLSDSSHVSRINY
SYCIDCAFGFVALKNSPLDKNLQILELMANSVNLLIQWHKNACSEPGNNYSVASNTSNSSLEECSKGTNFAMTLF
VKLGEALRKTSLARREEIRNKAVLALQKSFTIAEELDFTSTNCINFFNLVIFAMVDDLHEKMLEYSRRENAEREM
RSMEGTLKIAMDLLADVYMQYLKQIAGSSEFRTFWLGVLRRMDTCMKADLGPYGETKLQEVVPDFLRNMIRTMKE
REILVQKEGEDLWEITYIQIQWIAPSLKEELFPEEEM*
**>Carpa_LOC110823971**
MGRLKLQSGIKAIEEEPEDFDDTNSNRATLACMINTEVGAVLAVMRRNVRWGGRYMSGDDQLEHSLIQSLKVLRK
QIFSWQHQWHAINPALYLQPFLDVIRSDETGAPITGVALSSVYKILTLDVIDQNTVNVEAAMHLVVDAVTSCRFE
VTDPASEEVVLMKILQVLLACMKSKASAMLSNQHVCTIVNTCFRIVHQAGSKGELLQRLARHTMHELVRCIFLHL
ADIDNTECALVNRTTTVKQEIGGIDNDYAFGSKQIENGNANSEYDGQNSSANFASNTSLSLAATMVEDGTIGAGS
GQEAVSFDLHLMTEPYGVPCMVEIFHFLCSLLNVVEHMGMGPRSNTVAFDEDVPLFALGLINSAIELGGPSIRHH
PRLLNLIQDELFHNLMQFGLSMSPLILSMVCSIGPTIAAISVVFDHAEHEDVYQTCIDGFLAVAKISACHHLEDV
LDDLVVSLCKFTTLLNPSSVEEPVLAFGDDTKARMATVTVFTIANRYGDYIRTGWRNILDCILRLHKLGLLPARV
ASDAADESELSADIGHGKPITNSLSAAHMPSMGTPRRSSGLMGRFSQLLSLDTEEPRSQPTEQQLAAHQRTLQTI
QKCHIDSIFTESKFLQAESLLQLARALIWAAGRPQKGSSTPEDEDTAVFCLELLIAITLNNRDRIVLLWQGVYEH
IANIVQSTVMPCALVEKAVFGLLRICQRLLPYKENLADELLRSLQLVSKLDARVADAYCEQITLEVSRLVKANAA
HIRSQMGWRTITSLLSFTARHPEASEAGFDALLFIMLDGAHLLPANYILCTDAARQFAESRVGQAERSVRALDLM
AGSVNSLERWACETKETMGEEEVVKMSQDIGEMWLRLVQGLRKVCLDQREEVRNHALLSLQKCLTGVDGIQLAHG
LWLQCFDLVIFTVLDDLLEIAQGHSQKDYRNMEGTLTLAVKLLSKVFLQLLPELSQLTTFCKLWLGVLSRMEKYM
KVKVRGKKSEKLQETVPELLKNILLVMKTKGVLVQRSALGGDSLWELTWLHVNNIAPSLQAEVFPDQDMGQSQVK
LGETAGSLVSDSGHSSETVASEGAGSAG*
**>Carpa_LOC110820219**
MVHLNPHTGNDPFCGEFKGCPTKSSRGALACMVNSEIGAVLAVMRRNVRWGVHYMPDDDQLEHSLIHSLKELRKH
IFSWQNQWHNIDPAVYLQPFLDVIQSDETGAPITGVALSSVNKILTLDVLTLDTVNVGDAMQLIVDAVTSCRFEV
TDPASEEVVLMKILQVLLACMKSRAAAKLSNQHVCTIVNTCFRIVHQASSKGELLQRIARHTMHELVRCIFSLLP
DINISEYSLTNGSSSYVGDMVDKQLENGNVSIELDVQPSVGYASSALSGSMKPENKMDTNNICVSDETEATEKTE
KVMMEPFGVPCMVEIFHFLCSLLNFIDHVGIGPRSNPIAYDEDVPLFALGLINSAVELGGPSFSKHPKLLGLIQD
ELFRNLMQFGLSMSPLILSTVYSIVHNLYHHLRTELKVQLEAFFSCVLLKIAQSKHGYSYQQQEVALEAMVDLCR
QQTFVAEMYANFDCDITCSNVFEDLANLISKSAFPLNGPLSAMQILALDGLISMIQGIAERIGNELPASKEAVVD
NERYEAFWTLRCENYSDPNYWVPYVRKTKYIKRRLMVGADHFNRDPKKGLEFLQGMHLLPDKLDPQSVASFFRYT
SGLDKNLIGDFLGNHDEFCILVLHEFAKTFDFCDMNLDAALRLFLGTFRLPGESQKIQRVLEAFAERYYEQSPQI
LFNKDAALLLSYSLILLNTDQHNVQVKKKMTEEDFIRNNRHINGGKDLPRQFLSELYHSICDNEIQMIPDQGAGF
PVMTSGRWINILHKLKETTPYIVCDSRAFLDHDMFAILSGPTIAAISVVFDQVEQEDVLRTCVGGLLAIAKISAC
YHFGNVLDDLVVSLCNFTTLLTPLSVEEAILAFEEDTKASMATTAVFTIANKYGDYIHSGWKNVLDCVLSLHKLG
FLPXRMASDAAEDVEPSYEQEQRKPSTNPLKSSNVPSTAIPRKSSGIIGRFSQLLSFDMEETKLQPSEEQIAAHQ
RFREMIQNCHVDAIFTESKFLRAESLLQLVRALTLAAGRLRKGSSTMEEEDNCVFCLELLIAITLNNRDRIMLIW
QGVHEHMSNIVQSILMPCTLVERAVFGLLRICQRLLPYKENLIDELLKSLQLILKLDARVADAYCEPITQEVMRL
VKANAAHIRSHVGWRTIISLLSITARHPEASEAGFETLVFIMSEGAYLLPSNYILCLDAARQFAESRVGEVGRSI
SAIDMMAGSVVCLVRWSHETKSSVVDEAAVKVNQSIGEMWLRLVQGLRKLCLDQREEVRNHAILMLQRSLAVVDG
IHIPNSLWFQCFDMVIFTLLDDLLEIAQGNFPKDYRKMEGTLVLATKLMSKTFLQSLQ*
**>Carpa_LOC110818636**
MKKPRDRNREEEEEGGNQNGCISCCGRKKDAAKSKRKELGLSCMLNTEVGAVLAVIRRPQDPNTQYISLHDDHYD
SAVQQSLKSLRALIFNPQQEWRTIDPSIYLSPFLDVIQSDDIPAAATGISLSAILKIVKFEIFDEKTPGAKDAMN
SIVTGITGCRLEKTNQVSEESVMMKILQVLTGIMKHPASDLLTDQAVCTIVNTCFQVVQQSAGRGDLLQRNARFT
MHELIQIIFSRLPDFEVNDDNSESDTEDADGNALDSGYGLRSAVDIFHFLCSLLNVVELVETEGTSYHTADEDVQ
LFALVLINSAIELSGDGIGKRPKLLRMVQDDLFHHLIHYGTSSSPLVLSMICSTVLNIYHFLRRFVRLQLEAFFS
FVLLKNVALGSSMQLQEVGLEGMINFCRQSTFIIEAYANYDCDPPCRSLFEEVGKLLCKHSFPGSGPMTSIQIQA
FEGLVILIHNIADHVDKEDDSTPSGPYPVEITEYKPFWMEMPKEDLEEWVEHIRLRKAQKRKILIASNHFNRDEK
KGLEYLMLSQLVSDPPDPKCFAFFFRHTPGLDKMMIGEYLGNPDDFHIQVLREYTQTFEFSGMVLDSALRRYLET
FRLPGESQKIQRILEAFSDRFYDQQSLEMFASKDTVLILCYSLIMLNTDQHNPQVKRKMTEEEFIKNNRAINGGK
DLPREYLSELFHSISTHEITVFGQSGSVEMSPSRWIELMNRSKKVQPFILCDFDRRIGRDMFACIAGPSIAALSA
FFEHADEDELLHECIDGLISMARIAQYGLEDTLDELVASFCKFTTLLNPYATAEETLFAFSHDMKPRMATLAVFT
IANHFGDSIRGGWRNIVDCLLKLKRLKILPQSVTEFDITSSSADAPGHSKSESSVMSSNHDHKCSKRPGTSILSR
FSHFVSLESIEESVSLGMSEYEQNQKIIKQCQIGNIFSNSSSLPDESLLNLGRSMIFAAGGKGQKFSTPVEEEET
VGFCWDLIVAIALANIHRFQAFWPYFHDFLLGVAQFPLFSPIPFAEKAIVGLFKVCLRLLSSCQVERIPEENLFK

SINLMWKLDKEILDTCCEFIIQSVSNIINEYPANLQTQLGWKSVLHLLSVCGRHPETYDQSVEALIKLMSDATHV
SRTNYAYCIDCAFGFVALKNSPLEKNLKILDLLSNSVTLLIQWYKNACSEPGNSYSTTSSASVSSLEDYSKGPNF
GLNLFIKLGEALRKTSLARREEIRNRAIIALRKSFLLAEELELTSTNCINCFNLVIFAMVDDLHEKMLEYSRREN
SEREMRSMEGTLRIAMELLKDVYMQYLKQIAENPDFRTFWLGVLRRMDTCMKADLGTYGETLLKEVVPDLLRNMT
AKMKEEEILVPNEGDDLWEITYIQIQWIAPSLKEEIFPEDEM*

**>Theca_LOC18610338**

MGRLKLQSGIKAIEEEPEDCDTTCSSKATLACMINSEVGAVLAVMRRNVRWGGRYMSGDDQLEHSLIQSLKALRK
QIFLWQLQWHTINPAVYLQPFLDVIRSDETGAPITGVALSSLHKILTLDVIDQNTVNVEDAMRLVVDAVTSCRFE
VTDPASEEVVLMKILQVLLACMKSKASVMLSNQHVCTIVNTCFRIVHQAEKKGELLQRIARHTMHELVRCIFSHL
SNVDNTEHALVNRTGTAKQELGGIDNDYAFGAKKVENGNGTEYDGQASSGSFASNGSAGLVATAREESMVVAGNG
KATVPYDLHLMTELYGVPCMVEIFHFLCSLLNAAEHVGMGPRSNTLAFDEDVPLFALGLINSAIELGGPSFRRHP
RLLSLIQDELFRNLMQFGLSMSPLILSMVCSIVLNLYHHLRTELKLQLEAFFSCVILRLAQGKYGASYQQQEVAM
EALVDFCRQKTFMVEMYANLDCDITCSNVFEDLANLLSKSAFPVNCPLSAMHILALDGLIAVIQGMAERIGNGSV
SSEYAPVSLEEYTPFWMVKCDSYGDPSHWVPFVRRRKYIKRRLMIGADHFNRDPKKGLEFLQGTHLLPDKLDPQS
VACFFRYTAGLDKNLVGDFLGNHDDFCVQVLHEFAGTFDFQDMNLDTALRLFLETFRLPGESQKIQRVLEAFSER
YYEQSPQILVNKDAALLLSYSLIMLNTDQHNVQVKKKMTEEDFIRNNRHINGGNDLPREFLSELYHSICKNEIRT
TPEQGFGYPEMTPSRWIDLMHKSKKTAPFIIADSRAYLDHDMFAIMSGPTIAAISVVFDHAEHEDVYQTCIDGFL
AVAKISACHHLEDVLDDLVVSLCKFTTLLNPSSVEEPVLAFGDDTKARMATVTVFTIANRYGDYIRTGWRNILDC
ILRLHKLGLLPARVASDAADESELSADPSHGKPITNSLSSAHIQSIGTPRRSSGLMGRFSQLLSLETEEPRSQPT
EQQLAAHQRTLQTIQKCHIDSIFTESKFLQAESLLQLARALIWAAGRPQKGNSSPEDEDTAVFCLELLIAITLNN
RDRIVLLWQGVYEHIANIVQSTVMPCALVEKAVFGLLRICQRLLPYKENLADELLRSLQLVLKLDARVADAYCEQ
ITQEVSRLVKANATHIRSQMGWRTITSLLSITARHPEASEAGFDALLFIMSDGAHLLPANYGLCVDAARQFAESR
VGQAERSVRALDLMSGSVDCLARWANEAKEAMGEEDLAKMFQDIGDLWLRLVQGLRKVCLDQREEVRNHALLSLQ
KCLTAVDGIHISHGLWLQCFDLVIFTMLDDVLEIAQGHQKDYRNMEGTLILAMKLLSKVFLQLLYELSQLTTFCK
LWLGVLSRMEKYMKVKIRGKKSEKLQELVLELLKHMLLVMKTRGVLMQRSALGGDSLWELTWLHVNNIAPSMQSE
VFPDQDLEQSLPKHGETGGVVSGEMASVPSNETAAPEGASAGG*

**>Theca_LOC18601571**

MGDLTTQSGNNGFEGELVKPSRGAFACMINSEIGAVLAVMRRNVRWGVRYVADDDQLEHSLIHSLKELRKQIFSW
QHQWQNVDPVVYLQPFLDVIRSDETGAPITGVALSSIYKILTLDVLDLDTVNVGDAMHLIVDAVTSCRFEVTDPA
SEEVVLVKILQVLLACMKSKAATRLSNRHVCMIANTCFRIVHQATSKGELLQRIARHTMHELVRCIFSHLPEISS
PEHEMANGSSSSASEVVTQNSNHMLGSTLLENGNVGLDCDGPSSISDAFSPLVVNSATETDTSKIGESDGKEDAQ
HGEILMAAPFGIPCMVEIFQFLCSLLNVIEHIGIGPKSNPIAYDEDVPLFALGLINSAIELGGPSFSKHPKLLAL
IQDELFRNLMQFGLSMSPLILSTVCSIVLNLYYHLRTELKVQLEAFFSCVLLRLAQSKHGSSYQQQEVAMEALVD
LCRQQTFVAEMYANFDCDITCSNVFEDLANLLSRSAFPVNGPLSAMHILALDGLVSMIKGMAERIGNELPVSDEA
STDHEGYEAFWTLKCQNYSDASSWIPFVRKMKYIKRKLTIGADHFNRDPKKGLEFLQGIHLLPDKLDPESVASFF
RYTTGLDKNLIGDFLGNHDEFCVQVLHEFARTFDFHDMNLDSALRVFLGTFRLPGESQKIQRVLEAFSERYYEQS
PHILCNKDAALLLSYSLILLNTDQHNVQVKKKMTEEDFIRNNRLINGGKDLPREYLSELYHSICENEIQMIPDQG
AGFPVMTSSRWINIVHKSKESSPFIACDSRALLDHDMFAILSGPTIAAISVVFDQVEQEDVLQTCIDGFLAIAKI
SAHYHFGDVLDDLVVSLCKFTNLLTPLSVEEAIVSFGDDARARMATSAVFTIANSYGDYIHSGWRNILDCVLILQ
KLGLLPAHLASDAADEMESLSDHERGKPATSSLSTSQTVRAPTPQKSSSLIGRFSQLLSFDMEEPRLQPTEEQLA
TQQNLREIVEKCHIDNIFMESKFLQSESLLQLVGAIVFAAGRFSKGSGIVEDEDTAVFCLDLLIAITLNNRDRIM
IIWQNVYKHIADIVQKKSTPCTLVEKAVFGLLKICQRLLPYKENLTDELLKSMQLILKLDARVADAYCEPITQEV
MRLVKANATHIRSHLGWRTIISLLFITARHPEASEFGFEALAFIMFEGAHLMPSNYVLCVDAAREFAESRVGEVD
RSIHSLDMMAGSVVCLVRWSYETKNAAEEAAIQVSQDIGEMWLRLVQGLRALCLDQREEVRNHAILMLQRSLAGV
DGIHLPNAMWFQCFDLVIFTLLDDLQEIAEGSSSKEYRKMEGTLVLATKLMSKAFLQLLQDLSQQPSFCKLWLGV
LNRMERYMKVKFRGKCSEKIHELVPELLKNTLLVMKTTGILMPSDDIGGDSFWQLTWLHVKNIVPSLQSEVFSEQ
ELEQMQAKHKNKAEIPETDGVVLVPSKETTA*

**>Theca_LOC18588627**

MEEEQNRCPSCGEKKKDTGKSRRKELGLSCMLNTEVGAVLAVIRRPIDPTAQFFSPQEEHYDSSIQQSLKSLRGL
IFNPHQEWRTTDPCIYLSPFLDVIQSDDIPAAATGVALSAILKILKLEIFDEKTPGAKEAINFVVSGITSCRLEK
TDPVSEDAVMMKILQVLTGILRHDASVLLTDQAVCTIVNTCFQVVQQSVTRGDLLQRSARYTMHELIQIIFSRLP
DAGDNEGENSESETEDIDENPGYGTRCAVDIFQFLCSLLNVVEVVENDGSTWHTVDEDVQLFALVLINCAIELSG
DGIGKHPKLLRMIQDDLFHHLIHYGTCSSPLVLSMICSTVLNIYHFLRRFIRLQLEAFFSYVLLKVAALGASLQL
QEVALEGIINFFRQPTFVIEAYVNYDCDPICRNLFEEVVKLLCKHAFPGTGPMTTLQVQAFEGLVIMIHNISDNI
DKEDDSSTSEPYPIEITEYRPFWVDKPKDDLATWVEYIRVRKAQKKKILIAGNHYNRDEKKGLEYLKHCQLVSDP
PNPKAFAFFFRFTPGLDKNMIGEYLGDPDEFHLEVLKEFTGAFEFKGMILDSALRTYLETFRLPGESQKIQRILE
VFSERFFDQQSSDIFVAKDSVFVLCYSLIMLNTDQHNPQVKKKMTEEEFIRNNRLINGGQDLPREYLSELFDSIS
NHAITLFGQSGAVEMNPSRWVELMNRAKLMQSYVLCDFDRRLGRDMFACVAGPTIAALSAFFEHADEDEMLHECI
EGLISVATIAQFGLADTLDELVASFCRFTTLLNPYASAEETLFAFSNDMKPRMATLAVFTIANNFGESIRGGWRN
IVDCLLKLKRIKLLPQSVIEFDVASTSSSDATEAFKSESGVIFPNHDPKFSKRQTSGMVSRFSHFLSIDSMEESI
SLGMSEFEQNLKIIKQCRIGSIFGNSSNLPIEALLNLGRSLIFAAAGKGQKFSTPIEEEETAGFCWDLIIAISLS
NIHRFQVYWPSFHDYLLAVAQLPLFSPIPFAEKAMVGLFKVCLKLLASYQVDKIPEELIFKSINLMLDKEVLDMC
CEYIIQSVSKILIEYPANLQTQLGWKSTLHLLQVAGRHPETYDQAVETFIMLMSDAFHISRINYAFCIDCAFGFI

ALRNSPVEKNLKILDLMSDSVNWLIQWYKTAHSDPGSSYSVASNTSTSSWEDNSKAIGSSNFTVNLFIKLGEALR
KSSLARREEIRNCAVASLKRGFQLAEELEFSSTSCINCFNLVIFAMVDDQHEKMIEYSRRENAEREMRSMEGTLK
ISMELLTDVYLQYLKVIAENPGFRTFWLGILRRMDTCMKADLGEYGETNLQEVVPDLLRKMIRNMQEKEILVPKD
GDDLWEITYIQIQWIAPSLKEELFPDEL*
**>Gosra_LOC105779596**
MGRLKLQSGIKAIEEEPEDYDMTYSNKATLACMINSEIGAVLAVMRRNVRWGGRYMSSDDQLEHSLIQSLKALRK
QIFLWQNQWHTINPAAYLQPFLDVIQSDETGAPITGVALSSVHKILTLDVIDQNTANVEEAMRLVVDSVTSCRFE
VTDQASEEVVLMKILHVLLACMKSKASVMLSNQHVCTIVNTCFRIVHQAEKKAELLQRIARHTMHELVRCIFSHL
LNVENTKHALVNRSGTAKLELGAIDNDYAFGAKQAENGNGSEYDDQASSGSFASNSVGLVGTVREESMAVAGNGK
ETVPYDSLLMTEPYGVPCMVEIFHFLCTLLNTVEHVGMDPRSNTLTFDEDVPLFALGLINSAIELGGPSFRRHPR
LLSLIQDELFRNLMQFGLSMSPLILSMVCSIVLNLYHHLRTELKLQLEAFFSCVILRLAQGKYGASYQQQEVAME
ALVDFCRQKTFMVEMYANLDCDITCRNVFEDLANLLSKSAFPVNCPLSAMHILALDGLIAVIQGMAERIGNGSVS
SEQAPVILEEYLPFWMVKCDNYADPVHWVPFVRRRKYIKRRLMIGADHFNRDPKKGLEFLQGTHLLPDKLDPQSV
ACFFRYTAGLDKNLVGDFLGNHDEFCVQVLHEFAGTFDFQDMNLDTALRLFLETFRLPGESQKIQRVLEAFSERY
YEQSPQILANKDAALLLSYSLIMLNTDQHNVQVKKKMTEEDFIRNNRHINGGNDLPRDFLSELYRSICKNEIRTT
PEQGFGYPEMTPSRWIDLMHKSKKTAPFIVADSRAYLDHDMFAIMSGPTIAAISVVFDHAEHEHVYQTCIDGFLA
VAKISACHHLEDVLDDLVVSLCKFTTLLNPSSVEEPVLAFGDDAKARMATVTVFTIANRYGDYIRTGWRNILDCI
LRLHKLGLLPARVASDAADESELSADPGHGKPITNSLSSAHLQSIGTPRRSSGLMGRFSQLLSLDTEEPRSQPTE
QQLAAHQRTLQTIQKCHIDSIFTESKFLQAESLLQLARALIWAAGRPQKGSSSPEDEDTAVFCLELLIAITLNNR
DRIVLLWQGVYEHIANIVQSTVMPCALVEKAVFGLLRICQRLLPYKENLADELLRSLQLVLKLDARVADAYCEQI
TQEVSRLVKANATHIRSQMGWRTITSLLSITARHPEASEAGFDALLFIMSDGAHLLPANYVLCIDAARQFAESRV
GQAERSVRALDLMSGSVDCLARWTREAKEAMGEDDAGKMSQDIGDLWLRLVQGLRKVCLDQREEVRNHALLSLQK
CLTGVDGIHLSHGLWLQCFDLVIFTMLDDLLEIAQGHQKDYRNMEGTLILATKLLSKVFLQLLHELSQLTTFCKL
WLGVLSRMEKYMKVKVRGKKSEKLQELVLELLKNILLVMKTRGILMQRSALGGDSLWELTWLHVNNIVPSLQSEV
FPDQGPELKHGETGCAVSGETVSVPSNETTTLEGAGAGS*
**>Gosra_LOC105798157**
MGDLITQSQKNGFNGELVKPCKGAFACMINSEIGAVLAVMRRNVRWGVRYVSDDDQLEHSLIRSLKELRKQIFSW
QHQWQNVDPVVYLLPFLDVIKSDETGAPITGVALSSIYKILTLDVLDLDTVNVEEAMHLIVDAVTSCRFEVTDPA
SEEVVLMKILQVLLACMKSKAAPRLSNKHVCMIANTCFRIVHQASSKGELLQRIARHTMHELIRCIFSHLPEIGS
TEHEMSNGRSLYASGVQVGAQNGNHVVGSTMVENGNVGISCDGPSSLSDASSPPIVKSGPETDSSNFGESDGTED
AQRADVLMSAPFGIPCMVEIFQFLCSLLNVIENIGIGPKSNPVAYDEDVPLFALGLINSAIELGGPSFGKHPKLL
AIIQDDLFHNLMQFGLSMSPLILSTVCSIVLNLYHHLRWELKVQLEAFFTYVLLKLVQSKHGSSYQQQEVAMEAL
VDLCRQQTFVAEMYANYDCDISCSNVFEDIANMLSKSAFPVNGPLSAMHILALDGLVSTVKGMTERIGYELPIPN
ESSTDHEGYEAFWTSKCQNYSDPSSWIPFVRKMKHLKRKLAIGADHFNRDPKKGLEFLQGIHLLPEKLDPESVAS
FFRYTVGLDKNLIGDFLGNHDEFCVQVLHEFVKTFNFKDMNLDSALRVFLGTFRLPGESQKIQRVLEAFSERYYE
QSPHILCNKDAALVLSYSLILLNTDQHNVQVKKKMTEEDFIRNNRLINGGKDLPREYLSELFHSICENEIQMIPD
QDAGVPVMTSSRWINVVNKSKGSTPFIACDSRALLDHDMFTILSGPTIAALSVVFDQVEQEDVLQMCIDGFLAIA
KLSAHYHFGDVLDDLVVSLCKFTNLSTPLSIEEAIVSFGDDSRARMATKTVFTIANSYGDCIHSGWTNILDCVLS
LHKLNLLPARLASDTADETESSSDHERGKSATSSLRRLKTAPAAIPRKSSSLIGRFSQFLSFDIEEPTLYPTEEE
LAAQRRLREIVQECHVDNIFMESKFLQAESLLQLVEAIILAASRFSKGTGVMEDEDTAVFCLELLIVITLNNRDR
ILIIWQKVYKHISDIVQQKAVPCALVEKAVFGLLKICQRLLPYKENITDELLKSMQLILKLDARVADAYCEPITQ
EVMHLVKANATHIRSHLGWRTIISLLYITARHPEASEFGFEALAFIMSEGAHLLPSNYVLCVDAAREFAESRVAE
VDRSISALDMMAGSVVCLVRWSYETKNAVEEEAAMKVSRDIGEMWLRLVQGLRAVCLDQREEVRNHAILMLQSSL
GGVDGIQIPNAMWFQCFDHVIFTLMDDLLEIAEESSPKEYRKIEGTLVLATKLLSKAFLQSLQDLSQQPSFCKLW
LGVLDRMERYVKVKFRGKRSEKIHELVPELLKNTLLVMKTTGILVPSDDLGGESFWQLTWLHVKNIVPSLQSEVF
SEQELEQVQGKHQKKTGIPSTDGAVLVPSNETAA*
**>Gosra_LOC105780543**
MGDVTTQSRNNGFIGEIIKPSRGAFTCMINSEIVAVLAVMRRNVRWGVHYVADDDHLEHSLIHSLKELRRKIFSW
QHQWQNVDPVVYLQPFLDVIQSDETGAPITGVALSSIYKILTIDVLDLDIVNVRVAMHMIIDAVTSCRFEVTDPA
SEEVVLMKILQVLLACMKSKAATRLSNKHVCMIANTCFRIVHQASSKGELLQRIARHTIHELVRCIFSHLPKICS
TEHEMVNGSSLVASGVVGTQNENHMNGSTQLKSGNVGIDESENKEDAQNGEILIMAPFGIPCMVEIFQFLCSLLK
VTENIDIGPKSNMIVHDEDVPLFALGLINSAIELGGSLFSKHPKLLAVIQDELFYNLMQFGLSMSPLILSTVCSI
VLNLYYHLRTELKVQLEAFISCVLLKLVQSKHGSSYQQQEVAMEALVDLCSQQMFVAEMYANFDCDITYSNVFED
IAYMLSKSAFPVNEPLAAMHILALNGLVSMIKGMAKRTCNELSISEEDSTNYEGYEAFWTFKCLNYNDPSSWIPF
VRKMKHIKRKLTIGADHFNRDPKKGLEFLQGIHLLPDKLDPESVASFFRYTNGLDKNLIGDFLGNHGEFCVHVLH
EFSRTFDFHDMTLDSALRVFLGTFRLPGESQKIQRVIEAFAERYYEQSSHIFCDRDAALLLSYSLIMLNTDQHNA
QVKKKMTEEDFIRNNRLINGGKDLPRKYLSELYHSICENEIQMIPDQGAGFPTMTLSRWINVIYRSKESAPFILC
DSRVPLYHDMFAILSGPTIAAISVVFDQAEQENVLQTCIDGFLAIAKISTHYRFGDILDDLVVSVCKFTTLLTPL
SFEDAIISFGDGPRARMATTTVFTIANRYGDCIHSSWRSIFDCVLSLYKLRLLPPCLANDAAGETDSSSDYVWSK
PDTSSLLISRTTPAAIPRRSFSLFGRFSQLLSFEVEEPLLHPTHEQLAAQQFLREIVLKCHIDNIFMDSKFLQAE
SLLQLVEAIILATVQFSKGSDIIEDEDSAVFCLELLISITLNNRDRILIIWKNVYKFISDIVQQKAMPCALVEKA
VFGLLKICQRLLPYKENLTDELLKSMQLILRLDAQVADAYCEPITQEVMRLVKANASHIRSHSGWRTIISLLCIT
AQHPEASEFGFEALAFIMFDGAHLLPSNYVLCVDAAREFAESWVGQIDRSISSLEMMASSVVCLVRWSYEAKNTV

NEEAAMKVSKDLGEMWLRLLQALKVVCLDQREEVRNHAILMLQRSFAGVGGIYIPNTIWFQCFNLVVFTLMDDLL
EIALESSSKEYKKMEGTLVLATKLMLKAFLQSLEELSQQASFCKLWLEVLDRMERYMKFKFRGKHSEKIHESIPE
LLKNTLLVMKTTGILMPSDDVGRDSLWQLTWLNVKKILPSLQSEVFLEPELEQMQAKHKKIYGIPATNETMLIPS
NETTT*

**>Gosra_LOC105769188**

MLKSKRKELGLSCMLNTEVGAVLAVIRRPIDPTSQFISPQEDYFDSSLQQSLKSLRGFIFNPHQEWRTIDPCIYL
SPFLDVVQSDGIPAAATGVALSAILKILKLGIFDEKTPGAKDAINIVVTGITSCRLEQTDPVSEDAVMMKILQVL
TATMRHNASVLLTDQAVCTIVNTCFQVVQQSVTRGDLLQRTARHTMHELIQIIFSRLSDVDVNEGESSESETDDI
DENSGYGIRCAVDIFQFLCSLLNVVEVVDNTEGPICHTADEDVQLFALVLINCAIELSGDEIGKHPKLLRMVQDD
LFHHLIHYGTCSTPLVLSMICSTVLNIYHFLRRFIRLQLEAFFSFVLLRIAAPGVSLQLQEVALEAIINFFRQPT
FVIEAYVNYDCDPVCRNIFEDVGKLLCKQAFPGNGPTSTLQVQAFEGLIIMIHSISTNIDKEEDSTPSDPYPIEI
KEYRPFWVEKPKDDLETWVEYIRVRKAQKRKILIGASHFSRDEKKGLEYLKHCQLVSDPPNPKSFAFFFRYTPGL
DKNMIGDYLGDPDELHIQVLKEFTATFEFHGMILDSALRTYLETFRLPGESQKIQRILEAFSERFFDQQSSEIFV
TKDSVFILCYSLIMLNTDQHNPQVKKKMTEDEFIRNNRAINGGQDLPREYLSELFHSISNNAITLFGQSGPVEMN
PSRWIELMNRAKLMQPYILCDFDRRLGRDMFACVAGPTISALSAFFEHADEDEMLHECIEGLISLARIAQYGLAD
TLDELVASFCKFTTLLNPYASAEETLFAFSNDMKPRMATLALFTIANNFGRSLRGGWRNIIDCLLKLKRLRLLPQ
SVIEFDVASSSDAPEGSKSESGVIFPNHDSKFSKNTAGVVSRFSHFLTLDSMEESISLGMSEFEQNLKIIRQCQI
GSIFGNSFQLPVEALMNLGRSLIFAAAGKGQKFSTPIEEEETVGFCWDLIIAISLANLHRFPVFWPSYSDYLLAV
AQFPLFSPLPFAEKGIIGLFKVCLKLLASYQADKTPEELIFKSINLMWKLDKEILDTCCEYITQSVSKILTEYPA
NLQTQLGWKSVLHLLSITGRHPETYDQAVEALIMLMSDAFHISKINYAYCIDCAFGFVALKNSPLEKNLKILDLM
SDSVKLLIQWYTTASDQTNNYSFSSNTSASSSDDNPKGIESVNFIMSLFVKLGEALRKTSLARREEIRNHAILAL
RRGFKLAEDLQLSSSRCISCFNLVIFAMVDDQHEKMIEYSRRENAERETRSMEGTLKLSMELLTDVYLQYLNQIM
ENPGFRTFWLGVLRRMDTCMKADLGEYGETKLHEVVPELLRKMITKMQETGILVRKEEDDLWDITYIQIQWIAPS
LKEELFPDEI*

**>Durzi_LOC111281800**

MGRLKLQSGIKAIEEEPGDYDSTYSNKATLACMINSDIGAVLAVMRRNVRWGGRYMSGDDQLEHSLIQSLKVLRK
QIFLWQNQWHTINPAVYLRPFLDVIRSDETGAPITGVALSSVHKILTLDVIDQNTVNVEDAMRLVVDAVTSCRFE
VTDPASEEVVLMKILQVLLACMKSKASVMLSNQHVCTIVNTCFRIVHQAEKKGELLQRIARHTMHELVRCIFSHL
SNVDNTEHALVNRIGTAKQELGGIDKDYAFGAKRPENGNGSEYDDQVSSGSFASNGSTGLVATVREESIVVAGSG
KETLPYDLHLMTELYGVPCMVEIFHFLCSLLNAVEHVGMGPHTNTLAFDEDVPLFALGLINSAIELGGPSFRRHP
RLLSLIQDELFRNLMQFGLSMSPLILSMVCSIVLNLYHHLRTELKLQLEAFFSCVILRLAQGKYGASYQQQEVAM
EALVDFCRQKTFMVEMYANLDCDITCRNVFEELANLLSKSAFPVNCPLSAMHILALDGLIAVIQGMAERIGNGLV
SSEQAPVSLEEYTPFWMVKCDNYGDPSHWVPFVRRRKYIKRRLMIGADHFNRDPKKGLEFLQGTHLLPDKLDPQS
VACFFRYTAGLDKNLVGDFLGNHDEFCVQVLHEFAGTFDFQDMNLDTALRLFLETFRLPGESQKIQRVLEAFSER
YYEQSPQILANKDAALLLSYSLIMLNTDQHNVQVKKKMTEEDFIRNNRHINGGNDLPREFLSELYQSICKNEIRT
TPEQGFGYPEMTPSRWIDLLHKSKKTAPYIVADSRAYLDHDMFAIMSGPTIAAISVVFDHAELEEVYQTCIDGFL
AVAKISACHHLEDVLDDLVVSLCKFTTLLNPSSVEEPVLAFGDDTKARMATVTVFTIANRYGDYIRTGWRNILDC
ILRLHKLGLLPARVASDAADESELSVDPGHGKPITNSLSSAHIQSIGTPRRSSGLMGRFSQLLSLDTEEPRSQPT
EQQLAAHQRTLQTIQKCHIDSIFTESKFLQAESLLQLERALIWAAGRPQKGSSSPEDEDTAVFCLELLIAITLNN
RDRIVLLWQGVYEHIANIVQSTVMPCALVEKAVFGLLRICQRLLPYKENLADELLRSLQLVLKLDARVADAYCEQ
ITQEVSRLVKANATHIRSQMGWRTITSLLSITARHPEASEAGFDALFFIMSDGAHLLPANYVLCLDAARQFAESR
VGQAERSVRALDLMSGSVDCLARWALGAKEAMGEEDAAKMFQDIGDLWLRLVQGLRKVCLDQREEVRNHALLSLQ
KCLTGVDGIHISHGLWLQCFDLVIFTMLDDLLEIAQGHQKEYRNMEGTLIHAMKLLSKMFLQLLHELSQLTTFCK
LWLGVLSRMEKYMKVKVRGKKSEKLQELVLELLKNILLVMKTRGVLMQRSALGGDSLWELTWLHVNNIAPSLQSE
IFPDQDPEQSLLKHGETGSVVSGETVSVPSNEMPTPEGAGAGG*

**>Durzi_LOC111286997**

MGRLKLQPGIKAIQEEPEDCDTTCSNKATLACMINSEVGSVLAVMRRNVRWGGRYILGDDQLEHSLIQSLKALRK
QIFLWQHQWHTINPALYLQPFLDVIRSDETGAPITGVALSSVHKILILDVIDQNTVNVEDAMCLVVDAVTSCRFE
VTDPASEEVVLMKILQVLLACMKCKASLMLSNQHVCTIVNTCFRIVHQAEKKSELLQRIARHTMHELMRCIFSHL
SSVDNTEHALVNTIGTAKQELGGMDNNYAFGAKQVENGNGSEYDGQASSGSFASNGSAGLGATVKEESIIVAGNG
KETVPYDLHLMTEPYGVPCMVEIFHFLCSLLNAVEHAGMGPYSNTLAFDEDVPLFALGLINSAIELGGPSFRRHP
RLLSLIQDELFRNLMQFGLSMSPLILSMVCSIVLNLYHHLRTELKLQLEAFFSCVILMLGQGKCGASYQQQEVAM
EALVDFCRQKTFMVEMYANLDCDITCSNVFEDLANLLSRSAFPVNGPLSAMHILALDGLIAVIQGMAERIGNGSV
SSEQAPVSLEEYTPFWMVKCDNYSDPSYWVPFVRRRKYIKRRLMIGADHFNRDPKKGLEFLQGTHLLPDKLDPQS
VACFFRYTAGLDKNLVGDFLGNHDEFCVQVLHEFAGTFDFQDMNLDTALRLLLETFRLPGESQKIQRVLEAFSGR
YYEQSSHILANKDAALLLSYSLIMLNTDQHNVQVKKKMTEEDFIRNNRHINGGNDLPREFLSELYHSICKNEIRT
TPEQGFGYPEMTPSRWIDLMHKSKKTAPFIVADSRAYLDHDMFAIMSGPTIAAISVVFDHAEHEDVYQTCIDGFL
AVAKISACHHLEDVLDDLVVSLCKFTTLLNASSVEEPVLAFGDNTKARMATITVFTIANRYGDYIRTGWRNILDC
ILRLHKLGLLPARVASDAADESELCADPIHGKPITNSXIFSSHTVHRKPRRSSGLMGRFSQLLSLDTEEPRSQPT
EQQLAAHQRTLQTIQKCHIDSIFTESKFLQAESLLQLARALIWAAGRPQKGSSSPEDEDTAVFCLELLIAITLNN
RDRIVLLWQGVYEHIAKIVQSTVMACALVEKAVFGLLRICQRLLPYKENLADELLRSLQLVLKLDARVADAYCEQ
ITQEVSRLVKANATHIRSQMGWRTITSLLSFTARHPEASEAGFDALLFIMSDGAHLLPANYVLCIDAVRQFAESR
VGQAERSVRALDVMSCSVDCLARWVNEAKEAMGEEDAAKMFQDIGDLWLRLVQGLRKVCLDQREEVRNHALLSLQ

KCLTGVDGIHISHGVWLQCFDLVIFTMLDDLLEIAQGHQKDHRNMEGTLILAMKLLSKVFLQLLHELSQLTTFCK
LWLGVLSRMEKYMKVKVRGKKSEKLQELVLEVLKNTLLIMKTKGVLIQRSALGGDSLWELTWLHVNNIAPSLQSE
VFPDQDPEQSLLKHGETGVISGETVSVPSNETAATEEAVAGV*
**>Durzi_LOC111291690**
MGDLTTKSRSNGFNGEFVKPSRGAFACMINSEIGAVLAVMRRNVRWGVHYVADDDQLEHSLIHSLKELRKQIFSW
QHQWKNVDPVVYLQPFLDVIQSDETGAPITGVALSSIYKILTLDVLDLDTVNVGAALHLIVDAVTSCRFEVTDPA
SEEVVLMKILQVLLACMKSKAATRLSNKHVCMIANTCFRIVHQASSKGELLQRIARHTMHELVRCIFSHLPEISC
TENEMADRSSLSALGVVGTQNGNHMLGSIQLENGNVGVDGVGPSSISDASSPPIVNSATEMDTGNIGESDRKEDA
QHGEIMMMAPFGIPCMLEIFQFLCSLLNVIENIGIGSKSNPIAYDEDVPLFALGLINSAIELGGPSFSKHPKLLA
LIQDELFRNLMQFGLSMSPLILSTICSIILNLYYHLRTELKVQLEAFFSCVLLKLAQSKHGSSYQQQEVAMEALV
DLCRQQTFVAEMYANFDCDITCSNVFEEIANLLSKNAFPVNGPLSAMHILALDGLVSMIKGMAERINNELPVSDE
ASTDREGYEAFWTLKCQNYSDPNSWIPFVRKMKYIKRKLTIGADHFNRDPKKGLEFLQEIHLLPDKLGPESVASF
FRYTTGLDKNVIGDFLGNHDEFCVQVLHEFARTFDFRDMNLDSALRVFLGTFRLPGESQKIQRVLEAFAERYYEQ
SPYILCNKDAALLLSYSLILLNTDQHNVQVKKKMTKEDFIRNNRQINGGEDLPRDYLSELYHSICENEIQMIPDQ
GAGFPMMTSSRWINVMHKSKESTPFIACDSRALLDHDMFAILSGPTIAAISVVFDQVEQEDVLQTCIDGFLAIAK
ISAHYHFGDVLDDLIVSLCKFTTLLTPFSVEEFIVSFGDDARARMATTTVFTIANSYGDYIHSGWRNILDCVLSL
HKLGLLPAVASDAADEMESLSDHERGKPATSSLSTSQTAPVAIPWKSSSLIGRFSQLLSFDMEEPRLQPTDEQLA
AQQSLREIVQKCHIHNIFMESKFLQAESLLQLVGAIIVAAGRFSKASGIVEDEDTAVFCLELLIAITLNNRDRIL
IIWQNVYKHISDIVQQKAMPCTLVEKAVFGLLKICQRLLPYKENLTDELLKSMQLILKLDARVADAYCEPITQEV
MRLVKANATHIRSHLGWRIIISLLFITARHPEASEFGFQALEFIMSEGAHLLPSNYVLCVDAAREFAESRVGEVD
RSISALDLMAGSVVCLVRWSYETKNAVEEAAMKVSQDIGEMWLRLLQGLRAVCLDQREEVRNHAILLLQSSLAGV
DGIHVPNAMWFQCFDLVIFTLMDDLLEIAQKSSSKEFRKMEGTLVLATKLMSKAFLQSLQDLSQQSSFCKLWLGV
LNRMERYMKVKFRGKHSEKIHELVPELLKNTLLVMKTTGILMPSDDIGGDSFWQLTWLHVKNIVPSLQSEVFSEQ
ELEKMQAKHKKTAGVPVTDGAVLVSSNETTT*
**>Durzi_LOC111284518**
MEEQQNSCLPCADKKKDMNKSKRKELGLSCMLNTEVGAVLAVIRRPIDPTAQLFSPQEDYFDSSIQQSLKSLRGL
IFNPHQEWRTIDPCIYLSPFLDVIQSDDIPAAATGVALSAVLKILKLDIFDEKTPGAKEAINIVVTGITNCRLEK
TDPVSEDAVMMKILQVLTGIMRHNASVLLTDQAICTIVNTCFQVVQQSVTRGDLLQRSARHTMHESIQIIFSRLP
DVDVNEGENSESETEDDENSGYGIRCAVDIFQFLCSLLNVVEVVETEGSTCHTADDDIQLFALVLINCAIELSGD
GIGKHPKLLRMVQDDLFHHLIHYGTCSSPLVLSMICSTVLNIYHFLRRFIRLQLEAFFSFVLLRIAAPGASLQLQ
EVALEAIINFFRQPKFVIEAYANYDCDPICRNIFEEIGKLLCKQAFPGTGPMSTLQVQAFEGLVIMIHNISDNID
KEDDSSTSEPYPVEITEYRPFWVDKPKDDLETWVEYIRVRKAQKKKIFIAGNHFNRDEKKGLEYLKHCQLVCDPP
NPKAFAFFFRYTPGLDKSMIGDYLGDPDELHLQVLKEFTRTFEFKGMILDSALRTYLETFRLPGESQKIQRILEV
FSERFFDQQSSEIFVTKDSVFILCYSLIMLNTDQHNPQVKKKMTEDEFIRNNRAINGGQDLPREYLSELFQSISN
HAITLFDQSGSVEMNPSRWIELMNRAKLMQSYVLCDFDRRLGRDMFASVAGPTIAALSAFFEHADEDEMLHECIE
GLISLARIAQFGLADTLDELVASFCKFTTLLNPYASAEETLFAFSNDMKPRMATLAVFTIANNFGKSLRGGWRSI
VDCLLKLKRLRLLPQSVIEFDVASTSSSDVPEALKSESGVIFPNHDPKFFKRQTSGMGSRFSHFLSLDSMEDSIS
LGMSEFEQNLKIIKQCQIGSIFGNSYNLPNEALLNLGRSLIFAAAGKGQKFSTPIEEEETVGFCWDLIIAISLSN
IHRFQVFWPSFHDYLLAVAQFPLFSPIPFAEKAIVGLFKVCLKLLASYQADKIPEELIFKSINLMWKLDKEILDT
CCEFIIQSVSKILIEYPANLQTQLGWKSVLHLLSVTGRHPETYDQAVEALIMLMSDACHISRINYAYCIDCAFGF
VALKNSPLEKNLKILDLMSDSTKLLIQWYITACSDPGNAYSISSNASTSSLEDNAKALGSINFTMSLFVKLGEAL
RKTSLARREEIRNHAILSLRKCFKLAEGLELSSTRCINCFNLVIFAVVDDQHEKMIEYSRRENAERETRSMEGTL
KLSMELLTDVYLQFLNQIAKNPGFRTFWLGVLRRMDTCMKADLGEYGETKLQQVVPDLLRKMIRKMQEEEILVRK
DGDDLWDITFIQIQWIAPSLKEELFPDEM*
**>Durzi_LOC111303223**
MEEQQNKKDMGKSKRKELGMSCMLNTEVGAVLAVIRRPIDPTAQFISPQENYFDSSIQQSLKSLRGFIFNPHQEW
RTINPCIYLSPFLDVIQSDYIPAAATGLALSAILKILKLEIFDEKTPEANEAINSVVTGITGCRLEKTDTVSEDA
VMMKILQVLTGIMRHNASVLLTDQAVCTIVNTCFQVVQQSVTRGDLLQRCARYTMHELIQIIFSRLPDVEVSEGE
NSESETDDVDENSGYGIRCAVDIFQFLCSLLNVVEVVDTEGSTWHTSDEDVQLFALVLINCAIELSGDGIGKHPK
LLRMIQDDLFHHLIHYGTCSSPLVLSRICSTVLNIYHFLRRFIHLQLEAFFSFVLLRIASPGASMQLQQVALEAI
INFFRQPTFVIEAYANYDCNPICRNVFEEVGKLLCKQAFPGTGPMTTLQVQAFEGLVIMIHNISDNIDKEDDSST
SEPFPVEITEYRPFWVDKPKDDLETWVEYIRVRKAQKKKILIAGNHFNRDEKKGLEYLKHCLLVSDPPDPKAFAF
FFRYTPGLDKSMIGDYLGDPDEFHLEVLKEFTGTFEFKGMILDSALRTYLETFRLPGESQKIQRILEVFSERFFD
QQSSEIFVTKDSVFILCYSLIMLNTDQHNPQVKKKMTEEEFIRNNRAINGGQDLPREYLSELFHSISNHAVTVFG
QSGSVEMNPSRWIELMNRAKSMQSYVLCDFDRRLGRDMFACVAGPTIAALSAFFEHADEDEMLHECIECLISVAR
IAQYGLADTLDELVASFCKFTTLLNPYASAEETLFAFSNDMKPRMATLAVFTIANNFGDALQGGWRNIVDCLLKL
KRLRLLPQSVIEFDVASTSSSDVPEALKSESGVIFPNHDPKLFKRQTSGMVSRFSHFLSLDTMEVSISLGISEFE
QNLKIIKQCRIGRIFQNSSNLPTEALLNLGRSLIFAAAGKGQKFSTPIEEEETVGFCWDLIIAISLSNIYRFQVF
WPSFHDYFLAVAQFPLFSPIPFAEKAIIGLFSICLKLLASYQADKMPEELIFKSINLMWKLDKEIFDTCSKFIIQ
SVSKILIEYPANLQTQLGWKSTLHLLSVTGRHPETYDQAVEILIMLMSDAFHISRINYAYCIDCAFGFVALKNSP
LEKNLKILDLMSDTVNLLIQRYKIEYSNPGNNYSIGSNANTSSLEDNSKAIGSINFTLSLFVKLGEALRKTSLAR
REEIRNHAIASMRRGFQLAEDLELSSTSCINCFNLVIFAMVDDLHEKMIEYSRRENAEREMRSMEGTLKISIELM

TDVYLQFLKLISENPGFRTFWLGVLRRMDTCMKADLGEYGETKLQEVVPDLLRKMIRNMQEKEILVPKDGDDLWE
ITYIQIQWIAPSLKEELFPDEL*
**>CCACVL1_06544**
MGRLKLQPGIKAIEEEPEDCDATHSNKATLACMINSEVGAVLAVMRRNVRWGGRYMSGDDQLEHSLIQSLKALRK
QIFLWQHQWHTINPAVYLQPFLDVIRSDETGAPITGVALSSVYKILTLDVIDQNTVNVEDAMRLVVDAVTSCRFE
VTDPASEEVVLMKILQVLHACMKSKASVMLSNQHVCTIVNTCFRIVHQAEKKGELLQRIARHTMHELVRCIFSHL
SNVDNTEHALVNRNSNANQELGGIDNDYAFGAKQAENGNGSEYDGQASSGSFASNGSSGLVATMREESMIVAGNG
KETVPYDLHLMTEPYGVPCMVEIFHFLCSLLNAVEHVGMGPRSNTLAFDEDVPLFALGLINSAIELGGPSFRRHP
RLLSLIQDELFRNLMQFGLSMSPLILSMVCSIVLNLYHHLRTELKLQLEAFFSCVILRLAQGKYGASYQQQEVAM
EALVDFCRQKTFMVEMYANLDCDITCSNVFEDLANLLSKSAFPVNCPLSAMHILALDGLIAVIQGMAERIGNASV
GSEHKPVSLDEYTPFWMVKCDNYSDPSHWVPFVRRRKHIKRRLMIGADHFNRDPKKGLEFLQGTHLLPDKLDPQS
VACFFRYTAGLDKNLVGDFLGNHDEFCVQVLHEFAGTFDFQDMNLDTALRLFLETFRLPGESQKIQRVLEAFSER
YYEQSPQILANKDAALLLSYSLIMLNTDQHNVQVKKKMTEEDFIRNNRHINGGNDLPREFLTELYHSICKNEIRT
TPEQGYGYPEMTPSRWIDLMHKSKKSAPFIVSDSRAYLDHDMFAIMSGPTIAAISVVFDHAEHEDVYQTCIDGFL
AVAKISACHHLEDVLDDLVVSLCKFTTLLNPSSVEEPVLAFGDDTKARMATVTVFTIANRYGDYIRTGWRNILDC
ILRLHKLGLLPARVASDAADESEVSADPSHGKPITNSLSSAQIQSIGTPRRSSGLMGRFSQLLSLDTEEPRSQPT
EQQLAAHQRTLQTIQKCHIDSIFTERRPQKGNSSPEDEDTAVFCLELLIAITLNNRDRIVLLWQGVYEHIASIVQ
STVMPCALVEKAVFGLLRICQRLLPYKENLADELLRSLQLVLKLDARVADAYCEQITQEVSRLVKANATHIRSQM
GWRTITSLLSITARHPEASEAGFDALLFIMSDGAHLLPANYVLCVDSARQFAESRVGQAERSVRALDLMSGSVDC
LVRWANEAKETMGEEDAAKMLQDIGDLWLRLVQGLRKVCLDQREEVRNHALLSLQKCLTGVDGIQISHGLWLQCF
DLVIFTMLDDLLEIAQGHQKDYRNMEGTLILAMKLLSKVFLQLFHELSQLTTFCKLWLGVLSRMEKYMKVKVRGK
KSEKLQELVLELLKNTLLVMKTRGVLIQRSALGGDSLWELTWLHVNNIAPSLQSEVFPDQDPDQSLLKHGETSGV
VSGETVSVSSNETAAPEGAVAGG*
**>CCACVL1_13722**
MGDLATQSRSNGFNGNFVKPSRGAFACMINSEIGAVLAVMRRNVRWGVRYVSDDDQLEHSLIHSLKELRKQIFSW
QHQWQNVDPVVYLKPFLDVIQSDETGAPITGVALSSILKILTLDVLDLDTVNVGDAMHLIVDAVTSCRFEVTDPA
SEEVVLMKILQVLLACMKSKAATRLSNKHVCMIANTCFRIVHQASSKGELLQRIARHTMHELVRCIFSQLPEISS
TEHETANGSSSSATEVVAQNRNYMHGSAQLENGNVGIDHDGSSSHSNASSPPILKSATEMDASEIGDTDGEMLAP
FGVPCMVEIFQFLCSLLNVIDHIGIGPKSNPIAYDEDVPLFALGLINSAIELGGPSFSKHPKLLALIQDELFRNL
MQFGLSMSPLILSTVCSIVLNLYYHLRTELKMQLEAFFSSVLVRLAQSKYGSSYQQQEVAMEALVDLCRQQTFVA
EMYANFDCDITCSNVFEDLANLLSKSAFPVNGPLSALHILALDGLVSMIKGMAERIGNELPVSDEASINGEGYEA
FWTLQCQNYSDPSSWIPFVRKIKHIKRKLMLGADHFNRDPKKGLEFLQGIHLLPDKLDPVSVASFFRYTTGLDKN
LIGDFLGNHDEFCVQVLHEFARTFDFHDMNLDSALRVFLGTFRLPGESQKIQRVLEAFAERYYEQSPHILCNKDA
ALLLSYSLIMLNTDQHNVQVKKKMTEEDFIRNNRLINGGKDLPREYLSELYHSICENEIQMIPDQGAGFPVMTSS
RWINVVHKSKESTPFITCDSRALLDHDMFAILSGPTIAAISVVFDQAEQEDVLQTCIDGFMAIAKISAHYHFGDV
LDDLVVSLCKFTTLLTPLSVEDAIVTFEDDARARMATTTVFTIANSYGDYIHSGWRNILDCVLSLHKLGLLPARL
ASDAADEMESSSDHERGKPATNSSSTSQTAPVATPRKSSSLIGRFSQLLSFDMEEPRVQPTEEQLAAQQRLREIV
QKCHIDNIFMESKFLQSESLLQLVGAIILAAGRFRKGSGFIEDEDSAVFCLELLIAITLNNRDRIMIIWQNVYKH
IADIVQQRATPCTLVEKAVFGLLKICQRLLPYKENLTDELLKSMQLILKLDARVADAYCEPITQEVMRLVKANAT
HIRSHLGWRIIISLLFITARHPEASESGFEALAFIMSEGAHLLPSNYVLCVDAAREFAESRVGEVERSISALDMM
AGSVVCLVRWSNETKNASEEDSTKVSQDIGEMWLRLVQGLRALCSDQREEVRNHAILMLQRSLAGVDGIHLPNAM
WFQCFDLVIFTLLDDLLEIAQGSSSKEHRKMEGTLVLATKLMSKAFLQSLQDLSQQPSFCKLWLGVLNRMERYMK
VRFRGKRSEKIHELVPELLKNTLLVMKTTGILMPSDDIGGDSFWQLTWLHVKNIVPSLQSEVFSEQELEQMQAKH
KRTEGIPATDGAVLVPSNETTA*
**>CCACVL1_09433**
MGKSKRKELGLSCMLNTEVGAVLAVIRRPDPTSQFMSPQEDYGDTQIQHSLKALRGLIFNPPQEWRAIDPCIYLS
PFIDVIQSDEIPATATGVALSAILKVLKLEIFNEKTPGAKDAINFIVSGITSCRLEKTDPASEDVVMMKILQVLA
GIMRHSCSFLLTDQSVCTIVNTCFQVVQQSATRGNLLQRSGRYTMHELIQIIFSRLPDVDENEGESSGSETDDMD
ENSGYGIRCAVDIFQFLCSLLNVVELVETEVSPWHTADEDIQLYALVLINCAIELSGDGIGKHPKLLRMIQDDLF
HHLIHYGTCSSPLVFSMICSTVLNIYHFLRRFIRLQLEAFFTFVLLRVGALGATLQLQEVAVEAIINFLRQPTFV
IEAYTNYDCDPICRNIFQEIGKLLCKHAFPGTGPLTTLQVQAFEGLSIMIHNISENIDKEDDSSTSEPYPVEITE
FRPYWVEKQKDDLETWVEYIRFRKVQKKKMLIAGNHFNRDVKKGLEYLKQSQLVSEPPNPKAFAFFFRYTPGLDK
SMIGEYFGDPDDFHLQVLREFTATFEFQGMILDSALRTFLETFRLPGESQKIERILEVFSERFFDHQSSEIFVTK
DVVFIFCYSLIMLNTDQHNPQVKKKMTEEEFIRNNRAINGGKDLPREYLSELFNSISNHAISLDGQSGQVDMNPN
RWIELINRAKIIQPYLLSDFERRIGRDMFASVAGPVVATLSAFFEHADDDEMLHECIEGLISIARIAQYGLSDTL
DELVASFCKFTTLLNPYASAEETLFAFSHDMKPRLATLAVFTIANNFGDSLRGGWRNIIDCLLKLKRLKLLPQSV
VEFDASDVTDASKSDSGVIFPNQDAKFSKRQHSGMVGRFSHFLSLDSMEDSISMGMSEFEQNLKVIKQCRIGSIF
GSSYNLPTDALMNLGRSLIFAAAGKGQKFSTPVEEEETVAFCWDLIIAISLANVHRFQVFWPSFHENLLAVAQFP
LFSPIPFVEKGIIGLFKVCLKILASYQVDKMPEELIFKSINLMWMLDKEILESCREIIIQSVSKILIEYPANLQT
PLGWKSVINLLSISGRHPDTYDQAVETLIMLMSDAFHISPMNYAYCIDCAFGIIAMKTSPLDKTLKILDLMSDTV
NLLVQWHQNNYTDAGNNNYSIASNTSTSSMDDTSKATGSNNLAMNLFIKLGEAFRKSSLARREEKRNKAVSALGK
SFELAENLELSSTNCITCFNLVIFAMVDDLHEKMIDYSRRDNAEREMRSMEGTLKLAMELLTDVYLQFLKQIAEN

PGFRTFWLGVLRRMDTCMKADLGEYGETKLQEVVPDLLRKMIKSMQEKEILVSKDGDDLWEITYIQIQWIAPSLK
EELFPENEM*
**>COLO4_27976**
MGRLKLQPGIKAIEEEPEDCDATHSNKATLACMINSEVGAVLAVMRRNVRWGGRYMSGDDQLEHSLIQSLKALRK
QIFLWQHQWHTINPAVYLQPFLDVIRSDETGAPITGVALSSVYKILTLDVIDQNTVNVEDAMRLVVDAVTSCRFE
VTDPASEEVVLMKILQVLHACMKSKASVMLSNQHVCTIVNTCFRIVHQAEKKGELLQRIARHTMHELVRCIFSHL
SDVDNTEHALVNRNSNAKQELGGIDNDYAFGAKQAENGNGSEYDGQASSGSFASNGSSGLVATMREESMIVAGNG
KATVPYDLHLMTEPYGVPCMVEIFHFLCSLLNAVEHVGMGPRSNTLAFDEDVPLFALGLINSAIELGGPSFRRHP
RLLSLIQDELFRNLMQFGLSMSPLILSMVCSIVLNLYHHLRTELKLQLEAFFSCVILRLAQGKYGASYQQQEVAM
EALVDFCRQKTFMVEMYANLDCDITCSNVFEDLANLLSKSAFPVNCPLSAMHILALDGLIAVIQGMAERIGNASV
GSEHKPVSLDEYTPFWMVKCDNYSDPSHWVPFVRRRKYIKRRLMIGADHFNRDPKKGLEFLQGTHLLPDKLDPQS
VACFFRYTAGLDKNLVGDFLGNHDEFCVQVLHEFAGTFDFQDMNLDTALRLFLETFRLPGESQKIQRVLEAFSER
YYEQSPQILANKDAALLLSYSLIMLNTDQHNVQVKKKMTEEDFIRNNRHINGGNDLPREFLTELYHLICKNEIRT
TPEQGYGYPEMTPSRWIDLMHKSKKSAPFIVSDSRAYLDHDMFAIMSGPTIAAISVVFDHAEHEDVYQTCIDGFL
AVAKISACHHLEDVLDDLVVSLCKFTTLLNPSSVEEPVLAFGDDTKARMATVTVFTIANRYGDYIRTGWRNILDC
ILRLHKLGLLPARVASDAADESEVSADPSHGKPITNSLSSAQIQSIGTPRRSSGLMGRFSQLLSLDTEEPRSQPT
EQQLAAHQRTLQTIQKCHIDSIFTERRPQKGNSSPEDEDTAVFCLELLIAITLNNRDRIVLLWQGVYEHIASIVQ
STVMPCALVEKAVFGLLRICQRLLPYKENLADELLRSLQLVLKLDARVADAYCEQITQEVSRLVKANATHIRSQM
GWRTITSLLSITARHPEASEAGFDALLFIMSDGAHLLPANYVLCVDSARQFAESRVGQAERSVRALDLMSGSVDC
LARWANEAKETMGEEDAAKMLQDIGDLWLRLVQGLRKVCLDQREEVRNHALLSLQKCLTGVDGIQISHGLWLQCF
DLVIFTMLDDLLEIAQGHQKDYRNMEGTLILAMKLLSKVFLQLFHELSQLTTFCKLWLGVLSRMEKYMKVKVRGK
KSEKLQELVLELLKNTLLVMKTRGVLIQRSALGGDSLWELTWLHVNNIAPSLQSEVFPDQDPDQSLLKHGEPAGV
VSGETVSVSSNETAAPEGAVAGG*
**>COLO4_12228**
MGDLATQSRSNGFNGNLVKPSRGAFACMINSEIGAVLAVMRRNVRWGVRYVSDDDQLEHSLIHSLKELRKQIFSW
QHQWQNVDPVVYLKPFLDVIQSDETGAPITGVALSSILKILTLDVLDLDTVNVGDAMHLIVDAVTSCRFEVAQNR
NYMLGSAQLENGNVGIDHDGSSSHSNASSPPVLKSATEMDTSEIGDTDGEMLAPFGVPCMVEIFQFLCSLLNVIE
HIGIGPKSNPIAYDEDVPLFALGLINSAIELGGPSFSKHPKLLALIQDELFRNLMQFGLSMSPLILSTVCSIVLN
LYYHLRTELKMQLEAFFSSVLVRLAQSKYGSSYQQQEVAMEALVDLCRQQTFVAEMYANFDCDITCSNVFEDLAN
LLSKSAFPVNGPLSALHILALDGLVSMIKAMAERIGNELPVSDEASINGEGYEAFWTLKCQNYSDPSSWIPFVRK
IKHIKRKLMLGADHFNRDPKKGLEFLQGIHLLPDKLDPVSVASFFRYTTGLDKNLIGDFLGNHDEFCVQVLHEFA
RTFDFHDMNLDSALRVFLGTFRLPGESQKIQRVLEAFAERYYEQSPHILCNKDAALLLSYSLIMLNTDQHNVQVK
KKMTEEDFIRNNRLINGGKDLPREYLSELYHSICENEIQMIPDQGAGFPVMTSSRWINVVHKSKESTPFITCDSR
ALLDHDMFAILSGPTIAAISVVFDQAEQEDVLQTCIDGFLAIAKISAHYHFGDVLDDLVVSLCKFTTLLTPLSVE
DAIATFEDDARARMATTTVFTIANSYGDYIHSGWRNILDCVLSLHKLGLLPARLASDAADEMESSSDHERGKPAT
NSSSTSQTAPVATPRKSSSLIGRFSQLLSFDMEEPRVQPTEEQLAAQQRLREIVQKCHIDNIFMESKFLQSESLL
QLVGAIILAAGRFRKGSGFIEDEDSAVFCLELLIAITLNNRDRIMIIWQNVYKHIADIVQQRATPCTLVEKAVFG
LLKICQRLLPYKENLTDELLKSMQLILKLDARVADAYCEPITQEVMRLVKANATHIRSHLGWRIIISLLFITARH
PEASESGFEALAFIMSEGAHLLPSNYVLCVDAAREFAESRVGEVERSISALDMMAGSVVCLVRWSNETKNASEED
SIKVGQDIGEMWLRLVQGLRALCSDQREEVRNHAILMLQRSLAGVDGIHLPNAMWFQCFDLVIFTLLDDLLEIAQ
GSSSKEHRKMEGTLVLATKLMSKAFLQSLQDLSQQPSFCKLWLGVLNRMERYMKVRFRGKRSEKIHELVPELLKN
TLLVMKTTGLLMPSDDIGGDSFWQLTWLHVKNIVPSLQSEVFSEQELEQMQAKHKRTEGIPATDGAVLIPSNETT
A*
**>COLO4_34896**
MEAEQNKCPSCGDKKKDMGKSKRKELGLSCMLNTEVGAVLAVIRRPDPTSQFMSPQEDYGDTQIQHSLKSLRGLI
FNPPQEWRAIDPCIYLSPFIDVIQSDDIPASATGVALSAILKILKLEIFNEKTPGAKDAINFIVSGITSCRLEKT
DPVSEDAVMMKILQVLAGIMRHNCSFLLTDQSVCTIVNTCFQVVQQSATRGDLLQRSGRYTMHELIQIIFSRLPD
VEDNEGESSGSETDDMDENSGYGIRCAVDIVQFLCSLLNVVELVETEVSSWHTADEDIQLFALVLINCAIELSGD
GIGKHPKLLRMIQDDLFHHLIHYGTCSSPLVFSMICSTVLNIYHFLRRFIRLQLEAFFTFVLLRVGALGATLQLQ
ELAVEAIINFLRQPTFVIEAYTNYDCDPICRNIFQEIGKLLCKHAFPGTGPLTTLQVQAFEGLSIMIHNISENID
KEDDSSTSEPYPVEITEFRPFWVEKQKDDLETWVEYIRFRRVQKKKMLIAGNHFNRDVKKGLEYLKHSQLVSEPP
NPKAFAFFFRYTPGLDKSMIGEYFGDPDDFHLQVLREFTATFEFQGMILDSALRTFLETFRLPGESQKIERILEV
FSERFFDHQSSEIFVTKDVVFIFCYSLIMLNTDQHNPQVKKKMTEEEFIRNNRAINGGKDLPREYLSELFNSISN
HAISLDGQSGQVDMNPNRWIELINRSKIINPYLLTDFERRIGRDMFASVAGPIVATLSAFFEHADDDEMLHECIE
GLISIARIAQYGLSDTLDELVASFCKFTTLLNPYASAEETLFAFSHDMKPRLATLAVFTIANNFGDSLRGGWRNI
IDCLLKLKRLKLLPQSVVEFDASDVTDASKSEFGVIFPNQDPKFSKRQHGMVGRFSHFLSLDSMEDSISMGMSEF
EQNLKVIKQCRIGSIFGSSCNLPTDALMNLGRSLIFAAAGKGQKFSTPVEEEETVAFSWDLIIAISLANIHRFQV
FWPSFHENLLAVAQFPLFSPIPFVEKGIIGLFRVCLKILASYQVDKMPEELIFKSINLMWMLDKEILESCREIII
QSVSKILIEYPANLQTPLGWKSVINLLSISGRHQDTYDQAVETLIMLMSDAFHISPMNYAYCIDCAFGFIAMKTS
PLDKTLKILDLMSDTVNLLVQWHQNNYTDAGNNNYSIASNNSTSSMDDTSKATGSNNLSMNLFIKLGEAFRKSSL
ARREEKRNKAVSALGKSFELAENLELSSTNCITCFNLVIFAMVDDLHEKMIDYSRRDNAEREMRSMEGTLKLAME
LLTDVYLQFLKQIAENPGFRTFWLGVLRRMDTCMKADLGEYGETKLQEVVPDLLRKMIKSMQEKEILVSKDGDDL
WEITYIQIQWIAPSLKEELFPEM*

**>Aqusi_92.109**
MGRLKLLSGIKAIEEEPEDFEATLTNKTSLACMINSEVGAVLAVMRRNVRWGGRYMSGDDQLEHSLIQSLKALRK
QIFSWQHQWHTINPAVYLQPFLDVIRSDETGAPITGVALSSVYKILTLDVIDQNTVNVEDAMHLVVDSVTSCRFE
VTDPASEEVVLMKILQVLLACMRSKASVVLSNQHVCTIVNTCFRIVHQAGSKGELLQRIARHTMHELVRCIFSHL
QNVDITEHALVRPSVVKREVGGTDNNYALGVKQMENGNGGSEYDTPSSTLTVPNTFSSGASPIEESTLIGGNGKD
TASYDLHIMTEPYGVPCMVEIFHFLCSLLNVVESMGMGPRSNTVAFDEDVPLFALGLINSAIELGGHSFRNHSRL
LSLIEDELFSNLMQFGLSMSPLILSIVCSIVLNLYNHLRTELKLQLEAFFSCVILRLAQGKNGASYQQQEVAMEA
LVDFCRQKTFMVEMYANLDCDITCSNVFEDLANLLSKSAFPVNCPLSAMHILALDGLIAVILGMAERIGNGSIGS
EHAPVTLEEYTPFWMVKCDNYNDPMHWVPFVRRRKYIKRRLMIGADHFNRDPKKGLEFLQGTHLLPDKLDPQSVA
CFFRYTAGLDKNLVGDFLGNHDEFCVQVLHEFAGTFDFQDMNLDTALRLFLETFRLPGESQKIQRVLEAFSERYY
EQSPQILANKDAALLLSYSLIMLNTDQHNVQVKKKMTEEDFIRNNRHINGGNDLPREFLSELYHSICKNEIRTTP
EQGHGYPEMTPSRWIDLMHKSKKTAPFIVSDAKPYLDHDMFAIMSGPTIAAISVVFDHAEHEDVYQTCIDGFLAV
AKISACHHLEDVLDDLVVSLCKFTTLLNPSSVEEPVLAFGDDTKARMATVTVFTIANRYGDYIRTGWRNILDCIL
RLHKLGLLPARVASDAADESEPSADPGQGKPMATSLSSAHIQSMGTPRRSSGLMGRFSQLLSLDTEEPRSQPTEQ
QLAAHQRTLQTIQKCHIDSIFTESKFLQAESLLQLARALIWAAGRPQKGNSSPEDEDTAVFCLELLIAITLNNRD
RIVLLWQGVYEHIANIVQSTLMPCALVEKAVFGLLRICQRLLPYKENLADELLRSLQLVLKLDARVADAYCEQIT
HEVSRLVKANATHIRSQMGWRTITSLLSITARHPEASEAGFDALLFIMSDGAHLLPANYVLCVDASRQFAESRVG
QAERSVRAVDLMAGSVECLAGWACEAKEAMGEDDAAKMVQDIGDMWLRLVQGLRKVCLDQREEVRNHALLSLQKC
LTGVDNIHLSHGLWLQCFDLVIFTMLDDLLEIAQGHSQKDYRNMEGTLILAMKLLLKVFLQLLDELSQLTTFCKL
WLGVLSRMEKYMKVKVRGKKSEKIQEMVLELLKNILIVMKTRGVLIQRSALGGDSLWELTWLHVNNIAPLLQSEV
FPDQEGEQAPHNEGEVGGGTLSGETIAVPSKEAPASEDSGSGG*
**>Aqusi_3.680**
MGHLNPQSVCHVFQGELKDHPAKPSGGALALMINSEIGGVLAVMRRNVRWGVHYMADDDQLDHSLIHSLKELRKQ
LLSWQLHWPSISPALYLQPFLDVIRSDETGAPITGVALSSVYKILSLDVLDLETVNVGAAMHLIVDTVTSCRFEV
TDPGSEEVVLMKILQVLLACMKSKAAASLSNQHVCNIVNTCFRVVHQAGSKGELLQRIARHTMHELVRCIFSQLP
HISCSDNAMANKTRSSGSKMTSADDGEYKAENAQLKSANNHGAPDGQSVCPGITEPELATEIRASHTGEGGHGFE
DIENGDESMKEPFGIPCMVEIFQFLCSLLNAMEHFGVDPKSNPMAYEEDVPLFALGLINSVIELGGSSFSKHRKL
LVVIQDELFRNLMQFGLSMSPLILSTVCSIILNLYYYLRTELKVQLEAFFSCVLLRLAQSKHGFSYQQKEVAMEA
LVDLCRQPSFIAEIYTNFDCDITCSDLFEELVNMLSKSAFPVNGPLSAIHILALDGLVSVVKGMSERIGTVLPVS
YETLPDYVGYEAFWTLKCENFSDPDHWVPFVRKMKHIKKKLMVGTDHFNQDPKKGLEFLQEMHLLPAKLDAKSVA
SFFRYTVGLDKNLTGDFLGKHDEFCIEVLSEFAGTFDFRDMNLDTALRLFLGTFRLPGESQKIQRVLEAFAERYY
QQSPHIFSNKDAALLLSYSIILLNTDQHNVQVKRKMTEEDFIRNNRNINGGKDLPREFLSELYHSICENEIQMIP
EQGAGFPVMTSGQWITVMHKSREISPFITWDSRAILDHDMFALLSGPIVAAISVVFDQAEQEEILQTCTDGLLSI
AKISAHYYFDGVLNDLVVYLCKFTTLLTPLSAEEALFTFGEDARARMATTAVFTLANNYGDYINSGWRNIFDCVL
GLQKHGLLPASLARDAADDVEFSFELEQGKPAAAGIFSELEQGKPAATSLSASHIPSVVTTRKSSSLIGRFSLFL
SFDVEEPNVQPTEEQLAVERQIREIIQNCHIDRIFTESKFLQAESLLQLVKALISVAGGLQKSSSSVEYEDSAVF
CLELLIAITLNNRDRIMLIWKGVYEHIANIVQYTLMPCTLVEKAVFGLLKICQRLLPYKENLTDELLKSLQLILK
LDARVADAYCEPITQEVTRLVKANATHISSNVGWRTIVSLLSITARHPEASESGFEALTFIMSDGAHLLPSNFVL
CVDAARQFAGSRVGQVDRSISALDMMADSVVCLTRWSNEAKNAGGDEAVKKVSQDIGEMWLRLVQALKKACLDQR
EEVRNHSILMLQRSLAGVDGIQLPSPMWLQCFDHVIFTLLDEVLDVGQGNSAKEYKKMEDTLVLAMKLMSKAFLQ
LMRDLSLQPSFCKLWLGVLNRMEKYMKVKIRGRRSEKIHELVPELLKNTLLVMKTTGILMPTDDIGGDSFWKLTW
LHVKNVVPSLQQEVFPEHEVVQLLAQYKDPPLNVTDGEVLVSSSETKV*
**>Aqusi_sc277.24**
MVEDEPKRCPSCGEKPKSGAKSKRKELTVSCMLNTEVGAVLAVIRRPPDPNLQYVAPQEELYDSVILQSLKSLRS
LIFNPHQEWRQIDPSIYISPFLDVIQSDDIPAAATNGALSSILKILELQVFDEKTPGAREAMNTIVWGLTSCRLE
KSDPLVEDAAMMKILQVLTGIMKHSASVLLTDQAVCTAVNTCFQVLQQSANRSDLLQRSARHTMHDLVQIIYSRL
SDLDGYEGDDHDGESTNAYVGYGVGCAVDIFQFLCSLLNVMQTVESTEGSTFQTSDEDVQIFGLVLINCAIELCG
DGIGRHPKLLGMIQDDLFHHLIHYGTCSSPLLLSMICSTVLNIYHFLRRFIRLQLEAFFSYVLLRTAASGSSVQL
QEVALEGIINFCRQPTFIVEAYVNYDCDPLCQNVFEEVGKLLCRHSFPGSGPLTSLQLQAFEGLVILVHNISDGC
FKEESSPFEAPQPVYVREYKPFWEEKPKEDFDTWVEHIRLRKAQKKKILIAANHFNREDKKGLDYLKLSHLVGDP
PEPKAFAFFFRYVPGLDKNMIGDYLGDPDDFHLQVLKEFTGTFEFSGMTLDAALRTYLESFRLPGESQKIQRILE
AFSEKFYDQQSSQIFVSKDAVFILCYSVIMLNTDQHNPQVKKKMTEEEFIRNNRAINGGQDLPREYLSDLFHSIS
TNAITLFGQNGLAMEMTQSQWIELINHTRAVQPLFLCDFDPRLGRDMFACIAGPVTAALSAFFEHVDEDALLHEC
IEGIISVSRVAQYGLEDILDELVASFCKFTTLLNPYASAEETLFAFSNDLKPRMATLAVFTIANSFGNSIRGGWR
NIIDCLLKLKKLKLLPQSVTEEAISALSDHEVVRSESGAVVPGHDPPFTTQATSSIVSRFSHFLSLDSMEESVSL
GMSEFEQNVKVIKQCQIGSIFSNTSNLPNDALLSLGRSLIFAAAGKGQKFSTPIEEEETVGFCWDLVTIISAANV
HRFVDFWPAFSDYLIGVAQLPLFSPIPFAEKGIISLLKVSMRLLSSDQSDKLAEDFIFKSINLMWKLDKEILDTC
SEFITQSVSKILSQYPANLQTQLGWKSALHLLSVTGRHPETYDQAVDTLILLMSEATHLSRINYAYCIDCAFGFI
ALKNSPLEKNLTLLELMSESTSSLTQWYRVGLSEQAMYSMSSTSSTASMEDSLRSPSYIMNLFVKLGEAFRKTSL
ARREEVRNHAVISLRKSFELAEALDFSSTNCMNCINLVIFAMVDDLHEKMLEYSRRDSSEREMRSMEGTLKIAME
LLTDVFLQFLKQIAENPGFRTFWLGVLRRMDTCMKADLGAYGATKLQEVVPDLLRRMIKRMQETGILEPKEGEDL
WEITYIQIQWIAPSLKDELFPDEF*
**>Citsi_LOC102627469**

MGRLKLQSGIKAIEEEPEEYDATYSNKATLSCMINSEVGAVLAVMRRNRSVRWGGQYMSGDDQLEHSLIQSLKTL
RKQIFSWQHPWHTINPAAYLQPFLDVIRSDETGAPITSIALSSVYKILSLDVIDQNSINVEEAMHLVVDAVTSCR
FEVTDPASEEVVLMKILQVLLACMKSKASIVLSNQHVCTIVNTCFRIVHQAGNKGELSQRIARHTMHELVRCIFS
HLPDVDNSEHALVNGVTAVKQEIGGLDTDYAFGGKQLENGNGGSEYEGQQSFANLVSPSGVVATMMEENMNGSST
GKDSVSYDLHLMTEPYGVPCMVEIFHFLCSLLNISEHMTMGPRSNTIALDEDVPLFALRLINSAIELGGPAIRRH
PRLLSLIQDELFRNLMQFGLSMSPLILSMVCSIVLNLYHHLRTELKLQLEAFFSCVILRLAQSRHGASYQQQEVA
MEALVDFCRQKTFMVEMYANLDCDITCSNVFEDLANLLSKSAFPVNCPLSAMHILALDGLIAVIQGMAERIGNAS
VSSEQSPVTLEEYTPFWMVKCDNYSDPNHWVPFVRRRKYIKRRLMIGADHFNRDPKKGLEFLQGTHLLPDKLDPQ
SVACFFRYTAGLDKNLVGDFLGNHDEFCVQVLHEFAGTFDFQDMNLDTALRLFLETFRLPGESQKIQRVLEAFSE
RYYEQSPQILANKDAALLLSYSLIMLNTDQHNVQVKKKMTEEDFIRNNRHINGGNDLPREFLSELYHSICKNEIR
TTPEQGVGFPEMTPSRWIDLMHKSKKTAPFIVADSKAYLDHDMFAIMSGPTIAAISVVFEHAEHEEVYQTCIDGF
LAVAKISACHHLEDVLDDLVVSLCKFTTLLNPAAVEEPVLAFGDDTKARMATVSVFTIANRYGDFIRTGWRNILD
CILRLHKLGLLPARVASDAADESELSADPSQGKPITNSLSSAHMPSIGTPRRSSGLMGRFSQLLSLDTEEPRSQP
TEQQLAAHQRTLQTIQKCHIDSIFTESKFLQAESLLQLARALIWAAGRPQKGNSSPEDEDTAVFCLELLIAITLN
NRDRIVLLWQGVYEHIANIVQSTVMPCALVEKAVFGLLRICQRLLPYKENLADELLRSLQLVLKLDARVADAYCE
QITQEVSRLVKANATHIRSQMGWRTITSLLSITARHPEASEVGFEALLFIMSDGTHLLPANYVLCIDSARQFAES
RVGQAERSVRALELMSGSVDCLARWGREAKESMGEDEVAKLSQDIGEMWLRLVQALRKVCLDQREDVRNHALLSL
QKCLTGVDGIHLPHGLWLQCFDMVIFTMLDDLLEIAQGHSQKDYRNMEGTLILAMKLLSKVFLQLLHELSQLTTF
CKLWLGVLSRMEKYMKVKVRGKKSEKLQEIVPELLKNTLLIMKTRGVLVQRSALGGDSLWELTWLHVNNIVPSLQ
SEVFPDQDSDQPQLKQSDNGGGLVSDEMGSIPSNESAASESAET*
**>Citsi_LOC102624619**
MRMGHLNLPSGTGSNAFYAGVKDPLIKPSGGAFACMINSEIGAVLAVMRRNVRWGVRYMADDEQLEHSLIHSLKE
LRKQIFLWQNQWHKVDPAVYLQPFLDVIQSDETGAPITGVALSSVYKILILDVLDLDTVNVGEAMHLIVEAVTSC
RFEVTDPASEEVVLMKILQVLLACMKSKAAVKLSNQHVCNIVNTCFRVVHQASSKGELLQRIARQTMHELVRCIF
SHLPHIDCLEQSSALGSRSDNGNKVGLMEKEITSGSKPLENGNVSVERDGQSSVEANNGETTVEMGSTENGEKIM
MEPFGVPCMVEIFHFLCSLLNAIENMGIGPRGNPIADDEDVPLFALSLINSAIELGGSSIGKYPRLLVLIQDELF
RYLMQFGLSMSPLILSTVCSIVLNLYHHLRVELKAQLEAFFSCVLLRLAQSKHGSSYQQQEVAMEALVDLCRQQS
FMSEMYANFDCDITCGNLFEDLTNLLSKSAFPVNGPLSAMHVLALDGMISMVQGMAERISNEFPAPEGATVDPEE
YNAFWTLKCSDYSDPNNWIPFVRKMKYIKRKLMVGADHFNRDPKKGLEFLQGMHLLPDKLDPQSVALFFRYTVGL
DKNLIGDFLGNHDEFCVQVLHEFAGTFNFRGMNLDTALRLFLGTFRLPGESQKIQRVLEAFAERYYEQSSDILSD
KDAALLLSYSLILLNTDQHNAQVKKKMTEEDFIRNNRSINGGKDLPREYLAELYHSICENEILMIPEQGAGSPVM
TSSRWINVLHKSREATPFIVCDSRALLDHDMFIILSGPTVAAMSVIFDQVEREDVLQRCVDGFLAVAKLSTFYHF
GDILDDLVVSVCKFTTLLTPLSVEEAVLALGDDTKARMALTTLFTIANRYGDYIHSGWKNILDCVLSLHKLGLLP
ARLVSDAADDMEPSSDQEQEKPATSSVSTSHVTPVATPRKSSSLIGRFSQLLSFDMEEPRLQPSEEELAAHQRTR
DIIQNCHIDSIFSESKFLQAESLLDLVKALILASGRLRKGSSSGEDEDTGVFCLELLIAITLNNRDRIMLIWHGV
YEHIANIVQSTVMPSMLVEKAVFGLLRICQRLLPYKENLTEELLKSLQLILKLDARVADAYCEPITQEVMRLVKA
NSTHIRSHVGWRTIISLLSITARHPEASEAGFEALAFIMSEAAHLLPSNFILCVDAARQFAESRVGEVDRSVSAL
ELMAGSVVSLVRWSSEAKNAVGEEAAIKLSQDIGEMWLRLVQGLKKVCLDQREEVRNHAVLALQRSLAAVDGIRL
PNALWFQCFDMVIFTLLDDLLEIAQASSPKDYRNIDGTLVLAMKLMSKAFLQQLQDLSQQPSFCKLWLGVLDHMD
KYMKLKLRGKRSDKIHELIPELLKNNLLVMKTTGILLPTDDIGGDSFWQLTWLHVKKISPSMQSEVFPDHELEQL
KAKLVTTGGTSATDGSVIVQSDENTACR*
**>Citsi_LOC102625158**
MVELHKEEDDNKCATYHDNKKDMDKYKRKQLGLSCMLNTEVGSVLAVIRRPLDAHYVQEDTFESAVVQSLKSLRS
LIFNPQQEWRTVDPSIYLSPFLDVVQSDDIPAAATGVALSAILKILKLEIFDEKTPGVKDAINIVVTGITSCQLE
KTDPISEDAVMMRILQVLIAIMRHRASILLTDEAVCTIVNTCFHVVQQSASRGDLLQRSARYTMHELIQIIFSRL
PDIEVKSGEGSESDTEDVDMDANLGSGYGIRSAVDIFHFLCSLLNVVELVEGEGSRTSDVDVQLFALVLINSAIE
LSGDAIGKHPKLLRMVQDDLFHHLIHYGARSSPLVLSMICSTVLNIYHFLRRFIRLQLEAFFGFVVLRVAASGNS
HQLQEVALEGIINFCRQPTFLIEVYVNYDCDPLCRNVIEEIGKLLCKHSFPVSGPLTSSQIQAFEGLVILIHNIA
ESIDKEGDTSPSGPYPVEITEYKPFWEEKPNDDSDTWVEYVRLRKAQKRKSLIAGNHFNRDEKKGLEYLKLCQLV
SDPPDPKALAFFFRFTQGLDKNMIGDYLGDADEFHIQVLKEFTETFEFAGMTLDNALRTYLETFRLPGESQKIQR
ILEAFSDRFFDQQTSEIFVAKDSVYIFCYSLIMLNTDQHNPQVKKKMTEEEFIRNNRGINGGKDLPREYLSELFH
SIASNAISVFGQSGQIVDMNPSRWIELINRSKTMLPFILCDFDRRLGRDMFASIAGPAVAALSAFFDHADEDDML
QECIEGLISISRIAQYGLEDTLDELLASFCKFTTLLNPYATAEETLFAFSNDMKPKMATLAVFTLANNFGNSIRA
GWRNIVDCLLKLKRLKLLPQSVIEFDISTTDAPSHSRAESGVVFPAYDPTSGNRRSSGMISRFTHFLSLDSPEDS
ISLGMNEFEQNLKVIKQCQIGNIFSNSTNLPLEALQNLGRSLIFAAAGKGQKFSTPVEEEETVGFCWDLIIAIAI
ANNNRFQAFWPSFHDYLLLVTQFPLFSPIPFAEKAMVGLFKVCLRLLSSYQSDKLPEELIFKSINLMWKLDKEIL
DTCSQLITQSVSKIIIEYPANLQSAVGWKSVLHLLSVTGRHPDTHEQAVETLIMLISDGTHISKATYAYCIDCAF
SFVALKNSPLEKNLKILDLLSDSVNLLIQWYKNAWSESGNNYSIASSTSTSSLEDYKGLNSLNFAVNLFIKLGEA
LRKTSLARREEIRNHAVLALQKCFTLAEDLDFSSINCINCFNLVIFAMVDDLHEKMIEYSRRENAEREMRSMEGT
LKNAMELLANVFLQFIKQIAESPGFRTFWLGVLRRMDTCMKADLGPYGETKLQETIPDLLRNMITMMKEREILAP
KEDEDLWEITYIQIQWIAPSLKEELFPDEI*
>**Anaoc_0003s0363**

MGRLKLQSGIKAIEEEPEDFDATYSNKTTLACMINAEVGAVLAVMRRNRNVRWGGQYMSGDDQLEHSLIQSLKTL
RKQIFSWQHPWHTINPAVYLHPFLDVIRSDETGAPITGIALSSVYKILTLDVIDQDTVNVEEAMHLVVDAVTSCR
FEVTDPASEEVVLMKILQVLLACMKSKASVMLSNQHVCTIVNTCFRIVHQAGSKGELLQRIARHTMHELVRRIFS
HLPDIGNTEHALVNGVAAVKQEIGGVDNDYACGSKQMENGNGNSEYEGHQSSANFVSATGVVANVVDENTIGAGS
GKEAFSYDLHLMTEPYGVPCMVEIFHFLCSLLNASEHIATGLRSNTLAFDEDVPLFALGLINSAIELAGPAFRCH
PRLLSLIQDELFRNLMQFGLSMSPLILSMVCSIVLNLYHHLRTELKLQLEAFFSCVILRLAQSRYGASYQQQEVA
MEALVDFCRQKTFMVEMYANLDCDITCSNVFEDLANLLSKSAFPVNCPLSAMHILALDGLIAVIQGMAERIGNGL
LSSEQPPVTLEEYTPFWMVKCDNYSDPNHWVPFVCRRKYIKRRLMIGADHFNRDPKKGLEFLQGTHLLPDKLDPQ
SVACFFRYTAGLDKNLVGDFLGNHDEFCVQVLHEFAGTFDFQDMNLDTALRLFLETFRLPGESQKIQRVLEAFSE
RYYEQSPQILVNKDAALLLSYSLIMLNTDQHNVQVKKKMTEEDFIRNNRHINGGSDLPRDFLSELYHSICKNEIR
TTPEQGAGFPEMNPSRWIDLMHKSKKTAPFIVADCRTYLDHDMFAIMSGPTIAAISVVFDHAEHEEVYQTCIDGF
LAVAKISACHHLEDVLDDLVVSLCKFTTLLNPSSVEEPVLAFGDDTKARMATVTVFTIANRYGDYIRTGWRNILD
CILRLHKLGLLPARVASDAADESELSSDPSHGKPLTNSLSSAHMPTMGTPRRSSGLMGRFSQLLSLDTEEPRSQP
TEQQLAAHQRTLQTIQKCHIDSIFTESKFLQAESLLQLARALIWAAGRPQKGSSSPEDEDTAVFCLELLIAITLN
NRDRIVLLWQGVYEHIASIVQSTVMPCALVEKAVFGLLRICQRLLPYKENLADELLRSLNLVLKLDARVADAYCE
QITQEVSRLVKANATHIRSQMGWRTITSLLSITARHPEASEAGFDALLFIMSDGAHLVPANYVLCVDASRQFAES
RVGQAERSVRALDLMADSVDCLARWSHEAKEALGEDEAAKLSQDIGDMWLRLVQGLRKVCLDQREDVRNHALLSL
QKCLTGVDEIHLPHSLWLQCFDMVIFTMLDDLLEIAQGHSQKDYRNMEGTLILAMKLLSKVFLQLLHDLSQLSTF
CKLWLGVLSRMEKYMKVKVRGKKNEKLLEIVPDLLKNTLLVMKARGVLVQRSALGGDSLWELTWLHMNNIVPSLQ
SEVFPDQDSEQSQNKQCQTVGGSMSDQMTSVPSNETLAS*

>**Anaoc_0018s0976**

MGHVNMESGSNVMRKDFEYFSVKPSRGVLACMINSEIGAVLAVMRRNRNVRWGVRYVVDDDQLEHSLIHSLKELRKQ
IFLWQNQWHTIDPAVYLQPFLDVIQSDETGAPITGVALSSVYKILTLDVLDLDTVNVGDAMLLMVDAVTSCRFEV
TDPASEEVVLMKILQVLLACMKSKAAVKLSNQHVCNIVNTCFRVVHQASSKGELLQRIARHTMHELVRCIFSHLP
DIDGPEHSVVDRNWSGTGEKVGMEKDLTFGNKLPENGNVSVENDGQSSTEYMSSASQMNTNKVDTSSGTEAAQND
EKIMMEPFGVPCMVEIFHFLCSLLNVSENMGVGPRPNPIAYDEDVPLFALGLINSAIELGGSSFSKHPTLLALIQ
DELFRNLMQFGLSMSPLILSTVCSIVLNLYHHLRTELKAQLEAFFSCVLLRIAQSKYGSSYQQQEVAMEALVDLC
RQQTFMTEMYANFDCDITYSNLFEDLVSLLSKSAFPVNGPLSAMHVLALDGLISMIQGMAERMSNELLAPEGASI
DPEEYNAFWSLKCENYGDPNHWVPYVRKMKHIKRGLMVGADHFNQDCKRGLEYLQGIHLLPDKLDPQSVAFFFRY
TTGLDKNLIGDFLGNHDEFCVKVLQEFAGTFNFHGMNLDTALRLFLGTFRLPGESQKIQRVLEAFAERYYEQSPD
ILTNKDAALLLSYSLIMLNTDQHNVQVKKKMTEDDFIRNNRHINGGKDLPREYLSELYHSICENEILMIPEQAVC
FPAMTSSRWINVMHKSRETIPFITSDSTALLDHDMFAILSGPTIAAMSVVFDQVEQEDILQRCIDGFLAIAKIST
CCHFGDVLDDLVVSLCKFTTLLTPLSVEEAVLAFGDDTKARMALTTVFTIANRYGDYIHSGWKNILDCILSLHEL
GLLPDRLARDAADDMEPSSDMERAKPATGSLSALHVTPVATPRKSSGLISRFSQLLSFDIEEPRLQPSEEEIAAK
QRTHEIIQNCHIDSIFMESKFLQTESLMELVKALILAAGRLQKGSSSMEDEDTGVFCLELLIAITLNNRDRIMLI
WQGVYEHIASIVQSTVMPCKLVEKAVSGLLRICQRLLPYKENLTDELLKSLQLILKLDARVADAYCENITQEVMR
LVKANATQIRSHVGWRTIISLLSITARHPEASEAGFEALAFIMTDGVHLLPSNYILCVDAARQFAESRVGEIDRS
VSALDLMGGSVFCLVRWSTETKNTVEEEAAKKLSQDIGEMWLRLVQGLKKVCLDQREKVRNHAVLVLQRSLAAVD
GIHLPSALWFQCFDTEIFTLLDNLLEIAQANSPKDYRNIDGTLVLAMKLMSKAFLQSLEDLSKQPSFCRLWLGVL
NHMDNYMRVKLRGKQSEKIHELVPELLKNSLLVMKTTGILLPTDDIGGDSFWQLTWLHVKKIVPSLQLEVFPDEE
VEKMKAKPKKAREVPAPDGTDPVPSSENAESR*

>**Anaoc_0010s1629**

MGHVNMGSGSNAICKDFKYCPVKPSRGGLACMINSEIGAVLAVMRRNRNVRWGVHYVADDDQLEHFLIHSLKELRKH
IFLWQNQWHTIDPAVYLKPFLDVIQSDETGAPITGVALSSVYKILTLDVLDLDTVNVGAAMYLIVDAVTSCRFEV
TDPASEEVVLMKILQVLLACMKSKAAVKLSNQHVCNIFNTCFRIVHQASTKGELLQRIARHTMHELVRCVFSHLP
DINSPEHFLVNRGWSGAGAGAGEKVGVEKDLAFGNRPLENGNVSVENDGQSSNECMSNASEMDANKVESSSGSEA
AQNGKKIMMEPFGVPCMVEIFHFLCSLLNVIENMGVGPRSNPIAYDEDVPLFALGLINSAIELGGSSFCKHPTLL
VLIQDELFRNLMQFGLSMSPLILSTVCSIALNLYHHLRTELKAQLEAFFSCVLLRIAQSKHGSSYQQQEVAMEAL
VDLCRQQTFMTEMYANFDCDITSGNLFEDLVNLLSKSAFPVNGPLSAMHVLALDGLISMIEGMAERMSNELSAPE
GASIDPDEYDAFWRLKCEDYGDPSYWVPFVRKMKYIKRRLMVGADHFNRDSEKGLEYLQGMHLLPDKLDPQSVAF
FFRYTTGLDKNLIGDFLGNHDEFCVQVLQQFAGTFNFHGMNLDTALRLFLGTFRLPGESQKIQRVLEAFSERYYE
QSPDILSNKDAALLLSYSLILLNTDQHNVQVKKKMTEDDFIRNNRQINGGKDLPREYLSELYHSIFENEILMIPE
QGVSFPAMTSSRWINVMHKSKETTPFILCHSTVLLDYDMFAILSGPTVAAMSVVFDQVEQEGILQRCIDGFLAIA
KISACYHFGDVLDDLVLSLCKFTTLLTPLSLEEAILALGDDTKARMAMTTVFTIANRYGDYIHSGWKNILDCVLS
LHKLGLLPAHLARDAADDMEPSSDVEQGKPTTGSLSTLNVTPVATLQKSSGLIGRFSQLLSFDIEEPRLQPSEEE
IAAKQRTHEVIQNCHIDSIYIESKFLQADSLVELVKALILAAGRLHKGSSSVEDEDTGVFCLELLIAITLNNRDR
IILIWQGVYEHIASIVQSTVMPCKLVEKAVFGLLRICQRLLPYKENLTDKLIKSLQLILKLDARVADAYCEHITQ
EVMRLIKANATHIRSHAGWRTIISLLSITARHPDASEAGFEALAFIMSERVHLLPSNYILCVDAARQFAESRVGE
VDRSLSALDLMGGSVVCLVRWSTETKNMVGEEAAMKLSQDIGEMWLRLVQGLKKVCLDQREQVRNHAVLVLQTSL
AAVDEIHLPNTLWFHCFDMVILKLLDDLLEIAQANSPKDYRNIDGTLVLAMKVMSKAFLQSLQDLSQQPSFCKLW
LGVLNHMERYMKLKLRGKRSEKIHELVPELLKNNLLVMKTAGILLPTDDIGGDSFWQLTWLHVKQIVPSLELEVF
PDHEVAQIKAKHIKTGAVPAPDGSDVVPLGESTESR*

**>Anaoc_0019s0415**
MGELYGEEDNENSKMEGKKDMEKCKRKELGLSCMLNTEVGSVLAVIRRPLDPSSQYIQEESFDSSLVQSLKSLRP
LIFNPHQEWRTVDPSIYLSPFLDVIQSGDVPAAATGVALSAVLKIFKVGIFDEKTPGVKDAINSVVTAVTSCQLE
KTDRVSEDAVMMRILQVFIHVMKHPASIFLTDDAVCIIVNSCFQVVQQSANRGDLLQRSARYTMHELMQVIFSRL
PEIEVKSGEASESDSEDVDFDRNLDSGYGVRSAVDIFHFLCSLLNVVQILETEGYHSAEEDVQLFALVLINSAIE
LSGDAIAKHTKLLRMIQDDLFHHLIHYGEGSSPLVLSMICSTVLNIYHFLRRFIRLQLEAFFSFVLLRIAARGRS
LQLQEVALEGIINFCRQPTFIVEAYVNYDCDPLCRNILGEIGRLLCKHSFPGTAPLTSSQMQAFEGLVILIHTIA
EKVDKEGDSTPSGPYPVEITEYRPFWEEVPKGDDLNAWVQSVRLRKAQKKKVLIASNHFNRDENKGLEYLKICQL
VSDPPDPKAYAFFFRYTHGLDKNFIGDFVGDPDEFHIQVLKEFTETFEFSGMVLDSALRIYLETFRLPGESQKIQ
RILEAFSEKFYDQQSSEIFVSKDSVLVLCYSLIMLNTDQHNPQVKKKMTEEEFIKNNRAINGGKDLPREYLSELF
HSIANNEITLLDQSGRSIEMNPSRWVELINRSRSMEPFILCDFDRRLGIDMFACIAGPSAATLCAFFEHADDELL
EECIEGLISISRIAQYGLQDTLDELLASFCKFTTLLNPYATPEETMFAFSNDMKPKMATLAVFTLANNFGDSIRG
GWRNIVDCLLKLKKLMLLPQSVVEFDTSDVPSHSRSESGVLFRTHEAKYGIGSRQTSGMISRFTHFLSLDSPEDA
ISHGMNELEQNLKVIKQCRIGNIFSNSSNLAVEALVNLGRSLIFSAAGKGQKFSTPVEEEETVGFCWDLITAIAL
ANINRFQTFWFYFHEYLLDVAQFPLFSPTPFAEKAIVGLFKVCLRLLSSYQIDKLPEELIFKSINLMWKLDKEIL
ETCCQFITQSVSKIIVEYPANLQSLLGWKTVLHLLSVTGRHPETYDQGVETLIMLMSDATHVSRTNYAYCIDCAF
SFVALKNSPLEKNLKILDLMSESVNLLINWYKNACLESGTNISVASSSSASSTEDSKSHASLIINLFIKLGEALR
KTTLARREELRNHAVSSLHRSFTLAEQLDLSSTNIINCFNLEIFAMVDDLHEKMVEYSRRENAEKEMRSMEGTLK
LAIEFMTDVYLQYLKQLVESPGFRTFWLGLLRRMDTCMKADLGPYGDTKLQEVIPELLTKMITKMKEMEILVPKE
GDDLFEITYIQIQWIAPSLKEDLFPNDFCI*
**>Pisve_LOC116120515**
MGRLKLQSGIKAIEEEPEDYDTTYSNKATLACMINSEVGAVLAVMRRNRNVRWGGQYMSGDDQLEHSLIQSLKTL
RKQIFSWQHPWHTINPAVYLQPFLEVIRSDETGAPITGIALSSVYKILTLDVIDPDTVNVEEAMHLVVDAVTSCR
FEVTDPASEEVVLMKILQVLLACMKSKASVMLSNQHVCTIVNTCFRIVHQAGSKGELLQRIARHTMYELVRRIFS
HLPDIDNTEHALVNGVAAVKQEIGGVDYDYAFGSKQMENGNGNSEYDGQQSSANFVSATGVVANAMDENTIGAGT
GKEAVSYDLHLMTEPYGVPCMVEIFHFLCSLVNASEHMAMGLRSNTLAFDEDVPLFALGLINSAIELAGPAIRCH
PRLLSLIQDELFRNLMQFGLSMSPLILSMVCSIVLNLYHHLRTELKLQLEAFFSCVILRLAQSRYGASYQQQEVA
MEALVDFCRQKTFMVEMYANLDCDITCSNVFEDLANLLSKSAFPVNCPLSAMHILALDGLIAVIQGMAERIGNGL
LSSEQTPVTLEEYTPFWMVKCDNYSDPNHWVTFVRRRKYIKRRLMIGADHFNRDPKKGLEFLQGTHLLPDKLDPQ
SVACFFRYTAGLDKNLVGDFLGNHDEFCVQVLHEFAGTFDFQDMNLDTALRLFLETFRLPGESQKIQRVLEAFSE
RYYEQSPQILVNKDAALLLSYSLIMLNTDQHNIQVKKKMTEEDFIRNNRHINGGSDLPRDFLSELYHSICKNEIR
TTPEQGTGFPEMTPSRWIDLMHKSKKTAPYIVADCRAYLDHDMFAIMSGPTIAAISVVFDHAEHEEVYQTCIDGF
LAVAKISACHHLEDVLDDLVVSLCKFTTLLNPSSVEEPVLAFGDDTKARMATVTVFTIANRYGDYIRTGWRNILD
CILRLHKLGLLPARVASDAADESELSSDPSHGKPLTNSLSSAHMPSMGTPRRSSGLMGRFSQLLSLDTEEPRSQP
TEQQLAAHQRTLQTIQKCHIDSIFTESKFLQAESLLQLARALIWAAGRPQKGNSSPEDEDTAVFCLELLIAITLN
NRDRIVLLWQGVYEHIASIVQSTVMPCALVEKAVFGLLRICQRLLPYKENLADELLRSLNLVLKLDARVADAYCE
QITQEVSRLVKANATHIRSQMGWRTITSLLSITARHPEASEAGFDALLFIMSDGAHLVPANYVLCIDASRQFAES
RVGQAERSVRALDLMADSVDCLSRWSHEAKDAIGEYEAAKLSQDIGDMWLRLVQGLRKVCLDQREDVRNHALLSL
QKCLTGVDEIHLPHSLWLQCFDMVIFTMLDDLLEIAQGHSQKDYRNMEGTLILAMKLLSKVFLQLLHDLSQLSTF
CKLWLGVLSRMEKYMKVKVRGKKSEKLLEIVPDLLKNTLLVMKARGVLVQRSALGGDSLWELTWLHVNNIVPSLQ
SEVFPDQDSEQSQNRQSEAGGGLGSDQMTSVPSSETLAS*
**>Pisve_LOC116112061**
MGHVNMESGSSAICKDFKSCPVKPSRGALACMINSEIGAVLAVMRRNVRWGVRYVTDDDQLEHTLIHSLKELRKQ
IFLWQNQWHTIDPAVYLQPFLDVIQSDETGAPITGVALSSVYKILTLDVLDLDTVNVGDAMHLIVDAVTSCRFEV
TDPASEEVVLMKILQVLLACMKSKAAVKLSNQHVCNIVNTCFRVVHQASSKGELLQRIARHTMHELVRCIFSHLH
DVDSPEQSLVDRSWSGAGEKVGLEKDLTFGNKPLENGNVSVENDGQSSTEYMSNANEMDANKVEISSGMEAAQNG
EKIMMEPFGVPCMVEIFHFLCSLLNVIENMGVGPRPNPIAYDEDVPLFALGLINSAIELGGSSFSKHPTLLALIQ
DELFRNLMQFGLSMSPLILSTVCSIVLNLYHHLRTELKAQLEAFFSCVLLRIAQSKHGSSYQQQEVAMEALVDLC
RQQTFMTEMYANFDCDITSGNLFEDLVNLLSKSAFPVNGPLSAMHVLALDGLISMIQGMAERMSNELPAPEGASI
DPEEYNAFWSLKCEDYGDPYYWVPFVRKMKYIKRRLMVGADHFNRDSKKGLEYLQGMHLLPDKLDPQSVAFFFRY
TTGLDKNLIGDFLGNHDEFCVQVLQEFAGTFNFQGMTLDTALRLFLGTFRLPGESQKIQRVLEAFSERYYEQSPD
ILTNKDAALLLSYSIILLNTDQHNVQVKKKMSEDDFIRNNRQINGGKDLPREYLSELYLSICENEILMIPDQGVG
FPAMTSSRWINVMHKSKETTPFIMCDSTAFLDHDMFAILSGPTIAAMSVVFEQVEQEDILQRCIDGFLAIAKIST
CCHFGGVLDDLVLCLCRFTALLNPLSVEEAVLAFGDDTKARMAMTTVFTIANRYGDYIHCGWKNILDCVLSLHKL
GLLPARLARDAADDMEPSSDVEQGKPATGSLSTSHVTPLATSRKSSGLIGRFSQLLSFDIEEPTLQPSEEEIAAK
QRTHEIIQNCHIDSIFTESKFLQAESLMELVKALILAAGRLHKGSISMEDEDTGVFCLELLIAITLNNRDRIMLI
WQGVYEHIASIVQSTVMPCKLVEKAVFGLLRICQRLLPYKENLTDELLKSLHLILKLDARVADAYCEHITQEVMR
LVKANATHIRSHAGWRTIISLLSITARHPEASEAGFEALAFIMSDGVHLLPSNYILCVDAARQFAESRVGEVDRS
VSALDLMGGSVVCLVRWSTETKNTVGEEAALKLSQDIGEMWLRLVQGLKKVSLDQREEVRNHAVLVLQRSLAAVD
GIHLPNALWFQCFDMAIFTLLDDLLEIAQANSPKDYRNIDGTLVLAMKLMSKAFLQSLQDLSQQPSFCKLWLGVL
NHMEGYMKLRLRGKRSENIHELVPELLKNNLLVMKTAGILLPTDDIGGDSFWQLTWLHVKKIAPSLLSEVFPDHE
VEQLKAKHIKAGEGPAPDGSEVVQSGESTESR*
**>Pisve_LOC116127888**

MGEEDNEDNKMEGKKDMEKFKRKELGLSCMLNTEVGSVLAVIRRPHDPSTQYIQEDGYDSSLVQSLKSLRSLIFN
PQQEWRTVDPAIYLSPFLDVIQSGDIPAAATGVALSAVLKIFKVGIFDEKTPGVKDAINSVVTAITSCQLEKTDR
VSEDAVMMRILQVFIYIMKHPASVLLTDNAVCIIVNSCFQVVQQSANRGDLLQRSARYTMHELIQIIFSRLPEIE
VKSGEASESDSEDFDSDRNLDSGYGVRSAVDIFHFLCSLLNVVQIMETEGSHSADEDVQLFALVLINSSIELSGE
AIGKHPKLLRMIQDDLFHHLIHYGEGSSPLVLSMICSTVLNIYHFLRRFIRLQLEAFFAFVLFRVAARGRSLQLQ
EVALEGIINFCRQPTFIVEAYVNYDCDPVCRNIVEEIGKLLCKHSFPGAGPLTSSQIQAFEGLVILIHTIAENID
KEGDSSPSGPYPVEITEYRPFWEEMPKDDGLEAWVHSVRQRKTQKRKIVIAGHHFNRDENKGLEYLKLSHLVSDP
PDPKAYAFFFRYTHGLDKNFIGDFLGDPDEFHIQVLKEFTETFEFSGMILDNALRTYLETFRLPGESQKIQRILE
AFSERLYDQQSSEIFVSKDSVFILCYSLIMLNTDQHNPQVKKKMTEEEFIKNNRAINGGKDLPREYLSELFHSIA
NNEITLFGKSGCPIDMNPSRWIELMNRTRSMKPFTLCDFDRRLGRDMFASIAGPSVAALSAFFEHADEDELLQEC
IEGLISISRIAQYGLEDTLDELLASFCKFTTLLNPYASAEETLFAFSNDMKPKMATLAVFTLANNFGDSIRGGWR
KIVDCLLKLKRLMLLPQSVIEFDTSSTSDVPPHSRTESGVVFTTHETKYGIGSRHTSGIIGRFTHFLSLDSPEDA
MSLGMNELEQNLKVIKQCRIGNIFSNSSNLPAEALLNLGRSLIFSAAGKGQKFSTPVEEEETVGFCWDLITAIAL
ANINRFQTFWFYFHEYLLDVAQFPLFSPVPFAEKAIIGLFKVCLRLLSYYQVDKLPEELIFKSINLMWKLDKEVL
ETCCPFITQSVSKIIVEYPANLQSSLGWKTVLHLLSVTGRHPETYDQGVETLIMLMSDATHVSRTNYAYCIDCAF
SFVALKNTPLEKNLKILDLMSDSVNLLIQWYKNACLESGTSVSVASISSAFSFEDTKSLASLNFAINLFIKLGEA
LRKTSLARREELRNHAVLSLHRSFTLAEDLDISSTNIINCFNLVIFAMVDDLHEKMVEYSRRENAEKEMRSMEGT
LKLAMEFMADVYLQYVKQLVESPGFRTFWLGLLRRMDTCMKADLGPYGDTKLQELIPELLTKMIAKMKEMEILVP
REGDDLWEITYIQIQWIAPSLKDDLFPNDFCI*

**>Aceya_EZV62_017872**

MGRLKLQSGIKSIEEEPEDYETTYSNKDTLACMINSEVGAVLAVMRRNVRWGGQYMSGDDQLEHSLIQSLKSLRK
QIFSWQHPWHTINPAIYLQPFLDVIRSDETGAPITGIALSSVYKILTLDVIDQNTVNVEEAMHLVVDAVTSCRFE
VTDPASEEVVLMKILQVLLACMKSKASVMLSNQHVCTIVNTCFRIVHQAGSKGELLQRIARHTMHELVRRIFSHL
LDIDNTEHALVNGVSAIKQEIGGRDNDYAFGNKQVENGNGSSEQDGQLSSANFVSNASTGVVATMMEENTIGAVT
DAASYDLHLMTEQYGVPCMVEIFHFLCSLLNVSEHMGIGPRSNTIAFDEDVPLFALGLINSAIELGGPAIRRHPR
LLSLIQDELFRNLMQFGLSMSPLILSMVCSIVLNLYHHLRTELKLQLEAFFSCVILRLAQSRYGASYQQQEVSME
ALVDFCRQKTFMVEMYANLDCDITCSNVFEDLANLLSKSAFPVNCPLSAMHILALDGLIAVIQGMAERIGNGLVS
SEYTPVTLEEYTPFWMVKCDNYSDPNHWVQFVRRRKYIKRRLMIGADHFNRDPKKGLEFLQGTHLLPDKLDPQSV
ACFFRYTAGLDKNLVGDFLGNHDEFCVQVLHEFAWTFDFQDMNLDTALRLFLETFRLPGESQKIQRVLEAFSERY
YEQSPQILANKDAALLLSYSLIMLNTDQHNVQVKKKMTEEDFIRNNRHINGGSDLPRDFLTELYSICKNEIRTTP
EQGAGFPEMNPSRWIDLMHKSKKTAPFIVADCRAYLDHDMFAIMSGPTIAAISVVFDHAEHEDVYQTCIDGFLAV
AKISACHHLEDVLDDLVVSLCRFTTLLNPSSVEEPVLAFGDDTKARMATVTVFTIANRYGDYIRTGWRNILDCIL
RLHKLGLLPARVASDAADESELSADPSHGKPITNSLSSAHMPTMGTPRRSSGLMGRFSQLLSLDTEEPRSQPTEQ
QLAAHQRTLQTIQKCHIDSIFTESKFLQAESLLQLAKALIWAAGRPQKGNSSPEDEDTAVFCLELLIAITLNNRD
RIVLLWQGVYEHIANIVQSTVMPCALVEKAVFGLLRICQRLLPYKENLADELLRSLQLVLKLDARVADAYCEQIT
QEVSRLVKANATHIRSQMGWRTITSLLSITARHPEASEAGFDALLFIMTDGAHLVPANYVLCVDSSRQFAESRVG
QAERSVRALDLMSGSVACLAGWGREAKEAMGEDEASKLSQDIGEMWMRLVQGLRKVCLDQREDVRNHALLSLQKC
LTSVDEIHLPHGLWLQCFDMVIFTMLDDLLEIAHGHSQKDYRNMEGTLILAMKLLSKVFLQLLHDLSQLTTFCKL
WLGVLSRMEKYMKVKVRGKKSEKLQEIVLELLKNTLLVMKTRGVLVQRSALGGDSLWELTWLHVNNIVPSLQSEV
FPDQDLEQSQYKQSEIGGGLVSDEIGSGSETAASEGTGTGG*

**>Aceya_EZV62_015425**

MGHLNPRSGSDAFYKEKLKDCSIKLSRGAIACMINSEIGAVLAVMRRNVRWGVRYEADDDQLEHSLIHSLKELRK
QIFLWQNQWHNIDPAVYLQPFLDVIRSDETGAPITGVALSSVYKILTLDVLDLDTVNVGDAMHIIVDAVTSCRFE
VTDPASEEVVLMKILQVLLACMKSKVAVKLNNQHVCNIVNTCFRVVHQASSKGELLQRIARHTMHELVRCIFSYL
PDIDCPEQSSVAQSQSDDSEKVRVSVLEKDHTSESKPFENGNVSAESQSSTGHKSSAPTYKLASEVDTNNGEIGG
MMEAAQNGEKIMMEPFGVPCMVEIFHFLCSLLNVIEHMGGGTRSNPMAYDEDVPLFALGLINSAIELAGSSFSRH
PKLLALIQDELFRNLMQFGLSMSPLILSTVCSIVLNLYHHLRIELKVQLESFFSCVLLRLAQSKYGSSYQQQEVA
MEAIVDLCRQQTFMTEMYANFDCDIISNNLFEDLANLLSKSAFPVNGPLSAMHVLALDGMISLIQGMAERVSNEL
PTPEGASVDPEEYNAFWNLKCKDYSDPHSWVPFVRKMKYIKRRLAVGADHFNRDPKKGLEFLQGINMLPDKLDPQ
SVAFFFRYTTGLDKNLIGDFLGNHDEFCVQVLQKFAGIFDFRDMNLDTALRLFIGTFRLPGESQKIQRVLEAFAE
RYYEQSQDILTNKDAALLLSYSLILLNTDQHNVQVKKKMTEEDFIRNNRHVNGGKDLPREYLSELYYSICENEIL
MIPDQGSGFQEMTSSRWISVLHKSREATPFIVCDSGAHLDHDMFAILSGPTIAAMSVVFDQVEQEDILRRCVDGF
LAIAKVSTCYHFDEVLDDLVLSLCRFTTLLTPLSVEEALLAFGDDTKARMVLTAVFTIANRYGDHIHSGWKNILD
CVLSLHKLGLLPARLASDAADDMEPSSDLERGKPATSSVSTSRITPVVTPRKSSGLIGRFSQLLSFDMEEPRSQP
DEEELAAHQRAREIMQNCHIDSIFTESKFLQAESLLQLVRALILAAGQLRKGSSYMDDEDTGVFCLELLIAITLN
NRDRIMLIWQGVYEHIANIVQSIVMPCMLVEKAVFGLLRICQRLLPYKENLTDELLKSLQLILKLDARVADAYCE
QITQEVMRLVKANATHIRSHAGWRTIISLLSITARHPEASEAGFEALAFIMSEGTNILPSNFILCVDAARQFAES
RVGEVDRSVSAIELMGGAVVCLVRWSSETKNTVGEEAAIKLSQDIGEMWLRLVQALKKVCLDQREEVRNHAIQVL
QRSLAAVDGIHLPNVLWFQCFDMVIFTLLDDLLEIAQGNSPKDYRNIDGTLVLAMKLMSKAFLQLLQDLSLQPSF
CKLWLGVLNHMERYMTLKLRGKRSEKIHELIPELLKNNLLVMKTTGILMPTDDIGGDSFWQLTWLHVKKISPSIE
SEVFPEDELEQLKAKHTQTGGMPAAVGFVVVPPGENTACR*

**>Aceya_EZV62_012732**

MANLHGEEEGDDNNCSSCRDKKRHTATTRKKELGLGLSCMLNTEVGAVLAVMRRPLDSGTQYIQEDTFDSSIMQS
LKSLRSLIFSPQQEWRLLDPSLYLSPFLDVIQSDDIPPTATGIALSAVLKILKLEIFDEKTPGARDAINSAVTGI
TSCQLEKTDPISEDAVMMRILQVLTSIMKHPASVLLSDQAVCTIVNTCFQVVQQSASRGDLLQRSARYTMHELIQ
IIYSRLPDIEVKSGEASESDTEDVDVDSNLDSGYGVRSAVDIFHFLCSLLNVVEIVETEGSTFQTADEDVQLFAL
VLINTAIELSGDAIGKHSKLLRMIQDDLFHHLIHYGDGSSPLVLSMICSTVLNIYHFLRRFVRLQLEAFFSFVLL
RVSTLQSSLQLQEVALEGIINFCRQPTFLIEIYVNYDCDPLSRNVMEEIGKLLCKQSFPGGGPLTSSQIQAFEGL
VIMIQNIADNIDKEEDSSPSGPYPVEITEYRPFWEEKPKEDLDTWVEYARIRKVQKRKILIAANHFNRDEKKGLE
YLKFCQLVSDPPDPKALAFFFRYTPGLDKSMIGDYLGDPDEFQIQVLKEFTRTFEFVGMILDNALRTYLETFRLP
GESQKIQRILEAFSERFYDEQSSDIFVTKDAVFILCYSLIMLNTDQHNPQVKKKMTEDEFIRNNRAINGGKDLPR
EYLSELFHSIATNAITLFGQSGQPIEMNPSRWIELINRSKAAEPFILSDFDRRLGRDMFACIAGPSVAAVSAFFE
HADEDELLHECIEGLISIARIAQYGLEDTLDELLASFCKFTTLLNPYASTEETLFAFSHDMKPRMATLAVFTLAN
NFGDSIRAGWRNIVDCLLKLKRLKLLPQSVIEFDPASDVQSDSTNESGALLPTHDPNFRKHQNSGIISKFTHFLS
LENPEESLSLGMNEFEQNLKVIKQCQIGNIFSNTSKLPYDALLNLGRSLIFAAAGKGQKFSTPVEEEETVGFCWD
LIIAIAMANINRFHIFWPSFHDYLLGVAQFPLFSPIPFAELAIVGLFKVCLKLLGSYQADRFPEELIFKSINLMW
KLDKEVLDTCCQYITQSVSTIIIEYPANLQSQLGWKTVLHLLSVTGRHPETYNQAVETLIMLLSDGTHVSRINYA
YCIDCAFGFVSLKNSPLEKNLKLLDLMSDSVSSLIQWYQNACSEPGINYSIASNISTSSMDDNKGLNSINLPMNL
FVKLGEALRKTSLARREEIRNHAVSSLHKSFKLAEELDFSSINCINCFNLVIFSMVDDLHEKMIEYSRRENAERE
MRSMEGTLKIAMELLTDVYLHFLKQIAETQEFRTFWLGVLRRMDTCMKADLGPYGETKLQEFIPELLRKMITKMQ
EMEVLVQREDDDLWDITNIQIQWIAPSLKEELFPEEF*

**XansoGNOMa**

MGRLKLQSGIKAIEEEPEDYETTYSNKATLACMINSEVGAVLAVMRRNRNVRWGGQYMSGDDQLEHSLIQSLKSL
RKQIFSWQHPWHTINPAIYLQPFLDVIRSDETGAPITGIALSSVYKILTLDIIDHNTVSVEEAMHLVVDAVTSCR
FEVTDPASEEVVLMKILQVLLACMKSKASIMLSNQHVCTIVNTCFRIVHQAGSKGELLQRIARHTMHELVRRIFS
HLLDVDNTEHALVNGVSTVKQEIGGRDNDYAFGNKQVENGNGSSEYDGQLSSATFVSNATTGVVATMMEENTIGA
GTGKDVSYDLHLMTEPYGVPCMVEIFHFLCSLLNVAEHMGIGPRSNTIAFDEDVPLFALGLINSAIELGGPAIRR
HPRLLSLIQDELFRNLMQFGLSMSPLILSMVCSIVLNLYHHLRTELKLQLEAFFSCVILRLAQSRYGASYQQQEV
SMEALVDFCRQKTFMVEMYANLDCDITCSNVFEDLANLLSKSAFPVNCPLSAMHILALDGLIAVIQGMAERIGNG
LVSSEHTPVTLEEYTPFWMVKCDNYNDPNHWVPFVRRRKYIKRRLMIGADHFNRDPKKGLEFLQGTHLLPDKLDP
QSVACFFRYTAGLDKNLVGDFLGNHDEFCVQVLHEFAWTFDFQDMNLDTALRLFLETFRLPGESQKIQRVLEAFS
ERYYEQSPQILANKDAALLLSYSLIMLNTDQHNVQVKKKMTEEDFIRNNRHINGGGDLPREFLSELYHSICKNEI
RTTPEQGSGFPEMTPSRWIDLMHKSKKTAPFIVADCRAYLDHDMFAIMSGPTIAAISVVFDHAEHEDVYQTCIDG
FLAVAKISACHHledvlddlvvSLCKFTTLLNPSSVEEPVLAFGDDTKARMATVTVFTIANRYGDYIRTGWRNIL
DCILRLHKLGLLPARvasdaadesevsadPSHGKPITNSLSSAHMPSMGTPRRSSGLMGRFSQLLSLDTEEPRSQ
PTEQQLAAHQRTLQTIQKCHIDSIFTESKFLQAESLLQLARALIWAAGRPQKGNSSPEDEDTAVFCLELLIAITL
NNRDRIVLLWQGVYEHIANIVQSTVMPCALVEKAVFGLLRICQRLLPYKENLADELLRSLQLVLKLDARVADAYC
EQITQEVSRLVKANAAHIRSQMGWRTITYLLSITARHPEASEAGFDALLFIMSEGAHLLPANYVLCVDSSRQFAE
SRVGQAERSVRALDLMSGSVDCLARWGCEAKEAMGEDEASKLSQDIGEMWLRLVQELRKVCLDQREDVRNHALLS
LQKCLTGVEEIHLPHGLWLQCFEMVIFTMLDDLLEIAQGHSQKDYRNMEGTLILAMKLLSKVFLQLLHDLSQLTT
FCKLWLGVLSRMEKYMKVKVRGKKSEKLQEIVPDLLKNTLIVMKTKGVLVQRSALGGDSLWELTWLHVNNIVPYL
QSEVFPDQDLEQSQHKQSEIGGGLVSDEIGSGSETAASEGTGTGG*

**>XansoGNOMb**

MGRLKLQSGIKAIEEEPEDYETTYSNKATLACMINSEVGAVLAVMRRNRNVRWGGQYMSGDDQLEHSLIQSLKSL
RKQIFSWQHPWHTINPAIYLQPFLDVIRSDETGAPITGIALSSVYKILTLDIIDHNTVSVEEAMHLVVDAVTSCR
FEVTDPASEEVVLMKILQVLLACMKSKASIMLSNQHVCTIVNTCFRIVHQAGSKGELLQRIARHTMHELVRRIFS
HLLDVDNTEHALVNGVSTVKQEIGGRDNDYAFGNKQVENGNGSSEYDGQLSSATFVSNATTGVVATMMEENTIGA
GTGKDVSYDLHLMTEPYGVPCMVEIFHFLCSLLNVAEHMGIGPRSNTIAFDEDVPLFALGLINSAIELGGPAIRR
HPRLLSLIQDELFRNLMQFGLSMSPLILSMVCSIVLNLYHHLRTELKLQLEAFFSCVILRLAQSRYGASYQQQEV
SMEALVDFCRQKTFMVEMYANLDCDITCSNVFEDLANLLSKSAFPVNCPLSAMHILALDGLIAVIQGMAERIGNG
LVSSEHTPVTLEEYTPFWMVKCDNYNDPNHWVPFVRRRKYIKRRLMIGADHFNRDPKKGLEFLQGTHLLPDKLDP
QSVACFFRYTAGLDKNLVGDFLGNHDEFCVQVLHEFAWTFDFQDMNLDTALRLFLETFRLPGESQKIQRVLEAFS
ERYYEQSPQILANKDAALLLSYSLIMLNTDQHNVQVKKKMTEEDFIRNNRHINGGGDLPREFLSELYHSICKNEI
RTTPEQGSGFPEMTPSRWIDLMHKSKKTAPFIVADCRAYLDHDMFAIMSGPTIAAISVVFDHAEHEDVYQTCIDG
FLAVAKISACHHLEDVLDDLVVSLCKFTTLLNPSSVEEPVLAFGDDTKARMATVTVFTIANRYGDYIRTGWRNIL
DCILRLHKLGLLPARVASDAADESEVSADPSHGKPITNSLSSAHMPSMGTPRRSSGLMGRFSQLLSLDTEEPRSQ
PTEQQLAAHQRTLQTIQKCHIDSIFTESKFLQAESLLQLARALIWAAGRPQKGNSSPEDEDTAVFCLELLIAITL
NNRDRIVLLWQGVYEHIANIVQSTVMPCALVEKAVFGLLRICQRLLPYKENLADELLRSLQLVLKLDARVADAYC
EQITQEVSRLVKANAAHIRSQMGWRTITYLLSITARHPEASEAGFDALLFIMSEGAHLLPANYVLCVDSSRQFAE
SRVGQAERSVRALDLMSGSVDCLARWGCEAKEAMGEDEASKLSQDIGEMWLRLVQELRKVCLDQREDVRNHALLS
LQKCLTGVEEIHLPHGLWLQCFEMVIFTMLDDLLEIAQGHSQKDYRNMEGTLILAMKLLSKVFLQLLHDLSQLTT
FCKLWLGVLSRMEKYMKVKVRGKKSEKLQEIVPDLLKNTLIVMKTKGVLVQRSALGGDSLWELTWLHVNNIVPYL
QSEVFPDQDLEQSQHKQSEIGGGLVSDEIGSGSETAASEGTGTGG*

**>Xanso_XS14G07389**

MGNLNPQSGSDAFYKEKLKDCPIKPSRGAIACMINSEIGAVLAVMRRNVRWGVRYVADEDQLEHSLIHSLKELRK
QIFLWQNQWHNIDPAVYLQPFLDVIRSDETGAPITGVALSSVYKILTLDVLDLDTVNVGDAMYLIVDAVTSCRFE
VTDPGSEEVVLMKILQVLLACMKSKAAVKLSNQHVCNIVNTCFRVVHQASSKGELLQRIARHTMHELVRCIFSHL
PDIDCPEQSSAVQSQSDAGEKSVLEQDHTSESKPFENGNVSVESQSSTGHVSSAPTNKLASEVDTNNGDIGGVME
ATQNGEKIMMEPFGVPCMVEIFHFLCSLLNVIEHMGAGSRSNPIAYDEDVPLFALGLINSAIELGGSSFSRHPKL
LALIQDELFRNLMQFGLSMSPLILSTVCSIVLNLYHHLRIELKVQLEAFFSFVLLRLAQSKYGSSYQQQEVALEA
LVDLCRQQTFMTEMYANFDCDIISGNLFEDLANLLSKSAFPVNGPLSPMHVLALDGMISMIQGMAERVSNELPTP
EGVSVDPDEYNSFWTLKCKDYSDPHSWIPFVRKIKYIKRKLAVGADHFNRDPKKGLEFLQGINLLPDKLDPQSVA
FFFRYTTGLDKNFIGDFLGNHDEFCVEVLHKFAGTFDFHGMNLDTALRLFIGTFRLPGESQKIQRVLEAFAERYY
EQSQDILTNKDAALLLSYSLILLNTDQHNVQVKKKMTEEDFIRNNRQINGGKDLPREYLSELYHSICENEILMVP
DQGSGFPEMTSSRWINVMYKSREATPFIVCDSGAHLDHDMFGILSGPTIAAMSVVFDQVEQEDILRRCIDGFLAI
AKVSTCYHFDEVLNDLVLSLCRFTNLLTPLSVEEALVAFGDDTKARMALTAVFTIANRYGDHIHSGWKNILDCVL
SLHKLGLLPAHLASDAADDMEPSSDLERGKPAISSVSTSRITPVVTPRKSSGLIGRFSQLLSFDMEEPSSQPDEE
EVAAHQRAREIMQNCHIDSIFTESKFLQAESLLQLVTALILAAGQLRKGSSYMEDEDTGVFCLELLIAITLNNRD
RIMLIWQGVYEHIANIVQSTVMPCTLVEKAVFGLLRICQRLLPYKENLTDEllkslqlvlklDARVADAYCEQIT
QEVMRLVKANATHIRSHAGWRTIISLLSITARHPEASEAGFEALAFIMSEGAHILPSNFILCVDAARQFAESRVG
EVDRSVSALELMGGSVICLVRWSSETKNAVGEEAAIRLSQDIGEMWLRLVQGLKKVCLDQREEVRNHAILLLQRS
LAAVDGSHLPNALWFQCFDMVIFTLLDDLLEIAQGNSPKDYRNIDGSLVLAMKLMSKAFlqllqdlslqPSFCKL
WLGVLNHMESYMKLKLRGKRSEKIHELIPELLKNNLLVMKTTGILLPTDDIGGDSFWQLTWLHVKKISPSMQSEV
FPDDELEQLKAKHTKTGGIPAPDESVVVPSGENPACG*
**>Xanso_XS04G12756**
MANLHGEEEEDDNNCSSCREKKKHMAKYKKKVLGLSCMLNTEVGAVLAVIRRPIDYIQEETFDSSILQSLKSLRS
LIFNPQQDWRNLDPSLYLSPFLDVIQSDDIPAAATGVALSAVLKILKLEIFNENTPGARDAINSAVTGITSCQLE
KTDPVSEDAVMMRILQVLTGIMKHPTSVLLTDQAVCTIVNTCFQVVQQSASRGDLLQRSARYTMHELIHIIYSRL
PDIEVKSGDTSESDTEDVDVDNNLDSGYGVRSAVDIFHFLCSLLNVVEIVETEGSTSQTADEDVQLFALVLINTA
IELSGDVIGKHTKLLRMIQDDLFHHLIHYGAGSSPLVLSMICSTVLNIYHFLRRFVRLQLEAFFAFVILRISTLG
SAPQLQEVALEGIINFCRQPTFLIEIYVNYDCDPLCRNVMEEIGKLLCKQSFPGGGPLTSLQIQAFEGLVIMIQK
IADNIDKEGDSSPSGPYPIEITEYRPFWEEKPKEDLDTWVDYLRVRKVQKRKILIAANHFNRDEKKGLQYIKFCQ
LVSDPPDPKAFAFFFRYTHGLDKNMIGEYLGDPDEFHIQVLKEFTRTFEFAGMNLDNALRTYLETFRLPGEAQKI
QRILEAFSDRFYEEQSLDIFVSKDAVYIFCYSLIMLNTDQHNPQVKKKMTEEEFIRNNRAINGGKDLPREYLSEL
FHSIATHAITIFGQSGQPIEMNPSRWIELMNRSKAVEPFILSDFDRRLGRDMFACIAGPSVAAVSAFFEHADEDD
VLHECIEGLISIARIAQYGLEDTLDELLASFCKFTTLLNPYASTDETLFAFSNDMKPRMATLAVFTLANNFGDSI
RAGWRNIVDCLLKLKRLKLLPQSVIEFENADSDVQSQSKTESGILLPTHDPKFGKHQNSGIINKFTHFLSLENPE
DSLSLGLNEFEQNLKVIKQCQIGNIFSNTSKLPSDALLNLGRALIFAAAGKGQKFSTSVEEEETVGFCWDLITAI
SVANISRFQILWPSFHDYFLGVVQFPLFSPIPFAEKAIVGLFKVCLKLLASYQADKFPEELIFKSINLMWKLDKQ
VLDTCCQYITQSVSKIIIEYPANLQSQLGWKSVLHLLSVTGRQPETYDQAVETLIMLMSDATHVSQTNYAYCIDC
AFGFVALKNSPLEKNLKLLDLMSDSVDSLVQWYKNACSEPGNNYSIASNTSSSSLEDNKGLNSLNFSMNCFVKLG
EALRKTSLARREEIRNQAVISLHKSFKLAEELDFSSINCINCFNLVIFAMVDDLHEKMIEYSRRENAEREVRSME
GTLKNAMELLADVYLQFLKEIAEAQEFRTFWLGVLRRMDTCMKADLGEYGDTKLQEYIPDLLRKMITKMQEMEIL
VRKDDDDLWDITYIQIQWIAPSLKEELFPEEI*
**>Dimlo_026160**
MGRLKLQSGIKAIEEEPEDYETTYSNKATLACMINSEVGAVLAVMRRNRNVRWGGQYISGEDQLEPSLIQSLKSL
RKQIFSWQHPWHTINPAIYLQPFLDVIRSDETGAPITGIALSSVYKILTLDVIDQNTVNVEEAMHLVVDAVTSCR
FEVTDPASEEVVLMKILQVLLSCMKSKASVMLSNQHVCTIVNTCFRIVHQAGSKGELLQRIARHTMHELVRHIFS
HLLDVGNTEHALVNGVSAVKQEVVGRDNDYAFGNKQVENGNGSSEYDSQLSSANFVSNVSTGVVATMMEENTIGA
GAGKDAVSYDLHLMTEPYGVPCMVEIFHFLCSLLNVSEHMGIGPRSNTIAFDEDVPLFALGLINSAIELGGPAIR
HHPRLLSLIQDELFRNLMQFGLSMSPLILSMVSSIVLNLYHHLCTELKLQLEAFFSCVILRLAQSRYGASYQQQE
VAMEALVDFCRQKTFMVEMYANLDCDITCSNVFEDLANLLSKSAFPVNCPLSAMHILALDGLIAVIQGMAERIGN
GLLTSEQNPVTLEEYTPFWMVKCDNYSDSNHWVPFVRRRKYIKRRLMIGADHFNRDPKKGLEFLQGTHLLPDKLD
PQSVACFFRYTAGLDKNLVGDFLGNHDEFCVQVLHEFAGTFDFQDMNLDTALRLFLETFRLPGESQKIQRVLEAF
SERYYEQSPQILANKDAALLLSYSLIMLNTDQHNVQVKKKMTEEDFIRNNRHINGGSDLPREFLSELYHSICKNE
IRTTPEQGAGFPEMTPSRWIDLMHKSKKTAPFIVADCRAYLDHDMFAIMSGPTIAAISVVFDHAEHEDVYQTCID
GFLAVAKISACHHLEDVLDDLVVSLCKFTTLLNPSSVEEPVLAFGDDTKARMATVTVFTIANRYGDYIRTGWRNI
LDCILRLHKLGLLPARVASDAADESELSADPSHGKPITNSLSSAHMPSMGTPRRSSGLMGRFSQLLSLDTEEPRS
QPTEQQLAAHQRTLQTIQKCHIDSIFTESKFLQAESLLQLAKALIWAAGRPPKGNSSPEDEDTAVFCLELLIAIT
LNNRDRIVLLWQGVYEHIASIVQSTMMPCALVEKAVFGLLRICQRLLPYKENLADELLRSLQLVLKLDARVADAY
CEQITQEVSRLVKANATHIRSQMGWRTITSLLSITARHPEASEVGFDALLFIMSEGAHLLPANYVLCVDSSRQFA
ESRVGQAERSVRALDLMSGSVDCLARWGCEAKETIGEDEALKLSQDIGEMWLRLVQGLRKVCLDQREDVRNHALL
SLQKCLTGVDEIHLPHGLWLQCFDMVIFTMLDDLLEIAQGHSQKDYRNMEGTLILAMKLLSKVFLQLLHDLSQLT
TFCKLWLGVLTRMEKYMKVKVRGKKSEKLQEIVPDLLKNTLLVMKTRGVLVQRSALGGDSLWELTWLHVNNIVPS
LQSEVFPDQDLEQSRHKQSEISGGLVSDDIGSGSETAVSEGTGMGG*
**>Dimlo_022708**

MLETGINMGHLNLQSQNDAFCRGLKGSPVKPSRGAIACMINSEIGAVLAVMRRNVRWGVRYLTDDDQLEHSLIHS
LKELRKQIFLWQNEWHNIDPAVYLHPFLDVIRSDETGAPITGVALSSVYKILTLDVLDLDTVNVGDTMHLIVDAV
TSCRFEVTDPASEEVVLMKILQVLLACLKSKAAVKLNNQHVCNIVNTCFRVVHQASSKSELLQRIARHAMHELVR
CIYSHLPDIDCQSSVAQSHSDAVEKVSVLEKGHTSESKPFENGNASVESQSSTGQISSAPTDKLANEVDTYNGEI
GGVMEATQNGEKIMMEPFGVPCMVEIFHFLCSLLNVIEHMGAGPRSNPIAYDEDVPLFALGLINSAIELGGSSFS
KHPKLLALIQDELFRNLMQFGLSMSPLILSTVCSIVLNLYYHLRIKLKVQLEAFLSCVLLRLAQSKYGSSYQQQE
VAMEALVDLCRQQTFMIEMYANFDCDISSGNLFEDLANLLSKSAFPVNGPLSAMHVLALDGMISMIQGMAKRVSN
ELPTPEEVSVDPEDYNAVWTSKCKDYSDPHSWIPFVRKMKYIKRRLAVGADHFNRDSKKGLEFLQGSNLLPDKLD
PQSVAFFFRYTNGLDKNHIGDFLGNHDEFCIEVLHKFAGTFDFHGMNLDTALRLFLGTFRLPGESQKIQRVLEAF
AERYYEQSQDILTNTDAALLLSYSLILLNTDQHNAQVKKKMSEDDFIRNNRHINGGKDLPREYLSELYHSICENE
ILMIPDQGAGFQEMTSSRWINVMHKSREATAYIVCDSGAHLDHDMFAILSGPTIAAMSVVFDQVEQEDILRRCVD
GFLAIAKVSTCYHFDDVLDDLVLSLCRFTTLLSPLSVEEALLAFGDDTKARMALTAVFTIANRYGDHIHAGWKNI
LDCVLSLHKLGLLPARLASDAADDVEPSSDLEQGKPATSSVSTSRITPVVTPRKSSGLIGRFSQLLSFDMEETRS
QPDEEDLAAQQRAREIMQNCHIDSIFAESKFLQAESLLHLVKALILAAGQLRKGSSYTEDEDTGVFCLELLIAIT
LNNRDRIMLIWQGVYEHIANIVQTTVMPCMLVEKAVFGLLRICQRLLPYKEKLTDELLKSLQLVLKLDARVADAY
CEHITQEVMRLVKANATHIRSHTGWRTIISLLSITARHPEASEAGYEALAFIMSEGAHILPSNFILCVDAARQFA
ESRVGEVDRSVSALELMGGSVVCLVRWSSETKNAVGEEAAIKLSQDIGEMWLRLVQGLKRVCLDQREEVRNHAIL
VLQRSLAAVDGIHLPNALWFQCFDMVIFILLDDLLEIAQGNSPKDYRNIDGTLVLAIKLLSKAFLQSLQDLSLQP
SFCKLWLGVLNHMEKYMKLKLRGRRSEKIHELIPELLKNNLLVMKTTGILLPTDDIGGDSFWQLTWMHVKKISPS
VQSEVFPDDELEQLKAKHTKTGGFPAPDGSVVVPSGENADSR*

**>Dimlo_019311**

MANLHGEEEDDNNCSSCGGKKKDLNKFKKKQLGVSCMLNTEVGAVLAVIRRPLDSGTTQYIQEDIIDSSILHSLK
SLRSLIFSPQQQWSTLDPSLYVSPFLDVIQSDFIPAAATSVSLAAILKILKLEIFDDKTPGAREAMNSIVTGITS
CRLEKTDPISEDAVMMRILQVLAGIMKHPASVLLTDQAVCTIVNTCFQVVQQSASRGDLLQRSARYTMHELIHTI
YSRLPEVEVKSGDASESDTEDVDVDSILDSGYGVRSAVDIFHFLCSLLNVVEIVETEGSTTQASDEDVQLFALVL
INTAIELSGEAIGKHTKLLRMIQDDLFHHLIHYGASSSLLVLSMICSTVLNIYHFLRRFIRLQLEAFFSFVLLRI
PASRASLHLQEVALEGIINFCRQPTFMIEIYVNYDCDPLCRNVMEDIGKLLCKQSFPGSGPLTGAQIQAFEGLVL
MIHNIADNIDKEDDSSPSGPYPVEITEYRPFWEEKPKQDLDTWVEYVRLRKAQKRKILIAANHFNRDEKKGLEYL
KFCKLVSDPPAPKVFAFFFRYTPGLDKNMIGDYLGDPDELHIQVLKEFTRTFEFAGMILDNALRTYLETFRLPGE
SQKIQRILEAFSERFYDEQSSEIFVSKDAVFILCYSLIMLNTDQHNPQVKKKMTEEEFIRNNRAINGGKDLPREY
LSELFHSIATNAITLFGQSGRPIEMNPSRWIELMNRSKAVQSFIMSDFDRRLGRDMFASIAGPSVAAVCAFFEHA
DEDELLHECIEGLISIARIAQYGLEDTLDELLASFCKFTTLLNPYATSEETLFAFSNDMKPRMATLAVFTLANNF
GDSIRAGWRNVVDCLLKLKRLKLLPQSVIEFENFAADVQSQSKNELDALTPTQDTKCEKSRNSGIIGKFTHFLSL
ESPEDSISLGMNEFEQNLKVIKQCQIGNIFSNTSKLPSDALLNLGRSLIYAAAGKGQKFSTPVEEEETVGFCWDL
IVVIAMANISRFQTFWSSFHDYLLGVTQFPLFSPMPFAEMAIVGLFKVCLKLLDSHQVDKLPEELIFKSINLMWK
LDKDILDTCCQYIMQSVSKIIIEYPANLQSQLGWKSVLHLLSVTGRHPETYDQAVESLVVLLSDTAYVSRINYAY
CIDCAFGFVALRNSPLEKNLKLLDLMSDSVNSLIQWYRNACSEPGNNYSIASNTSTSSLEDNKGQSSLNLTMNLF
VKLGEALRKTSLARREEIRNHAVISLHKAFTLAEELDFSSSNCISCFNLIIFAMVDDLHEKMLEYSRRENAEREM
RSMEGTLKIAMDLLADVYLQFLKQIAETQEFRTFWLGVLRRMDTCMKADLGPYGETKLQEFIPDLLRKIITKMQE
TEILAPREDDDLWDITYIQIQWIAPSLKEALFPEDV*

**>AzainGNOM_ctg18078**

MGRLKLQSGIKAIEEEPEDYDTTYSNKATLSCMINSEVGAVLAVMRRNRSVRWGGQYMSGDDQLEHSLIQSLKTL
RKQIFSWQHPWHTINPAVYLQPFLDVIRSDETGAPITSIALSSLYKILTLDVIDQNTVNVEEAMHLVVDAVTSCR
FEVTDPASEEVVLMKILQVLLACMKSKASVVLSNQHVCTIVNTCFRIVHQAGNKGELLQRIARHTMHELIRRIFS
HLPDVGNSEHALVNGVGAVKQEISGLDNEYAFGGKQMDNGNGGSEYDGQQSSANLVSATGVVATMAEENTNGAAT
GKDSLSYDLQLMTEPYGVPCMVEIFHFLCSLLNVSEHMTMGPRSNTMALDEDVPLFALRLINSAIELAGPAIRHH
PRLLSLIQDELFRNLMQFGLSMSPLILSMVCSIVLNLYHHLRTELKLQLEAFFSCVILRLAQSRYGASYQQQEVA
MEALVDFCRQKTFMVEMYANLDCDITCSNVFEDLANLLSKSAFPVNCPLSAMHILALDGLIAVIQGMAERIGNGS
VSSEQSPVTLEEYTPFWMVKCDNYNDPNHWVPFVRRRKYIKRRLMIGADHFNRDPKKGLEFLQGTHLLPDKLDPQ
SVACFFRYTAGLDKNLVGDFLGNHDEFCVQVLHEFAGTFDFQDMNLDTALRLFLETFRLPGESQKIQRVLEAFSE
RYYEQSPQILVNKDAALLLSYSLIMLNTDQHNVQVKKKMTEEDFIRNNRHINGGSDLPREFLSELYHSICKNEIR
TTPEQGAGFPEMTPSRWIDLMHKSKKTAPFIVADSRAYLDHDMFAIMSGPTIAAISVVFDHAEHEEVYQTCIDGF
LAVAKISACHNledvlddlvvSLCKFTTLLNPSSVEEPVLAFGDDTKARMATVTVFTIANRYGDYIRTGWRNILD
CILRLHKLGLLPARVasdaadeselsadpsQGKPITNSLSAAHMPSMGTPRRSSGLMGRFSQLLSLDTEEPRSQP
TEQQLAAHQRTLQTIQKCHIDSIFTESKFLQAESLLQLARALIWAAGRPQKGNSSPEDEDTAVFCLELLIAITLN
NRDRIVLLWQGVYEHIANIVQSTVMPCALVEKAVFGLLRICQRLLPYKENLADELLRSLQLVLKLDARVADAYCE
QITQEVTRLVKANATHIRSQMGWRTITSLLSITARHPEASEAGFDALLFIMSEGAHLLPSNYVLCIDASRQFAES
RVGQAERSVRALDLMAGSVDCLARWCCEAKEAMGEDEVTKLSQDIGEMWLRLVQALRKVCLDQREDVRNHALSSL
QKCLTGVDEIQLPNGLWLQCFDMVIFTVLDDLLEIAQGHSQKDYRNMEGTLILAMKLLSKVFLQLLHELSQLTTF
CKLWLGVLSRMEKYMKVKVRGKKSEKLQEIVPDLLKNTLLVMKTRGVLVQRSALGGDSLWELTWLHVNNIVPSLQ
SEVFPdqdsdqsqhkqsdMVGSLVSDEMGSVASNETAASEGAEP*

**>AzainGNL1_ctg9988**

MGQLNLHSESNVSCRGIKDRPIMPSRAAFACMINSEIGAVLAVMRRNVRWGVRYMADDDQLEHSLIHSLKELRKQ
IFLWQNQWHNVDPAVYLQPFLDVIRSDETGAPITGVALSSVYKILALDVLDLDTMNVGEAMHLIVDAVTSCRFEV
TDPASEEVVLMKILQVLLACMKCKAAVKLSNQHVCNIVNTCFRVVHQASSKGELLQRIARQTMHELVRCIFSHLP
YIDCPEESFVNGSRSDIGAKVGLTDKECTSGSKLLENGNISVGQDVQLSIDMSSAPTAKVTSEMDANNADFTTEM
EAAVNGEKIMMEPFGVPCMVEIFHFLCSLLNVIEHMGIGPRSSPIADDEDVPLFALSLINSAIELGGSSISKYPR
LLSLIQDELFHYLMQFGLSMSPLILSTVCSIVLNLYHHLRLELKAQLEAFFSCVLLRLAQNKHGSSYQQQEVAME
ALVDLCRQQSFRTEMYANFDCDITSGNLFEDLTNLLSKSAFPVNGPLSAMHVLALDGMISMVQGMAERISNEMPA
LEGASADPEEYNSFWTLKCEDYNDPSCWIPFVRKMKYIKKKLMVGADHFNRDPKKGLEFLQGMHLLPDKLEPHSV
ALFFRYTTGLDKNLIGDFLGNHDEFCVQVLREFAGTFNFHGMNLDTALRLFLGTFRLPGESQKIQRVLEAFAERY
YEQSPHILSNKDAALLLSYSLIMLNTDQHNVQVKKKMTEEDFIRNNRHINGGKDLPREYLSELYHSICENEILMI
PDQGTGFPAMTSSRWINVLHKSREATPFITCDSSGLLDHDMFAILSGPMIAAMSVVFDQVEREDVLQRCVDGFLA
IAKLSTCYQFGHVLDDLVVSLCKFTTLLTPLSIEDAVLAFGDDTKAKMAMTTVFTIANRYGDYIHSGWKDILDCV
LSLHKLGLLPARLASDAADDMEPSDPEREKPttgsvstshvtsvATPRKSSSLIGRFSQLLSFDMEEPRSQPSEE
ELAAHQRTQEIIQNCHIDSIFTDSKFLQAESLLELVKALILAAGRPRVGNNFAEDEDTGVFCLELLIAITLNNRD
RIMLIWEGVYEHIANIVQSTVMPCALVEKAVFGLLRICQRLLPYKENLTNEllkslqlilklDARVADTYCEHIT
QEVMRLIKANATHIRSHAGWRTIISLLSITARHPEASDAGFEALAFIMSEGAHLLPSNFIVCVDAARQFAESRVG
EVERSVSALELMAGSVICLVRWPSEAKNAVGEEAAMRLSQDIGEMWLRLVQGLKKVCLDQREEVRNHAILVLQRS
LAAADGIHLPNALWFQCFDVVIFTLLDDLLEIAQGHSPKDYRNIDGTLVLAIKLMSKAFlqllqdlspqpsFCKL
WLGVLNHMEKYMKLKLRGKRSEKIHELIPELLKNNLLVMKTTGILLPTDDIGGDSFWQLTWLHVKKISPSLQSEI
FPDDELEEMKAKLIKTAGAPAPEKSVVVPPDENTACR*

**>AzainGNL2_ctg12079**

MREVYEEEDDDKQLTSGRERKKDVIKYKRKELGISCMLNTEVGSVLAVIRRPLDPNTQYLQEDTFESAVVQSLKS
LRSLIFNPQQEWRTVDPSIYLSPFLDVIQSDDIPAAATGvalsavlkvlklEIFDEKTPGVKDCINSVVTGITSC
QLEKTDPISEDAVMMRILQVLTAIMKHPASILLTDEAVCTIVNTCFQVVQQSASRGDLLQRSARYTMHELIQIIF
SRLPDIEVKSGEGSESDTEDVDLDSNLDSGYGVRSAVDIFHFLCSLLNVVEMVETESSCTADVDVQLFALVLINS
AIELSGDAIGKHSKLLRMIQDDLFHHLIHYGSASSPLVLSMICSTVLNIYHFLRRFIRLQLEAFFSFVLLKVAAF
GSSLQLQEVALEGIINFCRQPTFLIEVYVNYDCDPFCRNVLEEIGKLLCKHSFPGTGPLTTSQIQAFEGLVILIH
NIAENIDKEGDTSPSGPYPVEITEYRPYWEDKPRDDLETWLEYIRVRKAQKKKIMIAGNHFNRDEKKGLEYLKLC
QLVSDPPDPKAIAFFFRYTQGLDKNMMGDYLGDPDEFHIQVLKEFTETFEFAGMILDNALRTYLETFRLPGESQK
IQRILEAFSERFFDQQSSEIFAGKDTVFILCYSLIMLNTDQHNPQVKKKMTEEEFIRNNRAINGGKDLPREYLSE
LFHSIANNAITLFGQSGLPVEMNPSKWIELINRSKSVQPFVLCDFDRRLGRDMFASIAGPSVAAFSAFFEHADED
ELLQECIEGLISTARIAQYRLEDTLDELLASFCKFTTLLNPYASAEETLFAFSNDMKPKMATLAVFTLANNFGDS
IRAGWRNIVDCllklkrlkllPQSVVEFDSSTSDAPSNLKNESGIVFPTHDPKFGNRQSSGMIGRFTHFlslesp
edslslGMNEFEQNLKVIKQCQIGNIFSNSTNLPYEALLNLGRSLIYAAAGKGQKFSTPVEEEETVGFCWDLIIA
ITLANINSFPTFWPSFHDFLLGVCQCLFKVCLKLLASYQPDKYPEELIFKSINLMWKLDKDILDTCCQFITQSLS
KIIVEYPANLQSAVGWKSVLHLLSITGRHPDTYDQAVETLIMLMSDATHVSRASYAYCIDCAFGFVALKNSPLEK
NLKILDLLVDSVNLLIQWYRNACSEsgnnysaasntstssVEDNKGINSLNFAVNLFIKLGEALRKTSLARREEI
RNHAVLSLRKSFALAEDLDFSSTNCINCFNLVVFAMVDDLHEKMVEYSRRDNAEREMRSMEGTLKIAMELLTDVY
LQFLRQISESSGFRTFWLGVLRRMDTCMKADLGVYGETKLQEVIPDFLRKMITNMKEKEILVPKEGEDLWEITYI
QIQWIAPSLKEELFPDEI*

**>Cepfo_24675**

MGRLKMQSGINAIEEEPEDLDATYSNKETLACMINSEVGAVLAVMRRNVRWGGRYMSGDDQLEHSLIQNLKALRK
QIFSWQHQWHTINPCLYLQPFLDVIRSNETGPPITGVALSSVYKILTLDVIDKSTVNVEDSMHMLVDAVTSCRFE
VTDPASEEVVLMKMLQVFLACVKSKASIMLSNLHVCTVVSTCFRIVHQARTKGELLQRIARHTMHELVRCIFGHL
PDVDNREHELVNGVRTVGQEIGGLDNEYAFGSRQMENGNSSMDSDGQPIFSSNASTGWVTSGMEENTIGAVPYGA
PCMVEIFHYLCSLLNDVEQTELGPRSNTIAFDEDVPLFALSLINSAIEVGGPSIRRHPRLLSLIQDELFCNLMQY
GLSMSPLILSTVCSIVLNLYHQLRTELKLQLEAFFSCVILRLAQSRYGASYQQQEVAMEALVDFCRQKTFMVEMY
ANLDCDITCSNVFEDLANLLSKSAFPVNCPLSSMHILALDGLIAVIQGMAERIGNTSVGSEQGRVNLEEYTPFWM
VKCGNYSEPSYWVPFVRRRKCIKRRLMIGADHFNRDPKKGLEFLQVTHLLPDKLDPQSVACFFRYTAGLDKNLVG
DFLGNHDEFCVQVLHEFAGTFDFQDMNLDTALRLFLETFRLPGESQKIQRVLEAFSERYYEQSPQILAHKDAALL
LSYSLIMLNTDQHNVQVKKKMTEEDFIRNNRLINGGNDLPREFLSELYHSICKNEIRTTPEQGTGFPEMTPSRWI
DLLHKSKKSAPFIVSDSRTYLDHDMFAIMSGPTIAAISVVFDHAEHEDVYQTCIDGFLAVAKISACHHLEDVLDD
LVVSLCKFTTLLNSSSVEEPVLAFGDDTKARMATVTVFNIANMYGDYIRTGWRNILDCILRLHKLGLLPARVASD
AADDSELSADPGHGKPLTNTSSSAHLPSVGTPRRSSGLMGRFSQLLSLDTDEPRSQPTEQQLAAHQRTLQTIQTC
HIDSIFTESKFLQAESLLQLARALIWAAGRPQKGSSSPEDEDTAVFCLELLIAITLNNRDRIVLLWQGVYEHIAN
IVQSTVMPCALVEKAVFGLLRICQRLLPYKENLADELLRSLQLVLKLDARVADAYCEQITHEVSRLVKANASHIR
SQMGWRIIASLLSITARHPEASEAGFGALLFIMSDGAHLLPANYVLCVDASRQFAESRVGQAERSLRALDLMAGS
VDLLARWAHEAKEANGEEETAKMSQDIGEMWLRLVQGLRKVCLDQREEVRNHALLSLQKCLTGVDGIDLPHGLWL
QCFDQVIFTMLDDLLEIAQGHSQKDYRNMEGTLILAMKLLSKVFLQLLYDLSQLTTFCKLWLGVLSRMEKYMKVK
VRAKKSEKLLEIVPELLKNTLCVMKAKGVLMQRSALGGDSLWELTWLHVNNIVPSLQAEVFPDQDWEQSQLKQCE
TGVGMMSGETGSVASSEILVASEGAVTGS*

**>Cepfo_21055**

MGVLSLQSGNDAFCGELKDCPIKPSRGALACVINSEIGAVLAVMRRNVRWGVRYVADDDQLEHSLIHSLKELRKQ
IFSWQSQWDNINPALYLQPFMDVIRSEETGAPITGVALSSVYKILTLDVFDIDTVNVGDAMHLIVDAVTSCRFEV
TDPASEEVVLMKILQVLLACMKSKAAGKLSNQHVCNIVDTCFRVVHQAGSKSELLQRIARHTMHELVRCIFSRLS
DFSNPELDLANGSRAYFGEEMGVADKDHSSGSRQQENGNVSVEYDAQSSIGYEPNAPKNKLVGEKDTENSEVGSG
MVAAENDQRIMMEPFGIPCMVEIFHFLCSLLNVIEHVGIGPRSNPIAYDEDVPLFALGLINSAIELGGPSFSKHR
NLLILIQDELFHNLMQFGLSMSPLILSTVCSIVLNLYFYLRTELKVQLESFFSCVLLRLAQSKHGSSYQQQEVAM
EAVVDLCRQQSFMTEMYANFDCDITCSNVFEDIGNLLSKSAFPVNGPLSAMHILALDGLISMIQDLAERIGNECP
VPEEPLADPESYKAFWTLRCENYDDPSFWIPFVRKMKHIKKKLMTGADHFNRDPKKGLEFLQVVHLLPDTLDAKS
VASFFRYTAGLDKNLIGDFLGNHDEFCVQVLHEFSGTFDFRGMNLDTALRLFLGTFRLPGESQKIQRVLEAFAER
YYEQSPQEFCNKDATLLLSYSIILLNTDLHNAQVKKKMTLEDFIRNNRSINGGKDLPREFLSELYHSICENEIQM
IPEQVVFDQVEQEDVLQRCIEGFLAIAKISAYYHLDEVLNDLVVSLCNFTTLLTPLSVQEVVLAFEDEEKSRMST
TTVFMIANRYGDYIRSSWKNILDCILSLHKLGLLPARVASDAVDDIEFTSDLERGKPATSSISTLHMPPLAAPRK
SSGLIGRFSSILSFDMEEPRSQPSEEQLAAHQRTRETIQNCHIDSIFTESKFLQAESLLQLVKALISASSRFHLG
SNTMEDEDTLVLCLELLITITLNNRDRIMIIWHAVYEHISNIVQSIVMPCVLVEKAVFGLLRICQRLLPYKENLT
DELIKSLQLILKLDARVADAYCENITHEVLRLVKANASHIRSHVGWRIIISLLSITARHPEASETGFEALVSVMS
EGAHLLPCNYILCIDAARQFAQSRVAENNRSLSAIDLMAGSVACLARWSCETKNSVGEEAAMRVKQDIGEMWLRL
VQGLREVCLDQRQEVRNHAVLMLQRSLAGVDGINFPNGLWFQCFDSVIFTLLDDLLEIAQGSSPKDYRDMEGTLV
LAMKLMSKSFLQSLQNLSQQPSFCKLWLGVLNRMERYMKVKFRGKRSEKIQELIYELLKNTLLVMKTTGILMPSN
DIGGDSFWQLTCLHVKNISPSLELEVFPKNELEELKTDRMKTGENAVIDGSDLVPSSETTA*

**>Cepfo_22974**

MAEDQNNCSSCRKKERDMSKSRKKELGLSCMLNTEVGAVLAVIRRPPDLTTQYISPQEETVDSGILNSLKSLRAL
IFNPQQEWRTIDPSIYLSPFLDVIQSDDIPAPATGVALSAILKILNLEIFDEKSPGAKDALHSVVIGITSCRLER
TDTVSEDAVIMRILQVLIGIMKHRSSVLLTDHSVCTVVNTCFQVVQQSASRGDLLQRSARYTMHELIQIIFSRLP
EIQVKSEDESESDTEDVDIDKNIDSGYGVRSAVDIFQFLCSLLNVVQVVETEGSTSHTADEDVQFFALVLINSAI
ELSGDAIGKHPKLLRMIQDDLFHHLIHYGICSSSLVLSMICSTVLNIYHFLRRFIRLQLEAFFTFVLFRVSSLRG
ASQLQEVVIEGIVNFCRQPTFTIEVYVNYDCDPVCRNVFEEIGKMLCRHAFPGPGPLTALQIQAFEGLVIMIHSI
ADNIDKEDDPHPRGPYSTEITEYKPFWEEKSQDDIETWLEHIRMRKAQKKKLLVAENHFNRDEKKGLEYLKHCHL
VKDPPNPKAYAFFFRYTPGLDKNMIGDYLGDPDEFRIQVLKEFTETFEFSSMILDNALRTYLETFRLPGESQKIH
RILEAFSERFYEQQSSDIFVSKDSVFILCYSLIMLNTDQHNPQVKKKMTEEDFIRNNRAINAGRDLPREYLSELF
LSISNRAITIFGKSGPIEMNSGRWIELMNRSKILQPFIICDFDHRLGRDMFACIAGPTVAAFSAFFENSDEEEIL
HECIDGLISVARIAQYGLEDTLDELLASFCKFTTLLNPYASAEETLYAFSNDLKPRMATIAVFSIGNNFGYSIRG
GWRNIVDCLLKLKRLKLLPQSVIEYDAASTSSSDVPAHSRNASGLNFTYHDPKFANCQSSGMINRFSHFLSLESM
EDSVSLGISEFERNLKVIKQCRIGSIFSNSSNLPDDAFLHLGRSLIFAAAGKGQKFSTPIEEEETVGFCWDLITA
IALVNIKRFQAFWPSFHGYLLAVADFPLFSPIPFAEKAIIGLLKVCLKFLTTQQPEKQLEELIFKSINLMWKLDK
EILDTCCEVIMQSTRKIIIEYPVILQSQLGWKTVLHLLSVTGRHPETYDQAVEALIMLMSDATNVSQTNYPYCIE
CSFSFVALKNSPLEKNLKILDLMSDSVNFLIHWYKSQHLDHETNFSVASITSTSSLEDNSRGFGSSNFPMNHFIK
LGEALRKTSLARREEIRNHAVLSLQRSFTLAQDLDFSSTSILSCFNLVIFAMVDDLHEKFVEYSRRENAEREMRS
MEGTLRIAMELLTEVYLMYLRQIAESPGFRTFWLGVLRRMDTCMKTHLGHYGETRLQELVPGLLSKMIAMMKEKD
ILVPREEDDLWEITYIQIQWIAPSLKKELFPEADF*

**>Aveca_019952**

MGRLKMQPGIKAIEEEPEDIDASYSNKSTLACMIDSEIAAVLAVMRRNVRWGGRFMSGDDQLEHSLIQNLKALRK
QIFSWQHQWHNINAAVYLQPFLDVIRSDETGAPITSVALSSVYKILTLDVIDQNTANVEDAMHVVVDAVTSCRFE
VTDPASEEVVLMKILQVFLACMKSKASVMLSNQHVCTIVNTCFRIVHQAGTKGDLLQRIARHTMHELVRCIFLHL
PDVDNTERAIVNGVSTAGQEVGGLDNEYAFVSRQTDNGNGSSENDGQPIFSSNASAGLATSVMDENIIGAVPYGV
PCMVEIFHFLCSLLNVVEHVGMGPRSNTVAFDEDVPLFALSLINSAIELGGSSIRHHPRLLSLIQDELFRNLMQF
GLSMSPLILSMVCSIVLNLYHHLRTELKLQLEAFFSCIILRLAQSRYGASYQQQEAAMEALVDFCRQKTFMVEMY
ANLDCDITCSNVFEDLANLLSKSAFPVNCPLSSMHILALDGLIAVIQGMAERIGNGMVSLEQAPVNLEEYTPFWM
VKCDNYNDPVHWVPFVRRRKYIKRRLMIGADHFNRDPKKGLEFLQGTHLLPDKLDPQSVACFFRYTAGLDKNLVG
DFLGNHDEFCVQVLHEFAGTFDFQDMHLDTALRLFLETFRLPGESQKIQRVLEAFSERYYEQSPQILANKDAALL
LSYSLIMLNTDQHNVQVKKKMTEEDFIRNNRHINGGNDLPREFLSELYHSICKNEIRTTPEQGAGFAEMTPSRWI
DLMHKSKKTAPYIVSDSRAYLDHDMFAIMSGPTIAAISVVFDHAEHEDVYQTCIDGFLAVAKISACHHLEDVLDD
LVVSLCKFTTLLNPSSVEEPVLAFGDDTKARMATVTVFTIANRYGDYIRTGWRNILDCILRLHKLGLLPARVASD
AADESEVSPDPGHGKPLTNSLSSAHMPSIGTPRRSSGLMGRFSQLLSLDTEEPRSQPTEQQLAAHQRTLQTIQKC
HIDSIFTESKFLQAESLLQLARALIWAAGRPQKGNSSPEDEDTAVFCLELLIAITLNNRDRIVLLWQGVYEHIAN
IVQSTVMPCALVEKAVFGLLRICQRLLPYKENLADELLRSLQLVLKLDARVADAYCEQITQEVTRLVKANATHIR
SQMGWRTITSLLSITARHPEASEAGFDALLFIMSDGAHLLPANYVLCVDASRQFAESRVGQVERSLRALDLMAGS
VDCLSRWAREAMETMGEEETAKMSQDIGEMWLRLVQGLRKVCLDQREDVRNHAILSLQKCLTGVDEINLPHAVWL
NCFEMVIFTMLDDLLEIAQGHSHKDYRNMEGTLILAMKLLSKVFLQLLQVLSQLTTFCKLWLGILNRMEKYIKAK
VRGKKSEKLQEIVLDLLKNTLVAMKTRGVLVQRSTLGGDSLWELTWFHVNNIASSLQAEVFPDQDQEQSQPKQGE
TGGGSMSGETGSVASSENEPSEAPGTGG*

**>Aveca_019955**

MGRLKMQSGVDENEGERGGIDATCNKATLAYIINSEIAAVLTVMRRNVRWGVRYISEDDQLEHSLIQNLKALRKL
VFSWQHEWDNISPTMYLQPFLDVIRSDETGAPITSVALSSVYKILTLDVIDQNTINVKDAMHSVVDAVTSCRFEV

TDPASEEVVLMKILQVFLASMKSKASVVLSNQHVCTIVNTCFRIVHQASTKGELLQRIARQTMHELVRCIFSHLS
DVDDSEGALVNGVSAVGQEIGGLDNEYAFGSKLMENGNVSSEKDGQSQHSNASTGFDTGVMEENIIEAVPYGIPC
MVEIFHFLCSLLNVVEHVGSGRRSNTIAFEEDVPLFALSLINSTIELGGPSFHRHPTLLSLIQDELFHNLMQFGL
SMSPLILSMVCNIVLNLYHHLRTELKLQLEAFFSCVILRLAQSKYGASYQQQEVAMEALVDFCRQKSFMVEMYAN
LDCDISCSNVFEELTNLLSKSAFPVNCPLSSMHILALDGLIAVIQGMAERIDSGSVSLEQVPVNLEEYSPFWTVK
CGNYNDPAHWVPFIRQRKYIKRRLLVGADHFNHDKKKGLEFLQGTHLLPDKLDPHSVACFFRYTAGLDKNLVGDF
LGNHDEFSVQVLHEFAQTFDFQDMSLDMALRLFLETFRLPGESQKIQRVLEAFSERYYEQSPQILANKDAALLLS
YSLIMLNTDQHNAQVKKKMTEEDFIRNNRRINGGSDLPREFLSELYHSICKNEIRTTPEQGAGLAEMTSSRWINL
MHKSKESSPFVVFDSGAYLDHDMFAIMSGPAIAAISVVFDHAEHEVVYKSCIDGFLAVAKISACYHLQDVLDDLV
VSLCKFTTLLDSSSVEEAVLAFADDPKAMMATVAVFTIVNKYGDTIRIGWRNILDCILRLHKLGLLPARVASSDS
AEETELCADPGHGKPITKSLSSSHTSMGTPKKSSGLMGRFSQLLSLDTEEPRLQPSEEQLAAHHYTRQTIEKCNI
DSIFTESKFLQAESLLQLVRALISAAGRPQKGNIYLEDEDIAVFCLELLIAITLNNRDRIALLWEGIYEHIANIV
QSTVMPCALVEKAVFGLLRICQRLLPYKENVADELLRSLQLVLKLDARVADAYCEQITQEVTRLVKANAPQIRSQ
IGWRTITTLLSITARHPEASEAGFDALVFIMSNGAHLLPANYVLCVDASRQFAESRVDEVGRSLHALDLMAGSIG
CLARWAHEAKETMGEEDTIKMLQNIGEMWLRLVQGLRKVCLDQRTEVRNHAILSLQKCLTGAEGMNLPQVLWLQC
FETVIFTLLDDLLEIARTPYQKEYRNMEGTLILAMKLMSKAFVQFLRDLSQLATFCNLWLGVLSQMENYMKVKVR
GKKSYELQETVPELLKNILIVMKTWGVLVRGAPGKDTLWESTWIHVNNIAPSLQSEVFPDQESEQSLKKGETGSA
TSGEAVASQVSGMVG*

**>Aveca_014651**

MEKIQAEEESNSSKKKELSKWKRKELGVSCMLNTEVGAVLAVIRRPPDPTTHYLSFQDETVDSGILNSLKSLRAL
IFNPQQEWRTIDPSIYLSPFLDVIQSDDIPASATGVALSAVLKIINLEIFDEKTPGAKDAMHSIVTGITSCRLEK
TDSVSEDAVMMRILQVLTGVMKHPASVLLTDHSVCTVVNTCFQVVQQSASRGDLLQRSATYTMHELIQTIFSRLP
DFEAKSGEELDSDTEDADSGGQFDSGYGIRCAVDIFHFLCSLLNVVDVVETEGSTYHTADDDVQLFALVLINSAI
ELSGDAIGRHPKLLRMIQDDLFHHLIHYGSCSRPLVLSMICSTVLNIYHFLRRFVRLQLEAFFIYVSFRVAALGG
AGQLQEVALEGIINFCRQPSFVVEVYANYDCDPMCRNVFEEIGKLICRHSFPGCGPLTPLQIQAFEGLMVVIHNI
ADNIDRENDLRPTGPFSVEITEYRAFWEEKPKDDSETWLDYVRMRKTQKKKIMVAENHFNRDDKKGLEYLKLCQL
VADPPDPKAYAFFFRYSPRLDKTAIGDFLGDPDEFHVKVLKEFTDTFEFSGMILDSALRAYLETFRLPGESQKIQ
RILEAFSERFYDQQSSDIFASKDSVCILCYSLIMLNTDQHNPQVKKKMTEEEFIRNNRQINDGKDHPREYLSELF
YSISNREITLSGQSGPVEMNPAKWIELMNRSKSMQPFIVCDFDRRLGRDMFACIAGPSVAALCAFFDHVDEDEML
NECIEGLISVARIAQYGLEDTLDELLASFCKFTKLLNPYASPEETLFAFSNDLKPRMATLAVFTIANNFGESIRG
GWRNIVDCLLKLRRLKLLPHSVIEFENASASSSDLRSESGIIFPNHDPKFGKHHNSGMLSRFSHFLSLDSAEESL
SLGMNEFEQNLKMIKKCRIGNIFSNSSNLPDDGLLNLGRSLIFAAAGKGQKFSTPIEEEETVGFCWDLIVAISLA
NVNRFQAFWPSFHDNLLGVAQFPLFSPIPFAEKAIVGLLKICLKVLATYQPDNLPEELIFKSVNLMWKLDKEILD
TCCEDITQSVSKILIEYAANLQTQLGWKSVLHLLSVTGRHQETYDQAVESLIVFMSDATHLSRINYAYCIDCAFG
FVALKNSPLEKNLKILDLLSESVNLLIQWNKNHYSDPGTNFSNASSTSTSSMEESLKGFGNFTTTLFVKLGEALR
KTSLARREEIRNHAITSLRRSFTLAEELGFSSTSIINCFNLVIFAMVDDLHEKMLEYSRRENAEREMRSMEGTLK
IAMELLSDVYIQYLKLISESPGFRTFWLGVLRRMDTCMKADLGEFGKTKLQETIPELLQKIITKMKDTEILVPKE
NDDLWEITYVQIQWIAPSLKDELFPEVAI*

**>Triwi_LOC119991987**

MGRLKLQSGIKAIEEETEEFETTYSNKSTLACIINSEISAVLAVMRRNVRWGGRYIAGDDQLEHSLIQSLKALRK
QIFSWQHQWHAINPAVYLQPFLDVIRSDETGASITGVALSSVYKILTLDIVDQNTFNVDDAMHLVVDAVTSCRFE
VTDPASEEVVLMKILQVLLACMKSKASVMLSNQHVCTIVNTCFRIVHQAGTKGELLQRIARYTMNELVRCIFAHL
PDLNNSEHAILNGVSAVKQETDGPENEYSFGAKQMDNGNVNSEQSSATFASDSTGMVTTMLEENSVGAASKDAVS
YDLQLMTEPYGIPCMVEIFHFLCSLLNLVEHTGMGSRSNTVAFDEDVPLFALSLINLAIELGGPAICHHPRLLSL
IQDELFRNLMQFGLSMSTLILSMVCSIVLNLYQHLRTELKLQLEAFFSCVILRLAQSRYGSSYQQEVAMEALVDF
CRQKTFMVEMYANLDCDITCGNVFEDLANLLSKSAFPVNCPLSAMHILALDGLIAVIQGMAERVGNGLVSSEHTP
VHLEEYTPFWMVKCDNYSDPSHWVPFVRRRKYIKRRLMIGADHFNRDPKKGLEFLQGTHLLPDKLDPRSVACFFR
YTAGLDKNLVGDFLGNHDEFCVQVLHEFAGTFDFQDMNLDTALRLFLETFRLPGESQKIQRVLEAFSERYYEQSP
HILANKDAALLLSYSLIMLNTDQHNIQVKKKMTEEDFIRNNRHINGGNDLPREFLTELYHSICKNEIRTTPEQGA
GFPEMTPSRWIDLMHKLKKTSPFIVADSRAYLDHDMFAIMSGPTIAAISVVFDHAEHEEVYQTCIDGFLAVAKIS
ACHHLEDVLDDLVVSLCKFTTLLNPSSVEEPVLAFGDDMKARMATVTVFTIANRYGDYIRTGWRNILDCILRLHK
LGLLPARVASDAADESELSADHTHGKPLTNSLSSAHMQSIGTPRRSSGLMGRFSQLLSLDTEEPRSQPTEQQLAA
HQRTLQTIQKCHIDSIFTESKFLQAESLLQLARALIFAAGRPQKGSSSPEDEDTAVFCLELLIAITLNNRDRIFL
LWQGVYEHIANIVQSTVMPCALVEKAVFGLLRICQRLLPYKENLADELLRSLQLVLKLDARVADAYCEQITQEVS
RLVKANSSHIRSQMGWFVITSLLSITARHPEASEVGFDALLFIMSDGSYLLPANYILCVDAARQFAESRVGQAER
SVRALDLMSGSVNCLAKWTHEAKQTVGEEEVAKLSQGIGEMWLRLVQGLRKVCLDQRDDVRNHALLSLQKCLTGV
EGFHLQHGLWLQCFDLVIFTMLDDLLEIAQGHSQKDFRNMEGTLTHALKLLSKVFLQLLYDLSQLTTFCKLWLGV
LGRMEKCMKVKVRGKKSEKLQELVPELLKNTLLVMKTKGVLVQRNALGGDSLWELTWLHVNNIAPSLQSEVFPDQ
DFEQSQLKQGETVGGLVDETINSSNDTLVTEGSGTGG*

**>Triwi_LOC120016528**

MGHLNPQSGTNAFTVDLKDSHIRSSQGALACMINSEIGAVLAVMRRNVRWGARYVSDDDQLEHSLIHSLKELRKQ
IFSWHNQWHNINPALYLQPFLDVIQSDETGAPITGVALSSIYKMLTVDVLDFNTVNVGEAMHLIVDAVTSCRFEV
TDPGSEEVVLMKILQVLLACMKSKASVKLSNQHVCNIINTCFRVVHQASSKGELLQRIARHTMHELVRCIFSHLP

DISNEGLANGSSSYCVEVVDAPKKDRTGVGKQLKNGNTSVKYDSQQSRENASGTLVDMLSSEKDDACKIETANGF
QADQNDEKLMMEPFGVPCMVEILHFLCSLLNVIEHIDIGPRSNPIAYDEDVPLFALGLINSAIELGGPSFSRHPK
LLSLIQDELFRNIMKFGMSMSPLILSMVCSIVLNLYQHLRTELKVQLEAFFSYVLLRIAESKHGSFYQRQEVAME
ALVDLCRQQTFIAEMFANFDCDIAGCNVFEDLASLLSKNAFPVNEPLSTMHILALDGLISMIKGIAERIGQESPP
TEEPSTEEYKAFWAMTCENYDDPYTWVPFVRKMKYTKRRLAVGADHFNRDPKKGLEFLQGMNLLPDKLDPQSVAS
VFRYTAGLDKNLIGDFLGNHDDFCVKVLHEFAGTFDFRGISLDTALRLFLGTFRLPGESQKIQRVLEAFAERYYE
QSPHILSNKDAALLLSYSLIMLNTDQHNVQVKKKMTEEDFIRNNRHINGGNDLPREYLSELYQSICENEIQMIPE
QGTGLPAMTPSRWINVMHKSQNTTPYIVCDFKALLDYDMFAILSGPTIAAMSVVFDQAEHEDVLRTCTDGFLAIA
KISAHYRSGDVLDDLVVSLCKFTNFLTPLSVDDAILAFGDDTKARMATSTVFTIANSYGDYIRSGWKNILDCVLS
FHKLGLLPAHLASDATDYVESSAELEPAKLTSPLKRSQTPSVATPRKSSGLMGRFSQFLSFDMEEPQLQPTEEQL
VAQHHTRETIQSCHIDSIFMESKFLQSESLLQLVRALILAAGRLRKGGSAVEDESTAVFCLELLIAIALNNRDRI
MLIWPVVFDHISNIVRSTIMPCNLVEKAVFGLLKICQRLLPYKENLTEELLKSLQLILKLDARVADAYCEPITHE
VLRLIRANATHIRSHMGWCIIISLLSITARHPEASEVGFEALVFIMSDGAHLLPSNYVLCIDAARQFAESRLGEV
DRSVAAVDMMATSITCLVHWSSETRNTIGEEAAMKVSQDIGEMWLRLVQALRKVCLDQREEVRNHAILMLQRSLA
GLDGTHLPAALWFQSFDMVIFSLLDDLLEIVQANSPKDYRKMEGTLVLAMKLMSKSFLQVLQDLSQQPSFCKLWL
GVLNHMEKYMKVKFRGKRSEKIHELVPELLKNTLLVMKTTGILMPSNDIGEDGFWQLTWLHVENICPSIKSEVFP
DHELELIQSKHIQGGGNPVPDGSMLVPSSETTA*

**>Triwi_LOC119996657**

MGHLNPQGTNAFPEDLMDSHVRFSQGALACMINSEIGAVLAVMRRNVRWGVHYVSDDDQLEHSLVHSLKELRKQI
FSWHSQWHNINPALYLQPFLDVIQSDETGAPITGVALSSIYKILTLDILDFNTVNVGEVMYLIVDAVTSCRFEVT
DPGSEEVVLMKILQVLLACMKSKASVKLSNQHVCNIINTCFRVVHQASSKGELLQRIARHTMHELVRCIFSHLPD
VSSEGLANGSSSYCCEEVDPPNNDSTGVGKQLENGNVNVKYDSQQSRENDSGTLVDMLSSEKDDAYKIETANGFQ
ADQNDEKLMAEPFGVPCMVEILHFLCSLLNVVEHIDIGSKSNPIAYDEDVPLFALGLINAAIELGGPSFNRHPKL
LSLIQDQLFRHLMKFGMSMSPLILSMVCSTVLNLYQHLRTELKVQLEAFISYVLLRIAESKHGSFYQRQEVAMEA
LVDLCRQKTFISEMFANFDCDIAGCNVFEDLASLLSKNAFPVNEPLSAMHILALDGLIAIIKGIAERIGQESPPT
EKPSTNNEEYKAFWTMTCENYHDPYTWVPFVRKMKYTKRRLAVGADHFNRDPKKGLEFLQGINLLPDKLDPYSVA
SVFRYTAGLDKNLIGDFLGNHDDFCVKVLHEFACTFDFHGMSLDTALRLFVGTFRLPGESQKIQRVLEAFAERYY
EQSPHILSNKDAALLLSYSLILLNTDQHNVQVKKKMTEEDFIRNNRRINGGNDLPREYLSELYQSICENEIQMIP
EQGTGFPAMTPSRWINVMHKSQNTTPYIVSDFIALLDYDMFAILSGPTIAAMSVVFDQAEHEDVLQTCTDGFLAI
AKISAHYRSGDVLDDLVVSLCKFTNFLTPLSVDDAILAFGDDTKARKATSTVFTIANRYGDYIRSGWKNILDCVI
SFHKLGLLQAHLTTTSDATDDVEPSAELERAKLISYLKRSQTSSVATPRKSSGLMGRFSQFLSFDTEEPQLQPTE
EQLAAQHCTRETIQSCHIDSIFMESKFLQAESLLQLVSALILAAGRLRKGVSAVEDENTAVFCLELLIAIALNNR
DRIMLIWPGIFDHISNIVQSTVMPCNHVEKAVFGLLKICQRLLPYKENLTEELLKSLQLILKLDARVADAYCEPI
THEVLRLVRANATHIRSHMGWRTIISLLSITARHPEASEVGFEALVFIMSDGAHLLPSNYVLCIDAARQFAESRL
GEVDRSVIAIDMMATSINCLVRWSSETRNSIGEEAAMKVSQDLGEMWLRLVQALRKVCLDQREEVRNQAILMLQR
SLAGVDGIQLPAVLWSQCFDTVIFTLLDNLLEIAEANSPKDYRKMECTLVLAMKLMSKSFSQVLQDLSKQPSFCE
LWLGVLNHMEKFMKVRFRGKRGEKIHESVPELLKNTLLVMKTTGILTPSNDIGVNSFWQLTWLHVKNICPSMQSE
VFPDHELEHMHIKAEGIPVPDGSMLVPSSETAA*

**>Triwi_LOC119996659**

MGHLNPQGTNAFPEDLMDSHVRFSQGALACMINSEIGAVLAVMRRNVRWGVHYVSDDDQLEHSLVHSLKELRKQI
FSWHSQWHNINPALYLQPFLDVIQSDETGAPITGVALSSIYKILTLDILDFNTVNVGEVMYLIVDAVTSCRFEVT
DPGSEEVVLMILQVLLACMKSKASVKLSNQHVCNIINTCFRVVHQASSKGELLQRIARHTMHELVRCIFSHLPDV
SSEGLANGSSSYCCEEVDPPNNDSTGVGKQLENGNVNVKYDSQQSRENDSGTLVDMLSSEKDDAYKIETANGFQA
DQNDEKLMAEPFGVPCMVEILHFLCSLLNVVEHIDIGSKSNPIAYDEDVPLFALGLINAAIELGGPSFNRHPKLL
SLIQDQLFRHLMKFGMSMSPLILSMVCSTVLNLYQHLRTELKVQLEAFISYVLLRIAESKHGSFYQRQEVAMEAL
VDLCRQKTFISEMFANFDCDIAGCNVFEDLASLLSKNAFPVNEPLSAMHILALDGLIAIIKGIAERIGQESPPTE
KPSTNNEEYKAFWTMTCENYHDPYTWVPFVRKMKYTKRRLAVGADHFNRDPKKGLEFLQGINLLPDKLDPYSVAS
VFRYTAGLDKNLIGDFLGNHDDFCVKVLHEFACTFDFHGMSLDTALRLFVGTFRLPGESQKIQRVLEAFAERYYE
QSPHILSNKDAALLLSYSLILLNTDQHNVQVKKKMTEEDFIRNNRRINGGNDLPREYLSELYQSICENEIQMIPE
QGTGFPAMTPSRWINVMHKSQNTTPYIVSDFIALLDYDMFAILSGPTIAAMSVVFDQAEHEDVLQTCTDGFLAIA
KISAHYRSGDVLDDLVVSLCKFTNFLTPLSVDDAILAFGDDTKARKATSTVFTIANRYGDYIRSGWKNILDCVIS
FHKLGLLQAHLTTTSDATDDVEPSAELERAKLISYLKRSQTSSVATPRKSSGLMGRFSQFLSFDTEEPQLQPTEE
QLAAQHCTRETIQSCHIDSIFMESKFLQAESLLQLVSALILAAGRLRKGVSAVEDENTAVFCLELLIAIALNNRD
RIMLIWPGIFDHISNIVQSTVMPCNHVEKAVFGLLKICQRLLPYKENLTEELLKSLQLILKLDARVADAYCEPIT
HEVLRLVRANATHIRSHMGWRTIISLLSITARHPEASEVGFEALVFIMSDGAHLLPSNYVLCIDAARQFAESRLG
EVDRSVIAIDMMATSINCLVRWSSETRNSIGEEAAMKVSQDLGEMWLRLVQALRKVCLDQREEVRNQAILMLQRS
LAGVDGIQLPAVLWSQCFDTVIFTLLDNLLEIAEANSPKDYRKMECTLVLAMKLMSKSFSQVLQDLSKQPSFCEL
WLGVLNHMEKFMKVRFRGKRGEKIHESVPELLKNTLLVMKTTGILTPSNDIGVNSFWQLTWLHVKNICPSMQSEV
FPDHELEHMHIKAEGIPVPDGSMLVPSSETAA*

**>Triwi_LOC119996549**

MEGNKTKKELTKSKRKELALSCMLNTEVGAVLAVIRRPLDHTSQYFSPQEDNFDANILQSLKSLRSILFNPQQEW
RAIDPSIYLSPFLDVIQSDDIPAAATGVALSAVLKILKLEIFDEKTPGARDAINAVVTGITSCRLERTDHASEDA
VMIRILQVLTAIMKHNASVLLTDHAVCTIVNTCFQVVQQSARRGDLLQRSARYTMHELIQVIFSRLPDIEVKDGE

ESESDTEDVDIGSTLDSGYGVRCAVDIFHFLCSLLNVVEVVGTEGHSAQTSDEDVQLFALNLINSAIELSGDAIG
KHPKLLRMIQDDLFHHLIHYGTCSSPFVLSMICSSVLNIYHFLRRFIRLQLEAFFAFVLLKNAATGSPYRLQEVA
LEGIINFCRQPEFIIEIFVNYDCDPTCRNLFEEIGKLLCRNSFPAGGPLTTLQMQAFEGLLIMIHNIADSVDKED
DSSVAGTYPIDIHEYRPFWEEKSGDSENLVEHSRIRKAQKRKILIAGHHFNRDEKKGLAYLTHCHLVSDPPDPKA
FALFFRFTPGLDKNLIGDYLGDPDEFHIQVLKQFTETFQFAGMILDPALRTYLETFRLPGEAQKIQRILEAFSER
FYDQQSSEIFVSKDCVYILCYSLIMLNTDQHNPQVKKKMTEEEFIRNNRQINGGNDLPREYLSELFHSISTNAIT
LFVQSGVPIEMNPSRWIELMNRSKIMQPFILCDFDPRLGRDMFACIAGPSIAALSAFFEHADEDEMLHECIEGLI
SVARIAQYGLEDTLDELLASFSKFTTLLNPYASAEETLFAFSHDMKPRMATLAVFTIANNFADSIRGGWRNIVDC
LLKLKRLKLLPQSVIEFDASSSPPSDNIPPDMRSESGVIFPSQDHKFGNPKSSMSRFSHFLSLECMEDSLSLGMN
EFEQNLKIIKQCQIGSIFSNSSNLPDDALLNLGRSLIYSAAGKGQKFSTPIEEEETVGFCWDLIIAIVLANIQRF
QTFWPYFHDYLLGVAQFPLFSPVPFAEKAIIGLFKVCLKILASCRVDKLPEELIFKSINLMWKLDKEILDTCCES
ITQSVSKIIIEYPANLQTYLGWKSVLHLLSVTGRHPETYDQGVETFIMLMSDSTHVSRINYAYCIDCAFAFVALR
NSPLDKNLKILDLLADSVNLLIQWYKSACSESGNNFSIVTNTSNSSLEENSKGLGSNFTMNFFVKLGEAFRKTSL
ARREEIRNHAVSSLQKSFILAEELDISSMNCINCFNLVIFAMVDDLHEKMVEYSRRENAEREMRSMEGTLKIAIE
MMTDVYLQFQKQIASNPGFRTFWLGLLRRMDTCMKADLGEYGETRLQETVPDLLRKMITKMKEEEILVQKEDDDD
DLWDITYIQIQWIVPSLKEELFPEQEI*

**>Poptr_LOC18107297**

MGRLKLNTGIKSIEEEPEERDAAVESNRADLLAYSINSEVSAVLAVMRRNVRWGGRYISGDDQLEDSLIQSLKTL
RKQIFSWQNPWHTINPALYLQPFLDVIRSDETGAPITGVALLSVYKILTLDVIDENTVNVEDAMQLVVDAVTSCR
FEVTDPSSEEMVLMKILQVLLACMKSKASVMLSNQHVCTIVNTCFRIVHQAGSKSELLQRISRHTMHELVKCIFS
HLPDVESAEQTLVNGVTSHKHEIGGLDNDYAFGSKQMENGNGNSELDGQASTVSFGSNASTALVAREENAIGTGG
GKDGLPFDLHLMTEPYGVPCMVEIFHFLCSLLNVVEHIGMGPRSNTIAFDEDVPLFALGLINSAIELGGPSIRCH
PRLLSLIQDELFRNLMQFGLSVSPLILSMVCSIVLNLYHHLRTELKLQLEAFFSCVILRLAQSRYGASYQQQEVA
MEALVDFCRQKTFMVEMYANLDCDITCSNVFEELANLLSKSAFPVNCPLSAMHILALDGLIAVIQGMAERIGNGS
VSSEQGPVNLEEYTPFWMVKCDNYSDPNHWVPFVRRRKYIKRRLMIGADHFNRDPKKGLEFLQGTHLLPDKLDPQ
SVACFFRYTAGLDKNLVGDFLGNHDEFCVQVLHEFAWTFDFQDMNLDTALRLFLETFRLPGESQKIQRVLEAFSE
RYYEQSPQILANKDAALLLSYSLIMLNTDQHNVQVKKKMTEEDFIRNNRHINGGNDLPREFLTELYHSICKNEIR
TTPEQGFGYPEMTPSRWIDLMHKSKKTAPFILSDSRAYLDHDMFAIMSGPTIAAISVVFDNAEHEDVYQTCIDGF
LAVAKISACHHLEDVLDDLVVSLCKFTTLLNQSSVEEPVLAFGDDAKARMATVTVFTIANRYGDYIRTGWRNILD
CILRLHKLGLLPARVASDAADESELAADPVHGKPITNSLSSVHMQSMGTPRRSSGLMGRFSQLLSLDTEEPRSQP
TEQQLAAHQRTLQTIQKCHVDSIFTESKFLQAESLLQLARALIWAAGRPQKGNSSPEDEDTAVFCLELLIAITLS
NRDRIVLLWQGVYEHIANIVQSTVMPCALVEKAVFGLLRICQRLLPYKENLADELLRSLQLVLKLDARVADAYCE
QITQEVTRLVKANATHIRSLMGWRTITSLLSITARHPEASEAGFDALLFIMTDEAHLLPANYVLCVDAARQFSES
RVGQAERSVRALELMAGSVNCLARWSHDAKETMGEEESAKLSQDIGEMWLRLVQGLRKVCLDQREEVRNHALLSL
QKCLTGVDEINLPHGLWLQCFDLVIFTMLDDLLEIAQGHQKDYRNMEGTLIIAVKLLSKVFLQLLNELAQLTTFC
KLWLGVLSRMEKYLKVKVKGKKNENLQETVPELLKNTLLAMKSRGVLVQRSALGGDSLWELTWLHVNNIAPSLQA
EVFPDQDREQSHHKLGETGGSLVSDETDSVSSKESVHAEVAGTGG*

**>Poptr_LOC18107296**

MGRIKLQSGIKVIEEEPEECGTSESYRATIVCMINAEAGTVLAVMRRNVRWGVHYMSGDDQLEDSLIQSLKTLRK
QIFSWQNPWHTINPALYLQPFLDVIRSDETGAPITGVALLSVYKILTLDVIDENTVNVEDAMQSVVDAVTGCRFE
VTDPASEEVVLMRILQVLLACMKSKASVMLSNQHVCILVNTCFRIVHQAGSKSELLQRISRHTMHELIRFIFSHL
PDVENTNQALVDGVATVKQESGGLGNDYALGSTQMENGNCNSELGGQASSVRFGSDASTGLAAIEENAIEENAIG
AGRGKDALPFDLHLMTLPYGVPCMVEIFHFLCSLLNVAEHIGKGPRSNTIAFYEDVPLFALGLINSAIELGGPSI
HCQPRLLSLIQDELFSNLMQFDLSTSPLILSMVCSIALNLYHHLRTELKLQLEAFFSFVILRLAQSRYGASYQQQ
EVAMEALVDFCRQKTFMVEMYANLDCDMTCNNVFEDLINLLSKSAIPMDCPLSSMHVLALDGLIAVIQGIVGRIG
NGSDGSEPAPVNIDEYTSFWTAKCDSYSDPNQWVPFIRQRKHIKRRLMVGADHFNRDPKKGLEFLQGTHLLPEKL
NPQSVACFLRYTSGLDKNLIGDFLGSHDEFSIQVLHEFAQTFDFEDMSLDNALRLLLETFRLPGESQKIQRVLEA
FSERYYEQSPQVLANKDAALLLSYSLILLNTDRHNAQVKKKMTEEDFIRNNRRINGGNDLPREFLSELYHSICKN
EIRTTPEQGFGIPEMTPSRWIDLMHKSKRTAPFILSDSRADLDHDMFAIMSGPTVDAISVVFDHAELEDIYQTCI
DGYLALAKISACHHLVDVLDAVVVSLCKFTTLLNPSSVEESVLAFGDDARAMMATVTVFTIANRYGDYIRTGWRN
ILDCVLRLYKLGLLPARVASDATEESELSADPVHGKTISNSLPSVHMQSMETPRRYTGLMGRFSQFLALDIEEPR
SQPTEQLAAHQRTLQTIQECHVGSIFTESKFLQAESLWELAQALILAAGHPQKGNSLPEDEDTAVICLELLITIT
LNNRDRILLLWQGVYEHIANIVQSTVMPCSLVEKAVFGLLRICQRLLPYKENLADELLRSLQLVLKLDARVADAY
CEQITQEVGRLVKANAAHIRSLMGWRTITSLLSITARHPDAAEAGFDALFFIMCDGSHLLPANYVLCVDAARQFA
ESRVGQAERSVCAVDLMGDSVSCLVRWSQDAKEAMGEEESAKLSEDIGEMWLRLVQGLRKVCLDQREEVRNRSLL
SLQKCLTGVDGIKLPHDLWLQCFDLVIFTMLDDLHEIAQGHQSNYRNMEGTLVIAVKLLSKVFLQLLPEIAQLTT
FCKLWVGVLSRMEKYLKVKVKGKKNEILQETVPELLKNTLLVMKSWGVLVQRSASGGDSLWELTWVHVNSIAPSL
QAEVFPDQGQEQSLHKLDETGCVPSNEPVHAECAAGTGG*

**>Poptr_LOC7458753**

MESTGGEEEERKLCKSYQQKRKRKEITSACMLNTELGAVLAVMRRPQQEPSFYTSDSQSDTNILNSLKSLRSLIF
NPQQEWQTIDPSIYISPFLDVVQGENIPAAATIVALSAILKILRLEIFNEKTPGARDAITCIVTGVSTCRLETTN
QACEEAVMLRILQVLTAVMRHSASILLSNHAVCTLVNTCFQVVQQSANRSDLLQRSARYAMHEVLQVIFSRLLEI
EVKPEQCESDTEDVDDGTDSDSGYGVRCAVDIFHFLCSLLNVVQVVETEGSTSHAADEDVQLFSLVLINSCIELG

GPEIGKHPKLLRMIQDDLFHHLIHYGTRSAPLLFSMICSIVLNIYHFLKRFIRLQLEAFFRFVILRVASTGSSVH
LQEVAVEAIINFLRQSSFIMEVYVNYDCHPTCLSVFEEIGKLLCKLAFPGAVPLTTIQVQAFEGLVIIMHTIAEN
IDNQGDSCPFGPYPVEITEYRPFWEEKSKDDLDLETWVEDSRIRRTQKKKLLIARDHFNRDEKKGLEYLKLCQLV
SDPADPKDIAMFFRYTPELDKNMMGDYLGDPDEFHLRVLREFAETFRFSGVILDTALRTYLAAFRLPGESQKIQR
ILEAFSDRFYDQQSSDIFASKDAVFILCYSLIMLNTDQHNPQVKKKMTEEEFIRNNRAINGGQDLPREYLSELFQ
SIATNPIAVFGQSGLLVEMNPGRWMELMNQSKVMQLYIQCDFDRQLGRDMFACVAGPSIAALSAFFEHSDEDEMF
HECIEGLMSVARIAQYGLEDTLDELIASFSKFTTLLNPYASAEETLFAFSNDMKPKMATLAIFTIANSFGDSIRA
GWRNIVDCLLKLKRLKLIPESAIDFDNAASANLSTESGVISPSHDPKFGDNQTPNAISRFSQFLSVESMEDSLSL
GMSEFERNLKVIKQCRIGSIFSNSSTLPEDTVLNLGRSLIFAAAGKGQKFSTPVEEEETVGFCWDLVTVIALANI
HRFQTFWPSFHDYLLVVAQFPLFSPIPFAEKGILCLFKVCLKLLSSSRADKISEELIFKSINLMWKLDKEILDTC
CESITNTISKILIGYPENLQTHLGWKSALHMLSVSGRHLETYEQGVETLITLMSDTGHVSRINYAYCIDCAFGFV
ALKNSPLEKNLKILDLLSDSVNLLIHWYRNYSDPGSNHSMVSNASNSSVEDIIKGSGNYTMNLFIKLGEAFRKTS
LARREEMRNHAIASLQKSFTLAEELDFSPVNCINCFNLVIFAMVDDLLEKMVEYSRRENAEREMRGMEGTLKLAM
ELLTDVYMLFLKQIAASPGFRTFWLGVLRRMDTCMKADLGVWGETKLQQIVPSLLRRMITKMKEEEILVQKEGDD
LWDITDIQIQWIAPSLKEELFPDEI*

**>Salbr_026992**

MGRLKLNTGIKSIEEEPEEHDAAVESNRVDLLAYVINSEVSAVLAVMRRNVRWGGRYISGDDQLEDSLIQSLKTL
RKQIFTWQNPWHTIDPALYLQPFLDVIQSDETGAPITGVALLSVYKILTLDVIDQYTLNVEDAMQLVVDAVTSCR
FEVTDPASEEMVLMKILQVLLACMKSKASVMLSNQHVCTIVNTCFRIVHQAGSKSELLQRISRHTMHELVKCIFS
HLPDVESAERTMVNGVASLKQEIGGMDNDYAFGSKQMENGNGNSELDGQASTVTFGSNASTALVAREENAIGTGG
GKDALPFDLHLMTEPYGVPCMVEIFHFLCSLLNVVEHSGMGTRSNTIAFDEDVPLFALGLINSAVELGGPSIRCH
PRLLSLIQDELFRNLMQFGLSMSPLILSMVCSIVLNLYHHLRTELKLQLEAFFSCVILRLAQSRYGASYQQQEVA
MEALVDFCRQKTFMVEMYGNLDCDITCSNVFEELANLLSKSAFPVNCPLSAMHILALDGLIAVIQGMAERIGNGS
VSSEQGPVHLEEYTPFWMVKCDDYSDPNHWVPFVRRRKYIKRRLMIGADHFNRDPKKGLEFLQGTHLLPDKLDPQ
SVACFFRYTAGLDKNLVGDFLGNHDEFCVQVLHEFAWTFDFQDMNLDTALRLFLETFRLPGESQKIQRVLEAFSE
RYYEQSPLILANKDAALLLSYSLIMLNTDQHNVQVKKKMTEEDFIRNNRHINGGNDLPREFLTELYHSICKNEIR
TTPEQGFGYPEMTPSRWIDLMHKSKKTAPFILSDSRAYLDHDMFAIMSGPTIAAISVVFDNAEHDDVYQTCIDGF
LAVAKISACHHLEDVLDDLVVSLCKFTTLLNSSLVEEPVLAFGDDAKARMATVTVFTIANRYGDYIRTGWRNILD
CILRLHKLGLLPARVASDAADESELTADPVHGKPIANSLSSVHMQSMGTPRRSSGLMGRFSQLLNLDTEEPRSQP
TEQQLAAHQRTLQTIQKCHVDSIFTESKFLQAESLLQLARALIWAAGRPQKGNSSPEDEDTAVFCLELLIAITLN
NRDRIVLLWQGVYEHIANIVQSTVMPCALVEKAVFGLLRICQRLLPYKENLADELLRSLQLVLKLDARVADAYCE
QITQEVTRLVKANATHIRSLMGWRTITSLLSITARHPEASEAGFDALLFIMTDAAHLLPANYVLCVDAARQFSES
RVGQAERSVRALELMAGSVNCLARWSHDAKETMGEEESAKLSQDIGEMWLRLVQGLRKVCLDQREEVRNHALLSL
QKCLTGVDEINLPHGLWLQCFDLVIFTMLDDLLEIAQGHQKEYRNMEGTLIIAVKLLSKVFLQLLNELAQLTTFC
KLWLGVLSRMEKYLKVKVKGKKNENLQETVPELLKNTLLVMKSKGVLVQRSALGGDSLWELTWLHVNNIAPSLQA
EVFPDQDREQSHHKLGETGGSLVSDETDSVSSKESVHAEVVGTGG*

**>Salbr_026991**

MGRIKLQSGIKVIEEVPEECGTSESYRATIVCMINAEAGTVLAVMRRNVRWGVHYMSSDYQLEDSLIQSLKMLRK
QIFSWQNPWHTINPSLYLQPFLDVIRSDETGAPITGVALLSVYKILTLDVIDQNTVNVEDAMQSVVDAVTGCRFE
VTDPASEEVVLMRILQVLLACMKSKASVMLSNQHVCTIVNTCFRIVHQAGSKSELLQRISRYTMHELIRFIFSHL
PDVKNTRQALVDEVNTVKQESGGLDNDYAFGSTQIENGNGNSELGGQASSVRFLSDASTGLAAIEENATGAGSGK
DALPFDSHLMTVPYGVPCMVEIFHFLCSLLNAVEHIGKGSRSNTIAFDEDVPLFALGLINSAIEVGGPSIRCQPR
LLSLIQDELFSNLMQFGLSMSPLILSMVCSIVLNLYHGLRTELKLQLEAFFSFVILRLAHSRYGASYQLQEVAME
ALVDFCRQKTFMMEMYANFDCDMTCNNVFEDLTNLLSKSAIPMNCPLSSIHILALDGLIAVIQGIVGRIGIGSVG
SEPAPVNIDEYTSFWTVKCDNYSDPYQWVPFIRRRKHIKRRLMVGADHFNRDPKKGLEFLQGTHLLPEKLNPQSV
ACFLRYTSGLDKNLIGDFLGSHDEFSIQVLHEFAGTFDFEDMSLDNALRLLLETFRLPGESQKIQRVLEAFSERY
YDQSPQVLANKDAALMLSYSLILLNTDRHNAQVKKKMTEEDFIRNNRRINGGNDLPREFLSELYHSICKNEIRTT
PEQGFSYPEMTPSRWIDLMHKSKRMAPYILSDSRAHLDHDMFAIMSGPTIDAISVVFDHAELEDIYQTCIDGFLA
IAKISACHHLVDVLDSVVVSLCKFTTLLNPSSVEESVLAFADDAKAMMATVTVFTIANRYGDYIRTGWRNILDCV
LSLYKLGLLPARVASDAAEESELSADPVHGKTISNSLPSVHMQSMEIPRRYTGLMGRFSQFLALDIEEPRSQPTE
QLAAHQHTLQTIQECHVGSIFTESKFLQAESLLELARALILAASHPQKGNSLPEDEDTAVICLELLITVTLNNRD
RILLLWQGVYEHIANIVQSTIMPCSLVEKAVFGLLRICQRLLPYKENLADELLRSLQLVLKLDARVADAYCEQIT
QEVGRLVKANAAHIRSLTGWRTITSLLSITARHPDAAEAGFDALFFIMSDGSHLLPANYVLCLDAARQFAESRVG
QAERSMCAVDLMGDSVGCLARWSQNAKEAMGEEESAKLSEDIGEMWLRLVQGLQKVCLDQREQVRNHSLLSLQKC
LTGVDGINLPHDLWLQCFDLVIFTMLDDLHEIAQGRQNNYRNMEETLVIAVKLLSKVFLLLLPEIAQLTTFCKLW
VGVLSRMEKYLKVKVKGKKNEILQETVHELLKNTLLVMKSWGVLVQSSASGGDSLWELTWLHVNTIAPSLQAEVF
PDLDPERSPHKLDETGYFPSNESVHAEFAGTGG*

**>Salbr_028175**

MESTGGEEEERKPCKSYQQKKKRKEITSACMLNTELGAVLAVMRRPQQEPSIYTSESQSDTNIINSLKSLRSLIF
NPQQEWQNIDPSIYISPFLDVVQGENIPAAATIVALAAILKILRLEIFNEKTPGARDSITCIVTGISTCRLETTN
QACEEAVMLRILQVLTAVMRHSASILLSDHAVCTLVNTCFQVVQQSANRSDLLQRSARYAMHEVLQVIFSRLLEI
EVKPEQCESDTEDFEDGTDSDSGYGVRCAVDIFHFLCSLLNVVQVVETEGSTSHAADEDVQLFSLVLINSCIELG
GDEIGKHPKLLRMVQDDLFHHLIHYGTRSAPLLFSMICSIVLNIYHFLKRFIRLQLEAFFRFVILRVASTGSSVH

LQEVAVEAIINFLRQPSFIMEVYVNYDCHPTCLNVFEEIGKMLCKLAFPGAAPLTTIQVQAFEGLVIIMHTIADN
IDNQRDSCPFGSYPVEISEHRPFWEEKSKDDLDLKTWVEHSRVRRTQKKKLLITRDHFNRDEKKGLEYLKLCQLV
SDPADPKDIAMFFRYTPELDKNMIGDYLGDPDEFHLQALREFAETFKFSGMILDTALRTYLAAFRLPGESQKIQR
ILEAFSDRFYDQQSSDVFASKDAVFILCYSLIMLNTDQHNPQVKKKMTEDEFIRNNRAINGGQDLPREYLSELFQ
SISNNPIAVFGQSGVLAEMDPGRWMELMNQSKVMQLYIQCDFDRQLGRDMFACVAGPSIAALSAFFEHSDEDEMF
HECIEGLMSVARIAHYGLEDTLDELISSFSKFTTLLNPYASAEETLFAFSNDMKPKLATLAIFTIANCFGDSIRA
GWRNIVDCLLKLKRLKLIPESAIDFDIAASANLSTESGVISPSHDPKCGDNQTPNAISRFSQFLSVESMEDSLSL
GMSEFERNLKIIKQCRIGSIFSNSSTLPEATVLNLGRSLIFAAAGKGQKFSTPVEEEETVGFCWDLVTAIALANI
HRFQTFWPSFHDYLLVVAQFPLFSPIPFAEKAILGLLKVCLKLLSSSRADKISEELIFKSINLTWKLDKEILDTC
CECITNTISKILIEYPENLQTHLGWKSALHMLSVSGRHLETYEQSVETLITLMSDSGHVSRINYAYCIDCAFGFV
ALKNSPLEKNLKILNLLSDSVNLLIHWYRNYSDPGSNHSMVSNASSSSVEDSIKGSSNYTMNLFIKLGEAFRKTS
LARREEMRNHAIASLHKSFTLAEELEFSPVNCINCFNVIIFAMLDDLHEKMVEYSRRENAEREMRSMEGTLKLAM
ELLTDVYLLFLKQIAASPGFRTFWLGVLRRMDTCMKADLGVWGETRLQQIVPYLLRRMIMKMKEAEILVQKEGDD
LWDITDIQIQWIAPSLKEELHCRT*

**>Kanob_002465a**

MGRLKLQSGIKAIEEEPEDLDNFYSNKTTLACMINAEVGAVLAVMRRNVRWGGRYMSGDDQMEHSLIQSLKALRK
QIFSWQHPWHTINPSVYLQPFLDVIRSDETGAPITGVALSSLYKILTLDMIDQHTVTVEDAMHLVVDAVTSCRFE
VTDPASEEVVLMKILHVLLACMKSKASIMLSNQHVCTIVNTCFRIVHQAGSKGELLQRIARHTMQELVRCIFSHL
PDVDNTECALVHGVNPVQIGGPDNDYGFGSKQMENSNGSSGVDGQASCVSLGSNASMVEGTSVMEESTMGVNNKK
DVLPYDLHLMTEPYGIPCMVEIFHFLCSLLNVVEHVGMDPRSNTIAFDEDVPLFALSLINSAIELGGPSICHHPR
LLGLIQDELFRNLMQFGLSMSPLILSLVCSIVLNLYHHLQTELKLQLEAFFSCVILRLAQSRFGASYQQQEVAME
ALVDFCRQKTFMVEMYANLDCDITCSNVFEDLANLLSKSAFPVNCPLSAMHILSLDGLIAVIQRMAERIGNGSVT
SEQPPVNIEGYIPFWMVKCDDYNDPNHWLPFVHRRKYIKRRLMIGADHFNRDPKKGLEFLQGTHLLPDKLDPQSV
ACFFRFTAGLDKNLVGDFLGNHDEFCVQVLHEFAGTFDFQDMNLDTALRLFLETFRLPGESQKIQRVLEAFSERY
YEQSPHILANKDAALLLSYSLIMLNTDQHNVQVKKKMTEEDFIRNNRHINGGNDLPREFLSELYHSICKNEIRTT
PEQGAGFPEMTPSRWIDLMHKSKKTAPFIVSDSKAYLDHDMFAIMSGPTIAAISVVFDHAEHEDVYHTCIDGFLA
VAKISACHHLEDVLDDLVVSLCKFTTLLNPSAVEEPVLAFGDDTKARMATVTVFTIANRYGDYVRTGWRNILDCI
LRLHKLGLLPARVASDAADESEVSPDPGHGKPLTNSLSSVHMQAIGTPRRSSGLMGRFSQLLSLDSEEPRSQPTE
QQLAAHQRTLQTIQKCHIDSIFTESKFLQAESLLQLARALIWAAGRPQKGNNSPEEEDTAVFCLELLIAITLNNR
DRIVLLWQGVYEYIANIVQSTVVPGALVEKAVFGLLRICQRLLPYKENLADELLRSLQLVLKLDARVADAYCEQI
TQEVSRLVKANATHIRSQMGWHTITSLLSITARHPEASEAGFDALVFIMSDGALLLPANYVLCVEAARQFAESRV
GQAERSVHALDLMAGSLNCLAQWFQEAKQAMGEEEAAKSSQDIGEMWLRLVQGLRKVCLDQREEIRNHALLSLQK
CLMGVDGINLPHVLWAQCFDHVIFTMLDDLLEIAQGHSQKDYRNLEGTLILAVKLLSKIFLRLLHDLSQTTFCKL
WLGVLSRMEKYVKIKVRGKKSEKLHEVVPELLKNMLLVMKTRGVLVHRSALGGDSLWELTWLHVNNIAPSLQSEV
FPNQNWEQSQPKQGEQDYSLVQPIVLNSFSDSPSLLRKTHRGFRTESSEICLIHAD*

**>Kanob_002465b**

**MGRLK**LQSGLEAVKEECGTSCSKENSLACMINAEIGAVLAVMRRNVRWGGRYGDDQLDHSLIHSFKLLRKQLFSW
RNPWHTIDPSLYLQPFLDVIRSHETGAPITGVALSSVYKILTLDVIDQNTVNVKDAMNLVVDAVTGCRFEVTDPA
SEEVVLMKILQVLLACVKSKALVLLSNQHVCTIVNACFRIVHQAGTKGELLQRIARHTMHELVRCIFSHLADIGH
HECALVNGLDAVKGKISATNNDYAFGSKKLENGNNGSEVDGKPFPATLGPSVSTVQVASVVEENMIGADSSCDDF
AYDVHLMTEPFGLPCMVEIFHFLCSLLNVVEHVGMDPKLNTIAFDEDLPLFALGLINSAIELGGLSIHHHPRLLS
LIQDELFHNLMQFGLSRSPLILSMVCSIVLTLYDNLRTELKLQLEAFFCCVILRHAQSRYGVSYQQQEVAMEALV
DFCRQSTFMVEMYANFDCDITCSNLFEDIANLLSKSAFPVNSQLSAIHILALDGLIAVIQRISERIGKGSASSEQ
ASLNIEEYTPFWTMKCDNYGDPNKWVPFVRRRKYIKRRLLIGADHFNHDPKKGLEFLQGAHLLPDKLDPKSVACF
FRCTAGLDKNLIGDFLGNHDEFCVQVLHEFAGTFDFQDMNLDHALRLFLETFRLPGESQKIQRVLEAFSERYYEQ
VPHILANKDAVLLLSYSIILLNTDQHNVQVKKKMTEEDFIRNNRCINGGDDLPRDFLSELYQSICRNEIRTTPEK
GVGFPEMTPSRWIYLMHKSKNTAPFIVLDSNAHIDHDMFAIMSGPTVAAISVVFDHAEHEDVYQTCIDGFLSVAK
VSACHHLEDVLDDLVVSLCKFTNLLNPSFEELVLAFGEDTKARMATVAVFTIANMYGDYIRTGWKNILDCILRLH
KVGLLPAHVVTDTVDESELFVDSDQGKPPINPLPSVKMQSVGTIRKSAGLIGRVSQLLSLDTEEPISQPTEQELA
VYQSTLQTVQKCHIQSIFTESKFLQAESLVNLAKALVRAAGQSERGNSSLEDEDATVFCLELLIAITLNNRDRIL
LLWQGVYEHIAAVVQSTVLPCALVEKAVFGLIRICQRLLPYKENLADELLRSLQLVLKLDARVADAYCEQITQEV
SRLVKANATHIRSLTGWRIVISLLSVTARHPEASDAGFDALFFIMSNGAHLLPANYVLCVDASRQFAESRVGQAE
RSMCALDLMADSVDCLVRWSNEAKEAMREEEAVKLSQNIGDMWLSLVQGLRNTCLDPREEVRNHSLLSLQKCLIV
VDGIKFPHDLWLQCYDLVIFATLDDLLEIAQGRSQKVYKNMEGTLVIAIKLLSKLFLQLPNELEQSTTFSKLWFG
VLERMEKYMKVKVRGKKSDKLQEIIPELLKNILLVMKTRGVLVQSSAVSNSLWEPTWVHVNNIAPSLQSEVFPAQ
ELERSEHKQGETGSVPLIKSVASEGIGTGS*

**>Rhiap_sc34.1**

MGRLKLQSGIKAIEEEPEDLDNFYSNKTTLACMINAEVGAVLAVMRRNVRWGGRYMSGDDQMEHSLIQSLKALRK
QIFSWQHPWHTINPAVYLQPFLDVIRSDETGAPITGVALSSLYKILTLDIIDQNTVNIEDAMHLVVDAVTSCRFE
VTDPASEEVVLMKILHVLLACIKSKASIMLTNQHVCTIVNTCFRIVHQAGTKGELLQRIARHTMQELIRCIFFHL
PDVDNTEHALVHGVNSVQVQIGGPDNDYTFGSKQMENSNGSSGVDGQASCVSLGSNASMVEGTSVMEESTMGVSN
KRDGLPYELHLMTEPYGIPCMVEIFHFLCSLLNVVEHVGMDPRSNTVAFDEDVPLFALSLINSAIELGGPSICHH
PRLLGLIQDELFRNLMQFGLSMSPLILSLVCSIVLNLYHHLRTELKLQLEAFFSCVILRLAQSRFGASYQQQEVA

MEALVDFCRQKTFMVEMYANVDCDITCSNVFEDLANLLSKSAFPVNCPLSAMHILSLDGLIAVIQRMAERIGNGL
VTPEQPPVNIEEYTPFWMVKCDDYNDPNHWVPFVRRRKYIKRRLMIGADHFNRDPKKGLEFLQGTHLLPDKLDPQ
SVACFFRFTAGLDKNLVGDFLGNHDEFCVQVLHEFAGTFDFQDMNLDTALRLFLETFRLPGESQKIQRVLEAFSE
RYYEQSPHILANKDAALLLSYSLIMLNTDQHNVQVKKKMTEEDFIRNSRRINGGNDLPREYLSELYHSICKNEIR
TTPEQGAGFPEMTPSRWIDLMHKSKKTAPFIVSDSRAYLDHDMFAIMSGPTIAAISVVFDHAEHEDVYQTCIDGF
LAVAKISACHHLEDVLDDLVVSLCKFTTLLNPSSVEEPVLAFGDDTKARMATVTVFTIANRYGDYVRTGWRNILD
CILRLHKLGLLPARVASDAADESEVSPDPGHGKPLTNSLSSVHMQSIGTPRRSSGLMGRFSQLLSLDSEEPRSQP
TEQQLAAHQRTLQTIQKCHIDSIFTESKFLQAESLLQLARALIWAAGRPQKGNSSPEDEDTAVFCLELLIAITLN
NRDRIVLLWHGVYEYIANIVQSTLMPGALVEKAVFGLLRICQRLLPYKENLADELLRSLQLVLKLDARVADAYCE
QITQEVSRLVKTNATHIRSQMGWHTITSLLSITARHPEASEAGFDALVFIMSDGAHLLPANYVLCVDAARQFAES
RVGQAERSVHALDLMAGSLNCLVWWSQEAKQAMGEEEAAKSSQDIGEMWLRLMQGLRKVCLDQREEIRNHALLSL
QKCLTGVDGINLPHGLWPQCFDHVIFTMLDDLLEIAQGHSQKDYRNMEGTLILAVKLLSKIFLRLLHDLSQTTFC
KLWLGVLSRMEKYVKIKVRGKKSEKLHEVVPELLKNILLVMKTRGVLVHRSALGGDSLWELTWLHINNIAPSLQS
EVFPNQNWEQSQPKQGEQGGSLVSDEASPIPSSEPVALEGGGTRS*

**>Rhiap_sc34.2**

MGRLKLQSGFEAVKGEECGTSCSKENSLACMINAEIGAVLAVMRRNVRWSGRYGDDQLDHSLIQSFKSLRKQLFS
WRNPWHTIDPSLYLQPFLDVIRSDETGAPITGVALSSVYKILTLDVIDRNTVNVKDAMHLVVDAVTSCRFEVTDP
ASEEVVLMKILQVLLACVKSKASVLLSNQHVCTIVNACFRIVHQAGTKGELLQRIARHTMHELVRCIFSHLADIG
HEECALVNGLDAVKQQIGATNNDYAFGSKKLENGNNGSEFDGKPSPANLASNVSTVQVASGVEGNMIQAGSSHDD
FSYDVHLMTEPFGVPCMVEIFHFLCSLLNVVEHVGMDPKLNTIAFDEDLPLFALGLIYSAMELGGPSMRHYPRLL
SLIQDELFRNLMQFGLSRSPLILSMVCSIVLTLYDNLRTELKLQLEAFFCCVILRHAQSRYGASYQQQEVAMEAL
VDFCQQSTFMVEMYANFDCDITCSNVFEDIANLLSKSAFPVNSQLSATHILALDGLIAVIQRIAERIGNGLASSE
QAPLNIEEYSPFWTMKCDKYGDPNEWVPFVRRRKYIKRRLLIGADHFNHDPKKGLEFLQGAHLLPDKLDPKSVAC
FFRYTAGLDKNLIGDFLGNHDEFCVQVLHEFSGTFDFQDMNLDHALRLFLETFRLPGEAQKIQRVLEAFSERYYE
QVPHILANKDAVLLLSYSLILLNTDQHNVQVKKKMTEEDFIRNNRSINGGNDLPREFLSELYQSICRNEIRTTPE
KGVGFPEMTPSRWIYLMHKSKNTAPFIVSDSNAHIDHDMFAILSGPTIAAISVVFDHAEYEDVYETCIDGFLSVA
KVSACLHLEDVLDDLVVSLCKFTNLLNPSFEELVLAFGEDTKARMATMAVFTIANTYGDYIRAGWKNILDCILQL
HKVGLLPAHVATDAADESELFVDSEQGRPPINSLPSVKMQSVGTMRRSAGLIGRVSQLLSLDTEEPRSQPTEQEL
AVYQSTLQTVQKCHIDSIFTESKFLQAESLVNLAQALIRAAGKSERGNSSLEDEDTAVFCLELLIAITLNNRDRI
LLLWQGVYDHIAAVVQSTVLPCALVEKAVFGLIQICQRLLPYKENLADELLRSLQLVLKLDAHVADAYCEHITQE
VSRSLTGWRIVISLLSVTARHPEASDAGFDALFFIMSNGAHLLPANYVLCVDASRQFAESRVGHVERSVCALDLM
ADSVDCLIRWFNEAHEAMGEEEATKLSQDIGDMWLRLVQGLHNTCLDPREEVRNHSLLSLQKCLIVVDGIKFPHS
LWLQCCDLVIFTMLDNLLEIAQGRSQKEYKNMEGTLVMAIKLLSRLFLQLLNELGKLTTFSKLWLEVLARMEKYM
KVKVRGKKSDKLQEIIPELLKNILLVMKTRGVLVQGSAVSNSLWEPTWVHANNIAPSLQSVVFPAQELEQSEHEQ
GETASVPSSKSIASEGIGTGS*

**>Rhiap_sc6.10**

MGGEEDETESPSSLQEKKKRKELGLACMLNTEVGAVLAVLRRPQFPTAATFLSQEENYDTPTLQALKSLRALIFN
PQQEWKTIDPSTYLSPFLDVIQSDDVSAQATGVALSAILKIFRLEIFDEKTPRARDVMHSVVAGITSCRLEKTDP
ISEDAVIMRILQVLTAIMKHSASVLLADQAVCTVVNTCFQVVQQSASRGDLLQRSARYTMHELIQIIFSRLPDVE
IKPGEDSESDTEDVDIDDGLESGYGIRCAVDIFHFlCLLLNVVEVVETEGYTYHTADENVQLFALVLINSAIELS
GDAIGKHPKLLRMVQDDLFHHLIHYGTCSSPLVLSMICSTVLNIYHFLRRFVRLQLEAFFGFVLLKVAAYEGSVQ
LQEVALEGIINFCRQPTFALEAYANYDCDPTCRNIFEEIGKLICKLSFPRASPLNTIQVQAFEGLLIIVHNIADN
VDKEGDLIPCGPYPVDIAEYRPFWEEKPKDNMETWVEYLRVRKVQKRKTLIAGNHFNLDEKKGLEYLKLCQLVSD
PPDPKAFAIFFRYTPRLDKNMIGDYLGDPDEFHIQVLKEFTATFHFSGMILDTALRTFVATFRLPGESQKIERIL
EAFSEKFYEQQSSDIFASKDAVFILCYSLIMLNTDQHNPQVKKKMTEDEFIRNNRAINGGQDLPREYLSDLFHSI
ANHAITLSGQSGPVEMNPARWMELINRSKVTQPLIHCDFDRRIGRDMFACISGPSVAALSAFFEHADEDEIFHEC
IEGLISVARIAQHGLQDTLDELIALFSKFTTLLNPYASAEETMFAFSHDMKARMATLAVFTIANSFGESIHGGWR
NIVECLLKLKRLKLLPQSVVEFDSTLGLSSDNPEHLKNESGLIFPSHDAKFGKCHASVMISRFSHFLSIDSMEDS
VSLGMSEFEQNLKIMKQCRIGSIFSNSSKLPDVTLLNLGRSLIFAAGGRGQKFSTPVEEEETVGFCWDLIVTISM
TNIHRVQDFWPSFHDCLLGVTQLPLFSPIPFAEKGIIGLFTVCTDLLSSFRPEKYSEEHIFKSINLMWKLDKEIL
DTCCEFITNSVSKILMEYAANLQSHLGWKTVLHLLSVSGRHAETYNQGVQTLISLISDTARLSPTNYAYCVDCAF
GFVALKNSPAEKNVKILDLLADSVNVLIQWYTEYSDPGSNHSMASNTSNSSLDDNSKGFGSHNFVFTLFVKLGEA
FRKTSLARREEIRNHAILSLTKSFSLAQGLDFSPVNCTNCCNLVVFAMVDDLHEKMLEYSRRENAEREVRSMEGT
LKLSMELLTDVYLQFLKQISVSPGFRTFWLGVMRRMDTCMKADLGAYGETKLQYVVPELLRKMITKMKQEEILER
QEGDDLWEITYIQIQWIAPALKEELFPDEEI*

**>Jatcu_LOC105630489**

MGRLKLQSGIKAIEEEPEECDSSYSNKATLACMINAEVGAVLAVMRRNVRWGGRYMSGDDQLEHSLIQSLKALRK
QIFSWQHPWHTINPAVYLQPFLDVIRSDETGASITGVALSSVYKILTLDVIDQNTVNVEDAMHLVVDAVTSCRFE
VTDPASEEVVLMKILQVLLACMKSKASVTLSNQHVCTIVNTCFRIVHQAGSKGELLQRIARHTMHELVRCIFSHL
PDVDNTEHALVNGVSPAKQEIGGLDNDYTFGRKQMENGNSSSEFDGQVSSVSFSSSASTGLVATVMEENTIGASG
GKDALPYDLHLMTEPYGVPCMVEIFHFLCSLLNVVEHMGMGPRSNTIAFDEDVPLFALGLINSAIELGGPSIRHH
PRLLSLIQDELFRNLMQFGLSMSPLILSMVCSIVLNLYHHLRTELKLQLEAFFSCVILRLAQSRYGASYQQQEVA
MEALVDFCRQKTFMVEMYANLDCDITCSNVFEDLANLLSKSAFPVNCPLSAMHILALDGLIAVIQGMAERIGNGS

VSSEQAPVNLEEYTPFWMVKCNNYSDPSHWVPFVRRRKYIKRRLMIGADHFNRDPKKGLEFLQGTHLLPDKLDPQ
SVACFFRYTAGLDKNLVGDFLGNHDEFCVQVLHEFAGTFDFQGMNLDTALRLFLETFRLPGESQKIQRVLEAFSE
RYYEQSPQILANKDAALLLSYSLIMLNTDQHNVQVKKKMTEEDFIRNNRHINGGNDLPREFLSELYHSICKNEIR
TTPEQGVGFPEMTPSRWIDLMHKSKKTAPFIVSDSIAYLDHDMFAIMSGPTIAAISVVFDHAEHEDVYQTCIDGF
LAVAKISACHHLEDVLDDLVVSLCKFTTLLNPSSVEEPVLAFGDDTKARMATVTVFTIANRYGDYIRTGWRNILD
CILRLHKLGLLPARVASDAADESELSADPGHGKPITNSLSSVHMQSMGTPRRSSGLMGRFSQLLSLDTEEPRSQP
TEQQLAAHQRTLQTIQKCHVDSIFTESKFLQAESLLQLARALIWAAGRPQKGNSSPEDEDTAVFCLELLIAITLN
NRDRIVLLWQGVYEHIANIVQSTVMPCALVEKAVFGLLRICQRLLPYKENLADELLRSLQLVLKLDARVADAYCE
QITQEVSRLVKANATHIRSIMGWRTITSLLSITARHPEASEAGFDAILFIMNDGAHLLPANYVLCVDAARQFAES
RVAQAERSVRALDLMAGSVDCLSRWSDEAKEAMGEEEAAKLLQDIGEMWLRLVQGLRKVCLDQREEVRNHALLSL
QKCLTGVDGINLPHGLWLQCFDLVIFTMLDDLLEIAQGHSQKDYRNMDGTLIIAVKLLSKVFLQLLHDLSQLTTF
CKLWLGVLSRMEKYLKVKVRGKKSEKLQEVVPELLKNTLLVMKTKGVLVQRSALGGDSLWELTWLHVNNIAPSLQ
AEVFPDQEWELSQHKQGETGGGLVSDETGSVPSNGSVASEG*

**>Jatcu_LOC105633077**

MGRLKLQSGIKAMEEESVGCDSSYSNKATLACMINAEVGAVLAVMRRNVRWGGRYVSGDDQLDHPLIQSLKALRK
QIFSCQHPWHTINPAVYLQPFLDVIRSDETSAPITGVALSSVYKILTVDVIDQNTVNVKDAMHLVVDAVTSCRFE
VTDPATEEVVLMKILQVLLACMKSKASATLSNQHVCTIVNTCFRIVHQAGTKGELLQRIARHTMHELVRCIFSLL
PDVDNKEYAMVNGVSPTKQEIRGLDNDYSFSSKQMENGNNGSEFDGQASSVSFGSSASSDLVATVMEENTIGVHS
GKNALPHDLHLMTQPYGLPCMVEIFHFLCSLLNVVDHMGVGPRSNTIAFDEDVPLFALGLINSAIELGGPYIRHH
PKLLSLIQDELFSNLMQFGLSMSPLILSMVCSIVLNLYHHLRAELKLQLEAFFSCVILRLAQSRYGASYQQQEVA
MEALVDFCRQKTFMVEMYANLDCDITCGNVFEDLANLLSKSAFPVNCPLSAMHILALDGLIAVIQGMAEGIDNGS
VGSEQAPLNLEEYTPFWMEKCDNYSDPTHWVPFVLQRKHIKRRLMIGADHFNRDPKEGLKFLQGTHLLPNKLHPQ
SVACFFRYTAGLDKNVIGDFLGNHDEFCVQVLHEFARTFDFQGMNLDDALRLFLETFRLPGESQKIQRVLEAFSE
RYYEQSPQILANKDAALLLSYSLIMLNTDQHNVQVKKKMTEEDFVRNNRRINGGNDLPREFLSELYHSICKNEIR
TTPEQGAGFPVMTSSRWISLIHNSKKTSPFIVYDSKAYLDRDMFAIMSGPAIAAISVVFDHAEHEDIYQTCIDGF
LAVAKISACHHFEHVLDDLVVSLCKFTTLLDPSYPEEHVLAFGDDTKARMATVALFTIANRYGDYIRTGWRNIME
CILRLHKLGLLPACVASDVADESETSADPGHGKPITNPLPSAHMQSTGAPKKSSGLMGRFSQLLSLETEEPISQP
TEEELAAQRRTLETIQKCHVDSIFAESKFLQAESLEQLAQALIWAAGGSQRGNSFPEDEDTAVFCLELLIAITLN
NRDRIVRLWQSVYEHIANIVQSTVMPCALVEKAVFGLLRICQRLLPYKENLADELLRSLQLVLKLDARVADAYCE
QITVEVSRLVKANATHIRSLMGWRTITSLLSITARHPEASEPGFDALSFIMSDGVHLLPANYVLWVDAARQFAES
RVAEAARSVHALDLMTGSVDCLVRWSHEAKESLGEEEAAKLSQDIGEMWLRLVQGLRKVCLDQREEVRNHALLSL
QKCLTVVDGINLPPGLWLQCFDLVIFTLLDDLLENAQGHTHKGYRNMDGTLIIAVKLLSKVFLRLLHDLSQLTTF
SKLWLEVLSRMEKYLKVKVRGRKSEKLHEAVPELLKNTLLVMKTSGVLVQSSASGGDSLWELTWQHVNKIAPSLQ
SEVFPDQELEKLQPKNAESGGGIMSDETGSVPPNVSVASGG*

**>Jatcu_LOC105638327**

MEEDEENKLHSASWREKKRRKELSLSCMLNTELSALLAVIRRPIDPTASKIQEDNFDTSVLHSLKSLRALIFNPQ
QEWRTIDPSVYISPFLDVIQSDDIPAVATNVALSAISKILKLQIFDEKTPGAKDAINFIVTGITSCRLERTDATN
EDAVMMRILQALTCIIKHRASVLLTDHAVCTIVNTCFQVVQKSANRADLLQRGARYSMREMIEIIFARLQDVEVK
LEEDSESDTEDIDIDSDMDSGYGVRCVVDIFHFLCSLLNVVEIVETEGVQTHACDENVQIFGLILINSAVELSGD
AIGKHPKLLRMIQDDLFHHLIHYGVSSSPLVLSMICSTVLNIYHALRRFIRVQLEAFFRFVLLRAASHGASQLQE
VALEGIINFCRQPAFIAEMYVNYDSDPICQNIFEEIGKLLCKLSFPGYSPLSCVQIQAFEGLLIIAHTIADNVDK
EDDSTPYGPYPVEITEYKPFWEEKSRDDYEAWVEHLRLRKAQKRKISIAGDHFNRDEKKGLEYLKLCQLVSDPAD
PKAIAMFFRYTPGLDKATIGDYLGDPDDFHLQVLKEFTQTFRFSGMVLDTALRTYLATFRLPGESQKIERILEAF
SDRFYDQQSSDIFATKDAVYIFCYSLIMLNTDQHNPQVKKKMTEEEFIRNNRAINAGQDLPREYLSELFNSIAAN
AIALFGKSGPVEMNPGNWIELINRSRVTQPYILCNFDRRIGRDMFAAIAGPTVAAISSFFEHADEDEMLHECIGG
LISVSRIAQYGLEDILDELLASFSKFTTLLNPYASAEETLFAFSNDLKPRMATLAVFSIANNFGDSIRGGWKNII
DCLLKLKRLKLLPQSVVEFDDTPASSSGGHGHRRNESNLSSSHDPRFGTRRGAGMTSRFSHFLTLESMEDSISLG
MSEFEQNLKIIKQCRIGSIFNSSYNLPDDSLHNLGRSLIFAAGGKGQKFSSPIEEEETVGFCWDLIVSISIVNIN
RFQNFWPSFHDNLLGVVQFPLFSPIPFAEKGILGLLKICVRLLSVLRPEKLPEEVLFKSITLMWKLDKEILDTCC
NFITKSMGKILTEYPANLQTPLGWKSCLHLLSVSGRHPETYDQAVDALITFLSDGTYVSRTNYACCIDCAFGFVA
LKNSSSEKSLKILDLLSDSVNSLIQWYKEYLDSGSNFSNASSTSNSSMDESAKNFASPNFAVTLFVKLGEAFRKT
SKARREEIRNHAIFCLQRSFELAQELDFSPLNCISCFNLVIFATVDDLHEKILEYSRRENAEKELRSMEGTLKLA
MELLTDVYLLFLKPISISPGFRTFWLGVLRRMDTCMKADLGEYGETKLQEVVPDLLRKIITKMKEDEILVQKEDD
DLWDITYIQIQWIAPSLKEDLFPELEF*

**>Lus10026627**

MGRLKMQPGINSIEEEPEDLEAAYVNKTTLACMINAEVGSVLAVMRRNVRWGGRYMSGDDQLDHSFIQSLKALRK
QIFTWQHPWNTINPAVYLQPFLEVIRSDEIGAPITGVALSSVYKILTLDVIDKNTANVEEAMHLVVDAVTSCRFE
PTDPASEEVVLMKFLQVLLACMKSKASVMLSNQHVCMIVNTCFRIVHQSGNKGELLQRMARHTMHELIRCIFAHL
PDVDITERPLVNGVSTSKSEVGGPDNNHALASKQTENGQSTSVSLLSDSTAIVASPMEESSIGGGSGTDAFSYDM
HLMTEPYGVPCMVEVFHFLCSLLNAAEHMGTGPRSNTIAYEEDVPLFALGLINSAIELAGPSLCHHPRLLTLIHD
ELFRNLMQFGLSMSPLILSMVCSIVLSLYHHLRSEIKLQLEAFFSCVILRLAQSKFGSSYQQQEVAMEALVDFCR
QKTFMVEMYANLDCDITCSNVFEELANLLSKSAFPVNSPLSAMHILALDGLIAVIQGMAERIGNESENPEQTPVV
LEEYTPFWMVKCDGYSDPDHWVPFVRRRKYIKRRLMIGADHFNRDPKKGLEFLQGTHLLPEKLDPQSVACFFRYT

AGLDKNLVGDFLGNHDEFCVQVLHEFAWTFDFQDMNLDTALRLFLETFRLPGESQKIQRVLEAFSERYYEQSPQV
LANKDAALLLSYSIIMLNTDQHNVQVKKKMTEEDFIRNNRHINGGQDLPREFLGELYHSICKNEIRTTPEQGAGF
AEMNPSRWIDLMHKSKKTPPYIVSDSRAHLDRDMFAIMSGPTIAAISVVFDHTDQEEVYQACIDGFLAIARISAC
HHLEDVLDDLVVSLCKFTSLLNPSFIEEPVLAFGDDAKARMATVTVFTIANRYGDYIRTGWRNILDCILRLHKLG
LLPARVASDAADESETSAEPGHGKPPISSSMSSVQMQSAGTPRQSSGLMGRFSQLLSLDSEEPRSQPTEQQLAAH
QRTLQTIQTCHVDSIFTESKFLQADSLLQLARALIWAAGRPQKGNNSPEDEDTAVFCLELLIAITLNNRDRIGLL
WQGVYEHIANIVQSTVMPCALVEKAVFGLLRICQRLLPYKENLADELLRSLNLVLKLDARVADAYCDQITQEVSR
LVKANATHIRSQMGWRTITSLLSITARHPEASEAGFDALLFIMTGSAHLLPANYILCIDAARQFSESRVGQSERS
VRALDLMAGSVDCLTQWSKEAENPMRDNIGEMWLRLVQGLRKVCLDQREEVRNHALLSLQKCLMGVEEINLPHGL
WLQCFDMVIFTMLDDLLEIAQGHSQKDYRNMEGTLIVALKLLSKVFLQLLDELAELTTFCKLWLGVLTRMEKYMK
VKIRSRKSDKLQESVPELLKNILLVMKSKGVLVQRTALGGDSLWELTWLHMNNIAPSLQSEVFAGHELEQSRQVK
SGSDETGDSPAVGSDSAGTGG*

**>Lus10030432**

MGRLKMQPGINSIEEEPEDLEAAYANKTTLACMINAEVGSVLAVMRRNVRWGGRYMSGDDQLDHSFIQSLKALRK
PIFTWQHPWNTINPAVYLQPFLEVIRSDEIGAPITGVALSSVYKILTLDVIDKNTANVEEAMHLVVDAVTSCRFE
PTDPASEEVVLMKFLQVLLACMKSKASVMLSNQHVCMIVNTCFRIVHQSGNKGELLQRMARHTMHELIRCIFAHL
PDVDITEHPLVNGVSTSKPEVGGPDNNHALASKQTENGQSTSVSLLSDSTAIVASPMEESSIGGGSGTDAFSYDM
HLMTEPYGVPCMVEVFHFLCSLLNAAEHMGTGPRSNTIAYEEDVPLFALGLINSAIELAGPSLCHHPRLLTLIHD
ELFRNLMQFGLSMSPLILSMVCSIVLSLYHHLRSEIKLQLEAFFSCVILRLAQSKFGSSYQQQEVAMEALVDFCR
QKTFMVEMYANLDCDITCSNVFEELANLLSKSAFPVNSPLSAMHILALDGLIAVIQGMAERIGNESENPEQTPVV
LEEYTPFWMVKCDSYSDPDHWVPFVRRRKYIKRRLMIGADHFNRDPKKGLEFLQGTHLLPEKLDPQSVACFFRYT
AGLDKNLVGDFLGNHDEFCVQVLHEFAWTFDFQDMNLDTALRLFLETFRLPGESQKIQRVLEAFSERYYEQSPQV
LANKDAALLLSYSIIMLNTDQHNVQVKKKMTEEDFIRNNRHINGGQDLPREFLGELYHSICKNEIRTTPEQGAGF
AEMNPSRWIDLMHKSKKTPPYIVSDSRAHLDRDMFAIMSGPTIAAISVVFDHTDQEEVYQACIDGFLAIARISAC
HHLEDVLDDLVVSLCKFTSLLNPSFIEEPVLAFGDDAKARMATVTVFTIAPIANRYGDYIRTGWRNILDCILRLH
KLGLLPARVASDTADESETSAEPGHGKPPISSSMSSVQMQSAGTPRQSSGLMGRFSQLLSLDSEEPRSQPTEQQL
AAHQRTVQTIQTCHVDSIFTESKFLQADSLLQLAKALIWAAGRPQKGNNSPEDEDTAVFCLELLIAITLNNRDRI
GLLWQGVYEHIANIVQSTVMPCALVEKAVFGLLRICQRLLPYKENLADELLRSLNLVLKLDARVADAYCDQITQE
VSRLVKANATHIRSQMGWRTITSLLSITARHPEASEAGFDALLFIMTGSAHLLPANYILCVDAARQFSESRVGQS
ERSVRALDLMESSVDILTQWSKEAENPMRDNIGEMWLRLVQGLRKVCLDQREEVRNHALLSLQKCLTGVEEINLP
HGLWLQCFDMVIFTMLDDLLEIAQGHSQKDYRNMEGTLIVALKLLSKVFLQLLDELAELTTFCKLWLGVLTRMEK
YMKVKIRSRKSDKLQESVPELLKNILLVMKSKGVLVQRAALGGDSLWELTWLHMNNIAPSLQSEVFAGHELEQSR
QVKSGPDRTSDSPAVGSESAGTGG*

**>Lus10037140**

MQSGIKAIEEEPVAYSNKTTLACMINAEVGAVLAVMRRNVRWGSRYMSGDDQLEHSLIQSLKGLRKQIFAWQQPW
NSVNPADYLQPFLEVIRSAETGAPITGVALSSVYKILTLDVIDQNNVNVEDSMHLVVEAVTGCRFELTDPASEEV
VLMKVLQVLLACMKSKASVMLSNQDVCTVVNTCFRIVHQAGNKGELLQRMARHTMHELIRCIFSHLPDVDKTEQA
LVNGVSTVEPEIGGLDSNNAFGSKQVENGNSNTELDGQPTSLSLSSNSTAMVSSTMEDSSIGAGNGSDAFSYDQQ
LMKEPYGVSCMVEIFHFLCSLLNAVEHMGMDPRSNTIAFDEDVPLFALGLINAAIELAGPSIRYHPRLLTLIQDE
LFQNLMQFGLSMSPLILSMVCSIVLSLYHHLHSKIKLQLEAFFSCVVLRLAQSRYGASYQQQEVAMEALVDFCRQ
KTFMVEMYANLDCDITCSNVFEELANLLSKSAFPVNCPLSAMHILALDGLIAVIEGMAERIENRPGNPEQVTVSL
EEYTPFWMVKCDNYKDHTHWVPFVRRRKYIKRRLMIGADHFNRDPKKGLEFLQGTHLLPEKLDPQSVACFFRYTA
GLDKNLVGDFLGNHDEFCVQVLHEFAGTFDFEDMNLDTALRLFLETFRLPGESQKIQRVLEAFSERYYEQSPQIL
ANKDAALLLSYSIIMLNTDQHNVQVKKKMTEEDFIRNNRHINGGQDLPREFLTELYYSICKNEIRTIPDQGAGFA
EMNPSRWIDLMLKSRKTAPYIVCDSIADLDHDMFAIMSGPTIAAISVLFDHAETKETYQTCIKFTTLLNPSSIEE
PVLAFGDDVKARMATVTVFTIANRYGDYIRTGWRNIVDCVLRLHKLGLLPARVASDAADELETSTELTQGKPVTN
PVSSVQMQQAGTPRKSSGFMGRFSQLLSLDVEEPRLQPTGQELAAHQRTVETIQQCHVDSIFTESKFLQAESLSQ
LAKALIWAAGRPQKGASSPEDEDTSVFCLELLIAITLNNRDRIILLWQGVYEHIASIVQSTVMPCALVEKAVFGL
LRICQRLLPYKENFADELLRSLNLVLKLDARVADAYCEQITQEVTRLVKANATHIRSLMGWRTITSLLSITARHP
EASDAGFDALLFIMSGGAHLLPANYILCVDAARQFSESRVGQSERSLHALDLMSGSVDCLTQWSNEAKAAMGDEE
AAKLLQDIGEMWLRLVQGVRKVCLDQREEVRNHALLSLQKCLANGLDLPPSLWLQCFDTVIFTMLDDLLEIAQGH
SQKEYRNMEGTLIVAIKLMSKVFLQLLQDLAQQLTTFCKLWLGVLGRTEKYMKVKIRGKKSEKLQETIPELLKNI
LLIMKTTGVLVPSSDSLWELTWLHMNNIAPSLKSEVFPEQELEQKPAVSDETGASSS*

**>Lus10036789**

MQSRIKAIEEEPVAYSNKSTLACMINVEVGAVLAVMRRNVRWGSRYMSGDDQLEHSLIQSLKGLRKQIFSWQQPW
NSVNPADYLQPFLEVIRSAETGAPITGVALSSVYKILTLDVIDQNTVNVEDAMHLVVEAVTGCRFELTDPASEEV
VLMKVLQVLLACMKSKASVMLSNQDVCTVVNTCFRIVHQAGNKGELLQRMARYTMHELIRCIFSHLPDVDKTEQA
LVNGVSTVEPEIGGLDSNNAFGSKQVENGNSNTELDGQPTSLSLSSNSTAMVSSTMEDSSIGAGNGSDAFSYDQQ
LMKEPYGVPCMVEIFHFLCSLLNAVEHMGMDPRSNTIAFDEDVPLFALGLINAAIELAGPSIRYHPRLLTLIQDE
LFQNLMQFGLSMSPLILSMVCSIVLSLYHHLYSEIKLQLEAFFSCVVLRLAQSRYGASYQQQEVAMEALVDFCRQ
KTFMVEMYANLDCDITCSNVFEELANLLSKSAFPVNCPLSAMHILALDGLIAVIEGMAERIENEPGNPEQVPVSL
EEYTPFWEVKCDNYNDPTQWVPFVRRRKYIKRRLMIGADHFNRDPKKGLEFLQGTHLLPEKLDPQSVACFFRYTA
GLDKNLVGDFLGNHDEFCVQVLHEFACTFDFEDMNLDTALRLFLETFRLPGESQKIQRVLEAFSERYYEQSPQIL

ANKDAALLLSYSIIMLNTDQHNVQVKKKMTEEDFIRNNRHINGGQDLPREFLTELYYSICKNEIRTIPDQGAGFA
EMNPSRWIDLMLKSRKTAPYIVCDSIADLDHDMFAIMSGPTIAAISVVFDHAEHEEVYQTCINGFLAVANISACH
HLEDVLDDLVVSLCKFTTLLNPSSIEEPVLAFGDDAKARMATVTVFTIANRYGDYIRTGWRNIVDCILRLHKLGL
LPARVASDAADELETFPELTQGKPVTNPVSSVQMQQAGTPRKSSGFMGRFSQLLSLDVEEPRLQPTGQELAAHQR
TVETIQQCHVDSIFTESKFLQAESLSQLAKALIWAAGRPQKGASSPEDEDTSVFCLELLIAITLNNRDRIILLWQ
GVYEHIASIVQSTVMPCALVEKAVFGLLRICQRLLPYKENFADELLRSLNLVLKLDARVADAYCEQITQEVTRLV
KANATHIRSLMGWRTITSLLSITARHPEASDAGFDALLFIMSGGAHLLPANYILCVDAARQFAESRVGQSERSLH
ALDLMSGSVDCLTQWSNEAKAAMGDEEAAKLLQDIGEMWLRLVQGVRKVCLDQREEVRNHALLSLQKCLTNNLDL
LPGLWLQCFDTVIFTMLDDLLEIAQGHSQKEYRNMEGTLIIAIKLMSKVFLQLLQDLAQQLTTFCKLWLGVLGRT
EKYMKVKIRGKKSEKLQETIPELLKNILLIMKTTGVLVRPLEFWSRAGPACGS*

**>Lus10021771**

MEGTEADEEGNNKYTSSTNKKSSKKKRIELGISCMLNTEVASVLAVLRRPPPDPTSAQFLSQEDQYDTALVHSFK
SLRTLIFNPPQDWQTMDPAVYLTPFLDVIQSDEIPSAATSIALYSVLKLLSVGIFDDRTTGARDAINSVVMGITG
CRLERTDPVSEDAVMMRILQILTAIMKHQAAILLTDQAVCTVVNTCFQVVQQSASRGDLLQRSARYTMHELVRMI
FAKLQDIEVNPGEESDTDMDDADFDGNLDSGYGIRCAVDIFHFLCSLLNVVEVVESEGNLYRTSDEDVQLFALVL
INSAFELSGDVIGSHPKLLRMIQDDLFHHLIHYGAPSSPLILSMICSTVLNIYHFLRRFVRFQLEAFFKYVLLRV
AANGVSTQLQEVALEGIIDFCRQPTFIIEAYVNYDCDPLCINMFEETAKLLCKLSFPGASPATSLQAQAFEGLVG
IIHNIAENINREGDSTPSGPYPVHLTEYSKFWQDRPLENNDASVAYMRIRKAQKRKILIASNHYSRDEKKGLEYL
RLSLMFPNPPDPKDFAFFFRFTPGLDKTMIGDYLGDPDEFHIQVLKEFSCTFGFSGMVLDTALRTYLATFRLPGE
SQKIQRIMEAFSDQFYEQQSSDIFASKDAVFVLCYSLIMLNTDQHNPQVKKKMTEEEFIRNNRAINAGQDLPREY
LSELFQSISSKAIALFGQTGQQEMNPGLWIELMGASRYLQPFTLYDFDRRLGRDMFASVAGPAVAALTAFFEQSD
EDEMFHECVEALISVARIVQYGLEDTLDEMLASLCKCTTLLNPYASAEETLFAFSHDLKPRMAVLVVFSIANNFG
CGIRGAWRPILDCLLKLKKLKLLPSSVVELDNSFAPSIDSSSDHSRNDGGLIFPSHESRSGGGRNRQSSTMIGSF
SHFLSLDSMEESLSLGLNEFEQNQKVIKQCQIGSIFTNSWNLPDETLAYLGRTLIFASAGKGQKFATPIEEEETI
EFCWDLIVALSFANINRLHNFWPAYHENLFEAIHFPLFSSIPFAEKVVSALFRICLKLLSSARLEKLPEELIFKS
INMMWQIEKQILDTCCESITKSVSMILFEYPANLQSFLGWKTVLHLLSVSGRHQETYDQGVETLIMLISDQTLNI
LDLLAESVSLQIHWYKNFSDPGSNFSATSTGSNNFFQDATSFSVSLFLKLGEAFRKTSLARREEIRNHAIKSLQK
SFSLAEELDFTPANCLNLFNLVVFAMVDDLHEKLIEYSRREGAEREMRSMEGTLSLAMELMTEVYLQFLKPIASS
PGFRTFWLGVLRRMDTCMKADLGEYGETKLQEVIPGLLKKMIIKMKEDEILVQKEDDDLFEITYIQIQWIAPSLK
EELFPELEI*

**>Lus10034615**

MEGTEAEEEGNTKYTSSTNRKSSKKKRIELGISCMLNTEVASVLAVLRRPPPDPTAAQFLSQEDQYDTALVHSFK
SLRTLIFNPPQDWQTMDPAVYLTPFLDVIQSDEIPSAATSIALYSVLKLLSVGIFDERTTGARDAINSVVMGITG
CRLERTDPVSEDAVMMRILQILTAIMKHQAAILLTDQAVCTVVNTCFQVVQQSASRGDLLQRSARYTMHELVRMI
FAKLQDIEVNPGEESDTDMDDADFDGNLDSGYGIRCAVDIFHFLCSLLNVVEVVESEGNLYRTSDEDVQLFALVL
INSAIELSGDVVGSHPKLLRMIQDDLFHHLIHYGASSSPLILSMICSTVLNIYHFLRRFVRLQLEAFFKYVLLRV
AANGVSTQLQEVALEGIVDFCRQPTFIIEAYVNYDCDPLCINMFEETAKLLCKLSFPGASLATSLQAQAFEGLVG
IIHNIAENINREGDSTPSGPYPVHLTEYSKFWQDRPLENNDASVAYMRIRKVQKRKILVASNHYSRDEKKGLEYL
RLSLMFPNPPDPKDFAFFFRFTPGLDKTMIGDYLGDPDDFHIQVLKEFSCTFGFSGMVLDTALRTYLATFRLPGE
SQKIQRIMEAFSDQFYEQQSSDIFATKDAVFVLCYSLIMLNTDQHNPQVKKKMTEEEFIRNNRAINAGQDLPREY
LSELFQSISSKAIALFGQTGQQDMNPGLWIELMGASRYLQPFTLYDFDRRLGRDMFASVAGPAVAALTAFFEQSD
EDEMFHECVEALISVARIVQYGLEDTLDEMLASLCKCTTLLNPYASAEETLFAFSHDLKPRMAVLAVFSIANNFG
CGIRGAWRPILDCLLKLKKLKLLPSSVVEIHNSLAPSVDSSSDHSRNDGGLIFPSHESKSSGGSNRQSSTMIGRF
SQFLSPDSMEESLSLGLNEFEQNQKVIKQCQIGSIFTNSWNLSDETLAYLGRTLIFASAGKGQKFATPIEEEETI
EFCWDLIVALSFANINRLHNFWPAYHENLFEAIHFPLFSSIPFAEKVIIALFRICLKLLSSARVEKLPEELIFKS
INMMWQIEKQILDTCCESITKSVSKILFEYPANLQSFLGWKTVLHLLSVSGRHQETYDQGVETLIMLVSDQTHVS
PVNYPFCIDCAFGFVALKNSPLEKNLKILDLLAESVSLLIHWYKNFSDPGSNFSTASTGSNNSFQDAASFSVSLF
LKLGEAFRKTSLARREEIRNHAIKSLHKSFSLAEELDFTPANCLNLFNLVVFAMVDDLHEKLIEYSRREGAEREM
RSMEGTLSLAMELMTEVYLQFLKPIASSPGFRTFWLGLLRRMDTCMKADLGEYGETKLQEVIPGLLKKMIIKMKE
DEILVQKEDDDLFEIAYIQIQWIAPALKEELFPELEI*

**>Manes_LOC110625274**

MGRLKLQSGIKAIEEEPEDCDSSYSNKATLACMINAEVGAVLAVMRRNVRWGGRYMSGDDQLEHSLIQSLKALRK
QIFSWQHPWHTINPSVYLQPFLDVIRSDETGAPITGVALSSVYKILTLDVIDQNTVNVEDAMHLVVDAVTSCRFE
VTDPASEEVVLMKILQVLLACMKGKASVTLSNQHVCTIVNTCFRIVHQAGTKGELLQRIARHTMHELVRCIFSHL
PDIENTEHALINGVSPAKQEIAGLDNDYTFVSKQIENGSSELEGQTSSVSYGSSASTGLVATVMEESTIGGSSGK
DALPYDLQLMTEPYGVPCMVEIFHFLCSLLNVVEHMGMGPRSNTIAFDEDVPLFALGLINSAIELGGPSIHRHPR
LLSLIQDELFRNLMQFGLSMSPLILSMVCSIVLNLYHHLRNELKLQLEAFFSCVILRLAQSKYGASYQQQEVAME
ALVDFCRQKTFMVEMYANLDCDITCSNVFEELANLLSKSAFPVNCPLSAMHILALDGLIAVIQGMAERIGNGSFN
SEQAPVNLEEYTPFWMVKCDNYSDPNHWVPFVRRRKYIKRRLMIGADHFNRDPKKGLEFLQGTHLLPDKLDPQSV
ACFFRYTAGLDKNLVGDFLGNHDEFCVQVLHEFAGTFDFQGMNLDTALRLFLETFRLPGESQKIQRVLEAFSERY
YEQSPQILANKDAALLLSYSLIMLNTDQHNVQVKKKMTEEDFIRNNRHINGGNDLPREFLSELYHSICKNEIRTT
PEQGAGFPEMTPSRWIDLMLKSKKTAPFIVSDSKAYLDHDMFAIMSGPTIAAISVVFDHAEHEDVYQTCIDGFLA
VAKISACHHLEDVLDDLVVSLCKFTTLLNPSLLEEPVLAFGDDPKARMATVTVFTIANRYGDYIRTGWRNILDCI

LRLHKLGLLPARVASDAADESEVSADPGHGKPISNSLSSAHVQSMGTPRRSSGLMGRFSQLLSLDTEEPRSQPTE
QQLAAHQRTLQTIQKCHVDSIFTESKFLQAESLLQLARALIWAAGRPQKGNSSPEDEDTAVFCLELLIAITLNNR
DRIVLLWQGVYEHIANIVQSTVMPCALVEKAVFGLLRICQRLLPYKENLADELLRSLQLVLKLDARVADAYCEQI
TQEVSRLVKANATHIRSLMGWRTITSLLSITARHPEASEAGFDALLFIMSDGAHLVPANFVLCVDAARQFAESRV
AQSERSVRALDLMAGSVDFLTRWSHEAKETMAEEEAAKLSQDIGEMWLRVVQGLRKVCLDQREDVRNHALLSLQK
CLRGVEGINLPHGLWLQCFDLVIFTMLDDLLEIAQGHSQKDYRNMEGTLIIAVKLLSKVFLQLLHDLAQLTTFCK
LWLGVLSRMEKYLKVKVRGKKSEKLQEVVPELLKNTLLVMKAKGVLVQRSALGGDSLWELTWLHVNNIAPSLQSE
VFPDQDWEQSEHKPAETVGNLVLDETGSVPSNGSVASEGSGAGG*

**>Manes_LOC110625243**

MGRIKQQSGIKAIEEEPEDCDSSYSNKATLACMINAEVGAVLAVMRRNVRWGGRCMSGDDQLEHSLIQSLKALRK
QIFSWQHPWHTINPSVYLQPFLDVIRSDETGAPITGVALSSVYKILTLDVIDQNTVNVEDAMHLVVDAVTSCRFE
VTDPASEEVVLMKILQVLLACMKGKASVTLSNQHVCTIVNTCFRIVHQVGTKGELLQRIARHTMHELVRCIFSHL
PDIENTEHALVNGVSPAKQEIAGLDNDYTFGSKQIENGSSELEGQTSSVSYVLSASTGLVATVMEESTIGGSSGK
GALPYDLQLMTEPYGVACMVEIFHFLCSLLNVVEHMGMGPRSNTVAVDEDLPLFALGLINSAIELGGPSFHRHPR
LLSLIQDELFRNLMQFGLSISPLILSMVCSIVLNLYHHLRTELKLQLEAFFSCVILRLAQSKYGALYQQQEVAME
ALVDFCRQKTFMVEMYANLDCDITCSNVFEDLANLLSKSAFPVNCPLSTMHVLALDGLIAIIQGMAERLGNGSFN
SDEPPVNLEEYTPFWMVKCDNYSDPDHWVTFVRRRKYIKRRLMIGADHFNRDPKKGLEFLQGTHLLPDKLDPQSV
ACFFRYTAGLDKNLVGDFLGNHDEFCVQVLHEFAGTFDFQGMNLDTALRLFLETFRLPGESQKIQRVLEAFSERY
YEQSPQILANKDAALLLSYSLIMLNTDQHNVQVKKKMTEEDFIRNNRHINGGNDLPREFLSELYHSICKNEICTT
PEQGTGFPEMIPSRWIDLMLKSKKTAPFIVSDSKAYLDHDMFAIMSGPTIAAISVVFDHAETVDVYQTCIDGFLA
VAKISACHQLEDVFDDLVVSLCKFTTLLNPSSAEEPVLAFGDDPKARMATVTVFTIANRYGDYIRAGWRNILDCI
LRLHKLGLLPARVASDAADESEVSSDPGHGKPITNSLSSAHTQSFGIPRRSSGLMGRFSQLLSLDTEEPRSQPTE
QQLAAHQHMLQTIQKCHVDSIFTESKFLQAESLLQLARALIWAAGHPQKGNNTPEDEDTAVFCLELLIAITLNNR
DRIALLWQGVYEHIANIVQSTVMPCALAEKAVFGLLRICQRLLPYKENLADELLRSLQLVLKLDARVADAYCEQI
TQEVSRLVKVNATHIRSLMGWRTITSLLSITARHPEASEAGFDALLFIMSDGAHLVAANYVLCIEAARQFAESRV
AQAERSMRSLDLMAGSIDFLTRWSHEAKEAMEEEEAAKLLQDIGEMWLRLVQGLRKVCLDHREDVRNHALLSLQK
CLRGVDGINLPHGFWLRCFDLVIFTMLDDLLEIAQGHSQKDYRNMDGSLIIAVKVLSKVFLQLLPDLAQLTTFCK
LWLGVLSRMEKYLKVKVRGKKCEKLQEVVPELLKNTLLVMKAKGVLVQRSALGGDSLWELTWLHVNNIAPSLQSE
VFPDRDWERTQHEHGETGESLVATESGSLTLNGPVASEDFGAGG*

**>Manes_LOC110609669**

MRHLYLESGTNSFHGEFSDCPVKAIKGALACMVNSEIGAVLAVMRRNVRWGIRYVADDDQLEHSLIHSLKELRKQ
IFSWQHKWHNIDPTIYLQPFLDVIKSDETGAPITGVALSSVYKILTLDVLDVYTVNVAKAMCLIVDAVTTCRFEV
TDPASEEVVLMKILQVLLACMKSKASVKLSNQHVCNIVNTCFRVVHQASSKGELLQRIARHTMHELVRCIFSHLP
DMDKNEDAVANGSSIVSREVDTLEKDNAFGSKQLENGNSTAQSNSQPSILDTPSNLVDMIESGKDDSKIEIGNGM
EAFDNGEKLNLEPFGIPCMVEIFHFLCSLLNVIKNIEVDLRSNTLAYDEDVPLFALGLINLAIELGGPSFSKHPA
LLSLIQGELFRNLMQFGLSMSPLILSTVCSIVLNLYHHLRTELKVQFEAFFSCVLLRVAESKHGSLYQLQEVAME
ALVDLCRQQTFMAEMYANFDCDITCINVFEDLANLLSRSAFPVNGPLSSMHIVALDGLVSMVKCMADRIGNEMSL
PEEASADLEGYKEFWTVKCENYSDPNFWIPHIRKMRHIKRKLMVGVDHFNRDPKKGLEFLQGMHPLPEKLNPQCV
ASLFRYTVGLDKNLIGDFLGNHDEFCIQVLQEFAGTFDFRGMNLDTALRIFLGTFRLPGESQKIQRVLEAFAERY
YEHSPQILANKDAALLLSYSLILLNTDQHNAQVKKKMIEEDFIRNNRCTNGGKDFPQEYLSELYRSICENEIQMI
PEQRAGVPLMTSGCWINVLHKSKITSPFIVSDSRELLDYDMFTILSGPAIAAMSVVFDHTENEEVLKICVDGFLA
VAKISSSYHFDNVLDDLVVSLCKFTDQLTPLSVDDAILAFEEDIKARMATTTVFTIANKYGDYIRAGWENILDRV
LSFHKLGLLPARLINDATDDMELSSDMERAKPAPSSSLPSQAPSGTNPRKSSGLMGRFSQLLFFDVEELRSLPIE
EQLAAHQLSSETIKSCHIDSIFIESKFLQAVSLLQLVKSLILVASRLSKGTSPVEDEGSAVFCLELLITITLNNR
DRIMLIWKEVYDHISNIVQSTKMPCTLVEKAVFGLLRICQRLLPYKENLTDELLKSLQLILKLDARVADAYYEQI
TQEVIHLVKANASHIKSHLGWRTITSLLSITSRHPEASQTGFETLAFIMSEGAYLLPSNYVLCVDAARQFAESRL
GDVDQSVSALDMMAGSIACLARWSSEAEITMGQEADTKVSQDIGEMWLRLVQGLRKVCLDHREEVRNHAILMLQR
CMAGVDEIHLPMALWFQCFDLVIFSLVDDLLDIAQQSSPKNYRKMEETLALAMKLMVKAFLQRLQDLSQQPSFCR
LWLGVLDHMERFVQVKFQGKHSEEIYELVLEFLKNILLVMKTTGVLVPSSDIGKDSFWQLTWLQVKNICPTLQSE
VFSDEELQQL*

**>Manes_LOC110610972**

MEKIVEEEEEHSISSWRDKKRRKEVSLSCMLNTELSAVLAVIRRPQDSSGSMIPQEDTFDVGILHSLKSLRALIF
NPQQEWRTVDPSVYISPFLDVIQSDDIPAMATNVALAAIAKILKLHIFDEKTPGAKDAINSVVMGITSCRLERTD
TVTEDAIMMRVLQALTSIIKHRASVLLTDQAICTIVNTCFQVVQKSTNRADLLQRGARFAMREMIEIIFARLQNV
EVRIGEDSESDTEDIDIDSTMDSGYGIRCVIDIFHFLCSLLNVVEVVEIDGVPSHAVDENVQIFGLVLINSAIEL
SGDVIGKHPKLLRMIQDDLFHHLIHYGTSASPLMLSMICSTVLNFYHSLRRFLRVQLEAFFEFVLLKAAAAASSY
QLQEVALEGIINFCRQPCFIIEMYVNYDCDPTCRNIFEGIGKLLCKLSFPGSSPLSCMQVQAFEGLLIIIHNIAD
HVDREDDSSPSGPYPVEITEYKPFWEEKPREYSDSWVEYLRFRKAQKKKIMIAGDHFSRDEKKGLEYLKLCQLIS
DPPDPKAFAIFFRHTPGLDKNSIGDYLGDPDDFHMQVLKDFTETFMFSGMILDTALRTYLATFRLPGESQKIQRI
LEAFSERFYDQQSSDTFASKDAVFILCYSLIMLNTDQHNPQVKKKMTEEEFIRNNRAINGGQDLPREYLTELFQS
IATNAITLFGQSGPVEMNPGTWVELMNRSKFIQPFILCDYDRRLGRDMFACVAGPSVAALSSFFEHTDDDEMLRE
CIGGLISVARVAQYDLEDILDELLASFSKFTTLLNPYASAEETLFAFSNDMKPRMATLAVFTIANNFGDSIRGGW
RNIVDCLLKLKKLKLLPQSVIEFDETAASSSDGLRNKRNESGISSSCDSRVSTRRSAGMISRFSHFLSLESMEES

ISLGMGEFEQNLKVIKQCRIGGIFNNSCNMPDDALLNLGRSLIFAAGGKGQKFSNPVEEEETVGFCWDLIIAICL
VNINRFHNIWPSFHENLLGVAQFPLFSPIPFAEKAILGLFKVCIRLLSSPRAEKLAEEALFKSINLMWKLDKEIL
DTCCEFIIKSISKIIIEYPANLQTNLGWKSCLHLLSVTGRHAETYEQGVYTLITLMSDGTYVSRINYAYCVDCAF
GFVALKNSPLEKNLKILDLLSDSVKLLIQWYKEYSDSGSNFSAASSASNSSMDDSSKGFASPNFAMNFFIKLGEA
FRKTSLARREEIRNHTILALQKSFALADELDFSPLNCINCFNLVIFAMVDDLHEKMVEYSRRDGAEKEMRSMEGT
LKLAMELLASVYLQFLKPITMSPGFRTFWLGVLRRMDTCMKADLGDYGETKLQEVIPDLLRKIITKMKEEEILVP
KEDDDLWDITYIQIQWIAPSLKEELFPDLEL*
**>Clach_001936**
MGRMRLHTDIRAIEEEPEEIDSSQSEKTSLACMINAEVGAVLAVMRRNVRWGGRYMSGDDQLEHALIQALKTLRR
QIFSWQQSWQSINPIAYLQPFLDVIISDETGAAITGIALSSLHKIITLDFIDQNTENVEESMQQIVDSVTSCRFE
VTDSSSEEIVLMKILQVLLACMKSKASSMLSNQNVCTIVNTCFRIVHQAETKGELLQRISRHTMHELVRCIFLHL
PRFHTTDRAIINGTSNPLENGTKKTDSESEKSIGMMEANTIGIFNTNGLPIDIHLMNEPYGVPCMVEIFHFLCSL
LNVVENAGMDPKSNTVSFDEDVPLFALGLINSAIDLGGSSIRRHPRLLSLVQDELFCSLMQFGLSSSALILSTVC
SIVLNLYNYLRSELKLQLAAFFSCVIIRFAQSRYNATYQQQEVAMEMLVDFCRQKSFMVEMYANLDCDITCGNVF
QELANLLSKNAFPVNCPLSAMHILSLDGLIAVLIGMAERVGNASTICDSSPVGIDEYTPFWMVKCDDYSDPNHWV
PFVRKRKYIKRRLMIGADHFNRDPKKGLEFLQGTHLLPDKLDPQSVACFFRYTAGLDKNLVGDFLGNHDEFCVKV
LHEFAGTFDFQDMNLDTALRLFLETFRLPGESQKIQRVLEAFSERYYEQTPHILANKDAALVLSYSLIMLNTDQH
NVQVKKKMTEEDFIRNNRHINGGNDLPRDYLSELYHSICKNEIRTTPEQGIMYAEMTPSRWIDLMSKSKKTAPFV
MPDSKPYLDHDMFAIMSGPTIAAISVVFDHAEQEDVYQTCIDGFFAVAKISACHHLEDVLDDLVVSLCKFTTLLN
SSSVEEPVLAFGDDTKARMATVTIFTIANRYGDFIRTGWRNILDCILRLHKLGLLPARVASDAADETEASSDIGQ
GKSVSNPLTSIPIQSIGTPRRSSGLMGRFSQLLSLDTEEPRSQPTERQLAAHQRTLQTIQKCHVDTIFTESKFLQ
ADSLLQLARALIWAAGRPQKGGTSPEDEDTAVFCLELLIAITLNNRDRIALLWQGVYEHIANIVQTTVMPCALVE
KAVSGLLRICHRLLPYKESLADDFLRFLQLVLKLDARVADAYSEQITHEISRLIKANASHIRSQMGWRTIASLLS
FTARHPEASEVGFDALSFVMNDGAYLVPVNYIYCVDSARHFAESRVGITDRSSRALDLMASSVESLVKWARETRE
AMTEEEAAKMSMSIGEMWLRVAQGLRKVCLDQREDIRNHALSALEKCLMSEERIKVHNNLLLQCFDMIIFTMLDD
LVEISQGNSQKDYKNMEATLTRALTVVTGFYLQVVKGLVEATTFSKVWLGVVSRMEKYMKVKVRGKRSEKLMEAV
PELVKRLLLAMKKMRVLGPRIGLSGDGPWDLTCVHVNNIAPHLISEIFPEFKPPQTDTAPSLTVL*
**>Clach_019647**
MGRMRLHTDIRAIEEEPEELDSSQSEKTSLACMINAEVGAVLAVMRRNVRWGGRYMSGDDQLEHALIQSLKTLRR
QIFSWQQSWQSINPIAYLQPFLDVIISDETGAAITGIALSSLHKIITLDFIDQNTENVEESMQQIVDSVTSCRFE
VTDSSSEEIVLMKILQVLLACMKSKASSMLSNQNVCTIVNTCFRIVHQAETKGELLQRISRHTMHELVRCIFLHL
PRVHTTDRAIINGTSNPLENGTKKTDSESEKSIGMMEANTIGIFNTNGLPIDIHLMNEPYGVPCMVEIFHFLCSL
LNVVENAGMDPKSNTVSFDEDVPLFALGLINSAIDLGGSSIRRHPRLLSLVQDELFCSLMQFGLSSSALILSTVC
SIVLNLYNYLRSELKLQLAAFFSCVIIRFAQSRYNATYQQQEVAMETLVDFCRQKSFMVEMYANLDCDITCGNIF
QELANLLSKNAFPVNCPLSAMHILSLDGLIAVLIGMAERVGNASTICDSSPVGIDEYTPFWMVKCDDYSDPNHWV
PFVRKRKYIKRRLMIGADHFNRDPKKGLEFLQGTHLLPDKLDPQSVACFFRYTAGLDKNLVGDFLGNHDEFCVKV
LHEFAGTFDFQDMNLDTALRLFLETFRLPGESQKIQRVLEAFSERYYEQTPHILANKDAALVLSYSLIMLNTDQH
NVQVKKKMTEEDFIRNNRHINGGNDLPRDYLSELYHSICKNEIRTTPEQGIMYAEMTPSRWIDLMSKSKKTAAFV
MPDSKPYLDHDMFAIMSGPTIAAISVVFDHAEHEDVYQTCIDGFFAVAKISACHHLEDVLDDLVVSLCKFTTLLN
SSSVEEPVLAFGDDTKARMATVTIFTIANRYGDFIRNGWRNILDCILRLHKLGLLPARVASDAADETEASSDIGQ
GKSVSNPLTSIPIQSIGTPRRSSGLMGRFSQLLSLDTEEPRSQPTERQLAAHQRTLQTIQKCHVDTIFTESKFLQ
ADSLLQLARALIWAAGRPQKGGTSPEDEDTAVFCLELLIAITLNNRDRIALLWQGVYEHIANIVQTTVMPCALVE
KAVSGLLRICHRLLPYKESLADDFLRFLQLVLKLDARVADAYSEQITHEISRLIKANASHIRSQMGWRTIASLLS
FTARHPEASEIGFDALSFVMNDGAYLVPVNYIYCVDSARHFAESRVGLTDPSSRALDLMAASVESLVKWARETRE
AMTEEEAAKMSMSIGEMWLRVAQGLRKVCLDQREDIRNHALSALEKCLMSEERIKVHNNLLLQCFDMIIFTMLDD
LVEISQGNSQKDYKNMEATLTGALTVVTGFYLQVVKGLVEATTFSKVWLGVVSRMEKYMKVKVRGKRSEKLMEAV
PELVKRLLLAMKKMRVLGPRSGLSGDGPWDLTCVHVNNIAPHLISEIFPEFKPPQTDTAPSLTVL*
**>Clach_003699**
MGHATVKTKINPYHYDLNSHAGRFVKGSFACMINSEIGAVLAVMRRNVRWGVQYVSDVDKLEHNLIHSLKELRKK
IFSWHNDWEKMDPSFFLQPFLDVIRSDETGAPITGVALSSVYKILTLDVFDTDSAPVEEAMHMIVDAVTTCRFEV
TDSASEEVVLMKILQVLLACMKNKASVRLSNQHVCNMVNTCFRVVHQASSKGELLQRIARHSMHELVRCIFSRLP
SWNSHQTTMLYVSQINGNLEVPSENGTYVGPPNEEMSPSTSNTKNTSSENNVDGNTKKERNIMLEPFGVPCMVEI
FHFLCSLLNMAETFHSTNSKSNPVAYDEDVPLFALGLINSAIELGGASLSNHPDLLSLIQGELFRNLLQFGLSLS
PLILSTVCSIVLNLYHTLRRHLKVQLESFFSCVLLRVAQGKNNASYQQQEVAMEALVDLCRQPCFMAEMYANFDC
DTPCNNIFEEVVNLLSKCAFTVNGPLTAMHIVALDGLLSMVHDIADRTENENPSFEDNSADFDDLKDFWEMKCSD
YDDPAVWVPHIRKLWHIKRRLMVCADHFNSNPKKGLQVFQEMNLVSNHRDPNSVANLFRYTFGLDKILIGDFLGN
HDEFSIKVLRAFAETFNFPGVTLDTALRMFLGTFKLPGESQKIQRILEAFAERYYDQSPDTLASKDATLLLSYSI
IMLNTDHHNAQVKKKMTEEDFIRNNRLINDGKDLPREYLSELYHSICENEIQINPDQGSISSPMTLGRWINVMHK
ANIASPYISCQSEEIIDCNILPILLGPTIAALSVVFDQAEHEEVLQSCVDGFSSIAKIAAFYHYDKTLDDLVVSL
CKFSSFPSGIPHDAIFVFGSDIKARMATIAAFSISNHYGDYLRLGWKNLLDCTLSLHKLGLLPRFMDPDGSDPVS
VSASEHAKPPSVPPMSKKPSGLMGRFTQLLSFDSEEAVTIPTEEELAAHQTARQAIQRCNIHSVFQESKFLLADS
LTSLVSSILDAARRAEDGTAGVFCLELLIVVTLNNRDRIMLIWQNVYDHIARVVQSTNTPCALVEKAVFGLLKIC
QRLLPYNEKLSEDLLKSLQLILKLDARVADAYCDPITQEVMRLVKVNALHITSPIGWRSIISLLSITARHVEASE

AGFETLKFIMSDGAYLFPSNFLSCVDAATQFAESRLGEDTRSLTAVDLIAGSVPSVRKWSATQETEEMWLRLVQA
MKKACLDDREEVRNHAILMLQRVISGVDGVQVGDRFWCGCFDSVIFPILDRFLEVGLESSLKEFIRLEKSLVLGT
KLMTKAFLQLLQDLSLHSSFCKLWIGVLNRMESYMKFKHRGKRSEKIHELIPEVLKNILLVMKTTGVLTKSDDSF
WHLTWLHVNNICPSLQPEVYPPEDS*
**>Clach_025206**
MLELIACKGRREALISNMFNTELAAVLAVIRRPSESNTPSSAQDNADTTLLLFSLKSLRPIVFNPQNEWHSIDPA
IYLSPFLDVIQHDDVPAAATGASLSSVKRILDHRIFDDKTPGARDAVNSLVLTGIPSCRLEKTDPVTQDAIMMKI
LLVLESIMKHPASVHLGDSAVCTLINACFQIMQHSSTGGDLLQRMAKNTMLEIIRVVFSKLPDINVNEMARNDES
GYGIACMVDIFQFLRSLLNASELNENDRTSNQDDLQLFALEMLNSIIELSGDAIKNHAKLLKILVNDLFYNLIHY
GARCRPLALAIISNIALNVYKFLRKFVRLQLEAFFEYVLLKAASSGSSIPLQEVALESIINFCRQPTFILEAYVN
YDCEPTCPNIFESIGKLLCKLSYYGAIPFHSLQAQAYEGMEVLFHTIATNIDKEDDPTTSGPYPIEIKEYRPFWV
EKQREDHEAWVEHIRVRRQQKKAIVVAGNHYNRDERKGLEYLRIRQLVPEPPEAKAVAMFFRFTPGLDKTFIGEY
LGDPEEFHLDVLREFTNTFEFSGMVLDTAFRTYLASFRLPGESQKIQRIMEAFSEMYFEHCDLFVTKDALLVLCY
SLIMLNTDQHNPQVRKKMTEEEFIKNNRQINGGKDLPREYLSQLFHSIASNGITFFGQSATLVEMNSSKWIELVN
RSESMLPFVVCEFDRRLGRDMFASIAGPAIASLSASFEQSEDDDVLHACAEGVMSMTRITQYGLEDILDELVAIF
SKYTTLLNSYSTPDEILFGFSNDTKPKLATIVIFTISNNFGSSIRDGWGSIVECLLKLKRLKLLPSSVNELNDEP
ASFDSSSFASLESREDMVALGMGEMEQNMKVIKQCRIGNIFTSSWKMCEESLISLSRSLVFAAGGKGQKVLTPME
EEDNITFCWDLIVVTTSSNIHRFQSFWPTIHECLLQVLQLPLFSPVPFAEKAIVRLFEICLKLLSYPNTLKASEE
FIFKSINLMWKIDKEILDMSFDVIIGYVIKILVDYSSSIKTPLGWKLTLQLLSITGRHPETYDRGVETLIKLMST
GTHLSTMNYIYCIDCAFGFVALRNSPLETNLKILDLLAGSAELLVKWYRELGDAGFNYSGASNLSTSSSDESNKV
PNLAITFLLKLGEAFRKTSLARRQEIRNHAIVALKRSFMMIQEMDFTYGNCFNCFNLIVFAMVDDLHEKMVEYSR
RENAEKEMRSMEETLKLAIELQTDLFLVCLKPLASNSGFRTFWLGVLRRMDTCMKADLGMYGQTKLQEVVPGLLK
KMIVQMQEDNVLVPTENDDLWEITYIQIQWIAPSLKRVLFPEIDL*
**>Ricco_LOC8281853**
MGRLKLQPGIKSIEEEPEECDSSYSNKATLACMINAEVGAVLAVMRRNVRWGGRYMSGDDQLEHSLVQSLKSLRK
QIFSWQHPWHTINPAVYLQPFLDVIRSDETGAPITGVALSSVYKILTLDVIDQNTVNVEDAMHLVVDAVTSCRFE
VTDPASEEVVLMKILQVLLSCMKSKASVTLSNQHVCTIVNTCFRIVHQAGSKGELLQRIARHTMHELVRCIFSHL
PDVDNTEHALVNGVSTVKQEIGGMDNDYTFVNKQSENGNSSSELDGQTSSVSFGSSVSTGLVPTVTEENTIGGSG
KDALPYDLHLMTEPYGVPCMVEIFHFLCSLLNVVEHMGMGPRSNTIAFDEDVPLFALGLINSAVELGGPSIRHHP
RLLSLIQDELFRNLMQFGLSMSPLILSMVCSIVLNLYHHLSTELKLQLEAFFACVILRLAQSRYGASYQQQEVAM
EALVDFCRQKTFMVEMYANLDCDITCSNVFEDLANLLSKSAFPVNCPLSAMHILALDGLIAVIQGMAERIGNGSV
SSEQAPVNLEEYIPFWMVKCDNYGDPDHWVPFVRRRKYIKRRLMIGADHFNRDPKKGLEFLQGTHLLPDKLDPQS
VACFFRYTAGLDKNLVGDFLGNHDEFCVQVLHEFAGTFDFQGMNLDTALRLFLETFRLPGESQKIQRVLEAFSER
YYEQSPQILANKDAALLLSYSLIMLNTDQHNVQVKKKMTEEDFIRNNRHINGGNDLPREFLSELYHSICRNEIRT
TPEQGAGFPEMTPSRWIDLMLKSKKTAPFIVSDSRAYLDHDMFAIMSGPTIAAISVVFDHAEHEDVYQTCIDGFL
AVAKISACHHLEDVLDDLVVSLCKFTTLLNPSSVEEPVLAFGDDTKARMATVTVFTIANRYGDYIRTGWRNILDC
ILRLHKLGLLPARVASDAADESELSTEPGQGKPITNSLSSVHMQSMGTPRRSSGLMGRFSQLLSLDTEEPRSQPT
EQQLAAHQRTLQTIQKCHVDSIFTESKFLQAESLLQLARALIWAAGRPQKGNSSPEDEDTAVFCLELLIAITLNN
RDRIVLLWQGVYEHIANIVQSTVMPCALVEKAVFGLLRICQRLLPYKENLADELLRSLQLVLKLDARVADAYCEQ
ITQEVSRLVKANATHIRSLMGWRTITSLLSITARHPEASEAGFDALLYIMSDGAHLMPANYVLCVDAARQFAESR
VAQAERSVRALDLMAGSVDCLARWSHEAKEAMGEEEAAKLLQDIGEMWLRLVQGLRKVCLDQREEVRNHALLSLQ
KCLTVVDGINLPHGLWLQCFDLVIFTMLDDLLEIAQGHSQKDFRNMDGTLIIAVKLLSRVFLQLLHDLAQLTTFC
KLWLGVLSRMEKYLKVKVRGKKSEKLQEVVPELLKNTLLAMKAKGVLVQRSALGGDSLWELTWLHVNNIAPSLQS
EVFPDQDWEQSQHKQGETIGSLASDGTGSVPSNGSVASEGSGMGG*
**>Ricco_LOC8283443**
MGHLNLQSEINSFQREFCDCPVISTKGAVACMVNSEIGAVLAVMRRNVRWGVRYVTDDDQLEHTLIHSLKELRKQ
IFSWQHKWHSIDPAIYLQPFLDVICSDETGAPITGVALSSVYKILTLDLLDVNTVNVAEAMHLIVDAVTTCRFEV
TDPASEEVVLMKILQVLLACMKSKASVKLSNQHVCNIVNTCFRVVHQASSKGELLQRIARHTMHELVRCIFSHLH
DIENNEDKLTSGSSSIDREVDTLVKDKTSGSKQPENGEIGVEGDGQLSIGDAPGVRMGKRESGKDENKIEVSNGM
ESAENGEKLMMEPFGVPCMVEIFHFLCSLLNVVEHIEVGPRSNPIAYDEDVPLFALGLINSAIELGGPSFRKHPA
LLCLIQDELFRNLMQFGLSMSPLILSTVCSIVLNLYHHLRIELKVQFESFFSCVLLRIAQSKHGSSYQLQEVAME
ALVDLCRQQAFMAEMYANFDCDITCSNLFEDLANLLSKSAFPVNGPLSAMHVVALDGLISMIKCMADRMGNELSL
SEETSVDLEGHNSFWTMKSESNTDPNYWIPHVRKMRSIKRTLMIGVDHFNRDPKKGLEFLQGMHLLPEKLQPQSV
ASFFRYTAGLDKSLIGDYLGNHDDFCIQVLQEFAGTFDFRGMSLDTALRLFLGTFRLPGESQKIQRVLEAFAERY
YEQSPQVLADKDAALVLSYSLILLNTDQHNVQVKKKMTEEDFIRNNRRTNGGKDFPREYLSDLYRSICENEIQMI
PEQGAGLPLMTSGRWINVLHKSKITSPFIFCGSRALLDYDMFIILSGPTIAAMSVVFYQTEHEEVLNSCVDGFLA
IAKFSASYHLDEVLDDLVVSLCKFTTHMTSLSVDDAILTFGDDTKARMATTTVFTIANRYGDYIRSSWKNILDCV
LSFHRLGLLPAQLASDAADDIELSSDLERVKPSPVSSLSHTPSGTTPRKSSGGLMGRFSQLLSFDMEEPRSLPTE
EQIAAHQLTRETIHSCHIDSIFTESKFLQAESLLQLVRSLILAASRLGKGTSPMEDEGAAAFCLELMIAITLNNR
DRIMLIWQDVYEHISNVVQSTIMPCTLVERAVFGLLKICQRLLPYKENLSDELLKSLQLILKLDARVADAYCEQI
TQEVMRLVKANASHIRSHVGWRTITSLLSITARHPEASETGFETLTFIMSNGAYLLPSNYILCVDAARQFAESRL
GDVDRSVSALNMMAGSVVCLTRWSSEAKIAVGQEAAMKVSQDIGEMWLRLVQGMRKVCLDHREEVRNHAILMLQR
SMAGVDGIHLPNALWFQCFDLVIFTLLDDLLDISLESSPKNYRKMEETLVLAMKLMTKAYLQQLHDLSQQPSFCR

LWLGVLNRMERYMKVKFRGKHSEKIYELVPELLKNILFVMKTTGVLLPSDDIGGDSFWQLTWLHVKNICPSLQSE
VFPDHELEQIHAEQNVPAQSSESTV*
**>Ricco_LOC107261358**
MEKIEEEEEDKSWQERKRRKEVSLSCMLNTELSAVLAVIRRPHDPTNSVFHQEEGYDTSILHSLKSLRDLIFNPQ
QEWRTIDPSVYISPFLDVIQSDGIPATATNVALSAISKILKLQFFDEKTPGAKEAVNSIVTGITSCRLERTDPST
EDAVMMRILQALTSIIKHRTSVLLTDHAVCTIVNTCFQVVQQSTHRADLLQRGAKFAMREMIEIIFARLQDFEVK
SGEESESDTEDIDIGSNMDSGYGVRCVVDIFHFLCSLLNVVDIVESEGFSSQASDQNIQIFGLVLINSAVELSGD
TIGKQPKLLRMIQDDLFHHLIHYGISSSTLVLSMICSTVLNIYHSLRSFIRVQLEAFLGFVLLRTAGAGSPSQLQ
EVALEAIINFCRQPSFIVEMYVNYDCDPICRNIFEEIGKLLCKLSFPGSSPLSYVQIQAFEGLLIIIHNIADNID
KDDDSSPSGPYPVKITEYIPFWEEKPKEDFETWVEYLRLRKAQKRKVLIAGDHFNRDEKKGLEYLRLCQLVSDPA
DPKAFAIFFRFTPGLDKSMIGDYLGDPDEFHMLVLKEFTETFRFSGMILDNALRTYLATFRLPGESQKIQRILEA
FSERFYDQQSSDIFASKDAVFILCYSLIMLNTDQHNPQVKKKMTEEEFIRNNRAINGGQDLPRDYLSELFQSIAA
HAITLFGQSGPVEMNPGSWIELMNRSRVMQPFILGDYDRRIGRDMFACIAGPSIAALSSFFEHADEDEMLHECIG
GLVSVARITQYELEDILDELLASFSKFTTLLNPYASAEETLFAFSNDLKPRMATLAVFTIANNFGDSIRGGWRNI
VDCLLKLKRLKLLPQSVVEFDDTSASSSDVPGHKRNESSISLSHDPKFGNRRSAGMMNRFSPFLTIESMEDSISL
GMSEFEQNLKVIKQCRIGSIFTNSINLPDDGLLNLGRSLIFAAGGKGQKFSTPIEEEETVGFAWDLIVAVSMVNM
HRFLNFWPSFHDNLLGVAQFPLFSPVPFAEKAILGLFKICVKLLSSNRTERLPEELIFKSINLMWKLDKEILDTC
CESITKSVSKILTDYPANLQTSLGWKTCLHLLSVTGRHPETYDQGVDTLIQMVSDGTHVSRMNYAYCIDCAFGYI
ALKNSPLEKNLKILELLADSVNLLIQWYKEYADTGSNYSITSSTSNSSFEDSKGLGSPNFAITLFVKLGEAFRKT
SLARREEIRNQAILSLQKSFSLSKELDFSPLNCISCFNLVIFAMADDLHEKMIEYSRRENAEREMRSMEGTLKLA
MELLTDVYLKFLKPITMSPGFRTFWLGVLRRMDTCMKADLGEYGETRLQEVIPDLLRRIITKMKEEEILVPTEDD
DLWDITYIQIQWIAPFLKEELFPEEEI*
**>Hevbr_LOC110645704**
MGRLKLQSGIKAIEEEPEECDSSYANKATLACMINAEVGAVLAVMRRNVRWGGRYISGDDQLEHSLIQSLKALRK
QIFSWQHPWNTINPAVYLQPFLDVIRSDETGAPITGVALSSVYKILTLDVIDQNTVNVEDAMHLVVDAVTSCRFE
VTDPASEEVVLMKILQVLLACMKSKASVTLSNQHVCTIVNTCFRIVHQAGTKGELLQRIARHTMHELVRCIFSHL
PDVDDTEHALVNGVSPAKQEIAGLDNDYTFGSKQIENGSSSELDGQTSSVSYGSSASTGLVATVMEESTIGGSSG
KDALPYDLHLMTEPYGVPCMVEIFHFLCSLLNVVEHMGMGPRSNTIAFDEDVPLFALGLINSAIELGGPSIRRHP
RLLSLIQDELFRNLMQFGLSMSPLILSMVCSIVLNLYHHLRTELKLQLEAFFSCVILRLAQSKYGALYQQQEVAM
EALVDFCRQKTFMVEMYANLDCDITCSNVFEDLANLLSKNAFPVNCPLSAMHILALDGLIAVIQGMAERIGNGSV
NSEQAPVNLEEYTPFWMVKCDNYSDPNHWVPFVRRRKYIKRRLMIGADHFNRDPKKGLEFLQGTHLLPDKLDPQS
VACFFRYTAGLDKNLVGDFLGNHDEFCVQVLHEFAGTFDFQGMNLDTALRLFLETFRLPGESQKIQRVLEAFSER
YYEQSPQILANKDAALLLSYSLIMLNTDQHNVQVKKKMTEEDFIRNNRHINGGNDLPREFLCELYHSICKNEIRT
TPEQGAGFPEMTPSRWIDLMHKSQKTAPFIVSDSRAYLDHDMFAIMSGPTIAAISVVFDHAEHEDVYQTCIDGFL
AVAKISACHHLEDVLDDLVVSLCKFTTLLNPSSVEEPVLAFGDDTKARMATVTVFTIANRYGDYIRTGWRNILDC
ILRLHKLGLLPARVASDAADESEVSADPGHGKPITNSLSSAHMQSMGTPRRSSGLMGRFSQLLSLDTEEPRSQPT
EQQLAAHQRTLQTIQKCHVDSIFTESKFLQAESLLQLARALIWAAGRPQKGNSSPEDEDTAVFCLELLIAITLNN
RDRIVLLWQGVYEHIANIVQSTVMPCALVEKAVFGLLRICQRLLPYKENLADELLRSLQLVLKLDARVADAYCEQ
ITQEVSRLVKANATHIRSLMGWRTITSLLSITARHPEASEAGFEALLFIMSDGAHLLPANYVLCVDAARQFAESR
VAQAERSVRALDLMAGSVDCLTRWSHEAKEAMEEEEAAKLSQDIGEMWLRLAQGLRKVCLDHREDVRNHALLSLQ
KCLTGVDGINLPHGLWLQCFDLVIFTMLDDLLEIAQGHSQKDYRNMDGTLIIAVKLLSKVFLQLLHELAQLTTFC
KLWLGVLSRMEKYLKVKVKGKKSEKLQEVVPELLKNTLLVMKTKGVLVQRSALGGDSLWELTWLHVNNIAPSLQS
EVFPDQHWEQSQHKQGETGGSLVSDETGAVPSNGSLPSEGSGTGG*
**>Hevbr_LOC110663786**
MGHLNLESGANSFHREFNDCPVKATKGALACMVNSEIGAVLAVMRRNVRWGVRYAADDDQLEHSLIHSLKELRKQ
IFSWQHKWHNIDPAIYLQPFLDVIQSDETGAPITGVALSSVYKILTLDVLDVDTVNVAEAMHLIVHAVTTCRFEV
TDPASEEVVLMKILQVLLACMKSKASVKLSNQHVCNIVNTCFRVVHQASSKGELLQRIARHTMHELVRCIFSHLH
EMDKYEDTLVNRSSSVFREVDAMDKDNTLGSKQLENGNVGVESDGQPSIVDTPDNHMGKLESGKDDNKIDVSNGM
EAADNGEKLMMEPFGVPCLVEIFHFLCSLLNVIEHIEVGPRSNPIAYDEDVPLFALGLINSAIELGGPSFSKHPS
LLSLIQGELFRNLMQFGLSMSPLILSTVCSIVLNLYHHLRTELKVQFEAFFSCVLLRIAQSKHGSSYQLQEVAME
ALVDLCRQQTFMAEMYANFDCDITCSNVFEDLANLLSKSAFPVNGPLSAMHIVALDGLVSMIKGMADRIGSELSL
SEEASVDREGYKAFWTTKCENYSDPNFWIPHVRKMRHIKRKLMVGVDHFNRDPKKGLEFLQGMHLLPEKLDPQSV
ASLFRYTAGLDKNLIGDFLGNHDEFCVQVLQEFAGTFDFRGMNLDTALRLFLGTFRLPGESQKIQRVLEAFAERY
YEESPQILANKDAALLLSYSLILLNTDQHNVQVKKKMTEEDFIRNNRRTNGGNDFPREYLSELYRSICENEIQMI
PEQGAGFPLMTSGCWINVLHKSKITDPFIICDSRVLLDHDMFAILSGPTIAAMSVVFDQTEHEEVLKTCVDGFLA
IAKVSSSYHFDDILDDLIVSLCKFTAHLTPFSVDDAILAFGEDTKARMATTTVFTIVNSYGDYIRSGWNNILDCV
LSFHKLGLLPARLINDTVDDMELSSDLERAKPASSSSLISRTPSGTTPRKSSGLMGRFSQLLSFDMEEPRSLPTE
EQLAAHQLTRETIQSCHIDSIFSESKFLQAESLLQLVKSLILAASRLSKGTGPVDDEGASVFCLELLIAITLNNR
DRIMLIWQEVYEHISNVVQSTIMPCTLVEKAVFGLLRICQRLLPYKENLTDELLKSLQLILKLDARVADAYCEQI
TQEVMRLVKANATHIRSHVGWRTITSLLSITARHPEASETGFETLAFIMSDGAHLLPSNYVFCVDAARQFAESRL
GDVDRSVSALDMMAGSVVCLARWSSEAQIAVGQEAAMKLSRDIGEMWLRMVQGLRKVCLDHREEVRNHAIVMLQR
SLVGVDMIHLPIALWFQCFDLVIFTLLDDLLDISLESSPKNYWKMEETLVLAMKLMAKAFLQQLQDLSQQPSFCR

LWLGVLNRMERYMKVKFRGKRSEKIHELVPELLKNILLVMKTTGVLKPSDDIGGDSFWQLTWLHVKNICPPLQSE
VFQDHELEQMQAKNMKSGVPAPDETLTVPSSERTV*
**>Hevbr_LOC110653973**
MEKVAEEEEYKFYSWKEKKRRKELSLSCMLNTELSAVLAVIRRPQGSTGSMLPHEDSFDISILQSLKSLRALIFN
PQQEWRTIDPSVYISPFLDVIQSDDIPAMATNVALSAISKILKLHIFDEKTPGAKEAINSIVTGITSCRLERTDP
VTEDAIMMRILQALTATIKHKASILLADHAVCTIVNTCFHVVQKSANRADLLQRGARYAMREMIEVIFARLQDVE
VKLGEESESDTEDIDIDSNMDSGYGVRCVVDIFHFLCSLLNVEEVVEVDGVSSRTIDENVQIFGLILINSAVELS
GDTIGKHPKLLRMIQDDLFHHLIHYGVSSSPLMLSMICSTVLSIYHSLRRFIRVQLEAFFEFVLLKAAATGSSYQ
LQEVALEGIINFVRQPSFIVEMYVNYDCDPICRNIFEEIGKLLCKLSFPGSSPLSSMQVQAFEGLLITIHNIADN
VDNEDDSSPSGPYPVEITEYRPFWEEKPNEELDTWVEYLRFRKAQKKKILIAGDHFNRDEKKGLEYLKICQLVSD
PADPKAFAVFFRYTPGLDKSIIGDYLGDPDEFHMQVLKEFTETFKFSGMILDTALRTYLATFRLPGESQKIQRIL
EAFSERFYDQQSSDTFATKDAVFILCYSLIMLNTDQHNPQVKKKMTEDEFIRNNRAINAGQDLPREYLSELFHSI
ATNAITLFGQSGPVEMNPGSWIELMNRSKVIHPLILCDFDRRLGRDMFACIAGPSVAAISSFFEHADDDEMLHEC
IGGLISVARVAQYDLEDILDELLASFSKFTTLLNPYASAEETLFAFSNDLKPRMATLAVFTIANNFGYSIRGGWR
NVVDCLLKLKRLKLLPQSVIEFDETSTSSDGLRHKRNESGISPSYDPKFGCRRSSGMVSRFSHFLTLESMEDSIS
LGMSEFEQNLKIIKQCRIGSIFNNSCNLPDDAMLNLGRSLIFAAGGKGQKFSTPIEEEETVGFCWDLIVAISFVN
INRLHNLWPSFHENLLGVAQLPLFSPIPFAEKAIVGLFKICVRLLSSPRAERLPEESIFKSINLMWKLDKEILDT
CFDTCCEFIIKSVSKILTEYPANLQTILGWKSSLHLLSVTGRHPETYELGVNTFITLMSDATYVSRINYAYCVDC
AFGFVALKNSPLEKNLKILDILSDSVNLLNQWYKDYSDSGSNFSIASSTSNSSMDDSSKGFTTPNFAMNLFIKLG
EAFRKTSLARREEIRNHTILSLQKSFSLAEELDFSPLNCINCFNLVIFAMVDDLHEKMVEYSRRDNAEKEMRSME
GTLKLAMELLTSVYLQFLKPITMSPGFRTFWLGVLRRMDTCMKADLGDYGETKLQEVVPDLLRKIITKMKEEEIL
VPKEDDDLWDITYIQIQWIAPSLKEELFPDLEI*
**>Verfo_054407**
MGRLKLQSGIKAIEEEPEDCDPSYSNKATLACMINAEVGAVLAVMRRNVRWGGRYMSGDDQLEHSLIQSLKALRK
QIFSWQHPWHTINPAVYLQPFLDVIRSDETGAPITGVALSSVYKILALDVMDQNTVNVEDAMHLVVDAVTSCRFE
VIDPASEEVVLMKILQVLLACMKSKASVSLSNQHVCTIVNTCFRIVHQAGSKGELLQRIARHTMHELVRCIFSHL
PDVDNTEHALVNGVSPVKQEIGGIDNDYSFGSKQMENGNNSSEFDGQGSSVNFGSSASTGLVATVMEENTIGGSS
GKDALPYDLHLMTEPYGVPCMVEIFHFLCSLLNVVEHMGMGPRSNTIAFDEDVPFFALGLINSAVELGGPSIRRH
PRLLSLIQDELFRNLMQFGLSMSTLILSMVCSIVLNLYHHLRTELKLQLEAFFSCVILRLAQSRYGASYQQQEVA
MEALVDFCRQKTFMVEMYANLDCDITCSNVFEDLANLLSKSAFPVNCPLSAMHILALDGLIAVIQGMAERIGNGS
VSSEQAPVKLEEYTPFWMVKCDNYSDPNHWVPFVRRRKYIKRRLMIGADHFNRDPKKGLEFLQGTHLLPDKLDPQ
SVACFFRYTAGLDKNLVGDFLGNHDEFCVQVLHEFAGTFDFQGMNLDTALRLFLETFRLPGESQKIQRVLEAFSE
RYYEQSPQILANKDAALLLSYSLIMLNTDQHNVQVKKKMTEEDFIRNNRHINGGNDLPREFLSELYHSICKNEIR
TTPEQGAGFPEMTPSRWIDLMLKSKKTAPFIVSDSKAYLDHDMFAIMSGPTIAAISVVFDHAEHEDVYQKCIDGF
LAVAKISACHHLEDVLDDLVVSLCKFTTLLNPASVEEPVLAFGDDTKSRMATVTVFTIANRYGDYIRTGWRNILD
CILRLHKLGLLPARVASDAADESELAADPGHGKPITNSLSSVHMQSMGTPRRSSGLMGRFSQLLSLDTEEPRSQP
TEQQLAAHQRTLQTIQKCHVDSIFTESKFLQAESLLQLARALIWAAGRPQKGNSSPEDEDTAVFCLELLIAITLN
NRDRIVLLWQGVYEHIANIVQSTVMPCALVEKAVFGLLRICQRLLPYKENLADELLRSLQLVLKLDARVADAYCE
QITQEVSRLVKANATHIRSLMGWRTITSLLSITARHPEASEAGFDALLFIMSDGAHLLPANYVLCVDAARQFAES
RVAQAERSVRALDLMAGSVDCLARWSHEAKEAMSEEEAAKLSQDIGEMWLRLVQGLRKVCLDQREEVRNHALLSL
QKCLTGVDGINLPHGLWLQCFDLVIFTMLDDLLEIAQGHSQKDYRNMDGTLIIAVKLLSKVFLQLLHDLSQLTTF
CKLWLGVLSRMEKYLKVKVRGKKSEKLQEVVPELLKNTLLVMKTKGVLLQRSALGGDSLWELTWLHVNNIAPSLQ
SEVFPDQDWEQSQHKQGETVGLVSDETGSVPSNGSVASEGSGTGG*
**>Verfo_053260**
MGHLNLQSETDSLHRKFNECPVKATKGALACMVNSEIGAVLAVMRRNVRWGVRYVADDDQLEHALIHSLKELRKQ
IFSWQNKWQNIDPTIYLQPFLDVIRSDETGAPITGVALSSVYKILALDVLDVDTVNVAEAMHLIVDAVTTCRFEV
TDPASEEVVLMKILQVLLACMKSKASVKLSNQHVCNIINTCFRVVHQASSKGELLQRIARHTMHEVVRCIFSHLP
DMDMYEDTLPNGSPSTGREDDAMYKDNTSQSKQLENGNVRVESDGQTSIGDAPSDLMDKLESGEDGNKLEVSDGM
DAAGSGEKLMMEPFGVPCIVEIFHFLCSLLNVSEHIEVVPRSNPIAYDEDVPLFALGLINSAVELAGPSFSKHPA
LLSLIQGELFRNLMQFGLSMSPLILSTVCSIVLNLYHHLRTELKVQFEAFFSCVLLRIVQSKHGSSYQLQEVAME
ALVDLCRQQSFMAEMYANFDCDITCSNVFEDLANVLSKSAFPVNGPVSAMHIVALDGLISMIKGMIDRIGNELSL
PEEASEDLEGYKAFWTMKCENYIDPKCWIPHVRKMRQIKKRLMVGVDHFNQDPKKGLEFLQGMHLLPDNLDPQSV
ASFFRYTAGLDKNLIGDFLGNHGEFHIMVLQEFAGTFDFRGMNLDTALRLFLGTFRLPGESQKIQRVLEAFAERY
YEQSPQILADKDAALVLSYSLILLNTDQHNVQVKKKMTEEDFIRNNRRTNGGKDFPREYLSDLFRSICESEIQMI
PEQGSGFPLMTSGRWINVLHKSKITGPFIVCDSRVLLDYDMFTILSGPTIAAMSVVFDQTEHEEVLKTCVDGFLA
IAKISASYHFDDVLDDLVVSLCKFTAHLTQLSVEDAILAFGDDTKARMATTTVFTIANRYGDYIHSGWKNILDCV
MSFHMLGLLPARLASDAADDTELSSDLEKAKPAPGSSSTSHPPSGITPRKSSGLMSRFSQLLSFDMEEPRSLPTE
EQLVAHQHTRETIQSCHIDSIFADSKFLQAESLLQLVRSLVLAANCLSKGTSPVEDEGAAVFCLELLIVITLNNR
DRIMLIWQEVYEHISNVVQSTIMPCTLVEKAVFGLLRICQRLLPYKENLTDELLKSLQLILKLDARVADAYCEQI
TQEVMRLVKANATHIRSHMGWRTITSLLSITARHPEASEIGFETLAFIMTDGAHLLPSNYVLCVDAARQFAESRL
GDVDRSVSALDMMAGSVVCLARWSSEAKIAAGHEAAMKVSQDIGEMWLRLVQGLRNVCQDHREEVRNHAILMLQR
SMAGVDAIHLPNALWFQCFDLVIFALLDELLDISLESSPRNFRKMEETLVLAMKLMTKAFLQQLQDLSQQPSFCR
LWLGVLSRMERYMKVKFRGKHSERIHELVPELLKNILLVMKTTGVLLPSDDIGGDSFWQLTWLHVKNICPSLQSE

VFPDNELEQLQAKHIKTGGVPAPDETLLVPSSETTV*
**>Verfo_035762**
MEGEDEENKSRSLLWQEKKRRKEVSLSCMLNTELSALLAVIRRPHDPTAALLQEDCFDTSLLQCLKSLRALIFNP
QQEWRTIDPSVYISPFLDVIQSDDIPAVATNVALSAISKILKQQIFDEKTPGAKDAINSIVTGITSCRLERTDPS
NEDAVMMRILQALTSVIKHRASILLTDHAVCTIVNTCFQVVQQSANRADLLQRGARYSMREMIEIIFARLQDVEV
KLGEDSESDTEDIDIDDMDSGYGFRCVVDIFHFLCSLLNVIEIVETEGFSAHTCDENVQIFGLILINSAIELSGD
AIGKHPKLLRMIQDDLFHHLIHYGMSSSPLVLSMICSTVLNIYHALRRFIRVQLEAFFGFVLLKAASSGASQLQE
VALEGIINFCRQPTFIVEMYVNYDCDPICRNIFEEIGKLLCKLSFPGSSPLSCVQIQAFEGLLILAHTLADHVDK
ENDSTPSGPYPVQITEYKPFWEEKPKEDYDTWVEYLRLRKAQKKKIMIAGDHFNRDEKRGLEYLKLCNVVSDPVD
PKAFAIFFRYTQGLDKAMIGDYLGDPDDFHLQVLKEFTETFKFSGMILDTALRTYLATFRLPGESQKIQRILEAF
SERFYDQQSSDIFASKDAVFILCYSLIMLNTDQHNPQVKKKMTEEEFIRNNRAINAGQDLPREYLSELFNSIATN
AITLFGQSGPVEMNPGSWIELINRSKVMQPYILCDFDRRIGRDMFASIAGPSVAAISSFFEHADEDEMLHECIGG
LISVARIGQYDLEDILDELLASFSKFTTLLNPYASAEETLFAFSNDLKPRMATLAVFTIANNFGDSIRGGWRNIV
DCLLKLKRLKLLPQSVVEFDDTSASASDGLGHRRNESGVSTSHDPKIGSRRSAGMISRFSHFLTLESMEDSISLG
MSEFEQNLKIIKQCRIGSIFNNSYNLPDDALLNLGRSLIFAAGGKGQKFSTPVEEEETVGFCWDLIVSISLVNIN
RFQNFWPSFHENLLAAIQFPLFSPIPFAEKAMLGLFKICIRLLSSLRAEKLPEEVLFKSITLMWKLDKEILDTCC
DFITKSVGRILTEYPANLQTSLGWKSCLHLLSVTGRHQDTYDQGVDTLIALMSDGAYISRTNYAYCIDCAFGFIA
LKNSPLEKNLKILDLLSDSVNSLIQWYKEYSDPGSNVSAASSTSNSSLDESSKGFGSPNFAVNLFVKLGEAFRKT
SLARREEIRNHAITSLKKSFALAEDLDFSPLNCINCFNLVIFAMVDDLHEKMIEYSRRDNAEREMRSMEGTLKLA
MELLTAVYLQFLKPITMSPGFRTFWLGVLRRMGTCMKADLGEYGETKLQEVVPDFLRKIITKMKEDEILVQKEGD
DLWDITFIQIQWIAPSLKEELFPELEF*
**>Eucgr_LOC104433426**
MGRLKMPPGINAIEEEPEEYDSSCSNKAILGCVINTEIAAVLSVMRRNVRWGGRYMSGDDQLEHSLIHSLKALRK
QIFSWQHQWHTVNPILYLQPFLDVIRSDETGAPITGVALSSVYKILTIDVIDQTTANVEDAMHLVVDSVTSCRFE
VTDPASEEVVLMKILQVLLACMKSKASVMLSNQHVCTIVNTCFRIVHQAGSKGELLQRIARHTMHELVRCIFSHL
PDVDNTERALVNGGNKVNQEIRGPDGEYAFGTQQLDNGNGGSEHDGQLSSGSFASNSSAAPMSGMMDEIGAGAGK
DAVPIDLHMMTEPYGVPSLVEIFHFLCSLLNLFEHTGMGPRTNTIAFDEDVPLFALGLINSAIELGGPSIRRHPR
LLSLVQDELFRNLMQFGLSLSPLILSMVCSIVLNLYQHLRTELKLQLEAFFSCVILRLAQSRYGASYQQQEAAME
ALVDFCRQKTFMVEMYANLDCDITCSNVFEDLANLLSKSAFPVNCPLSSMHILALDGLIAVIQGMAERIANGSLS
SEQAPVALDEYTPFWLVKCEDYGDPNHWVPFVRRRKYIKRRLMIGADHFNRDPKKGLEFLQGTHLLPDKLDPQSV
ACFFRYTAGLDKNLVGDFLGNHDEFCVQVLHEFAWTFDFQDMNLDTALRLFLETFRLPGESQKIQRVLEAFSERY
YEQSPQILVNKDAALLLSYSLIMLNTDQHNVQVKKKMTEEDFIRNNRHINGGSDLPRDFLSELYHSICKNEIRTT
PEQGAGYPEMNPSRWIDLMHKSKRTAPFIISDSRAYLDHDMFAIMSGPTIAAISVVFDHAEQEEVYQTCIDGFLA
VAKISACHHLEDVLDDLVVSLCKFTTLLNPSSVEEPVLAFGDDAKARMATITVFTIANRYGDYIRTGWRNILDCI
LRLHKLGLLPARVASDAADETEVSTEPGHGKPVANSLAAAHMPSMGTPRRSSGLMGRFSQLLSLDTEEPRSQPTE
QQLAAHQRTLQTIQKCHIDSIFTESKFLQAESLLQLARALIWAAGRPQKGNSSPEDEDTAVFCLELLIAITLNNR
DRIVLLWQGVYDHIANIVQSTVMPSALVEKAVFGLLRICQRLLPYKENLADELLRSMQLVLKLDARVADAYCEQI
TQEVSRLVKANATHIRSQMGWRTITSLLSITARHPEASEAGFEALLFIMSDGAHLLPANYVLCVDASRQFAESRV
GQAERSVRALDLMSGSVDSLARWAREAREAMAEDEVAKMSLDIGEMWLRLVQGLRKVCLDQREEVRNHALLSLQR
CLTGVDGIQLPHNLWIQCFDVVIFTMLDDLLEIAQGQSQKDYRNMEGSLILAMKLLSKVFLQLLHDLSQLTTFCK
LWLGVLSRMEKYMKVKVRGKKSEKLQELVPELLKNTLLVMKTRGVLVQRSALGGDSLWELTWLHVNNIAASLQSE
VFPDQELQQAETQGDQVSEETVSAHPNESLANNGN*
**>Eucgr_LOC104440351**
MAEEQPQRASEEEDSSQCKDTRDNPDPRSKRKELGLSCMLNTEVGAVLAVIRRPLDPAGPFLPTHEDQCDSSIAH
SLKSLRSLIFNPQQEWRALDPSIYLSPFLDVIQSDEVPATATGVALSSILKMLKLEMFDEKTLGAKDAINSIVTA
TVNCRLERTDPGSEVAVMMRILQVLASIMKHRASVFLSDQAVCTIVNTCFQVVQQSASRGDLLQRNARYTMHELI
QIIFSRLPEIEVRDGDNNSESETDDVGGNLDSGYGVRCTVDIFHFLCSLLNVVEIVETEGNTSHTTDEDVQLFAL
VLINTAIDLSGDAIAKHPKLLRMIQDDLFHHLIHYGMSSSQLVLAMICSTILSIYHFLRRFIRLQLEAFFSFVLL
KVATFATSIQLQEVAIEGIINFCRQPSFIFEAYVNYDCNVTCRNVFEEMGKLLCKHAFPTSTPLTSLQIQAFEGL
VIVIHNIADNINWEENSSTLARHAVEITEYRPFWVEKFEDEDENWVDSARMRKAQKRKMMIAGNHFNHDERKGLE
YLKLSRLVSDPIDPKAFAFFFRYTPGLDKNLIGDYLGDPDEFHVQVLKEFTDTFEFAGMILDTALRTYLETFRLP
GEAQKIQRVLEAFSEKFFDQQSTEIFVSKDAVFVLCYSLIMLNTDQHNPQVKKKMTEEEFIRNNRGINEGSDLPR
EYLSELFYSISNNAIKLYGQSGLSIEMSPSRWIELMDRSKTIHPYVQCNFNSTLGRDMFATIAGPSVATLCACFE
HADEDEFLHECVEGLFSIARIAQYGLEDTLDELLASFCKFTTFLNPYASPEETLFAFSHDLKPRMATLAVFTIAN
NFGSSIRGAWRNIIDCLLKLKRLKLLPQSVVETDASSNSSSDLAMDTRCESGIVNPNQDFRPGAQPNYGMMGRFS
HFLSLDSPEESVSLGLSEFEQNMKIIRQCRIGNIFTNSSSLPNETILNLGRSLIFAAAGKGQKFSTPVEEEETVG
FCWELIVAIALANLHRFSTFWVSFHDNLLTVAQFPQFSPVPFAEKAMLSLLKVCLKLLAVPQQDSIPEELIFRSI
NLMWKLDKEILDTCCEVITKSVSRMLIEFPANLQTQLGWKSALHLLSITGRHPETYDQGVETLIMLMSDGTHVSR
MTYAYCIDCAFGFVALKNSPLEKNLKILDLLSESVNLLIQWHKNGLSDAGSSYSASSFEDNLKGLGSSNFTLNLF
VKLGEALRKTSLARREEIRNHAILSLQRSFTLAEDLEFSSTNCINCFNLVIFAMVDDLHEKMLEYSRRENAEREM
RSMEGTLKIAMEFLTNVYLQFLKQISESPGFRTFWLGVLRRMDTCMKANLGPYGESAILKELVPDLLRKIITKMK
ESEILVQKEDDDLWDITYIQIQWISPLLKEMFPEEDT*
**>Syzol_LOC115671905**

MGRLKMPPGINAIEEEPEDYDSSCSNKATLGCVINTEIAAVLAVMRRNVRWGGRYMSGDDQLEHSLIQSLKALRK
QIFSWQHQWHTVNPILYLQPFLDVIRSDETGAPITGVALSSVYKILTLDVIDQTTANVEDAMHLVVDSVTSCRFE
VTDPASEEVVLMKILQVLLACMKSKASVMLSNQHVCTIVNTCFRIVHQAGSKGELLQRIARHTMHELVRCIFSHL
PDVDNTEHALVNGGNKVKQEVRGPDSEYAFGTQQLDNGNSGSEHDGQLSSGSFASNSSTAPIPGMMDEIGAGAGK
DAVPVDLQMMTEPYGVPSLVEIFHFLCSLLNLFEHTGMGPRSNTIAFDEDVPLFALGLINSAIELGGPSIRRHPR
LLSLVQDELFRNLMQFGLSMSPLILSMVCSIVLNLYQHLRTELKLQLEAFFSCVILRLAQSRYGASYQQQEVAME
ALVDFCRQKTFMVEMYANLDCDITCSNVFEDLANLLSKSAFPVNCPLSSMHILALDGLIAVIQGMAERIANGSLS
SEQAPVALDEYTPFWLVKCDDYGDPNHWVPFVRRRKYIKRRLMIGADHFNRDPKKGLEFLQGTHLLPDKLDPQSV
ACFFRYTAGLDKNLVGDFLGNHDEFCVQVLHEFAGTFDFQDMHLDTALRLFLETFRLPGESQKIQRVLEAFSERY
YEQSPQILVNKDAALLLSYSLIMLNTDQHNIQVKKKMTEEDFIRNNRHINGGSDLPRDFLIELYHSICKNEIRTT
PEQGAGYPEMNPSRWIDLMHKSKRTAPFIISDSRAYLDHDMFAIMSGPTIAAISVVFDHADHEEVYQTCIDGFLA
VAKISACHHLEDVLDDLVVSLCKFTTLLNPSSVDEPVLAFGDDAKARMATITVFTIANRYGDYIRTGWRNILDCI
LRLHKLGLLPARVASDAADETEVSAEPGHGKPVANSLAAAHMPSMGTPRRSSGLMGRFSQLLSLDTEEPRSQPTE
QQLAAHQRTLQTIQKCHIDSIFTESKFLQAESLLQLARALIWAAGRPQKGNSSPEDEDTAVFCLELLIAITLNNR
DRIVLLWQGVYEHIANIVQSTVMPCALVEKAVFGLLRICQRLLPYKENLADELLRSMQLVLKLDARVADAYCEQI
TQEVSRLVKANATHIRSQMGWRTITSLLSITARHPEASEAGFDALLFIMSDGAHLLPANYVLCVDASRQFAESRV
GQAERSVRALDLMAGSVDSLARWACEAREAMAEDEVAKMSQDIGEMWLRLVQGLRKVCLDQREEVRNHALLSLQR
CLTGVDGIQLPHNLWSQCFDLVIFTMLDDLLEIAQGQSQKDFRNLEGTHILAMKLLLKVFLQLLHDLSQLTAFCK
LWLGVLSRMEKYMKVKVRGKKSEKLQELVPELLKNTLLVMKTGGVLVQRSTLGGDSLWELTWLHVNNIAPSLQSE
VFPDEELPQAETQGNQVSEETVSAHSNESLASNGS*
**>Syzol_LOC115690464**
MAEELTQPAPEEEDSSQCKDRDKPGPRSKRKELGLSCMLNTEVGAILAVIRRPLDPAGPFLPTQEDHYDSSIAHS
LKSLRTLIFNPQQEWRAIDPSIYLSPFLDVVQSDEVPAAATGVALSSILKMLKLEMFDEKTHGAKDAINSIVTAT
VNCRLERTDSGSEEAVMMRILQVLAGIMKHRASVFLSDQAVCTIVNTCFQVVQQSASRGDLLQRNARYTMHELIQ
IIFSRLPEIDGRDGDNNSESETDDVGGNLDSGYGVRCTVDIFHFLCSLLNVVEIVETEGNTSHTTDEDVQLFALV
LINSAIDLSGDAVAKHPKLLRMIQDDLFHHLIHYGMSSSQLVLAMICSTILSIYHFLRRFIRLQLEAFFSFVLLR
VAMFATSIQLQEVAIEGIINFCRQPSFIFEAYVNYDCNVTCRNVFEEMGKLLCKHAFPTSNPLTSLQIQAFEGLV
IVIHNIADNINCEENSGTLARHAVEITEYRPFWVEKFEDEDENWVDSARIRKAQKRKMMIAGNHFNRDERKGIEY
LKLSRLVSDPMDPKAIAFFFRYTAGLDKNMIGDYLGDPDEFHVRVLKEFTDTFEFAGMILDTALRTYLETFRLPG
EAQKIQRVLEAFSERFFDQQSAEVFVSKDAVFVLCYSLIMLNTDQHNPQVKKKMTEEEFIRNNRGINEGSDLPRE
YLSELFYSISSNAIRLYGQSGISIEMSPSRWIELMNRSKTVHPYVQCKFNSTLGRDMFATIAGPSVATLCACFEH
ADEDEFLHECVEGLFSIARIAQYGLEDTLDELLASFCKFTTFLNPYASAEETLFAFSHDLKPRMATLAVFTIANN
FGSSIRGAWRNIIDCLLKLKRLKLLPQSVVETDASSTSSSDLAMDTQCESGIVNPNQDFRPGAQHNYGMMGRFSH
FLSLDSAEESLSLGLSEFEQNVKIIRQCRIGNIFTNSSSLPDEAILNLGRSLIFAAAGKGQKFSTPVEEEETVGF
CWELIVAIALANIHRFSIFWASLHENLLTVAQFPLFSPIPFAEKAMLSLLKVCLKLLAEPQQDSISEELIFRSIN
LMWKLDKEILDTCGEVITKSVSRILIEYPANLQTQLGWKSVLHLLSVTGRHPETYDQGVETLIMLLSDGTHVSRT
TYAYCIDCAFGFVALKNSPLDKNLKILDLLSESVNLLIQWHRNGLSDVGNNYSTASIEDNLKGLGSSNFSLNLFI
KLGEALRKTSLARREEIRNHAILSLQRSFALAEDLEFTSTNCINCFNLVIFAMVDDLHEKMLEYSRRENAEREMR
SMEGTLKIAMELLASVYLQFLKQISESPGFRTFWLGVLRRMDTCMKANLGPYGESAMLKELVPDLLRKIITKMKE
SEILVQKEDDDLWDITYIQIQWISPLLKELFPEEDI*
**>Corci_B2628**
MGRLKMPPGINAIEEEPEDYDSSCSNKASLGCVINTEIAAVLAVMRRNVRWGGRYMSGDDQLEHSLIQSLKALRK
QIFSWQHQWHTVNPILYLQPFLEVIRSDETGAPITGVALSSVYKILTLDVIDQTTANVEDAMHLVVDSVTSCRFE
VTDPASEEVVLMKILQVLLACMKSKASVMLSNQHVCTIVNTCFRIVHQAGSKGELLQRIARHTMHELVRCIFSHL
PDVDNTERALVNGGNKVKQEVRGPDNEYAFGTQQLDNGNSGSENDGQLSSGGFASNSSTAPISGMMDDIGAGAGK
DAAPLDLHMMTEPYGVPSLVEIFHFLCSLLNLFEHTGMGPRSNTIAFDEDVPLFALGLINSAIELGGPSIRHHPR
LLSLVQDELFRNLMQFGLSMSPLILSMVCSIVLNLYQHLRTELKLQLEAFFSCVILRLAQSRYGASYQQQEVAME
ALVDFCRQKTFMVEMYANLDCDITCSNVFEDLANLLSKSAFPVNCPLSPMHILALDGLIAVIQGMAERIANGSLS
SEQAPVALDEYTPFWLVKCDDYGDPNHWVPFVRRRKYIKRRLMIGADHFNRDPKKGLEFLQGTHLLPDKLDPQSV
ACFFRYTAGLDKNLVGDFLGNHDEFCVQVLHEFAGTFDFQDMNLDTALRLFLETFRLPGESQKIQRVLEAFSERY
YEQSPQILVNKDAALLLSYSLIMLNTDQHNVQVKKKMTEDDFIRNNRHINGGSDLPRDFLSELYHSICKNEIRTT
PEQGSGFPEMNPSRWIDLMHKSKRTAPFIISDSRAYLDHDMFAIMSGPTIAAISVVFDHAEHEEVYQTCIDGFLA
VAKISACHHLEDVLDDLVVSLCKFTTLLNPSPVEEPVLAFGDDAKARMATITVFTIANRYGDYIRTGWRNILDCI
LRLHKLGLLPARVASDAADETEVSAEPGHGKPVANSLAAAHMPSMGTPRRSSGLMGRFSQLLSLDTEEPRSQPTE
QQLAAHQRTLQTIQKCHIDSIFTESKFLQAESLLQLARALIWAAGRPQKGNSSPEDEDTAVFCLELLIAITLNNR
DRIVLLWQGVYDHIANIVQSTVMPCALVEKAVFGLLRICQRLLPYKENLADELLRSMQLVLKLDARVADAYCEQI
TQEVSRLVKANATHIRSQMGWRTITSLLSITARHPEASEAGFDALLFIMSDGAHLLPANYVLCVDASRQFAESRV
GQAERSVRALDLMSGSVDSLARWACEAREAMAEDEVAKMSLDIGEMWLRLVQGLRKVCLDQREEVRNHALLSLQR
CLTGVDGIQLPHNLWIQCFDVVIFTMLDDLLEIAQGQSQKDFRNMEGTLILAMKLLSKVFLQLLHDLSQLTTFCK
LWLGVLSRMEKYMKVKVRGKKSEKLQELVPELLKNTLLVMKTRGVLLQRSALGGDSLWELTWLHVNNIAPSLQSE
VFPDQELQQAETQGNQVSEETVTSHSNESLANNGS*
**>Corci_C2224**

MAEELTQRASEEEDSSQCKDRDKPDSQSKRKELGLSCMLNTEVGAVLAVIRRPFDPAGPFLPTHEDQFDSSVAHS
LKSLRSLIFNPQQEWRALDPSIYLSPFLDVIQSDEVPATATGIALSSILKMLKLEMFDEKTHGAKDAINSIVTAT
VNCRLERTDPGSEEAVMMRILQVLASIMKHRASVFLSDQAVCTIVNTCFQVVQQSASRGDLLQRNARYTMHELIQ
IIFSRLPEIEVREGDNNSESDTDDVGGNLDSGYGVRCMVDIFHFLCSLLNVVEIVETEGNASHTTDEDVQLFALV
LINSAIDLSGDAIMKHPKLLRMIQDDLFHHLIHYGMSSSQLVLAMICSTILSIYHFLRRFIRLQLEAFFGFVLLR
VAMFETSIQLQEVAIEGIINFCRQPSFIFEAYVNYDCNVTCRNVFEEMGKLLCKHAFPTSNPLTSLQIQAFEGLV
IVIHNIADNINWEENSSMLARHAVEITEYRPFWVEKFEDEDENWVDSARIRKAQKRKMMIAGNHFNRDSKKGLEY
LKLSHLVSDPMDPKAFAFFFRYTPGLDKNMIGDYLGDPDEFHVQVLKEFTDTFEFGGMILDTALRTYLEMFRLPG
EAQKIQRVLEAFSEKFFDQQSAEVFVNKDAVFVLCYSLIMLNTDQHNPQVKKKMTEEEFIRNNRGINEGSDLPRE
YLSELFYSISNNAIKLYGQSGHSIEMSPSRWIELMDRSKTIHPYVQCNFNSTLGRDMFATIAGPSVATLCACFEQ
ADEDEFLHECVEGLFSIARIAQHGLEDTLDELLASFCKFTTFLNPYASAEETLFTFSHDLKPRMATLAVFTIANN
FGSSIRGAWRNIIDCLLKLKRLKLLPQSVVEADPSSPLLSDLAMDTRCESSIANPNQDFRPGAQHNYGMMGRFSH
FLSLESPEESVSLGLSEFEQNMKIIRQCRIGNIFTNSSSLPNEAILNLGRSLIFAAAGKGQKFSTPVEEEETVGF
CWELIVTIALANVHRFSTFWVSFHDNLLMVAQFPLFSPVPFAEKAMLSLLKVCLKLLAVPQQDSIPEELIFRSIN
LMWKLDKEILDMCCEIITKSVSRMLIEYPANLQTQLGWKSVLHLLSVAGRHPETYDQGVETLIALLSEGTHVSKM
TYAYCIDCAFGFVALKNSPLDKNLKILDLLSESVNLLIQWHRNGSSDGGSSYSGSSFEDNLKGLGSSNFSLNLFI
KLGEALRKTSLARREEIRNHAILSLQRSFTLAEDLEFTSTNCINCFNLVIFAMVDDLHEKMLEYSRRENAEREMR
SMEGTLKIAMELLTNVYLQFLQQISESPGFRTFWLGVLRRMDTCMKANLGPYGESAVLTELVPDLLRKIITKMKE
SGILVQKEDDDLWDFTYIQIQWISPLLKELFPEEDT*

**>Rhoar_LOC115755598**

MGRMKMPPGINAIEEEPEDYDSPFANKATLGCVINTEIAAVLAVMRRNVRWGGRYMSGDDQLEHSLIQSLKALRK
QIFSWQHQWHTVNPILYLQPFLDVIRSDETGAPITGVALSSVYKILTLDVIDQTTANVEDAMHLIVDSVTSCRFE
VTDPASEEVVLMKILQVLLACMKSKASVMLSNQHVCTIVNTCFRIVHQAGSKGELLQRIARHTMHELVRCIFLHL
PDVDNTEHALVNGGNKVKQEVRGPDSEYAFGTQQLDNGNSGSELDGQLSSGSFTSNSSTTPISGMMDEFGAGAGK
DAVPVDLQMMTEPYGVPSLVEIFHFLCSLLNLFEHTGMGPRSNTIAFDEDVPLFALGLINSAIELGGPSIRRHPR
LLSLVQDELFRNLMQFGLSMSPLILSMVCSIVLNLYQHLRTELKLQLEAFFSCVILRLAQSRYGASYQQQEVAME
ALVDFCRQKTFMVEMYANLDCDITCSNVFEDLANLLSKSAFPVNCPLSSMHILALDGLIAVIQGMAERIANGSLS
SEQAPVALDEYTPFWLVKCDDYGDPNHWVPFVRRRKYIKRRLMIGADHFNRDPKKGLEFLQGTHLLPDKLDPQSV
ACFFRYTAGLDKNLVGDFLGNHDEFCVQVLHEFAGTFDFQDMNLDTALRLFLETFRLPGESQKIQRVLEAFSERY
YEQSPQILVNKDAALLLSYSLIMLNTDQHNVQVKKKMTEEDFIRNNRHINGGSDLPRDFLTELYHSICKNEIRTT
PEQGAGYPEMNPSRWIDLMHKSKRTAPFIISDSRAYLDHDMFAIMSGPTIAAISVVFDHAEHEEVYQTCIDGFLA
VAKISACHHLEDVLDDLVVSLCKFTTLLNPSSVEEPVLAFGDDAKARMATITVFTIANRYGDYIRTGWRNILDCI
LRLHKLGLLPARVASDAADETEVSAEPGHGKPVANSLAAAHMPSMGTPRRSSGLMGRFSQLLSLDTEEPRSQPTE
QQLAAHQRTLQTIQKCHIDSIFTESKFLQAESLLQLARALIWAAGRPQKGNSSPEDEDTAVFCLELLIAITLNNR
DRIVLLWQGVYEHIANIVQSTVMPCALVEKAVFGLLKICQRLLPYKENLADELLRSMQLVLKLDARVADAYCEQI
TQEVSRLVKANATHIRSQMGWRTITSLLSITARHPEASEAGFDALLFIMSDGAHLLPANYVLCVDASRQFAESRV
GQAERSVRALDLMGGSVDSLARWACEAREAMPEDEVAKMSEDIGEMWLRLVQGLRKVCLDQREDVRNHALLSLQR
CLTGVDGIQLPHILWSQCFDVVIFTMLDDLLEIAQGQSQKDYRNMEGTLILAMKLLSKVFLQLLHDLLQLTTFCK
LWLGVLSRMEKYMKVKVRSKKSEKLQELVPELLKNTLLVMKTRGVLVQRSALGGDSLWELTWLHVNNIAPSLQSE
VFPDQELQQPETQGNQVSEETVSAHSNESLASNGS*

**>Rhoar_LOC115729910**

MAEELTQRASEEEDSSQCRDRAKHDPRSKRKQLGLSCMLNTEVGAVLAVIRRPLDPAGPFLPTQEDQYDSSIAHS
LKSLRTLIFNPQQEWRVMDPSIYLSPFLDVVQSDEVPAAATGIALSSILKMLKLEMFDEKTLGAREAINSIVTAT
VNCRLERTDSGSEEAVMMRILQVLSSIMKHRASVFLSDQAVCTIVNTCFQVVQQSASRGDLLQRNARYTMHELIQ
IIFSRLPEIEGRDGDNNSESETDDVGGSLDSGYGVRCTVDIFHFLCSLLNVVEIVEAEGNTSHATDEDVQLFALV
LINSAIDLSGDAIAKHPKLLRMIQDDLFHHLIHYGMSSSQLVLAMICSTILSIYHFLRRFIRLQLEAFFGFVLLR
VAMFATSTQLQEVAIEGIINFCRQPSFIFEAYVNYDCNVTCRNVFEEMGKLLCKHAFPTSNPFTSLQVQAFEGLV
IVIHNIADNINCEENSSTIARHAVEITEYRPFWVEKFEDEDENWVESARMRKAQKRKMMIAENHFNRDERKGIEY
LKLSRLVSDPMDPKAFAFFFRYTPGLDKNMIGDYLGDPDEFHVQVLKEFTDTFECAGMILDTALRTYLETFRLPG
EAQKIERILEAFSERFYDQQSAEVFVSKDAVFVLCYSLIMLNTDQHNPQVKKKMTEEEFIKNNRGINGGSDLPRE
YLSELFYSISNNAIKLYGQSGLSVEMSPSRWIELMNRSKNAHPYVECKFNSTLGRDMFATIAGPSVATLCACFEH
ADEDEFLHECVEGLFSIARIAQYGLEDTLDELLASFCKFTTFLNPYATAEETLFAFSHDLKPRMATLAVFTIANN
FGSSIRGAWRNIIDCLLKLKRLKLLPQSVVETDASSTSSSDLAMDTRCESGVVNPNQDFRPGTQQNYGMMGRFSH
FLSLETAEESLSLGLSEFEQNVKIIRQCRIGNIFTNSSSLPDEAILNLGRSLIFAAAGKGQKFSTPVEEEETVGF
CWELIVAIALANVHRFSTFWASFHDNLLTVAQFPLFSPVPFAEKAMLSLLKVCLKLLAVPQQDSIAEELIFRSIN
LMWKLDKEILDTCCEVITKSVSRMLIEYPANLQTQLGWKSVLHLLSVTGRHPETYDQGVETLIMLLSDGTHVSRM
TYAYCIDCAFGFVALKNSPLDKNLKILDLLSESVNLLIQWHRNGLSDPGSSYSASSFEENLKGLGSSNFSLNLFV
KLGEALRKTSLARREEIRNHAILSLQRSFALAEDLEFTSTNCTNCFNLVIFAMVDDLHEKMLEYSRRENAEREMR
SMEGTLKIAMELLTDVYLQFLKQISESPGFRTFWLGVLRRMDTCMKANLGPYGESEILKELVPDLLRKIITKMKE
SEILVQKEGDDLWEITYIQIQWISPLLKELFPEEDG*

**>Rhoar_LOC115729907**

MAEELTQRASEEEDSSQCRDRAKHDPRSKRKQLGLSCMLNTEVGAVLAMIRRPLDPAGPFLPTQEDQYDSSIAHS
LKSLRTLIFNPQQEWRVMDPSIYLSPFLVVVQSDEVPAAATGIALSSILKMLKLEMFDEKTLGAREAINSIVTAT

VNCRLERTDSGSEEVVMMRILQVLSSIMKHRASVFLSDQAVCTIVNTCFQVVQQSASRGDLLQRNARYTMHELIQ
IIFSRLPEIEGRDGDNNSESDTDDVGGSLDSGYGVRCTVDIFHFLCSLLNVVEIVEAEGNTSHATDEDVQLFALV
LINSAIDLSGDAIAKHPKLLRMIQDDLFHHLIHYGMSSSQLVLAMICSTILSIYHFLRRFIRLQLEAFFGFVLLR
VAMFATSTQLQEVAIEGIINFCRQPSFIFEAYVNYDCNVTCRNVFEEMGKLLCKHAFPTSNPFTSLQVQAFEGLV
IVIHNIADNINCEENSSTIARHAVEITEYRPFWVEKFADEDENWVESARMRKAQKRKMMIAENHFNRDERKGIEY
LKLSRLVSDPMDPKAFAFFFRYTPGLDKNMIGDYLGDPDEFHVQVLKEFTDTFECAGMILDTALRTYLETFRLPG
EAQKIERILEAFSERFYDQQSAEVFVSKDAVFVLCYSLIMLNTDQHNPQVKKKMTEEEFIKNNRGINGGSDLPRE
YLSELFYSISNNAIKLYGQSGLSMEMSPSRWIELMNRSKTVHPYVECKFNSTLGRDMFATIAGPSVATLCACFEH
ADEDEFLHECVEGLFSIARIAQYGLEDTLDELLASFCKFTTFLNPYATAEETLFAFSHDLKPRMATLAVFTIANN
FGSSIRGAWRNIIDCLLKLKRLKLLPQSVVETDASSTSSSDLAMDTRCESGVVNPNQDFRPGTQQNYGMMGRFSH
FLSLETAEESLSLGLSEFEQNVKIIRQCRIGNIFTNSSSLPDEAILNLGRSLIFAAAGKGQKFSTPVEEEETVGF
CWELIVAIALANVHRFSTFWASFHDNLLTVAQFPLFSPVPFAEKAMLSLLKVCLKLLAVPQQDSIAEELIFRSIN
LMWKLDKEILDTCCEVITKSVSRMLIEYPANLQTQLGWKSVLHLLSVTGRHPETYDQGVETLIMLLSDGTRVSRM
TYAYCIDCAFGFVALKNSPLDKNLKILDLLSESVNLLIQWHRNGLSDPGSSYSASSFEENLKGLGSSNFSLNLFV
KLGEALRKTSLARREEIRNHAILSLQRSFALAEDLEFTSTNCTNCFNLVIFAMVDDLHEKMLEYSRRENAEREMR
SMEGTLKIAMELLTDVYLQFLKQISESPGFRTFWLGVLRRMDTCMKANLGPYGESEILKELVPDLLRKIITKMKE
SEILVQKEGDDLWEITYIQIQWISPLLKELFPEEDG*
**>Pungr_LOC116199793**
MGRLKMPPGIKAIEEEPEDYESCSEKATLACMINSEVAAVLAVMRRNVRWGGRYMSGDDQLEHSLIQSLKALRKQ
IFSWQNQWETVNPTLYLQPFLDVIRSDETGAPITGVALSSVYRILTLDVIDQNNANVEEAMHSVVDAVNSCRFEV
TDPASEEVVLMKILQVLLACMKSKASVMLSNQHVCNIVNTCFRIVHQAGSKGELLQRIARHTMHELVRCIFSHLP
EVDNTERALVNGGNKVKQEVTGPDNEYAFNSRQLENGNSGSEYDGQLSSGSFASNASGVPMAGMMDDSAIGAGKD
AVPLDLHLMNEPYGVPCMVEIFHFFCTLLNALEHIGVGPRSNTIAFDEDVPLFALGLINSAIELGGPSIRSHPRL
LGLIQDELFRNLMQFGLSASPLILSMVCSIVLNLYQHLRTELKLQLEAFFSCVILRLAQSKYGASYQQQEVAMEA
LVDFCRQKTFMVEMYANLDCDITCSNVFEDLANLLSKSAFPVNCPLSSMHILALDGLIAVIQGMAERIGNGSGSS
EQAPVKLDEYTPFWMVKCDNYGDPNHWVPFVRRRKYIKRRLMIGADHFNRDPKKGLEFLQGTHLLPDKLDPQSVA
CFFRYTAGLDKNLVGDFLGNHDEFCVQVLHEFAGTFDFQDMNLDTALRLFLETFRLPGESQKIQRVLEAFSERYY
EQSPQILANKDAALLLSYSLIMLNTDQHNVQVKKKMTEEDFIRNNRHINGGNDLPREFLSELYHSICKNEIRTTP
EQGAGFPEMTPSRWIDLMHKSKKTAPFIVSDSRAYLDHDMFAIMSGPTIAAISVVFDHAEHEDVYQTCIDGFLAV
AKISACHHLEDVLDDLVVSLCKFTTLLNPSSVEEPVLAFGDDSKARMATVTVFTIANRYGDYIRTGWRNILDCIL
RLHKLGLLPARVASDAADESELSPADPGHGKPIANPSSAAHMPSMGTPRRSSGLMGRFSQLLSLDTEEPRSQPTE
QQLAAHQRTLQTIQKCHIDSIFTESKFLQAESLLQLARALIWAGGRPQKGSSSPEDEDTAVFCLELLIAITLNNR
DRIVLLWQGVYEHIANIVQSTVMPCALVEKAVFGLLRICQRLLPYKENLADELLRSMQLVLKLDARVADAYCEQI
TQEVSRLVKANAAHIRSQMGWRTITSLLSITARHPEASEAGFDALLFIMSDGAHLLPANYVLCAEASRQFAESRV
GQAERSVQSLDLMAGSVDCLARWAREAKEAMAEEDVAKMSLDIGEMWLKLVQGLRKVCLDQREEVRNHALLSLQR
CLTGVDEIQQSHSLCLQCFDPVIFTMLDDLLEISQGQSQKDYRNMESTLILAMKLLSKVFLQLLHELSQLTNFCK
LWLGVLSRMEKYTKVKVRGKKSEKLQELVPELLKNTLLVMKTKGVLMQRSALGGDSLWELTWLHVNNIAPSLQSE
VFPDQEQEQLKQADEKTGTSIPSEAVVAKVAGTAG*
**>Pungr_LOC116209960**
MGCPNRPSGAREVKEGYEDCSFTSSDKHATFACIINSEVGAVLAVMRRNVRWGGHYLSGNDQREHSLIQSLMALR
KQIFYWQHQWNDTDPRFYLQPFLDIIQSDETGAPITGVALSAVYKILSVDMIDQYTVNAESALQLVVDAVTSCRF
EVTDPASEEVVLMKILQVLLACMKSKASTVLSNQHVCSIVNTCFRIVHQAGTKGELLQRIARNTMHELVRCIFSR
LQDFDKTYHPVPSGAITVKQEVGGVHIEDSTGRNQLENGSSISNYDGQLSPGTVTSDAAGVPVTPLPSKTVSRDD
DGKDATSSGLHMQKEHYGIPCMVEIFQFLSSLLTAVEHIGMGTRANSIAFDEDVPIFALGLINSALELSGSSLHH
HPSLLHLIRDEVFQNFIQFGLSMSPLVLSMVFSIIILNLYHHMHTELKLQLEAFFSCVILRLAQGGNGHSYPQEEV
AMEALVDFCRQKEFVIDMYVNLDCDVTCSNVFEDIVNLLSKSAIPEDSSLSSMHTLALDGLIAVIKGMTERIGSG
SINSKGTLLNSEEHTPIWTVRCDDYSDPNHWVSFVQWRKLLKKSVMAGVDQFNTNPKKGLEFLQQAHLLPDKLEP
QSVSCFFRYTPGLDKNLIGDFLGNHDQFCIQTLHAFAGTFDFRGMNLDNALRLFLETFRLPGESQKIQRILEAFS
ERYYEQSPQILANKDAALLLSYSLILLNTDQHSSQVKRKMTENEFIRNNRLINGGNDLPREYLLDLYHSICNKEI
CFTDKGAASMEMTSSRWLDLIDKSKKTDPFIVSDCKAYLDRDMFAIVSGPTVAAISVVFELAEEDDVYQTCINGF
LDVAKVSACYHFEDVLDDLVVSLCGFTILLNLSSFDEPVLGEDMKARMATEAVFTIANRYGDSIRASWRNILDCI
LRLHKLGLLPAHMASDAADESEHSANAGNRRSTLSSSPNAQSQKTGSPQGYVGFVSRFSRLFSFDTDEPRSEPTE
QQLAAHRQARQIIEKCHIESIFTESKFLQAESLLQLAQALIWASEQAAKQNGSPEDEDTAVFCLELLIAITLNNR
DRIGLLWQCVYEHITSIVQSIVMPCALVEKAVFGLLRICQRLLPYKEDLADELLRSLQLVLKLDARVADAYCEQI
TQEINCLVKSNAVHIRSQMGWRVITSLLSISARHPEAAEAGFDSLFFIMSGGAHLLPVNYLLCMDASRQFAESRV
GGAERSLLALDLMAGSLDCLVQWTHEAKKAMEEQDAARFCQDIGEMWLRLVQGLKRVCSDQREEIRNHALLSLQN
CLIGADGIDLPHYLWLQCFNSVIFSMLDDLIEITQVRSYKDYRNMEGTLILAVKLLSKAFLQLLHWLSRVETFGQ
LWVDVLTRMDKYMKVKVRGKKSEKLLELVPELLKNTLLVMKARGILTPRSSTSQGDVLWEVTWTNVNNVDPSLRS
EVFSDQELEQQSQQAESGADSISNGTSVSTSG*
**>Pungr_LOC116200818**
MEEAKDQNPEDDQIIAKQRRKELGLSCMLNAEVGAVLAVIRRPSDHYISPQEESFDSSILHSLKSLRALIFNPQQ
EWRMIDPSVYLSPFLDVVQSDDVPAVATGVALSALLKILKLGIFDEKTPGAKDAINSVVTGIVNCRLGKTDPGFE
EAVLMRILQVLSNTMRHRASILLTDHAVSTVVNTCFQVVQQSASRGNLLQRNARYAMHELIQIIYSRLPEIEVKE

GDNSESDTDDVCSLSESGYGVRCMVDVFQFLCSLLNVVEVVETDGATSQTADEDVQIFALALINSAIELSGDEIG
RHPKLLRMIQDDLFHHLIHYGTSSSQLVLAMICSTVLSIYHFLRRFIRLQLEAFFGYVLFRVALHGTSTQLQEVA
TEGIINFCRQPAFILESYVNYDCNIMCRNVFEEIGKLLCRHAYPTSSPMTSLQIQAFEGLVIMIHNIADNINMED
PSSFEPHTMEITDYRPFWEEKPEEEEENWVDLVRLRKAQKRKIMIAGNHFNRDDKKGLDYLKFSKLVSEDALDPK
GFAFFFRFTPGLDKDAIGDYLGDPDEFHIKVLKEFTNTFEFSGMVLDTALRTYLQTFRLPGESQKIHRIIEAFSD
RFYDQQSSEIFASKDAVFVLCYSIIMLNTDQHNPQVKKRMTEEDFIRNNRAINGGNDLPREYLSELFHSISTNEI
KIFNHSGLPQEMGPNRWLELMNRSKSANPFFQCNFDGRVGRDMFATIAGPSVATLSACFEHAEEDEFLHECVEGL
FSVGRIAQYGLEDTLDELLATFCKFTTLLNPYASSEETLFAFSNDLKSRMATLAVFTIANSFGFAIRGGWRNIVD
CLLKLKRLKLLPQTVIELDTDSDVSSSAGSESIFSNQDPEFGGSRQNSGMIGRFSHFLAVDSAEDSLTLGASEFE
QNLKIIKQCRIGNIFAGSKNLPDEGLLHLGRSLIFSAAGKGQKFSTPVEEEETVGFCWELLNAIITANIHRFSMF
WPAFHENLLAVAQFPLFSPIPFAEKAVLCLLKVCLRLISIRRNDILPEELIFRSITLMWKLDKEILETCSESIFK
LISRMLVEFPDNIQSQIGWKSVLHLLVVTGRSPETFDQGVETLITILSDTNHLSRINYVCCVDCAFGFIGLKSCP
VEKKIKVLELLTDTINLLIQWHRHGFSDPGNNYSTTSNTSSSSLEDHSKALGTSNLALNMFVKLAEFLKNTSLAR
REEVRNHAIQDLGRIFAPANNLDLSSTSCIKCFNLVVFSMVDDLHEKVVVYSRRENVESELRSWEESLKLAMEVL
TDIYLHFLNLIWESPGFRTFWLGVLRRMDTCMKADLGPYGEPGLQKLVPELLRRMITKMKEDNILLPRDGDDLWE
ITYIQIQWIAPFLKEELFPDL*

**>Medtr_LOC25489342**

MGRLKLQTGINSIEEEEPEECDAACPSRTTLSCMINSEIGAVLAVMRRNVRWGSRYMSGDDHLEHTLIQSFKALK
RQIFSWNHHQWQAINPTLYLQPFLDVIRSDETGAAITGVALSSVYKILTLDVIDHNTVNVEDAIHLVVDAVTSCR
FEVTDTSSEEVVLIKILQVLLACMKSKASVMLSNQHVCTIVNTCFRIVHQAGNKGESLQQISRYTMHELVRCIFS
HLQDVDNTDHALVNGSSNLKQETGGGLNNEYAFGSRKLENGSMSSEYDNQPLPSNFASNAASVVTVTMMDENTAI
ALTGKESASYDVHLMTEPYGVPCMVEIFHFLCSLLNVIENMELGPRSNTIAFDEDVPLFALTLINSAIELGGPSI
HRHPRLLSLIQDELFHNLMQFGLSMSPLILSMVCSIVLNLYHHLRTELKLQLEAFFTCVILRLAQSRYGASYQQQ
EVAMEALVDFCRQKTFMVDMYANFDSDITCSNVFEDLANLLSRSAFPVNCPLSAMHILALDGLIAVIQGMAERIA
NGSANSEYSPVNLEEYIPFWMVKCENYGDPNHWVTFTRRRKYIKRRLMIGADHFNRDPKKGLEFLQGTHLLPDKL
DPQSVACFFRYTAGLDKNLVGDFLGNHDEFCVQVLHEFAGTFDFQDMNLDTALRLFLETFRLPGESQKIHRVLEA
FSERYYEQSPHILANKDAALVLSYSMIMLNTDQHNVQVKKKMTEEDFIRNNRHINGGSDLPREFLTEIYHSICKN
EIRTTPEQGVGFPEMTPSRWIDLMHKSKKTAPFIVSGSKAYLDHDMFAIMSGPTIAAISVVFDHAEHEDVYQTCM
DGFLAIAKISACHHLEDVLDDLVVSLCKFTTLLNPSLVEEPVLAFGDDMKARMATVTVFTIANRYGDYIRTGWRN
ILDCILRLHKLGLLPARVASDAADESELSAETVHGKPIVNSLSSAHMQSIGTPRRSSGLMGRFSQLLSLDTEEPR
SQPTEQQLAAHQRTLQTIQKCHIDSIFTESKFLQAKSLEQLAKALIWAAGRPQKVNSTPEDEDTAVFCLELLIAI
TLNNRDRIGILWPGVYDHISNIVQSTVMPCALVEKAVFGLLRICQRLLPYKENIADELLRSLQLVLKLDARVADA
YCEQITQEVSRLVKANASHIRSQLGWRAITSLLSITARHIEASEAGFDALLFIMSDGAHLLPANYVLCVDTARQF
AESRVGQAERSVRALDLMTGSVNCLAQWTSEGKEAMEDEQMTKLSKDIGDMWLILGQGLRKVCLDQREEVRNHAL
SSLQKCLIGADDIYLPYGKWLDCFDLVIFTVLDDLLEISQGHSQKDYRNMEGTLILAVKLLSKVFLQSLPVLSQL
TTFCKLWLGVLTRMEKYMKVKVRGKRSEKLQETVPDLLKNSLLAMKMRGILAQRSALGGDSLWELTWLHVNNISP
SLQLEVFPEQDSEHSQYKQGESVGGLLPDEKVSVPSSEIPNLEDAGIVG*

>**Medtr_LOC25489344***

MGHLNMQMPAGVNGIEDQSQQFGAGYPNKTTIACMINAEIGSVLAVMRRNVRWGVHYMSDDDQSEHFLVQSLKTL
RRQIFSWQNQWHSINPALYLQPFLDVIQSDETGAPITGVALSSVYKILTLDVIDQNTVNVVDTMDLVVDAVTSCR
FEVTDPGSEEVVLMKILQVLLACVKSKASVILSNQHICTIVNTCFRIVHQAGTKGELLQRIARYTMHELVRCIFS
HLQEIDIAEPALINGSTTSKQETRELNNEHASESRQFENGSLNSAYDGQPSSTGLASSTAYVGTVTVMPENTTTA
SSDKENDIRELQLMTEPYGVPCMVEIFHFLCSLLNVVEHMGMNPRSNTLTFDEDVPLFALTLINSAIELGGPSFR
HHPRLLSLIQDEVFHILMQFGLSMSPLVLSLVCSIVLNLYHHLRTELKLQLEAFFSCVILRLAQSKYGASYQQQE
VLMEALVDLCRQKTFMVEMYVNFDCDISCSNVFEDIANLLSKSAFPVNNPLSSMHILALDGLIAVMQGMADRIGN
GFVNSEHSPVNFEEYIPFWLEHCENFGDPNDWVPFVRRRKYFKRRLMIGADHFNHDAKKGLEFLQGAHLLPDKRD
PQSVACFFRFTAGLDKNVIGDFLGNHDEFCVQVLREFARTFDFQDMTLDTALRIFLETFRLPGESQKIHRVLEAF
SERYYEQSPHILANKDAALVLSYSIIMLNTDQHNVQVKKKMTETDFIRNNRHINGGNDLPREFLSEIYHSICKNE
IRTTPESGFGSPGMTPSRWISLIHKSKTTAPYIVSDSRAYLDYDMFVILSGPTIAAVSVVFDNAENEEVYQTCID
GFLAIAKISAYYHLENVLDDLIVCLCKFFTILDPLSVEESVLAFGDDTKARMVTETVFTIASRYGDYIRTGWRNI
LDCIFRLHNLGLLPTHLASDAVDESEPSTKTGRGKLNSNSFSSTHLQSISTPKRSSGLLSRFSQLLSLATDEPRA
IPTEEQLAAHHQAVQTIQKCHIDSIFTESKFLQAESLLQLVRSLINAGAQSQKGSKKSQNEDTSVFCLELLVAIT
LNNRDRIGLIWKGVYEHISNIVKSTVMPCALVEKAVFGLLRICHRLLPYKENMADELLKSLQLILKLDALVADVY
YDQITQEVSRLVKANASRIRSHIGWRTITSLLSVTARHLEASEVGFDTLSFIMSDGNYLLPANYILCIDAARQFA
ESRVGLVDRSIGSLDLMVGSFNCLQKWTSDAKQAAKEEEVAKMLQDIGEMWLRLVQGLRKVCLDQREEVRNHALL
SLQKCMTEAGKTHLPGDLWLQCFDQVIFTLLDDLLDISQERSQKDYRNIEETLTLALKLLFKVFLLLLQELSQLE
EAFCKLWVGVLSHAEKYMKVKVRGKRSEMLQLPVPELLKNALVVMKSGGILVESSGLAESNLWELTWLHMKNIAP
SLQSEVFQEHDAEQLQQKQIETVGDLESDAKISVPSNETLGQDGAGIG*

**>Medtr_LOC11438427**

MAHAKLQMQTDINAMEEEAGQCGVGYLSRTSVACMINAEIGAVLAVMRRNVRWGVHYMSDDDQLEHSLVQSLKAL
RREIFSWQSQWHVINPVLYLQPFLDVIQSDETGAPITGVALSSVYKILTLDVIDQNTVNVGDSMHLVVEAVTSCR
FEVTDPGSEEVVLMKILQVLLACVKSKASVMLSNQHICTIVNTCFRIVHQAGTKSELLQRIARYTMHELVRTIFS
HLQDIDVTEHALVNGSTALKEEIDGQNIEHNSMHNQLENGSLISASDSQSVSTDIASNTVSDVAAVIVDANTATS

SGKETDLNKQLMNEPHGIPCMLEIFRFLCSLLNVGEHMGMSPRSNTIAFDEDVPLFALTLINSAIELGGSSFHLH
PRLLSLIQDELFCNLMQFGLSMSPLVLSMVSSIVLNLYHHLRTEIKFQLEAFFSCVILRLAQSKYGASYQQQEVA
MEALVDFCRQKAFVVEMYANFDCDITCSNIFEDIANLLSKSAFPVNSPLSSMNILALDGLIAVIQGMAERIGNGS
LSSEHSVVNLEEYTPFWLEKCENFNDPNDWVPFVGRRKHFKKRLMIGADHFNRDTKKGLQFLQGTHLLPDKLDPQ
SVAFFFKYTTGLDKNLIGDYLGNHDEFCVQVLQEFARTFDFNDMALDTALRIFLETFRLPGESQKIQRVLEAFSE
RYYEQSPHILANKDAALLLSYSIIMLNTDQHNSQVKKKMTEEDFVRNNRRINGGNDLPREVLSELYHSICKNEIR
TTPEQGSAFPEMTPSRWIYLIHKSKNTAPFIVSDCRAHLDYDMFSIMSGPTVAAISVVFDNAETEEVYQTCMDGF
LAVAKVSAYYHLESILDDLVVSLCKFVTILDPLSPEESILAFGEDTKARMATETVFTIANRYGDYIRTGWRNILD
CILKFHKLGLLPAQMANDAAEESEPSTETGNGKRYANSLSSSQLLSVNTPKRSSGFISRFSQLLYLGAEETRSEP
SEEQLAAQQCSLQTIQKCHIESIFTESKFLQAESLLHLVKALKSAGVRPKKGNGTSEDEDTSVFCLELLVAITLN
NRDRIELLWQDVYEHISNIVQSTVMPCTQVEKAVFGLLRICHRLLPYKENMTDELLRSLQLVLKLDARVADTYYE
QITQEVSNLVKANASHIRSQLGWRTITSLLSITARHLESSEAGFDALFFIMSDGAHILPSNFALCVDAAKQFAES
RVGQVERSVVALDLMAGSINCFEKWANDAKQATTEEMAKMLQNIEDMWLRLVQGLKKLCMDQREEVRNHALLSLQ
NCLTASVGIHLPHDLWLQCFDQVIFTVLDDLLESSQTHSPKDYRNMEGTLILALKLLSKVFLLLLQDLSQSTDFS
KLWLNVLNRFEIFMKVKIRGRRSEKFQELVPELMKNTLLVMKASHVLEQSSSSGDEKSLWELTWVHINNIAPSLQ
SEVFPEQEAKQLEQEKAEQVGDRGPAENVSAPSRAGNELS*
**>Medtr_LOC11437178**
MDSIEDFTHEELEDDRNQKKKDTTKSKRRQLGLSCMLSTEVGAVLAVIRRYPEFTPVYNISSPEESYDSSIINSL
RSLRSLIFNPQQEWRTIDPSIYLTPFLDVIQSDDVPASATGVALSSILKILKFQVFDDRTPGAKEAMESVVVGIT
SCRLEKTDPVSEDVVMMKILQVLAGIMHHKASVLLSDQAVCTLVNACFQVVQQSVGRGDLLQRSARYTMHELIQA
VYARLPEIEGNDREGDSESDVEDVDDNAGLKSGYGVRAAIDIFHFLCSLLNVVSVVEADGSTTHTADEDVQIFAL
VLINSAIELSGDKIGNHPKLLRMVQDDLFHHLIYYGTWSSSFVLSMICSTVLNAYHFLRKFLRFQLEAFFGHVLI
RVATLGSTMQLQEVAVEGIINFCRQPTFIAEVYANYDCNPLCRNIFEEVGRLLCKHSFALTGHLTSLHIQAFEGL
LIMIHNIADNIDKIDNRTPLGPYTTQLIEYKPFWEEKEKDDDLEAWVEHVRITKVQKKKLLIAANHFNRDNKKGL
EYLKHAKLISDPPDPKSYAYFFRFTPGLDKKALGEYLGDPDSFYLEVLREFTETFHFNGMVLDTGLRFFLESFWL
PGESQKIERVLDAFSNRFYDQQSSDLFASKDTVLILCYSLIMLNTDHHNAQVKKKMTEEEFIKNNRTINSGQDLP
REYLSELFQSITNNAIVMKQTELSLDMTQSKWTQLINQSKVMQSFVQCEFDRRMCRDMFACIAGPSVAALSSFFE
HADEEELMHECIEGLFSVARIAQYGLEDTLDELITSFCKFTTLLNPYASTEETMFTFSHDMKPRLATVAVFTLAN
DFRDSIRGGWKNIVDCLLKLKRLRLLPQSTIDFDAPANAPTMPESSGVVSPTANHKSGTHHRFPSVMTRLSLLSQ
ENSEDGLTLGSEFEQNLKMIKLCRIGSIFGSSSNIPKECLQNLGRSLIFAAAGKGQKFSTPVEEEETVGFCWDLI
TALSLANVHRFHIFWPNFHEYLLTVAQFPMFSPIPFAEKALLALLKVCLKLFSTPREDKLAEELIFKSITLMWKL
DKEILDMCYEIIAQLMSKIVIQYPANLQTQVGWKSVLNLLSVAWRHPETYDLAIEALISLFSDATNVTRTNYAYL
IDCAFGCFLAKNSPIEKKKKILDLLADSTNLLVQWQRTQYTDPGSNVSVVSNTSSSSAEDNFRGPSSANYNMNLF
VKLGEIFRRTSLSRQEEIRNHAVYSLHKSFNLAEEMLYTSSNCISYFNLVIFAMVDELHEKMLEYSRRDNAEREM
RSMEGTLKLAMELLSAMYLQSLRQITESPGFRAFWLGVLRRMDTCMKAELGHYGPSSLSEIIPDLLKKIITQMKD
EGILEPKEDDDMWEITYIQIQWICPRLKDELFPL*
**>Tp_L195_g002385**
MGRLKLQAGINSIEEEEPEECDAACPNRTTLSCMINSEIGAVLAVMRRNVRWGSRYMSGDDHLEHSLIQSFKALR
RQIFSWNHHQWQAINPTLYLQPFLDVIRSDETGAAITGVALSSVYKILTLDVIDHNTVNVEEAIHLVVDAVTSCR
FEVTDTTSEEVVLIKILQVLLACMKSKASVMLSNQHVCTIVNTCFRIVHQAGNKGESLQQISRYTMHELVRCIFS
HLQDVDNTDHALVNGSSNLKQETGGLNNEYAFGSRKLENGSVNSEYDNQPLSANFASNSASAVTVTMMDENTAIA
LTGKESASYDVHLMTEPYGVPCMVEIFHFLCSLLNVIENMELGPRSNTIAFDEDVPLFALTLINSAIELGGPSIH
RHPRLLSLIQDELFHNLMQFGLSMSPLILSMVCSIVLNLYHHLRTELKLQLEAFFSCVILRLAQSRYGASYQQQE
VAMEALVDFCRQKTFMVDMYANFDSDITCSNVFEDLANLLSRSAFPVNCPLSAMHILALDGLIAVIQGMAERITN
GSADSEYSPVNLEEYIPFWMVKCENYGDPNHWVPFTRRRKYIKRRLMIGADHFNRDPKKGLEFLQGTHLLPDKLD
PQSVACFFRYTAGLDKNLVGDFLGNHDEFCVQVLHEFAGTFDFQDMNLDTALRLFLETFRLPGESQKIHRVLEAF
SERYYEQSPHILANKDAALVLSYSMIMLNTDQHNVQVKKKMTEEDFIRNNRHINGGSDLPREFLSEIYHSICKNE
IRTTPEQGVGFPEMTPSRWIDLLHKSKKTAPFIVSGSKAYLDHDMFAIMSGPTIAAISVVFDHAEHEEVYQTCMD
GFLAIAKISACHHLEDVLDDLVVSLCKFTTLLNPSLVEEPVLAFGDDMKARMATVTVFTIANRYGDYIRTGWRNI
LDCILRLHKLGLLPARVASDAADESELSSETVHGKPIVNSLSSAHMQSIGTPRRSSGLMGRFSQLLSLDTEEPRS
QPTEQQLAAHQRTLQTIQKCHIDSIFTESKFLQAKSLEQLAKALIWAAGRPQKVNSTPEDEDTAVFCLELLIAIT
LNNRDRIGILWPGVYDHISNIVQSTVMPCALVEKAVFGLLRICQRLLPYKENIADELLRSLQLVLKLDARVADAY
CEQITQEVSRLVKANASHIRSQLGWRAITSLLSITARHIEASESGFDALLFIMSDGAHLLPANYVLCVDTARQFA
ESRVGQAERSVRALDLMTGSVNCLTRWTSEAKEAMEEEQMSKLSKDIGDMWLILGQGLRKVCLDQREEVRNHALL
SLQKCLTGADDIYLPYGKWLDCFDLVIFTVLDDLLEISQGHSQKDYRNMEGTLILAVKLLSKVFLQSLPVLSQLT
TFCKLWLGVLTRMEKYMKVKVRGKRSEKLQETVPDLLKSSLLAMKMRGILAQRSALGGDSLWELTWLHVNNISSS
LQLEVFPEQDSEHFQHKQGELVGGLVPDEKVSVSSSEIASLEDAGIVG*
**>Tp57577_23676**
MGNVKLQMQTGINAMEEEAGQSGAGYPSRTTVACMINAEIGAVLAVMRRNVRWGVHYMSDDDQLEHSLVQSLKAL
RRQIFSWHSQWHVINPVLYLQPFLDVIQSDETGAPITGVALSSVYKILTLDVIDQNTVNIGDAMHLVVDAVTSCR
FEVTDPGSEEVVLMKILQVLLACVKSKASVMLSNQHICTIVNTCFRIVHQAGTKSELLQRIARYTMHELVRSIFS
HLQDIDVTEHALENGSAALEEEIDGQKNEHNSANSQLENGSLISASDSQSVSTGVASSTLSDVAAIIVDVNAAAS
SGKETDPNEQLMNEPYGIPCMVEIFHFMCSLLNVVEHMGMSPRLNTIAFDEDVPLFALTLINSAIELGGSSFHRH

PKLLSLIQDELFGNLMQFGLSMSPLVLSMVSSIVLNLYHHLRTELKLQLEAFFSCVILRLAQSKYGASYQQQEVA
MEALVDFCRQKAFVVEMYANFDCDITCSNIFEDIVNLLSKSAFPVNSPLSTMNILALDGLIAVIQGMAERIGNGS
LSSEHSIVNLEEYTPFWLEKCENFTEPNNWVPFVGRRKQFKKRLMIGADHFNRDTKKGLQFLQGTHLLPEKLDPQ
SVAFFFRYTTGLDKNLIGDYLGNHDEFCVEVLHEFARTFDFNDMALDTALRIFLETFRLPGESQKIQRVLEAFSE
RYYEQSPHILANKDAALLLSYSIIMLNTDQHNSQVKKKMTEEDFVRNNRRINGGNDLPREVLSGLFHSICKNEIR
TTPEQGSTFPEMTPSRWIYLIHKSKSTAPFIVSDCKAHLDYDMFTILSGPTVAAISVVFDNAETEVVYQTCMDGF
LAVAKVSAYYHLDSILDDLVVSLCKFVTILDPLSPEESILAFGEDIKARMATETVFTIANRYGDYIRTGWRDILD
CILKFHKLGLLPSQMANDAEESEPSTETENGKRYANSLASSQLVSVNTPKRSSGLISRFSQLLYLGAEETKSEPT
EEQLAAQQSSLQTIQKCHIGSIFTESKFLQADSLLHLAKALKSAGVKPKKGSEDEDTSVFCLELLVAITLNNRDR
IELLWPDVYEHISNIVQSTVMPCAQVEKAVFGLLRICHRLLPYKENITDELLKSLQLVLKLDARVADTYYEQITQ
EVSNLVKANASHIRSQLGWRTITSLLSITARHLESSEAGFDALFFIMSDGAHILPANFVLCIDAAKQFAESRVGQ
VERSVVALDLLAGSVNCLEKWTNDAKQVMEEEEVAKMLQDIGDMWLRLVQGLKKLCLDQREEVRNHALLLLQNCL
TGSVGIHIPHELWLQCFDQVIFTVLDDLLEISQTHSPKDYRNMEGTLVLAVKLLSKVFLLLLQDLSQSTDFSKLW
LGVLSRFEIFMKVKIRGRKSEKFQELVPELLKNTLLVMKAGRVLERSSSSGDENSLWEQTWLHINNIAPSLQSEV
FPEHDSKQLEQKQTEQVGDRGPAENVSVPSNETAGEDGTGNGIG*

**>Tp57577_2954**

MGQLKMQMPAGLNNGIEDQFEQCDAGYPNKTTIACMINAEIGSVLAVMRRNVRWGVHYMSDDDQSEHFLVQSLKT
LRRQIFSWQDQWHSINPALYLQPFLDVIQSDETGAPITGVALSSVYKILTLDVIEQNTVNVIDTMDLVVDAVTSC
RFEVTDPGSEEVVLMKILQVLLACVKSKAKASVILSNQHICTIVNTCFRIVHQAGTKGELLQRIARYTMHELVRC
IFSHLQEIDITESPKQETCELNNEHASASRQFENGSPNSAYGGQPLSPSIASSTAYVGTVTAMPENTTTASSDKE
SDLRELQLMTEPYGVPCMVEIFHFLCSLLNVVEHMGVNPRSNTLTFDEDVPLFSLSLINSAIELGGPSFRHHPRL
LRLIQDEVFHILMQFGLSMSPLVLSMVCSIVLNLYHHLRTELKLQLEAFFSCVILRLAQSKYGASYPQQEVLMEA
LVDLCRQKTFMVEMFVNFDCDISCSNVFEDIANLLSKSAFPVNNPLSSMHILALDGLIAVMQGMADRIGNGFVSS
EHSPVNFEEYIPFWLEQCENFGDPNEWVPFVRRRKYFKRRLMIGVDHFNRDAKKGLKFLQGTHLLPDKLDPQSVA
CFLRFTAGLDKNVIGDFLGNHDEFCVQVLREFSRTFNFQDMTLDTALRIFLETFRLPGESQKIHRVLEAFSERYY
EQSPHILVNKDAALVLSYSIIMLNTDQHNVQVKKKMTEADFIRNNRHINGGEDLPREFLSEIYQSICKNEIRTTP
ESGFGSPGMTPSRWISLIHKSKATAPYIVSDSRAYLDYDMFVILSGPTIAAVSVVFDNAENEEVYQTCIDGFLAI
AKISAYYHLENVLDDLIVCLCKFFTILDPLSVEESVQAFGDDTKARMVTETVFTIASRYGDYIRTGWRNILDCIL
RLHKLGLLPTYLASDAVDESEPSTKTGHGKLKSNSLSLPRLQSISTPKRSSGLLSRFSQLLSLATDEPRAIPTEE
QLAAHHHAVQTIQKCHIDSIFTESKFLQAESLLQLVRSLINAGAQSQKRKKISQFEDASVFCLELLVAITLNNRD
RIGLIWKSVYEHISNIVKSTVMPCALVEKAVLGLLRICHRLLPYKENMADDLLKSLQLILKLDALVADAYYEQIT
WEVSRLVKANASHIRSHLGWRTITSLLSVTARHLEASEVGFDTLSFIMSDGTHLLPANYILCIDAARQFAESRVG
LVGRSIGSLDLMVGSFNCLQKWTGDAKQAAKEEEVEKMLQDIGEMWLRLVQGLRKVCLDQREEVRNHALLSLQKC
MTEAVETRLPCDLWLQCFDQVIFTLLDDLLDIAEEHSPKDYPNIEETLILAMKLLFKVFLQLLQELSQLETFSKL
WVGVLSHAENYVKAKVRGRRIEKLQFIVPELLKNTLVVMKSRGILVENSDLWELTWLHMKNIVPSLQSEVLQEQL
DQKQIETVAKLEHDSNISVPSNETLGQDGAVIIS*

**>Tp57577_31612**

MDSIDDFTNEELEDDHHNHKKKTTSKSKRRQLGLSCMLNTEVGAVLAVIRRYPEFGPLYNISSPEETYDSSIISS
LRSLRSLIFNPQQEWRTIDPSIYLSPFLDVIQSDDVPASATGVALSSILKILKFQVFDEKTPGAKEAMESIVAGI
TSCRLEKTDPVSEDAVMMKILQVLAGIMHHKASILLNDQAVCTLVNACFQVVQQSVNRGDLLQRSARYTMHELIQ
VVFSRLPEIEVYDREGDSESDIEEEEEDDDDDDDDDHIHHLESGYGIRSAIDIFQFLCSLLNVVSIVEADGSTTH
TADEDVQIFALVLINSAIELSGDKIGNHPKLLRMVQDDLFHHLIYYGTWSSSFVLSMICSTVLNAYHFLRRSIRF
QLEAFFGHVLIRIATLGTTMQLQEVAVEGIINFCRQPTFIVEVYANFDCDPCSRNIFEEVGRLLCKHSFALNGHV
TSLHIQAFEGLLIVIHNIADNIDKVYDRTPLSPYSTQLNEYKPFWEEMERDDNDIEAWVEHVRTTKVQKKKLLIA
GNHFNRDNKKGLEYLKHAKLISDPPDPKAYAYFFRYTPGLDKKAIGEYLGDPDNFYLEVLKEFTETFYFNGMVLD
NALRFFLESFRLPGESQKIQRVLEAFSDRFYDYSSSDLFASKDTVLILCYSLIMLNTDQHNPQVKKKMTEEEFIR
NNRAINQGQDLPREYLSELFQSISTNAIISAENFVSLDMTQSKWIQLINRSKDSEDFVQCEFDRRMCRDMFACIA
GPSVAALSSFFEHADEEELMHECIEGLFSVARIAQYGLEDTLDELITSFCKFTTLLNPYASTEETMFTFSHDLKP
RLATVAVFTLANDFRNSIRGGWKNIVDCLLKLKRLRLLPPSTIDFDVPADAPTTPGSGAISPTDDHKFGTHRFPS
VMTRLSLLSQENSEDGLQIGSEFEQNLKMIKMCRIGSIFGSSSNIPKECLQNLGRSLIFAAAGKGQKFSTPVEEE
ETVGFCWDLITAISLSNVHRFQIFWPTFHDFLLSVAQFPMFSPIPFAEKALLCLLKVCLKLFSTPREDKLAEEFI
FKSITLMWKLDKEILDTCHDVISQTMSKIVVEYPANLQTQIGWKSVLNLLSVTWRYPETYDLGIEALISLLSQGA
NVTRANYAYCIDCAFGCFLAKNSPIEAKKKILDLLADSTNLLVQWHRTQFSDPGSNISIASITSSSSMEDNFRGS
NSANYNMNLFVKLGEAFRRTSLSRQEEIRNHAVYSLHKSFNLAEEMLYISSNCINYFNVIFAMVDELHEKMLEYS
RRENAEREMRSMEGTLKLSMELLSAMYLTSLKQISESPGFRAFWLGVLRRMDTCMKADLGQYGPSSLSEIIPDLL
KKIITQMKDEGILEPRENDDMWEITYIQIQWICPPLKDELFPL*

**>Phvul_001G235300**

MGRLKLQAGINAIEEEEPEECDAAYPDKTTLACMINSEIGAVLAVMRRNVRWGGRYMSGDDQLEHSLIQSFKTVR
RQIFSWHHHQWQAINPALYLQPFLDVIRSDETGAPITGVALSSVYKILTLDVIDQNTVNVEDAMHLVVDAVTSCR
FEVIDPSSEEVVLMKILQVLLACMKSKASIMLSNQHVCTIVNTCFRIVHQAGSKGELLQQIARYTMHELVRCIFS
HLQDVGNTDHALVNGSTNLKQETGGLDNDYAFGSRQLENGSMSSEYDNQSLSSNSAPNVSSVVKATVMDENTAIT
ISCKDGVPYDMHLMTEPYAVPCMVEIFHFLCSLLNVVEHTGMGPRSNTLAFDEDVPLFALTLINSAIELGGPSIC
RHPRLLSLIQDELFHNLMQFGLSMSPLILSMVCSIVLNLYHHLRTELKLQLEAFFSCVILRLAQSRYGASYQQQE

VAMEALVDFCRQKTFMVDMYANFDCDITCSNVFEDLANLLSKSAFPVNCPLSAMHILALDGLIAVIQGMAERIAN
GSVSSEYSPVNLEEYTPFWMVKCENYNDPNHWVPFVRRRKYIKRRLMIGADHFNRDPKKGLEFLQGTHLLPDKLD
PQSVACFFRYTAGLDKNLVGDFLGNHDEFCVQVLHEFAGTFDFQDMNLDTALRLFLETFRLPGESQKIHRVLEAF
SERYYEQSPHILANKDAALVLSYSMIMLNTDQHNVQVKKKMTEEDFIRNNRLINGGNNLPREMLSEIYHSICKNE
IRTTPEQGVGFPEMTPSRWIDLMHKSKKTAPFIVSDSKAYLDHDMFAIMSGPTIAAISVVFDHAEQEDVYQTCMD
GFLAIAKISACHHLEDVLDDLVVSLCKFTTLLNPSSVEEPVLAFGDDMKARMATVTVFTIANRYGDYIRTGWRNI
LDCILRLHKLGLLPARVASDAADESELSAETVNGKPIMNSLSSAHMQSIGTPRRSSGLMGRFSQLLSLDTEEPRS
QPTEQQLAAHQRTLQTIQKCHIDSIFTESKFLQAESLLQLARALVWAAGRPQKGNSTPEDEDTAVFCLELLIAIT
LNNRDRIGILWHGVYEHISNIVQSTVMPCALVEKAVFGLLRICQRLLPYKENIADELLRSLQLVLKLDARVADAY
CEQITQEVSRLVKANASHIRSQLGWRTITSLLSITARHIEASEAGFDALLFIMSDGAHLLPANYVHCIDTARQFA
ESRVGQAERSVRALDLMAGSVNCLARWTSEAKEAMEEEQVSKLSQDIGEMWLRLVQGLRKVCLDQREEVRNHALL
SLQKCLTGADGIYLPHSMWLQCFDLVIFTVLDDLLEIAQGHSQKDYRNMEGTLILAMKLLFKVFLQLLPELSQLT
TFCKLWLGVLSRMEKYMKVKVRGKRSEKLQETVPELLKNSLLVMKMRGILAQRSALGGDSLWELTWLHVNNISPS
LQLEVFPEQDSEHLQHKQGEPIGGLVPDDKGSVPSSETASREDAGIVG*
**>Phvul_008G231400**
MGHVKLQTQTGINAIEEGSGQCNAGYPDKITVACMINAEIGAVLAVMRRNVRWGVHYMSDDDHLEHSLVQSLKSL
RKQIFSWQNQWHVISPTLYLQPFLDVIQSDETGAPITGVALSSVYKILTLDVIDQNTVSVGDAMHLVVDAVTSCR
FEVTDPGSEEVVLMKILQVLLACVKSKASMMLSNQHICTIVNTCFRIVHQAGSKSELLQRIARYTMHELVRCIFS
HLQDIDNTELTLVNGSTALKKEIGGLNNEHNSANRQLENGSLTSSSNGQSLSTGIASSTVSDVAATVVDEDSSIS
SSGKEADLNELQLINEPYGIPCMVEIFHFLCSLLNVAEHMGMSPRSNTIAFDEDVPLFALTLINSAIELGGPSFH
RHPRLLSLIQDELFRNLMQFGLSMSPLVLSMVCSIVLNLYHHLRTELKLQLEAFFSCVILRLAQSKYGASYQQQE
VVMEALVDFCRQKTFMVEMYANFDCDLTCSNVFEDIANLLSKSAFPVNSPLTSMHIIALDGLIAVMHGMAERIGN
GSLSSEQSPVNLEEYTPFWQEKCEKFCDPKNWVPFVCRKKYFKKRLMIGADHFNRDIKKGLEFLQGSHLLPDKLD
AQSVACFLRYTAGLDKNLIGDFLGNHDEFCIEVLHEFARTFDFEDMMLDTALRLFLETFRLPGESQKIQRMLEAF
SERYYEQSAQILANKDAALLLSYSIIMLNTDQHNSQVKKKMTEEDFIRNNRRINGGNDLPREFLSELYHSICKNE
IRTTPEQGSGFPEMTPSRWIYLKHKSKKSAPFIVSDSKAYLDYDMFSILSGPTIAAISVVFDNAENAEVYQTCMD
GFIAVAKISAYYHLENILDDLVVSLCKFVTVFDPLAVEESILAFGDDMKARMATETVFTIANRYGDFIRTGWRNI
LDCILKFHRLGLLPARMASDAAEESEVSSQTEDGGKRNTNTLSLSRLPSVNTPKRSSGLMSRFSQLLYLGAEEPK
SVPTEEQLVAQQCTLQTIEKCHIDCIFSESKFLQAESLLQLAKAITSAGGRPKKGNSTSEDEDTSVFCLELLVAI
TLNNRDRIELLWQGVYEHIANIVQSTVMPCALVEKAVFGLLRICHRLLPYKENITDELLRSLQLVLKLDARVADA
YYEQITQEVSHLMKANASHIRSHFGWRTITSLLSITARHLEAAEAGFDALMFIMSDGAHLLPANYVLCIDAARQF
AESRVGQVERSVMALDLMSGSVSCLEKWTNDAKQATKEEEVAKVLQDIGDMWLRLVQGLKKLCLDQREEVRNHSL
LSLQNCLTGSVAINLPHSLWLQCFDQVIFSVLDDLIEISQTQSQKDYRNIEGTLVLALKLLCKVFLQLIQYLSEL
PDFSILWLAVLSRLETYMKVKFRGRRSEKLHELVPELLKNTLLVMKTGQVLVRSSSEDGSSLWELTWLHINNIAP
SMQSEVFPEQDSELLQKKQTEKVEGLVSDENKSLSSNETEGQDGPGIS*
**>Phvul_001G235400**
MQMQTDLNAIEDQYGQCEAKYPNKTAIACMINAEISAVLAVMRRNVRWGVHYMLDDDQSEHFLVQSLKTLRRQIF
SWQNQWHTINPALYLQPFLDVIRSDETSAPITGVALSSVYKILTLDVIDQHTVNVGDSMHLVVDSITSCRFEVTD
PGSEEVVLMKILQVLLACVKGKASVMLSNQHICTIVNTCFRIVHQAGTKGELLQRIARYTMHELVRSIFSHLQNI
DNTKCGFINGSATLKQVNNGLNNEHALASRELENERLNSAQDAQPLSTITGSGTATVVTANVVDENTVIASSGNK
TEPHELQLMTDGVPCMVEIFHFLCSLLNVTEHMGVNPRSNTMTFDEDVPLFALTLINSAIELGGPSFRHHPGLLS
LIQDELFYNLMQYGLSVSPLVLSMVCSIVLNLYEHLRQELKLQLEAFFSCVILRLAQSKYGSSYQQQEVVMEALV
DFCRQKTFMVEMYTNFDCDISYSDVFEDIANLLSKSAFPVNNPLSSIHILALDGLIVVMQGIAERLCSGSLSSEQ
SPVNFEEYTPFWMEKCDSFADPNGWVPFFRRRKYIKRKLMIGADHFNRDIKKGLEFLQGAHLLPNKPDPKSVACF
LRYTAGLDKNLIGDFLGNHEEFYVQVLHEFAKTFDFHDMTLDTALRVLLESFRLPGESQKIHRVLEAFSERYYEQ
SPDILANKDAALVLSYSIILLNTDHHNVQVKKKMTEEDFIRNNRRINDGNDLPQEFLSEIYHSICKNEIRTTLEP
GFGFPEMTPSRWISLTHRANKTAPYIVSDSRAYLDYDMFVLLSGPTIAAISVVFDNAENKEVYQRCMDGFLVAAK
ISAYYHLENVLDDLVVCLCKFVTVLDPLSADDSVLAFGDDTKVRIATETVFTIANRYGDYIRTGWRNILDCILIF
YKLGLLPTRLASDAADESQVTTEIGHGKSNSNSLLSSNLQYSAPKRSSGLISRFSQLLSLGSEEIRPIPTKEQLA
AHQQATQAIQKCHIDSIFTESKFLQAESLLNLERALINAGAQHLKGNRMSEDEESSVFCLELLVAITLNNRDRVI
LLWKGVYDHISNIIQSTVMPCALVEKAIFGLLRICHRLLPYKENITDELLKSLQLVLKLDALVADAYYEQITQEV
GFLVKANASHIRSQSGWWTITALLSITSRHLESSEAGFDALLFIMSDGAHLLPANYILCVDAARQFAESRVGQVD
RSIIALDLMVSSVNSLERWSSNAKKAAKEEEVEKMLQDIGEMWFRLVQGLGKVCLDQREEVRNHALLSLQKCLKG
AVGTHLPCDLWLTCFDKVIFIVLEDLLEIAESHSQKVYPNMEGTLVIALKLMSEVFLQLLQELSQLAAFCELWEG
MLSCMEKCAKMKFKGRRSEKLQVLVPELLKNTLLVMKSRGILVQTNNGSGENSLWEVTWLYMKNIAPSLHSVVFP
EQDPEQLQHKQVKAVESLGPDANISDPSNEKDGQGVEIS*
**>Phvul_004G161900**
MDNVEDFMHDDENNKKGTSKSKRRQLGLSCMLNTEVSAVIAVVRRPECIPLYNISVTEDAYDSSLITSLRSLRSL
IFNPQQNWRTIDPSIYLIPFLDVVQSDDVPAAATGVALSAVLRILKFEVFDEKSPGAREGMESVVSGITSCRLEK
TDPASEDAVMMKILQVLTGIMNHRASTLLSDQSVCTLINTCFQVVQQSASRGDLLQRSARYTMHELIQVVFKRLP
EIEAKDREGDSDSDMEDGDEGGGLETGYGVRCAIDIFHFLCSLLNVVSIVEADGSTSHTADEDVQIFALVLINSA
IELSGDEIGKHPKLLRMIQDDLFHHLIYYGTWSSSFVLSMICSTVLNAFHFLRRSIRFQLEAFFSYVLFRVASFG
STIALQEVAIEGIINFCRQPTFIVEAYANFDCDPCCRNVFEEIGRLLCKHSFALNGHVTSLHIQAFEGLLIMIHN

IADNIDKDGSTGSLGAYSIQLTEYKPFWEEMDKEDDLEGWVEHVRLRRLQKKKLLIAANHFNRDNKKGLEYLKHA
KLISDPPDPAAHAHFFRYTPGIDKKTIGEFLGDPDPFYLIVLKEFTDTFHFQGMSLDTALRFYLESFLLPGESQK
IQRVLEAFAERFYDHQSSDMFASKDTVLILCYSLIMLNTDQHNPQVKKKMTEEEFIRNNRAINAGHDLPREYLSE
LFQSISTCAFSLSQTTVSLDMSPSRWIQLINRSKVVQPFTPCDYDRRICRDMFASIAGASVAALSSFFEHADEEE
LLHECIEGLFSVARICQYGLEDTLDELITSFCKFTTLLNPYASIEETMFSFSHDMKPRMATVAVFTVANYFRDSI
QGGWKNIVDCLLKLKRLKLLPQHVIDLEASVDVPTTPESGGVMSPTDDHKVGCQRVSSMMTRFSHLSSEGMDDGL
TLGSEFEQNTKMIKLCKIGSIFSNCSNIPRECLQSLGRSLIYAAAGKGQKFSTPVEEEETVEFCWDLIGAISLAN
VHRFQIFWPSFHDNLLSVAQFPMFSPIPFAEKAIVILLKVCVKLFSAPREDKQVEEFIFKSITLMWKLDKEILDT
CHDVISQSISRILIEYPANVQTQIGWKSMLNLLSVAWRHPETYDAGIEALIALFSDGTNISRSNYAYCIDCAFGC
FLAKNSPIDKKKKILDLLADSVNLLVQWQRSQYSDPGSNVSAASYTSSSSIEENSRGPSSGNYIMNLFVKLGEAF
RKTSLSRQEEIRNHAVCSLQKSFNLAEDLLFISSNCINYFNLVIFAMVDELHEKMLEYSKRENAEKETRSMEGTL
KLAMELFSDMYLQSLKQITESPGFRTFWLGILRRMDTCMRADLGEYGLSTLKEIIPDLLRKIITQMKEEGILEPR
EEDDMWEITYIQIQWICPRLKDELFPL*

**>Vigan_LOC108329051**

MGRLKLQAGINAIEEEEPEECDAAYPNKTTLACMINSEIGAVLAVMRRNVRWGGRYMSGDDQLEHSLIQSFKTVR
RQIFSWHHHQWQAINPALYLQPFLDVIRSDETGAPITGVALSSVYKILTLDVIDQNTVNVEDAMHLVVDAVTSCR
FEVTDPSSEEVVLMKILQVLLACMKSKASIMLSNQHVCTIVNTCFRIVHQAGSKGELLQQIARYTMHELVRCIFS
HLQDVGNTDHALVNGSTNLKQETGGLDNDYAFGSRQLENGSISSEYDNQSSSTNSAPNVSSVVKATVMDENTAIT
ISCKDGVPYDMHLMTEPYGVPCMVEIFHFLCSLLNVVEHTGMGPRSNTLAFDEDVPLFALTLINSAIELGGPSIC
RHPRLLSLIQDELFHNLMQFGLSMSPLILSMVCSIVLNLYHHLRTELKLQLEAFFSCVILRLAQSRYGASYQQQE
VAMEALVDFCRQKTFMVDMYANFDCDITCSNVFEDLANLLSKSAFPVNCPLSAMHILALDGLIAVIQGMAERIAN
GSVSSEYSPVNLEEYTPFWMVKCENYNDPNHWVPFVRRRKYIKRRLMIGADHFNRDPKKGLEFLQGTHLLPDKLD
PQSVACFFRYTAGLDKNLVGDFLGNHDEFCVQVLHEFAGTFDFQDMNLDTALRLFLETFRLPGESQKIHRVLEAF
SERYYEQSPHILANKDAALVLSYSMIMLNTDQHNVQVKKKMTEEDFIRNNRLINGGNDLPREMLSEIYHSICKNE
IRTTPEQGVGFPEMTPSRWIDLMHKSKKTAPFIVSDSKAYLDHDMFAIMSGPTIAAISVVFDHAEQEEVYQTCMD
GFLAIAKISACHHLEDVLDDLVVSLCKFTTLLNPSSVEEPVLAFGDDMKARMATVTVFTIANRYGDYIRTGWRNI
LDCILRLHKLGLLPARVASDAADESELSAETVHGKPIMNSLSSAHMQSIGTPRRSSGLMGRFSQLLSLDTEEPRS
QPTEQQLAAHQRTLQTIQKCHIDSIFTESKFLQAESLLQLARALVWAAGRPQKGSSTPEDEDTAVFCLELLIAIT
LNNRDRIGILWHGVYEHISNIVQSTLMPCALVEKAVFGLLRICQRLLPYKENIADELLRSLQLVLKLDARVADAY
CEQITQEVSRLVKANASHIRSQLGWRTITSLLSITARHIEASEAGFDALLFIMSDGAHLLPANYVLCVDTARQFA
ESRVGQAERSVRALDLMAGSVNCLARWTSEAKESMEEEQVSKLSQDIGEMWLRLVQGLRKVCLDQREEVRNHALL
SLQKCLTGADDIYLPYSMWLQCFDLVIFTVLDDLLEIAQGHSQKDYRNMEGTLILAMKLLFKVFLQLLPELSQLT
TFCKLWLGVLSRMEKYMKVKVRGKRSEKLQETVPELLKNSLLVMKMRGILAQRSALGGDSLWELTWLHVNNISPS
LQLEVFPEQDSEHLPHKQGEPIGGLVTDDKGSVPSSETASREDAGVVG*

**>Vigan_LOC108338084**

MGHVKLQTQTGINAIEEASGQCDAEYPDKTTVACMINAEIGAVLAVMRRNVRWGVHYMSDDDQLEHSLVQSLKAL
RKQIFSWQNQWHVINPALYLQPFLDVIQSDETGAPITGVALSSVYKILTLDVIDQNTVNVGDAMHLVVDAVTSCR
FEVTDPGSEEVVLMKILQVLLACVKSKASMMLSNQHICTIVNTCFRIVHQAGTKSELLQRIARYTMHELVRCIFS
HLQDIDNAELALNNGSTALQKEIVGINNDHNSPNRQLDNGNLTSASNGQPLSTGIAPSTVSDVAATVLDEDTAIA
SSDQVADLNELQLVNEPYGIPCMVEIFHFLCSLLNVAEHMGNSPRSNTIAFDEDVPLFALTLINSAIELGGPSFH
QHPRLLSLIQDELFRNLMQFGLSMSPLVLSMVCSIVLNLYHHLRTELKLQLEAFFSCVILRLAQSKYGASYQQQE
VVMEALVDFCRQKTFMVEMYANFDCDLTCSNVFEDIANLLSKSAFPVNSPLTSMHILALEGLIAVMHGMAERIGN
GSLSSEQSPVNLEEYTPFWQEKCENFCDPKNWVPFVSRRKYFKKRLMIGADHFNRDTKKGLEFLQGTHLLPDKLD
PQSVACFFRYTAGLDKNLIGDFLGNHDEFCIEVLHEFAKTFDFKDMMLDTALRLFLETFRLPGESQKIQRVLEAF
SERYYEQSPEVLANKDAALLLSYSIIMLNTDHHNSQVRNKMTEEDFIRNNRRINGGNDLPRDFLSELYHSICKNE
IRTTPEQGSGFPEMTPSRWIYLKHKSKKSDPFIVSDSKAYLDYDMFSILSGPTIAAISVVFDNAENAEVYQTCMD
GFIAVAKISAYYHLENILDDLVVSLCKFVTVFDPLAVEESILAFGEDMKARMATETVFTIANRYGDFIRTGWRNI
LDCILKFHRLGLLPTRIASDAAEESELSTETEDGGKRTTNALSLSRLPSVNTTKRSSGLMSRFSQLLYLGAEEPK
SVPSEEQLAAQQCTLQTIQKCHIDSIFSESKFLQAESLLQLAKAITSAGVRPKKGNSTSEDEDTLVFCLELLVAI
TLNNRDRIELLWQGVYEHISNIVQSTVMPCALVEKAVFGLLRICHRLLPYKENITDELLRSLQLVLKLDARVADA
YYEQITQEVSHLMKANASHIRSHLGWRTITSLLSITARHLEAAEAGFDALLFIMSDGAHLLPANYVLCIDAAKQF
AESRVGQVERSVMALDLMSGSVSCLEKWTNDAKQATKEEELAKVLQDIGDMWLRLVQGLKKLCLDQREEVRNHAL
LSLQKCLTGSVGINLPHNLWLQCFDQVIFSVLDDLIEISQTQSQKDYRNIEGTLVLALKLLCKVFLQLIQFLSEL
PDFSTLWLAVLSRLETYMKVKIRGRRSEKLHELVPELLKNTVLVMKTGQVLVRSSSVDGSSLWELTWLHINNISP
SMQAEVFPESDSKDLQKKQTEKVEGLVSDENNSVSSNETAGQDGPGIS*

**>Vigan_LOC108329296**

MQMQTDFNAVEDQYELCEAGYPNKTAIACMINAEISAVLAVMRRNVRWGIHYISDDDQSEHYLVQSLKTLRRQIF
SWKNQWRAINPALYLQPFLDVIRSDETSAPITGVALSSVYKILTLDVIDKHTVKVGDTMHLVVDSVTSCRFEVTD
PGSEEVVLMKILQVLLACVKGKASVMLSNQHVCTIVNTCFRIVHQAGTKGELFQRIARYTMHELVRSIFSHLQNI
DNTKCGFINGSATLKQVTNGLNNDHALASRKLENERLNSARDAQPLSTITASGAAPVVAANAVDENTAIASSGNE
TDPHELLLMTHGVPCIVEIFHFLCSLLNVTEHMGVNPRSNTMTFDEDVPLCSLTLINSAIELGGPSFCHHPRLLC
LIQDELFFNLMQFGLSMSPLVVSMVCSIVLNLYHHLRQELKLQLEAFFSCIILRLAQSKYGASYQQQEVVMEALV
DFCRQKTFMVEMYANFDCDISCSDIFEDIVDLLSKSAFPVNNPLSSMHILSLDGLIAVMQGISERIGSESLSSKQ

SPVNFEEYTPFWMEKCDSFDDPNDWVPFVCQRKYIKRKLMIGADHFNRDIKKGLEFLQGAHLLPNKPDSQSVACF
LRYTAGLDKNSIGDFLGDHDEFYIQVLHEFAKTFDFHDMTLDTALRVFLETFRLPGESQKIHRVLEAFSERYYEQ
SPDILANKDAALVLSYSIILLNTDHHNVQVKKKMTEEDFIRNNRHINGGNDLPREFLSEIYHSICKSEIRTTLEP
GFGFPEMTPSRWISLMHEANKTSPYIVSDSRAYLDYDMFVLLSGPTIVAISVAFDNAENKEVYQRCMDGFLVAAK
ISAHYRLENVLDDLVVCICKFITILDPLSVEDSVLALGDDTKVRIATETVFTIANRYGDYIRTGWRSILDCILIF
YKLGLLPPRLASDAADESQVTTEIGHGKSNSYSIVSSNLQYIAPKRSSGLISRFSQLLSLGSEDIQSIPTKEQLA
AHQQATQAIQKCHIDSIFTESKFLQAESLLNLERALINAGAQHLKGNRIREDEESSVFCLELLVAITLNNRDRVV
LLWKGVYEHISNIIQSTVIPCALVEKAIFGLLRICHRLLPYKENITDELLRSLQLVLKLDALVADAYYEQITREV
GCLVKANASRIRSQSGWRTITALLSITSRHLESSEAGFDALLFIMSDGAHLHPANYILCVDAARQFAESRVGQVD
RSIIALDLMASSVNSLEKWSSNAKKTAKQQEVEKMLQDIGEMWFTLVQGLSKVSLDQREEVRNHALLCLQKCLKG
TVGTHLPRDLWLTCFDQVIFTVLEDLLEIAQLHSQKVYPNIEGTLVIALKLVFEVFLQLLLELSQLASFCKLWEG
VLSCMEKCVKMKIKGRRSEELQELVPELLKYTLLVMKSGDILVQINNGSGENSLWELTWLHMKNIAPSLQSEVFP
EQDPEQLQHRQVKAVESLGSDANISDPSNEKEGNDVVEIS*
**>Vigan_10g017100**
MDNVQDFMHDVVNKKGTTKSRRRQLGLSCMLNTEVSAVIAVVRRPECIPLYNLSAPDETYDSSLISSLRSLRSLI
FNPQQEWRTVDPTIYLTPFLDVIISDDVPAAATGVALSAVLKILKFEIFDEKTSGAREGIESVVSGITSCRLEKT
DPVSEDAVMMKILQVLTGIMNHRASDLLSDQSVCTLINTCFQVVQQSASRGDLLQRNARYTMHELIQAVFKRLPE
IEAKDREGDSESDMEDGDEGGGLESGYGIRCAIDIFHFLCSLLNVVSIVEADGSTTHTADEDVQIFALVLINSAV
ELSGDEIGKHPKLLRMIQDDLFHHLIYYGTWSSSFVLSMICSTVLNAFHFLRRSIRFQLEAFFSYVLFRVASFGS
TIALQEVAIEGIINFCRQPTFIVEAYANFDCDPSCRNVFEEIGRLLCKHSFALNGHVTSLHIQAFEGLLIMIHNI
ADNIDKDGSSGSVGAYSIQLPEYRPFWEEMDREKYLEAWVEHVRLRRLQKKKLLIAANHFNRDNKKGLEYLKHAK
LISDPPDPKAHGHFFRYTPGIDKKTIGEFLGDPDPFYLKVLKEFTDTFYFQGMSLDTALRFYLESFLLPGESQKI
QRILEAFAERFYDHQSSDMFASKDTVLILCYSLIMLNTDQHNPQVKKKMTEEEFIKNNRAINAGQDLPREYLSAL
FQSISTCAFSLSQTTVSLDMSPSRWIQLINRSKVAQPFTPCDYDRRICRDMFACIAGPSVAALSSFFEHADEEEL
LHECIEGLFSVARICQYGLEDTLDELITSFCKFTTLLNPYASIEETMFSFSHDIKPRMATVAVFTVANYFRDSIQ
GGWKNIVDCLLKLKRLKLLTQPVIDFDVVAATTPESGGAMSPTDGHKVGYQRIASMITRFSNLSSEGMEDGLTLG
SEFEQNSKMIKLCKIGSIFSNCSNIPKECLQSLGRSLIFAAAGKGQKFSTPVEEEETVEFCWDLIGAISVANVHR
FQVFWPSFHEYLLSVAQFPMFSPIPFAEKAMLILLKVCLKLFSAPREDKLVEEFIFKSITLMWKLDKEILDTCHD
VIAQTISKVLIEYPANVQTQIGWKSVLNLLSVAWRHHDTYDVGIEALIALFSDYAYISRTNYAYCIDCAFGCFLA
KNSPIDKKKKILDLLADSVTLLIQWHRGQYSDPGTNASASSYNSSSSIDENSRAPSSGNYNMNLFVKLGEAFRRT
SLSRQEEIRNHAVYSLQKSFNLAEDLLFISSNCINYFNLVIFAMVDELHEKMLEYSRRENAERETRSMEGTLKLA
MEQLSDMYLQSLKQITESPAFRTFWLGILRRMDTCMKADLGQYGASTLSETIPDLLRKIITQMKEEGILEPREDD
DMWEITYIQIQWICPPLKDELFPL*
**>Arahy_LOC112733642**
MGRLKLQAGINAIEEEEPEDCDATCPNKTTLSCMINSEIGAVLAVMRRNVRWGSSYMSGDDQMEHSLIQSLKALR
RQIFSWHHQWHAINPTLYLQPFLDVIRSDETGAPITGVALSSVYKILTLDVIDQNTVNVEDAMHLVVDAVTSCRF
EVTDPSSEEVVLMKILQVLLACMKSKASVMLSNQHVCTIVNTCFRIVHQAGTKGELLQRIARHTMHELVRCIFSH
LQDVDNTDHALVNGSSNLKQETGGLNNEYAFGSRQLENGSLSSEYDNQTLPTNYAPSAASVATGTRMDENTAIAI
SGKDGVPYDMHLMTEPYGVPCMVEIFHFLCSLLNVVEHMGVGPRSNTIAFDEDVPLFALTLINSAIELGGPSIRC
HPRLLGLIQDELFRNLMQFGLSMSPLILSMVCSVVLNLYNHLRTELKLQLEAFFSCVILRLAQGRYGASYQQQEV
AMEALVDFCRQKTFMVDMYANFDCDITCSNVFEDLANLLSKSAFPVNCPLSAMHILALDGLIAVIQGMAERIDNG
SVSSEYSPVNLEEYNAFWMVKCENYGDPNHWVPFVRRRKYIKRRLMIGADHFNRDPKKGLEFLQGTHLLPDKLDP
QSVACFFRYTAGLDKNLVGDFLGNHDEFCVQVLHEFAGTFDFQDMNLDTALRLFLETFRLPGESQKIHRVLEAFS
ERYYEQSPHILANKDAALVLSYSMIMLNTDQHNVQVKKKMTEEDFIRNNRHINGGNDLPREFLSEIYHSICKNEI
RTTPEQGVGFPEMTPSRWIDLMHKSKKTAPFIVSDTKAYLDHDMFAIMSGPTIAAISVVFDHAEHEEVYQTCTDG
FLAVAKISACHHLEDVLDDLVVSLCKFTTLLNPSSVEEPVLAFGDDMKARMATVTVFTIANRYGDYIRTGWRNIL
DCILRLHKLGLLPARVASDAADESELSSETVHGKPVTNSLSSAHMPSIGTPRRSSGLMGRFSQLLSLDTEEPRSQ
PTEQQLAAHQRTLQTIQKCHIDSIFTESKFLQAESLLQLARALIWAAGRPQKGNSTPEDEDTAVFCLELLIAITL
NNRDRIGILWQGVYEHISNIVQSTVMPCALVEKAVFGLLRICQRLLPYKENIADELLRSLQLVLKLDARVGDAYC
EQITQEVSRLVKANATHIRSQLGWRTITLLLTNTSSHIEASEAGFDALLFIMADGAHLLPANYAFCLDTARRFAE
SRVGQAERSIRALDVMAGSVNCLARWTSEAKEAQDEEQAAKMLQEFGEMWLRLVQGLRKVCLDQREDVRNHALLC
LQNCLTGADGIYVPHGRVLQCFDIVIFTLLDDLLEIAQGHSQKEYRNMEGTLILAMKFLSKVFLQLLPDLSQLST
FCKLWLGVLSRMEKYMKVKIRGKRSEKLQETVPELLKNSLLVMKMKGILVQRSALGGDSLWELTWLHVNNISPSL
QLEVFPEQDSDHLQKKQGEAVGGLVPDEMGSVPSSETESLEDAGVAG*
**>Arahy_LOC112791591**
MGHLKMQMETGINAIEDQSGYTNRNSIACMINAEIGAVLAVMRRNVRWGVHYMADDDQLEHSLVKSLKELRRQIF
SWQGQWHSINPVLYLQPFLDVIKSDETGAPITGVALSSVYKILTLDVIDQNTVNIEDAMHLVVDAITSCRFEVTD
PGSEEVALMKILQVLLACVKCKASTTLSNQHICTIVNSCFHIVHQAGTKSELLQRIARYTMHELVRCIFSHLQDI
ENTEHLLINERSPLKQETSGQNNESTVFSRLLENGSQNSSYDGEPLGISSGIASAVTSTVMDENSAFASNGNETD
PHELQLMSEPYGVPCMVEIFHFLCSLLNVVEHMGMNPRSNTIAFDEDVPLFALTLINSAIELGGPFFRCHPRLLS
LIQDELFRNLMQFGLSMSPLVLSMVCSIVLNLYHHLRKELKLQLEAFFSCVILRLAQSKYGASYQQQEVAMEALV
DFCRQKTFMVEMYANYDCDITCSNVFEDITNLLSKSAFPVNNPLSSMHILALDGLIAVMQGMAERIGNGYVSSER
SPVNLEQYIPFWLEKCENYGDPNDWVPFVFRRKYFKQRLMIGADHFNRDVKKGLEFLQGTHLLPDKLDPQSVACF

LRYTAGLDKNLIGDYLGNHDEFCVQVLHQFARTFDFQGITLDKALRIFLESFRLPGESQKIHRVLEAFSERYFEQ
SPDVLVNKDAALVLSYSIIMLNTDHHNVQVKKKMTEEDFIRNNRLINGGKDLPREFLSEIYHSICKNEIRTTPEP
GFGFPEMSPSRWISLMHKSRKTAPFIVSDSRADLDYDMFAILSGPAIAAISVVFDNAENEEIYQTCIDGFLAVAK
ISAHYHLANVIDDLVVCLCKFITILDPFSVEESVQALGDDTKVRMALETVFAIANSYGDYIRAAGWRNILDCILR
LHKLGLLPTRLVSDAAEESELSAESGHGKPNSNLSSSTQLPSVSTPKRSSGLMSRFSQLLYLGTEEPRSVPTAEE
LVAHQQAVQIIQKCHIDTVFTESKFLQAESLLLLARALINAGGQPRKGNSISGDEDTSVFCLELLVAITLNNRDR
IGLLWKDVYEHISNIVQSTVTPCILIEKAVFGLLRICHRLLPYKESIADELLRSLQLVLKLDARVADIYYEQITQ
EVSRLVKANASHVRSQLGWRTITSLLSFTARHQESSEAGFDTLLFIMSDGAHLLPANYVLCVDTARQFAESRLGQ
VERSVVALDLMAGSVNCLEKWTSDAKQAAKDEEVEKMLKDIVEMWLRLVQGIKKVCLDPREEVRNHALLSLQTCL
TGAVGIHVPPDLWLQCFDQVIFTAVDDLLEIAEEYYQKEYRNMETLILVLKLLSKVFLQSLSKLSQLTDFSKLWV
AVLDRTEKYTKAKIRGRRSEKLQELVPELLKNTLLVMKSQGILLPSSGAGENSLWELTWLHMKNIAPSLQSEVFP
APELQQALHKQIEAVDANTSVSSNETVSQDGAGVGS*

**>Arahy_LOC112765119**

MGHVKLLSQTGINSIEEEKGHCNDGYPNKTTVACMINAEIGAVLAVMRRNVRWGVHYMSDEDQVEHSLVQSLKIL
RRTIFYWKDPWHAINPVRYLQPFLDVIQSDETGAPITGVALSSVYKILTLDVIDQNTTNVGDAMHLVVDAVTSCR
FEVTDPGSEEVVLMKILQVLLACVKSRASVMLSNQHICIIVNTCFRIVHQAGTKGELLQRIARYTMHELVRCIFS
HLKDIDNVEHSLVNGSNGLNQETGGQNNEHNSTNRQLENGSLNSASNRQSYSTGIASSTESVLTVGVVDDITAID
SSGKEVDLDELDLMKEPYGIPCMVEVFHFLCSLLNVVEHVGTSTRLNTIAFDEDVPLFALTLINSAVELGGSSFH
RHPRLLSLIQDELFHNLMEAGLSMSPLVVSMVCSIVLNLYHHLRTEIKLQLEAFISCVILRLAQSKYGASYQQQE
VAMEALVDFCRQKTFMMEMYANLDCDITCSNVFEDIANLLSKSAFPVNSPLSSLNILALDGLIALMQGMAERIGN
ESLSSEHPHVNLEQYSSFWQEKCENFSDPNNWVPFVCRRKYFKKILMIGADHFNRDVKKGLEFLQGTHLLPDQPH
AQSVAYFLRFTTGLDKNLIGDFLGNHDEFCVQVLQEFARTFDFKDTTLDTALRLFLETFRLPGESQKIQRVLEAF
SERYYEQSPQILANKDAALLLSYSIIMLNTDHHNVQVKKKMTEEDFIRNNRRINGGNDLPRGFLSEIYHSICRNE
IRTTPEQGSGFPEMTPSRWIYLIHKSRKTAPFIVSDSRGYLDNDMFAILSGPTIAAISVVFDNAENEEVYQTCMD
GFLAVAKISAYYHLENVLDDLVVSLCRFVTILDPLSVEEPILVFAYDTKARMATNTVFTIANKYGDYIRTGWRNI
LDCILRFHKLGLLPANMANDATEESEHPTETLHEKRNAKSLSSRSLRSISTPKRSSGLMSRFSQLLLLGSEEPRP
IPTKEQVAAQQSTLQTIQKCHIDSIFSESKFLQAESLLQLARALITAGGRLKKGNITFDDEDSSVFCLELLITIT
LNNRDRIELLWHGVYEHISNIVHLTVMPCALVEKAVFGLLKVCHRLLPYKENFTDELLRSLQLVLKLDARVADAY
YEKITQEVSHLVKANASHIRSQLGWRTITSLLYITARHLEASDAGFDALLFIMSDGAHLLPSNYVLCIDVARQFA
ESRVGQVERSVVALDLMVGSANCLEKWTSDAKKEMKEEDVAKMLQDIGEMWLRLVQELKEVCLDQREEVRNHALL
SLQNCLTGSLGTHIQHDLWLRCFDQVIFAVLDELVETYQTRSQKEYRNMEGSLIIALKLLCKVYLQLLPSISEST
KFTKLWLGVLSRLEVYMTVRVRGRRSEKIQELVPELLKNTLLVMKAHDILVRSSSGGDNSFSELTWKHINNISPS
LQSEVFPEQDSEQLENKQAKTVEDLGPNNENISIPSNERAGQDGAGIG*

**>Arahy_LOC112749204**

MAKSIKRLSAMDFLNGEEDYNHQHNQRSKSISRSKRKQLALSCMLNTEVGAVLAVIRRSPEFNPLYSSPEDTYDS
SIVTSLRSLRSLIFNPTQEWRSIDPSIYLSPFLDVILNDDMPAAATAVALSSILKILKFEVFDERTPGAREGMEA
IVTAITGCRLEKTDPVSEDAVMMKILQVLAGTMHHRASILLSDPAVCTLVNACFQVVQQSVNRGDLLQRNARYTM
HELIQVVFSRLPELEKKGGRRHLRSGEYSGSETDDDDDDDEDDDGDGGEVTGYGIKCAIDIFHFLCSLLNVVSVI
ENPDGTLPHTADENVQIFALLMINSALELSGDVIGRHPRLLRMIQDDLFHHLIYYGSWSSSFVLSMICSTVLNAY
HFLRRFLRFQLEAFFLNVLTRLASFGSSISIQEVAGEGIINFCRQPTFIIEAYVNFDCDPCCRNVFEDFGRLLCK
HSFAGSGSSTSMQIQAFEGLLILIHHIADNIDKDDSGPLGAYAVQLNEYIPFWEDMERDEDDLDTWVEHVRMRKY
QKKKLLIAANHFNRDNKKGLEYLKHAKLISDPPDPKGYAYFFRYTPGLDKKIIGEYLGDPDEFYLKVLKEFTETF
HFNGMVLDTALRFYLESFLLPGESQKIQRILEAFSDRFYDQQSSDLFASKDTVLILCYSLIMLNTDQHNPQVKKK
MTEEEFIRNNRAINAGQDLPRDYLSELFQSISSYAIALDTSTVSLDMTPSRWIQLINRSKVSQNYIQCDYDRRLC
RDMFACIAGPSVAALSSFFEHNDEEELLHECIEGLFSVARIAQYGLEDTLDELITSFCKFTTLQNPYASPEETLY
TFGHDLKPRLATVAVFTIANMFRNGIRGGWKNIVDCLIKLKRLKLIPQSVLDLEQSADAATTPESATSPTEDHRF
SSHQMGTIMQRFTLLSPDNTDDGINLSNESDRNIKMIKLCQIGSIFSTCSNIPIESLHSLGRSLISAAAGKGQKY
STQHDEEETIEFCWELLSAVATVNAHRFHMFWPNFHEYLLAVVQFQMFSPVPFAEKALLGLLKICLKLFSQPVDD
KISEEAIFKSITLMWKLDKEILDTCHDIISQTASRIVTDYAANLQTPFGWKSLLNLLSVAWRYPETYDLGVDALV
TLFSDGTHVSRTNYAFCIDCAFGCFLAKNSPADKKKKMLDILADSVNLLIHWHRTLYSDPGSNFSISSSTSSSSN
EETSRSLANYNMNLFVKLGEAFRRTTLSRQEEIRNHAVHSLHKSFLLAEELLFISSNCINYFNLVIFAMVDELHE
KMLEYSRREGAEKEMRSMEGTLKLAMELFTDMFLQSLRQIAESPGFRTFWLGILRRMDTCMKADLGHYGPSSLSE
IIPHLLKKIITQMKEQEILVPKEDDDMWEITYIQIQWICPAIKDDLFPM*

**>Abrpr_LOC113846820**

MGRLKLQAGINAIEEETEECDVAYPNKTTLACMINSEIGAVLAVMRRNVRWGGRYMSGDDQLEHSLVQSLKSLRR
QVFSWQHPWHTINPALYLQPFLDVIQSDETGAPITGVALSSIYKILTLDVIDQNTVNVEDAMHLVVDAVTSCRFE
VTDPASEEVVLMKILQVLMACMKSKASIMLSNQHVCTIVNTCFRIVHQAGTKGELLQQIARYTMHELVRCIFSHL
QDIDNTQHALVNGNSTLKQETGDLNNEYAFGNRQLENGSMSTEHVNQPLSTNFSPNASVVTTTVVDENTAIAING
KEAVPYDMHLMTEPYGVPCMVEIFHFLCSLLNVVEHIGMGPRSNTIAFDEDVPLFALTLINSAIELGGPSICSHP
RLLSLIQDELFRNLIQFGLSMSPLILSMVCSIVLNLYHNLRTELKLQIEAFFSCVILRLAQSRYGASYQQQEVAM
EALVDFCRQKTFMVDMYANFDCDITCSNVFEDLANLLSKSAFPVNCPLSAMHILALDGLIAVIQGMAERIGNGSL
SSEHSPVNLEEYTPFWMVKCENYSDPNHWVPFVRRRKYIKRRLMIGADHFNRDPKKGLEFLQGTHLLPDKLDPQS
VACFFRYTAGLDKNLVGDFLGNHDEFCVQVLHEFAGTFDFQDMNLDTALRLFLETFRLPGESQKIQRVLEAFSER

YYEQSPHILANKDAALVLSYSIIMLNTDQHNVQVKKKMTEEDFIRNNRHINGGNDLPREFLSELYHSICKNEIRT
TPEQGVGFPEMTPSRWIDLMHKLKKTAPFIVSDSKAYLDHDMFAIMSGPTIAAISVVFDHAEHEEVYQTCMDGFL
AVAKISACHHLEDVLDDLVVSLCKFTTLLNPSSVEEPVLAFGDDMKARMATVTVFTIANRYGDYIRTGWRNILDC
ILRLHKLGLLPARVASDAADESELSAETGHGKPIGNALSSAHMPPIGTPRRSSGLMGRFSQLLSLDTEEPRSQPT
EQQLAAHQRTLQTIQKCHIDSIFTESKFLQAESLLQLARALIWGARRSQKGNSTPEDEDTAVFCLELLIAITLNN
RDRIVLLWQGVYEHISNIVQSTVMPCALVDKAVFGLLRICQRLLPYKENIADELLRSLQLVLKLDARVADAYCAQ
ITLEVSRLLKANASHIRSQLGWRTITSLLSITAKHLEAYEAGFDALLFIMSDEAHLMPANYVLCVDTARQFAESR
VGQAERSVRALDLMAGSVNCLVQWTSEAKEAIEQEQVSRLFQDIGEMWLRLVQGLRKVCLDQREEVRNHALLSLQ
RCLTGADGIDLPHGLWLQCFDQVIFTVLDDLFEIAQGHSQKDYRNMEGTLILAMKLMSKVFLQLLQDLSQLTTFC
KLWLGVLSRMEKYTKVKVRGKRSEKLQELVPELLKNILLVMKTKVILVQRSALGGDSLWELTWLHVNNIAPSLQA
EVFPDQDFDHLQHKQVESSVGNLMPDEVNYTPSNETENREDAGIVG*

**>Abrpr_LOC113849046**

MGRLKLQAGINAIEEEEPEECDPAYPNKTTLACMINSEIGAVLAVMRRNVRWGGRYISGDDQLEHSLIQSFKTVR
RQIFSWHHHQWQAINPALYLQPFLDVIRSDETGAAITGVALSSVYKIMTLDVIDQNTVNVEDAMRLVVDAVTSCR
FEVTDPSSEEVVLMKILQVLLACMKSKASIMLSNQHVCTIVNTCFRIVHQAGSKGELLQQIARHTMHELVRCIFS
HLQDVDNTDHALVNGSSNLKQETGGLNNEYAFGSRQLENGSMNSEFDNQPLATNFAPNATSVVKATVMDENTALA
ISGKEGVPYDMHLMTEPYGVPCMVEIFHFLCSLLNVVEHTGMGPRSNTLSFDEDVPLFALTLINSAIELGGPSIC
RHPRLLSLIQDELFHNLMQFGLSMSPLILSTVCSIVLNLYHHLRTELKLQLEAFFSCVILRLAQSRYGASYQQQE
VAMEALVDFCRQKTFMVDMYANFDCDITCSNVFEDLANLLSKSAFPVNCPLSTMHILALDGLIAVIQGMAERIAN
GSVSSEYSPVNLEEYTPFWMVKCENYNDPNNWVPFVRRRKYIKRRLMIGADHFNRDPKKGLEFLQGTHLLPDKLD
PQSVACFFRYTAGLDKNLVGDFLGNHDEFCVQVLHEFAGTFDFQDMNLDTALRLFLETFRLPGESQKIHRVLEAF
SERYYEQSPHILANKDAALVLSYSMIMLNTDQHNVQVKKKMTEEDFIRNNRHINGGNDLPREFLSEIYHSICKNE
IRTTPEQGFGFPEMTPSRWIDLMHKSKKTAPFILSDSKAYLDHDMFAIMSGPTIAAISVVFDHAEQEEVYQTCMD
GFLAIAKISACHHLEDVLDDLVVSLCKFTTLLNPSSVEEPVLAFGDDMKARMATVTVFTIANRYGDYIRTGWRNI
LDCILRLHKLGLLPARVASDAADESELSAETVHGKPIMNSLSSAHMQSIGTPRRSSGLMGRFSQLLSLDTEEPRS
QPTEQQLAAHQRTLQTIQKCHIDSIFTESKFLQAESLLQLARALIWAGRPQKGNSTPEDEDTAVFCLELLIAITL
NNRDRIGILWQGVYEHISNIVQSTVMPCALVEKAVFGLLRICQRLLPYKENVADELLRSLQLVLKLDARVADAYC
EQITQEVSRLVKANASHIRSQLGWRTITSLLSITARHIEASEAGFDALLFIMSDGAHLLPANYVLCVDTARQFAE
SRVGQAERSVRALDLMAGSVNCLVRWTSEAKETIEEEQVSKLSQDIGEMWLRLVQGLRKVCLDQREEVRNHALLS
LQKCLTGADGIYLPYGLWLQCFDLVIFTVLDDLLEIAQGHSQKDYRNMEGTLILAMKLLSKVFLQLLPELSQLTT
FCKLWLGVLSRMEKYMKVKVRGKRSDKLQEIVPELLKNSLHAMKMKGILAQRSALGGDSLWELTWLHVNNISPSL
QLEVFPEHKQGESIGVLRPDEIGSVPSSETASREDAGIVG*

**>Abrpr_LOC113846103**

MGNVKLQMQTSVNAVEEEPGQCDAGFPNKTTVACMINAEIGAVLAVMRRNVRWGVHYMSDDDQLEHSLVQSLKAL
RRQIFSWQNQWNVINPALYLQPFLDVIQSDETGAPITGVALSSVYKILTLDVIDQNTVNVGNAMHLVVDAVTSCR
FEVTDPGSEEVVLMKILQVLLACVKSKASVMLSNQHICTIVNTCFRIVHQAGTKSELLQRIARYTMHELVRCIFS
HLQDIGNPECVLVNGSTALKQETFGLNNEHNSANRQLENGSLNSASDGQLLSTGFASSTVTNVTTTVVDEGTAIP
SSGKDTELNELQLMNEPYGIRCMVEIFHFLCSLLNVVEHMGMSPRSNTIAFDEDVPLFALTLINSAIELGGPSFC
RHPTLLSLIQDELFRNLMQFGLSMSPLVLSMVCSIVLNLYHHLCTEVKLQLEAFFSCVILRLAQSKYGASYQQQE
VAMEALVDFCRQKTFMVEMYANFDCDITCSNVFEDIANLLSKSAFPVNSPLSSMHILALDGLIAVMQGMAERIGN
GSLSSEQSPVNLGEYTPFWLENCDNFSDPNNWVPFVCRRKYFKKRLMIGADHFNRDIKKGLEFLQGAHLLPDKLD
PQSVAFFFRYTAGLDKNLIGDFLGNHDEFCVQVLHEFARTFDFKDMTLDTALRIFLETFRLPGESQKIQRVLEAF
SERYYEQSPHILANKDAALLLSYSIIMLNTDQHNMQVKKKMTEEDFIRNNRRINGGNDLPREFLSELYHSICKNE
IRTTPEQGSGFPEMTASRWIYLMHKSKKSAPFIVSDCRSYLDYDMFSILSGPTIAAISVVFDNAENAEVYQTCMD
GFLAVAKISAYYHLENILDDLVVSLCKFVTIFDPLSVEESILAFADDMKARMATETVFTIANRYGDYIRTGWRNI
LDCILKFHKLGLLPARMASDAAEESELSTETEHGGKSNANSLSLSQLPSVSPPKRSSGLMSRFSQLLYLGTEEPR
LVPTEEQLAAQQCTLQTIQKCHIDSIFTESKFLQAESLMQLAKALTSAGVRPKKGSSASEDEDTLVFCLELLVAV
TLNNRDRIELLWLCVYEHISNIVQSTVMPCALVEKAVFGLLRICHRLLPYKENIADELLRSLQLVLKLDARVADA
YYEQITQEVNHLVKANASHIRSQLGWRTITSLLSITARHLEASEAGFDALLFIMSDGAHLLPANYVLCVDAARQF
AESRVGQVERSVVALDLMSGSINCLLKWTNDAKQATKEEEVAKMLQDIGDMWLRLVQGLKKLCLDQREEIRNRAF
VSLQNCLTGSVGIHLPHGLWLQCFDQVIFPVLDDLIEVSQTHSQKDYRNMEGTLILALKLLCKVFLQLIQDLSQL
TGFSKLWLGVLSRLEVYMKVKVRGRRSEKLQELVPELLKNILLVMKAGRVLVWSSSVDGNSLWELTWLHINNIAP
SLQAEVFPEQDSELLQHDQTEKDGGLGQDGNVTVPSNETLGQAGPGIG*

**>Abrpr_LOC113866795**

MDNVEDFMNGYENKKYSNKTKRKQLGLSCMLNTEVGAVLAVIRRPEFNPLYNISSSEESNDSSLVSSLRSLRSLI
FNPQQEWRTVDPSIYISPFLDVIQSDDVPATATGVALSSILKILKFGVFDHKSPCAREVMESVVSGITSCRLEKT
DPASEEAVMMKILQLLSAIMNHRASTLLTDQAVCTLVNTCFQVVQQSASRGDLLQRSARYTMHELIQAVFTRLAE
IETREREEDSESDMEDEDEGMEYGYGVRCAIDIFHFLCSLLNVVSIVDSDGSTSHTADEDVQIFALVLINSAIEL
SGDEIGRHPKLLRMIQDDLFHHLIYYGIWSSSFVLSMICSTVLNAYHFLRSFIRFQLEAFFGYVLLRIATSGSTV
PLQEVAIEGIINFCRQPTFIVEIYANSDCDPCCRNVFEDVGRLLCKHSFPMNGHLTSLNIQAFEGLLIVIHNIAD
NIDKDGSTGPLGPYSIQLAEYKPFWEEIDREDDLEAWVEHVRMRRMQKKKLLIAANHFNRDNKKGLQYLKHAKLV
SDPPDPKAYAYFFRYTPGLDKKTIGEYLGDPDPFYLQVLKEFTETFHFQGMVLDTALRFYLESFRLPGESQKIQR
VLEAFAERFFDHQSSDIFASKDTVLILCYSLIMLNTDQHNPQVKKKMTEEEFIRNNRAINAGQDLPREYLSELFQ

SISNYAIALAQTTVSLDMTPSRWIQLINRSKVTSPFTQCDYDRRMCRDMFACIAGPSVAALSSFFEHADDEELLH
ECIEGLFGVARICQYGLEDTLDELITSFCKFTTLLNPYASIEETMFTFSHDLKPRMATVAVFTIANYFRDSIQGG
WKNIVDCLLKLKRLKLIPQSVIDFEASDVPTTPVASGSQRVASMLTRFSLLSSDNMDDGLSLGSEFEQNLKMIKM
CKIGSIFSNCSSIPKECLQSLGRSLIFASAGKGQKFSTPVEEEETIEFCWDLITAIALANVHRFHIFWPNFHEYL
LAVAQFPMFSPIPFAEKALLGLLKVCLKLFSAPRDDKLAEELIFKSITLMWKLDKEILDTCHEIISQTISKIVIE
YPANLQTQIGWKSVLNLLSVVWRYPETYDLGIEALISLFSEGTLSRANYSFCIDCAFGCFLAKNSPIDKKKKILD
LLADSANLLVQWQRNQYSDPGSNVSTASYTSNSSMEDNSRGPASANYNMNLFVKLGEAFRRTSLSRQEEIRNHAV
ISLHKSFNLAEELLYVSSNCINYFNLVIFAMVDELHEKMLEYSRRENAERETRSMEGTLKLAMELLTDMYLQSLR
QITESPAFRTFWLGILRRMDTCMKADLGQYGPSTLSEIVPDLLKKIITQMKDEGILEPREDDDLWEITYIQIQWI
CPPLKDELFPL*

**>Cajca_LOC109801720**

MGRLKLQAGINAIEEEEPEECDASYPNKTTLACMINSEIGAVLAVMRRNVRWGGRYMSGDDQLEHSLIQSFKTVR
RQIFSWHHHQWQAINPALYLQPFLDVIRSDETGAPITGVALSSVYKIMTLDVIDQNTVNVENAMHLVVDAVTNCR
FEVTDPSSEEVVLMKILQVLLSCMKSKASIMLSNQHVCTIVNTCFRIVHQAGSKGELLQQIARYTMHEIVRCIFS
HLQDVGNTDHALVNGSSNLKQETGGLDNEYAFGSRQLENGSMSSEYDNQSLSTNSAPNAASVVKATVMDENTAIT
ISGKDGVPYDMHLMTEPYGVPCMVEIFHFLCALLNVVEHTGMGPRSNTLAFDEDVPLFALNLINSAIELGGPSIC
RHPRLLSLIQDELFHNLMQFGLSVSPLILSMVCSIVLNLYHHLRTELKLQLEAFFSCIILRLAQSRYGASYQQQE
VAMEALVDFCRQKTFMVDMYANFDCDITCSNVFEDLANLLSKSAFPVNCPLSAMHILALDGLIAVIQGMAERIAN
GSVSSEYSPVNLEEYTPFWMVKCENYDDPNHWVPFVRRRKYIKRRLMIGADHFNRDPKKGLEFLQGTHLLPDKLD
PQSVACFFRYTAGLDKNLVGDFLGNHDEFCVQVLHEFAGTFDFQDMNLDTALRLFLETFRLPGESQKIHRVLEAF
SERYYEQSPHILANKDAALVLSYSMIMLNTDQHNVQVKKKMTEEDFIRNNRHINGGNDLPREMLSEIYHSICKNE
IRTTPEQGVGFPEMTPSRWIDLMHKSKKTAPFIVSDSKAYLDHDMFAIMSGPTIAAISVVFDHAEQEEVYQTCMD
GFLAIAKISACHHLEDVLDDLVVSLCKFTTLLNPSSVEEPVLAFGDDMKARMATVTVFTIANRYGDYIRTGWRNI
LDCILRLHKLGLLPARVASDAADESELSSETVHGKPIMNSLSSAHMQSIGTPRRSSGLMGRFSQLLSLDTEEPRS
QPTEQQLAAHQRTLQTIQKCHIDSIFTESKFLQAESLLQLARALTWAAGRPQKGNSTPEDEDTAVFCLELLIAIT
LNNRDRIGILWQGVYEHISSIVQSTVMPCALVEKAVFGLLRICQRLLPYKENIADELLRSLQLVLKLDARVADAY
CEQITQEVSRLVKANASHIRSQLGWRTITSLLSITARHIEASEAGFDALLFIMSDSAHLVPANYVFCVDTARQFA
ESRVGQAERSVRALDLMAGSVNCLAQWTSEAKEAMKEDQVSKLSQDIGEMWLRLVQGLRKVCLDQREEVRNHALL
SLQKCLTGADGIYLPYSLWLQCFDLVIFTVLDDLLEIAQGHSQKDYRNMEGTLILAMKLLSKVFLQLLPDLSQLS
TFCKLWLGVLSRMEKYMKVKVRGKRSEKLQETVPELLKNSLLVMKMRGILAQRSALGGDSLWELTWLHVNNISPS
LQLEVFPEQDSEHLQHKQAESIGVSVPGEKGSVPSSETASREDAGIVG*

**>Cajca_LOC109808001**

MGHVKLQTQTGINAIEEGSEQFDAGYPNKTTVACMINAEIGAVLAVMRRNVRWGVHYMSDDDQLEHSLVQSLKAL
RRQIFSWQNQWHVISPALYLQPFLDVIQSDETGAPITGVALSSVYKILTLDVIDENTVNVGDAMHLVVDAVTSCR
FEVTDPGSEEVVLMKILQVLLACVKSKASVMLSNQHICTIVNTCFRIVHQAGTKSELLQRIARYTMHELVRCIFS
HLQDINNTELALINGETALKQETGGLSNEHNSANRQLENGSLNSASDGQPLSTGIASSTVSDVAGTVVDEDTAIA
SNGKGTDPSELQLMKEPYGIPCMVEIFHFLCSLLNVVEHMGVSPRSNTIAFDEDVPLFALTLINSAIELGGASFH
RHHRLLSLIQDELFRNLMQFGLSMSPLVLSMVSSIVLNLYHHLRTELKLQLEAFFSCVILRLAQSKYGASYQQQE
VAMEALVDFCRQKTFMVEMYANFDCDITCSNVFEDIANLLSKSAFPVNSPLSSMHILALDGLIAVMQGMAERIGN
GSLSSEQYPVNLEEYTPFWLEKCENFSDPNNWVPFVCRRKYFKRRLMIGADHFNRDTKKGLEFLQGSHLLPDKLD
PQSVACFFRYTAGLDKNLIGDFLGNHDEFCVEVLHEFARTFDFKDMMLDTALRLFLETFRLPGESQKIQRVLEAF
SERYYEQSPHILANKDAALLLSYSIIMLNTDQHNSQVKKKMTEDDFIRNNRRINGGSDLPREFLTELYHSICKNE
IQTTPEQGSGFPEMTPSRWIYLMHKSKKSAPFIVSDSEAYLDYDMFAILSGPTIAAISVVFDNAENAEVYQTCMD
GFIAVAKIAAYYHLENILDDLVVSLCKFVTVFDPLSVEESILAFGDDTKARMATETVFTIANRYGDYIRTGWRNI
LDCILKFHKLGLLPARMASDAAEESELSTETEHDGGKRNTNSLSLSHLPSVSTPKKSSGLMSRFSQLLYLGTEEP
RSVPTEEQLAAQQSTLQTIQKCHIDSIFSESKFLQADSLLQLAKALTSAGVRSKKGNSSFEDEDTVVFCLELLVA
ITLNNRDRIELLWQGVYEHISNIVQSTVMPCALVEKAVFGLLRICHRLLPYKENITDELLRSLQLVLKLDARVAD
AYYEQITQEVSHLMKANASHIRSHLGWRTITSLLSITARHLEAAEAGFDALVFIMSDGAHLLPANYVLCVDAAKQ
FAESRVGQVERSVMALDLMAGSVSCLEKWTNDAKQAAKEEEVAKMLQDIGDMWLRLIQGIKKLCLDQREEVRNHA
LLSLQNCLTGSVGVNLSHSLWLQCFDQVIFTVVDDLLEISQTHSQKEYRNIEGTLVLALKLLCKIFLQLIQALSQ
LTGFSELWLAVLSRLESYMKVRVRGRRSEKLQELVPELLKNILLVMKATRVLVRSSSVDGNSLFELTWQHINNIA
PSLKSELFPEQDTEKVEGVRPDENNSVSSKETAGQDGPGIG*

**>Cajca_LOC109801940**

MGHLKMQLQTGLNGTEDQYEHCGAGNLNKTTIACMINAEIGAVLAVMRRNVRWGVHYMSDDDQSEHFLVQSLKAL
RRQIFSWQNEWNAINPALYLQPFLDVIQSDETGAPITGVALSSVYKILTLDVIDQNTVNVGDAMHLVVDAITSCR
FEVTDSGSEEVVLMKILQVLLACVKGKASVMLSNQHICTIVNTCFRIVHQAGTKGELLQRIARYTMHELVRCIFS
HLQNIDNRECAFVNGSATLKQETNGVNNEHALASGQLENGRLNSTYEAQPLSTGIAFSTATVVTEAVMDENASIV
TSVNGIDPHELQVMIEPYGVPCMVEIFHFLCSLLNVAEHMGVNPRSNTITFDEDVPLFALTLINSAIELGGASFR
RHPRLLSLIQDELFRNLMQFGLSMSPLVLSMVCSIVLNLYHHLRMELKLQLEAFFSCVVLRLAQSKYGASYQQQE
VVMEALVDFCRQKTFMVEMYANFDCDISCSNVFEDIANLLSKSAFPVNNPLSSIHILALDGLIAVMQGMAERIGS
GSLGSEQSPVNFEEYSPFWMEKCDNFGDPNDWVPFVRRRKYIKRRLMIGADHFNRDDKKGLEFLQGTHLLPNKLD
PQSVACFLRYTAGLDKNLIGDFLGNHDEFCVQVLHEFSRTFDFEDMTLDMALRVFLESFRLPGESQKIHRVLEAF
SERYYEQSPHILANMDAALVLSYSIILLNTDHHNVQVKKKMTEEDFIRNNRHINAGNDLPREFLSEIYHSICKNE

IRTTPEPGFGFPEMTPSRWISLMHKSKQTAPFIVSDSRAYLDYDVFVTLSGSTIAAISVVFDNAENEEVYQACMD
GFLVVAKISAYYHLENVLDDLVVCLCKFITILDPLSVEESVLAFGDDTKARMATETVFTIANRYGDYIRTGWRNI
LDCILIFHKLCLLPPQLASDAADESQVTTETGDGKPNSNLLPSTHLRYTTPKRPSGLMGRFSQLLSLGVEETRSV
PTEEQLVAHQQAAQTIHKCHIDSILGESKFLQAESLLHLARALINAGAQPLKGNRISEDEDTSVFCLELLIAITL
NNRDRVGLLWTGVYEHISNIVQSTVMACALVERAIFGLLRICHRLLPYKENITDELLRSLQLVLKLDARVADAYY
EQITQEVTRLVKANASHIRSQFGWRTISSLLSITARHLEASEAGFDALRFIMSDGAHLLPANYILCVDVARQFAE
SRLGLVDRSIIALDLMAGSVICLEKWSNNAKQAVKEEEVEKVLQDIGEMWFRLVQGLRKVCLDQREEVRNHALLS
LQKCFTGAVGTHLPRDLWITCFDQVIFTVLDDLLEIVQSHSQKDYRHIEGTLIIALKLLFNVFLQLLLELSQLEA
FCKLWEGVLSRTEKSLQVKIWGRRSEKLQELAPELLKNTLLVMKSEGILVQNSGSGENSLWEMTFLHMKNIAPSL
QFEVFPEQDSEQXWHKPIETVGSLGPDANIFVPSNEKWAKRVMLSVKSTCSA*
**>Cajca_LOC109813450**
MEDAEDLTNESESSSSKRKRRELGLSCMLNTEVGAVLAVIRRPEFTSPYNMAPSEEHYDSSIVTSLRSLRSLIFN
PNQEWRTVDPSIYLSPFLDVIQSDDVPASATGVALSSLLKILKFQVFDDKSPGSREAMESVVSGITSCRLEKTDP
VSEDAVMMKILQVLTGIMHHRASTLLTDQSVCTLVNTCFQVVQQSATRGDLLQRSARYTMHELIQVVFTRLPDID
EDREDSESDMEDGDEGGGLESGYGVRCAIDIFHFLCSLLNVVSIVEADGSGSHTADEDVQIFALVLINSAIELSG
DEIGKHPKLLRMIQDDLFHHLIYYGTWSSSFVLSMICSTVLNAYHFLRSFIRYQLEAFFGYVLIRIASFGSTIPL
QEVAVEGIINFCRQPTFIMEVYANYDCDPFCRNVFEDAGRLLCKHSFALNSHLTSLHIQAFEGLLIIIHNIADNI
DKDGPACDLGSYTIQLSEYRPFWEEMDREDDLEAWVEHVRMRRLQKKKLLIAANHFNRDNKKGLEYLKHAKLVSD
PPDPKAYAYFFRYTPGVDKKAIGEYLGDPDTFYLQVLKEFTDTFHFQGMVLDTALRFYLESFWLPGESQKIQRVV
EAFSERFFDHQSSDMFASKDTVLILCYSLIMLNTDQHNPQVKKKMTEEDFIRNNRAINAGQDLPREYLSELYQSI
STCAFALSQTTVSLDMSPSRWIQLINLSKVDPTFTQCDFDRRICRDMFACVAGPVVAALSSFFEHAEEEELLHEC
IEGLFSVARICQYGLEDTLDELITSFCKFTTLLNPYASIEETMFTFSHDLKPRMATVAVFTIANYFRDSIQGGWK
NIVDCLLKLKRLKLLPQSVIDFEASADVPATPESGATSPTDDHKFGSQRIASMLSRFSLLSSDSMDDGLTIGSEF
EQNLKMIKMCKIGSIFIHSSNIPKESLQNLGRSLIFAAAGKGQKFSTPVEEEETVEFCWDLVTAISIANVNRFHI
FWPNFHEYLLAVAQFPMFSPIPFAEKAILGLLRVCLKLFSAPREDKLAEELIFKSITLMWKLDKEILDTCHEVIS
QTISKIIIDYPANVQTQFGWKSVLNLLSVAWRHPETYEAGIEALIALFSDGSCISRTNYTFCIDCAFGCFLAKNS
PVDKKKKILDLLADSVNLLVQWHRNLYTDPGSNASASSYNSNSSLEDNNRGPSANYNMNLFVKLGEAFRRTSLSR
QEEIRNHAVYSLHKSFNLAEDLLFTSSNCINYFNLVIFAMVDELHEKMLEYSRRENAERETRSMEGTLKLAMELL
SDMYLQSLRQITESPAFRTFWLGILRRMDTCMKADLGQYGASTLSDVIPDLLRKIISQMKEQGILEPKEEDDMWE
ITYIQIQWICPPLKEELFPL*
**>Proal_LOC114732932**
MGRLKLQAGINAIEEEPEEYDDAYPSKTTLACMINSEIGAVMSVMRRNVRWGGRYMSGDDQLEHSLIQSLKTLKR
QIFSWQHQWHAISPALYLQPFLDVIRSDETGAPSTSVALSSVYKILTLDVIDQNTVNVEDAMHLVVDSVTSCRFE
VTDPASEEVVLMKILQVFVACMKSKASVMLSNQHVCTLVNTCYRIVHQAGAKSELLQRIARHTMHELIRCIFSHL
QDVGNTEHALVNGSSTLKQEIGALNNEYAFGSKQLENGSMNSEYDNQSLPTDSAAGAATVIDENTAIASSGKGTV
PYDMHLMTEPYGVPCMVEIFHFLCSLLNAEHMDMGPRSNTIAFDEDVPLFALNLINAAIELGGPSIRQHPRLLSL
VQDELFRSLMQFGLSTSALILSMVCSIVLNLYHHLRAELKLQLEAFFSCVILRLAQSRYGASYQQQEVAMEAVVD
FCRQKTFMVDMYANFDCDITCSNVFEDLANLLSKSAFPVNCPLSSMHILALDGLIAVIQGMAERIGNGSNISEHS
PVNLEEYTPFWMVKCENYNDPNHWVPFVRRRKYMKRRLMIGADHFNRDPKKGLEFLQGTHLLPDKLDPQSVACFF
RYTAGLDKNLVGDFLGNHDEFCVQVLHKFAGTFDFQDMNIDTALRLFLETFRLPGESQKIQRVLEAFSERYYEQS
PHILANKDAALVLSYSLILLNTDQHNVQVKKKMTEEDFIRNNRHINGGNDLPREFLLELYHSICKNEIRTTPDQG
AGFPEMTPSRWIDLMHKSKKTAPFIVSDSRAYLDHDMFAIMSGPTIAAISVVFDHAEHEEVYQTCVDGFLAVAKI
SACHHLEDVLDDLVVSLCKFTTLLNPSSVEEPVLAFGDDTKARMATVTVFSIANRYGDYIRTGWRNILDCILRLH
KLGLLPARVASDAADESELSTETGHGKPIANSLSSAHMPSIGTPRRSSGLMGRFSQLLSLETEEPRSQPTEQQLA
AHQRTLQTIQKCHIDSIFTESKFLQAESLLQLARALIWAAGFPQKGSSTPEDEDTAVFCLELLIAITLNNRDRIG
TLWQGVYEHISNIVQSTVMPCALVEKAVFGLLKICQRLLPYKESIADELLRSLQLVLKLDARVADAYCEQITQEV
NRLVKANASHIRSQLGWRTITSLLSITAKHLEASEVGFEALLFIMSDGAHLLPANYVLCVEAARHFAESRVGQAE
RSVRALDLMAGSVDCLVQWTSEATQTMGEEEVSKLSQDIGEMWLRLVQGLRKVCLDQREEVRNHAILSLQKCLTG
VDGIYLSHSVWLQCFDLVIFTVLDDLLEIAQGQSQKDYRNMEGTLIFAMKLLSKVYLQLLQDLSQLSTFCKIWLG
VLSRMEKYMKVKVRGKKSEKLQELVPELLKNTLLVMKTRGILVQRSALGGDSLWELTWLHVNNIAPSLQSEVYPE
QDSEHLQHEKGGSNEGLEPVEVGSVSPNDTAKREDSALVG*
**>Proal_LOC114748431**
MGHLKLQAQAAISAIEEEPKQYDAGYPNRTTVACMINAEIGAVLAVMRRNVRWGVHYMSDDNQLEHSLVQSLKVL
RRQIFMWRNQWHAINPALYLQPFLDVIRSDETGAPITGVALSSVYKILTLDVIDQNTVNVGDTMHLVVDAVTSCR
FEVTDPASEEVVLMKILQVLLACVKSKASVVLSNQHICTIVNTCFRIVHQAGTKGELLQRIARYTMHELVRCIFS
HLQDVGSTDHALLNGSTAMTQETRQLRNGNFNAANDNQSLSACITSNADENPTIATGGKETVLHELHPMSERYGV
PCMVEIFHFLCSLLNVVEHMGMSPRSNTIAFDEDVPLFALTLINSAIELGGHSFCHHPRLLSLIQDELFRNLMQF
GLSMSPLVLSMVCSIVLNLYHHLRTELKLQLEAFFSCVILRLAQSRYGASYQQQEVVLEALVDFCRQKSFTMEMY
ANFDCDITCSNVFEDIANLLSKTAFPVNTPLSSMHILALDGLIAVMQGMAERIGIGSLSLESSQLNLKEYTPFWT
EKCDNFSNPNHWVSFVRRRKYLKRKLMIGADHFNRDPKKGLEFLQGTKLLPDKLDPQSVACFFRYTTGLDKNLIG
DFLGNHDEFCVQVLHEFARTFDFEDMTLDNALRLFLETFRLPGESQKIQRVLEAFSERYYEQSPQILANKDAALL
LSYSIIMLNTDQHNVQVKKKMTEEDFIRNNRRINGGNDLPREFLSDLYHAICKNEIRTTPEQGVGFPEMTPSRWI
YLMQKSRKTAPFIMSDSRSYLDHDMFAILSGPTIAALSVVFDNAEHEEVYQTCKDGFLIVAKISAYYHLDNVLDN

LVVSLCKFITILDPSLDDEPVLAFGDDIKARMATETVFTIANRYGDFVRTGWRNILDCILRLHKLGLLPAHMASD
AAEELSAEMGLAKPNANSLLSTHLPSSTPKRSSGLMSRFSQLLYLGAEEPKSEPTEEQLIAHQHSLETIQKCHID
SIFTESKFLQAESLLQLTKALISATGRPQKGNNTSDDEDTAVFCLDLLVAITLNNRDRIGLLWQGVYEHISAIVQ
SSVMPCALVEKAVFGLLRICQRLLPYKENIADELLRSLQLVLKLDARVADAYCEQITQEVGRLVKSNASHIRSQL
GWRTIASLLSITARHPEASEAGFDALLFIMSDGAHLLPANYVLCVDAARQFAESRVGQSERSIRALDLMSASVNC
LAKWTGDAKQAMQEEQVSKMLQDIGDMWLRLVQGLKKLCLEQREEVRNHALFSLQKCLSGAVDIDLTHGLWLQCF
DLVIFPVLDDLLEIAQAQSQKDYRNMEGTLILALKLLSKVFLQLFRDLSQSTAFTKLWLAVLGRMEKYVKVKIRG
RRSEKLQELVPELLKNTLLVMKSGGLLVQSSVLSGNSLWELTWQHVNNIDPTLQSEVFPEQDSEHLQPRKGETAG
DLRPGENVSIPSKETTGQDGAGIG*
**>Proal_LOC114759906**
MASGEEDRDKRRGSTKFKRKELGVSCMLSTEVGAVLALVRRPPDFNSPYLPADDNYESASVNSLRSLRSLIFTLH
QEWRTVDPSIYLTPFLEVIQADDVPASATSVALSSVLKILKLGIFDERTPGAKDALDAVANGITCCRLEKTDPVS
EDAVMMRILQVLTAIMSHRASLLFSDQAVCTIVNTCFQVVQQSANRGDLLQRTARYTMHELIQIVFSRLAQIEVK
DGEESSESDTDDVGVLESGYGVRCAIDIFHFLCSLLNVMEVIETEGPNAHSVDEDVQIFSLILINSAIELSGDAI
GKHPRLMRMIQDDLFHHLIYYGTWSSSLVLSMICSTVLNAYHFLRSFLRLQLEAFFGYVLLRVAALANTVQLQEV
AVEGIINFCRQPTFIAEAYANYDCDPFCRNVFEDIGKLLCKHSFAGNGPLTSLQIQAFEGLVIIIHNIADNIDKE
DESSPLGPYSNHITEYSPFWEEKEKDGKDLEAWVEHVRTRKLQKRKMLIAGNHYIRDDKKGLEYLKHSQLISDPP
DPKAFAFFFRYTPGLDKNLIGEYLGDPDDFHLQVLREFAATFYFTGMALDTALRFYLETFRLPGESQKIQRVLEA
FSERFYDQQSADIFASKDTVLILSYSLIMLNTDQHNPQVKKKMTEEDFIKNNRAINEGQDLPREFLSELYHSILS
NPLLAPTSTNVDMNPSKWIQLINRSKVVQPFIQCDFDRRMCRDMFACIAGPAVAALSTFFEHADEDDLLHECIEG
LFSVARIAQYGLEDTLDELISSFCKFTTLLNPYASTDEILYLFNNDLKPRMATVAVFTVANNFGDSIRGGWKNII
DCLLKLKKLKLLPQSVIDFELTPTQSADGSGVVFRVRNSKLGSRQVSNIASHSSFSSQDGKDNLIIGSEFEQNVN
MIKQCRIGAIFSKSSNIPEESLHNLGRSLIFAAAGKGQKFNTPIEEEETVGFCWDLITAITLVNVHRFQAFWPQF
HEYLLTVAQFPIFSPIPFAEKATAALCRISLRLFSSPQADKFPEELIFKSINLMWMLDKEILETCGGVISQSVIQ
LVSEYPGNLQTHLGWKTALHLLSITGRHPETYDQSSEALITLMSDGANFSRINYSYCVDCAFSFIALRNSPLEKN
LKILDLLSESVNLLIQWQKNQYSDPGSNSSGVSNNSSSSLDENPRGGSSNFTMNLFVKLGEAFKKTSLARREEIR
NYAVQSLRKSFDLADELPFTSANYISCLNLVIFAMVDDLHEKMLEYSRRENSEREMRSMEGTLKISMELMTDVYL
QFLKPLSESSGFRTFWLGILRRMDTCMKADLGQYGASTVQDLVPDLLRRIITRMKEKEILVQREGDDLWEITYFQ
IQWIAPALKDELFPD*
**>CercaGNOMa_sc32**
MGRLKLQAGINAIEEEPEESDPSYSNKTTLACMINSEIGAVLAVMRRNVRWAGRYMSGDDQLEHSLIQSLKALRR
QIFSWQHQWHTINPALYLQPFLDVIRSDETGAPITGVALSSVYKILTLDVIDQNTVNVEDAMHLVVDAITSCRFE
VTDPASEEVVLMKILQVLLACMKSKASVMLSNQHVCTIVNTCFRIVHQAGTKGELLQRIARYTMHELVRCIFSHL
QDVDNTEHALVNGSSTLKQEVGGLNNEYAFGSRQLENGSMSSEYDNQPLSTNFASATVGTANVMDENIAIPGSGK
ETIPYDMHLMTEPYGVPCMVEIFHFLCSLLNVVEHMGMGPRSNTIAFDEDVPLFALTMINSAVELGGPSICRHPR
LLSLIQDELFRNLMQFGLSMSPLILSMVCSIVLNLYHHLRTELKLQLEAFFSCVILRLAQSRYGSSYQQQEVAME
ALVDFCRQKTFMVDMYANFDCDITCSNVFEDLANLLSKSAFPVNCPLSSMHILALDGLIAVIQGMAERIGNGSVS
SKHSPVNLEEYTPFWMVKCENYGDPNNWVPFVRRRKYIKRRLMIGADHFNRDPKKGLEFLQGTHLLPDKLDPQSV
ACFFRYTAGLDKNLVGDFLGNHDEFCVQVLHEFAGTFDFQDMNLDTALRLFLETFRLPGESQKIQRVLEAFSERY
YEQSPQILANKDAALLLSYSLIMLNTDQHNVQVKKKMTEEDFIRNNRHINGGNDLPREFLTELYHSICKNEIRTT
PEQGVGFPEMTPSRWIDLLHKSKKTAPFIVSDSRAYLDHDMFAIMSGPTIAAISVVFDHAEHEEVYQTCMDGFLA
VAKISACHHLEDVLDDLVVSLCKFTTLLNPSSVEEPVLAFGDDTKARMATVTVFTIANRYGDYIRTGWRNILDCI
LRLHKLGLLPARVASDAADESELSAETGHGKPITNSLSSAHMPSIGTPRRSSGLMGRFSQLLSLDTEEPRSQPTE
QQLAAHQRTLQTIQKCHIDSIFTESKFLQAESLLQLARALIWAAGRPQKGNSTPEDEDTAVFCLELLIAITLNNR
DRIELLWQGVYEHISNIVQSTVMPCALVEKAVFGLLRICQRLLPYKENIADELLRSLQLVLKLDARVADAYCEQI
TQEVSRLVKANASHIRSQLGWRTITSLLSITARHLEASEAGFDALLFIMSDGTHLLPANYVLCVDAARQFSESRV
GQADRSVRALDLMAGSVNCLAQWTSEAKESMEEDEVAKLSQDIGEMWLRLVQGLRKVCLDQREEVRNHALLALQR
CLTGADEIYLPHSLWLQCFDMVIFTVLDDLLEIAQGHSQKDYRNMEGTLILAMKLLSKVFLQLLQDLSQLTTFCK
LWLGVLSRMEKYMKVKVRGKRSEKLQELVPELLKNTLLVMKTRGILVQRSALGGDSLWELTWLHVNNIAPSLQPE
VFPEQDLEHLQHKQGESVGGLVPDEKGSIPSNETVSHEDAGIVG*
**>CercaGNOMb_sc32**
MGHLKLQMQTGINAIEEEPEQSDASYPNKTTVACMINAEIGAVLAVMRRNVRWGVHYLSDDDQLEHSLVQSLKAL
RRQIFFWQNQWHAINPALYLQPFLDVIQSDETGAPITGVALSSVYKILTLDVIDQNTVSVGHTMHLVVDAVTSCR
FEVTDPGSEEVVLMKILQVLLACVKSKASIMLSNQHICTIVNTCFRIVHQAGTKGELLQRIARYTMHELVRCIFS
HLQEVENKEQILVNGGTTLKQEVGGQNNVHAFASRELENGNLRSAQNSQSLSTNSASSAGTVVTATLMDENTAVS
TSGKETDPHDLYLMNEPYGIPCMVEIFHFLCSLLNVVEHMGLGPRSNTIAFDEDVPLFALTLINSAIELGGPSIR
HHPKLLSLVQDELFRNLMQFGLSTSPLVLSMVCSIVLNLYHHLRTELKLQLEAFFSCVILRLAQSRYGASYQQQE
VAMEALVDFCRQETFMVEMYANFDCDITCSNVFEDLANLLSKSAFPVNGPLSSMHILALDGLIAVMQGMAERIGN
GVVNSELSRVNLEEYTPFWVEKCENFNDPNCWVPFACRRKYLKRRLMMGVDHFNRDPKKGLEFLQGAHLLPNKLE
PQSVACFFRYTAGLDKNLIGDFLGNHDEFCIQVLNEFARTFDFQDMTLDNALRLFLETFRLPGESQKIQRVLEAF
SERYYEQSPRILANMDAALLLSYSIILLNTDQHNVQVKKKMTEEDFIRNNRRINGGNDLPREYLSELYHSICKSE
IRTTPEKGGGFPEMTPSRWIYLVNKSRETAPFMVSDSRAYLDHDMFAILSGPTIAAISVLFDNAEYEEVYQTCMD
GFLAVAKISACYHLENVLDDLVVSLCKFTTILDPSSVDEPILAFGDDAKAKMATETVFTIANRYGDYIRTGWRNI

LDCILRLHKLGLLPARMASDAAEESELSAETGHGKSNTNSLSSAHMPSSISTPRKSSGLISRFSQLLSLGTEEPR
SQPTEEELAAHQHTLQTIQKCHIDSIFTESKFLQAESLSQLAKALINAAGPPQKGNNSPEDEDTAVFCLELLIAV
TLNNRDRIGLLWQGVYEHIFNVVQSTMMPCALVEKAVFGLLRICQRLLPYKENIADELLRSLQLVLKLDARVADA
YYEQITQEVSRLVKANASHIRSQLGWRTITSLLSITARHLESSEAGFDALLFIMSDGAHLLPANYVLCVDAARQF
AESRVGYEERSMQALDLMAGSVNCLTKWAGDAKQAMQEEEEEVAKMLQDIGEMWLRLVQGLRKVCLDQREEVRDH
ALLSLKKCLTGAVVIYLPVGLWLVCFDSVIFTLLDDLLEIAQAHSQKDYRNMEGTLILALKLLSKVFLQLLQDLS
QLTTFNKLWLGVLSQMEKYMKVKIRGRRSEKLQELVPELLKNTLLVMKTRGILVQGSAVGGDNLWELTWLHLNKI
APSLQSEVFTKLDAENLQHKQGEAVGDLGPDEKVSVPSNETVGQDGGVIG*

**>CercaGNL2_sc515**

MENANEGEDEQNNRDLTKSKRKELGISCMLNTEVGAVLAVIRRRPLEINSNSPYYISSPDEICDSSILNSLRSLR
SLIFNPQQEWRTIDPSIYLSAFLDVVQSDDVSSAATGVALSAILKILKLEIFDDKSPGAKEAMDVVVRGITSCRL
EKTDSVSEDAVMMKILQVLTGIMNHKASTLLTDHAVCTIVNTCFQVVQQSASRGDLLQRTARYTMQELIQVVFSR
LPEIEVKDEEDEDSESESEMDDVGGNSDSGYGVRCAIDIFHFLCSLLNVVEVVENTEGPTAHTADEDVQIFALVL
INSAIELSGDAIGKHPKLLRMIQDDLFHHLIHYGSWSSSLVLSMICSTVLNAYNFLRRFIRLQLGAFFGYVLVRV
SAFGSTIQLQEVAVEGIINFCRQSTFLIEAYANYDCDPLCPNVFEEVGKLLCKHTFVPTGPLSSLQIQAFEGLVI
IIHNLADSIDKEDESSSLGLHSMSITEYRPFWEEKPKEDVETWVEYVRVRKIQKRKILIAGNHFNRDDKKGLEYL
KHAQLISDPPDPKAFAFFFRYTPILDKNMIGEFLGDPDDLHLQVLKEFTETFQFSGMILDTALRTYLATFRLPGE
SQKIQRILEAFSDKFYEQQSSDTFASKDTVLILCYSLIMLNTDQHNPQVKKKMTEEEFIRNNRGINAGQDLPREY
LSEFFHSISTNAIAIFSQGVMQLDMNPSKWVELINRSKIVQPFIQCDFDRRMCRDMFACIAGPSVAALSSFFEHA
DEDELLHECIEGLFSVARIAQYGLEDTLDELITSFCKFTTLLNPYASSEETLFVFSHDLKPRMATLAVFTIANNF
GNSIRGGWKNVIDCLLKLKRLKLLPQSVFEFEIDGSPSEDVRTTPESGVIFPAHDPKFSNRHVFGISSRFSQFIS
LDSSDDGLTLGSELEQNLKIIKQCRIGNIFRNSSNIHEESLDNLGRSLIFAAAGKGQKFSTPIEEDETVGFCWDL
ITAIGLANVHRFQTFWSNFHEYLITVAQFPMFSPVPFAEKAIVGLLRICLSLLSTPQDDKLPEELIFKSINLMWK
LDREILDTCSEFITQSISSLVNEHSANLQTQLGWKSALHLLSVTGRHPECYDQGVDALITLMSDGTNISRINYPY
CIDCAFGFVALRNSPLEKNQKILDLLSDSVHLLIQWHNSQYPDTGSYTSLTSTTSNSSLEDSSRGLGSANFTINL
FVKLGEAFRKTSLSRREEIRNYAVKSLQKSFAMAEELVFTSALCINCFNLVIFAMVDDVHEKMLEYSRRDNAERE
MRSMEGTLKIAMELLTDVYLQFLKQISENPGFRTFWLGILRRMDTCMKADLGHYGATTLNESVPDLLRRIITKMK
ENELLEPIEGNDMWETTYIQIQWICPGLKDELFPPSSKDT*

**>Treor_270070**

MGRLKLQTGIKAIEEEPEDCDGCSNKATLACMINSEIGAVLAVMRRNVRWGGRYMSGDDQLEHSLIQSLKALRKQ
IFTWQHQWHTINPAVYLQPFLDVIRSDETGAPITGVALSSVYKILTLDVIDQNTVNVEDAMHLLVDAVTSCRFEV
TDPASEEVVLMKILQVLLACMKSKASVMLSNQHVCTIVNTCFRIVHQAGTKGELLQRIARHTMHELVRCIFSHLP
DVGNSESALVNGIDSIDRESSGLNSEYAFGSRQLENGSMTSEYDGQALSTNLASNASVGAAAVGMDEDAIGTGTG
KDAIPYDLRLMTEPYGVPCMVEIFHFLCSLLNVVENVGMGPRSNTVAFDEDVPLFALGLINSAIELGGASISHHP
RLLCLIQDELFRNLMQFGLSVSPLILSMVCSIVLNLYHHLRTELKLQLEAFFSCVILRLAQSRYGASYQQQEVAM
EALVDFCRQKTFMVEMYANLDCDITCSNVFEDLANLLSKSAFPVNCPLSSMHILALDGLIAVIQGMAERVGNGSV
GSEYTPVSLEEYTPFWMVKCDNYNDPSHWVPFVRRRKYIKRRLMIGADHFNRDPKKGLEFLQGTHLLPDKLDPQS
VACFFRYTAGLDKNLVGDFLGNHDEFCVQVLHEFAGTFDFQDMNLDTALRLFLETFRLPGESQKIQRVLEAFSER
YYEQSPQILANKDAALLLSYSLIMLNTDQHNVQVKKKMTEEDFIRNNRHINGGNDLPRDFLSELYHSICKNEIRT
TPEQGAGFPEMTPSRWIDLMHKSKKTAPFIVSDSRAYLDHDMFAIMSGPTIAAISVVFDHAEHEEVYQTCIDGFL
AVAKISACHHLEDVLDDLVVSLCKFTTLLNPSSAEEPVLEFGDNTKARMATVTVFTIANRYGDYIRTGWRNILDC
ILRLHKLGLLPARVASDAADESELSADTGHGKPLTNSLSSAHMPSMGTPRRSSGLMGRFSQLLSLDTEEPRSQPT
EQQLAAHQRTLQTIQKCHIDSIFTESKFLQAESLFQLARALIWAAGRPQKGNSSPEDEDTAVFCLELLIAITLNN
RDRIVLLWQGVYEHISNIVQSTVMPCALVEKAVFGLLRICQRLLPYKEKLADELLRSLQLVLKLDARVADAYCEQ
ITQEVSRLVKANASHIRSQLGWRTITSLLSITARHPEASEAGFDALLFIMSDGIHLLPANYVICVDASRQFAESR
VGQAERSVRALDLMAGSVDCLTRWASEAKEAMGEEEAVRMSQDIGEMWLRLVQALRKVCLDQREEVRNHALLSLQ
KCLTGVDGVHLPHGLWLQCFDLVIFTMLDDLLEIAQGHSQKDYRNMEGTLILAMKLLPKVFIQLLPDLSQLTTFC
KLWLGVLSRMEKYIKVKVRGKKSEKLQELVPELLKNMLIVMKTRGVLVQRSALGGDSLWELTWLHVNNIAPSLQS
EVFPDQSSEQSPNGEAAGDLVSDETDQVPSTQTTSSEGAAG*

**>Treor_079340**

MGHLNPQVSSNSFSEELRESQFKHSRTAIACMINSEIGAVLAVMRRNVRWGVRYVGDDDQLEHSLIHSLKELRKQ
IFSWQLDWHNIDPAVYLQPFLDVIQSDETGAPITGVALSSVYKILTLDILDLNTLNVGDAMHLIVDAVTSCRFEV
TDPASEEVVLMKILQVLLACTKSKASVKLSNQHVCNIVNTCFRVVHQASSKGELLQRIARHTMHELVRCIFSHLP
DIGNVEDALSNGSSSFVSKQVNEPDNDCKLGSKQTKHGTGGGEHDGQPTENASSSPKGTISSGTSENKIDVSNGI
EAVQSGEEVMMEPFGVPCMVEIFHFLCSLLNIIEHVEFGPRSNPIAYDEDVPLFALGLINSAIELAGPSIHKHPK
LLSIIQDELFHNLMQFGLSMSPLILSTVCSIVLNLYHHLRSELKVQLEAFFSCVILRLAQSKHGSSYQQQEVALE
ALVDFCRQPTFMAEMYANYDCDITCGNIFEDFANLLSKSAFPVNGPLSAMQILALDGLISMINGMAERVGSELPA
SEEVSVDLEDFKSFWMMKCENYNIPNYWVPFVRKIKYIKKKLTIGADHFNRDPKKGFEFLQGMHLLPDELDPQSV
AFFFRYTPGLDKNLVGDYLGNHDKFCVQVLHEFASSFDFSGMNLDTALRLFLGTFRLPGESQKIQRVLEAFAERY
YEQSPHIFFNKDAALLLSYSLIMLNTDQHNVQVKKKMTEEDFIRNNRRINGGNDLPREFLSELYHSICENEIQMV
PMQSAGFPVMTSSRWINVMHKSKNTPPFIICDSGVLLDYDMFRILSGPTTAAISVVFEQVEQADVLQTCIDGFLA
VAKISACYRFGDVLDDLVVSLCKFTTLLSPISVEEAIETFGDDTRAKMATTTVFTIANRYGDYIHSGWKNILDCV
LRLHKLGFLPPRLTSDAVDEDTSSDVEQGKPAANSFSTTQTPPLVTQRKSSGLMGRFSQLLSFDMEEPKLEPTEE

QIASQRRTRETIQNCHIDSIFTESKFLQAESLSQLVTALILAAGRLRKGTSTMEDEDTAVFCLELLIAITLNNRD
RIMLIWQGVYEHIADIVQSTVMPCPLVEKAVFGLLKICQRLLPYKENLTDELLKSLQLILKLDARVADAYCEHIT
QEVLRLVKANAAHIRSHMGWRTIISLLSITARHPEASEVGFEALATIMSDGAHLLPSNYVLCVDAARQFAESRVG
EVDRSVSALDLMAGSVVCLLRWSSETKHTVGEEAAMKVYQDIGEMWLRLVQGLRKLCLDQREEVRNHAILVLQRS
LAGVDGIDIPNTLWLLCFDGVIFTVLDDLLEIALGKSPKEYRNMEGSLVLAMKLMSKVFLQLLLDLCKLPSFCKL
WLGVLNRMERYIKAKLRGKRSDKIDELVPELLKNTLLVMKTTGILMPSDTIGADSFWQLTWLHVKNISPSLQSEV
FSSLELEQLQTKHVKPEGITAPEGTVLVQSE*

**>Treor_095630**

MEKEKDVPKSKRKELGLSCMLNTEVGAVLAVIRRPSDSSSHHLTSPEDSNDNALLQSLKSLRTLIFNPQQEWRSV
DPSIYLSPFLDVIQSDDIPAAATAVALSALLKIVKLEIFDDKSPGAAEAISAIVTAITNCRLEKSDPVSEDAVMM
KILQVLTGLMNHPASALLTDHAVCTIVNTCFQVVQQSAGRGDLLQRGARYTMHELIQIIFSRLPEIEANEAAESD
VDEAEELSVSGEFVSGYGIRSAVDVFHFLCSLLNVVELVETDGSTLQTADEDVQLFALVLINSAVELSGDEIGRH
PKLLRMLEDDLFHHLIHYGTCSSPLVLSMICSTVLNTYHFLRRFLRLQIEAFFAFVLLRISTTQGVPLQLQEVSL
EGIINFCRQPTFIIEMYENYDCDPHCRNVFEEIGKLLCKHSFPVSSPLTSLKIQAFEGLMIIIHNIADNIDRQDD
DSSTSFEPYPVEVTEFRPFWEDNTTNKSTNWVDFVRFRKAQKRRLLIAGYHFNRDEKKGLEYLKLSKLSPDPPDP
KAFAFFFRFTPGLDKSIVGDYLGDPDEFNLKVLKEFTETFQFSGMILDHALRTYLETFRLPGESQKIQRILEAFS
DRFYDQQTSDIFASKDAVFILCYSLIMLNTDQHNPQVKKKMTEEEFIRNNRAINGGKDLPKEYLSELFNSISNNA
ITIFGQSGQSVEMNPSRWIELMNRSRIVQPYKLSNFEHRQLGRDMFASIAGPSVATLCAFFEQADEEDLLHECVE
GLFSIAKIAQYGLEETLDELLASFSKFTTLLNPYASAEETLFAFSHDLKPKIATLAVFTIANNFRNSIRGGWKNI
VDCLLKLKRLKLLPQSVIEFDAASSTPSSDPQATSESGLIFPSNDPKFGTRQASGMISRFSHFLSLESMEESLSL
GVSEFEQNLKIIKQCQIGSTFSRTSNLPEDALINLGRSLIFSAAGKGQKFSTPVEEEETVGFCWDLIVSITLANI
QRFQTFWPAFNDHLLAVAQFPTFSPVPFAEKGIIGLFKVCLKLLGTYQPDELPEEYIFKSINLMWKLEKEILDTC
CDVITKSVNKILTEFPANLKTQLGWKSVLHLLSITGRNPETYEQGVETLIMLMSDGTHLSRINYAYCIDSAFSFV
ILRNSPLEKNIKILDLLANSVGHLVQWYKSQCSELASNSFSVASSASNSSLEDNLRGPGLLNFYMNLFIKLGEAL
RKTSLARREEMRNAAVLALRKSFSLTEELDLTPNNCVNCFNLVIFAMVDDLHEKMLEYSRRENAERETRSMEGTL
KIAMELLTEVFLKLMKQISNSPGFRTFWLGVLRRMDTYMKADLGEYGESKLPQAVPDLLRKMITEMKENEILVQK
EDDDLWEITYIQIQWIAPSIKDELFPEESY*

**>Paran_256170**

MGRLKLQTGIKAIEEEPEDCDGCSNKATLACMINSEIGAVLAVMRRNVRWGGRYMSGDDQLEHSLIQSLKALRKQ
IFTWQHQWHTINPAVYLQPFLDVIRSDETGAPITGVALSSVYKILTLDVIDQNTVNVEDAMHLLVDAVTSCRFEV
TDPASEEVVLMKILQVLLACMKSKASVMLSNQHVCTIVNTCFRIVHQAGTKGELLQRIARHTMHELVRCIFSHLP
DVGNSESALVNGIDSINRERSGLNNEYAFGSRQLENGNMTSVYDGQALSTNLASNASVGAAAAGMDEDAIGTGTG
KDTIPYDLRLMTEPYGVPCMVEIFHFLCSLLNVVENVGMGPRSNTVAFDEDVPLFALGLINSAIELGGASISHHP
RLLGLIQDELFRNLMQFGLSVSPLILSMVCSIVLNLYHHLRTELKLQLEAFFSCVILRIAQSRYGASYQQQEIAM
EALVDFCRQKTFMVEMYANLDCDITCSNVFEDLANLLSKSAFPVNCPLSSMHILALDGLIAVIQGMAERVGNGSV
GSEYTPVSLEEYTPFWMVKCDNYNDPSHWVPFVRRRKYIKRRLMIGADHFNRDPKKGLEFLQGTHLLPDKLDPQS
VACFFRYTAGLDKNLVGDFLGNHDEFCVQVLLEFAGTFDFQDMNLDTALRLFLETFRLPGESQKIQRVLEAFSER
YYEQSPQILANKDAALLLSYSLIMLNTDQHNVQVKKKMTEEDFIRNNRHINGGNDLPRDFLSELYQSICKNEIRT
TPEQGAGFPEMTPSRWIDLMHKSKKTAPFIVSDSRAYLDHDMFAIMSGPTIAAISVVFDHAEHEEVYQTCIDGFL
AVAKISACHHLEDVLDDLVVSLCKFTTLLNPSSAEEPVLEFGDNTKARMATVTVFTIANRYGDYVRTGWRNILDC
ILRLHKLGLLPARVASDAADESELSADTGHGKPLTNSLSSAHMPSMGTPRRSSGLMGRFSQLLSLDTEEPRSQPT
EQQLAAHQRTLQTIQKCHIDSIFTESKFLQAESLFQLARALICAAGRPQKGNSSPEDEDTAVFCLELLIAITLNN
RDRIVLLWQGVYEHISSIVQSTVMPCALVEKAVFGLLRICQRLLPYKEKLADELLRSLQLVLKLDARIADAYCEQ
ITQEVSRLVKANASHIRSQLGWRTITSLLSITARHPEASEAGFDALLFIMSGGVHLLPANYVICMDASRQFAESR
VGQAERSVRALDLMAGSVDCLTRWASEAKEAMGEEEAVRMSQDIGEMWLRLVQGLRKVCLDQREEVRNHALLSLQ
KCLTGVDGVHLPHGLWLQCFDLVIFTMLDDLLEIAQGHSQKDYRNMEGTLILAMKLLPKVFLQLLPDLSQLTTFC
KLWLGVLSRMEKYIKVKVRGKKSEKLQELVPELLKNTLIVMKTRGVLVQRSALGGDSLWELTWLHVNNIAPSLQS
EVFPDQSSEQSPNGEAAGDLVSDETDQLPSTQTISSEGAAG*

**>Paran_321370**

MGHLNPQVLCNSFSGEWRESQFNHSRTAIACMINSEIGAVLAVMRRNVRWGVRYVGDDDQLEHSLIHSLKELRKQ
IFSWQLDWHNIDPAVYLQPFLDVIQSDETGAPITGVALSSVYKILTLDILDLNTLNVGDAMHLIVDAVTSCRFEV
TDPASEEVVLMKILQVLLACTKSKASVKLSNQHVCNIVNTCFRVVHQASSKGELLQRIARHTMHELVRCIFSHLP
DNGNVEDALSNGSSSFVSKQVSEPDNDCKLGSKQTKHGTGGGEHDGQPTENASSSPKGTISSGTSENKIDVSNGI
EAVQSGEEVMMEPFGVPCMVEIFHFLCSLLNIIEHVEFGPRSNPIAYDEDVPLFALGLINSAIELAGPSIHKHPK
LLSIIQDELFHNLMQFGLSMSPLILSTVCSIVLNLYHHLRSELKVQLEAFFSCVILRLAQSKHGSSYQQQEVALE
ALVDFCRQPTFMAEMYANYDCDITCGNIFEDFANLLSKSAFPVHGPLSAMQILALDGLISMIHGMAERVGSELPA
SEEVSVDLEDFKSFWMMKCENYNIPNYWVPFVRKIKYIKKKLTIGADHFNRDPIKGFEFLQGMHLLPDELDPQSV
AFFFRYTPGLDKNLVGDYLGNHDKFCVQVLHEFASSFDFSGMNLDTALRLFLGTFRLPGESQKIQRVLEAFAERY
YEQSPHIFFNKDATLLLSYSLIMLNTDQHNVQVKKKMTEEDFIRNNRRINGGNDLPREFLSELYHSICENEIQMV
PMQSAGFPVMTSSRWINVMHKSKNTPPFIICDSGVFLDYDMFGILSGPTTAAISVVFEQVEQADVLQTCIDGFLA
VAKISACYRFGDVLNDLVVSLCKFTTLLSPISVGEAIETFGDDTRAKMATTTVFTMANRYGDYIHSGWKNILDCV
LRLHKLGFLPPCLTSDAVDEDTSSDVEQGKPAANSLSMTQTPPLVTQRKSSGLMGRFSQLLSFDMEEPKLEPTEE
QIASQRRTRETIQNCHIDSIFTESKFLQAESLSQLVTALILAAGRLRKGTSTMEDEDTAVFCLELLIAITLNNRD

RIMLIWQGVYEHIADIVQSTVMPCPLVEKAVFGLLKICQRLLPYKENLTDELLKSLQLILKLDARVADAYCEHIT
QEVLRLVKANAAHIRSHMGWRTIISLLSITARHPEASEVGFEALASIMSDGAHLLPSNYVFCVDAARQFAESRVG
EVDRSVSALDLMAGSVVCLGRWSSETKHTVGEEAAMKVNQDIGEMWLRLVQGLRKLCLDQREEVRNHAILVLQRS
LAGVDGIDIPNTLWLLCFDGVIFTVLDDLLEIALGKSPKEYRNMEGSLVLAMKLMSKVFLQLLLDLCKLPSFCKL
WLGVLNRMERYVKAKLRGKRSDKIDELVPELLKNTLLVMKTTGILMPSDTIGADSFWQLTWLHVKNISPSLQSEV
FSSLELEQLQTKHVKPEGITAPEGTVLVQSE*
**>Paran_058240**
MEKEKDLPKSKRKELGLSCMLNTEVGAVLAVIRRPSDSSSHYLTSPEDSNDIALLQSLKSLRTLIFNPQQEWRSV
DPSIYLSPFLDVIQSDDIPAVATAVALSALLKIVKLEIFDDKSPGAAEAISAIVTAITNCRLEKSDPVSEDAVMM
KVLQVLTGLMNHPASALLTDHAVCTIVNTCFQVVQQSAGRGDLLQRSARYTMHELIQIIFSKLPEIEANEASESD
IDEAEDLSISGDFVSGYGTRSAVDVFHFLCSLLNVVELVETDGSTLQTADEDVQLFALVLINSAVELSGDEIGRH
PKLLRMVEDDLFHHLIHYGTCSSPLVLSMICSTVLNTYHFLRRFLRLQIEAFFAFVLLRISTTQGVPLQLQEVSL
EGIINFCRQPTFIIEMYENYDCDPHCRNVFEEIGKLLCKHSFPVSSPLTSLKIQAFEGLMIIIHNIADNIDHQDD
DSSTFFEPYPVEVTEFRPFWEDNTTNKSTNWVDFVRFRKAQKRRLLIAGYHFNRDEKKGLEYLKLSKLSPDPPDP
KAFAFFFRFTPGLDKNIVGDYLGDPDEFNLKVLKEFTETFQFSGMILDHALRTYLETFRLPGESQKIQRILEAFS
DRFYDQQTSDIFASKDAVFILCYSLIMLNTDQHNPQVKKKMTEEEFIRNNRAINGGKDLPKEYLSELFNSISNNA
ITIFGQSGQSVEMNPSRWIELMNRSRIVQPYKLSNFEHRQLGRDMFASIAGPSVATLCAFFEQADEEDLLHECVE
GLFSIAKIAQYGLEETLDELLASFSKFTTLLNPYASAEETLFTFSHDLKPKIATLAVFTIANNFGNSIRGGWKNI
VDCLLKLKRLKLLPQSVIEFNAASSTPSSDPQATSDSGLISPSNDPKFGTRQASGMISRFSHFLSLESIEESLSL
GVSEFEQNLKIIKQCRIGSTFSRTSNLPEDALINLGRSLIFSAAGKGQKFSTPVEEEEAVGFCWDLIVSITLANI
QRFQTFWPAFNDHLLAVAQFPTFSPVPFAEKGIIGLFKVCLKLLGTYQPDELPEEYIFKSINLMWKLEKEILDTC
CDVITKSVNKILTEYPANLKTQLGWKSVLHLLSITGRNPEAYEQGVETLVMLMSDGTHVSRINYAYCIDSAFSFV
ILRNSPLEKNIKILDLLANSVGHLVQWYKSQCSELASNSFSVASSASNSSLEDNLRGPGLLNFYMNLFIKLGEAL
RKTSLARREEMRNAAVLALRKSFSLTEELDLTPNNCVNCFNLVIFAMVDDLHEKMLEYSRRENAERETRSMEVTL
KIAMELLTDVFLKLMKQISNSPGFRTFWLGVLRRMDTYMKADLGEYGESKLPQAVPDLLRKMITEMKENEILVQK
EDDDLWEITYIQIQWIAPSIKDELFPEESY*
**>Frave_LOC101300393**
MGRLKLQTGIKSIEEETEECDATNSHKSTIACIINSEIGSVLAVMRRNVRWGGRYMSGDDQLEHSLIQSLKALRK
QIFSWQHQWHTINPAVYLQPFLDVIRSDETGAPITGVALSSVYNILTLDVIDQNSVNVDDAMHMLVDAITSCRFE
VTDPASEEVVLMKILQVLLACMRSKASVMLSNQHVCTIVNTCFRIVHQAGTKGELLQRIARHTMHELVRCIFSHL
PDVHSTESALVNGNNTVKREIAGVNNEYAFGSRQLENGSINSEYDLQQLSTIPASNGSSGLAASGMDDTTIGASG
GKEAVQYDLHLMTEPYGVPCMVEIFHFLCSLLNVSEHMGMGPRSNTIAFDEDVPLFALVLINSAIELGGASIQHH
PKLLNLVQDELFRNLMQFGLSTSPLILSMVCSIVLNLYHHLRTELKLQLEAFFSCVILRLAQSRYGASYQQQEVA
MEALVDFCRQKNFMVEMYANLDCDITCSNVFEELANLLSKSAFPVNCPLSSIHILALDGLIAVIQGMAERVGNGS
VSSAHTPVNLEEYTPFWMVKCDNYSDPNHWVPFVRRRKYIKRRLMIGADHFNRDPKKGLEFLQGTHLLPEKLDPQ
SVACFFRYTAGLDKNLVGDFLGNHDDFCVQVLHKFAGTFDFQDMNLDTALRLFLETFRLPGESQKIQRVLEAFSE
RYYEQSPLILANKDAALLLSYSIIMLNTDQHNVQVKKKMTEEDFIRNNRHINGGSDLPRDFLAELYHSICKNEIR
TTPEQGAGYPEMTPSRWIDLMHKSKKNAPFIVSDSRAYLDHDMFAIMSGPTIAAISVVFDHAEHEEVYQTCIDGF
LAIAKISACHHLEDVLDDLVVSLCKFTTLLNPSSVEEPVLAFGDDTKARMSTVTVFTIANRYGDYIRTGWRNILD
CILRLHKLGLLPARVASDAADESEFSADAGPGKPIPNALSSVQLATVGTPRRSSGLMGRFSQLLSLDTEEPRSQP
TEQQLAAHQRTLQTIQKCHIDGIFTESKFLQAESLLQLARALIWAAGRPQKGNSSPEDEDTAVFCLELLIAITLN
NRDRIVLLWQGVYEHISNIVQSTVMPCALVEKAVFGLLRICQRLLPYKENLADELLRSLQLVLKLDARVADAYCE
QITLEVSRLVKANASHIRSQLGWRTITSLISITARHPEASEAGFDTLSFIMSDGTHLMPTNYNLCVDASRQFAES
RVGQTERSLTALDLMAGSVDCLVRWAHEAKKATNEEEAVKMSQDIGEMWLRLVQGLRKVCLDQREEVRNHALSLL
QKCLTEVDGIPLPHGLWLPCFDLVIFTMLDDLLEIAQGHSQKDYRNMEGTLISAMKLLSKVFLQLLSDLSQLTTF
CKLWLGVLSRMEKYMKAKVRGKKSDKLQEQVPELLKNTLVVMNSKGVLVQRSALGGDSLWELTWLHVNNISPSLK
SDVFPDQTLEQSETKTGETGGGLVSDEAGKVAPTETMSSELSGTGD*
**>Frave_LOC101300670**
MGRVKLQSGIKAGGEPKDSNGHYSDKATLACIINSEIGTVLAVMRRNVRWGGRYASDDDQREHSLIHSLKALRKK
IFSCQHQLNTISPAVYLQPFLDVIRSDETGAPITGVALASVYNILTLDVIDQNSVNLEDAMHLLIDAITGCRFEV
TDPASEEVVLMKILQVLLACMKSKASFMLSNQQVCTIVNTCFRIVHQAATKGELLQRIARHTMHELVRCIFSHLP
DICSTENALSNGSTSSKEETLGLNNEHALDSRQLDNGSISAEYDSQLLSTNPALHASPGVVESGMGEKPTGASDG
KDAGQNDLNLMTEPFGVPSMVEIFQFLCSLLNVAEHIGLGPKSNTIAFDEDVPLFSLTVVNSAIELGGSSIQRHP
KLLNLVRDELFQNLMQFGLSTSPLILSMVCSIVLNLYHHLRTELKLQLEAFFSCVILRLAESRYGASYQQQEVAM
EALVDFCRQKNFMVEMYANLDCDVTCSNIFEDLANLLSKSAFPVNSPLSSIHILALDGLIAVIQGMTERVDNGSS
SLSDTPVTLEEYTPFWMVKCDNYSDPNHWVPFVRRMKYIKRRLMTGAEHFNHDPKKGLEFVQGIHLLPEKLDPQS
VAWFFRYTAGLDKNLIGDFLGNHDEFCIQVLHKFAGTFDFQDMNLDTALRIFLETFRLPGESQKIQRVLEAFSER
YYEQSPLILADKDAALLLSYSTILLNTDHHNVQVKKKMTEEDFIRNNRHINGGEDLPREFLSELYHSICKNEIRT
TPEQGAGFPEMNPSRWIDLIHKSKKNASFIVSDSRAYLDQDMFSIVSGPTIAAISVVFDNAEQEEVYQTCLDGFL
AVAKISACYHLEDVLDDLVVSLCKFTTLLNSSYLDEPELEFGDDIKARMSTVMVFTIANRYGDYIRTGWRNIIDC
ILRLHKLGLLPAHVVQDAADESEFPSETDHGKPVTNSQPTIHTPSVSTPKKSSGFMGRFSQLLYLDTEEPKPQPT
EEELAAHQCTVQTVQECHIDSIFTESKFLQAESLLQLSQALIWAAGQLQKGNKSAEDEDTVVFCLELLIAITLNN
RDRIMLLWQGVYEFISNIVQSTFMPSALVEKAVFGLLRICQRLLPYKENLADELLRSLQLVLKLDARVADAYCEQ

ITQEVSRLVKANASHIRSQLGWRIITSLLSITARHPDASEAGFDALFFIMSGGAHLLPANYVLCVDASRQFAESR
VGQVDRSVCALDLMAGSVDCLARWAREAKLSMNEEDAVKMSQDIGEMWLRLVQGLRKVCLDQREVVRNHALLLLR
NCLTGVDGIPLPYGMWLQCFDLVIFTVLDDLLEIAQVHSQKEYRNMEGTLILAMKLLSKVFLQLLPELSHLTTFY
KVWLGVISRMGNYMKVKVGGRKSEKLQDEVPELLKNTLLVMNLRGVLVQKSGSGEDGLWEQTWLHVNKIAPSLQA
EVLAQILVQSHTEQGETEGVSDEKDPLLPTETIPSGASGSGG*
**>Frave_LOC101305794**
MGRVKLQTGIKAIEEENATNSHKSTIACIINSEIGSVLAVMRRNVRWGGRYMAGDDQLEHSLIQSLKALRRQIFS
WQHQWHTINXAVYLQPFLDVIRSDETGAPITGVALSSVDNILTVDVIHQNTVNVDDAMHMLVDAITSCRFEVTDP
ASEEVVLMKILQVLLACMRSKASVMLSNQHVCTIVNTCFRIVHQAGTKGELLQRIARNTMHELVRCIFSHLPDVH
RTESALVNGNNTVKREVAGVNNEYAFGSRHLENGSLNSEYDLQQLSTIPASNDSSVRAASGIDETTIGAFGRKEA
VQYDLHLMTEPYGVPCMVEIFHFLCSLLNVSEHMGMGPRSNTIAFDEDVPLFSLVLINSVIELGGVSIQHHPKLL
NLVRDELFRNLMQFGLSTSPLILSMAYSIVLNLYHHLRTELKLQLEAFFSCVILRLAQSRYGASYQQQEVAVEAL
VDFCRQKNFMVEMYANLDCDITCSNLFEELANLLSKSAFPVNCPLSSFHILALDGLIAVIQGIAERVGNGSVSSA
HTPVNLEEYTPFWMVKCDNYSDPNHWVPFVRRRKYIKRRLTIGADHFNHDPKKGLEFLQGTHLLPDKLDPQSVAC
FFRYTAGLDKNLVGDFLGNHDEFCVQVLQKFAGTFDFQDMNLDTALRLFLETFRLPGESQKIQRVLEAFSERYYE
QSPLILANKDAALLLSYSIIMLNTDQHNVQVKKKMTEEDFIRNNRHINGGSDLPREFLAELYHSICKNEIRTTPE
QGAGYPEMTPCRWIGLMHKSKKNAPFIVSDSRAYLDHDMFAIISGPTIAAISVVFDHAEHEEVYQTCIDGFLAIA
KISACHHLEDVLDDLVVSLCKFTTLLNPSSVEEPVLAFGDDTKARMSTVTVFTIANRYGDYIRTGWRNILDCILR
LNKLGLLPARVASNAADESEFSADAGPGKPITNALSSVQMATMGTPRRSSGLMGRFSQLLSLDTEEPRSQPTEQQ
LAAHQRTLQTIQKCHIDGIFTESKFLQPDSLLQLARALIGATGRPQKGNSSPEDDGTAVFCLELLIAITLNNRDR
IVLLWQGVYEHISNIVQSTVMPCALVEKAVFGLLRICQRLLPYKENLADELLRSLQFVLKLDARVADAYCEQITL
EVSRLVKANASHIRSQLGWHTITSLISITARPEASEAGFDTLSFIMSDGTHLLPTNYILCVDASKEFAESRVGQV
DRSVCALDLMAGSVDCLARWAREVKLSMNEEDAVKMSRDIGEMWLRLVQGLRKVCLDQREVVRNHALLLLRNCLT
GVDGIPLPYGLWLQCFDLVIFTVLDDLFEIAQVHSQKEYRNMEGTLILAMQLLSKVFLQLLPDLSHLTTFCKVWP
GVISRMGNYMKVKVGGRKSEKLQDEVPELLKNTLLVMHLRGVLVQKSGSDEDGLWEQTWLHVNKIAPSLQAEVLA
QILVQSHPQEVLAVVGRVQ*
**>Frave_LOC101307629**
MENHESNGKTKQATKLKRKELGLSCMLNTEVGTVLAVIRRPSDSNSHVFGAAPEDNVDPSLLQSLKSLRALIFNP
QQEWRTIDPTIYISPFLDVIQSDFVHAAATRVALSATLKILKLGVFDEKTPGAKDAINTVVNAITTCRLEKTDPV
SEDAVMMHILQLLTGVMNHRAAVLLSDQAVCTIVNTCFQVVQQSTSRGDLLQRTAKYTMHDLIHIIFSRLPEVDI
HDEGSETDTEDVDDGGLDFGYGMRCAVDIFHLLCSLLNVVEVVETDGSTVQTADEDVQLFALVLINTAIELSGDG
IGTHSKLLRMMKDDLFHHLVHYGTCSSPVVFSMICSTVLNMYHFLRRHIRLQLESFFTFEIFRVASAPGATTQLQ
EVALEGIINFCRQATFVVEVYVNYDCDPLCYNVFEEIGKLLCRQCFPVGAPLSSLQIQAFEGLVIMIHNIADSID
KENDTSPSGPYPVEVTEYTPFWEDQVKEYLETWVEFVRLRKAQKKKIMIAGHHYNRDEKKGMEYLRLTKLVSDPP
DPKSVAYFFRYTPGLDKMLIGDYLGDPDEFHIQVLREYTETFGFAGMNLDSGLRTYLETFRLPGESQKIERILEV
FSKNFYEQQPTDIFANKDTVMILCYSLIMLNTDQHNPQVKKKMTEDEFIRNNRAINGGKDLPRDYLSDLFHSISN
TAIAVFGQSGLQLEMNPNRWIELMNRTITVQPFFVCEFDRRLGRDMFACIAGPSVAALSAVFEHAEEEELLHECI
EGLFSVARIAQYGLEDTLDQILATFSKFTTLLNPYASAEETLFAFSKDLKPRMATLAVFTIANNFGESIRGGWRN
IVDCLLKLKRLKLLPQSAIEFDAAASTSSENLTSESGVIFPAQDAKFGGRQPSGMINRFSHFMSLESAEDAVSLG
ISEFEQNLKVIKQCRIGNIFSNSSKFPEDALLNLGRSLIYAAAGKGQKFSTAVEEEETVGFCWDVIVAITSANSQ
RFHTFWPSFHDYLIVVAQFPMFSPIPFAEKAIAGLFKICFKILGTFQPEKIPEELIFKSINLMWKLEKEILDSCC
DFITQSLNKILTEYPANLQTQLGWKSVLHLLQVSGRHTETYELGVETLVRFMSDGTHVSRINYAYCIDCAFSFIA
LKNSPLEKNIKILDLLAESVNLLVEWYRNYYSDPANSYSAHSNSSLDEARGIGSSCSFAMNLFVKLGEVLRKSSL
ARREEMRNHAVLSLKKSFRIAEALDFTPVNCINCFNLVVFAMVDDLHEKMLEYSRREHAEREMRSMEATLKIALK
LLTEVYLQFLIQISQSPGFRTFWLGVLRRMDTCMKADLGGYGESTVLQELVPDLLRQMITMMKEREILVQKEDDD
LWEITHIQIQWIAPAIKDELFPEENL*
**>Prupe_LOC18785443**
MGRLKLQSGIKAIEEEPEDCDATYSNKATLACIINSEIGSVLAVMRRNVRWGGRYTSGDDQLEHSLIQSLKALRK
QIFSWQHQWHTINPAVYLQPFLDVIRSDETGAPITGVALSSVYNILTLDVMDQNSVNVEEAMHLLVDATTSCRFE
VTDPASEEVVLMKILQVLLACMKSKASVMLSNQHVCTIVNTCFRIVHQAGTKGELLQRIARHTMHELVRCIFSHL
PDVNDTERALLNGSNTVTQEIAGLNNEYSFGNRQLENGNLSSGYDGQPLSTNPASNSSSGLVASVIDENKIGDST
GKDAVQYDLHLMTEPYGVPCMVEIFHFLCSLLNISEHMGMGPRSNTIEFDEDVPFFALVLINSAIELGGSYIQNH
PKLLSLVQDELFRNLMQFGLSTSPIILSMVCSIVLNLYHHLRTELKLQLEAFFSCVILRLAQSRYGASYQQQEVA
MEALVDFCRQKTFMVEMYANLDCDITCSNVFEELANLLSKSAFPVNCPLSSIHILALDGLIAVIQGMAERVGNGS
VSSEHTPVHLEEYTPFWMVKCENYSDPTDWVPFVRRRKYIKRRLMIGADHFNRDPKKGLEFLQGTHLLPDKLDPQ
SVACFFRYTAGLDKNLVGDFLGNHDEFCVQVLHKFAGTFDFQDMNLDTALRLFLETFRLPGESQKIQRVLEAFSE
RYYEQSPLILANKDAALLLSYSLIMLNTDQHNVQVKKKMTEEDFIRNNRHINGGSDLPREFLSELYHSICKNEIR
TTPEQGAGYPEMTPSRWIDLMHKSKKNAPFIVSDSRAYLDHDMFAIMSGPTIAAISVVFDHAEHEEVYQTCIDGF
LAVAKISACHHLEDVLDDLVVSLCKFTTLLNPSSVEEPVLAFGDDAKARMATVTVFTIANRYGDYIRTGWRNILD
CILRLHKLGLLPARVASDAADESEFSADTGPGKPISNSLSSVHIPSIGTPRRSSGLMGRFSQLLSLETEEPRSQP
TEQQLAAHQRTLQTIQKCHIDSIFTESKFLQAESLLQLARALIWAAGRPQKGNSSPEDEDTAVFCLELLIAITLN
NRDRIVLLWQGVYEHISSIVQSTVMPCALVEKAVFGLLRICQRLLPYKENLADELLRSLQLVLKLDARVADAYCE
QITQEVSRLVKANASHIRSQLGWRTITSLLSITARHPEASESGFDALFFIMSEGTHLLPANYALCVDASRQFAES

RVGQAERSICALDLMAGSVDCLARWAREAKQARNEEEVVKMSQDIGEMWFRLVQALRKVCLDQREDVRNHALSLL
QKCLTGVDGIPLPHNLWLQCFDVVIFTMLDDLLEIAQGHSQKDYRNMEGTLILAMKLLSKVFLQLLPDLSQLTTF
CKLWLGVLSRMEKYMKVKVRGKKSEKLQDQVPELLKNTLLVMNLKGVLVQRSALGGDSLWELTWLHVNNIAPTLQ
SEVFPDQISEQSETKQGENGGSLVSDETGTLLPTEMVSATGG*

**>Prupe_LOC18784935**

MGRLKLQNGIKAIDEEPGNCDATYSKKDTLACIINSEIGAVLAVMRRNVRWGGRYISGDDQLEHPLIQSLKVLRK
QIFSWQHQLHTINPAAYLQPFLDVIRSDETGAPITGVALSSVYNILTLDVIDQNSVNVEDAMHLLVDAITGCRFE
VTDPASEEVVLMKILQVLLACMKSKASVILSNQHVCTIVNTCFRIVHQAGTKGELLQRIARHTMHELVRCIFSHL
PDVQNTERALSNGNNTINREIAGINNEYPSGSRQLENGNVSSEFDSQLLSTNPALNASSGLVESGMDEKTTGASS
GKETVQYDSRLMAEPFGVPCMVEIFNFLCSLLNVVEHIGMGPRSNTISFDEDVPLFALGLVNSAIELGGSSIQNH
PKLLSLVQDELFQNLMQFGLSMSPLILSMVCSIVLNLYHHLRTELKLQLEAFFSCVILRLAQSRYGASYQQQEVA
MEAVVDFCRQKTFMVEMYANLDCDITCSNAFEDLANLLSKSAFPVNFPLSSIHILALDGLIAIIQGMAERSGNGS
VSSAETLTNLEEYTPFWLMKCDDYSDPNHWVPFVRRRKYIKRRLMIGADHFNHDPKKGLEFLQGTHLLPDKLDPE
SVACFFRYTSGLDKNLVGDFLGNHDEFCIQVLHEFAGTFDFQDMNLDTALRLFLETFRLPGESQKIQRVLEAFSE
RYYEQSPQILANKDAALLLSYSIIMLNTDRHNVQVKKKMTEEDFIRNNRHINGGDDLPREFLSELYHSICKNEIR
TTPEQGASFPEMTPSRWIDLIHKSKKNAPFIVSNFRPHLDQDMFAIMSGPTIAAISVVFDHAEHEEIYQTCIDGF
LSVAKIAACYHLEDVLDDLVVSLCKFTTLLNPSVDEPVLAFGDDPKARMSTVTVFTIANTYGDYIRTGWRNILDC
ILRLHKLGLLSACVASEAAGDSEVSADTGHGNPITNSLSSVHMPSVSTPRRSSGLMGRFSQLLSLDTEEPRSQPT
EEELAAHQRTLQTVQKCHIDGIFSDSKFLQAESLLQLAQALIWAGGRPHKGSSSPEDEDTGVFCLELLIAITLNN
RDRIMLLWQIVYEHISNIVQSTVMPCALVEKAVFGLLRICQRLLPYKENLADELLRSLQLVLKLDARVADAYCEQ
ITQEVGRLVKANASHIRSQLGWRIITSLLSITARHPEASEAGFDALFFIMSDGTHLLPANYVLCVDASRQFAESR
VGEVDRSVCALDLMAGSVDCLARWVCEAKQSMNDEEAVKMSQDIGKMSQDIGEMWLRLVQGLRKVCLDQREEVRN
HALSLLRKCLTGVDGIPLPPGLWLQCFDMVIFTMLDDLLEIAQRHSPKDYRNMEGTLILALKLLSKVFLQLLPDL
SQLTTFCKLWLGVLSRMEKYMKVKIGGKKSDKLRDQVPELLKNTLLVMILRGVLVERSDLGDDSLWELTWRLVNN
IAPSLQSEIFRDPILEQSETKQGETGGVSEATGTLLPTDTTSAEGSGS*

**>Prupe_LOC18770144**

MEKPSDPSNDENNGKSNQDKKLKRKELGLSCMLNTEVGAVLAVIRRPLDPTSHCFAAPDDAFDPSLLQPLKSLRA
LIFNPQQEWRTIDPSIYLFPFLDVIQSDEVHAAATRVALSATLKILKLGIFDEKTPGAKDAINTVVTAITTCRLE
RTDPVSEDAVMMNILQVLTGIMNHRASVLLSDQAVCTIVNTCFQVVQQSSSRGDLLQRCARYTMHELIQIIFSRL
PEINFRDGDSSASTDTEDADADEGNLDSGYGIRCAVDIFHFLCSLLNVVQVVETDQGSSVQTADEDVQLFALVLI
NSAIELSGDGIGSHSKLLRMMKDDLFHHLVHYGTRSSPLVFSMICSTVLNMYHFLRRFIRVQLEAFFTFVLFRVA
GPGVSIQLQEVALEGIINFCRQLTFVVEVYVNYDCDPLCHNVFEEIGKLLCKQSFPVSNPLTTIQIQAFEGLVIM
IHNIAESIDREHDTSPSGPYPIEITEYAPFWEDKPKDDSEAWVQFVRVRKAQKRKILIAGHHFNRDEKKGMEYLK
LYNLVSDPPDPKTLAFFFRYTPGLDKTMIGDYLGDPDEFHIKVLEEFTGTFGFAGMNLDSALRTYLETFRLPGES
QKIQRILEAFSESFYEQQPADLFVNKDTVFILCYSLIMLNTDQHNPQVKKKMTEEEFIRNNRAINGGKDLPREYL
SELFQSISNNAITLFGQSGLPVIMNPSRWIELMNRSKTVQPFILCDFDRRLGRDMFACIAGPSVAAISAFFEQAE
EEELLHECIEGLFSVARIAQYGLEDTLDELIATFSRFTTLLNPYASAEETLFAFGKDLKPRMATLAVFTIANNFG
ESIGGGWRNIVDCLLKLKRLKLLPQSVIDHDAASTSSSDAQATSESGIIFPAHDPKFGGHRASGMISRFSHFLSL
ESPEDALSLGMSEFEQNLKVIKQCRIGNIFSTSSHFPEDSLLNLGRSLIYAAAGKGQKFSTPIEEEETVGFCWDL
IVVIALANVHRFQAFWPNFHDYLLAVAQFPMFSPIPFAEKAIVGLFKVCLKLLGTYRPDRVPEELIFKSINLMWK
LEKEILDTCGELITQSVNKILIEYPANLQTQLGWKSVLHLLSVSGRHQDTYEQGVETLIMVMSDGTHVSRTNYAY
CIDCAFGFIALKNSPSEKNLKILDLLSDSVNLLIQWCRNQYSDPGNNYSVASNTSNSSLEDSKGFGSNNFAMNLF
VKLGEAFRKTSLARREDIRNHAVLSLQKSFKLAEELEFTPTNCINSFNLVIFAMVDDLHEKMLEYSRRESAEKEM
RSMEGTLKIALELLADVYLQFLIPISQCSGFRTFWLGVLRRMDTCMKADLGAYGESTLPELIPDLLRKMITEMKE
KEILVQKEDDDLWDITHIQIQWIAPSIKEELFPE*

**>Cansa_LOC115707648**

MGRLKMQSGIKAIEEEPEECDGYSIKTTLACMINSEVGAVLAVMRRNVRWSGRYMSGDDQLEHSLIQSLKALRKQ
IFTWQNNWRAINPSVYLQPFLDVIRSDETGAPITGVALSSVYKILTLDIIDQDTEHVEDAMNLLVDAVTSCRFEV
TDPASEEVVLMKILQVLLGCMKSKASVMLSNQHVCTIVNTCFRIVHQAGSKGELLQRIARHTMHELVRCIFSHLP
DVGNSESALVNGISNMNESSGLNNEYAFGSRQLENGNMASEYDGQALSMNLASNTSVGAAAVGMIDDTLGAGNGK
DSLPYDLSPTYGVPCMVEIFIFLCSLLNVVENVGMGPRSNTLAFDEDVPLFALGLINSAIELGGPSIRHHPRLLG
LIQDELFRNLMQFGLSTSPLILSMVCSIVLNLYHHLRTDLKLQLEAFFSCVILRLAQSRYGASYQQQEVAMEALV
DFCRQKTFMVEMYANLDCDITCSNIFEDLANLLSKSAFPVNCPLSSMHILALDGLIAVIQGMAERVSNGSVGSEY
TPVNFEEYTPFWMVKCDNYSDPSHWVPFVRRRKYIKRRLMIGADHFNRDPKKGLEFLQGTHLLPDKLDPQSVACF
FRYTAGLDKNLVGDFLGNHDDFCVQVLHEFAWTFDFQDMNLDTALRLFLETFRLPGESQKIQRVLEAFSERYYEQ
SPLILANKDAALLLSYSLIMLNTDQHNVQVKKKMTEEDFIRNNRHINGGNDLPRDFLIELYHSIIKNEIRTTPEQ
GAGFPEMTPSRWIDLMHKSKKTAPFIVSDSRAYLDHDMFAIMSGPTIAAISVVFDHAEHEEVYQTCINGFLAVAK
ISACHHLEDVLDDLVVSLCKFTTLLNPSSVEEPVLAFGDDTKARMATVTVFTIANSYGDYIRTGWRNILDCILRL
HKLGLLPARVASDAADESELTADSGHGKPLTNSLSSAHMPPMGTPRRSSGLMGRFSQLLSLDTEEPRSQPTEQQL
AAHQRTLQTIQKCHIDSIFTESKFLQAESLLQLARALIWAAGRPPKGNSSPEDEDTAVFCLELLIAITLNNRDRI
VLLWQGVYDHISNIVQSTVMPCALVEKAVFGLLRICQRLLPYKENLADELLRSLQLVLKLDARVADAYCEQITQE
VSRLVKANASHIRSQLGWRTITSLLSITARHPEASEAGFDALLFIMSDGAHLLPANYVLCVDASRQFAESRVGQA
ERSVRALDLMAGSVDCLTRWASEAKESMAEEDAVRMSQDIGEMWLRLIQGLRKVCLDQREEVRNHALLSLQKCLI

GVDGVHLPHTLWLQCFDMVVFTMLDDLLEIAQGHSQKDYRNMEGTLILAMKLLPKVFLRLLPDLSQLTTFCKLWL
GVLSRMEKYIKVKVRGKKSEKLQELVPELLKDTLIVMKTSGVLVQRSALGGDSLWELTWLHVNNIAPSLQSEVFP
DQCLEQSHNGDAAGDLASDETGQVPLTETASSEVAAGGG*
**>Cansa_LOC115704658**
MGQLKLKTGIKAMEEPLPECDRYSSKATLSCVINSEIGAVLAVMRRNVRWGVHYMSNEDQLEHSLIQSLKALRKQ
IFTWQHRWHTINPSVYLQPFLDVIRSDETGAPITGVALSSVYKILTLDVIDRNTVNVEDAMHLLVDAVTSCRFEV
TDPASEEVVLMKILQVLLACMKSKASVMLSNQHVCTIVNACFRIVHQAGTKGELLQRIARHTMDELVRCIFSHLP
DVDNSEMSPISGVNTLTRSSGLNSEPAFGSRQYENGNMTSEYDGQTLSANLVSNTYVHAGAVGLDDETVETGSGK
DSSSYDLILMTKPYGVLCMMEIFHFLCSLLNVVENMGMDPRSSTIAIDEDVPLFALSLINSAIQLGGPSIRHHPT
LLCFIQDELFHNLMQFGLSTSPLILSMVCSIVLNLYHHLRTELKLQLEAFFSCVILRLAQSRYGASYQQQEVAME
ALVDFCRQETFMVEMYANLDCDIPCSNVFEDLADLLSKSAFPVNSPLSSMHILALDGLIAVIQGMGERAANASVG
SECTPFNLKEYTPYWMEKCDNYSDPSHWVPFVRRRKYIKKRLMIGADNFNRDPKKGLEFLQGAHLLPDKLDPQSV
ACFFRYTTGLDKNLIGDFLGNHDEFSIQVLHEFAWTFDFQDMNLDTALRLFLEKFRLPGESQKIQRVLEAFSERY
YEQSPHVLANKDAALLLSYSLILLNTDQHNVQVKKKMTEEDFIRNNRKINGGNDLPREFLSELYHSILKNEIRTT
PEQGASFPEMTPSRWIDLILKSQKTAPFILSDCGAYLDHDMFAIVSGPTIAALSVVFDNAEDEEVYQTCIDGFLA
VAKISACHHLEDVLDDLVVSLCKFTTLLNPSSFEEPMLEFGDDTKARMATATVFTIANSYGDYIRTGWRNILDCI
LKLHNLGLLPARVASDASDELELSADRGHNKPITNSLSSGHMPPVNLPKRPSGLMGRFSQFLSLDTEEPKSQPTE
EQLAAHQRTLQTIQKFHIDSIFSESKFLQAESLLQLAQALILAAGRLQKGSSSPEDEDGAVFCLELLIAITLTNR
DRIVLLWQGVYDHISNIVQSTVMPCALVEKAVFGLFRICQRLLPYKENLADELLRSLQLVLKLDARVADAYCEQI
TQEVGRLVKANASHIRSQLGWRTITSLLSITARHPEASESGFDALLYVMSDGAHLLPANYVLFVDAARQFAESRV
GQAERSVCALDLMADSVDCLSRWASEAKKAMGEEEAVRMSQDIGEMWLRLVQGFRKVCLDQREVVRNHALLSLQK
CLTVVDSLHLSHGLWLQCFDLVIFSMLDDLHEIAQGQAQKDYRNMEGTLILAMKLLPKVFLRLLPDLSQLTTFVK
LWLGVLNRMEKYMRVKIRGRKSEKLHELVPELLKNTLIVMKTRGVLVPRSAPGGESLWEMTWSQVNNIAPSLQSE
VFPDQSLEQSYHGEAAADLVSDESDKVPSTQMSSFEGAAGGG*
**>Cansa_LOC115716528**
MEKEKEQQKSKRKELGLSCMLNTEVGALLAVIRRPADPNSHYLGSPEESQDNAILQSLKSLRALIFNPQQQWCTI
DPSIYLSPFLEVIQSDDIPASATGVALSALLKILKLEFFNLKTPAAAEAIATIVTSITNCRLEKTDSISEDAVMM
KILQVLSAIMNHSASALLTDQAVCVLVNTCFQVVQQSIGRGDLLQRSARYTMHELIQKIFSKLPEIETVTSIEGS
RLGEEEEEGGEFEGESGYGIRCAVDVFHFLCSLLNVVELVENSGSEVGNNTSFCGDEDVQVFALVLINSAIELSG
DGIGKHPKLLRMVQDDLFHHLIHYGTCSSPLVLSMICSTVLNTYHFLRRFLRLQIEAFFNFVLLRVPTTQGVSLQ
LQEVSLEGIINFCRQPTFIIEIYENYDCDPLCRNIFEDIGKLLCKHSFPVTSPLTSLKIQAFEGLMIIIHNIADN
IDRQYDNDDDNDNDSRTLFEPYPVEISEFRPFWEDKPRNDSINWVDFLRFRKAQKRKLLVTSYHFNCDEKKGLEY
LQLSKLSPDPHDPKALAFFFRYTPGLDKSMVGDYLGDPESFHLQVLKEFTETFEFSSMILDHALRTYLETFRLPG
ESQKITRILEAFSDRFYEQQTSDIFASKDAVFILCYSLIMLNTDQHNPQVKKKMTEEEFIRNNRAINGGKDLPKE
YLSELFNSISNNAITIFGQSGQIVEMNPSRWIELINRSRIIQPYKLSNFDDDKRIGRDMFGSIAGPSVATLCAFF
EHGDEEEILHECVEGLFSIARIAQYGLVETLDELLASFSKFTTLLNPYASAEETLFAFSHDLKPKIATLAVFTIA
NNFGNSIRGGWKNIVDCLLKLKRLKLLPQSVIEFDAANFGTTSSDSQATSEQGLSTPNDSKVVNVSRFSHFLSLE
SMEDSLSLGVSEFEQNLKIIKQCRIGNIFSKTSNLPEDALINLGRSLIFSAAGKGQKFSTPVDEEETVGFCWDLI
IAITLANLHRFQTFWHAFNDHLLAVAQFPTFSPVPFAEKGIVGLFKVCLKLLGTYQQEKTPEEMIFKSINLMWKL
EKEILDTCCEFITKSVNKILTEFPANLKTQLGWKSVLHLLSITGRQPETYDQAVETLIMLISDGTHISKMNYTYC
IDSAFSFLALRNSPLEKNIMILDLLASTVTQLVQWYKTQCLELSNSNNFSATSNASNSSFEENLRAPSSLNFFMS
LFIKLGEALRKTSLARREEIRNAAVMALQKSFMLAEELEFTPNNCINCFNLVIFAMVDDLHEKMFEYSRRENAER
ETRSMEGTLTIAMELLCEIFLKLMTQISNSPGFRTFWLGVLRRMDTYMKADLGEYGVSKLPGVVPDLLRKMITEM
KENGILVQKEDDDLWEITYIQIQWIAPSIKDELFPEESI*
**>Zizju_LOC107413409**
MGRLKLQTGIKAIEEEPEDFDDTYSNKTTLACMINSEIGAVLAVMRRNVRWGGRYVSGDDQLEHSLIQSLKALRK
QIFSWQHQWHTINPAVYLQPFLEVIRSDETGAPITGVALSSVYKILTLDVIDQNTVNVEDAMSLLVDAVTSCRFE
VTDPASEEVVLMKILQVLLACMKGKASIMLSNQHVCTIVNTCFRIVHQAGTKGELLQRIARNTMHELVRCIFSHL
SDVNNTKGALVNGTNTSKQEIAGLNNEHAFQSRQLDNGNMSSEYDGQSLSLNLASNTSMGAEAAGLDGDTIGAGT
GKSTVPYDLHLMTEPYGVPCMVEIFHFLCSLLNVVEHMGMDPRSNTIAFDEDVPLFALGLINSAIELGGPSIHQH
PRLLSLIQDELFRNLMQFGLSMSPLILSMVCSIVLNLYHHLRTELKLQLEAFFSCVILRLAQSRYGASYQQQEVA
MEALVDFCRQKTFMVEMYANLDCDITCSNVFEDLANLLSKSAFPVNCPLSSMHILALDGLIAVIQGMAERVSNGS
AGSEYTPVTLEEYTPFWMVKCDNYNDPDHWVLFVRRRKYIKRRLMIGADHFNRDPKKGLEFLQGTHLLPDKLDPQ
SVACFFRYTSGLDKNLVGDFLGNHDEFCVQVLHEFAGTFDFQDMNLDTALRLFLETFRLPGESQKIQRVLEAFSE
RYYEQSPHILVNKDAALLLSYSLIMLNTDQHNVQVKKKMTEEDFIRNNRHINGGNDLPREFLSELYHSICKNEIR
TIPEQGAGFPEMTPSRWIDLMHKSKKSAPYIVSDSRPYLDHDMFAIMSGPTIAAISVVFDHAEHEEVYQTCIDGF
LAVAKISACHHLEDVLDDLVVSLCKFTTLLNPSSVEEPVLAFGDDAKARIATVTVFTIANRYGDYIRTGWRNILD
CILRLHKLGLLPARVASDAADETELSADTGQGKPLTNSLSSAHLPSMGTPRRSSGLMGRFSQLLSLDTEEPRSQP
TEQQLAAHQRTLQTIQNCHIDSIFTESKFLQAESLLQLARALIWAAGRPQKGNSSPEDEDTAVFCLELLIAITLN
NRDRIVLLWQGVYDYIANIVQSTVMPCALVEKAVFGLLRICQRLLPYKENLADELLRSLQLVLKLDARVADAYCE
QITQEVSRLVKANASHIRSQLGWRTITSLLSITARHPEASEAGFDALFFIMSDGAHLLPANYVLCVDASRQFAES
RVGQAERSVRALDLMAGSVDCLAQWACEGKEAMGEEEAVKMTQDIGEMWFRLIQGLRKVCLDQREDVRNHAILSL
QKCLTRVDGIHLSHGLWLQCFDLVIFTMLDDLLEITQGHPKDYRNMEGSLVLAMKLLPKVFLQLLPDLSQLTTFC

KLWLGVLGRLEKYMKVKIRGKKSEKLQELVPELLKNTLLVMKTKGVLVQRSALGGDSLWELTWLHVNNIAPSLQS
EVFPDQNLEHSHSKQGEAGEDITSDEMGHVPSTEGAAGGG*
**>Zizju_LOC107413408**
MGRLKLQTGIKAIEEEPEDCDATCSNKATLACMINSEIGAVLAVMRRNVRWGGRYMSGDDQLEHSLIQSLKALRK
QIFSWQHQWQTINPAVYLQPFLDVIRSDETGAPITGVALSSVYKILTLDVIDQNTVNVEDAMHLLVDAVTSCRFE
VTDPASEEVVLMKILQVLLACMKGKASVMLSNQHVCTLVNTCFRIVHQAGTKGELLQRIARHTMHELVRSIFSHL
PDIDNTEGALVNGVNTVKREIDGLNNEYAFGSRQLENGNMSTEYDGQSLSTNLASNASIVAGMDEDTIRVGTGKG
NDPYDLHLMTEPYGVPCMVEIFHFLCSLLNAVENIGLGPKSNTIAFDEDVPLFALGLINSAIEVSGPSIRHHPRL
LTLIQDELFRSLMQFGLSMSPLILSVVCSIVLNLYHHLCTELKLQLEAFFSCVILRLAQSRFGASYQQQEVAMEL
VDFCRQKTFMVEMYANLDCDITCSNVFEDLANLLSKSAFPVNCPLSSMHILSLDGLIAVIQGMAERIGNGLVSSE
QSPMNIEEYTPFWMVQCDSYADPNHWVPFVRRRKYIKRRLMIGADHFNRDPKKGLEFLQGTHLLPDKLDPESVAC
FFRYTAGLDKNLVGDFLGNHDEFCVQVLHEFAWTFDFQDMNLDTALRLFLETFRLPGESQKIQRVLEAFSERYYE
QSPQILANKDAALLLSYSLILLNTDQHNVQVKKKMTEEDFIRNNRRINGGNDLPRDFLSELYHSICKNEIRTIPE
QGAGFSEMTPSRWIDLMHKSKKTAPFILSDSKAYLDHDMFAIMSGPTIAAISVVFDHAEHEEVFQTCINGFLAIA
KISACHHLEDVLDDLVVSLCKFTTLLNPSSVEEPVLAFGDDSKARMATVTVFTIANKYGDYIRTGWRNILDCTLR
LHKLGLLPARVASDAADESEPSVDTGHGKPLTNSLSTAHMPSVGTPRRSSGLMGRFSQLLSLDTEEPRSQPTEQQ
LAAHQRTLQTIQKCHIDSIFTESKFLQAESLSQLARALTWAAGRPQKGSNSPEDEDTAVFCLELLIAITLNNRDR
IVLLWQGVYEHIASIVQSTVMPCALVEKAVFGLLRICQRLLPYKENLADELLRSLQLVLKLDARVADAYCEQITQ
EVSRLVKTNASHIRSQMGWRTITSLLSITARHPEASEAGFDALLFIMSDGAHLLPANYVLCVDASRQFAESRVGQ
AERSVHAVDLMAGSVDCLARWACEGKEAMGEEEAMKMTQDIGEMWLRLVQGLRKVCLDQREEVRNHALLSLQTCL
TRMDGIHLPYGFWLQCFDLVIFTLLDDLLDIIAQGSSQKDYRNMEGTLIFALKLLSKVFLQLLPDLSQLTTFCKL
WLGVLNRMEKYAKVKVRGKKSEKLQELVPELLKNTLLVMKTRGVLVQRTALGGDSLWELTWLNVNKIAPSLQSEV
FPDQNSEQSEGKQGEPLEGLLSNESSDVPATKATLSEGAAGGG*
**>Zizju_LOC107431830**
MEKARDLKQENDREMTKSKRRELGISCMLNAEVGSVLAVIRRQPDANSLYVTAEQDNFDSALLQSLKSLRALIFN
PQQEWRGMDPSVYLSPFLDVIQSDDVPAIATGVSLSALLKILKLGVFDEKTPGAKDAINSVVTAITSCRLEKTDP
VSEDAVMLKILQVLIGTMCHPASVWLTDQAACTLINTCFQVVQQSASRGDLLQRSARYTMHELVQVMFSRLAEVE
EASESDTEDADIGGDLDSGYGVRSAVDIFHFLCSLLNVVELVEGDGSPLRNADEDIQLFALVLINSAVELSGDEI
GKHPKLLRMVQDDLFHHLIHYGVCSSPLVLSMICSTVLNVYHFLRRFCRLQIEAFFEYVLLKVSTMGAPLALQEV
AVEGIVNFCRQPKFIVEVYINYDCDPLCHNVFEEIGKLLCKHSFPVNSPLNALRIQAFEGLMVLIHNIADNIDRE
DDSTPHGPYPVEISGYRPFWEDKPKEDLETWVEYFRVRKAQKRKLLIAGHHYNRDEKKGLDYLKLSHLVSDPPDP
KAYAFFFRYTNGLDKNSIGDYLGDPDEFHVNVLKEFTETFQFAGMILDTALRTYLETFRLPGESQKIQRILEAFA
DRFYDQQTGDIFATKDAVFILCYSVIMLNTDQHNPQVKKKMTEEEFIRNNRAINAGKDLPREYLSELFQSIANNA
ITLFGQSGGPMEMNPSRWIELINRSKLVEPFTLCDFDRRLGRDMFACIAGPSVAALCAFFEHSDEEDLLHECLEG
LFSVARIAQYGLEDTLDELLASFCKFTTLLNPYASAEETLFAFSNDLKPKMATLAVFTIANNFGNSIRGGWRNIV
DCLLKLKRLKLLPQSVIEFDAASTLSSDNPATSESGVIFPPHDPAFANRHNSGMISRFSHFLSLESTEDAVSLGV
SEFEQNLKVIKQCRIGNIFSNTSNLPDDALLNLGRSLIFAAAGKGQKFSTPVEEEETVGFCWDLITAVTLANIQR
FQTFWGNFHDYLLAVAQFPLFSPIPFVEKAILGLFKVCLKLLGTYQPEKLPEELIFKSINLMWKLDKEILDTCCE
FITQSANRILIEYPANLQTQLGWKSALHLLSVTGRHPETYEIGVETLIMLLSDGTHVSRLNYAYCIDCAFGFVAL
RNSPLDKNLRILELLSDTVNKLVQWYKNQCADSGSNYSISSNTSSSSLEENSKIFGSGNFAMNLFVKLGEALRKT
SLARREEIRNQAVLSLQKSFLLAEELDFTPANCINCFNLVIFAMVDDLHEKMLEYSRRDNAEKEMRSMEGTLKLA
MELLTDVYLRYLKNLSQSPGFRTFWLGVLRRMDTCMKADLGDFGESKILQELIPELLRKIITEMKEKKILVKNEG
DDLWDITHIQIQWIAPSVKEELFPEETL*
**>Rosch_LOC112182803**
MGRLKLQSGIKAIEEEPEECDATISHKATLACIINSEIGSVLAVMRRNVRWGGRYMSGDDQLEHSLIQSLKALRK
QIFSWQHQWHTINPTVYLQPFLDVIRSDETGAPITGVALSSVYNILTLDVIDQNSVNVDDAMHLLVDAITSCRFE
VTDPASEEVVLMKILQVLLACMRSKASVMLSNQHVCTIVNTCFRIVHQAGTKGELLQRIARHTMHELVRCIFSHL
PDVHNTESALVNGNNTVKREIAGVNNEYAFGSRQLENGNINSEYDLQQLSTYPASNGSSGLVASGIDENTIGASG
GKEAVQYDLQLMTEPYGVPCMVEIFHFLCSLLNVSEHMGIGPRSNTIAFDEDVPLFALVLINSAIELGGASIQHH
PKLLNLVQDELFRNLMQFGLSTSPLILSMVCSIVLNLYHHLRTELKLQLEAFFSCVILRLAQSKYGASYQQQEVA
MEALVDFCRQKSFMVEMYANLDCDITCSNVFEELANLLSKSAFPVNCPLSSIHILALDGLIAVIQGMAERVGNGS
VSSAHTPVNLEEYTPFWMVKCDNYSDPNHWVPFVRRRKYIKRRLMIGADHFNRDPKKGLEFLQGTHLLPDKLDPQ
SVACFFRYTAGLDKNLVGDFLGNHDEFCVQVLHKFAGTFDFQDMNLDTALRLFLETFRLPGESQKIQRVLEAFSE
RYYEQSPLILANKDAALLLSYSIIMLNTDQHNVQVKKKMTEEDFIRNNRHINGGSDLPRDFLTELYHSICKNEIR
TTPEQGAGYPEMTPSRWIDLMHKSKKNAPFIVSDSRAYLDHDMFAIMSGPTIAAISVVFDHAEHEEVYQTCIDGF
LAIAKISACHHLEDVLDDLVVSLCKFTTLLNPSSVEEPVLAFGDDTKARMSTVTVFTIANRYGDYIRTGWRNILD
CILRLHKLGLLPARVASDAADESEFSADAGPGKPITNSLSSVHMPTVGTPRRSSGLMGRFSQLLSLDTEEPRSQP
TEQQLAAHQRTLQTIQKCHIDGIFTESKFLQAESLLQLARALIWAAGRPQKGNSSPEDEDTAVFCLELLIAITLN
NRDRIVLLWQGVYEHISNIVQSTVMPCALVEKAVFGLLRICQRLLPYKENLADELLRSLQLVLKLDARVADAYCE
QITLEVSRLVKANASHIRSSLGWRTITSLISITARHPEASEAGFDTLSFIMSDGTHLLPANYSLCVDASRQFAES
RVGQTERSVIALDLMAGSVDCLVRWAHEAKQATNEDEAVKMSQDIGEMWLRLIQGLRKVCLDQREEVRNHALSLL
QKCLTEVDGIPLPQGLWLQCFELVMFTMLDDLLEIAQGHSQKDYRNMEGTLISAMKLLSKVFLQLLSDLSQLTTF

CKLWLGVLSRMEKYMKVKVRGKKSDKLQDQVPELLKNTLVVMNLKGVLVQRSALGGDSLWELTWLHVNNISPSLQ
SEVFPDQTLEQSETKTGETGGSLVSDEAGKVAPTETMSSEVSGTGG*
**>Rosch_LOC112182802**
MGRLKVQSGIKAVEGELKDSDGIYSDKATLACIINSEIGTVLAVMRRNVRWGGRYASDDDQMEHSLIQSLKALRK
KIFSWQHQLNSISPAVYLKPFLDVIRSDETGAPITGVALASVYNILTLDVIDQNSVNVEDAMHLLVDAITGCRFE
VTDPASEEVVLMKILQVLLACMKSKASFMLSNQQVCTIVNTCFRIVHQAGTKGELLQRIARHTMHELVRCIFSHL
PDIRSTENALSNGSNSLKQETSGLNNEHALESRQLDNGSISSEYDSQLLSTNPALHASPGIVESVMGEKTTGASD
RKDAVQNDLNLMTEPFGVPSMVEIFQFLCSLLNVVEHIGSGPKSNPIAFDEDVPLFALTLVNSAIELGGSSIQHH
PKLLSLVRDELFQNLMQFGLSTSPLILSMVCSIVLNLYHHLRTELKLQLEAFFSCVILRLAQSRYGASYQQQEVA
MEALVDFCRQKNFMVEMYANLDCDITCSNIFEELANLLSKSAFPVNSPLSSFHILALDGLIAVIQAMTERVENGS
YSLSHTPVTLEEYTPFWMVKCDNYSDPNHWLPFVRQTKYIKRRLITGADHFNHDPKKGLEFLQGIHLLPEKLDPQ
SVACFFRYTAGLDKNLIGDFLGNHDDFCIQVLHKFAGTFDFQDMNLDTALRLFLETFRLPGESQKIQRVLEAFSE
RYYEQSPLILADKDAALLLSYSIIMLNTDQHNVQVKKKMTEEDFIRNNRHINGGEDLPREFLSELYHSICKNEIR
TTPEQGAGFPEMTPSRWIDLMHKSKKNAPFIVSDSRAYLDQDMFSIVSGPTIAAISVVFDNAEQEEVYQTCLDGF
LAVAKISACYHLEDVLDDLVVSLCKFTTLLNSSFHDEPELEFGDDIKARMSTVTVFTIANRYGDYIRTGWRNIID
CILRLHKLGLLPARVAQDAADESEFPSETDHGKPITNSQSKIHTPSVSTPKKSSGFMGRFSQLLSLDTEEPKPQP
TEEELATRQRTVQTVEECHIDSIFTESKFLQAESLLQLSQALIWAAGRLQKGNNSAEDEDTVVFCLELLIAITLN
NRDRIMLLWQGVYDFISNIVQSTVMPCALVEKAVFGLLRICQRLLPYKENLADDLLRSLQLVLKLDARVADAYCE
QITLEVSRLVKANASHIRSQLGWRTITSLLSITARHPDASEAGFDALFFIMSGGTHLLPANYVLCVDASRQFAES
RVGQADRSVCALDLMAGSVDCLARWACEAKLSMNEEDSVKMSQDIGEMWLRLVQGLRKVCLDQREVVRNHALSML
RNCLTGVDGIPLPYGLWLQCFDLVIFTVLDDLLEIAQVHSQKEYRNMEGTLILAMKLLSKVFLQLLPDLSHLTTF
CKLWLGVISRMENYMKVKVGGRKSEKLQDEVPELLKNTLLVMNLRGVLVQKSGSGEDGLWEQTWLNVNRIAPSLQ
AEVLAQILVQSHTEQGETGVSAEKGSSTETIPSGGSGCGG*
**>Rosch_LOC112190848**
MENHDKKQAIKLKRKELGLSCMLNTEVGTVLAVIRRPSETVFGAAPDDSIDPSLLQSLKSLRALIFNPQQEWRTI
DPSIYLSPFLDVIQSDFVQASATRVALSATLKILKLGVFDEKTPGAKDAINTVVNAITTCRLEKTDPVSEGAVMM
NILQLLTGVMNHRASMLLSDQAVCTIVNTCFQVVQQSTNRGDLLQRNARYTMHELIHIIFSRLPEVEMSHEDEET
DNEDVDDGGLDFGYGMRCAVDIFHFLCSLLNVVEVVETEGSTVQTADEDVQLFALVLINTAIELSGDGIGTHSKL
LRMMKDDLFHHLVHYGTCSSPVVFSMICSTVLNMYHFLRRYIRLQLESFFAFVLFRVAGAPGATTQLQEVALEGI
INFCRQPTFIVEVYVNYDCDPFCHNVFEEIGKLLCKQSFPVGGPLSSLQIQAFEGLVVMIHNIADSIDKENDTSP
SGPYPVEITEYTPFWEDEAKEDSEIWVGFVRLRKAQKKKILIAGHHYNRDEKKGMEYLKLTKLVSDPPDPKALAF
FFRYTPGLDKMLIGDYLGDPDEFHIEVLREYTETFGFAGMNLDSGLRTFLETFRLPGESQKIERILEVFSENFYE
QQPTDIFANKDTVLILCYSLIMLNTDQHNPQVKKKMTEDEFIRNNRAINGGKDLPREYLSELFHSISHNAFTTFG
LQVEMNPSKWIELMNRTKTVQPYFLCEFDRRLGRDMFACIAGPSVAALSAVFEHAEEEELLHECIEGMFSVARVA
QYGLEDTLDELLATLSKFTTLLNPYASAEETLFAFSKDLKPRMATLAVFTIANNFGESIRXGWRNIVECLLKLKR
LKLLPQSAIEFDAASTSSENLKSESGIIFPAHDPRFGGRPAGMINRFSHFMSLESAEDAVSLGISEFEQNLKVIK
QCRIGNIFSNTSNFPEVALLNLGRSLIYAAAGKGQKFSTAIEEEETVGFSWDLIVAITLANNHRFQTFWPHFHDY
LIVVAQFPMFSPIPFAEKAIVGLFKICLTILETHRADKIPEELIFKSINLMWKLEKEILDTCCHFITQSLNRILI
EYPANLQTQLGWKSVLHLLSVSGRHPECYELGVETLISLMSDGTHVSRTNYAYCIDCAFGFIALKNSPLEKNIKI
LDLLAESVNFLIQWYRNHYSDPGNSYSNSSLEDAKGFGSSCSFAMNLFVKLGEAFRKSSLARREEMRNHAILCLQ
KSFKLAEELDFTPINCINCFNLVVFAMVDDLHEKMLEYSRRENAEKEMRGMEGTLKVALKLLTEVYLQFLKQISQ
SPGFRTFWLGVLRRMDTCMKADLGGYGESTTLQELVPDLLRQMITVMKEKEMLVQKEDDDLWEITHIQIQWIAPA
IKEELFPEENL*
**>Morno_LOC21408061**
MGRLKLQSGIKAIEEEPEDCDSYSNKATLACMINSEIGAVLAVMRRNVRWGGRYMSGDDQLEHSLIQSLKVLRKQ
IFTWQHHWHTINPAVYLQPFLDVIRSDETGAPITGVALSSVYKILTLDVIDQNTVNVEDAMHLLVDAVTSCRFEV
TDPSSEEVVLMKILQVLLACMKSKASVMLSNQDVCTIVNTCFRIVHQAGSKGELLQRMARHTMHELVRCIFSHLP
DVGNSESALVNGIDTINRESSGLNNEYAFGSRQLENGNTTSDYDGQALSTNLTSNASVGPGGMDEDAIGTGKDTV
PYDLRLMTEPYGVPCMVEIFHFLCSLLNVVEQVGMGPKSNTIAFDEDVPLFALGLINSAIELGGPSIRYHPRLLS
LIQDELFRNLMQFGLSMSPLILSMVCSIVLNLYHHLRTELKLQLEAFFSCVILRLSQSRYGASYQQQEVAMEALV
DFCRQKTFMVEMYANLDCDITCSNVFEDLANLLSKSAFPVNCPLSSMHILALDGLIAVIQGMAERVGNGSVGSEH
TPVTLDEYTPFWMVKCDNYSDPSYWVPFVRRRKYIKRRLMIGADHFNRDPKKGLEFLQGTHLLPDKLDPQSVACF
FRYTAGLDKNLVGDFLGNHDEFCVQVLHEFAGTFDFQDMHLDTALRLFLETFRLPGESQKIQRVLEAFSERYYEQ
SPEILANKDAALLLSYSLIMLNTDQHNVQVKKKMTEEDFIRNNRHINGGNDLPREFLSELYHSICKNEIRTTPEQ
GAGFPEMTPSRWIDLMHKSRKAAPFIVSDSRAYLDHDMFAIMSGPTIAAISVVFDHAEHEEVYQTCIDGFLAVAK
ISACHHLEDVLDDLVVSLCKFTTLLNPSSVEEPVLAFGDDTKARMATVTVFTIANRYGDYIRTGWRNILDCILRL
HKLGLLPARVASDAADESELSADTGHGKPLTNSLSSAHMPPMGTPRRSSGLMGRFSQLLSLDTEEPRSQPTEQQL
AAHQRTLQTIQKCHIDSIFTESKFLQADSLLQLAKALIWAAGRPQKVGSSPEDEDTAVFCLELLIAITLNNRDRI
VLLWQGVYEHIAGIVQSTVMPCALVDKAVFGLLRICQRLLPYKENLADELLRSLQLVLKLDARVADAYCEQITQE
VSRLVKANAPHIRSQLGWRTITSLLSHTARHPDASEAGFDALLFIMSDGAHLLPANYVLCVDASRQFAESRVGQA
ERSVRALDLMTGSVDCLARWASEAKEAMGEEEAVRMSQDIGEMWLRLVQGLRKVCLDQREEVRNHALLSLQKCLT
TGVDGIHLPHGLWLECFDMVIFTMLDDLLEIAQGHSQKDYRNMEGTLILAMKLLPKVFLQLLPDLSQLTTFCKLW

LGVLSRMEKYIKVKVRGKKSEKLQELVPELLKNTLLVMKTRGVLVQRSALGGDSLWELTWLHVNNIAPSLQAEVF
PDQSLEEPSHGDEVGGDLVPDETDRVPSANTTSPEGPAGGGLDVV*
**>Morno_LOC21408060**
MGRLKSGIKTIEEREDCDGYSSNATLACMINSEIGAVLAVMRRNVRWGVRYMSGDDELEHSLIQSLRALRKTIFT
WKSHWNTIDPAEYLVPFLDVIRSDETGAPITGVALSSVYKILTLDVIDQNTVSVKDAMHLLVDAVTSCRFEVTDP
VSEEVVLMKILQVLLACMKSKASVVLSNQHVCTIMNACFRIVHQAGSKGELLQRIARHTMHELVRCIFSHLPDVG
NSEKALVDGVDTGDYKSTGLSTEFELGNRQLENGNMMSEYDGQTLPANHALNASVGQPAIGMDENPIGTGAGKDT
VTYDLPMMTEPFGVPCMVEIFHFLCSLLNVVEHVGMGPRSNTIAFDEDVPLFALGLINSTIELVGPSICHHPRLL
SLIQNELFRNLMQFGLSMSPLTLSMVCSIVLNLCHQLRAELKFQLEAFFSCVILRIAQCKYGASYQLQEVAMEAL
VDFCRQKTFMVEMYANFDCDITCSNVFEELANLLSKSAFPVNSPLSSMHIFALDGLIAVIQEMAERVSNGSVGLE
QTSVNLEEYTPFWMVKCDNYKDASQWVPFVRRRKYIKKRLMIGADHFNHDPKKGLEFLQGTHLLPDKLDPQSVAC
FFRYTPGLDKNLVGDFLGNHDEFCIQVLHEFARTFDFEDMSLDTALRLFLETFRLPGESQKIQRVLEAFSERYYE
QSPHILVDKDAALLLSYSLILLNTDLHNAQVKRKMTEEDFIRNNRRINGGNDLPREFLSELYHSICQNEIRTTPE
QGPGFLEMTPSRWIELIHKSKKTAPFIVSDSTAYLDQDMFAIISAPTIAAISVVFDQAEHEEVYQTCIDGFLAVA
EISACHHLEDVLDDLVVSLCKFTTLLNPSSLEESVLAFGDDTKARMATVTVFTISNSYGDYIRAGWRNIFDCILR
LHKLGLLPARVASDAADEPELPTDNGHGKPLKNSLSSSNMSSVTTPKRSYGLMGRFSQLLSLDTEEPRIQPTEEE
LAARQRTLQTIQQCHIDSIFTESKFLQAESLLQLVRALIWAAGRPQKVGSSLDDEDSAVFCLELLIAITLNNRDR
IVLLWQCVYEHIAGIVQSTVMPCALVEKAVFGLLRICQRLLPYKENIADELLRSLQLVLKLDARVADAYCEQITQ
EVSRLVKANAPHIRSQLGWRTITSLLSITARHPEASDAGFDTLLFIMSDGAHLLSANYVLCVDASRQFAESRVGQ
AERSVVAVDLMVGSINCLAQWACEAKDTMGEEEALRMCQDVGEMWFRVVQGLRKVSLDPREEVRNHALLSLQKCL
KGVDGIHFPHGLWLQCFDMGIFTLLDDLLEIVQGQSLKEYRNMEGTLVLAMKLLTKVFLQLLPDLSQLTTFGKLW
LGVLSRMENYIKVKVRGRKSEKLQKLIPELLKNTLLVMKTTGVLAHKNALGGDSLWELTWLHANNIAPSLQSELF
PDQSLEEPSHGGKAGGDVVSDETGPHATLTASTEVPSGGG*
**>Morno_LOC21403706**
MDKHNELPKSKRKELSLSCMLNTEVGAVLAVIRRPSDPNSYGHLSTAADDTIDSALIQSLKSLRALVFNPQQEWR
TVDPLIYLSPFLDVIQSDDVPAAATGIALSALLKILKLGVFDEKTPGVRQAISAVVAAITACRLEKTDPFFEDAV
MMKILQVLTETMHHPASALLSDQYVCIIVNTCFQVVQQSIGRGDLLQRSARYAMHELIQIIFSQLPEIEARDGEG
AFETDVEDADLNGYTDSRYRIRCAVDIFHFLCSLLNVVEQLETEGSTLRTAEEDIQFFALVLINSAVELSGDGIG
RHPKLLRMVQDDLFHHLIHYGTCSSPLVLSMICCTVLNAYHFLRRFIRLQIEAFFGFVLLRISTTQGVPLQLQEV
SLEGIINFCRHPTFIIETYVNYDCDPHCRNVFEEIGKLLCKHSFPVSSPLTSLKIQAFEGLMVIINNIADNIDRE
DDSTPFEPYPIEISEYIPFWEDKPSDDSMNWVDFLRLSRAQKRKLLIAAYHFNHDEKKGLEYLKFSKLSPDPPDP
KSYAFFFRYTPGLDKNMLGDYLGDPDEFHIQVLKEFTETFEFTGMVLDTALRNYLESFRLPGESQKIQRILEAFS
DRFYELKTEDIFESKDAVLILCYSLIMLNTDQHNPQVKKKMTEDEFIRNNRAINGGKDLPREYLSELFNSISNNA
ITIFSQSGQSVEMNPSRWIELMNRSRTVQPFSLSNFDRRLGRDMFSSIAGPAVASLYAFFEHADEEELLHECVEG
LFSIARIAQYGLEETLDELLASFCKFTTLLNPYASAEETLFAFSHDLKPKIATLAVFTIANNFANSIRGGWKNIV
ECLLKLKRLKLLPQSVIEFDATSPTTSGSGATFLSNETKFGSRQASGMINRFSHFLSMESIEESLSLGVSEFEQN
LKIIKQCRIGNIFSNTSSLHEDALINLGRSLIFAAAGKGQKFSTPVEEEETVGFCWDLIIAITLANIHRFQSFWP
AFHDHFLAIAQFPMFSPIPFAEKAIVGLFKVCLKILGTYQPEKSPEELIFKSINLMWKLEKEILDTCCTFITQSV
NKILIEFPANLNSQLGWKSVLHLLSITGRHSETYEQGVETLITLISAETHVSKLNYAYCIDCAFSFVALRNSPLE
KNIKILDLLANSVSKLVQWYKTQCLELENSYSVTSSASSSSLEDNNFKGHGNSLNLFMNLFIKLGEALRKTCLAR
REEIRNHAVLALQKSFQLAEELDFTPSNCINCFNLVIFAMVDDLNEKMLEYSRRENAERETRGMEGTLKIAMEFL
TEVYLRYVKQISSCPGFRTFWLGVLRRMDTYMKADLGEYGESKLPETVPDLLRKIIKEMKEKEILVPKEGDDLWD
ITYIQIQWIAPSLKEELFPEESF*
**>Rharu_RR24768**
MGRLKLQTGIKAIEEEPEDFDATYCSKANLACMINSEIGAVLAVMRRNVRWGGRYVSGDDQLEHSLIQSLKAVRR
QIFSWQHKWHTINPAVYLQPFLDVIRSDETGAPITGVALSSVYKILTLDVIDQNTVNVEDAMNLLVDAVTSCRFE
VTDPASEEVVLMKILQVLLACMKSKASDMLSNQHVCTIVNTCFRIVHQAGTKGELLQRIARNTMHELVRCIFSHL
SDVNKTEGALVNGISTIKHEVAGLTNEQAFPTRELDNGNMSSEYDGHPLSMNLASNTSMGAEAAGMDEDIIGVGT
GTNAVPYDLHLMTEPYGVPCMVEIFHFLCSLFNVVEHIGMDPRSNTIAFDEDVPLFALGLINSAIELGGPSIRHH
PRLLSLIQDELFRNLMQFGLSMSPLILSMVCSIVLNLYHHLRTELKLQLEAFFSCVILRLAQSRYGASYQQQEVA
MEALVDFCRQKTFMVEMYANLDCDITCSNVFEDLANLLSKSAFPVNCPLSSMHILALDGLIAVIQGMAERIGNGT
GGLEYTPVNLEEYTPFWMVKCDNYNDPNHWVMFVRRKKYIKRRLMIGADHFNRDPKKGLEFLQGTHLLPDKLDPQ
SVACFFRYTAGLDKNLVGDFLGNHDEFCVQVLHEFAGTFDFQDMNLDTALRLFLETFRLPGESQKIQRVLEAFSE
RYYEQSPHILANKDAALLLSYSLIMLNTDQHNVQVKKKMTEEDFIRNNRHINGGNDLPREYLSELYHSICKNEIR
TIPEQGTGFSEMTPSRWIDLMHKSKKSVPFIVSDSKAYLDHDMFAIMSGPTIAAISVVFDHAEHEEVYQTCIDGF
LAVAKISACHHLEDVLDDLVVSLCKFTTLLNPSSVEEPVLAFGDDAKARIATVTVFTIANRFGDYIRTGWRNILD
CILRLHKLGLLPARVASDAADESELSADTGHGKPLTNSLSTTHMPSVGTPRRSSGLMGRFSQLLSLDTEEPRSQP
TEQQLAAHQRTLQTIQNCHIDSIFTESKFLQAESLLQLARALIWAAGRPQKGNSSSEDEDTAVFCLELLIAITLN
NRDRIVLLWQGVYDHIASIVQSTIMPSALVEKAVFGLLRICQRLLPYKENLADELLRSLQLVLKLDARVADAYCE
QITQEVSRLVKANASHIRSQLGWRTITSLLSITARHPEASESGFDALLFIMSDGAHLLPANYVLCVDASRQFAES
RVGQAERSVRSIDLMAGSVDCLSRWASKAKEAMGEEEATKMTQDIGEMWLRLVQALRKVCLDQREEVRNHALLAL
QKCLTSVEGIHLPHALWLQCFDLVIFTMLDDLLEIAQGHSQKDYRNMEGTLVIAMKLLPEVFLQLLAHLSQLTTF

CKLWLGVLSRMEKYMKVKVRGKKSEKLQELVPELLKDTLLVMRTRGVLVQRSALGGDSLWELTWLHVNNIAPSLQ
NEVFPDQDLEQSQSKEGEARNDGISDETGHVYSTS*
**>Rharu_RR24766**
MGRLKLQSGIKAIEEEPEDCDATCSNKATLACMINSEVGAVLAVMRRNVRWGGRYMSGDDQLEHYLIQSLKALRK
QIFSWQHQWHTINPAVYLQPFLDVIQSDETGAPITGVALSSVYKILTLDVIDQNTVNVEDAMHLLVDAVTSCRFE
VTDPASEEVVLMKILQVLLACMKSKASVMLSNQHVCTIVNTCFRIVHQAGSKGELLQRIARHTTHELVRCIFSHL
ADVDNTDGALVNGVNTVKRDIDGLNNQYALRSRQLENGNMSTKHDNQSVSTTLPSNASIGSVAAGMDEDTIGVGT
GKSTDPYDLHLMSEPYGVPCMMEIFHFLCSLLNVVENIGLSPRSNTIAFDEDVPLFALGLINSAIELSGPSIRHH
PRLLTLIQDELFCNLMQFGLSMSSLILSVVCSIVLNLYHHLRTELKLQLEAFFSCVVLRLAQSRFGASYQQQEVA
MEALVDFCRQKTFMVEMYANLDCDITCSNVFEDLANLLSKSAFPVNCPLSSMHILALDGLIAVIQGIAERIGNGL
VGSEHSPVNLEEYTPFWMVTCDNYNDPNHWVPFVRRRKYIKRRLMIGADHFNRDPKKGLEFLQGTHLLPDKLDPQ
SVACFFRYTAGLDKNLVGDFLGNHDEFCVQVLHEFAGTFDFRDMNLDTALRLFLETFRLPGESQKIQRVLEAFSE
RYYEQSPRILVNKDAALVLSYSIILLNTDQHNVQVKKKMTEEDFIRNNRRINGGNDLPREFLSELYNSICQNEIR
TIPEQGAGFPEMTPSRWIDLMHKSKKTAPFIVSDCKAYLDQDMFAIMSGPTIAAISVVFDHAEHEEVYETCINGF
LAVAKISACHHLEDVLDDLVVSLCKFTTLLNPSSVEEPVLAFGDDSKARMATVTVFTIANKYGDYIRTGWRNILD
CILRLHRLGLLPARVASDAADESELSADTGHGKPLMNSLSTAHMPSMGTPRRSSGLMGRFSQLLSLDTEEPRSQP
TEQQLAAHQRTLQTIQKCHIDSIFTESKFLQAESLLQLARALTWAAGQPQKGNSSPEDEDTAVFCLELLIAITLN
NRDRIVLLWQSVYEHIASIVQSTILPSALVEKAVFGLLRICQRLLPYKESLADELLRSLQLVLKLDARVADAYCE
QITQEVSRLVKANASHIRSHLGWRTITSLLSITARHPEASEAGFDALLFIMSDGAHLLPANYVLCVDASRQFAES
RVGQAERSVCALDLMGGSVDCLARWACEGKEAMGEEEAVRMTQDIGEMWLRLVQGLRKVCLDQREEVRNHALLSL
QKCLTRVDGIHLPHALWLQCFDLVIFTMLDDLFEIAQGHSQKDYRNMEGSLILAMRLLPKVFLQLLPDLSQLTTF
CKLWLGVLSRLEKYMKVKVRGKRSEKLHELVPELLKNTLLVMKTSGVLVQRSALGGDSLWELTWLHVNNIASSLQ
SEVFPDQDSEQLESKQGETMEGLTSNAASDVPPTEATLSEVAAGGG*
**>RharuGNL2_LG7**
MEEEQDHKQEHEREKELSKYKRRELGISCMLNAEVGAVLAVIRRPQDTNSHYAAPSDHDTVDSSLIQSLKSLRSI
IFNPQQEWRTMDPSIYLSPFLDVIQSDDIPAIATGVSLSALLKILKLGVFDEKTPGAKDAINSVVTAITSCRLEK
TDPVSEDAVMLKILQVLTGTMSHPASVSLTDHAVCTLINTCFHVVQQSVSRGDLLQRNARYTMHELIQIIFSRLT
EVEEACESDTEDGDIGGDLDSGYGGRCAVDIFQFLCSLLNVVELVEADGSPSPTADEDIQLFALVLINSAIELSG
DEIGKRPKLLRMVQDDLFHHLIHYGACSSPLVLSMICSTVLNIYHFLRRFCRLQIEAFFDFVLLRVSTMGDSLQL
QEVALEGIVNFCRQTKFIVEIYVNYDCDPLCRNVFEEIGKLLCKHSFPTNGPLTTLKVQAFEGLIVLIHNIADNI
DREDDSSPYGPYPVEVSEYRPFWEDKPKEDLETWVEFvrvrkaqkrkvvIAGHHYNRDEKKGLDYLKLSQLVSDP
PDPKAYAFFFRYTPGLDKNSIGDYLGDPDELHLNVLKEFTETFQFSGMILDTALRTYLETFRLPGESQKIQRILE
AFADRFYEQQTTEIFATKDSVFILCYSVIMLNTDQHNPQVKKKMTEEEFIRNNRAINAGKDLPREYLSELFQSIA
NNAITLFGQTGATIEMNPSRWVELINRSKTVQPFTLCDFDRRMGRDMFACIAGPSVAALSAFFEYADEEDLLHEC
LEGLFSVARIAQYGLEDTLDELLASFCKFTTLLNPYASAEETLFAFSNDLKPKMATLAVFTIANSFGNSIRGGWR
NIVDCllklkrlkllPQSVIEFDsastsssdapassesGVIFPAHDSTFRSRQNSGMISRFSHFLSLESTEDALS
LGISEFEQNLKIIKQCRIGNIFSNTSSLPDDALLNLGRSLIFAAAGKGQKFSTPVEEEETVGFCWDLIIAVTLAN
IQRFQSFWPSFHDYLLTVAQFPLFSPIPFAEKAIIGLFKVCLKILGTYQPEKLPEDLIFKSINLMWKLDKEILDT
CCEFITQSINKILIEYPANLQTQLGWKSALHLLSVTGRHPETYEQGVETLIMLLSDGFHVSRLNYAYCIDCAFGF
VALRNSPLEKNLKILDLLSDTVNMLVQWSRNQCTEPGnnyssasntstssLEENFKGFAPGNFAMNLFVKLGDAL
RKTSLARREEIRNQAVLSLRKSFLLAEELEFTPANSMNCFNLVIFAMVDDLHEKMLEYSRRDNAEREMRSMEGTL
KIAMELLTDVYLQFLSQISQSPGFRTFWLGVLRRMDTCMKADLGDYGQSKMQDLIPELLTKIITSMKEKEILVQK
EGDDLWEITYIQIQWIAPTVKDQLFPDQDF*
**>MDP0000129069**
MGRLKLQTGIKAIEEEPEDCEAANSNKATLACIINSEIGSVLAVMRRNVRWGSRYISGDDQLEHSLIQSLKALRK
QIFSWQHQWHTINPVVYLQPFLDVIRSDETGAPITGAALSSXYNILTLDVIDQNSVNVEDAMNLLVDAITSCRFE
VTDPASEEVVLMKILQVLLACMKSKASVMLSNQHVCTIVNTCFRIVHQAGTKGELLQRIARHTMHELVRCIFSHL
PDVHDTERALLNGSTTRKQEIAGLNNEYAFGSRQLENGNLSSGYDGQPLSTSPPSNASSGLVAPGIDESVLTVST
EKEAVQYDLHIMTEPYGVPCMVEIFHFLCSLLNVSEHMGVGPRSXTITFDEDVPLFALVMINSAIELGGSHIQNH
PKLLSLVQDELFRNLMQFGLSTSPLILSMVCSIVLNLYHHLRTELKLQLEAFFSCVILRLAQSKYGASYQQQEVA
MEVLVDFCRQKKFMVEMYANLDCDITCSNVFEELANLLSKSAFPVNCPLSSIHILALDGLIAVIQGMAERVGNGS
VSAHTPVNLEEYTPFWMVKCDNYSDPSNWVPFVRRRKYIKRRLMIGADHFNRDPKKGLEFLQGTYLLPDKLDPQS
VACFFRYTAGLDKNLVGDFLGNHDEFCVQVLHKFAGTFDFADMNLDTALRLFLETFRLPGESQKIQRVLEAFSER
YYEQSPLILANKDAALLLSYSLIMLNTDQHNVQVKKKMTEEDFIRNNRHINGGSDLPREFLTELYHSICKNEIRT
TPEQGAGYPEMTPSRWIDLMHKSKKNAPFIMSDSRAYLDHDMFAIMSGPTIAAISVVFDHAEHEEIYQTCIDGFL
AVAKXSACHHLEDVLDDLVVSLCKFTTLLNPSSVEEPVXAFGDDTKARMATVTXFTIANRYGDYIRTGWRNILDC
ILRLHKLGLLPARVASDAADESEFSADTGPGKPITNSLSSVHVSSIGTPRRSSGLMGRFSQLLSLETEEPRSQPT
EQQLAAHQRTXQTIQKCHIDSIFTESKFLQAESLLQLARALVWAAGRPQKGNSSPEDEDTAVFCLELLIAITLNN
RDRIDLLWQGVYEHISTIVQSTVMPCALVEKAVFGLLRICQRLLPYKENLADDLLKSLQLVLKLDARVADAYCEQ
ITQEVSRLVKANASHIRSHGGWRTITSLLSITARHPDASEAGFNALFFIMSEGTHLLPANYVVCVDASRQFAESR
VGQAERSVCALDLMAGSVDCLARWACEAKQAVNEEEAVKMSEDIGEMWFRLVQALRKVCLDQREDVRNHALSLLQ
KCLTGVDGIPLLHGLWLQCFDVVIFTMLDDLLEIAQGHSQKDYRNMEGTLILAMELLSKVFLQLLPDLSQLTTFC

KLWLGVLSRMEKYMKVKVRGKKSEKLQDQVPKLLKNTLLVMNLKGVLVQRSALGGDSLWELTWLHVNNIAPSLQS
EVFPDQGSEQPETKLGENRGLVSDEKGNVLPTEMASPEVSGTAN*

**>MDP0000427621**

MGRLKLQTGIKAIEEESEDFEATNSNKATLACIINSEIGSVLAVMRRNVRWGGRYISGDDQLEHSLVQSLKALRK
QIFLWQHQWHTINPVVYLQPFLDVIRSDETGAPITGVALSSVYNILTLDVIDQNSLNVEDAMNLLVDAITSCRFE
VTDPASEEVVLMKILQVLLACMKSKASVMLSNQHVCTIVNTCFRIVHQAGTKGELLQRIARHTMHELVRCIFSHL
PDVHDTEPTLSNGSTTRKQEIAGLNNEYAFGSRQLENGNVSSGYDSQPLSTSPASNASSGLVAPGIDENALAVST
GKEAVQYDLLLMTEPYGVPCMVEIFHFLCSLLNVSEHMGMGPKSNTIAFDEDVPLFALVLINSALELGGSHIQNH
PKLLSLVQDELFRNLMQFGLSTSPLILSMVCSIVLNLYHHLRTELKLQLEAFFSCVXLRLAQSXYGASYQQQEVA
MEXLVDFCRQKXFMVEMYANLDCDITCSNVFEELANLLSKSAFPVNCPLSSIHILALDGLIAVIQGMAERVGNGS
VSAHTPVXLEEYTPFWMVKCDNYSDPSNWVPFVRRRKYIKRRXMIGADHFNRDPKKGLEFLQGTYLLPDKLDPQS
VACFFRYTAGLDKNLVGDFLGNHDEFCVQVLHKFAGTFDFTDMNLDTALRLFLETFRLPGESQKIQRVLEAFSER
YYEQSPLILANKDAALLLSYSLIMLNTDQHNVQVKKKMTEEDFIRNNRHINGGSDLPREFLTELYHSICKNEIRT
TPEQGAGYPEMTPSRWIDLMHKSKKXAPFIMSDSRAYLDHDMFAIMSGPTIAAISVVFDHAEHEEIYQTXIDGFL
AVAKXSACHHLEDVLDDLVVSLCKFTTLLNPSSVEEPVXAFGDDTKARMATVTXFTIANRYGDYIRTGWRNILDC
ILRLHKLGLLPARVASDAADEXEFSADTGPGKPIXNSLSSVHVXSIGTPRRSSGLMGRFSQLLSLETEEPRSQPT
EQQLAAHQRTXQTIQKCHIDSIFTESKFLQAESLLQLARALXWAAGRPQKGNSSPEDEDTAVFCLELLIAITLNN
RDRIDLLWQGVYEHISXIVQSTVMPCALVEKAVFGLLRICQRLLPYKENLADDLLKSLQLVLKLDARVADAYCEQ
ITQEVSRLVKANASHIRSHGGWRTITSLLSITARHPDASEAGFNALFFIMSEGTHLLPANYVVCVDASRQFAESR
VGQAERSVCALDLMAGSVDCLARWACEAKQAVNEEEAVKMSEDIGEMWFRLVQALRKVCLDQREDVRNHALSLLQ
KCLTGVDGIPLLHGLWLQCFDVVIFTMLDDLLEIAQGHSQKDYRNMEGTLILAMELLSKVFLQLLPDLSQLTTFC
KLWLGVLSRMEKYMKVKVRGKKSEKLQDQVPKLLKNTLLVMNLKGVLVQRSALGGDSLWELTWLHVNNIAPSLQS
EVFPDQGSEQPETKLGENRGLVSDEKGNVLPTEMASPEVSGTAN*

**>MDP0000410984**

MGRLKLQTGIKAIEEESEDFEATNSNKATLACIINSEIGSVLAVMRRNVRWGGRYISGDDQLEHSLVQSLKALRK
QIFLWQHQWHTINPVVYLQPFLDVIRSDETGAPITGVALSSVYNILTLDVIDQNSLNVEDAMNLLVDAITSCRFE
VTDPASEEVVLMKILQVLLACMKSKASVMLSNQHVCTIVNTCFRIVHQAGTKGELLQRIARHTMHELVRCIFSHL
PDVHDTEPTLSNGSTTRKQEIAGLNNEYAFGSRQLENGNVSSGYDSQPLSTSPASNASSGLVAPGIDENALAVST
GKEAVQYDLLLMTEPYGVPCMVEIFHFLCSLLNVSEHMGMGPKSNTIAFDEDVPLFALVLINSALELGGSHIQNH
PKLLSLVQDELFRNLMQFGLSTSPLILSMVCSIVLNLYHHLRTELKLQLEAFFSCVXLRLAQSXYGASYQQQEVA
MEXLVDFCRQKNFMVEMYANLDCDITCSNVFEELANLLSKSAFPVNCPLSSIHILALDGLIAVIQGMAERVGNGS
VSAHTPVXLEEYTPFWMVKCDNYSDPSNWVPFVRRRKYIKRRXMIGADHFNRDPKKGLEFLQGTYLLPDKLDPQS
VACFFRYTAGLDKNLVGDFLGNHDEFCVQVLHKFAGTFDFTDMNLDTALRLFLETFRLPGESQKIQRVLEAFSER
YYEQSPLILANKDAALLLSYSLIMLNTDQHNVQVKKKMTEEDFIRNNRHINGGSDLPREFLTELYHSICKNEIRT
TPEQGAGYPEMTPSRWIDLMHKSKKXAPFIMSDSRAYLDHDMFAIMSGPTIAAISVVFDHAEHEEIYQTXIDGFL
AVAKXSACHHLEDVLDDLVVSLCKFTTLLNPSSVEEPVXAFGDDTKARMATVTXFTIANRYGDYIRTGWRNILDC
ILRLHKLGLLPARVASDAADEXEFSADTGPGKPIXNSLSSVHVXSIGTPRRSSGLMGRFSQLLSLETEEPRSQPT
EQQLAAHQRTXQTIQKCHIDSIFTESKFLQAESLLQLARALXWAAGRPQKGNSSPEDEDTAVFCLELLIAITLNN
RDRIDLLWQGVYEHISXIVQSTVMPCALVEKAVFGLLRICQRLLPYKENLADDLLKSLQLVLKLDARVADAYCEQ
ITQEVSRLVKANASHIRSHGGWRTITSLLSITARHPXASEXGFXALFFIMSEGTHLLPXNYXXCVDASRQFAESR
VGQXERSVCALDLXAGSVDCLARWAXEAKQAVNEXEAVKMSXDIGEMWFRLVQALRKVCLDQREDVRNHALSLLQ
KCLTGVDGIPLXHGXWXQCFDVVIFTMLDDLLEIAQGHSQKDYRNMEGTLILAMXLLSKVFLQLLPXLSQLTTFC
KLWXGVLSRMEKYMKVKVRGKKSEKLQDQVPELLKNTLLVMNLKGVLVQRSALGGDSLWELTWLHVNNIAPSLQS
EVFPDQGSEQSATKLGENGGLVSEEKGNVLPTEMASTEVSGTAGPATAAKPDFRGFRHEIAVDLEGRRTIG*

**>MDP0000802196**

MGRLKLQSGIQAIGEEPGECDATYPNHATLACIINSEIGTVLAVMRRNVRWGGRYMSGDDQLEHPLIQSLKALRK
QIFSWKHHMHSVNPSEYLQPFLDVIRSDETGAPITGVALSSVYNILTLDVIDQNSLNVEDAMHLLVDAITGCRFE
VTDPASEEVVLMKILQVLLACMKSKASIILTNQHVCTIVNTCFRIVHQAGTKGELLQRIARHTMHELVRCIFSHL
PDVQPAERALPNGSXTINGEIAGKNNEHASGSRQLENGNMISEFDNQLLSTNPALNTSSGLVESGMDEKTAGASS
GKEAGQYDVQLMAEPFGVPCMVEIFNFXCSLLNVVEHIGMGPRSNTIAFDEDVPLFALGLVNCAIELSGSSIRHH
PKLLSLVQDELFQNXMQFGLSTSPLILSMVCSIVLNLYHHLRTELKLQLEAFFSCVILRLVQSRYGASYQQQEVA
MEAVVDFCRQKTFMVEMYANLDCDITCNNVFEELANLLSKSAFPVSLPLSSIHILALDGLIAIIQGMAERAGNGS
VSAVQAPPTNLDEYTPFWLMKCDNYSDPNHWVPFVRRRKHIKRRLMIGADHFNHDPKKGLEFLQGTHLLPDKLDP
ESVACFFRYTAGLDKNLVGDFLGNHDEFCIQVLNKFAGTFDFQDMHLDTALRLFLETFRLPGESQKIQRVLEAFS
ERYYEQSPQILANKDAALLLSYSIIMLNTDQHNVQVKKKMTEEDFIRNNRHINGGDDLPREFLSELYHSICKNEI
RTTPEQGAGYPEMNPSRWIDLIHKSKKSAPFILSDTRAYLDQDMFAIMSGPTIAAISVVFDHAEHEEIYQTCIDG
FLSVAKIAACYHLEDVLDDLVVSLCKFTTLLNPSSYEEPVIAFGDDPKARMSTVTVFTFANTYGDYIRTGWRNVL
ECILRLHKLGLLSAFVASEAAGDSESSADTGHGKPITNSLSSVPMPPVSTPRKSSGFMGRFSQLLSLDTEEPRSQ
PTEEELAAHQRTLQTIQKCHIDGIFSESKFLQAESLLQLAQALIWAGGRPQKGSSSPEDEDTAVFCLELLIAITL
NNRDRIMLLWQIVYEHISNIVQSTVMPCALVEKAVFGLIQICQRLLPYKEDLADELLRSLQLVLKLDARVADAYC
EQITQEVGRLVKANASHIRSPLGWRIVTSLLSITARHXEASEAGFDALFFIMSEGTHLLPANYVLCVDASRQFAE
SRVGEVERSVCALDLMAGSIDCLARWACDAKQSMNEEVALKMSQDXGQMWLRLVQGLRKVCLDQREEVRNHALSI
LRRCLTGVDGIPLPHGLWLQCFDMVIFTMLDDLLEIAQGHSPKEYRNMEGTLILALKLLSKVFLELLPELSQLTN

FSKLWLGVLNRMENYMKVKVGGRKCEKLRDQVPELLKNTLLVMILRGVLVEKSDGGDDSLWELTWLHVNKIAPLL
QSEISRDPILEQSDTKQQGESGRVSDXTGTLIPTETAAAEGSSSGG*
**>MDP0000427619**
MGRLKLRSGIQEIEEEHGECDATHSNHATLACIINSEIGTVLAVMRRNVRWGGRYVSGDDQLEHPLIQSLKALRK
QIFSWKHHMHSVNPSAYLRPFLDVIRSDETGAPITGVALSSVYXILTLDVIXQNSLNVEDAMHLLVDAITGCRFE
VTDPASEEVVLMKILQVLLACMKSKASXILTNQHVCXIVNTCFRIVHQAGTKGELLQRIARHTMHELVRCIFSHL
PDVHHSEHALPNGSNTIKGEIAGQNNEHASGSRQLENGNMISEFDXQLLSXNPALXXSSGLVESGXDEKTAGXSS
XKEAXQYDVQLMAEPFGXPCMVEIXXFLCSLLNVVEHIGMGPRXNTIAFDEDVPLFAXGLVNCAIELXGSSIRHH
PKLLSLVQDELFQNXMQFGLSTSPLILSMVCSIVLNLYHHLRTELKLQLEAFFSCVILRLVQSRYGASYQQQEVA
MEAVVDFCRQKTFMVEMYANLDCDITCNNVFEELANLLSKSAFPVXLXLSSIHILALDGLIAIIQGMAERAGXGS
VSAVQAPPXNLDEYTPFWXMKCDNYSDPNHWVPFVRRRKHIKRRLMIGADHFNHDPKKGLEFLQGTHLLPDKLDP
ESVACFFRYTAGLDKNLVGDFLGNHDEFCIQVLNKFAGTFVFQDMHLDTALRLFLETFRLPGESQKIQRVLEAFS
ERYYEQSPQILANKDAALLLSYSIILLNTDQHNVQVKKKMTEEDFIRNNRHINGGDDLPLEFLSELYHSICKNEI
RTTPEQGAGYSEMNPSRWIDLIHKSKKNAPFIESDTRAYLDQDMFAIMSGPTIAAISVVFDHAEHEEIYQKCIDG
FLYVAKIAACYHFEDVLDDLVVSLCKFTTLLNPSSYEEPVLAFGDDPKARMSTVTVFTIANTYGDHIRTGWRNVL
DCILRLHKLGLLSAFVASEAAGDSEFSADTGHGKAITNSLSSVPMPPLSTSRKSSGLMGRFTQLLSLNTEEPRSQ
PTEEELVAHQRTLQTIQKCHIDGIFSESKFLQAKSLLQLAQALIWAGERPQKGSSSPEDEDTAVFCLELLIAVTL
NNRDRIMLLWQIVYEHISNIVQSTVMPCALVEKAIFGLLRICQRLLPYKEDLADDLLRSLQLVLKLDARVADAYC
EQITQEVGRLVKANASHIRSQLGWHITTSLLSITAPHPEASEAGFDALFFIMSEGAHLLPANYVFCVDASRQFAE
SRVGEVARSVCALDLMAGSVDCLSRWASDANQSMNGEEAVKMSQDIGQMWLRLVQGLRKVCLDQREEVRNHALSM
LRKCLTGVDGIPLPHGLWLRCFDTVIFTMLDDLLEIAQGQSPKEYRYMEGTLILALKLLSKVFLELLPELSQLTT
FSKLWLGVLNRMENYMKVKVGGRKCEKLRDQVLELLKNTLIVMILRGVLLEKGDVGDDNLWELTWLHVNKIAPSL
QSEVSRDPFLEQSETKQGESGRVSDATGTILPTETIPAEGSGGGDAFFLPGGWLRLESPVVDDCRGLKSHSTWKL
GTCKGKTGGTRKNQITYVIVRQGELGPLTYAPPQPPKW*
**>MDP0000565847**
MEKDPISEENGKSSRNKQTKFKRKELGLSCMLNTELGAVLAVIRRPLNPTSHCFAAPEDTIDPSLLQSLKSLRAL
IFNPXQEWRTIDPSIYLSPFLDVIQSEDVHAAATRVALSATLKIIKLGVFDEKTPGAKEAINTMVTSITTCRLER
TDSVNEDAVMLSILQVLTGIMNHRASVLLSDQSVCTIVNTCFQVVQQSANRGDLLMRSARYTMHELIQIIFSRLP
EIDFRDDESLASTDTEDADADDGTMDSGYGIRCAVDIFHFLCSLLNVVQVVETDQGSTVQTANEDVQLFALVLIN
SAIDLSGDGIGSHSKLLRMMKDDLFHHLVHYGTRSSPLVFSMICSTVLNMYHFLRRFIRLQLEAFFTFVLFRVSA
PGVSIQLQEVALEGIINFCRQLTFVVEVYVNYDCDPLCHNVFEEIGKLLCKLSFPVASPLTTLQIQAFEGLVIMI
HNIADSIDKENDTSPSGPYPIEIREYTPFWEDKPKDDSEAWVQFVRLRKAQKRKILIAGHHFNCDEKKGMEYLKL
SNLVSDPPDPKTLAYFFRYTPGLDKTMIGDYLGDPDEFHIKVLKEFTGAFGFSGMNLDSALRTYLETFRLPGESQ
KIQRILEAFSESFFEQQSADVFVNKDTVFILCYSLIMLNTDQHNPQVKKKMTEEEFIRNNRAINGGKDLPREYLS
ELFQSISNNAITLFGQSGMTVLMNPSRWIELMNXSKTXQPFILCDFDRRLGRDMFACIAGPSVATISAFFEQAEE
EELLHECIEGLFSVARIAQYGLEDTLDELIATFSRFTTLLNPYASAEETLFAFSKDLKPRMATLAVFTIANNFGE
SIRGGWRNIVDCLLKLKRLKLLPQNVLEFDASTSSDAQPTSESGVIFPGHDPKFGGRQASGMISRFSHFLSLESP
EDAVSLGMSEFEQNLKVIKQCRIGNIFXTSSNFPEDSLLNLGRSLIYAAAGKGQKFSTPVEEEEAVXFCWDLMVA
ISLANVHRFQAFWSSFNDYLLAVAQFPMFSPIPFAEKAIVGLFKVCLKLLGTYQPDRVPEELIFKSINLMWKLEK
EILDTCSESITQSLNKILVEYPANLQTQLGWKSVLHLLSVSGRHPDTYEQGVETLIMVMSDGTHVSRTNYAYFID
CAFGFIALKNSPSEKNLKILDLLADSVNLLIQWYRNHYSDPGNNYSIASNTSTSSLEDSKGFGSNNFTMNLFVKL
GEAFRKTSLARREEIRNHAVLSLXXSFQLAEELEFTPTNCINCFNLLIFAMVDDLHEKMLEYSRRENAEKEMRGM
EGTLKIALELLTDVYLQFLIPISQCSGFRTFWLGVLRRMDTCMKADLGSYGESKLPEVVPELLRKLITEMKEKEI
LVPKEGDDLWEITHIQIQWIAPSIKEELFPEESL*
**>Pyrxb_LOC103935740**
MGRLKLQTGIKAIEEEPEDCEATNSNKATLACIINSEIGSVLAVMRRNVRWGSRYISGDDQLEHSLIQSLKALRK
QIFSWQHQWHTINPVVYLQPFLDVIRSDETGAPITGVALSSVYNILTLDVIDQNSVNVEDAMNLLVDAITSCRFE
VTDPASEEVVLMKILQVLLACMKSKASVMLSNQHVCTIVNTCFRIVHQAGTKCELLQRIARHTMHELVRCIFSHL
PDVHDTERALLNGSATRKQEIAGLNNEYAFGSRQLENGNVSSGYDSQPLSTSPASNASSGLVAPRIDESVLTVST
EKEAVQYDLHLMTEPYGVPCMVEIFHFLCSLLNVSEHMGMGSRSNTITFDEDVPLFALVMINSAIELGGSHIQNH
PKLLSLVQDELFRNLMQFGLSTSPLILSMVCSIVLNLYHHLRTELKLQLEAFFSCVILRLAQSKYGASYQQQEVA
MEVLVDFCRQKNFMVEMYANLDCDITCSNVFEELANLLSKSAFPVNCPLSSIHILALDGLIAVIQGMAERVGNGS
VSAHTPVNLEEYTPFWMVKCDNYSDPSNWVPFVRRRKYIKRRLMIGADHFNRDPKKGLEFLQGTYLLPDKLDPQS
VACFFRYTAGLDKNLVGDFLGNHDEFCVQVLHKFAGTFDFADMNLDTALRLFLETFRLPGESQKIQRVLEAFSER
YYEQSPLILANKDAALLLSYSLIMLNTDQHNVQVKKKMTEEDFIRNNRHINGGSDLPREFLTELYHSICKNEIRT
TPEQGAGYPEMTPSRWIDLMHKSKKNAPFIMSDSRAYLDHDIFAIMSGPTIAAISVVFDHAEHEEIYQTCIDGFL
AVAKISACHHLEDVLDDLVVSLCKFTTLLNPSSVEEPVLAFGDDTKARMATVTVFTIANRYGDYIRTGWRNILDC
ILRLHKLGLLPARVASDAADESEFSADTGPGKPMSNSLSSVHVPSIGTPRRSSGLMGRFSQLLSLETEEPRSQPT
EQQLAAHQRTLQTIQKCHIDSIFTESKFLQAESLLQLARALIWAAGRPQKGNSSPEDEDTAVFCLELLIAITLNN
RDRIDLLWQGVYEHISNIVQSTVMPCALVEKAVFGLLRICQRLLPYKENLADDLLKSLQLVLKLDARVADAYCEQ
ITQEVSRLVKANASHIRSHGGWRTITSLLSITARHPEASESGFDALFFIMSEGTHLLPTNYALCVDALRQFAESR
VGQAERSVCALDLMAGSVDCLARWAREAKQAVNEDEAVKMSQDIGEMWFRLVQALRKVCLDQREDVRNHALSLLQ
KCLTGVDGIPLPHVLWLQCFDVVIFTMLDDLLEIAQGHSQKDYRNMEGTLILAMKLLSKVFLQLLPGLSQLTTFC

KLWLGVLSRMEKYMKVKVRGKKSEKLQDQLPELLKNTLLVMNLKGVLVQRSALGGDSLWELTWLHVNNIAPSLQS
EVFPDQGSEQSETKLGENGGLVSEEKGNVLPTEMASTEVSGTAG*
**>Pyrxb_LOC103942956**
MGRLKLQTGIKAIEEESEDFEAMNSNKATLACIINSEIGSVLAVMRRNVRWGGRYISGDDQLEHSLVQSLKALRK
QIFSWQHQWHTINPVVYLQPFLDVIRSDETGAPITGVALSSVYNILTLDVIDQNSLNVEDAMNLLVDAITSCRFE
VTDPASEEVVLMKILQVLLACMKSKASVMLSNQHVCTIVNTCFRIVHQAGTKGELLQRIARHTMHELVRCIFSHL
PDVHDTEHALSNGSTTRKQEIAGLNNEYAFGSRQLENGNVSSGYDSQPLSTSPASNAPSGLVAPGIDENALAVST
GKEAVQYDLHLMTEPYGVPCMVEIFHFLCSLLNISEHMGMGPRSNTIAFDEDVPLFALVLINSALELGGSHIQNH
PKLLSLVQDELFRNLMQFGLSTSPLILSMVCSIVLNLYHHLRAELKLQLEAFFSCVILRLAQSRYGASYQQQEVA
MEALVDFCRQKNFMVEMYANLDCDITCSNVFEELANLLSKSAFPVNCPLSSIHILALDGLIAVIQGMAERVGNGS
VSAHTPVNLEEYTPFWMVKCDNYSDPSNWVPFVRRRKYIKRRLMIGADHFNRDPKKGLEFLQGTYLLPDKLDPQS
VACFFRYTAGLDKNLVGDFLGNHDEFCVQVLHKFAGTFDFADMNLDTALRLFLETFRLPGESQKIQRVLEAFSER
YYEQSPLILANKDAALLLSYSLIMLNTDQHNVQVKKKMTEEDFIRNNRHINGGSDLPREFLTELYHSICKNEIRT
TPEQGAGYPEMTPSRWIDLMHKSKKSAPFIMSDSRAYLDHDMFAIMSGPTIAAISVVFDHAEHEEIYQTCIDGFL
AVAKISACHHLEDVLDDLVVSLCKFTTLLHPSSVEEPVLAFGDDTKARMATVTLFTIANRYGDYIRTGWRNILDC
ILRLHKLGLLPARVASDAADESEFSADTGPGKPVSNSLSSVHVPSIGTPRRSSGLMGRFSQLLSLETEEPRSQPT
EQQLAAHQRTVQTIQKCHIDSIFTESKFLQAESLLQLARALIWAAGRPQKGNSSPEDEDTAVFCLELLIAITLNN
RDRIDLLWQGVYEHISNIVQSTVMPCALVEKAVFGLLRICQRLLPYKENLADDLLKSLQLVLKLDARVADAYCEQ
ITQEVSRLVKANASHIRSHGGWRTITSLLSITARHPDASEAGFNALFFIMSEGTHLLPANYVLCVDASRQFAESR
VGQAERSVCALDLMAGSVDCLARWACEAKQAVNEEEAVKMSEDIGEMWFRLVQALRKVCLDQREDVRNHALSLLQ
KCLTGVDGIPLLHGLWLQCFDAVIFTMLDDLLEIAQGHSQRDYRNMEGTLILAMELLSKVFLQLLPDLSQLTTFC
KLWLGVLGRMEKYMKVKVRGKKSEKLQDQVPKLLKNTLLEMNLKGVLVQRSALGGDSLWELTWLHVNNIAPSLQS
EVFPNQGSEQPETKLGENGGLVSDEKGNVLPTEMVSPEVSGTAN*
**>Pyrxb_LOC103935729**
MGRLKLQSGIQAIGEEPGECDATYSNHATLACIVNSEIGTVLAVMRRNVRWGGRYMSGDDQLEHPLIQSLKALRK
QIFSWKHHMHSVNPSDYLQPFLDVIRSDETGAPITGVSLSSVYNILTLDVIDQNSLNVEDAMHLLVDAITGCRFE
VTDPASEEVVLMKILQVLLACMKSKASIILTNQHVCTIVNTCFRIVHQAGTKGELLQRIARHTMHELVRCIFSHL
PDVHPAERALPNGSNTINGEIAGKNNEHVSGSRQLENGNMISEFDNQLLSTNPALNASSGLVESGMDEKTTGASS
GKEAGQYDVQLMAEPFGVPCMVEIFTFLCSLLNVVEHIGMGPRSNTIAFDEDVPLFALGLVNCAIELGGSSIRHH
PKLLSLVQDELFQNLMQFGLSTSPLILSMVCSIVLNLYHHLRTELKLQLEAFFSCVILRLVQSRYGASYQQQEVA
MEAVVDFCRQKTFMVEMYANLDCDIICNNVFEELANLLSKSAFPVSLPLSSIHILALDGLIAIIQGMAERAGNGS
VSAVQAPPTNLDEYTPFWLMKCDNYSDPNHWVPFVRRRKHIKRRLMIGADHFNHDPKKGLEFLQGTHLLPDKLDP
ESVACFFRYTAGLDKNLVGDFLGNHDEFCIQVLNKFAGTFDFQDMHLDTALRLFLETFRLPGESQKIQRVLEAFS
ERYYEQSPQILANKDAALLLSYSIIMLNTDQHNVQVKKKMTEEDFIRNNRHINGGDDLPREFLSELYHSICKDEI
RTTPEQGAGYPEMNPSRWIDLIHKSKKSAPFIVSDTRAYLDQDMFAIMSGPTIAAISVVFDHAEHEEIYQTCIDG
FLCVAKIAACYHLEDVLDDLVVSLCKFTTLLNPSSYEEPVIAFGDDPKARMSTVTVFTFANTYGDYIRTGWRNVL
DCILRLHKLGLLSAFVASEAAGDSESSADTGHGKPITNSLSSVPMPPVSTPRKSSGFIGRFSQLLSLDTEEPRAQ
PTEEELAAHQRTLQTIQKCHIDGIFSESKFLQAESLLQLAQALIWAGGRPQKGSSSPEDEDTAVFCLELLIAVTL
NNRDRIMLLWQIVYEHISNIVQSTVMPCALVEKAVFGLIRICQRLLPYKEDLADELLRSLHLVLKLDARVADAYC
EQITQEVGRLVKANASHIRSPLGWRIVTSLLSITARHPEASEAGFDALFFIMSEGTHLLPANYVLCVDASRQFAE
SRVGEVERSVCALDLMAGSVDCLARWACDSKQSMNEEETLKMSQDIGQMWLRLDIGQMWLRLVQGLREVCLDQRE
EVRNHALSILRRCLTGVDGIPLPHGLWLQCFDMVIFTMLDDLLEIAQGHSPKEYRNMEGTLILALKLLSKVFLEL
LPELSQLTTFSKLWLGVLNRMEKYMKVKVGGRKCDKLRDQVPELLKNTLLVMFLRGVLVEKSDGGDNSLWELTWL
HVNKIAPSLQSEISRDPILEQSETKQGESVRVSDVTGTSIPTETAAAEGSSSGG*
**>Pyrxb_LOC103942957**
MGRLKLQSGIQEIEEEHGECDATHSNHATLACIINSEIGTVLAVMRRNVRWGGRYLSGDDQLEHPLIQSLKTLRK
QIFSWKHHMHSVNPSAYLRPFLDVIRSDETGAPITGVALSSVYNILTLDVINQNSLNVEDAMHLLVDAITGCRFE
VTDPASEEVVLMKILQVLLACMKSKASVILTNQHVCIIVNTCFRIVHQAGTKGELLQRIARHTMHELIRCIFSHL
PDVHHAEHALPNGSNTINGEIAGQNNEHASGSRQLENGNMISQFDSHLLSTNPALHASSGLVESGIDEKTAGGSS
RKEADQYDVHLMAEPFGIPCMVEIFDFLCSLLNVVEHIGMGPGYNTIAFDEDVPLFALGLVNCAIELGGSSIRHH
PKLLSLVQDELFQNLMQFGLSTSPLILSMVCSIVLSLYHHLRTELKLQLEAFFSCVILRLVQSRYGASYQQQEVA
MEYVVDFCRQKTFMVEMYANLDCDITCNNVFEELANLLSKSAFPVNLLLSSIHILALDGLIAIIQGMAERAGNDS
VSAVQAPPMNLDEYTPFWLMKCDNYSDPTHWVPFVRRRKHIKRGLMIGADHFNHDPKKGLEFLQGTHLLPDKLDP
ESVACFFRYTAGLDKNLVGDFLGNHDEFCIQVLNKFAGTFDFQDMHLDSALRLFLETFRLPGESQKIQRVLEAFS
ERYYEQSPQILANKDSALLLSYSIILLNTDQHNVQVKKKMTEEDFVRNNRHINGGDDLPREFLSELYHSTCKNEI
RTTPEQGAGYPEMNPSRWIDLIHKSKKNAPFIESDTRAYLDQDMFAIMSGPTIAAISVVFDHAEHEEIYQKCIDG
FLSVAKIAACYHFEDVLDDLVVSLCKFTTLLNSSSYEEPVLAFGDDPKARMSTMTVFTIANTYGDHIRTGWRNVL
DCILRLHKLGLLSAFVAGEAAGDSEFSADTGHGKAITNSLSSVPMPPLSTPRKSYGLMGRFTQLLSLDTEEPRSQ
PTEEELVVHQRTLQTIQKCHIDSIFSESKFLQAKSLLQLARALIWAGGRPQKGSSSPEDEDTAVFCLELLIAVTL
NNRDRIMLLWQIVYEHISNIVQSTVMPCALVEKAVFGLLRICQRLLPYKENLADDLLKSLQLVLKLDLDARVADA
YCEQITQEVRRLVKANASHIRSQLGWRITTSLLSITAPHPEASEAGFDALFFIMSEGAHLLPANYVFCVDASRQF
AESRVGEVERSVCALDLMAGSVDCLSRWASDANQSMNEEEAVRMSQDIGQMWLRLVQGLRKVCLDQREEVRNHAL
SMLRKCLTGADGIPLPHGLWLQCFDTVIFTMLDDLLEIAQGHSPKEYRYMEGTLILALKLLSKVFLELLPELSQL

TTFSKLWLGVLNRMENYMKVKVGGRKCEKLRDQVPELLKNTLIVMILRGVLVEKSDVGDDSLWELTWLHVNKIAP
SLQSEVSRDPILEQSETKQGESGRVSDATGTMLPTETVPAEGSSGGG*
**>Pyrxb_LOC103929457**
MEKDPISDENGKSSRNKQTKLRRKELGLSGMLNTELGAVLAVIRRPLNPTSHCFAAPEDTIDPSLLQSLKSLRAX
XXXXXXXXRTIDPSIYLSPFLDVIQSEDVHAAATHFALSATLKIIKLGVFDEKTPGAKEAINTMVTSITTCRLER
TDSVNEDAVMLNILQVLTGIMNHRASVLLSDQSVCTIVNTCFQVVQQSVNRGDLLMRSARYTMHELIQIIFSRLP
EIDLRDDESLASTDTEDADADDGNMDSGYGIRCAVDIFHFLCSLLNVVQVVETDQGSTVQTANEDVQLFALVLIN
SAIDLSGDGIGSHSKLLRMMKDDLFHHLVHYGTRSSPLVFSMICSTVLNMYHFLRRFIRLQLEAFFTFVLFRVSA
PGVSIQLQEVALEGIINFCRQLTFVVEVYVNYDCDPLCHNVFEEIGKLLCKLSFPVASPLTTLQIQAFEGLVIMI
HNIADSIDRENDTSPSGPYPIEIREYTPFWEDKAKDDSEAWVQFVRLRKAQKRKILIAGHHFNCDEKKGMEYLKL
SNLVSDPPDPKTLAYFFRYTPGLDKTMIGDYLGDPDEFHIKVLKDFTGAFGFSGMNLDSALRTYLETFRLPGESQ
KIQRILEAFSESFFEQQSADVFVNKDTVFILCYSLIMLNTDQHNPQVKKKMTEEEFIRNNRAINGGKDLPREYLS
ELFQSISNNAITLFGQSGMTVLMNPSRWIELMNRSKTVQPFILCDFDRRLGRDMFACIAGPSVATISAFFEQAEE
EELLHECIEGLFSVARIAQYGLEDTLDELIATFSRFTTLLNPYASAEETLFAFSKDLKPRMATLAVFTIANNFGE
SIRGGWRNIMDCLLKLKRLKLLPQNVLEFDAASTSSDAQPTSESGVIFPAHDPKFGGRQASGMISRFSHFLSLES
PEDAVSLGMSEFEQNLKVIKQCRIGNIFSTSSKFPEDSLINLGRSLIYAAAGKGQKFSTPVEEEEAVGFCWDLIV
AISLANVHRFQAFWSSFNDYLLAVAQFPMFSPIPFAEKAIVGLFKVCLKLLGTYQPDRVPEELIFKSINLMWKLE
KEILDTCSESITQSVNKILVEYPANLQTQLGWKSVLHLLSVSGRHPDTYEQGVETLIMVMSDGTHISRTNYAYFI
DCAFGFIALKNSPSEKNLKILDLLADSVNLLIQWYRNHYSDPGNNYSIASSTSTSSLEDSKGFGSNNFTMNLFVK
LGEAFRKTSLARREEIRNHAVLSLQKSFQLAEELEFTPTNCINCFNLLIFAMVDDLHEKMLEYSRRENAEKEMRG
MEGTLKIALEILTDVYLQFLIPISQCSGFRTFWLGVLRRMDTCMKADLGSYGESKLPEVVPELLRKLITEMKEKE
ILVPKEGDDLWETTHIQIQWIAPSIKEELFPEESL*
**>Betpe_c0365.g0008**
MGRLKLQAGINAIEEEPEDCDATYSNKATVACMINSEIGAVLAVMRRNVRWGGRYMSGDDQLEHSLIQSLKALRK
QIFSWQHQWHSINPSMYLQPFLDVIRSDETGAPITGVALSAVHKILTLDVIDQNTVNVEDAMHLVVDAVTSCRFE
VTDPASEEVVLMKILQVLLACMKSKASVMLSNQHVCTIVNTCFRIVHQAGSKGELLQRIARHTMHELVRCIFSHL
PDVDNMECALVNGTNAIKQEIGGLNNEYVLGSRQVENGNMGSDYEGQPLSTNFASNASTVSVATSIDENTIGSSN
GKDAVSYDSHLMTEPYGVPCMVEIFHFLCSLLNVVEHMGMGAKSNSLAFDEDVPLFALGLINSAIELGGPSICRH
PRLLSLIQDELFRNLMQFGLSMSPLILSMVCSIVLNLYHHLRTELKLQLEAFFSCVILRLAQSRYGASYQQQEVA
MEALVDFCRQKTFMVEMYANLDCDITCSNVFEDLANLLSKSAFPVNCPLSSMHILALDGLIAVIQGMAERIGNAP
DSSEHNPVNLEEYTPFWMVKCDNYSDPSHWVPFVRRRKYIKRRLMIGADHFNRDPKKGLEFLQGTHLLPDKLDPQ
SVACFFRYTAGLDKNLVGDFLGNHDEFCVQVLHEFAGTFDFQDMNLDTALRLFLETFRLPGESQKIQRVLEAFSE
RYYEQSPQILANKDAALLLSYSLIMLNTDQHNVQVKKKMTEEDFIRNNRHINGGSDLPREFLSELYDSICKNEIR
TTPEQGAGFPEMTPSRWIDLMHKSKKTAPFIVSDSRAYLDHDMFAIMSGPTIAAISVVFDHAEHEEVYQTCIDGF
LAVAKISACHHLEDVLDDLVVSLCKFTTLLNPSSVEEPVLAFGDDTKARMATVTVFTIANRYGDFIRTGWRNILD
CILRLHKLGLLPARVASDAADESELSGDSGHGKPLPNSLSSAHMPSIGTPRRSSGLMGRFSQLLSLDTEEPRSQP
TEQQLAAHQRTLQTIQKCRIDGIFTESKFLHAESLLQLARALIWAAGRPQKGNSSPEDEDTAVFCLELLIAITLN
NRDRIVLLWQGVYEHISNIVQSTVMPCALVEKAVFGLLRICQRLLPYKENLADELLRSLQLVSKLDARVADAYCD
QITQEVSRLVKANASHIRSQLGWRTITSLLSFTARHPDASEAGFDALVFIMSDGAHLLPANYLLCVDASRQFAES
RVGQAERSVRALDLMAGSVNCLARWASEAKEAMGVEETVKMSQDIGEMWLRLVQGLRKVCLDQREEVRNHALLSL
QKCLTGVDEIHLPHGLWSQCFDGVIFTMLDDLHDIAQGHSQKDYRNMEGTLILAMKLLSEVFLQLLHDLSQLATF
CKLWLGVLSRMEKYMKVKVRTKKSEKLQELVPELLKKTLLFMKTTGVLVQRSALGGDSLWELTWLHVNNISLSLQ
SEVFPDQDSEPLHHKQGQTGGGPVSDETGSVPLNETVTSEGAGTRG*
**>Betpe_c0281.g0095**
MGHINPQFGINNDERKESPAKPSKGALACMVNSEIGAVLAVMRRNVRWGVRYVADDDHLEHSLIHSLKELRKHIF
SWQHQWHKIDPAVYLQPFLDVIRSDETGAPITGVALSSVYKILSLDILDLDTVNVGDAMHLIVDAVTSCRFEVTD
PASEEVVLMKILQVLLACMKSTTSVRLSNQHVCNIVNTCFRVVHQASSKGELLQRIARHTMHELVRCIFSHLLIY
LPDINNMEYALANGSGSSVHKEVGSTDMEDHTSENKQLENGNAGVEYDGQSSTYASKAPMSASASGRSDAGTYEN
ITEVSNGMEATKNDEKVMTEPFGVPCMVEIFHFLCSLLNVIEHVEFGPRSNPIAYDEDVPLFALGLINSAIELAG
PSISKHPKLLAIVQDELFHNLMQFGLSMSPLILSTVCSIVLNLYHYLRTELKLQLEAFLSCVLLRLAQSKHGSSY
QQQEVAMEALVDLCRQQTFMAEMYANFDCDITCSNVFEDLANLLSKSAFPVNGPLSAMHILALDGLISMVQGMAE
KIGNEPATSEASIDDEGYKAFWTMTCKNYSDPNCWVPFVRKVKYIKRKLMIGADHFNRDPKKGLEFLQGMHLLPE
KLDPLTVACFLRYSAGLDRNLVGDYLGNHDEFCVQVLHEFARTFDFQDMNLDTALRLFLGTFRLPGEAQKIHRVI
EAFSERYYEQSPRILMDQDVALVLSYSLILLNTDQHNSQVKKKMTEEDFIRNNRKINGGQDLPREFLSELYHSIC
ENEIQMIPDQSTGFPVMTSSRWINVMHKSKKVSPFIICDSRELLIYDMFAILSGPTIAAISVVFDQVEHEDVLQT
CIDGFLSVAKIAACYHLNDVLDDLIVSLCKFTTLLTPLSVEEAIVALGDYTKVRMATAAVFTIANSYGDYIRSGW
KNIMDCVLSLHKLGLVPTGLASDAVHDMEPSSDPERGKPATTSLSISNVPSVPTRKSSGLMGRFSQLLSFDMEEP
RTQPTEEQLEAQQRTRGTIQTCHIDSIFTESKFLQADSLVQLVKSLILAADRLRKGTVSVEDEDTAVFCLELLIA
IMLNNRDRIMLIWQGVFEHISDIVQSTIMPCTLVEKAVFGLLRICQRLLPYKENLTDELLRSLQLVLKLDARVAD
AYCEHITQEVMRLVKANATHIRSHVGWRTIISLLSITARHPEASEVGFEALAFIMSDGAHLLPSNYVLCVDAARQ
FGESRVGEVDRSVSALDMMAGSVVCLVRWSCETKNAVGEEAATKVNQDIGEMWLRLVQGLRKVCLDQREEVRNHA
ILMLQRSLAGVDGIHIPNSLWLQCFDVVIFTLLDDLEEIALGNSRKDYRSMEGTLVLAVKLMSKVFLQLLLDLWQ

LPSFCKLWLGVLNHMERYMNMKFRGRRSEKIQELIPELLKNTLFVMKSTGLLTPSDTIGGDSFWQLTWLHVKNIS
PSLQSEVFPAPELEQLQKKQITASGVPAPDGTVLVQSSETTA*
**>Betpe_c1159.g0023**
MEKSSDQNHDAEENNCPSCREKKKDSAAKSRRKELGLSCMLNTELGAVLAVIRRPPDPASQYFSPDEHYFDSAIL
HSLKSLRALIFNPQQEWRTIEPSVYLSPFLDVIQGDDIPAAATGVALSALLKILRLEIFDDKTPGAKEAINSIVS
GITSCRLEKTDPVSEDAVMMKILQVLTAIMKHRASLLLTDHAVCTIVNTCFQVVQQSASRGDLLQRNARYAMHEL
IQTIFSRLPDIEVNDGEDSESDSDDADVGGNLDSGYGVRCAVDIFYFFCSLLNVVEVVETEGSTSHTVDEDVQLF
ALVLINSAIELSGDAIGKHPKLLRMIQDDLFHHLIHYGTRSSPLVLSMICSTVLNMYNFLRRFIRLQLEAFFAYV
LLRVAALGSAMPLQEVALEGIINFCRQPTFIIEVYLNYDCDPLGRNLFEEIGKLLCKDLFPVASPATTLQVQAFE
GLAAMIHNIADNVDREDDLSPFGPCPLEIAEYSPFWEEKSQDELESWVEFVRIRKAQKKKILIAANHFNRDEKKG
LEYLKLSNLVSDPPDPKAHAFFFRYTPGLNKNAIGDYLGDPDQLHIEVLKEFTDTFKFSGMILDTALRTYLESFR
LPGESQKIQRILEAFSEKFYDQQSSDVFASKDAVFILCYSLIMLNTDQHNPQVKKKMTEDEFIRNNRAINAGKDL
PREYLSELFHSISNHAITLFGQSGVSLEMTPSRWVELINRSKLAQPFILCDFDRHLGRDMFACIAGPSVASLSAF
FEHIDEEEVLHKCVEVLFSIARIAQYGLEDTLDELLASFCKFTTLLNPYASAEETLFAFSNDMKPKMATLAVFTV
ANNFGESIRGGWRNIVDCLLKLKRLKLLPQSVIEFDTASTSSPDVPKESESAIIFPTRDKFGSLRASGTISRFSH
FLSLDSMEDSLSLNVSEFEQNLKIIKQCRIGNIFSNSSNLPDDALFNLGRSLIFAAAGKGQKFSTPVEEEETVGF
CWDLVIAIAMANIQKFQAFWPNFHEYMLAVAQFPMFSPIPFAEKAIIGLLRLSLKLLGTYRPEKFPEELIFKSIN
LMWKLDKEILDTSCEFITQSVSKILIEYPANLQSQLGWKSVLHLLSVTGRHSETYDQAVETLIMLMSDGTHVSRT
NYAYCIDCAFGFVALKNSPLEKNLKILDLMADSVNLLIQWYRNHYSEPGNNFSIASNTSNSSAEDNSRGLGSTNF
VMNLFLKLGEALRKSSLARREEMRNHAILALQKSFTLAEELDFTSTNCINYFNLIIFAMIDDLHEKMLEYSRRDN
SEREMRSMEGTLKIAMELLADVYLQFLKQISQSAGFRTFWLGVLRRMDTCMKADLGEHGKSKVEDLIPDLLKKMI
TQMKEKEILVQKEGDDLWEITYIQIQWIAPALKEELFPEVIF*
**>Jugre_LOC108991852**
MGRLKLQTGINSIEEEPEDCDTAYSNKATLSCMINSEIGAVLAVMRRNVRWGGRYMSGDDQLEHSLIQSLKALRK
QIFSWPHQWHTINPAVYLQPFLDVIRSDETGAPITSVALSSVHKILSLDVIDQNTVNVEDAMHLVVDSVTSCRFE
VTDPASEEVVLMKILQVLLSCMKSKASVMLSNQHVCTIVNTCFRIVHQAGSKGELLQRIARHTMHEIVRSIFSHL
PDVDNTERSLVNGSNTVKKEIGGLNNEYSLGSSQLENGNMGSDYDGQPLSTNFTTNVSTGSLATGTDEKTIGSVN
GKETVSYDLHLMTDGVPCMVEIFHFLCSLLNVVEHVGNDPRSIAYEEDVPLFALGLINSAIELGGPSIRHHPRLL
SLIQDELFRNLMQFGLSTSPLILSMVCSIVLNLYHHLRTELKLQLEAFFSCVILRLAQSRYGASYQQQEVAMEAL
VDFCRQKTFMVEMYANLDCDITCTNIFEDLANLLSKSAFPVNCPLSSMHILALDGLIAVIQGMAERIGNTSAGSE
HAPVSLEEYTPFWMVKCDNYSDPNHWVPFVRRRKYIKRRLMIGADHFNRDPKKGLEFLQGTHLLPDKLDPQSVAC
FFRYTAGLDKNLVGDFLGNHDEFCVQVLHEFAWTFDFQDMNLDTALRLFLETFRLPGESQKIQRVLEAFSERYYE
QSPQILANKDAALLLSYSLIMLNTDQHNVQVKKKMTEDDFIRNNRHINGGSDLPREFLSELYHSICKNEIRTTPE
QGAGFPEMTPSRWIDLMHKSKKTAPFIVSDSRAYLDHDMFAIMSGPTIAAISVVFDHAEHEEVYQTCIDGFLAVA
KISACHHLEDVLDDLVVSLCKFTTLLSPSSVEEPVLAFGDDMKARMATVTVFTIANRYGDYIRTGWRNILDCILR
LHKLGLLPARVASDAADELEVSADTGHGKPLTSSLSSAHMPSMGTPRRSSGLMGRFSQLLSLDTEEPRSQPTEQQ
LAAHQRTLQTIQKCHIDSIFTESKFLKAESLLQLARALIWAAGRPQKGNSSPEDEDTAVFCLELLIAITLNNRDR
IVLLWQGVYEHISNIVQSTVLPGALVEKAVFGLLRICQRLLPYKENLADELLRSLQLVLKLDPRVADAFSEQITQ
EVNRLVKANASHIRSQLGWRTITSLLSNTARQPEASEAGFDALLFIMSDGAHLSPANYVLCVDSSRQFAESRVGQ
AERSVRALDLMAGSVNCLARWMSEAKEAMGEEEALRTSQDIGEMWFRLVQGLRKVCLDQREEVRNHALLSLQKCL
TGVDELHLPHGLWSQCFDTVIFTMLDDLLEIAPGHSPKDYRNMEGTLILVMKLLSKVFLQLLNDLSQLTTFCKLW
LGVLSRMEKYMKVKVRGKRSEKLQELVPELLKNTLLVMKTRGVLVQRSALGGDSLWELTWLHVNNISLSLQSEVF
PDQDSEPPQHKQSQTGGVPISHETGSGPPNEAVASESAGTGG*
**>Jugre_LOC109006351**
MGHLNPQFGTNSFNGERKESPAKPSRGALACIINSEIGAVLAVMRRNVRWGVRYVADDDNLEHPLIHSLKELRKQ
IFSWQHLWNNIDPFLYLQPFLDVIRSDETGAPITGVALSSVYKILSLEILGLDTVNVGEAMHLIVDAVTSCRFEV
TDPASEEVVLMKILQVLLACMKSTASVWLSNQHVCNIINTCFRVVHQASSKGELLQRIARHTMHELVRCIFSHLP
DIEKTEHALANGSRSCVHEEVGSPGKGHTSESEQLENGNGGIEYGGQSTAYASTAPMSTSASGMSDGGTNENITE
CSNRMEPTKNDEKHMMDPFGIPCMLEIFHFLCSLLNVIEHIEFGPKSNPIAYDEDVPLFALGLINSAIELGGPYI
SKHPQLLAVVQDELFHNLMQFGLSRSPLILSTVCSIVLNLYHHLRTELKVQLEAFLSCVLLRLAQSKHGSSYQQQ
EVAMEALVDLCRQQTFMAEMYANFDCDITCSNVFEDLANLLSKSAFPVNGPLSAMHILALDGLICMVHGMAERLG
NEPPSLKASFDDEGYEAFWTMTCENYSDPNFWVPYVRKLKYLKRNLMVGADYFNRDPKKGLEFLQGMHLLPDKLD
PKSVACFLRYTTGLDKNVVGDFLGNHDEFCVQVLQEFAWTFDFQDMNLDTALRLFLGTFRLPGESQKIQRVLEAF
AERYYEQSPHILINKDAALVLSYSLIMLNTDQHNAQVKKKMTEEDFIRNNRRINGGKDLPREFLSELYHSISENE
IQMIPDQSAGFPVMASSCWINVMHKSKKTSPFIVCDSREPLTYDMFAILSGPTIAAMSVIFDQVEREDVLQTCID
GFLSVAKIAACYHLDDVLDDLVVSLCKFTTLLTPLSIEEAIVSFGDNSKARMATTAVFTIANRYGDYIHSGWKNI
LDCVLSLYNLGLLPANLASDAEYETDAPSDPDRAKPATTLSASHMPSLAPSRKSSGLMGRFSQLLSFDMEEPRSQ
PTEEQLAAQQHTRETIQKCHIDSIFTESKFLQAESLLQLVKALILAAGRLRKVTGSIEDEDTAVFSLELLITITL
NNRDRIMLIWQGVYEHISDIVQSTILPCTLVEKAVFGLLKICQRLLPYKENLTDELLRSLQLVLKLDARVADAYC
EHITQEVLRLVKSNATHIRSHLGWRTIISLLSITARHPEASEVGFEALAFIMSDGAHLLPSNYALCVDAARQFAE
SRVGEIDRSVSALDLMAGSVVCLVTWSSESKNSVDEEASMKVNQDIGEMWLRLVQGLRKVCLDQREEVRNHAILM
LQRSLTGVDGIHLPNTLWLQCFDLVIFVLLDDLLEIVLGQSPKDYRNMEGTLGLAVKLMSKVFLHLLLDLWRLPS

FCKLWLGVLNRMEKYMNMKLRGRRSEKIHELIPELLKNTLLVMKTTGLLTPSDAIGGDSFWQLTWLHVKNISPSL
QLEVFPAHELEQLQTKQVKAAGIPAPDGTVLVQSSETTA*
**>Jugre_LOC108985951**
MGHLKPDSGTISFNGERKDSPAKNFRGALACMVNSEIGAVLAVMRRNVRWGVRYVADDDKLEHSLIHSLKELRKK
IFSWQNKWHNVDPAVYLQPFLDVIQSDETGAPITGVALSSVYKILSLDILNLDTVNVGDAMHLIVDAVTSCRFEV
TDPASEEVVLMKILQVLLACMKSTASVRLSNQHVCNIVNTCFRVVHQASSKGELLQRIARHTMHELVRCIFSRLP
DMDSAEHALANGNRSCVNTEVGSPEKDHTLESEQLQNGFGGVEFGGQSSTYASKAPMSTSASGMSDGGMNASSGV
EPTKNDEKLLMEPYGVPCMVEIFHFLCSLLNAIEHIEFSPQSNPITYDEDVPLFALGLINSAIELGGPYVSKHPQ
LLAIVQDELFRNLMQYGLSMSPLILSTVCSIVLNLYHHLRTELKVQLEAFLSCVLLRLAQSKHGSSYQLQEVAME
ALVDLCRRQTFMAEMYANFDCDITCSNMFEDLANLLSKSAFPVNGPLSAMHILALDGLISMVQGMAERLGNEPLT
SEPSLDDEGYKEFWTKTCENYSDPNCWVPFVRKLKYIKRKLMVGADHFNRDPKKGLEFLQGMHLLPEKLDPQSVA
CLFRYTTGLDKNLVGDFLGNHDEFCVQVLHEFARTFDFHDMSLDTALRLFLGTFRLPGESQKIQRVLEAFAERYY
EQSPHILINKDAALLLSYSLILLNTDQHNAQVKKKMTEEDFIRNNRRINGGMDLPRGFLSELYRSICENEIQMIP
DQNAGVPVIASSRWINVIYKSRKSSPFIVCDSIELLTYDMFTILSGPTIAAMSVIFDQVEQEDVLQTCIDGFLSV
AKIAACYHCDDVLDDLVVSLCKFTTLVTPLSIEEAIVSLGDDSKARMATTTVFTIANRYGDYIHSGWKNILDCVL
SLYKLGLLPASLTSDAEYDLEPSSDPERGKPATASLSASRMPSVATPRKSSGLMGRFSQFLSFDMEEPSSQPTEH
QLAGQQRTRETIQNCHIDSIFMESKFLQAESLVQLVKALILAAGRLRKVTGSIEDEDTVVICLELLIAITLNNRD
RIMLIWKGVYEHISDIVQSTIMPCMLVEKAIFGLLKICQRLLPYKENLTDELLRSLQLVLKLDARVADAYCEHIT
QEVMRLVKANATHVRSHMGWRTIISLLSITARHPEASEVGFEALAFIMCDGAHLLPSNYVLCVDAARQFAESRVG
EVDLSVSALDMMAGSVVCLVTWSCETKNDVDKEAAVKVSQDIGEMWLRLVQGLRKVCLDQREEVRNHAILMLQRS
LAGVDGIQLPNTLWLQCFDLVIFTLLDDLLETVLGKSPKDYRNMEGTLVLAMKLMSKVFLHLLLDLWRLPSFCKL
WLGVLNRMERYMNIKFRGRRSEKVHELIPELLRNTLLVMKTTGLLTPSDTVGGDSFWQLTWLHVKNISPSLQSEL
FPAFELEQQQSKQIKAAGIPAPDGTVLVRSSETTA*
**>Jugre_LOC108992663**
MARDQYQDTDENDYTSSRETKKDSAARSRRKDLGISCMLNTELGAVMAVIRRPPLDPTSQFFVPEENVDPTILQS
LKSLRALLFNTQQEWRTVDPSIYLTPFLQVIQGDDIPAAATGIALSAILKVLTLEIFDEKTPGVKEGINSIVIGI
TTCRLEKIDSTSEDAVMLKILQVLTGVMKHRTSVLLTDHAVCTVVNTCFQVVQQSASRGDLLQRSARYTMHELIQ
IIFSRLPEIEVKQGEDSESESDGHDVDVGGNLDSGYGVRCAVDIFHFLCSLLNVVQVVETESSSFQAADEDVQLF
ALVLINSAIEWSGDEIGRHPKLLRMIRDDLFHHLIHYGIRSSPLVLSMICSTVLNIYHFLRRFIRLQLEAFFVYV
LLRVAALGSPLPLQEVALEGIINFCRQPTFIIEAFVNYDCDPLCRSIFEDTGKLLCKNTFPVASPATASQVQAFE
GLATMIHNIADNIDREDDLSPFDPYPAEITEYRPFWEEKSQDHELESWVEFVRVRKAQKKKILIAANHFNRDEKK
GLDYLKLSKLVSDPPDPKAHAFFFRYAPGLDKNMIGDYLGDPDQFHVLVLKEFTDTFNFSGMILDTALRTFLETF
RLPGESQKIQRILESFSEKFYDQQSTDTFATKDAVFILCYSLIMLNTDQHNPQVRKKMTEEEFIRNNRAINGGQD
LPREYLSELFCSISNNAITLFGQPGVQVDMNPSRWIELMNRSNYTQPLVLSNFDRRLGRDMFACIAGPSVASLSA
FFEHTDEEEFLHECVEVLFSIARIAQYGLEDTLDELLASFCKYTTLLNPYASAEETLFVFSNDMKPKMATLAVFT
IANNFGESIRGGWGNIVDCLLKLKRLKLLPQSVIEFDTASPSSSDIPKESESAIIFPARNPKFGSRQASSTISRF
SHFLSLDSTEDSLSLNMSEFEQNLKIIKQCHIGNIFSNSSNLPDEALFNLGRSLIFAAAGKGQKFSTPIEEEETV
GFCWDLVIAIALANIQRFQEFWPNFHDYMLAVTQFPMFSPIPFAEKAIICLSKVCLKLLSTYRHEKFPEELIFKS
INLMWKLDKEILDSCCELITQSASKILIEYPANLQTQLGWKSVLHLLSVTGRHPETYDQGVETLIMLLSDGTHVS
RINYAYCIDCAFGFIALKNSPLDKNLKILDLMADSVNLLIQWYRNQFSESGNSFSVASNTSNSSMEDNSKGIGSL
NFAMNLFLKLGEAFRKTSLARREEMRNHAILSLKRSFTLAEELDLTSTNCINCFNLVLFAMVDDLHEKMLEYSRR
ENSEREMRSMEGTLKISMELLTDVYLQFLKQISASPGFRTFWLGVLRRMDTCMKADLGEFGKSTVQELIPDLLRK
MITQMKEKEILVRKEGEDLWEITYIQIQWIAPSLKDELFPEEIF*
**>Quesu_LOC112039782**
MGRLKLQTGINSIEEEPEDCDVNYSNKATLACTINSEVGAVLAVMRRNVRWGGRYMSGDDQLENSLIQSLKALRK
QIFSWQHPWHAINPALYLQPFLDVIQSEETGAPITGVALSSVHKILSLDVIDQNTVNVDDAMHLVVDAVTSCRFE
VTDPASEEVVLMKILQVLLACMKIKASVMLSNQHVCTIVNTCYRIVHQAGSKGELLQRIARHTMHELLRCIFSRL
PDVENTEHMNGGNTIKQEIGGLNNEYTLGSGNFENGNMSSDYDGQPLSTNFASNASMGLVATAIDENIGGSGGGK
EAVSYDLHMMTGPYGVPCMVEIFNFLCSSLNVNEHMGMGPRSNTIAFDEDGPLFSLGLINSAIELGGPSIRRHPR
LLSLIQDELFRNLMQFGLSMSPLILSMVCSMVLNLYNHLRTELKLQLEAFFSCVILRLAQSRYGATYQQQEVAME
ALVDFCRQKTFMVEMYANLDCDITCSNIFEDLANLLSKSAFPVNCPLSSMHILALDGLIAVIQGMAERIGNGLVG
SENSPVNLEEYTPFWMVKCDNYSDTRHWVPFVRRRKYIKRRLMIGADHFNRDPKKGLEFLQGTHLLPDKLDPQSV
ACFFRYTAGLDKNLVGDFLGNHDEFCVQVLHEFAWTFDFQDMNLDTALRLFLETFRLPGESQKIQRVLEAFSERY
YEQSPQILANKDAALLLSYSLIMLNTDQHNIQVKKKMTEEDFIRNNRHINGGNDLPRDFLSELYHSICKNEIRTT
PEQGAGFPEMTPSRWIDLMHKSKKTAPFIVSDSRAYLDRDMFAIMSGPTIAAISVVFDHAEHEEVYQTCIDGFLA
VAKISACHHLEDVLDDLVVSLCKFTTLMNPSSAEEPVLAFGDDTKARKATVAVFTIANNYGDYIRTGWRNILDCI
LRLHKLGLLPARVASDAADESELSTDTGHGKPLPNSLSSAHMPSMGTPRRSSGLMGRFSQLLSLDTEEPRSQPTE
QQLAAHQRTLQTIQKCHIDSIFTESKFLQAESLLQLAKALMLAAGRPQKGNSSPEDEDTAVFCLELLIAITLNNR
DRIVLLWQGVYEHIANIVQSTVIPCALVEKAVFGVLRICQRLLPYKENLSDELLKSLQLVSKLDARVADAYCEQI
TQEVSRLVKANASHIRSQFGWRTITSILSNTARHPEASEAGFDALLFIMSDGAHLLPANYVLCVETSRNFAESRV
GQAERSVRALDLMAGSVNCLARWASEAKEAMGVEEALKMSQDIGEMWLRLVQGLRKVCLDQRDEVRNYALLSLQK
CLTGVDEIHLPHGLWLQCFEMVIFTVLDDLLEIAQGQSQKDYRNMEGTLILAMKLLSKVFLQLLHDLSQLSIFCK

LWLGVLSRMEKYMNVKVRGKRSEKLHELVPELLKNTLLVMKTRGVLVQRSALGGDSLWELTWLHVNKISVSLQSE
VFPDQDLEPSQHKHSQIGGGLVSDETGSSPLNETAASESAGIRG*
**>Quesu_LOC111985027**
MEDQNQDAYENNTRERKKIKTKRKELGLSCMLNTEVSALLAVIRRPPEPTSPYSDDHFDSSALHSLKSLRALIFN
PQQEWHNVDPAIYISPFLHVIQGDDVPAAATAVALYALLKILNLEIFDEKTKGAKEAINSIVTAIAGCRLEKTDS
STEDAVMMRILQVLTAIMKHCASVLLTDHVVCNIVNTCFQVVQHSAGRGDLLQRNARYALHELIQIIFSRLPHLA
IKDSESESESESGSESENGTDDSSGGDADANLNSGYGIRSAADIFRFLCSLLNVVRVVEMEGSTFQDPDEDVQHF
ALVLINSAIELSGEEIGRHSKLLRMIQDDLFHHLIHYGTCSSPLVLSMICSTVLNIYHFLRRFIRLQLGAFFAIV
LLRVVPLASSISLQEVALEGLINFCRQPTFIIEVYVNYDCDPLSRSLFEEIGKLLCKNSFPMTSPVTTLQVLAFE
GLATMIHNIADNIDKEDDSSPIGPYPVVITEYSPFWEEKSKDELESWVDFVRIRKAQKKKILIAASHFNRDEKKG
LEYLKFSKLVSDPPDPKAYAFFFRYTPGLDKNMIGDYLGDPSHFNIEVLREFTDTFKFSGMVLDIALRSYLETFR
LPGEAQKIQRILEAFSERFYDEQSSDVFVSKDAVFILCYSVIMLNTDQHNPQVKKKMTEEEFIRNNRAINGGQDL
PREYLSELFHSISNKAITLFGQSGLSVEMNPSKWIELINRSKLVQPLILCDFDRRLGRDMFACIAGPAVASLCAF
FEHTDEEELLQECTEMLISIARIAQYGLEDTLDELLASFCKFTTLLNPYASTEETLFAFSNDLKPKMATLAVFTI
ANNFGESIRGGWRNIVDCLLKLKRLKLLPQSVIEFDSASASSSDVPKEPESGIIFPSHDPKFGSRQASGTISRFS
HFLSLESMEDSLSLNLSEFEQNLKIIKQCRIGNIFSNSSNLPDDALLNLGRSLIFAAAGKGQKFSTPVEEEETVG
FCWDLVIAIALANIHRFQTFWTNFHDYMLSVAQFPLFSPIPFAEKAILGLFKVCLKLLGTYRVDKIPEELIFKSI
NLMWKLDKEILDTCCEFITQSVSKVLIEYPANLQTQLGWKTVLHLLSVTGRHSETYDQGVEALIMLMSDGTHVSR
TNYVYCIDCAFSFVALKNSPLENNLKILDLMADSVNLLTQWYKNHQAENNLSVSSNTSNSSLEENSKTLGSSNFT
LNLFLKLGEALRKSSLARREEMRNHAIVALQKSFSLAQDLDFSPTSCMNCFNLVIFAMVDDLHEKMLEYSRRENS
EREMRSMERTLEIAMELLTNVFLQFLKQISEGPGFRTFWLGVLRRMDTCMKADLGQYGTSKIEELVPDLLKKMIT
QMKEKGILVQKEGADLWEITYIQIQWIAPSLKEELFPEEMF*
**>Quelo_LOC115954940**
MGRLKLQTGINSIEEEPEDCDVNYSNKATLACTINSEVGAVLAVMRRNVRWGGRYMSGDDQLENSLIQSLKALRK
QIFSWQHPWHAINPALYLQPFLDVIQSEETGAPITGVALSSVHKILSLDVIDQNTVNVDDAMHLVVDAVTSCRFE
VTDPASEEVVLMKILQVLLACMKIKASVMLSNQHVCTIVNTCYRIVHQAGSKGELLQRIARHTMHELLRCIFSRL
PDVENMEHINGGNTIKQEIGGLNNEYTLGGGNFENGNMSSDYDGQPLSTNFASNASIGLVATAIDENIGGSGGGK
EAVSYDLHMMNGPYGVPCMVEIFNFLCSSLNVNEHTGMGPRSNTIAFDEDGPLFSLGLINSAIELGGPSIRRHPR
LLSLIQDELFRNLMQFGLSMSPLILSMVCSMVLNLYNHLRMELKLQLEAFFSCVILRLAQSRYGATYQQQEVAME
ALVDFCRQKTFMVEMYANLDCDITCSNIFEDLANLLSKSAFPVNCPLSSMHILALDGLIAVIQGMAERIGNGLVG
SENSPVNLEEYTPFWMVKCDNYSDTRHWVPFVRRRKYIKRRLMIGADHFNRDPKKGLEFLQGTHLLPDKLDPQSV
ACFFRYTAGLDKNLVGDFLGNHDEFCVQVLHEFAWTFDFQDMNLDTALRLFLETFRLPGESQKIQRVLEAFSERY
YEQSPQILANKDAALLLSYSLIMLNTDQHNIQVKKKMTEEDFIRNNRHINGGNDLPRDFLSELYHSICKNEIRTT
PEQGAGFPEMTPSRWIDLMHKSKKTAPFIVSDSRAYLDRDMFAIMSGPTIAAISVVFDHAEHEEVYQTCIDGFLA
VAKISACHHLEDVLDDLVVSLCKFTTLMNPSSAEEPVLAFGDDTKARKATVAVFTIANNYGDYIRTGWRNILDCI
LRLHKLGLLPARVASDAADESELSTDTGHGKPLPNSLSSAHMPSMGTPRRSSGLMGRFSQLLSLDTEEPRSQPTE
QQLAAHQRTLQTIQKCHIDSIFTESKFLQAESLLQLAKALMLAAGRPQKGNSSPEDEDTAVFCLELLIAITLNNR
DRIVLLWQGVYEHIANIVQSTVIPCALVEKAVFGVLRICQRLLPYKENLSDELLKSLQLVMKLDARVADAYCEQI
TQEVSRLVKANASHIRSQFGWRTITSILNNTARHLEASEAGFDALLFIMSDGAHLLPANYALCVDTSRQFAESRV
GQAERSVRALDLMAGSVNCLARWASEAKEAIGVEEALKMSQDIGEMWMRLVQGLRKVCLDQREEVRNYALLSLQK
CLTGVDEIHLPHGLWLQCFEMVIFTVLDDLLEIAQGQSQKDYRNMEGTLILAMKLLSKVFLQLLHDLSQLTIFCK
LWLGVLSRMEKYMNVKVRGKRSEKLHELVPELLKNTLLVMKTRGVLVQRSALGGDSLWELTWLHVNNISVSLQSE
VFPDQDSEPSQHKHSQIGGGLVSDETGSSPLNETVASESAGIRG*
**>Quelo_LOC115976920**
MGHLNPQFGTNSSNGLREESPIKPSRGALACMINFEIGAVLAVMRRNVRWGVRYVADDDQLEHSLIHSLKELRKQ
IFSWQHQWQYIDPAVYLRPFLDVIRSDETGAPITGVALSSVYKILSLDILDLDTVNVGDAMHLIVDAVTSCRFEV
TDPGSEEVVLMKILQVLLACMKSKASVRLNNQHVCNIVNTCFRVVHQASSKGELLQRIARHTMHELVRCIFSHLP
DIDTTEHALTNGSRSYVREEVGLADEDHTSESKQLENGNVGIESDGQSSTYASKDHMGTSDGGMNDSGMSEKITE
VSNGVEASKNDESLLMEPFGVPCMVEIFHFLCSLLNVSEHIEFGPRSNPIAYDEDVPLFALGLINSAIELAGPSI
SKHPQLLAIVQDGLFLNLMQFGLSMSPLILSTVCSIVLNLYHHLRTELKVQLEAFLSCVLLRLSQSKHGLSYQQQ
EVAMEALVDFCRQQTFMAEMFANFDCDVTCSNVFEELVNLLSKSAFPVNGPLSAMHIVALDGLISMVQGMAESVG
NEPFTSEPAVDDEGYTPFWRMECESYSDPNCWVPFVRKMKYIKKRLMVGADHFNRDPKKGLEFLQGMNLLPEKLD
PQSVAHVFRYTTGLDKNLIGDYLGNHDDFCIQVLHEFARTFDFQDMNLDIALRLFVGTFRLPGESQKIQRVLEAF
AERYYEQSPHILINKDAALLLSYSLILLNTDQHNSQVKKKMTEEDFIRNNRKINGGKDLPREVLSELYHSICENE
IQMVPDQSTGSPMMTSSCWINVIHKSQKSSPFIICDSRDLLNHDMFAILSGPTIAAMSVVFDQAEQEDVLQTCVD
GFLAVAKIAASYHLDDVLDDLVVSLCKFTTLLTPLSVEEAFFALGDDIKARMATTTVFTIANRYGDYIRSSWKNI
LECVLSLHKLGLLPARLASDAVDDIEPPSDLDRGKPATTSLATPNMPSITTRKSSGLMGRFSQLLSFDVEEPRSQ
PTEEELASQQRTRETIQTFHIDSIFTESKFLQAESLLQLVKALILAAGQLRKGTSYYEDESTAVFCLELLIAITL
NNRDRIMLIWQGVYEHISDIVQSTIMPCTLVEKAVFGLLKICQRLLPYKENLTDELLRSLHLVLKLDARVADAYC
EHITQEVLRLVKANSAHIRSNAGWRTIISLLSITARHPEASDVGFEALAFIMSDGTHLLPSNYVLCVDAARQFAE
SRVGEVDRSVSALDLMAGSVVCLVSWSCEMNSVGEEAAKKVNQDIGEMWLRLVQGLRKVCLDQREEVRNHSILML
QRSLAGVDGIHLSNALWLQCFDQVIFTLLDDLLEITVGNSPKDYRNMEGTLVLAMKLMSKVFLQLLLDLWQLPSF

CKLWLGVLNRMERYMSAKFRGRRSEKIHELIPELLKNTLFVMKTTGLLVPSDTIDGDSFWQLTWLHVKNITPSLQ
SEVFPPQELEQLQAKQLKEAGIPAPDGAVLVQSSETTV*
**>Quelo_LOC115981000**
MEDQNQDAYENNNRERKKIKTKRKELGLSCMLNTEVSALLAVIRRPPEPTSPYYFSSDDHFDSAALHSLKSLRAL
IFNPQQEWHNIDPAIYISPFLHVIQGDDVPAAATGVALYALLKILNLEIFDEKTKGAKEAINSIVTAIAGCRLEK
TDSSTEDAVMMRILQVLTAIMKHCASVLLTDHVVCNIVNTCFQVVQHSAGRGDLLQRNARYALHELIQIIFSRLP
DLAIKDSESEPESESGSGSESETGMDDSSGGDADADLNSGYGIRSAVDIFHFLCSLLNVVRVVEMEGSTFQDPDE
DVQHFALVLINSAIELSGEEIGRHSKLLRMIQDDLFHHLIHYGTCSSPLVLSMICSTVLNIYHFLRRFIRLQLEA
FFAVVLLRVVPLASSMSLQEVALEGLINFCRQPTFIIEVYVNYDCDPLSRSLFEEIGKLLCKNSFPMTSPVTTLQ
VLAFEGLATMIHNIADNIDKEDDSSPIGPYPVVITEYSPFWEEKSKDELESWVDFVRIRKAQKKKILIAASHFNR
DEKKGLEYLKFSKLVSDPPDPKAYAFFFRYTPGLDKNMIGDYLGDPSHFNIEVLREFTDTFKFSGMVLDIALRSY
LETFRLPGEAQKIQRILEAFSERFYDEQSSDVFVSKDAVFILCYSVIMLNTDQHNPQVKKKMTEEEFIRNNRAIN
GGQDLPREYLSELFHSISNKAITLFGQSGVSVEMNPSKWIELINRSKLVQPLILCDFDRRLGRDMFACIAGPTVA
SLCAFFEHTDEEELLQECTEMLISIARIAQYGLEDTLDQLLASFCKFTTLLNPYASTEETLFAFSNDLKPKMATL
AVFTIANNFGESIRGGWRNIVDCLLKLKRLKLLPQSVIEFDSASTSSSDVPKEPESGIIFPSHDPKFGSRQASGT
ISRFSHFLSLESMEDSLSLNLSEFEQNLKIIKQCRIGNIFSNSSNLPDDALLNLGRSLIFAAAGKGQKFSTPVEE
EETVGFCWDLVIAIALANIHRFQSFWTNFHDYMLSVAQFPLFSPIPFAEKAILGLFKVCLKLLSTYPVDKIPEEL
IFKSINLMWKLDKEILDTCCEFITQSVSKVLIEYPANLQTQLGWKSVLHLLSVTGRHSETYDQGVEALIVLMSDG
THVSRTNYVYCIDCAFSFVALKNSPLEKNLKILDLMADSVNLLTQWYKNHHAENNSSVSSNTSNSSLEENSKSLG
SSNFTLNLFLKLGEALRKSSLARREEMRNHAIVALQKSFSLAQDLDFSPTNCMNCFNLVIFAMVDDLHEKMLEYS
RRENSEREMRSMERTLEIAMELLTNVFLQFLKQISEGPGFRTFWLGVLRRMDTCMKADLGQYGTSKIEELVPDLL
KKMITQMKEKGILVQKEGKFYELYELDAEIGHEELDWKMTSESNLHARIAILSGQVGISESGIDDAVQKLVAHG*
**>FagsyGNOM_chr9**
MGRLKMQSGINSIEEEPEDCDVTYSNKATLACMINSEVGAVLAVMRRNVRWGGRYMSGDDQLENSLIQSLKALRK
QIFSWQHQWHTINPAMYLQPFLDVIQSEETGAPITGVALSSVHKILTLDVIDQNTVNVDDAMHLVVDAVTSCRFE
VTDPASEEVVLMKILQVLLACMKIKASVMLSNQHVCTIVNTCFRIVHQAGSKGELLQRIARHTMHELLRCIFSHL
PDVDNTEHALVNGSNTIKQEIGGLNNDYALGSRQLENGNVSSDYDGQPLSTNFSSTGLIATAIDENTSGPGSGKE
QVSYDLHMMTGPYGVPCMVEIFHFLCSLLNVIEHMGMGPRSNTIAFDEDVPLFALGLINTAIELGGPSIRHHPRL
LSLIQDELFRNLMQFGLSMSPLILSMVCSIVLNLYHHLRTELKLQLEAFFSCVILRLAQSRYGASYQQQEVAMEA
LVDFCRQKTFMVEMYANLDCDITCSNIFEDLANLLSKSAFPVNCPLSSMHILALDGLIAVIQGMAERIGNGLVGS
ENSPVNLEEYTPFWMVKCDNYSDPRHWVPFVRRRKYIKRRLMIGADHFNRDPKKGLEFLQGTHLLPEKLDPQSVA
CFFRYTAGLDKNLVGDFLGNHDDFCVQVLHEFAWTFDFQDMNLDTALRLFLETFRLPGESQKIQRVLEAFSERYY
EQSPQILANKDAALLLSYSLIMLNTDQHNVQVKKKMTEEDFIRNNRHINGGNDLPRDFLSELYHSICKNEIRTTP
EQGASFPEMTPSRWIDLMHKSKKTAPFIVSDSRAYLDRDMFAIMSGPTIAAISVVFDHAEHDEVYQTCIDGFLAV
AKISACHHledvlddlvvSLCKFTTLLNPSSVEEPVLAFGDDTKARKATVTVFTIANNYGDYIRTGWRNILDCIL
RLHKLGLLPARVASDTADESELSADTGHGKPLTNSLSSAHMPSIGTPRRSSGLMGRFSQLLSLDTEEPRSQPTEQ
QLAAHQRTLQTIQKCHIDSIFTESKFLQAESLLKLARALMWAAGRPQKGNSSPEDEDTAVFCLELLIAITLNNRD
RIGLLWQGVYDHISNIVQSTVMPCALVEKAVFGLLRICQRLLPYKENLADELLRSLSLVTKLDARVADAYCEQIT
QEVSRLVKANASHIRSQLGWRTITSLLSFTARHLEASEAGFDALLFIMSDGAHLLPANYVLCVDVSRQFAESRVG
QAERSVRALDLMAGSINCLARWASEAKEARGEEEAVKMSQDIGEMWLRLAQGLRKVCLDQREEVRNHALLLLQKC
LTGVDEIHLPHGLWLQQCFEMVIFTMLDDLLEIAQGQSQKDYRNMEGTLILAMKLLSKVFLQLLHDLAQLSTFCK
LWLGVLSRMEKYMRVKVRGKKSEKLQELVPELLKNTLLVMKTRGILVQRSALGGDSLWELTWLHVNNISLSLQSE
VFPDQDLEPSQHKHGQTGGGHVSDETGFIPLNETMASESAGTGG*
**>FagsyGNL1_chr8**
MGHLNPQFGTNSSNGQREEAPTKPSGGALACMINSEIGAVLAVMRRNVRWGVRYVADDDQLEHSLIHSLKELRKQ
IFSWQHKWHNIDPAVYLQPFLDVIRSDETGAPITGVALSSVYKILSLDILDLDTVNVGDAMHLIVDAVTSCRFEV
TDPGSEEVVLMKILQVLLACMKSKASVRLSNQHVCNIVNTCFRVVHQASSKGELLQRIARHTMHELVRCIFSHLP
DIDSTEHALTNGSRSYVREEVGSADKDHTCESKQLENGNAGVESDGQSSSYASKDHMGTSASGTNDSGTSANLTE
VSNGVEATKNDEKLMMEPFGVPCMVEIFHFLCSLLNVIEHIEFGPRSNPIAYDEDVPLFSLGLINSAIELAGPSI
SKHPQLLAIVQDGLFHNLMQFGLSMSPLILSTVCSIVLNLYHHLRTELKAQLEAFFSSVLLRLSQSKHGSSYQQQ
EVALEALVDLCRQQTFMAEMFANFDCDITCSNVFEELVNLLSKSAFPVNGPLSAMHIVALDGLISMVQGMAESIG
SEPPTSEATVDDEGYTAFWRMACESYSDHNCWVPFVRKMKYIKRRLMVGADHFNRDPKKGLEFLQGMNLLPEKLE
PQSVAYLFRYTAGLDKNLVGDFLGNHDEFCIQVLHEFARTFDFQDMNLDSALRLFLGTFRLPGESQKIQRVLEAF
AERYYEQSPHILSNIDAALLLSYSLILLNTDQHNAQVKKKMTEEDFIRNNRRINGGKDLPREILSELYHSICENE
IQMVPDQGSGFPMMTSSRWITVMHKSQKTSPFIICDSRDLLNYDMFSILSGPTIAAMSVVFDQAEQEDVLQTCID
GFLAVAKIAACYHlddvlddlvvSLCKFTTLLTPLSVDEAFVALGDDIKARMATTTVFTIANRYGDYIRSSWKNI
LDCVLSLHMLGLLSSRLASDAVDDMEPSSDSDRGKPASTSLATPHMPSITPRKSSGLMGRFSQLLSFDVEEPRSQ
PTEEQLAAQQRTHETIQTCHIDSIFTESKFLQAESllqlvqalilaaGRLRKGTSSIEDEDTAVFCLELLIAITL
NNRDRIMLIWQGVYELISDIVQSTIMPSTLVEKAVFGLLKICQRLLPYKENLTDELLRSLQLVLKLDARVADAYC
EHITQEVLRLVKSNATHIRSNAGWRTIISLLSITARHPEASEVGFEALAFIMSDGTHLLPSNYVLCVDAARQFAE
SRVGEVDRSVSALDLIAGSVVCLVRWSCETKNAVGEEAAKKVNQDIGEMWLRLVQGLRKVCLDQREEVRNHAILM
LQRSLAGVDGIHLPNALWLQCFDLVIFTLLDDLLEIAVGNSPKDYRNMEGTLVLAMKLMSKVFLQLLLDLWRLPS

FCKLWLGVLNRMERYMNTKFRGRRSEKIHELIPELLKNTLLVMKTTGLLVPSDTIGGDSFWQLTWLHVKNITPSL
QSEVFPPNELEQLQTKAVGIPAPDGTVLVQSSETTV*
**>FagsyGNL2_chr12**
METASDENHNTDENDGASIRDKKKAKRKELGLSCMLNTEVSAVLAVIRRPLDSTSQYYFSPDDHFDSATLQSLKS
LRALLFNPQQEWRNIDPSIYLSPFLHVIQGDDVPAAATCVALSAILKILKLEIFDEKTQGAKDAINSIVTAITSC
RLEKTDSVTEDAVMMKILQVLTGIMKHRTSVLLTDHTVCTIVNTCFQVVQQSSSRGDLLQRNARYALHELIQIIF
SRLPDIEVKEGHDSESDTDDADVGANLDSGYGIRCAVDIFHFLCSLLNVVEVVEMEGSTSHSADEEVQHFALVLI
NSAIELSGDEIGRHPKLLRMIQDDLFHHLIHYGTCSSPLVLSMICSTVLNIYHFLRRFIRLQLEAFFVFVLLKVA
ARGSSVSLQEVALEGLINFCRQPTFIIEVYVNYDCDPLSRSLFEEIGKLLCKNSFPMASPVTTLQVQAFEGLTTM
IHNIADNIDKEGDSGPFGPYPVEITEYSPFWEEKDKDELESWVEFVRIRKAQKKKILIAANHFNRDEKKGLEYLK
FSKLVSEPQDPKAYAFYFRYTPGLDKNMIGDYLGDPDQFHIEVLREFTDSFKFAGMILDFALRTYLETFRLPGES
QKIQRILEAFSEKFYDQQSSDVFVSKDAVFILCYSLIMLNTDQHNPQVKKKMTEEEFIRNNRAINGGQDLPREYL
SELFHSISNKAITIFGQSGVPVEMNPSRWIELINRSKLVQPFILCDFDRRLGRDMFAYIAGPSVASLSAFFEHTD
EEELLHECTEMLFSIARIAQYGLEDTLDELLASFCKFTTLLNPYASTEETLFAFSNDLKPKMATLAVFTIANNFG
ESIRGGWRNIVDCllklkrlkllPQSVIEFDSASTSSSDIPEESESGIIFPAHDPKFGSRKTSGTITRFSHFlsl
esiddslslnlseFEQNLKIIKQCRIGNIFSNSSNLPDDALLNLGRSLIFAAAGKGQKFSTPVEEEETVGFCWDL
VIAIALANIQRFQTFWPNFHDYMLSVAQFPMFSPIPFAEKAIQGLFKVCLKLIGTYRLEKLPEELIFKSINLMWK
LDKEILDTCCELITQSVSKILIEYPANLQTQLGWKSVLHLLSVTGRHPETYDQGVETLIMLMSDGTHVLRTNYAY
CIDCAFSFVALKNSPLEKNLKILDLMAESVNSLIQWYKNHSTEsgnnfsvssnsssednsRGQGSSNFGLNLFLK
LGEALRKSSLARREEMRNYAIMALQKSFILAVELDFSPTNCMNCFNLVIFAMIDDLHEKMLeysrrenseremrs
merTLEIGMELLTDVYLQFLKQISQGPGFRTFWLGVLRRMDTCMKADLGEYGTSKIEELIPYLLRKMITQMKEKE
ILVQKEGDDLWEITYIQIQWIAPSLKDELFPEEFF*
**>Morru_MR3G014762**
MGRLKLQTGINAIEEEPEECDATHSYKATLACMINSEIGAVLAVMRRNVRWGGRYMSGDDQLEHSLIQSLKALRK
QIFSWQHQWHTINPAVYLQPFLEVIRSDETGAPITGVALSSVHKILSLDVIDQNTVNVEDAMHLIVDAVTSCRFE
VTDPASEEVVLMKILQVLLACMKSKASVMLSNQHVCTIVNTCYRIVHQAGSKGELLQRIARHAMHELVRCIFSHL
PDVDNMEHALVNGRNTIKQEIGGLNNDYALGNRQSENGNMGSDCDVQPVSTNIVSTVSVATGVDDNTIGSGNGKE
AVSFDLQLLTEPYGVSCMVETFHFLCTLLDSVEHMGMGSRSDTMAFDEDVPLFALGLINSAIELSGPSILKHRRL
LSLIQDELFRYLMQFGLSMSPLILSMVCSTVLNLYHHLRTELKLQLEAFFSCVILRLAQSRYGASYQQQEVAMEA
LVDFCRQKTFMVEMYANLDCDITCSNVFEDLANLLSKSAFPVNCPLSSMHILALDGLIAVIQGMAERIGNAPVGS
EHAPVSLEEYTPFWMVKCDNYGDPNHWVPFVRRRKYIKRRLMIGADHFNRDPKKGLEFLQGTHLLPDKLDPQSVA
CFFRYTSGLDKNLVGDFLGNHDEFCVQVLHEFAGTFDFQDMHLDTALRLFLETFRLPGESQKIQRVLEAFSERYY
EQSPQILANKDAALLLSYSLIMLNTDQHNVQVKKKMTEDDFIRNNRHINGGGDLPREFLSELYQSICKNEIRTIP
EQGAGFPEMTPSRWIDLMHKSKKTTPFIVSDSRAYLDHDMFAIMSGPTIAAISVVFDHTEHEEVYQTCIDGFLAV
AKISACHHLEDVLDDLVVSLCKFTTLLNPSSVEEPVLAFGDDMKARMATVTVFTISNRYGDYIRTGWRNILDCIL
KLHKLGLLPARVASDAADESEVSADTGHGKPLTSSLSSAHMQSMGTPRRSSGLMGRFSQLLSLDTEEPRSQPTEQ
QLAAHQRTLQTIQKCHIDSIFTESKFLQAESLLQLARALIWAAGRPQKGNSSPEDEDTAVFCLELLIAITLNNRD
RIVLLWQGVYEHISNIVQSTVMPGALVEKAVFGLLRICQRLLPYKENLADELLRSLQLVLKLDARVADAYCEQIT
QEVSRLVKANASHIRSQLGWRTITSLLSITARHPEASEAGFDALLFIMSDGAHLLPANYVLCVDSARQFAESRVG
QAERSVRALDLMADSVHCLARWANEAKEAMGEEEAAKTSLDIGEMWLRLMQGLRKVCLEQREEVRNHAILSLQKC
LTGVDEINLPHGLWLQCFDMVIFTMLDDLLEIAQGHSQKDHRNMEGTLILALKLLSKVFLQLLNDLSQLTTFCKL
WLGVLGRMEKYMKVKVRGKRSEKLQELVPDILKNTLLVMKTRGVLLQRSALGGDSLWELTWLHVNNISVSLQSEV
FPDQDSEPPPHKQGQTGVGPVSDEMGSVAPYESAASESVGTGS*
**>Morru_MR2G025548**
MGNLNPQFGPNSFNGERKESPARPSGGALACMVNSEIGAVLAVMRRNVRWGVRYVEDDDKLEHSLIHSLKELRKQ
IFSWQHQWNNIDPAVYLQPFLDVIQSDETGAPITGVALSSVYKILSLDIVDLNTVNAGDAMHLIVDAVTSCRFEV
TDPASEEVVLMKILQVLLACVKSTASVRLSNQHVCNIINTCFRVVHQASSKGELLQRIARQTMHELVRCIFSHLP
DIDNAGHPLANGSRSQVHEEVATNKERTSESKQLENGNAGVEHGGQSSMNASKASTSACGMSESGTNESITEATT
AMEPTNNDEKLMMEPFGVPCMVEIFHFLCSLLNVIENVEFGPRSNPIAYDEDVPLFALGLINSAIELGGPYISRH
PQLLAIVQDELFRNLMLFGSSMSPLILSTVCSIVLTLYHHLRTELKVQLEAFLSCVLLRIAQSKHGSSYQQQEVA
MEALVDLCRQQTFVAEMYANFDCDITCNNLFEDLANLLSKSAFPVNGPLSAMHILALDGLISMVQGMAEWQGHEP
PSSDASNDEGYKAFWTMTCENYSDPTYWVPFVHKSKYIKRKLMIGADHFNRDPKKGLEFLQGMHLLPEKLDAQSV
ACFFRYTAGLDKNLVGDFLGSHDEFCVEVLHEFAITFDFRDMNLDTALRLFLGNFRLPGESQKIQRVLEAFAERY
YEQSTDILNNKDAALLLSYSLILLNTDQHNAQVKKKMTEEDFIRNNRRINGGKDLPREYLSELYHSICENEIQMV
PDQSAAFPVMASSRWINVMHKSKKTSPFILCDSRDSLNYDMFAILSGPTIAAMSVVFDQVEQEDILQTCIDGFLS
VAKIAACYHLDDVLDDLVVSLCKFTTLLSPLSVEEFIISLGDETKARMATATVFTIANRYGDYIRTGWKNILDCV
LSLHKVGLLPASLASDAEYDMEPSSDPEQGKPATTSFSTSHMPSVAIPRKSSGLMGRFSQLLSFDVEEPRSQPTE
EQLAAQQRSRETIQTCHIDSIFTECKFLQAESLMQLVKALVLAADQLRKGTGSTEDEDTAVFCLELLITITLNNR
DRIMLIWQGVYEHISDIVQSTSMPCTLVEKAVFGLIKICQRLLPYKENLTDELLRSLQLVLKLDARVADAYCEHI
TQEVMHLVKANAAHIRSHVGWRTIISMLSITARHPEASEVGFEALTFIMADAAHLLPSNYILCVDAARQFAESRV
GEVNRSVSALDLMAGSVVCLVRWSCETKNAGDEEAAMKANQDIGEMWLRLVQGLRKVCLDQREEVRNHAILMLQR
SLAGVDGIHLPNILWLQCFDLVIFTLLDDLLEISLGKSPKDYRNMEGTLVLAMKLMSKVFLQLLLDLWRLPSFCK

LWLGILNRMERYMNIKFRGRRSEKIHELIPELLKNTLLVMKTTGLLTPSDTIGGDSFWQLTWLHVKNISPALQLE
VFPARELEQPHARHIKTGGTPAPDGTVLVQASDTTA*
**>Morru_MR0G007998**
MTDSAAKSRRKELGISCMLNTEVGAVLAVIRRPPLEPAPQHFSLEDHFDSGILHSLKSLRALLFNPHQEWRTLEP
SIYLTPFLNIIQSDDIPGTATGVALSAILKILRLEIFDQKTPGAKDALNSVVTGITSCRLEKSDPVSEDAVMLKI
LQVLTGVMKHRASVLLSDHEVCTIVNTCFQVVQQSASRGDLLQRSARYTMHELIQIIFSRLPDVEINDGDDLESE
SDTEYGDVGGDLDSGYGVRCAVDIFHFLCSLLNVVEVVETEGSTFHTADEDVQLFALVLINSAIELSGDGIGRHP
KLLRMIQDDLFHHLIHYGTRSSPLVLSMICSTVLNVYHFLRRFIRLQLEAFFTYVLHRVARLGSPIPLQEVALEG
IINFCRQPTFMIEVYANYDCEPLSRSLFEETGKLLCKNSFPMASTVTTLQVEAFEGLATMIHNIADNINREDDSI
PCGPYLVEITKYSPFWEERSKDELESWVEFVRLRKAQKKKILIAANHFNRDEKKGLEYLKLSKLVSDPPDPKAHA
YFFRYTTGLNKNQIGDYLGDPDQFHIEVLKEFTNTFTFSGMILDTALRTYLETFRLPGEAQKIQRILEAFSERFY
DQQSSNVFASKDAVFILCYSLIMLNTDQHNPQVRKKMTEEEFIRNNRAINGGQDLPREYLSELFHSISNHAITLF
GHSGVELEMNPSRWIELMNRSKLTQNFIPCNFDRRLGRDMFACIAGPSVASLSAFFEHTDEEELLHESIEVMFSI
ARIAQYGLEDTLDELLASFCKFTTLLNPYASAEETLFVFSNDMKPKMATLAVFTIANNFGKSIRGGWINIVDCLL
KLKRLKLLPQSVIEFDTAFTSSEIPKESESAIIFPARNPKFGSHSRSRQASGTVSRFSHFLSLESVEDSLSLNVS
EFEQNLKIIKQCRIGNIFSNSSTLPDDALFNLGRSLIFAAAGKGQKFNTPVEEEETVGFCWDLVVAIALANIQRF
PAFWPNFHDYMLAVAQFPTFSAVPFAEKAIECLLKISLKLLSTCRHERFPEELIFKSINLMWKLDKEILDTCCEF
ITQSVNKILTEYPANLQTQLGWKSVLHLLSVTGRHPETYEQGVETLIMLMSDGTHVSRTNYAYCIDCAFGFVALK
NSPLEKNLKILELMADSVNLLSQWHRNPGSESGNNFSIASSTSNSSIEDLRSPVSSNVAINLFVKLGEALRKTSL
ARREELRNHAILALRKSFTVAEDLDFTSNNCINCLSHVIFAMIDDLHEKMLEYSRRENSEREMRSMEGTLKMAMQ
LLTDVYLQFLKRISQSPGFRTFWLGVLRRMDTCMKADLGEHGQSTVEELVPHLLRKMILEMKEREILVKSEGDDL
WEITYIQIQWIAPSLKDELFPEEIF*
**>Carfa_001430**
MGRLKLQAGINAIEEEPEDCEATYSNKATVACMINSEIGAVLAVMRRNVRWGGRYMSGDDQLEHSLIQSLKALRK
QIFLWQHQWHSINPAMYLQPFLDVIRSDETGAPITSVALSAVHKILTLDVIDQNTVNVEDAMHLVVDAVTSCRFE
VTDPASEEVVLMKILQVLLACMKSKASVMLSNQHVCTIVNTCFRIVHQAGSKGELLQRIARHTMHELVRCIFSHL
PDVDNMECALVNGTSTIKQEIGGLNNEYALGSRQVENGNMGSDYDGQPLSTNFASNASTVSVATGINENTIGSGN
GKEAVSYDSHLMTEPYGVPCMVEIFHFLCSLLNVVEHMGMGPKSNTIAFDEDVPLFALGLINSAIELGGPSICRH
PRLLSLIQDELFRNLMQFGLSMSPLILSMVCSIVLNLYHHLRTELKLQLEAFFSCVILRLAQSRYGASYQQQEVA
MEALVDFCRQKTFMVEMYANLDCDITCSNVFEDLANLLSKSAFPVNCPLSSMHILALDGLIAVIQGMAERISNAS
VGSEHAPVNLEEYTPFWMVKCDNYSDPSHWVPFVRRRKYIKRRLMIGADHFNRDPKKGLEFLQGTHLLPDKLDPQ
SVACFFRYTAGLDKNLVGDFLGNHDEFCVQVLHEFAGTFDFQEMNLDTALRLFLETFRLPGESQKIQRVLEAFSE
RYYEQSPQILANKDAALLLSYSLIMLNTDQHNVQVKKKMTEEDFIRNNRHINGGSDLPREFLIELYHSICKNEIR
TTPEQGTSFPEMTPSRWIDLMHKSKKTAPFIVSDSRAYLDHDMFAIMSGPTIAAISVVFDHAEHEEVYQTCIDGF
LAVAKISACHHLEDVLDDLVVSLCKFTTLLNPSSVEEPVLAFGDDTKARMATVTVFTIANRYGDYIRTGWRNILD
CILRLHKLGLLPARVASDAADESELSGDSGHGKPLPNSLSSAHMPSMGTPRRSSGLMGRFSQLLSLDTEEPRSQP
TEQQLAAHQRTLQTIQKCHIDSIFTESKFLHAESLLQLARALIWAAGRPQKGNSSPEDEDTAVFCLELLIAITLN
NRDRIVLLWQGVYDHISNIVQSTVMPCALVEKAVFGLLRICQRLLPYKENLADELLRSLQLVSKLDARVADAYCE
QITQEVSRLVKANASHIRSQLGWRTITSLLSFTARHPEASEAGFDALVFIMSDGAHLLPANYFLCVDTSRQFAES
RVGQAERSVRALDLMAGSVNCLARWASEAKEATGVEETVKMSQDIGEMWLRLVQGLRKVCLDQREEVRNHALLSL
QKCLTGVDEIHLPHGLWSQCFDGVIFTMLDDLHDIALGHSQKDYRNMEGTLILAMKLLSEVFLQLLNDLSQLATF
CKLWLGVLSRMEKYMKVKVRSKKSDKLQQLVPELLQKTLLDMKTRGVLVQRSALGGDSLWELTWLHVNNISLSLQ
SEVFPDQDLEPLHHKQGQTGGGPVSDETVSVPLKETVTSEGAGTGG*
**>Carfa_007592**
MEHINPQFGTNNDERKESPAKPSRGALACMVNSEIGAVLAVMRRNVRWGVRYVADDDHLEHSLIHSLKELRKRIF
SWQHQWHKIDPAVYLQPFLDVIQSDETGAPITGVALSSVYKILSLDILDLDTVNVGDAMHLIVDAVTSCRFEVTD
PASEEVVLMKILQVLLACMKSAASVRLSNQHVCNIVNTCFRVVHQASSKGELLQRIARHTMHELVRCIFSHLPDI
NNIEHALANGSGSSVRKEVGSTDEDHTSENKQLENGNAGVEYDGQSSTYASKAPMSASASGTSDSGMHENITEVS
NGMEGTKNDEKVMMEPFGVPCMVEIFHFLCSLLNVIEHVEFGPRSNPIAYDEDVPLFALGLINSAIELAGPSINK
HPKLLAIVQDELFHNLMQFGSSMSPLILSTVCSIVLNLYHHLRTELKVQLEAFLSCVLLRLAQSKHGSSYQQQEV
AMEALVDLCRQQTFMAEMYANFDCDITCSNVFEDLANLLSKNAFPVNGPLSAMHILALDGLISMVQGMAEKIGDE
PATSEASMDDEGHKAFWTMTCENYSDPNCWVPFIRKVKYIKRKLMIGADHFNRDPKKGLEFLQGMHLLPEKLDPL
SVACFFRYTAGLDRNLVGDYLGNHDEFCVQVLHEFARTFDFREMNLDTALRLFLGTFRLPGESQKIHRVLEAFSE
RYYAQSPRILVNKDVALVLSYSLIMLNTDQHNTQVKKKMTEEDFIRNNRRINAGKDLPREFLSELFHSICENEIQ
MIPDQSTGFPAMASSRWINVMHKSKTVSPFIICDSRELLIHDMFAILSGPTIAAISVVFDQVEHEDVLQTCIDGF
LSIAKIAACYHLHDVLDDLIVSLCKFTTLLTPLSVEEAIVALGDYTKVRMATATVFTIANSYGDYIRSGWKNILD
CVLSLHQLGLVPTSLASDAVHDMEPSSDPEQGKRATSLSISNVPSVPTRKSSGLMGRFSQLLSFDMEEPRTQPTE
EQLAAEQRTRETIQTCHIDSIFTESKFLQAESLLQLVKSLILAADRLRKGTSSIEDEDTAVFCLELLIAITLNNR
DRIMLIWQGVYEHITDIVQSTIMPCTLVEKAVFGLLKICQRLLPYKENLTDELLRSLQLVLKLDARVADAYCEHI
TQEVMRLVKANATHIRSHVGWRTIISLLSITARHPEASEVGFEALAFIMSDGAHLLPSNYVLCVDAARQFGESRV
GEVDRSVSALDMMAGSVVCLVRWSCETTIAVGEEAATKVNQDIGEMWLRLVQGLRKVCLDQREEVRNHAILMLQR
SFAGVDGSHLPNTLWLQCFDVVIFALLDDLVEIALCNSPKDYRNMEGTLVLAMKLMSKVFLQLLLDLWRLPSFCK

LWLGVLNRMERYMNIKFRGRRSEKIHELIPELLKNTLFVMKTTGLLTPSDTIGGDSFWQLTWLHVKNISPSLQSE
VFPAPELEQLQKKQIKAAGVPSPDGTVLVQPSETTA*
**Carfa_003276**
MEKSSDQNHDAEENNCPSCREKKKGSAAKSRRKELGVSCMLNTELGAVLAVIRRPPDPASQYFSPDEHYFDSAIL
QSLKSLRALIFNPQQEWRTIDPSIYLSPFLDVIQGDDIPAAATGVALSAILKILRLEIFDEKTPGAKESINSIVN
GITSCRLEKTDPVSEDAVMMKILQVLTATMKHPASLLLTDHAVCTIVNTCFQVVQQSASRGDLLQRSARYAMHEL
IQTIFSRLPDIEVKDGEDSESDSDDVDVGSNSDAGYGVRCAVDIFYFFCSLLNVVEVVETEGSTSHTVDEDVQLF
ALVLINSAIELSGDAIGKHPKLLRMIQDDLFHHLIHYGTRSSPLVLSMICSSVLNIYNFLRRFIRLQLEVFFAYV
LLRVAALGSAMQLQEVALEGIINFCRQPTFVIEVYVNYDCDPLGRNLFEEIGKLLCKDSFPVASPVTTLQVQAFE
GLAAMIHNIADSVDREDNSSPFGPYPLEITEYSPFWEEKSKDELESWEEYVRIRKAQKKKILIAANHFNRDEKKG
LEYLKLSRLVSDPPDPKAHAFFFRYTPGLNKNTIGDYLGDPDELHIEVLKEFTDTFKFSGMILDTALRTYLETFR
LPGESQKIQRILEAFSEKYYDQQSSDVFASKDAVFILCYSLIMLNTDQHNPQVKKKMTEDEFIRNNRAINAGKDL
PREYLSELFHSISNNAITLFGHSAVSLEMTPSRWIELINRSKIAQPFIVCDFDRHLGRDMFACIAGPSVASLSAF
FEHIDEEELLHKCVEVLFSIARIAQYGLEDTLDELLASFCKFTTLLNPYASAEETLFAFSNDMKPKMATLAVFTV
ANNFGESIRGGWRNIVDCLLKLKRLKLLPQSVIEFDTASTSSPDVPKESESAIIFPTRDKFGSRRASGTISRFSH
FLSLDSMEDSLSLNVSEFEQNLKIIKQCRIGNIFSNSSNLPDDALFNLGRSLIFAAAGKGQKFSTPVEEEETVGF
CWDLVIAIAMANIQKFQAFWPNFHEYMLAVAQFPMFSPIPFAEKAIIGLLRLSLKLLATYRSEKFPEELIFKSIN
LMWKLDKEILDTCCEFITQSVSKILIEYPANLQTQLGWKSVLHLLSVAGRHSETYDQAVETLIMLMSDGTHVSRT
NYAYCIDCAFGFVALKNSPLEKNLKILDLMADSVNLLIQWYKNHYSEPGNNFSIASNTSNSSAEDNSRGLGSASF
VMNLFLKLGEALRKSSLARREEMRNHAILALQKSFTLAEELDFTSTNCINCFNLIVFAMIDDLHEKMLEYSRRDN
SEREMRSMEGTLKTAMELLTDVYLQFLKQISESYGFRTFWLGVLRRMDTCMKADLGEYGKSTIEELVPDLLKKMI
TLMKEKEILVQKEGDDLWEITYIQIQWIAPSLKEELFPELAF*
**>CaseqGNOM_sc309**
MGRLKLQAGINSIEEEPEECDGTYSNKATLACMTNSEIGAVLAVMRRNVRWGGRYMSSDDQLEHSLIQSLKALRK
QIFLWQNQWHSINPAMYLQPFLDVIRSDETGAPITGIALSAVHKILTLDVIDQNTVNVEDAMRLVVDAVTSCRFE
VTDPASEEVVLMKILQVLLACMKSKASVMLSNQHVCTIVNTCFRIVHQAGSKGELLQRIARHTMHELVRCIFSHL
PDVDNMECALVNGTSTINQGVGGLNNEYAMGTQQVENGNMASDYEGQPLSTNFASNASAGSVATGVDENMIGSGN
GKEAVSYDLHLMTEPYGVPCMVEIFHFLCSLLNVVENIGPGTKSNSMAFDEDVPLFALGLINSAIELGGPSICRH
PRLLSLIQDELFRNLMQFGLSMSPLILSMVCSIVLNLYHHLRMELKLQLEAFFSCVILRLAQSRYGASYQQQEVA
MEALVDFCRQKTFMVEMYANLDCDITCSNVFEDLANLLSKSAFPVNCPLSSMHILALDGLIAVIQGMAERISNAS
VGSEHAPVNLEEYTPFWMVKCDNYSDPNHWVPFVRRRKYIKRRLMIGADHFNRDPKKGLEFLQGTHLLPDKLDPQ
SVACFFRYTAGLDKNLVGDFLGNHDEFCVQVLHEFAGTFDFQGMNLDTALRLFLVTFRLPGEAQKIQRVLEAFSE
RYYEESPQILANKDAALLLSYSLIMLNTDQHNVQVKKKMTEEDFIRNNRHINGGGDLPREFLSELYHSICKNEIR
TTPEQGAGFPEMTPSRWIDLMHKSKKTAPFIVSDSRAYLDHDMFAIMSGPTIAAISVVFDHAEHEEVYQTCIDGF
LAVAKISACHHledvlddlvvSLCKFTTLLNPSSVEEPVLAFGDDSKARMATVTVFTIANRYGDYIRTGWRNILD
CILRLHKLGLLPARVasdaadeselsadsGHGKPLPNSLSANQMPSIGTPRRSSGLMGRFSQLLSLETEEPRSQP
TEQQLAAHQRTLQTIQKCRIDSIFTESKFLHAESLLQLARALIWAAGRPQKGNSSPEDEDTAVFCLELLIAITLN
NRDRIVLLWQGVYEHIANIVQSTVIPCALVEKAVFGLLRICQRLLPYKENLADELLRSLQLVLKLDARVADAYCD
QITQEVSRLVKANASHIRSQLGWRTITSLLSITARHLEASEAGFDALSFIMSDGAHLLPANYVLCVDASRQFAES
RVGQAERSLRALDLMAGSVNCLAGWASETKEAMGMEEALKMSQDIGEMWLRLVQGLRKVCLDQREEVRNHALLSL
QKCLTGVDEIHLPHALWLQCFDGVIFTMLDDLHEIAQGHTQKDYRNMEGTLILAMKLLSKVFLQLLHDLSQLTTF
CKLWLGVLSRMEKYMRMKVRGKKSEKLQELVPELLKNTLLVMKTRGVLVQRSALGGDSLWELTWLHVNSISSSLQ
SEVFPDQDSGSSVHKQVQTGEVSISNETGSLPVNETVTSGGSGTGG*
**>CaseqGNL1_sc190**
MGAINSEIGTNNNERKDSTGRPSRGALACMVNSEIGAVLAVMRRNVRWGVRYMADDDHLEHSLIHSLKDLRKHIF
SWQHQWHKIDPAVYLQPFLDVIRSDETGAPITGVALSSVYKILSLDILDLDTVNAGDAMHLIVDAVTSCRFEVTD
PASEEVVLMKILQVLLACMKSTASVRLSNQHVCNIVNTCFRVVHQASSKGELLQRIARHTMHELVRCIFGHLHDI
VNTEHALANGSRSSVGEEVGSADGVHIVEDKQLENGNAGVEYDGQSSSFVPNAPMSASAGGVSDIRANENITEVS
NGVEPTKNEEKVMMEPFGVPSLVEIFHFLCSLLNVVEHTEFGPRSNPIAYDEDVPLFALGLINSAIELAGPSIGK
QPQLLAIVHDELFHNLMQFGLSMSPLILSTVCSIVLNLYYHLRTELKVQLEAFLSRVLLRLVQTKYGSSYHQQEV
AMEALVDLCRQQTFMAEMYANFDCDITCSNVFEDLANLLSKSAFPINGPLSAMHILALDGLISMVQGMAESIGNE
PASSEASIDDDGYRSFWTMTCENYSDPNCWVPFIRKVKYIKRKLMIGADHFNRDPKKGLEFLQGMHLLPEKLDPQ
SVACFFRYTSGLDKNHVGDFLGSHDEFCVQVLHEFARTFDFRDMNLDAALRLFMGTFRLPGESQKIQRVLEAFAE
RYYEQSPHILANKDAALLLSYSLILLNTDQHNTQVKKKMSEEDFIRNNRGINGGKDLPREFLSELYYSICENEIQ
MIPDQSTGFPAMASSCWINVMNKCKTVSPFIVCDSREPLIYDMFAILSGPTIAAISVVFDLVEQEDVLQTCIDGF
LSVAKIAASYHLTDVLDDLIVSLCKFTTLLTPLSLEEAIVALGDDTKVRMATTTVFTIANSYGDYIHSGWKNILE
CVLSLHKFGLLPSSLASDAVHDTEPPSDPEPEKHATTSLSTSNVPPVPTRKSSGLIGRFSQLLSFDMEEPRVQPT
EEELAAQQRTRDTIRTCHIDSIFTESKFLQAESLLQLVKALILAAGRLRKGTSGVEDEGTTVLCLELLVAITLNN
RDRIMLIWQGVYEHISDIVQTTIMPCTLVEKAVFGLLKICQRLLPYKEDLTDELLRSLQLVLKLDARVADAYCEH
ITQEVLRLVKANATHIRSHAGWRTIISLLSITARHPEASEVGFEALAFIMTNGAHLLPSNYVLCVDAARQFAESR
VGGVDRSVSALDMMAGSIVCLGRWSLetkttageeaatkVNQDIGEMWLRLVQGLRKVCLDQREEVRNHAVLMLQ
RSLAGVDGIHLPNSLWLQCFDLVIFTLLDDLVEIALGNSPKDFRNMEGTLVLAMKLMSKVFlqllldlwqlPSFC

KLWLGVLNRMERYMSIKVRGRRSENIHELIPELLKNTLFVMKTAGLLIPSDTIGGDSFWQLTWLHVKNISPSLQS
EVFPASELEQLQTKQIKVAEVPRSDGSVLVQSSETKA*
**>CaseqGNL2_sc144**
MDESTCQNPDTDEKNSPSCRPKKKSSAAESRRKELGLSCMLNTEVGAVLAVIRRPPEPTSQYFFPDEHYFDASIL
QSLKSLRAVIFNPEQVWRNIKPSVYLSPFLDVIQGDDIPAAATGVALSAVLKMLRLEIFDEKTPGAKEAINSVVN
AISSCRLEKTDPTAEDAVMMKILQVLTAVMKHRSSSLLSDRAVCTIVNMCFQLVQlaasrsdlllrsarssMHEL
IQTIFSYLPDFEVKDGEDSESDIDDVSAGGNSDAKYGVRCAIEIFYFLCSMLNVIAVEETEWSASHNVDEDLQLF
ALVLINSAIEIGGNGMGKHPKLLRMIQDDLFQHLVHFGMHASPFVLSMICSTVLNIYHFlrrlglngl*lirigM
KLRYGDKNAICFGRFILLQLQALFAYVLLRVAAPGNPIPLQEVALEGIINFCRQPTFIVEVYVNYDCDSICRNLF
EEIGKLLCKEAFPMASPVTPLQVQAFEGLAAVNHTIADNVDKEDDSSPFVPYPVEITHYSPFWEEKSMDELESWV
EFVKLRKAQKKKILIAANHFNRDEKQGLEYLKLSKLVSDPPDPKAHAFFFRYTPGLDKSMIGDYLGDPDQFHIEV
LKEFTDTFNFSGVILDTALYTFLETFRLPGESQKIQRILEAFSEKFYDQQSSDLFASKDAVYIFCYSLIMLNTDQ
HNPQVRKKMTEDEFIRNNRAINAGQDLPRGYLSELFHSISNNAIAVFSQSGAPLEMNPSRWIELINRSKLSRPFV
QCDFDRCLGRDMFACIAGPAVASLSAFFERIEEEELLHKCVEVVFSIARISQYGLEDTLDELLASFCKFTTLLNP
YASAEETLFAFSNDMKPKMATLTVFTIANNFGGTIRGGWRNIVDCllklkrlkllPQSVIEFDTASSSSPDDIKE
SESVTSFPARNPQLGSRCASGTISRFSHFlsldsvddslNVSEFEQNLKVIKQCRIGNIFSNSSNLPDDALFNLG
RSLIFAAAGKGQKFSTSVEEEETVGFCWDLVIAIAMANIHKFQAFWPNFHEYILAVAQFPSFSPIPFAEkailal
lklclklla THRPEKLPEELIFKSINLMWKLDKEILDTCCEYITQSVGKILIEYPANLQTQLGWKSVLHLLSVTG
RHPETYDQAVETLIMLMSDGIHVSRANYAYCIDCAFGFVAIKNSPLEKSLRILDLMSDSVNLLIQWYRNHCPEPG
nnfsiasstsnssaddnsRGLGSSNFAMNFFLKLGEALRKTSLSRREEMRNHAILALQKIFTRAEELDFTSTNCL
NCFNLVIFAMVDDLHEKILEYSRRDNAEREMRSMEGTLKIAMELLTDVYLQFLKQVSESPGFRTFWLGVLRRMDT
CMKADLGEYGKSTIEELIPDLLKKMITQMKEREILVQKEGDDLWEITYIQIQWIAPSLKEELFPEEIF*
**>Cucsa_LOC101204638**
MGRLKLQTGINAIEEEPEECDVTYTNKTALACMINSEIGAVLAVMRRNVRWGGRYMSGDDQLEHSLIQSLKSLRK
QIYSWQHPWHTINPAVYLQPFLDVVRSDETGAPITGVALSSVYKILTLDMIDQNTVNAGDSVHLIVDAVTCCRFE
LTDPASEEMVLMKILQVLLACMKSKASIMLSNQHVCTIVNTCFRIVHQAATKGELLQRIARHTVHELVRCIFSHL
SEINTTERALVNGNSSSKQEAGRGANDDYVLGNRLLENGNVGHEFDGQSSSNNFDSNPSSGLMATGMEENLLEDG
SAKDTVPFDFHLMNEPYGVPCMVEIFRFLCSLLNLVEHMELGARSNTMAFDEDVPLFALGLINSAIELGGPSFRH
HPRLLSLIQDELFRNLMQFGLSTSSLILSMVCSIVLNLYHHLRTELKLQLEAFFSCVILRLAQSRYGASYQQQEV
AMEALVDFCRQKTFMVEMYANLDCDITCSNVFEDLANLLSKSAFPVNCPLSSMHILALDGLIAVIQGMAERIGNG
AGLENTPVNLEEYTPFWMVKCENYSDPTQWVPFVRRKKYIKRRLMIGADHFNRDPKKGLEFLQGTHLLPDKLDPK
SVACFFRYTAGLDKNLVGDFLGNHDEFCVQVLHEFAWTFDFQDMNLDTALRLFLETFRLPGESQKIQRVLEAFSE
RYYEQSPQILVNKDAALLLSYSLIMLNTDQHNVQVKKKMTEEDFIRNSRHINGGNDLPRDFLSELYHSICKNEIR
TTPEQGNGFPEMTPSRWIDLMHKSKKSSPFIVSDSKAYLDRDMFAIMSGPTIAAISVVFDHAEHEEVYQTCIDGF
LAVAKISACHHLEDVLDDLVVSLCKFTTLMNPSSVEEPVLAFGDDTKARMATMTVFTIANRYGDFIRTGWRNILD
CILRLHKLGLLPARVASDAADESELSADAGHGKPLTSSLSAAHIQSIGTPKRSSGLMGRFSQLLSLDSEEPRSQP
TEQQLAAHQRTLQTIQKCNIDSIFTESKFLQAESLLQLAQALIWAAGRPQKGNSSPEDEDTAVFCLELLIAITLN
NRDRIVLLWPGVYDHISNIVQSTVMPCALVEKAVFGLLRICQRLLPYKENLADELLRSLQLVLKLDARVADAYCE
QITQEVSRLVKANASHIRSPSGWRTITSLLSITARHPEASEAGFDALLFIVSDGAHLLPANYTLCIDASRQFAES
RVGQAERSLRALDLMAGSVDCLGRWAKEGKEAAREEEAIKMSQDIGDMWLRLVQGLRKICLDQREEVRNQALLSL
QKCLTGVDEINLPHDLWLQCFDLVIFTMLDDLLEIAQGHSQKDYRNMEGTLILAMKLLSKVFLLLLQDLSQLTTF
CKLWLGVLSRMEKYAKAKVRGKRSEKLQELVPELLKNNLLVMKTKGVLVQRSALGGDSLWELTWLHVNNISPSLQ
SEVFPDQDSNRVLGQGEKGGLTSSEANSVSSTEKVASDNAGTGG*
**>Cucsa_LOC101207938**
MGRPKLHTGINAIEEEQEECDVTFTYMNKTALACMINSEIGAVLAVMRRNVRWGGRYMSGDDQLEHSLIQSLKSL
RKQIYSWQHPWHTINPAVYLQPFLDVIRSDETGAPITGVALSSVYKILTLDMIDQNTVNAGESVHLIVDAVTCCR
FELTDPASEEMVLTKILQVLLACMKSKASIMLSNQHVCTIVNTCFRIVHQAATKGELLQRIARHTVHELVRCIFS
HLSEINTTEHALVNGNTSSKQEAGRGANDDYALGSRLLENGNLDHEFDGQSPSTNLDSKPSSGLMVTGMEENLLE
DGSVKDTVPFDFHLMNEPYGVPCMVEIFRFLCSLLNLVEHMELGARSNTMAFDEDVPLFALRLINSAIELGGPSF
RHHPRLLSLIQDELFRNLMQFGLSTSSLILSMVCSIVLNLYHHLRTELKLQLEAFFSCVILRLAQSRYGASYQQQ
EVAMEALVDFCRQKTFMVEMYANLDCDITCSNVFEDLANLLSKSAFPVNCPLSSMHILALDGLIAIIQGMAERIG
NGTGLENTPVNLEEYTPFWMVKCENFSDPIEWVPFVRRKKYIKRRLMIGADHFNRDPKKGLEFLQGTYLLPDKLD
PKSVACFFRYTAGLDKNLVGDFLGNHDEFCVQVLHEFAWTFDFQDMNLDTALRLFLETFRLPGESQKIQRVLEAF
SERYYEQSPQILVNKDAALLLSYSIILLNTDQHNVQVKKKMTEEDFIRNSRHINGGNDLPRDFLSELYHSICKNE
IRTTPEQGNGFPEMTPSRWIDLMHKSKKSSPFIVSDSKVYLDRDMFAIMSGPTIAAISVVFDHAEHEEVYQTCID
GFLAVAKISAWHHLEDVLDDLVVSLCKFTTLVNPSSVEEPVLAFGDDIKARMATMTVFTIANRYGDFIRTGWRNI
LDCILRLHKLGLLPARVASDAADESELSSDAGHGKPLSSSLSVAHIQSIGTPKRSSGLMGRFSQLLSLDSEEPRS
QPTEQQLAAHQRTLQTIQKCNIDSIFTESKFLQAESLLQLAQALIWAAGRPQKGNSSPEDEDTAVFCLELLIAIT
LNNRDRIVLLWPGVYDHISNIVQSTVMPCALVEKAVFGLLRICQRLLPYKENLADELLRSLQLVLKLDARVADAY
CEQITQEVSRLVKANASHIRSPSGWRTITSLLSITARHPEASEAGFDALLFIVSDGAHLLPANYTLCIDASRQFA
ESRVGQVERSLRALDLMAGSVDCLRRWAKEGKEAVREEEVIKISQDIGDMWLRLVQGLRKVCLDQREEVRNQALL
SLQKCLAGVDEIRLPHDLWLQCFDLVIFTVLDDLLEIAQGHSQKDYRNMEGTLILAVKLLFKVFLLLLQDLSQLT

TFCKLWLGVLSRMEKYAKAKVRGKRSEKLQELVPELLKNNLLVMKTKGVLVQRSALGGDSLWELTWLHVNNISPS
LQSEVFPGQDSNFELGQGEKSGLTSSEANSVTSSDKVVSDNAGTGG*
**>Cucsa_LOC101210509**
MDKRSSDDEDEKDSPKSKRRELGLSCMLNTEVGSILAVIRRPPSELNSPYISTIDETYDSTIQQSLKSLRALIFH
PQQKWRTIDPSIYISPILDVIQSDDIPAAATGVALSALLKIIKVEIFDEKTPGAKDAINLIVIGITNCKLEKTDL
VTEDAVMMKVLQVLAGLMNHRASYLLNDQSVCTIVNTCFNVVQQSASRGDLLQRTARYTMNELIQIIFSRLPEIE
VRDGEESESDTEDADLGGSLDSGYGIRCVIDVFHFLCSLLNVVEIMVEVGDGGLGSRTADEDVQLFALVLINSAV
ELSGDAIGKHPKLLRMVQDDLFHHLIHYGAASNPLVLSMICSTVLNIYHFLRRFVRLQLESFFVYVALKLASFGN
STQIQEVALEGIINFCRQSSFILEFYVNYDCDPLRWNLLEEIGKLLCKLSFPTGSPLTTLNIQAFEGLVIVIHNI
AEKLDKHKEETCGGGGNLRVYPAQVDEYIPFWEEKSKEDLDLEDWLRYVRVRKAQKKKILIAGHHFNRDEKKGLA
YLKLSLLVSDPPDPKAYAYFFRYTHGLDKQFVGEYLGDPGQFHVKVLAEFTETFEFTGMILDTALRTYLETFRLP
GEAQKIHRILEAFSERFYELQSSNTFASKDTVFVLCYSLIMLNTDQHNPQVKKKMTEDEFIRNNREINAGKDLPR
DYLSELFHSISNNAIILSPQSGLQLDMNPSKWVELMNRSKIIQPFMSYDFDPRLGRDMFGCIAGPSVASLAAFFE
HADEDEMLNECIEGLFSIAKITQYGLEDTLDELLAMFCKFTTLLNPYASAEETLFVFSHDMKPKLATLAVFTIAN
NFGDTIRGGWRNIVDCLLKLKRLKLLPQSVIDFEVASTSSNDVARSDSGVIFPSQDPKFCTQQSSGMVSRFSQFL
SLDSMEDSLTLNLNEYEQNLKFIKQCRIGNIFSNSSNILDEALLNLGRSLIFAAAGKGQKFSTPIEEEETVGFCW
DLIITMTMANLYRFQVFWPNFHEYLQTVVQFPLFSAIPFAEKAVLGLFKVCLRLLSTYQPDKIPEELIFKSINLM
WMLDKEILDTCFESITQSVSKILIEYPANLQSQIGWKSLLHLLSATGRHPETYDQGVETLIMLMSDASHITRTNY
TFCIDCAFSYVALKNSPLDKNLKILDALSDSVNFLVQWYRNYCAESGNSFSVASNASSSSLDEKGLGSSNFALTL
FLKLGEALRKTSLARREEIRNHAITSLKKSFVLAEELDFPPTNCIGCFNNIIFAMVDDLHEKMLEYSRRDNAERE
ARSMDGTLKISMELLTDVYLVYLKQISESPGFRTFWLGVLRRMDTCMKADLGSYGESSLKDLIPELLRKIITTMR
EKEILVKKEGEDLWEITYIQIQWIAPGIKDELFPEECF*
**>Cucma_LOC111490452**
MGRLKLQTGINSIEEEPEECDVTYTNKTALACMINSEIGAVLAVMRRNVRWGGRYMSGDDQLEHSLIQSLKSLRK
QIYSWQHPWHTINPAVYLQPFLDVIRSDETGAPITGVALSSVYKILTLDMIDQNTVNAGDAVHLIVDAVTCCRFE
ITDPASEEMVLMKILQVLLACMKSKASIMLSNQHVCTIVNTCFRIVHQAASKGELLQRIARHTVYELVRCIFSHL
SEVNTTERALVNGNSSSKQEAGRGANDDYVLGSRLLENGNLGHEFDGQSSSNNFDSNSPSGLMATGMEENLLEDG
SVKDTVPFDFHLMKEPYGVPSMVEIFRFLCSLLNLVEHMELGARSNTMAFDEDVPLFALGLINSAIELGGPSFRH
HPRLLSLIQDELFRNLMQFGLSTSSLILSMVCSIVLNLYHHLRTELKLQLEAFFSCVILRLAQSRYGASYQQQEV
AMEVLVDFCRQKTFMVEMYANLDCDITCSNVFEDLANLLSKSAFPVNCPLSSMHILALDGLIAVIQGMAERIGNG
TGLENTPVNLEEYTPFWLVKCENYSDPNQWVPFVRRKKYIKRRLMIGADHFNRDPKKGLEFLQGTHLLPDKLDPK
SVACFFRYTAGLDKNLVGDFLGNHDEFCVQVLHEFAWTFDFQDMNLDTALRLFLETFRLPGESQKIQRVLEAFSE
RYYEQSPQILVNKDAALLLSYSLIMLNTDQHNVQVKKKMTEEDFIRNSRHINGGNDLPRDFLSELYHSICKNEIR
TTPEQGNGFPEMNPSRWIDLMHKSKKSSPFIVSDSKAYLDRDMFAIMSGPTIAAISVVFDHAEHEEVYQTCIDGF
LAVAKISACHHLEDVLDDLVVSLCKFTTLLNPSSVEEPVLAFGDDTKARMATMTVFTIANRYGDFIRTGWRNILD
CILRLHKLGLLPARVASDAADESELSADTGHGKPLTSSLSAAHIQSIGTPKRSSGLMGRFSQLLSLDTEEPRSQP
TEQQLAAHQRTLQTIQKCNIDSIFTESKFLQAESLLQLAQALIWAAGRPQKGSSSPEDEDTAVFCLELLIAITLN
NRDRIVLLWPGVYDHISNIVQSTVMPCALVEKAVFGLLRICQRLLPYKENLADELLRSLQLVLKLDARVADAYCE
QITQEVSRLVKANASHIRSPSGWRTITSLLSITARHPEASEAGFDALLFIVSDGAHLLPANYTLCIDASRQFAES
RVGQAERSLRALDLMAGSVDCLARWAREGKEAAREEEAIKMSQDIGDMWLRLVQGLRKVCLDQREEVRNQALLSL
QKCLTGVDEVYLPHGLWLQCFDLVIFTMLDDLLEIAQGHSQKDYRNMEGTLILAMKLLSKVFLLLLQDLSQLTTF
CKLWLGVLSRMEKYAKAKVRGKRSEKLQELVPELLKNTLLVMKTKGVLVQRSALGGDSLWELTWLHVNNISPSLQ
SEVFPDQDSDRLLGQGEKGGQTSSESNSSSTEKVAPDSVGTGG*
**>Cucma_LOC111476959**
MGRLKLQTGINAIEEEPEECDVTYTNKTALACMINSEIGAVLAVMRRNVRWGGRYMSGDDQLEHSLIQSLKSLRK
QIYTWQHPWHTINPAVYLQPFLDVIRSDETGAPITGVALSSVYKILTLDMIDQNTVNAGDSVHLIVDAVSCCRFE
LTDPASEEMVLMKILQVLLACMKNKASIMLSNQHVCTIVNTCFRIVHQAATKGELLQRIARHTVYELVRCTFSHL
SEISTTEHALVNGNSSSKQEAGRGSNDDYVLGSRLLENGNLGHEFDGQLSSNNFDSNSSSAPLATGMDENLLEDS
NVKDTVPFDLHLMNEPYGVPCMVEIFRFLCSLLNLVEHMELGGRSNTMAFDEDVPLFALGLLNSAIELGGPSFRH
HPRLLSLIQDELFQNLMQFGLSTSSLILSMVCSIVLNLYHHLRTELKLQIEAFFSCVILRLVQTRYGASYQQQEV
AMEALVDFCRQKTFMVEMYANLDCDITCSNVFEDLANLLSKSAFPVNCPLSSMHILALDGLIAVIQGMAERIGNG
TGIENTPVNLEEYTPFWMVKCENYNDPNQWIPFVRRKKYIKRRLMIGADHFNRDPKKGLEFLQGTHLLPDKLDPK
SVACFFRFTAGLDKNLVGDFLGNHDEFCVQVLHEFAGTFDFQDMNLDTALRLFLETFRLPGESQKIQRVLEAFSE
RYYEQSPQILVNKDAALLLSYSLILLNTDQHNVQVKKKMTEEDFIRNSRHINGGNDLPRDFLSELYHSICKNEIR
TIPEQGNGFPEMTPSRWIDLMNKSKKSSPFIVSDSKAYLDRDMFAIMSGSTIAAISVVFDHAEHEEVYQTCIDGF
LAVAKISACHHLEDVLDDLVVSLCKFTTLMNPSSVEEPVLAFGDDTKARMATTTVFNIANMYGDFIRSGWRNILD
CILRLHKLGLLSARVASHAADESELSSDTGHGKPLTSSLSAAHIPSVGTPKRSSGLMGRFSQLLSLDNEEPRSQP
TEQQLAAHQRTLQTIQKCNIDSIFTESKFLQAESLLQLAQALIWAAGRPQKGNSSPEDEDTAVFCLELLIAITLN
NRDRIVLLWPGVYDHISNIVQSTVMPCALVEKAVFGLLRICQRLLPYKENLADELLRSLQLVLKLDARVADAYCE
QITQEVSRLVKANASHIRSPSGWRTITSLLSITARHPEASEAGFDALLFIVSDGAHLLPANYTLCIDASRQFSES
RVGQAERSLRALDLMAGSVDCLARWAREGKEAAREEEVIKMSQDIGDMWLRLVQGLRKVCLDQREEVRNQALLSL
QKCLTGVDEIHLPHGLWLQCFDLVIFTMLDDLLEIAQGHSQKDYRNMEGTLILAMKLLSKVFLLLLQDLSQLTTF

CKLWLGVLNRMEKYAKAKVKGKRSEKLQELVPELLKNTVLVMKTKGVLVQRSALGGDSLWELTWLHVNNISPSLQ
SEVFPDQDSNRLLGQGEKGGPTSSEANSVSSAEKVAADSAGTGG*
**>Cucma_LOC111465412**
MEKRFSRHEDGKESSKSKRRELGLSCMLNTEVGAVLAVIRRPPSELNAPYMCTADDTYDSSIQQSLKSLRALIFQ
PQQKWRTIDPSIYISPILDVIQSDDIPAAATGVALSALLKIIKVEIFEEKTPGAKDAINLIVLGITNCKLEKTDV
VTEDAVMMKILQVLAGLMNHRASYLLNDQSVCTIVNTCFNVVQQSASRGDLLQRTARYTMNELIQIIFSRLPEIE
FRDGEDSESDTEDADLGGSMDSGYGIRCVIDVFHFLCSLLNVVETAEMVDGLGSRTADEDVQLFALVLINSAVGL
SGDAIGKHPKLLRMIQDDLFHHLIHYGARSNPLVLSMICSTVLNIYHFLRRFVRLQLEAFFIYVVLRLASFGNST
QIQEVALEGIINFCRQSSFILEFYVNYDCDPFRSNLFEEIGKLLCKLSFPMGSPLTTLQIQAFEGLVIMIHNIAE
KLEKEESSGGSSRVYPAQINVYRPFWEEKSKDDLQYENWLDYIRVRKAQKKKVLIAGHHFNRDEKKGLAYLKLCQ
LVSDPPDPKAYAFFFRYTNGLDKQLIGEYLGDPDQFHVRVLAEFTDTFDFTSMILDTALRTYLETFRLPGEAQKI
HRILEAFSERFYILQSSNTFASKDTVFVLCYSLIMLNTDQHNPQVKKKMTEDEFIRNNREINAGKDLPRDYLSEL
FHSISNHAIILSPQSGLQFDMNPSKWVELMNRSKIIQPFMLCDFDHRLGRDMFACVAGPSVASLAAFFEHADEDE
MLNECIEGLFSIAKITQYGLEDTLDELLAIFSKYTTLLNPYASTEETLFAFSHDLKPKLATLAVFTIANNFGDSI
RGGWRNIVDCLLKLKRLKLLPQSVIDFEVASTTSNDMAKSDSGVIFPSQDPKFGTQQSSGMAGRFSQFLSLDSME
DSLSLNLNEFEQNLKFIKQCRIGSIFSSSSSLHDEALLNLGRSLIFAAAGKGQKFSTPVEEEETVGFCWDLITTM
SLANVYRFQVFWPSFHEYLQTVVQFPLFSAIPFAEKAVLGLFKVCLKLLSTYQPEKHPEELIFKSINLMWMLDKE
ILDTCFESITQSVSKIIIEYPANLQTAIGWKSLLHLLSATGRHPETYEQGVETLIMLMSDGTHITRTNYTFCIEC
AFSYVALKNSPLDKNLRILDLLSDSVNFLVQWYRNYCAESGNSYSVASNASSSSVEDKGLGSNFALTLFIKLGEA
LRKTSLARREEIRNHAVISLKKSFMLAEELDFTSTNCINFFNLVVFAMVDDLHEKMLEYSRRDNAKREARSMEGT
LKISMDLLTDVYLLFLKQISESSGFRTFWLGILRRMDTCMKADLGSYGESSLKDLVPELLRKIITKMREKEILMK
KEGDDLWEITYIQIQWIAPAIKEELFPEESF*
**>Momch_LOC111010039**
MGRLKMQTGINAIEEEPEEFDVTYTNKTALACMINSEIGAVLAVMRRNVRWGGRYMSGDDQLEHSLIQSLKALRK
QIYSWQHPWHTINPAVYLQPFLDVIRSDETGAPITGVALSSVYKILTLDMIDQNTVNAGDAVHLIVDAVTCCRFE
LTDPASEEMVLMKILQVLLACMKSKASIXLSNQHVCTIVNTCFRIVHQAGSKGELLQRIARHTVHELVRCIFSHL
SEINTTEHALVNGNSSSKQEAGRGANDDYVLGSRLLENGNLGHEFDGQSSSNNFEINSSSGLMATGMEENLLEDG
STKDTVPFDFHLMNEPYGVPCMVEIFRFLCSLLNLVEHMELGARSNTMAFDEDVPLFALGLINSAIELGGPSFRH
HPRLLSLIQDELFRNLMQFGLSTSSLILSMVCSIVLNLYHHLRTELKLQLEAFFSCVILRLAQSRYGASYQQQEV
AMEALVDFCRQKTFMVEMYANLDCDITCSNVFEDLANLLSKSAFPVNCPLSSTHILALDGLIAVIQGMAERIGNG
TGLENSPVNLEEYTPFWMVKCENYSDPSQWVPFVRRKKYIKRRLMIGADHFNRDPKKGLEFLQGTHLLPDKLDPK
SVACFFRYTAGLDKNLVGDFLGNHDEFCVQVLHEFAWTFDFQDMNLDTALRLFLETFRLPGESQKIQRVLEAFSE
RYYEQSPQILVNKDAALLLSYSLIMLNTDQHNVQVKKKMTEEDFIRNSRHINGGNDLPRDFLSELYHSICKNEIR
TTPEQGNGFPEMTPSRWIDLMHKSKKSSPFIVSDSKAYLDRDMFAIMSGPTIAAISVVFDHAEHEEVYQTCIDGF
LAVAKISACHHLEDVLDDLVVSLCKFTTLLNPSSVEEPVLAFGDDTKARMATMTVFTIANRYGDFIRTGWRNILD
CILRLHKLGLLPARVASDAADESELSADTGHGKPLTSSLSAAHIQSIGTPKRSSGLMGRFSQLLSLDTEEPRSQP
TEQQLAAHQRTLQTIQKCNIDSIFTESKFLQAESLLQLAQALIWAAGRPQKGNSSPEDEDTAVFCLELLIAITLN
NRDRIVLLWPGVYDHISNIVQSTVMPCALVEKAVFGLLRICQRLLPYKENLADELLRSLQLVLKLDARVADAYCE
QITQEVSRLVKANASHIRSPSGWRTITSLLSITARHPEASEAGFDALLFIVSDGAHLLPANYTLCIDASRQFAES
RVGQAERSLRALDLMAGSVDCLARWAREGKEAAAREEEAIKMSQDIGDMWLRLVQGLRKVCLDQREEVRNQALLS
LQKCLIGVDEIHLPHGLWSQCFDLVIFTMLDDLLEIAQGHSQKDYRNMEGTLILAMKLLSKVFLLLLQDLSQLTN
FCKLWLGVLSRMEKYAKAKVRGKRSEKLQELVPELLKNTLLVMKTKGVLVQRSALGGDGLWELTWLHVNNISPSL
QSEVFPDQDSDHILGQGEKGGVTSSEANPVSSTEKVASDSAGTGG*
**>Momch_LOC111025550**
MGRLKLQTGINAIEEEPEECDVTYTNKTALACMINSEIGAVLAVMRRNVRWGGRYMSGDDQLEHSLIHSLKALRR
QIYSWQHPWHTINPAVYLQPFLDVIRSDETGPPITGVALSSVYKILTLDMIDQNTVNAGDAVHLIVDAVTCCRFE
LTDPASEEMVLMKILQVLLACMKSKASIXLSNQHVCTIVNTCFRIVHQAGSKGELLQRIARHTVHELVRCIFSHL
SRINTTERALVGGNSSSKQEAGTAANDDHVLGSRLLENGNLGHAFDCQSSSNNFQSNSSSGPMATGMEENLLEDG
STKDTVPFNFHLMNEPYGVPCMVEIFRFLCSLLNLVEHMELGARSNTMAFDEDVPLFALGLINSAIELGGPSFRH
HPRLLSLIQDELFRNLMQFGLSTSSLILSMVCSIVLNLYHHLRTELKLQLEAFFSCVILRLAQSRYGASYQQQEV
AMEALVDFCRQKTFMVEMYANLDCDITCSNIFEDLANLLSKSAFPVNSPLSSTHILALDGLIAIIQGMAERIGNG
TGLENSPVNLEEYTPFWMVKCENYSDPNQWVPFVRRKKYIKRRLMIGADHFNRDPKKGLEFLQGTHLLPDKLDPK
SVACFFRYTAGLDKNLVGDFLGNHDEFCVQVLHEFAWTFDFQDMNLDTALRLFLETFRLPGESQKIQRVLEAFSE
RYYEQSPQILVNKDAALLLSYSLIMLNTDQHNVQVKKKMTEEDFIRNSRHINGGNDLPRDFLSELYHSICKNEIR
TTPEQGNSFPEMTPSRWIDLMLKSKKSSPFIVSDSKAYLDRDIFAIMSGPTIAAISVVFDHAEHEEVYKTCIDGF
LGVAKISACHHLEDVLDDLVVSLCKFTTLLNPSSVEEPVLAFGDDTKARMATMTVFTIANRYGDIIRTGWRNILD
CILRLHKLGLLPARVASDAADESELSTDTGHGKPLTSSLSAAHIQSIGTPKRSSGLMGRFSQLLSLDTEEPRSQP
TEQQLAAHQRTLQTIQKCNIDSIFTESKFLQAESLLQLAQALIWAAGRPQKGNSSPEDEDTAVFCLELLIAITLN
NRDRIVLLWPGVYDHISTIVQSTVMPCALVEKAVFGLLRICQRLLPYKENLADELLRSLQLVLKLDARVADAYCE
QITQEVSRLVKANASHIRSPSGWRTITSLLSITARHPEASEAGFDALLFIVSDGAYLLPANYTLCIDASRKFAES
RVGQAERSLRALDLMAGSVDCLARWAREGKEAAREEEAIKMSQDIGDMWLRLVQGLRKVCLDQREEVRNQALLSF
QKCLISVDEIHLPHGLWSQCFDLVIFTMLDDLLEIAQGHSQKDYRNMEGTLILAMKLLSKVFLLLLQDLSQLTNF

CKLWLGVLSRMEKYAKAKVRGKRSEKLQELVPELLKNTLLVMKTKGVLVQRSALGGDGLWELTWLHVNNISPPLQ
SEVFPDQDSDHLLGQGEKGGVTSSEVNSVSWTERVASDSAGTGG*
**>Momch_LOC111024228**
MTKRSDEEDDRDSSKSKRRELGLSCMLNTEVGAILAVIRRPPSELNSPYISPIDDTFDSSIQQSLKSLRALIFHP
QQKWRTIDPSIYISPILDVVQSDDIPAAATGVALSALLKIIKVEIFNEKTPGAKDAINLIVLGITNCKLEKTDLI
TEDAVMMKILQVLAGLMNHRASYLLNDQSVCTIVNTCFNVVQQSANRGDLLQRTARYTMHELIQIIFSRLPEIEV
RDGEDSESDTEDADLGGSLDSGYGIRCVIDVFHFLCSLMNVVEMGEMMDGLASRTADEDVQLFALVLINSAVELS
GDAIGKHPRLLRMVQDDLFHHLIHYGARCNPLVLSMICSTVLNIYHFLRRFVRLQLEAFFVYVALRSASYGNSTQ
IQEVALEGIINFCRQSSFILETYVNYDCDPLRWNLFEDIGKLLCKLSFPAGSPLSSLQIQAFEGLVIMIHNIAEK
LDKLEGGPSRGSSLRVYPAHVSEYRPFWEERSKEDLELEGWLNYVRVRKAQKKKILIAGHHFNRDEKKGLAYLKL
SLLVPDPPDPKAYAFFFRYTYGLDKQFVGEYLGDPDPFHVRVLAEFTGTFEFTGMILDTALRTYLETFRLPGEAQ
KIHRILEAFSERFYDLQSSNTFVNKDAVFVLCYSLIMLNTDQHNPQVKRKMTEDEFIRNNREINGGKDLPRDYLS
ELFQSIANNAIILAPQSGLQLDMNPSKWIELTNRSKIIQPFVLCDFDIRLGRDMFACIAGPSVASLAAFFEHADE
DEMLNECIEGLFSIAKITQCGLEDTLDELLALFSKFTTLLNPYASAEETLFAFSHDLKPKLATLAVFTIANNFGD
SIRGGWRNIVDCLLKLKRLKLLPQSVIEFELASTAPHDLAKSESGVIFPSQDPKFCTQQSSGMASRFSQFLSLDS
MEDSLSLNLNEFEQNLKFIKQCRIGNIFSNSSNLLDDALLNLGRSLIFAAAGKGQKFSTPIEEEETVGFCWDLII
TMSLANVHRFQVFWPSFHEYLQAVIQFPLFSTIPFAEKAVLGLFKVCLKLLSTYQPEKYPEELIFKSINLMWMLD
KEILDTCFEAITQSVSKILIEYPANLQSTIGWKSLLHLLSATGRHPETYDQGVETLIMLMSDGTHITRTNYTFCI
DCAFSYVALKNSPLEKNLKILDLLSDSVNFLIQWYRNYCAESGNSFSVSSNASSSSLEDKGLGSSNFALTLFVKL
GEALRKTSLARREEIRNHAIVSLKKSFFLAEELDFSPTNCINCFNLVIFAMVDDLHEKMLEYSRRDNAEREARSM
EGTLKISMDLLTDVYLLFLKPISESAGFRTFWLGVLRRMDTCMKADLGAYGESSLKDLVPDLLRKLITNMREKEI
LVQKEGDDLWETTYIQIQWIAPSIKEELFPEESF*
**>Benhi_LOC120080467**
MGRLKLQAGINAIEEEPEECDVTYTNKTALACMINSEIGAVLAVMRRNVRWGGRYMSGDDQLEHSLIQSLKSLRK
QIYSWQHPWHTINPAVYLQPFLDVIRSDETGAPITGVALSSVYKILTLDMIDQNTVNAGDSVHLIVDAVTCCRFE
LTDPASEEMVLMKILQVLLACMKSKASIMLSNQHVCTIVNTCFRIVHQAATKGELLQRIARHTVHELVRCIFSHL
SEINTTERALVNGNSSSKQEAGRGANEDYVLGSRLLENGNLGHEFDGQSSSNNFDPNPSSGLMTTGMEENLLEDG
SVKDTVPFDFHLMNEPYGVPCMVEIFRFLCSLLNLVEHMELGARSNTMAFDEDVPLFALGLINSAIELGGPSFRH
HPRLLSLIQDELFRNLMQFGLSTSSLILSMVCSIVLNLYHHLRTELKLQLEAFFSCVILRLAQSRYGASYQQQEV
AMEALVDFCRQKTFMVEMYANLDCDITCSNVFEDLANLLSKSAFPVNCPLSSMHILALDGLIAVIQGMAERIGNG
TGLENTPVNLEEYTPFWMVKCENYSDPVQWVPFVRRKKYIKRRLMIGADHFNRDPKKGLEFLQGTHLLPDKLDPK
SVACFFRYTAGLDKNLVGDFLGNHDEFCVQVLHEFAWTFDFQDMNLDTALRLFLETFRLPGESQKIQRVLEAFSE
RYYEQSPQILVNKDAALLLSYSLIMLNTDQHNVQVKKKMTEEDFIRNSRHINGGNDLPRDFLSELYHSICKNEIR
TTPEQGNGFPEMTPSRWIDLMHKSKKSSPFIVSDSKAYLDRDMFAIMSGPTIAAISVVFDHAEHEEVYQTCIDGF
LAVAKISACHHLEDVLDDLVVSLCKFTTLMNPSSVEEPVLAFGDDTKARMATMTVFTIANRYGDFIRTGWRNILD
CILRLHKLGLLPARVASDAADESELSADTGHGKPLTSSLSAAHIQAIGTPKRSSGLMGRFSQLLSLDSEEPRSQP
TEQQLAAHQRTLQTIQKCNIDSIFTESKFLQAESLLQLAQALIWAAGRPQKGNSSPEDEDTAVFCLELLIAITLN
NRDRIVLLWPGVYDHISNIVQSTVMPCALVEKAVFGLLRICQRLLPYKENLADELLRSLQLVLKLDARVADAYCE
QITQEVSRLVKANASHIRSPSGWRTITSLLSITARHPEASEAGFDALLFIVSDGAHLLPANYTLCIDASRQFAES
RVGQAERSLRALDLMAGSVDCLGRWAREGKEAAREEEAIKMSQDIGDMWLRLVQGLRKVCLDQREEVRNQALLSL
QKCLTGVDEIHLPHDLWLQCFDLVIFTMLDDLLEIAQGHSQKDYRNMEGTLILAMKLLSKVFLLLLQDLSQLTTF
CKLWLGVLSRMEKYAKAKVRGKRSEKLQELVPELLKNNLLVMKTKGVLVQRSALGGDSLWELTWLHVNNISPSLQ
SEVFPDQDSNRVLGQGEKGGVTSSEANSVSTEKVASDNAGTGG*
**>Benhi_LOC120079606**
MEKRSSDDEEEKDSPKSKRRELGLSCMLNTEVGAVLAVIRRPPSDLNSPYISTIDDTYDSSIQQSLKSLRALIFH
PQQKWRTIDPSIYISPILDVIQSDDIPAAATGVALSALLKIIKVEIFNEKTPGVKDAINLIVLGITNCKLEKTDL
VTEDAVMMKILQVLAGLMNHRASYLLNDQSVCTIVNTCFNVVQQSASRGDLLQRTARYTMNELIQIIFSRLPEIE
VRDGEDSESDTEDADLGGSLDSGYGIRCVIDVFHFLCSLLNVVEIVEIGDGGLGMGGGSRTADEDVQLFALVLIN
SAVELSGDAIGKHPKLLRMVQDDLFHHLIHYGASSNPLVLSMICSTVLNIYHFLRRFVRLQLEAFFVYVALKLAS
FGNSTQIQEVALEGIINFCRQSSFILEFYVNYDCDPLRWNLFEEIGKLLCKLSFPTGSPLTTLNIQAFEGLVIMI
HNIAEKLDKHKEESCGGNGSLRVYPAQVTEYRPFWEEKSKEDLELEDWLNYVRVRKAQKKKILIAGHHFNRDEKK
GLAYLKLSQLVSDPPDPKAYAFFFRYTHGLDKQFIGEYLGDPDQFHVKVLAEFTDTFEFTGMILDTALRTYLETF
RLPGEAQKIHRILEAFSERFYDLQSSNTFASKDTVFVLCYSLIMLNTDQHNPQVKKKMTEDEFIRNNREINAGKD
LPRDYLSELFHSISNNAIILSPQSGLQLDMNPSKWVELMNRSKIIQPFMLCDFDPRLGRDMFACIAGPSVASLAA
FFEHADEDEMLNECIEGLFSIAKITQYGLEDTLDELLALFSKFTTLLNPYASAEETLFVFSHDLKPKLATLAVFT
IANNFGDSIRGGWKNIVDCLLKLKRLKLLPPSVIDFEVASTSSNDVARSESGVIFPSQDPKFCSQQSSGMASRFS
QFLSLDSMEDSLTLNLNEYEQNLKFVKQCRIGNIFSSSSNIHDEALFHLGRSLIFAAAGKGQKFSTPVEEEETVG
FCWDLIITMTMANLYRFQVFWPSFHEYLQAVVQFPLFSAIPFAEKAVLGLFKVCLRLLSTYQPEKLPEELIFKSI
NLMWMLDKEILDTCFESITKSVSKIIIEYPANLQSQIGWKSLLHLLSATGRHPETYDQGVETLIMLMSDGTHITR
TNYTFCIDCAFSYVALKNSPLEKNLKILDSLSDSVNFLVQWYRNYCAESGNSFSVASNASSSSLDDKSLGSSNFS
LTLFLKLGEALRKTSLARREEIRNHAIASLKKSFVLAEELDFPPTNCISCFNNIIFAMVDDLHEKMLEYSRRDNA
EREARSMEGTLKISMEVLTDIYLLYLKQISESAGFRTFWLGVLRRMDTCMKADLGSYGESSLKELVPDLLRKIIT
NMREKEILIKKEGEDLWEITYIQIQWIAPGIKDELFPEESL*

**>Datgl_137S20405**
MGRLKLQTSINAIEEEPEECDVSYSNKATLSCMINSEIGAVLAVMRRNVRWGGRYMAGDDQLEHSLIQSLKALRK
QIFSWQHPWHTINPAVYLQPFLDVVRSDETGAPITGVALSSVYKILTLGMIDENTVNVGDAVHLVVDAVTSCRFE
VTDPGSEEMVLMKILQVLLACMKNKASVMLSNQHVCTIVNTCFRIVHQAGTKGELLQRIARHTVHELVRCIFSHL
SDVDNTEHALVNGTSAINHETAGLNNEYAFGSRQPENGNTASEFDGHPSSNNFTSNASTDLGTMMDANALGSVSG
KEAVPFDLHLMTEPYGVPCMVEIFRFLCSLLNVVEHMEIGPRSNTIAFDEDVPLFALGLINSAIELGGPSFCHHP
RLLSLIQDELFRNLMQFGLSMSPLILSTVCGIVLNLYHHLRTELKLQLEAFFACVILRLAQSRYGASYQQQEVAM
EALVDFCRQKTFMVEMYANLDCDITCSNVFEDLANLLSKSAFPVNCPLSAMHILALDGLIAVIRGMAERIGNGIV
GLEHAPVNLDEYTPFWMVKCDNYRDPSYWVPFVRRRKYIKRRLMIGADHFNRDPKKGLEFLQGAHLLPDKLDPQS
VACFFRYTAGLDKNLVGDFLGNHDEFCVQVLHEFAGTFDFQDMNLDTALRLFLETFRLPGESQKIQRVLEAFSER
YYEQSPQILANKDAALLLSYSLIMLNTDQHNVQVKKKMTEEDFIRNSRHINGGGDLPREFLSELYHNICKNEIRT
TPEQGAGFPEMNPSRWIDLMHKSKKTAPFIVSDSRAYLDHDMFAIMSGPTIAAISVVFDHAEHEEVYQTCIDGFL
AVAKISACHHLEDVLDDLVVSLCKFTTLLNPSSVEEPVLAFGDDTKARMATVTVFTIANRYGDFIRSGWRNILDC
ILRLHKLGLLPARVASDAADESELSADSGHGKPITNSLSAAQMQSIGTPRRSSGLMGRFSQLLSLETEEPRSQPT
GQQLAAHQRTLQTIQKCHIDSIFTESKFLQAESLLHLARALIWAAGRPQKGNSSPEDEDTAVFCLELLIAITLNN
RDRIVLLWQGVYEHISNIVQSTVMPGALVEKAVFGLLRICQRLLPYKENLADELLRSLQLVLKLDARVADAYCDQ
ITQEVSRLVKANASHIRSPSGWRTITSLLSITARHPEASDAGFDALSFIMSDGAHLLPANYVLCIDASRQFAESR
VGQAERSLRALDLMAGSVDCLARWMRDGKEAVGEEEVVKISQDIGEMWLRLVQGLRKVCLDQREDVRNHALLSLQ
KCLTGVDDIHLPHGFWLQCFDLVIFTMLDDLLEIAQGHSQKDYRNMEGTLILATKLLSRVFLQLLQDLSQLTTFC
KLWLGVLSRMEKYSKAKVRGKRSEKLFELVPELLKNTLLVMKSKGVLIQRSALGGDSLWELTWLHVNNISPSLQS
EVFPDQDSEQSLHHQGKTGVDLTSNEMNFGHLNETVASEGAGTGG*
**>Datgl_891S35907**
MGHPVPHCGAILVGGEPKEYPPRFSRSTLACMVNSEIGAVLAVMRRNVRWGVRYVAEDDHLEHSLIHSLKELRKQ
IFSWQHDWHNTNPTKYLKPFLDVIQSDETGAPITGVALSSIYKILNLDILDLHTVNVGEAMHLIIDAVTSCRFEV
TDPASEEVVLMKILQVLLACMKSTASAKLSNNHVCNIVNTCFRVVHQASSKGELLQRVARHTMHELVRCIFSHLP
EVDDTEEVIGLDEDRMHGSIQLSNGNISIDNGQPSIACTLNAPAGTVAIEKDDEKVATSNEIGTAQNGDNIMMEP
FGVPCMVEILHFLCSLLNVVEQTEFGPRSSLMAYDEDVPLFALGLINLAIELAGPSIRKHPKLLSIVQDELFRNL
MQFGLSMSPLILSTVCSIVLNLYHLLRAELKVQLEAFFSCVLLRIAQSKHGSSYQQQEVALEALVDLCRQQTFIA
EMYANFDCDITSSNVFEDLANLLSKSAFPVNGPLSAMHILALDGLMSMIQGMAERLGNELPTSEEPSEDLDRYKA
FWMTKCENYDDPNYWVPFVRKMKHLKRRLMIGADHFNRDPKKGLEFLQAVHLLPDKLDPQSVACYFRYTVGLDKN
LVGDFLGSHDEFCIQVLHEFARMFDFQDLNLDTALRLFLGTFRLPGESQKIQRVLEAFAERYYEQSPHILMNKDA
ALLLSYSIIILNTDQHNQQVKRKMTEEDFIRNNRRINGGSDLPREFLSELYHSICENEIQMIPEQSTTFPEITSS
RWINIIHKAKKMTPYIICDSRALLDYDMFAILSGPTIAAMSVVLDQVEQEDLLKTCIDGFLAVAKIAACYHLDDV
LDDLVVSLCKFTTLLTPLSVEEAMTSFCDDTKAKLATTAVFTIANKYGDYIRSGWKNVLDCVLNLQKLGLLPVHL
ISDVVNDMEPSLDSDRGKPATSSLSSSNIPSTATPRKSSGLMGRFSQLLSFDVEEPKSQPTEEELAAHKQSREII
QNCHIDSIFTESKFLQAQSLSQLVKALILAAGRHRKGTNSMEDEDSAVFCLELLIAITLNNRDRIMLIWQGVYEH
IYNIVQSTLMPCVLVEKAVFGLLKICQRLLPYKDNLTDELLRSLQLVLKLDARVADAYFEHITQEVVRLVKANST
HIRSHAGWRTIISLLSITARHREASEVGFEALSFIMSDGAHLMPSNYVLCVDAARQFAESRVGEVERSISAIDLM
LGSVVCLVRWSSEAKNAVGVEASVKVLQDIGEMWLRMVQGLKKVCLDQREEVRNHAILMLQRSMAGVDGIHLPNS
VWLQCFDLVIFTLLDDLLEISQGNSPNDYRNMEGTLFLAMKLMSKVFLQLLQDLWRIPSFCKLWLAVLNLMERYM
KVKVRGRRNERIQELVPELLKNTLLVMKTTGILVPGDAVVGDSFWQLTWLHVKNISPSLQSEVFPDNELETLQMK
HIKAGGIPTSDGTGTVVVQPSQTTA*
**>Datgl_574S11525**
MQKTTEENDNATNCPSCKKHKRKDSAMCKRKELGLSCMLNTEVGAVLAVIRRPPESNSQLLSPPEEILNSSIQQS
LKSLRSLIFNPQQEWRAIDPAVYLSPFIDVIQCDDVPAVATGVALSAILKIIKLQIFDEKTPGAKDAMNSIVRGI
TSCRLEKTNSASEDAVMMKILQVLSGIMKHNASILLNDQSVCTIVNTCFQVVQRSASRGDLLQRSARYTMHELIQ
IIFSRLPEIEVRDDGESSESDSEDGNLDFGYGVRCAVDIFHFLCSLLNVVEVVETDGSMSKTADEDVQLFALVLI
NSAIELSGDGIGKHPKLLRMVQDDLFHHLIYYGTCSNALVLSMICSTVLNIYQFLRRFIRLQLEAFFSFVLLRVA
SFASSIQIQEVALEGIINFFRQQSFIFEIYVNYDCDPLCRNVFEEIGKLLCKLSFRMGNPLTSLQIQAFEGLILL
IHNIAEKIDTEDNSRSFRPSLTKITEYIPFWEDKSKEDFDGWIDCIRLRRAQKKKLLIAGHHFNRDEKKGLQYLK
LSKLVDDPPDPKAFAFFFRYTPGLDKSMIGDYLGDPDEFNIRVLTEFTENFEFTDMILDTALRTYLETFRLPGES
QKIQRILEAFSEKFYSQHSSEIFASKDAVFILCYSLIMLNTDQHNPQVKRKMTEEEFIRNNREINGGKDLPREYL
SELFHSISNNAITLFGHSGITLEINPSRWIELINRSKITQPYVMCDFDRHLGRDMFACIAGPCVAALSAFFEQTD
EDEMLNECIEGLFSIARIAQYGLEDTLDELLASLCKFTTLLNPYASAEETLYAFSNDLKPKMATLVVFTIANNFG
ESIQGGWRNIVDCLLKLKRLKLLPQSVVEFDVAFPSSSDLSETESGVIFSSRDFKFGHRTINRFSQFLAFDSMED
SLSLNLNEFEQNLKVIKQCRIENIFGNSSNLPDDSLLNLGRSLIFAAAGKGQKFSTSVEEEETVGFCWDLIIAMT
LANINRFQSFWPSFHDYLQAVAQFPLFSPIPFAEKAILGLLKVCLKLISAYQPNKIFEELIFKSINLMWMLDKEI
LDTCCEFITQYVSKILIEYPANLQSQLGWKSVLHLLSVTGRHPETYDQGAETLIELMSDGVHVMRTNYAYCIDCA
FSFVALKNSPLEKSLKLLDLMSDSVNLLLQWYKNHYTETGNNFSIASNTSTSSLEGLGSSNFAMNLFIKLGESLR
KTSLARREEIRNHAIFALQKSFALAEELEFTPTNCINCFNLVVFAMVDDLHEKMLEYSRRENSEREMRSMEGTLK
NAMELLTDVYLKFLRQISQSTGFRTFWLGVLRRMDTCMKANLGEYGESKMQELVPDLLRKMITRMREEEILMDRE
GSDDLWDTTCIQIQWIAPSLKDEFFPEFNI*
**>Begfu_1501S08048**

MGRLKLPSGINAIEEEPEEFDVTNSSKNTLACMINSEVGAVLAVMRRNVRWGGRYMSGDDQLEHSLIQSLKALRK
QIFSWQHPWHTINPAVYLQPFLDVIRSDETGAPITGVALSSVYKILTLDMIDQNTSNVGDAVQLVVDAVTSCRFQ
VTDAGSEEMVLMKILQVLLACMKNKASVMLSNQHVCTIVNTCFRIVHQAGNKGELLQRVARHTVQEIVRCIFSHL
SDIENTEHALVNGTSTITQETGFPNNDHALGTRQLENGNTGADFDGPPSSNIISSNISTDLVSSMMEENALGSGG
KEGVPFDLHLMTEPYGVPCMVEIFRFLCSLLNVVEQMEELGPRSNTMALDEDVPLFALGLINAAIELGGSSFSHH
PRLLCLIQDELFRNLMQFGLSMSPLILSTVCSIVLNLYHHLRTELKLQLEAFFSCVILRLAQSRYGSSYQQQEVA
MEALVDFSRQKTFMVEMYANLDCDITCTNVFEDLANLLSKSAFPVNCPLSSMHILALDGLIAVIQGMAERIGNGM
VGLEYHPVRLDEYTPFWMVKCDNYSDPNSWVPFVRCRKYIKRRLMIGADHFNRDPKKGLEFLQGTHLLPDKLDPQ
SVACFFRYTAGLDKNLVGDFLGNHDEFCVQVLHEFAGTFDFQDMNLDTALRLFLETFRLPGESQKIQRVLEAFSE
RYYVQSPQVLANKDAALLLSYSIIMLNTDQHNVQVKKKMTEEDFIKNSRHINGGSDLPREFLSELYHSICKNEIR
TTPEQGIGFPEMNPSRWIDLMHKSKKTAPFIVSDSRAYLDHDMFAIMSGPTIAAISVVFDHAEHEEVYQTCIDGF
LAVAKISACHHLEDVLDDLVVSLCKFTTLLNPSSVEEPVLAFGDDTKARMATVTVFTIANRYGDYIRTGWRNILD
CILRLHKLGLLPARVASDAAEDSELTADTSQGKPITSSLSAAHMQSIGTPRRSSGLMGRFSQLLSLDTEEPRSQP
TEQQLAAHQRTLQTIQKCHIDSIFTESKFLQAESLLHLARALIWAAGRPQKGGSSPEDEDTAVFCLELLIAITLN
NRDRIVLLWQGVYEHISNIVQSTVMPCALVEKAVFGLLRICQRLLPYKENLADELLRSLQLVLKLDARVADAYCE
QITQEVSRLVKANASHIKSLSGWRTISALLSITARHPEASEAGFDALSFIMSDGLHLLPANYVLCVDAARQFAES
RVGQAERSLRALDLMGGSIDCLARWALEGKEAVGEEEAVKISHDIGEMWFRLVQGLRKVCLDQREEVRNHALISL
QKCLTGVCQIHLPHSFWLQCFDLVIFTMLDDLLEIAQAQKDFRNMEGTLILAIKLLSRVFVQLLQDLSQLTTFCK
LWLGVLSRMEKYIKAKVRGKKSERLVEMVPELLKNMLILMKTKGLLVQRSALGGDSLWELTWLHVNTISPSLQSE
VFPNQDSDPSEHKHPETVEDPASDDLKSDPSREMAGSEGVESGG*

**>Begfu_4464S35236**

MGRLKLPSGINAIEEEPEEFDVMNSTKNTLACMINSEVGAVLAVMRRNVRWGGRYMSGDDQLEHSLIQSLKGLRK
QIFSWQHPWHTINPAVYLQPFLDVIRSDETGAPITGVALSSVYKILTLDMIDQNTSNVGDAVHLVVDAVTSCRFQ
VTDAGSEEMVLMKILQVLLACMKNKASVMLSNQHVCTIVNTCFRIVHQAGNKGELLQRVARHTVQEIVRCIFSHL
SDIENTEHALVNGTSTITQETGFPNNDHALGTRQLENGNTGADFDGPPSSNIISSNISTDLVSSMMEENALGSGG
KEGVPFDLHLMTEPYGVPCMVEIFRFLCSLLNVVEQMEELGPRSNTMALDEDVPLFALGLINAAIELGGSSFSHH
PRLLCLIQDELFRNLMQFGLSMSPLILSTVCSIVLNLYHHLRTELKLQLEAFFSCVILRLAQSRYGSSYQQQEVA
MEALVDFSRQKTFMVEMYANLDCDITCTNVFEDLANLLSKSAFPVNCPLSSMHILALDGLIAVIQGMAERIGNGM
VGLEYHPVRLDEYTPFWMVKCDNYSDPNSWVPFVRCRKYIKRRLMIGADHFNRDPKKGLEFLQGTHLLPDKLDPQ
SVACFFRYTAGLDKNLVGDFLGNHDEFCVQVLHEFAGTFDFQDMNLDTALRLFLETFRLPGESQKIQRVLEAFSE
RYYVQSPQILANKDAALLLSYSIIILNTDQHNVQVKRKMTEEGFIQNSRLINGGSDLPREFLSELYHSICKNEIR
TTPEQGIGFPEMNPSRWIDLMHKSKKTAPFIVSDSRAYLDHDMFAIMSGPTIAAISVVFDHAEHEEVYQTCIDGF
LAVAKISACHHLEDVLDDLVVSLCKFTTLLNPASVEEPVLAFGDDTKARMATVTVFTIANRYGDYIRTGWRNILD
CILRLHKLGLLPARVASDAAEDSELSSDTSLGKPITSSLSAAHMQSIGTPRRSSGLMGRFSQLLSLDTEEPRSQP
TEQQLAAHQRTLQTIRKCHIDSIFTESKFLQAESLLHLARALIWASGRPQKGGSSPDDEDTAVFCLELLIAITLN
NRDRIVLLWQGVYEHISNTVQSTVMPCALVEKAVFGLLRICQRLLPYKENLADELLRSLQLVLKLDARVADAYCE
QITQEVSRLVKANASYIKSPSGWRTISALLSITARHPEASEAGFDALSFIMFDGLHLFPANYVLCVDAARQFAES
HVGQAERSLRALDLMGGSIDCLARWAIEGKEAVGEEEAVKISHDIGEMWFRLVQGLRKVCLDQREEVRNHALISL
QKCLTSVFQIHLPHSFWSQCFDLVIFTMLDDLLEIAQAQKDFRNMEGTLILGIKLLSRVFVQLLQDLSQLTNFCK
LWLGVLSRMEKYTKAKVRGKKSERLIEMVPELLKNMLILMKTKGLLVQRSALGGDSLWELTWLHVNTISPLLQSE
IFPDQDSDLSENKQQEAVEDTAAKDLKSEPSREMAGSEGGESGG*

**>Begfu1692S08615**

MENAEDDNNGDSKENNCPSCEEKKKGSGSVASRGKELGLSCMLNTEVGAVLAVIRRPSDLNLQFVSSPEENLDSS
VQQSLKSLRSIIFNPQQEWQSMDPSIYLSPFLDVLQSEDVPAAATGVALSTVLKIINLQVFNEKTPGAKRAMNSI
VSAITSCRLEKTDPVSEDAVMMKILQVLTAIMKHNASILLNDQAICTIVNACFQVVQQSSSRGDLLQRAARYTMH
ELIQIIFSCLPEIEDSTGEFSESDTDDVDIVGNSENGYGIRCAVDIFHFLCSLLNVVTTVEIDGCSANTADEDIQ
LFALVLINSAIELSGDSIGKHPKLLRMAQDDLFHHLIHYGTCSNPLVLSMICSTVLNLYHFLRSFIRLQLEAFFG
FVLLRVATLGGTIQVQEVALEGIINFFRQPSFIVEVYINYDCDPLCRNVFELIGKLLCRLAFPTKSPLNSLQIQA
FEGLIVMIHNIAERIDKDGDSSSFHLQPVETTDYRPFWEEKSKDEVDDWFEFLRVRRAQKRKILIAAHHFNRDVK
KGLEYLKLSHLISDPPDPKAYAFFFRYTPGLDKILIGDYLGDPDEFHISILKEFTFTFDFTGMILDTALRMFLET
FRLPGESQKIQRILESFSEKFYDQQTSEIFATKDSVFILCYSLIMLNTDQHNPQVKKKMTEEEFIRNNRAINGGK
DLPRQYLSELFSSISNNAITLFGNSGPSLDMNPSRWIELINRSKITQPYLMCDFDRLLGRDMFACIAGPTVAAIS
SFFEHAYEDEMLNECIDGLFSIARIAQYGLEDTLDELLAAFSKFTTLLNPYASAEETLFAFSNDMKSKLATLAIF
TIANNYGESIRGGWRNIVDCLLKLKRLKLLPQSVVEFDSASTDSPSTESGVIFPSQDSKLGNRQASGATSRFSHF
LALDNIEDSLSLNLSEFEQNLNVIKQCRIGNIFSNSSNLPDESLLNLGRSLIFAAAGKGQKFSTPIEEEETVGFC
WDLIIAMVLANINRFHTFWPNFHDYLQAVAQFPLFSPVPFAEKAMLGLFKVCLRLLSSFRQEKLSEELIFKSINL
MWMLDKEILDTCCEFLTKSVSRILIEYPANLQTQLGWKSVLHLLSVTGRQPETYDQGIETLITLMSDGIHVSRTN
YMYCVDCAFSFVALKNSPLDKNLKLLDLMSDSVNLLTQWSKNYYSDPGNGYSMTSNTSNSSVEDSKTNSTNFGMN
LFVKLGEALRKTSLARREEMRNHAISALHKSFTLAEELEFPPNSCINCFNLVIFAMVDDLHEKMLEYSRRENSEK
EMRSMENTLKIAMELLANVYLQFLKKMSPSSGFRTFWLGVLRRMDTCMKADLGEYGESKMQELVPDLLRKMITKM
REEEILVQQEDDDLWEITYIQIQWIAPSLKEEMFPEESL*

**>CitlaGNOM_chr7**

MGRLKLQAGINAIEEEPEECDVTYTNKTALACMINSEIGAVLAVMRRNVRWGGRYMSGDDQLEHSLIQSLKSLRK
QIYSWQHPWHTINPAVYLQPFLDVIRSDETGAPITGVALSSVYKILTLDMIDQNTVNAGDSVHLIVDAVTCCRFE
LTDPASEEMVLMKILQVLLACMKSKASIMLSNQHVCTIVNTCFRIVHQAATKGELLQRIARHTVHELVRCIFSHL
SEINTTERALVNGNSSSKQEAGRGANDDYVLGSRLLENGNLGHEFDGQSSTNNFDSNPSSGLMATGMEENLLEDG
SVKDAVPFDFHLMNEPYGVPCMVEIFRFLCSLLNLVEHMELGARSNTMAFDEDVPLFALGLINSAIELGGPSFRH
HPRLLSLIQDELFRNLMQFGLSTSSLILSMVCSIVLNLYHHLRTELKLQLEAFFSCVILRLAQSRYGASYQQQEV
AMEALVDFCRQKTFMVEMYANLDCDITCSNVFEDLANLLSKSAFPVNCPLSSMHILALDGLIAVIQGMAERIGNG
TGLENTPVNLEEYTPFWMVKCENYSDPVQWVPFVRRKKYIKRRLMIGADHFNRDPKKGLEFLQGTHLLPDKLDPK
SVACFFRYTAGLDKNLVGDFLGNHDEFCVQVLHEFAWTFDFQDMNLDTALRLFLETFRLPGESQKIQRVLEAFSE
RYYEQSPQILVNKDAALLLSYSLIMLNTDQHNVQVKKKMTEEDFIRNSRHINGGNDLPRDFLSELYHSICKNEIR
TTPEQGNGFPEMTPSRWIDLMHKSKKSSPFIVSDSKAYLDRDMFAIMSGPTIAAISVVFDHAEHEEVYQTCIDGF
LAVAKISACHHledvlddlvvSLCKFTTLMNPSSVEEPVLAFGDDTKARMATMTVFTIANRYGDFIRTGWRNILD
CILRLHKLGLLPARVasdaadeselsadTGHGKPLTSSLSAAHIQSIGTPKRSSGLMGRFSQLLSLDSEEPRSQP
TEQQLAAHQRTLQTIQKCNIDSIFTESKFlqaesllqlaqaliwaaGRPQKGNSSPEDEDTAVFCLELLIAITLN
NRDRIVLLWPGVYDHISNIVQSTVMPCALVEKAVFGLLRICQRLLPYKENLADELLRSLQLVLKLDARVADAYCE
QITQEVSRLVKANASHIRSPSGWRTITSLLSITARHPEASEAGFDALLFIVSDGAHLLPANYTLCIDASRQFAES
RVGQAERSLRALDLMSGSVDCLGrwaregkeaareeeaIKMSQDIGDMWLRLVQGLRKVCLDQREEVRNQALLSL
QKCLTGVDEIQLSHDLWLQCFDLVIFTMLDDLLEIAQGHSQKDYRNMEGTLILAMkllskvflllqdlsqlttf
CKLWLGVLSRMEKYAKAKVRGKRSEKLQELVPELLKNNLLVMKTKGVLVQRSALGGDSLWELTWLHVNNISPSLQ
SEVFPDQDSNRALGQGEKGGLTSSEANSVSSTEKVASDSAGTGG*

**>CitlaGNL2_chr 11**

MDKRSSDDEEEKDSPKSKRRELGLSCMLNTEVGAVLAVIRRPPPDLNSPYISTIDDTYDSSIQQSLKSLRALIFQ
PHQKWRTIDPSIYISPILDVIQSDDIPAAATGVALSALLKIIKVEIFNEKTPGVKDAINLIVLGITNCKLEKTDL
VTEDAVMMKILQVLAGLMNHRASYLLNDQSVCTIVNTCFNVVQQSASRGDLLQRTARYTMNELIQIIFSRLPEIE
VRDGEDSESDTEDADLGGSLDSGYGIRCVIDVFHFLCSLLNVVEMVEMGDGLGSRTADEDVQLFALVLINSAVEL
SGDAIGKHPKLLRMVQDDLFHHLIHYGASSNPLVLSMICSTVLNIYHFLRRFVRLQLEAFFVYVALKLASFGNST
QIQEVALEGVINFCRQSSFILEFYVNYDCDPLRWNLFEEIGKLLCKLSFPTGSPLTTLNIQAFEGLVIVIHNIAE
KLDKDKEESRGGNGNLRVYPAQVNEYKPFweekskedleledwLQHVRVRKAQKKKILIAGHHFNRDEKKGLAYL
KLSQLVSDPPDPKAYASFFRYTHGLDKQFIGEYLGDPDQFHVKVLAEFTETFEFTGMILDTALRTYLETFRLPGE
AQKIHRILEAFSERFYELQSSNTFASKDTVFVLCYSLIMLNTDQHNPQVKKKMTEDEFIRNNREINAGKDLPRDY
LSELFHSISNNAIILSPQSGLQLDMNPSKWVELMNRSKIIQPFMLCDFDPRLGRDMFACIAGPSVASVAAFFEHA
DEDEMLNECIEGLFSIAKITQYGLEDTLDELLALFSKFTTLLNPYASAEETLFVFSHDLKPKLATLAVFTIANNF
GDSIRGGWRSIVDCllklkrlkllPQSVIDFEVAATSSNDVARSESGVIFPSQDPKFSTQQSSGMASRFSQFLSL
DSMEDSLTLNLNEYEQNLKFVKQCRIGNIFGNSSNILDEALLNLGRSLIFAAAGKGQKFSTPVEEEETVGFCWDL
IITMTMANLYRFQVFWPSFHEYLQTVVQFPLFSAIPFAEKAVLGLFKVCLRLLSTYQPEKLPEELIFKSINLMWM
LDKEILDTCFESITQSISKIIIEYPANLQSQIGWKSLLHLLSATGRHPETYDQGVETLIMLMSDGTHITRTNYTF
CIDCAFSYVALKNSPLDKNLKILDLLSDSVNFLVQWYRNYCAEsgnsfsvasnassssIDDKGLGSNFSLTLFLK
LGEALRKTSLARREEIRNHAIASLKKSFVLAEELDFPPTncincfnniifAMVDDLHEKMLEYSRRDNAEREARS
MEGTLKISMELLTDIYLLYLKQISESAGFRTFWLGVLRRMDTCMKADLGSYGESSLKELVPDLLRKIITNMREKE
ILMKKEGEDLWEITYIQIQWIAPAIKDELFPEESF*

**>Kalfe_61s0170**

MGRMKMQSGIKSIEEDEPTDSETASSSKEGLSCMINSEIGAVLAVMRRNARWGGRYMTGDDQLEHSLIQSLKALR
RQIFSWQNQWATVDPALYLQPFLDVIRSDETGAPITGVALSSVYKILTLDVLDMNTVNVEDAMHFVVDAVTSCRF
EVTDPASEEVVLMKILQVLLACMKSKASVMLSNQHVCTIVNTCFRVVHQAGTKGELLQRFARHTMHELVRCIFAH
LSEVFNMDWALTNGSSAKQEVGSLHDDNSSGTKQLENGNGSSEYDGAPSSGSYPSNASSGLIAAGMEENVGTGEG
KDSVSYDLHHMTEPYGVPCMVEIFHFLCSLLNVSEHPGLDPRSNTIVFDEDVPLFALGLINSAVELGGSSIGKHP
RLLNLIQDELFRNLMQFGLSMSPLILSMVCSIVLNLYQHLRTELKLQFEAFFSCVILRLAQSRYGASYQQQEVAV
EALVDFCRQKSFMVEMYANLDCDITCSNVFEDLANLLSKSAFPVNSPLSSMHILALDGLIAIIQGMAERVSNGSL
GLEHSPVTLEEYTPFWMVKCDNYSDPTHWVPFVRRRKYIKRRLMIGADHFNRDPKKGLEFLQGTHLLPDKLDPQS
VACFFRYTAGLDKNLVGDFLGNHDEFCVQVLHEFAGTFDFQDMNLDTALRLFLETFRLPGESQKIQRVLEAFSER
YYEQSPHILANKDAALLLSYSLIMLNTDQHNVQVKKKMTEEDFIRNNRHINGGSDLPREFLSELYYSICKNEIRT
IPDAGSGFPEMNPSRWIDLMHKSKKTAPFIVADDRALLDHDMFAIMSGPTIAAISVVFDHAEYEDVFQTCIDGFL
AIAKISACHHLEDVLDDLVVSLCKFTTLLNPISVEEPVLAFGDDTKARMATVAVFTIANRYGDYIRTGWRNILDC
ILRLHKLGLLPARVASDAADDSELSSEPGQGKPLSSSLSSAHLQSLGTPRRSSGLMGRFSQLLSLDTEEPRSQPT
EQQLAAHQRTLQTIQKCHIDSIFTESKFLQANSLLQLARALIWAAGRPQKGTGSPEDEDTAVFCLELLIAITLNN
RDRIVLLWQGVYEHIVNIVQVTVMPCALIEKAVFGLLRICQRLLPYKESLADELLKSLQLVLKLDARVADAYCEQ
ITQEVSRLVKANATHIRSQMGWRIITALLSITSRHPEASDSGFDALVFIMSDGAHLSPANFVLCVDAARQFAESR
VGQAERSLRALDLMASSMVCLLQWAAEAKDVATGEDAAKLSQDIGDIWLRLSQGLRKVCFDQREEVRNHALLSLQ
RCLTDADDLHFAHTLWLQCFDQVIFTMLDDLLELAQGHSQKDFRNMEGTLLLAMKLLSKVFLQLLLDLPQQTTFC
KLWLGILSRMEKYTKVKVRGKRSEKLQELVTELLKNNLVTMKTKGILAPRSALGGDSLWELTWLHVHNIAPSLQS
EVFPEQGSDPSQQTHPAESDSVPARDTAASEG*

**>Kalfe_43s0155**

MGLMKMPSSIEEEQTEDAETVSSSDAGLACMINSEIGAVLAVMRRNARWGGRYMSGDDQLEHSLIQSLKALRRRI

FSWKNQWATVNPALYLQPFLDVIRSDETGAAITGVALSSVYKIMTLDVLDINTVNVVDDMHLVVDAVTSCRFEVT
DPASEEVVLMKILQVLLACMKSKASVMLSNQHVCTIVNTCFRVVHQAGTKGELLQRFGRHTMHELVRCIFAHLSE
VLNMDWALTNGSIGKQEDGSLDDDNSSGTKQLENGNISSEYDGAPSSGSYPSNASSGLIAAGMEENVGTWEGKDS
VFYDSHHLTEPYGVPCMVEIFHFLCSLLNVSEHPGLDARSNTIVFDEDVPLFALGLINSAVELGGPSIGKHPRLL
NLIQDELFRNLMQFGLSMSPLILSMVCSIVLNLYQHLRTELKLQLEAFFSCVILRLAQSRYGASYQQQEVAVEAL
VDFCRQKSFMVEMYANLDCDITCTNVFEDLANLLSKSAFPVNSPLSSMHILALDGLIAIIQGMAERVSNGSLGLE
NSPVTLEEYTPFWMVKCDNYRDPTHWVPFVRRRKYIKRRLMIGADHFNRDPKKGLEFLQGTHLLPDKLDPQSVAC
FFRYTAGLDKNLVGDFLGNHDEFCVQVLHEFAGTFDFQDMNLDTALRLFLETFRLPGESQKIQRVLEAFSERYYE
QSPHILANKDAALLLSYSLIMLNTDQHNVQVKKKMTEEDFIRNNRHINGGGDLPREFLSELYYSICKNEIRTIPD
AGSGFPEMNPSRWIDLMHKSKKTAPFIVADDRALLDHDMFAIMSGPTIAAISVVFDHAENEDVFQTCIDGFLAIA
KISACHHLEDVLDDLVVSLCKFTTLLNPISVEEPVLVFGDDAKARMATATVFTIANRYGDYIRTGWRNILDCILR
LHKLGLLPARVASDAADDSGLSSEPGQGKPLSSSLSSAHLQSLGTPRRSSGLMGRFSQLLSLDTEEPRSQPTEQQ
LAAHQRTLQTIQKCHIDSIFTESKFLHANSLLQLARALIWAAGRPQKGTGSPEEEDTAVFCLELLIAITLNNRDR
IILLWQGVYEHIVNIVQTVMPCALIEKAVFGLLKICQRLLPYKENLADELLKSLQLVLKLDSRVADAYCEQISQ
EVCRLVKANAAHIRSQMGWRTITALLSVTARHLEASEPGFDALVFIMSGGAHLSPVNFVLCADAARLFAESRIGQ
AERSVRALDLMAGSMVWLTQWALKAKEAATVEDAVKLSQDIGEMWARLSQGLRKVCFDQREEVRNHALLCLQRCL
IDADDLHFTHALWVQCFDQVIFTMLDDLLELAQGHSQKDFRNMEVTLVLAMKLLSKVFLQLLHDLPQSTTTFCKL
WLGILSRMEKYTKVKVRGKRSEKLQELVPEALKNILVAMKTKGILAQRNALGGDSLWELTWLHVNNIASDLQSEV
FPEPCSDPSQQTLPAESDRVASEG*

**>Kalfe_1222s0075**

MGLLTSQSEMDTLHQESKECSSQGAFACIINTEISAVLAVMRRNVRWGVRYLTDDNQMEHPLIQSLKNLRKEIFS
WQNQWNSTNPAAYLQPFLDVIQSDETGAPITGVALSSIYKILTLDILHIDTVNVEDAMALIVDAVTSSRFEVTDP
ASEEVVLMKILQVLMACMKSKASKKLSNQHVCNIVNACFRVVHQASSKGELLQRIARHTMHEMVRCIFSCLPDIA
IEEHTHRKEGQLSTSDEAGSQNIGDVTNNREQGGGDVSNASTGKAEPIVDNGVQMEDASNEEQLMTKPFGIPCMV
EIFHFLCSLLNVIENIESGPRSNPIAYEEDVPLFSLGLINSAIELGGAFFARHPKLLLLIQDELFQNLMQFGLSM
SPLILSTVCSIVLNLYHHMRTELKLQLEAFFSFVLLRLAHNKHGYSFQQQEVAMEAIVDLCRQPSFVTEIYANYD
CDITCNNVFEDISNLLSKSAFPVNSPLSVVNLLALDGLIAIVQGMANRVGDENPSAEQVLVNSEGYESFWRVKCE
SYHDLDLWIPCVRKMKYIKRRLMVGADHFNRDSKKGLQFLQEMHLLPEKLDPQSVAYFFRYTISLDKNVIGDFLG
NHDDFCVQVLQEFAKTFDFEGMSLDGALRLFLGTFRLPGESQKIQRVLEAFAERYYEQCPHILSNKDAALLLSYS
IIMLNTDQHNAQVKRKMTEEDFIRNNRCINAGKDLPRDFLSELYHSISENEIQMLPDQAGSGLPVMTSSRWVSLL
QNSKKCSQFILCDSTARLDYDMFTILSGPSIAAISVVIDNMEHEEVIQTCINGFMNIAKISTCYHLNDILDDLVV
SLCKFTTLLDPSSAEESILALVEDIKARMATTSVFAIANKYGDCVRTGWKNIVDCVLYLHKLRLLRLADGTADNG
KPSSESEIAPAVRSQSAPPVLTSSSSRKSSGLMGRFSQLFSFEAEEPIQQSTLEELESHQQALQIIQSCQIGSIF
MESKFLQADSLLQLVQALILAAGRLQKGPYSPEDEDTAVFCLGLLIEITVNNRDRIMLLWQSVYEHIANVVQSTV
MPCTLVEKAVFGLLQICQRLLPYKENLTDELLRSLRLVLKLDARVADAYCEHITQEVMHLVKGNAPQIRSHVGWR
TIISLLSITAQHPEASEVGFETLVYIMFDAAHLSPANYVLCVDAARQFAESRVGKIDRSVRALDLMADSFVCLIR
WSNEAKQAVGDEAAAKVYQDIGEMWMRLVQGLKRSGLDRREQVRNHAILVLQRCLIGAEGIYLPQELWLQCFDTV
IFALLDDLLNVGQGSSVKDYRNMEGTFVLAVKLLSKVFLQLLQDDSAQKYLTSLWEGYLNRMESFMKAKFRGKWS
ENVHEVVPELLKNTLLIMKAKGILVASDAAAAGKDSSLWQLTWSHVKQIAPSLQTEIFSSDESEQMQRGNEVKTG
SMTPSDGAVLVPTTNEIKA*

**>Kalfe_96s0081**

MGLVTTQSKMDSLHHKSRDCASHGAYACIINTEISAVLAVMRRNVRWGVRYLGDDIQMEHPLIHSLKKLRKEIFS
WQNQWESTNPAAYLQPFLDVIQSDETGAPITGVALSSVYKILTLDILNEHTANVEDAMALIVDAVTSCRFEVTDP
ASEEVVLMKILQVLMACMKSKASKKLNNQHVCNILNACFRVVHQASSKGELLQRIARHTLHEMVRCIFSRLPDIA
TEEHTNHTEGLLSTSDEPVSSNSLHFSDEQNTEDVKNYREQSGGDASNASTGEAEPKVDNGSQMEDVSNEEQLMT
KPFGTPCMVEIFHFLCSLLNVIENIESGSRSNPIAYEEDVPLFSLGLINSAIELGGDSFARHPKLLALIQDELFQ
NLMQFGLSMSPLVLSTVCSIVLNLYYHMRTELKMQLEAFFSFVLLRVAHNKHGYSYQQQEVAMEAIVDLCRRPSF
VTEIYANYDCHITCNNVFEDIANLLSKSAFPVNSPLSAVNLLALDGLIAILQGMANRGGDENPFADQILLAKGYK
SFWTEICESYHDLDLWIPSVRKMKYIKRRLMVGADHFNRNSKKGLQFLQELHLLPEKLDPQSVAYFFRCTISLDK
NVMGDFLGSHDDFCVQVLEEFAKTFDFEGMSLDGALRLFLGTFRLPGESQKIQRVLEAFAERYYEQCPHILSNKD
AALVLSYSIILLNTDQHNLQVKRKMTEEDFIRNNRCINAGKDLPREFLSELYQSISENEIQMLPDQGAELPVMTS
SRWISLLHSSKKCSPFILCDSTARLDYDMFTVLAGPFIAAISVVIDNTEHEEVIQTCIDGFMNIAKISACYHLND
ILDNLVVSLCKFTTLLDPSSVEESIIALVEDNKARMATTSVFTIANKYGDCIRAGWKNIVDCVLYLHKLGLLRLA
DGTADDGKPSPESEKSALAVSSQSAPPALTSSTSRKSSGLIGRFSQLFSFDPEEPIPRTTLEELESQQQALQTIQ
SCQIGSIFMESKFLQADSLLQLVQALILAAGRLQKGFYSREDEDTAVFCLGLLIETTVNNRDRIMLLWQYVYEHV
ANVVQSTVMPCTLVEKAVFGLLQICQRLLPYKENLTDELLKSLRLVLKLDARVADVYCEHITQEVMHLVKANAPQ
IRSHVGWRTIISLLSITAQHPEASEAGFDTLFYIMSDAAHLSPANYILCVDAARQFAESRVGMIDRSVRALDLMA
DSFVCLIRWSSEAKQAVGDDATAKVYQDIGEMWMRLVQGLKRLGLDQREQVRNYGILLLQRCLIGVEGISLPQEL
WLQCFDTVIFALLDDLLKLGQGSFLKDYRNMEGTFVLAIKLLSKVFLQLLQNDPAPKYLINLWEGYLDRMESYMK
AKFRGRRSDNVHQVVPELLKNTLLIMKAKGILVPNDAAANGKDDDLWRLTWSHVNNISPSLQLAIFSSQELEQLQ
SSDDIRTVSVPPPSDGAVLVPANKITRPEILVQRKLSHSLGFVQTLTHWENL*

**>Kalfe_31s0117**

MDEEEDAGAYNGTHPRRFRARDRGLSCMLNTEISAILASIRRVPPEHQGSQLLMSQSDDRYDTHAVQSIKSLRPL

IFNPPQEWRTVDPSVYLVPFLDVVQSDDITATATLVALQSVLKILRLHFFDEKTPGARDAINAAVIAVTGCRLEK
TDHVTEDSVLIKVLQVLVAVMDHPASVFLTDNAVCTAVNTCFHVVQTTHRGGSMNKNARNIMQEMIQIVFSRLPD
IESADSDTEDADIGGSYGIGYGVRCTVDIFHLMCNLLNVAEVGVENTSDDMHTENIAPDEDMQLFGLVMINYAIE
LGGEAIRYHPKLVRMVQDDLCHHLVHYGMRSKPLVLSVISSTILNIYHFLRRSIRLQLEAFFFFVLFRVARMGSL
VLQEVAVEAIINFVRQPTFIFEAYANYDCDPIRCNVFEEIGKLLCRQATPHVPPTAPQTTVAAAHVNFTSLHTQA
LEGVLLLIHTIADSISKDDPPSSFEPYPVQITDFSPYWEEKAKNDDEMDEWVEHVRIRKAQKKKLRIAANIFNRE
EKKGLDYLKAAGLISEPPDPKAYAYFFRYMPGLDKNMIGDYLGDPGAFHLQVLKDFTDTFHFPGVILDTALRTYL
ETFRLPGEAQKIQRILEAFSDRFYDQQSLELFVSKDAVYILCYSLIMLNTDQHNPQVKKKMTEEEFIRNNRGING
GQDLPREYLSQLFRSISSHPISIFTQTGLPTEMNPSKWVQLICQSKIAEPFVPCEIFDGKLSRDMFAAISGPSVA
LLSVIFVRTNEHEILNDCIEGFFSVVKVAQHGLYDTMDEVVACFYKATRLLSPQEIDLASVHDLRKRLATLALFT
VANDYGHSIRGGWRDIVEYLVFLKRQKLIPQILDEPDAASAELESHPRFGSSTLHNPYERTLGNRASSSMIGRFS
HFLSMETVEDSLSLDGEYEQIQKMIQQCRIGNIFKNSSSLPEESLLCLGRALIFAAGGKAHKFTTSVEEEDTVGF
CWELLLSLTSVNIDRFPSFWPPFQDFLLQVAQFPLFSPMPFAEKAIVSLMKICLKLLSSYREEKNSEETIFKTIN
LMWKLDKEILDICGEVITQTVSKILVEYPGNLQTQLGWKSVLHLLSVTGRHPETYAQGVEALITLMSEKTYITKY
NYAYCIDCAFGFVALKNSPIEKNVKILDLMSNSVNLLVQWNKNYSDPGSSFSMPSNASSSSFDDNPKALGGSSLA
MNLFVKLGEALRKTSLARREEVRNQAVAALHKSFILAQELEMQSTNCIGSFNLVIFAMVDDLHEKMLEYSRRENA
EREMRSMEGTLKIAMELLTDVFLQYVKQIAENPGFRTFWLGVLRRMDTCMKADLGAYGEASLQEMVPGLLRKIIT
TLKENEILVKVEGDDLWEITYIQIQWIAPALKEELFPDEEL*
**>SedalGNOM_ctg13**
MGRMKMQSGIKSIEEEPVDLETGSYSKAGLACMINSEIGAVLAVMRRNARWGGRYMSGDDQLEHSLIQALKALRR
QIFSWENQWQTVDPALYLQPFLDVIRSDETGAPITGVALSSVFKILTLDVLDMNTVNVEDAMHLVVDAVTSCRFE
VTDPASEEVVLMKILQVLLACMKSKTSVVLSNQHVCTIVNTCFRVVHQAGTKGELLQRFARHTMHELVRCIFAHL
SEVGNMDWALTNGSNGKQEVGSLDDDKSPATRQLENGNGSTEYDGPLSSGGFPSNASSGLIAAGMEENVGAGEGK
DSVSYDLRHMTEPYGVPCMVEIFHFLCSLLNVSEHPGLDPRSNTIVFDEDVPLFALGLINSAVELGGSSIGKHPR
LLSMIQDELFRNLMQFGLSMSPLILSMVCSIVLNLYQHLRTELKLQLEAFFSCVILRLAQSRYGASYQQQEVAVE
ALVDFCRQKSFMVEMYANLDCDITCSNVFEDLANLLSKSAFPVNSPLSSMHILALDGLIAIIQGMAERVGNGSIG
IEHSPVTLEEYTPFWMVKCDNYSDPSQWTPFVRRRKYIKRRLMIGADHFNRDPKKGLEFLQGTHLLPDKLDPQSV
ACFFRYTSGLDKNLVGDFLGNHDEFCVQVLHEFAGTFDFQDMSLDTALRLFLETFRLPGESQKIQRVLEAFSERY
YEQSPHILANKDAALLLSYSLIMLNTDQHNVQVKKKMTEEDFIRNNRHINGGSDLPREFLSELYYSICKNEIRMI
PDAGAVFPEMNPSRWIDLMNKSKKTAPFIVANDRALLDHDMFAIMSGPTIAAISVVFDHAEHEDVFQTCIDGFLA
IAKISACHHLEDVLDDLVVSLCKFTTLLNPISVEEPVQAFGDDTKARMATVTVFTIANRYGDFIRTGWRNILDCI
LKLHKLGLLPARVTSDASDDTELPLEPGQGKPLTGSLSSAHLQALGTPRRSSGLMGRFSQLLSLDTEEPRSQPTE
QQMAAHHRTMQTIQKCHIDSIFTDSKFLQANSLLQLARALIWAAGRPQKGTGSLEDEDTAVFCLELLIAITLNNR
DRIALLWQGVYEHIVNIVQVTVAPCLLIQKAVFGLLRICQRLLPYKENLADELLKSLGLILKLDARVADEYCAQI
TQEVYRLVKANANQIRSQMGWRTITALLSITARHPEASEPGFDALVFIMSDGTHLSPVNFAVCVDAARQFAESRV
GQAERSVRALDLMASSMVSLSRWTLETKETTSAEDALKISQDLGEMWLRLSQELRKVCFDQREEVRNHALLSLQR
CLTAADEIHFAHTLWLQCFDEIIFTMLDNMLELVLGHSQKDFRNMEGTLILAMKLLSKVFLQLLDELPHLTTFCK
LWLGILSRMEKYTKVKVRGKKSEKLQELIPELLKNILVTMKTKGILAQRNALGGDSLWELTWLHVNNIAPYLQSE
VFPNASSQGHPPADSVPISSNDTPAPLEG*
**>SedalGNL1a_sc317**
MGLLPPPPEMVNIPQESRECSIHGAAACVINTEISAVLAVMRRNVRWGVRYLAVDNQLEHPLIQSLKNLRKEIFS
WKNQWDTTNPAAYLQPFLDVIQSDETGAPITGVALSSVYKILTLDILNIYTGKVEDAMALIVDAVTSCRFEVTDP
ASEEVVLMKILQVLLACMKTKASKKLTNQHVCNIVNACFRVVHQASSKGELLQRIARHTLHEMVRCIFSCLPDIA
DHEHTHDTESQFSTSDEVSFSDRLHGSGSKQIENVNGSSASMGEAEPNADNGYLMENASNEETLMSEPFGIPCMF
DIFHFLCSLLNVIENIESGPRSNPIAYEEDVPLFSLGLINSAIELGGASFARHPKLLMLIQDELFRNLMQFGLSM
SPLILSTVCSIVLNLYQHMRTELKLQLEAFFSFVLLRLAHNNHGYSYQQQEVAMEAIVDLCRQTSFVTEMYVNYD
CDITFNNVFEDIVNLLSKNAFPVNSPLSAVNLLAMDGLTAIIQGITHRIGEENPSVEQTLLNCEGYESFWNVKCE
RYDDIDILIPFVRKMKYIKRRLMVGADHFNRDSKKGLQFLQEMRLLPEKLDSQSVACFRYTISLDKNVIGDFLGN
HDEFCVQVLQEFAKTFDFEGMSLDGALRLFLGTFRLPGESQKIQRVLEAFAERYYDQCPHILSNKDAALLLSYSI
IMLNTDQHNAQVKRKMTEEDFIRNNRCINAGKDLPRDFLSELYHSISENEIQMLPDQGAGLPVMTSSRWASLLQN
SKKCSPFIICNSTARLDYDMFTVLSGPSIAAISVVIDNMEQEEVIQTCIDGFLNIAKISACYHlnnvlddlvvSL
CKFTNLLDPSSTEESILALVEDHKVRMATSSVFIIANNYGDCIRTGWKNIVDCVLYLHKLGLLRLPVGTADNGKQ
LSEsekspaplalsssssrksssGLMGRFSQLFSFEAEEPILQSTMEELESYQLAVQTIQSCQIGNIFLESKFLQ
ADSlvqlvqalilaaGRLQKGHYSPEDEDTAVFCLELLIEITVNNRDRIMLLWQSVYEHIANVVKSTVMPCTLVE
KAVFGLLQICQRLLPYKENLTDEllkslklilklDARVADAYCEHITQEVMHLVKANARKIRSHVGWRTIISLLS
ITAQHPEASEVGFETLYIMADSAYLSPANYVLCVDAARQLAESRVGNMDRSIRAADLMADSLVCLKRWSNeakqa
veeeaaakvYQDIGEMWMRLVQGLKRLGLDRREQMRNYAILVLQRSLIAVEGIYLSHELWLQCFDTVIFSLLDDL
VNVIQGGSHRDYRNIEGTLVLAVKLLSKAFLQLLQDDLSPKDLTTLWEGYLNRMDSCMKTKFRGKRSENIHEVVP
ELLKNTLLVMKSKGILVPNDAAAGKDNLWQLTWLHVKKISPALQSEIFSSQELEQMKSDEMKSGSMPPSESAALV
QADEIKA*
**>SedalGNL1b_sc25**
ASNVLPTPQLKMNTFHQGSRKCTSQGTFARIINSEISAILAVMRRNVRWGVRYLAVDNEREHPLIQSLKKLRKEI
FSWKVQWDMVNPAVYLQPFLDVIKSDETGAPITVVALSSVYKLLTLDLFDSDFEQVEDAMALIVDVVTNYQFDVA

DPASEEVVMMKILQVLLACMKGTASKKLNNQHVCNIVDMCFRVVHQASSESELLQRFARHTLHEVVKCIFSCLPE
IASEEHGQHSEDQIMMSDEPGSSSKLHVSDSQKIENVRNYKEQPLHGSNDSTGKAEPEVDNGAQTKYSSNEEQLM
KKPFGIPCMIETFHFLCSLLNVTENLESGPRPNPIAYEEDVPLFALGLINSAVELGGGYFTQHPKLLKLIQDDLF
QNLVRFGQSISPLILSTVCSIVLNLYHQMRSELKLQLEAFFSFVLLRLAHDRHGYSYQQQEVAMEAIVDLCRQPS
FATEIYVNYDCDITCNNIFEDIVNLLSKSAFPVNNPLRTVNLLALDGLVAILQCMADTVGYDNPFPEQTVVSSEG
YESFWKLKCENYNDLYLWMPFVQKMKNIKRILMMGADHFNRDSKKGLQFLQEMCLLPKKLDPLSVACFFRHTVGL
DKTAIGDFLGNHDEFSVQVLQEFARTFDFEGMGLDGALRLFLGTFRLPGESQKIQRVLEAFAERYYEQCPHVLSN
KDAALVLSYSIILLNTDQHNAQVKRKMTEEDFirnnrninagnnLPREFLSELYHSISKNEIQMLPDQGTGLPAM
TSSRWVVLMQNSKKCNPYIVCESIARLNHDMFTVLSGPSIAAISVVIDNMEQEEVIKTCIEEGFMNIASIAARYH
FDGILDDLVASLCKFTTLLDPISAEESVLALLEDYKARLATSAVFSIANKYGDCIQTGWRNIVDCVLYLHKIGLF
RLADGAAANEMTLAESGKAAHAITLQAATPAVTSTSSQKASGLMGRFSQLFSFEAEDPIQKSTLEESESHQLALQ
IVQSWQIGSIFIESKFLQADSLLQLVKALILTAERFHKGPYSPEDEASAIFCLGVLIEVTVNNRDRIMLIWQSVY
EHIASVVQSIVMPCALVEKAVFGLLQICQRLLSYKENLTDELLRSLQLVLKLDARVADVYCEQITQEVMHLVKAN
ATRIKSHVGWRTIVSLLSITAQHPEASEVGFETLSYIMSGAAHISPVNYVLCVDAARQFVESRVGMIDRSVRAID
LMVDSFSCLLQWSNEAKQTIVVDGITTKVYQDIGDMWMRLVQGLKKSSLDQREQVRNYAILALQRCLIGAEGIHL
PPELWLQCFDTVIFALLDDLLKVSQGSLFKDYRNLEGTFVLAIKLLSKVFLQLLQYESAHKYLANLWEGYLNRMG
NYMEAKFRGRRCKNVQEVVPELLKNTLLVMKAKEILVPKSATSNDSLWEMTWSHIKDIAPSLQSEIFPNQELEQI
QGNEIRTGSTPPSEGVPTNEINA*

**>SedalGNL2_ctg425**

MDEEEDPNSYGHHQRRFRPRGLSCMLNTEVSAILASIRRVPPEHQGSQLLMSQSDDRYDTHAVQSIKSLRSFIFN
PPDEWRNVDPCVYLMPFLDVIQSDDMTATATLVALQSVLKILKLHFFDEKTPGAREAVNAAVHAVTGCRLEKTDH
VMEDSVLIKVLQVLVAVMDHPASIFLTDNAVCTAVNTCFHVVQTTHRGGSLNKNARNNMQEMIQIIFSRLPDIES
TDSDTEDADIGGSYDVGYGVRCTVDIFHLMCNLLNVAEVTVEITADDMHTDNISADEDMQHFGLVMINYAIELGG
EGIRNHPKLLRMVQDDLCHHLVHYGMRSKPLVLSVISSTILNIYHFLRKSIRLQLEAFFFYVLFRVARIGSLVLQ
EVAVEAIINFVRQPTFIFEAYVNYDCDPLRCNVFEEIGKLLCRqatphvpptapqttvaaahvNFTSLHTQALEG
VLLVIHTLADSIAKEDEPRSFDPYPVRITEYSPYWEEMVRTDEDVNHMDEWVEHVRIRKAQKKKLRIAANIFNRE
EKKGLDYLKSAGLISDPPDPKAYAYFFRYLPGLDKNVIGDYLGDPGAFHLQVLKDFTDTFHFPGVILDTALRTYL
ETFRLPGEAQKIQRILEAFSDRFYDQQSLELFVSKDAVYILCYSLIMLNTDQHNPQVKKKMTEEEFIRNNRGING
GQDLPREYLSQLFRSISSHPISLFTQTGMPKEMNPSKWVELICQSKITQPFITCDVFDGKLSRDMFAVISGPSVS
LLSVIFVRTNEQEILNHCIEGFFSIVKIAQHGLYDTIDEIVTCFYKATRLLSPQELDLASVHDHRKRLATLALFT
VANDYGHGIRGGWRDIIEYLVFLKRQKLIPQFHDEPEAAASTELESQPRSSSSTFQHSYERTLGNRTTSSMIGRF
SHFLSMETVEDSLSLDGEYEQIQKIIQQCRIGNIFKNSANLPEESLLSLGRALIFAAGGKAHKFTTSAEEEDTVE
FCWDLLLTLTLVNIDRFPAFWPNFHDFILQVAQFPLFSPMPFAEKaiiallkvclkllsscREEKISEDTIFKSI
NLMWKLDKEILDICGDAITQTVSKILVEYPGNLQTQLGWKSVLHLLSVTGRHPESYPQGVEALITLMSEGTHITK
YNYAYCIDCAFGFVALKNSPIDNNVKLLDLMSNSVNLLVQWNKNYSDPGGGFSMGINSSFDDNPKAMGGSNFAMN
LFVKMGEALRKTSLARREEVRNQAVAALHKSFILAQELEMPPTNCISCFNLVIFAMVDDLHEKMLEYSRRENAER
EMRSMEGTLKIAMELLTDVFLHYVKHISESPGFRTFWLGVLRRMDTCMKADLGAYGEASLKEMVPDLLRKIIKTM
KENEILMKVEGDDLWEITYIQIQWIAPALKEELFPDEEL*

**>Rhocr_CCG027441**

MGRMKMQSGIKSIEEEPEDSETGSSSKAGLACTINSEIGAVLAVMRRNARWGGRYMSGDDQLEHSLIQSLKALRR
QIFSWQNQWHTVNPALYLQPFLDVIRSDETGAPITSVALSSVYKILTLDVLEMNTVNVEDAMHLVVDAVTSCRFE
VTDPASEEVVLMKILQVLLACMKSKTSVMLSNQHVCTIVNTCFRVVHQAGTKGELLQRFARHTMHELVRCIFAHL
SEVGNMDWALTNGSSAKQEVGSLDDDKPSATKQLENGNGSSEYDGPPSSGSYPSNASSGLIAAGMEENVGAGEGK
DSVSYDLHNMTEPYGVPCMVEIFHFLCSLLNVSEHAGLDPRSNTIVFDEDVPLFALGLINSAVELGGSSIGKHPR
LLSLIQDELFRNLMQFGLSMSPLILSMVCSIVLNLYQHLRTELKLQLEAFFSCVILRLAQSRYGASYQQQEVAVE
ALVDFCRQKSFMVEMYANLDCDITCSNVFEDLANLLSKSAFPVNSPLSSMHILALDGLIAIIQGMAERVSNGSLG
LEHSPVSLEEYTPFWMVKCDNYSDPTRWVPFVRRRKYIKRRLMIGADHFNRDPKKGLEFLQGTHLLPDKLDPQSV
ACFFRYTAGLDKNLVGDFLGNHDEFCVQVLHEFAGTFDFQDMSLDTALRLFLETFRLPGESQKIQRVLEAFSERY
YEQSPHILANKDAALLLSYSLIMLNTDQHNVQVKKKMTEEDFIRNNRHINGGNDLPREFLSELYYSICKNEIRMI
PDAGAVFPEMNPSRWIDLMNKSKKTAPFIIANDRALLDHDMFAIMSGPTIAAISVVFDHAEHDDVFQTCINGFLA
IAKISACHHLEDVLDDLVVSLCKFTTLLSPISVDEPGLAFGDDTKARMATVAVFTIANRYGDYIRTGWRNILDCI
LRLHKLGLLPARVTSDAADDSELSVEHGQGKPLTSSLSSTHLQSLGTPRRSSGLMGRFSQLLSLDTEEPRSQPTE
QQEAAYKRTMQTIQKCHIDSIFTESKFLQANSLLQLARALIWAAGRPQKGTGSSEDEDTAVFCLELLIAITLNNR
DRISLLWQGVYEHIVNIVQVTVMPCALIEKAVFGLLRICQRLLPYKENLADELLKSLQLILKLDARVADKHCEQI
TQEVSRLVKANVTQIRSQMGWRTITALLSITARHPEASEPGFDALVFIMSDGAHLSPVNFVLCVDAARQFAESRV
GQAERSVRALDLMASSMVCLSQWALEAKEAALVEDVAKLSQDIGEMWLRLSQGLRKVCFDQREEVRNHALLSLQR
CLTGADDLGFAHALWLQCFDQVIFTMLDDLLELAQGHSQKDFRNMEGTLILAMKLLSKVFLQLLHDLPQLTTFCK
LWLGILSRMEKYAKVKVRGKKSEKLQELIPELLKNILVTMKTKGILAQRSALGGDSLWELTWLHVNNIAPSLQSE
VFPDQVSDPSQDNPPTESDPCSVNGTTAPLEG*

**>Rhocr_CCG023694**

MGLLTPQSEMDTLHQELRDDSMQGAFACIINTEISALLAVMRRNVRWGVRYFAIDNKLEHPLIQSLKRLRKEIFL
WQNQWDSTNPAAYLQPFLDVIQSDETGAPITGVALSSVYKILTLDILNIYTGKVEDAMALIVDAVTSCRFEVTDP
ASEEVVLMKILQVLLACMKSKASKKLNNQHVCNIVNACFRVVHQASSKGELLQRIARHTLHEMVRCIFSCLPDIA

IDEHTCHSEGQLSTSDEAGSSDRLHLSGSQKIENDNGSDASTGKEDPKVDNGALTEDVSNEEQLMAKPFGIPCMV
EIFHFLCSLLNVIENIESGPRSNPIAYEEDVPLFSLGLINSAIELGGASFARHPKLLMLIQDELFHNLMQFGLSM
SPLILSTVCAIVLNLYHHMRTELKLQLEAFFSFVLLRLAHNKHGYSYQQQEVAMEAIVDLCRQPSFVTEMYVNYD
CDITCNNVFEDIANLLSKNAFPVNSPLCAVNLLALDGLTAIVQGIANRVGDENPFTEQTLVNSEGYESFWTVKCE
SYDDLDLWIPFVRKMKYIKRRLMVGADHFNRDSKKGLQFLQEMHLLPEKLDPRSVACFFRYTISLDKNVIGDFLG
NHDEFCVQVLQEFAKTFDFEGMSLDGALRLFLGTFRLPGESQKIQRVLEAFAERYYEQCPHILSNKDAALLLSYS
IIMLNTDQHNAQVKRKMTEEDFIRNNRCINAGKDLPKEFLSELYHSIGENEIQMLPDQGAGLPVMTSSRWVSLLQ
NSKKCSPFIICDSTSRLDYDMFTVLSGPSIAAISVVIDNMEQEEVIQACVNGFMNIAKISACYHLNDLLDDLVVS
LCKFTTLLDPSSAEESILALVEDHKARLATTSVFTIANNYGDYIRTGWKNIVDCVLYLHKLGLLRLADGTTDNEK
HSSESEKSSQSALPALSSSSSRKSSGLMGRFSQLFSFEAEEPIPQSTLEELESHQLALQTIQSCQIGNIFLESKF
LQADSLLQLVQALILAAGRLQKGPYSPEDEDTSVFCLGLLIEITVNNRDRIMLLWQSVYEHITNIVQSTVMPCTL
VEKAVFGLLQICQRLLPYKENLTDELLKSLRLVLKLDARVADAYCEHITQEVMHLVKANAPQIRSHVGWRTIISL
LSITAQHPEASEVGFETLVYIMSDTAHLSPANYVLCVDAARQLAESRVGNIERSVRALDLMADSFICLIRWSNEA
KQAVGDELAAKVYQDIGEMWMSLVQGLKRLSLDKREQVRNYAISLLQRCLIGVEGISIPQELWLQCFDAVIFALF
DDLIKVNRERSHKDYRNMEGTIVLAIKLMSKVFLQLLQDDSAQKFLINLWEGYLSRMDSCMEAKIRGRRSENIQE
VVPELLKNTLLIMKSKGILVPNDAATGKDNIWQLTWSHVKKIAPSLQSEIFSSQELEQTQSNEIKSGSMPSSDGA
VLVQANEIKA*

**>Rhocr_CCG028218**

MDIEEEDPNYNAHGHQQHRIRFKSGRDRGRGGGLSCMLNTEVSAILASVRRVPPEHQGSQLLMSQSDDRYDTHAV
QSIKSLRSFIFNPPHEWRNVDPCIYLIPFLDVIQSDDMTATATLVALQSVLKILKLHFFYHNTPGAREALNAAVH
AVTACRLEKTDHVTEDSVLIKVLQVLVAIMDHPASIFLTDNAVCTAVNTCFHVVQTTHRGGSMNKNARNTMQEMI
QIIFSRLPEIDSTDSDTEDADIGGCYDAGYGVRCTVDIFHLMCNLLNVAEVTVEITADDMHSDNIAADEDMQLFG
LLMINYAIELGGEGIRNHPKLLRMVQDDLFHHLVHYGMRSKPLVLSVISSTILNIYHFLRRSVRLQLEAFFFYVL
FRVARVGSLVLQEVAVEAIINFVRQPTFIFEAYVNYDCDPLRCNVFEEIGKLLCRQATPHVPPTAPQTTVAAAHV
NFTSLHTQALEGVLLVIHTIADSILKDDEPSDYEPYPVKITEYSPYWEEKVRTDDDIDEWVEHVRIRKAQKKKLR
IAANIFNREEKKGLDYLKAAGLISDPPDPKAYAYFFRYMPGLDKNVIGDYLGDPGAFHLQVLKDFTDTFHFPGVI
LDTALRTYLETFRLPGEAQKIQRILEAFSDRFYDQQSLELFVTKDAVYILCYSLIMLNTDQHNPQVKKKMTEEEF
IRNNRGINGGQDLPREYLSQLFRSISSHPISLFTQSGLPAEMNPSKWVEVICQSKFSEPFVPCDVFDGKLSRDMF
AVISGPSVALLSVIFVRTNEQEILNHCIEGFFSIVKVAQHGLYDTIDEIVTCFYKATRLLSPQEIDLASVHDHRK
RLATLALFTVANDYGHSIRGGWRDIVEYLVFLKRQKLIPQFLDEPETAAASAELESQPRSGSSMFHHSYERTLGN
RATSGMIGRFSHFLSMDTTEDSLSLDGEYEQIQKIIQQCRIGNIFNNSANLPEDSLLSLGRALIFAAGGKAHKFT
TSVEEDDTVGFCWDLLLTLTSVNMDRFPSFWPCFHDFLLQVAQFPLFSPMPFAEKAIISLLKICLKLLSSCREEK
ISEETIFKSINLMWKLDKEILDICGEAITQNVSKILVEYPANLQTHLGWKSVLHLLSVTGRHPESYPQGVEALIT
LMSEGTHITKYNYAYCIDCAFGFVALKNSPVDKNMKLLDLMSSSVNLLVQWNKNYSDPGSGGFSMGSNASASLFD
DNPKALGGPNFAMNLFVKMGEALRKTSLARREEVRNQAVAALHKSFILAQELEMPPTNCISCFNLVIFAMVDDLH
EKMLEYSRRENAEREMRSMEGTLKIAMELLTDVFLHYVKHISESPGFRTFWLGVLRRMDTCMKADLGAYGEASLQ
EMVPDLLRKIITIMKENQILMKVEGDDLWEITYIQIQWIAPALKEELFPDEEL*

**>HamviGNOMa_ctg1436**

MGRLKLQSGIKSIEEEPEECDTTYFNKAALACMINSEVGAVLAVMRRNVRWGGRYMSGDDQLEHSLIQSLKALRK
QIFSWQHQWHAINPAVYLQPFLDVIRSDETGAPITGVALSSVYKILTLDVLDLNTVNVEDAMHLVVDAVTSCRFE
VTDPASEEVVLMKILQVLLACMKSKASVMLSNQHVCTIVNTCFRVVHQAGSKGELLQRMARHTMYELVRCIFAHL
PDVDTTECALVNGGSAVKQEVXXLDNDYAFGNKQLENGNGSSEYDGQRSSGGLVSNASTGLVATGMDENIIGSGN
GKEAVPYDLHLMTEPYGVPCMVEIFHFLCSLLNVVEHMGMGPRSNTIAFDEDVPLFALGLINSAIELGGPSICRH
PRLLSLIQDELFRNLMQFGLSMSPLILSMVCSIVLNLYQHLRTELKLQLEAFFSCVILRLAQSRYGASYQQQEVA
MEALVDFCRQKTFMVEMYANLDCDITCSNVFEDLANLLSKSAFPVNCPLSAMHILALDGLIAVIQGMAERVGNGS
LSTEQAPVNLEEYTPFWMVKCDNYNDPNHWVPFVRRRKYIKRRLMIGADHFNRDPKKGLEFLQGTLLLPDKLDPQ
SVACFFRYTAGLDKNLVGDFLGNHDEFCVQVLHEFAWTFDFQDMNLDTALRLFLETFRLPGESQKIQRVLEAFSE
RYYEQSPQILANKDAALLLSYSLIMLNTDQHNVQVKKKMTEEDFIRNNRHINGGNDLPREFLSELYQSICKNEIR
TTPEQGAGFPEMTPSRWIDLMHKSKKTAPFIVADSRALLDHDMFAIMSGPTIAAISVVFDHAEHEDVYQTCIDGF
LAVAKISACHHLEDVLDDLVVSLCKFTTLLNPSSVEEPVLAFGDDTKARMATVTVFTIANRYGDYIRTGWRNILD
CILKLHKLGLLPARVASDAADDSELSADPGHGKPLTNSLSSAHMQSLGTPRRSSGLMGRFSQLLSLDTEEPRSQP
TEQQLAAHQRTLQTIQKCHIDSIFTESKFLQADSLLQLARALIWAAGRPQKGNSSPEDEDTAVFCLELLIAITLN
NRDRIVLLWQGVYEHIANIVQSTVMPGALVEKAVFGLLRICQRLLPYKENLADELLRSLQLVLKLDARVADAYCE
QITQEVSRLVKANATHIRSQMGWRTITSLLSITARHPEASEAGFDALLFIMSDGVHLLPANYVLCLDASRQFAES
RVGQAERSVRALDLMAGSVSCLSRWAHETKEAMGEEDAAKMSQDIGEMWLRLIQGLRKVCLDQREEVRNHALLSL
QRCLTLLDGLHLPHSLWLQCFDLVIFTMLDDLLEIGHSQKDYRNMEGTLILAMKLLSKVFLQLLNDLSQLTTFCK
LWLGVLSRMEKYVKVRVRGKRCEKLQDLVPELLKNILLVMKTRGVLVQRSALGGDSLWELTWLHANNIAPSLQSE
VFPDQDLDQSLQKQGVTGGSPVPEETVTVPKNEKVASEGTRTGG*

**>HamviGNOMb_ctg1436**

MGRLKLQSGIKSIEEPEECDTTYTNKAALACMINSEVGAVLAVMRRNVRWGGRYISGDDQLELSLIQSLKALRKQ
IFSWEHQWHAINPAVYLQPFLDVIRSDETGAPITGVALSSVYKILTLDVLDLNTVNVKDAMHLVVDAVTSCRFEV
TDPGSEEVVLMKILQVLLASMKSKASVMLSNQHVCTIVSTCFRVLHQAGSKGELLQRIARHTMYELVRCIFAHLP
DVDTTECTLVNGGSAVKQKVSRLDDDYAFENKQLENGNGSSEHDGQPSSGSLVSNASTGLVATGMDENIIGSGNG

EAVLYDLHLMTEPYGVPCMVEILHFLCSLLNVVEHMGMGPRSSIIAFDEDVPLFALGLINSAIELGGPSICQHPR
LLSLIQDELFHNLMQFGLSMSPLILSMVCSIVLNLYQRLRTELKLQLEAFISYVILRIAQSRYGASYQQQEVAME
ALVDFCRQKTFMVEMYANLDCDITCSNVFEDLANLLSKSAFPVNCPLSDMHILALDGLIAVIQGIAERLGNGSLS
SEQAPVNLEEYTPFWMVKCDNYSDPNHWVPFVRRRKYIKRRLMIEADHFNRDPKKGLEFLQGTHLLPDKLDPQSV
ACFFRYTAGLDKNLVGDFLGNHDEFCVQVLHEFARTFDFQDMNLDTALRLFLETFRLPGEAQKIQRVLEAFSERY
YEQSPQILANKDASLLLSYSLILLNTDQHNVQVKKKMTEEDFVRNNRRINGGNDLPREFLSELYHSICKNEIRTI
PEQGASFPEMTPSRWIDLMQKSKKTAPFIVADSRALLDYDMFAIMSGPTIAAISVVFDHAEHEDIYQTCIDGFLA
VAKISACHHLEDVLDDLVVSLCKFTTLLNPSSVEEPVLAFGDDTKARMATVTVFTIVNRYGDYIRTGWRNILDCI
IKLHKLGLLPARVASDAADDSELSADPGHGKPLTNSLSSSNMQSLGTPRRSYGLMSRFSQLLSLDTEEPRSQPTE
QQLAAHQLTLQTIQKCHIDSIFTESKFLQADSLlQLAQLAIWAAGQPQKGNTSPEDEDTAVFCLELLITITLNNR
DRIVLLWQGVYEHITNIVQSTVMPGALVEKAVFGLLRICQRLLPYKENLADELLRSLQLVLKLDARVADAYCEQI
TQEVSRLVKANATHIRSQMGWRTITSLLSITARHPEASEAGFDALLFIMSDGVHLLPANYVLCLDASRQFAESRV
GQAEQSVLALDLMAGSVACLSRWAHETKEAMGEEDAARMSQDIGEMWLRLIQGLRKVCLDQREEVRNHALLSLQR
CLTGVDGLHLPHSLWLQCFDLVIFSMLDDLLEIAWGHSQKDYRNMEGTLILAMKLLSKVFLQLLHDLSQLTTFCK
LWLGVLSQMEKYMKVTLRGKRSEMLQDLVPELLKNTLLVMKTRGVLVQRSALGGDSLWELTWLHVNNIAPSLQSE
VFPDQDPHQSQQKQGVTGASPLSEETVTVPKNEKVVSEGTCT*
**>HamviGNL2_ctg1273**
MEKANDQNQEGGSDNKCSSCRHKKKEVTRQKRKELGLSCMLNTEVGAVLAVIRRPSELNSQFMSSPEENYDSSIL
QSLKSLRALIFNPQQEWRTIDPSIYLSPFLDVVQSDDVPAAATGVALSAILKILKLEIFDEKTPGARDCINSVVT
AITSCRLEKTDPISEDAVLMRILQVLIGIMKSRASVLLTDHAVCTIVNTCFQVVQQSASRGDLLQRNARYIMHEL
IQTIFSRLPEIEVKDEDNSETDTEDADIGSNSDSGYGGRCAVDIFHFLCSLLNVVEVMEMEGSISRTADEDVQLF
ALVLINSAIELSGDGIGKHPKLLRMIQDDLFHHLIHYGTCSSPLVLSMICSTVLSIYHFLRRLIRLQLEAFFTFV
LFRNASIGSSIQLQEVALEAIISFCRQPTFIFEIYVNYDCDPIFQDVFEEIGKLLCKHAFPLGNPLTSLQIQAFE
GLVVMIHTIADNKDREDNNSSSLGMYPIEITEYKPFWEVKSKNDDDLDAWVEFVRVKKAQKRKIRIAGNHFNRDE
KKGLDFLKISSLVPDPPDPKAFAYFFRYTPRLEKGMIGDYLGDPDEFHIQVLKEFSGTFEFSGMVLDTALRTYLE
TFRLPGEAQKIQRILEAFSEKFYDLQSSEIFVSKDAVFILCYSLIMLNTDQHNPQVKKKMTEEEFIKNNRANNGG
KDFPREYLSELFHSISNNAITIFGQSGLPIEMNPSRWIELINRGKVMEPFIQCDSDRRLGRDMFAAIAGPSIAAL
SAFFEHADEDEMLNECIEGLFSVARIAQCGLEDTLDELIASFCKFTTLLNPYASAEETLYAFSNDLKPRMATLAV
FTIANSFGESILGGWRNIVDCLLKLKRLKLLPQSVVEPDNASAPLSELETHSRSESGVIFPAHDLSFGSSRQSSG
MIGRFSHFLTLDAAEDSLTLCGSEFEQNLKIIQQCRIGNIFSSSSSLPEEPLLNLGRSLIFAAAGKGQKFSTPVE
EEETVGFCWDLIITIALANIHRFTTFWSSFHDYLLGVAQFPLFSPVPFAEKAILGLFKVCLKILASCQSDKHSEE
LIFKSINLMWKLDKEILDTCSEFITQSVSKILTECPANLQTQLGWKSVLHLLSITGRHTETYDQGVETLIMLMSD
GIHVSRINFAYCIDCAFGFVALKNSPLEKNLKILDLMSDSVNLLIQWYKNGHSDPGSNFSIASNTSSSSVEDNLK
GIGSSNFTMNLFVKLGDALRKTSLARREEIRKQAILALRKTFSLAEELDLTSTNCINCFNLIIFAMVDDLHEKMV
EYSRRENSEKEMRSMEGTLKLAMEFLTDMYLQYLKPLSESSGFRTFWLGMLRRMDTCMKADLGDYGDSKLQELVP
ELLKKMITTMQEKEILVQKEGEDLWEITYIQIQWIAPSLKEQLFPDEF*
**>Vitvi_LOC100247152**
MGRLKLQSGIKSIEEEPEDCESTSSNKAALACMINSEVGAVLAVMRRNVRWGGRYMSGDDHLEHSLIQSLKALRK
QIFSWQHQWHTINPAVYLQPFLDVIRSDETGAPITGVALSSVYKIVTLDVLCLNTVNVEDAMHLVVDAVTSCRFE
VTDPASEELVLMKILQVLLACMKSKVSVMLSNQHVCTIVNTCYRIVHQAATKSELLQRIARHTMHELVRCIFSHL
PDVGNTEHALVNRGSSVKLEGSGQDNEYNFGNKQLENGNGASEYDGQPSSVSFASNSSTGLVGSMLDENTVGAGN
GKEATPYDLHLMTEPYGVPCMVEIFHFLCSLLNVVEHMGMGSRSNTMAFDEDLPLFALGLINSAIELGGLSIRRH
PRLLSLIQDELFRNLMQFGLSTSPLILSMVCSIVLNLYQHLRTELKLQLEAFFSCVILRLAQSKYGASYQQQEVA
MEALVDFCRQKTFMVEMYANLDCDITCSNVFEDLANLLSKSAFPVNCPLSAMHILALDGLIAVIQGMAERIGNGS
LGSEQSPVNLEEYTPFWMVKCDNYSDPSVWVPFVCRRKYIKRRLMIGADHFNRDPKKGLEFLQVTHLLPDKLDPQ
SVACFFRYTAGLDKNLVGDFLGNHDEFCVQVLHEFAGTFDFQDMNLDTALRLFLETFRLPGESQKIQRVLEAFSE
RYYEQSPQILANKDAALLLSYSLIMLNTDQHNVQVKKKMTEEDFIRNNRHINGGNDLPRDFLSELYHSICKNEIR
TTPEQGAGFPEMTPSRWIDLMHKSKKTAPFIVADSRAFLDHDMFAIMSGPTIAAISVVFDHAEHEEVYQTCIDGF
LAVAKISACHHLEDVLDDLVVSLCKFTTLLNPSPGEESVQAFGDDTKARMATVTVFTIANRYGDYIRTGWRNILD
CILRLHKLGLLPARVASDAADDSELSADPGQGKPITNSLSSAHMPSIGTPRRSSGLMGRFSQLLSLDTEEPRSQP
TEQQLAAHQRTLQTIQKCHIDSIFTESKFLQSDSLLQLARALIWAAGRPQKGNSSPEDEDTAVFCLELLIAITLN
NRDRIKLLWQGVYEHISNIVQSTVMPCALVEKAVFGLLRICQRLLPYKENLADELLRSLQLVLKLDARVADAYCE
QITQEVSRLVKANATHIRSQMGWRTITSLLSITARHPEASEAGFDALLFIMSDGAHLLPANYVLCVDAARQFSES
RVGQAERSVRALDLMAGSVVCLSHWALEAKQAMAEEELSKMSQDIGEMWLRLVQGLRKVCLDQREEVRNHALISL
QRCLSGVEGFQLPHSLWLQCFDMVIFTMLDDLLDIAQGHSQKDYRNMEGTLSLAMKLLSKVFLQLLNDLAQLTTF
CKLWLGVLSRMEKYMKVKVKGKRSEKLPELVPELLKNTLLVMKTRGVLVQRSALGGDSLWELTWLHVNNIAPTLQ
SEVFPDQGLDQPRDKKDETGRSLVSDEMGSVPSNETVVSEGGRTG*
**>Vitvi_LOC100255904**
MEKTRDDNGEEEEEENKPKNVRELRRRELGLSCMLNTEVGAVLAVIRRPSDTNSQFLAAQEDNFDSSLLHSLKSL
RALIFNPQQEWRTIDPSIYLAPFLDVVQSDDVPATATGVALSAIFMILKLEIFDEKTPGARDAVNSIVTGITTCR
LERTDPISEDAILMKILHVLTEIMKKRASILLTDHAVCTIVNTCFQVVQQSAIRGDLLQRGARHTMQELIQTIFS
RFPEINKDRDKSLNDFQDGDINNEMESGYGVRSAVDIFSFLCSLLNVVEVVDMDGSQVQTADEDVQLFALVLINS
AIQLSGDEIGKQPKLLRMIEDDLFHHLIHYGMCSSPLVLSMICSTVLNIYNFLRRFIRLQLEAFFTFVSFRVAVQ

ASSSLQLKEVALEAVINFCRQPTFIFEAYANYDCHIIFRDVFEEIGRLLCKHAFPTGSPLSTLQIQAFEGLVIMI
HNISDHVDGEHDSSSSGPYPVEITEYRPFWDENFKANDSEDWADHARLRKAQKRKIKIAGDHFNRDEKKGLDYLK
ISHLVPDPPDPKPFAYFFRYTPGLDKSMIGDYLGSPDELNLQVLKEFTETFNFSGMILDNALRTYLETFRLPGES
QKIQRILEAFSERFYDQQSSEIFVSKDAVFILCYSLIMLNTDQHNPQVKKKMTEEEFIKNNRAINGGKDLPREYL
SELFQSISNKAISLFGQSGQPVEMNPSRWIELINRSRNSEPFIKCDFDHRLGRDMFAAIAGPTVAALSAIFEHSD
EDEIINECMQGLISVARIAQYGLQDTLDELLASFCKFTTLLNPYASAEETLYAFSNDLKARMATLAVFTIANNFG
HSIKGGWRNIVDCLLKLRRLKLLPQSVVEPEIASTSSSDLQMHTRSDSGVIFPSYESSFDSNCQTSSMMSRFSHF
LSMETADDSLTLGGSEFERNLKIIQQCRIGNIFSNSSKLPDDALPNLGRSLIFAAAGKGQKFSTPIEEEETVGFC
WDLLISIALANVHRFSTFWPYFHDYLLAVAQFPLFSPVPFVEKAMLGLFKICLKVLSSYQSDKLLEELIFKSINL
MWKLDKEILDTCCEYITQSVSKILIEYPANLQSQLGWKSVLQLLSITGRHPETYDQAVETLIMLMSDGFHVSRSN
YPFCIDCAFGFIALKNSPLEKNLKLLDLLADSVNLLIQWSKNGYSDPGNNLSIASSTSSSSFEDNSKFNPSNFSM
NLFIKLGEALRRTSLARREEIRNHAIVALQRSFTLADELDFTSVSCINCFNLVIFAMVDDLHEKMVEYSRRDNLE
REVRSMEGTLKIAMELLTDVFLQFLKPLTENSGFRTFWLGVLRRMDTCMKADLGGYGETTLQELIPDLLRKMITI
MKEKQILVQKEGEDLWEITYIQIQWIAPSLKDELFPDEDL*

**>Vitri_LOC117929271**

MGRLKLQSGIKSIEEEPEDCESTSSNKAALACMINSEVGAVLAVMRRNVRWGGRYMSGDDHLEHSLIQSLKALRK
QIFSWQHQWHTINPAVYLQPFLDVIRSDETGAPITGVALSSVYKIVTLDVLCLNTVNVEDAMHLVVDAVTSCRFE
VTDPASEELVLMKILQVLLACMKSKVSVMLSNQHVCTIVNTCYRIVHQAATKSELLQRIARHTMHELVRCIFSHL
PDVGNTEHALVNRGSSVKLEGSGQDNEYNFGNKQLENGNGASEYDGQPSSVSFASNSSTGLVGSMLDENTVGAGN
GKEATPYDLHLMTEPYGVPCMVEIFHFLCSLLNVVEHMGMGSRSNTMAFDEDLPLFALGLINSAIELGGLSIRRH
PRLLSLIQDELFRNLMQFGLSTSPLILSMVCSIVLNLYQHLRTELKLQLEAFFSCVILRLAQSKYGASYQQQEVA
MEALVDFCRQKTFMVEMYANLDCDITCSNVFEDLANLLSKSAFPVNCPLSAMHILALDGLIAVIQGMAERIGNGS
LGSEQSPVNLEEYTPFWMVKCDNYSDPSVWVPFVCRRKYIKRRLMIGADHFNRDPKKGLEFLQVTHLLPDKLDPQ
SVACFFRYTAGLDKNLVGDFLGNHDEFCVQVLHEFAGTFDFQDMNLDTALRLFLETFRLPGESQKIQRVLEAFSE
RYYEQSPQILANKDAALLLSYSLIMLNTDQHNVQVKKKMTEEDFIRNNRHINGGNDLPRDFLSELYHSICKNEIR
TTPEQGAGFPEMTPSRWIDLMHKSKKTAPFIVADSRAFLDHDMFAIMSGPTIAAISVVFDHAEHEEVYQTCIDGF
LAVAKISACHHLEDVLDDLVVSLCKFTTLLNPSPGEESVQAFGDDTKARMATVTVFTIANRYGDYIRTGWRNILD
CILRLHKLGLLPARVASDAADDSELSADPGQGKPITNSLSSAHMPSIGTPRRSSGLMGRFSQLLSLDTEEPRSQP
TEQQLAAHQRTLQTIQKCHIDSIFTESKFLQSDSLLQLARALIWAAGRPQKGNSSPEDEDTAVFCLELLIAITLN
NRDRIKLLWQGVYEHISNIVQSTVMPCALVEKAVFGLLRICQRLLPYKENLADELLRSLQLVLKLDARVADAYCE
QITQEVSRLVKANATHIRSQMGWRTITSLLSITARHPEASEAGFDALLFIMSDGAHLLPANYVLCVDAARQFSES
RVGQAERSVRALDLMAGSVVCLSHWALEAKQAMAEEELSKMSQDIGEMWLRLVQGLRKVCLDQREEVRNHALISL
QRCLSGVEGFQLPHSLWLQCFDMVIFTMLDDLLDIAQGHSQKDYRNMEGTLSLAMKLLSKVFLQLLNDLAQLTTF
CKLWLGVLSRMEKYMKVKVKGKRSEKLPELVPELLKNTLLVMKTRGVLVQRSALGGDSLWELTWLHVNNIAPTLQ
SEVFPDQGVDQPRDKKDETGRSLVSDEMGSVPSNETVVSEGGRTG*

**>Vitri_LOC117925203**

MEKTRDDNGEEEEEENKPKNVRELRRRELGLSCMLNTEVGAVLAVIRRPSDTNSQFLAAQEDNFDSSLLHSLKSL
RALIFNPQQEWRTIDPSIYLAPFLDVVQSDDVPATATGVALSAIFKILKLEIFDEKTPGARDAVNSIVTGITTCR
LERTDPVSEDAILMKILHVLTEIMKKRASILLTDHAVCTIVNTCFQVVQQSAIRGDLLQRGARHTMQELIQTIFS
RFPEITKDRDKSLNDFQDCDINNEMESGYGVRSAVDIFSFLCSLLNVVEVVDMDGSQVQTADEDVQLFALVLINS
AIQLSGDEIGKQPKLLRMIEDDLFHHLIHYGMCSSPLVLSMICSTVLNIYNFLRRFIRLQLEAFFTFVSFRVAVQ
ASSSLQLKEVALEAVINFCRQPTFIFEAYANYDCHIIFRDVFEEIGRLLCKHAFPTGSPLSTLQIQAFEGLVIMI
HNISDHVDGEHDSSSSGPYPVEITEYRPFWDENFKANDSEDWADHARLRKAQKRKIKIAGDHFNRDEKKGLDYLK
ISHLVPDPPDPKPFAYFFRYTPGLDKSMIGDYLGSPDELNLQVLKEFTETFNFSGMILDNALRTYLETFRLPGES
QKIQRILEAFSERFYDQQSSEIFVSKDAVFILCYSLIMLNTDQHNPQVKKKMTEEEFIKNNRAINGGKDLPREYL
SELFQSISNKAISLFGQSGQPVEMNPSRWIELINRSRNSEPFIKCDFDHRLGRDMFAAIAGPTVAALSAIFEHSD
EDEIINECMQGLISVARIAQYGLQDTLDELLASFCKFTTLLNPYASAEETLYAFSNDLKARMATLAVFTIANNFG
HSIKGGWRNIVDCLLKLRRLKLLPQSVVEPEIASTSSSDLQMHTRSDSGVIFPSYESSFDSNCQTSSMMSRFSHF
LSMETADDSLTLGGSEFERNLKIIQQCRIGNIFSNSSKLPDDALPNLGRSLIFAAAGKGQKFSTPIEEEETVGFC
WDLLISIALANVHRFSTFWPYFHDYLLAVAQFPLFSPVPFVEKAMLGLFKICLKVLSSYQSDKLLEELIFKSINL
MWKLDKEILDTCCEYITQSVSKILIEYPANLQSQLGWKSVLQLLSITGRHPETYDQAVETLIMLMSDGFHVSRSN
YPFCIDCAFGFIALKNSPLEKNLKLLDLLADSVNLLIQWSKNGYSDPGNNLSIASSTSSSSFEDNSKFNPSNFSM
NLFIKLGEALRRTSLARREEIRNHAIVALQRSFSLADELDFTSVSCINCFNLVIFAMVDDLHEKMVEYSRRDSLE
REVRSMEGTLKIAMELLTDVFLQLLKPLTENPGFRTFWLGVLRRMDTCMKADLGGYGETTLQELIPDLLRKMITI
MKEKQILVQKEGEDLWEITYIQIQWIAPSLKDELFPDEDL*

**>VITMro_g023260**

MGRLKLQSGIKSIEEEPEDCESTSSNKAALACMINSEVGAVLAVMRRNVRWGGRYMSGDDHLEHSLIQSLKALRK
QIFSWQHQWHTINPAVYLQPFLDVIRSDETGAPITGVALSSVYKIVTLDVLCLNTVNVEDAMHLVVDAVTSCRFE
VTDPASEELVLMKILQVLLACMKSKVSVMLSNQHVCTIVNTCYRIVHQAATKSELLQRIARHTMHELVRCIFSHL
PDVGNTEHALVNRGSSVKLEGNGQDNEYNFGNKQLENGNGASEYDGQPSSVSFASNSSTGLVGSMLDETTVGAGN
GKEAAPYDLHLMTEPYGVPCMVEIFHFLCSLLNVVEHMGMGSRSNTMTFDEDLPLFALGLINSAIELGGLSIRRH
PRLLSLIQDELFRNLMQFGLSTSPLILSMVCSIVLNLYQHLRTELKLQLEAFFSCVILRLAQSKYGASYQQQEVA
MEALVDFCRQKTFMVEMYANLDCDITCSNVFEDLANLLSKSAFPVNCPLSAMHILALDGLIAVIQGMAERIGNGS

LGSEQSPVNLEEYTPFWMVKCDNYGDPSVWVPFVCRRKYIKRRLMIGADHFNRDPKKGLEFLQVTHLLPDKLDPQ
SVACFFRYTAGLDKNLVGDFLGNHDEFCVQVLHEFAGTFDFQDMNLDTALRLFLETFRLPGESQKIQRVLEAFSE
RYYEQSPQILANKDAALLLSYSLIMLNTDQHNVQVKKKMTEEDFIRNNRHINGGNDLPRDFLSELYHSICKNEIR
TTPEQGAGFPEMTPSRWIDLMHKSKKTAPFIVADSRAFLDHDMFAIMSGPTIAAISVVFDHAEHEEVYQTCIDGF
LAVAKISACHHLEDVLDDLVVSLCKFTTLLNPSPGEESVQAFGDDTKARMATVTVFTIANRYGDYIRTGWRNILD
CILRLHKLGLLPARVASDAADDSELSADPGQGKPITNSLSSAHMPSIGTPRRSSGLMGRFSQLLSLDTEEPRSQP
TEQQLAAHQRTLQTIQKCHIDSIFTESKFLQSDSLLQLARALIWAAGRPQKGNSSPEDEDTAVFCLELLIAITLN
NRDRIKLLWQGVYEHISNIVQSTVMPCALVEKAVFGLLRICQRLLPYKENLADELLRSLQLVLKLDARVADAYCE
QITQEVSRLVKANATHIRSQMGWRTITSLLSITARHPEASEAGFDALLFIMSDGAHLLPANYVLCVDAARQFSES
RVGQAERSVRALDLMAGSLVCLSNWALEAKQAMAEEELSKMSQDIGEMWLRLVQGLRKVCLDQREEVRNHALVSL
QRCLSGVEGFQLPHSLWLQCFDMVIFTMLDDLLDIAQGHSQKDYRNMEGTLSLAMKLLSKVFLQLLNDLAQLTTF
CKLWLGVLSRMEKYMKVKVKGKRSEKLPELVPELLKNTLLVMKTRGVLVQRSALGGDSLWELTWLHVNNIAPTLQ
SEVFPDQGLDQPRDKKDETGRSLVSDEMGSVPSNETVVSEGGRTGG*

**>VITMRO_g058050**

MGRLKLQSGIKSIEEEPEDCESTSSNKAALACMINSEVGAVLAVMRRNVRWGGRYMSGDDHLEHSLIQSLKALRK
QIFSWQHQWHTINPAVYLQPFLDVIRSDETGAPITGVALTSVYKIVTLDVLCLNTVNVEDAMHLVVDAVTSCRFE
VTDPASEELVLMKILQVLLACMKSKVSVMLSNQHVCTIVNTCYRIVHQAATKSELLQRIARHTMHELVRCIFSHL
PDVGNTEHALVNRGSSVKLEGSGQDNEYNFGNKQLENGNGASEYDGQPSSVSFASNSSTGLVGSMLDETTVGAGN
GKEAAPYDLHLMTEPYGVPCMVEIFHFLCSLLNVVEHMGMGSRSNTMAFDEDLPLFALGLINSAIELGGLSIRRH
PRLLSLIQDELFRNLMQFGLSTSPLILSMVCSIVLNLYQHLRTELKLQLEAFFSCVILRLAQSKYGASYQQQEVA
MEALVDFCRQKTFMVEMYANLDCDITCSNVFEDLANLLSKSAFPVNCPLSAMHILALDGLIAVIQGMAERIGNGS
LGSEQSPVNLEEYTPFWMVKCDNYGDPSVWVPFVCRRKYIKRRLMIGADHFNRDPKKGLEFLQVTHLLPDKLDPQ
SVACFFRYTAGLDKNLVGDFLGNHDEFCVQVLHEFAGTFDFQDMNLDTALRLFLETFRLPGESQKIQRVLEAFSE
RYYEQSPQILANKDAALLLSYSLIMLNTDQHNVQVKKKMTEEDFIRNNRHINGGNDLPRDFLSELYHSICKNEIR
TTPEQGAGFPEMTPSRWIDLMHKSKKTAPFIVADSRAFLDHDMFAIMSGPTIAAISVVFDHAEHEEVYQTCIDGF
LAVAKISACHHLEDVLDDLVVSLCKFTTLLNPSPGEESVQAFGDDTKARMATVTVFTIANRYGDYIRTGWRNILD
CILRLHKLGLLPARVASDAADDSELSADPGQGKPITNSLSSAHMPSIGTPRRSSGLMGRFSQLLSLDTEEPRSQP
TEQQLAAHQRTLQTIQKCHIDSIFTESKFLQSDSLLQLARALIWAAGRPQKGNSSPEDEDTAVFCLELLIAITLN
NRDRIKLLWQGVYEHISNIVQSTVMPCALVEKAVFGLLRICQRLLPYKENLADELLRSLQLVLKLDARVADAYCE
QITQEVSRLVKANATHIRSQMGWRTITSLLSITARHPEASEAGFDALLFIMSDGAHLLPANYVLCVDAARQFSES
RVGQAERSVRALDLMAGSLVCLSNWALEAKQAMAEEELSKMSQDIGEMWLRLVQGLRKVCLDQREEVRNHALVSL
QRCLSGVEGFQLPHSLWLQCFDMVIFTMLDDLLDIAQGHSQKDYRNMEGTLSLAMKLLSKVFLQLLNDLAQLTTF
CKLWLGVLSRMEKYMKVKVKGKRSEKLPELVPELLKNTLLVMKTRGVLVQRSALGGDSLWELTWLHVNNIAPTLQ
SEVFPDQGLDQPRDKKDETGRSLVSDEMGSVPSNETVVSEGGRTGG*

**>VITMro_g386820**

MDKTRDDNGEEEEEEENKPKNVRELRRRELGLSCMLNTEVGAVLAVIRRPSDTNSQFLAPQEDNFDSSLLHSLKSL
RALIFNPQQEWRTIDPSIYLAPFLDVVQSDDVPATATGVALSAILKILKLEIFDEKTPGAKDAVNSIVTGITSCR
LERTDPISEDAILMKILHVLTEIMKKRASILLTDYAVCTIVNTCFQVVQQSAIRGDLLQRGARHTMQELIQAVFS
RFPEINKDRDKSLNDFQDGDINNEMESGYGVRSAVDIFSFLCSLLNVVEVVDMDGSQVQTADEDVQLFALVLINS
AIQLSGDEIGKQPKLLRMIEDDLFHHLIHYGMCSSPLVLSMICSTVLNIYNFLRRFIRLQLEAFFTFVSFRVAVQ
SSSSLQLKEVALEAVINFCRQPTFIFEAYANYDCHIIFRDVFEEIGRLLCKHAFPTGSPLSTLQIQAFEGLVIMI
HNISDHVDGEHDSSSSGPYPVEITEYRPFWDENFKANDSEDWVDHARLRKAQKRKLKIAGNHFNRDEKKGLDYLK
ISHLVPDPPDPKPFAYFFRYTPGLDKSMIGDYLGSPDELNLQVLQEFTETFNFSGMILDNALRTYLETFRLPGES
QKIQRILEAFSERFYDQQSSEIFVCKDAVFILCYSLIMLNTDQHNPQVKKKMTEEEFIKNNRAINGGKDLPREYL
SELFQSISNKAISLFGQSGQPVEMNPSRWIELINRSRNSEPFIKCDFYHRLGRDMFAAIAGPTVAALSAIFEHSD
EDEIINECMQGLISVARIAQHGLQDTLDELLASFCKFTTLLNPYVSAEETLYAFSNDLKARMATLAVFTIANNFG
HSIKGGWRNIVDCLLKLRRLKLLPQSVVEPEIASTSSSDLQMHTRSDSGVIFPSYESSFDSNCQNSSMMSRFSHF
LSLETADDSLTLGGSEFERNLKIIQQCRIGNIFSNSSKLPDDALPNLGRSLIFAAAGKGQKFSTPIEEEETVGFC
WDLLISIALANVHRFSTFWPYFHDCLLAVAQFPLFSPVPFVEKAMLGLFKICLKVLSSYQSDKLLEELIFKSINL
MWKLDKEILDTCCEYITQSVSKILIEYPANLQSQLGWKSVLQLLSITGRHPETYDQAVETLIMLMSDGFHVSRSN
YPFCIDCAFGFIALKNSPLEKNLKLLDLLADSVNLLIQWYKNGYSDPGNNLSIASSTSSSSFEDNSKFNPSNFSM
NLFIKLGEALRRTSLARREEIRNHAIVALQRSFTLADELDFTSVSCINCFNLVIFAMVDDLHEKMVEYSRRDNLE
REVRSMEGTLKIAMELLTDVFLQFLKPLTENPGFRTFWLGVLRRMDTCMKADLGGYGETTLQELIPDLLRKMITI
MKEKQILVQKEGEDLWEITYIQIQWIAPLLKDDLFPDEDL*

**>VITMro_g138390**

MDKTRDDNGEEEEEEENKPKNVRELRRRELGLSCMLNTEVGAVLAVIRRPSDTNSQFLAPPEDNFDSSLLHSLKSL
RALIFNPQQEWRTIDPSIYLAPFLDVVQSDDVPATATGVALSAILKILKLEIFDEKTPGAKDAVNSIVTGITSCR
LERTDPISEDAILMKILHVLTEIMKKRASILLTDYAVCTIVNTCFQVVQQSAIRGDLLQRGARHTMQELIQAVFS
RFPEINKDRDKSLNDFQDGDINNEMESGYGVRSAVDIFSFLCSLLNVVEVVDMDGSQVQTADEDVQLFALVLINS
AIQLSGDEIGKQPKLLRMIEDDLFHHLIHYGMCSSPLVLSMICSTVLNIYNFLRRFIRLQLEAFFTFVSFRVAVQ
SSSSLQLKEVALEAVINFCRQPTFIFEAYANYDCHIIFRDVFEEIGRLLCKHAFPTGSPLSTLQIQAFEGLVIMI
HNISDHVDGEHDSSSSGPYPVEITEYRPFWDENFKANDSEDWVDHARLRKAQKRKLKIAGNHFNRDEKKGLDYLK
ISHLVPDPPDPKPFAYFFRYTPGLDKSMIGDYLGSPDELNLQVLQEFTETFNFSGMILDNALRTYLETFRLPGES

QKIQRILEAFSERFYDQQSSEIFVCKDAVFILCYSLIMLNTDQHNPQVKKKMTEEEFIKNNRAINGGKDLPREYL
SELFQSISNKAISLFGQSGQPVEMNPSRWIELINRSRNSEPFIKCDFYHRLGRDMFAAIAGPTVAALSAIFEHSD
EDEIINECMQGLISVARIAQHGLQDTLDELLASFCKFTTLLNPYVSAEETLYAFSNDLKARMATLAVFTIANNFG
HSIKGGWRNIVDCLLKLRRLKLLPQSVVEPEIASTSSSDLQMHTRSDSGVIFPSYESSFDSNCQNSSMMSRFSHF
LSLETADDSLTLGGSEFERNLKIIQQCRIGNIFSNSSKLPDDALPNLGRSLIFAAAGKGQKFSTPIEEEETVGFC
WDLLISIALANVHRFSTFWPYFHDCLLAVAQFPLFSPVPFVEKAMLGLFKICLKVLSSYQSDKLLEELIFKSINL
MWKLDKEILDTCCEYITQSVSKILIEYPANLQSQLGWKSVLQLLSITGRHPETYDQAVETLIMLMSDGFHVSRSN
YPFCIDCAFGFIALKNSPLEKNLKLLDLLADSVNLLIQWYKNGYSDPGNNLSIASSTSSSSFEDNSKFNPSNFSM
NLFIKLGEALRRTSLARREEIRNHAIVALQRSFTLADELDFTSVSCINCFNLVIFAMVDDLHEKMVEYSRRDNLE
REVRSMEGTLKIAMELLTDVFLQFLKPLTENPGFRTFWLGVLRRMDTCMKADLGGYGETTLQELIPDLLRKMITI
MKEKQILVQKEGEDLWEITYIQIQWIAPLLKDDLFPDEDL*

**>Sanal_SaT02071**

MGRLKLQSGIKSIEEEPEDCDAAYSNRAALACMINSEVGAVLAVMRRYVRWGGRYISGDDQLEHSLIQSLKALRRQIFSW
QHQWHTINPVVYLQPFLDVIRSDETGAPITGVALSSVHKILSLDVLDLNTVNVDDAMHLVVDAVTSCRFEVTDSISEDVV
LMKILQVLLACMKSKASVVLSNQHVCTIVNTCFRVVHQAGTKGELLQRIARHTMHELVRCIFSHLSNVDNTERALMNPGN
NLKQESGRQQSDYAFGSKQLVNENGTSGLADENTIMAGDRKDVTPYDLHLMTEPYGIPCMVEIFHFLCSLLNVVEHMGVG
PRSDAKPFDEDVPLFALGLINSAIELGGASIRHHPRLLSLIQDELFRNLMQFGLSMSPLILSMVCSVVLNLYHHLRTELK
LHLEAFFSCVILRLAQSRYGASYQQQEVAMEALVDFCRQKSFMVEMYANLDCDITCSNVFEDLANLLSKSAFPVNCPLSS
MHILALDGLIAVIQGMAERIGNGPFGLEQTPVNLEEYTPFWMVKCDNYADPNFWVPFVRRRKYIKRRLMIGADHFNRDPK
KGLEFLQGTHLLPDKLDPQSVACFFRYTAGLDKNLVGDFLGNHDEFCVQVLHEFAGTFDFQDMNLDTALRLFLETFRLPG
ESQKIQRVLEAFSERYYEQSPQILVNKDAALVLAYSLIMLNTDQHNVQVKKKMTEEDFIRNNRHINGGSDLPREFLSELY
HSICKNELRTTPEQGAGFPEMTPSRWIDLMHKSKKTAPFIVSDSRAYLDHDMFAIMSGPTIAAISVVFDHAEHEDVYQTC
IDGFLAVAKISACHHLEDVLDDLVVSLCKFTTLLNPSSMEEPVLAFGDDTKARMATVTVFTIANRYGDYIRTGWRNILDC
ILRLHKLGLLPACVASDAADDQELSAETGHGKPLTNSLSSFQLPAIGTPRRSSGLMGRFSQLLSLETEEPRSQPTEQQLA
AHQRTLQTIQKCHIDSIFTESKFLQADSLLQLARALIWAAGRPQKGNSSPEDEDIAVFCLELLIAITLNNRDRIVLLWQG
VYDHITSIVQSTVMPCALVEKAVFGLLRICQRLLPYKENLADELLRSLQLVLKLEPRVSDAYCEQITQEVSRLVKANATH
IKSSMGWHTITYLLLITAPHPEASEAGFDALVFIMSDGAHLLPANYVLCNDAARQFAESRVGQAERSIRALDLMAGSVAC
LIQWAQESKRAAGEEEAAKMFQDIGEMWLRLVQGLRKVSLDHREDVRNHAVLSLQRCLTGVDGLQLSHGLWLQCFDLVIF
TMLDDLLEIAQGHSQKDYRNMEGTLVLAMKLLSKIFLQLLHNLSQLTTFCKLWLGVLSRMEKYMKVKVRGKRCEKLQDLV
PELLKNNLIVMKTRGILVQRSALGGDSLWELTWLHANNIAPSLQSEVFPDQDSEVSLRKWGDTAEDLVSDRMGSASSTET
HSSEGIHAGG*

**>Sanal_SaT03034**

MGCPNMPFGINAFDGGTKEHTGKPSRGAIACMINSEITAVLAVMRRNVRWGLRYMEDDDQLEHSLIQSLKELRKKIFSWQ
HQWHTINPIMYLQPFLDVIQSDETGAPITGVALSSVNKILNLDLLDLQTVNVQDAMHLTVDAVTSCRFEVTDPGSEEVVL
MKILQVLLACMKSNASVKLSNQHVCNIVNTCFRIVHQASGKGELLQRIARQTTHELVRCIFLHLHDIDSSEHALANGGMS
YVSKEVDAADGGHSIGNKQLENGNANSEHHGQSSVGFVSNASSVTLGGKACENNIEVNDEMEPGQNGRHVMMEPYGIPCM
LEIFQFLCSLLNVIENIGIGPRSNPMAYDEDVPLFALGLINSAIELGGTTIKQHPKLLALIQEELFRNLMQFGLSPSPLI
LSMVCNIVLNLYHHLRTELKLQLEAFLSCVLLRVAQSRHGSSYQQQEVAMEALVDLCRQQAFATEMYANFDCDISCTNVF
EELASLLSKSAFPVNGPLSSMHVLALNGLTAMIQGMAERTEKVSKELPTAELGSIDFEGYKTIWMEKCESYGDPNYWIPF
VRKRKYFKRKLMIGADHFNQDPKKGLEFLQGMNLLPDKLEPSSVAYFFRYANGLDKNLIGDYLGNHDEFCIQVLHEFAGT
FDFQDMNLDTALRIFLETFRLPGESQKIQRVLEAFAERYYDQSPQILINKDAALLLSYSIILLNTDQHNAQVKKKMSEED
FIRNNRRINGGGDLPRELLSELYHSICENEIRLAPEQGAGSPVMTSNHWIGLPHRARKSSPYIICDTIEHLDCDMFAIFS
GPTIAAICVVFDNVEHENVLQTCTDGFLAIAKISACYHLHDVLDDLVVSLCKFTTLLSPLSVEDTILAIGDDTKVRLALT
TVFSIANRFGDYIRSGWKNILDCVLSLQKLGLLPARLASDAADELELSSESERSRPATSSLSTSHMQSVATPRRSSSLMG
RFSQLLSFDMEEPRSQPTDEEFAAHQRTLQTVQNCNIDSIFTESKFLQADSLLQLFRALVLAAGRPRKGYHSLEVEGTAV
FCLELLITVTVNNRDRIMLLWRDVYEHISNIVQSIVMPCTLVKKTVFGLLRICQRLLPYKENLTDELLKSLQLILKLDAR
VADAYCEHITQEVMRLVKSNVTHIRSHTGWRAIISLLSITARHPEASEAGFETLAFIMTDGVHLLPANYVLCMDAACEFA
ESWVGDVDRSVCALDLMAGSIVCLVRWSCETKEAVEEKDAMKLSEDIGEMWLKLVQGLKKVCLDRRIEVRNHALSMLQRS
LIGAEGISFPHDLWLRCFDSVIFGLLDDLLEIVRGNSPKDYRNMEGTVILALKLLSKVFLQFLIDLSRLPSFCRLWVGVL
NRIEECMKVKFRGRQSEKMHELVPELLKNTLLVMKSSGILVPSDATGGDSFWQLTWLHVKNIDPLLQSEVFPDHELEQSQ
AKDMSTGGGPQSDETLAVQSSETTA*

**>Sanal_SaT05437**

MENTVDEINNCLSCQEKKKDVAALKKKELGLSCMLNTEVGAVLAVIRRTPDANSQYIPSIEDLADSAILHSLKSLRVLIF
NPQQEWRMVDPSIYLSPFLDVIQSDDVPAAATALALSAILKIVKLNIFDDKTPGAKDAINSVVKGITNCRIQKTDSFMEE
GTLMRILQLLVELMRARASILLTDHAVCTVVNTSFQVVQRATRIGDLLQRTARHTLHELVQTIFKRLPDIKVKDGENADS
NAEDTNIGKNSDSDSGFGIPCIVDIFRFLCSLLNVVNVVETEGSTSHQSDEDVQVFALVLINSAIEMSGEGIGNHPKLLQ
MIQDDLFHHLIHYGTSSGSLVVSMICSTVLSIYHFLRRSIRLQLEAFFTYVLFRIAATGTTIQRQEVAVEGIINFCRQPT
FTVEAYINYDCDPICQNVFEEIMKFLCKHAFPAGTSLTSLQLQAFEGLITTIHAIADCIDKGGGEEEENNVNPMGPYLIE
IREYRPLWEEKSRDDLDLGTWVNFVRLRKAQKRKMMIAGDHFNRDEKKGFEYLKFSHLVSDPADPKCFAFFFRYTPRLNK
NMIGEYLGDPDEFHLQVLKEFTETFNFSGMILDTALRAFLKAFRLPGESQKIQRILEAFSDKFYDQESSQIFESKDTVLI
LCYSLIMLNTDQHNPQVKKKMTEDEFIRNNRSINGGKDLPRDFLSELFQSISNNAITLFSQSGTARGMNANQWIELMNRS
KVMHPFIVCNFDRQIGRDMFAAIAGPSVAVLCAIFEHADDDEILQECVEGLFSVARIAQFGLVDTLDEILASFCKFTTLL
NPYVTAEETLYVFSNDMKPRMAILAVFTIANNFENSIRGGWRNIVDCLLKLKRLKLLPQSVIEPDAAAITGPSSESLHSH

AGSESGIIFPNHDPTFGSSSSNHQPLGLMGRFSHFLAMENIMEDSLTLGGSEFEQNLKIIQQCRIGNIFTNTGGLPEEPL
LNLGRSLIFAAAGRGQKFSTPVEEEETVGFCWDLVTTITLANTHRFLTFWPSYHDYLLAVAQFPLFSPIPFAEKAIVGLF
KVCLKLLNSHHTDNRFSEELIFKSINLMWKLEKEILDTCCEFITQTVCKILIEYPSNLHGQLGWKSVLHLLSITGRHPEM
YELAVATLATFMSEGTHVSRTNYSYCIECAFGFVALKNSPLEKNLKILDLMSDSVNLLIQWYRSGCSDTGSNNFSGSSCS
SMEDSKGGFPNFTVHLFVKLGEALRKTSLARREEIRNHAIMSLRKSFTLAEELEFTSTHCISSFNLVIFAMVDDLHEKML
EYSRRDNSEREVRSMEGTLKIAMELLTDVFLQFLEPIAECPEFRTFWLGVLRRMDTCMKADLGPYGESKLVELVPFWLKS
MITTLKEKDILVQKEGEDLWEITYIQIQWIAPHLKEELFPEEEL*

**>Maole_005233**

MGRLKLQSGIKSIEEEPEDCDMAYSNKAALACMVNSEVGAVLAVMRRYVRWGGRYMSGDDKLEHSLIQSLKALRK
QIFSWQHQWHTINPVVYLQPFLDVIRSDETGAPITGVALSSVYKILTLDVLDTSTVNVEDAMHLVVDAVTSCRFE
VTDPASEEVVLMKILQVLLACMKSKASVVLSNQHVCTIVNTCFRVVHQAGTKGELLQRIARHTMHELVRCIFSHL
PDVDNTEGALINASSSIKQEIGGQQNDYTFGSKQLENGNVSYDGQQSSAIFSSSGLVDDNTIGAGSRKEAASYDL
HLMTEPYGVPCMVEIFHFLCSLLNVVEHIGMGPRSDAIEFDEDVPLFALGLINSAIELGGPSVRQHPRLLSLIQD
ELFRNLMQFGLSMSPLILSMVCSIVLNLYHHLRTELKLQLEAFFSCVVLRLSQSRYGASYQQQEVAMEALVDFCR
QKSFMVEMYANLDCDITCSNVFEDLANLLSKSAFPVNCPLSAMHILALDGLIAVIQGMAERIGNGSFGLEQAPVN
LEEYTPFWMVKCDNYTDPNHWVPFVRRRKYIKRRLMIGADHFNRDPKKGLEFLQGTHLLPDKLDPQSVACFFRYT
AGLDKNLVGDFLGNHDEFCVQVLHEFAGTFDFQDMNLDTALRLFLETFRLPGESQKIQRVLEAFSEKYYEQSQQI
LVNKDAALLLSYSLIMLNTDQHNVQVKKKMTEDDFIRNNRHINGGNDLPREFLAELYHSICKNEIRTTPEQGAGF
PEMTPSRWIDLMHKSKKTAQFIVADSRAYLDHDMFAIMSGPTIAAISVVFDHVEHEDVYQTCIDGFLAVAKISAC
HHLEDVLDDLVVSLCKFTTLLNPSSVEEPVLAFGDDTKARMSTVTVFTIANRYGDYIRTGWRNILDCILRLHKLG
LLPARVASDAADDSELSADTGQGKPLTNSLSSSHMPSIGTPRRSSGLMGRFSQLLSLETEEPRSQPTEQQLAAHQ
RTLQTIQKCHIDSIFTESKFLQSDSLLQLARALIWAAGRPAKGNSSPEDEDTAVFCLELLIAITLNNRDRIVLLW
QGVYEHIANIVQSTVMPCALVEKAVFGLLRICQRLLPYKENLADELLRSLQLVLKLDARVADAYCEQITQEVSRL
VKANATHIRSPMGWHTITYLLLITAQHPEASEAGFDALVFIMSDGAHLLPANYVLCIDAARQFAESRVGQAERSI
RALDLMAGSVACLAQWAQETRDAVGEEEAAKVSQDIGEMWLRLVQGLRKVCLDHREDVRNHALMSLQRCLTVVDG
LYLPHGLWLQCFDLVIFTMLDDLLEIAQGHSQKDYRNMEGTLILAMKLLSKVFVQLLHNLAQLTTFCKLWLGVLS
RMEKYMKVKVRGKRCEKLQELVPELLKNNLLVMKTKGILVQRSALGGDSLWELTWLHANNIAPSLQSEVFPDQDT
DQHKQQETGEMLVSDQMGSEKVALEVCLIQIFEFSDEVDERSPHL*

**>Maole_012409**

MGCMNLLSGVNAFDGGSKEHTGSPSSGAIACMINSEIGAVLAVMRRNVRWGIHYMADDGQLEHSLIQSFKDLRKQ
IFLLQHQWHTINPTLYLQPFLDVIQSDETGAPITGVALSSVYKILTLDILDLHTVNVEDAMHLIVDAVTSCRFEV
TDPASEEVVLMKILQVLLACMKSKASVKLSDQLVCNIVNTCFRIVHQASSKGELLQRIARHTMNELVRCIFSHLP
DIDSTEHAPANGGRSHVKKEFGMVDEARMVGSEQLENGNVSVEYDGQPTVEFASNASTVTLMGKAGENRIEVDEG
IESAQNGRHVMMEPYGIPCMVEIFHFLCSLLNVIEHIEIGPRSNPIAYDEDVPLFALSLINSAIEVAGASIRQHP
KLLTLIQNELFHNLMQFGFSMSPLILSMVCSIVLNLYHHLRTELKLQLEAFVSCVLLRLAQAKHGASYQQQEVAM
EALVDLCRQQTFTAEMYANFDCDLSRSNVFEDLASLLSKSAFPVNVPLSAMHILALNGVIAMIQGMAERAEMVGN
ELPNAERSIDVEGYKPFWTEKCENYSNPSYWIPFVRKMRYIKRKLMIGADHFNRDPKKGLEFLQGMNLLPDKLDA
QSVACFFRYSIGLDKNLVGDFLGNHDEFCIQVLHEFAGTFDFRDMNLDTALRIFLETFRLPGESQKIQRALEAFA
ERYYEQSPQILINKDAALLLSYSLILLNTDQHNVQVKKKMTEDDFIRNNRHINGGSDLPREFLSELYHSICENEI
RMAPGQAAGFPGMTSSHWIGLLHQSRKSTPYITCDCESIPHLDYDMFSILSGPTIAAISVVFDHAEHDDVLQTCV
DGFLAIAKISTCYHLYDVLDDLVVSLCKLTTLLSPLSFEDSLLAIGDDTKAMMATTTVFTIANRYGDYIRSGWKN
ILDCVLSLHKLGLLPARLASDAADGLEPSPDLELSRPATSSLSLSWRPSMATPRKSSGLMGRFSQLLSFDMEEPS
SQPTEEQLAVHQRTLQTIQSCHIDSIFTESKFLQADSLLQLSRALILAAGQPRKGNNSLEDEDVAVFCLELLIAI
TVNNRDRIMLLWEDVYEHVSNIVQSIIMPCTLVEKAVFGLLHICQRLLPYKENLTGELLKSLQLILKLDARVADA
YCEHITQEVLRLVKANAAHIQSHMGWRAIISLLSITARHPEASEAGFEALAFIMSDGAYLLPENYGLCMDAASEF
AESWVGHVDRSVCAMDLMAGSIDCLVRWSCETKEAVGGEAAAKLCQDIGEMWLRLVQGLKKVCLDRRVEVRNHAL
LMLQRCLIVGEGIHLQSDLILRCFDLVIFTLLDDLLEIVRGNSPKDYRNMEGTVVLAMKLLSKVFLQFLQDLSEL
PSFCKLWVGVLNRIEECTKAKFRGRKSEKIHELVPELLKNTLLVMKTSGVLSKTDAIGGDSFWQLTWLHVKNIAP
TLQMEIFPDHESEQLQMNIKTGGNPTPDGTGIVPSGETTA*

**>Maole_013225**

ACPEKKKDSAKLKRKKLGLSCILNTEVGAVLAVLRRIPELNSSYMYPSDDYTDPEIIHSLKSLRALIFNPHQEWR
MIDPSVYFSPFLEVIQGDNVQSSATAVALSALHKIFKLDIFDEKTPGAKEAVNSVVAGILNCRLEKCQPTTRDEI
LMRIIQLLNGLMHSRASTLLSDHSVCTIVTTYFQVVQRTAGRGNLLQREARYAMHELIQTIFSRLPEMEVKDEEN
SESDTEDVDVGSNLGSGYGVRCVVDIFHFLCSLLNVIVETDGSMAPTDEEDIQVFAMVLINSVIELSGDGIAKHP
ALLRMIQDDLFHHLIHYGTSSSPLILSMICSIVLNVYHSLRRYIRLQLEAFFTFVLFRIAVPGSLVQIQEIAVEG
IITFCRQPTFAIETYINYDCDPMCHNLFKEIGKLLCKYAFPTGIPLTSLQIQAFEGLVATIHNIADDVEKEEKSS
SFGPYPVDITEYRPIWEEKSKEDSDLGTWVDFVRTRKAQKKKMMIAGNHFNLDEKKGLEYLKISHLVSDPADPKV
LAFFFRYTPRLDKKMIGDYLGDPAEFNIQVLKEFTGTFDFSGMILDIALRNFLETFRLPGESQKIQRILEAFSER
FYDQQPTELFASKDAMFIFCYSLIMLNTDQHNPQVKKKMTEEDFIRNNRAINGGKDLPREYLSELFQSISNNAIT
LFTQSGLPTEMNPKRWIELINRSKVMHPFIMCDFDHLLGRDMFASIAGPSVATLSAVFEHADDEEILHECIEGLF
SVARIAKYGLEDILDEVVSLLCKFTTLLNPYAAAEEILYVFSNDMKPRMATLAVFTIANNFESSIRGGWRSIVDC
LLKLKRLKLLPQSVVEPNTTTTSSSDLHTHTRSESGIIFPNHDPKFGGKRQSSGMIGQFSHFLSMESMEDSLTLG
GNEFEQNLKIIQQCQIGNIFAKSCNLPDETLQNLGRSLIFAAAGKGQKFSTPIEEEETVGFCWDLITAITLANTH

RFSTFWPSYHDYLLAVAQFPLFSPVPFAEKAIVGFFKVCVKLLSLNQPNKFSEELIFKSINLMWKLEKEILDTCY
EFITQSVSKILTEHPKNLQSQLGWKSTLHLLSITGRHTETYDQAVETLIKLMSDGTYVSRANYPYYIDCAFGFVA
LKNSPLEKNMKILDLMSDTVNLLIQWSKCGFSDPGSSTSNVSSSSTEDSSRGFGLYNLIVNLFVKLGEALRKTSL
ARREEIRNHAILALQRSFTLAEELCFTSTNCINCFNLVIFAMVDDLHEKMLEYSRRENSEREMRSMEGTLKIAME
LLTDVFLQFLRPISGSPGFRTFWLGVLRRMDTCMKADLGEYGGSKLPELVPDWLRKMITTMKEKEILTQIEGEDL
WEITNIQIQWIAPSLKEELFPEEI*
**>Maole_014154**
DTTELKRKKLGLSCILNTEVGAVLAVLRRIPESNSPYMSPSEDYTDPEIVHSLKSLRALIFNPQQEWRMIDPSVY
FSPFLEVIQGDNVQSNATEVALSALHKIFKLDMFDEKTPGAKAAINSVVAGIMNCRLEKCQPTTRDEILMRILQL
LNGLLHSRASTLLNDKAICTIVTTYFQVVRRTASRGNLLQREAKYAVHELIQIIFSRLPEMEAKDGENSESDTED
ADIESNLGSGYGVRCLVDIFHFLCSLLNAIVETDGSVAPTADEDIQVFSMVLINSAIELSGDGIAKHPDLLRMIQ
NDLFHYLIHYGTIPGTLVQIQEIAVEGIITFCRQPTFTIETYINYDCDPMFQNLFEEIGKLLCKHAFPTGSPFTS
LQIQAFEGLVATIHSIADNIEKEEKSSFSGPYSVDITEYRPIWEEKSKEDCDLGTWVDFVRTRKVQKRKMMIAGN
HFNLDVKKGLEYLKASHLVSDPADPKVLAFFFRNTPRLDKNMIGDYLGDPAGFNIQVLKEFTETFDFSGMILDTA
LRTFLETFRLPGESQKIQRILEAFSDKFYVQQPSQLFVSKDAVFIFCYSLIMLNTDLHNPQVKKKMTEEEFIRNN
RAINGGKDLPREYLSEFFHSISNNAITLFSQSGLPIEMNPNKWIELINRSKVMHPFIACDFDHRLGRDMFASIAG
PSVAALSAIFEHADDEEILHECIEGLLSVARIARYGLEDTLDEVLSLLCKFTTLLNPHASAEEILYVFSNDMKPR
MAILAVFTIANTFECSIQGGWRNIVDCLLKLKRLKLLPQSIVEPNATTTSTSDLHTHTKSESGIIFPSHDPKFGD
NRCHHSSGIIGQFSHFLSMESMEDSLTQGGSELEQNLKTIQQCRIGNIFAKSCNLPDESLQNLGRSLIFAAAGKG
QKFSTPIEEEETIGFCWDLVTTIALANSHRFPTFWPSYHDYLLAVAQFPLFSSIPFAEKAIVSFFKVCIKLLTLN
QPNKFSEELIFKSINLMWKLEKEILDTCCEFITQSVSKILTEHPANLQSQLGWKSTLHLLSITGRHPETYDQAVE
TLITLMSDGTYVSQANYPYHVDCAFGFVALKSSPLEKATKILDLMSDSVNWLVQWNKSGYADPGSNMISNVSSSF
MEDNSRGFGLSNLTVSLFVKLGEALRKASLARREEIRNHAILALQRSFTLAEELCFTSTNCINCFNLVIFAMVDD
LHEKMLEYSRREDSEREIRSMEGTLKIAMELLTDVFLQFLRPISGSPGFRTFWLGVLRRMDTCMKADLGEYGDSE
LQALIPEWLRKMITTMKEKGILSQREGEDLWDITHIQIQWIAPSLKEELFPEEM*
**>EchplGNOMa_LG8**
MGWQRMQTPINSIEEEPEDMEISHSNKAALACMINSEISAVLAVMRRNVRWGSRYVAGDDQMEHSLIQSLKTLRK
QIFKWQPQWQSINPVLYLQPFLDVIRSDETGAPITGVALSSVYKILTLDVLDLNTVNIEDTMHLIVDSVTSCRFE
VTDPASEEVVLMTVLRVLLSCMKCKASVKISNQNVCTIVNTCFRIVHQSNMKSELLQRMARFTMHELVRCIFSHL
PNIDHSEKSLVTGISALQPEVAGLGPDSLINKIENGNSNSESNGEVSGALASTVENVVQTEGKDAVSYDLDLKTK
PYGIPCMVEIFHFLCSLLNVVEHPGLGPRGNTIAFDEDMPLFALSLITSAIELGGPAISKHPSLLSLIQDELFNN
LLQFGLSMSPLILSMVCSIVLNLYHHLRSELKLQIEAFFSCVILRLAQSRFGSSYQQQEVAMEALVDFCRQKTFM
LEMYANLDCDITCSNIFEELANLLSKSAFPVNTPLSMMHILALDGLIAVIQGMAERISNGSFDQEPSPLKLAEYT
PFWKVQCENYNDPNYWVPFIRRRKYIKRRLMIGADHFNRDPKKGLEFLQGTHLLPEKLDPQSVACFFRYTAGLDK
NLVGDFLGNHDEFCVQVLNEFAGTFDFQDMSLDTALRLFLETFRLPGESQKIQRVLEAFSERYYDQTPHIFANKD
AALLLSYSIIMLNTDQHNVQVKKKMTEEDFIRNNRHINGGNDLPREFLAELYHSICKNEIRTTPEQGPGFAEMNP
SRWIDLMHKSRKSAPYIVSDYRAYLDHDMFAIMSGPTIAAISVVFDHAENEDVYQTCRDGFLAIAKISACHHLED
VLDDLVVSLCKFTTLLSPSSVEEPVLAFGDDTKARMATVTVFTIANRYGDFIRTGWRNILDCILRLHKLGLLPAR
VASDAADDSDMSPDPAQGKPLTNSLSSSHLQASGTPRRSSGLMGRFSQLLSLDAEEPRSQPSDQQLAAHQRTLLT
IQKCRIDSIFTESKFLLSDSLLQLARALIWAAGRPQKGNSSLEDEDTAVFCLELLIAITINNRDRMELLWQGVYE
HIVNIIQSTVMPCALVEKAVFGLLRICQRLLPYKENIADELLRSLQLVLKLDARVADAYCEQITQEVSRLVKANA
SHVRSQMGWRTIASLLSITARHPEASEAGFDALLYIMSEGAHLSPTNYVICIDAARQFAESRVGQAERSIHAVDL
MVGSLKCLVQWSQKSKEAIADAETVKLSPDIGEMWLRLVQGLRKVCLDQREEVRIHALQSLRTCLLGVDGMLLPN
GLWFQCFDLVIFTMLDDLIEIAQAQKDYRNMEGTLVHTLKLLVEVFLQLLPELSQFTAFCKLWRGILSRMEKYMK
AKVRGKKSDKLQELVPELLENALLVMKAKGVLAPRSALGGDGLWELTWLHVNNIVPSLQPKIFPDQELESNQGDS
G*
**>EchplGNOMb_LG3-261**
MEYLNHRGSFTPSKEGSCTSKATLACMINSEIGAVLAVMRRNVRWGAHYGVDDDQLEHALIYSFKKLRKEIFTWH
QQRNLDPVVYLQPFLDVIQSDETSAPITGVALSSVYKMLTLDVIDVDSSNLSNPLHLIIDAVTSCRFEVTDPASE
EVVLMKILQVLLACMKCKASVALSNRHVCNMVNTCFRIVHQASSKGELLQRIARHTMHELVRCIFSHLPNIEELP
CPLTQSPEDAKKYSHAELSSKTSSNEVTMEVNSDSNVNEETTEIGNVNDNEADSMTDPYGIPCMVEIFHFLCSLL
NVMEHMEVGPRSNPIAYDEDVPLFALGLINSAIELGGSSFERHPKLLSILQGDLFQNLMRFGLSISPLILSTVCS
IVLNLYHNLRSKLKLQIEAFISGVLLRVAQSKHGASYQQKEVALEALVDFCRQHTFMTEMFANFDCDISRNNIFE
DVTNSLSKSTFPVNSPLSAMNTVALDGLIAMIQGMAERVSHDSSVPEQISLDLQRYSPFWTDKCSDYTDSSEWIP
FVHKMKLIKRKLAIGVDHLNRDLKKGLEYLQGIRLLPEKLDPQSVACLLRYTAGIDKTLIGDYLGSHDEFSIEVL
HEFAQTFDFSNMNLDTALRIFLETFRLPGESQKIQRVLEAFAERYYEQSPNILANKDAALLLSYSLIMLNTDQHN
SQVKKKMTLEDFIRNNRRINGGNDLPRDLLTELFHSICENEIRMTPDKGVVVQVITPGHWIGLVHRSKQVTPFIS
SGYVSYLDYDMFTILSGPTIAAISVVFDHVEQENVLQTCIDGFLAIAKVSASYNFDDVLDDLVVSLCKFTTLLSP
TYLEEFVLAFGDDAKAKMATEAVFTIANRYGDHIRSGWKNILDCILSMHKLGLLPARLATDAASDLDSSPDQDLG
KSASSSSAATHVLSANLSRKSSGLMGRFSQLLYFDREEPAPEPTEEDVAARQRALQAIRNCHIDSIFADSKFLQA
DSLLQLVRSLVAAADRSRKGMNSFEDEESAVFCLELLIVFALNNRDRIMLLWQCVYEHIANVVQSTIMPCALVEK
AIFGLIRICQRLLPYKENLTDELLKSLQLVLKLDARVADAYCEQITQEVMNLVKSNAMQIRSHTGWRTIISLLSI
TARHPEASEAGFETLSFVMSDGLHLLPANYVFCINAARQFAESRVGNVDRSVKSLDLMAGSLVCLNRWFLKTKEA

EGQEAADKMSQDIWEMWLRLVQGLRKVCLDQREEVRNHAILMLQHSLTGVEIHIPNDIWLQCFDMVIFTLLDDVL
EMSEQNTPKKNRNIEGTFVLSLKLLSKAFLQALQDLCQLPSFCKLWLKALSYMDSCMKLTFRGKRCEKIHELVPE
LLKNILLVMKTNGILVPNDAIREDSLWQQTWLHVKKISPSLQEEVFPSGELDLQKNDSQEGSDLLPDESALATPN
ETTA*
**>EchplGNOMc_LG3-262**
MEYLNHRGSFTPSKEGSCTSKATLACMINSEIGAVLAVMRRNVRWGAHYGVDDDQLEHALIYSFKKLRKEIFTWH
QQRNLDPVVYLQPFLDVIQSDETSAPITGVALSSVYKMLTLDVIDVDSSNLSNPLHLIIDAVTSCRFEVTDPASE
EVVLMKILQVLLACMKCKASVALSNRHVCNMVNTCFRIVHQASSKGELLQRIARHTMHELVRCIFSHLPNIEELP
CPLTQSPEDAKKYSHAELSSKTSSNEVTMEVNSDSNVNEETTEIGNVNDNEADSMTDPYGIPCMVEIFHFLCSLL
NVMEHMEVGPRSNPIAYDEDVPLFALGLINSAIELGGSSFERHPKLLSILQGDLFQNLMRFGLSISPLILSTVCS
IVLNLYHNLRSKLKLQIEAFISGVLLRVAQSKHGASYQQKEVALEALVDFCRQHTFMTEMFANFDCDISRNNIFE
DVTNSLSKSTFPVNSPLSAMNTVALDGLIAMIQGMAERVSHDSSVPEQISLDLQRYSPFWTDKCSDYTDSSEWIP
FVHKMKLIKRKLAIGVDHLNRDLKKGLEYLQGIRLLPEKLDPQSVACLLRYTAGIDKTLIGDYLGSHDEFSIEVL
HEFAQTFDFSNMNLDTALRIFLETFRLPGESQKIQRVLEAFAERYYEQSPNILANKDAALLLSYSLIMLNTDQHN
SQVKKKMTLEDFirnnrringgnDLPRDLLTELFHSICENEIRMTPDKGVVVQVITPGHWIGLVHRSKQVTPFIS
SGYVSYLDYDMFTILSGPTIAAISVVFDHVEQENVLQTCIDGFLAIAKVSASYNFddvlddlvvslCKFTTLLSP
TYLEEFVLAFGDDAKAKMATEAVFTIANRYGDHIRSGWKNILDCILSMHKLGLLPARLATDAASDLDSSPDQDLG
KSASSSSAATHVLSANLSRKSSGLMGRFSQLLYFDREEPAPEPTEEDVAARQRALQAIRNCHIDSIFADSKFLQA
DSLLQLVRSLVAAADRSRKGMNSFEDEESAVFCLELLIVFALNNRDRIMLLWQCVYEHIANVVQSTIMPCALVEK
AIFGLIRICQRLLPYKENLTDEllkslqlvlklDARVADAYCEQITQEVMNLVKSNAMQIRSHTGWRTIISLLSI
TARHPEASEAGFETLSFVMSDGLHLLPANYVFCINAARQFAESRVGNVDRSVKSLDLMAGSLVCLNRWFLKTKEA
EGQEAADKMSQDIWEMWLRLVQGLRKVCLDQREEVRNHAILMLQRSLTGVEIHIPNDIWLQCFDMVIFTLLDDVL
EMSEQNTPKKNRNIEGTFVLSLKLLSKAFLQALQDLCQLPSFCKLWLKALSYMDSCMKLTFRGKRCEKIHELVPE
LLKNILLVMKTNGILVPNDAIREDSLWQQTWLHVKKISPSLQEEVFPSGELDLQKNDSQEGSDLLPDESALATPN
ETIA*
**>EchplGNL2_LG4**
MARPDEPCLTKYRRRQLAISCMLNTEVGSVLAVIRRAPDINSQFYPYADDYCDPNILQSLKSLRAVIFNPQQEWR
TIDPSMYITPFLDVIQSDDVPAAATGVALAAFQKIMKFNVFDDKSPGAREIINLSVSAITGCRLEKTDHISEDAV
MMKILMSLAAIMRHKASYLLTDESVCTLVNVCFQVVQQSAGRGDLLQRTGRFIMHDMIQMIYSRLPEIEVHEWEN
EESDTEECDSDSRFGIRSAVDIFHFLCSLLNVVEVVNVEGLISQTSDDNVQLFALVLISSAIELSGDSIGKHPKL
LRMVQDDLFHHLIHYGTCSSPLILSMICSTVLNIYHIFRRSVRLQLESFFSFVLLKVASLSTTQLQEVALEGITN
FCRQPKFIIEVYVNYDCDPTLHSVFEEMGKILCKNAFPGGGALTSLQVQAFEGLVSIIHNIADNIDKDDDTSTSE
PYPVEITEYRPYWEEKSLDDFDFETWVDWVRVKMVQKRKLLIAGNHYNRDQKKGLDYLKFSKLISDPPDPKAYAM
FFRYAPKLDKTTIGEYLGDPEPFHLEVLREFTNTFEFNGMVLDSALRTYLESFRLPGESQKISRILEAFSERFFD
QQASETFVSKDAVMVLCYSVIMLNTDQHNPQVKKKMTEEEFLKNNRGINDGEDLPREYLSELFRSIATCAIKLKD
ESNAAPVEMNPNRWIQLINRSKIIKPFSLCNFERRLGRDMFASIAGPSVATLSAIFEQSDDEEILHECIEASFSV
ARIAQYGLEDTLDEIICSFCKFTTLLNPYASPEETLYAFSHDMKPRMATLSVFTIVNTFKSSIRGGWKTIIECLL
KLKRLKLLPQSLIEFDPTSEESATHGEDNEDGGGFAPHDSKFPKQRRSSGIIGRFSNFLSMDNVEDSLSLGMTEF
EQNFKIIQQCHIGSIFSGSSGLSDETILNLGRSLISAAAGKGQKFSTAIEEEETVAFCWDLIYILASSNIHRFVN
FWPQYHDFIIDAAQFPMFSPIPFAEKAIMALMKICVRHLSLDQPDKITEELIFKSINLMWKLDKEILDSCCEFLM
QSISKILTDYPSKLQTHLGWKTIMHLLSVTGRAPENYDQGVNSLISLMSEGTHIMKTNYAYCIDCAFGFVALKNS
PLDKNLKLMDLMAESVNLLVQWCKAAYSDPGSSVSMSSNASNSSQDEQFKAINYANFTMNLFIKLGEALRKTSLA
RREEIRNHAITSLHKSFTLAEELCFTPSNCINCFNLVIFAMVDDLHEKMLEYSRRENAERETRSMEGTLKLAMEA
LTEVYLQFLRPIAESPNYRQFWLGILRRMDTCMKADLGEYGESKMKTVIPELLKKIILTMKEKEILVPREGDDLW
EITYIQIQWIAPSVKEELFPDV*
**>CRO_018704**
MGRLRLQTSIKAIEEEPEDCDTTSSSKAALACMINSEIGAVLAVMRRNVRWGGRYVSGDDQLEHSLIQSLKTLRK
QIFLWQNQWQTVNPVLYLQPFLEVIRSDETGAPITSVALSSMYKILTLDVLDLNTVNIEDGMNLVVDAVTSCRFE
LTDPASEEVVLMKILQVLLACMKSKASVVLSNQHVCTIVNTCFRVVHQAGTKGELLQRIARHTMHELVRCIFSHL
PDIDNTEWSLVKGVNSVKHEVSGIDTDYSLSNKSENGSGISDYNGQPSSGAHASTGLLSSGMDDGTAKTDNGKDA
VPYDLHLMTEPYGVPCMVEIFHFLCSLLNVVENMGMGPRSNSIAFDEDVPLFALGLINSAIELGGPAIRYHPRLL
SLVQDELFRNLMQFGLSMSPLILSMVCSIVLNLYQHLRTELKLQHEAFFACVILRLAQSRHGGSYQQQEVAMEAL
VDFCRQKTFMVEMYANLDCDITCGNVFEELANLLSKSAFPVNSPLTSMHILALDGLIAVIQGMAERIGNGSSNSE
HVPVNLVEYTPFWMLKCENYSDPDHWVPFVRRRKYIKRRLMIGADHFNRDPKKGLEFLQGTHLLPEKLDPQSVAC
FFRYTAGLDKNLVGDFLGNHDEFCVQVLHEFAKTFDFQDMNLDTALRLFLETFRLPGESQKIQRVLEAFSERYYE
QSPQILANKDAALLLSYSLIMLNTDQHNVQVKKKMTEEDFIRNNRHINGGNDLPREFLSELYHSICKNEIRTIPE
QGAGFAEMTPSRWIDLMHKSKKSAPYIVSDSRAYLDHDMFAIMSGPTIAAISVVFDHAEHEDVYQTCIDGFLAVA
KISACHHLEDVLDDLVVSLCKFTTLLNPSSVEEPVLAFGDDCKARMATVTVFTIANRYGDFIRTGWRNILDCILR
LHKLGLLPARVASDAADDTDMSSEPGHGKPLTNSLSSVQMQSIGTPRRSSGLMGRFSQLLSLDTEEPRSQPTEQQ
LAAHQRTLQTIQKCHIDSIFTESKFLLAESLLQLARALIWAAGRPQRGSSSPEDEDTAVFCLELLIAITLNNRDR
IGLLWQGVYEHIANIVQSTVMPCALVEKAVFGLLRICQRLLPYKENLADELLRSLQLVLKLDARVADAYCEQITQ
EVSRLVKANATHIRTQLGWRTIASLLSITARHPEASEAGFDALLFIMSDGAHLSQANYVLCIDAARQFAESRIGQ
ADRSVRAVDLMACSVACLARWVEDTKSGTGEAEPLKLSQDMGEMWLRLAQALRKVCLDQREEVRNHALSSLQMCL

TGVDGIHLPHVLWLQCFDMVIFTMLDDLIEIAQGHSQKDYRNMEGTLVLALKLLIKVFLQLLPELSQLTTFCKLW
LGVLSRMEKYVKVKVRGKRSEKLQALVPELLKNTLLVMKTKGVLVQRSALGGDSLWELTWLHVNNIVPSLQSELF
PDKDADVSEINQEIGASPTSNETDPA*

**>CRO_021518**

MGCLNQQTAVNDLAGEPKGGCFRSSRVALACMVNSEIGAVLAVMRRNVRWGVRYADDDLVEHSLIHSFKELRKKI
FSWKHQWHVIDPIVYLKPFLDVIQSDETGAPITGVALSSVYKILTLEILDSDTVNVENALHQIVDAVTSCRFEVT
DPASEQVVLMKILFLLACVKKKHPVHLSNHHVCNIVNTCFRIVHQASSKGELLQRIARHTMHELIRCIFSHLPDI
DDKRPSLAEGSRSADKEVFVPTDNQSFIDKQSISSNGKAKSDDQVPSVFVASGISTLTFPGKVDEEATSFAQKEV
VENGGSSMMDPYGVPCMVEIFHFLCSLLNVMEHIETGPRSNPIAYDEDVPLFALGLINTAIELGGASFGCHPKLL
ALIQEELFYNLMQFGLSMSPLILSTVCSIVLNLYHHLRNKLKLQLEAFFSGVLLRVAGSKHGSSYQLQEVAMEAL
VDFCRQHFFMSEMYVNFDCDISCSNVFEDLANLLSKSAFPVNSPLSAMNTLALDGLIAMIQGIAERIGHESSPMD
RAALDLLEYEPFWTARCENHNDPSNLVPFAHKMKRIKRKLTIGVDHFNRDPKKGLEYLQGVHLLPDKNDPQSVAC
FFRYTAGLDKNLVGDFLGSHDEFCIQVLHEFSKTFDFRDMNLDTALRIFLETFRLPGESQKIQRVLEAFAERYYE
QAPDILVNKDAALLLSYSLIMLNTDQHNAQVKKKMTEEDFIRNNRRINGGDDLPREFLSELYYSICENEIRMVPE
QAVGIGTMMRSHWIGLLHKSKQSAAYVFCDSGPHLDYEMFAILSGPTIAAISVVFDHVDQDEILQSCVDGFLDIA
KISASYNLIDVLDDLVVSLCKFTNLLLPPFVDEAFILAFGDDSKARMATTAVFTIANRYGDHIRSGWKNIVDCIL
SLNKMGLLPARLASDAADDLESSSDQNLVRPASSSPSAAAVPMPPTRKPSGLMGRFSQLLYLDTEEPAPEPTEDQ
LAARQRTVQTIQDCHLDSIFAESKFMQAESLTQLVQALLFAAVRPRKVNNLVEDEVTAVFCLDLLITITLNNRDR
IMLLWQGVYEHIANIVQSTVMPCALVEKAVFGLLRICQRLLPYKENLTDELLKSLQLILKLDARVADAYCEQITQ
EVMHLVKANAMQIRSHMGWRTIISLLSITARHPEASEAGFETLSFIMTDGAHLLPANYVLCVSAARQFAESRVGS
IDRSIISLDLMAGSLDCLIRWFYKTKEAVEEEAGLKMAQDIWEMWLRLGQGLRKSLCGPERRGQKSCYSDVAEVL
NWSGDDAVPSELWSHCFDLVVFTLLDDLLEITLQQSPKDYRNMEGTLVLSLKLLSKVFLQSLQDISQSASFSKLW
LGVLGCMERYMKVKFRGKRSEKIHELVPELLKNTLLVMKSSGILVPADAVTADSFWQLTWLHVKNIAPSLQLEVF
SSDELEQLQKQHSKSGNNPIQDGMLVPASETAA*

**>CRO_013169**

MDINQQEDETKKTKNKRKELGLSCMLNTEVGAVLAVIRRAPDSQFYHSPEDHFDCSVLQSLKSLRTIIFNPQQEW
RFIDPSVYLSPFLDVIQSDDVPASATGVALSSILKILRLEIFDHKTPCAGEIMNAIVTAITGCRLEKTDPVTEDA
VMMRILQVLVAVMKHRAHILLTDHSVCTVVNTCFQVVQQSANRGDLLQRSARHTMQELIQIIYYRLPEIEVGDCE
DSESETEDSHLDSGFGIYSAVDIFQFLCSLLNVVHVGPEIEGYPSQTPDLDVQMFALVLINSAIELSGDSIGKHP
KLLRMIQDDLCHHLIHYGSHCNPLVLSMICSTILNIYHFLRRSVRLQLEAFFSIVLLRAANLTNSPQLQEVATEA
IVNFCRQTTFAVEVYVNYDCSPIFMNVFEETAKLICKNAFPAGGTLTSLQLQAFEGLLITIHNIADKVDKEDDST
PSGPYPLEITDYVPFWEEKSKDDDALDDWINFLRLKKAQKRKVLIAGNHFSQDDKKGLEYLKISRLISDPPDPKA
YAIFFRFTPGLDRTMIGDYLGDPDEFHIQVLKEFTDTFEFGGMLLDSALRCYLESFRLPGEAQKIHRILEAFSER
FYDQQSSELFVSKDSVFILCYSLIMLNTDQHNPQVKKKMTEEEFIKNNRSINGGNDLPREYLSELFNSIATTAIA
INSQSGIRVEMNPNRWVQLINRAKIMKPFVLCDFDRRLGRDMFAAIAGPALATVAAIFEHCDEEEIIQECIEVLF
SIARIAQYQLEDTLDELICLFCKFTTLLNPYASTDETLYAFSNDMKPRMATLAVFTIANGFKDTIRGSWRNIIDC
LLKLKKLKLLPQSVIDPDASDLSADCSGAILSKQDSKVGDKKQGTGLIGRFSQFLSMESVEESLNLGLSEFEQNM
RIIQQCRIETIFSKSSNLPEESLWNLGRTLIFAAAGKGQKFSTPIEEEDTVGFCWDLIVTISYANTHRFSSFWPQ
YHEYLLAVAQFPLFSPIPFAEKAIVAQFKICLKLLSTSQSDKNGEELIFKSINLMWKLEKEILDTCSDFLIQSIC
KILTEYPANMSSSLGWKTVLHLLSITGRHPESYDQGVESLISLMSDASVISRTNYAFCIDCAFTFIALKNSPVEK
NIKIMDLMSESVKSLVQWYRTGYAEADSNVSVASNASSSSTEDNAKAVSSSSFVMNLFLKVGEAFRKTCLARREE
IRNHAVLSIQRSFMFAEELCFTPANCIYCFNSLIFAMVDDIHEKMLEYSRRDNAGRETRSMEGSLKIAMEVLTEV
FMQLLKPLSESPHFKAFWMGLLRRMDTCVKADLGEYGKTTMEDVVADLLKKLISSMKEKEMLVPKEGDNLWEITY
VQIQWIAPSLKEELFPDT*

**>Cofar_LOC113716666**

MGRLRLQSNINAIEEEPEDCETTSSNKAALACMVNSEIGAVLAVMRRNVRWGGRYVSGDDQLEHSLIQSLKTLRK
QIFVWQHQWQTINPVLYLQPFLEVIRSDETGAPITGVALSSVYKILTLDVLDLSTVNIEEAMNLVVDAVTSCRFE
ITDPASEEVVLMKILQVLLACMKGKTSVVLSNQHVCTIVNTCFRVVHQAGTKGELLQRIARHTMHELIRCIFSHL
PDVDNTECSLVKGGTPIKHEVGGIDADYSFGSKSENGSEYNGQLGNVASDISTGLNVMGNSAGKTDSGKDSVPYD
SHLMTEPYGVPCMVEIFHFLCSLLNVVEHMGMGPKANSIAFDEDVPLFALGLINSAIELGGPAITHHPRLLSLVE
DELFRNLMQFGLSTSSLILSMVCSIVLNLYQHLRTELKLQLEAFFSCVILRLSQSRYGASYQQQEVAMEALVDFC
RQKTFMVEMYANLDCDITCGNVFEELANLLSKSAFPVNSPLSAMHILALDGLIAVIQGMAERIGNGSFNSEPAPV
NLEEYTPFWMLKCENYNEPDKWVPFVRRRKYIKRRLMIGADHFNRDPKKGLEFLQGTHLLPEKLDPQSVACFFRY
TAGLDKNLVGDFLGNHDEFCVQVLHEFAKTFDFQDMNLDIALRLFLETFRLPGESQKIQRVLEAFSERYYEQSPQ
ILANKDAALLLSYSLIMLNTDQHNVQVKKKMTEEDFIRNNRHINGGNDLPREFLSELYHSICKNEIRTTPEQGAG
FAEMTPSRWIDLMHKSKKSAPYIVSDSRAYLDHDMFAIMSGPTIAAISVVFDHAEHEEVYQTCIDGFLAVAKISA
CHHLEDVLDDLVVSLCKFTTLLNPSSVEEPVLAFGDDCKARMATVTVFTIANRYGDFIRTGWRNILDCILRLHKL
GLLPARVASDAADDSEMSPEPGQGKPLTNSLSSVQVQAIGTPRRSSGLMGRFSQLLSLDTEEPRSQPTEQQLAAH
QRTLQTIQKCHIDSIFTESKFLLAESLLQLARALIWAAGRPQKGSSSPEDEDTAVFCLELLIAITLNNRDRIGLL
WQGVYEHIANIVQSTVMPSALVEKAVFGLLRICQRLLPYKENLADELLRSLQLVLKLDARVADAYCEQITQEVSR
LVKANATHIRSQLGWRTIASLLSITARHPEASESGFDALLFIMSDGAHLTPANYVLCIDAARQFAESRVGQADRS
VRAIDLMAGSVACLTRWDEDAKGAMAEAEALKLSQEIGEMWLRLAQALRKVCLDQREEVRNHALLSLQTCLTGVE
GIHLPHALWLQCFDMVIFTMLDDLTEIAQGHSQKDYRNMEGTLVLALKLLTKVFLLLLHELSQLTTFCKLWLGVL

SRMEKYMKVKVRGKKSEKLQELVPELLKSTLLVMKTKGVLVQRSALGGDSLWELTWLHVNNIVPSLQSELFPGNE
LEQSDSKLGEMGKNTLPNETGSA*
**>Cofar_LOC113718983**
MGRLRLQSNINAIEEEPEDCETTSSNKAALACMVNSEIGAVLAVMRRNVRWGGRYVSGDDQLEHSLIQSLKTLRK
QIFVWQHQWQTINPVLYLQPFLEVIRSDETGAPITGVALSSVYKILTLDVLDLSTVNIEEAMNLVVDAVTSCRFE
ITDPASEEVVLMKILQVLLACMKGKTSVVLSNQHVCTIVNTCFRVVHQAGTKGELLQRIARHTMHELIRCIFSHL
PDVDNTECSLVKGGTPIKHEVGGIDADYSFGSKSENGSEYNGQLGNVASDISTGLNVMGNSAGKTDSGKDSVPYD
SHLMTEPYGVPCMVEIFHFLCSLLNVVEHMGMGPKANSIAFDEDVPLFALGLINSAIELGGPAITHHPRLLSLVE
DELFRNLMQFGLSTSSLILSMVCSIVLNLYQHLRTELKLQLEAFFSCVILRLSQSRYGASYQQQEVAMEALVDFC
RQKTFMVEMYANLDCDITCGNVFEELANLLSKSAFPVNSPLSAMHILALDGLIAVIQGMAERIGNGLFNSEPAPV
NLEEYTPFWMLKCENYNEPDKWVPFVRRRKYIKRRLMIGADHFNRDPKKGLEFLQGTHLLPEKLDPQSVACFFRY
TAGLDKNLVGDFLGNHDEFCVQVLHEFAKTFDFQDMNLDIALRLFLETFRLPGESQKIQRVLEAFSERYYEQSPQ
ILANKDAALLLSYSLIMLNTDQHNVQVKKKMTEEDFIRNNRHINGGNDLPREFLSELYHSICKNEIRTTPEQGAG
FAEMTPSRWIDLMHKSKKSAPYIVSDSRAYLDHDMFAIMSGPTIAAISVVFDHAEHEEVYQTCIDGFLAVAKISA
CHHLEDVLDDLVVSLCKFTTLLNPSSVEEPVLAFGDDCKARMATVTVFTIANRYGDFIRTGWRNILDCILRLHKL
GLLPARVASDAADDSEMSPEPGQGKPLTNSLSSVQVQAIGTPRRSSGLMGRFSQLLSLDTEEPRSQPTEQQLAAH
QRTLQTIQKCHIDSIFTESKFLLAESLLQLARALIWAAGRPQKGSSSPEDEDTAVFCLELLIAITLNNRDRIGLL
WQGVYEHIANIVQSTVMPSALVEKAVFGLLRICQRLLPYKENLADELLRSLQLVLKLDARVADAYCEQITQEVSR
LVKANATHIRSQLGWRTIASLLSITARHPEASESGFDALLFIMSDGAHLTPANYVLCIDAARQFAESRVGQADRS
VRAIDLMAGSVACLTRWDEDAKGAMAEAEALKLSQEIGEMWLRLAQALRKVCLDQREEVRNHALLSVQMCLTGVE
GIHLPHALWLQCFDMVIFTMLDDLTEIAQGHSQKDYRNMEGTLVLALKLLTKVFLLLLHELSQLTTFCKLWLGVL
SRMEKYMKVKVRGKKSEKLQELVPELLKNTLLVMKTKGVLVQRSALGGDSLWELTWLHVNNIVPSLQSELFPGNE
LEQSDSKLGEMGKNTLPNETGSAQEEWHP*
**>Cofar_LOC113697893**
MGCLNQQTAVNAVVRGPKDGSFRSSRVALACMVNSEIGAVLAVMRRNVRWGVLYADDDQVEHSLIQSFKELRKKI
FLWQHQWHSIDPVVYLKPFLDVIQSDETGAPITGVALSSVYKILTLEILDSETVNVDKALHLIVDAVTSCRFEVT
DPASEEVALMKILQVLLACMKNKASVHLSNHHVCNIVNTCFRIVHQASSKGELLQRIARHTMHELVRCIFSHLPD
IGDRPHALAEGSRMSAETEVHVPKDTHILEGTQGVNANGSAKFDGKALPLKEASGNPAFTSASKVDEEIMTLGSV
EVAQNGADSMMDPYGIPCMVEIFQFLCSLLNVMEIESGVRSNPIAYDEDVPLFALGLINTAIELAGASFGDHAKL
LVLIQEDLFYNLMQFGLSMSPLILSTVCSIVLNLYHHLRTKLKLQLEAFFSRVLLKIAGSKHGSSYQQQEVAMEA
LVDFCRQPFFMAEMYANFDCDISCSNVFENLANLLSKSAFPVNSPLSALNTLALDGLIAMIEGISERIGHESSAP
ERSPIQPEEYRPFWTVTCDNYGEPNCWVPFVYKKKQIKKKLMIGADHFNLDPKKGLEFLKGVNLLPDTRDTRSVA
CFFRYTTGLNKNLVGEFLGSHDEFCIQVLKEFAQSFDFQDMNLDTALRIFLETFRLPGESQKIQRVLEAFAERYY
EQSPDILVNKDAVFVLSYSLIILNTDQHNSQVKKKMTEEDFLRNNRAINGGSDLPREFLSQLYHSICENEIRMIP
EQAAAASVMSRSHWISLVHRAKQSAPYIFCDSGPHLDYDMFAILSGPTIAAISVVLDHVERDNVLHTYIDGYLTM
AKISASYNSVDSLDDLVVSLCRFTNLLIPSLNNEEFILAFGDDAKARMATVAVFTIANRYGDQIRSGWKNILDCI
LSLNKLGLLPARLASDAVDDSEVSSDQNLVRPPSSSPTTSTPSPVAPSRKSSGLMGRFTQLLYLDTEEPESYPTQ
EQVAARQRSLQTIQDSHIDSIFAESKFLQAESLSQLVQALLLAAGRPRKANNSMEVEVTAVFCLELLIAITLNNR
DRIMLLWQGVYEHIASVVQSTVMPCALVEKAVFGLLRICQRLLPYKENLTDELLKSLQLILKLDARVADAYCEQI
TQEVMHIVKANAMQIRSHMGWRTIISLLSITARHPEASEAGFETLSFIMSEGAHLSPANYVLCLTAARQFAESRV
GNVDQSIRSLDLMAGSLDCLVRWFYKTKEAMGEEAALKMAQDIWEMWLRLVQGLRKVCVDQREEVRNHAILMLQR
CLTGAEVMHIPVGLWSQCFDLVIFTLLDDLVEIAQQHSAKDYRNMEGTLVLSLKLLSKVFLQLLHDVSQLASFSE
LWSRVLSCMERYMKVKFRGKRSEKIHELVPELLKNTLLIMKSSGLLVPSEEDNFWQLTWSSVKNLAPSLQLEVFS
SDELEQLQQKTQTNGCSPIQDGNVIVPPSETRA*
**>Cofar_LOC113692407**
MEDQQEGGNNNNMTKIKRKELGLSCMLNAEVGAVLAVIRRIPEANPQFHHPSEENYDSTILQSLKSLRSLIFNAE
QEWRKIDPSLYLSPFLDVIQSDDIPAAATGVALSSILKILRLDIFDEKTPGAKDVMDATVTAITACRLEKTDPVS
EEAVMMRILHVLTAVMRHKASVLLTDHAVCTVVNTCFQVVQQSAHRGNLLQRSARYTMQELIQIIYYRLPDIEVN
DWENSESDTEDSNLDSGFGIRSAVDIFQFLCSLLNVVEIAESEVLHAHTTDGDVQLFALALINSAIELSGDSIGK
HPKLLRMVQDDLFHHLIYYGTYSSPLVLSMICSTVLNIYHFLRRSVRLQLEAFFSFVLFRAANLGSSPQLQELAT
EAIVNFCRQSTFIIEVYVNYDCDPIFGNAFEEIGRLLCKHAFPTGGPLTSLQVQAFEGLVLVIHNIAENIDKEDD
TCPSGPYPVEITEYTPFWEEISKDDGLEDWVTFVRVRKAQKKKLLIAGNHFNQDDKKGLEYLKISHLIPDPPDPK
AFALFFRFTPRLDKTMIGDYLGDPDEFHIQVLNEFTNTFVFTGMILDSALRNYLETFRLPGESQKIHRILEAFSE
RFYDQQSSELFVSKDAVFVLCYSLIMLNTDQHNPQVKKKMTEEEFIRNNREINGGSDLPREYLSELFHSISSKAI
TLFGQSGTPAEMNPSRWIQLVNRAKLMKPYVLCNFDRRLGRDMFAAIAGPSVATLAAIFEQTEEEEILHECIELL
FAIARIAQYELEDNLDELICSFCKFTTLLNPYASAEETLYAFSNDMKPRMATLAVFTIANTFKNSLRGSWRTIVE
CLLKLRRLKLLPQSVVEFEIASDASIGTKHDKNESAASISTRNTKFNHIRRSSGLIGRFSHFLSLESVEESLNLG
VSEFEQNQNIIQQCHIENIFTTSSSLPDESLLNLGRALIYASAGKGQKFSTPVEEEETVAFCWDLIVIISCVNMH
RFSTIWPHYHDYLLGVAQFPLFSPIPFAEKAIVAQMKICLKILSAYQLDKVSEELIFKSINLMWKLEKEILDTCS
EFLVKSVSRILNEYPANLQSSLGWKTVLHLLSVTGRHPETYDQGVESLINLMSDGMHISRTNYASCIECAFGFVA
LKNSPVEKNIKIMDLMSDSVKLLVQWFRSGYSDPGSNVSTGSNASASSTEDNARSGSSASFIMNLFVKLGEALRR
TCLARREEVRNHAVISLQRSFILAEELCFTPANCIYCFNSLIFAMVDDIHEKMLEYSRRENAERETRSMEGSLKR

AMEVLVEVYSQLLKPISESPHFRPFWMGLLRRMDTCMKVDLGGYGKSDMQEVIPDLLKRIISIMKENELLIPKEG
DDLWEITYVQIQWIAPSLKDELFPGAP*
**>Cofar_LOC113695788**
MEDQQEGENNNNMTKIKRKELGLSCMLNAEVGAVLAVIRRIPEANPQFHHPSEENYDSTILQSLKSLRSLIFNTE
QEWRKIDPSLYLSPFLDVIQSDDIPAAATGVALSSILKILRLDIFDEKTPGAKDVMDATVTAITACRLEKTDPVS
EEAVMMRILHVLTAVMRHKASVLLTDHAVCTVVNTCFQVVQQSAHRGNLLQRSARYTMQELIQIIYYRLPDIEVN
DWENSESDTEDSNLDSGFGIRSAVDIFQFLCSLLNVVDIAESEVLHAQTTDGDVQLFALALINSAIELSGDSIGK
HPKLLRMVQDDLFHHLIYYGTYSSPLVLSMICSTVLNIYHFLRRSVRPQLEAFFSFVLFRAANLGSSPQLQELAT
EAIVNFCRQSAFINEVYVNYDCDPIFGNAFEEIGRLLCKHAFPTGGPLTSLQVQAFEGLVLVIHNIAENVDKEDD
TSPSGPYPVEITEYTPFWEEISKDDGLEDWVTFVRVRKAQKKKLLIAGNHFNQDDKKGLEYLKISHLIPDPPDPK
AFALFFRFTPRLDKTLIGDYLGDPDEFHIQVLNEFTNTFVFTGMILDSALRNYLETFRLPGESQKIHRILEAFSE
RFYDQQSSELFVSKDAVFVLCYSLIMLNTDQHNPQVKKKMTEEEFIRNNREINGGSDLPREYLSELFHSISSKAI
TLFGQSGTPVEMNPSRWIQLVNRAKLMKPYVLCNFDRRLGRDMFAAIAGPSVATLAAIFEQTEEEEILHECIELL
FAIARIAQYELEDNLDELICSFCKFTTLLNPYASAEETLYAFSNDMKPRMATLAVFTIANTFKNSLRGSWRTIVE
CLLKLRRLKLLPQSVVEFEIASDASIGTKHDKNESAASISTRNTKFNHIRRSSGLIGRFSHFLSLESVEESLNLG
VSEFEQNQNIIQQCHIENIFTTSSSLPDESLLNLGRALIYASAGKGQKFSTPVEEEETVAFCWDLIVIISCVNMH
RFSTFWPHYHDYLLGVAQFPLFSPIPFAEKAIVAQMKICLKILSAYQLDKVSEELIFKSINLMWKLEKEILDTCS
EFLVKSVSRILNEYPANLQSSLGWKTVLHLLSVTGRHPETYDQGVESLINLMSDGMHISRTNYASCIECAFGFVA
LKNSPVEKNIKIMDLMSDSVKLLVQWFRSGYSDPGSNVSTGSNASTSSTEDTARSGSSASFIMNLFVKLGEALRR
TCLARREEVRNHAVISLQRSFILAEELCFTPANCIYCFNSLIFAMVDDIHEKMLEYSRRENAERETRSMEGSLKR
AMEVLVEVYSQLLKPISESPHFRPFWMGLLRRMDTCMKADLGGYGKSDMQEVIPDLLKRIISIMKENELLIPKEG
DDLWEITYVQIQWIAPSLKDELFPGAP*
**>Cofca_T00032293001**
MGRLRLQSNINAIEEEPEDCETTSSNKAALACMVNSEIGAVLAVMRRNVRWGGRYVSGDDQLEHSLIQSLKTLRK
QIFVWQHQWQTINPVLYLQPFLEVIRSDETGAPITGVALSSVYKILTLDVLDLSTVNIEEAMNLVVDAVTSCRFE
ITDPASEEVVLMKILQVLLACMKGKTSVVLSNQHVCTIVNTCFRVVHQAGTKGELLQRIARHTMHELIRCIFSHL
PDVDNTECSLVKGGTPIKHEVGGIDADYSFGSKSENGSEYNGQLGNVASDISTGLNVMGNSAGKTDSGKDSVPYD
SHLMTEPYGVPCMVEIFHFLCSLLNVVEHMGMGPKANSIAFDEDVPLFALGLINSAIELGGPAITHHPRLLSLVE
DELFRNLMQFGLSTSSLILSMVCSIVLNLYQHLRTELKLQLEAFFSCVILRLSQSRYGASYQQQEVAMEALVDFC
RQKTFMVEMYANLDCDITCGNVFEELANLLSKSAFPVNSPLSAMHILALDGLIAVIQGMAERIGNGSFNSEPAPV
NLEEYTPFWMLKCENYNEPDKWVPFVRRRKYIKRRLMIGADHFNRDPKKGLEFLQGTHLLPEKLDPQSVACFFRY
TAGLDKNLVGDFLGNHDEFCVQVLHEFAKTFDFQDMNLDIALRLFLETFRLPGESQKIQRVLEAFSERYYEQSPQ
ILANKDAALLLSYSLIMLNTDQHNVQVKKKMTEEDFIRNNRHINGGNDLPREFLSELYHSICKNEIRTTPEQGAG
FAEMTPSRWIDLMHKSKKSAPYIVSDSRAYLDHDMFAIMSGPTIAAISVVFDHAEHEEVYQTCIDGFLAVAKISA
CHHLEDVLDDLVVSLCKFTTLLNPSSVEEPVLAFGDDCKARMATVTVFTIANRYGDFIRTGWRNILDCILRLHKL
GLLPARVASDAADDSEMSPEPGQGKPLTNSLSSVQVQAIGTPRRSSGLMGRFSQLLSLDTEEPRSQPTEQQLAAH
QRTLQTIQKCHIDSIFTESKFLLAESLLQLARALIWAAGRPQKGSSSPEDEDTAVFCLELLIAITLNNRDRIGLL
WQGVYEHIANIVQSTVMPSALVEKAVFGLLRICQRLLPYKENLADELLRSLQLVLKLDARVADAYCEQITQEVSR
LVKANATHIRSQLGWRTIASLLSITARHPEASESGFDALLFIMSDGAHLTPANYVLCIDAARQFAESRVGQADRS
VRAIDLMAGSVACLTRWDEDAKGAMAEAEALKLSQEIGEMWLRLAQALRKVCLDQREEVRNHALLSLQTCLTGVE
GIHLPHALWLQCFDMVIFTMLDDLTEIAQGHSQKDYRNMEGTLVLALKLLTKVFLLLLHELSQLTTFCKLWLGVL
SRMEKYMKVKVRGKKSEKLQELVPELLKSTLLVMKTKGVLVQRSALGGDSLWELTWLHVNNIVPSLQSELFPGNE
LEQSDSKLGEMGKNTLPNETGSA*
**>Cofca_T00039609001**
MGCLNQQTAVNAVVRGPKDGSFRSSRVALACMVNSEIGAVLAVMRRNVRWGVLYADDDQVEHSLIQSFKELRKKI
FLWQHQWHSIDPVVYLKPFLDVIQSDETGAPITGVALSSVYKILTLEILDSETVNVDKALHLIVDAVTSCRFEVT
DPASEEVALMKILQVLLACMKNKASVHLSNHHVCNIVNTCFRIVHQASSKGELLQRIARHTMHELVRCIFSHLPD
IGDRPHALAEGSRMSAETEVHVPKDTHILEGTQGVNANGSAKFDGKALPLKEASGNPAFTSASKVDEEIMTLGSV
EVAQNGADSMMDQYGIPCMVEIFQFLCSLLNVMEIESGVRSNPIAYDEDVPLFALGLINTAIELAGASFGDHAKL
LVLIQEDLFYNLMQFGLSMSPLILSTVCSIVLNLYHHLRTKLKLQLEAFFSRVLLKIAGSKHGSSYQQQEVAMEA
IVDFCRQPFFMAEMYANFDCDISCSNVFEDLANLLSKSAFPVNSPLSALNTLALDGLIAMIEGISERIGHESSAP
ERSPIQPEEYRPFWTVTCDNYGEPNCWVPFVYKKKQIKKKLMIGADHFNLDPKKGLEFLKGVNLLPDTRDTRCVA
CFFRYTTGLNKNLVGEFLGSHDEFCIQVLKEFAQSFDFQDMSLDTALRIFLETFRLPGESQKIQRVLEAFAERYY
EQSPDILVNKDAVFVLSYSLIILNTDQHNSQVKKKMTEEDFLRNNRAINGGSDLPREFLSQLYHSICENEIRMIP
EQAAAASVMSRSHWISLVHRAKQSAPYIFCDSGPHLDYDMFAILSGPTIAAISVVLDHVERDNVLHTCIDGYLTM
AKISASYNSVDSLDDLVVSLCRFTNLLIPSLNNEEFILAFGDDAKARMATVAVFTIANRYGDQIRSGWKNILDCI
LSLNKLGLLPARLASDAVDDSEVSSDQNLVRPPSSSPTTSTPSPVAPSRKSSGLMGRFTQLLYLDTEEPESYPTQ
EQVAARQRSLQTIQDSHIDSIFAESKFLQAESLSQLVQALLLAAGRPRKANNSMEDEVTAVFCLELLIAITLNNR
DRIMLLWQGVYEHIASVVQSTVMPCALVEKAVFGLLRICQRLLPYKENLTDELLKSLQLILKLDARVADAYCEQI
TQEVMHIVKANAMQIRSHMGWRTIISLLSITARHPEASEAGFETLSFIMSEGAHLSPANYVLCLTAARQFAESRV
GNVDQSIRSLDLMAGSLDCLVRWFNKTKEAMGEEAALKMAQDIWEMWLRLVQGLRKVCVDQREEVRNHAILMLQR
CLTGAEVMHIPVGLWLQCFDLVIFTLLDDLVEIAQQHSAKDYRNMEGTLVLSLKLLSKVFLQLLHDVSQLASFSE

LWSRVLSCMERYMKVKFRGKRSEKIHELVPELLKNTLLIMKSSGLLVPSEEDNFWQLTWSSVKNLAPSLQLEVFS
SDELEQLQQKTQTNGCSPIQDGNVVVPPSETRA*
**>Cofca_T00041903001**
MEDQQEGGNNNNMTKIKRKELGLSCMLNAEVGAVLAVIRRIPEANPQFHHPSEENYDSTILQSLKSLRSLIFNTE
QEWRKIDPSLYLSPFLDVIQSDDIPAAATGVALSSILKILRLDIFDEKTPGAKDVMDATVTAITACRLEKTDPVS
EEAVMMRILHVLTAVMRHKASVLLTDHAVCTVVNTCFQVVQQSAHRGNLLQRSARYTMQELIQIIYYRLPDIEVN
DWENSESDTEDSNLDSGFGIRSAVDIFQFLCSLLNVVEIAESEVLHAHTTDGDVQLFALALINSAIELSGDSIGK
HPKLLRMVQDDLFHHLIYYGTYSSPLVLSMICSTVLNIYHFLRRSVRLQLEAFFSFVLFRAANLGSSPQLQELAT
EAIVNFCRQSTFIIEVYVNYDCDPIFGNAFEEIGRLLCKHAFPTGGPLTSLQVQAFEGLVLVIHNIAENIDKEDD
TSPSGPYPVEITEYTPFWEEISKDDGLEDWVTFVRVRKAQKKKLLIAGNHFNQDDKKGLEYLKISHLIPDPPDPK
AFALFFRFTPRLDKTMIGDYLGDPDEFHIQVLNEFTNTFAFTGMILDSALRNYLETFRLPGESQKIHRILEAFSE
RFYDQQSSELFVSKDAVFVLCYSLIMLNTDQHNPQVKKKMTEEEFIRNNREINGGSDLPREYLSELFHSISSKAI
TLFGQSGTPAEMNPSRWIQLVNRAKLMKPYVLCNFDRRLGRDMFAAIAGPSVATLAAIFEQTEEEEILHECIELL
FAIARIAQYELEDNLDELICSFCKFTTLLNPYASAEETLYAFSNDMKPRMATLAVFTIANTFKNSLRGSWRTIVE
CLLKLRRLKLLPQSVVEFEIASDASIGTKHDKNESAASISTRNTKFNHIRRSSGLIGRFSHFLSLESVEESLNLG
VSEFEQNQNIIQQCHIENIFTTSSSLPDESLLNLGRALIYASAGKGQKFSTPVEEEETVAFCWDLIVIISCVNMH
RFSTIWPHYHDYLLGVAQFPLFSPIPFAEKAIVAQMKICLKILSAYQLDKVSEELIFKSINLMWKLEKEILDTCS
EFLVKSVSRILNEYPANLQSSLGWKTVLHLLSVTGRHPETYDQGVESLINLMSDGMHISRTNYASCIECAFGFVA
LKNSPVEKNIKIMDLMSDSVKLLVQWFRSGYSDPGSNVSTGSNASASSTEDNARSGSSASFIMNLFVKLGEALRR
TCLARREEVRNHAVISLQRSFILAEELCFTPANCIYCFNSLIFAMVDDIHEKMLEYSRRENAERETRSMEGSLKR
AMEVLVEVYSQLLKPISESPHFRPFWMGLLRRMDTCMKVDLGGYGKSDMQEVIPDLLKRIISIMKENELLIPKEG
DDLWEITYVQIQWIAPSLKDELFPGAP*
**>GelseGNOM_sc8**
MGRLRLQSSIKAIEEEPEDCEATSSNKAALACMLNSEIGAVLAVMRRNVRWGGRYVSGDDQLEHSLIQSLKTLRK
QIFLWQHQWQTINPVLYLQPFLDVIRSEETGAAITGVALMSVYKILTLDLLDLNTFNIEDAMHLVVDTVTSCRFE
LTDLASEEVVLMKILQVLLACMKSKASVVLSNQHVCTIVNTCYRVVHQAASKGELLQRIARYTMNELVRCIFSHL
PDIDNTDRSLVKRGSSTKDEVAGIDTDYGFTGKSENGSGTSDGQLSSGVFASNASGGMLSGVMDDGALKTDDGKD
AIYDLHLMTEPYGVPCMVEIFHFLCSLLNAVENLGLGSRANSVAFDEDIPLFALGLINSAIELGGPEVRHHPRLL
NLVQDELFRNLMQFGLSTSPMILSMVCSIVLNLYQHLRTELKLQLEAFFSCVILRLAQSRYGASYQQQEVAMEAL
VDFCRQKTFMVEMYANLDCDITCSNVFEELANLLSKSAFPVNSPLSAMHILALDGLIAVIQGMAERIGNGSSNSE
PASVNLREYTPFWMLKCESYGDPDDWVPFVRRRKYIKRRLMIGADHFNRDPKKGLEFLQGTHLLPEKLDPQSVAC
FFRYTAGLDKNLVGDFLGNHDEFCVQVLHEFAKTFDFQDMNLDTALRLFLETFRLPGESQKIQRVLEAFSERYYE
QSPQILANKDAALLLSYSLIMLNTDQHNVQVKKKMTEEDFIRNNRHINGGNDLPRDFLSELYHSICKNEIRTTPE
QGAGFAEMTPSRWIDLMHKSKKSAPYIMSDSRAYLDHDMFAIMSGPTIAAISVVFDHAEHEYVVYQTCIDGFLAVA
KISACHHLEDVLDDLVVSLCKFTTLLNPSSVEEPVLAFGDDCKARMATVTVFTIANRYGDFIRTGWRNILDCILR
LHKLGLLPARVANDAADDSEMSSEPGQGKPLTNSLSSVQMQSIGTPRRSSGLMGRFSQLLSLDTEEPRSQPTEQQ
LAAHQRTLQTIQKCHIDSIFTESKFLLAESLLQLARALIWAAGRPQKGSSSPEDEDTAVFCLELLIAITLNNRDR
IQLLWPGVYEHIANIVQSTVMPCALVEKAVFGVLRICQRLLPYKENLADDLLRLLNLVLKLDARVADAYCEQITQ
EVSRLVKANASHIRSQLGWRTITNLLSFTARHLEASEAGFDALSFIMSDRSHLSPANYVLCLDAARQFAESRVGQ
ADRSVRAVDLMAGSVASLARWAEDTKGAMAEAESIKLSQDIGEMWLRLTQLLRKVCLDQREEVRNHALLSLQMCL
TGVDEIHLPHGIWLQCFDMVIFTLLDDLIEIAQGHSQKDYRNMEGTLALALKLLTKVLLQLLQELSQLTTFCKLW
LGVLSRMEKYMKVKVRGKRNEKLQELVPELLKNTLLVMKAKGVLAQRSALGGDSLWELTWLHVNNIVPSLQAELF
PDKDSEQSENNQEQKSI*
**>GelseGNL1_sc0**
MGCLNQQGTVNDLAGEPKGCSFRSSRVAIACMVNSEIGAVLAVMRRNVRWGVRYADDDQLEHSLIHSFKELRKKI
FSWQHQWQSIDPVVYLKPFLDVIQSDETGAPITGVALSSVHKILTLEILDSETVNVDNALHLIVDAVTSCRFEVT
DPASEEVVLMKILQVLLACMKNKASVYLSNHHVCNIVNTCFRIVHQASSKGELLQRIARHTMHELVRCIFSHLPD
IDCKHHSLAQGSRSSVDTEVYVPSENHTLEDKQSVNANGSRKSDGDAPSIVASGNPTVTTAGKVDEGTISGGQKE
VAQNGVSSMMDPYGLPCMVEIFHFLCSLLNVMEHIETGPRSNPIAHDEDVPLFALGLINTAIELGGAAFGHHPKL
LALIQEELFYNLMQFGLSMSPLILSTVCSIVLNLYHHLRAKLKLQLEAFFSGVLLRISGSKHGSSYQQQEVAMEA
LVDFCRQCSFMAEMYANFDCDISCSNVFEDLANLLSKSAFPVNSPLSAMNTLALEGLIAMIQGMAERIGHESSAS
ERASLDLQEYEPFWRVLCENHNDPGNWVPFVHKMKQIKRKLMIGVDHFNRDSKKGLEFLKAACLLPDKHDPESVA
CFFRYTTGLDKNLVGDFLGSHDEFCIQVLHEFAKTFDFRDINLDTALRIFLETFRLPGESQKIQRVLEAFAERYY
EQSPDILVNNDAALLLSYSLILLNTDQHNSQVKRKMTEEDFIRNNRRINGGNDLPREFLSELYHSICENEIRMIP
EKASSTQVMTQSHWIGLVHKSKQSAPYIVCESGPYLDYDMFALLSGPTIAAISVVFDNVEQDDVLQTCVDGFLAI
AKISASYNRIDVLDDLVVSLCKFTNLLLTSYVDEEFILAFGGDAKARMATSAVFTIANRYGDQIRSSWKNILDCI
LSLNKLELLPARLASDAADDLKPSTDQHLVRPPASSPSAATVSSTAPSRKTSGLMGRFSQLLYLDTEEPAPQPTE
EQLATRQSTLQTIQDCHIDSIFAESKFLQAESLTQLVQALLLAAGRPGKLNSSLEDEVTAVFCLELLITITLNNR
DRIMLLWQGVYEHIANVVQSTVMPCALVEKAVFGLLRICQRLLPYKENLTDELLKSLQLILKLDARVADAYCEQI
TQEVMHLVKANAMQIRSLMGWRTIISLLSITARHPEASEAGFETLSFIMTDGAHLLPANYVLCVNAARQFAESRV
GNIERSVKSLDLMAGSLDCLVRWFYKTKEAVGEEAALKMAQDIWEMWLRLVQGLRKVCVDSREEVRNHAILMLQR
CLTGVEVMHIPIDLWLQCFDLVIFTLVDDLLDIAQQHSPKDYRNMEGTLVLYLKLLSKAFLQSLQDLSQLTSFGK

VWLGVLSSMEKYMKVKFRGKRSEKIHELVPELLKNTLLVMKSSGILVPTDAASGDNIWQITWSNVKNIAPSLQLE
VFSSDELEQLQEQQSKSGFSPTQDGNVLVPQSETTT*
**>GelseGNL2_sc147**
MDNSQDGENMTKMRRKQLGLSCMLNTEVGAVLAVIRRAPESNSQFYHPSEENYDSSIVQSLKSLRSLIFNPQQEW
RSIDPALYLSPFLDVIQSDDVPASATNVALSSILKILRLEIFDEKSPGAKEIMNGAVTAITSCRMEKTDPVSEDA
VMMRILQLLVTIMRHRASVLLTDHVVCTVVNTCFQVVQQSANRGDLLQRTARFAMQELIQIIYSRLPEIEVGNWE
DSESDTEDSNLDSGFGIRSAIDIFQFLCSLLNVVQIAETDGFPAQIADVDIQIFALVLINSTIELSGDSICKHPK
LLRMIQDDLFHHLIHYGARCSPLVLSMISSIVLNIYHFLRRSVRLQLEAFLIFVLFRSANLSLSLQLQEVATEAI
INFCRQPTFIIEVYVNYDCDPICRNVFEEMTKLLCKHAFPAGGPLTSLQVQAFEGLLVIIHNIADNINKEDDSSP
SGPYPVEISEYSPFWEERSKDDGQENWINFLRMRKAQKKKILIAANHFNQDEKKGLEYLKISHLISDPPDPKAYA
MFYRYTPGLDKTMIGDYLGDPDEFHLQVLKEFTNTFEFSGMILDSALRSFLETFRLPGESQKIQRILEAFSEKFY
DQQSSELFASKDAVFILCYSVIMLNTDQHNPQVKKKMTEEEFIRNNRAINGGSDLPRDYLSELFHSILANAITLF
GQSGMTVEMNPNRWIQLVNHSKIMKPFVFCDFDRRLGRDMFAAVAGPVLATLAAIFEQTDEEEILHECIDLMFSI
ARIAQYELEDTLDELVCSFCKFTTLLNPYASAEETLYAFSNDMKPRMATLAVFTIANTFKNSTRGSWRTIVDCLL
KLKRLKLLPPSVIEPEIASDSPAEPKTEKSDSGALPPQDPRLSTRNRVSSVIGRFSHFLSMESVEESLNLGIGEY
EQNVKIIQQCHIESLFSTCSSLPDDSLLNLGRALIFAAAGKGQKFSTPVEEEETVAFCWDVIVTLSMKNTGRFST
FWPHYHDYLLAVAQFPLFSPIPFAEKAIIAQVKICLKLLSTYKFDKTPEELIFKSINLMWKLEKEILDTCSEFLI
QSVSKILNEYHANLHSSLGWKSVLHLLSVTGRHPETYDQGVATLIALMSDGIHISRTNYSFCIDCAFSFVALKNS
PVEKNIKIMDLMSDSVKLLVQWYRSGYSDPGSNVSIASNASTSSTEDNAKIVTSSSFIMNLFVKLGEALRKTCLA
RREEIRNHAVISLQRSFNLAEELCFTPANCVYCFNNLIFAMVDDLHEKMLEYSRRENSERETRSMEGTLKLAMEV
LTEVYLQLLMPISESPHFRPFWMGLLRRMDTCVKADLGVYGQTKMNSTIPELLKKIISAMKENDMLVPKEGDDLW
EVTYVQIQWIAPSLKDELFPDASSL*
**>Calgi_g003986**
MGRLRLQSSINAIEEEPEDCETTSSNKAALACVVNSEIGAVLAVMRRNVRWGGRYVSGDDQLEHSLIQSLKTLRK
QIFLWQHQWQTINPVLYLQPFLDVIRSDETGAPITSVALSSVYKILTLDTLDLNTVSIEDAMHLVVDAVTSCRFE
LTDPASEEVVLMKILQVLLACMKSKVSIVLSNQHVCTIVNTCFRVVHQAGTKGELLQRIARHTMHELVRCIFSHL
PDIDNTERSLVKGVNSAKHEVTTIDADYSLSSKSENGIGASAYDEQLSGAFASNASTGLISSGMDDGATKTVNGK
DAIPYDLHLMTEPYGVPCMVEIFHFLCSLLNVVENMGMGPRANSIAFDEDVPLFALGLINSAIELGGPAIRHHPR
LLSLVQDELFRNLMQFGLSMSPLILSMVCSIVLNLYQHLRFELKLQLEAFFSCVILRLAQSRHGGTYQQQEVVME
ALVDFCRQKTFMVEMYANLDCDITCGNVFEELANLLSKSAFPVNSPLSAMHILALDGLIAVIQGMAERIGNGSSN
SEPAPVNLEEYTPFWMLKCENYRDPDDWVPFVRRRKYIKRRLMIGADHFNRDPKKGLEFLQGTHLLPEKLDPQSV
ACFFRYTAGLDKNLVGDFLGNHDEFCVQVLHEFAKTFDFQDMNLDTALRLFLETFRLPGESQKIQRVLEAFSERY
YEQSPQILANKDAALLLSYSLIMLNTDQHNVQVKKKMTEEDFIRNNRHINGGNDLPREFLSELYHSICKNEIRTI
PEQGAGFAEMTPSRWIDLMHKSKKTAQYIVSDSRAHLDHDMFAIMSGPTIAAISVVFDHAEHEDVYQTCIDGFLA
VAKISACHHLEDVLDDLVVSLCKFTTLLNPSSVEEPVLAFGDDGKARMATVTVFTIANRYGDFIRTGWRNILDCI
LRLHKLGLLPARVASDAADDSDMSSEPGHGKPLTNSLSSVQMQSIGTPRRSSGLMGRFSQLLSLDTEEPRSQPTE
QQLAAHQRTLQTIQKCHIDSIFTESKFLLAESLLQLARALIWAAGRPQRGSNSPEDEDTAVFCLELLIAITLNNR
DRIGLLWQGVYEHIANIVQSTVMPCALVEKAVFGLLRICQRLLPYKENLADELLRSLQLVLKLDARVADAYCEQI
TQEVSRLVKANATHIRSQLGWRTIASLLSITARHPEASEAGFDALLFIMSEGAHLSQVNYVLCIDAARQFAESRI
GQADRSVRAVDLMAYSVSCLAQWADDTKSGTIEAEAVKLSQDIGEMWLRLAQALRKVCLDQREEVRNHALLSLQI
CLSGVDGIRLPHVLWLQCFDMVIFTMLDDLIELAQGHSQKDYRNMEGTLVLALKLLTKVFLQLLPELSQLTTFCK
LWLGVLSRMEKYLKVKVRGKRSEKLQELVPDLLKNTLHVMKTKGVLVQRSALGGDSLWELTWLHVNNIVPSLQSE
LYPDKEAEQSETNPGEIRASPTSNETDSARKE*
**>Calgi_g002145**
MGFLNYQTVTNDMAGEPTGGCFRSSRVALACMVNSEIGAVLAVMRRNVRWGVRYADDDQIEHSLIHSFKELRKKI
FSWKYQWHVVDPIVYLKPFLDVIQSDETGAPITGVALSSVYKILTLEILDSDTVNVDNALHQIVDAVTSCRFEVT
DPASEEVVLMKILQVLLACMKHKASSHLSNHHVCNIVNTCFRIVHQASSKGELLQRIARHTMHELVRCIFSHLPD
IDNKQTSLAEGTGSTSDKEVSMPIENQSLKDKQTINANGSAKPDDQASLFVASSISTVTLPGKVDEESTSPCPKE
VVLNGGNSMMDPYGVPCMVEIFHFLCSLLNVMEHIETGPRSNPIAYDEDVPLFALGLINTAIELGGAALGNHPKL
LVLIQEELFYNLMQFGMSMSPLILSTVCSIVLNLYHHLRSKLKLQLEAFFSGVLLRIAGSKHGSSYQQQEVAMEA
LVDFCRQHFFMSEMYANFDCDISCSNVLEDLVNMLSKSAFPVNSPLSAMNTLALDGLIAMIKGMAERIGHESSTP
EQFLLDLQEYEPFWTGRCENYNDPGNWVPFIQKMKKIKRKLITGVDHFNRDPKKGMEFLQGVHLLPDETDPQSVA
CFFRYTTGLDKNLVGDFLGSHDEFCIQVLHEFSKTFDFRAMNLDTALRIFLETFRLPGESQKIHRVLEAFAERYF
EQSTDILANKDAALVLSYSLIMLNTDQHNSQVKKKMTEEDFIRNNRRINGGNDLPREFLSELYHSICENEIRMVP
EQSTGAAVMMRSHWIGLVHRSKQSAAYIFCDSLPHLDYEMFAILSGPTIAAISVVFDHVDQDEVLQTCIDGFYAV
AKIAASYNLIDVLDDLVVSLCKFTNLLLPPFVDEEFILSFGDDTKARLATTAVFTIANRYGDHIRSGWKNIVDCI
LSLNKMGLLPARLATDAADELETSPDTHLVRAPASSPSSATVSMPPARKSSGLMGRFSQLLYLDTEEPAPQPTEE
QVAARQRTLQTIQDCHLDGIFAESKFLQAESLMQLVRALLSAAARPHKVNNSLEDEVSAVFCLELLIAITLNNRD
RIMLLWQGVYEHIANVVRSTVMPCALVEKAVFGLLRICQRLLPYKENLTDELLKSLQLILKLDARVADAYCEQIT
QEVMHLVKANAIQIRSHMGWRTIISLLSITARHPEASETGFETLSFIMTDGIHLLPANYVLCVNAARQFAESRVG
SIDRSIKSLDLMAGSLDCLLRWCHETKEAVAEEASLKMAQDIWEMWLRLVQGVRKVCVDQREEARNHAILILQRC
LIGVEVMHIPNELWSQCFDQVLFTLLDDLLEITQQHSPKDYRNMEGTLVLSLKLLSKVFLQSLQDISQLASFSKL

WLGVLSCMERYMKVKFRGKRSEKMHELVPELLKNTLLVMKSSGILVPVDPVRGDSFWQLTWLNVKNIAPSLQSEV
FSSDELDQLQKQHASGDSPIQDGSILIPTSETAA*
**>Calgi_g000739**
MENQQEEESKKKGMMSRNKRKELGLSCMLNTEVGAVLAVIRRAPDSQFYHPSDDHIDCSVLQSLKSLRTLIFNPQ
QEWRSVDPSIYLSPFLDVVQSDDIPAFATGVALSSILKIFRLEIFDQKTPGVGEIMNSTVTAITGCRLEKTDQVT
EDVVMMRILQVLVAVMKHKAYVLLTDHSVCTVVNTCFQVVQRSAGRDLLQRSARHTMQELTQIIYQRLPEIEVVD
GQSSESETEDTNLESGFGIYSAVDIFQFFCSLLNVVNDSDGNNSYTQNHNIPDFNIQMFALVLINSAMELSGDSM
GKHPKLLRMIQDDLFHHLIYYGTSCNPLVLSMICSTVLNIYHFLRRSVRLQLEAFFSFVLFKAANLDKAQIQEVA
IEAIVNFLRQPNFTIEVYVNYDCSPFFGNVFEETAKLICKNAFPVGGILTNLQLQSFEGLAMMIHNIADNVDKED
DYSTSGPYPVEVTNRTPFWEEKSKDDVGLEKWINLLRIKKVQKKRLLIAGNHFNQDYKKGLEYVELSGLVSGSDP
PNPKAYAMFFRYTPGLDKTMIGDYFGDPEEFHIQVLQEYTKTFEFSGMLLDSALRCFIESFRLPGESQKIQRILE
AFSERFYDQQSSDIFENKDAVFVLCYSLIMLNTDQHNPQVKKKMTEEEFIKNNRAINGGKDLPREYLSKLFHAIS
SKAFETLGQPGTTDTHMHPYRWVQLVNRSKIMNPYVECDFDRRLGRDMFAAIAGPSIATVAAIFEHSDDEDVTHE
CVETLFSIARIAQYQLQDTLDELISVFCKFTTLLNPYASADETLFAFSNDTKPRVATLAVFNIANTFKDSLRGSW
RNIIDCLLKLKKLKLLPFSAFDPDASDLSSRDSGRLSRTSTRDSESGSSSSTSKRLQQIGLIGRFPDFVSIESLE
ESLNLGLSEFEQNMKIIQQCHIERIFRGSCGLPEESLWNLGRYLIFAAGGKGQKFSTLVEEEDTVALCWDLIATI
SYANTHRFTSFWRQYHEYVLAVTQFPLFSAIPFAEKAMVAQLKICLKLLSVTHPDKQAEELIFKSINFMLELEKE
ILDTCSDLLLQAITKILREYRGNVQSSRGWKTILHLLYVTGRHPDSYDQGVESLISLMCDKSAMSRTNYGFCIEC
AFNFIALKNSSVDKNAKIMDLMCESVKVLVQWYRNEYVDPDSNASVHSTASTSSSGDDNSKGMSSTSTFIMNLFL
KVGEVLRKTCLARREEVRNHAVQSIQKIFTFAEELCFTPANSAFCFNSLIFAMTDDVHEKMLDYSKRDNFEKESL
SMENSLKNAMGVMTEVYLQLLKPLSENPRFKAIWMGLLRRMDACVKAEVGDTKEMQELVSNLLKKIIDSMKEKDM
LVPREGDDLWEMTYVQIEWIAPSLKEELFPDLAGLYIDQKSNP*
**>Chial_g3955**
MGRLRLQSNIKAIEEEPEDCETTSSNKAALACMINSEIAAVLAVMRRNVRWGGRYVSGDDQLEHSLIQSLKKLRK
EIFLWQHQWQTINPVLYLQPFLEVIRSDETGAPITGVALSSIYKILTLDVLDLSTVNIEDAMNLVVDAVTSCRFE
ITDPASEEVVLMKILQVLLACMKGKTSVVLSNQHVCTIVNTCFRVVHQAGTKGELLQRIARHTMHELVRCIFSHL
PDVDNTERSLVKGGTSIRHEVGGIDSDYSFGSKSENGSEYEGQPSSGASTGLLSGVMGDGGGKMDNGKDSVPYDF
HLMTEPYGVPCMVEIFHFLCSLLNVVEHVGMGPKANSLAFDEDVPLFALGLINSAIELGGAAIRHHPRLLSLVQD
ELFRNLMQFGLSMSPLILSMVCSIVLNLYQHLRTELKLQLEAFFSCVILRLSQSRYGASYQQQEVAMEALVDFCR
QKTFMVEMYANLDCDITCSNVFEELANLLSKSAFPVNSPLSAMHILALDGLIAVIQGMAERIGNGLFNSEPAPVK
LEEYTPFWMLKCENYNEPNNWVPFVRRRKYIKRRLMIGADHFNRDPKKGLEFLQGTHLLPEKLDPQSVACFFRYT
AGLDKNLVGDFLGNHDEFCVQVLHEFAKTFDFQDMNLDTALRLFLETFRLPGESQKIQRVLEAFSERYYEQSPQI
LANKDAALLLSYSLIMLNTDQHNVQVKKKMTEEDFIRNNRHINGGNDLPREFLSELYHSICKNEIRTTPEQGAGF
AEMTPSRWIDLMHKSKKTASYIVSDSRAYLDHDMFAIMSGPSIAAISVVFDHAEHEEVYQTCIDGFLAVAKISAC
HHLEDVLDDLVVSLCKFTTLLNPSSVEEPVLAFGDDSKARMATVTLFTIANRYGDFIRTGWRNILDCILRLHKLG
LLPARVASDAADESEMSPEPGQGKPLTNSLSSVQVQAIGTPRRSSGLMGRFSQLLSLDTEEPRSQPTEQQLAAHQ
RTLQTIQKCHIDSIFTESKFLLAESLLQLARALIWAAGRPQKGSSSPEDEDTAVFCLELLIAITLNNRDRIGLLW
PGVYEHIANIVQSTVMPCALVEKAVFGLLRICQRLLPYKENLADELLRSLQLVLKLDARVADAYCEQITQEVSRL
VKANATHIRSQLGWRTIASLLSITARHPEASEAGFDALLFIMSDGAHLSPANYVLCIDAARQFAESRIGQADRSV
RAVDLMGGSLACLTQWDNDAKGAMVEAEAMKLSQEIGEMWLRLAQALRKVCLDQREEVRNHALLSLQMCLMGVEG
NHLPYGLWLQCFDMVIFTMLDDLAEIAQGHSQKDYRNMEGTLVLALKLLTKVFLQLLNELSKLTTFCKLWLGVLS
RMEKYMKVKVRGKKSEKLQELVPELLKNTLLVMKTKGVLVQRSALGGDSLWELTWLHVNNIVPSLQSELFPDNDS
EQSDSKQDEMGKTAVPNETGSA*
**>Chial_g16800**
MGCLNQQTAGNALVIGPKNCSFRSSRVAIACMVNSEIGAVLAVMRRNVRWGIRYADDDQQEQSLIHSFKALRKKI
FLWQHEWHLIDPVVYLQPFLDVIQSDETGAPITGVALSSVYKILTLDILDSETVNVDKALHLIVDAVTSCRFEVT
DPASEEVVLMKILQVLLACMKNKASVHLSNHHVCNIINTCFRIVHQASSKGELLQRTARHTMNELVRCIFSHLPE
IDNIPDALAEGTRMSTETEVNVPTENHMVEEKQGVDANGITKSDGKAPSPEDASGDPSVTPDGKVDEDTKPLGQR
KVAQNGAESMMDPYGVPCMVEIFHFLCSLLNVMEHIENGPRSNPIAYDEDVPLFALGLINTAIELGGASFGDHPK
LLALIQEELFYNLMQFGLSMSPLILSTVCSIVLNLYHHMRAKLKLQLEAFFSRVLLRVAGSKHGSSYQQQEVAME
ALVDFCRQPFFMAEMYANFDCDISCSNVFEDLASLLSKSAFPVNSPLSALNILALDGLIAMIEGMAERIEHESSA
LERTSLELEDYRPFWTGTCDNYKEPSYWVPFVHQMKQIKKKLMIGVDHFNRDPKKGLEFLKGVNLLPDTHDPHSI
ACFFRFTTGLDKDLVGDYLGSHDEFCIQVLNEFAKSFDFRGMNLDTALRIFLETFRLPGESQKIQRVVEAFAEGY
YEQSPDIFANKDAALLLSYSLILLNTDQHNAQVKKKMTEEDFIRNNRRINGGNDLPRDFLSELYHSICEDEIRMI
PEQSSAASGITRSHWIGLVHRSKQSAPYIFCDSGPYLDCDMFTILSGPTIAAISVVFDHVEQDSVLQTCTDGYLA
LAKVSASYNFMDALDDLVVSLCRFTNLLLPSFVDEDFILAFGDDAKARMATVTVFTIANRYGDQIRSGWKNILDC
ILSLNKLGLLPARLASDAVDDLEPSSDQNLVRPPSASPTPSTPSTMAHSRKSSGLMGRFSQLLYLDAEEPEPQPS
QEQVAARQRSLQTIQDSHIDSIFAESKFLQAESLSQLVQALLLAAGRPRKASNYLEDEVTAVFCLELLITITLNN
RDRIMLLWQGVYDHIANVVQSTVMPCALVEKAVFGLLRICQRLLPYKENLTDELLKSLQLILKLDARVADAYCEQ
ITQEVMHLVKANAMQIRSHMGWRTIISLLSITARHPEASEAGFETLSFIMSDGAHLLPANYVLCVNAARQFAESR
VGNADRSVRSLDLMAGSLDCLVRWFYKTKEAVEEEAALKMAQDIWEMWLRLVQGLRKVCVDQREEVRNHAILILQ
RCLIGVEVMHIPGDLWSQCFDLVIFTLLDDLVELAQQRSAKDYRNMEGTLVLSLKLLSKVFLQSLQDLSRLPSFS

NLWSRVLSCMERFTKVKFRGKRSEKIHELVPELLKNTLLVMKSSGILVPSEGDGFWQLTWSNVRNIAPSLQLEVF
SSDELEQVQMAQTKICNPIQEGNVVVPRSETTA*

**>Chial_g8037**

MDNQQEEESKKKNMTKIKRKEQGFSCMLNAEVGAVLAVIRRIPEPNHQFYHPSEENYDSIIQQLKSLRSLLFNPE
QEWRKIDPSLYLSPFLDIVQSDDIPATATGVALSSILKILRLEIFDEKTTGAKDIMNATVSAITACRLEKTDPVY
EEAVMMKILHILTAIMRHRASVLLTDHAVCTVVNTCFQVVQQSAYRGNLLHRSARFTMQELIQIIYYRLPDIEVN
DWENSESDTEDDNLDSGFGIRSAVDIFHFLCSLLNVVEVAEPTEILHTQTTDGDVQIFALVLINSAIELSGDSIG
KHPKLLRMVQDDLFHHLIHYGTYSSPLVFSMICSTVLNIYHFLRRSVRLQLEAFFSFVLFRAANLGNSLQIQEVA
TEAVVNFCRQPTFLIEAYVNYDCDPIFGNAFEEIGKLLCKQAFPTGGPLTSLQVQAFEGLIVVIHNIAENIDKED
DTTPSGPYPVGIPDYKPFWEEISKDDGLEVWVNFVRVRKAQKKKILIAGNHFNQDDKKGLQYLKISQLIPDSPDP
KAFALFFRFTPRLDKTMIGDYLGDPDDFHIQVLKEFTTTFEFTGMILDSALRNYLETFRLPGESQKIQRILEAFS
ERFYDQQSSDIFASKDAVFILCYSLIMLNTDQHNPQVKKKMTEEEFIRNNRAINGGNDLPREYLSELFHSISSNA
ITLFGQSGVAVEMNPSRWIQLVHRAKLTKPFILCDFDRRLGRDMFAVIAGPSVATLAAIFEQSEEEQILHECVEL
LFSIARIAQYELEDTLDELICSFCKFTTLLNPYASAEETLYAFTNDMKPRMATLAVFTIANTFKNSIRGSWRTIV
DCLLKLKRLKLLPQSLVEFDTALDASTEAKHEKFESGAAFSTHNTKFNHIRRSSGLIGRFSQFLSMESVEESLNL
GVSEFEQNQNVIQQCHIEFIFTTSSSLPHESLLNLGRALIFAAAGKGQKFSTPIEEEETVAFCWDLIVIISSVNM
NRFSTFWPHYHDYLLGVAQFPLFSPIPFAEKAIVAQMKICLKLLSTFQFDKLPEELIFRSINLMWKLEKEILDTS
SDFLIQSVSRILNEYPANLQTSLGWKTVLHLLSVTGRHPETYDQGVESLINLMSDGTHISRTNYALCIECAFGFV
ALKNSPVEKNTKIMDLMSDSVKLLVQWYRNGYSDPGSNASVGSNASGSSAEDNAKPISSASFVMNSFIKLGEALR
KTCLARREEVRNHAVISLQRSFMLAEELCFTPANCIFCFNSLIFAMVDDIHEKMLEYSRRENAERETRSMEGSLK
RAMEVLTEVYLQLLKPISESPHFRPFWMGFLRRMDTCMKADLGEYGQSNMQDVVPDLLKKIISVMKKNDMLVPKE
GDDLWEITYVQIQWIAPSLKDELFPDAT*

**>Genau_EPS60000**

MGRVKPQPSIKAIEEEPEDCETTSSNMDALKCMINSEIGAVLAVMRRNARWGGRYVSGDDLLEHSLIQSLKTLRK
QIFFWQQDWQSVNPVLYLQPFLEVIRSDETGAPITGVALQSVYKILTLDVLDLNTLNIEDAMHLVVDAVTSCRFE
VTDPTSEEVVLTKILQVLLACVKSKASVVLSNQHVCTIVNTCFRIVHQAGSKSELLQRFSRHTMNELIRCIFSHL
PDVDRTEGSLVKGVNSLKNKTGGVDSDSTFVRSSENGNGTHDFDGQLSSGILSSASSGLINGPMDETNSRADNGK
DVVPYDLHLMSAPYGIPCMVETFHFLCSLLNVVEHTGMGSGPNSVAFDEDVPLFALGLINSAIELGGSVIRHHPR
LLNLVHDELFRNLIQFGLSMSPLILSTVCSIVLNLYHHLRTELKLQLEAFFSCVILRLAQSRYGASYQQQEVAME
ALVDFCRQKAFMVEMYANLDCDITCGNVFEDLANLLSKSAFPVNCPLSSMHILALDGLIAVIQGMAERIGNGSVA
PEPNPVSLVEYVPFWMVKCQNYDDPDQWVPFVRRRKYIKRRLMVGADHFNRDPKKGLEFLQGTHLLPEKLDPQSV
ACFFRFTAGLDKNLVGDFLGNHDEFCVQVLHEFAGTFDFQDMNIDTALRLFLETFRLPGESQKIQRVLEAFSERY
YEQSPLILANKDAALLLSYSLIMLNTDQHNVQVKKKMTEEDFIRNNRHINGGNDLPRDFLSELYQSICKNEIRTT
PEQGGGFAEMTPSRWVDLLHKSRKSPPYIVADSRTCLDHDMFAIMSGPTIAAISVVFDHAEYEDVYQTCIDGFLA
VAKISACHRLEDVLDDLVVSLCKFTTLLNPSSSVEEPVLAFGDDAKARLATVTVFTVANRYGDFIRTGWRNILDC
ILRLHKLGLLPARVASDAAEDSEVPTDSRHGSKIPTNSLSSAHLQAIGTPRRSSGLMGRFSQLLSLDTEEPRSQP
TEQQLAAHQRTLQTIQKCHIDSIFTESKFLEADSLSQLARALIWAGGRPHSKGSPEDEDTAVFCLELLIAITLNN
RDRIGILWAGVYEHIAGIVQSTVVACAVVEKAVFGLLRICQRLIPYKESLSDDLLKSLNLVLKLDARVADQYCDQ
ITQEVGRLIRANVAHIRSPVGWRTIGSLLSITARHPDASESGFEALSFVMAAEEEGGGAHLSPAGFVFCVDAARQ
FAESRIGPADRSIRAVDLIAGSKSCLIRWARDAGEGGMPESESVKLRQEIGEMWLRLLQALRKVCLDQREEIRTH
SLHSLQICLSAYDGGLRSQIFDLVIFPLVDDLAEVAAQNQKEYRNVEGSLIQAVKLLSEAFAEWLGELSESAGFG
KLWRNAMGRMEKCVKGKMKRGGEKAVQEAVQEAMRNCLAAMKAKGVLVAPPGLAVGDNGGVWEQTWLHVNKMFPA
MQSQVFPGHD*

**>Genau_EPS65079**

MRSIRMQRQLNGSSAQSEALFKPSGGALACIVNAEIGAVLAVMRRNVRWGVHYGHDDEQLEHSLIISFKELRKKI
FSWQNLWHTIDPVLYIQPFLDVVKSDETGAPITGVALSSIYNILILQILDSITVNVEKALHLIVDAVTTCRFEVT
DPASEEVVLMKILQVLLACMKSQTSNRLSNHHVCSIVNTCFRIVHQASSKSELLQRTSRHTMLELVRCIFQKLPQ
LDNKMDELSSGSKFRKLEIDSAVKVQTVREQPYGNELSFSEYEVKENETAHGEVDDAMTDPYGLPAMADIFHFLC
SLLNVMENMETGSKSNPIAYDEDVPLFALGLINSAIELTGPSLGNHPKLLRMIQGELFCNLMQFGSSKSPLILSM
VCSIVLNLYYHLRTELKLQLEAFFSGVLLRVAQSKYGSSYQQQEVAMEALIDFCRQPIFAAELYANFDCDISCSD
VFEGIGNLLSRSAFPVNNPLSAMNTLALDGLIALLQNMADRIGLDTSNIGKSSQELPENTSFWTLRCDDYDEPLH
WVPFVNYMKSIKKKLMTGVDHFNRDPKEGLEFLQVLHLLPQDLDPKSVACFFRYTVGLDKDLVGDFLGHHDEFSV
QVLHEFARTFDFHDMNLDAALRIFLETFRLPGESQKIVRVLEAFSESYFEQAPNILVNKDAAFVLSYSVIMLNTD
QHNVQVRNKMTEDAFINNNRKINGGKDLPREYLSELYHSICENEIRMVPDQVGSAALFTRSHWIGLIHRSKQTSP
YIASDAGSLLDPDMFAVLSGPAVAAVSVVFDHAEQEEVLNSCIDGYLAIAKLSASYNFGEVLNDLVISLCKFTTL
LHPPSGENSILYGGDDIKAKMATEAVFTVANRYSDHIRSGWKNIVECILSLHKIGLLPSRLVTDATDDLESFPLK
DQIRSPASTSPAVQVPATANSRKRSGIMGRFSLLLSLDAEEPMDQPSEDQLAARQRVLQIVQNCHIDSIFAESKF
LQAESLLELVNALVLAAERPLKGNHSLEDEDTAVFCLELLIAVTLNNRDRIMLLWQNVYEYIANVVQSIAMPCAL
VEKAVFGLLRICQRLLPYKENLTDELLKSLQLVLKLDARVADAYCEQITQEVMHLVKANAFQIRSHMGWRTIVSL
LSITARHPEASESGFETLSYIMSEGAHLSPANYVICINAARQFAESRVGHVDRSIKSLDLMADSVFCLGTWFQRS
KEAASEEVAKKMCQDILEMWMRLVQGLRRVCVDHRQEVRNHAIVLLQRCLAGVNGIRIPASLWLQCFDLVIFTLL
DELPEMVQQHSPKEYRSIEASMVFSLKLLSKTFLQSLVDLSQSTQFCQTWLKVLGFMERYMNLKFRGKRSEKIHE

LIPELLKNTLLVMKSGGILVPGDPVGGDSFWKLTWMHVKNIDPSLQSEAFPGEESAKTGCSPAIPDGNIVVPSNE
TAA*

**>Genau_EPS71757**

MAPLGGLKNKINGASLWPECVVKPSSGAIACMVNAEIGSVLAVMRRNVRWSVLYGADGELLEHSLVSSFKELRKK
IFSWQNLWNSVDPISYINPFLDVIKSDETDATVTGAALSSVYKILSMQILDSKTVNVESALRLIVDVVTTCRFEV
TDPASEEAVLMKILQVLSGCMKNQFSNTLSNHHVCSIVNTCFRIVHQASSRSELLQKASCHSMHELINSIFLRLP
HLDYGSSFRNMEADAVQIRAVVEKVQISGFSFSLPATIKAKAEEDDVFYGPDDSMMDPFGLPAMVEIFHFLCSLP
DVMMEEGNIGINGEDVPLFALGLINSIIELVGPSLGYHPKLLSLIQDKLFFNLMRFGFSTSSPLVLSAVCSIALN
LYYHLRTKLKLQLEAFISGVLFRVAEGNYGSSYRLQEVAMEALVDFCRQPTFVSELYANYDCDLSCNDLFEGIAD
LLSRNAVPVSSQLSSVNILALDGLIVLLHSMVDRIGLDTSDVGKSLQEIPEAKPFWTLRCHDYDELVHWVPFVHA
MKMTKKKLMIGVDHFNRDPQEGLEFLQELHLLPEKLDPGSVARFFRYAVGLDKNLVGEFLSRRDEFSTQVLRDFA
KSFDFHDMNLETALRTFLETFRLPGESQKILRVLEAFSESFYEQCSKIFVSKDVAFLLSYSLVMLNTDQHSLQVK
NRMTEEDFIANHRRINGGNDLPRDYLSEIYHSIRRNEISTLPDRVEPAATTLPSRSDWIGLLQRAKRASPYIVSD
AESLLDPAIFSVLSRPVISAVSAVFDRAAREEVLNSCIDGFLAMAELSVSYDNGEVINDLVLSLCKFAPTDVDSI
ETKVKMAIEAIFAVSNRYGDYVRSGWKNVVECISNLHKIGVLSSDDIVADAIDDSVRNRSSSVINWPSIRVVPAR
SSRNIPRKRSGIMGRFTLLLSRDSGERSTDERIAEKLEHVLRTVQNCNIDRIFAESKFLRPVSLSELANALLVEA
RLSFTEDESQTVFRLDLLIAVALVNCDRIMLVWPDIYGYVSDVVQSVLVPCDLVEKAIFGLLRVCKDVLPYQEDL
TDELLIKSLQLVLKLDPRVSDAYCERIASRVMDLMKANASRIRSHVTWRSIVSLLSITVRHPEASETGFEILSFI
MSGGSGAPLLSPANYVICINAGRLFVESSVSPVERCIKSLNLMSDSIFCLSTWFRRSKEEEEEASEEMWLEMWVR
LLQGLRIVSTDYREEVRNHAIVSLQKCLSGGIDGIRTPSDLWVQFLNQLIFTLLDELIEIGQRSRKQYGSVEVSM
VLALELVFEAFAEWVVSRGDLSQAFDEAWVKLLRLMERCLKMRFFRGKRSEKIHGVIHERLKKTLSVMKTLVNGG
DFDGFWLDVKDIVPSLESEHGSLFVSTNGTTV*

**>Genau_EPS59939**

MEDESSTSKYRRRDLGLSCMLNTEIGALLAVIRGADSNSACFFHHHDDGCDTWILQSLKSLRSILFNPQQEWRRI
NPSVYLSPFLDVIQSDDVPATATAVALQSVLKILRLGVFDDRSPGAREVINLAVTAVTGCRLEKTSPMSEDAVMM
RILQVLTAIMRHPASVLLTDHSACTLVNTCFQVVQQSASRGDSLQRSARYTMHELIQIIYTRLPGIEVNDWENSE
SDSEDNNMDSGYGIRSAIDIFHFLCSLLNVIDVVESDGMAYQTADENTQAFALYLINSAIELSGDSIGKHPKLLR
MIQDDLLHHLIHYGATSSPLVLSMICSTVLNIYHFIRRSIRLQLEAFFSFVLFRVMTPSKPVQLQEVAIEGIINF
CRQPTFTIEIYVNYDCDPTFRNIFEETAKLLCRYAFPTGGGGHTTSIQFQAFEGLAVILHIIADNIDREDDSSPS
GPYPVEITRYKPFWEDKEEQNQTSPENGEEETWVDLVRYRKVQKRKILIAGNHFNRDDKKGLEYLKISQLVSDPP
DAKAYATFFRYTPRLDKIMIGDFLGDPDSFHIIVLKEFTDTFEFSGMILDTALRTYLESFRLPGESQKIQRILEA
FSERFYDQQSSDLFVSKDAIMILCYSIIMLNTDLHNPQVKKKMTEEDFIRNNRSINGGSDLPRDYLSELFQSILS
NPIILDQSGGSLGMNPNKWIQLINRSKIIRPFIHCEFERRLGRDMFAAIAGPSVATVAAVFEHSDEEEILHECIE
ALFSIARISQYGLEDTLDELLCSFCKFTTLINPYASAEETLYAFCHDMKPKMAILAVFTIANAFKESIRGGWRTI
VECLLKLKRLKLLPQNMFEQDESRPPVLPNHDPKTGNRRHNSGIMARFQNVMSMESVEEALNLGLSEFEQNVKII
QQCRIGTMFSTSSNLPDDSLLNLGRCIIFSAAGKGQKFSTPIEEEEIVGFCWELITSLTLANTHRLATFWPRYHD
YLVEVAQFPLFSPIPFSEKGIIAVTKMCLKLLASDELGATEEIFFKSINLMWRTEKEILDTCCEFLLQSASKIIA
EHPGKLQTQLGWATALHLLSVTGRHSETYDSGVEALIKLMSGRSYITRVNYSSCVDCAFAFVALRNSKVENNVKI
MDLLAETANLLVEWYRATYLDPEAGASSNNSGGSSTEDGSSKHYNPVNLFQRLGESLRKTSLARREEVRNHAVVS
LHRSFISAEDLFFIPANCINSFYNIIFAMVDDLHEKMLQYSRRENSERETRSMEGTLKLSMEVLTDVYLNFVKPI
SESPNFRTFWLGILRRMDTCMKADLAEYGESNMPDVIPDLLRKIVAEMKEKGILRRDGDDDLWDTTFYQIQWIAP
ALTQELFPEGLGLNKP*

**>UtrgiGNOM_ctg749**

MGRLKPQPSIKAIEEEREDSDVNSSPMAALQCMINSEIGAMLAVMRRNVRWGGRYVSGDDQLEHSLIQSLKTLRR
QIFSWPQDWQSINPVLYLEPFLDIIRSDETGAPITGAALQYIYKILTLDILNVNTVNIEGAIHLVVDAVTSCRFE
ITDPTSEEVVLTKILQVLLACIKGKASAVLSNQHVCTIVNTCFRIVHQAGTKGELLQRIARHTMNELIRCIFSHL
PDVDHTEKSLVKGGNSVNCKVYNVDVDCNVGKSAENGSGPLDCDGHLSSGVFNSAPYGLISRPLDESNSRTENGK
YMLPYGMHLMAEPHGLPCMVEIFHFLCSLLNVVENNGMGSGANSMTFDEDVPLFALDLINTAIELGGSVIRHHPR
LLNLVQDELFQNLIQFGLSMSPLILSTICSIVLNLYQHLRTELKLQLEAFFSCVILRLTQSRYGASYQQQEVAME
ALVDFCRQKTFMVEMFANLDCDITCGNVFEDLSNLLSKSAFPVNCPLSSMHILALDGLIAVIQGMAERIGNGSVS
SELNPLNLVEYTPFWMVKCEDYGDFEHWVPFVRKRKYIKRRLMIGADHFNREPKKGLEFLQGTHLLPEKLDPQSV
ACFFRYTAGLDKNLVGDFLGNHDEFCVQVLHEFAKTFDFQDMNLDTALRLFLETFRLPGESQKIQRVLEAFSERY
YEQSPQILANKDAALLLSYSLIMLNTDQHNVQVRKKMTEDDFIRNNRHINGGNDLPREFLSELYHSICKNEIRTT
PEQGAGFAEMTPSRWIDLLHKSKKTSPYIVTDSKAYLDHDMFAIMSGPTIAAISVVFDHAEYEDVYQTCIDGFLA
VAKISACHHledvlddlvvSLCKFTTLLNPLSVEEPVLAFGDDAKARLATVTVFTIANRYGDFIRTGWRNILDCV
LRLHKLGLLPARVasdaaddsemssdaaYGKPLTNLMSSTHVQTAGTPRRSSGLMGRFSQLLSLDNEEPRSQPTE
QQLAAHQRTLQTIQKCHIDSIFTESKFLQADSLLQLARALIWAGGRPQKGNPSPEDEDTAVFCLELLIAITLNNR
DRIGLLWPGVYEHIANIVQSTIIACALVEKAVFGLLRICQRLLPYKENLADELLRSLQLVLKLDARVADQYCDQI
TREVSRLVKANATHIRSSVGWRTIGSLLSITARHPDASESGFEALSFIMADGAHLSPANFVFCIDAAKQFAESRI
GQPDRSVHAVDLMAGSSASLIRWAKDAREAMAEAESEKLFQDIGDMWLRLVQALRKVCLDQREEVRNHALLSLQT
CLTGMDEIHVPVGLWPQCFEMVIFSLLDDLADIAQGNAQTQKEYRNMEGTLVLSLKLLHKVFLHRLRELSESPGF
CKLWRNMISRIEKFIKLKIKKAERLQELAPELLKNTLVVMKTKGVLTPTSTLGGDNVWEQTWLNVNKIFPSLQAE
VFPNEDPQSPPASNLAEPTKGRTQK*

**>UtrgiGNL1_ctg 736**
MRSTPQQSKVNGVLEKPECPFRPSGGALACMVNSEIGAVLAVMRRNVRWGVQYGADDEQLEHSLIVSFKELRKKI
FLWQSHWYTVDPVLYIQPFLDIIKSDETGAPITGVALSSIYKILNLQILDSRTVNVDSALHLIVDAVTSCRFEVT
DPASEEVVLMKILQVLLACMKSQASNSLSNHHVCSIVNACFRIVHQASSKSELLQRTSRHTMHELVRCIFLRLPY
FHDGMEDITRRDRYPKIEVSSNQDVETVQNHSVKTELHIDGFHVAEYEKKNESGDGDDGSFIGNSSMMDPYGLPA
AIEILHFLCSLLNVMENIEISSRSNPIAYVEDVPLFALGLINSAIELSGPSLGNHPKLLAIIQEDLFCNLMQFGL
SMSPLILSTVCSIVLNLYHHLRTKLKLQLEAFISGVLLRVAQGKHGASYQQQEVAMEALIDLCRQPMFATDMYAN
YDCDISCTNVFEAIANLLSRSAFPVNSPLSVINTLALDGLIAVLESMADRIGPTASGDGEYSQELTEYKPFWTLG
CYDYEQPLHWVPFVRNMKKIKKKLVVGVDQFNTDPKKGLEALQATHLLPEKLDPKSVAFFFRFTFGLDKDVIGDF
LGSHDPFSIQVLQEFARTFDFKDMNLDVALRIFLETFRLPGESQKIHRVLEAFSESYYEQSPDILVNKDAALLLS
YSLIMLNTDQHNTQVKKKMSEEDFIRNNRKINGGKDLPQDFLSELYHSICENEIRMVPDQVGSAAVLRRSHWIGL
VHKAKQTTPYIASDGACHLDYDMFAVLSGTAVAAVSVVFDHAEQEVTLNSCIDGFLAIAKLSVSYNYGEVLNDMV
VSLCKFTTLLHPSFSEKSLLYFGDDIKAKLATEAVFTLVNRYGDHIRSGWRNIVECILSLQKIGLLPSHIVNDAT
DDLESLPLMDQTTLSGNHSPIFQAPAVTTSRKPSGIMGRFSLLLSLDGEQATTQPSEEQVAGRRRIVQMVQNCHI
DSIFAESKFLQADSLSQLVHALVMTAGSSLKGNISVEDEETSVFCLELLIAVTLNNRDRIMLLWQKVYEHIANVV
QSIAMPCALVEKAVFGLLQICQRLLPYKENLTDEllkslqlvlklDARVADAYCEQITQEVMHLVKANAAQIRSH
MGWRTIISLLSITARHPEASEAGFETLRYIMSDGAHLSPANYVLCISAARQFAESRVGQVDRSIKSLDLMADSVV
CLSAWYRKTKEAASGEAVEKMSQDILEMWMRVNQGLKRVCVDQREVVRNHAIMSLQRCSAGVDGIQVPTDILLQY
LDQVIFKLVDELLDMVLPHSLKDYRSMEGSMILALKLLTKAFLQSLNDISQSNAFSEIWLKVLRCLEKYLNLRFR
GKKSEKIHELVPELLKNTLLVMKASGILVPSDAVAGDGLWDFTWLHVKNIAPSLQSEVFSEEELKKLQENQAVSG
CSLIPDGNTLVSPNQTAA*
**>UtrgiGNL2_chr1**
MDSGEQDSSCSMGRSRRKELGLSCMLNTEVGALLAVIRESNSGFYHRPDDGCDPWILQSLKSLRALLFNPQQEWH
AIHPCVYLSPFLDVLQSDDVPaaataaaLQSVLKILQLEVFSESSPGAREVINLTVTAITSCRLEKTSPMAEDAV
MMRILHALVAIMKHSASVLLTDYSVCTIVNTSFQVVQQSANRGDSLQRSAKYTMHELIQIIFSRLPDIEVNDWEN
SESDSEDNNMDAGFGIRSAIDIFHFLCSLLNVVDILESDGMTYQTADENMQAFALYLINSAIELSGDSIGKHPKL
LRMIQDDLFHHLIHYGTTSSPLVLSMICSTVLNIYHFFRRSIRLQLEAFFSFVLFKVATAGRSPQLQEVAVEGII
NFCRQPTFTIEVYVNYDCDPTFKNVFEETGRLLCKYAFPTGGRMSSIQFQAFEGLALIFHTIADNIDREDDSGPS
GPYSVEISCYRPFWEEKLPESDGETWVEFVRYRKVQKRKILIAGNHFNRDDKKGLEYLKISQLISDPPDAKAYAL
FFRYTPKLDKTMIGDFLGDPDAFHLQVLREFTNTFEFSGMVLDTALRTYLETFRLPGESQKIQRILEAFSEKLYE
QQSSDLFVSKDAVMILCYSIIMLNTDQHNPQVKKKMTEEDFIRNNRAINGGNDLPREYLSELFQSISNNAISLDQ
SGVPVDMNPSRWIQLINRSKTIQPFILCDFERRLGRDMFAAIAGPSLATLAAVFEHSDEEEILHECIETLFSAAR
IAKYGLEDTLDEILCSFCKFTTLLNPYASSEETLYAFSHDMKPKMAILAVFTIANAFKDSIRGGWRTIFECllkl
krlkllPPSTFEQDNTEPPFKTTESGANLPTQDHKSRNRRHGSGIMAKFLNFTSMESVEEAMNLGLNELEQNMRI
IQQCRIGSIFSTSSNLPVESLLDLGRCIIFAAGGKGQKFNTLIEEEEIIVFCWELITSLTLANAYRLSTFWPRYH
DYMVEVTQFPLFSSQQFCGKAIISLMKMCLKLLSTNELGSSEEILFKSINFMWRTDKEILDICSEFLLQSAIKII
ADHPDKLQTQLGWASVLHLLSVTGRLPETYDRGVNSLIIFMSDATHITKINYPNCIDCAFGFIALKHSLIDNNLK
IMDLMAESAGLLVDWYRAAYMDPsgaggsnsngsateegsKFYNPINLFQRLGESLRKTSLARREEVRNHAVASL
QRSFILAEDLLFIPVNCVNSFYNVIFAMVDDLHEKMLQYSKKTEREAKSMEGTLKLAMEVLTEVYLHFIKPISES
PSFRTFWLGILRRMDTCMKADLGDFGESklpdlipdllkkiVIKMKEKGILRQDGHDDLWDTTYYQIKWIAPALT
DELFPET*
**>Sesin_LOC105159998**
MGRLRLQSSIKAIEEEPEDCEATSSNKAALECMINSEIGAVLAVMRRNVRWGGRYVSGDDQLEHSLIQSLKTLRR
QIFSWQNEWQSVNPSLYLQPFLDVIRSDETGAPITSVALSSIYKILTLDVLDVNTVNVEDAMHMVVDAVTSCRFE
VTDPASEEVVLTKILQVLLACMKSKASVMLSNQHVCTIVNTCFRVVHQAGTKGELLQRIARHTMHELVRCIFSHL
PDVDNTERSLVKGGNSMKNETGGLDADYSFSSKPENGGGTSEYDGQLSAGAFNASSGLISGMMDESNARTDNGKD
AVPYDFHLMTEPYGVPCMVEIFHFLCSLLNVVEQPGMGPRANSLTFDEDVPLFALGLINSAIELGGPAIRHHPRL
LNLVQDELFRNLMQFGLSMSPLILSMVCSIVLNLYQHLRTELKLQLEAFFSCVILRLAQSRFGASYQQQEVAMEA
LVDFCRQKTFMVEMYANLDCDITCGNVFEELANLLSKSAFPVNCPLSSMHILALDGLIAVIQGMAERIGNGSIGF
ERTPVYLEEYTPFWMVKCENYGDPDHWVPFVRRRKYIKRRLMIGADHFNRDPKKGLEFLQGTHLLPEKLDPQSVA
CFFRYTAGLDKNLVGDFLGNHDEFCVQVLHEFAGTFDFQDMNLDTALRLFLETFRLPGESQKIQRVLEAFSERYY
EQSPQILANKDAALLLSYSLIMLNTDQHNVQVKKKMTEEDFIRNNRHINGGNDLPREFLSELYYSICKNEIRTTP
EQGAGFAEMTPSRWIDLMHKSKKTSPYIVADSRAYLDHDLFXMSGPTIAAISVVFDHAEYEDVYQTCIDGFLAVA
KISACHHLEDVLDDLVVSLCKFTTLLNPSSVEEPVLAFGDDAKARMATVTVFTIANRYGDFIRTGWRNILDCILR
LHKLGLLPARVASDAADDSEMSSDPGHGKPLTNSLSSAHMQAIGTPRRSSGLMGRFSQLLSLDTEEPRSQPTEQQ
LAAHQRTLQTIQKCHIDSIFTESKFLQADSLLQLARALIWAAGRPQKGNTSPEDEDTAVFCLELLIAITLNNRDR
IGLLWQGVYEHIANIVQSTVVACALVEKAVXLLRICQRLLPYKENLADELLRSLQLVLKLDARVADQYCEQITQE
VSRLVKANATHIRTPMGWRTIASLLSITARHPDASESGFEALSFIMADGAHLSPANFILCVDAARQFAESRVGQT
DRSVHAVDLMAGSVACLVRWAQDAREAMAEAEAAKLCQDIGEMWLRLVQGLRKVCLDQREEVRNHALLSLQMCLT
GVDEIHLPLGFWPQCFEMVIFTMLDDLAEIAQGNPQTQKEYRNIEGTLVLALKLLTKVFLHLLQELSQLTSFCKL
WRNVIGRMEKYMKLKVKRGEKLQELVPELLKNTLLVMKTKGVLVPTSTLGGDNVWEQTWLHVNKIFPSLQSEVFP
NQDSEPLQSNQSDGGRSPISSETSAQTE*
**>Sesin_LOC105177279**

MTMNLLNQKEVNGLDEQPDYLFKLSGGAVACMVNSEIGAVLAVMRRNVRWGVHYASDDEQIEHSLIISFKELRKK
IFAWQKHWHTIDPVLYLQPFLDVIKSDETGAPITGVALSSVYKILNLQILDSKTVNVDNALHLIVDAVTSCRFEV
TDPASEEVVLMKILQVLLACMKNKASVSLNNHHVCSIVNTCFRIVHQASSKSELLQRISRHTMHELVRCIFSHLP
DLGDERHDLDHGNRPSPNIEDDGTVQNHTLEEKQYVDGYASADSDISKNENAHGKDSTSSAESLMMDPYGVPAMV
EIFHFLCSLLNVMENIEVGPRSNPIAYHEDVPLFALGLINSAIELGGAYFGNHPKLLALIQEELFYNLMQFGLSM
SPLILSTVCSIVLNLYHHLRTKLKLQLEAFISNVLLRIAQSKYGASYQQQEVAMEALIDFCRQPMFVIESYANYD
CDISCSNVFEGLANLLSRSAFPVNSPLSAMNTLALDGLTALVQGMAERIGHDSSSLGEASLELEEYKPFWTVRCH
DYSEPLHWVPFVHNMKNIKKKLMIGVDHFNRDPKKGLEFLQGLNLLPDNLDPRSVACFFRYTTGLDKNIIGDFLG
SHDEFCVQVLHEFARTFDFRDMNLDTALRIFLDTFRLPGESQKIQRVLEAFAESYFEQAPNILANKDAALLLSYS
LILLNTDQHNAQVKKKMSEEDFIRNNRKINGGDDLPRDFLTELYHSICENEIRMVPDQGGAGAVLTRSHWIGLTH
KAKQTSPFIVSESGSHLDFDMFTILSGPAIAAISVVFDHAEQEDVLQSCIDGFLAIAKLSASYNLNEVLDDLIVS
LCKFTTLLHPSFDENSILYFGDDTKAKMATVAVFTIANRYSDHIRLSWRNILDCILSLQKIGLLPARLASDATDE
LESSPDNDQVKISATPSPAPQVTVPSRKSSGLMGRFSLLLSLDAEEPAPQPSEEELAARQRTLQTIQNCHIDTIF
AESKFLQAESLSQLVRALVVAARRPLKGNNSLEDEDTAVFCLELLIAITLNNRDRIMLLWQNVYEHIASVVQSTV
MPCALVEKAVFGLLRICQRLLPYKENLTDELLKSLQLVLKLDARVADAYCEPITQEVMHLVKANAMQIRSHMGWR
TIISLLSITARHPEASESGFETLSYIMSDGAHLSPANYVLCVNAARQFAESRVGHVDRSVKSLDLMAGSVVCLVT
WFHQTKEAAGEEAAVKMSQDILEMWMRLVQSLRKVCVDHREEVRNHAIVLLQRCLTGVEGIHIPTDLWLQCFDLV
IFTLLDELPELAQQQSPKDYRSMEGSMIFSLKLLSKTYLQTLNDLSQSTSFCQLWLKVLGCMEKYMKMRFRGKRS
EKIHELVPELLKNTLLVMKTSGILVPSDPVGGDSFWQLTWLHVKNIAPSLQLEVFPSEELEKLQEKHGKAGCSPL
PDGNGIVPPNETTA*

**>Sesin_LOC105158259**

MKKSKRKELGLSCMLNTEVGALLAVIRRPSESNPSFYHLHEESCDASILQSLKSLRALLFNPQQEWRMVDPSVYL
APFLDVIQSDEIPAAATAVALQSILKILRLGIFDEKTPGAKDVINSAVTAITGCRLEKTNLIAEDAVMMRILQVL
TAIMRHRCSFLLTDHSACTVVNTCFQVVQQSASRGDLLQRSARYTMYELIQIIYSRLPDIEVKDWENSESDTEDN
NMDSGYGIRSAIDIFHFLCSLLNVVDIMETDGPTVQIADESTQVFALYLINSAIELSEDSIGKHPKLLRMIQDDL
FHHLIHYGASSSPLVLSMICSTVLNIYHFLRRSIRLQLEAFFSFVLFKVATPGSTLQIQEVAVEAIINFCRQPTF
IIELYVNYDCDPTFRNVFEETGKLLCKYAFPTGGVSTSIQVQAFEGLSIIFHYIADNIDKEDDSGPSGPYPVEIT
EYRPFWEEKPKEDDMDAWVDYLRIRKVQKRKILIAGSHFNRDEKKGLEYLKISQLISEPPNPQAYAYFFRYTPKL
DKTIIGNYLGDPDAFHLQVLKEFTETFEFAGMVLDTALRTYLETFRLPGESQKIQRVLEAFSDKFYDQQSSDIFA
SKDAVMILCYSIIMLNTDQHNPQVKKKMTEEDFIKNNRSINGGNDLPRDYLSELFQSIATNAISLDQPGSTVEMN
PNRWIQLVNRAKNIQPFILCDFDRRLGRDMFAAIAGPSIATLAAVFEHSDEEEILHECIEALFSIARIAQYGLED
TLDELLCSFCKFTTLLNPYASAEETLYAFSHDIKPKMAMMAVFTIANTFKDSIRGGWRTIIECLLKLKRLKLLPQ
SLFEMDNASNSVTEPPMEKLESGIMPSQDPKFGNRQHSSGIMARFLSFMSMESVEEALNLGLSEFEQNMTIIQQC
RIGSIFSTSSTLPEESLLSIGRCIIFAAAGKGQKFSTPVEEEETVAFCWELITAVALANTHRIPTYWPHYHDYLL
EVAQFPLFSPIPFSEKAIVSLMRICLKILASDQPEGSEELIFKSINLMWRMEKEILETCYEFLLQSAIKIITEYP
TKLQTQLGWMSVLHLLSVTGRHPETYDQGVEALMTLMSNTTQITRTNYTYCIDCAFAFVALRNSRIEKNVKIMDM
LADTASLLVEWYRAGYSDPGSSVSVTSNAGGSFIEEPARVYNPISFFQRLGESLKKTSLARREEVRNHAVISLKK
SFILAEDILFTPITCTNSFYNIIFAMVDDLHEKMLQYSRRENAERETRSMEGTLKLSMEVLTDVYLHFLKPISES
PDFRTFWLGILRRMDTCMKAELAEYGASKMPEVIPDLLRKIVTSMKEKEILTRAGEDDLWDTTFYQIQWIAPALT
DELFPE*

**>Erygu_LOC105967563**

MGRLRLQSSINAIEEEPEECEATSSNKSALGCMINSEIGAVLAVMRRNVRWGGRYVSGDDQLEHTLIQSLKTLRR
QIFSWQHEWQSVNPSLYLQPFLDVIRSDETGAPITGVALSSIYKILTLDVLDVNTVNVDDAMHLVVDAVTCCRFE
VTDPASEEVVLTKILQVLLACMKSKASVMLSNQHVCTIVNTCFRVVHQAGSKGELLQRIARHTMHELVRCIFLHL
PDVDNTEQSLVKGGSSVKNENAGLDTDYNFSGKSENGSGASEFDGQLSSGIYNSASSGLIGGMMDESIPRNDNGK
DAVPYDLHLMTEPYGVPCMVEIFHFLCSLLNVVEHTGMGLRANSLTFDEDVPLFALGLINSAIELGGPAIRHHPR
LLSLIQDELFRNLMQFGLSMSPLILSMVCSIVLNLYQHLRTELKLQLEAFFSCVILRLSQSRFGASYQQQEVAME
ALVDFCRQKTFMVEMYANLDCDITCGNVFEELANLLSKSAFPVNCPLSSMHILALDGLIAVIQGMAERIGNGSVG
FELTPVNLQEYTPFWMVKCENYGDPEHWVPFVRRRKYIKRRLMIGADHFNRDPKKGLEFLQGTYLLPEKLDPQSV
ACFFRYTAGLDKNLVGDFLGNHDEFCVQVLNEFAGTFDFQDMNLDTALRLFLETFRLPGESQKIQRVLEAFSGRY
YEQSPLILANRDAALLLSYSLIMLNTDQHNVQVKKKMTEEDFIRNNRHINGGNDLPRDFLSELYYSICKNEIRTT
PEQGAGFAEMTPSRWIDLMHKSRKTSPYIVSDSRAYLDHDMFAIMSGPTIAAISVVFDHAEYEDVYQTCIDGFLA
VAKISACHHLEDVLDDLVVSLCKFTTLLNPSSVEEPVLAFGDDAKARMATVTVFTIANRYGDFIRTGWRNILDCI
LRLHKLGLLPARVASDAADDSELSSDPGSHGKPLTNSLSSAHMQSIGTPRRSSGLMGRFSQLLSLDTEEPRSQPT
EQQLAAHQRTLQTIQKCHIDSIFTESKFLHADSLLQLARALIWAAGRPQKGSTSPEDEDTAVFCLELLIAITLNN
RDRIGLLWQGVYEHIANIVQSTVVACALVEKAVFGLLRICQRLLPYKENLADELLRSLQLVLKLDARVADQYCEQ
ITQEVSRLVKANATHIRSPMGWRTIASLLSITARHPDASESGFEALTFIMADGAHLSPANFVLCADAARQFAESR
VGQTDRSIQSVDLMAGSVSCLVRWAQDAREGTAEAEAAKLCQDIGEMWLRLVQGLRKVCLDQREEVRNHALLSLQ
MCLTGVDEIHLPLGLWPQCFEMVIFTMLDDLAEIAQGNPQTQKEYRNIEGTLVLALKLLTKVFLHLLNELSQLSS
FCKLWRNVIGRMEKYMKLKVKRGEKLLELIPELLKNTLLVMKTKGVLVPTSTLGGDNVWEQTWLHVNKIFPSLQS
EVFPNLDSEPLQSSPVLGESASDQTN*

**>Erygu_LOC105970904**

MSSLNQLNEVNGLVDRANCLVKPPGGALACMVNSEIGAVLAVMRRNVRWGVQYASDDEQIEHSLIMSFKELRKNI

FSWQNHWHTIDPVLYLQPFLDIIKSDETGAPITGVALSSVYKILNLQILDSETVNVDNALHLIVDAVTSCRFEVT
DPASEEVVLMKILQVLLACMKNKASISLNNHHVCSIVNTCFRIVHQASSKSELLQRISRHTMHELVRCIFSHLPD
LDDKSHELAHGSKSSPNTEDNVAEPSHTLQDKQYADGYTSVESDSKKDDSTHVKDGTSSADSSMMDPYGVPCMVE
IFHFLCSLLNVMENIEVGPRSNPIAYHEDVPLFALGLINSAIELGGASFGNHPKLLALIQDELFYNLMQFGLSMS
PLILSTVCSIVLNLYHHLRTKLKLQLEAFISTVLLRIAESKHGASYQQQEVAMEALIDFCRQPMFVTELYANYDC
DISCSNVFEGLANLLSRSAFPVNSPLSAMNTLALDGLIVLLHGMAERVGHDSSGPGEASLELQEFKPFWTLRCHD
YDEPLHWVPFVHNMKNLKRMLMTGVDHFNRDPKKGLEFLQGLHLLPDKLDPRSVACVFRYTMGLDKNLIGDFLGG
HDEFSVLVLHEFARTFDFQDMNLDTALRIFLETFRLPGESQKIQRVVEAFAESYFEQSSDILVNKDAALLLSYSL
ILLNTDQHNSQVKKKMSEEDFIRNNRNINGGNDLPRDFLSELYHSISENEIRMVPDQGGASGILTRSHWLGLTHK
AKQTSPYIVSDSGSHLDYDMFAILSGPAIAAISVVFDHAEQDYVLQSCIDGYLAIAKLSASYNFGEVLDDLVVSL
CKFTTLLHPSFNERSILYFGDDIKAKMATIAVFTIANRYSDHIRSGWRNILDCILSLQKIGLLPARLASDATDEL
EPPSSDNDQVKTSAAHSPPSQVPPATPARKSSGIMSRFSLLLSYDSEEPAPQPSEEQLAARQRTLQAIQNCHIDT
IFAESKFLQADSLLQLVRALVMAAGRPLKGNNNSLEDEETAVFCLELLIAITLNNRDRIMLLWQNVYEYIANVVQ
STVMPSTLVEKAVFGLLRICQRLLPYKENLTDELLKSLQLVLKLDARVADAYCEQITQEVMHLVKGNAMQIRSHM
GWRTIISLLSITARHPEASEAGFETLSYIMSDGAHLSPANYVLCSNAARQFAESRVGNVERSVRSLDLMSGSVIC
LVTWFYQTKEAAGEEAAIKMSEDILEMWMRLVQALRKVCTDHREEVRNHAITLLQRCLTGVDGIRIPTDLWLQCF
DLVIFTLLDELPEMSNQLSPKDYRFIEGSMILSIKLMSKVFLQSLRDLSRSTSFCQLWLKVLGCMERYMNMRFRG
RRSEKIHELVPELLKNTLLVMKTSGILVPSDPVGGDSFWELTWLHVQKIAPSVQLEVFPGEELEKLQENRAKSGC
SPLSEGNVLVPPNETTA*
**>Erygu_LOC105976035**
MADNNNEEEEDENNKIKSNRKELGLSCMLNTEVGALLAVIRRSPESNYHHHHHPDDACDPTIIQSLKSLRALLFN
PNQPWRSVDPSLYLSPFLDVIQSDDIPSVATAVTLQSILKILRLGIFDTKTPGAKDVINSAVTAITGCRLEKTNP
ITEDAVMMRILQVLTAIMRHEASILLTDNSVCTVVNTCFQVVQQSAGRGDLLQRNARHTMHELIQMIYSRLPDIE
VKDWENSESDTEDNNLDSRYGIRSAVDIFHFLCSLLNVVDIMETDGLTFQIADENTQVFALYLINSAIELSGDSI
GKHPKLLRMIQDDLFHHLIHYGASSSPLVLSMISSTVLNIYHFLRGSIRLQLESFFSFVLFRVAGPTSPLQLQEV
AVEGVISFCRQPSFIIEVYVNYDCDPTFRNVFEEMGKLLCKYAFPTGGLLTSIQVQAFEGLSVIFHYIADNIDKE
DDPSPSGPYPVEISGYRPFWEEKTKKDNDVNNDNWVDYVRVRKVQKRKILIAGSHFNRDDKKGLEYLKYSQLISD
PPDPKAYAVFFRYTPKLDKTMIGDYLGDPEEFHMRVLKEFTDTFEFSGMVLDTGLRTYLETFRLPGESQKIQRIL
EAFSERFYDQQSSEIFVSKDSVMILCYSIIMLNTDQHNPQVKKKMTEEDFIRNNRSINGGSDLPREYLSELFHSI
SGNAISLSDQPGKIIEMSPNRWIQLINRAKIIQPFIMCNFDRRLGRDMFACIAGPAVATVAAVFEQSDDEDMLHE
CIEALFSIARIAQYGLEDILDELLCSFCKFTTLLNPYASADETLYAFSHDMKPKMAILAVFTIANTFKDSIRGGW
RTIIECLLKLKRLKLLPQSIIVETENSPSDTPVEKPSQDPKFGNKRHSSGIMARFLNFMSMESVEEALNLGMGEF
EQNMTVIQQCRIGSIFTTSSTLPEDSLLNLGRCIIFAAAGKGQKFSTPVEEEETVGFCWELIGSIAVANTHRFSS
FWPYYNDYLLEVAQFPLFSPIPFAEKAIVSLMKTCLKILALDKMEGAEELIFKSINLMWRMEKEILETCCEFLLQ
SASTIVTEHPQKLQTQLGWMSVLHLLSATGRHPETYDQGVEALITLMSDKTHITKTNYSHCIDCAFAFVALRNSR
VEKNIKIMDLLAESASLLVEWYRTGYSDPGSCSSIASNASLSSSVEDTRNQNPISLFQRLGESLKKTSLARREEV
RNHAVISLQRSFLLAEELLFTPTNCINSFYNIIFAMVDDLHEKMLQYSRRENAERETRSMEGTLKVSMEVLTEVY
LHFLRPISESPNFRTFWLGLLRRMDTCMKAELGVYGDSNVPEIVPNLLRRIVTSMKEKGILSHAGEDDLWDTTFY
QIQWIAPALTDELFSEGLKTFL*
**>Liphi_05G047100**
MGRLRLQSSINAIEEEPEECETTSSIKTALGCMINSEIGSVLAVMRRNVRWGGRYVSGDDQLEHSLIQSLKTLRR
QIFSWQNEWHSVNPSLYLQPFLEVIRSDETGAPITGVALSSVYKILTLDVLDVNTVNVEDAMHLVVDAVTSCRFE
VTDPASEEIVLTKILQVLLACMKSKASVMLSNQHVCTIVNTCFRIVHQAGTKGELLQRIARHTMHELVRCIFSHL
PDVDNTEQSLVKGGSSINSETAGLDVDYSFSGKSGNGLGTSEYDGQFSSGVFNSASSGLIGGVMDENNATGDNGK
DAVPFDLHLMTEPYGVPCMVEIFHFLCSLLNVVEHTGMSLRANSITFDEDVPLFALGLINSAIELGGPAIRHHPR
LLSLIQDELFRNLMQFGLSASPLILSMVCSIALNLYQHLRTELKLQLEAFFFCVILRLAQSRYGASYQQQEVAME
ALVDFCRQKTFMVEMYANLDCDITCGNVFEELANLLSKSAFPVNCPLSSMHILALDGLIAVIQGMAERIGNGSVG
SELTPVNLEEYTPFWMVKCENYGDPDHWVAFVRKRKYIKRRLMIGADHFNRDPKKGLEFLQGTHLLPEKLDPQSV
ACFFRYTAGLDKNLVGDFLGNHDEFCVQVLHEFAGTFDFQDMNLDTALRLFLETFRLPGESQKIQRVLEAFSGRY
YEQSPQILANKDAALLLSYSLIMLNTDQHNVQVKKKMTEEDFIRNNRHINGGSDLPREFLSELYYSICKNEIRTT
PEQGAGFAEMTPSRWIDLMHKSKKTSPYIVSDSRAYLDHDMFAIMSGPTIAAISVVFDHAEYEDVYQTCIDGFLA
VAKISACHHLEDVLDDLVVSLCKFTTLLNPQSVEEPVLAFGDDAKARMATVTVFTIANRYGDFIRTGWRNILDCI
LRLHKLGLLPARVASDAAEDSEMSSDPTGHGKPLANSLSSAHMQSIGTPRRSSGLMGRFSQLLSLDTEEPRSQPT
EQQLAAHQRTLQTIQKCHIDSIFTESKFLQADSLLQLARALIWAAGRPQKGNTSPEDEDTAVFCLELLIAITLNN
RDRIGLLWQGVYEHIAGIVQSTVVGSALVEKAVFGLLRICQRLLPYKETLADELLRSLQLVLKLDARVADQYCEQ
ITQEVSRLVKANATHIRSPMGWRTIASLLSITARHPDASESGFEALSFIMADGAHLSPANFVLCVDAGRQFAESR
VGQTERSVRAVDLVAGSVSCLVRWAHDAREASMAEDESVKLREDIGEMWMKLVQSLRKVCLDQREEVRNHALLSL
QTCLTGVDGIGLPIAFWPKCFEIVIFTMLDDLAEIAQGNPQTQKEYRNIEGTLVLALKLLAEVFSHLLTELSQLS
SFCKLWRNVIGRMEKYLKLKVKKGEKLQELVPELLKNILLLMKTKGVLVPTSTLGGDNVWEQTWLHVNKIFPALQ
SELFPNQGSEVLQSNQDGEK*
**>Liphi_09G029200**
MKMSLLNQQIEVNGSAEQPNGLYKPSGGSYACMVNSEIGAVLAVMRRNVRWGVHYASDDEQIEHSLIISFKELRK
KIFSWQNHWHSIDPVLYLQPFLDVIKSDETGAPITGVALSSVYKILNLQILDSETVNVDNALNLIVDAVTSCRFE

VTDPASEEVVLMKILQVLLACMKNKASVSLNNHHVCSIVNTCFRIVHQASSKSELLQRISRHTMHELVRCIFSRL
PGLDARGHDLAHESRSSSNTEDGVATQSHTLEEKKNVDGYTSAESDKKKNENVHVKDSTISADPSMMDPYGVPSM
VEIFHFLCSLLNVMENIEVGPRSNPIAYHEDVPLFALGLINSAIELGGASLGNHPKLLVLIQDELFYNLMQFGLS
MSPLILSTVCSIVLNLYHHLRTKLKLQLEAFISGVLLRIAQSKYGASYQLQEVAMEALIDFCRQPTFVTELYANY
DCDISCSNVFEGLANLLSRSAFPVNSPLSAMNTLALDGLIALVQGMAERIGHDSSGSGEASLELEEYKPFWTLRC
DDYGEPRLWVPFAHNMKNIKRKLMVGVDHFNRDPKKGLEFLQGLKMLPDTLEPRSVACFFRYTVGLDKNLIGDFL
GSHDEFCVQVLHEFARTFDFRDMNLDTALRIFLETFRLPGESQKIQRVLEAFSESYFEQSHDILVNKDAALLLSY
SLIMLNTDQHNAQVKKKMSEQDFIRNNRKINGGNDLPRDFLSELYHSICENEIRMVPDQGGAAGVLTRSHWIGLI
HKAKQTSPFILSGSGSHLDYDLFAICSGPAIAAISVVFDHAEQEDTLQSCIDGFLAIAKLSASYNLGEVLDDLVV
SLCKFTTLLHPSFSEKSILYFGDDIKAKLATVAVFTIANRYSDHIRSGWRNILDCILSLHKIGLLPARLASDATD
DPDNDQVKISTTHSPASQVPVVTPSRKSSGIMGRFSLLLSLDSEEPAPQPSEEQLAARQRTLQTIQNCHIDSIFA
ESKFLQAESLLQLIRALVSAAGRPLKGNNFMEDEDMAVFCLELLIAITLNNRDRIMLLWQYVYEHIANVVQSTVM
PCALVEKAVFGLLRICQRLLPYKENLTDELLKSLQLVLKLDARVADAYCEQITQEVMHLVKANAMQIRSHMGWRT
IISLLSITARHPEASEAGFETLSYIMSDGAHMSPANYVLCVNAARQFAESRVGNVERSVKSLDLMSGSVVCLVTW
FYQTKEAAGEEAAVKMAQDILEMWMRLVQGLRKVCIDQREDVRNHAIMLLQRCLTGVDGINIPTDLWLQCFDLVI
FTLLDELPEMAQQQTPKDYRSMEGSMVLSLKLLSKAFLQSLIDLSRSTSFCQLWIKVLGCLEKYMSMRFGKRSE
KIHELIPELLKNTLLVMKTSGILVPSDPIGGDSFWQLTWLHVKNISPSLQLEVFPSEDGKAGCSPLPDGNVLVPT
KETPAA*
**>Liphi_08G097200**
MDNNKDDENKKSNLTKSKRTEIGLSCMLNTEVGALLAVIRRAPESGPGFYHDHYDASVLQSLKSLRALLFNPQQE
WRTIDPSVYVSPFLDVIQSDDIPAAATAVALQSVLKILRLGIFDEKTPGAKDMMNSSVTAITGCRLEKTNSITED
AVMMRILQVLTAIMRHRASILLTDQSVCILVNTCFQVVQQSAHRGDLLQRSARYTMHELIQIIYWRLPDMEVKDW
ENSESDTEDNNPDSRYGISSAVDIFHFLCSLLNVLDVMETDGLTFHIADENTQVFALYLINSAIELSGDSIGKHP
KLLRMIQDDLFHHLIHYGASSSPLVLSMISSTVLNIYHFLRKSIRLQLEAFFSFVLFRAASPGVPLQLQEVAMEG
IVNFCRQPTFIIEVYVNYDCDPTFRNVFEETGKLLCKYAFPTGGHLTSIQAQAFEGLAVIFHYIADNIDKEDDPS
PSGPYPVEISEYRPFWEEKPKEDDHEAWVDYVRVRKVQKRKILIAGNHFNRDDKKGLEYLKVSQLISDPPDPKAY
AIFFRYTPKLDKTMIGDYLGDPDEFHIQVLKEFSETFEFTGMVLDTALRTYLETFRLPGESQKIQRILEAFSEKF
YDQQSSDMFASKDAVMILCYSIIMLNTDQHNPQVKKKMTEEDFIRNNRSINGGSDLPREYLSELFNSISTNAISL
DQSGTVVVDMNPNRWIQLVNRAKTMQPYILCDFDRRLGRDMFAAIAGPSVATVAAVFEHSDDEEILHECIEALFS
IARIAQYGLEDILDELLCSFCKFTTLLNPYASAEETLYAFSHDMKPKMAILAVFTIANSFKDSIRGGWRTIIDCL
LKLKRLKLLPQSVIEPENASNLLPTEKLDSGGFFPSQDSKFGNKKRHSSGIMARFLNFMSMESVEEALNLGLSEF
EQNVSIIQQCRIGNIFCNSSSLPEDSLLNLGRCIIFAAAGKGQKFSTPVEEEETVGFCWELISTLAFANAHRFST
FWPHYHDYLLEVAQFPLFSPIPFSEKAIVVLMKMCLKLLASDKLEGAEELVFKSINLMWRMEKEILETCCESLLT
SASKIITEYPEKLQTPLGWMSVLHLLSVTGRHPETYDKGVETLIGLMSDGAHITRTNYSYCIDCAFAFVALRNSR
VDKNVKIMDLLAESARLLVEWYKAGYTDPGSFTSIASNASGTSSVEESARIYNPISLFQRLGESLKKTSLARREE
VRNHAVVSLQRSFVLAEDILFTPTNCINSFYNIIFAMVDDLHEKMLQYSRRENAERETRSMENTLKLSMEVLSDV
YLHFLKPISESPNFRTFWLGILRRMDTCMKADLGEYGESKMPDVIPDLLRRIVTSMKEKAILTRTGDDDLWDTTF
YQIQWIAPALTDELFPEGQKNNG*
**>Calam_14G038300**
MGRLRLQSSINAIEEEPEECEATSSNKAALGCMINSEIGAVLAVMRRNVRWGGRYVSGEDQLEHSLIQSLKTLRR
QVFSWQHEWQTVNPSLYLQPFLDVIRSDETGAPITGVALSSVYKILTLDVLDVNTVNVEDAMHLVVDAVTSCRFE
VTDPASEEVVLTKILQVLLACMKSKASVMLSNQHVCTIVNTCFRVVHQAGTKGELLQRIARHKMHELIRCIFSHL
PDVDNTERSLVKGGSSNKNETAGLDTDYSFSSKAENGSGTSEYDGQLSSGVFSSAASGLISSVMDESNARNDNGK
DVVPYDLHLMTEPYGVPCMVEIFHFLCSLLNVVEHTGMGLRANSITFDEDVPLFALGLINSAIELGGPAIRHHPR
LLSLIQDELFRNLMQFGLSMSPLILSMVCSIVLNLYQHLRTELKLQLEAFFSCVILRLAQSRYGASYQQQEVAME
ALVDFCRQKTFMVEMYANLDCDITCGNVFEDLANLLSKSAFPVNCPLSSMHILALDGLIAVIQGMAERIGNGSVG
SERTPVSLEEYTPFWMVKCENYGDPDHWVPFVRRRKYIKRRLMIGADHFNRDPKKGLEFLQGTHLLPEKLDPQSV
ACFFRYTAGLDKNLVGDFLGNHDEFCVQVLHEFAGTFDFQDMNLDTALRLFLETFRLPGESQKIQRVLEAFSGRY
YEQSPQILANKDAALLLSYSLIMLNTDQHNVQVKKKMTEEDFIRNNRHINGGNDLPREFLSELYYSICKNEIRTT
PEQGAGFAEMTPSRWIDLMHKSKKTSPYIVADSRTYLDHDMFAIMSGPTIAAISVVFDHAEYEDVYQTCIDGFLA
VAKISACHHLEDVLDDLVVSLCKFTTLLNPSSVEEPVLAFGDDAKARMATVTVFTIANRYGDFIRTGWRNILDCI
LRLHKLGLLPARVASDAADDSEMSSDPGHGKPLTSSLSSAHMQTIGTPRRSSGLMGRFSQLLSLDTEEPRSQPTE
QQLAAHQRTLQTIQKCHIDSIFTESKFLQADSLLQLARALIWAAGRPQKGNTSPEDEDTAVFCLELLIAITLSNR
DRIGLLWQGVYEHIAGIVQSTVVACALVEKAVFGLLRICQRLLPYKENLADELLRSLQLVLKLDARVADQYCEQI
TQEVSRLVKANATHIRSPMGWRTIASLLSITARHPDASESGFEALSFIMADGAHLSPANFVLCVDAARQFAESRV
GQTDRSVHAVDLMAGSVSCLVRWAQDAREAMAEAEAAKLCQDIGEMWLRLVQGLRKVCLDQREEVRNHALLSLQM
CLTGVDEICLPLGFWPQCFEMVIFTMLDDLAEIAQGNPQTQKEYRNIEGTLVLALKLLTKVFLHLLHELSQLSSF
CKLWRNVIGRMEKYIKLKVKRGEKLQELVPELLKNTLLVMKTKGVLVPTSTLGGDNVWEQTWLHVNKIFPSLQSE
VFPNQDSEPSQSNQADAGKSPTSSEIASAQNE*
**>Calam_09G131100**
MSLLNQQNEVNGSVEQPNCSFKLSGGALACMVNSEIGAVLAVMRRNVRWGVHYASDDEQIEHSLIISFKELRKKI
FSWQNHWHSIDPVLYLQPFLDVIKSDETGAPITGVALSSVYKVLNLQILNAKTVNVDNALHLIVDAVTSCRFEVT
DPASEEVVLMKILQVLLACMKSKASVNLNNHHVCSIVNTCFRIVHQASSKSELLQRISRHTMHELVRCIFLHLPD

LDDRRHELARGERSSPNIEDDAVAQSRTVEGKPKADGYTSAESENKKIENAYGEDSTFSVDSSMMDPYGVPSMVE
IFHFLCSLLNVMENIEIGPRSNPIAYHEDVPLFALGLINSAIELGGASFGNHPKLLALIQEELFYNLMQFGLSMS
PLILSTVCSIVLNLYHHLRTKLKLQLEAFISNVLLRIAQSKYGASYQQQEVAMEALIDFCRQPMFVTELYANYDC
DISCSNVFESLANLLSRSAFPVNSPLSAMNTLALDGLIVLVQGMAERIEHDSSSSAEASPELEEYSSFWTRRCDD
YGEPLHWVPFVHNMKNIKRKLMIGVDHFNRDPKKGLESLQGLKLLPDKLDPRSVACFFRYTMGLDKSIIGDFLGS
HDEFCVQVLHEFARTFDFRDMNLDTALRIFLETFRLPGESQKIQRVLEAFSESYFEQSPNIFVDKDAALLLSYSL
ILLNTDQHNAQVKKKMSEDDFIRNNRKINGGNDLPRDFLSELYHSICENEIRMVPDQGGAAAVLTRSHWIGLTHK
AKQASPFIVSDSGSHLDYDMFAILSGPAIAAISVVLDHAEQEDVLQSCIDGFLAIAKLSASYNLGEVLDDLVVSL
CKFTTLLHPSFNEKSILYFGDDVKAKMATVAVFTIANHYSDNIRLSWRNILDCILSLHKIGLLPARLASDATDEL
ESSPDNDQVKTSATQSPAFQAPATIPSRKSSGIMGRFSLLLSLDAEEPAPQPSEEQLAARQRTLQTIQNCHIDSI
FAESKFLQAESLSQLVRALVVAAGRPLKGNNSLEDEDTAVFCLELLIAITLNNRDRIMLLWQNVYEHIANVVQST
AMPCTLVEKAVFGLLRICQRLLPYKENLTDELLKSLQLILKLDARVADAYCEQITQEVMHLVKANAMQIRSHMGW
RTIISLLSITARHPEASEAGFETLSYIMSEGAHLSPANYVLCVNAARQFAESRVGHVDRSVRSLDLMSGSVVCLV
TWFYQTKEAAGEEAAVKMSQDILEMWMRLVQGLRKVCVDHREEVRNHAIMLLQRCLTGMDGIHIPINLWLQCFDL
VIFTLLDELPEMAQQQSPKDYRSMEGSMILSLKLLSKAFLQSLLALSQSTSFCQLWLKVLSCMERYMKIRFRGKR
SEKFHELVPELLKNTLLVMKTSGILVPSDPVGGDSFWQLTWLHVNNVAPSLQLEVFPSEELETLKEKHAKAGCSP
LPDGSVLVPPNETAA*

**>Calam_12G060000**

MDNMEEDENNKSSMPKSKRKDLGLSCMLNTEVGALLAVIRRAPESNPGFYHHLDENCDASILQSLKSLRALLFNP
QQEWRMIDPSVYISPFLDVIQSDDIPAAATAVALQSVLKILRLGIFDEKTPGARDVINSAVAAITCCRLEKTNPV
TEDAVMMRILQFLAAIMRHGASVLLTDHSVCTVVNTCFQVVQQSANRGDLLQRSAKYTMHELIQIIYSRLPDIEV
KDWENSESDTEDNNLDSGYGIGSAVDIFHFFCSLLNVVDIMESDGLTYPTTDENTQAFALYLINSAIELSGDSIG
KHPKLLRMIQDDLFHHLIHYGVSSSPLVLSMISSTVLNLYHFLRRSVRLQMEAFFSFVLFRVASPGNPLQLQEVA
VEGIINFCRQPNFIIEVYVNYDCDPTLRNVFEETGKLLCKYAFPTGAFLTSIQVQAFEGLAVIFHYIADNIDKED
DSSRSRPYPVEISEYRPFWEEQPKGDDNDNETWIDYVRFRKVQKRKILIAGSHFNRDDKKGLEYLKISQLISEPP
DPKAYAYFFRYTPKLDKKMIGDYLGDPDGFHIQVLKEFTNTFEFTGMILDTALRTYLETFRLPGESQKIQRIVEA
FSERFYDQQSSDFFANKDSVMICCYSIIMLNTDQHNPQVKKKMTEEDFIRNNRSINEGRDLPREYLSELYQSIST
NAISLDDSGAIVEMNPNRWIQLINRAKIIQPFILCDFDRRLGRDMFAAIAGPSVATVAAVFEHSDEEEILQECIE
VLFSIARIAQYGLEDTLDELLCSFCKFTTLLNPYASAEETLYGFSHDMKPKLAILAVFTIANTFKDSIRGGWRTI
IDCLLKLKRLKLLPQSIMEPDNASNSLSDVPTGKSEAGVIFPSQDPKFCNRQQNSGITARFLNFMSMESVEEALN
LGLSEFEQNMAIIQQCRIGSIFSTSSSLPEDSLLNLGRCIIFAAAGKGQKFSTPVEEEETVGFCWELISSMALAN
THRLSTFWPYYHEYLLEVAQFPLFSPIPFSEKAIVSLMKMFLKLLASDQLEGGSEELIFKSINLMWRMEKEILET
CCEFLLQSASKIITEYPRKLQTQLGWMSVLHLLSVTGRHPETYDRGVETLIQLMSDGIHITRTNYSYCIDCAFAF
VALRNSRVEKNVKIMDLLVESASLLVEWYKGGYSDPGSSSISIGSNGSSPSVEESGRIYNPISFFQRLGESLKKT
SLARREEVRNHAVMSLQRSFLLAEDLLFTPTSCINAFYNVIFAMVDDLHEKMLQYSRRENAERETRSMEGTLKLS
MEVLTHLYLHFLKPISESNNFRTFWLGILRRMDTCMKADLGEYGESKMPHVIPDLLKEIVMSMKEKEILRITAED
DLWDTTFYQIQWIAPALTDELFPEGLKNND*

**>FraexGNOMa_chr7**

MGRLRLQSSINAIEEEPEDCGITSPSEAALASMINSEIGAVLAVMRRNVRWGGRYVSGDDQLENSLIQSLKALRK
QIFSWQHQWQSVNPSLYLQPFLDVIRSDETGAPITGIALSSVYKIltldvldlntvdvedAMHLVVDAVTSCRFE
VTDPASEEVVLMKILQVLLACMKSKASVMLSNQHVCTIVNTCFRVVHQAGTKSELLQRIARHTMHELVRCIFSHL
QNVDNTERSLVKGGSSSKNEVCIFDNDYSFSSKSENGNRTSEYDGLPssgvfnstssssslvsGMMDETVGTDNGK
DAVPYDFHLFTEPYGTPCMVEIFHFLCSLLNVVEHVGMGPRANTISFDEDAPLFALGLINSAIQLGGPAIRNHPR
LLSMVQDELFRNLMQFGLSMSPLILSMVCSIVLNLYQHLRTELKLQLEAFFSCVILRLAQSCYGASYPQQEVTME
ALVDFCRQKTFMAEMFANLDCDITCSNVFEDLANLLSKSAFPDNCPLSSMHILALDGLIAVIQGMAERIGNGSIN
SELPPVNLEEYTPFWMVKCENYGDPDHWVPFVRKRKYIKRRLMIGADHFNRDPKKGLEFLQGTHLLPEKLDPQSV
ACFFRYTAGLDKNLVGDFLGNHDEFCVQVLHEFAGTFDFQDMNLDTALRLFLETFRLPGESQKIQRVLEAFAERY
YEQSPQILANKDVAFVLSYSLIIILNTDQHNAQVKKKMTEEDFirnnrdinggnnLPREFLSELYYSICKNEIRTM
PEQGAGFAEMTPSRWTDLMHKSKKTSPYIVADSRAYLDHDMFAIMSGPTIAAISVVFDYAEHEDVYQTCIDGFLA
VAKISACHHledvlddlvvSLCKFTTLLNPSSVEEPVLAFGDDAKARMATVTVFTIANRYGDFIRTGWRNILDCI
LRLHKLGLLPARvasdaaddsevssePGQGKLLTNSLSSAHMQSIGTPRRSSGLMGRFSQLLSLETEEPRSQPTE
QQLAAHQRTLQTIQKCHIDSIFTESKFlladsllqlaqaliwaaGRPQKGNSSPEDEDTAVFCLELLIAITLNNR
DRIGLLWQGVYEHISNIVQSTVVACALVEKAVFGLLRICQRLLPYKENLADELLRSLQLVLKLDARVADQYCEQI
TQEVSRLVKANATHIRSQMGWRTIASLLSITARHPDASEPGFDALSFIMTDGAHLSPANFIFCIDAARQFAESRL
GQTDRSLRAVDLMAGSVACLGQWARDARDAMAEVEAVKLTQDIGEMWLRLVQELRKVCLDQREEVRNHALVSLQM
CLTGVDEIHLPLGLWSQCFEMVIFTMLDDLAEIAQGNSQTQKECRNIEGTlvlalkllvkvflhllhelsplssF
CKLWRNVIGRMEKYMKLKVKRSEKLQELVPELLKNTLLVMKTNGVLVPTSTLGGDSIWEQTWLHVNKMFPSLQSE
IFPQGSETVLSNPGDLGRSAEDTETDDGLQHISNLF*

**>FraexGNOMb_chr15**

MGRLRLQSSINAIEEEPEDCGITSPSEAALASMINSEIGAVLAVMRRNVRWGGRYVSGDDQLENSLIQSLKALRK
QIFSWQHQWQSVNPSLYLQPFLDVIRSDETGAPITGIALSSVYKIltldvldlntvdvedAMHLVVDAVTSCRFE
VTDPASEEVVLMKILQVLLACMKSKASVMLSNQHVCTIVNTCFRVVHQAGTKSELLQRIARHTMHELVRCIFSHL
QNVDNTERSLVKGGSSSKNEVCIFDNDYSFSSKSENGNRTSEYDGLPssgvfnstssssslvsGMMDETVGTDNGK

DAVPYDFHLFTEPYGTPCMVEIFHFLCSLLNVVEHVGMGPRANTISFDEDAPLFALGLINSAIQLGGPAIHDHPR
LLSLVQDELFLNLMQFGLSMSPLILSMVCSIVLNLYQHLRTELKLQLEAFFSCVILRLAQSCYGASYPQQEVTME
ALVDFCRQKTFMAEMFANLDCDITCSNVFEDLANLLSKSAFPDNCPLSSMHILALDGLIAVIQGMAERIGNGSIN
SELPPVNLEEYTPFWMVKCENYGDPDHWVPFVRKRKYIKRRLMIGADHFNRDPKKGLEFLQGTHLLPEKLDPQSV
ACFFRYTAGLDKNLVGDFLGNHDEFCVQVLHEFAGTFDFQDMNLDTALRLFLETFRLPGESQKIQRVLEAFAERY
YEQSPQILANKDVAFVLSYSLIILNTDQHNAQVKKKMTEEDFirnnrdinggnnLPREFLSELYYSICKNEIRTM
PEQGAGFAEMTPSRWTDLMHKSKKTSPYIVADSRAYLDHDMFAIMSGPTIAAISVVFDYAEHEDVYQTCIDGFLA
VAKISACHHledvlddlvvSLCKFTTLLNPSSVEEPVLAFGDDAKARMATVTVFTIANRYGDFIRTGWRNILDCI
LRLHKLGLLPARvasdaaddsevssePGQGKLLTNSLSSAHMQSIGTPRRSSGLMGRFSQLLSLETEEPRSQPTE
QQLAAHQRTLQTIQKCHIDSIFTESKFlladsllqlaqaliwaaGRPQKGNSSPEDEDTAVFCLELLIAITLNNR
DRIGLLWQGVYEHISNIVQSTVVACALVEKAVFGLLRICQRLLPYKENLADELLRSLQLVLKLDARVADQYCEQI
TQEVSRLVKANATHIRSQMGWRTIASLLSITARHPDASEPGFDALSFIMTDGAHLSPANFIFCIDAARQFAESRL
GQTDRSLRAVDLMAGSVACLGQWARDARDAMAEVEAVKLTQDIGEMWLRLVQELRKVCLDQREEVRNHALVSLQM
CLTGVDEIHLPLGLWSQCFEMVIFTMLDDLAEIAQGNSQTQKECRNIEGTlvlalkllvkvflhllhelsplssF
CKLWRNVIGRMEKYMKLKVKRSEKLQELVPELLKNTLLVMKTNGVLVPTSTLGGDSIWEQTWLHVNKMFPSLQSE
IFPQGSETVLSNPGDLGRSAEDTETDDGLQHISNLF*

**>FraexGNOMc_chr8**

MRCRNGQNEAYGLVKPTCAFKPSGGALACIVNSEIGAVLAVMRRNVRWGVQYATDDEQMEHSLIKSFKELRKQIF
SWQHEWHSVDPVVYLQPFLDVIKSDETGAPITGVALSSVYKILNLEILELETVNVDNTLHLIVDAVTSCRFEVTD
PASEEVVLMKILQVLLACMKNRASVNLNNHHVCSIINTCFRIVHQASSKSELLQRISRHTMHDLIRCIFSHLPNI
DNTQHILAQGSRSYPNTEVENQTLEDKQYDNGHNSAESDNNKMEGDSNVAAYTIAVEMGDKIKEIEQGKATSPNG
DFSMMDPYGVPCMVEILHFLCSLLNVMENIETGPRTNPIAYDEDVPLFALGLINSAIELGGATFESHPKLLSLIQ
DELFYNLMQFGLSMSPLILSTVCSIVLNLYHYLRTKLKLQLEAFISCVLLRILQSKYGASYQQQEVAMEALVDFC
RQPTFVAEMFANYDSDISCSNVFECLSNLLSKSAFPVNSPLSAMNTLSLDGLVALVQGIADRIGHDSTALDEGEP
ELQEYYPFWTLQYESYGEPHQWVPFIHRMKHIKRKLMIGVDHFNREPKKGLEFLQALHLLPDPLDPQSVAHFFRY
TAGLDKAVVGDFLGSHDEFYIQVLHEFARTFNFQGLNLDAALRIFLETFRLPGESQKIMRVLEAFAESYYFQSPN
IFANADAALVLSYAMIMLNTDLHNAQVKKKMREEDFirnnrringgnDLPREYLSELYHSILDNEIRMVPEQAAG
VAVMTRSHWIGLVHKARETAPFFVSESGCDFDYDMFAILSGPSIAAISVVFKHAEQEDVLQSCIDGFLAIAKLSA
SYNNGEVLNDLVVSLCEFTTLMHPSFVEESILDFGDDTKAKMATVAVFTIANRYGDHIRSGWRNIVECILSLQKL
GLLPTRlasdaaddldssPVLGHatpaatsspasqvpatVPARKSSGLIGRFSLLLSLDteepapqpteeqlaaR
QRTLQMIQNCHIDSIFAESKFLQGESLAQLVHALILARGCSSKGNNYLEDEDTAVFCLELLIAITLNNRDRIMLL
WQNVYEHIANVVQSTLMPCALVEKAVFGLLRICQRLLPYKENLTDEllkslqlvlklDARVADAYCEQITQEVMH
LVKANAMQIRSHMGWRTIISLLSITARHPEASDVGFETLSFIMSDGSYLLPTNYVLCLNAARQFAESRVGHVDRS
VRSLDLMAGSIACLVRWFQKTKEAVDEEEAAIKMSQDILEMWMRLIQGLKKVCLDSREEVRNHAILSLQRCLTGS
DGIRIPNDLWLQCFDQVIFTLLDELLEAAQQNSPKEYRSMEGSMIVSLKLLSKAFLQSLQEISQSTSFCQLWLRV
LSCMDKYMKMKIRGKRSEKIHELVPELLKNTLLIMKSSGILVPSDPVGGDSFWQLTWLHVKNLAPSLELEVFPSe
eleklkekHVKPGCSPLPDGNILVPPNETTA*

**>FraexGNL2_chr1**

MDNTQVSKELGLSCMLNAEVGAVLAVIRRGPESTSQFHHHPDENCDVSVLQSLKSLRAFLFNPRQEWRRIDPSLY
LSPFLDVIQSDDIPATATAVALQSILKILRLGIFDERTVDAKGAINSAVTAVTGCRLERTNSVSEEAVMMKILQV
LTAIMRHRASVLLTDHAACTIVNTCFQVVQQLASRGDLLQRSARYTMHELIQIIYSRLPDIEVKDWENSDSDTED
NNLDLGYGIRAAIDIFHFLCSLLNVVDVMETEGLISQTADENVQLFALVLINSAIELSGDSIGKHPKLLRMIQDD
LFHHLIHYGASSSPLVLSMICSTVLNLYHFHRRSIRLQLEAFFSFVLFRIASPGSSLQLQEVAVEGLINFCRQPT
FIIEVYVNYDCDPTFRNVFEETGKLLCKYAFPTGGLATSIQIQALEGLVVIFHNIADHIDQENDPSPTGPYPVEI
SGYRPFWEEKSKDDDVETWVDYVRVRKVQKRKILIAGNHFNRDDKKGLEYLKISKLIADPPDPKAYAIFFRYTPG
LDKTVIGDYLGDPDDFHIKVLKEFSDTFQFTGMILDTALRTYLEAFRLPGESQKIQRILEAFSERFYDQQSSEIF
VTKDAVFILCYSLIMLNTDLHNPQVKKKMTEEEFIRNNRKINGGNDLPREYLSELFHSILTNAIVLQGQSGAGAP
VEMNPNRWIQLINRGMFIQPFILCEFNRRLGRDMFASIAGPSVATLAAIFEHSDEEEILHECIEAMFSIARIAQY
GLEDTLDELLSSFCKFTTLLNPYASAEETLYAFGNDMKPKMATLAVFTIANTFKNSIRGGWRTIIDCllklkrlk
llPQSVIEPDNASDSGSDFSANERLESGVVFPSHDPKFGGRHSSYGIMGRFLNFLSLESVEDALNNGVSEFEQNL
KFIQQCRIGSIFSTSASLPDESLLNLGRCLIFAAAGKGQKFSTPIEEEDTVGFCWDLICAIASANTHRFSTFWPH
YHDYLLEVTQLPLFSPTPFGEKAIVAQMKMCLKLLALGQFDKVEELLLKSIVLIWTMEKDILDTCCEFLVQSASK
IIMEYPEKLQTHLGWKSFLHLLSVTGRHPETYDKGVEALITLMSDETHISPTNYSCCIDCAFGFVALRNSSVEKN
TKIMDLMANSAKLLVQWHRSGYSDTGSYASFISNTSRSSAEENARSHNLMTFFQRLGESLRKTTLARREEVRNHA
VTSLQKSFIQAEELLFTPANCINSFYNVIFAMVDDLHEKMLEYSRRENAERETRSMEQTLKISMEVLTEVYLHFL
KPISESPSFRTFWLGILRRMDTCMKVELGVYGESKLPDAIPDLLRKIVTTMRNEQILIQKGEDDLWETTFYQIQW
IAPALTDELFPKGTG*

**>AndpaGNOM_chr19**

MGRLRTQSSIKAIEEEPEDCEATSSDKAAVECMINSEIGAVLAVMRRNVRWGGRYVSGDDQLEHSLIQSLKALRR
QIFSWQNDWQSVNPVLYLQPFLDVIRSDETGAPITGVALSSVYKILTLDVLEVNTINVEEAVHMVVDAVTSCRFE
VTDPASEEVVLTKILQVLLACMKSKASVMLSNQHVCTIVNTCFRIVHQAGTKGELLQRIARHTMHELIRCIFSHL
PNVDNTEQSLVKGVMSIKNESGGLNPDYSFNSISENGTGASEYDGQLSSGGFNSASSGLIGGVMDESIGLTGNGK
DSVPYDLHLMTEPYGVSCMVEIFHFLCSLLNVVEHTGVGGRANSITFDEDVPLFALGLINSAIELGGPAIRHHPR

LLSLVQDELFRNLMQFGLSASPLILTMVCSIVLNLYRRLRTELKLQLEAFFSCVILRLAQSRFEASYQQQEVAME
ALVDFCRQKDFMVEIYANLDCDITCGNVFEDLANLLSKSAFPVNCPLSSMHILALDGLIAVIQGMAERIGNGSVG
FEPNPVHLEEYTPFWMVKCENYGDPDYWVPFVRRRKYIKRRLMIGADHFNRDPKKGLEFLQGTHLLPEKLDPQSV
ACFFRYTAGLDKNLVGDFLGNHDEFCVQVLHEFAGTFDFQDMNLDTALRLFLETFRLPGESQKIQRVLEAFSERY
YEQSPQILANKDAALLLSYSLIMLNTDQHNVQVKKKMTEEDFIRDNRHINGGNDLPREFLSELYYSICKNEIRTT
PEQGAGFAEMTPSRWIDLMHKSKKTPPYIVADSRAYLDHDMFAIMSGPTIAAISVVFDHADKDVYQTCIDGFLAV
AKISACHHledvlddlvvSLCKFTTLLNPASVEEPVLAFGDDAKARMATVTVFTIANRYGDFIRTGWRNILDCIL
RLHKLGLLPARVASDAADDSEMAVDAGHGKPLTNSLSSAQMQTIGTPRRSSGLMGRFSQLLSLDTEEPRSQPTEQ
QLAAHQRTLQTIQKCHIDSIFTESKFLHADSLLQLARALIWAAGRPQKGNTSPEDEDTAVFCLELLIAITLNNRD
RIGLLWQGVYEHIANIVQSTVVACALVEKAVFGLLRICQRLLPYKENLADELLRSLQLVLKLDARVADQYCEQIT
QEVSRLVKANATHIQSPMGWRTIASLLSITARHPDASESGFEALSFIMADGAHLSPANFILCVDAARQFAESRVG
QADRSVHAVDLMAGSVSCLVRWAQDAREAMSESEAAKLCQDIGEMWLRLVQGLRKVCLDQREEVRNHALLSLQRC
LTGVDEICLPLGFWPQCFEMVIFTMLDDLAEIAQGNAQTQKEYRNIEGTLVHALKLLTKVFLHLLNELSQLSSFC
KLWRNVIGRMEKYMKLKVKRGEKLQELVPELLKNILIVMKAKGVLVPTSTLGGDNVWEQTWLHVNKIFPSLQSEV
FPNQDLDQATVGRSPVSNESVASQNEIAVRPHNGMSSDLLFIVILRSFFHCGTKFTQIFIT*

**>AndpaGNL1_chr2**

MAMNYVHQQEVNGSVDHSNCLIAPSGGAFACMVNSEIGAVLAVMRRNVRWGVHYASDDEQIEHSLITSFKELRKK
IFSWQNRWHSIDPVLYLQPFLDVIKSDETGAPITGVALSSVYKILNLQILDSKTVNVDNALHLIVDAVTSCRFEV
TDPASEEVVLMKILQVLLACMKNKASVSLNNHHVCSIVNTCFRIVHQASSKSELLQRISRHTMHELVRCIFPTFL
TWTIKDRTFLMRKALPPNXIVHLLKIQFLLQESVATQSHTLEEKQYVDGYASAKSDSHVLETRQGKDGTLTSDST
MMEPYGIPAMVEIFHFFCSLLNVMENIDIIPRSNPIAYHEDVPLFALGLINSAIELGGSSFGNHPKLLALIQEEL
FYNLIQFGLSMSPLILSTVCSIVLNLYHHLRTKLKLQFEAFVSSVLLRTAQSKYGASYQQQEVAMEALIDFCRQP
MFVPELYANYDCDISCTNVFEDLANLLSRSAFPVNSPLSAMNILALDGLIALVQGMADRVSHESLGSGEASQELG
EYKQFWTLRCNDYCDPHQWVPFVHHMKNIKKKLMIGVDHFNRDPKKGLEYLQGMKLLPEKLDPKSVALFFRYTIG
LEKNLIGDFLGSHDEFCVQVLHEFARIFDFRDMHLDTALRIFLETFRLPGESQKIQRVLEAFAESYYEQSPNILV
NKDAALLLSYSLIMLNTDQHNAQVKKKMSEEDFIRNNRKINGGNDLPREFLSELYHSICENEIRMVPDQGGAIGL
LTQSHWIGLTHKAKQTSPFIISGSGSQLDYDMFSILSGPAIAAISVVFDHAEQEDVLQLCIDGYLAIAKLSASYG
LGEVLDDLVISLCKFTTLLHPSLSEKLIMYFGDDTKAKMATVAVFTIANRYSDHIRSGWRNILDCILSLHKLGVL
PSRLATDATDELESSNDNDQVKKLGTNSLsspapaaapsRKSSGFMGRFSLLLSLDADEPAPQPSEEQLAARQRT
IQTIQNCHIDSIFAESKFLQAESLTQLVRALVMSAGRPLKGNNSAEDEETAAFCLELLIAITLNNRDRIMLLWQN
VYEYIASVVQSTVMPCTLVEKAVFGLLRICQRLLPYKENLTDELLKISATRLETRCRVADAYCEQITQEVMHLVK
ANAMQIRSHMGWRTIISLLSITARHPEASEAGFETLSYIMSEGVHLSPANYVLCVNAARQFAESRVGHVERSVKS
LDLMADSIGCLVKWFHQTKEAAGEEAAMKMAEDILEMWMRLVQSLRKVCVDHRKVRNHAILQLQRCLTGVDGIRI
PTELWLQCFNLKIFTLVEDLRAMAQQHSPKDFRSIEGSIVLSLKLLSKHSYNRLPDISQDQSFCQLWVKVLNCPE
KAHEHRFRGKRSEKIHELVPELLKNTLLVMKTSGILTPSDPVAGDSFWQLTWLHVKNIAPALHVEVFPTEELENM
QDGCSPMPDGNVHVSQNETAAEDIYI*

**>AndpaGNL2_chr10**

MAEDEQTRPKSRRKDLGISCMLNTEVGALLAVIRRGPESSPGFYHHPDDLCDATllqslrslrallFNPEWRMLD
PSVYITPFLDVVQSDDIPAAATSVALQAILKILRLGIFDKKTPGAKDAINTVVTAITGCRLEKTNPVAEEAIMMR
ILQVLTAVMRHPASVLLTDHSTCTVVNTCFQVVQQSATRGDLLQRSARYTMHELIQIIFSRLPEIevrdsedges
dsddneldsGYGIRSAVDIFHFLCSLLNVVGVMENEGVTIQVADENIQVFALYLINSAMELSGDSMGKHSKLLRM
IQDDLFHHLIHYGTCCSPLVLSMICSTVLNIYHFLRSSMRVQLEAFFSYVLLRVASPGSPIQLQEVAVEAIINFC
RQPSFVLEAYVNYDCDTTFRDVFEDTGKLLCRYAFPTGGILTSTQVQAFEGLSLILHYIADNIDNEEDHPTPNPT
DPYPVNISDPGDQPFWEEKPKEDDEEAWVEYLRIRKAQKKKILIARSHYSRDDKKGLEYLKYSRLVSDPPHPKAY
ASFFRYTQKLDKTMIGDFLGDPAEFNIQVLKEFSEMFDFTGMVVYTALRTYLETFRLPGESQKIQRILEAFSERY
YDQQSSDMFVSKDAVMILCYSIIMLNTDQHNPQVKKKMTEEDFIRNNRAINGGNDLPREYLSEIFHSISTNAISL
DRPGATTEINPNRWIQLVSRGKVIQPFIFCNFDRRLGRDMFAAIAGPSVATVASIFEQSDEEEILHECLEALFSI
SRIAQYGLEDTLDELLCSFCKFTTLLNPYASTEETLYAFSHDMKPKMAILSVFTIANAFKDSIRGGWRTIMDCll
klkklklIPQSIVDPDSYATPSEAESSSVIQDLKAGQQKRYSSGVMSRFLNFMSVenleeslnlglseFEQNVKI
IQQCRIGSIFTTSSTLPDESLLNLGRCIIFAAAGKVQKFSTSVEEEEIMAFCWELISAISLCNVHRFSSFWPHYH
EYLLEVVQLPLFSTIPFTEKAIVSLLRICLKLAASSQLEGSEEPLFKSINLMWRLEKEILDNCSEFLLQSVSKII
TDYPDQLQTQLGWMSVLHLLSVTGRHPETYDIGVEVLIALMSKLNNITRTNFSYCIDCAFAFVALRNSRVDKNIK
IMDLLAESASLLVEWYRAGYSDPGSYPSNVSGSSVEETTRIYNPISFFQRLGESLKKTSLARREEVRNHAVLSLQ
KSFMLAEDLLFTPNTCINAFYNVIFAMVDDLHEKMLQYSRRENAERETRSMEGSLVRSMKVMTEVYLHFLKPISG
SPNFRTFWLGILRRMDTCMKADLGEYGESRMSEVIPELLRKIVSAMKEKEILTPEGEDDLWDTTYYQIQWIVPSL
TDELFPLKE*

**>Am03g32310**

MGRLRLQSNIKAIEEEPEECEVATSSTKALLACTINSEIGAVLAVMRRNVRWGGRYVSGDDQLEHSLIQSLKTLR
KQIFSWQHQWHSVNPCLYLQPFLDVIRSDETGAPITGVALSSVYKILTLDVLDINTVSVEDAMHLVVDAVTSCRF
ELTDPASEEVVLTKILQVLLACMKSKASVMLSNQHVCTIVNTCFRVVHQAGTKGELLQRIARHTMHELVRCIFSH
LPDVDNTEQSLVKGGSSMKNETSGIDPFSSKPENGGGTSEYAGQLSSGLNSASSGLIGGMMDDNIVRTDNGKDAV
PYDLHLMTEPYGVPCMVEVFHFLCSLLNVVEHTGLGPRANSITFDEDVPLFALGLINSAVELGGPAIQHHPRLLS
LVQDELFRNLMQFGLSMSPLILSMVCSIVLNLYQHLRTELKLQLEAFFSCVVLRLAQSRYGASYQQQEVAMEALV

DFCRQKTFMVEMYANLDCDITCGNTFEDLANLLSKSAFPVNCPLSTMHILALDGLIAVIQGMAERIDKDSVSSRR
TSVNLEEYTPFWMVKCENYGDPDHWVPFVRRRKYIKRRLMIGADHFNRDPKKGLEFLQGTHLLPEKLDPQSVACF
FRYTAGLDKNLVGDFLGNHDDFCVQVLHEFAGTFDFQDMNLDTALRLFLETFRLPGESQKIQRVLEAFSERYYEQ
SPLILANKDAALLLSYSLIMLNTDQHNVQVKKKMTEEDFIRNNRHINDGNDLPRDFLSELYYSICKNEIRTTPEQ
GAGFAEMTPSRWIDLMSKSRKTSPFIVADSRAYLDHDMFAIMSGPTIAAISVVFDHAEHEDVYQTCIDGFLAVAK
ISACHHLEDVLDDLVVSLCKFTTLLNPSSVEEPVLAFGDDAKARMATVTVFTIAVSYGDFIRTGWRNILDCILRL
HKLGLLPARVASDAADDSDMSSDPAHGKPLTNSLSSAHMQSIGTPRRSSGLMGRFSQLLSLDTEEPRSQPTEQQL
AAHQRTLQTIQKCHIDSIFTESKFLLPDSLLQLARALIWAAGRPQKGNTSPEDEDTAVFCLELLIAITLNNRDRI
GLLWQGVYEHIANIVQTTVVACALVEKAVFGLLRICQRLLPYKQILADELLRSLQLVLKLDARVADQYCEQITQE
VSRLVKANAAHIQSPMGWRTIASLLSITARHPDASESGFEALSFIMTDGAHLAPANFVLCVDAARQFAESRVGQS
DRSVHAVDLMAASVTCLVQWAQDAREAMAEAEATKLCQDIGEMWLRLVQGLRKVCLDQREEVRNHALLSLQMCLT
GVDNIRLPLGLWPQCFEVVIFTMLDDLAEISQGNPQTQKEYRNIEGTLVLALKLLATVYLHLLHELSQMSSFCKL
WRNVIGRMEKYMKLKVKRSEKLQELVPELLKSTLLAMKDKGVLVPTNTLGGDNVWEQTWLHVNKMFPSLQSEIFS
SQDYSEPSVSNQSDAGTTLASETSA*

**>Am06g34280**

MSYLNQPSKGNGLVEKTNKPSGGALACLLNSEIGAVLAVMRRNVRWGVQYASDDEQIEHSLIIAFKELRKKIFLW
QNQWHSVDPLVYLQPFLEIIKSDETGAPITGVALSSVYKILNLEVLDSQTPNVDNALHLIVDAVTSCRFEVTDPA
SEEVVLMKILQVLLACMKNKASVSLSNHHLCSIVNACFRIVHQASSKSELLQRISRHTMHELIRCIFSHLPDIEN
ELNQGSRSSPSKEDDAETENHVSEEKQHANGYVIVESDSKKLDEIPNKKDRTSSADSSMMDPYGVHAMGEIFQFL
CSLLNVMDDIDIGPRSNPIAYHEDVPLFALGLINAAVEVGGASFGNHPKLLSLIQEELFYNLMQFGLSMSPLILS
TVCSIVLNLYHHLRSKLKLQLEAFISGVLLRIAESKYGASYQQQEVAMEALVDFCRQPMFVSELYANYDCDISCS
NLFEDLANLLSKSAFPVNNPLSAMNVLALDGLIALVLGMAQRIGHDSSGSVDASEELKEYKPFWIMRCNDYSEPL
HWVPYVHNLKHIKRKLMIGIDRFNKDPKKGLDYLQEVHLLPDKMDPRSVAYFFRYANGLDKNLVGDFLGSHDEFW
IQVLHEFARTFDFRNLNLDTALRVLLETFRLPGESQKIQRVLEAFSESYFEQSRDILVNKDAALLLSYSLIMLNT
DQHNAQVKKKMTEEDFIRNNRKINGGNDLPRDFLSELYHSICSNEIRMVPDQGAGVTVLSRSNWIGLVYKAKHTS
PFIISDSGSHLDYDLFAILSGPTIAAISVVLGNAEQEEVLRSCIDGYLSIAKLAASYNLGEVLDDLVVSLCEFTT
LLHPSFSEKSILYFADDEKAKMATVAVFTIANQYGDHIRSGWRNILDCILSLQKIGLLPARLASDATDDLESSTD
RDQVGTLATQSPASQVPATTPSRKSSGIMGRFSLLLSLDAEEPAPQPSEEQLAARQRTLQTMQSCHVDSIFTESK
FLQAESLSQLVRALVLAAGRSRKGNNSLEDEDMAVFCLELLIAITLNNRDRITLLWKNVYEHIAAIVQSTFMPCT
LVEKAVFGLLRICQRLLPYKEDLTDKLLDSLQLVLKLDARVADAYCEQITQEVMHLVKANAMQIKTHTGWRTIIS
LLSITARHPEASEAGFETLSYIMSEGAYLSPANYVLCLTAARQFAESRVGNIDRSMKSLDLMAGSVVCLVRWFSH
TKEASAEEAAIKVSQDILEMWMRLIQGLRKVCLDQREEVRNHAILLLQRCLTGVEGICIPTDMWLQSFDLVIFTL
LDELLEMAQQQSPKDYRSVEGSLVLALKLLSKAFLRSLKDLSQSASFCQVWLRVLSCLEKCMKMRFRGKRSEKIH
ELVPELLKTTLLVMKDNRILVPCDPIGGDSFWQLTWLHVKNIAPSLQLEVFSSEELEKIQEKPPKAGCSPLPDGS
VLVPPNETTA*

**>Am06g18620**

MDKTEHDENNKPYMKKSKRNDLGVSCMLNTEIGAVLAVIRRVPESNTGFYHHLEENCDPSILQSLKSLRALLFNP
QQEWRMVDPSTYISPFLDVLQSDDIPAIATAVTLQSILKILRLGIFDEKTPGAKDAINSVVTSITGCRLEKTNPI
AEDAVMMRILQVLTGIMRHRASILLTDQSACTVVNTCFQVVQQSSNRGDLLQRNARYTMHELIQIIYTRVPDIEV
SEWENSESDTEDNNLDSGYGIRSTVDIFHFLCSLLNVVEIMDGDGLTSQTPDENVQLFAMYLINSAIELSGDSIG
KHPKLLRMIQDDLFHHLIHYGASSSPLVLSMICSTVLNIYHFLRRSIRLQLEAFFSFVLFRVATPGSSLQLQEVA
VEGIINFCRQPTFIIEAYVNYDCDPTFRNVFEETGKLLCKYAFPTSSLLTSVQVQAFEGLAVFFHNIADSIDKED
DSSPSGPYPVEISGFRPFWEEKSKGDDIETWVDYVRVRKVQKRKILIAGSHFNRDDKKGLEYLKISKLIPEPPDP
KAYAVFFRYTPKLDKTMIGDFLGNPDEFHIQVLKEFTDTFEFTGMVLDTALRCYLETFRLPGESQKIQRILEAFS
EKFYDQQSSDIFVNKDSVMILCYSIIMLNTDQHNPQVKKKMTEDDFIRNNRAINGGSDLPREYLSDLFHSIATNA
IVLNEQSTAPVEMNPNRWIQVVNRAKIMNPFILCDFDRRLGRDMFATIAGPSVATLAAIFEHSDDEEILHECIEA
LFSIARITQYGLEDTLDELLCSFCKFTTLLNPYASAEETLYAFSNDMKPKMATLAVFTIVNTFKNSIKGGWRTIM
DCLLKLKRLKLLPPGVVEQDNSSNEDSDPQTERSESGVLFPSQDPKFNNKRHSAGIMGRFLNFMSMESVEEALNL
GLSEFEQNMNVIKQCRIGTIFSTTSSLPEDSLLNLGRCIIFAAAGKGQKFSTPIEEEETVAFCWELISSIAFANK
HRFPSFWPYYHDYLLEVTQFPLFSPIPFAEKAVVSLMKLCLRLLAADQLESSEELIFKSINLMWRMEKEILDTCC
EFLLSSVSKIITEHPEKLQTQLAWMSLLHLLSVTGRHAETYDQGVETLIQLMSEGNRISRANYSYCVDCSFTFVA
LRNSKVEKNMKIMDLLVDSATLLVEWYRAGYSDPGSSISMASNASGSSLEESGRIYNPISFFQRLGEFLKKTSLA
RREEVRNHAIISVQRSFILAEDLLFTPMNCINSFYNVIFAMVDDLHEKVLQYSRRENAERETRSMESSLKLSMEV
LTNVYLYFLKPISESPSFRTFWLGILRRMDTCMKADLGEYGASKMPEAIPNLLRKIVTNMKETEILTQTGEDDLW
DTTFYQIQWIAPALTDELFPEGLKNNE*

**>Hanim_22191**

MGRLRLQSSIKAIEEEPEDYEATSSNKAALECMINSEIGAVLAVMRRNVRWGGRYVSGDDQLEHSLIQSLKTLRR
QIFSWQNEWQSVNPSLYLQPFLDVIRSDETGAPITGVALSSVYKVLTLDVLDVNTVNVEDAMHMVVDAVTSCRFE
VTDPASEEVVLTKILQVLLACMKSKASVMLSNQHVCTIVNTCFRIVHQAGTKGELLQRIARHTMHELIRCIFSHL
PDVDNTERSLVKGGSSVKNETAGLDSDYSFSSKSENGGGTSEYDGQLSSGFMSGVMDENNPRTDNGKDAVPYDLH
LMTEPYGVPCMVEIFHFLCSLLNVVEHSGMGSRPNSITFDEDVPLFALGLINSAIELGGPAIRHHPRLLSLVQDE
LFRNLMQFGLSMSPLILSMVCSIVLNLYQHLRTELKLQLEAFFSCVILRLAQNRYGASYQQQEVAMEALVDFCRQ
KTFMVEMYANLDCDITCGNVFEDLANLLSKSAFPVNCPLSSMHILALDGLIAVIQGMAERIGNGSVGSEPSPVNL

EEYVPFWMVKCENYSDPDHWVPFVRRRKYIKRRLMIGADHFNRDPKKGLEFLQGTHLLPEKLDPQSVACFFRYTA
GLDKNLVGDFLGNHDEFCVQVLHEFAGTFDFQDMNLDTALRLFLETFRLPGESQKIQRVLEAFSERYYEQSPQIL
ANKDAALLLSYSLIMLNTDQHNVQVKKKMTEEDFIRNNRHINGGNDLPREFLSELYYSICKNEIRTTPEQGAGFA
EMTPSRWIDLMHKSKKTSPYIVSDSRAYLDHDMFAIMSGPTIAAISVVFDHAEYEDVYQTCIDGFLAVAKISACH
HLEDVLDDLVVSLCKFTTLLNPSSVEEPVLAFGDDAKARMATVTVFTIANRYGDFIRTGWRNILDCILRLHKLGL
LPARVASDAADDSEMSSDPGHGKPLTNSLSSAHMQSIGTPRRSSGLMGRFSQLLSLDTEEPRSQPTEQQLAAHQR
TLQTIQKCHIDSIFTESKFLQADSLLQLARALIWAAGRPQKGNTSPEDEDTAVFCLELLIAITLNNRDRIGLLWQ
GVYEHIANIVQSTVVACALVEKAVFGLLRICQRLLPYKENLADELLRSLQLVLKLDARVADQYCEQITQEVSRLV
KANATHIRSPMGWRTIASLLSITARHPDASESGFEALSFIMVDGAHLSPANLVLCVDAARQFAESRVGQTDRSVH
AVDLMAGSISCLVRWAQDAREAMAEAEAAKLCQDIGEMWLRLVQGLRKVCLDQREEVRNHALLSLQKCLTGVDEI
CFPVSFWPQCFEMVIFTMLDDLAEIAQGNPQTQKEYRNIEGTLVLALKLLTKVFLHLLTELSQLSSFCKLWRNVI
GRMEKYMKLKVKRGEKLQELAPELLKNTLLVMKTKGVLVPTSTLGGDNVWEQTWLHVNKIFPSLQSEVFPNQDSE
PQSDATRSPVSSETASAKNE*

**>Hanim_24118**

MTVSLLNQQKEVNGLAEQPNSLPKPSGGALACMVNSEIGAVLAVMRRNVRWGVQYASDDEQIEHSLIISFKELRK
KIFSWQNHWHTIDPVLYLQPFLDVIKSDETGAPITGVALSSVYKILNIQILDSRTVNVENALHLIVDAVTSCRFE
VTDPASEEVVLMKILQVLLACMKNKASVSLNNHHVCSIVNTCFQIVHQASSKSELLQRISRHTMHELVRCIFSLL
PDLDDKKHELHGSSSPNFKGDAAAQSHPLEAKQHVDDYSSAESDNKKNETAHGKDNTYNADSSMMEPFGVPCMVE
IFHFLCSLLNVMENIEIGHRSNPIAYHEDVPLFALGLINLAIELGGASFGNHPKLLALIQEDLFYNLMQFGLSMS
PLILSTVCSIVLNLYHHLRTKLKLQLEAFISSVLLRIAQSKHGASYQQQEVAMEALIDFCRQPMFVTELYANYDC
DISCGNVFEGLANLLSRSAFPVNSPLSAMNTLALDGLIALVQGMAERIGCDSSGAGEAPLELEEYKPFWTSRCND
YGEPLHWVPFVHYMKNIKRKLMIGVDHFNRDPKKGLEFLQGLKLLPDKLDPRSVACFFRYTVGLDKNLIGDFLGS
HDEFCVQVLHEFARTLDFRNMNLDTALRIFLEIFRLPGESQKIQRVLEAFSDSYFEQSPDILANKDAALLLSYSL
ILLNTDQHNTQVKKKMSEEDFIRNNRKINGGNDLPRDFLSELYHSICENEIRMVPDQGGVGAVLTRSHWIGLTHK
AKHTSPYIVSDSGSHLDYDMFAILSGPAIAAISVVFDHAEQEDVLQSCIDGYLAIAKLSASYNLGEVLDDLVVSL
CKFTTLLHPSFNEKSILYFADDIKAKMATVAVFTIANRYSDQIRSGWRNILDCILSLQKLGLLPARLASDATDEL
ESYPNNDQVKSSATHSSAPQVPAPAASRKSSGIMGRFSLLLSLDAEEPAPQPSEEQLAARQRTLQTIQNCHIDSI
FAESKFLQAESLSQLVRALVVAAGRPLKGNNSLEDEDTAVFCLELLIAITLNNRDRIMLLWQNVYEHIANVVQST
SMPCTLVEKAVFGLLRICQRLLPYKENLADELLKSLQLVLKLDARVADAYCEQITQEVMHLVKANAMQIRSHMGW
RTIISLLSITARHPEASEAGFETLSHIMSEGAHLSPTNYVLCVNAARQFAESRVGNVDRSVKSLDLMAGSVVCLV
TWFYQTKEAAGEEAAIKMSQDILEMWMRLVQGLRKVCVDHREEVRNHAIMLLQRCLTGVDGIHIPPDLWLQCFDL
IIFTLLDELPEMAQQHSPKDYRSIEGSMILSLKLMSKTFLQSLLDLSQTTSFCQLWLKVLSCLERYMKIRFRGKR
SEKIHELVPELLKNKLLVMKTSGILVPSDPVGGDSFWQLTWLHVKNISPSLQLEVFPSEELEKLQEKQAKSGCSP
LPDGNVLVPPNETTA*

**>Hanim_17220**

MDKFKRKELGLSCMLNTEVGALLAVIRRAPETNPGFYHHPDDNCDASILQSLKSLRALLFNPQQEWRTIDPSVYL
SPFLDVIQSDDIPASATALALQSILKILRLGICDEKTPGAKDVINSAVTAITGCRLEKTSPVTEDAVMMRILQVL
TAIMRHKASVLLTDQSACTVVNTCFQVVQQSANRGDLLQRSARYTMHELIQIIYSRLPDIEVKDWENSESDTEDN
NLDSGYGIRSAVDIFHFLCSLLNVVDIMEADGLTFHIADENIQVFALYLINSAIELSGDSIGKHPKLLRMIQDDL
FHHLIHYGASSSPLVLSMICSTVLNIYHFLRRSIRPQLEAFYSFVLFRAATPGNPLQLQEVAVEGIINFCRQPTF
IIEAYVNYDCDPTFRNVFEETGKLLCKYAFPIGGLSTSVQVQAFEGLAVIFHNIADNIDKEDDNSSPSGPFPVEI
SGYRPFWEEKSREDDVETWVDYVRVRKVQKKKILIAGSHFNRDDKKGLEYLKISQLISEPPDPKAYAIFFRYTPG
LDKTMIGNYLGDPDAFHIQVLKEFADTFEFTGMNLDTALRTYLETFRLPGESQKIQRILEAFSEKFYDQQSSDMF
ASKDSVMVLCYSIIMLNTDQHNPQVKKKMTEEDFIRNNRSINGGKDLPREYLSEIFQSISTNAISLDQSGTMVEM
NPNRWIQLVNRAKIIQPYILCDFDRRLGRDMFATIAGPSVATVAAVFEHSDEEELLHECIEALFSIARIAQYGLE
DTLDELLCSFCKFTTLLNPYASAEETLYAFSHDMKPKMAMLAVFTIANTFKDSIRAGWRTIIDCLLKLKRLKLLP
QSIFEPENGSNSVSDMPAEKLESGVAVPSQDTKFGNRRHSSGIMARFMNFMSMESVEEALNLGLSEFEQNMTIIQ
QCRIGSIFSTSSSLPLDSLLNLGRCIIFAAAGKGQKFSTPIEEEETVGFCWELIGSIGLANTQRFPIFWPYYHEY
LLEVAQFPLFSPIPFAEKAIVSLMKMCLKLLASDQLESSEELIFKSINLMWRMEKEILDTCFELLLQSVSKIITE
YPGKLQTQLGWMSVLHLLSVTGRHPDTYDQGVETLIKLMSEGTHITRTNYTYCVDCAFAFVALRNSRIDKNVKIM
DLLAESASLLVEWYRAGYSDPGSSTSIASNASGSSAEENVKIYNPISFFQKLGESLKKTSLARREEVRNHAVTSL
KRSFLLAEDILFTPTNCVNSFYNVIFAMVDDLHEKMLQYSRRENAERETRSMEGTLKLSMEVLTDVYLHFLKPIS
ESPNFRTFWLGILRRMDTCMKADLGEYGESKMPDVIPDLLRKIVLSMKEKEILIQTGEDDLWDTTFYQIQWIAPA
LTDELFPEGLKNND*

**>PrihuGNOMa_chr14**

MGRLRLQSSIKAIEEEPEDCETTSSSMAVLACTINSEIGSVLAVMRRNVRWGGRYVSGDDQLEHSLIQSLKTLRK
QIFSWQHQWQSVNPSLYLQPFLDVIRSDETGAPITGVALSAVYKILALDVLGINTINVEDAMHLVVDAVTSCRFE
VTDPSSEEVVLMKILQILLACMKSKASVMLSNQHVCTIVNACFRIVHQAGSKGELLQRISRHTMHELVRCIFSHL
PDVDTTEQSLVRGGSSLKTETAGIDSDHSIVRKSENDTGNSDYDGQLAPGVINPTSSGLINSLTDENTMRSDNEK
DVVPFDLHLMTEPYGLPCVVEIFHFLCSLLNVVEHVGMDSRANNITFDEDVPLFALGLINSAIELGGPAIHRHPR
LLNLLQDELFRNLMQFGLSVSPLILSMVCSIVLNMYQHLRSELKLQLEAFFSCVILRLAQSRFGASYQQQEVAME
ALVDFCRQKTFMVEMYANLDCDITCSNVFEDLANLLSRSAFPVNCPLSSMHILALDGLIAVIQGMAERVGNGSLS
SEPTPVYLEEYTPFWMVKCENFGDPDHWVPFARKKKYIKRRLMIGADHFNRDPKKGLEFLQGTHLLPEKLDPQSV

ACFFRYTAGLDKNLVGDFLGNHDEFCVQVLHEFAGTFDFKDMNLDTALRLFLETFRLPGESQKIQRVLEAFSERY
YEQSPQILANKDAALLLSYSIIMLNTDQHNIQVKKKMSEEDFIRNNRHINGGNDLPREFLSELYYAICKNEIRTT
PEQGTGFAEMTPSRWIDLMHKSNKTSPYIMADSRAYLDHDMFAIMSGPTIAAISVVFDHAEHEDVYQTCIDGFLA
VAKISACHHLEDILDDLVVSLCKFTTLLNPLSVEEPVLAFGDDAKARMATVTVFTIANRYGDFIRTGWRNILDCI
LRLHKLGLLPARVasdaadesemssdPVHGKPLTSSLSAAHIQSIGTPRRSSGLMGRFSQLLSLDTEEPRSQPTE
QQLAAHQRTLQTIQKCHIDSIFTESKFLHADSLLQLARALIWAAGRPQKGNSSPEDEDTAVFCLELLIAITLNNR
DRIGLLWQGVYDHIANIVQSTVVACALVEKAVFGLLRICQRLLPYKENLADELLRSLQLVLKLDARVADQYCEQI
TLEVSRLVKANATHIRSPMGWRTITSLLSITARHPDASESGFEALSFIMSDGSHLSPANFVLCADAARQFAESRV
GQTDRSVHAVELMVGSVSCLVRWAKDAREAMADVEAGKLCQDIGEMWLRLVQGLRKVCLDQREEVRNHALLSLQT
CLTGVDDIFLPPNLWPQCFEMVIFTTLDDLAEIAQGNPQTQKEYRNIEGTlvlalkllaavflHQLQELSQLPSF
CKLWRNVIGRMERYMKLKVRRSEKHQELVPDLLRNILFAMKAKGVHVPTSTLGGDNVWEQTWLHVNKIFPSLQSE
VFPNQDSYLGDCGVSSVSSE*
**>PrihuGNOMb_chr3-29**
MGRLRLQSSIKAIEEEPEDCETTSSSMAVLACTINSEIGSVLAVMRRNVRWGGRYVSGDDQLEHSLIQSLKTLRK
QIFSWQHQWQSVDPSLYLQPFLDVIRSDETGAPITGVALSAVYKilaldvlaidiiNVEDAMHLVVDAVTSCRFE
VTDPSSEEVVLMKILQILLACMKSKASVMLSNQHVCTIVNTCFRIVHQAGSKGELLQRIVRHTMHELVRCIFSHL
PDVDTTEQSLVRGGNSLKAETAGIDPDYsfgsksengsggsEYDGQLGPGVINSTSSGLINSLTDENTMRADNEK
DVPFDLHVMTEPYGLPCMVEIFHFLCSLLNVVEHSGMGFRANNIAFDEDVPLFALGLINSAIELGGPAIHGHPRL
LNLLQDELFRNLLQFGLSASPLILSMVCSIVLNMYQHLRTELKLQLEAFFSCVILRLAQSRFGASYQQQEVAMEA
LVDFCRQKTFMVEMYANLDCDITCSNVFEDLANLLSRSAFPVNCPLSSMHILALDGLIAVIQGMAERVGDGSITS
EPTHAYLEEYTPFWMVNCVNFGDPGHWVPFARKKKYIKRRLMIGADHFNRDPKKGLEFLQGTHLLPEKLDPQSVA
CFFRYTAGLDKNLVGDFLGNHDEFCVQVLHEFAGTFDFKDMNLDTALRLFLETFRLPGESQKIQRVLEAFSERYY
EQSPQILANKDAALLLSYSIIMLNTDQHNIQVKKKMNEEDFIRNNKHINGGNDLPREFLSELYYAICKNEIRTTP
EQGTGFAEMTPSRWIDLMHKSNKTSPYIVADSRAYLDHDMFAIMSGPTIAAISVVFDHAEVEDVYQTCIDGFLAV
AKISACHHLEDILDDLVVSLCKFTTLLNPLSVEEPVLAFGDDAKSRMATVTVFTIANRYGDFVRTGWRNILDCIL
RLHKLGLLPARVasdaadesemssdPVHGKPLTNSLSAAQMQSIGTPRRSSGLMGRFSQLLSLDTEEPRSQPTEQ
QLAAHQRTLQTIQKCHIDSIFTESKFLHADSLLQLARALIWAAGRPQKGNSSPEDEDTAVFCLELLIAITLNNRD
RIGLLWQGVYEHIANIVQSTVMACALVEKAVFGLLRICQRLLPYKENLADELLRSLQLVLKLDARVADQYCEQIT
LEVSRLVKANAANIQSPMGWRTIASLLSITARHPDASESGFEALSFVMSDGAYLSPANFVLCVDAARQFAESRVG
QTDRSVHAVDLMARSVSCLVRWAKDAREAMAEVEAGKLCHDIGEMWLGLVQGLQKVCLDQREEVRSHALLSLQTC
LTGVDGIRLPNLWPQCFETVIFTMLDDLAEIAQGNPQTQKDYRNIEGtlvlalkllatvflhllQELSQLPSFCK
LWRNVIGRMERYTKLKVKRSDKHQELVPELLKNILLAMKAKSVLVPTSTLGGDNIWEQTWLHVNKIFPSLQFEVF
PNQDSNPGDCGISSVSSE*
**>PrihuGNOMc_chr3-28**
MNCLNRQQEVDVLVEKLNGFSNPSGGTLACMLNSEIGAVLAVMRRNVRWGVHYAADEEQIEHSLIISFKELRKKI
FSWQNQWHRVDPVLYLQPFLDVIKSDETGAPITGVALSSVYKILNLDIIDSKTGNIDNALHPIVDAVTSCRFEVT
DPASEEVALMKILQVLLACMKTKASVSLNNNHVCGIVTTCFRIVHQASSKSELLQRISRHTMHELIRCIFLHLPS
IDGNCHRLAPGSQSHTEVTKEAAQTNDKTTEIVHEKIssmssdssmTDPYGIPAMVEIFHFLCSLLNVMEIIDIG
PRSNPIAYDEDVPLFALGLINSAIELGGASFGNHPKLLMLIQEELLYNLMHFGLSMSPLILSTVCSIVLNLYHHL
RTKLRLQLAEFISGVLLRISQSKYGASYQQQEVAMEALVDFCRQPMFVTELYVNYDCDISCSNVFEDLANLLSKS
AFPVNSPLSSMNTLALDGLIALVQGMAERIGHDSSGFGEASPSLELNDYEPFWTAKCNDYVRPHHWVPFVHNMKH
IKRKLLIGVDHFNRDPKKGLEYLQELHLLPSKLDPISVAYFFRYTAGLDKNLIGDFLGSHDEFCIQVLHEFARTF
EFRDMNLDTALRIFLEAFRLPGESQKIQRVLEAFSESYFEQSPHILVNKDAALLLSYSLIMLNTDQHNVQVKKKM
SEDDFIRNNRKINGGDDLPRDFLSDLYHSICENEIRMFPDQGGVAVFTRSHWIGLIHKAKHAAPFIFCDPGSYLD
YDMFAILSGPSVAAISVVFDHAEQEGVLQSCISGFLAIAKLSASYSFGKILDDLVVSLCKFTTLLNPLFNEKSIL
SFGNDIKARMATVGVFTIGNCYGDHIRSGWRNMLDCILSLQKMGLLPARLARDVADDLEAYPDNDHVKPSESPAS
QVPATIPSRKSSGIMGRFSLLLSLDaeepapqpseeqlaaRQVALQTVKNCHIDNIFAESKFLQADSLSQLVRAL
VVAAGRPLKGNSSLEDEVTAVFCLDLLISITLNNRDRIMLLWQNVYDHISNVVQSSVMPGTLVEKAVFGLLRICQ
RLLPYKENLTDEllkslqlvlklDARVADAYCEQITQEVTHLVKANAMQIRSHMGWRTIISLLSITARHPEASEA
GFETLSYVMSEGAHLLPTNYVLCLNAARQFAESRVGQVDRSVRSLDLMAGSVVCLVRWFNHTKEAVNEETAIKIS
QDILEMWMRLVRGLRKLCLDPREEVRNHAILLLQRCLASDGICLTTDLWLECFDPVIFTLLDELLETAQQSSPKD
YRSIEGSIILSLKVLAKAFLQSLRDISQSTSFCQLWLKVLSCMERHMKLRFRGKRSEKIHELVPELLKNVLLVMK
TSGILVPSDPVGGDSFWQLTWLHVKNITPDMQSEVFHREEMEKLQEKQPKGGCSPFPDGNVVVSPNETTA*
**>PrihuGNOMd_sc40239**
MNALNGQQEVDELIEKLNSFSNPSGGTLACILNSEIGAVLAVMRRNVRWGVHYAAVEEQMEHSLIISFKELRKKI
FSWQNHWHRVDPVLYLQPFLDVIKSDETGAPITGVALSSVYKILNLDIIDSQTVNIDNALHLIVDAVTSCRFEVT
DPASEEVALMKILQVLLACMKTKASVSLNNNHVCGILTTCFRIVHQASSKSELLQRISRHTMHELIRCIFSHLPY
IDGKRHRLAPGSLSNTNNLQDAVAAHTQSLQGNKHVNGYPSSESGNEPLAVSSDVPMDTVAVQTNEKTTEVVHGK
ISSLSADSSMTDPYGIPAMVEIFHFLCSLLNVMENIDIGPRSNPIAYDEDVPLFALGLINSAIELGGASFGIHPK
LLTLIQEELLYNLMHFGLSMSPLILSTVCSIVLNLYHHLRTKLRLQLAEFISGVLFRISQSKYGASYQQQEVAME
ALVDFCRQPMFVTELYVNYDCDISCINVFEDLANLLSKSAFPVNSPLSAMNTLALDGLIALVQGMAERIGQDSSG
FGEASPSLELQDYEPFWTAKCNDYVRPHHWVPFVHSRKHIKRKLMIGVDHFNRDPKKGLEYLQELHLLPSKLDPV
SVAYFFRYTTGLDKNLIGDFLGSHDEFCVQVLHQFARTFEFRDMNLDTALRIFLEAFKLPGESQKIQRVLEAFSE

SYYEQSPYILANRDAALLLSYSLIMLNTDQHNAQVKKKMSEDDFIRNNRKINGGDDLPRDFLSDLYHSICENEIR
MFPDQGGVAVFTRSHWIGLLHRAKHATPFIVCDRGSYLDHDMFAILSGPSVAAISVVFDHAEQESVLQSCVEGFL
AIAKLSAAYSFGKVLDDLVVSLCKFTTLLNPFFNEKSILIFGNDVKARMAAVGVFTIANCYGDHIRSGWRNMLDC
ILSLQKMGLLPARLARDATDELESYPDNDHVKPSESPASQVPAIIPSRKSSGIIGRFSLLLSLDaeepapqpsee
qlaaRHVALQTVKNCHIDNIFAESKFLQADSLSQLVCALVAAGRPLKGNSSFEDEETAVFCLDLLIAITLNNRDR
ITLLWQNLHEHIANVVQSSVMPGTLVEKAVFGLLRICQRLLPYKENLTDEllkslqlvlklDARVADAYCEQITQ
EVMHLVKANAMQIRSHTGWRTIISLLSITARHPEASEAGFETLSYVMSEGAHLLPTNYMLCLNAARQFAESRVGQ
VDRSVRSLDLMAGSVVCLVRWFNQTKETVNEETAIKISQDILEMWMRLVQGLRKVCLDPREEVRNHAIMLLQRCL
APDGIRVTTDMWIQCFDPVIFTLLDELLETAQQSSPKDYRSIEGSIILSLKVLAKAFLQSLQDISQSTSFYQLWL
KVLSCMEKHVKMRFRGKRSEKIHELVPELLKNIVLVMKTSGILVPSDPVGGDSFWQLTWLHVKNITPDLQSEVFP
REEMEKLQEKQPKGGCSPLPDGNVVVSPNETIA*

**>PrihuGNL2_chr3-24**

MTKAKRKELGVSCMLNTEIGAVLAVIRRAPDPNTGIYHHPDENFDASILQSLKSLRALLFNPQQEWRTVDPSLYL
SPFLDVIQSDDIPAAATAVALQSILKILRLGVFDEKTPGAKDVINSAVTAITGCRLQRTNPAAEEAVMMRILQVL
TLIMRHRASVLLTDHSVCTVVNTCFQVVQQSTGRGDLLQRNGRYTMHELIQIIYSRLPEIEVNDWENSESDTEDN
NMDTGYGIRSAIDIFHFLCSLLNVVEIMDTDGLTSDENIQHFALVLINSAIELSGDSIGKHPKLLRKIHDDLFHH
LIHYGANSSPLVLSMICSTVLNIYHFLRRSIRLQLEAFFSYVLLIVPNAGSPLQLQEVAIEAVVNFCRQPSFIVE
AYVNYDCDPTFRNVFEKTGKLLCKYAFPTGGALTSIQVQAFEGLVGIFHNIADNIDKEEESIPSGMHLVEISGYR
PFWEEKPKEENTDSWVDYVRVRKLQKRKILIAGSHFNRDDKKGLEYLKISRLISDPPDPKAHALFFRYTPRLDKT
MIGDYLGDPDDFHIQVLKEFTNTFEFSGMILDTALRTYLETFRLPGESQKIQRILEAFSEKFYDQQSSEIFVSKD
AVFVLCYSLIMLNTDQHNAQVKKKMTEEEFIRNNRAINGGSDLPREYLSELFHSIATNAIAIFNQSGTPAKMNPS
RWIQLINRSKIIQPYILCDFDRRIGRDMFATIAGPSVATLAAVFEHTDDEEILHECIEALFSISRIAQYGLEDTL
DELLSTFSKFTALLNPYATAEETLYAFSNDIKPKMATLAIFTIANTFKGSIRGGWRTIIDCllklkrlkllPQSV
LEPENDSDIVNSESQDEKLDSGGVVFPSNVPKVSHFRRSSGIMGRFVNILSMETVEEALNLGLSEFEQNMKIIQH
CRIGSIFTTSSILPDDSILDLGRCIIFTAAGKGQKFSTSIEEEETVAFCWDLINSISLANTHRFSKFWPQYHEFF
LEVAQFPLFSPIPFAEKAIlclmkmclkllASDLEGVEELIFKSINLMWRMEKEILDTCCEFLLQSVSKIITEYP
DKLLTQLGWMSVLNLLSVTGRHHETYDQGVETLIGLMSDEFHITRTNYSYCIDCAFSFVALKHSRVECNVKIMDL
LAESANLLVEWYRTGYSDPGSCISMASNtsissieestRIYNPVSFFQKLGESLRKTSLARREEVRNHAVISLQK
CFVLAEDILFTPMNCINSFYNIVFAMVDDLHEKMQQYSRRENAERETRSMENTLKLSMEVLTELCLHFLMPISES
PNFRTFWLGILRRMDTCMKAELGGYGESKIPDTIPDLLRKIVQRMKEKEILTLEGDDDLWDTTFYQIQWIAPALT
DELFPEGLKKKS*

**>Solyc05g013040**

MGRLRPQSSIKAIEEEPEDCETTSSNKTSIACMINSEVSAVLAVMRRNVRWGGRYVSGDDQLEHSLIQSLKTLRK
QMFSWQHSGQTISPALCLQPFLDVIRSDETGAPITGVALSSVFKILTLDILDLDAVNIEDAMHSVVDAVTSCRFE
VTDPASEEVVLMKILQVLLACMRSKTSVVLSNQHVCTIVNTCFRVVHQAGTKSEVLQRIARHTMHELVRCIFAHL
PEVDNIQHSIVRQHGSSKNEVAGIVNEYSLSSKSENGSGPSEYDSLPPSGGFTSASTGLLSSVTEEGMVMGDNGK
DSVPYDLHLMTEPYGVPCMVEIFHFLCSLLNVVEHVGMGPRANTMAFDEDVPLFALGLINSAIELGGPAICSHPR
LLSLVQDELFRNLMQFGLSMSPLILSMVCSIVLNLYQHLGTELKLQLEAFFSCVVLRLAQSRYGASYQQQEVAME
ALVDFCRQKSFMVEMYANLDCDITCSNIFEELANLLSKSAFPVNSPLSSMHILALDGLIAVIQGMAERIGNGSYS
SEYTPINLEEYSPFWMVKCENYSDPDHWVPFVRRRKYIKRRLMIGADHFNRDPKKGLEFLQGTHLLPEKLDPQSV
ACFFRFTAGLDKNLVGDFLGNHDEFCVQVLHEFAGTFDFQDMNLDTALRLFLETFRLPGESQKIARVLEAFSERY
YEQSPQILANKDAALLLSYSIIMLNTDQHNVQVKKKMTEEDFIRNNRHINGGNDLPRDFLSELYHSICNNEIRTT
PEQGAGFAEMNPSRWIDLMHKSKKTSPYIMCDSKAYLDHDMFAIMSGPTIAAISVVFDHAEHEDVYQTCIDGFLA
VAKISACHHLEDVLDDLVVSLCKFTTLLNPSLVEEPVLAFGDDAKARKATVTVFTIANKCGDFIRTGWRNILDCI
LRLHKLGLLPARVASDAADDSEASSDPGHGKPLPNSLTAAHMQSLGTPRRSSGLMGRFSQLLSIDTEEPRSQPTE
QQLAAHQRTLQTIQKCQIDTIFTESKFLLADSLLQLARALIWAAGRPQKGSSSPEDEDTAVFCLELLIAITLNNR
DRIALLWQGVYEHIAHIVHSTIMPCALIEKAVFGLLRICQRLLPYKENLADDLLRSLQLVTKLDARVTDAYCEQI
TQEVSRLVRANASHIRSQMGWRTITQLLSITARHPEASEAGFDVLGFIMSDGSHLSPANFVLCIDVARNFAESRV
GPADRPIRAVDLMTGSAACLAIWSKDTREAMAEAEALKLSQDIGEMWLRLVQGLRKVCLDQREVRNHALSSLQTC
LTGVDEMYLSHGLWLQCFDIVIFTMLDDLIELTSQKDYRNMEETLILALKLLTKVFLQLLHELSQLTTFCKLWLG
VLNRMEKYMKVKVRGKKSEKLQELVPELLKNTLVVMKSKGVLVQRGALGGDSLWELTWLHVNNIVPSLQAEVFPE
NELGHVESDQTDARGAAYDVADPS*

**>Solyc04g005560**

MGRLRIQSSIKAIEEEPEDCETTSSNKTAIACMVNSEVSAVLAVMRRNVRWGGRYVSGDDQLEHSLIQSLKTLRK
QIFSWQHQWQTINPALYLQPFLDVIRSDETGAPITGVALSSVFKILTLDFLDHNTVNVENAMHSVVDAVTSCRFE
VTDPASEEVVLMKILQVLLTCMRSKASVTLSNQHVCTIVNTCFRVVHQAGSKSELLQQTARHTMHELVRCIFSHL
PEVDDMQQSIVRRGSSTKNKVLDVDNEYSFNKPENGTGTSEYDSQPLSGTFTSSASTGLLNSVIDEGMVMNDNGK
DSVPNDLHLMTEPYGVPCMVEIFHFLCSLLNVVEHIGMGPRSNTTAFDEDVPLFALCLINSVIELGGPAIHSHPR
LLSLVQDELFQNLMIFGLSTSPVILSTVCSIVLNLYQHLRTELKLQLEAFLSCVVLRLAQSRYGASYQQQEVAME
ALVDFCRQKSFMVEMYANLDCDITCTNVFEELANLLSKSAFPVNSPLSAMNILALDGLIAVIQGMAERISNGCRS
QQNLINLEDYAPFWMVECESYSDPDHWVPFVRRRKHIKRRLMIGADHFNRDPKKGLEFLQGTYLLPEKLDPQSVA
CFFRYTAGLDKNLVGDFLGNHDEFCIQVLHEFAGTFDFEDMNLDIALRLFLETFRLPGESQKIARVLEAFSERYY
EQSPQILANKDAALLLSYSLIMLNTDQHNIQVKKKMTEEDFIRNNRNINGGNDLPREYLSELYHSICNNEIRTTP

EQSAGFAEMNPSRWIDLMHKSKKTPPYIVCDSRAYLDHDMFATMSGPTIAAISVVFDHAEHEDVYKACIDGFLAV
AKISACHHLEDVLDDLVVSLCKFTTLLNPSLVEEPVLAFGDDAKARMSTVTVFTIANEYGDFIRTGWRNILDCIL
RLHKLGLLPARVASDAADDSEAPSDTGHGKSLPNSLSSAHLQSIGTPRRSSGLMGRFSQLLSLDTEEPRSQPTEQ
QLAAHQRTLQTIQKCRIDSIFTESKFLLADSLLQLARALIWAAGRPQKGSSSPEDEDTAVFCLELLIAITLNNRD
RIRLLWQGVYEHIASIVHSTVMPCALIEKAIFGLLRICLRLLPYKENLADELLRSLQLVLKLDARVADAYCEQIT
QEVSRLVKANATHIRSQMGWRTITLLLSITARHPEASEAGFNALVFIMSDGAHLSPANYVLCIDAARNFAESRVG
PADRPARAVDLMAGSVASLACWSKDTREALAETEAVKLSQDIGDMWLRLVQGLRKVSLDQREVRNHALLSLQMCL
KGDHEIYLPNSVWLQFFDMVIFTMLDDLIELASQKDYRNMEETLVLALKLLAKVFHQLLHELSQLTTFCKLWLGV
LNRMEKCMKVKVRGKKSEKLQELVLELLKNTLTVMKSKGVLVKRSALGGDSLWELTWLHVNNIVPSLQSEVFPDN
DSENVQPDPVFSETNPNE*

**>Solyc02g067010**

MGCLNQQSEVNTPFSEPNDCTVKPSKGALACMVNSEIGAVLAVMRRNVRWGFHYAAADDQLEHPLIHSFKELRKN
VFSWKHHWNRVDPLLYLQPFLDVIQSDETGAPITGVALSSVYKFLTLEIIDSSIMNVEKALYQIVETVTSCRFEV
TDPASEEVVLMKILQVLLACMKSKASENLSNHHVCNIVNTCFRLVHQASAKSELLQRIARHTMHELVRHIFAHLP
NIISKAHEFDQQSRLCADSEAGEKQHDNGCVSAESTGKSAPAAVPSNASDKRDGTTDEKTQKEEIASNRENPMMD
PYGVPCMVEIFHFLCSLLNVMESIEIGSRSNPIAYDEDVPLFALGLINSAIEVSGASSGNHPELLALIQKDLFHN
LMRFGLSMSPLILSTVCSIVLNLYHHMRSKLKLQLGTFFSGVLLRIAQSKYGTYYQQQEVAIETLVDFCRQPMFM
PEMYANFDCDISCSNVFEDLANLLSKSSFPVNIPLSALNTLALDGLIAMMEGMAERISQDSFVSDQASIDLGEYR
SFWTEICKDYSDPNHWVPYLRKMKVIKRKLLIGVDHFNRDPKKGMDFLQGVHLLPEKRDPKSVACFFRYTTGLDK
NLIGDFLGSHEDFYIEVLHEFAGTFDFRGMNLDIALRIFLETFRLPGESQKIQRVLEAFAERYYEQSQNILADKD
AALLLSYSIIMLNTDQHNAQVKKKMTEEDFIRNNRRINGGNDLPREFLSELYRSICEDEIRITPDRGAGIPMMAP
SHWIGLVHKSRQTSPYIICDPGPYLDYDMFAMLSGPAIASISVVFDNVEQEDVWETCISGFLAIARIAAAYSFDD
VLNDLVVSLCKFTTLLLPSYVDEFTVAFAEDGKARLATLAVFTLANEYGDHIRSGWKNILDCILCLHKLGLLPTR
LFSDAADDLESTGDADPRRPTALFPSPSRFPSSTPSRKSSGLMGVFSQLLYLDEEPAPQPNEQQLAARQQTLQTI
QSCHVDSIFAESKFLQAESLLQLVRAVVLAAGKPRKRNNSLEDEETAVFCLELLIAITINNRDRIMLLWQVVYDH
IASVVHLTTMPSTLIEKAVFGLLRICQRLLPYKENLTDELLKSLQLILKLDARVADAFLEQITREVMHLVKANAM
QIRSHIGWRTIISLLSFTARHPEASETGFDTLVFIMADGAHLLPANYVLCLNVAAQFADSHVGNVDQSVRSLDLM
AGSLISLIRWSHQAKEALGQEAAVKMTQDITEMWLRLIQGLRKFCKDRREEVRDHAILMLQMCLTGVDGIHIPED
LWLQCFDQVIFTLLDELLNLAQPSFVKDYRSTEGAIVLALKLMFKMFLQSLNHLSQSTSFCKLWLGVLSLTERCM
KVKFKGKWSEKIPELISELLKNTLLVMKTSGILGPSNPVGGDSFWKSTWLHVHKICPSLQTEIFPTNEAEQSEKQ
HIQVGCSPLAEGNVIVSSGNSTA*

**>Solyc02g088290**

MGFPNQPSGGNTPISEPKDHTIKPSRGALECMVNSEIGAVLAVMRRNVRWGFRYAAEDDQLEYSLIQSFKELRKK
IFSWRHEWNNVDPLLYLRPFLDVIQSDETGAPITGVALSSVYKFLTLGIIESADMNVDKALHQIVDAVTSCRFEV
TDPASEEVVLMKILQVLLACMKSKASKNLTNHHVCNIVNTCFRLVHQASVKSELLQRIARHTMHELVRCIFVHLP
DIEKITIQGMELAHLWLSIQPTGKKQQEDNGCVDVAVGNDQTDEKTRKRDIACNGENPMMDPHGVPCMVEIFHFL
CSLLNVMESIEIGSRSNPIAYEEDVPLFALGLINSAIELGGAAFANHPKLLALIREELFHNLMRFGLSMSPLILS
TVCSIVLNLYHHLRFKLKLQLEAFFSGVLLKIAQSKHGASYQLQEVAMETLVDFCRQHMFVAEMYANYDCDISCS
NIFEELANLLSKSTFPVNSPLSALNTLALEGLIAMIQGMAERIGQDSLVSDQGSFNLDGFRPFWVEICKDYNDPD
HWVPFVHKMKQIKKKLLVGVDHFNRDPKKGMEYLQAVHLLPDKLDPKSVACFFRFSNGLDKNLVGDFLGSHEEFY
IQVLHEFSRTFDFQDMNLDTALRIFLETFRLPGESQKIHRVLEAFSERYYEQSPDVLANKDAALVLSYSLIMLNT
DQHNTQVKKKMTEEDFIRNNRRINGGNDLPREFLSELYHSICENEIRISSDRGADTPVLPPSHWIGLVHKSRQTS
PFIVCDHGPYLDYDMFAMLSGQTIASISVVLDHVEQEDVWQTCIDGFLAIAKISASYNFDDVLDDLVVSLCKFTT
LLLPSYTDEFIVTFAQDNKARLATLAVFTIANKYGDHIRSGWKNILECILSLHNFGLLPTRLFNDAADDVESTSD
AYKSKPVAASPSTPHVPSLAPSRKSSGLMGRFSQLLYLDAEEPAPQPNEKQLAALLQAESLSQLVRALVMAAGRP
HKGNISLEDEETAVFCLELLIAITINNRDRIMLLWQVVYEHIASIVQSTTMPCSLVEKAVFGVLRICQRLLPYKE
NLTDELLKSLQLILKLDARVADAFLEQITQEVMHLVKANAMQIRSNMGWRTIISLLSITARHPEASEAGFETLSF
IMADGAHLLPANYILCLNAASHFADSRIGSVDQAVRSLDLMAGSLVCLVRWSHKTKDALGEEAAIKMSQDITEMW
LRLVQGLRKFCLDWREEVRGHAILMLQRCLTVVEGIHISTDLWLQCFDQLIFTMLDELLELAPQGSLKDYRSIEG
AIFLSLKLMFKVFLQSLEHLSQLPSFCKLWLGLLDHTERCMKMKFKGRRSEKIPELVPELLKNTLLIMKTSGILM
PSDPVGGDSFWQLTWLHVHKICPSLQSEVFPSSELEQLQKQQIQAGCSPLSEGSVLVSPS*

**>Solyc03g070470**

MTKIKRKELGISCTLNTEVGAVLAVIRRSPEGNLHFYHPPEENYDSQISLSLKSLRSLIFNPQQEWLTIDPMIYL
SPFLDVVQSDDVPSAATAVALSSILKILKLEIFYHKSPGAREAINSAVTAVTGCRLEKTDPVSEDAVMMKILQAL
TAIMNHPSSVLLTDQSVCTVVNTCFQVVQQSANRSDLLQRSARYTMHELIHVVYQRLPEIEVKDWEDSESDTEDG
ILDSGYGILSAVDIFHFLCSLLNVVEVMETDGSTPQTSDENVQLFALVLINSAIELSGDSIGKHQKLLRMIQDDL
FHHLVHYGTSSNPIVASMICSIVLNIYHFLRRSVRLQLEAFFSFVLLKVASLANSLQLQEVAIEGMINFCRQPSF
IVEVYINYDCHPMFKNVFEEIGKSLCRHAFPTGGCLTSLQVQAFEGLAVIIHNIADNVDKEDDLTPSGPYPVEIS
EYRQFWEEKSKEDEEDLENWIDFIRVRMAQKRKILIAGNHFSRDEKKGLEYLKLSLLIPDPPDPKAYAMFFRYTP
GLNKVAIGDFLGDPDDFYLQVLKEFTETFEFMGMVLDTALRTYLETFRLPGESQKIQRILEAFAERFFDQQSSEI
FASKDAVLILCYSVIMLNTDQHNPQVKKKMTEDEFIRNNRGINGGQDLPREYLSEFFHSISVNAITLFGSSGAPV
EMNPSRWIQLINKSRKMKPFIFTNFDRRLGRDMFASIAGPTVSTLATIFEQSDEEEILHECVEALFSIARITKYG
LEDTLDELLCSFCKFTTLLNPYASSEETLYAFSNDMKPRMATLAVFTIANDFKKSIRGAWRTIVDCLLKLRKLKL

LPQSVVEPENASNSSSNPPGIHERCASGVVFPTQDVKFGSKAQNSGIIGRFSHFLSMETVEESLNLGVSEFEQNL
KVIQQCRIGSIFSNSSSLPDEPLLNLGRCLIFAAAGKGQKFSTPIEEEETVGFCWDLIVSIASSNTNRLLVFWPH
YNEYLLDVAQFPLFSPIPFAEKGIIALMKICLKLLSSFHSDKSPEELMFKSINLMWKLEKEILDTCSDFIVQSVT
TILTEYPANLQSQLGWKTVMHLLSVTGRHPETYEQGVEALINLMSDGFHISRLNYPYCIDCAFGFVALKNSPLEK
NMKIMDLMSDTVNLLVQWYKSGYTDPGSSTSINSSASSCSLEESSKALSSSNLTVTYFAKLGEAFRKTSLARREE
IRNHAVMSLQKSFALGEELYFSPANILSCFNLILFAMVDDLHEKMLEYSKRGNAEREARSMEGTLKLSMEVLTEV
YLQFLKPLSESPSFRAFWMGILRRMDTCMKADLGDCGESKLPHTIPVLLKKMVITMKQKEILVPGDDEDLWEMTH
VQIQWIAPSLTEELFSDV*

**>Cusau_000320**

MGRLRLQSNIKAIEEEPEECESTSSSKNDLACMINSEVSAVLAVMRRNMRWGGRFVSGDDQLEHSLIQSLKNLRR
LIFSWHHHQWHTINPALYLQPFLEVIRSDETGAPINGVALSSVYHILTLDVLELNTANIEDAMHLVIDAVTSCRF
DVTDPASEVLMKILQVLLAVMRSKVSVVLSDQHVSTIVNTCYRVFHQAGTKGELLQRIARHTMHELVRCIFSHLP
EVDHTQSSIVKRGSSAKHEVSALDGGYGYSSKLENGTASSEYDSQLPSGVDESMVMSEYGKDTVPYDLHQITEPY
GVPCMVEIFHFLCSLLNVVEHKGISPKSNTVPSDEDVPLFALGLINSAVELGGRAIRNHPRLLSLIQDELFRNLI
QFGLSTSSLILSMVCSIVLNLYQHLRTELKLQLEAFFSCVVLRLSQSRYGASYQQQEVAMEALVDFCRQKAFMVD
MYANLDCDITCGNVFEELANLLSKNAFPVNSPLSAMHILALDGLIAVIQGMAERVSNGSYNSDPHHLNFDEYTPF
WMVKCEDYNDPEHWVPFVRRRKYIKRRLTIGADHFNRDPKKGLEFLQGTHLLPEKLDPQSVACFFRYTAGLDKNL
VGDFLGNHDEFCVQVLHEFAGTFDFQDMNLDTALRLFLETFRLPGESQKIQRVLEAFSERYYEQSPHILANKDAA
LVLSYLIMLNTDQYNVQVKKKMTEEDFIRNNRHINGGNDLPRDYLRELYHSICSNEIRTTLEQGVGFTEMTPSRW
IDLMSKSRKTSPYIQCDSRAYLDHDMFAIMSGPTIAAISVVFDHAEHEDVYQTCIDGFLAVAKISSCHHLEDVLD
DLVVSLCKFTTLLNPSSVEEPVLAFGDDAKARMATVTVFTIANKYGDFIRAGWRNILDCILRLHKLGLLPARVAS
DDDSEMISSSEPGYGGKPPLTSSLSTAHMQSIGTPKRSSGLMGRFSQLLSLDSEEPRSQPTEQQLAAHQRTLQTI
QKCHIDSIFTESKFLLSDSLLQLARALIWAAGRPQKGSSSPEDEDTAVFCLELIAITLNNRDRIGLLWQGVYDHI
ANIVHSTVMPCALVEKAVFGLLRICQRLLPYKDDLGDELLRSLQLVLKLDARVADAYCEQITQEVSRLVKANPGH
IRSQTGWRTIASLLSITARHLEASEVGFDALMFIMSDGAHLSPANYVLCIDAARQFAESRVGQADRSVTAIDLMV
SISCLASWSRDAREAMAETEALKLSQDIGEMWLRLIQGLRKVCLDQREEVRNHALLSLQACLTGVDDIYLPHGLW
LQCFDMVIFTVCDDLVDISQGHSQKDYRNMEGTLVIALKVMTKVFLTLLPKLSQLTTFSKLWRGVLNRMEKYMKV
KVRNKRSEKLQEMVPELLKNCLVMRTSGVLVQRSGLGGDSLWDLTWLHVNNIMPSLKSEVFPDHDPDEQPEVSIQ
GLETDSSTVN*

**>Cusau_005101**

MGCLNQQFEANGICSEPKGGMGKPSRSALACMVNSEIGAVLAVMRRNVRWGIQYASSDDQLEHSLIQSFKELRKK
IFSWQHQWNVVDPLVYLQPFLDVVQSDETGAPITGVALSAIYKFLTLEILDSTTKNVDKALHQIVDAVTSCRFEV
TDSASEEVVLMKILQVLLACMKSKASIYLGDQHVCNIVNTCFRLVRQASAKSELLQRITRHTMHELVRCIFSHLP
DIESSEDVFVQRSRFCATPEVGMMMDVQTPGDGQSDNGSVNVESNHKTSSDEGSVTIPPLNVSPSQLTSKKNEQG
SQKESVIGGALSMMDPYGIPCMVEILHFLCSLLNVMDHMEIGPRSNPIAYDEDVPLFALGLINSAIELGGASFGR
HPKLLALIQQELFRTLMRFGLSMSPLILSTVCSIFLNLYQHLRDKLKLQLETFFSGVLLRIAQNKHGSSYQQQEV
ALEALVDFCRQNVFMTEMYANFDCDISCSNIFEELASLLSKSTFPVNSPLTAMNTIALDGLIAMVGGVCERIGHG
PVSEHVSLDSEEYKPFWTEECQDYNEPIHWVPYVSKMKHIKQKLMIGADHFNRDPKKGLQFVQGMHLLPGKLEPT
SVSCFLRYAPGLDKNLIGDFLGSHDEFCINVLHEFARTFDFAGMNLDIALRIFLETFRLPGESQKIQRVLEAFAE
RYYEQSPDILANKDAALLLSYSIIMLNTDQHNAQVKKKMTEEDFIRNNRRINGGNDLPREFLSDLYLSISESEIR
MTPDAGTVSTAVMMPSYWVGLAHRSNQTAPYIVCDTKPYLDKHLFSILSGPAVAAISVVFDHVEHEGILQTCVDG
FMAVAKISASYNFDQVLDDLVVSLCKFTNMLLPSYSEDFIIRYGDDSKARMATTAVFTIANEYGDHIRSGWKNIL
DCILSLHRFALLPPRILSDATDDQDSSSADVGKNTTTVDLPQVPPLTSNRKSSGLMGRFSQLLSLDIIDDPVDDP
SEDQIDARQRILNMINTCHIDSIFADSKFLQAESLLQLVQALVLAAGRPHKGKKEDHGGIAVFCLDALIAITLNN
RDRIMLLWQSVYKHIASIVQSTAMPCALVEKAVFGLLRICQRLLPYKENLTDELLMSLQLVLKLDARVADAYLPQ
ITEELMHLVKANAPQIRSLVGWRAIVSLLSITAQHPEASDAGFETLSFVMSGDGAHLLPANYILCVNAARQFAEH
RIGSVDRSVTSLNLMAGSVVCLVRWSSKTRESLADEATSKMCQDIWEMWLRLVQGLRKVCLDSREAVRNHAILML
QRCLTEVEGIHIPNDLWLQCFNQIIFTLTDDLVELTQRNNSAKDHSIEGSIVLSLKLMSKVFLQRLHDLHQLESF
RTLWSTVLDCSERCANVKLRGKRSGKINEFVPELLKNILLVMKTSGVLSPTEEGDSLWQLTWSRVEAIAPSLTPS
ELFSPAELVEEEHLQKQPVVASNQETTS*

**>Cusau_001637**

MTVINQQRKELGISCMLNTEVGAVLAVIRRCPEAATAHFFPSAERYETHISQSLKSLRSLIFNPRHEWRSLDPSS
YLLPFLDVVHGDDVPAAATGVALSSVLKILKFDVFDHKSPGAREAINSSVAAVTNCRLEKTDPASVDGIMMKILQ
VLGAIMAHPASVWLTDQAVCAAVSSCFQVVQQSAGCGGDLLHRTARFTMHDLICTVFSRLPEVMSAAAAGENPES
DDWEDAIAAEDSGGYGVRAALDIFRFLCSLLNVAAEGAGDSDESVQLFSLAVINSAVELGGDSMGMHPKMLRMIQ
DDLFHHLIHYGCSTGRPLVLSMICSIVLNLYHFLPGSVRLQLEAFFSFVLFKAASTGNVLTLQEVAIEGLVNFCR
QPRFVVEAYVNYDCDPTFRNVFEDTGRLLCRHAYPSGPALTSLQLQAFEGLVTVIHNVAENIDREEDSTPSGPYP
VEITEYTPFWEEDVDLDSWIDTVRVRKAQKRKILIAGDHFNRDCKKGLDYLKISSLVSETPDPKAYAMFFRYTPK
LDKVLIGEYLGDPDDFHLKVLKEFTDTFEFTGMVLDNALRTFLESFRLPGESQKIQRVLEAFAERFYDQQCSEFF
CGKDAVLVLCYSVIMLNTDQHNPQVKKKMTEDEFIKNNRAINAGQDLPKEYLSDLFHSISTNAIVFESSSSSWVE
MNPNQWVQLINRSKSMSPYIMCGFDRRLGRDLFASIAGPSVATLAAIFEQADVEEVLNDCIEALFSVARICQYGL
DDTLDELVCTFCKFTTLLNPYASAEETLYAFSNDMKPRMATLAVFTIANSFKNYIRGGWRTIVDCLLKLKRLKLL
PKSVLEPEVPSFSDRSRHERSSSTSSFGGKQHNASGGVMGRFTQFLSMENVEESLNLGVSEFENNIKIINQCHIG

NIFNTCSTLPEETLQSLGRSLIFSAAGKGQKFSTAIEEEETVGFCWDLIVAISSANIHRIATFWPHYHEYLLDVA
QFPLFSPIPFAEKAMVSLMKICLKLLSSSSPQADKAPEELIFKSINLMAKLEKEILDMCSEFIVQSVTTILNEYP
ANLQTLLGWKSVLQLLSVSGRHPEMYEQKVEAIINLMSNGSHITRTNYPFCIEAAFGFVALKNSPLEKNIKIMDL
MTNTVTLLVSWNRSGHSDPGSTPNSSSTDENKALSSSNFITTCFIKLGEAFRKTSLSRREEIRNHSVISLQKSFA
LAEELCFTPTTIVNCFKLVLFSMVDDLHEKMLEYSKRENAERETRGMEGTLKLSMEVLTEVYLGYLKQISESPDM
RGFWMGILRRMDTCMKADLGVYGESRLLDTIPELLKKMVLTMKEEEILVQKEDNDLWDMTNIQIQWIVPSLKEEL
FPDD*

**>Eucul_013014**

MGRLKLHSGIKSIEEEPEDCETNCSDKAALGCMINSEIGAVLAVMRRNVRWGGRYMSGDDQMEHSLIQSLKALRR
QLFSWQHQWHSIDPSVYLQPFLDVIRSDETGAPITGVALSSVYKILTLDMLDLQTVNVDEAMHLVVEAVTSCRFE
VTDPASEEVVLMKILQVLLACMKSKASVMLSNQHVCTIVNTCFRIVHQAGTKGELLQRIARHSMHELVRCIFSHL
PDVDTSESSMVKGGNPIKQDVGGADTNYSFRSSQLENGNGNSEYNGQLPSASFSSNASGGPVAAVMDENIVPPGN
GKDALSYDTHLMTEPYGVPCMVEIFHFLCSLLNVVEHMGMGPRSNTIAFDEDVPLFALGLINSAIELGGPSIRNH
PRLLSLVQDELFRNLMQFGLSVSPLILSVVCSIVLNLYNHLRTELKLQLEAFFACVILRLAQGRYGASYQQQEVA
MEALVDFCRQKTFMVEMYANLDCDITCSNVFEDLANLLSKSAFPVNCPLSAMHILALDGLIAVIQGMAERIGNGS
VNSEPALLNLEEYTPFWMVTCNNYSDPDYWVPFVRRRKYIKRRLMIGADHFNRDPKKGLEFLQGTHLLPDKLDPQ
SVACFFRYTAGLDKNLVGDFLGNHDEFCVQVLHEFAGTFDFQDMNLDTALRLFLETFRLPGESQKIQRVLEAFSE
RYYEQSPQILANKDAALLLSYSLIMLNTDQHNVQVKKKMTEEDFIRNNRHINGGSDLPRDFLCELYHSICKNEIR
TTPEQGAGFPEMNPSRWIDLMHKSKRTAPFIMSDSRAHLDRDMFAVMSGPTIAAISVVFDHAETEDVYQTCIDGF
LAIAKISACHHLEDVLDDLVVSLCKFTTLLNPSSVEEPVLAFGDDTKARMSTITVFTIANRYGDFIRAGWRNILD
CILKLHKLGLLPARVASDAADDSDISSESGQGKPLTSSVSSAHMPPMGTPRRSSGLMGRFSQLLSLDTEEPRLQP
TEQQLAAHQRTLQTIQKCHIDSIFTESKFLQADSLLQLGRALIWAAGRPQKGNSSPEDEDTAVFCLELLIAITLN
NRDRIGLLWQGVYEHISNIVQSTVMPCALVEKAVFGLLRICQRLLPYKENLADELLRSLQLVLKLDARVADAYCE
QITQEVSRLVKANATHIRSQMGWRTITSLLSITARHPEASEAGFDALLFIMLDRAHIIPANYVLCVDAARQFAES
RVGQADRSVRALDLMAGSVSCLARWARDSRDAMGEAEAAKTCNDIGEMWLRLVQGLRKVCLDQREEVRNHAISSL
QMCLTGVDEVRFPHGLWLQCFDLVIFTVLDDLLEISHGHSQKDFRNMEGTLLLALKLLSKVFLQLLHGLSQLTTF
CKLWLGVLSRMEKYTKAKVRGKKSEKLQELVPELLKNTLLVMKTRGLLLQRSALGGDSLWELTWLHVNNIAPSLQ
SEVFPDQEDPAQSPLIQREEGEDTTSNTAGSVQDETAAASGG*

**>Eucul_026230**

METQEEEDNPMKTTAKAKGKDLGVSCMLNTEIGAVLAAIRRVPDANHNLHYPFAHHLEEACDPAILQSLKSLRSI
IFNPCQEWRSIDPAIYLSAFLDVIQSDVPAAATAVALSSVLKILKLGIFDEKTPGARDAVNSAVVAITGCRLERT
DAASEDAVMMKILQVLSEIVKHRSSALLTDHAVCTIVNTCFQVVQQSAGRGDLLQRGARSVMHELIQTIYSRLPE
LEVTEWENSESDTEDSSLDSRFGIGSAVDIFHFLCSLLNVVQVVDLEGSAAQTPDEEVQLFALLLINNAVELSGE
CVGKHPKLLRMIQDDLFHHLIHYGTSSSPIVLSTICSTVLSIYHFLRSSIRLQLEAFFSFVLFRAASFGSSIQLQ
EVAVEGIISFCRQPTFVIEAYVNYDSHPIFQNVFEETGKLLCKHAFPAGGPFTGLQVQAFEGLLSIIHTIADNVD
KDDDPSPSGPYPVEITEYKPFWEEKSDEVDEEDGDLENWVESVRLRKAQKRKIMIAGNHFNRDEKKGLEYLKICH
LISDPPDPKAFAIFFRFTAGLDKVTIGDYLGDPDEFHIQVLKEFTETFEFNGMTLDAALRTYLETFRLPGESQKI
QRILEAFSERFYDQQSSEIFVSKDSVFILCYSLIMLNTDQHNPQVKRKMTEEEFIRNNRAINGGNDLPREYLSEL
FHSISTKAIKLFGQSGGGTVEMNPNRWIQLINRAKHMNPFVLCDFDRRLGRDMFAAIAGPSVATLSATFEYAVEE
ETLHECIEGLFSVARISQYGLEDTLDELIASFCKFTTLLNPYASAEETLYAFSNDMKPRMATLAVFTVANGFKDS
IRGGWRNIIDCLLKLKRLKLLPQSVIEPDESSYSPFSDARRSESGVIFPTHDLKSGSRRNSSGLIGRFSLFLSIE
SVEESLTLGVSEFEQNLKIIQQCRIGNIFSRSSSLPLEPLLNLGRSLIFAAGGKGQKFSTPVEEEETVEFCWDLI
QAISTANILRLSSFWPNYHDYILNIAQFPLFSPTPFAEKGILAIIKICLKLLSSFQIDKRLEELIFKSINLMWKL
EKEILDNCCEFITKSVNKILTEYAANLQTALGWKSVLHLLSVTGRHPETYDQGVETLIILMSDGTHISRLNYAYC
IDCAFGFVALKNSPVDKNIKLMDLMSESVNCLVQWYKNGFSDPGSSGSVRSNTSNSSMDDNPNKSFGSSNFTANL
FIKLGESLRKTSLARREEIRNHAIVSLQRSFRLAEELYLTPANCINCFNLVIFAMVDDLHEKMVEYSRRENAERE
TRSMEGTLKLSLELMAGVYLQLLKAVAGSGEFRTFWLGILRRMDTCMKADLGGYGDSSLQEVVPELLGKMIRTMK
EMGILEPREDDDMWEITYIQIQWIAPSLKEELFPDIG*

**>Lonja_GWHPAAZE005599**

MGRLKLQPGIKAIEEEPEDCDTTFSDRDKAALACMINSEIGAVLAVMRRNVRWGGRYMSGDDQLEHSLIQTLKAL
RRQIFSWQHQWHSINPVAYLQPFLDVIRSDETDAAPITGVALSSLYKILTLDMLDLNTVNVEEAMHLVVDGVTSC
RFEVTDPASEEVVLMKILQVLLACMKSKASVMLSNQHVCTIVNTCFRVVHQAGTKGELLQRIARHTMHELVRCIF
SHLPDVDNTELSLVNGGGSNKEVSGQHRDYSFGSKQLENGNGNSGYDGQASSASFAPNAGLVASGMDENIVPPGN
GKDTVPYDVHLMTEPYGIPCMVEIFHFLCSSLNDVEHLAMGPRSNTIAFDEDVPLFALGLINSAIELGGPSIRRH
PRLLSLVQDELFRNLMQFGLSTSPPILSMQQEVAMEALVDLCRQKAFMVEMYANLDCDITCSNVFEDLANLLSKN
AFPVNCPLSAMHILALDGLIAVIQGMAERIGNASVSSEPAPVNLDEYTPFWMVKCNNYSDPDHWVPFVRRRKYIK
RRLMIGADHFNRDPKKGLEFLQGTHLLPDKLDPQSVACFFRYTAGLDKNLVGDFLGNHDEFCVQVLHEFAWTFDF
QDMNLDTALRLFLETFRLPGESQKIQRVLEAFSERYYEQSPQILANKDAALLLSYSIIMLNTDQHNVQVKKKMTE
EDFIRNNRHINGGNDLPREFLTELYHSICKNEIRTTPEQGAGFPEMTPSRWIDLMHKSKKTAPFILSDSRAHLDR
DMFAIMSGPTIAAISVVFDHAELEDVYQTCIDGFLAVAKISACHHLEDVLDDLVVSLCKFTTLLTPLSVEEPVLA
FGDDAKARMATITVFTIANRYGDFIRAGWRNILDCILKLHKLGLLPARVASDAADDLEPSSEPGHGKPLTNSLSS
VHVPSVGTPRRSSGLMGRFSQLLSLDTEEPRLQPTEQQLAAHQRTLQTIQKCHVDSIFTESKFLQADSLLHLARA
LTWAAGRSQKGSSSPEDEDTAVFCLELLIAITLNNRDRIVLLWQGVYEHISNIVQSTVMPCALVEKAVFGLLRIC

QRLLPYKENLADELLRSLHLVLKLDARVADAYCEQITQEVSRLVKANATHIRSQAGWRTIASLLSITARHPEASE
AGFDALLFIMSEGAHLMPANFVVCVDAARQFAESRVGQADRSIRSLDLMGGSVSCLTRWAQEAKEAMGEVESVKM
SQDIGEMWLRLVQGLRKVCLDQREEVRNHSLSSLQMCLTGVDEVNIHTDYGCSVLIWSYSQSSKTCLNMEGTLIL
ALKLLSKVFLQLLTELSQLTTFCKLWLGVLSWMEKYSKVKVRGKRSEKLQEIVPELLKNTLLVMKRKGVLVQRSA
LGGDSLWELTWLHVNNIAPSLQSEVFMDQDCGQSQPNRGEGKESVSAEMEAA*

**>Lonja_GWHPAAZE028449**

MKAMGCLNQHSNINTLALELKNCSNRPSGGALALMVNSEIGAVLAVMRRNVRWGVRYMADDDQLEHSLIQSFKEL
RRKIFSWQHQWHSINPVVYLQPFLDVIQSDETGAPITGVALSSVYKFLTLEIFDLDTVNVDDALHLIVDAVTSCR
FEVTDPASEEVVLMKILQVLLACMKNKASLKLSNQHVCNIVNTCFRVVHQASSKGELLQRIARHTMHELIRCIFL
HLPDIDNTEVSVRDADHTSGSKQQNNGNASSEHDDKPYSVGVSLNVSTDTFEGDGNMIEVGQGKEAAQSGDHLMK
EPYGVPCMVEIFHFLCSLLNIIEENSGIGPRSNPIAYDEDVPLFALGLINSAIELGGASFAHHPKLLALIQDELF
RHLMQFGLSMSPLILSTVCSIVLNLYYHLRTKLKLQLEAFFSCVLLRIAQSKHGASYQQQEVAMETLVDLCRQQT
FMAEMYVNFDCDISCNNVFEELANLLSKSAFPVNSPLSTMHILALDGLIAMIQGMAERTFGELPASQNLEDYQAF
WTVKCDNYDDPSQWVPFAHKMKFIKRKLMIGADHFNRAPMKGLEFLQGMHLLPDKLDSQSVACFLRYTTGLDKNL
VGDFLGNHDEFCIDVLHEFAQTFDFSNMNLDTALRVFLETFKLPGESQKIQRVLEAFAERYYEQSPHILANKDAA
LLLSYSLIMLNTDQHNAQVKKKMTEEDFIRNNRHINGGNDLPRDYLCELYHSICMNEIRMTPETGAGFPVMTSSH
WIGLKHKSKQSAPFIVCDSRENLDYDMFRILSSPTIAALSVVYDHVEQEDVLQTCIDGFLAVAKIAARYNLDDVL
DDLVVSLSRMATVAVFTIANGYGDCVRGGWKNILDCILSLHRLGLVPTRLASDATDELDPAPELERGRAAAAPPV
SHKPTLITSKRSSSLMGRFSQFLYLDADEPAPQPTEEQLLAHQRSLQAVQNCHIDNIFTESKFLQAESLSQLSRA
LILAAGRSYKGNNSVENEDIAVFCLELLIAVTLNNRDRIMLLWQGVYEHIANVVQSAVMPCTLVEKAVFGLLRIC
QRLLPYKENLTDELLKSLQLVLKLDARVADAYCEQITNEVMRLVKANAMQIRSHAGWRTITSLLSITARHPEASE
VGFQTLEFIMSDGAHLLPSIKSLDLMANSLVCLVQWSCKTKEAIGEEASALAKMYQDIGEMWLRLVQGLRKICMD
QREEVRNHSILMLQRCLTGMDGIHIPNELWLQCFDQVIFTLLDEILQITEDNSPKEYRNMEMSLFLCLKLLSKVF
LQSMEDLYQSSSFCRLWVGVLGCMDRYMKVKLRGKRSEKIHELVPELLKNTLLVMKTRGILVENDSVGGESFWQL
TWLHVKNISPSLQLEVFPEHNKAGCSPLPDGNLVVQSSETTT*

**>Lonja_GWHPAAZE011710**

MKAMGSLNQHSDINTFAPELKNCSSRPSGGDLALMVNSEIGAVLAVMRRNVRWGVRYMADDDQLEHSLILSFKEL
RRKIFSWQHQWHLINAVVYLQPFLDVIQSDETGAPITGVALSSVYKFLTLEIFDLDTVNVDDALHLIVDAVTSCR
FEVTDPASEELVLMKILQVLLACMKNKASLKLSNQHVCNIVNTCFRVVHQASSKGELLQRIARHTMHELIRCIFL
HLPDIDNTEVSVRDADHTSGSKQQNNGNASSEHDDKTYSVGVSSNVSTDTSEGDGNIMEVGQGKEAAQSGDHLMK
EPYGVPCMVEIFHFLCSLLNIIEQHSGIGPRSNPIAYDEDVPLFALGLINSAIELGGASFAHHPKLLALIQDELF
RHLMQFGLSMSPLILSTVCSIVLNLYHHLRTKLKLQLEAFFSCVLLRIAQSKHGASYQQQEVAMETLVDLCRQQT
FMAEMYVNFDCDISCNNVFEELANLLSKSAFPVNSPLSAMHILALDGLISMIQGMAERTFSGLPASLSLEDYQPF
WMVKCNNYVDPSQWVPFAHKMKFIKRKLMIGADHFNRAPMKGLEFLQGMHLLPDKLDSQSVACFLRYTTGLDKNL
VGDFLGNHDEFCIEVLHEFTRTFDFSNMNLDTALRVLLESFKLPGESQKIQRVLEAFAERYYEQSPHILANKDAA
LLLSYSLILLNTDQHNAQVKKKMTEEDFIRNNRHINGGNDLPREYLCELYHSICMNEIRMTPETGAGFPVMTPSH
WIGLKHKSKQSAPFIVCDSRENLDYDMFRILSSPTIAALSVVYDHVEQEDVLQTCIDGFLAVAKIAARYNLDDVL
DDLVVPLCKFTTLLLPLSVEESILAFGDDTKARMATVAVFTIANKYGDYIRGGWKNILDCILSLHRLGLVPTHLV
SDATDELDPAPILERGRAAAAPPPVSHKPSLITSQRSSSLMGRFSQFLYLDADEPAPQPTEEQVLAHQRSLQAVQ
NCHIDNIFAESKFLQSDSLSQLAQALILAAGQSYKGNNSVENEDIAVVCLELLIAVTLNNRDRIMLLWQGIYEHI
ANVVQSAVMPCTLVEKAVFGLLRICQRLLPYKENLTDELLKSLQLVLKLDARVADAYCEQITNEVMRLVKANAMQ
IRSHVGWRTVTSLLSITARHPEASEVGFQTLEFIMSDGAHLLPANYVLCVNAARQFAESRVGNIELSIKSLDLMA
NSLVCLVRWACKTKEAVGEEALALAKMYQDISEMWLRLLQGLGKVCMDQREEVRNHSILMLQRCLTGMDGIHIPN
ELWLQCFYQVIFTLLDEILQIAEDNSPKEYRNMETSLFLCLKLLSKVFLRLMEDLYQLSSFCRLWVGVLGCMERY
MKAKLRGKRSEKVHELVPELLKNTLLVMKSRGILVEDDSVGESFWQLTWLHVKNISPSLQLEVFPEHNKAGCSPL
PDGNLVVQ*

**>Lonja_GWHPAAZE011712**

MVWQGRLSLHRSRPSGGNLALTVNSEIGAVLAVMRRNVSVRDADHTSGNTSEGDGNIMEAGQGKEAAQSGDHLMK
EPYGVPCMVEIFHFLCSLLNIIEQHSGIGPRSNPIAYDEDVPLFALGLINSAIELGGASFAHHPKLLALIQDELF
RHLMQFGLSMSSLILSTVCSIVLNLYHHLRTKLKLQLEAFFSCVLLRIAQSKHGASYQQQEVAMETLVDLCRQQT
FMAEIYVNFDCDISCNNVFEELTNLLSKSAFPVNGPLFAMHILALNGQFAMIPGMAERTFSGLPASLSLEDYQPF
WLVKCNNYDEPSQWVPFAHKMKFIKRKLIIGADHFNRAPMKGLEFLQGMHLLPDKLDSQSVACFIRYTTRLDKNL
VGDFLGNHDEFCIEVLHEFARTFDFSNMNLDTALRVLLESFKLPGELQKIQRVLEAFAERYYEQSPHILANKDAA
LLLSYSLILLNTDQHNAQVKKKMTEEDFIRNNWHINRGNDLPREYLCELYHSICMDEIRMTPETGAGFPVMTSSH
WIGLKHKSKQSAPFIVCDSRENLDFDMFRILSSPTIAALSVVYDHVEQEDVLQTCIDGFLAVAKIAARYRLDDVL
DDLVVSLCKFTTLLLPLSVEESILAFGDDTKARMATVAAFTIANKYGDYIRGGWKNILDCILGLHRLGLVPTHLA
SDATDELDPAPELDQFLYLDADEPAPQPMEEQLLAHQQSLQAVQNCHIDNFFTESKFLQAESLSQLARALILAAG
QSYKGNNSVENEDIAVFCLELLIAVTLNNGDRIMLLWQGIYEHIANVVQSAVMPCSLVEKAVFGLLRICQRLLPY
KENLTDELLKSLQLVLKLDARVADAYCEQITNEVMRLVKANAMKIRSHVGWRTITSLLSITARHPEASEVGFQTL
EFIMSDGAHLLPANYVLCVNAARQFAESRVGNIEQSTKSLDLMANSLVCLVRWSCKTKEAVGEEALALAKIYQDI
SEMWCLTGRDGIHILNELWLQCFDQVIFTLLDEILQIAQDNSPKEYRNMETSLFLCLKLLSKVFLRSMEDLYQLS
SFCRLWVGVLDCMERYMKAKLRGKRSEKVHELAPELLKNTLLVMKTRGILVENDSVGGESFWHLTWLHVKNISPS
LQLEVFPEHNIAGCSPLPDGNLVVQSSEMTT*

**>Lonja_GWHPAAZE011900**
MKAMGSLNQHSDINTFALELKNCSSRPSGGDLALMVNSEIGAVLAVMRRNVRWGVRYMADDDQLEHSLIQSFKEL
RRKIFSWQHQWHLINQVVYLQPFLDVIQSDETGEPITVVALSSVYKFLTLEIFDLDTVNVDDTLHLIVDVVTSCR
FEVTDPASEEVVLMKILQVLLACMKNKASLKLSNQHVSVRDADHTSRTKQQNNGNASSEHDDKTYSVGVSSNVST
DTSEGDGNIMEVGQGKEAAQSGDHLMKEPYGVPCMVEIFHFLCSLLNIIEQHSGIGPRSNPIAYDEDVPLFALGL
INSAIELGGASFAHHPKLLALIQDELFCHLMQFGLSMSPLILSTVCTIALNLYHNLRTKLKLQLEAFFSCVLLRI
AQSKHGASYQQQEVAMETLVHLCRQQTFMAEMYAMHILALDGLIAMIQGMAERTFGGLPASLSLEDYQPFWMVKC
NNYDDPSQWVPFAHKMKFIKRKLIIGADHFNRAPMKGLEFLQGMHLLPNKLDSQSVACFLWYTTGLDKNLVGDFL
GNHDEFCIEVLHEFARTFDFSNMNLDTALRVLLESFKFPGESQKIQRVLEAFAERYYEQSPHILANKDAALLLSY
SLLLLNTDQHNAQVKKMTEEDFIRNNRHINGGNDLPREYLCELYHSICMNEIRMTPKTGAGFPVMTSSHWIGLKH
KSKQSAPFIVCDSRENLDYDMFRILSSPTIAALSVVYNHVEQEDVLQTCIDGFLAVAKIAARYNLDDVLDDLVVS
LCLVPTHLASDATDELDPAPELERGQFLYLDADEPAPQPTEEQLLAHERSLQAVQNCHIDKKFTESKFLQAESLS
QLARALILAAGQSYKENNSVENEDIAVFCLELLIAVTLNNRDRILLLWQGIYEHIANVVQSAVMPCTLVEKAMFG
LLHICQRLLPYNENLTDELLKSLQLVLKLDACVADAYCEQITNEVMRLVKANAMKIRSHVGWRTITSLLSITARH
PEASEVGFQTLEFIMSYGAHLLPENYVLCVNAARQFAESRVGNIELSIKSLDLMANSLVCLVRWSCKTKVAIGQE
ALALAKMYQDISEMWLRLLQGLRKVCMDQREEVHNHSILMLQRCLTGMDGIHVPNELWLQCFDRVIFTLLDEILQ
IAEDNSPKEYRNMETSLFLCLKLLSKCFLQSMEDLYQLSSFCRLWVGVLGYMERYMKAKLRGKRSEKIHELLPEL
LKNALLVMKSRGILVENDSVVESF*
**>Lonja_GWHPAAZE01190**
MKAMGYLNQHSDINTFAPELKNCSSRPSGGDLALVVNSEIGAVLAVMRRNVRWGVRYMADDDQLEHSLIQSFKEL
RRKIFSWQHQWHLINPVMYLQPFLDVIQSDETGAPITGVALSSVYKFLTLEIFDLDTVNVDYALHLIVDAVTSCH
FEVTDPASEEVVLMKILQVLLACMKNKASLKLSNQHVSLQDVDHTSGSKQQNNGHASSERDEKTYSVGVSSNVST
DTFEGDENIMEVGQGKEAAQSGDHLMKELYGVPCMVEIFHFLCSLLNIIEQHSGIGPRSNPIAYDEDVPLFALGL
INSAIELGGASFAHHPKLLALIQDELFRHLMQFGLSMSPLILSTVCSIVLNLYHHLRSKLKLQLEAFFSCVLLRI
AQSKHGASYQQQEVAMETLVDLCRQQTFMAEMYVNFDCDISCNNVFEELANLLSKSAFPVNSPLSAMHILALDGL
IAMIQGMAERTFSGLPASLSLEDYQPFWMVKCNNYDDPSQWVPFAHKMKFIKRKLMIGADHFNRAPMKGLEFLQG
MHLLPDKLNSRSVACFLRYTTGLDKNLVGDFLGNHDEFCIEVLHEFAWTFDFSNMNLDTALRVLLESFKLPGESQ
KIQRVLEAFAERYYEQSPHILANKDAALLLSYSLILLNTDQHNAQVKKKMTEEDFIRNNRHINGGNDLPREYLCE
LYHSICMNEIRMTPETGAGFPVITSSHWIGLKHKSKQLAPFIVCDSRENLDYDMFKILSSPTIAALSVVYDHVEQ
EDVLQTCIDGFLAVAKIAARYNLDDVLDDLVVSLCKFTTLLLPLSVEESILAFGDDTKGRMATVAVFTIANKYGD
YIRGGWKNIIDCILSLHRLGLVPTHLASDATNELDPAPELERGRAAAAPPPVSHKPSLITSQRSSSLMGRFSQFL
YLDADEPAPQPTGEQLLAHQRSLQAVQNCHIDNIFTESKFLQAESLSQLARALILAAGQSYKGSNSFENEDIAVF
CLELLIAVTLNNRDRIMLLWRGIYEHIANVVQSAVMPCTLVEKAVFGLLRICQRLLPYKENLTDELLKSLQLVLK
LDARVADAYCEQITNEVMRLVKANAMQIRSHVGWRTITSLLSITARHPEASEVGFQTLEFIMSDGAHLLPANYVL
CVNAARQFAESRVGNIEQSIKSLDLMANSLVCLVRWSGKTKEAVGEEALAHAKMYQDISEMWLRLLQGLRKVCMD
QREEVRNHSILMLQRCINGMDGIHIPNEFWLQCFDQVIFTLLDEILQIAEDNSPKEYQNMEMSLFLCLKLLSKVF
LQSMEDLYHLSSFCRLWVPILGCMERYMKAKLRGKRSENVHELVPELLRNTLLVMKTRGILVENDSVGGESFWHL
TWLHVKNISPSLQLEVFPEHNIAGCSPLPDGNLVVQSSETTT*
**>Lonja_GWHPAAZE025641**
MGNNQDEEDETKNNKLNKIKKSKELGISCMLNTEVGAVLAVIRRKTDFYPQYIHPPDETFDSSILHSLKTLRSLI
FNPQQEWRRIDPSLYLSPFLDVIQSDDVPAAATGVALSSVLKIFRLQVFDEKTPGAKDAIGEAVTAVTGCRLEKT
DPVSEDAVMMRILLVLSAIMRHRASCLLTDHGVCTLVNSCFQVVQQSASRGDLLQRCARFTMHELIQIIYSRLPD
IEVRDWENSESDTEDSGLDSVYGIRSAVDIFHFLCSLLNVVEVFDVEGSSTAQTVDEDVQLFALVLINSAIELSG
DSIGKHPKLLRMVQDDLFHHLIHYGMCSTPLILSMICSTVLNIYHFLRSSVRLQLEAFFSFVLFRTANLASVQPQ
EVAVEGIINFCRQPTFIVEVYVNYDCDPTFRNVFEEIGKLLCKHAFPAVDRLTSLQIQAFEGLVLILHNIADNVD
KEDDSSPSGPYPVEISEYRPFWEEKTKEHEDFKTWVDFVRVRKAQKRKIMIASNHFNRDLKKGLDYLKICDLVSN
PPNPKAYAMFFHYTPGLDKTMIGDYLGDPDEFHIQVLKEFTDTFEFSGMILDTALRTYLETFRLPGESQKIQRIL
EAFSEKFYDQQSSEIFVSKDSVFILCYSLIMLNTDQHNPQVKKKMTEDEFIRNNRAINGGKDLPREYLSELFHSI
STNAITLFGQTGAPMEMNPSRWIQLINRAKIMKPFALCDYERCLGRDMFAAIAGPSVATLSAIFEQSDDEEMLHE
CIEALFSVSRIAQYGLEDTLDELLASFCKFTTLLNPYASAEETLYAFSNDMKPRMATLAVFTIANGFKDSIRGGW
RIIIDCLLKLKKLKLLPQSVVEPEIASASSLDNQMSEQSESGSIFPGQDIKKLGSKHHGASMMDRFSHFLSMESV
EESLNLGVSEFEQNLKVIQQCRIGSIFSNSSTLPEDPLLNLGRSLIFAAAGKGQKFSTPVEEEETVGFCWDLLLA
ISVANIHRFTTFWPHYNDFLLGVAQFPLFSPIPFAEKGIIALMKICLKLLASYQSEKLPEELIFKSINLMWKLEK
EILDTCCEFLTQSVSKILIEYPANLQTRLGWKSVLHLLSVTGRHQETYDQGVETLIMLMSDGIHISLTNYAYCID
CAFGFVALRNSPVDKNIKIMDHMADSVNLLVQWYKNGYSDPGITMSNMSSSSIEDNNMKALNSSNMTMNLFLKLG
ESLRKTSLARREEIRNHAIGSLQRSFALAEELYFMPTNCINCFNLVIFSMVDDLHEKMLEYSRRGNADRETRSME
GTLKLSMELLAEVYLQLLKPISESPDFRTFWLGILRRMDTCMKADLGYYGESTLQEIVPGLLRKMITTMKENEIL
VQREGDDLWEITYIQIQWIAPSIKEELFPEEM*
**>Dauca_LOC108218573**
MGRLKLQSGINAIEEEPEDSEASFPDKERSSLACMINSEIGVVLAVMRRNVRWGSRYTSGDDQLEHTLIQSLKAL
RRKIFSWQHEWYTINPIVYLQPFLDVIRSDETGAPITGVALSSLYKILSLDVLDLNTVNVEDAMHLVIDAVTSCR
FEVTDPASEEVVLMKILQVLMACMKSKASVMLSNQHVCTIVNTCFRVVHQAGTKGELLQRIARHGMHELVRCIFS
HLRDVDNTDTSLVKGGTSTLQEVGGLHSNYSTGSKQSENDNSTSDYNTQTSSGSFASNTSAGMVGVTGETIVTSS

NGKDIDTYDFHLMTEPYGIPCMVEIFQFLCSLLDVVEHMGMGPKANTIAFDEDVPFFALGLINAAIELGGSSIGR
HPRLLGLVQDKLFCYLMQFGLSMSPLILSMVCSIVLNLYQNLRTELKLQLEAFFSCVILRLAQSRYGASYQQQEV
AMEALVDFCRQKAFMVEMYANLDCDITCSNVFEDIANLLCKSAFPVNCPLSAMHILALDGLIAVIQGMAERIGNM
SAGSEPSPVHLEEYTPFWMVKCDNYNDPNHWVPFIRRRKYIKRRLMIGADHFNRDPKKGLEFLQGTHLLPDKLDP
QSVACFFRYTAGLDKNLVGDFLGNHDEFCVQVLHEFAWTFDFQDMNLDTALRLFLETFRLPGESQKIQRVLEAFS
ERYYEQSPLILANKDAALLLSYSIIMLNTDQHNVQVKKKMTEEDFIRNNRHINGGNDLPRDFLIELYHSICKNEI
RTTPEQGAGFPEMTPSRWIDLMNKAKKTAPFIISDSKAHLDRDMFAIMSGPTIAAISVVFDHAELDDVYQTCIDG
FLAVAKISACHHLEDVLDDLVVSLCKFTTLLNPTFAEEPVLAFGDDSKARMATITVFTIANRYGDFIRAGWRNIL
DCILKLHKLGLLPARVASDAADDSEHTAEPGHGKPPTNSLSSAHLQSVGTPRRSSGLMGRFSQLLSLDTEEPRLQ
PTEQQLAAHQRTLQTIQKCHIDNIFTESKFLQSDSLLHLARALIWAAGRPQKGNSSPEDEDTAVFCLELLIAITL
NNRDRIVLLWQGVYEHISNIVQSTVMPCALVEKAVFGLLRICQRLLPYKENLADELLRSLQLVLKLDARVADAYC
EQITQEVNRLVKANATHIRSPMGWRTITSLLSFTARHPEASEAGFDALMFIMSEGAHLVLANYVLCVDAARQFAE
SRVGQVDRSVLALDLMAGSVSCLARWAQEAKDVMGEEATKASHDIGEMWLRLVQGLRKVCLDQREEVRNRALLSL
RMCLTGVDDVNLPYGLWLQCFDMVIFTVLDDLLEIAQGHSQKDYRNMEGTLVHALKLLSKVFLQLLHDLSQLTTF
CKLWLGVLSRMEKYLKVKVRGKKSEKLQELVPELLKNTLIVMKTKGVLVQRSALGGDSLWELTWLHVNNIAPTLQ
SEVFSDQDSGQSWPNLGEVGENCVPSESVSVPHKTMVKN*
**>Dauca_LOC108218683**
MGCLKLQSGINAIKEEPEDSEASFPDKERSALACIINSEISVVLAVMRRNVRWGSRYTSGDDQLEHTLIQSLKAL
RRKIFSWQHEWYTINPIVYLQPFLDVIRSDETGAPITGVALSSIYKILSLNVLDLNTVNVEDAMHLVIDAVTSCR
FEVTDPASEEVVLMKILQVLMACMKSKASVILSNQHVCTIVNTCFRVVHQAGTKGELLQRIARHGMHELVRCIFS
HLRDVDNTDTSLVKGGTSTLQEVGGLHSDYSNGSKQSENDNSASDYNTQTSSGSFASNTSAGMVGATGETIVTSS
NGKDIDSHDFHLMTEPYGIPCMVEIFQFLCSLLDVVEHMGIGPKANTIAFDEDVPLFALGLINAAIELGGSSIGR
HPRLLGLVQDKLFCYLMQFGLSMSPLILSTVCNIVLNLYQNLRTELKLQLEAFFSCVILRLAQSRYGASYQQQEV
AMEALVDFCRQKDFMVEMYLNLDCDITCSNVFEDIANLLCKSAFPVNCPLSAMHILALDGLIAVIQGMAERIGNM
SAGSEPSPVHLEEYTPFWMVKCENYNDPNHWVSFIRRRKYIKRRLMIGADHFNRDPKKGLEFLQGTHLLPDKLDP
QSVACFFRYTAGLDKNLVGDFLGNHDEFCVQVLHEFAWTFDFKDMNLDTALRLFLETFRLPGESQKIQRVLEAFS
ERYYEQSPLILANKDAALLLSYSIIMLNTDQHNVQVKKKMTEEDFIRNNRHINGGNDLPRDFLIELYHSICKNEI
RMTPEQGAGFPEMTPSRWIDLMNKAKETAPFIISDSKAHLNRDMFDLDRDMFAIMSGPTIAAISVVFDHAELDDV
YQTCIDGFLAVAKISACHHLEDVLDDLVVSLCKFTTLLNPTFAEEPVLAFGDDSKARMATITVFTIANRYGDFIH
AGWRNILDCIVKLHKLGLLPARVASDAADDSEHTAEPGHGKPPTNSLSSAHMQSVGTPRRSSGLMGRFSQLLSLD
TEEPRLQPTEQQLAAHQRTLQTIQKCHIDNIFTESKFLQSDSLLHLARALIWAAGRPQKGSSSPEDEDTAVFCLE
LLIAITLNNRDRIVLLWPGVYEHISNIVQSAVMPCALVEKAVFGLLRICQRLLPYKENLADELLRSLQLVLKLDA
RVADAYFEQITQEVNRLVKANATHIRSPMGWRTITSLLSFTARHPEASEAGFDALMFIMSEGAHLVLANYVLCVD
AARQFAESRVGQVDRSVLALDLMAGSVSCLARWAQEAKDAMGEEATKASQDIGEMWLRLVQGLRKVCLDQREEVR
NHALLSLRMCLTGVGDLNLPYGLWLQCFDMVIFTVLDDLLEIAQGHSQKDYRNMEGTLVHALKLLSKVFLQLLHD
LSQLTTFCKLWLGVLSRMEKYLKVKVRGKKSEKLQELVPELLKNTLIVMKTKGVLVQRSALGGDSLWELTWLHVN
NIAPTLQSEVFSDQDSGQ*
**>Dauca_LOC108214229**
MGCPCDQSNSNSIGRGFDGCSSSPAGGAFALTVNSEIGAVLAVMRRNVRWGVNYMAEDDQLEHSLIHSFKELRKK
IFMWQSRWKTVNPVLYLKPFLDVIQSDETGAPITGVALSSVYKILTLEVIDLDSVNVDEAFQLIVDAVTSCRFEV
TDPGSEEVVLVKILQVLLACMKNKASVKLSNQHVCNIVNTCFRVVHQASSKGELLQRIARHTMHELVRCIFTCLS
NFDNLEKAYTQAGSSHVNNEVGTHDENHIFASKLEDNGHANLESDDKTTPDGVGHDSGDASEEKVDVNVVVPGQD
KVIAQAGGDRMSEPYGVPCVVEIFQFLCSLLNVVETTGVSSKSDSIAYDEDVPLFALGLINSAIEVCGSSIQHHP
KLLVLIQDDLFFNLTRFGLSTSPLILSVVCSIVLNLYHHLRIKLKLQLEAFISSVLLRIAKSKYGASYQQQEVAM
ETLVDLCRQPTFMSEMYVNYDCDISCSNVFEDLANLLSKSAFPVNRPLSAMHFLALDGICAMIHGMAERISSDSP
SSQQASISLTGYQSFWRLRCDNYDNPDEWVPFVHKTKFIKRKLMIGAEHFNRDPKKGLEYLQEVHLLPEKLDPES
VACFFRFTLGLDKIHVGDYLGSHDEFCIQVLHEFARTFDFEGMNLDIALRVFLETFKLPGESQKIQRVLEAFAER
YYEQSPHILCDKDAALVLSYSIILLNTDQHNAQVKKKMTEEDFIRNNRRINGGNDLPAEYLSELYHSISENEIRM
TPEQATGSPTMTASHWVGLMHKSKQTAAFISCDSRGQLDRDMFAILSGPTIAALSVVLDNVVQEDALQTCISGFL
CVAKIAASYHFNDLLNDLVVSLCKFTTLMLPLYGEESVIMLGDDTKARIATEAVFTIATRYGDYIRSSWKNIVDC
VLSLHKIGLLPARLASDATDDLELPFDPEKVKPDLTSSPSPLKLVSSTSRKSSSLIGRFSQLLYFDAEEPAPQPS
EEELAARQVCLEMVQKCQINSIFSESKFLQGDSLSQLVQALISAAGRSDKGINFEEDEETAIFCLELLIAVTLNN
RDRIMLLWQGVYEHITNVVQSAVMPCTLVEKAVFGLLRICQRLLPYKENLTDELLKSLQLVLKLDARVADAYCEP
ITQEVMRLVKANAMQIRSHAGWRTITALLSITARHPEASEVGFETLEFIMADSAHLLPANYVLCINAARQFAESR
VGNVDRSIKSLDLMSGSFVCLVRWSSKTNEALGEETASRMKQDIGEMWLRLVQGVSRICLDPREEVRNHAILMLQ
KSLTGVDGIDLVNELWLQCFDQVILTLLDDLLEIAKESSPKDYRNMEGSLVLSLKLLSKVFLHLLQRISQLSTFS
KLWLEVLSRFEGYMKVKFRGKRCEKIHEIVPELLKNTLLVMKTSGILVPDDDNLEGSLWQLTWLHVKNIAPSLQL
EIFPANETKQLQSNNTETGGS*
**>Dauca_LOC108212563**
MGCLSHQPEVNSFGRGFDDCSSRPSGGAFALIVNSEIGAVLAVMRRNVRWGASYMLDEDDQLDHSLINSFKQLRK
NIFVWQKHWRTINPVLYLKPFLDVIQSDETSVPITGVALSSVYKLLTLEVIDLDSVNVDEALHLIVDAVTSCRFE
VTDPASEEVVLMKILQVLLACMKNKASVKLSNRHVCNLVNTCFRVVHQASSKGELLQRIARHTMHELIRCIFTRL
SKFDSMEKAYIQACSSHVNNDVGTPDGNHIASKLEDNGHAKAASDTSKEKMGINVVVAGQDEEIAQAGGDRMTEP

YGVPCVVEIFQFLCSLLNVTENIGVGSGSNLIAYDEDVPLFALGLINSAIELSGASMQNHPKLLALIQDDLFFNL
TRFGLSTSPLILSTVCSIVLNLYHHLRMKLKLQLEAFFSSVLLRIAKSKYGASYQQQEVAMETLVDLCRQPTFMS
EMYVNYDCDISCTNVFENLANLLSKSAFPVNKPLSAMHVLALDGLCAMIQGMAERITSDIPSSRQASVSLIDYQS
FWRLRCDNYDNPDEWVPFVHRTKFIKRKLMIGAEHFNQDPKKGLEYLQEIHLLPDKLDPQSVACFFRFTNGLDKN
HVGDYLGSHDEFCIKVLQEFARTFDFEGMNLDIALRVFLETFKLPGESQKIQRVLEAFAERYYEQSPLILCDKDA
ALVLSYSIILLNTDQHNPQVKKKMTEEDFIRNNRRINGGNDLPREYLSELYHSISENEIRMIPEQANGSPTMTQS
HWVGLMHKSKQTAPFVSCDNRAQLDPDMFAILSGPAIAALSVVFDNVVQQDVLESCVGGFLNVAKIAASYQFNNL
LNDLVVSLYKFTTLMLPLSGEESVIMLGDDTKARMATEAIFTIANRYGDFIRSGWKNILDCLLCLQKLGLLPTRL
ASDATDDLEPPLDTERVKHASSSSPSPLKMASSTSRRSFSLIGRFSQLLNFDVEEPAPQPSEEELVARQICQQMV
QKCQIYSIFSESKFLQADSLSQLVQALISAAGRSNKGNNFPEDEDTAVFCLELLIAVTLNNRDRIMLLWKDVYEH
ITNVVQSAVMPCMLVEKAVFGLLRICQRLLPYKENLTDELLKSLQLVLKLDARVADAYCEHITQEVMRLVKANAM
QIRSHMGWRTITSLLSITARHPEASEVGFETLEFIMADGAHLLPANYVLCINASRQFAESRVGNVDRSLKSLNLM
SESVTCLVSWSSKTNETVGEEVASKLKQDIGEMWLRLVQGVRKVCLDPREKVRNHAISILERSLTGVDGIDILND
LWLLCFDQVIFALLDDLLELAKESSPKDYLKMEGSLVLSLKLLSKVFLQLLPRISQLTTFSNLWLKVLNHFKIYM
KLKFRGKRCEKIHELVPELLKNTLLVMKTCEILVPGDDNEDDSLWKLTWLHVKDIAPSLQLEIFPLNEMKQLQSN
SS*
**>Dauca_LOC108199672**
MSSARRKEIGRACMLSTEVGAVLAAIRRKNESHPHYFSQEDCYDTSILQSLKSLRTLIFSPQQAWGTIDPSIYLS
PFLDVVESDDVPAPATAVALAAITKIFRNNVFDEKTPGAKHAMAGAVTAITSCRLEKTDSACEDAVMMRILYVLT
ELVRHPASVLLTDHAICTIVNTCFQVVQQSANRGDLLQRSAMFTMHELVQIIFARLPDLDVRDGGNSESDDEEIY
DNRYGIRSAVDIFHFLCSLLNVVQVYHVEYETGQGTMPRQSLDEDVQLFALVLINSVVELSGDSIMNHPKLLRMV
QDDLFHHLIHYGTRCTPLILSMISSIVLNIYHFLRRSVRLQLEAFFSFVLLRAASLGNPVQVQEVAIEGIINFCR
QPTFITEAYVNYDTHPMFRNIFEDLGKLLCMHSFPANGPLTSLQVQAFEGLIITLQNIADSVDKEDSSSPLGEYP
IEVSEYRPFWEERPANGSESWVDFVRKRKAQKRKITIASNHYNRDEKKGLDYLKVSHLVSDPPVAKAFAMFFRFT
PGLDKTMIGDYLGDPSDFHLQVLKDFTETFEFTGMALDSALRTYLESFRLPGESQKIHRIIEAFSNQYYDHQTSD
TFQDGEVVLVLSYSLIMLNTDQHNPQVKKKMTEDEFIRNNRAINGGKDLPREYLSELFHSIATKEITVFGQNGMT
LEMNPNIWIQLINRSKKMKSFKVCDTDRDLGRDMFAVIAGPSVATLSAIFEHTEDEGMLHECTEGLFSIARISQH
GLEDTLDELLASLCKFTTLLNPYASADETLYGFSNDMKPRMATLAVFAIANSFKESIRGGWRNIVDCLLKLKRLK
LLPPNVVDPDNVIESSDSVADEDSESPGQRGLVGRFSHFLSMESVEESLHLGLSEFEQNQKVIQQCRIGYIFNNS
SSLSNDAMQNLGRSLIFAAAGKGQKFSTPVEEEETVALCWDLLLTTTLANVNRFATFWPNYHDYLLTVAQFPMFS
PIPFVEKGIIALIKVSSKILATYRPEKQSEELVFTSINLMWKLEKEILDTCFEPLTKQMSKILTEHPGNIQTQLG
WKSLLHLLSITGRHQETYEQAVEALIFLMSDGTHIYRTNYIYCIDCAFGFVALRNSPVETNKKIMDSMAESVKNL
VQWYTSGYSDPGSSTSTSSADDSSRALASSNFTVNLFVRLGEALRKTSLARREGIRNHAIASLQRSFTLAEELYF
SPSNIINCFNLVLFAMVDDLHEKMLEYSRRENAEKETRSMEGTLKLSMELLTNVYLQFLKPISESPGFKTFWLGI
LRRMDTCMKADLGVYGVPDLQQIVPDLLRKIIVAMKEKNLLVPTEGDDLWEITYIQIQWIAPSVKDELFPDV*
**>Cenas_5.1079**
MGRLKLQPGIKAIEEEPEDCETSFSDKDKAALACMINSEIGAVLAVMRRNVRWGGRYMTGDDQLEHSLIQSLKAL
RRQVFSWQHQWHTINPAVYLQPFLDVIRSDETGAPITGVALSSLYKILSLDVLDLNTVNVEDAMHLVVDAVTSCR
FEVTDPASEEVVLMKILQVLMACMKSKASITLSNQHVCTIVNTCFRVVHQAGTKGELLQRIARHAMHELVRCIFS
HLQDVDNTETSLVKGGSSTKQEVGVMHNDYSIGRKQLENGNGSSEYDSQATSGSFASNASSGMVGPTMDENIVTP
GNGKDAVPYDLHLMTEPYGVPCMVEIFHFLCSLLNVVENMGIGPRSNTIAFDEDVPLFALGLINSAIELGGPSIR
RHPRLLSLVQDELFRNLMQFGLSSSPLILSMVCSIVLNLYQHLRTELKLQLEAFFSCVILRLAQSRYGASYQQQE
VAMEALVDFCRQKSFMVEMYANLDCDITCSNVFEELANLLSKSAFPVNCPLSAMHILALDGLIAVIQGMAERIGN
ASSGSEPTPIKLEEYTPFWMVKCDSYDEPDHWVPFVRRRKYIKRRLMIGADHFNRDPKKGLEFLQGTHLLPDKLD
PQSVACFFRYTAGLDKNLVGDFLGNHDEFCVQVLHEFAWTFDFQDMNLDTALRLFLETFRLPGESQKIQRVLEAF
SERYYEQSPLILANKDAALLLSYSLIMLNTDQHNVQVKKKMTEEDFIRNNRHINGGNDLPREFLTELYHSICKNE
IRTTPEQGAGFPEMTPSRWIDLMHKSKKTAPFIVTDSRAHLDRDMFAIMSGPTIAAISVVFDHAEFEEVYQTCID
GFLAVAKISACHHLEDVLDDLVVSLCKFTTLLNPSSVDEPVLAFGDDAKARMATVTVFTIANRYGDFIRSGWRNI
LDCILKLHKLGLLPARVASDAADDSELSSEPGQGKPLPSSLSSGQMQSVGTPRRSSGLMGRFSQLLSLDTEEPRL
QPTEQQLAAHQRTLQTIQKCHIDSIFTESKFLQSDSLLHLARALIWAAGRPQKGNSSPEDEDTAVFCLELLIAIT
LNNRDRIVLLWQGVYEHISNIVQSTVMPCALVEKAVFGLLRICQRLLPYKENLADELLRSLQLVLKLDARVADAY
CEQITQEVSRLVKANATHIRSQMGWRTITSLLSITARHPEASEAGFDALLFIMSEGAYLMPTNYVLCVDAARQFA
ESRVGQVERSVRALDLMSDSVACLARWAQEAKEAMAEEAVKISQDIGEMWLRLVQGLRKVCLDQREEVRNHALLS
LQMCLTRADEVHLPYALWLQCFDLVIFTVLDDLLEIAQGHSQKDYRNMEGTLVHALKLLCKVFLQLLHDLSQLTT
FCKLWLGVLGRMEKYLKVKVRGKRSEKLQELVPELLKNILLVMKTKGVLVQRNALGGDSLWELTWLHVNNITPSL
QSEVFPDQDSGQSQSNQGKVGENHVSNESVMVPTESVG*
**>Cenas_1.4625**
MNKIKRRELGLSCMLNTEVGAVIAAIRRKSDMHPHFSSPEDHYDTPTIQSLKSLRSLIFTPLQEWRTMDPSLYLS
PFLDVIESDDVSAAATSVALSSILKIFRIDIFDEKTPGAKQAIESAVTAVTGCRLEKTDVNSEDAIMMRILLVLI
AIIRHPASALLTDHSVCTIVNTSFQVVQQSAHRGDLLQRGAMYTMHELIQTIFARLPDIEVKECENSESDTEEDT
GLDSRCGIRSAIDIFQFLCSLLNVVQVHHVENEMVPGVTHQTADEDVQLFALVLINSVIELSGDSIGNHPKFLRM
IQDDLFHHLIHYGMRCSPLILSMISSIILNIYHFLRRSVRLQLEAFFSFVLLRAASLGNSVQLQEVAVEGIVNFC
RQPSFIIEAYVNYDTHPIFRNIFEDVGKLLSMHSFPTNGPFTSLQVQAFEGLVVILHNIADNVDKEDDSSPSGPY

PVEVYDYQPFWEERERDPNNLESWINFVRVRKAQKRKILIASNHYNRDDKKGLEYLKISNLISDPPDPKAYAMFF
RYTPRLDKTMIGDYLGDPDEFHLAVLKEFTATFQFSGMTLDAALRTYLEAFRLPGESQKIHRVIESFSEQFYDQQ
SSEIFVSKDSVFVLCYSLIMLNTDQHNPQVKKKMTEDEFIRNNRAINGGKDLPREYLSELFHSIATKEITIFGQN
GATVDMNPSRWIQLINRCKKMDPFIMCDMDRRLGRDMFSVIAGPSVATLSAIFEHSDDEEMLHECTEGLFSIARI
AQYGLEDTLDELLASFCKFTTLLNPYASAEETLYAFSNDTKPRMATLAVFTIANCFKDSIRGGWRNIVDCLLKLK
RLRLLPQSVVEPENYPSSIDGDEKSESGVALPVQDAKLNKRGHGSGLMGRFSHILSVESVEESLNLGMSEFEQNL
KVIQQCQIGNIFSNSSSLPDEPLQNLGRSLIFAAAGKGQKFSTPVEEEETVGFCWDLLLTVSSVNIHRFSSFWPN
YHDYLLTVAQFPFFSPIPFVEKGIVALVKVCLKILASNRYEKLSEEFIFVSINLMWKLEKEILDTCSEFLTQSVN
KILMEYPANVQTQLGWKTLLHLLSITGRHQETYDQAVETLITLMSDGTHISRTNYAYCIDCAFGFVALRNSPVEK
NKKIMDMMAESVNNLVQWFRSGYSDPGSNVSTSSTDDNTKALASSNFTANLFVKLGESLRKTSLARREEIRNHAI
VSLQRSFALAEELYFSPSNIINCINLVIFAMVDDLHEKMLEYSRRENAERETRSMEGTLKLSMELLADVYLQFLT
PISESPGFKTFWLGILRRMDTCMKADLGVYGVSDMQNMVPELLRKIIVAMKAKDLLLPTEGDDLWEITYIQIQWI
APSVKDELFPDVA*

**>PlagrGNOM_chr7**

MGRQKLQSGIKSIEEEPEDCESTFDDRDKAALGCMINSEIGAVLAVMRRNVRWGGRYMSGDDQLEHSLIQSLKAL
RRQIFSWQHQWHTINPAVYLQPFLDVIRSDETGAPITGVALSSLYKILTLNVLDLNTVNVEDAMHLVVDAVTSCR
FEVTDPASEEVVLMKILQVLLACMKSKASVMLSNQHVCTIVNTCFRVVHQAGTKGELLQRIARHTMHELVRCIFS
HLPDVDNTDRSLVKGSRQEVGGLQADYTFGRVPLENGNGSSEHDGQASSSFASITSSGLVPAMDENIVVPGNGKD
TVSYDLHLMTEPYGVPCMVEIFHFLCSLLNVVEHMTMGPRANTIAFDEDVPLFSLGLINSAIELGGPSICRHPRL
LGLVQDELFRNLMQFGLSASPLILSMICSIVLNLYHHLRTELKLQLEAFFSCVILRLAQSRYGASYQQQEVAMEA
LVDFCRQKTFMVEMYANLDCDITCTNVFEDLANLLSKSAFPVNSPLSAMHILALEGLIAVIQGMAERISNASVNS
ELGPVNLEGYTPFWMVKCDNYSDPDHWVPFVRRRKYIKRRLMIGADHFNRDPKKGLEFLQGTHLLPDKLDPQSVA
CFFRYTAGLDKNLVGDFLGNHDDFCVQVLHEFAWTFDFQDMNLDTALRLFLETFRLPGESQKIQRVLEAFSERYY
EQSAQILANKDAALLLSYSLIMLNTDQHNVQVKKKMTEEDFIRNNRHINGGNDLPREFLSELYHSICKNEIRTIP
DQGAGFPEMTPSRWIDLMHKSKKSAPFIVSDSSAHLDRDMFAVMSGPTIAAISVVFDHAELEDVFQTCIDGFLAV
AKISACHHLEDVLDDLVVSLCKFTTLLNLSSVEEPVLAFGDDAKARMATITVFTIANRYGDFIRAGWRNILDCIL
KLHKLGLLPARVASDAADDSELSPEPGQGKPLTNSLSSAQIPSVGTPRRSSGLMGRFSQLLSLDTEEPRLQPTKQ
QLDAHQRTLQTIQKCHIDSIFTESKFLQADSLLHLARALIWAAGRPQKGNSSPEDEDTAVFCLELLIAITLNNRD
RIVILWQGVYEHISNIVQSTVMPCALVEKAVFGLLRICQRLLPYKENLADELLRSLQLVLKLDARVADAYCEQIT
QEVSRLVKANASHIRSQMGWRTITSLLSITARHPEASEAGFDALLFIMSDGAHLLPANYVLCVDAARQFAESRVG
QADRSLRSLDLMSGSVGCLRQWAQEAKEAMGEAEAAKTCQDIGEMWLRLVQGLRKVCLDQREEVRNHALLALQMC
LTGVDEVNIQHALWLQCFDLVIFTVLDDLLEISQGHSQKDFRNMEGTLILALKLLSKVFLQLLHELSQLTTFCKL
WLGVLSRMEKCIKAKVRSKRSEKLQELVPELLKNTLLVMKAKGVLVERSALGKECPRGDSLWELTWLHVNNIDSS
LQSEVFPDRDSGESLPGQVVASDETNPIPNETASPDAARTGG*

**>PlagrGNL1_chr1**

MGYPNQQSEEVEIAQQANDDSYRPPRATSAFMVNSEIGAVLAVMRRNVRWGVRYMANEDQLEHSLIRSFKELRRE
VFSWQHKWNAINPAVYLQPFLDVIQSDETGAPITGVALSSVYKFLTLEVLDVDTVNVADALHLIVDAVTSCRFEV
TDPASEEVVLMKILQVLLACMKNKASVKLSNQHVCNIVNTCFRVVHQASSKGELLQRIARHTMHELIRCIFSHLP
DIDSTEQAMTQGGQSRVNKEVGEPDEDQISGGQQQDNGYAISEYEEVPSGEIASIVSTGTLAVKMDETISEVGQS
RQHLMTEPYGVPCMVEIFHFLCSLLNAIERNGGDPGSNSIAYDEDVPLFALGLINSAIELGGPYLGHHPELLALI
QDELFRSLLQFGLSMSPLILSTVCSIILNLHHHLRIKLKLQLEAFFSRVLLSIAQSKHGASYQQQEVAMETLVDL
CRQQAFVDELYANFDCDISCSNVFEDLANLLSKSAFPVNSPLAAINILALDGLLAMIKGMAERIGSDLPISDNTS
LDLEDYESFWTLKCENYDNPNCWVPFIHKMKFIKRKLIVGADHFNRDPKKGLEFLQGMHMLPDKLDPSSVACFFR
YTTGLDKNLIGDYLGNHDEFCVDVLQEFARTFDFQGMNLDTALRVFLETFRLPGESQKIQRVLEAFAERYYEQSP
HILADKDAALLLSYSLILLNTDKHNAQVKKKMTEEDFIRNNRRINGGNDLPREYLSELYHSICENEIRMTPDQGV
GPLITHSNWVGLIHKSKEAAPFIVCDSGDHLSYDMFSILSGPTIAALSVVLDHVEQEDVLHTCIDGFLDVAKIAA
CYHLDDVLDGLVVSLCKFTTLLLPLSAEESVLAFGDDSKARMATTAVFTIANKYGDYIRSGWKNILECVLSLHKL
GLLPARLASDAADDLEPPSDLDQGKPALGSPQASSTPPLVTPKKSSSLMGRFSEFLYYDTEKPAPQPTEEQLQAQ
QRSLETVKDCHIDTIFTESKFLQSESLSQLVRAIIIAAGRSYKVNNPLEDEGTAVFCVELLITVTLNNRDRIMLL
WQGVYQHIANVVQSTVMPCTLVEKAVFGLIRICQRLLPYKENLTDELLKSLQLVLKLDARVADAYCENITQEVMR
LVKANARQIRSHLGWRTITSLLSITARHPEASEVGFETLEFIMSDGAHLLPANYVLCVNAARQFAESRVGDVARS
VKSLDLMAGSSVCLVRWSSSAKEAVGEEAAMKLYQDIGEMWLRLVQGLRKVCLDQREEIRNHAILMLQRCLTGLE
GIYLPNELWLQCFDLVVFTLVSDLLEIVQEKSPKEYRNMEGTLVLSLKLLSKVFLHSLQSISQMTSFCKLWLGVL
SCMERYMKVKFKGKRSEKIHELIPELLKNTLLVMKTSGILVPTDAVGGDSFWQLTWLNVKNIAPSLQLEVFPGNE
LEQLQTTTGGKTGVSPVADGKS*

**>PlagrGNL2_chr9**

MKGMTKNKRKELGLSCMLNTEVSSVLAVIRRKESSSQFIHPQDENFDSNVLHSLKSLRSLIFNPQQEWRMMDPSI
YLSPFLDVIQSDDVPASATGVALSSVLKIFRLGIFDEKTPGARDAMASAVSAVTGCRLEKTDPVSESAVMMRILQ
VLNAIMRHNASVLLTDHAVCTIVNTCFQVVQQSASRGDLLQRNARYTMHELIQTIFSRLPDIEVRDWEDSESDSE
DTVEDSGYGIRCAVDIFHFLCSLLNVVEVVEMEGSTSLTADEDVQLFALVLINSAIELSGDSIGKHPKLLRMVQD
DLFHHLIHYGTCSSPLVLSMICSTVLNIYHFLRRSVRLQLEAFSSFVLFKAASSGNSIQLQELAVEGIINFCRQP
TFIIEVYVNYDCDPIFRNVFEEIGKLLCKHAFPGTGPFTSLQLQAFEGLTVLVHNIADNVDKESDSSPCSPYPVE
VSQYRPFWEEKLREKLDLDTWVDHVRVRKAQKRKILIAANHFNRDEKKGLEYLKISQLISDPPDPKAYAMFFRYT

PRLDKAKIGDFLGDPDEFNLRVLKELTDTFEFSKMILDTALRTYLEAFRLPGESQKISRILEAFSEKFYDQQSSE
IFVSKDSVFVLCYSLIMLNTDQHNPQVKKKMTEEEFIRNNRAINGGNDLPRDYLSELFHSISTHAIAVFTESKAS
NEMNPSKWIQLINKAKSMNPFLVCDFDRRLGRDMFACIAGPSVSAISAVFDHADEEEIVHECIEGLFSVARIAQY
GLEDTLDELLASFCKFTTLLNPYASAEETLFAFSHDMKPRMAILAVFTVANTFKDSIHGGWRCIVDCLLKLKRLK
LLPQSLIEPGSSTSDGQTHRRSSSGLILPASGPSKRLNTSGLMGRFSHfLSMESVEESLSLGVSEFEQNMKVIQQ
CQIGRIFTDSSTLPHESLLNLGRSLIFAAGGKGQKFSTPVEEEETVGFCWDLILAITSANLLRFTSFWPTYHDYL
LTVAQFPLFSPVPFAEKGIIALLKFCLRLLSSYRADKLPEELLFKSINMMWKFEKEVLDTCCEFITQSVSRILNE
YPANLQTQLGWKSVLHLLSVTGRHSDTYDQGVETLIMLMSDGTHISRTNYAYCIDCAFGFVALRNSPVDKNVKLM
DLMAESVNLLVQWFRSGYSDPGsNASMNSSSSMDDISSKALASSNFTTNLFLKLGEALRKTSLARREEIRNQAIL
SLQKSFALGEELYFTPTNCINCFNLVIFAMVDDLHEKMLEYSRRDNAERETRSMEGTLKLSMELMTEVYLQFLKP
ISESPGFRTFWLGIMRRMDTCMKADLGEYGESKLQELVPELLKKMIKTMQESEILCKKEGDDLWEITYIQIQWIA
PALKDELFPDEV*
**>Cynca_LOC112507087**
MGRLKLQPGIKAIEEEPEDYENTASERDKAALGCMINSEIGAVLAVMRRNVRWGGRYMSGDDQLEHSLIQSLKAL
RRQIFSWQHPWHTVNPAVYLQPFLDVIRSDETGAPIRGVALSSLYKILSLDVLDLNTVNVEDAMHLVVDAVTSCR
FEVTDPASEEVVLMKILQVLLACMKSKASVMLSNQHVCTIVNTCFHVVHQAGTKGELLQRMARHTMHELVRCIFS
HLPDVDTTEHTFIKGNNTAKQEVSGLPGDYPFGGKQLENGNGGSELDSRAPSANFVSNVTGGMVPAMDENIVVPG
NGKDTVSYDLNLMTEPYGVPCMVEIFHFLCSLLNVVEHMAIGPRANTIAFDEDVPLFALGLINSAIELGGTAIRR
HPRLLSLVQDELFRNLMQFGLSTSPLILSMICSIVLNLYHHLRTELKLQLEAFFSCVILRLAQSRYGASYQQQEV
AMEALVDFCRLKSFMVEMYANLDCDITCSNVFEDLASLLSKSAFPVNCPLSAMHILALDGLIAVIQGMAERISNA
SVSSESVNLEEYTPFWMVKCDNYSDPDHWVPFVRRRKYIKKRLMIGADHFNRDPKKGLEFLQGTHLLPDKLDPQS
VACFFRYTAGLDKNLVGDFLGNHDEFCVQVLHEFAGTFDFQDMNMDTALRLFLETFRLPGESQKIQRVLEAFSER
YYEQSPQILANKDAALLLSYSLIMLNTDQHNVQVKKKMTEEDFIRNNRHINGGNDLPRVFLSELYDSICKNEIRT
IPDQSAAFPEMNPSRWIDLMHKSKKTAPFIVSDSRSHLDRDMFAVMSGPTIAAISVVFDCAELEDVFQTCIDGFL
AVAKISACHHLEDVLDDLVVSLCKFTTLLNSSSPEEPVLAFGDDPKARVATITVFTIANRYGDFIRAGWRNILDC
ILKLHKLGLLPARVASDAADDSELSSEPGQGKPLTNSLSSAHMPSVGTPRRSSGLMGRFSQLLSLDTEEPRLQPT
EQQLAAHQRTLQTIQKCHIDSIFTESKFLQADSLLNLARALIWAAGRPQKGNSSPEDEDTAVFCLELLIAITLNN
RDRIVILWQGVYEHIANIVQTTLMPCALVEKAVFGLLRICQRLLPYKENLADDLLRSLQLVLKLDARVADAYCEQ
ITQEVSRLVKANASHIRSQMGWRTITSLLSITARHLEASEVGFDALLFIMSDGINLMPSNFTTCVDAARQFAESR
VAHADRSIRALDLMSGSIGCLTQWAQEAQEAMGEAEAAKMCQDIGEMWLRLVQGLRKVCLDQREDVRNHALLALQ
MCLTGVDEVHIQHGLWLQCFDMVIFTVLDDLLEISQGHSQKDFRNMEGTLVLALKLLSKVFLQLLPELSQVTTFC
KLWLGVLSRMEKYLKVKVRGKKNEKLQELVPELLKNTLVVMKNKGVLVQRSALGGDSLWELTWLHVNNIAPSIQS
DVFPDQDSEQKQKPVSAENETEIR*
**>Cynca_LOC112512756**
MGCPHQQSEVVAVSRDTKDCSCTPPRASSAFMVNSEIGAVLAVMRRNVRWGVHYMADEDQLEHPLIKSFKEIRKK
VFSWQRQWHTINPIMYLQPFLDVIQSDETSAPITGVALSSIYKFLSLDVLDLETVNVADALHLIVDAVTSCRFEV
TDPASEEVVLMKILQVLLACMKNKASVKLSNQHVCNIVNTCFQIVHQASSKGELLQRTARHTMHELIRCVFMHLP
DVGDNKEQEFTLEGRSDDGGEVDIPDKDHSSTSKPQDNGYTGSKIESVGGALDLSTEMPEVKMDETTGDIGETKN
HLVTDPYGVPSMVEIFNFLCSLLNAVEHVGVGPRSNSLAYDEDVPLFALGLINSAIELGGAALSHHPEILALVQN
DLFHSLIQFGLSPSPLILSTVCSIVLNLYHLMRVKLKLQLEAFFSSVLMKIALNKHGASYQQQEVAMETLVDLCR
QPTFIHEMYANYDCDISCSNVFEDLANLLSKSAFPVSSPLSAIHVLALEGLIAMINGMAEKIGDEVPFLEPDAST
IEDHESFWKVKCENYENSDFWVPFIRKMKHIKKKLMIGADHFNQDPKKGLQFLQGMYLLPESLDPVSVACFLRYT
IGLDKNLIGDYLGNHDQFCVDVLQEFARTFDFQDMNLDIALRVFLETFRLPGESQKIQRVVEAFAERYYEQSPHI
LANKDAALLLSYSLIMLNTDQHNAQVKKKMTEEDFIRNNRRINGGNDLPREYLSELYHSICENEIRMTPELGVGF
PVMTHDNWVGLIHKSRQTAPFIICSTGERINNEMFAILSGPTVAALSVVLDLVEQEDVLQTCIDGFLDVAKIAGC
YHLDDVLDGLLVSLSKFTTLLLPMSIEESVLAFGDDTKARKATVAVFTIANTYGDYLRSGWRNILDCILSLHKLG
LLPARMASDAADDLESVSEPDLGKSPLVSPAGYTKPLLAPARKSSGLMGRFSEFLYYDTEKPAPQPSEEQVEARN
RATETVKDCHIDSVFTESKFLQSESLSNLAQALILAAGRSQKGNNSSIEDEDAAIFCLELLITVALNNRDRIMLL
WQGVYEYIANIVQSAVMPSTLVEKAVFGLIRICRRLLPYKEDLTDELLKSLQLILKLDARVADAYCEHITQEVMR
LVKANVGQIKSHMGWRTITSLLSITARHPEASEPGFETLEFIMLDGAHLLPANFVLCVNAARQFAESRVGDVARS
QKSLDLMAGSAVCLVRWSRETREAMGEEAEAAIRLYQDIGEMWLRLVQALRKVCIDPREEIRNHAILMLQRCLTG
LDGINLHDEMWVQCFDSVIFPLINDLLEIAREKSAKEYRNMETTLVLSLKLMSKSFLHSLPSISRLISFQKTWLG
VLTCMDQYEKVKFRGKRSEKIHELVPELLKNTLLVMKSTGILSPTNTAGSDGLWQETWLHLENVAPSLRSEVFPD
NEPETMGASPVPGVNLLVP*
**>Cynca_LOC112516397**
MAKQDDDQEEEVEEEEEEEDDDDYDYNEDDDDDDEKATRIKKSKELGLSCMLNTEVGAVLAVIRRRSDPTSQFMYQ
DDVMDSSLLHSLKSMRSLIFNPRQSWRLIDPSMYLSPFLDVISSDDVPAGATAVALFSVLKIMKLGMFDRKTPGA
KEAISNVVIAVTGCRLEKTHPITEAGIMMRILQVLAAVMRHPASVLLDDHSVCTIVNTCFQVVQQSSNQSDMLQR
IARHTMHDLIQIIFSRLPEIAVGDWENSESDTEDTTEETGYGIRCAVDIFHFLCSLLNVVGMVENEGSLALTADE
DVQVFALVLINSALELSGDSIGKHKKLLRMVQDDIFHHLIHYGTSSSPLVFSMICSTVLNLYYFHRKSIRLQLEA
FFLFVLFKISNQSYLIQLQELAVEGIINLCRQPTFIIEAYVNYDCDPICNNIFEEIGKSLCKHAFLSGNGAPTSV
QLQAFDGLMVLIHNIADNVDKQTNSRPTGPYPVEISEYIPFWEEKPKPKEDIDLETWIDSVRLRKVQKRKIMIAG
NHFNRDDKKGIDYLKISNLISDPPDPKGHAMFFRYTPGLDKTKIGEFLGDPGEFSLKVLKEFTDTFELSGLVLDT

ALRTYLETFRLPGESQKIQRILEAFSERFYDQQSSEIFVSKDAVFILCYSVIMLNTDQHNPQVKKKMTEEEFIKN
NRAINGGNDLPREYLSDLFQSIATKAITIFGQTGSPIEMNPSRWIQLINKSKLVQPYITCNFDRRIGRDMFATIA
GPTVATLSTIFEHTEDEEIIHECIEGLFAVARICQYGLEDTLDELISSFCKFTTLLNPYASVEETLFSFGHDLKP
RMATLAVFTVANNFKNSIRGGWRSIIDCLLKLKRLKLLPQPIVDVGTSSANGRPHRRTESESISNDQAGRINTSN
LMAAISHCLSMENVEESLNLGISEFEQNLKVIQQCKIGSIFSKSSNLPIEPLLNLGRSLIFAAAGKGQKFSTPLE
EEETVQFCWDLIQAMASCNVHRLASFWPAYNDYLLTVIQFPLFSPNPFAEKAITALMRICIKLLASKSNEKQSEE
LLFKSINLMWKLDREILDTCNEFIIQSITKILNKYSSNLLSQLGWKTVLHLLSITGRHPETYDRGVEALINLMAD
GTGISIANYPYCIDCAFGFVALRNSPVDKNMRLMDLMADSVKLIVDRLRNGYSDPGSNPSINSSASLDENMKYLA
STTNMSMNLFVKLGEALRKTSLARREEVRNYAVSSLQKSFMNAQELYFTPTNIINCFNLVIFAMVDDLHEKMLEY
STRENAERETRSMEGTLKLAMEVLVDVYLQYLKLLMESSGFKTFWLGILRRMDTCMKADLGTYGESKIKEQVPEL
LKKMITTMREKEVLVLQEGNDLWEITYIQIQWIAPELKDELFPDAA*

**>Lacsa_LOC111878063**

MGRLKLQPGIKSIEEEPEEFESTISEKDKAALGCMINTEIGAVLAVMRRNVRWGGRYMSGDDQLEHSLIQSLKSL
RRQIFSYQHAWHTVNPTVYLQPFLDVIRSDETGAPITGVALSSLYKILTLDVLDLNTVNVEDAMHLVVDAVTSCR
FEVTDPASEEVVLMKILQVLLACMKSKASVMLSNQHVCTIVNTCFRVVHQAGSKGELLQRMARYTMHELVRCIFS
HLPDVGNSENAYFNGNNIDKQEVSGISGDYTFGAKQMENGNSNSNSELDSQSSVPGMDENIVVPGNGKDTVPYDL
NLMTEPYGVPCMVEIFHFLCSLLNVVEHMAIGPRANTISFDEDVPLFALSLINSAIELGGTAISRHPRLLSLIQD
ELFRNLMQFGLSTSPLILSMICSIVLNLYHHLRTELKLQLEAFFSCVILRLAQSKYGASYQQQEVAMEALVDFCR
LKSFMVEMYANLDCDITCSNIFEDLASLLSKSAFPVNCPLSAMHILALDGLIAVIQGMAERISNAPVNYEFTPVC
LEEYTPFWMVKCENYNDPEHWVPFVRRRKYIKKRLMIGADHFNRDPKKGLEFLQGTHLLPDKLDPQSVACFFRYT
AGLDKNLVGDFLGNHDEFCVQVLHEFAGTFDFQDMNLDTALRLFLETFRLPGESQKIQRVLEAFSERYYEQSPLI
LANKDAALLLSYSLIMLNTDQHNVQVKKKMTEDDFIKNNRHINGGNDLPRDFLCELYDSICKNEIRTIPDQSAAF
PEMNPSRWIDLMHKSKKTAPFILSDSRSHLDRDMFAVMSGPTIAAISVVFDHAEVEDVFQTCIDGFLAVAKISAC
HHLEDVLDDLVVSLCKFTTLLNPSSAEEPVLAFGDDAKARMATITVFTIANRYGDFIRAGWRNILDCILKLHKLG
LLPARVASDAADDSELSSEPGHGKPLTNSLSSSHMPSIGTPRRSSGLMGRFSQLLSLDTEEPRLQPTEQQLAAHQ
RTLQTIQKCHIDSIFTESKFLQSDSLLHLARALIWAAGRPQKGNNSPEDEDTAVFCLELLIAITLNNRDRIVILW
QWVYEHIAGIVQTTVMPCALVEKAVFGLLRICQRLLPYKENLADDLLRSLQLVLKLDARVADAYCEQITQEVSRL
VKANANHIRSQMGWRTITSLLSITARHPEASEVGFDALLFIMSDGVNLIPANFITCVDAARQFAESRVAHADRSV
RALDLMSGSIACLTQWAHEAQQAMGETDSVKTCQDIGDMWLRLVQSLRKVCLDQREEVRNHALTALEMCLTGVDR
VNIQPGVWLQCFDMVIFTVLDDLLEISQGHSQKDFRNMEGTLVLGLKLLSKVFLQMLPELSQVTTFCKLWLGVLS
RMEKYLKVKVRGKKSEKLQELVPELLKNTLIVMKNKGVLAQRSALGGDSLWELTWLHVNNIAPLIQSQVFPDENG
NGTGTEESG*

**>Lacsa_LOC111879745**

MGCPNQLPEGVTAARDTKDCSPSSSSSSSSSSSSSSSTSSAFMVNSEIGAVLAVMRRNVRWGVQYMDNEDQLEHP
LIKSFKEIRKKIFSWQHQWHTINPILYLQPFLDVIQSDETSAPITGVALSSLYKFLTLEVLDLDTVNVSDALHSI
VDAVTSCRFEVTDPASEEVVLMKILQVLLGCMKNEASVKLSNQHVCNVVNTCFQIVHQAGAKGELLQRTARHTMH
ELIRCVFMHLPDVVNKEDEVTHEVNVPSNEQSSTSNGYTGSKIESVGGSLNLSAVVPEVKMDETSGEVGQTKNHV
VTEPYGVPSVVEIFNFLCSLLNAVEHSGIAPRSNSLAYDEDVPLFALGLINSAIELGGAALSQHPEILALVQNDL
FHSLIQFGLSPSPLILSTVCSIVLNLYHLMRVKLKLQLEAFFSSVLMKIALNKHGASYQQQEVAMETLVDLCRQP
TFIHEMYANYDCDISCSNVFEDLSNLLSKSAFPVNSPLSAIHVLALEGLIAMISGMAESIGDEAPPLEPDPSSIE
DHESFWKVKCENFENPDAWVPFIRKMKHIKKKLAIGSDHFNRDPKKGLQFLQGMYLLPETLDPISVACFLRYTTG
LDKNLVGDYLGNHDQFCIDVLQEFAGTFDFRDMNLDIALRVFLETFRLPGESQKIQRVVEAFAERYYEQSPDILA
NKDAALLLSYSLIMLNTDQHNAQVKKKMTEEDFIRNNRRINGGNDLPREYLSELYRSISENEIKMTPEHGVGFPV
MTHDNWVGLIHKSRQTAPFIVCNSGDNSGDRINHEMFAILSGPTVAALSVILDLVEQEDVLQVCIDGFLDVAKIA
GSYHLDDVLDGLLVSLSKFTTLLIPMSIEDSVLAFGDDSKARKATVAVFTIANSYGDYIRSGWRNILDCILSLHK
LGLLPARLARDAADDFDSVSDPDSGKSTVTPVSTKPALPPSRKPSGLMGRFSEFLYYDTEKPAPQPSEEQVEARN
RAMETVKDCHIENVFTESKFLQSESLSSLARALMLGAGPSGISNSGDDEDAAIFCLELLITVALNNRDRIMILWQ
GVYEYIANIVQSAVMPSALVEKAVFGLIRICRRLLPYKEDLTDDLLKSLQLILKLDARVADAYCEHITQEIMRLV
KVNVGQIKSHMGWRTITSLLSITARHPEASEPGFETLEFIMSDGAHLLPANFVLCVNAARQFAESRVGDVVRSVK
SVDLMAGSGVCLVRWSREAEEASLQVYEDIGEMWLRLVQGLKKVCLDSREEIRNHAVLMLQRCVTGLDGIYLEDE
MWLQCFELVIFGVVNDLLEIVEEKSVKEYRNMEGTLVLSLKLLSKAFLHSLPCISRLVSFEKIWVGVLNGMKRYM
EVKFRGKRSEKIYELVPELLKNTLLVMKATGILSLPDSDSGGGGDSGGVWNQTWVHVKTLAPLLQSEVFPDEEPE
PRTVGATTPVPEANLLVP*

**>Lacsa_LOC111920050**

MGCPSQESEAITLAQEPMDTSCTPLRSPSAFIINTEIGAVLSVMRRNVRWGIQYIEDEDQIDRSLIKSFKEIRRE
VFSWQTQWHTINPILYLQPFLDVIQSDETSAPITGVALSSIYKFITLEVLHLNTVNVADALHKIIDAVTSCRFEV
TDPASEEVVLVKILQVLLACMKNKASVKLSNQHVCDMVNTCFQIVHQASSKGELLQRIARQTLHELVRCIFSHLP
NVATPHVNEKENGYISSKTEGATNLDMLAVKVDERIGHTKEFLTTEPYGVPCMVEIFQFLCSLLTAVENSRVGPG
LRSNSLSYDEDVPLFSLGLINSVVELGGASLGQHPELLTLVENDLFHSLIQFGLSPSPLILSTVCSIVLNLYHLL
RTKLKLQLEAFFSSVLMRIAQNKHGAKYQQQEVAMETIVDLCRQPMFIYEMYTNYDCDISCSNIFEDLVNLLSKS
AFPVNTPLTAIHVLALEGLIAMTNCISEKITGNCEVQVSEHAVPDPENEPFWKVKCENYENPGCWVPYIHKTKHM
KKKLSVGSDHFNTDPKKGLEFLQGMRMLPDTLDPVSVACFLRYTTGLDKILIGDYLGNHDRFNVDVLLEFAKTFD
FHDMNLDIALRVFLETFRLPGEAQKIQRFVEAFADQFYEQSPDIFANKDAALLLSYSLILLNTDRHSPQVKKKMT

EEDFIRNNRRINGGNDLPREYLSELYSSICENEIRMTPAQGAGVMTHDNWVGLLHKSRQTVPYIVCDSGEQINNE
MFAILSGPTLAALSVVLDLVEQDDVLQTCIDGFLKVAKIAGCYQLDDVIDGLLVALSKFTNLLTPVSMEEPVFAF
GNDTKAMKATIGVFTIANTYGDYIRSGWRNTVDCILTLHKLGLLPTRLATDTANDSDQIEDPGKSVLLSPPPPAV
LASHSRKSSGLMGRFSDFLYYDMEKPAPQPSQEHLEAHKRAVETVKSCHVNNIFTESKFLQSDSLLNLTRALILS
VGQNKNEEASVFCLELLITVTLNNRDRIMLLWENVFGYISNIVQSAVMPSTLVEKAVFGLIRICQRLLPYKENLT
DDLIKSLQLILKLDARVADAYCEHITQEIMRLVKANARQIKSHLGWRTITSLVSITACHPEASDPGFQTLEFIMF
EGTHLLPANFVLCVNAARQFAESRVGETSRSLKSLDLMSGSAICLVKWSRATSEGEEGEAADKVYQDIGEMWLRL
LQSLKKVCLDPREEIRNHAVFTLQKCLTGLDGIHLEDEMWAQCFDLVVFTLASELLEIVQEKSTKEYRNMEGTAV
LCVKLLLNVFLYSLPSLCRLVSFCKVWIGVLSCLERFMKVKIRGKTSDKIHEIVMELLKESLVVMKGNGILVESD
SVGSDSFWQLTWLHVKNIAPNLQAEIFPVS*

**>Lacsa_LOC111897921**

MGKQNEEEEEVEEEEEDDDEDDDEDEEDDEDEEDDKTIKIRKSKELGISCMLNTEVGAVLAVIRRRSDPSSYIYL
DDALDSNILSSLKSLRSLIFNPRQSWRAVDPSIYLSPFLDVISSDDVPAAATAVALSSILKMLKLDMFDKKTPGA
REAVSNVVMAVTGCRLEKTHPVTEAAVMMRILQVLAAVMRHVASVLLNDHAVCTIVNTCFQVVQQAANQGDLLQR
TARHTMHDLIQIIFSRLLEIEVGDWENSESDTEDATDETGYGIRCAVDIFHFLCSLLNVVGIVDSDGSPSLTADE
DVQVFALVLINSALELSGDGIGKHKKLLRMVKDDLFHHLIHYGTSSSPLVFSMICSTVFNIYYFHRKSIRLQLEA
FFLFVLFKTANQGYSIQLQELAIEGIINFCRQPTFIIEAYINYDCDPMFNNIFEEIGKTLCKHAFLSGGTAPTSL
QFQSFEGLMVLIHNIADFVDKQTDSGTGPYPVEITDYKPFWEETPVEGTESETWIDSVRLRKAQKRKIMIAGSHF
NRDDKKGLDYLKISKLISDSPDPRGYAMFFRYTPGLDKVKIGDFLGDPGEFNLKVLKEFTESFELTGLVLDTALR
TYLETFRLPGESQKIHRILEAFSERFYNQQSSEIFVSKDAVFILCYSVIMLNTDQHNPQVKKKMTEEEFIRNNRA
INGGNDLPREYLSDLFQSIASKAITIFGQTGSPIEMNPSRWIQLINKSNLILPYVKCGFDRRIGRDMFAVIAGPS
VATLSAIFEHAEDEEIIHECIEGLFAIARICQYGLEDTLDELISSFCKFTTLLNPYASAEETLFSFGHDLKPRMA
TLAVFTVANNFKNSIRGGWRSIIDCLLKLKKLKLLPQSVVDVGSSTANGRLHRRGESESVISKDQTGKINTSNLM
AAISHCLSMENLEESLNLGISEFEQNLKVIQQCQIGSIFSKSSTLPLDPLLNLGRSLIFAAAGKGQKFNTPVEEE
ETVQFCWDLIQAMASCNVHRLTQFWPTYNEYLLTVVQLPTFSPIPFAEKAIVALIRICLKLLASRSNEKVAEELI
FKSINMMWKLEREILDTCNDFIIQSVTKILNKYSGNLLSQLGWKTVLHLLSITGRHPESYERGVKALINLMADGS
GISVTNYPSCIDCAFGFVALRNSPVDKNMKLMDLMADSVKLIVQWSRNGYSDPGSNSSINSSASIDDNTKALGTM
NMSMNLFVKVGEVLRKTSLARREEVRNYAVTSLHKSFINADELYFTPNNIMQCFNLVIFALVDDVHEKMLEYSTR
ENAERETRSMEGTLKLAMEFFVDVYLQFLKVLVESPGFKTFWLSMLRRMDSVMKADLGVYGESKMRELVPGLLKK
MIQTMQEKEILVPQEGNDLWEITYIQIQWIAPSLKDELFPDPV*

**>CEY00_Acc06361**

MGRLKLQSGIKAIEEEPEDCETTSSNKAALACMINSEIGAVLAVMRRNVRWGGRYVSGDDQLEHSLIQSLKALRK
QIFSWQHPWHTINPAVYLQPFLDVIRSDETGAPITGVALSSVHKILTLDVLDLDTVNVEDAMHSVVDAVTSCRFE
VIDPVSEEVVLMKILQVFLACMNSKASVMLTNQHVCTIVNTCFRIVHQAGTKGELLQRIARQTMHELVRCIFAHL
QDVNDTEGSLVKGTSSIKEEIGGIDSDYRFGSKQLENGNGSSAYDSQPSSASLASNVSNGLVTGVMDESTVRSSN
GKDTVPYDLHLMTEPYGVPCMVEIFHFLCSLLNIVEHMGMGPRSNTIAFDEDVPLFALGLINSAIELGGPSIRQH
PRLLSLVQDELFRNLMQLGLSMSPLILSMVCSIVLNLYNHLRNELKLQLEAFFSCVILRLAQSRYGASYQQQEVA
MEALVDFCRQKTFMVEMYANLDCDITCSSIFEDLANLLSKSAFPVNCPLSSMHILALDGLIAVIQGMAERISNGL
VNMELPPVNLEEYTPFWMVKCDNYSDPDHWVPFVRRRKYIKRRLMIGADHFNRDPKKGLEFLQGTHLLPDKLDPQ
SVACFFRYTAGLDKNLVGDFLGNHDEFCVQVLHEFAWTFDFQDMNLDTALRLFLETFRLPGESQKIQRVLEAFSE
RYYEQSPLILANKDAALLLSYSLIMLNTDQHNVQVKKKMTEEDFIRNNRHINGGNDLPREFLSELYHSICKNEIR
TTPEQGAGFPEMTPSRWIDLMHKSKKTAPFIVSDSKAYLDHDMFAIMSGPTIAAISVVFDHAEHEDVYQTCIDGF
LAVAKISACHHLGDVLDDLVVSLCKFTTLLNPSSVEEPVLAFGDDTKARMATVTVFTIANRYGDFIRTGWRNILD
CILKLHKLGLLPARVASDAADDSELSTDPGQGKPLTNSLSSAHMPSIGTPRRSSGLMGRFSQLLSLDTEEPRSQP
SEQQLAAHQRTLQTIQKCHIDSIFTESKFLQADSLLQLARALIWAAGRPQKGNSSPEDEDTAVFCLELLIAITLN
NRDRIVLLWQGVYEHISSIVQSTVMPSALVEKAVFGLLRICQRLLPYKENLADELLRSLQLVLKLDARVADAYCE
QITQEVSRLVKANATHIRSPMGWRTIASLLSITARHPEASEAGFDALLFIMSDGAHLLPANYVLCVDAARQFAES
RVGQADRSVRSLDLMAGSFACLSRWAQETKESLGEAEAVKMSQDIGEMWLRLVQGLRKVCLDQREDVRNHALLSL
QMCLTGVDEVRLPHSLWLQCFDLVVFTMLDDLLEIAQGNSQKDYRNMEGTLIIALKLLCKVFLQLLQELSQLTTF
CKLWLGVLSRMEKYMKVKGRVKKTEKLQELVPELLKNTLLVMKTRGVLVQRSALGGDSLWELTWLHVNNISPALK
PEVFPDQDSEQSQPNEGQIGGSSETVSASNEAVAPQE*

**>CEY00_Acc31499**

MGRLKLQSGIKAIEEEPEDCETTSSNKAALACMINSEIGAVLAVMRRNVRWGSRYVSGDDQLENSLIQSLKALRK
QIFSWQHPWHTINPAVYLQPFLDVVRSDETGAPITGVALSSVHKILTLDVLDLDTVNVEDAMHLVVDAVTSCRFE
VIDPVSEEVVLMKILQVLLACMKSKASVVLSNQHVCTIVNTCFRVVHQAGTKGELLQRIARQTMHELVRCIFAHL
QDVNDIEGSLVKGTSSIKQEIGGIDSDYRFGSKQLENGNGSSAYDSQPSSASLASNVSNGLVTGVMDESTVRSSN
GKDTVPYDLHLMTEPYGVPCMVEIFHFLCSLLNIVEHMGMGPRSNTIAFDEDVPLFALGLINSAIELGGPSIRQH
PRLLSLVQDELFRNLMQLGLSMSPLILSMVCSIVLNLYNHLRNELKLQLEAFFSCVILRLSQSRYGASYQQQEVA
MEALVDFCRQKTFMVEMYANLDCDITCSNIFEDLANLLSKSAFPVNCPLSSMHILALDGLISVIQGMAERISNGL
VNMELPPVNLEEYTPFWMVKCDNYSDPDHWVPFVRRRKYIKRRLMIGADHFNRDPKKGLEFLQGTHLLPDKLDPQ
SVACFFRYTAGLDKNLVGDFLGNHDEFCVQVLHEFAWTFDFQDMNLDTALRLFLETFRLPGESQKIQRVLEAFSE
RYYEQSSQILANKDAALLLSYSLIMLNTDQHNVQVKKKMTEEDFIRNNRHINGGNDLPREFLSELYHSICKNEIR
TTPEQGAGFPEMTPSRWIDLMHKSKKTAPFIVSDSKAYLDHDMFAIMSGPTIAAISVVFDHAEHVDVYQTCIDGF

LAVAKISACHHLEDVLDDLVVSLCKFTTLLNPSSVEEPVLAFGDDTKARMATVTVFNIANRYGDFIRTGWRNILD
CILKLHKLGLLPARVASDAADDSELSTDPGQGKPLTNSLSSAHMPSIGTPRRSSGLMGRFSQLLSLDTEEPRSQP
TEQQLAAHQRTLQTIQKCHIDSIFTESKFLQADSLLQLARALIWAAGRPQKGNSSPEDEDTAVFCLELLIAITLN
NRDRIVLLWQGVYEHISSIVQSTVMPSALVEKAVFGLLRICQRLLPYKENLADELLRSLQLVLKLDARVADAYCE
QITQEVSRLVKANATHIRSPMGWRTIASLLSITARHPEASEAGFDTLLFIMSDGAHLLPANYVLCVDAARQFAES
RVGQTDQSVRSLDLMAGSVTCLSQWAQETKETLGEAEAVKMSQDIGEMWLRLVQGLRKVCLDQREDVRNHALLSL
QMCLTGVDEVRLPHSLWLQCFDLVVFTMLDDLLEIAQGNSQKDYRNMEGTLIFALKLLCKVFLQLLLELSQLTTF
CKLWLGVLSRMEKYMKVKGRVKKTEKLQELVPELLKNTLLVMKTRGVLVQRSALGGDSLWELTWLHVNNISPALK
PEVFPDQDSEQSQPNEGQVGGSSETVSAPCEAVPPEE*

**>CEY00_Acc02326**

MGCPNQQSGDNAFLKHKTSSIRPSRGGLACMVNSEIGAVLAVMRRNVRWGVRYMADDDHAEHSLIQSFKELRKQI
FSWENRWHNVSPVLYLQPFLDVIKSEETSAPITGVALSAVYKILTLDILDVDYVNVDDALHLIVDAVTSCRFEVT
DPASEEVVLVKILQVLLACMKNKSSVKLSNQHVCNIVNTCFRVVHQATSKGELLQRIAHHTMHELIRCIFSHLLD
IDSGEHAPPQGGSLSIMEQVGAPDVDDTYQNKQPNNGNVNLESNCQPPLVGSASDISLGTSAGKMDESATEVGDR
REAAQNEGHVSMEPYGIPCMVEIFHFLCSLLNVMEHIEIGPRSNPIAYDEDVPLFALGLINSAIELGGSSIGRHP
KLLALVQDELFHNLMQFGLSMSPLILSIVCSIVFNLYHHLHTKLKLQFEAFFSCVLVRVAQSKHGASYQLQEVAM
ETLVDLCRQRAFMAEMYANFDCDISSSNVFEEIANLLSKSAFPVNGPLSAMHVLALDGLIAMIQGMADRIGNDSP
TPEQASLDLEEYKPFWTIKCENYDNPNHWVTFLREMKYRKRRLIIGVDHFNRDPKKGLELLQGMHLFPDKLDPKS
VAYFFRYATGLDKNLVGDFLGNHDEFCIQVLHEFAGTFDFRDINLDTALRVFLETFRLPGESQKIQRVLEAFAER
YYEQSPSILVDTDAALLLSYSLILLNTDQHNAQVKKKMSEEDFIRNNRRINGGNDLPREYLSELYHSICQNEIRM
TPEQGAGLPVMTPSNWIALTHKSKQAAPFVICDSGAHLDYDMFAILSGPAIASISVVFDQVEQEDVLQTCIEGFL
AIAKISACYHLDDVLDDLIASLCKFTTLLRPLSVEEFVLAFGEDTRARIAIVAVFTIANKYGDFIRSGWKSILDC
ILSLYKLGFLPLRLASDAADDLKPSSESERGKPPTSSAPASRMPSAAPSRKSSGLIGRFSQLLYLDTEDSVSEPS
EEQLAAHQHTLQMIQSCHIDSIFTESKFLQSESLLQLSRAIVFAAGRYRKGSNSLEDEEIAVFCMELLIAITLNN
RDRFILLWQGVYEHIASVVQTTVMPCALAEKAVFGLLRICQRLLPYKENLSDELLTSLQLILKLDARVADAYCEH
ITQEVMHLVKANAMQIRSHMGWRTITSLLSITARHPEASEPGFETLSFIMADGAYLLPANYVLCVNAARQFAESR
VGHVERSVKSLDLMAGSVICLAKWSCKTKEAAGEESMKLYQDIGEMWLRLVLGLRKVCLDQREEVRNHAILSLQR
CLTGVDGLLLPSESWLQCFDLVIFKLLDDLLEIAQENSPKDYRNMEGSLALSMKLSSKAFLQFLQDLSQSTSFCK
LWVGVLNRMERYMKVRFRGKWSEKIHELVPELLKNTLLVMKTKGILVPHDAITEDRFWLLTWLHVKNIAPFLQEE
VFPGNELEQLQAKHIKTGGSPVSNGNSYIQAGETTA*

**>CEY00_Acc09920**

MEETTDQNREEKNSKIKNMTKLKRKELGLSCMLNTEVGAVLAVIRRLSPEPNSHFLPPQEENYDTTILQSLKSLR
GLIFNPQQEWRTMDPSMYLSPFLDVIQSDDMPAAATGVALSSILKILRLGIFDEKTPGAKGSINMAVTAITGCRL
ERTALVAEDAVMMRILQVLAAIVHHPAAVLLTDQAVCTVVNTCFQVVQQSAGRGDLLQRSARYTMHELIQTIFSR
LPDIEVRDWENSESDTEDGSLDSGYGIRSAVDIFHFLCSLLNVVEVLDNSEGFGAQNTTADEDVQLFALVLINSA
LELSGDSIGSHPKLLRMVQDDLCHHLIHYGTCSSPLVLSMICSTFLNVYHFLRRSIRLQLEAFFSFVLFKVMSLG
NFIQLQELAIEGIINFCRQPTFTIEAYVNYDCDPTCQNVFEDVGKLLCKHAFPTNGALTSLQVQAFEGLVLIIHN
IADNIDKEEEDSSISAPYPVEISAYRPFWEEKSKDGNDLDTWVEFVRVRKKQKRKLLIAGNHFNRDEKKGLEYLK
ISNLVSNPINSKAVAMFFRHTPRLDKTMIGDYLGDPDDFHLQVLKDFTDTFEFSGMILDTALRTFLETFRLPGES
QKIQRILEAFSEKFYDQQSSEVFANKDAVFILCYSLIMLNTDQHNPQVKKKMTEDEFIRNNRAINDGKDLPREYL
SELFHSISTKAITIFGQSGLPVEMNPNRWIELMNRAKLVQPFVFCDFDHRLGRDMFSTIAGPSVATLLAIFEQND
DEEILHECIEGLLSVARIAQYGLEDTLDELLSSFCKFTTLLNPYASAEETLYAFSHDMKPRMATLAVFTISNSFG
ESIRGGWRNIVDCLLKLKRLKLLPQTEYHSATSFDPETHSQSESGVVLPIPGPKLGHGRHVSGLIGRFSHFLSME
SMEENLNLGVSEFEQNLKIIQQCQIGKIFSSSSKLPDEPLLNLGRSLIFAAAGKGQKFSTTVEEEETVGLCWDLV
LTIALVNTHRFTTFWPQYHDFLLGVAQFPLFSAVPFAEKAIVALMKLCLKLLSSHQIDKLPEELIFKSINLMWKL
DKEIIDTSCEFIAQSVSKLLTKYPENLQTQLGWKSALHLLSVTSRHPETYDVGVDTLIALMSDGTHVSRSNYAYC
IDCAFGFIALKNTPSDKNMKIMDLLADSVNLLIQWYKNGYSDPGSSISMVSNTSSSSVEDNLKYLGSSNFTVNLF
IKLGEALRKTSLARREEIRNHAISALKKSFTLAEELYFTPANCMNCFNLILFAMVDDLHEKMLEYSRRENAEREM
KSMEGTLKIAMELLTEVYLQFLKLISESSGFRTFWMGILRRMDTCMKAELGDYGSSKMQEIVPVLLKRMITTMKE
NDILVQQEGHELWETTYIQIQWIAPSLKEELFPDEL*

**>Camsi_LOC114306371**

MGRLKLQSGIKSIEEEPEDCEVISSNKAALACIVNAEVGAVLAVMRRNVRWGGRYMSGDEQLENSLIQSLKALRK
QIFSWQHPWPTINPAVYLQPFLDVIRSDETGAPITGVALSSVYKILSLDVLDLNTVNVEEAMHLVVDAVTSCRFE
VTDTSSEEVVLMKILQVLLACMKSKASVRLSNQHVCTIVNTCFRVVHQAIRKGELLQQLARNTMHELVRCIFSNI
PDVNNTEDSLVKGASSFKQELGGMDNDYNFGAKQLENGNGSSEHDSQLSSAGLASNAPSGLVTGMMDENTIRSGN
RMDAVPYDSHLMTEPYGIPCMVEIFHFLCSLLNAVEHNGMGPSPRPNSMAFDEDIPLFAFGLINSAIELGGASIR
QHPRLLSLVQDELFRNLMQFGLSMSPLILSMVCSIVLNLYHHLRAELKLQLEAFFCCVILRLAQSRYGASYQQQE
VAMEALVDFCRQKTFMLEMYTNLDCDITCSNVFEDLANLLSRSAFPVNCPLSSMHILALDGLIAVIQGMAERIGN
GSVGSEPAPVNLEEYTPFWMVKCDNYSDPDHWVPFVRRRKYIKRRLMIGADHFNRDPKKGLEFLQGTHLLPDKLD
PQSVACFFRYTAGLDKNLVGDFLGNHDEFCVQVLHEFAGTFDFDGMNLDTALRLFLETFRLPGESQKIQRVLEAF
SERYYEQSPQILANKDAALLLSYSLIMLNTDQHNVQVKKKMTEEDFIRNNRHINGGNDLPREFLSELYHSICKNE
IRTTPEQGAGFPEMNPSRWIDLLHKSKKTAPFIVSDSKAYLDHDMFAIMSGPTIAAISVVFDHAEHEDVYQTCID
GFIAVAKISACHHLEDVLDDLVASLCKFTTLLNPSSVEEPVLAFGDDSKARNATVTVFNIANWYGDFIHTGWRNI

LDCILKLHKLGLLPARVASDAADDSELSSSDPGHGKLLTNSLSSAHMPSIGTPRRSSGLMGRFSQLLSLDTEEPR
SQPTEQQLAAHQRTVQTIQKCHIDSIFTESKFLQADSLLQLARALILAAGRPQKGNSSPEDEDTAVFCLELLIAI
TLNNRDRIMLLWQCVYDHISSIVQSTVIPGALVDKAVFGLLRICQRLLPYKENLADELLRSLHLVLKLDARVTDA
YCEQITQEVSRLVKANATHIRSSTGWRTITSLLCITTHHPEASEAAFDALMFIMSDGDHLLPANYVLSVDAARQF
AESRVGQADRSVRALDLMVGSVSCLARWAQEAKESGETTEAVKMSQDIGEMWLKLVQGLRKVCLDQREEVRNHAL
VSLQTCLTGADGVHLSSNLWLHCFDLVIFTMLDDLLEISQGHSQKDYRNMEGTLILALKLLSKVFLQLLHELSQL
TTFCKLWLGVLSRMEKYMKVKVRGKKSEKLLELVPELLKNTLLVMKRKGVLVQRSALGGDSLWELTWLHVSNIAP
ALQSEVFPDQDLEQPQTNEAHVSGIHETVSTPNEAVNPED*

**>Camsi_LOC114271324**

MGCLNQQSGDSAFLGEPKDCIRPSRGGLACMVNSEIGAVLAVMRRNVRWGVPYMADDDHLEHSLIQSFKELRKQI
FSWQNRWHTINPVLYLQPFLDVIQSDETGAPITGVALSAIYKVLTLDILDIDTVNVDDALHLIVDAVTSCRFEVT
DPASEEVVLMKILQVLLACMKNKASVKLSNQHVCNIVNTCFRVVHQASSKGELLQRISRHTVHELIRCIFSHLPD
IDHGEHAPPQGGSSYIKEEVGASDVDHTSSNKQPNNGVVIVENNGLPPSVGFVVSLEESAPEEVAQGKEATTQNG
GHVLTEPYGIPCMVDIFHFLCSLLNAMEHFEIGPRSNPIAYDEDVPLFALGLINSAIELGGASFGRHPKLLELIQ
DELFHNLMRFGLSMSPLILSTVCSIVFNLYHHLRTKLKLQIEAFFSCVLLRIAQSKHGASYQQQEVAMETLVDLC
RQQAFVAEMYANFDCDISCSNVFEDLANLLSKSAFPVNGPLSAMHVLALDGLNAMIQGMADRIGNDSPAPEQAAS
SDLEEYNPFWTVKCENYDDPKHWVAFVHKMKFIKKRLMIGVDHFNRDPKKGLEFLQGMHLLPDELDPKSVAYFFR
YTTGLDKNLVGDFLGNHDEFCIQVLHEFAGAFDFQGMNLDTALRVFLENFRLPGESQKIQRVLEAFAERYYEQSP
HVLANKDAALLLSYSLILLNTDQHNAQVKRKMTEEDFIRNNRRINGGNDLPREYLTELYHSICRNEIRMNTEQGV
TGFPVMTPSHWIGLMHKSKQTAPFVICDNSGAHLDYDMFAILSGPTIAAISVVFDHVEQEDILQTCIDGFLAIAK
ISACYHLDHVLDDLVVSLCKFTTLLGPLSVEESIIAFGDDTKARMATVAVFTIANRYGDFIRSGWKNILDCILSL
HKLGFRPPPLASNTAGDLEPSSESDRGRPATSSSAPPSRMLSATPSRKSSGLMGRFPTLYLDTEEEPVAQPSEEQ
LAAQQRTLQMIQNCHIDSIFTESKFLQAESLLQLAQAVVLAAGRRRKGNNNALEDEEIAVFCLELLIAITLNNRD
RFMLLWQGVYEHIASVVQSTAMPCALVEKAVFGLLRICQRLLPYKENLTDELLKSLQLILKLDARVADAYCENIT
QEVMHLVKANAMQIRSQMGWRTITSLLSITARHPEASEAGFETLSFIMADGAYLLPANYVLCVNAARQFAESRVG
LIERSVRSLDLMAGSVVCLSKWSCKSKEAHGEEEATKMNQDIGEMWLRLLQGLRKVCLDQREEVRNQAILLLQRC
LTGVDGIVVLPNELWLQCFDLVIFTLFDDLLEIAQENSPKDYRNMESSLALSMKLSSKVFLQFLHDLSQSASFCK
LWLGVLNRMERYVKVKLRGKRSEKIHELVPELLKNTLLVMKAKGILVPSDAINGDSFWQLTWLHVKNIAPSLQVE
VFPGNELEQLHTKTAE*

**>Camsi_LOC114282320**

MEEVKGQSNGNNKTKTMTKLKRKELGLSCMLNTEVGAVLAVIRRAPEPNSHFLPPQEDNSDTSILQSLKSLRALI
FNPQQEWRTIDPQMYISPFLDVIQSDDIPAAATGVALSSILKILKLEIFDEKTPGARDAMNAAVTAITGCRLERT
APVSEDAVMMRILQVLAAIMKHRASVLLTDHAVCTVVNTCFQVMTGQSASRGDLLQRSGRYAMHELIQTIFTRLP
DIEVKDWENSESDTEDTNLDCGYGIRSAVDIFHFLCSLLNVVEVVETEGLIAQTADEDVQLFALVLINSALELSG
DSIGSHPKLLRMVQDDLCHHLIHYGTCSSPLVLSMICSTVLNIYHFLRRSIRLQLESFFTFVLFRVVNLGHFIQL
QEVAVEGIINFCRQPTFIIEVYVNYDCDPTFRNVFEDIGKMLCKHAFPTGGALTSLQVQAFEGLVVIIQTIADNI
DKDDDSSPSGPYPVEISEYRPFWEEKLTDDNELATWVEFVRMRKAQKRKILIAGNHFNRDEKKGLEYLKISKLLS
DPPDPKALAIFFRYTPRIDKTMIGDYLGDPDEFHIQVLKEFTNTFEFSGMILDTALRTFLETFRLPGESQKIHRI
LEAFSEKFYDQQASEVFANKDAVFILCYSLIMLNTDQHNPQVKKKMTEEEFIRNNRAINEGKDLPREYLSELFHS
ISMNAIKLFGQSGTPIEMNPSRWIELMNRAKIMHSFILCDFDRRLGRDMFATIAGPSVASLSAIFEQADDEEILH
ECIEGLFSVARIAQYGLEDTLDELLASFCKFTTLLNPYASSEETLYAFSHDMKPRMATLAVFTTANSFGESIRGG
WRNIIDCLLKLKRLKLLPQTIIEPDNGSASSSEHGTHSKSESCVLFPKHSRRGSGLVGRFSHFLSMESMEENLNL
GVSEFEQNLKVIQQCQVGNIFTTSSKLPDEPLMNLGRSLIFSAAGKGQKFSTPVEEEETVGFCWELILTVTLANT
HRFTTFWPQYHDFLLAVTQFPLFSPVPFAEKAIVALLKICLKLLSSYQPEKLPEELIFKSINLMWKLDKEIIDTS
CEFLAQSVSKIIIDYPANLHTQMGWKSVLHLLSVTGRHPETYDLAVETLISLMSDGNNVSRTNYAYCIDCAFGFI
ALKNTPSEKNMKIMDLMADSVNMLIQWYKSGYSDPGSSVSMVSNTSCSSLEENLKSLGSTSFTVNLFVKLGEALR
KTSLARREEIRNHAISALQKSFTLAEELYFTPANCINCFNLVIFAMVDDLHEKMLEYSRRENAEREMRSMEGTLK
IAMELLTEVYLQFLKPISENSGFRTFWLGILRRMDTCMKAELGEYGNSKVQEIIPDLLNKMITTMKENEILVQKE
GDDLWEITYIQIQWIAPSLKEELFPEEM*

**>Rhosi_10G0094900**

MGRLKLQPGIKAIeeepeeteSTFSNKAALACMINSEIGSVLAVMRRNVRWGGRYVSGDDQLEHSLIQSLKALRK
QVFSWQHPWHTVNPAVYLQPFLDVVRSDETGAPITSVALSSVHKILTLDVLDLNTANVDDAMHLVVDAVTSCRFE
VTDPASEEVVLMKILQVLLACMKSKASVMLSNQHVCTIVNTCFRVVHQAGTKGELLQRIARHTMHELVRCIFAHL
QDVDDTEGSLVKGTISMKQEVGAIDNDYNFGGKQLENGNGTSEYDREqssaslassdqsGLVYGGMDGTAVRSGN
GKDAVPFDLHLMTEPYGVPCMVEIFHFLCSLLNVIEHMGTGPRSNSLAFDEDVPLFALGLINSAIEIGGPSFRQH
PRLLTLVQDELFRNLMQFGLSMSPLILSMVCSIVLNLYHNLRTELKLQLEAFFSCVILRLSQSRYGASYQQQEAA
MEALVDFCRQKTFMVEMYANLDCDITCSNIFEELANLLSKSAFPVNCPLSSMHILALDGLIAVIQGMAERISNGL
DTTERASVNLEEYTPFWMVKCDNYSDHDHWVPFVRRRKYIKRRLMIGADHFNRDPKKGLEFLQGTHLLPEKLDPQ
SVACFFRYTAGLDKNLVGDFLGNHDEFCVQVLHEFAGTFDFQDMNLDTALRLFLETFRLPGESQKIQRVLEAFSE
RYYEQSPQILANKDAALLLSYSLIMLNTDQHNVQVKKKMTEEDFIRNNRHINGGNDLPRDFLTELYHSICKNEIR
TTPEQGAGFPEMTPSRWIDLMHKSKKTAPFIVSDSKSHLDHDMFAIMSGPTIAAISVVFDHAENEDVYQTCIDGF
QAVAKISACHHledvlddlvvSLCKFTTLLNPSSVEEPVLAFGDDTKARKATVAIFDIANRYGDFIRTGWRNILD
CILKLHKLGLLPARVASDAADDLEISSSEPGHGKPAASSLSSAHMPSAGTPKRSSGLMGRFSQLLSLDTEEPRSQ

PTEQQLAAHQRTLQTIQTCHIDSIFTESKFLQADSLLQLAKALIWAAGRPQKGNTSPEDEDTAVFCLELLIAITL
NNRDRIVLLWQGVYEHISNIVHSAMTPSALVEKAVFGLLRICQRLLPYKENLADELLRSLQLVQKLDPRVADTYC
LQITQEVSRLVKANAAHIRSPTGWRTITSLLSITARHPDASDAGFDALSFVMADGAHLSPANYVLCADAARQFAE
SRVGQADQSVRSLDLMAGSVACLARWAQETKDSLGEEAEAVKMSQDIGEMWLRLVQGLRKVCLDQREEIRNHALL
SLQTCLTGVDEVNVPQTLWFQCFELVIFTMLDDLVEISQGHSQKDYRNMEGTLVFALKLLSKVFLQLVKELSQLN
TFCKLWLGVLSRMEKYMKVKVRGKKSEKLQELVPELLKNTLLVMKSKGVLVQRGALGGDSLWELTWLNVNNFAPD
LQLEVFPDQDSDQSRRSEGEAVGIGGTVSAQNEAIAASQPQTPGI*
**>Rhosi_08G0072200**
MGCPNQQSVGNAFMGEPKNCSVRPYRGGFACMVNSEIGAVLAVMRRNVRWGVRYIEEDDHIEHSLVQSFKKLRKQ
IFSWQNRWDTISSILYLQPFLDVIQSDETGAPITGVALSSVYKILTLDILDTDYVNVDDALHLIVDSVTSCRFEV
TDPASEEVVLMKILQVLLACMKNKSSVKLGNQHVCNIINTCFRVVHQASSKSELLQRIARHTMHELIRHIFSRLP
DIDGGEHAPTQGGSLYIKGEVGASDVDHTHENKQLNNGSGSLENKGSPVVGVASNVSLDATEVGEVKEASQSGGH
VLMEPYGISCMVEIFHFLCSLLNVMERVEIGPRSNPIAYDEDVPLFALGLINSAIELGGSCIGQHPKLLDLIKDE
LFNNLMQFGLSMSPLILSTVCSIGFNLYYHLRSKLKLQLEAFFSSVLVRIAQSKHGASSQLQEVAMETLVDLCRQ
QEFVAEMYANFDCDISCTNVFEDIANLLSKSAFPVNGPLSAIHVLALDGLIAMIQGMAERVENDFPAPEIAPLDH
EEYKPFWTVKCENRDDPNQWVAFQRERKQIKRKLMVGVDHFNRDPKKGFEFLQRMHLLPNKLDKKSVAYFFRYTT
GLDKNLVGDFLGNHDEFCIEVLHEFVGLFNFQEMNLDTALRVFLETFRLPGESQKIHRVLEAFAERYYEQSPHIL
ADKDAAFVLSYSLILLNTDQHNAQIKKKMTEEDFIRNNRRINGGNDLPREYLSELYHSVCQTEIRLTPDQGSGFP
IMTRSHWIGLMHDSKPASPFVSCDFGARLDYDTFAIISGPTIAAISVVFDHVEREDVLETCIEGFLSIAKISAYH
HLDQVLDDVVVSLCKFTTLLHPSSVEEFVPAFGDDTKARMATVAVFNIANRYGDLIRSGWKNIVDCVLSLHRLGF
LPRVASDGADDPKPSSDSDRGGPATSSTPPSRIPATVPSRKSSGLIGRFSQLLYLDTEDPMPQPSEEQLAAHQNI
LQIIQSCHIDSIFTESKFLQADSLVHLSRAIVLAAGRDRKRTNSAEEEEIAVFCLELLIGITLNNRDRFTLLWQG
VYDHIANVVHSTLVPCSLAEKAVFGLLRICQRLLPYKETLTDELLKSLQLILKLDARVADAYCEHITQEVMHLVK
GNAMQIRSHVGWRTITSLLSITARHPEASELGFETLSFVMADGAYLLPANYVLCVNAARQFAESRVGHVERSVKS
LDLMAGSVVCLVKWSFKTKEASMLEAKKLSEDLGEMWMRLVLGLRKVCLDPREEVRNHAILSLQRCLTGVNEVIL
PSESWVQCLDLVILKLLDDLLEIALENSPKDYGNIERSLSLSMKLLSKAFLQFLQNLSKSESFCKLWVGVLNCME
RYMKLKFRGKRSEKIHELVPELLKNTLLVMKTNGILAPGDDISGDSFWQLTWLHVKNIAPSLQGEVFPGKELEQL
AKKHLKAADSPISEGNELVPSGETTA*
**>Rhosi_03G0244100**
MLHFQREKTKGKTKNMAKLKGKELGLSCMLNTEVGAVLAVMRRRMPEPNSHFHPPQDDNCDTDVLQSLKSLRGFI
FNPQQEWRKIDPSIYLSPFVDVIQSDEVPAAATGVALSSILKILKLEIFDEKTPGAKETMNSAVTAITGCRLERT
APVAEDAVMMRILQVLTAIMHHRASVLLTDHSVCTVVNTCFQVVQQSASRGDLLQRSARYTMHELIQTIFSRLPD
IEVRDWENSESDTEDSSLDSGYGIRATVDIFHFLCSLLNVVDIMESTEGMGSQTADEDVQLFALVLINSALELSG
ESIGSHPKLLRMVQDDLCHHLIHYGACSSPLVLSMICSTFLNIYHFLRRSIRLQLEAFFSFVLFKVMSLGNIIQI
QDVTIEGIITFCRQPTFPIEVYVNFDSDPLCRNVFEEIGKLLCKHAFPAGGALTTLQVQAFEGLVLIIHNIAENI
DKEENSSIISGPYPVEISEYRPFWEEKSREGNDLETWVEYVRMRKAQKRKLLIAGNHFNRDEKKGLEYLRVSNLV
SDDPKAYAMFFRYTPRLDKTMIGDYLGDPDEFHIRVLKEFTYTFEFSGMILDTALRTYLESFRLPGESQKIQRIL
EAFSEIFYDQQSSEVFANKDAVFVLCYSLIMLNTDQHNPQVKKKMTEEEFIRNNRAINDGKDLLREYLSELFHSI
STNAITIFGQSGLPVEMNPSRWIELVNRAKTVQPFEVCDFDCRIGRDMFATIAGPSVATLSAIFEHTDEEGILHE
CIEGLFSVARIAQFGLEDILDELLASFCKFTTLLNPYASAEETLHAFSHDMKPRMATLAVFTIANSFGESIKGGW
RNIVDCLLKLKRLKLLPQSIVEPGLASNSSSDMDIHTIRSESGVVFPSQGLKQGHGRQSSGIIGRFSHFLSMESV
EESLTLGVSEFEQNLKVIQQCQVGNIFSNSSKLPDEPLLNLGRSLIFAAAGKGQKFSTPVEEEETVGFCWDLVVI
IASCNTERFTTFWPQYHDFFLAVAQFPLFSPVPFAEKAIVALMKIALKLLSSYRADRLPEELIFKSINLMWKLDK
EIIDTCCEPITLSIKKIITEYPGNLQTQLGWKSVLHLLSITGRHPGTYDMGVDTLIALMSDGTNVSRTNYAYCID
CAFSFVALKNTPSEKNMKIMDLMADSVNFLIQWYRSGYSDPGSNASMVSNASSSSLEDNMKSLGSSNFAVNLFIK
LGEALRKTSLARREEIRNHSISALQKSFTLAEDLYFTSSNCMNCFNLVIFAMVDDLHEKMLEYSRRENAEREMRS
MEGTLKIAMELLTEVYLQFLKPLSESSGFRNFWMGILRRMDTCMKAELGDYGNSIVQEIVPDLLKRMISTMKENE
ILVQREGDALWETTYIQIQWIAPSLKEELFPEEL*
**>Diool_1472.179**
MGRLKLQSGIKAIEEEPEDSETSPSSKAALACMINSEIGAVLAVMRRNVRWGGRYMSGDDQLEHSLIQSLKALRK
QIFSWHTINPAVYLQPFLDVVRSEETGAPITGVALSSVYKILTLDVLDLNTVNIEDAMHLVVDAVTSCRFEVTDP
GSEEVVLMKILQILLACMKSKASIMLSNQHVCTVVNTCFRIVHQAGTKGELLQRVARHTMHELVRCIFSHLKDVN
STEGSLVKGTSSIKHEIGGIDSDYNFGSKQLENGNGSSEYDSQPSLASLASSACSGLVTGVIDDIAVRSGNGKDS
VPYDHLMTEPYGVPCMVEIFHFLCSLLNVVEHMGMGSRSNTIAFDEDVPLFALGLINSAIELGGPSIGQHPRLLG
LVQDELFRNLMQFGLSVSPLILSMVCSIVLNLYHHLRTELKLQLEAFFSCVILRLAQSRYEASYQQQEVAMEALV
DFCRQKTFMVEMYANVDCDITCSNVFEDLANLLSRSAFPVNCPLSSMHILALDGLIAVIQGMAERITNGLVTSEQ
APVNLDEYTPFWMVKCDNYSDPDQWVPFVRRRKYIKRRLMIGADHFNRDPKKGLEFLQGTHLLPEKLDPQSVACF
FRYTAGLDKNLVGDFLGNHDEFCVQVLHEFAGTFDFQDMNLDTALRLFLETFRLPGESQKIQRVLEAFSERYYEQ
SPQILANKDAALLLSYSLIMLNTDQHNVQVKKKMTEEDFIRNNRHINGGSDLPREFLSELYHSICKNEIRTTPEH
GAGFPEMTPSRWIDLMHKSKKTASFIVSDSKAYLDHDMFAIMSGPTIAAISVVFDHAEHEDVYQTCIDGFLAVAK
ISACHHLEDVLDDLVVSLCKFTTLLNPSSVEEPVLAFGDDTKARMATVTVFTIANRYGDFIRTGWRNILDCILKL
HKLGLLPARVASDAADDSELSSEPGHGKSLTNSLSAAHMQSIGTPRRSSGLMGRFSQLLSLDTEEPRSQPTEQQL
AAHQRTLQTIQKCHIDSIFTESKFLHADSLLQLARALIWAAGRPQKGNSSPEDEDTAVFCLELLIAITLNNRDRI

VLLWHGVYEHISNIVQSTVMPCALVEKAVFGLLRICQRLLPYKENLADELLRSLQLVLKLDARVADAYCEQITQE
VSRLVKANATHVRSPMGWRTITSLLSITARHPEASEAGFDALLFIMSDGDHLLPANYVLCIDAARQFAESRGQAD
RSLRALDLMARSVTCLAQWAQDSKGALGEVDAVKMSQDIGEMWLRLAQGLRKVCLDQREEVRNHALLSLPDVFDR
SGWHPPSPWLVVTVFRPRHIHRAR*

**>Diool_124.634**

MGFLNQQHGGNTLIGESKYSSAAHYRSGLACMVNSEIGAVLAVMRRNVRWGIRYMTDDDQFEHSLIQSFKELRKQ
IFSWQNQWHTISPVLYLQPFLDVIKSDETGAPITGVALSSVYKFLSLEILDIDTVNVDDALHQVVDAVTSCRFEV
TDPGSEEVVLMKILQVLLACMKNKASVKLSNQHVCNIVNTCFRVVHQASSKGELLQRIARHTMHELVRCIFSHLC
DIANVKHAPSQGGRSYTKEVSALDEDCSLESKQLDNGSMGLENNSQPPPVEVTSNDSSGVLVGKMDECKTEVGQE
EDAENGGHVMMEPYGIPCMVEILHFLCSLLNVMDHVEIGLGPNPIAYDEDVPLFALGLINSAIELGGASIGDHPK
LLSLIQDELFHNLMQFGLSMSPLILSTVCSIVHNLYHHLRTKLKLQLEAFFSCVLLRIAKSKHGASYQQQEVAME
TLVDLCRQQEFVSEMYANFDCDISCSNVFEDLANLLSKSAFPVNGPLSAMHTLALDGLIAMIQGMASRVGNVSQP
PEQASLDLEEYKPFWMVKCESYDNPGHWVAFLRKMKYIKRRLMVGADHFNRDPKKGLEFLQGIHLLPDKLDPKSV
ACFFRYTTGLDKNLVGDYLGNHDEFCIQVLHEFARTFDFQDMTLDTALRVFLETFRLPGESQKIQRVLEAFAERY
YEQSPHILANKDAALLLSYSLILLNTDQHNAQVKKKMTEDDFIRNNRRINGGSDLPREYLSELYRSICENEIRMT
PEEGAGFPVTIPSNWISLMHKMKQAAPFVISASGAHLDYDMFAILSGPTIAAISVVFDNVEQEDILQSCIEGFLA
IAKIAACYHLDDVLNDLVVSLCKFTTLLHPLPAEETLLAFGNDNKARMATGAVFTIANRYGDFIRSGWKNILDCI
LSLHKLGFLPTRLVSDAADELEPSSELERGRPATSSGPAARILSTAHSRKSSGLMGRFSQLLYLDSEEPASQPSE
EELAAHQRILQMIQDCHIDSIFAESKFLQAESLLQLSQALILAAGRRRKGNNFLEDENIAVFSLELLIAVALNNR
DRIMLLWQGIYEHIASVVQSIVMPCALVEKAVFGLLRICQRLLPYKEDLTDELLKSLQLILKLDARVADAYCEHI
TQEVMHLVKANAMQIRSHIGWRTITSLLSITARHPEASETGFETLSFIMSGGAHLLPANYVLCVNAARQFAESRV
GHVERSIRSLDLMAGSIVCLSKWSCKTKEASGEEAMKMYQDIGEMWLRLVQGLKKVCLDQREEVRNHAILSLQQS
FTGIDGNYLPIELWFQCFDLVIFALLDDLLEIAQENSPKDYRNMEGSLALSIKLLLKVFLQFMHDLYQSASFCKL
WLGVLNRMEKYMEVKFRGKRGEKIHELVPELLKNALLVMKARGILVPSDVSKDSFWQLTWQHVKKISPSLQVEVF
PGNELEQLQVKQIQTGGTPVSDENIVVPSGQTTV*

**>Diool_968.73**

MEEVKDQKNQEEEEEEEEEEVNKKKTRKMARLKGKEVGLSCMLNTEIGAVLAVMRRPHDPNSPFLHAQDDHYDTS
ILQSLKSLRALIFNPQQEWRTMDPSMYLSPFLDVIQGDDIPASATALALSSILKILKLEIFDDKTPGARDAINSV
VTAVTCCRLERTVPIAEDVVMMRILQLLTIIMRHPASILLTDHAVCTIVNTCFQVVQQSANRMDLLQRGARYAMQ
DLVQTIFSQLPDIEVNDWDNSESDTEDNNLDSGYGIHSAVDIFHFLCSLLNVAENVDSEGVAAQTADEDVQLFGM
VLINCTLEQSGNSIGSHPKLLRMVQDDLCHHLIHYGTRSSPLVLSMICSTFLNIYHFLRRSIRPQMEAFFTFVLF
TVVGSANFAQLHEVAIEGIVNFCRQPTFAIEAYVNYDCDPVCCNVFEEIGKVLCKHAFPTVGALTSLHAQAFEGL
IVIIHNIADNIDKEGDSSPSGPYPVEVSKYTPYWVEKSKDENDLEGWVDFVRVRKAQKRKILIAGNHFNRDEKRG
LEYLKFSNLVSDPPDPKAFAIFFRYTPRLDKTKIGDYLGDPGEFNLRVLKEFADTFKFSGMILDTALRTYLEAFR
LPGESQKIQRILEPFSEKFYDQQTSEVFADKDAVLVLCYSLIMLNTDQHNPQVKKKMTEEEFIKNNRAINAGKDL
PREYLSELFHSISTNAITVFSQSGLPVEMNPTRWVELMNRAKLMQPFVSCDFDRRLGRDMFASIAGPSVAVLTAI
FEQADEEEILHECIEGLLSVARISQYRLEDTLDELLASFCKFTTLLNPYASAEETLYAFGHDMKPRLATLAVFTI
ANNFGESIRGGWRNIIDCLLKLKRLKLLPQSIVEPDNFSISSFDHETNPRSESGVLFRGQSLKRGHARKGSSMIG
RFSQFLSMENVEENLNLGVSEFEQNLKVIQQCQVGNIFSSSSKLPEEALLNLGRSLIFAAAGKGQKFSTPIEEEE
TVGFCWDLVQIIVFANIHRFTTFWPQYHDYLLAVTQFPLFSPIPFAEKAVIALIKIGLKLLASYQTEKLPEELIF
KSINFMMKLDKEIIETCSDFLAQSVSIMITEHPANLQTALGWKSVLHLLSILGRLPQAYDRGVEALITLMSDGSH
VSRVNYNYCIECAFGYIALKNTPLEKNLKIMDLMSDSVNLLVQWYKSDQYSDPGSTVSMSSATSGSCVEDNSKAI
ITFFVKLGEALRKTSLARREEIRNHAILSLQKSFKLAEELSFTPANCIYFFNLVVFAMVDDLHEKMLEYSRRENA
EREMRSMEGTLRIAMELLTEAYLQFLKTIYESSNFRTFWLGILRRMDTCMKADLGEYYGQSRVQEIVPELLKKMI
TTMKEKHILVRKEGDDLWETTYIQIQWIAPSLKDDLFPEEEF*

**>Diool_92.671**

MEEVKDQKNQEEEEEEEEQVNKQKTRKMARLKGKELGLSCMLNTEIGAVLAVMRRPHDPNSPFLHAQDDHYDTSI
LQSLKSLRALIFNPQQEWRTMDPSMYLSPFLDVIQGDDIPASATALALSSILKILKLEIFDDKTPGAREAINSVV
TAVTCCRLERTVPIAEDVVMMRILQLLTIIMRHPASILLTDHAVCTIVNTCFQVVQQSANRMDLLQRGARYAMQD
LVQTIFSQLPDIEVNDWDNSESDTEDSNFDSGYGIHSAVDIFHFLCSLLNVAENVDSEGVAAQAADEDVQLFGMV
LINCTLEQSGNSIGSHPKLLRMVQDDLCHHLIHYGTRSSPLVLSMICSTFLNIYHFLRRSIRLQMEAFFTFVLFT
VVGSANFAQLHEVAIEGIVNFCRQPTFAIEAYVNYDCDPVCCNVFEEIGKVLCKHAFPTVGALTSLQAQAFEGLI
VIIHNIADNIDKEGDSSPSGPYPVEVSKYTPYWVEKSKDENDLEGWVDFVRVRKAQKRKILIAGNHFNRDEKRGL
EYLKFSNLVSDPPDPKAFAIFFRYTPRLDKTKIGDYLGDPGEFNLRVLKEFADTFKFSGMILDTALRTYLEAFRL
PGESQKIQRILEPFSEKFYDQQTSEVFADKDAVLVLCYSLIMLNTDQHNPQVKKKMTEEEFIKNNRAINAGKDLP
REYLSELFHSISTNAITVFSQSGLPVEMNPTRWVELINRAKLMQPFVLCDFDRRLGRDMFVSIAGPSVAVLTTIF
EQADEEEILHECIEGLLSVARISQYRLEDTLDELLASFCKFTTLLNPYASAEETLYAFGHDMKPRLATLAVFTIA
NNFGESIRGGWRNIIDCLLKLKRLKLLPQSIVEPDNFSTTSFDHETNPRSESGVLFRGQSLKRGHARKGSSMIGR
FSQFLSMENVEENLNLGVSEFEQNLKVIQQCQVGNIFSSSSKLPEEALLNLGRSLIFAAAGKGQKFSTPIEEEET
VGFCWDLVQIIVFANMHRFTTFWPQYHDYLLAVTQFPLFSPIPFVEKAVIALIKIGLKLLASYQTEKLPEELIFK
SINFMMKLDKEIIETCSDFLAQSVSIMITEYPTNLQTALGWKSVLHLLSILGRLPQAYDRGVETLITLMSDGSHV
SRVNYNYCIECAFGYIALKNTPLEKNLKIMDLMSDSVNLLVQWYKSDQYSDPGSTVSMSSATSGSCMEDNSKAII
TFFVKLGEALRKTSLARREEIRNHAILSLQKSFKLAEELSFTPANCIYFFNLVVFAMVDDLHEKMLEYSRRENAE

REMRSMEGTLRIAMELLTEAYLQFLKTIYESSNFRTFWLGILRRMDTCMKADLGEYYGQSRVQEIVPKLLEKMIT
TMKEKHILVRKEGDDLWETTYIQIQWIAPSLKDDLFPEEEF*
**>Rorgo_13293**
MGRLKLQSGIKAIEEEPEECESISSTKAALACMINSEIGAVLAVMRRNVRWGGRYVSGDDQLEHSLIQSLKALRR
QIFSWQHPWHTVNPSVYLQPFLDVIRSDETGAPITGIALTSVQKILTLDVLDLNTVNVEDAMHLVVDAVTSCRFE
VTDPVSEEVVLMKILQVLLACMKSKASVMLSNQHVCTIVNTCFRVVHQAGTKGELLQRIARHTMHELVRCIFAHL
QDVNDTEGSMVKGTSSFKQELGGMDNDYNFGGKQLENGNGSSAYDSQQSSASLASNASSGLVSGVMDEITLRSGN
GKDVVPYDLHLMTEPYGVPCMVEIFHFLCSLLNVVEHMGIGPRSNTISFDEDVPLFALGLINSAIELGGPSIRQH
PRLLSLVQDELFRNLMQFGLSMSPLILSMVCSIVLNLYHHLHTELKLQLEAFFSCVILRLAQSRYGASYQQQEVA
MEALVDFCRQKTFMVEMYANLDCDITCSNVFEDLANLLSRSAFPVNCPLSSMHILALDGLIAVIQGMAERISNEL
VSMELAPVHLEEYTPFWMVKCDNYSDPDHWVPFVRRRKYIKRRLMIGADHFNRDPKKGLEFLQGTHLLPDKLDPQ
SVACFFRYTAGLDKNLVGDFLGNHDEFCVQVLHEFAGTFDFQDMNLDTALRLFLETFRLPGESQKIQRVLEAFSE
RYYEQSPQILANRDAALLLSYSLIMLNTDQHNVQVKKKMTEEDFIRNNRHINGGSDLPREFLTELYQSICKNEIR
TTPEQGAGFPEMNPSRWIDLMHKSKKTAPFIVSDSKAYLDHDMFAIMSGPTIAAISVVFDHAEHEDVYQTCIDGF
LAVAKISACHHLEDVLDDLVVSLCKFTTLLNPSSVEEPVLAFGDDAKARMATVTVFTIANRYGDFIRTGWRNILD
CILKLHKLGLLPARVASDAADDSELSPDPGQGKPITNSLSSAQMPSIGTPRRSSGLMGRFSQLLSLDTEEPRSQP
TEQQLAAHQRTLQTIQKCHIDSIFTESKFLQADSLLQLARALIWAAGRPQKGNSSPEDEDTAVFCLELLIAITLN
NRDRIVLLWQGVYEHISSIVQSTVMPCALVEKAVFGLLRICQRLLPYKENLADELLRSLQLVLKLDARVADAYCE
QITQEVSRLVKANATHIRSPMGWRTVASLLSITARHPEASESGFDALLFIMSDGAHLLPANYVLCLDAARQFAES
RVGQADRSLRSLDLMADSVACLARWAQETKESLGEAEAAKMSQDIGEMWLRLVQGLRKVCLDQREEVRNHALLSL
QTCLTGVDEVCLPPSLWLQCFDLVIFTMLDDLLEIAQGHSQKDYRNMEGTLVFALKLLSKVFLQLLHELWQLTTF
CKLWLGVLSRMEKYMKVKVRGKKSEKLQELVPELLKNTLSEMKIKGVLVQRSALGGDSLWELTWLHINNIAPALQ
TEVFPDEDMEQLPPNDSQIGGSLETVSSPNEAVTLEQ*
**>Rorgo_03512**
MGCPYQQSGDNAFLGGHKPCSIRSSRGGLACMVNSEIGAILAVMRRNVRWGVRYMEDEGHVEHSLIQSFKELRKQ
IFSWQNRWHAINPVLYLQPFLDVIKSDETSAPITGVALSSVYKILTLDIVDIDCANVDDALHQVVDAVTSCRFEV
TDPASEEVVLVKILQVLLACMKNKSSVKLSNQLVCNIVNTCFRVVHQASSKGELLQRIARHTMHELVRCIFSHLP
DIDCGEHAPSQGGSLCIEEEVGAPDADHSSGNKELNNGSVSLESTGQPPLVGSASSISLGTLAGKMDEGATDTGS
KEAGENEVHVLMESYGIPCMVEIFHFLCSLLNAVEHIEIGPRSHPIAYDEDVPLFALGLINSAIELGGSSIGRHP
KLLALVEDELFHSLMQFVLSMSPLILSVVCSIVFNLHHHLRSKLKLQFEAFLSCVLLRVAQSKHGASYQLQEVAM
ETLVDLCRQREFMAEMYANFDCDISCSNVFEDIANLLSKSAFPVNGPLSAMHVLALDGLIAMIQGIADRIGNDSP
AQEQASLDLQEYKPFWTVKCEDYDDPYKWVAFLREMKFIKRRLIVGVDHFNREPKKGLEFLQGMHLLPDKLDPES
VAYFFRYTTGLDKNLVGDFLGNHDEFCIQVLHEFAGTFDFRDMNLDTALRVFLETFRLPGESQKIQRVLEAFAER
YYEQSPQILVDKDAALLLSYSLIMLNTDQHNAQVKKKMSEDDFIRNNRRINGGKDLPREYLSELYHSIYKNEIRM
TPDQGASFPFMTASNWIALMHRSKQAAPFVICDSGAHLDYDMFSILSGPAIASISVVFDHTEQEDVLQACIEGYL
AIAKISACYRLDDVLDDLIVSLCKFTTLPRPLSVEESILTFGEDTRARMATVAVFTIANRYGDFIRTGWKSILDC
ILSLYKLGFLPLCLACDAADDLKPSSDSEQVKPATSSAPAAPSRKSSGLIGRFSQLLYLDSEDPEPQPSEEQLIA
HQRTLQMIQSCHIDSIFAESKFLQAESLLQLSRAIVLAAGRYRKGSNSLEDDEIAVFCMELLIGITLNNRDRFML
LWQGVYEHIASVVQSIVMPCSLAEKAVFGLLRICQRLLPYKENLSEELLKSLQLILKLDARVADAYCENITQEVM
HLVKANAMQIRSHMGWRTITSLLSITARHPEASEAGFETLSFIMADGAYLLPANYVLCVNAARQFAESRVGHVER
SVKSLDLMEGSVICLAKWSSKTKEASGEEAMKLYQDIGEMWLRLVLGLRKVCLDQREEVRNHAILSLQQCLTGVD
GLLLPNESWLQCFDLVIFKLLDDLLEIAQETSPKDYRNMEGSLALSMKLSSKAFLQFLQDLSQSTSFCKLWVGVL
NRMERYMKVRFRGKRIEKIHELVPELLKNTLLVMKTKEILVPSDAIGGDSFWQLTWLHVKNIAPSLQEEVFPGNE
LEQLQAKHIKTGASPVSDGNSYIPAGETAA*
**>Rorgo_08858**
MEEESKNQNREENLEKSKTMTKLKRKELGLSCMLNTEVGAVLAVIRRWSPDPNSHFLPPPEENYDIAIIQSLKSL
RGLIFNPQQEWRTMDPSMYLSPFLDVIQSDDVPAAATGLALSSILKILRLEVFDSKTPGAKEAINGAVTSITGCR
LERVAPVAEDAVMMRILQVLAAIVKHHAAVLLTDHAVCTVVNTCFQVVQQSANRGDLLQRSARYTMHELIQTIFA
RLPDIEVRDWENSESDTEDTNLDSGYGIRSAVDIFHFLCSLLNVAEVVVHSGDGMTTQTTSSDEDVQLFALVLIN
SALELSGDSIGAHPKLLRMVHDDLCHHLIHYGTCSSPLVLSMICSTFLNIYHFLRRSIRFQLEAFFSYVLFKVMS
LGNIIQLQEVAIEGMINFCRQPTFTIEAYVNYDCDPLCRNVFEDVGKLLCKHAFPTGGILTSLQVQAFEGLVLIV
HNIADSIDKEDDSSISGPYPVEILEYTPFWEEKSKDGVDLDTWVEFVRVRKSQKRKLLIAGNHFNRDEKKGLEYL
KISNLVSDPPDPTAFAMFFRYTPGIDKTMIGDYFGDPDEFHLKVLREFTNTFQFSGMIIDTALRTYLETFRLPGE
SQKIARVVEVFSEKFYDQQSSEVFANKDAVYILCYSLIMLNTDQHNAQVKKKMTEEEFIRNNRAINDGKDLPREY
LSELFYSISTNAIAIFEQSIVPVDMKPSRWIELMNRAKLVQSLILCDFERRLGRDMFATIAGPTIATLSAIFEQM
DEEETLHECLEGLFSVARIAQYGLEDTLDELLACFCKFTTLLNPYASAEETLYAFSHDMKPRMATLAVFTIANSF
GESIRGGWRNIVNCLLKLKRLKLLPQQSVIEPELAPAAPFDTQTHSKPEPNGVFPGKHGHGRQSSGTIGRFSHFM
SMESVEENLNLGLSEFEQNLKVIHQCQVGKIFSNSSNLPDEPLLNLGRTLIFAAAGKGQKFSTPVEEEETVGFCW
ELILAIALANTHRFTHFWPQYHDFLLGVAQFPLFSPVPFAEKAIVALMKLCLKLLSSHQINKIPEELVFKSINLM
WKFDKEIIDTSCEFLAQSISKIITKYGENLQTQLGWKSVLYLLSVTGRHPETYDIGVEALIELMSDGTHVSRTNY
GYCIDCAFGFIALKNTPSEKNMKIMDLMADSVKFLIQWYKSGYSDPGSTASNTSSSSIEDNTKALGSSNFTVNLF
IKLGEALRKTSLARREEIRNHAIFALQKSFTLAEELYFTPSNCINCFNLVIFAMVDDLHEKMLEYSRRENAEREM

RSMEGTLKISMELLTQVYLQFLKPISESSGFRTFWLGILRRMDTCMKAELGEYGSSIVQEIVPSLLKRMITTMKE
NEILAQKEGDDLWEITYIQIQWIAPTLKEELFPEEL*

**>Clear_17208**

MGRLKLQSGIKAIEEEPEDCESTSSNKAALACMINSEIGSVLAVMRRNVRWGGRYVSGDDQLEHSLIQSLKALRK
QVFSWQHPWHTINPSVYLQPFLDVIRSDETGAPITGVALTSVHKILSLDVLDLNTVNVEDAMHLVVDAVTSCRFE
VTDPASEEVVLMKILQVLLACMKSKASVMLSNQHVCTIVNTCFRVVHQAGTKGELLQRIARQTMHELVRCIFAHL
QDVNDTEGSLVKGTSSTKQEFVGIDSDYNFGGKQLENGNGSSEYDSQKSSVSLASSAQSGLLTGVMDDSTGRSGN
GKDAVPYDLHLMTEPYGVPCMVEIFHFLCSLLNVIEHMGMGPRSNTIPFDEDVPLFALGLINSAIELGGPSIRQH
PRLLSLVQDELFRNLMQFGLSMSPLILSMVCSIVLNLYHHVRAELKLQLEAFFSCVILRLAQSRYGASYQQQEVA
MEALVDFCRQKTFMVEMYANLDCDITCSNVFEDLANLLSKSAFPVNCPLSSMHILALDGLIAVIQGMAERISNGL
VTMEPASVNLEEYTPFWMVKCDNYNDPDHWVPFVRRRKYIKRRLMIGADHFNRDPKKGLEFLQGTHLLPDKLDPQ
SVACFFRYTAGLDKNLVGDFLGNHDEFCVQVLHEFAGTFDFQDMNLDTALRLFLETFRLPGESQKIQRVLEAFSE
RYYEQSPLILANKDAALLLSYSLIMLNTDQHNVQVKKKMTEEDFIRNNRHINGGNDLPRDFLTELYHSICKNEIR
TTPEQGSGYPEMTPSRWIDLMHKSKKTAPFVVSDSKAYLDHDMFAIMSGPTIAAISVVFDHAEHEDVYQTCIDGF
HAVAKISACHHLEDVLDDLVVSLCKFTTLLNPSSVEEPVLAFGDDAKARKATVAVFEIANGYGDFIRTGWRNILD
CILKLHKLGLLPARVATDAADDYSELPSDPAGHGKPPLTNSLSSAHMPSSIGTPRKSSGLMGRFSQLLSLDTEEP
RSQPTEQQLASHQRTIQTIQTCHIDSIFTESKFLQADSLLQLARALIWAAGRPQKGGNTSPEDEDTAVFCLELLI
AITLNNRDRIVLLWQGVYEHIASIVQSTVMPGALVEKAVFGLLRICQRLLPYKENLADELLRSLQLVLKLDARVA
DAYCEQITQEVSRLVKANATHIRSPMGWRTITSLLSITARHPEASEAGFDALCFIMSDGAHLLPANYVLCADAAR
QFAESRVGQADRSVRSLDLMAGSVACLARWAQESSKETLGEADTVKMSQDLGEMWLRLVQGLRKVCLDQREEVRN
HALLSLQMCLTGVDEVNLPHGLWLQCFDLVIFTMLDDLHEIAQGHSQKDYRNMEGTLVFALKLLSKVFLQLLQEL
SQLTTFCKLWLGVLGRMEKYMNVKVRGKRSEKLQELVPELLKNTLLVMKTKGVLVQRSALGGDSLWELTWFNVNK
IAPGLQSEVYPDQDSEPHESDTGGNPETVSAPNEEEA*

**>Clear_18756**

MGCPNQQSGDNAFIGEPKNCSIRPPGGGLACMVNSEIGGVLAVMRRNVRWGVRYMADDDHLEHSLIQSFKELRKQ
IFTWKNRWDTINPVLYLQPFLDVIQSEETGAPITGVALSSVYKILTLDILDMAYANVDDALHLVVDAVTSCRFEV
TDPASEEVVLMKILQVLLACMKNKASIKLSNQHVCNIVNTCFRVVHQASSKGELLQRIARHTMHELVRCIFSHLP
NIDHREDAPSQGGGLYNKEEVGAPDGDHTHGNKQLNSGNVSLENNGQPPLVGFGSNVSLGALAGKTDESATEVGG
QEAAQNGEHVLVEPYGIPCMVEIFHFLCSLLNVMDRIEIGPRSNPIAYDEDVPLFALGLINSAIELGGASIGQHP
KLLALIKDELFHNLMQLGLSMSPLILSTVGSIVFNLYHHLRTKLKLQLEAFFSCVLLRIAQSKQGASYQLQEVAM
ETLVDFCRQQEFVAEMYANFDCDISCSNVFEDIANLLSRSAFPVNGPLSAMHVLALDCLIVIVQGMADRMGNDSP
APEQASLNLEEYKPFWTVKCENYDDPNHWVAFLREKKFIKRRLMIGVDHFNREPKKGLEFLQGMHLLPDKLDPKS
VACFFRYTTGLDKILVGDFLGNHDEFCIQVLHEFVGTFDFQGMNLDTALRVVLETFRLPGESQKIQRVLEAFAER
YYEQSPRILADKDAALLLSYSLILLNTDRHNAQVKKKMTEEDFIRNNRRINGGNDLPREYLSELYHSICRNEIRM
TPEQGAGFPVLTSSHWVGLMHESKQATSFVICDSGAHLDYDMFAIISGPTIAAISVVFDHAEQEDVLQTCIEGFL
AVAKISACYHLANVLDDLVVSLCKFTTLLHPLSVEESVLAFGDDVKATKATAAVFTIANRYGDFIRSGWKNILDC
ILSIHKLGFLPPRLVNDAADDVKPSSDSDKGRASRMPSTAPSRKSSGLIERFSQLLYFDTEDPVPQPSEEQLAAQ
QHTLQMIQSFHIDSIFTDSKFLQAESLLQLSRAIVLASERYRKGANSLEEDEGIAVFCLELLIGITLNNRDRFML
LWQGVYDHIASVVQSTVMPCALAEKAVFGLLRICQRLLPYKENLADELLKSLQLILKLDARVADAYCENITQEVM
HIVKANAMQIRSHVGWRTITSLLSITAQHPEASEAGFETLSFVMADGAYLLPANYVLCANAARQFAESRVGNVER
SVKSLDLMAGSVVCLAKWSFKTKEASGEEATRLCKDIAEMWLRLVLGLRKVCLDQREEVRNHAILSLQRCFTLGD
GIILPNELWLQYFDLVIFKLLDDLLEIAQEHSPKDYRNMEGSLALSMKLSSKAFLQLLQDLSQSTSFCRIWVGVL
NRMERCMKVKFRGRRSEKIHELVPELLKNTLFVMKTNGILVPSDAVGGDSFWKLTWLHVKNIAPSLQIEVFPVNE
LEQLQTKHLQTVGSSPISDGNLLVQSGETTA*

**>Clear_06455**

MEEQENGNNKSKMKNMGRLKGKELGLSCMLNTEVGALLAAIRRRMLEPNSHFLPQEESFDTTILQSLKSLRGLIF
NPQQEWRKIDPSIYLAPFLDVITSDEVPAAATGVALSSILKILKLEIFDEKTPGVKEAMNSAVTAITGCRLERTA
PVAEDAVMMRILQVLNGMMQHRASVLLTDHSVCTIVNTCFQVVQQSASRGDLLQRSARYTMHELIQTIFSRLPDI
EVKDWENSESDTEDTATLDSGYGIQSAVDIFHFLCSLLNVVEVVESSDTADEDVQLFALVLINSALELSGESIDS
HPKLLRMVQDDLCHHLIHYGTCSSQLVLSMICSTFLNIYHFLRRSIRLQLEAFFSFVLFKVMSLGNIIQLQEVAI
EGIITFCRQPTFTIEAYVNFDCDPLCRNVFEDAGKLLCKHAFPAGGSGNAGLTSLQVQAFEGLVLIIHNIAENIE
KEENRSSSLSGGQYPVEVSAYRPFWEEKSREGSDLDAWVEFVRERKARKRKLLIAGNHFNRDEKKGLEYLKVSNL
VSDDPKAYAMFFRYTPRLDKTLIGDYLGDPDEFHIEVLKEFTNTFEFSGMILDTALRTFLETFRLPGESQKIQRI
LEAFSEKFYDQQSSEIFTNKDAVFILCYSLIMLNTDQHNAQVKKKMTEEEFIRNNRAINDGKDLPREYLSELFHS
ISSNAITIFGQSGLPVEMNPSRWIELMNRAKTSQPFVLCEFDRCIGRDMFATIAGPSVATLSAIFEQTEEEEILH
ECIEGLFSVARIAQFGLEDTLDELLASFCKFTTLLNPYASTEETLYAFSNDMKPKMATLAVFTIANNFGESIGGG
WRNIVDCLLKLKRLKLLPQSIIEPENSSPSSFDNEKNSRHDLKLGHERHSSGIIGRFSHFLSMESIEENLNLGVS
EFEQNLKIIQQCQVGSIFSNSSKLPDETLLNLGRSLIFAAAGKGQKFSTPIEEEETVGFCFELVLIIASSNSHRF
TTFWPQYHDFLLGVTQFPLFSPVPFAEKAIVALIKICLKLISSYQPDKTPEELIFKSINLMWKLDKEIIDTCCEF
ITQSISKIITDYAQNLQTQLGWKSVLHLLSVTGRHPDTYDIAVETLMALMSDGAYISRANYAYCIDCAFGFVALK
NTPSEKNMKILDLMADSVNFLIQWYKSGYSDPGSSVSMVSNTSSSSIEDLKSIGSSSFAVNLFIKLGEALRKTSL
ARREEIRNHAILALQKSFLLAEELYFSPTNCMNCFNLVIFAMVDDLHEKMLEYSRRENAEREMRSMEGTLKIAME

LMTEVYLQFLRPISEGSGFRNFWLGILRRMDTCMKAELGEYGNSSVQEIVPDLLKKMIVMMKENEILVAREGDAL
WETTYIQIQWIAPLLKEELFPDEL*
**>Aegco_05G0197980**
MGRLKQPSGINAIEEEPQDGETTTSKNAALACMINSEIAAVLAVMRRNVRWGVRYMSGDDQLEHSLIQSLKALRK
QIYGWKHPWHTINPAVYLQPFLDVIRSDETGAPITGVALSSVYKILTLDVLDLNTVNVKEAMLLVVDAVTSCRFE
VTDPASEEVVLMKILQVLLACIKSKTSVVLSNQQVCTVVNTCFRVVHQAGTKGELLQRIARQTMYELVRRVFLHL
SDVKNSEHSQVKGASPDKQEVGVIDNSYSFSSNASENSNGSSEHESQTFSVNLSSNAPSGLATGMIDEHTVQSSN
GKAAVQYDLHVMNEPFGVPCMVEIFQFLCSLLNVVEHMGLGPGSNTIAFDEDMPLFALGLINSAIELGGPSIREH
PRLLSLVQDELFRNLMQFGLSTSSLILSMVCSIVLNLYHHLRFELKLQLEAFFSCVILRLAQSRYGASYLQQEVA
MEALVDFCRQKNFMVEMYVNLDCDITCCNIFEDLANLLSKSAFPVKCPLSSMHILALDGLISVIQGLAERISNGL
GTSELDPVNLEEYIPFWMVRCDNYADPDQWVSFVRRRKYIKRRLMIGADHFNRDPKKGLEFLQGTHLLPDKLDPQ
SVACFFRYTAGLDKNLVGDFLGNHDEFCVQVLHEFARTFDFEGMNLDTALRLFLETFRLPGESQKIQRVLEAFSE
RYYEQSPETLADKDAALVLSYSLIMLNTDQHNAQVKKKMTEEDFIRNNRRINGGNDLPREFLSELYHSICKNEIR
TTPEQGMGFPEMTPSRWVDLMNKSRKTPQFIVSDSNAYVDHDMFAIMSGPAIAAISVVFDHAEHEEVYQTCIDGF
LAIAKISSCHHLEDVLDDLVVSLCKFTTLLNPEVPVLAFGDDMKARMATITVFTIANTYGDFIRTGWRNILDCIL
KLHKLGLLPARVASDAADDTELSSDHGQGKPLTNSLSSSSLSSVSTPRRSSGLMGRFSQLLSLDTEEPRSQPTDQ
QLAAHQCTLQTIQKCHIDSIFTESKFLQADSLLQLARALIWSAGRPQKGGSSPEDEDTAVFCLELLIAITLNNRD
RIGILWQGVYDHITNIVQSTVMPCALVEKAVFGLLRICQRLLPYKENLADELLRSLQLVLKLDARVADAYCEQIT
QEVSRLVKANASHIHSPMGWRTITSLLSITARHPEASEAGFDALFFIMSDGAHLSLANFVLSIDSARQFAESRVG
QVDRSVKALDLMATSVLCLTRWAQEAKTSLGEAEAFKMSQNIGEMWLRLVQALRKVCLDQREEVRNHALFSLQTC
LMGGDEIYLSNGFWLQCFDLVIFTVLDDLLEIAEGQSQKDYRNIEGTLILALRLLSKVFLQLIHELSQLTTFCKL
WLGVLGRMEKYMKAKVRGKKSDKLQELVPELLKNTLLVMKTRGVLVQRSALGGDSLWELTWLHVKNVSPDLQNEV
FSDQDFAQTLPGHESVPSIDETVTPEVARVEG*
**>Aegco_14G0264380**
MGRLNKPSGIKAIEEEPEDRETTLSNNAALACMINSEIAAVLAVMRRNVRWGGQYVSGDDQFEHSLIQSLKALRK
QIYGWKDPWHTVNPAVYLQPFLDVIRSDETGAPITGVALSSVYKILTLDVLDLNTVNVKEAMLLVVDAVTSCRFE
VTDPSSEEVVLMKILQVLLACIKSKTSVVLSNQQVCTIVNTCFRVVHQAGTKGELLQRIARQTMYELVRRVFLYL
SGVNNSERLMLKGAGPIKQEVGVMDNSYSFSSKASENSNGSSEYDSQTFSVSLSSNALSTGMIDEYTVQSSNSKA
AAPYDLHIMNEPFGIPCMVEIFQFLCSLLNVVENMDLGPGSNTIAFDEDMPLFALGLINSAIELSGPSIREHPRL
LSLVQDELFRNLMQFGLSLSPLILSMVCSIVLNLYHHLHTELKLQLEAFFSCVILRLAQSRYGAYYLQQEVAMEA
LVDFCRQKAFMVEMYINLDCDITCCNVFEDLANLLSRNAFPVNCRLSSMHILALDGLISVIQGMAERISNGSVTS
GQGPVNLEEYIPFWMVKCDNYSDPQQWVPFVRRRKFIKRRLMVGADHFNRDPKKGLEFLQGTHLLPDKLDPQSVA
CFFRYTTGLDKNLVGDFLGNHDEFCVQVLHEFAGTFDFEGMNLDTALRLFLETFRLPGESQKIQRVLEAFSERYY
EQSPETLADKDAALVLSYSLIMLNTDQHNAQVKKKMTEEDFIRNNRRINGGSDLPRKFLSELYHSICKNEIRTTP
EQGMGFPEMTPSRWVDLMNKSRKTPRYIASDSNAYVDHDMFAIMSGPAITAILAVFDRAENEEVYQTCIDGFLAI
AKISSCHHLEDVLDDLVVSLCKFATHLNPLFMDVPVLAFGDDAKARMATITVFSIANTYGDFIRTGWRNILDCIL
KLHKLGLLPARVASDAADDTELSSDSGQGKPLTSSLSSSNASSVGTPRRSSGLMGRFSQLLSIDTEEPRSQPTDQ
QLAAHQRTLQTIQKCRVDSIFTESKFLQADSLLHLARALIWAAGRPQKGGTSPEDEDTAVFCLELLIAITLNNRD
RIGILWQGVYDHITNIVQSTVMPCALVEKAVFGLLRICQRLLPYKENLADELLRSLQLVLKLDARVADAYCEQIT
QEVSRLVKANAPHIRSPMGWRTITSLLSITARHPEASEAGFDALSFIMSDGAHLSPANFVLCVDSARQFSESRVG
LVDRSVKALDLMASSVLCLSKWAQEAKGSLGEAEAFKMSQNIGEMWLRLVQALRKVCLDQREEVRNHALFSLQLC
LICWDKINISNGFWLQCFDLVIFTVLDDLLEIAEGQSQKDYRNMEGTLTLALKLLLKVFLQLIHELSQVTTFCKL
WLGVLGRLEKYMKVKVRGKRSEKLQELVPELLKNTLFVMKTRGVLVQRSALGGDSLWELTWLHVKNISPNLQNEV
FPDKDLDQTSLGHKTVSSTDETITPESARVES*
**>Aegco_02G0321210**
MEAENKDGTMTKLKRKELGFLCMLNTEIGAVLALIRRAPDLQNPYFLPNQDDTYDSTVLQSLKSLRSLMFSLEQE
WRTVDPSMYLGPFLDVIQSDSIPAPATGVAISSVLKILKLDVFDEKTLGAKQAINSVVTAITSCRLERTAPTAEE
AIMMRILQVLTAVMKHRASVLLTDQAVCTIVNASFQVLQQSTAHGDLLQRSARYTMHELIQTIFSYLPDIEVKDN
ENSESDTEENNLDAGFSIRSAVDIFHFLCSLLNVVEVVDAEGLNAQTADDDVQIFTLVLINSALEQSGDSIGTHP
KLLKMVQNDLCHHLIHYGKCSNPLVLSMICSTFLNIYYFLHRSVRLQLEAFFSFVLFRIANLGHYFHLQEVALEG
IINFCRQPTFIIEAYVNYDCDPIFRNIFEDVGKILCKLAFPTGGPLNSLQIQALEGLVVIIHNIADNVDKENDPT
PSGPYPVEISEYRPFWEEKSKDEMDLETWVDFIRVRKVQKRRISIAVNHFNHEKKKGLEYLKISKLISDDPDPKG
YAVFFRYTIGLDKTMIGDYLGDPDEFHLRVLKEFTDTFDFSGMILDTALRTFLETYRLPGESQKIQRILEAFSEK
FYEQQSSDIFANKDVVLILCYSLIMLNTDQHNHQVKKKMTEEEFIRNNRGINDTKDLPREYLSELFHSISSNGIV
SFGQTGMIIEMNPNRWLELMNRSKLLNPFVLCDFDRRLGRDMFSTIAGPSVSTLSSIFEQTDEEELLHECIEGLF
SIARIAQYGLEDILDEILTSFCKFTTLLNPYASAEETLYAFSHDMKPKLATLAVFTIANVFGESIRGGWRNIVGC
LLKLRRLKLLPQSMLELDISSYSTDSPKTESLFPGQKTGHSRMSSGLIARFSQFFSMETVEENLNLGLSEFEQNM
KIIQQCQIGRVFSNSSKMSNESLINLGRSLIFCAAGKGQKFSTPVEEEETVEFCWDLLSIISVSNTHRFMSFWPQ
YHDFLLGVVQFPLFSPVPFTEKGITAIVKVCLKLLSAPNPDKRPEELIFKSINLMWKLDKEIVETCGSFLAESIS
KIVTEYPGNLHSSLGWKSVLHLLSLTGRHPDTYELGVKTLITLMSDENSISRTNFAYCTDCAFNFIALKNTPLET
NMRIMDLMSDSVNYLIHWSKSGYSDPGSSSSVEDMRSSSSSHYTVDLFIRLGESLRKTSLARREEVRNHAIVSLR
KSFSLAADLNFPPANCINCFNLVVFAMVDDLHEKMLEYSRRENAEREMRSMEGTLKMAMELLTEVFLQFLKTISE

SSGFRTFWLGILRRMDTFMKAELGEYGQSRIQEIVPNLLSKMIATMKKNQILVPREGDDLWEITYIQIQWIAPGL
KDELFPDEL*
**>Prive_BG8809307**
MGRLKQSSGIKSIEEEPEDVETTFSNNSALACMINSEITAVLAVMRRNVRWGGRYMSGDDQLEHSLIQSLKALRK
QIYSWKHPWHTINPSVYLQPFLDVIRSDETGAPITGVALSSVYKILTLDVLDLNTVNVKEAMLLVVDAVTSCRFE
VTDPASEEVVLMKILQVLLACVKSKTSIVLSNQQVCTIANTCFRVVHQASTKGELLQRIARQTMYEIVRRIFSHL
SDVHKSGNFQVKGAGPYKQEGPIDSSNSFSSQASENSNGSSEYDGQSFTVNPLSTGMTDEHTFQSSSGKAAVPYD
PHVMNESFGVPCMVEIFQFLCLLLNSVEHVELGSGSNTVAFDEDMPLFALGLINSAIELGGSSIREHPRLLSLVQ
GDLFHNLMHFGLSSSPLILSMVCSIVLNLYHHLRSELKLQLEAFFSCVILRLAQSRYGASYLQQEVAMEALVDFC
RQTNFMVEMYVNLDCDITCCNVFEDLANLLSRSAFPVNCPLSSMHILALDGLISVIQGMAERITNGLVNSDLGPV
SLDEYIPFWMVKCDNYSNPDQWVPFVRRRKYIKRRLMIGADHFNRDPKKGLEFLQGTHLLPDKLDPQSVACFFRY
TAGLDKNLVGDFLGNHDEFCVQVLNEFAGTFDFEGMNLDIALRLFLETFRLPGESQKIQRVLEAFSERYYEQSPE
TLADKDAALVLSYSLIMLNTDQHNAQVKKKMTEEDFIRNNRRINGGSDLPREFLSELYYSICKNEIRTTPEVAMG
FPEMTPSRWVDLMNKSRKTPQFIVSDSNAYVDHDMFAIMSGPAIAAISVVFDHAEHEEVYQTCIDGFLAIAKISS
CHRLGDVLDDLVVSLCKFTTLLNPSFMEVPVLAFGDDIKARMATITVFSIANTYGDFIRTGWRNILDCILKLHKL
GLLPARVASDAADDLELSSEPGQGKPLANSLSSSNLSSIGTPRRSSGLMGRFSQLLSLDTEEPRSQPTDQQLAAH
QRTLQTIQKCHIDSIFTESKFLQADSLLQLARALIWAAGRPQKGGSSPEDEDTAVFCLELLIAITLNNRDRIGIL
WQGVYDHITGIVQSTVMPCALVEKAVFGLLRICQRLLPYKENLADELLRSLQLVLKLDARVADAYCEQITQEVSR
LVKANASHIRSPIGWRTITSLLSITARHPEASEAGFDALFFIMSEGVHLSPANFVLCIDSARQFSESRVGQVDRS
VKALDLMASSVLCLTRWAREAKVSLGESEASKMFQNIGDMWLRLVQALRKVCLDQREEVRNHSLFSLQMCLTCWD
EIHLPSGLWLQCFDLVIFTVLDDFVEIADGQSQKDYRNIEGTLILAIKLLSKVFLQLINELSQLSTFCKLWLGVL
GRMEKYMKVKVRGKKSEKLQELVPELLKNTLMVMKTKGVLVQRSTLGGDSLWELTWLHVKNISPDLQNEVFSDQG
MEQTLPSSETVPSTNETSTANML*
**>Prive_BG8816056**
METENNSDMMTKQKRKELGFSCMLNTEIGAVLAVIRRAPDLQNPYFIPNQEENCDSTILQSLKSLRSLVFNPQQE
WRTIDPSMYLSPFLDVIQSDDIPAPATGVAMSSILKILKLDMFDEKTPGAKEAINAVVTAITCCRLERTAPVAEE
AIMMRILLALIAVMKHRASVLLTDQAVCTIVNTSFQVVQQSSTRGDLLQRSARYTMHDLIQTIFSYLPDIEVKDW
DNSELDTEENNSDSGFGIRSAVEIFRFLCSLLNVVEIGDAEGLNAQTADEDVQIFALGLINSALEQSGDSIGTHP
KLLKMVQDDLCHHLIHYGTCSSPLVLSMICSTFLNIYHFLRRSVRLQLEAFFSFVLFRVVNLGHYLHLQEVAVEG
IINFCRQPTFIIDAYVNYDCDPIFRNVFEDCGKILCKLAFPTGGPLTSLQIQAFEGLVGIIHNIADTIDNENDST
PSGPYPVEISEYRPFWEEKSKREMDLETWVDYVRMRKIQKKKISIAANHFNRDEKKGLEYLKISKLISDEPDPIA
YAVFFRYTVGLDKTMIGDYLGDPDEFNLRVLKEFTNTFDFSGMILDIALRTFLETFRLPGESQKITRVLEAFAER
YYENQSSDIFANKDSVFILCYSVIMLNTDQHNNQVKKKMTEEEFLRNNRGINDKKDLPREYLSELFHSISSNAIT
LFGQTGMIIEMNPTRWVELMNRSKIMNPFVLSNFDRRIGRDMFSTIAGPSVSTLSSIFEQTDEEELLHECIEGLF
SIAQIAQYGLEDILDEILASFCKFTTLLNPYASADETLYAFSHDMKPRLATLAVFTIANILGESIRGGWRNIVDC
LLKLKRLKLLPQSLIEPDNSSDSTAENELLFPGQKTGNNKISYGLVGRFSNFLSMETVEENLNLGLSEFEQNLKM
IQHCHIGQVFSNSSKLPEESLLNLGRSLIFSAAGKGQKFSTPIEEEETVTFCWDLILIISLSNIHRFMSFWPQYH
DFLLGVAQFPLFSPVPFAEKALAVLFKACLKLVSTYNPDKYPEELIFKSINLMWKLDKEAVETCGMFLTQSIKKI
ITEYSGNLHTSLGWKSCLHLLSLIGRHPDTYELGVETLIVLMSDGHSISRVNYAYCIDCAFSFVALKNTPLETNM
RIMDLMSDSVTHLIQWYKSGYSDPGSGVSISSLEDTRTNSSSNFTVTLFIKLGESLRKTSLARREEIRNHAIVSL
QKSFNLAEELYFTPTNCINCFNLVLFAMVDDLHEKMLEYSRRENAEREMRSMEGTLKIAMELLTEVYLQFLRTIS
ESSGFRTFWLGILRRMDTCMKAELGEYGTSSVQEIVPNLLKKMITTMKNYEILVQREGEDLWEITYIQIQWIAPS
LKEELFPDEF*
**>Cac_g011358**
MGRLKLQSGIKAIEEEPEDCEMPSSNKAALACMINSEIGAVLAVMRRNVRWGGRFLSGDDQLEHSLVQSLKALRK
QIFSWQHQWHTVNPSVYLQPFLDVIQSDETGAPITGVALSSVYKILTLDALDLNTVNVEDAMHLVVDAVTSCRFE
VTDPASEEVVLMKILQVLLACMKSKASVMLSNQHVCTIVNTCYRIVHQAGTKGELLQRIARHTVHELVRCIFSHL
PDVDITERSLVKGGSYVKQEVGGIDNDYSFGGKQLDNGNGSSEYDSQLPSGSLESNVPGGLVTGVINENAIGTNN
WKDATQYDMHLMTEPYGVPCMVEIFHFLCSLLNVVENMGMAPRSNTIAFDEDVPLFALGLINSAIELGGHSIRRH
PRLLSLVQDELFRCLMQFGLSMSPLILSMVCSIVLNLYHHLRTELKLQLEAFFSCVILRLAQSRHGASYQQQEVA
MEALVDFCRQKTFMVEMYANLDCDITCSNVFEDLANLLSKSAFPVNCPLSAMHILALDGLIAVIQGMAERIGNGS
VGSELSPVNLEEYTPFWMVKCDNYSDPDHWVPFVRRRKYIKRRLMIGADHFNRDPKKGLEFLQGTHLLPDKLDPQ
SVACFFRYTAGLDKNLVGDFLGNHDEFCVQVLHEFAGTFDFQDMNLDTALRLFLETFRLPGESQKIQRVLEAFSE
RYYEQSPQILANKDAALLLSYSLIMLNTDQHNVQVKKKMTEEDFIRNNRHINGGNDLPREFLSELYHSICKNEIR
TTPEQGAGFPEMTPSRWIDLMHKSKKTAPFIVSDSRAYLDRDMFAIMSGPTIAAISVVFDHAEHEDVYQTCIDGF
LAVAKISACHHLEDVLDDLVVSLCKFTTLLNPSSAEEPVLAFGDDTKARMATVTVFTIANRYGDFIRTGWRNILD
CILKLHKLGLLPARVASDAADDSELSSDTGHGKPLTNSLSSAHMPSMGTPRRSSGLMGRFSQLLSLDTEEPRSQP
TEQQLAAHQRTLQTIQKCHVDSIFTESKFLQADSLLQLARALTVAAGRPQKGNSSPEDEDTAVFCLELLIAITLN
NRDRIVLLWQCVYEHISSIVQSTMMPGALVEKAVFGLLRICQRLLPYKENLADELLRSLNLVQKLDARVADAYSE
QITQEVNRLVKANATHIRSPMGWRIITSLLSMTARNPEASEAGFDALLFIMSDGAHLLRANYVPCIEAARQFAES
RVGQADRSVRVLDLMAESVSCLARWAHETKESLGEAEAVKIYQDIREMWLRLVQELRKVCLDQREEVRNHALVSL
QICLTGVDGVHFPHGLWLQCFDLVIFTMLDDLLEIAQGHSQKDYRNMEGTLILALKLLSKVFLQLLHELSQLTTF

CKLWLGVLSRMEKYMKVKVRGKKSEKLQELVPELLKNTLLVMKTRGVLVQRSALGGDSLWELTWLHVNNISPSLQ
SQVFPDQESEPSQPNKGEMGGSLVANETVSAPNETVTAGG*
**>Cac_g006307**
MGCLNQQSGVNAFVGEPKDYFMMPSRGALACMVNSEIGAVLAVMRRNVRWGVRYSTDDDQLEPSLIQSFKELRKK
IFSWQHQWHAINPVVYLQPFLDVIKSDETGAPITGVALSSIYKILTLEILDINTVNVADALHLIVDAVTSCRFEV
TDPASEEAVLMKILQVLLACMKNKASAKLSNQHVCNIVNTCFRIVHQASSKGELLQRIARHTMHELIRCIFSHLP
DINDTGDALAQGGRLCVNKEVGALDKDPASGTKKEENGYVSSEYDGSLPSAGVASTTVSTGTVAGKMDENIIEVG
QGMDGAQNGGHAMMEPYGVPCMVEIFHFLCSLLNVTEQIEIGPRSNPIAYDEDVPLFALGLINSAIELGGSSIGH
HPKLLALIQDELSYNLMQFGLSMSPLILSTVCSIVLNLYHHLRAKLKLQLEAFFSCVLLKIAQSKHGALYQQQEV
AMETLVDLCRQQMFMTEMYANFDCDISCSNVFEDLANLLSKSAFPVNSSPLSAMHILALDGLIAMIQGMAERVGN
ESPASEQASLDLEEYKPFWTVKCENYGDPSHWVPFVHRRKYIKKRLMNGADHFNRDPKKGLEFLQAMRLLPDKLD
PKSVACFIRYTTGLDKNLIGDFLGNHDEFCVQVLYEFAETFDFQGMNLDTALRVFLETFRLPGESQKIQRVLEAF
AGRYHEQSSNILVNKDAALLLSYSLILLNTDQHNVQVKKKMTEEDFIRNNRHINGGNDLPREYVSELYHSICKNE
IRMTPEQGTGFPLMTPSHWIGLMHKSKQATPFIICDSKGQLDYDMFVILSAPTIAAISVVFDHVEQEDILQTCMD
GFLAIAKISACYHLDDVLNNLVVSLCKFTTLLLPLSVEESVLAFADDTKARMATITVFTIANRYGDHIRSGWKNI
LDCILSLHKLGLLPSRLASDAADDFESPSELERGRPATSSTPASHIPSVTIPRKSSGLMGRFSQFLYYDAEEPTP
QPTEEQLAAHQHALQTIQSCHIDSIFTESKFLQSESLLQLAQALILAAGQPNKGNNFPEGGDTGVFCLELLIAIT
LNNRDRIMLLWQGVYEHISNIVQSTAMPCAVVEKAVFGLLRICQRLLPYKENLTDELLKSLQLVLKLDARVADAY
CEHITQEVMRLVKANAMQIRSQMGWRTITSLLSITARHPEASEAGFDTLVFIMSDGAHLLPANYVLCVNAARQFA
ESRVGHVDRSIRSLDLMSGSVVCLVNWSSKTKEAVGEEAAAKLYQDIGEMWLRLVQGLRKVCLDQREEVRNHAVL
LLQRCLTGVNGIHLPNELWLQCFDLVVFTLLDDLVEIAQESSPKDYRNMEGTLALSVKLLSKVFLLLVQDLSQLT
SFCKLWLGILNRMERFMKLKFRGKRSEKIHELVPELLKNTLLVMKTRGILVQSDAVGGDSFWQLTWLHVKNISPS
LQLEVFPGNEMDQLQDKHIITGNSPVPDGNLLSPSSETTA*
**>Cac_g011770**
METDDDVENNKKNMSRLKRKELGVSCMLNTEVGAVLALIRRPSDSNTQFVSPPDDNYDSSISQSLKSLRALIFNP
QQEWRTIDPSIYLSPFLDVIQSDDVPATATGVALSSILKILKLQIFDEKTTGARDAINSTMSAITSCRLEKTDSV
SEDAVMMRILQVLTAIMKHRASVLLTDHAVCTIVNTCFQVVQQSASRGDLLQRSARYTMHDLIQIIFSRLPDIEV
KDWENSESDTEDGNLDSGYGIRSAVDIFHFLCSLLNVMEIPETEMLTPQSADEDVQLFALVLINCAIELSGDSIG
RHAKLLRMIQDDLFHHLIHYGTCSSPLVLSMICSTVLNIYHFLRRSIRLQMEAFFSFVLFRAASLANSIQLQEVA
VEGIINFCRQPTFALEVYVNYDSDPICCNVFEEIGKLLCKHAFPMGGHLTSLQIQAFEGLVLIIQGIADNIDKED
IPGPFGPYPVEISEYTPFWEEKSKDNDDSEAWVDFVRIRKAQKRKLLIASNHFNRDEKKGSEYLKISELVSDPPD
PKAFAMFFRYTPRLDKTMIGDYLGDPDEFHIQVLKEFTDTFEFSGMILDTALRTFLETFRLPGESQKIQRILEAF
SEKFYDQQSSEIFVSKDSVFILCYSLIMLNTDQHNPQVKKKMTEEEFIRNNRAINGGKDLPREYLSELFQSISTN
AITLFGQSGLAIEMNPGRWIELINRAKHMKPFISCDFDCRLGRDMFATIAGPSVAALSAIFEQADDEEILHECIE
GLFSVARIAHYGLEDTLDELLSSFCKFTTLLNPYASAEETLYAFGNDMKPRMATLAVFTIANSFGNSIRGGWRGI
VECLLKLKRLKLLPQSVVDHENASALSPDLQTHSRSESGVIFSTHDPKFSNGRHISGLIGRVSHFLSMESVEEAL
NLGVSEFEQNLKIIQQCRIGNIFSSSSSLPDDALLNLGRSLISAAAGKGQKFSTPVEEEETVGFCWDLILTISVA
NIQRFSTFWPHYHDYLLGVAQFPLFSPVPFAEKAIFTLIKVCLRLLASYRSDKFPEELIFKSINLMWKLDKEILD
TCCEFITQSVNRILIEYPANLQTQLGWKTVLHLLSITGRHPETYDQAIETLIMLMSDGTHVSRINYAYCIDCAFG
FVALKNSPLDKNLKIMDLMSDSVNLLIQWYKNEYSDPGSNASIVSNTSSSSIEDNSKALGSANLTMNLFAKLGES
LRKTSLARREEIRNHAIFSLQKSFTLAEELYFTPSNCINCFNLVIFAMVDDLHEKMLEYSRRESAEREMRSMEGT
LKIAMELLTEVYLQFLRPISESPGFRTFWLGILRRMDTCMKADLGEYGESKLQEMVPDLLRKMISTMKEKEILVQ
SEGDDLWEITYIQIQWIAPSLKEELFPDEL*
**>Nyssi_012948**
MGRLNLQSGIKAIEEEPEDCETPSSNKTALACMINSEIGAVLAVMRRNVRWGGRYMSGDDQLEHSLVQSLKALRK
QIFSWQHQWHTVNPSVYLQPFLDVIQSDETGAPITGVALSSVYKILTLDALDLNTVNVEDAMHLVVDAVTSCRFE
VTDPASEEVVLMKILQVLLACMKSKASVMLSNQHVCTIVNTCYRIVHQAGTKGELLQRIARHTMHELVRCIFSHL
PDVDITEQSLVKGGSYVKQEVGGIDNDYSFGGKQFDNGNGSSEYDSQLSSGSLESTVPSGPVTGVINENAIGTGN
WKDATQYDLHLMTEPYGVPCMVEIFHFLCSLLNVVEHMGMGPRTNTIAFDEDVPLFALGLINSAIELGGHSICRH
PRLLSLVQDELFRCLMQFGLSMSPLILSIVCSIVLNLYHHLRTELKLQLEAFFSCVILRLAQSRYGASFQQQEVA
MEALVDFCRQKTFMVEMYANLDCDITCSNVFEELANLLSKSAFPVNCPLSAMHILALDGLIAVIQGMAERIGNGS
VSSELSPVHLEEYTPFWMVKCDNYSDPDHWVPFVRRRKYIKRRLMIGADHFNRDPKKGLEFLQGTHLLPDKLDPQ
SVACFFRYTAGLDKNLVGDFLGNHDEFCVQVLHEFAGTFDFQDMNLDTALRLFLETFRLPGESQKIQRVLEAFSE
RYYEQSPQILANKDAALLLSYSLIMLNTDQHNVQVKKKMTEEDFIRNNRHINGGNDLPREFLSELYHSICKNEIR
TTPEQGASFPEMTPSRWIDLMHKSKKTAPFIASDSRAYLDRDMFAIMSGPTIAAISVVFDHAEHEDVYQSCIDGF
LAVAKISACHHLEDVLDDLVVSLCKFTTLLNPSSVEEPVLAFGDDTKARMATVTVFTIANRYGDFIRTGWRNILD
CILKLHKLGLLPARVASDAADDSELSSDTGHGKPLTNSLSSAHMPSMGTPRRSSGLMGRFSQLLSLDTEEPRSQP
TEQQLAAHQRTLQTIQKCHIDSIFTESKFLQPDSLLQLARALTGAAGRPPKGNSSPEDEDTAVFCLELLIAITLN
NRDRIVLLWQCVYEHISSIVQSTMMPGALVEKAVFGLLRICQRLLPYKENLADELLRSLNLVQKLDSRVADAYSE
QITQEVNRLVKANATHIRSPMGWRTITSLLSMTARNPEASEAGFDALLFIMSDGAHLLRANYVLCIDAARQFAES
RVGQADRSVRVLDLMADSVACLARWAQETKESLGDQEAVKMHQDIREMWLRLVQELRKVCLDQREEVRNHALLSL
QMCLTGVDGVHLPHGLWLQCFDLVIFTMLDDLLEIAQGHSQKDYRNMEGTLILALKLLSKVFLQLLHELSQLTTF

CKLWLGVLSRMEKYMKVKVRGKKSEKLQELVPELLKNTLLVMKTRGVLVQRSALGGDSLWELTWLHVNNISPSLQ
SQVFPDQESELLQPNQGS*
**>Nyssi_012781**
MGRLKLQSGIKAIEEEPEDCETPSSNKAALACMINSEIGSVMAVMRRNVRWGGRYMSGDDQLEHSLIQSFKALQK
QIFSWQHQWHTIDPAVYLQPFLDVIRSDETGAPITGVALTSVYKILTLDVLDLNTVNVEEAMHLVVDAVTSCRFE
VTDTASEEVVLMKVLQVLLACMKSEASVMLNNQHVCTLVNTCSRIVHQAGTKGELLQRIARHTMHELIRCIFSHL
LDVDITDRSLVKGGTSVKHEVSGFGNDYSFGVKQLDNRNGSSEFDHQPSSGSLVSSDPSGLVTGVMDGNTIGTGN
GKDTTPYDFHLMTEPYGVPCMVEIFHFLCSLLNVVEHMGMDPRSNTIAFDEDVPLFALGLISSAIELGGASICRH
PRLLSLVQDELFCRLMHFGLSMSPLILSMSRYGASYQQQEVTMEALVDFCRQKTFMVEMYANLDCDITCSNVFED
LANLLSKSAFPVNCPLSSMHILALDGLFAVIQGMAERIGNGSVSSEPSPVNLEEYTPFWNVKCDDYGDPDHWVSF
VRRRKYIKRRLMIGADHFNRDPKKGLEFLQGTHLLPDKLDPQSVACFFRYTAGLDKNLVGDFLGNHDEFCIQVLH
EFAGTFDFQDMNLDTAMRLFLETFRLPGESQKIQRVLEAFSERYYEQSPQILANKDAALLLSYSLIMLNTDQHNV
QVKKKMTEEDFIRNNRHINGGGDLPREFLSELYHSICKNEIRTTPEQGAGFPEMTPSHWIDLMHKSKRTAPFIVS
DSRAYLDHDMFAIMSGPTIAAISVVFDHAEYEDVYQTCINGFLAVAKISACHHLEDVLDDLVVSLCKFTTLLNPS
SVEEPVLAFGDDTKARMATITVFTIANRYGDFIRTGWRNILDCILKLHKLGLLPACVASDAADDSELSSDTGHGK
PLTNSLSSAHMPSLGTPRRSSGLMGRFSQLLSLDTEEPRSQPTEQQLAAHQRTLQTIQKCHIDSIFTESKFLQAD
SLLQLAQALIWAAGRPQKGNSSEDEDTAVFCLELLIAITLNNRDRIVLLWQGVYEYISKIVQSTVMPCALVEKAV
FGLLRICQRLLPYKENLVDELLRSLQLVLKLDARVFDAYCEQITQEVSHLVKANATHIRSQIGWRTITSLLSNTA
RQPEASEVGFDALLFIMSDGAHLLLANYDLCVDAARQFAESRVGQADRSVRALDLMAGSVTCLARWAQETKKSLG
EVGAVKMYQDIGEMWLRLVQGLRKVCLDQREEVRNHALLSLQMCLTGVDGIHLPHGLWLQCFDLVIFTMLDDLLE
IAQGHFQKDYRNMEGTLILALKLLQKVFLQLLRELSQLTTFCKLWLGVLSRMEKYMKVKVRGKRSEKLQELVPDL
LKNTLLVMKTRGVLVQRSALGGDSLWELTWLHMNNIAPSLQSEVFPVQDSEQSQLNQGEMGGSLVSNETGSAHNE
TVTPEAFRTGG*
**>Nyssi_000634**
MGCLNRQSGVNSFVGEPKDSFMISSRGALACMVNSEIGAVLAVMRRNVRWGVRYSTDDDQLEPSLIQTFKELRKK
IFSWQHKWHTISPVVYLQPFLDVIKSDETGAPITGVALSSIYKILTLEILDINTVNVADALHLIVDAVTSCRFEV
TDPASEEVVLMKILQVLLACMKNKASVKLSNQHVCNIVNTCFRIVHQASSKGELLQRIARHTMHELIRCIFSHLP
EIDNTGYVLAQGGRLYVNNEVGVPDADPASGSKQQENGHVSSEYDGQMPSVGVASTVSTGTLVGKMDENIIEVGQ
AMDAAQNGGHAMMEPYGLPCMVEIFHFLCSLLNVAEQTEIGPRSNPIAYDEDVPLFALGLINSAIELGGSSIGHH
PKLLALIQDELSYNLMQFGLSMSPLILSTVCSIVLNLYHHLRAKLKLQLEAFFSCVLLRIAQSKHGALYQQQEVA
METLVDLCRQQMFMTEMYANFDCDISCSNVFEDLANLLSKSAFPVNSPLSAMHILALDGLIAMVQGMAERIGNES
PASEQASLDLEEYKPFWTVKCENYGDPSHWVPFVHKMKYIKKRLMNGSDHFNRDPKKGLEFLQGMRLLPDKLDPR
SVACFIRYTTGLDKNLIGDFLGNHDEFCVQVLHEFAGTFDFQDMTLDTALRVFLETFRLPGESQKIQRVLEAFAG
RYHEQSPNILANKDAALLLSYSLILLNTDQHNVQVKKKMTEEDFIRNNRQINGGNDLPREYLSELYHSIVKNEIR
MTPEQGAGFPLMTPSHWIGLMHKSKQATPFIICDSNVQLDYDMFVILSGPTIAAMSVVFDHVEQEDILQTCIDGF
LAIAKISACYHLNDVLDDLVVSLCKFTTLLLPLSVEESVLAFADDTKARMATITVFTIANRYGDHIRSGWKNILD
CILSLHKLGLLPSRLASDAADDFEPPSELERGRPATSSTPASHTPTMTIPRKSSGLMGRFSQFLYYDAEEPTPQP
TAEELAAHQHALQTIQNCHIDSIFTESKFLQSESLLQLAQALILAAGQPNKGNNFPEGGDTMVFCLELLIAITLN
NRDRIMLLWQGVYEHISNIVQSTAMPCALVEKAVFGLLRICQRLLPYKENLTDELLKSLQLVLKLDARVADAYCE
HITQEVMRLVKANAMQIRSQIGWRTITSLLSITARHPEASEAGFETLVFIMSDGAHLLPANYVLCVNAARQFAES
RVGHVHRSIRSLDLMAGSVVCLVNWSCKTKEAVGEEAAAKMYQDIGEMWLRLVQGLRKVCLDQREEVRNHAVLLL
QRCLTGVERIRLPNELWLQCFDLVIFTLLDDLIKIAQESSPKDYRNMEGTLALSVKLLSKVFLQLVQDLSQLTSF
CKLWLGVLNRMERLMKVKFRGKRSEKIHELVPELLKNTLLVMKTRGILVQRDAVGGDSFWQLTWLHVNNISPSLQ
LEVFPGNELEQLQDKHIITGGSPVPDGNLLSPSSETTA*
**>NyssiGNL2**
MTRLERKELGISCMLNTEVGAVLALIRRPTDPNSQFIAPPDENYDSFISQSLKSLRAFIFNPQQEWRTIDPSIYL
SPFLDVIQSDDLPAAATGVALSSILKILKLQIFDEKTTGARDAINSAMSAITSCRLEKTDSVSEDAVMMRILQVL
TAIMKHQASVLLTDHAVCTIVNTCFQVVQQSASRGDLLQRSARYTMHDLIQIIFSRLPDIEVNDCENSESDTEDS
NLDSGYGIHSAVDIFHFLCSLLNVMEVPAAEVLTPQSADEDVQLFALVLINSAIELSGDSIGKHPRLLRMIQDDL
FHHLVHYGTCSSPLVLSMICSTVLNIYHFLRRSIRLQLEAFCSFVLLRAASLGNSNQLQEVAVEGIINFCRQPAF
AIEVYVNYDSDPICHNVFEEIGKLLCKHAFPAGGHLTSLQVQAFEGLVVIIQSIADNIYKEDISGPFGSYPVEIP
EYRPFWEENSKDNDDSEAWVDFVRVRKAQKRKLSIAGNHFNRDEKKGSEYLKISQLVSDPPDPKALAMFLHYTPK
LDKTMIGDYLGDPDEFHIQVLKEFTDSFEFSGMILDTALRTFLETFRLPGESQKIQRILEAFSEKFYDQQSSEIF
VSKDSVFILCYSLIMLNTDQHNPQVKKKMTEEEFIRNNRAINGGKDLPREYLSELFHSISTHAITLFGQSGLPIE
MNPNRWIELINRAKIMKPFILCDFDRRLGRDMFATIAGPSVAALSAIFEHADEEEILHECIEGLFSVARIAHYGL
EDTLDELLASFCKFTTLLNPYASAEETLHAFGNDMKPRMATLAVFTIANSFGNSIRGGWRGIVDCMLKLKRLKLL
PQSVVDHENTSASSPDLQTHSRSESGVIFSAHDPKLGNGRHIFGLVGRFLNFQSMESVEEALNLGVSEFEQNLKI
IQQCRIGNIFSNSSNLPDDAVLNLGRSLIFAAAGKGQKFSTPVEEEETVGFCWDLILTISVANIHRFSTFWPHFH
EYLLGVTQFPLFSPVPFAEKAILALIKVCLKLLASYRSDKVPEELIFKSINLMWKLDKEILDTCCEFITQSVNRI
LIEYPANLQTQLGWKSVLHLLSVTGRHPETYDQGVETLIMLMSDGTHVSRTNYAYCIDCAFSFVALKNSPLEKNL
KIMDMMSDSVNLLIHWYKNEYSDPGSNASIVSNTSSSSSIEDNSKALGSANFTMNLFVKLGESLRKTSLARREEI
RNHAIFSLHKSFTLAEELYFTPSNCINCFNLVIFAMADDLHEKMLEYSRRENAEREMRSMEGTLKIAMELLTEVY

LQFLRPISESPSFRTFWLGILRRMDTCMRPDLGEYGESKLQEIVPDSLTKMISTMKEKEILVQNEGEDLWEITYI
QIQWIAPSLKEELFPDELLEGI*
**>Hyque_10G111800**
MGRLKLQSGIKSIEEEPEDCEVSSSSHKAALACMINSEIGAVLAVMRRNVRWGGRYMSGDDQLEHSLIQSLKALR
KQIFTWQHQWHTVNPAVYLQPFLDVIRSDETGAPITGVALSSVYKILTLDVLDLDTVNVEDAMQLVVDAVTSCRF
EVTDPASEEVVLMKILQVLLACMKSKASAMLSNQHVCTIVNTCYRVVHQAGTKGELLQRIARHTMHELVRCIFSH
LPNVDNTEHTLVKGVNVVKQEVGHIDNDYSFGDKQLENGNGSSEYDSQSHSGSLVSNAPSGLVTGVMDEYAIGSG
NGKDATPYDLQLMTEPYGVPCMVEIFHFLCSLMNVGEHMGMGPRSNTIAFDEDVPLFALGLINSAVELGGPSISQ
HPRLLSIVQDELFRSLMQFGLSTSPLILSMVCSIVLNLYQHLRTELKLQLEAFFCCVILRLAQSRYGASYQQQEV
AMEALVDFCRQKTFMVEMYANLDCDITCSNVFEDLANLLSKSAFPVNCPLSAMHILALDGLIAVIQGMAERIGNG
SVSSELSPVNLEEYIPFWMVKCDNYSDPDHWVPFVRRRKFIKRRLMIGADHFNRDPKKGLEFLQGTHLLPDKLDP
QSVACFFRYTAGLDKNLVGDFLGNHDEFCVQVLHEFAWTFDFQDMNLDTALRLFLETFRLPGESQKIQRVLEAFS
ERYYEQSPLILVNKDAALLLSYSIIMLNTDQHNVQVKKKMTEEDFIRNNRHINGGNDLPREFLAELYNSICKNEI
RTTPEQGASFPEMNPSRWIDLMHKSKKTAPFIISDSRAYLDHDMFAIMSGPTIAAISVVFDHAEHEDVYQTCIDG
FLAVAKISACHHLEDVLDDLVVSLCKFTTLLNPSSVEEPVLAFGDDTKARMATVTVFTIANRYGDFIRTGWRNIL
DCILKLHKLGLLPARVASDAADDSELSSDPGHGKPLTNSLSSAHMQSMGTPRRSSGLMGRFSQLLSLDTEEPRSQ
PTEQQLAAHQRTLQTIQKCHIDSIFTESKFLQSDSLLQLARALIWAAGRPQKGTSSPEDEDTSVFCLELLIAITL
NNRDRIVLLWQGVYEHISSIVQSTVMPCALVEKAVFGLLRICQRLLPYKENLADELLRSLQLVLKLDARVADAYC
DQITQEVTRLVKANAAHIRSPMGWRTITSLLSHTARHPEASEAGFDALLFIMSDGSHLLPANYILCIDAARQFAE
SRVGQADRSVRAVDLMAGSVACLARWTQETKDSLGEVEGVKMSQEVGDMWLRLVQGLRKVCLDQREEVRNHALLS
LQMCLTGVDGVHLPHILWLQCFDQVIFTMLDDLLEIAQGHSQKDYRNMEGTLVFALKLLSKVFLQLLHELSQLTT
FCKLWLGVLSRMEKYMKVKVRGKRSEKLQELVPELLKNTLLVMKTRGVLVQRSALGGDSLWELTWLHVNNIAPSL
QSEVFPDQNSEQSEPDGLESSESGSSPSEPVAP*
**>Hyque_13G093400**
MGCLNQQSGELKDCDTRPSRGALACMVNSEIGGVLAVMRRNVRWGVRYVIDDDQLEHPLIQSFKELRKKIFSWQQ
QWHTINPVVYLKPFLDVIKSDETGAPITGVALSSVYKILSLEIIDLNTVNVAEALHLIVDAVTSCRFEVTDPASE
EVVLMKILQVLLACMKNKASVKLSNQHVCNIVNTCFRVVHQASSKGELLQRIARHTMHELVRCIFLHLPDIDSTE
HESAQAGRSYISKEGGATDNGFVSSEIDVQPASFESTGPIGTLAGNGSVVEVGQGKEADQNGAHPMMEPYGVPCM
VEIFHFLCSLLNMMEHMEIGPRSNPIAYDEDVPLFALGLINSAIELGGDSFGHHPKLLTLVQDELFRNLMRFGLS
MSPLILSTVCSIVLNLYHHLRTKLKLQLEAFFSSVLLRIAQSKQGASYQQQEVAMETLVDLCRQQMFMTEMYANF
DCDISCSNVFEDLANLLSKSAFPVNNPLSSMHILALDGLIAMIQGMGERITNEPPASEQSSSHLEDYKPFWTVKC
ENYGDPNDWVPFVHSMKYIKRRLMIGADHFNGDPKKGFEFLQGVHLLPDKLEPQGVACFFRYTTGLDKNLIGDFL
GNHDEFCVRVLHEFAGTFDFREMNLDTALRVFLETFRLPGESQKIQRVLEAFAGRYYEQSPHILANRDAALLLSY
SLILLNTDHHNAQVKKKMTEEDFIRNNRRINGGNDLPREYLSELYHSICENEIRMTPDQGAGSPVVTPSNWIGLM
HKSKETAPFIVCDSRARLDYDMFAILSGPTIAAISVVFDHVDQENVLQTCIDGFFAIAKISASYHLEDVLDDLVV
SLCKFTTLLLPLSVEESILAFTEDTKARMATIAVFTIANRYGDYIRSGWKNILDCILSLHKLGLVPSRLASDAAD
DLESSSDPDRGRPATTSTPPPMATPRKSFGLMGRFSQLLYLDTEEPAPQPSEEQLAAHQHTLQTIQNCHIDSIFA
ESKFLQAESLSQLARALILAGGRPNKGNNSMQDEDTRVFCLELLIAISLNNRDRIMLLWEGVYEHISSIVQSTSM
PCALAEKAVFGLLRICQRLLPYKENLTDELLKSLQLVLKLDARVADAYCEHITQEVMRLVKANAMQIRSHMGWRT
ITSLLSITARHPEASEAGFETLVFIMSDGAHLLPANYVLCVNSARQFAESRVGDVDRSVRSLDLMAGSVLSLVKW
SCKTKEAAEEEAAAKMYQDIGEMWLRLVQGLRKVCLDQREEVRNHAILSLQRCLTGVDGISLAHNLWLQCFDLVI
FTLLDDLVEIAQENSPKDYRNMEGTLVLSMKLLSKVFLQLLQNLSGSASFCKLWLGVLNNMERFMKVKFRGRRNE
KIHELIPELLKNTLFVMKTREILVHSSDSVGGDSFWELTWLHVKNIAPSLQSEVFPPSELEKKPIKTEGSPVPDG
IVLVPSSETTA*
**>Hyque_18G092600**
MTRLNRKDQGVSCMLNTEVGAVLAVIRRSPEPTSQYISPVEDNYDTSLLQSLKSLRSIIFNPQQEWRNVDPSTYL
LPFLDVIQSDDVPANATGASLSSILKILKLEIFDETTHGAREAINSAVTAITGCRLEKTDPVSEDAVMMRILQVL
TAIMKHRSSDLLTDHAVCTIVNTCFQVVQQSANRGDLLQRSARHAMHEIIQIIFSRLPEIEVRDWEDSESDTEDT
NLDSGYGIRSAVDIFHFLCSLLNVVEVVETTEVLTAQSADEDVQLFALVLVNSAIELSGESIGKHRKLLRMIQDD
LFHHLVHYGTCSSPLVLSMICSTVLSIYHFLRRAVRLQLEAFFSFVTFRVASPGNATQLQEVAIEGLINFCRQPT
FILEAYVNYDCEPICRDVFEEIGKLLCKQAFPTGGHLTSSQVQAFEGLVVVIQNIADNIDKEDDSSNSGPYPVEI
TEFRPFWEEKSIDGNDLETWVDYVRVRKAQKRKILIAGNHFNRDEKKGLEYLKLSKLVSDPPVPKGFALFFRYTP
RLDKTMIGDYLGDPDEFHIQVLKEFTDTFEFSGMILDTALRTFLETFRLPGESQKIQRILEAFSEKFYDQQSSEI
FVSKDSVFILCYSLIMLNTDQHNPQVKKKMTEEEFIRNNRAINGGQDLPREYLSELFYSISTKAITLFGQSGSPV
EMNPSRWIELIHRAKTTNPYITCAFDRRLGRDMFATIAGPSVAAFSAIFEQADEEEILHECIEGLFSVARIAQYR
LEDTLDELLASFCKFTTLLNPYASAEETLYAFSNDMKPRMATLAVFTIANNFGESIRGGWRNIVDCMLKLKRLKL
LPQSIVEFENLSTSSSDLQAQSTSESGVIFPNQDPKFGSGQLSSGLIGRFSHFLSMESVEESLNLGISEFEQNQK
VIQQCRIGNIFKNSSNLPEDALLNLGRSLIFAAAGKGQKFSTPVEEEETVGFCWDMILAIALANIERFSSFWPQY
HEYLLGVAQFSLFSPIPFAEKAVLALMKVCLKIVSSYQTDKSPEELIFKSINLMWKFDKEILDTCSDSITQSVSK
IIFEYPANLQTQLGWKSILHLLSVTGRHPETYDRGVETLITLMSDGSHVSRTNYAYCIDCAFGFVALKNSPLEKN
LKIMDLMADSVNLLIQWYKSGYSDPGSTVSVMSSTSTSSLDESSKALGSANFTMNLFVKLGESLRKTSLARREEI
RNHAIFSLQKSFTLAEELYFTPTNCINCFNLVIFAMVDDLHEKMLEYSRRDSAEREMRSMEGTLKNAMELLTEVY

LQFLKPISESPGFRTFWLGIMRRMDTCMKADLGEYGESKLQEIVPELLSKMITTMKEKEILAQKEGDDLWEITYI
QIQWIAPSLKEELFPEEL*
**>Amacr03g15130**
MGRRLNLQSGIKAIKEEPEESEEVSSSSNKAALSCMINSEIGAVLAIMRRNVRWGGRFMSSGEDQFDQHSLIQSM
KSLRKQVFMWQQQQWCNINPVLYLQPFLDVIRSDETGAPITSVALSSVYKILTLDGVLDLNTTNIEDAMHLVVDS
VTSCRFEVTDPASEEVVLTKILQVLLACMKSKASVVLSNQHVCTVFNTCFRIVHQATTKGELLQTMAAHTMRELV
RCIFLHLPQVDVESMENSVNNGNSMQAEIGGIGNEYSFSSGQINNGNNSEYDDGQLPTANFGSNVSVGMGAGVVD
ENSTGNGKDAGAYDLHVTMAPYGVPSMFEIFQFLCSLLNVVDQGVISRSNGLAIDEDAPLFALGLINSAIELGGA
SFVRHPRLLNLIQDELFRNLMLYGLSPSPLILSMVCSVVLNLYHHLRMELKLQLEAFFSCVILRLAQSRYGASYQ
QQEVAMEALVDFCRQKTFMVEMYANLDCDITCSNVFEDLANLLSKSAFPVNSPLSSMHILALDGLIAVIQGMAER
IGNGSLGLEQAPVSLEEYSPFWMVKCDNYSDPNHWVPFVRRRKYIKRRLMIGADHFNRDPKKGLEFLQGTHLLPE
KLDPQSVACFFRYTAGLDKNLVGDFLGNHDEFCVQVLHEFAQTFDFQDMNLDTALRLFLETFRLPGEAQKIQRVL
EAFSERYYEQTSHILANKDAAFVLSYSLIMLNTDLHNVQVKKKMTEEDFIRNNRRINAGEDLPREFLCELYHSIC
KNEIRTTPEQGAGFAEMTPSRWIDLLHKCKKTPPYILADSRAYLDHDMFAIMSGPTIAAISVVFDNAEYEEVYRT
CIDGFLAIAKISACHHLEDVLDDLVVSLCKFTTLLNPSSVEEPVLVFGDDMKARMATVTVFTIANRYGDYIRTGW
RNILDCILRLHKLGLLPARVASDAADDSELSSDLSQGKPLTSSLSASHFPSMGTPRRSSGLMGRFSQLLSLDTEE
PRSQPTEQQLAAHQRTLQTIQKCHIDGIFTESKFLQAESLLQLAKALIWAAGRPQKGAGSPEDEDTAVFCLELLI
AITLNNRDRIVLLWQGVYEHIANIVQSTVMPCALVEKAVFGLLRICQRLLPYKENLADELLRSLKLVLKLDARVA
DAYCEQITQEVTRLVKANATHIRSQMGWHTITSLLSVTARHPEASESGFDALVFIMSDGAHMLPSNFILCVESAR
QFAESRVGQSDRGGQSDRSIRALDLMAGSVSCLVKWVSETKQAVEEDVVKMSQDIGEMWLRLVQALRKVCLDQRE
EVRNHAVSSLQKCLTEMDEVHLAHNLWLQCFDMVIFTMLDDLLEIAQGHSQKDYRNMEGTLTIAIKLLSKVFLQL
LPELSQLTTFCKLWLGVLSRMEKITKVKIRGKRSEKLQDLVLELLKNILVEMKNRGVLAQRSALGGDSLWELTWL
HVNNISPSLQSEVFPSAVADESTNSIHTEAVPLDS*
**>Amacr12g11020**
MSCLELQPGSLACMVNTEIGAVLAVMRRNVRWGNRFTSVDDQQEHSLVQSLKTLRKKMFTWHAQWHSVNPVMYLQ
PFLDVIQSEETGAPITSVALSSVYKILMLDMFDMNTIGIHNTMHLIVDSVTNCHFEVTDPTSEEVVLMKILQVLL
ACVKSKVSVALSNQHICNIVDTCFRVVQQTSAKSELLQRIARHSMHELVRSIFSRLPEVDSTCYKHAGGGNLTND
EIQQVNGDFSFAAPQSGPKDSTPQLDQLAIAYSTSSPSTALVDGPTDGSATKEVAQYDLHLMTQKFGVPCMVEVL
HFLCSLLNISEHVDQVRKSNSLAFDEDVPLFSLTLINSAIELGGSAFHRHPRLLNLIQDNLFCSLMQFGSTTSPL
ILSMVCSIVLNLYLHLRSKLKLQLEAFFSSVILRLAQSKYVASYQQQEVAMEALVDFCRQKTFVMELYANYDCDI
SSRNIFEELVTLLSKSAFPVNSPLSTMHILALEGLIAVIQGIAERIGDEQVGRAKVPLAFEEYKPFWLFKCDSYE
EPVHWVPFLRKMKFIKQQLLTGADHFNRDPRKGLEFLQKIRLLPEKLDAQSVAFFLRFTCGLDKNVVGDFLGNHE
EFNVQVLDQFAQTFDFKGMTLDTALRIFLETFRLPGEAQKIQRVLEAFSARYYEQSPHIFANKDAALLLSYSLIL
LNTDQHNAQVKRKMTEEDFIRNNRRINGGNDLPRELLSGLYHSICENEIRISPEQGSGPCELRQSHWIDLMHKTQ
NTAPYIVAACKPYLDDDMFAIISGPTVAAISVVFDHAEEEDAYQLCIDGFLAAAKIAAYFHLDSVLDDVVSSLCK
FTTLLNPSYYEDFLFAYSTDTRARMATSAVFTIANSYGDCIRTGWQNIINCILRFFEFGFLSSFSVSDVADEPAC
SAVPGNVKPLANSISLPHMPTVAPKKPSGLMGRFSQLLSFEVEEPISQPSEEELAAQQCILEIVQKCQINNIFLE
CKFLQADSLMHLVGAVVAEQAQTETRYPEDEVNAVFCLELLIAITLNNRDRISLLWQSVYEHISNIVQSVAVPCA
LVEKAVFGLLRICRRLLPYKENLVDELLRSLQLVLKLDARVADAYCEQITQEVGQLVRTNAHCIRSQMGWRVILS
LLAITARHPDASEAGFEALCFIMGDGAHLLPSNFILCVDGARQFAESRVGEVERSVHALDLMSGSLTCLVQWARE
SKEVMNEDAAAAAMFTQNIGEMWLRLVQGLRKLCLDDREAVQNHSLLSLQKCLSGVQEIHLSHNQWVQCFDVVIF
PLLDDFLEIAQGPSSKTYRNMEAVLVVAIKLLSKVFLLSINEVMQLPRFENLWMSVISQMESYMKVKIRGKKSEK
LQELVPELIKNSLLIMKAGGVLKPENGSSGDTSSLWELTWSHVNKLAPSLQSEVFTDAELDKASARGQTSQGNIE
ALQVSTCVSVVSTPTKIVAQEG*
**>Amacr14g03290**
MSRLIKKEVGISCSINTEVGAVLAALRKSYDEEIYHDTTIQQQSLKSLRSLLFNPQQEWQLVDPSIYLSPFLDII
QSDDIPASATSIALSSLIKLLKLPVFQDKTPGAQQAMNSLVMAATSCRLERTDPLSEEAILMKVLQLLNAVMTHP
ASRLIRDESVCRVVNTAFQVVQQSSNRGNLLQRTAKFVMLEICQTVFSRLNDIQVKEGENSESDSEDMDDNDIVN
TGFSLRCVVDIFQFLCSLLNVVQTVEIDGVVTQCNDEDIQILALVLINTALELSGDGFSKHPKLRRMIRDDLFHH
LIHYGTRTSALILSMICSTVLNSYHFLRGSLRLQLEAFFNFVLFRVAKFGSSMQYQEVALEGIINLCRQPDFLIE
TYMNYDCALTCSNVFEDIGKLLCKLAFPLTGPLTSLQLQAFEGLIIVIHAIADKVKRGDEVSIEPLPVQHSECSP
IWEEPVRKQGEFETWIRNLQLKKAQKRKVLIAGTHFNEDEKKGLEYLKISSLIPDPPDAKAFAYFFRFTPGLDKT
AIGDYVGDPEDFHLQVLNEFTCTFDFTGVVLDTALRTYLETFRLPGESQKIHRILEAFSERFFNQQSSEVFETKD
AVFILCYSLIMLNTDQHNPQVKKKMTEDEFIRNNRAINNGKDLPREYLSELFQSIATHAIAVFAETTLLVEMNPS
RWIELMSRSKYIQPFIACDFDHRLGRDMFASIAGPTLAALSAIFEQADDEDVLNECIEGIVQVAKIARHGLDDIL
DELVASLCKFTTLLNPYASSEETLFSFANDIKSRMATLAAFTIANKFGESIRGAWRNIIDCLLKMKRLKLLPSSL
VDYGTLVSKEGESHSRSMSEDLNLGNGRGSSSTAASRLSNLLLESNEDSLVLGGEFEQSLKVIEQCQIDKIFRNS
SKLPVDTLLNLGRSLIFAAAGKGQKFTTAVEEEETVIFCWDLVGAIAYANIDRYSTLGPAYNEYIIAVTQFPLFS
PVPFAEKGAIILFKLCVKLLSTYQTDKYPEEIMFKSINMMWKLDKEILDTCSDIIIQTVCKILTKYPGNVQTQIG
WKTLLHLLSSNGRLPENYDRAVEAIIQLLSDEKNLSGTNYAYCMDCAFGFVALRTSLVEKNILIMDLMANSVNLL
IQWSREHSDPGSSISNTSITSSSSMEDGIKSSNFTLNLFVKLGEALRKTSLARREEVRNHAVASLKKSFELAVGL
AFTSTNCINCLNYVVFAMVDDLHEKMLDYSRRDNAEREMRSMEGTLKNAMELLTDLFLLYLKSICDNPGFRTFWL

GLLRRMDTCMKADLGAHGPSKMQDVVPDLLKKMIAAMKDNEVLVPKENDDLWEITYIQIQWIAPAIKGELFPDVP
S*
**>Spiol_LOC110785052**
MGRLKMQSGIKSIEEEPEESEVTSSDKAALSCMINSEIGAVLAIMRRNVRWGGRYMSGDDQLEHSLIQSLKALRK
QVFMWQQQWHNINPVLYLQPFLDVIRSDETGAPITSVALSSVFKILTLDVLDLNTANIEDAMHLVVDAVTSCRFE
VTDPASEEVVLTKILQVLLACMKCKASVVLSNQHVCTIVNTCFRIVHQATTKGELLQRMAAHTMRELVKCIFSHL
PLVEGVEHSVVNNGSINSEFGGLANEFNFPAKQFDNGNGNSEYDGLLPTATFESNAPAGMVAGVVDENGDGSGKE
SKTYDLHVMTEPYGVPSMFEIFNFLCSLLNVVDHIGIGPRSNGLAIDEDAPLFALGLINSAIELGGASFLRHPSL
LNLIQDELFRNLMLFGLSPSPLILSMVCSIALNLYHHLRMELKLQLEAFFACVILRLAQSRYGASYQQQEAAMEA
LVDFCRQKTFMVEMYANLDCDITCSNVFEDLANLLSKSAFPVNSPLSSMHILALDGLIAVIQGMAERIGNGSLGL
EQAPVSLEEYAPFWMVKCDSYNDPNHWVPFVRRRKYIKRRLMIGADHFNRDPKKGLEFLQGTHLLPEKLDPQSVA
CFFRYTAGLDKNLVGDFLGNHDEFCVQVLHEFAQTFDFQDMNLDTALRLFLETFRLPGESQKIQRVLEAFSERYY
EQSSHILANKDAAFLLSYSIIMLNTDLHNAQVKKKMTEEDFIRNNRHINNGEDLPREYLSELYHSICKNEIRTTP
EQGAGFAEMNPSRWIDLMHKCKKTAPYIVADTRAYLDHDMFAVMSGPTIAAISVVFDHAEHEEVYRTCIDGFLAV
AKISACHHLEDVLDDLVVSLCKFTTLLNPSSVEEPVLAFGDDMKARMATVTVFTIANLYGDYIRTGWRNILDCIL
RLHKLGLLPARVASDAADESELSSDPGQGKPLTNSLSASHLPSMSTPRRSSGLMGRFSQLLSLDTEEPRSQPTEQ
QLAAHQRTLQTIQKCRIDSIFTESKFLQAESLLQLARALIWAAGRPQKGAGSPEDEDTAVFCLELLIAITLNNRD
RIVLLWQGVYEHIANIVQSTVMPCALVEKAVFGLLRICQRLLPYKENLADELLRSLQLVLKLDARVADAYCEQIT
QEVTRLVKANATHIRSQMGWRTITSLLSVTARHPEASESGFDALVFVMSEGAHMLPCNFILCVDSARQFAESRVG
QSDRSIRSLDLMAGSVGCLVKWVSETKEAVEEDAVKMSQDIGEMWLRLVQALRKVCLDQREEVRNHAISSLQKCL
TEMNEVHLEHSLWLQCFDMVIFTMLDDLLEIAQGHSQKDYRNMEGTLTVAIKLLSKVFLQLLSELSQLTTFCKLW
LGVLSRMEKFTKAKIRGKKSEKLQEIVLELLKNTLVEMKNSGVLAQRSALGGDSLWELTWLHVNNISPSLQSEVF
PSAEVDESSNETPVEPVAAES*
**>Spiol_LOC110804592**
MGRLKLQSFVNTIEEEPEESEVTSPNKAALSCMINSEIGAVLAIMRRNVRWGGRYMSGDDQLEHSLIQSMKALRK
QIFMWQQQWHNINPVLYLQPFLDVIRSDETGAPITSVALSSVYKIFTLDVLDINTTNIEDAMHLVVDAVTSCRFE
VTDPASEEVVLTKILQVLLACMESKASVVLSNQHVCTIVNTCFRIVHQATTKGELLHRMAVHTMRELVRCIFSHL
PEVESMEHFVVNRGSMKSEIGGLDNEFNFSAKQFNNGNGSSEYDGHLPSANFGSTATVSRSAGVVDENSGGSEME
SRAYNLDVMTEPYGVPSMFEIFRFLCSLLNVVDHIGISSRSNGMEIDEDAPLFALGLINSAIELGGASFLRHPSL
LDLIQDELFHNLMLFGLSPSPLILSMVCSIALNLYHHLRMDLKLQLEAFFSCVILRLAQSRYGASYQQQEAAMEA
LVDFCRQKTFMVEMYANLDCDITCSNVFEDLANLLSKSAFPVNCPLSSMHILALDGLIAVIQGMAERIGDGSLDL
EQAPVSLEEYAPFWMVKCNNYSDSNHWVPYVRRRKYIKRRLMIGADHFNRDPKKGLEFLQGTHLLPEKLDPQSVA
CFFRYTAGLDKNLVGDFLGNHDEFCVQVLHEFAQTFDFQDMNLDIALRLFLETFRLPGESQKIQRVLEAFAERYY
EQSSHILANKDAAFILSYSLIMLNTDLHNVQVKKKMTEDDFIRNNRRINGGEDLPGEYLLELYHSICKSEIRTTP
EQGAGFAEMNPSRWIDLMHKCKKTAPYIVPDSRAYLDHDMFSIMSGPMIAAISVVFDHAEHEEVYRTCIDGFLAI
AKISACHHLEDVLDDLVVSLCKFTTLLNPSSVEEPVLAFGDDMKARMATVTVFTIANRYGNYIRSGWRNILDCIL
RLHKLGLLPARVASDAADESELSSDPGKGKPFMNSLSASHLPSMGTPRRSSGLMGRFSQLLSLETEEPRSHPTDQ
QLAAHQRTLQTIQKCHIDSIFTESKFLQAESLLQLARALIWAAGRPQKGAGTLEDEDTAVFCLELLIAITLNNRD
RIVLLWQGVYEHIANIVQSTVMPCALVEKAVFGLLRICQRLLPLKENLADELLKSLQLVLKLDARVADAYCEQIT
QEVTRLVKANSTHIRSQMGWRTITSLLSVTARHPEASEPGFDALVYMMSEGAHMMPCNFILCVDTARQFAESRVG
QSDRSIRALDLMAGSVGCLVKWVCETKDALEMSQDIGEMWLRLVQALRKVCLDQREEVRNHAVLSLQKCLTEMDE
VHLAHSIWLQCFDMVVFTMLDDLLEISQGHSQKDYRNMEGTLTFAVKLLSKLFLQLLTQLSQLTTFCKLWLGVLS
RMEKYMKVKIRGKRSEKLQNIVLELLKNTLVEMKNRGILAQRSALGGDSLWELTWLHVNNITPSLQSEVFPCAGA
DGSSPVEPVITES*
**>Spiol_LOC110776164**
MTSLELQPASFPSMVNTEIGAVLAVMRRNVRWGTRFTFADDQQDHSLILSLKTLRKKLFTSHAHWHDIDPVTYLQ
PFLSVIQSEETGAPITSVALSSVYKILTLDIFDMETINIQDTMHLIVDAVTSCHFEVTDPASEEVVLMKILQVLL
ACVKCKAAVALSNQHVCNIVDTCFRVVQQTSAKGELLQRIARHSMHELVRSIFSRLPEILCHELDSGGNLINHEI
QQVNSDPIFVAPQPDSKDNTPTDSKLTETCSTSSSSAPFVDCPSNGTTTMDTVQYESHLMNQQFGVPCMVEILHF
LCSLLNVKEHADMNRKSNSLAFDEDVPLFALGLINSAIELGGSSFQRHPRLLKLIQDNLFCNLMQFGMSMSPLIL
SMVCSIVLNLYQHLRTKLKLQLEAFFSSVILSLAQSRYGASYQQQEVAMEALVDFCRQRTFIMEMYANYDCDLTS
RNIFEELVNLLSKSAFPVNTPLSTMHVLALEGLVAVIQGMAERTADERVDLGNTSITLEEYKPFWLFKCNGYEEP
GHWVPFLRQTKYIKQRLMAGADHFNRDPKEGLEFLQSTHLLPEKLDAQSVACFLRYTTGLDKNLVGDFLGNHEEF
YVQVLNEFAQTFDFKGMNMDTALRIFLETFRLPGEAQKIQRVLEAFSARYYEQSPHILANKDAALLLSYSIILLN
TDQHNSQVKKKMTEEDFIRNNRLINDGNDLPRELLSELYSSICENEIRITPEQGSGSCEIRLSRWIDLMQRSQNT
SPYIAAAYKPYLDDDMFAIVAGPAVAAISVIFDHAEDEDVYQLCIDGFSAASKIAAYLHHDSVLDDIVISLCKFT
TLLNPSYLEEFLLAYSNDTRARISTTTVFTIANRYGDCIGTGWQNIINCILRFYEFGFLSAFAVCDAADEPALSA
LPEHVKPLANTLSSPHIQISVAPKKPSGLMGRFSQLLSFETEEPISQPSEEELAARLCTQEIVQKCQINNIFTES
KFLQADSLMHLVRAIVGAEQIQAETGYPEDEANATFCLELLIAITLNNRDRISLLWQSVYEHISNIVQSTMAPCA
LVEKAVFGLLRICQRLLPYKENLIDELLKSLQLVLKLDARVADAYCEQITQEVGQLVKTNANRIRSQMGWRIILS
LLAITARHPDASEAGFEALSFIMSEGAHLLPSNFVLCVDGARQFAESRVGEVERSVHALDLMSGALTCLVRWARE
SQEVIDKDDATMVVQDIGEMWLRLVQGLRKLCLDDREAVRNHALSSLQKCLSGAQEIHLSQSLWLQCFDVVVFPM
LNDFVEIVQGQSSKTYKNMEAMLVVAIKLLSKVFLLSINEVMQLPTFCELWMTVLRQMETYMKVKIRGKKSEKLQ

ELVPELLKNSLLIMKTRGVLRPESGSDGDSLWELTWSHVNKIVPSLQAEVYSEADLDRGQHNQGNKEASTCVVLP
PSKPVALEG*
**>Spiol_LOC110777245**
MTKYMKKEIGISCTINTEVGAVLAALRNSFELPAADDNYHHTVIQQQSLKSLRSLIFNPQQEWQFIDPSIYLSPF
LDVIQSDDVPAAATNIALSSLIKLLKLQLFDEKTPGAQEAMNSLVIATTSCRLERTDPLSEEGILMKVLLLLNGI
MMHPASCLLKDESVCRVVNTAFLVVQQASHRGNLLQRTAKYVMLEISHTVFSKLNNIEAKEGDNNTESDADDNDI
VNDGFSLRCVEDIFQFLCSLLNVVQTVETDGIISQSNDEDIQILALMLINTALELSGDGFSKHPKLCRMIQDDLF
HHLIHYGTRSSALILSMICSTVLNSYNFLRRSLRLQLEAFLTFVLYRVAKNNSSMQYQEVALEGIINLCRQSTFL
IEIYVNYDCSFTCCNVFEEMGKLLCKLAFPMTTPLTSLQLQAFEGLIIIIHAIADKVNKGGDGVSFEPFPVGITE
YSPFWEEPLRKPDDLEKWVHILRLRKAQKKKLLIAGTHFNQDEKKGLEYLKISQLIPNPPDAKAFAYFFRFTPGL
NKTAIGDYLGDPDEFHLQVLKEFTNTFDFTGFALDAALRAYLESFRLPGESQKIHRILEAFSERFFNQQSSEIFE
TKDAVFILCYSLIMLNTDQHNPQVKKKMTEEEFIRNNRAINNGKDLPREYLSELFQSIATHAIAIFAQAALLVDM
NPSRWVELISRSRLMKPFTVCDFDHRLGRDMFGCIAGPTVAALSAIFEHAEDDEVLNECIEGIVSVAKIAQHGLE
DILDELVASLCKFTTLLNPYASAEETLYLFANDLKPRMATLAVFSIANKFGESMRGAWRNIVDCMLKLKRLKLLP
QSVVDYDTLVTAEGESHSRNIIEDSKYGSCRGNNSIAAGQLSHFLLESNEDSLVLGGSEFEQNLKVIQQCKIGNI
FSNSSKLPAETLLNLGRTLIFAAAGKGQKFTTALEEEETVTFCWDLVGAIAYANIDRFSTFWPAYNEYILAVTTF
PLFSPVPFAEKAAIIFFKLSVKLLSSNQTDKYAEEVLFKSISLMWQLDKEILDTCCEIIIQTVCKILTEYPGNIQ
TPIGWKTLLNLLSANGRLPENYDKAVETVIQLMSDGKNMSRTNYAFCIDCAFSFVALRTSLVEKNIEMMDLMANS
VKLLIQWSKGYSDPGSSISNASITSSPEDSMRLDYTVPLFVKLGEALRKTSLARREEVRNQAIISLRKSFELALD
LDMTSANCISCIDIVIFAMVDDLHEKMLDYSRRENAAREMRSMEGTLKYAMELLTDTFLLYLKPISENPGFSKFW
LSLLRRMDTCMKADLGVHGPSRMQEVVPELLKKMIAVMKTTEILVPRENDDLWENTYIKIQWIAPSITRELFPNI
PS*
**>Betvu_LOC104897851**
MGRLKLQSGIKAIEEEPEESEVTSSHKDALSCMINSEIGAVLAIMRRNVRWGGRYMSGEDQLEHSLIQSLKALRK
QVFMWQQQWHNINPVLYLQPFLDVIRSDETGAPITSVALSSVYKILTLDVLDLNIANIEDAMHLVVDAVTSCRFE
VTDPSSEEVVLTKILQVLLACMKSKASVVLSNQHVCTIVNTCFRIVHQATTKGELLQRMAAHTMRELVRCIFSHL
SEVESVEYSSVNGGPMKPEIGGLDNEYANEFNNGNRSSEYDGQLPSTNFGSNVPVGVGGGVVDENGAGNGKETRA
YDLHVMSEPYGVPSMFEIFHFLCSLLNVVDNIGISPRSNGLAIDEDAPLFALGLINSAIELGGASFLRHPTLLNL
IQDELFHNLMLFGLLPSPLILSMVCSIALNLYHHLRMELKLQLEAFFSCVILRLAQSRYGASYQQQEVAMEALVD
FCRQKTFMVEMYANLDCDITCGNVFEDLANLLSKSAFPVNSPLSSMHILALDGLIAVIQGMAERIGNGSLGLEQA
PVSLEEYTPFWMVKCDSYSDPNHWVPFVRRRKYIKRRLMIGADHFNRDPKKGLEFLQGTHLLPEKLDPQSVACFF
RYTAGLDKNLVGDFLGNHDEFCVQVLHEFAQTFDFQDMNLDTALRLFLETFRLPGESQKIQRVLEAFSERYYEQT
SHIMANKDAAFLLSYSLIMLNTDLHNAQVKKKMSEEDFIRNNRRINGGEDLPRELLLELYRSICKNEIRTTPEQG
AGFAEMNPSRWIDLMHKCKKTAPYIVADSRAYLDHDMFAIMSGPTIAAISVVFDHAEHEEVYRTCIDGFLAVAKI
SACHHLEDVLDDLVVSLCKFTTLLNPSSVEEPVLAFGDDMKARMATVTVFTIANRYGDYIRTGWRNILDCILRLH
KLGLLPARVASDAADDSELSSDPGQGKPLTNSLSASHLPSMGTPRRSSGLMGRFSQLLSLDTEEPRSQPTEQQLA
AHQRTLQTIQKCHIDSIFTESKFLQAESLLQLARALIWAAGRPQKGAGSPEDEDTAVFCLELLIAITLNNRDRIV
LLWQGVYEHIANIVQSTVMPCALVEKAVFGLLRICQRLLPYKENLADELLRSLQLVLKLDARVADAYCEQITLEV
TRLVKANATHIRSQMGWRTITSLLSVTARHPEASESGFDSLVFMMSEGAHMLPSNFILCVEATRQFAESRVGQSD
RSIRALDLMAGSVSCLVKWVLETKEAVEEDAVKMSQDIGEMWLRLVQALRKVCLDQREEVRNHAVSSLQKCLTEM
DEVHLAHSLWLQCFDVVIFTMLDDLLEISQGHSQKDYRNMEGTLTIAMKLLFKVFLQLLPELSQLTTFCKLWLGV
LSRMEKYMKVKIRGKRSEKLQDVVSELLKNTLIEMKNRGVLAQRSALGGDSLWELTWLHVNNISPSLQSEVFPSE
GVDVSSNTPPAEPVALES*
**>Betvu_LOC104906035**
MSCLEMESGSLACMLNTEISAVLAVMRRNVRWGSRFTSVDDQQEHTLIQSFKALRKKVFTWHAHWHTINPVMYLQ
PFLAVIRSEETGAPITSVALSSVLKILTLDLFHIHTVNIQDTMHLIVEAVTTCHFEVTDPASEEVVLMKILQVLL
ACVRSKASLALSNQHVCNIVETCFRIVQQTASKGELVQRIARHSMHELVRSIFSRLPEANRTYHELTSGGNLINH
EIQQVNSDPNLVAPEPGFNDNTTNYSEVFVGCSTSSSIPLVDGMTNGTMDDVLHYDSFLMNQQFGVPCMVEVLRL
LCSLLNITEHVDMSRKSNSLAFDEDVPLFALGLINSAIELGGPSFHRHPRLLNLIQDDLFCNLMQFGLSMSPLIL
SVVCSIVLNLYQHLRVMLKLQLETFFSSVILRLAQSSYGASYQQQEVAMEALVDFCRQKTFIMEMYANFDCNLTS
RNIFEELVNVLSKSAFPVNSTLSTMHILALEGLIAVIQGMAGRTKEERVRLGNTPTTLEVYKPYWLFKCDSYEEP
SHWVPFLRQMKYIKQRLMTGADHFNLGPKKGLEFLQGTHLLPEKLDAQSVACFLRYTGGLDKNLVGDFLGDHEEF
CVQVLNEFAQTFDFKGMSLDNALRIFLETFRLPGESQKIQRVLEAFSARYYEQSPHIFANKDAALVLSYSLILLN
TDRHNAQVKKKMTEEDFIRNNRRINDGNDLPRELLSELYNSICEHEIRITPEQGSGSCEMRQSHWMDLMQKTRNT
SPYIVAVDKPYLDDDMFALIAGPTVAAISVVFDHAEDEDVYQLCIDGFLAAAKIAAHFHLDSVLDDIVISLCKFT
TLLNPSYFEEFLLAYSNDTRARMATMTVFTVSNKYGDYIRTGWQNILTCILRFYEFGFLSAFSVCDTADEPGFSA
ILGMAGHVKPLLNSLPLPHIQSSVAPKKPSGLMGRFSQLLSFEIEEPSAQPSEEEVFAQQRIEETIQKCQINNIF
IESKFLQADALMHLVCAIVAAEQIQMEASYSENEVNAAFCLELLIAITLNNRDRISLLWQSVYEHISNIVQSTTV
PCALVEKAVFGLLRICQRLIPYKEDLIDELLKSLQLVLKLDARVADAYCEQITQEVGQLVKTNANRIRSQMGWRV
ILSLLAITARHPEASEAGFEALSFIMSEGAHLLPSNFVLCVDGATQFAGSRVGEVERSVHALDLMSGSLSCLVRW
ERESKELVNADEVAMVSKDIGEMWLRLAQGLRKLCLDDREVVRNHALFSLQKCLSGVQEIYLSHLWLQCFDVVIF
PLLDDFVEIAQGKSSKAYRNLEAMLLVAIKLLSKVFLLSLNEVMQLPTFNNLWMAVLSQMETYMKVKIRGKKSEK

LRELVPELLKNSLLIMKTGGMLIPGSGSGGDSLWELTWSHVNKIAPSLQAEVFPNADLDRGQHNQGKKEAIQASA
VSIPSKTVALEG*
**>Betvu_LOC104902176**
MVYHEKKISMSRYIRKEIGISCTINTEVGAVLAALRNIYETSTTTEDYYHDTTIQQQSLKSLRTLIFNPQQQWQY
IDPSIYLSPFLDVIQSDDVPASATSIALSSLIKLLKVQVFDEKTPGAQEAMNSLVMASTSCRIERTDQMREEGIL
MKVLQLLNGIMTHPASCLLKDESVCRVINTAFQVVQQSSSRGNLLHRTAKFVMLEISQTVFSRLNDIQVKEGENS
ESDSEELDDKNEIVNSGFSLRCVVDIFQFLCSLLNVVQTVEIDGIVSQSTDENIQILGLVLINTALELSGDGFAN
HPKLRRMMRDDLFHHLIHYGTRSSALILSMICSTVLNSYHFLRGSLRLQLEAFFTFVLFRVAKFGSSMQYQEVAL
EGIINLCRQPTFLIETYVNYDCAFMCCNMFEEMGKLLCKLAFPMTSPLTSLQLQAFEGLIIIIHAIADKLNKGGE
REGVSFEPFPVQISEYSPYWEEPVRKEDDLENWVYTLRLRKAQKRKLLIAGTHFNQDEKKGLEYLKISQLIPDPP
NAKSFAYFFRFTPGLDKTAIGDYLGDPDEFHLEVLNVFTSLFDFTGAVLDTALRTYLETFRLPGESQKIHRILEA
FSERFFNQQSSELFETKDAVFILCYSLIMLNTDQHNPQVKKKMTEEEFIRNNRAINNGKDLSREYLSELFQSIAS
NAIAIFAQANLLVEMNPSRWIDLISRSKYMTPFIACDFDHRLGRDMFGSIAGPTVAALSAIFEHADDDEVLNECI
EGIISVARIARHGLEDILDELVASLCKFTTLLNPYASAEETLYIFANDLKPRMATLAVFTVANKFGGSIRGAWRN
IIECLLKLKRLKLLPQSVVDYDTLVSAEGGSHSRSTSEDSKFGSGRGNNSIAAGRLSNLLLESNEDSLVLGGSEF
EQNLKVIQQCKIGKIFSDSVKLPAETLLNLGRSLIFAAAGKGQKFTTAVEEEETVTFCWDLVGAIAYANIDRFST
FWPAYNDYIIQVTQFPLFSPVPFAEKAALILFKLCVKLLSSPQSDKYPEEIMFRSINLMWKLDKEILDTCCEIII
QTVCKILIEYPGNVQTQIGWKTLLHLLSANGRLPENYDQAVEAIIQLLSDGKNLSRINYTYCIDCAFGFVALRTS
LVEKNIVIMDLMAKSVNLLIQWYRDQCEHESSMSNASMTSSSSLEDGKYGNFTLNLFVKLGEALRKTSLARREEV
RNQAVASLKKSFELAQELDLTSTNCINCFNYVIFAMVDDLHEKMLDYSRRENAEREMRSMEGTLKNAMEFLTDIF
LLYLKSISENPGFRTFWLGLLRRMDTCMKADLGVHGPSRMQEVVPDLLKKMIAAMKDREILVPKENDDLWEITYI
QIQWIAPSMKGELFPDVPS*
**>Drosp_00009154**
MGRIKLQPGIKAIEEEPKGCDDIPSNRAALACMINSEIGAVLAIMRRNVRWGGRYTSGDDQLEHGLIQSLKALRR
NVFMWEDQWGEISPVLYLEPFLEVIRSDDTGAPITGIALSAVYKILTRDVFDLRTVDVDKGMLMVVDAVTSCRFE
ETDPASEEVVLTKILQVLLACMKSKASVVLSNQHVCTIVNTCFRVVHQAGTKSELLQRIARHTMHELIRCIFAHL
PEVESIEHSLVNGGISIKSEDGMFLGEMNETTNGIGLGKDGGALENAHLMTEPYGVPCMVEIFHFLCSLLNAVEH
IGATSRSNTIAFDEDAPLFALGLINSAIELGGASFLRHARLMNLIQNDLFLNLMQFGLSMSPLILSMVCSIVLNL
YHHLRTELKLQLETFFSCVILRLAQSKYGASYPQQEVAMEALVDFCRQKTFMVEMYANLDCDITCNNVFEDLANL
LSKSAFPVNCPLSSMHLLALDGLMAVIQGMAERVSDESMGLESAPVTLEEYTPFWMVKCGNYKDPNYWVPFVRHR
KYIKRRLMIGADHFNRDPKKGLEFLQGTHLLPEKLDPQSVACFFRYTAGLDKNLVGDFLGNHDEFCVQVLYEFTK
TFDFQDMNMDTALRLFVDTFRLPGEAQKIQRVIEAFAERFYEQSPHILADKDAAFLLAYSIIMLNTDLHNAQVKK
KMTEEEFIRNSRCVNIPREYLSELYHSICKNQLRTTPEQGTGPPEMNPSRWIDLVHRSKKTAPYIVADDGAYLDH
DMFAIMSGPTIAAISVVFDHAEHEDVYQTCIDGFMAIAKISACHHLEDVLDDLVVSLCKFTTLLNPSSVDEPVLT
FGDDTKARMATVAVFTIANRYGDYIRTGWRNILDCILRLHKLGLLPARVATDAADEMELPLEPGHGKPLPSTLTS
AHLASMGTPRRSSGLMGRFSQLLSLETEEPRSQPTEQQLAAHQRTLQTIQTCHIDSIFTESKFLQSESLLQLARA
LIWAAGRPQKSTGSPEDEDTAVFCLELLIAITLNNRDRIKLLWQGVYEHIASIVQSTVMPSALVEKAVFGLLRIC
QRLLPYKENLADELLRSLQLVLKLDARVADAYCEQITQEVSRLVKANANHIRSQMGWRNITSLLSVTARHPEASD
AGFEALQFIMSDGAHLSPLNFTLVVDTSRQFAESRVGPADRSLQASDLMAGSGMWLVRWANESKGASKEDAAKLT
QDIGDMWLRLVQAVRKVCLDQREEVRNHAVSSLKKCLTGMEDVLLPHALWLQCFDVVIFTLLDDLLEIAQGHSQK
DYRNIESTLTLAMKLLSAVFLQLLSVLSQLTTFCKLWLGVLNRMEKYVKVRIRGKKSEKLQELVIELLKNMLVVM
KTKGVLAPRSSLGGDSLWELTRLHVKNISPSLQSDVFGAEGPEQNYMGQLETTQVPEGLVSSPDRAVVSQG*
**>Drosp_00004000**
MQVYGAQVAMRQPFTVDSPTVVMQFGADMLEEHKEWNGHHEITAILCTINTEIGTLLAAMRRNVRWGGRFVSGDN
QLEDPLVYSLKDLRKKLLNLEDQWLNVSPVTYLQPFLDVIQSDQTSAIITGVALSSVYKILTLDAFDLDTENIAD
TMHMIVDTVTSCRFEVTDPVSEEQVLMKILQVLLACMKSKASVVLSNRHICLIVSSCFRVVRRAGTKRELLRQIA
RDTMHELVKCIFSRLPEIDSVNLEPTDENGLIQPEYGGLGIPLTFMDRVTADNDSSARTGSSFPSSLIVGAVDDH
STGAILENEAYSYDFQLMQQPYGLPCMVEVFQFLCSLLDIAEHEGKYPKSNTLAFDEDVPLLALGLINSAIKFGG
PSFRKHPRLLSLIQDDLFFNLIQCGSSTSPLILSMVCSIVLNLYYYMRAEIKLQLEAFFSYVIVRLAQSRYGASY
QQQEIAMEALVDFCRQKAFMVEMYANFDSEITSSNVFEDLVNLLSKSAFPVNSPLSALNTVALDGLIAVVQAMAE
RTGDAPLSFESPPAHLEEYTPFWMIKCEDYKNPSNWVPFIRHTKYIKRKLTIGADHFNKSPKRGFEYLQSLNLLP
EELDPKSVACFFRYTAGLDKYLIGDFLGDRDEFCVEVLLEFVRTFDFHEMSLDTALRIFLESFRLPGEAERIMKV
LEAFAEIYYEQSHHQLFFSKDAIFLLSLSMIMLNTDLRSAQVKKKMTEEEFVRTNRHINGGNDIPRDYLSELYHS
IRKNEIVTTPELGTGFPDITPSLWIDLMLKSGKAPPYILADNRPYFDYDMFSKMAGPTIAAISVIYDHAEGESVC
RTCIDGFMAVAKVSATYHLADPLDDLVISLCKFTGLLNPLRMEEFLVLFSERRKVRMAITTLFTLANSYGDYIRK
GWRNILDCVLRLHEFGLLPTHVTIDSVDGSALSSDTVDEKASPKSSSSAHFPSTALPRRQSGLFELVSQLLSLDK
EEPGPAEEVLAAHQRTKETIKHCHVINIFSESKFLQADSLVQLAGAIIAAGRSRETLSAEGNSSAVFCLELLITI
ALNNRDRISLLWSSVYEHICHIVQMTSIPCALVEKAVFGLLNISQRLMPYKEELVDELLRSLQLVLKLDARVVDA
YCKQITQEVGRLVKANAGVIRSQMGWRTILSLLAITARHPEASGTGFDTLSFIMSDGAYLLPSNFVLCADAAKQF
AESRVGPAERSIQALDLMADSVNSLVRWANDTKEASLMEDAVKMSQDIGEMWLKLVQGFRKICLDSREEVRNHSL
LSLQKCLVVTHDFCISHEVWLQCFDLVIFSMLDELFGIAQGYSQKIHHGIEGTLVIAMKLLYKVFMLSLDEPVQS
ASFSKLWLGVLDQTEKYMKIKIRGKRSEKLEELLPELLKNTLLVMKIGGVLEPDSSAGGELMWDLTWSHVNNIAP
SLLGQIFSDSGSEPLQQHAASEAAQVAETVASNLPIEV*

**>Drosp_00017218**
MGRVEMQFDSNIVGQHDDPSISPLKTTSVACKINTEVGTVLAVMRRNVRWGSHFLSADNKLEHPLIHSFKNLRKQ
IFTWQTQWHNINPVLYLQPFLDVIQSDETGAPITGVALSSVYKILTLDVFDLDTVNVGNAMHFIVDAVTSCRFEV
TDPASEEVVLMKILQVLLACMKSKASAVLTNQHVCTIVDTCFRVVHQAGTKSELLQRIARHTMHEVVKCIFFCLP
DISSMVPGAAVDHNLTVQENGKLGNAHNYTHGLSADNISVRYDGEADCATAPSSAAASSSDENSSGPGKAKSADS
DDAHLMKQPYGLPCMIEVLQFLCSLLDIVEHVGSDLKSNTLAFDEDVPLFALGLINSAIELGGSSFRRHPRLLRL
IQDDLFFNLMRFGLSRSPLILSMVCSIVLNVWSHLRTEIRLQIEAFFSSIILRVAESRYGASYHQQEVAMEALVD
FCRQKAFMVEMYVNFDSEITCSNVFEDLVNLLSKSAFPVNSPLSGLNTVALDGLIAVIQAMAERVEDAPPFIEIT
PVCLEEYTPFWMTKCDDYNDPRKWIPFTRQRKHIKKKLMMGADHFNRGSKKGFEFLQASHLLPKELDPQSVACFF
RYTAGLDKNLIGDYLGSHDEFAVAVLHEFARTFDFHDMSLDTALRIFLETFRLPGESQKIQRVLEAFAESFYEQS
RQIFVNKDAALLLSYSTIMLNTDCHNSQVKKKMTEEDFIRNNRRINGGEDLPREYLSELYHSICSNEIRTTRELA
GFQDTTPSLWIDLLLKSKKAPPYIMVDNRPYLDHDMLAIMVGPTIAAISVIFDHADDEAVCQTCVEGFLAVAKVS
AMYHLANVLDDLVISLCKFTTLLTPSYAEEFMVLFSEDTKAMMATTTLFTIANAFGDHIRTGWKNILDCVLRLHE
FGLLPSRVATDSGDESSLSINTDDGKTSAKSSSSQLPSTGLPRRPSGLMGRFSQFLSFDTEEPKSEPTDEEVAAR
KRTLETIEQCHINSIFRDSKFLQADSLLQLAGAIMAAGRSSREKTSAESSEAAIFCLELLITITLNNRDRISLLW
SNVYEHIAHIVQSTPVPCNLVEKAVFGLISICQRLLPYKENLVDELLRSLQLVLKLDARVADAYCEQITQEIGHL
VKANATHIRSQMGWRIVLSLLAMTARHPEASEAGFDTLSIVMGDGAHLLPSNFVLCVDAARQFAESRVGQAERSV
HALNLMAGSVDSLVRWAHETKGATMVEDATKISQDIGEMWLRLVQGFRKACTDSREEVRNHALLSLQKCLVVVRD
VCIPHELWLQCFDLVIFPVLDELFGITQDYSQKTYRNVEGTLIIAMKLLSKIFLLSIDEPMDPTTFGKLWLGVLG
QMEKFIKVKIRGKRSEKLQEMLPELLKNTLLVMKTGGVLSRGNAAGGEVLWDLTWSHVNNIVPSLLSEIFPDTSS
DTGSDRPPKHDGEPEAPKDAISLTPTRAVEPVASEG*
**>Drosp_00008616**
MSNIKRRRKIDVSCVLNTELCAVLAALRRSCDPSTTDETYSNNDVSLQQSLRSLRSLIFNPRQEWRNLDPSIYMS
PFLDVIQSDEVPALATGVALSSIVKILRLEIFDERTLGAREIMNVLVGAATGCRLENSETSYVEAIMMKILQLLL
GIMSHPASILLSDDAVCTVVNTCFHVVQQVQRKGDLLQRNGKNTMHEIIQIVYSRLPEIKCEKKQDEKASESDDE
DLDADNDVDSGFGVGCAVDIFHFLCSLLNVVEIFETEGMSSQRTDEDVQVFGLVLINTALELSGDVIAKHPKLLR
MIQDDLFHHLIHYGTRSSPLVLSMICSTVLSIYHFLRSSVRLQLEAFFTFVLLKIVAQGSSVQLQEVALEGVINF
CMQPAFLFEAYLNYDCSFLCGNLLEEIGRLLCKHAFPMDNPLTSIQVQAFEGLMLLIHAIADNVDKAGSVGSCGP
YEVEVSVYTPFWEESPMGEKDTDGWVESLRVKKNQKKKIFIAGSHFNLDDKKGLEYLKLSNLIPDSPEAKAFAYF
FRFTPGLDKTMIGDYLGDPDEFHIHVLKEFTQTFDFSGMLLDSALRTYLETFRLPGESQKIQRVLEAFSEQFYEQ
QSSNLFESKDVVYILCYSLIMLNTDQHNPQVKKKMTEDEFVRNNRAINGGKDLPREYLSELFQSIANNAITVFGQ
AGLPAEMNPNRWIELMNRSKAMEPFVTCEFNHKLARDMFANIAGPSVAVISAVFEHAEEDDILNECIEGLISVAR
IAQFGLEDILDELIASFCKFTTLLNPYASPEETLYIFANDLKPRMATLAIFTVANKFGGSIRDAWRNIVDCLLKL
KRLKLLPQSVVDPDALLADEPSDSTPRSARAGLQNGTAFGSKSQKASSISRFSHFLAMENGEDSLILGGSEFEQN
LKVIQQCKIGSIFSNSAFLPEDALLNLGRSLIFAAAGKGQKFSTAVEEEETVGFCWDLIVVLGFVNFHRYMTIWP
AYHEYMISVAQFPMFSPVPFAEKALIVLLKLCLKILSSYQSENNLEELIFKSINLMWKYDMEILETCSNAIMQTV
CKILNDYPANVQTQLGWKSLFHLLSVTGRRPDSYDQGVETIINLLLDGTHVSRTNYAYCIDCAFAFVALKNSPVD
KNVKILDLMANSVKLLVEWYRSFSDPGTTMSNASSSSTEENTMRSGNFTMNLFVKLGEALRKTSLARREEVRNHA
VSSLKKSFIFANELDLTPANCISSFAIVIFTMVDDLHEKMLEYSTRANAEKEMRSMEGTLKNAMEVLTDVFLHYL
RLIAENPGFRSFWLGVLRRMDTCMKADLDEHGPSKMEEVVPELLKKIIAAMKENDILVPRENDDLWEITYIQIQW
IAPSLKGELFPEV*
**>Suaar_00002098**
MGRLKLQSAINSIEEEPEEAEVTSSNKAALSCMVNSEIGAVLAIMRRNVRWGGRYMSGDDQLEHSLIQSLKALRK
QVFMWQQPWHNINPVLYLQPFLDVIRSDETGAPITSVALSSVYKILTLDAVDLNTTNIEEAMHLVVDAVTSCRFE
VTDPASEEVVLTKILQVLLACLKSKASVVLSNQHVCTIVNTCFRIVHQATSKGELLQRMAAHTMRELVRCIFSHL
SEVEGIDHSVVNNGSMKSEIGGLDNEFNFSTKQLNNGNINSEFDGQLPTAHFGSNVPVGMGGVIDENGSGDGKES
RAYDLNVMTEPYGVPCMYEIFHFLCSLLNVDDHIGFSARSNGLAIDEDAPLFALGLINSAIELGGASFLRHPSLL
NLIQDELFRNLMLFGLSPSPLILSMVCSIALNLYHHLRVELKLQLEAFFACVILRLAQSRYGASYQQQEAAMEAL
VDFCRQKTFMVEMYANLDCDITCSNVFEDLANLLSKSAFPVNSPLSSMHILALDGLIAVIQGMAERIGNGSLGLE
QAPVSLEEYTPFWMVKCDNYNDPNHWVPFVRRRKYIKRRLMIGADHFNRDPKKGLEFLQGTHLLPEKLDPQSVAC
FFRYTAGLDKNLVGDFLGNHDEFCVQVLHEFAQTFDFQDMNLDTALRLFLETFRLPGESQKIQRVLEAFSERYYE
QSSHVLANKDAAFLLSYSIIMLNTDLHNVQVKKKMSEEDFIRNNRRINGGEDLPREYLLELYHSICKNEIRTTPE
QGSGFAEMNPSRWIDLMHKCKKTAPYIVADSRAYLDHDMFAIMSGPTIAAISVVFDHAEHEDVYRTCIDGFLAVA
KISACHHLEDVLDDLVVSLCKFTTLLNPSSVEEPVLAFGDDMKARMATVTVFTIANRYGDYIRTGWRNILDCILR
LHKLGLLPARVASDAADDSELSSDPGQGKPITNSLSASHLPSMGTPRRSSGLMGRFSQLLSLDTEEPRSQPTEQQ
LAAHQRTLQTIQKCHIDTIFTKSKFLQAESLLQLARALIWAAGRPQKGAGSPEDEDTAVFCLELLIAITLNNRDR
IVLLWQGVYEHIANIVQSTVMPCALVEKAVFGLLRICQRLLPYKENLADELLRSLQLVLKLDARVADAYCEQITQ
EVSRLVKANATHIRSQMGWRTITSLLSVTARHPEASESGFEALVFMMSEGAHMLPSNFLLCIDAARQFAESRVGQ
TDRSIRSLDLMAGSVSCLVKWVHEAKEAAEEDAVKMSQDIGEMWLRLVQALRKVCLDQREEVRNHALSSLQKCLT
EMDEVHLAHSLWLQCFDMVIFTMLDDLLEISQGHSQKDYRNMDGTLTIAMKLLSKVFLQLLPELSQLTTFCKLWL
GVLSRMEKYMKVKIRGKKSEKLQDLVLELLKNTLLEMKNRGVLAQRSALGGDSLWELTWLHVNNISPSLQSEVFP
SAVVEESSDPTPAEPVASES*
**>Suaar_00025615**

MSYSEMQPSALACMINTEIGTVLAVMRRNVRWGSRFTSADDQQEHSLIQSLKSLRKEMFTWHGQWHNINPVMYLQ
PFLDVIQSEETGAPITSVALSSVYKILTLDLFDTNTINIQDTMQLIVDAVTTCHFEVTDPASEEVVLMKILQVLL
ACVKSKASVALSNQHICNIVDTCFRVVQQTRTKSELLQRIARHSMHELVRSLFARLREVDNSYLELGSGDNLMNH
EVNSNSILVTPQPGSTDETAKCGELPLKSSFPSSSIPLLDGPSNGTAAMNAVQHDLNHAHQPFGAPCMIEIFDFL
CSLLNVIEHVGMTRKSNSVAFDEDVPLFALGLINSAIELGGSAFLRHRRLLVLIQDNLFCNLMHFGLSRSPLILS
MVCSIALNLYQHLRTELKLQYEAFFSSVILSLPQSRYGASYQQHEVAMEALVDFCRQETFIMEMYANYDCDLTSR
NVFEELVNLLSRSAMPVNSPLSTMHILALEGLIAVIQGMSKRTGDEHVNLGSTAITFDEYKPFWLSNCDSYEEPR
HWVPFLRQMKHIKQCMMTGAEHFNRAPKKGFEYLQAIHLLPEKLDAKSVACFLRYTAGLDKQLVGDYLGNHEAFC
VQVLNEFARTFDFKGMNMDTALRIFLETFRLPGEAQKIQRVLEAFSDTYYEQSPHILANKDAALLLSYSLILLNT
DQHNSQVKKKMTEGDFIRNNRCINGGNDLPHELLSELYNSICENEIRINADQSSGSCEMGQSHWIDLMQKARNAS
PYIIAPNKPYLNDDMFSIVASPIVAAISVVFDHAEDERVYQLCIDGFLAAANTVSFFHLDSLLDDIIVSLCKVTN
LLNPSYFEEFVLAYSNDLRARMATVTVFTIANKYGDCIRAGWKNILTCILRFYEFGLISAAAVCDAADEHTFLAV
HGHVKPPESSFSSSQMQISVAPKKPSGLMGRFSQLLSFETEEPKSEPSEEEVAARQFTLEIIPKCEINCIFAESK
FLQADSLMNLVSAIVAAEQIETETGYPEDEVNAAFCLDLLIAITLNNRDRISLIWKSVYDHISKIVQSTTVPCAL
VEKAVFGLLKICQRLLPYKEDLADELLRSLHLVLKLDARVADAYCEHITQEVGQLVKTNANCIRSQTGWRVILSL
LAITARHPDASETGFESLSFIMSDGAYLMPSNFILCVDGARQFAESHVGEVERSVRALDLMSGSLTRLVWWAHES
QEVVDEDTAAMLSQDIGEMWLRLVQGLRKLCVDDREAVRNHALLSLQKCVSGVQELHLSHTLWLQCFDAVIFPML
DNFLELAQGHPSITYRNVEAMLIVAVKLLSKVFLLSINEVMKLPTFCNLWMVILNKMETSMKVKIRGKKSSKLQE
LVPELLKNSLLIMKSVGMLTPDPGSGSGEVSFWELTWSHVNKIAPSLQDEVFPDAELDNGHHSNQASACVLSTPT
QTVALVDTT*

**>Suaar_00007839**

MTIKKEIGIACTINTEVGVVLAALRNGYETSSTTEDYYHDTTIQQQSLKSLRALIFNPQQEWQYTDPSIYLSPFL
DIIQSDDVPasatsialssliklklsVFNEKTPGAKEAMNTVVTATTSCRLERTDPLSEEGILMKILKLLNCIM
THPASCLLKDESVCKVLNTTFQVVQQSSNRGNLLQRSAKFVMLEISQTVFSKLNDMEVKEGensesdsedldend
IVTNNGFSLRCVVDIFQFLCSLLNVVQTIEIDGMISQTNDEDIQILALVLINTALELSGDAFSKHPKLCRMIRDD
LFHHLIHYGTRSSALILSMICSTVLNSYHFLRGSLRLQLEAFFNYVLFRVAKYGSSTQYQEVALEGIINLCRQPS
FLIETYVNYDCAFTCCNMFEDMGKLLCKLAFPMSSPLTSLQLQAFEGLIIIIHGIADKVNKGEDEVSFQLPWEIP
EYSPFWEEPVSQEEDLEGWAKTLRLRKAQKRKLLIAGTHYNEDEKKGLDYLKISHLIPNPPDAKAFAYFFRFTPG
LNKTAIGDYLGDPDEFHLQVLNEFTNTFDFTGVVLDTALRTYLEAFRLPGESQKIHRILEAFSERFFNQQSSEVF
ETKDAVFILCYSLIMLNTDQHNPQVKKKMTEEEFIRNNRAINNGKDLPREYLSELFQSISSHAIAIFAQGNMLVE
MNPSRWVELMSRSRYIGPFRACEFDHRLGRDMFSSIAGPVVAALSAIFEQADDDEVLNECIEGMISVARIARHGL
EDVLDELVASLCKFTTLLNPYASAEETLYIFANDLKPRMATLAVFNIANKFGEFIRGAWRNIIECLLKLKRLKLL
PQSVVDYDNLVSAEGGSHSRSMSGDSKLGSSRGNASLTTTRLSNLLLESNEDSLVLGGSEFEQNLRVIQQCEIGS
IFSNSSKLPVETLLNLGRSLIFAAAGKGQKFTTAVEEEETVTFCWDLVGAIACANVDRFSTFWPTYNDYIISVTQ
FPLFSPVPFAEKAAAILYKLCVKLLSSYQADRNAEEVMFKTINLMLKLDKEILDTCNDIIIQSVCKILTEYSGNV
QTQIGWKTLLHLLSANGRLPENYDLAVEAIIQLLAEGKDMSRTNYAYCIDCAFGFVALRTSLVEKNIQIMELMAN
SVNLLIQWYKDSSDPCNGTSNASMASSSSTTEENTKPSNFTLHLFVKLGEALRKTSLARREEVRNQAVASLKKSF
ELALGLEFTSTNCINCFNYVIFAMVDDLHEKMLDYSRRENAEREMRSMEGTLKNAMEMLTDVFLIYLKSISENSG
FRTFWLGLLRRMDTCMKADLGVHGPSRMQEVVPHLLKKMIAAMKDTQILVPKENDDLWEITYIQIQWIAPSIKGE
LFPDISS*

**>Poram_023444**

MGRLKLQSGIKAIEEEPDESDVTSSDKSALSCMINSEIAAVLAIMRRNVRWGGRYMSGDDQLEHSLIQSLKALRK
QVFTWQQPWHSINPVLYLQPFLDVIRSDETGAPITSVALSSVYKILTLDVLDLNTANVENAMHLVVDAVTSCRFE
VTDPASEEVVLTKILQVLLACMRSKASVVLSNQHVCTIINTCYRIVHQAATKGELLQRMAAHTMRELIRCIFSHL
PEVENVDQALANGTNSVKSEVGGLDNDYNVAAKQLNNGNANAEYDGQLPPTVGVGAGAVDENGAGSGKEAKPYDL
QLMAEPYGVPCMLEIFNFLCTMLNDLDHFGIGPGSNALAIDEDGPLFALGLINSAIELGGPSFVRHPRLLNLIQD
ELFRNLMVFGLSPSPLILSMVCSIVLNLYHHLRMELKLQLEAFFSCVVLRLAQSRYGASYQQQEVAMEALVDFCR
QKTFMVEMYANLDCDITCTNVFEDLANLLSKSAFPVNSPLSSMHILALDGLIAVIQGMAERIGNGSLGLEQASVS
LEEYSPFWMVKCDNYGDPNHWVPFVRRKKYIKRRLMIGADHFNRDPKKGLEFLQGTHLLPEKLDPQSVACFFRYT
AGLDKNLVGDFLGNHDEFCVQVLHEFAQTFDFQDMNLDTALRLFLETFRLPGESQKIQRVLEAFSERYYEQSPQI
LANKDAALLLSYSLIMLNTDQHNVQVKKKMTEEDFIRNNRHINGGNDLPREFLSELYHSICKNEIRTTPEQGAGF
AEMTPSRWIDLMHKCRKTPPYIVADSRAYLDHDMFAIMSGPTIAAISVVFDHAEHEEVYQTCIDGFLAVAKISAC
HHLESVLDDLVVSLCKFTTLLNSSSVEEPVLAFGDDAKARMATVTVFTIANRYGDYIRQGWRNILDCILRLHKLG
LLPARVASDAADDSELSADPGQGKPLTNSLSATHLPSMGTPRRSSGLMGRFSQLLSLDTEEPRSQPTEQQLAAHQ
RTLQTIQKCHIDSIFTESKFLQAESLLQLARALIWAAGRPQKGAGSPEDEDTAVFCLELLIAITLNNRDRIVLLW
QGVYEHIANIVQSTVMPCALVEKAVFGLLRICQRLLPYKENLADELLRSLQLVLKLDARVADAYCEQITQEVSRL
VKANATHIRSQMGWRTITSLLSVTARHPEASDAGFDALVFIMSEGAHLLPSNFILCVDAARQFAESRVGQADRSI
RALDLMSGSVGCLVKWFQETKEAVREEDAIKMCQDIGDMWLRLVQALRKVCLDQREDVRNHALSSLQRCLTEIEE
VHLAHNLWLQCFDVVIFTMLDDLLEIAQGQSQKESRNMEGSLMIAMKLLSKVFLRLLPELSQLTTFCKLWLGVLS
RMEKYMKAKIRGKRSEKLQELVLELLKNMLLVMKNSGVLVQRSALGGDSLWELTWLHVNNISPSLQSEVFPNGGA
DERASTPAEPVAAES*

**>Poram_037122**

MGRVNLQSGMKSIGEESVNSEVTMSNKTAFSCMINSEIAAVLAIMRRNDRWGGEYMWGDDEFEHPLVQSLKALRK
QVFTWQQPWHSINPVLYLQPFLDVIQSDETSPPITSVALSSVYKFLTIDMLDLNTANVEDAMHLVVDSVTKCRFE
KVDPVSEEVVLMKILQVLLACMKSRASIVLRDHHVFTIVKTCFQIVTQAASKGELLQRLAVNTTRDLIICIFSHL
PAVDNMDQSLINGKISVKSELNDENGVYENGEEAKPYDLHLMMEPYGVPCMFEVFHFLCSLLNDPDHFGIGPGSN
SLEADEDAPLFALGLINSALELGGPSFGRHPRLLNLIQNELFRNLIAFGLSPSPLVLSTVCSIVLNLYHHLRKEL
KVQLEAFFSHVILRLAQNSNGASYLQHEVAMEAIVDLCRQKTFMADMYANYDCEITCGNVFEEIVDLLSESVSSV
NSPLSSMHILAIDGLVAVIQGMAERINNGSHGLEWDPVSFEEYAPFWTVKCDDYTDPNHWVPFVHLRKYIKSRLT
EGADHFNRNPKKGFEFLQREHLLPEKLDPQSVAYFLRYAPGLDKNVVGDFLGNPDEFSVQVLHAFAETFDFQDTA
LDTALRLFLETFRLPGEAQKIHRVLEAFSERYYEQCKNIFANKDAAFVLSYSVILLNTDLHNAQVKKKMTEEDFI
RNNRRINGGNDLPREFLSGLYRAICKNEIQIRTSPKPGAGFAEMTSPGLWIDLVLKSKKTAPYIETDSWVHLDRE
MFAIMCRPAVAAISKVFDHADPEEEVYKVCIDGFLAIARVSAFHRVENVCDDLVVALCKSTTLLDSSSHEDPDPI
LAFGDDPKAKTAVLTLFTIATSYGDYIRLEWRSILDCILSLHKLHLLPACLASDPPGPTSTVKPTSKSSPGLLSR
FTKLLSFDTEEIWSQPTEEQLAAHRRAAETIQKCHIESIFTDSKFLQPESLLKLIEALIKTAQKKEEDEETAVFC
LELLIRITLTNRDRIVLLWQAVYEHIARIVGSTNVPCVLVEKAVFGLLRICQRLLPYKETLADELLRSLQLVLKL
DARVADAYCEPITQEVGHLVRASASHIGSQFGWRMIVSLLSVTARHTEASEAGFDTLAFVMSDGAHLVPANFVLC
MDSARQFAESRVGGADRSIAALNLMSGSVGCLVKWARGETREAAVRDEDPNPIPRNSAVERSDEAILPQNPVFEG
SDVGILPQNPVIEGSNAVVLPLNSAIQKDQFQLGKDAVKMPQDVGVMWLKLLQALRKVCLDQREEVRNHALSLLQ
RCMIDVDGAHLEDGLWIRCFDDVIFATLDDLLEIAQGQSQKEYRNIEGTLMLAMKLLSKVFLHRLPKLSHLPTFR
KLWLGVLTRVDNYTRVKIRGRTSKKLQELLLELLKNMLLVMKNGDVFLGHGESLWDPTWLHVNKISPTLQSEIFP
SVVSDGTDLPLAVKPWLQKVDGIVSV*
**>Poram_026381**
MTRFIKREIGVSCTINTEVGAVLAALRTSSEPSSNPEDHYFQDISIQQSLKSLRTLIFNPQQEWRFIDPNIYLSP
FLDIIQADDIPAQATGVALSSLLKLLRLEIFDDKTPGAKEAMNALVTATTNCRLERTDPVSEEGVLMKILQLLTG
IMTHPASRLLKDEAVCTVINTCFQVVQQSATRGNLLQRTGKHAMLEITQMVFSRLNELEVKEGENSESDSEDLDD
DDEDTIDSGFGLRCVVDIFHFLCSLLNVVQTVENEGVVSQTTDEDIQWFALVLINTALELSGDGFANHPKLLRMV
RDDLFHHLIHYGTRSSAPILAMICSTVLNVYHFLRESIRLQLESFFNFVLIRIASFGGLVQFQEVALEGIINLCR
QPNFLIEAYVNYDCAFICNNTFEEIGKLLCKLAFPMGSPLTSLQLQGFEGLVIVIHTIADNINKVNREEVSFDPY
PVEITEFSPFWEEPTRDEDDDEDEEDDEDEDDEDLETWVNELRLRKAQKRKIFIAGTHFNQDEKKGYEYLRISHL
VPEPPDPKAYAYFFRFTPKLNKTAIGDYLGDPDEFHLQVLQEFTNTFDFTGMILDNALRTYLDTFRLPGESQKIQ
RILEAFSDRYYDHQTSELFETKDAVLIICYSLIMLNTDQHNPQVKKKMTEEEFIKNNRAINNGKDLPREYLSELF
QSIANNAITVFAKAGVTAEMNPSRWIELMGRSRVVQPFVACEYDHRLGRDMFASIAGPTVASLSAVFEHADDDDI
LNECIEAVVSVARIARHGLQDILDELVASFCKFTTLLNPYASTEETLYIFANDMKPRMATLAVFTIANRFGEAIR
GAWRNIVECLLKLKRLKLLPQSVVELDIPAPASEGGGGHKRSESGSMPGSGKGTSGRFSNLLLEGHEDSLVLGGS
EFEQNLKVIQQCRIGNIFRNSSTFPLETLLNLGRSLIFAAAGKGQKFTTSAEEEETVGFCWDLVAVIGFANIDRF
ATFWPSYNEYILSVAQFPLFSPIPFAEKAVLVLFKLCVKLFSSYKPDKYPEELIFKSINLMWKFDKEVLETCADF
ILQSVCTILTDYPGNVQTHLGWKTLLHLLSVTGRHPETYDQGVEVLIQLLSDVEHMSTTNYAYCIDCAFGFVALR
SSLIEKNILIMDLMANSVNLLIQWYRTYSDPGNITNSNASISSNEDNPKFGNLTMNLFVKLGEALRKTSLARREE
VRNHAVISLKKSFQLAEGLELTSTNCINCFNFVIFAMVDDLHEKMLDYSRRDNAEREMRSMEGTLKIAMELLTDI
FLQYLKIISDNPGFRTFWLGLLRRMDTCMKADLGVHGPSRMQEVVPELLKKMIAAMKDTNILVPKENDDLWEITY
IQIQWIAPSIKGELFPDVS*
**>Simch_030709**
MGRLKLQSGIKAIEEEPEEYEVSPSNKAALACMINSEIGAVLAIMRRNVRWGGRYMSSNDQLEHSLIQSLKALRK
QVFAWQQQWHNVNPVLYLQPFLDVVRSDETGAPITGVALSAVYKILTLDVLDQSTVNVEDAMHLVVDSVTSCRFE
VTDPSSEEVVLTKILQVLLACMKSKASVVLSNQHVCTIVNTCFRIVHQAGTKEVDNTEQALVNGSNPVKQEISGM
DNDYDFGAKEFNNGNGNSEYDGQLSSAGFGSSAPSGMIAGSLDENGVGNGREAKPYDLHLMTEPYGVPCMVEIFH
FLCSLLNVVEHVGMGPRSNTLAFDEDVPLFALGLINSAIELGGPSFLRHPRLLILIQDELFRSLMQFGLSPSPLI
LSMVCSIVLNLYHHLRTELKLQLEAFFSCVILRLAQSRFGALYQQQEVAMEALVDFCRQKTFMVEMYANLDCDIT
CSNVFEDLANLLSKSAFPVNSPLSSMHILALDGLIAVILGMAERIGNGSLGLEQAPVNLEEYTPFWMVKCDNYSD
PNHWVPFVRRRKYIKRRLMIGADHFNRDPKKGLEFLQGTHLLPEKLDPQSVACFFRYTAGLDKNLVGDFLGNHDE
FCVQVLHEFAQTFDFQDMNLDTALRLFLETFRLPGESQKIQRVLEAFSERYYEQSPHILANKDAALLLSYSLIML
NTDQHNVQVKKKMTEEDFIRNNRHINGGNDLPREYLSELYHSICKNEIRTTPEQGAGFAEMNPSRWIDLMHKCKK
TAPYIVADSRAYLDHDMFAIMSGPTIAAISVVFDHAENEEVYQTCIDGFLAVAKISACHHLEDVLDDLVVSLCKF
TTLLNPSSVDEPVLAFGDDTKARMATVTVFTIANRYGDYIRTGWRNILDCILRLHKLGLLPARVASDAADDSELS
TEPGQGKPLTSSISSVQLPSMGTPRRSSGLMGRFSQLLSLDTEEPRSQPTEQQLAAHQRTLQTIQKCHIDSIFTE
SKFLQAESLLQLARALVWAAGRPQKGAGSPEDEDTAVFCLELLIAITLNNRDRIVLLWQGVYEHIANIVQSTVMP
CALVEKAVFGLLRICQRLLPYKENLADELLRSLQLVLKLDARVADAYCEQITQEVSRLVKANATHIRSQMGWRTI
TSLLSVTARHPEASESGFDALVFIMSEGAHMLPSNFVLCIETARQFAESRVGQADRSIRSLDLMANSVTCLMRWA
RETKDAMKEEDVVKVSQDIGEMWLRLVQALRKVCLDQREDVRNHALSSLQKCLTEIDSVHLSHGLWLQCFDVVIF
TMLDDLLEIAQGHSQKDYRNMEGTLMFAMKLLSKVFLQLLSELAQLTTFCKLWLGVLSRKEKYMKVKIRGKRSEK
LQELVLELLKNTLLVMKNKGVLVQRSALGGDSLWELTWLHVNNIAPSLQSEVFPSAGTTQVNDSSSHGESLSAEN
*
**>Simch_029603**

MGCPEMPSGVNATKEEREGRDVSPARKAALICMINTEIGAVLAVMRRNVRWGSHFMSSDDQLDHSLIQSLKKLRK
QIFTWHNQWHNINPVLYLQPFLDVVRSEETGAPITSVALSSVYKILTLDVFDLNTVNIEDALHLIVDAVTSCRFE
VTDPASEEVVLMKILQVLLACMRSRVAVVLSNQHVCTIVDTCFRVVHQAGTKGELLQRIARHSMHELVRCIFSRL
PEVYSRDCALENGSGLFKHEIGKLDHDHSFIDGPRVGGNVISESDSKPSSTSIASGLSTNENQQGSEKDEAVQYD
SHLMKQPYGLPCMVEILQFLCSLLNVVEHVGMGAKSNTLAFDEDVPLFALGLINSAIELGGSSFRKHPRLLNLIQ
DDLFYSLMQFGLSTSPLILSMVSSIVLNLYHHLRTELKLQLEAFFSSVILRLAQSRYGASYQQQEVAMEALVDFC
RQKTFIVEMFANFDCDMASRNVFEELVNLLSKSAFPVNSELSAMHVLALEGLIAVIQGMAERTAEGPISFGNAPA
TVEEYRPFWLVNCENYSDPSVWVPFVRQMKYIKRKVTVGADHFNRGGKKSLEFLQEKHLLPEKLDPQSVACFFRY
TGGLDKNLIGDFLGNHDEFCIQVLHEFVRTFDFQDMTLDTALRIFLETFRLPGEAQKIQRVLEAFSERYYEQSPE
ILANKDAALLLSYSLILLNTDQHNVQVKKKMTEEDFIRNNRKINGGNDLPREFLSELYHSICNNEIRTTPDQGAG
FPEMNPSRWIDLVHKSKKVSPYIIADSKPYLEQDMFTTIAGPTIAAISVIFDHAEDEDVYRTCIDGFLSVAKIAA
SYHLESVLDDLVISLCTFTTLLNPSYFEEFILSFSDDPKARMSTTAVFTIVNKYGDYIRKGWRNILDCILRLHEL
GLLSMSVASDAGDDSTASADTGHGKFTTNSVSSSRLPSINMTKRSSGLMGRFSQLLSFETEEPKLQPTEEELAAR
ECALETIQKCQIKNIFTESKFLQADSLLHFAQAIMAAGRSRKESGYAEDENTAIFCLELLIAITLNNRDRISLLW
EAVFEHISNIVQSTVTPCPLVEKSVFGLLRICQRLLPYKENLADDLLKSLQLVLKLDARVADAYCEQITQEVGRL
VKANANHIRSQLGWRIISSLLAITARHPEASDAGFDALSYIMTDGAHILPSNFVLCIDAARQFAESRVGEVERSV
HALDLMAGSVACLVRWALETKEAAPEEDATKLSRDIGEMWLRLVQGLRKVCLDNREEVRNHAILSLQKCMSGVHE
VRLPKNLWLECFDLVMFPMLDDLLEISQGQSQKTYRHMDSTLVVALKLMSRVFLLSIPESVNSEKFTELWAGVLT
RMENYMKVKIRGKKSEKLQELVPELLKNTLLVMKTGGLLMEGGDLGGNTLCEMTWSRVSSMAPSLKSEVFPDAES
GQAHRAPAPPVETVASEG*
**>Simch_015694**
MPRRDDDDDDRKTKRKKREIGVSCMLNTGVSSVLAVLRRVSDIPDDGAGSYHQDNIslqqslkslrslIFNPQQE
WRNIDPSIYLSPFMDVVQSDDVPAAATGVALSSVLKVLRLEIFDEKTPGVKEAMNALVVAATGCRLERTDPLYEE
AVLMRIVQLLAGIMQHPASMLLSDESVCTVVNTCFQVVRQSAATPRGNLLQKTAKYTMHEIITSVYSRLPDIEVK
EGENSESDSEDLDSDDSLLGYSLRSAVDIFHFLCSLLNVLPAAAIAVPDVVADGGYSPPSADEDIQMFALVLINT
ALELSGDSFAKHPKLLRMVKDDLFHHLIHYGTRSGPLILSMICSTVLNLYHFLRRSIRFQLEAFFTYVFLKVGAL
GSSLQLQEVALEGIINFCRQPTFLIESYVNYDCSFICCNVVEEMGKLFCRQAFPTGNALTSLQLQAFEGIVIMIH
AIAEKVDKEDESNSIGQYLVEIKDGSSFWEEITDSSDKSEGWVDAVRVKKAQKKKVIIAGTHFNLDEKKGLEYLK
ISKLVHNPPDAKSYAYFFRFTSWLDKTALGDFLGDPDEFHMQVLKEFTNLFDFTGMILESALRTYLETFRLPGES
QKIQRILEAFSERFYDLQQSELFESKDAVLILCYSLIMLNTDQHNPQVKRKMTEEEFIRNNRAINNGKDLPREYL
SELFHSIATHAITLSGQLGNAVEINPNHWIEIMNRAKVFQPFMLVDFDHQLGRDMFASVAGPCVATLSAIFEHAD
DDEILNECIDALISVARIAQYGLQDILDELVASFCKFTTLLNPYASTEETLYIFANDLKPRMATLAVFTVANKFG
ESIRGAWRNIVDCVLKLKKLKLLPQSVVDEAENPSHSRSESGSGLYGNDSKSGSRGSNSGMMGRFSHFLTMDSTE
DSLVLGGSEFEQNLKVIQQCGIGNIFRNSANLPIDSLLNLGRPLIFSAAGKGQKFSTAVEEEETVGFCWDLILTI
AFANIHRLAIFWPSYHEYMISVAQFPMFSPIPYAEKATVVLFKLCLKILSSYQSDKYLEELIFKSINKMLKFDKE
IIDTCSDFIMQSVSKIINEYPANVQTQLGWTSLLRYLQAMGRHPDNYEQGIETLIMLMSDGTHLSRTNFAPCIDC
AFNFVALKNSPIEKNSILLDQMASSVNLLVKWYKNYSDPGMSSSSISSTSSAEDHNPKFGNFITNLFVKLGDFLR
KMCLARREEIRHQAIAALRKSFQLAEELEFTSTNCINCFSYVIFAMVDDLHEKMLEYARRDNAERETRSMEGTLK
SAMEFLTDVFLHYLKQISDNPSFRPFWLGVLRRMDTCMKADLGEHGPSRLQEVVPDLLKRIIAAMKETDILVPKE
NDDLWEITYIQIQWIAPSLKGELFPDLDA*
**>FagtaGNOMa_chr5-18.6**
MGRLKLQPGIKSIeeepeegeptpSNKAALACMINAEIGAVLAIMRRNVRWGGRYMSGDDQLEHALIQSLKALRK
QVFLWQNQWHNINPVLYLQPFLDVVQSDETGAPITGVALSTVYKILTLDVLDTSTVNVEDAMHLVVDSVTSCRFE
VTDPASEEVVLMKILQVLLACMKSKASVMLSNQHVCTIVNTCFRVVHQAGTKGELLQRIARHTMHELIRCIYAHL
REVDNSGSALINKGNSLQSEVLSNCILVQSAPEHFDNGNGTSEYDGQFPSVTSLSGTPSGLVTAGGMHENLNGYG
DVKEVGGYDSHLMREPYGVPCMVEIFHFLCSLLNVVEHIGMGPRSNTLAFDEDVPLFALGLINSAVELGGAAFLR
HPRLLNLIQDELFRNLMQFGLSMSPMILAMVSSIVLNLYHHLRTELKLQLEAFFSCVVLRLAQSRYGASYQQQEV
AMEALVDFCRQKTFMVEMYANLDCDITCGNVFEDLANLLSKSAFPVNSPLSSMHILALDGLIAVIQGMAERIGNG
SLGLEKAPVNLEEYTPFWMVKCDNYGDPNHWVPFVRRRKYIKRRLMIGADHFNRDPKKGLEFLQGTHLLPDKLDP
QSVACFFRYTAGLDKNLVGDFLGNHDEFCVQVLHEFAGTFDFQDMNLDTALRLFLETFRLPGESQKIQRVLEAFS
ERYYEQSPHILANKDAALVLSYSIIMLNTDQHNVQVKKKMTEDDFIKNNRHINGGSDLPREYLSELYQSICNNEI
RTTPEQGAGLPEMTPSRWIDLMHKSRKTATYILADSRAYLDHDMFAIMSGPTIAAISVVFDHAEHEDVYQTCIDG
FLAIAKISACHHledvlddlvvSLCKFTTLLSPLSVEEPVLAFGDDIKARMATVTVFTIANRYGDYIRTGWRNIL
DCILRLHKLGLLPARVTSDAADDSELPTDPGHGKPLTNSLSATHLPSMGTPRRSSGLMGRFSQLLSLETEEPRSH
PTEQQLAAHQRTLQTIQKCHIDSIFTESKFLQAESLLQLARALVWAAGRPQKGAGSPEDEDTAVFCLELLIAITL
NNRDRIVLLWQGVYEHIANIVQSTVMPCALVEKAVFGLLRICQRLLPYKENLADELLRSLQLVLKLDARVADAYC
EQITQEVSRLVKANATHIRSQTGWRTITSLLSVTARHPEASEAGFDALTFIMSDGAHMLASNFILCLDTARQFAE
SRVGQVDRSIRALDLMANSVACLVQWANeakeesesakISHDIGEMWLRLVQALRKVCLDQREEVRNHSLSSLQN
CLTGADGVHLSHNLWLQCFDMVIFTMLDDLLEIANGHSQKDYRNMEGTLIIAMKLLSKVFlqllqelsqlTTFCK
LWLGVLSRMEKYMKVKIRGKRSEKLQELVPELLKNSLTVMKTSGVLVQRSALGGDSLWELTWLHVKNIDPTLQAE
VFATAGSDYQSETVSPDGAVSSPTNETVAASES*
**>FagtaGNOMb_chr5-43.3**

MGHVETQPGICSIVDGCKQQTDMSSKMAALACTINTEIGTVLAVMRRNVRWGGRYGSGDNQLEHSLIQSLKALRK
QIFSWKNQWQDINPILYLQPFLDVIQSDETGAPITGVALSSVYKILTLDVFDLNNVSIKDAIHSIVDAVTSCRFE
VTDPASEEVVLMKILQVLLACMKNKASSVLSNLHVCTIVDTCFRVVQQAGTKSELLQRFARQTMHELVRCVFSRL
PDMEKSDELISEENPITQEIVRLNNKKYSTGQFDTVGFSSDNHSVSFASSGSADVGTRSEDENLDVTGKERQPEA
DDSDLMKLPYGVPCLVEILHFLCSLLNVAEHVGMISKSNTLAFDEDVPLFALGLINSAIELGGSSFVRHPRLLSL
IQDDVFRHLMQFGVSMSPLILSMVCSIVLNLYHHLRTELKLQIEVFFSCVILRLAHSRSGASYQQQEVAMEALVD
FCRQKTFMMEMFANFDCDISSSNVFEDLVNLLSKSAFPVNSPLSAMHILALDGLVALVQGLAERTGEGPLGLDNA
PTLEQFTPFWMIKCEDYSDPRCWVPFVCQMKYIKRRLMIGADHFNHGPKKGWEFLQVANLLPKELDAKSVACFFR
YTAGLDKNLIGDFLGNHDPFCVQVLHEFARTFNFQDMSLDTALRLFLETFRLPGESQKIQRVLEAFSERYYEQSP
EILANKDAALLLSYSIIMLNTDQHNAQVKKKMSEEDFIRNNRCINGGKDLPRDFLSDLYHSICKDEIRMIPEEGA
GFPDMTSSRWFYLMHKSDDAAPYIVPDNRPYLDKDMFAIVAGPAIAAISVVFDHAEDDDVYQTCIDGFLAIAKIA
AFYHIDSVLDDLVISLSKFTTLLNPAHVDEFLLSFSEDSKARMAMLAVFTIVNKYGDCIRNSWRNILDCILRLHE
YGLLFARVANDTTDDAVLSSEPENSKPIKNSSVTAIPKRSSGLIGRFSQLLSFDSEESIPQPTEEEVAARQRTSE
IVQKCQINNIFLDSKFLQADSLIQLAQAIMAAGQSLKESDSSEDEAGAVFCLDLLIGITLNNRDRISLLWQGVYE
HISNIVQSTVMPCSLVEKAIFGLLSICQRLLPYKENlvdellkslqlvlkldARVADAYCEQITQEVGHLIKANA
TRIRSQMGWRIVLSLLAITARHPEASEPGFDALSFVMGEGAYLLPSNFSLCVDAARQFAESRVGEVERSVHALDL
MGSSVSCLVRWARETKESMTEEESGRMSQDISEMWLRLVQGLRRTCLDNREEVRNRALLSMQKCLTVVHEIHLTC
TLWLECFDTVIFSLLDELLEIAIGPSHKNYRNMDttlmlsmklltkVFLLSIDEVTKLTTFSKLWLDVLTRMENY
LKVKLKGRRSEKLQELVPELLKNTLLVMKSGGVLRQESASLWELTWSHVNNMAPSLQFEIFPDVGPDQSPNKADA
SPPLATSTPIQAVNAEG*

**>FagtaGNOMc_chr6-36.2**

MDHVKMQSDVTSSRDRFKDGMDMASRKDALACAVNSEIVTVLAVMRRNVRWGGRFTLGDDLLEHSLIQSLKALRK
QIFTWRDQRQNISAILYLQPFLDVIQSDETGAPITGVALSSVYKILSLDVFDLDTIYIKDAIHSIVEAVTSCRFE
VTDPASEEVVLMKILQVLLACVKSKASNVLSNQHVCTIVDTCFRVVHQAGTKSELLQCIARHTMHELIRCIFSHL
LDLESLGKDLNNDENLSQQEVLHLWPIVLAILYGIFYQVLWVFGLEVYFLCIECNEYPSFVQIDRLNHRNTNTAD
IDDSYLMKLPYGVPCMVQILHFLCSLLNVAEHVDTDSKTNTLAFDEDVPLFALALINSAIELGGSAFRKHPKLLS
LIKDDIFCHLMQFRPSMSPLILSMVCSIVLNLYHHMRTDLKLQIEVFFSSIILRLAQSRYGASYEQQEVAMEALV
DFCRQKTFVMEMFANFDCDISCTNVFEELVKMLSKSAFPVNNPLSTMNILALDGLVVVVQQMAERAGEGRIGLNS
VPNLEKYAPFWMIKCNDYSDPSCWVPFVFQMRYIKRRLIIGVDHFNRGPKKGLEFLQVANLLPKEPDAESVAYFF
RYIAGLDKNLIGEFLGNHDQFSIQVLHEFAQTFNFQGMSLDAALRLFLETFRLPGESQKIQRVLEAFSERYYEQS
PQILANKDAALVLSYSVILLNTDQHNSQVKKKMSEEDFIRNNRCINGGKDLPREFLSDLFHSICKNEIRMIPEQS
AGFPDMTSSHWLDLMHKSENAALYIVPDNKPYLDQDMFPIVAGPTIASISVVFDHAEDDFSYQTCIDGFLAVAKI
AAFYHVHTVLDDLVISLCKFTTLLNHAYREEFLLSFAEDSKAKIATLAVFTIANKYGDCIRNSWRNILDCILRLH
EFGLLSACIANDVTDDTILSSEPGNGKPLKNSSSMGLPKRSSGLIGRfsqllsfeseesvsqPTEEELAARQLTF
EIVQKCQINNIFLDSKFLQAESLLQLARAIMAAGQTLKGTDDEAGAVFCLELLIGITLNNRDRISLLWQGVYEHI
SIIVQSTLMPCALVEKAIFGLLSICQRLLPYKENLVDDLIKSLQLVLKLDARVADTYCEQISREVGSLIKSNVTR
IRSHMGWRTVLSLLANTARRLEASEAGFETLSFIMNEGAHLLPSNFSLCVDAARQFAESRVGQVERSVHALDLMS
SSVTCLVRWAYETKGLITVEEATKMSHDIGEMWLRLTQCLKRTCLDNREEVRDHAILSMQKCLTGVYEIHLSSKL
WLECFNTMIFTLLDDFLEISTLQSRKDYQNMEDTLVLAIKLLSKVFLLSINELTQVTAFSELWMNVLTRMEKYMK
VKIKGRKSEkllelvpellkttllvMKSEGLLIQPSASDNFWEMTWSHVMNIAPSLQAELFPDAVSDHHSLHNKD
VEETNPNQAVSLASLAPIQIVASEG*

**>FagtaGNL2a_chr5-51.6**

MSQRRKMNEAGVPCMLNTEIGAVLAVLRKGATELSVGPAEESPYQDVslqqslrslrTLIFNPRQNWRNMDPSIY
LSPFLDVILSDEVPAVGTMVALAAILKIVKIEIFNDKTPGARHAMNAIVVATTGCRVEKTANHVQEAILMRILQV
LTAAMNCGASILLSDDAVCTIVNTCFNVLGNPSISDLLKRTAHSTMHEIIRNVYTRLPEIDLDDGFNhsededdd
fdygddddhdsgyGVRCTVDIFQFLCSLMNVVETVEAVMDGSYMTTSVQRTNEEVQMVALVMMNTAIELSGEVLG
KHPRLLRMIQDDLLHHLLHYATRSSPLVLSMICSTVFNIYNFLRRHLRLQLEAYFVFVLFRVASPPYPHQMQEVA
LEALVNFLRHHTFLIDMYVNFDCSFRCTNLFEEMGKLLCKQSFLTSSPMTSLQVQAFEGLLLMIHTIAENVDKES
AKNRDSLGPYTAVEIKDYEPFWEDAARDDLEMESWLDTLRLRKAQKKKLSMAGNHFNQDEKKGLEYLKLSNLVPY
DPPDPKAFAYFFRFTPRLNKTALGDYLGDPDSFHLEVLEEFTATFEFSGMLLDCALRSYLETFRLPGESQKIHRI
LEAFSERFYEQQSSDLFETKDAVFVLCYSLIMLNTDQHNPQVKRKMTEDEFIRNNRAINGRKDLPREYLCELFQS
IANNAITMFGQLGPIELTPSRWIELINQSKTTKSFTLCKFEPPWIGRDMFATIAGPCVAALTAIFEHTEEDEITN
ESVEGLISVARIAQYGLEDILDELLASMCKFTSLLNPYASTEETLYIFANETKPRVATLAVFTITNRFAHSIRAA
WRDIVDCllklkrlkllPQSVVQIPTHSRSVSGNLPNGSDPRAPNRRHSSSMVGKFSSLLPMDGNEDSLVLMGSE
FEQNVKIIQQCNIGDIFNNSNMKAESLTNLGRTLIFAAAGRGQKFSTALEEEETVTFCWDLILKASLANIGKIST
FWPACHDYILTVTQPPQFSPVPFAEKAFLALFKIVLSLLSSNRIEKSSEETLFISINLMWKLDMEILDTCCDLIY
SVVSKIINDYPANVQTQLGWKSLLHLLSVAGRHPEAYNLGVETLVELLSKGTYVSKTNYTYCIDCAFGFVALRNS
PIEKTVRILDLLAKSVDLLVYWCKNFSDAGISNSSMEDNNMMRSPNYMVNLFFKLGEVFRKTSLARREELRNHAI
LALKRSFVLAEDLEFTSVQCMSCFNYIVFAMVDDLHEKMIEYSTRENAEKEEMSMENTLGHALELMEEVFVRYLK
LMAVNAAFNAFWMGFLRRLDTCMKADLGEYGESRMQELVPEILKRVITEMKDQGVLVPREDDDLWELTYVQIQWI
APSLGESNLM*

**>FagtaGNL2b_chr6-47.9**

MSMRRKMKEAGMPCMLNTEIGAVLAVLRKGATELSVGPAEDSPYQDIslqqslrslrTLIFHPQQDWKNMDPSIY
LSPFLDVILSDEVPAVGTMValaailkiikiEIFNDKTPGAKDAMSAIVIATTGCRVEKTANHVQEAILMRILQV
FTAAMNCPASILLTDDAVCTIVNTCFNVLQQSNISDLLKRTAHSTMHEIIRNVYCRLIEIdaqgsggdqdesdde
dfdadnnsesRGYGVRCAVDIFQFLCSLLNVVETVEIDGYVSQKTNEEVQVIALVMMNTALELSGGVLWKHQKLL
RMIQDDLMHHLIHHGTFTSPLVLSMICSIVLNIYNFLRRHTRLQLESFFVFVLSRVATAGYSTQMQEVALEAIVS
FCRQPTFLIEMYVNYDCSFMCWNLFEEMGKLLCRQAFPKAGPMASSQVQAFEGLLLTVHTIADSVDRDCSVKGSL
GPYPIEIREYEPFWEDVARDDVDMEEWLDNlrlrkaqkkllMAGNHFNQDEKKGLEYLKLSNLVPQDPPDALAF
SYFFRFTPRLDKTAIGDYLGDPDSFHLQVLKEFTCTFEFNGMLLDSALRTYLETFRLPGESQKIQRILEAFSERF
YEQQSSETFESKDAVFILCYSLIMLNTDQHNPQVKKKMTEDEFIKNNRAINGGKDLPREYLSELFQSISNNAITM
FGQTGPIEMTHSRWIELINRSKSIKPYVLCTFDPRLGRDMFATIAGPCVAALSAIFEHTDEDDVMNESIEGLVSV
ARIAQHGLEDILDEllaslckltttllNPYASTEETLYIFANEVKPRMATLAVFTIANRFGASIRGAWRDIVDCll
klrrlkllPQSVIEAPQSSSSVDhrgsharsgsgsIPYGSEMRAPNRRHSSSMVGKFAPFLPMDGNEDSLVLMGS
EFEQNLKIIEQCKIGSIFNNSDLPEEALVHLGRSLIFAAAGRGQKFSTAIEEEETVSFCWDLILTTSLSSINKIS
LFWPAYHDNMLAVAQFPLFSPVPFAEKGILILFKLAVRLLSTYRIDKYPEELIFKSINLMWKFDKEILDTCCDLI
MQSVSKILSDNPENVQTPLGWKSLLHLLSVTGRHPEAYDLGVETLINLLSGANYVTRTNYAYCVDCAFGYVALKN
SPVDKNYKILDLLANSVKLLIQWNRsfsdpessisstsigsntsteeMMMRNSSYMINLFVKLGEALRKTSLARR
EEVRNEAIFALNKSFKLARELDLTSAHCINCFNYIIFAMVDDLHEKMVEYSRRENAEREMRSMETTLKYAMELLT
DVFLQYLKPISENTAFKPFWLGLLRRMDTCMKVDLGEHGASKMQELVPLLLTRIITEMKDEGILVPREDDDLWEI
TYIQIQWIAPSLGDTSSF*

**>Diaca_4337**

MGRLKMQPGIKSIEEEPEETEVTPSNKAALSCMINSEIGAVLAIMRRNVRWGGRYMSGDDQLEHSLIQSLKAFRK
QVFTWQQQWHNINPVLYLKPFLDVIRSDETGAPITSVALSSVYKILTLDVLDLTTTNIEDAMDLVVDAVTSCRFE
VTDPSSEEVVLTKILQVLLACMKSKASVVLNNQHVCTIVNTCFRIVHQASTKGELLQRMAAHTMRELVRCIFSHL
PEVQSSEHTLVNGGGYSMKSEIGGLDNEYNFPANQDNNGVGNSEYDGSLPNTNFGSNASAVVGGVMDETSSVNGR
DTTSYDLQVMTEPYGVPSMVEIFNFLCSLLNVVDHIGIGPRSNGSAYDEDGPLFALGLINSAIELGGNSFLQHPS
LLHLIRDELFRNLMLFGLSPSPLILSMVCSIVLNFYHHLRMELKLQLEAFFSCVILRLAQSRYGASYQQQEVAME
ALVDFCRQKTFMVEMYANLDCDITCGNVFEDLANLLSKSAFPVNSPLSSMHILALDGLIAVIQGMAERIGNGSLG
LEQAPVSLGEYTPFWMVKCDSYTDPNHWVPFVRRRKYIKRRLMIGADHFNRDPKKGLEFLQGTHLLPEKLDPQSV
ACFFRYTAGLDKNLVGDFLGNHDEFCVQVLHEFAQTFDFQDMNLDTALRLFFETFRLPGESQKIHRVMEAFSERY
YEQSSQVLANKDAAFILAYSLIMLNTDLHNVQVKKKMTEEDFIRNNRRINGGEDLPREYLLELYHSICKNEIRTT
PEQVAGFAEMNPSRWIDLMHKCKKSAPYIVADSRAYLDHDMFAIMSGPTIAAISVVFDHAEHDEVYRTCIDGFLA
VAKISACHHLEDVLDDLVVSLCKFTTLLNPSSVEEPVLAFGDDSKARNATVTVFTIANRYGDYIRTGWRNILDCI
LRLHKLGLLPARVASDAADELELAADPGHGKPLTNSMSATHLPSMGTPRKSSGLMGRFSQLLSLDTDEPSSQPTE
QQLAAHQRTLQAIQTCRIDSIFTESKFLQAESLLQLARALIWAAGRPQKGVGSTEDEDTAVFCLELLIAITLNNR
DRIVLLWQGVYEHIANIVQTTVMTGALVEKAVFGLLRICQRLLPYKENLADELLRSLQLVLKLDARVADSYCEQI
TKEVNRLVRANASHIRSQMGWRTITSLLSVTARHSDASKPGFDALVFIMSEGAHLLPSNFILCVDAARQFAESRA
GDTDSSIRALDLMSGSVSCLVKWVHETKETVKEEDAIKMSQDIGEMWLRLVQALRKVCLDQREEVRNHSLSSLQR
CLTEIDEVHLAHSLWLQCFDMVIFTMLDDLLEIAQGHSQKDYRNMEGTLIISMKLLSKVFLQLLPELSQLTTFCK
LWLGVLSWMEKYTKVKIRGKKSEKLHDVVLELLKNMLEEMKRRGVLAQRSALGGDSLWELTWLHVNNISPTLQSE
VFPPAGAVDETAPSPTEPIPSDR*

**>DiacaGNL1_sc4**

MHCLEMQPGVDVHGQGQVACEVTSNKSSLACIINTEIGAVLAVMRRNVRWGSRFVSADDQLEFSLIQSLKLLRKK
IFSWSPQWHTIDPVLYLQPFLDVIRSEETGAPITSVALSSVYKILTVDAFSVDTINIQDAMHLIVDSVTSCHFEV
TDPASEEVVLMKILQVLLACVKSKASSVLTNQHVCTIVDTCFRVVQQTGTKGELLQRIARHSMHELVRTIFSRLP
EFDSTYSELGSDGSFTDPQIQMPNGNPESVTLQSGSENHTPQNAELPSADHNSTSAPTVTSLQEMETIDTAQIDP
HLMNQPFGIPCMVEILQFLCSLLNVTEHTGMGPKSNTVAFDEDVPLFALGLINSAIELGASSFHKHSRLLKLIQD
DLFWNLMQFGLSASSLILSMVYSIVLSLYIHMRAKLKLQLEAFFSSVILRLAESKYGASYQQQEVAMEALVDFCR
QKAFMTEMYANFDCDLTSRNIFEDLVNLLSRSAFPVNSPLSTLHILALEGLIAVVQGMAERIENGHSVTDNSVFK
LEEYKPFWFVKSEDYEQPSHWVPFVRQMKHIKQRLVIAADHFNQGPKKGVEFLQQTHLLPEKLDPQSVACFLRCT
AGLDKNLVGDYLGNHDEFCVQVLHAFARTFDFEGMNLDIALRIFLETFRLPGESQKIHRVLEAFSERYYEQSPDI
LADKDAALVLSYSLIMLNTDQHNVQVKKKMKEEDFVRNNRRINGGNDLPREYLSELYHSICENEIRMTPEQGSGF
PEMTNSRWIDLLQKSKITAPYILAVSKPYLDKEMYSMIAGPTVAAISVVFDHADDEDVYQLCIDGYLATAKIAAY
FHLESVLDDMVISLSKFTTLLNPSYLEEFLFTFSSDAKAKMATTAVFTIANLYGDYIRTGWQNLLDCILQFYEFG
FLRSGFLYDASDDPAAPGHAKPFTNTLSSPQISAPVVSKKSSGFMGRFSQLLSFDTEEPISQPTEEELAAQQEIH
EIIQKCQISNIFTESKFLQADSLMELVRAIMAAGQIQKETSYLVNEATAAFCLELLIAITLNNRDRISLLWQAVY
GHISNIVQSTSMPCSLIEKAVFGLLRICQRLLPYKENLVDDLLSSLQLVLKLDARVADAYCEQITQEVGQLVKAN
ANHIRSQTGWRVVLSLLAITARHPEASEAGFNAVSYIMSDGAHLLPSNFVLCIEAARQFAESRVGDEKRSIHALD
LMSGSVSCLGRWALETREGINEEDAAMLSKDIGQMWLRLVQALQKLCLDNREDVRNHALLSFQKCLSGVQEINLS
KQLWLQCFDLVIFPMLDAFLEIAQTQSSKMYRNIEAALLITMKLLSKVFLLSINEIMHLPTFSDLWMAVISQMEN
YMSVKIRGRRSEKLQELVPELLKNSLLVMKAAGVLVPGSDFDKDGTLWELTWSRVNKIAPSLQAEVFTEADLGNR
KSSQFKNETIQESVQVGSTPIETVASEG*

**>Diaca_14696**

MTRFIKKQIGISCTINTEVGAVLAALRNGYDISAPDEFHDITLQQSLKSLRSLIFNPQQEWHLIDPSIYLSPFLD
VIQSDDVPAVATGVALSSVLKLLKLEIFDERTPGAKEAMNALVIATTSCRLERTDPITEEGVLMRILLLLAGIMT
HPAACLLKDESVCAVINTSFQVVQQSSNRGNLLQRIAKYNMHEIMQTVFSRLNEMEVKEGENSESESEEDDDDEV
INSGFGLRCVVDIFQFLCSLLNVVQTVEVDGVVSQSYDEDIQIFALVLINTSLELSGDAFAKHPKLLRMVRDDLF
HHLVHYGIRCSPVVLSMICSTALNAYHFLRGSIRLQLEAFFVFVLFRIATYGGSLQFQEVAMEGIINLCRQPTFL
IEIYINYDCAFTCCNIFENMGKLLCKQAFPVSNPLTSLQLQAFEGLIIIIHRIADKINTGENEASFDPYPVEINE
YSPFWEEPKRNEDDLETWVDALRLKKAQKRKILIAGTHFNQDEKKGLDYLKICGLVPVPPDAKSFAYFFHFTPGL
DKTALGDYLGDPDEFHLQVLNEFTGTFDFTGMILDSALRTYLESFRLPGESQKIQRILEAFSERFFEQQSSEMFE
TKDAVFILCYSLIMLNTDQHNPQVKKKMTEEEFIRNNRAINNGKDLPREYLSELFHSIATHPISIFAQAGINSEM
NPSRWIELISRSKVVQPYISCDFDHRLGRDMFATIAGPVVASLSAIFEHADDDEVLNECVEALVSVSRIARHGLE
DILDELVASFCKFTNLLNPYASAEETLYIFANDIKPRMATLAVFTIANKFGDSIRGAWRNIVDCLLKLKRLKLLP
PSVVDFDSTTGTTTAGGSHTRSESVFGSGRATGRFSHLLLEGNEETLVLGGSEFEQNLKVIQQCKIGSIFNNSSN
LPEETLLNLGRSLIFAAAGKGQKFTTAVEEEETVGFCWDLVGSIGLANVDRFATFWPAYNEYMLSVTQFPLFSPV
PFAEKAVVVLFKLCVKILSTYQTDKYPEELIFKSINLMWKLDNQILETCADFIMQSVCRILNDYPGNIQTNLGWK
TLLPMLSLFGRHPETHDLAVDTIIGLLSDGTHLNKFNYPYCIECAFNFVALRTSVVEKNVQMMSLMANSVKLLVH
WYRDFSDPGNSTSSTSITSTSSSSTEDSLKLSGNFILNLFVKLGESLRKTSLARREEVRNQAVASLKRSFEFAED
LDLTSTNCINCFNYIIFAMVDDLHEKMLDYSRRENAEREMRSMEGTLKNAMELLTDVFLQYLKIMADNPAFRTFW
LGLLRRMDTCMKADLGPHGASRMQDVVPQLLKKMIAAMKDAQVLVPKENDDLWEITYIQIQWIAPSIKNELFPDI
A*

**>CargiGNOMa_sc8504**

MGRLKLQSGIKAIeeepeeseLTLSNKAAVSCMINSEIAAVLAIMRRNVRWGGRYMSGDDQLEHSLIQSLKALRK
QVFTCQQPWNNINPVLYLQPFLDVIRSDETGAPITSVALSSVYKILTLDVLGLNTANVEDAMHLVVDAVTSCRFE
VTDPASEEVVLTKILQVLLACMRSKASVVLSNQHVCTIVNTCFRIVHQATTKGELLQRMAAHTMRELIRCIFSHL
PEVENVDQAFVNGNSSVKSEVGGLDNDYNFSAKQLNNGNANSDYDGQLHAVGMGASAVDENGTGNGKEAKPYDLH
LMTEPYGVPCMVEIFHFLCSLLNDLDHFGIGPGSNALAIDEDAPLFALGLINSAIELGGASFVRHPRLLNLIQDE
LFRNLMVFGLSPSPLILSMVCSIVLNLYHHLRMELKLQLEAFFSCVILRLAQSRHGASYPQQEVAMEALVDFCRQ
KTFMVEMYANLDCDITCSNVFEDLANLLSKSAFPVNSPLSSMHILALEGLIAVIQGMAERIGNGSLGLEEASVSL
EEYTPFWMVKCDNYSDPNHWVPFVRRRKHIKKKLMIGADHFNRDPKKGLEFLQGTHLLPEKLDPQSVACFFRYTA
GLDKNLVGDFLGNHDEFCVQVLHEFAQTFDFQDMNLDTALRLFLETFRLPGESQKIQRVLEAFSERYYEQSPQIL
VDKDAALLLSYSLIMLNTDQHNVQVKKKMTEEDFirnnrringgnDLPREFLSELYHSICKNAIRTIPDQGARYP
EMTPSRWIDIMHKCKKSAPYIVADSRAYLDRDMFAIMAGPTIAAISVVFDHAEHEEVYQTCIDGFLAVAKISACH
PLENVLDDLVVCLCKFTTLLSSSAVEEPVLAFGDDTKARMATVTIFTIANHYGDYIHEGWTNILDCILRLHKLGL
LPARVasdaaddselsadPGQGKPLTNSLSAAHLPSMGTPRRSSGLMGRFSQLLSLDTEEPRSQPTEQQLAAHQR
TLQTIQKCHIDSIFTESKFLQAESLLKLAQALRGAAGRPQKGAGSPEDEDTAVFCLELLIAITLNNRDRIVLLWQ
SVYEHIANIVQSTVMPCALVEKAVFGLLRICQRLLPYKENLADELLRSLQLVLKLDARVADAYCEQITQEVSRLV
KANATHIRSQMGWRTITSLLSVTARHPEASDAGFDALVFIMSEGAHLLPSNFILCIDAARQFAESRVGQADRSIR
ALDLMSGSVSCLVKWVRETKEAVREEDAIKMSQDIGDMWLRLVQALRKVCLDQREEVRNHALSSLQKCLTEIEEV
HLAHNLWLQCFDVVIFTMLDDLLEIAQGQSQKDYRNMEGTLMIAMKLLSKVFLRLLTELSHLTTFCKLWLGVLSR
MEKYMKVKIRGKRSDKllelvlellknmllvmknGGVLVQRSALGGDSLWELTWLHVNNISSSLQSEVFPSAGAD
ETTSTPGEAVAAES*

**>CargiGNOMb_sc1597**

MGCLKLQTGINAIeeepeeseVTLPNKAALSCMINSEIAAVLAIMRRNVRWGGRYMDDQSGHSLIQSLKALRKQV
FTWQQPWHSINPVLYLQPFLDVIRSDETGAPITSVALSSVYKILTLRVLDLNTTNVEDTMHLVVDAVTSCRFEVT
DPASEEVVLTKILQVLLSCMRSKASVVLSNQHVCTILHTCFQIVHQATTKGELLQRTAAHTMRELVRCIFSQLPK
VGNVDQALVDGHSSMKSEVGGLDIDYDFLTEQLNTGNANSEYNQLPAVDLGAGAVDENVPGDGKEAKPHDLYLMT
EPYGIPCMVEIFHFLCSLLNDLDHVGIGPGSNALAIDEDAPLFALGLINSAIELGGASFVRHSRLLNLIQDELFH
NLIVFGLSPSPLILSMVCSIVLNLYHHLRMELKLQLEAFFSCVILRMAQSSDEAYDQRQEVAMEALVDFCRQKTF
MVEMFANLDCDITCSNVFEdlanllsknassldsPLSMHIHALDGLIAVIQGIAERVGNGFLGVEQASVSLEEYT
PFWRVKCDNYGDPNQWVPFVLRRKGIKKELMAGAAHFNRDPKKGLELLQRRHLLPGKLDAQSVACFLRYAAGLDK
KVIGDFLGNHDEFCVQVLHEFAQTFDFQDMNLDTALRLFLEAFIPPGESQKIHRVLEAFSERYYEQSPQILVNKD
AAFVLSYSIIMLNTDQHNVQVKKKMTEEDFIRMNRQINGGNDLPREFLSELYHSICNNEIRTIPKQETDFVEMTP
SKWIDLMHKCKNTAPYIVTDSRAYLDHDMFAIIAHPTICSISEVFHHAEDEEVYQTCIDGFLAVAKISACLHLKD
VLNDLVTSLCKSTALLNSFSVEELVLAFGDDTKARMATVTVFTIANRYGDYINSGWRDILACILRLHKLGLLPAR
VTSDAANDSELSADPVQGKPLTNPLSTTHLPFMGTPRRSSGLTARFSQLLSFDTEEPRSQPTEQQLVAHQCTLQT
IQECNIDSIFTESKFlqaesllqlaqalrlaaGRPQKGAGSPEDEDTAVFCLELLIAIMLHNRDRIVLLWQDVYE
HIANIVQSTVMPCALVEKGVFGLLRICQRLLPYKETLADELLRSLHLVLKLDARVADAYCEQITREVSRLVKANA
THIRSQMGWRTITSLLSVTARHPEASDAGFDALVFIMAEGAYLLPSNFILCIDAARQFAESRVGQADRSIHALDL
MSGSVSCLVKWVReakeaageedakkMSQDIGDIWLRLVQALRKVCLDQREEVRNHALSSLQKCLTEIEEVHPAH
NLWFQCFDVVIFTMLDDLLEIAQGQSHKDYSNMEGTLTIAMKLLSKVFLQLLPELSHLTTFCRLWLCVLSRMEKY
MKVKIRGKRSDKLQEFVLELLKNILLVMKNGGVLVQRSAVGGDSLWELTWLHVNNISPSLQSEVFPSAGDDETAS
TLVKQWLQQVDGMIST*

**>CargiGNL2_sc1566**

MTRYIKREIGVSCTINTEVGAVLAALRSSHDVSNPEDHYYQDVSIQQSLKSLRSLIFNPQQEWRYVDPNIYLSPF
LDIIQADDVPTVATGVAlssllkllrlEIFDDKTPGAKEAMNALVTATTNCRLERTDPVSEEGVLMKILQLLTGI
MTHPASCLLRDEAVCTVINTCFQVVQQSATRGNLLQRTAKCTMHEITQMVFSRLNDIEIRSGensesdsedlddd
dedmdsGFGLRCVVDIFHFLCSLLNvvqtvetegvvtqtNDEDIQWFALVLINTALELSGDGFANHPKLLRMIRD
DLFHHLIHYGTRSSAPILSMICSTVLNVYHFLRRSIRLQLEAFFNYVLIRIASFGGLVHFQEVALEGIINLCRQP
NFLIEAYVNYDCAFICNNIFEEIGKLLCKLAFPMGNPLTSLQLQAFEGLVIVIHTIADNINKGKDEISFGPYPVE
ITEYSSFWEELEMDEDGLETWVSELRLRKAQKRKILIAGTHFNQDEKKGYEYLRISHLVSEPPDPKDYAYFFRFT
PKLNKTAIGDYLGDPDEFHLQVLKEFTNTFDFTGMILDNALRTYLDTFRLPGESQKIQRILEAFSDRYYDHQTSD
LFETKDAVLIICYSLIMLNTDQHNPQVKKKMTEEEFIKNNRAINNGKDLPREYLSELFHSIATNAITVFAKAGTT
AEMNPSRWIELIGRSRVVQPFIACEYDHRLGRDMFAGIAGPAVASLSAIFEHADDDDVLNECIEAVVSVARIARH
GLEDILDELIASFCKFTTLLNPYASTEETLYIFANDMKPRMATLAVFTIANKFGEAIRGAWRNIVDCllklkrlk
llPQSVVELEMLSSEGGSTGGSHRRSESGNMPGSGRGTSGRFSNLLLERDEDSLVLGGSEFEQNLKVIQQCRIGN
IFRNSSTFPVETLLNLGRSLIFAAAGKGQKFTTAVEEEETVGFCWDLVAVIGFANIDRFAAFWPAYNEYMISVAQ
FPVFSPIPFAEKAvvvffklcvkllSSYQSDKYPEELIFKSISLMWKFDKEILDTCADFILQSVWTILTEYPGNV
QTQLGWKTLLHLLSVTGRHPETYDQGVEILIQLLSDVTHMSTMNYPYCIDCAFGFVALRSSLIEKNIKIMDLMAN
SVNLLIQWYRSYSDPGnnvsnasissnssTEDNPKFGNLTMNLFVKLGEALRKTSLARREEVRNHAVVSLKKSFQ
LAEGLEFSSTNCINCFNYVIFAMVDDLHEKMLDYSRRENAEREMRSMEGTLRIAMELLTDVFLQYLKIISENPGF
RTFWLGLLRRMDTCMKADLGMHGPSRMQDVVPELLKKMIAAMKDIEILVPKENDDLWEITYIQIQWIAPSIKGEL
FPDVL*

**>Troal_0.675**

MGRLKLQSGIKAIEEEPEECDSLSNRAALACMVNSEVGAVLAVMRRNVRWGGRHMAGDDQLEHSLIQSLKALRKQ
IFSWQHQWNTINPAVYLQPFLDVIRSDETGAPITGVALSSVHRILTLDVLDLNTVNVEDAMHLVVDAVTSCRFEV
TDPASEEVVLMKILQVLLACMKSKASVMLSNQHVCTIINTCFRVVHQAGTKGELLQRIARHTMHELVRCIFSHLP
DIDNTRRALTDGGGSSAKQEVGGLAKDYTFGNKQLENGNSGSEYDGQPPSVGFASNASTCPVASVVDDNTLVAGN
EEEAAPSDLHLMTEPYGIPCMVEIFHFLCSLLNIVEHNGMGPRSNTIAFDEDVPLFALGLINSAIELGGPSICQH
PKLLALIQDELFSNLMQFGLSMSPLILSMVCSIVLNLYHHLGTELKLQLEAFFSCVILRLAQSRHGASYQQQEVA
MEALVDFCRQKTFMAEMYANLDCDITCSNVFEDLANLLSKSAFPVNCPLSAMHILALDGLIAVIQGMAERIGNGL
PVSEQAPVDLEEYTPFWTVKCENYGDPYHWVPFVRRRKYIKRRLMIGADHFNRDPKKGLEFLQGTHLLPDKLDPQ
SVACFFRYTAGLDKNLVGDFLGNHDEFCVQVLHEFAWTFDFQDMNLDTALRLFLETFRLPGESQKIQRVLEAFSE
RYYEQSPQILVNKDAALLLSYSLIMLNTDQHNVQVKKKMTEEDFIRNNRHINGGGDLPRDFLSELYHSICKNEIR
TIPEQGASFPHMTPSRWIDLMRKSKKTAPFIVCDSRAFLDHDMFAIMSGPTIAAISVVFDNAEHEDVFQTCVDGF
LAIAKISACHHLEDVLDDLVVSLCKFTTLLNPSSVEEPVLAFGDDTKARMATITVFTIANRYGDYIRTGWRNILD
CILRLHKLGLLPARVASDAADDSELSADPGHGKPVASSLSASHIPSMGTPRRSSGLMGRFSQLLSLDTEEPRSQP
TEQQLAAHQRTLQTIQKCHVDSIFTESKFLQADSLLQLARALIWAAGRPQKGNSSPEDEDTAVFCLELLIAITLN
NRDRIRLLWQGVYEHISNIVQSTVMPCALVEKAVFGLLRICQRLLPYKEDLADDLLRSLQLVLKLDARVADAYCE
HITQEVMRLVKTNATHIRSQMGWRTITSLLSITARHPEASEAGFDALMFIMSDGVHLSPANYILCVDASRQFAES
RVGQADRSVRALDLMAGSAACLARWSRETKEAVGEEAATKMSQDIGEMWLRLVQGLRKVCLDQREEVRNHALLSL
QKCLTGVDGVHFPHASWLQCFDLVVFTMLDDLLEIAQGHSPKDYRNMEGTLVLAMKLLSKVFLQFLHDLSQLTTF
CKLWLGVLSRMEKYMKAKVRGKRSEKLQELVPELLKNTLLVMKTRGVLVQRSALGGDSLWELTWLHVNNIAPSLQ
SEVFPDQDLEQLQHKQRETGGSPMSDGTGFVPLSETTAEEAKQ*

**>Troal_13.1500**

MGRLKLQSGIKAIEEEPEDCDSTSNRGALACMVNSEVGAVLAVMRRNVRWGGRFMAGDDQLEHSLIQSLKVLRKQ
IFSWQHQWHTIDPAVYLQPFLDVIRSDETGAPITGVALSSVYKILTLDVLDLKTVNVEDAMHLVVDAVTSCRFEV
TDPASEEVVLMKILQVLLACMKSKASVMLSNQHICTIINTCFRVVHQAGTKGELLQRIARHTMNELVRCIFSHLP
DIDNTECALANGVGYSVKQEVGGLAKDYTSGNKQLENGNSSSEYDCQPPSLGFVSNASSDPVASVIDENTIGASP
NDLHLMTEPYGVPCMVEIFHFLCSLLNIVEHNGMDPRSNTIAFDEDVPLFALGLINSAIELGGPSICQHPKLLAL
LQDELFSNLMQFGLSMSPLILSMVCSIVLNLYHHLRTELKLQLEAFFSCVILRLAQSRHGASYQQQEVAMEALVD
FCRQKTFMAEMYANLDCDITCSNVFEDLANLLSKSAFPVNCLLSAMHILALDGLIAVIQGMAERISNGSPVSEQA
PVDLEEYTPFWTVKCENYGDPNHWVPFVRRRKYIKRRLMIGADHFNRDPKKGLEFLQGTHLLPDKLDPQSVACFF
RYTAGLDKNLVGDFLGNHDEFCVQVLHEFAWTFDFQDMNLDTALRLFLETFRLPGESQKIQRVLEAFSERYYEQS
PLILANKDAALLLSYSLILLNTDQHNVQVKKKMTEEDFIRNNRHINGGSDLPRDFLSELYHSICKNEIRTTPEQG
GGVPHMTPSRWIDLMRKSKKTAPFIMCDSRAFLDHDMFAIMSGPTIAAISVVFDHAEHEDVFQTCVDGFLAVAKI
SACHHLEDVLDDLIVSLCKFTTLLNPSSVDEPVLAFGDDTKARMATVTVFTIANRYGDYIRTGWRNILDCILRLH
KLGLLPARVASDAADDSELSVDPGHGKPVASSLSASHLPSMGTPRRSSGLMGRFSQLLSLDTEEPRSQPTEQQLA
AHQRTLQTIQKCHVDSIFTESKFLQADSLLQLARALIWAAGRPQKGNSSPEDEDTAVFCLELLIAITLNNRDRIG
ILWSGVYDHISNIVQSTVMPCALVEKAVFGLLRICQRLLPYKENLADELLRSLQLVLKLDARVADAYCEHITQEV
MRLVKTNAMHIRSQMGWRTITSLLSITARHLEASEAGFDALMFIMSDGVHLLPANYVLCVDASRQFAESRVGQAD
RSVRALDLMSGSVACLARWSRETKEAVGEEAATKVSQDIGEMWLRLVQGLRKVCLDQREEVRNHAILSLQKCLTG
VDGVHLPHALWLQCFDLVVFTMLDDLLEITQGHSPKDFRNMEGTLVLAMKLLSKVFLQLLHDLSQLTTFCKLWLG
VLSRMEKYMKAKVRGKRSEKLQELVPELLKNTLLVMKTMGVLMQRSALGGDSLWELTWLHVNNIAPSLQSEVFPD
QDLEQLQHKQSETSGSPFSDGTGFVPSSEMTAEEGVQAGG*

**>Troal_12.117**

MGSLKLQSGIEADVEETGHCMPREYALACMVNSEIGAVLAVMRRNIRWGGRYLADDDQLEHSLILSLKSLRKQIF
SWDSQWHTVNPSIYLQPFLDVIRSEEIGAPITGVALSSIYKILALDILDPTIVNVKDSVHLLVDAVTSCRFEVTD
PTSEEVVLTKILQVLLACMKSKASVLLNNQHVCTIVNTFFRIVHQSGSKGELLQRIARHTMHELVRCIFSRFPDI
HHTEQPSCNGDGPSIKLEVGGLDQDYTFGVKQSDSVNDSLKNASIAAGSEGSLVGENAVGAGSMEEAVYDMQQVM
REPYGVPCTVEIFQFLCSLLNIVEHMGVGARSNPTEFDEDVPLFALGLINSAIELAGPSIPRHPKLLALIQDELF
CNLMQFGLSMSPLILSMVCSIILNLYLNLRTELKLQLEAFFSCVILRLTQSRHGGASYQQQEVLMEALIDFCRQK
AFMAEMYANFDCDISCSNVFEDLANLLSKSVFPVNSPLSSMHVLALEGLVAVVQGMGERIDSPSPFSEQTSFDLE
EYTPFWMVKCENYGDPNCWVGFLRQRKHIKRTLMTSASYFNRDPKKGMEFLQGTHLLPDKLDPQSVACFFRYTVG
LDKNVVGDFLGSHDEFWVRVLHEFVQTFDFRDMSLDTALRLFLETFRLPGESQKIQRVVEAFSERYYEQSPQILA
NKDAALLLSYSVIMLNTDQHNVQVKKKMTEEDFIRNNRHINGGKDLPREFLLELYHSICKNEIRMTPKQGAGFAE
VTLSRWINLLNKSKNAAPFITCNSRPFLDRDMFAILSGPILAALSVVYDHAEQEDVCQMCVDGFLAVAKISASHH
FDDVLDDLVVSLCKFTTLLSTYHTEEFILAFANDSKAMMATVTVFAIANTFGDFIRTGWRSILDCILSLHKLGLV
PALVASNSTIDNDLSSDPMRGIDVTSSHSVSHVTVRSTTPPRRSSGLMGRFSQLLSLDTEEPRIETTEEEIAAHQ
RIHETIQKCCVSSLFTESKFLLSDSLLQLVQALILVSDPPRKGNYSSPEDIDTSVFCLELLITVTLYNRDRIVLL
WPCVYEHIACTIKSTTTPCVLVEKAVFGLLHICQRLLPYKENLVDELLRSLQLILKLDARIADAYCEHITQEVAD
LVKANAVCIRSQMGWWTITSLLSITARHPEAYEVGFEALAFITSDGAHLLPINYTLCMDALQQFAESRVGPTERS
MHALDLMAGSLACFVRWGLSPQEVGENWLKLLQGLRKVCLDQREEVRNHAVLSLRRCMIGLEGFCFLPATRVQCF
DLVIFTLLDDMLEIAQGQSSRNYRNMEGTLVHALKLSSKMFLQLVGDLSPITSFSKLWLGILGRTENYAKAKIRG
KTSERLRELILELLKNTLLVMKTKGLLMKPSTATGSNGDPLWELTWSHVNNIAPSLKSELFQDQEQELAHLQKGM
ASPAFPDNDTGFLKS*

**>Troal_17.673**

MTKTREHNQLEEEQEDEEEEGDHSSCSQKKKKIKRKELSLSCMLNTEVGAVLAVIRRVPDQNSHFTSPPDENYDS
SVLHSLKSLRALIFNPQQEWRTIDPSLYLSPFLDVIQSDEIPPSATGVALSAILKILKIDIFDEKTPGARDAINS
VVSGITNCRLEKTDPVSEDAVLMRVLQVLIGIMKNRASVLLTDHAVCTIVNSCFQIVQQSASRGDLLQRNARYTM
HELIQTIFSRLPEIEVQHGESSDTDTEDADIASNLESGYGIRCAVDIFHFLCSLLNVVEFVEMEGSTAQTSDEDV
QLFALVLINSAIELSGDGIGKHPKLLRMIQDDLFHHLIHYGACSSSLVLSMICSTVLNIYHFLRRSIRLQLEAFF
AFVLFRVATPGSLTQLQEVALEGIISFCRQPTFIIEMYVNYDCDPICQNVFEEIGKLLCKNAYPMGNPLTSLQIQ
AFEGLIVMIHNIADNIEEDDNSSSTGIHPAEIPEYRPFWVEISKNTDDLDTWVEFIRVRKVQKKKIKIAGNHFNR
DEKKGLEYLKIANIVMDPPDPKAFAFFFHYTPGLDKNMIGDYLGDPDEFNIQVLQEFTETFEFSGLILDTALRTY
LETFRLPGESQKIQRILEAFSEKFYDQQSSEIFVSKDAVFILCYSLIMLNTDQHNPQVKKKMTEEEFIRNNRAIN
GGNDLPREYLSELFLSISNNAITLFAQSGSSIEMNPSRWVDLINRSKVMEPFILCNFDRRIARDMFASIAGPSVA
SLSAIFEHAEDDEILHECVEGLFSIARIAQFGLEDTLDELLASFCKFTTLLNPYASAEETLYAFSNDLKARMATL
SVFTIANKFGDSIRGGWRNIVDCLLKLKRLKLLPQSVVEPDTASTSSSDFLMHSRSESGVIFPSHDPRFGSGRQS
SGLIGRFSHFLSLESAEDSLAHAGSEFEQNLKIIQQCRIGNIFSNSSNMPDDSLLNLGRSLIFAAAGKGQKFSTP
VEEEETVGFCWELLITITLANLHRFQMFWPSFHDYLLVVAQFPLFSPCPFAEKAIVGLFRVCLKLLASYQSEKLS
EELIFKSINLMWKLDKEILDTCCECITQSVSKILTEYPANLQSQLGWKSVLHLLSVTGRHPETYDQGVEALIMLM
SDGTHVSRTNYAYCIDCAFGYAALKNSPLEKSSKILDLMADSVNLLIQWYRNGYSDSGSNASTMSNTSNSSLEDG
PKALNSFNFAMNLFIKLAEALRKSSLVRREEIRNHAVLALQRSFTLADELDFTSTNCINCFNLVIFAMVDDLHEK
MVEYSRRENAEREMRSMGGTLKVAMELLTEMFLQFLKQISQSSGFKTFWLGVLRRMDTCMKADLGDYGSSRLQEV
VPELLKKMIMAMKEKEILVQKEGDDLWEFTYIQIQWIAPSLKEELFPEEKF*

**>Tetsi_007283**

MGRLKLQSGIKAIEEEPENCDFTSNGGALACMVNSEVGAVLAVMRRNVRWGGRFIAGDDQLEHSLIQSLKALRKQ
IFSWRHHWHMIDPAVYLQPFLEVIRSDETGAPITGVALSSVYKILTLDVLDLKTVNVEDAMHLVVDAVTSCRFEV
TDPASEEVVLMKILQVLLACMKSKASVMLSNQHICTIINTCFRVVHQAGTKGELLQRIARHTMHELVRCVFSHLP
DIDNTECALANGGGYSVKQEVDGLAKDYTSGNKQLDNGNSSSEYDCQPPSLGFASNASSDPVASEIDENTIRASP
NDLHLMTEPYGVPCMVEIFHFLCSLLNIVEHIGMDPRSNTIAFDEDVPLFALGLINSAIELGGPSICQHPRLLAL
IQDELFSNLMQFGLSMSPLILSMVCSIVLNLYQHLRTELKLQLEAFFSCVILRLAQSRHGASYQQQEVAMEALVD
FCRQKTFVAEMYANLDCDITCSNVFEDLANLLSKSAFPVNCLLSAMHILALDGLIAVIQGMAERISNRSPIPEQA
PIDLEEYIPFWSVKCENYGDPDHWVPFVLRKKYIKRRLMIGADHFNRDPKKGLEFLQGTHLLPDKLDPQSVACFF
RYTAGLDKNLVGDFLGNHDEFCVQVLHEFAGTFDFQDMNLDTALRLFLETFRLPGESQKIQRVLEAFSERYYEQS
PQILVNKDAALLLSYSLIMLNTDQHNVQVKKKMTEEDFIRNNRHINGGSDLPRDFLSELYHSICKNEIRTTPEQG
GGFPHMTPSRWIDLMRKSKKTAPFIMCDSRAFLDHDMFAIMSGPTIAAISVVFDHSEHEDVFQTCVDGFLAVAKI
SACHHLEDVLDDLVVSLCKFTTLLNPSSAEEPVLAFGDDIKARMATITVFTIANAYGDYIRTGWRNILDCIIRLH
KLGLLSARVASDAADDSELSADPWHGKPVASSLSASHMPSMGTPRRSSGLMGRFSQLLSLDTEEPRSQPTEQQLA
AHQRTLQTIQKCHVDSIFTESKFLQADSLLQLARALTWAAGRPQKGNSSPEDEDTAVFCLELLIAITLNNRDRIG
ILWPGVYEHISNIVQSTVMPCALVEKAVFGLLRICQRLLPYKENLADELLRLLQLVLKLDARVADAYCEHITQEV
MRLVKTNAMHIRSQMGWRTITSLLSITARHPEASEAGFDALMFIMSDGAHLLPANYILCVDASRQFAESRVGQAD
RSVCALDLMAGSVTFLARRSREMTEAVGEEAATKITQDIGEMWLRLVQGLRKVCLDQREDVRNHAILSLQKCLTG
VDGLHLPHASWLQCFDQVIFTMLDDLIEIAQGHSPKDYRNMEGTLVLAMKLMSKVFLQLLHDLSQLTTFCKLWLG
VLSRMEKYIKAKVRGKRSEKLQELVPELLKNTLLVMKTRGVLVQRSALGGDSLWELTWLHVNNIGPSLQSEVFSD
QDLEQLQHKQSQTGGSTFSDGTGFVPSSEMTAEEGVQAGG*

**>Tetsi_000870**

MGSLKMQSGIEADVVEHGHCMPRECALACMVNSEIGAVLAVMRRNIRWGGRYMADDEQLEHSLILSLKSLRKQIF
SWDNQWHTVNPSIYLQPFLDVIRSEEIGAPITGVALSSIYKILALDILDPTTVNVKDSVHSVVDAVTSCRFEVTD
PTSEEVVLTKILQVLLACMKSKASVLLNNQHVCTIVNTFFRIVHQSGSKGELLQRIARHTMHELVRCIFSRLPDF
HHTEQPPSNGDGPSIKLEVGGLDQDYSFGVKQSDSANDSMKNASIAAGSEGSLVGENAVGAGRMEEAVNDMQQIM
REPYGVPCTVEIFQFLCSLLNIVEHTGMGPRSNPAEFDEDVPLFALGLINSAIELAGPYIPQHPKLLALIQDELF
CNLMQFGLSMSPLILSMVCSIILNLYLNLRTELKLQLEAFFSCVIFRLTQSRHGGASYQQQEVLMEALVDFCRQK
PFMSEMYANFDSDISCSNVFEDLANLLSKSVFPVNSPLSSMNVLALEGLIAVVQGMCERIDSPSPFSEQTSFNIE
EYTAFWTVKCENYGDPNCWVGFLRQRKLIKKTLMTSANYFNSDPKKGMEFLQGTHLLPDKLDPQSIACFFRYTAG
LDKNVIGDFLGSHDEFWVRVLHEFVQTFDFRDMSLDTALRILLETFRLPGESQKIQRVVEAFSERYYDQSPHILA
NKDAALLLSYSVILLNTDQHNVRVKKKMTEEDFIRNNRHINGGKDLPREFLLELYHSICKNEIRMTPKQGAGFSE
VTLNHWIYLFYKSKDAAPYITCNSRPFLDRDMFAILSGPTLAALSVVYDDTEQEDVCQMCVDGFLAVAKISASYH
FVDVLDDLVVSLCKFTTLLSTYHTEEFTLAFANDSKAMMATETVFSIANTFGDYIRAGWRSILDCILSLHKLGLV
PALVASNSTIDTDLSSDPMPGISVTSSQSVSHVPVRSTPPRRSSGLMGRFSQLLSLDTEEQSIEPTEEEIAAHQR
THETIQKCCVSSLFTESKFLQSDSLLQFVQALILVSDPPHKGKYSSPDDVDTSVFCLELLITITLYNRDRIVLIW
PCVYEHIAGTVKSTSTPCILVEKAVFGLLHICQRLLPYKENLADELLRSLQLILKLDARIADAYCEHITQEVADL
VKANAVCIRSQMGWRTIISLLSITARHPEASEVGFEALAFITSDGAHLLPTNYTLCMDALQQFAESRVGPTERSM
HSLDLMAGSLTCFTRWGLSPQELGENWLKLVQGLRKVCLDQREEIRNHAVLSLRRCTTGLEGFCFLPAMWVQCFD
IVIFTLLDEMLDIAQGQSSRNYRNMEGTLVNALKLSSNMFLQLVGDFFPIASFSKLWLGILVRMENYLKAKIRGR
ASERLQELIPELLKNTLLVMKAKGLLVKPSSVTGSNGDPLWELTWSHVNNIDPSLKSELFQDQEQELAHLQKETA
SPAFPDNDTSLIKL*

**>Tetsi_015395**

MGRLKLQSGIKSIEEEPEECDSTSNRVALACMVNSEVGAVLAVMRRNVRWGGRHMAGDDQLEHSLIQSLKALRKQ
IFSWQHQWHTINPAVYLQPFLDVIRSDETGAPITGVALSSVYKILTLDVLDLNTVNVEDAMHLVVDAVTSCRFEV
TDPASEEVVLMKILQVLLACMKSKASVMLSNQHVCTIINTCFRVVHQAGTKGELLQRIARHTMHELVRCIFSHLP
DIDNMGRALTDGGMSSAQQEVGGLAKDYTFGNKQLENGNSGSEYDGQTPSVGFASNASTYPVANVVEENTLVASN
GNEATPSDLHLMTEPYGIPCMVEIFHFLCSLLNIVEHTGMGPRSNTIAFDEDVPLFALGLINSAIELGGPSICQH
PKLLALIQDELFSNLMQFGLSMSPLILSMVCSIVLNLYHHLRTELKLQLEAFFSCVILRLAQSRHGASYQQQEVA
MEALVDFCRQKTFMAEMYANLDCDITCSNVFEELANLLSKSAFPVNCPLSAMHILALDGLIAVIQGMAERIGNGL
PVSEQAPVDLEECTPFWTVKCENYGDPFHWVPFVRRRKYIKRRLMIGADHFNRDPKKGLEFLQGTHLLPDKLDPQ
SVACFFRYTAGLDKNLVGDFLGNHDDFCVQVLHEFAGTFDFQDMNLDTALRLFLETFRLPGESQKIQRVLEAFSE
RYYEQSPQILVNKDAALLLSYSIIMLNTDQHNVQVKKKMTEEDFIRNNRHINGGNDLPRDFLTELYHSICKNEIR
TTPEQGASFPQMTPSRWIDLMRKSKKTAPFIVCDSRAFLDHDMFAIMSGPTIAAISVVFDNAEHEDVFQTCVDGF
LAVAKISACHHLEDVLDDLVVSLCKFTTLLNPSSVEEPVQAFGDDTKARMATITVFTIANRYGDYIRTGWRNILD
CILRLHKLGLLPARVASDAADDSELSTDPGHGKPVASSLSASHIPSMGTPRRSSGLMGRFSQLLSLDTEEPRSQP
TEQQLAAHQRTLQTIQKCHVDSIFTESKFLQADSLLQLARALIWAAGRPQKGNSPEDEDTAVFCLELLIAITLNN
RDRIRLLWQGVYDHISNIVQSTVMPCALVEKAVFGLLRICQRLLPYKEDLADDLLRSLQLVLKLDARVADAYCEH
ITQEVMRLVKTNATHIRSQMGWRTITSLLSHTARHPEASEAGFDALMFIMSDGAHLSPANYILCVDASRQFAESR
VGQADRSVRALDLMAGSVACLARWSRETKEAVGEEAATKMLQDIGEMWLRLVQGLRKVCLDQREEVRNHALLSLQ
KCLIGVDGIHLPHASWLQCFDLVVFTMLDDLLEIAQGHSLKDYRNMEGTLVFAMKLLSKVFLQFLHDLSQLTTFC
KLWLGVLSRMEKYMKAKVRGKRSEKLQELVPELLKNTLLVMKTRGVLLQRSALGGDSLWELTWLHVNNIAPSLQS
EVFPDQDLEQLQHKQRETGGSPVSDGTGFVPETTAEEGVLAGL*

>**Tetsi_012649**

MAKANQREEEEEEEECDHSSCPQKKKKITILKRKELGLSCMLNTEVGAVLAVIRRVPDLNSHLSPSHEENHDSSV
LHSLKSLRALIFNPQQEWRSIDPSIYLSPFLDVIQSDDIPASATGVALSAILKILKLEIFDEKTPGARDAINSVV
NGITSCRLEKTDPVSEDAVLMRVLQVLIEIMKNSASVMLTDHAVCIVVNSCFQIVQQSASRGDLLQRNARYAMHE
LIQTIFSRLPDIEVQHGENSETDTEDADIVSNLDSGYGIRCVVDIFHFLCSLLNVVEFVEMDGSTSQTSDEDVQL
FALVLINSAIELSGDGIGKHPKLLRMIQDDLFHHLIHYGTCSSSLVLSMICSTVLNIYHFLRRSIRLQLEAFFTF
VLFRVATPGILTQLQEVALEGIISFCRQPTFIIEMYVNYDCDPVCQNVFEEIGKLLCKNAFLTGNPLTSLQIQAF
EGLIVMIHNIADRIDEDDNSSSISIHPAQIPEYRPFWVEISKNTDDLETWVEFIRVRKAQKRKIKIAGNHFNRDE
KKGLEYLKIANIVPDPPDSKAFAFFFRYTPGLDKIMIGDYLGDPDEFNIQVLKEFTETFEFSGLILDTALRTYLE
TFRLPGESQKIQRILEAFSERFYDQQSSEIFVSKDAVFIICYSLIMLNTDQHNPQVKKKMTEEEFIRNNRAINGG
NDLPREYLSELFHSISNKAITLFGQSGSPIEMNPSRWVDLINRSKVMEPFILCNFDGRIARDMFASIAGPSVASL
SAIFEHAEDDEILHECIEGLFSIARIAQFGLEDTLDELLASFCKFTTLLNPYATAEETLYAFSNDLKARMATLSV
FTIANKFGDSIRGGWRNIVDCLLKLKRLKLLPQSVVEPDTASTSSSDFLLHSRSESGVIFPSHDPRFGSGRQSSG
LIGRFSHFLSLESTEDSLAHAGSEFEQNLKIIQQCRIGNIFSNSSNMPDDSLLNLGRSLIFAAAGKGQKFSTPVE
EEETVGFCWELLITITLANLHRFQLFWPSFHDYLLVVVQFPLFSPCPFAEKAIVVLFRVCLKLLASYQSQKLSEE
LIFKSINLMWKLDKEILDTCCECITQSVSKILTEYPANLQSQLGWKSVLHLLSVTGRHPETYDQGVEALIMLMSD
GTHVSRTNYAYCIDCAFGYAALKNSPLEKSSKILDLMADSVNLLIQWYKNGFSDPGSNASNMSNSSNSSLEDGSK
SHNSFNFTMNLFIKLAEALRKSSLVRREEIRNHAVLALQRSFTLADELEFTSSNCINCFNLVIFAMVDDLHEKMI
EYSRRENADREMRSMGGTLKVAMELLTEMFLQFLKQISQSSGFRTFWLGILRRMDTCMKADLGDYGSSILQEVVP
ELLKKMIMAMKEKEILVQKEGDDLWEFTYIQIQWIAPSLKEELFPEEEF*

**>Nelnu_LOC104605370**

MGRLKLQSGIKAIEEEPEDYDSTSNRGALACMVNSEVGAVLAVMRRNVRWGGRYMAGDDQLEHSLIQSLKTLRKQ
IFSWQHQWNTMNPAVYLQPFLDVIRSDETGAPITGVALSSVYKILTLDVIDLNTTNVEDAMHLVVDAVTSCRFEV
TDPASEEVVLMKILQVLLACMKSKASVVLSNQHVCTIVNTCFRVVHQAGTKGELLQRIARHTMHELVRCIFLHQP
DIDNTDRSLSRGMSSVKQEVCGPEKDHSFGVKQLDNGNGSSESDGQLPSVGFASNSSMGLVGTLMDENTIGVVNG
KDASPNDLNLMTEPYGVPCMVEIFHFLCSLLNIAEHIGMGPRSNSMAFDEDVPLFSLGLINSAIELGGPSIRKHP
KLLSLIQDELFRNLMQFGLSMSPLILSMVCSIVLNLYLHLRSELKLQLEAFFSCVILRLAQSRHGASYQQQEVVM
EALVDFCRQKTFMAEMYANFDCDITCSNVFEDLANLLSRSAFPVNCPLSAMHILALDGLIAVIQGMAERIGNGSP
VVQASSELEEYTPFWTVKCENYADPNHWVPFVRRRKYIKKRLMMGADHFNRDPKKGLEFLQGTHLLPDKLDPQSV
ACFFRYTAGLDKNLVGDFLGNHDEFCVQVLHEFAGTFDFQDMNLDTALRLFLETFRLPGESQKIQRVLEAFSERY
YEQSPQILANKDAALLLSYSLIMLNTDQHNVQVKKKMTEEDFIRNNRHINGGNDLPREFLSELYHSICKNEIRTT
PEQGAGFPEMTPSRWVDLMRKSSKTSPFIMCDSRAFLDHDMFAIMSGPTIAAISVVFDHAEHEEVFQTCVDGFLA
VAKISACHHLEDVLDDLVVSLCKFTTLLNPSSVEEPVLAFGDDTKARMATVTVFTIANRYGDYIRTGWRNILDCI
LRLHKLGLLPARVASDAADDLELCSDPSQGKPVTSSLSASQMPSMGTPRRSSGLMGRFSQLLSLDTEEPRSKPTE
QQLEAHQRTLQTIQKCHIDSIFTESKFLQADSLSYLARALIWAAGRPQKGNNSPEDEDTAVFCLELLIAITLNNR
DRIILLWQLVYEHISNIVQSTMMPSALVEKAVFGLLRICQRLLPYKENLADELLRSLQLVLKLDARVADAYCEQI
TQEVMRLVKANATHIRSQIGWRTITSLLSITARHPDASEAGFEALIFIMYDGAHLSPANYILCIDASRQFAESRV
GQAERSIRALDLMAGSFACLAQWSRETKETAGEGAAKLFHDIGEMWLRLVQGLRKVCLDPREEVRNHAILSLQRC
LTGVEGLSLSLSLWSQCFDLVIFTMLDDLLEIAQGHSPKDYRNMEGTLILAMKLLAKVFLQLLHDLSQDKTFCKL
WLGVLNHMEKYMKAKVRGKRSEKLQELVPELLKNVLLVMKTKEILVHRSALGGDGLWELTWLNVNKIAPSLQSEV
FPGQELEQVRHKHNDAGGSTASDGTGFIPSSEALTTSESTRTGG*

**>Nelnu_LOC104612314**

MGRLKLQSGIKAIDEEPEECNSSSSRGASACMVNSEVGAVLSVMRRNVRWGGRYMLGDDQLEHSLIQSLKALRKK
IFSWHHEWNTMNPAVYLQPFLDVIRSDETGAPITGVALSSVYKILTLDVLDLNTTNVKDAMHLVVDAVTSCRFEV
TDPSSEEVVLMKILQVLLACMKSKASVVLSNQHVCTIVNTCFRIVHQAGSKGELLQRIARHTMHELVRCIFSHLP
DIENMEHLPSRGIYSVKQEVGGADKDYNFGGKQLDNGNGSSEYDSQLPSVGFSSNASTGLMGSLMDENTIGASNG
KDTASNDLMTEPYGVPCMVEIFHFLCSLLNIAEHIGMGPRSNSMAFDEDVPLFALGLINSAVELGGPSIRKHPKL
LALMQDELFRNLMQFGLSMSPLILSMVCSIVLNLYLHLRSELKLQLEAFFSCVILRLAQNRHGASYQQQEVAMEA
LVDFCRQKAFMAEMYANLDCDITCSNVFEDLANLLSKSAFPVNCPLSAMHILALDGLIAVIQGMAERIGNGSSVN
EHASSDLEEYTPFWTVKCDNYEDSNHWVPFVRRRKYIKRRLMMGADHFNRDPKKGLEFLQGTHLLPDKLDPQSVA
CFFRYTAGLDKNLVGDFLGNHDEFCVQVLHEFAGTFDFQDMNLDTALRLFLETFRLPGESQKIQRVLEAFSERYY
EQSPQILANKDAALLLSYSLIMLNTDQHNVQVKKKMTEEDFIRNNRHINGGNDLPREFLSELYHSICKNEIRTTP
EQGAGFPEMTPSRWVDLMRKSRKTAPFIVCDSRAFLDHDMFAVMSGPTIAAISVVFDHAEHEEVFQTCVDGFLAV
AKISACHHLEDVLDDLVVSLCKFTTLLNPSSGEEPVVAFGDDMKARMATVTVFTITNRYGDYIRTGWRNILDCIL
RLHKLGLLPARVANDAVDDSEFCADPGQGKPVTSSLSASQIPSMGTPRRSSGLMGRFSQLLSLDTEEPRSKPTEQ
QLEAHQRTLQTIQKCHIDSIFTESKFLQAESLLDLARALIWAAGRPHKGNTSPEDEDTAVFCLELLIAITLNNRD
RITLLWQFVYDHISNIVQTTVMPCALVEKAVSGLLRICQRLLPYKENLVDDLLRSLQLVLKLDARVADAYCEQIT
QEVMRLVKANATHIRSQMGWRTITSLLSITARHPEASEAGFEALTFVMYDGAHLIPTNYILCVDASRQFAESRVG
QVDRSLRALDLMAGSLTCLLQWSQKTKEAVGEAATKLSHDIGEMWLRLVQGLRKVCLDPREEVRNHAILSLQRCL
TGAEGLCLSRSLWLQCFDVVIFTMLDDLLEIAQGHSPKDYRNMEGTLALAMKLLAKVFLQLLHLLSQENTFCKLW
LGVLNHMEKYAKAKVRGKKSEKLQELVPELLKNTLLVMKTREILVQRSALGGDSLWELTWLNVNKIAPSLQSEIF
PGQELEQVHHKHNDTASSTPSDPSSEALEASESARTVGSQQVLMRWAT*

**>Nelnu_LOC104585650**

MARTRDPTQEAEDEEEQDRLSYRQRRKKPARVERKELGLSCMLNTEINAVLAVLRRITDSNSLDENYDSSLFNSL
KSLRALLFKPKQEWRLIDPSIYLSPFLDVIQSDDIPIQATGIALLTILKILNSDIFDEKTPGAGDAINAVVIAVT
NCRIEMIDTTSEDIILTRVLQLLSGIMKNRISFLLTDHAVCTIVNTCFQVVQKSASRGDLLQRNARHTMYELIQT
IFNRLQELDVRDGKNESNVNDADFGYDLDYSGYGARSATDIFHFLCSLLNVVEVMDSEGLTTPNSEDNFQLFAFV
LINSTIELSGEVIPKHPKILRMIQENLFHSLIHYGTGSSPLFLSIICSTVLNIYHFLRRYIRFQLEAFFTFVLLR
VASEETLPQLQEVALEGIISFCRHPKFIIEMYVNYDCDPLCRNVFEEVGKLLCKHAFPQGSSLNGLQIQAIEGLI
IMIQNIADHIVDDESSTSAGSYPVEVTEYIPFWVEKSNRNEDLDTWIEYMRVRKAQKRKIMIAANHFNRDEKKGL
EYLKVAHLVSEPPDPKAFAYFYRYTPGLDKNTIGDYLGDPDEFHLEVLKEFTGTFDFSGMILDTALRTFLETFRL
PGESQKIHRILEAFSEKFYDQQSSELFVSKDAVFILCYSLIMLNTDQHNPQVKKKMTEEEFIRNNRAINGGKDLP
REYLSDLFQSIATNEITLFGQSGSSGEMDLKWSDIINRAKVMEPFILSNFDRRLGRDMFATIAGPAVASICATFE
QSDEDEILHECVEGIFAVARIAQFGLGDVLDELIASFCKFTTLLNPYASAEETLYAFSNDLKPRMATLAVFTIGN
KFGNSIRGGWRNIVECLLKLKRLKLLPQSVVEHNITSSTDYATNAKIDSGVILPPSDNTISTTGLITRFSHFLSL
ECSEDSLTLPGCEFEQNLKVIQQCRIGNIFSGSSSLPEESLQNLGRSLIFAATGKGQKFSTPVEEEETVEFCWDL
ITTITLANIHRFLVFWPSFHDHLIVVAQFPLFSPCPFAEKAIAGLMKVCLKLLSSYQSEKLSEELIFKSIALMIK
LDKEILETCCENITHVVSRIIIEFPATVQTPVGWKTILHMLSYAGRLPETYDQGIEALITLMSDETHMSRTNYAS
CIECAFGFACLKIIPLEKSWKILDLMASSVSLLVQWCRSGYSDPGSGLSSSSNSSLEDNMKPLNSFNFTMNLFLR
VADSLRKTSLVRREEIRNHAVLSLQKSFTFAEDLGFTPANCTSCFNLIIFAMVDDLHEKLIEYTMRDNSEKEVRS
MVGTLKLAMELLTEVFLQFLKPISLTSGFRTFWLGVLRRMDTCMKADLGNIVETRLQVMVPDLLRKIITTMKEKE
ILVQRDGDDLWEITYIQIQWIAPSLKEELFPEEDF*

**>Mint05g10540**

MGRLRLQTGIKAIEEEPEEFDCSSNGGALACMVNSEVGAVLAVMRRNVRWGGRYMTGDDQLEHSLIQSLKALRKQ
IFSWQNQWQSINPAAYLQPFLDVIRSDETGAPITSVALSSVYKILTLDILHLNTVNVEHAMPLVVDAVTSCRFEV
TDPASEEVVLMKILQVFLACMKSKASVVLSNQHVCNIVNTCFRIVHQAGSKGELLQRIARHTMHELVRCIFSHLP
EIDKTERLLANGGRYVKQEVGGLEKDYTFSSKQLDNGNDSSEHDGQMPSVGLASNASMGLRVSLTEENTFGAGNG
KEASPTDLHLMTEPYGVPCMVEILHFLCALLDNIEHIGMGPRSNSMASHEDVPLFALGLINSAIELGGPSIRQHP
KLLALIQDELFRNLMQFGLSMSPLILSIVCGIVLNLYYHLRSELKLQLEAFFSCVILRLAQSRHGASYHQQEVAM
EALVDFCRQKTFMAEMYANFDCDITCSNAFEDLANLLSKSAFPVNYPLSAMHILALDGLIAVIQGMAERISNGPP
VLEQVPVDLEEYTPFWTVKCENYADPNHWVPFVRRRKYIKRRLMIGADHFNRDPKKGLEFLQGTHLLPDKLDPQS
VACFFRYTAGLDKNLVGDFLGNHDEFCVQVLHEFAGTFDFEDMNLDTALRLFLETFRLPGESQKIQRVLEAFSER
YYEQSPQILANKDAALVLSYSLIMLNTDQHNVQVKKKMTEEDFIRNNRHINGGNDLPREFLSELYHSICKNEIRT
TPEQGAGFSEMTPSRWIDLMRKSKKTAPFVMCDSRPFLDHDMFAIMSGPTIAAISVVFDHAESEDVFQTCVDGFL
AVAKISACHHLEDVLDDLVVSLCKFTTLINPSSVEEPVLAFGDDTKARMATVTVFTIANKYGDYIRTGWRNILDC
ILRLHKLGLLPARVASDAADDLEVAADPCHGKPVTSSLSASQLPSMGTPRRSSGLMGRFSQLLSLDTEEPRSQPT
EQQLAAHQRTLQTIQKCHIDSIFTESKFLQADSLLHLARALIWAAGRSQKGNSSPDDEDTAVFCLELLIAITLNN
RDRIGLLWQGVYEHISNIVQSTVMPCALVERAVFGLLRICQRLLPYKENLADELLRSLQLVSKLDARVADAYCEN
ITQEVMRLVKANATHIRSQMGWRTITNILSITARHPEASEAGFEALMFIMSDGAHLSPANYILCVDTTRQFAESR
VGQAERSIRALDLMAGSVTCLGRWSCETKEAAGEEAAAKMSQDIGEMWLRLVQGLRKVCFDQREDVRNHALSSLQ
KCLTGVGGFALLHTSWLQCFDQVIFTMLDELLELAQGHSPKDYRNIEGTLILAMKLLSKVFLQLLNDLVQLTTFC
KLWLGVLSRMEKYMKAKIRGKKSEKLQELVPELLKNTLLVMKTRGVLVCRSALGGDSLWELTWLHVNNIAPSLQS
EVFPNQELEQLHKQSETEGVPVSDGSGSVPSGETLTASEEGGQTLVC*

**>MintSC289g00040**

MEIRSLKLQYGVEGDKEKPGHCIPRGCALACMVNSEIGAVLAVVRRNVRWGGRYMADDDQLEHSLVQSLKILRKQ
IFLWDNQWHTVNPSIYLRPFLDVIRSDEIGAPITGVALSSVYKILTLGILDPNTINVGEAMHLIVDAVTGCRFEV
TDPASEEVVLMKILQVLLACMKCKASILLNNQHVCTIVYTCFRIVHQAGTKGELLQRIARTTMHELIRCIFSRLP
YIHVEKPSDGGGPSSQTEVGGVHSEYTFEVKQSASANGSFKHASISVGSEWSLPAASAASDGSIDETVNDMQQIM
REPYGVPCMVEVFHFLCSLLNIAEQMEMSPRSNQTEFDEDVPLFALGLINSAIELAGSTIKHHPKLLALIQDDLF
HNLMQFGMSLSPLILSMVCNIILNLYLHLYDELKLQLEAFFSCLTWRLTHSRHGSASYQQHEVLMEALVDFCRQK
TFMAEMYTNFDCDISFSNVFEDLANLLSKSSFPVNSPLSSMHILALEGLLAVVQGLAERTGNALPIKDQTLFGLA
DYTPFWTVKCENHTDPNCWVEFVRQKKHLKRVLMTSADHFNRDPKKGLEFLQGMHLLPDKLDPSSVACFLRYTPG
LDKNVVGDLLGSHDEFWVQVLHEFARTFNFQDRSLDTALRLFLESFRLPGESQKIQRVLEAFSERYYEQSPHILA
NKDAALLLSYSIIMLNTDRHNVQVKKKMTEDDFIRNNRRINGGNDLPREFLSELYHSICKNEIRMTPGQGADFAE
VTSSRWINLLHKSQGATPFITYDARPFLDHDMFAILSGPTLAALSVVYDHAEHEDVYQMCNDGFLAVAKISASHH
FDEVLDDLVVSLCKFTTLLNPSQPEELILTFTDDAKAMMATVTVFTIANTYGDSIHTGWRSILECILTLHKLGLV
PALVGRDSTGENELFSDSMPGKPVKNSISVSHVPAKSVITVQRSSGLMSRFSRLLSFDAEEPSLEPSDEEVEARQ
RAHETIKKCRINGIFTESKFLQSNSLLHLVQALIWVSGKSQKGKYLSHEDEDASVFCLELLITITLYNRDRIVLL
WPSVHEHIAGIVQSTAMPSALVERAVFGLLHICRRLLPYKENLGDDLLRSLQLVLKLDARVADAYCEHITQEVTD
LVKTNAGCIRSQMGWRTITSLLSITARHPEASEVGFEALAFVMYDGAHLSPDNCILFIDASRQFAESRVGPSERS
IHALDLMAGCLTCLKKWEVSPQEIGEIWMRLVRGLRKVSLDRREEVRNHAVLSLQRCTTGCEGFPIPPLIWMQCF
DEVIFTMLDDMLEVTQGQSSKDYRNMEGTLIHAVKLSSKVFLQLVNDLSPTESFSRVWLGMLGRMEKYTKVKIRG
KKSEKLQELIPELLKNTLLVMKTKGVLVKPSSATGSDSLWELTWPHVSNIASSLKSQVFPDQEPDNQWKETANSA
SPESGTGFIES*

**>Mint05g17010**

MAKARDQNEGEEEEENNSSSCGPKVKKLTKRKKKELGISCMLNTEVGAVIAVMRRSSNTTTDPYSQFVSPSEEIY
DSTLLYSLKSLRSLIFNPQREWRTVDPSIYLSPFLDVIQSDEIPASATGVALSAILKILKLEIFDEKTPGAKEAI
NSIITGITNCRLEITDPASEDAVLMRVLQVLAGIVKNRASVLLNDHSVCTIVNTCFQVVQKSASRGDLLQRNARH
TMHELIQTIFARLPEIDQVREDGEQSTDQTGFDDYDGNLDSGGYGARCAIDIFHFLCSLLNVVEITETDGSSTQN
SDEDVQLFALVLINSAIELSGDGIGNHPKLLRLIQDDLFYHLIYYGTRSSPLVLSMICSTVLNIYHFLRRSIRLQ
LEAFFVFVLFRVASWGNSPQLQEVALEGIISFCRQPTFIVEMYVNYDCDPLCRNIFEDIGRLLCKQAFPLSSPLT
SLQIQAFQGLIVIVHNLADNVEEENDYSSSKGSYPLEISEYKPFWIEKCDNGDDLESWIKFVRLRKAQKKKILIA
GNHFNRDEKKGLEYLKITNLVSDPPEPKSLAYFYRYTPGLDKIAIGDYLGDPGELHIQVLKEFTATFDFSGMILD
TALRTFLETFRLPGESQKIHRILEEFSEKFYDQQSSEIFVSKDAVFILCYSLIMLNTDQHNPQVKKKMTEEEFIR
NNRAINGGKDLPREYLCDLFQSIANNAITLFGQSGSSLEIDPRRWIELISKAKVMEAFILCDFECQLATDMFASI
AGTSVATLAAIFEHTDEDDILHECVEGMFSVAKISQFGLEDTLDELLASFCKFTTLLNPYASAEETLYAFSNDLK
PRMATLAAFTIGNKFGKSIRGGWRNIVDCLLKLKRLKLLPQSVSEPNIASTSSTDHPTHSKSESVVLFPPVPSDH
VYGSSRQSSGMIGRFSQFLSLESSEDSMVHSGSEFEHNLKIIQQCQIENILNNSTNLSEESLLNLGRSLIFAAAG
KGQKFSTPVEEEETVGFCWDLITTITLANIHRFSTFWPFFHDYQLLVAQFPLFSPCPFAEKAIVGLFKVCLKLLA
SCQSEKISEELIFKSINLMWKLDKEILDSCCELIVQALGKIFTEYPANLQTQTGWKSALHLLSVAGRHPETNDQG
VEALIMLMSDGIHVSRMNYAYIMDCAFGYAALKIIPLDKSWQLIDLMSDSTNLLVQWFKNGYSDPGSNISTTSNS
SEDGSKTFNSSSFTTNLFVKLADALRKSSLVRREEIRNHAVSALRRSFKLADELGFTSSNCINVFNLVIFAMVDD
LHEKMVEYSRRENAEREMRSMEGTFKMAMELLTEVFLQFLEPLSQNSGFRTFWQGVLRRMDTSMKVNLGDNQSRL
QELVPVLLKKMIIEMKEKEILVRQEGNDLWEITYVQIQWIVPSLKEELFPEEEL*

**>Aquca_014_00263**

MGRLKIQPGINAIEEEPEDCEHTPTKGALACMVNSEIGAVLAVMRRNVRWGGRYMAGDDQLEHSLIQSLKSLRKQ

IFSWQHQWNTINPVLYLQPFLDVVRSDETGAPITGVALSSVYKILSLDVIDLDTVNVEDAMHLVVDAVTSCRFEV
TDPASEEVVLMKILQVLLACMKSKAAIVLSNQHVCTIVNTCFRVVHQAGAKGELLQRIARHTMHELVRCIFSHLP
NVDNREGSLANGNASASKQEFGGVHKDAGSKQLDNGNSSSEFDGQPPSFGFASTAYANSVESITTENTIGTINDK
EASPDDLLLMTEPYGVPCMVEIFHFLCSLLNIIEHIGMGPRSNTIAFDEDVPLFALGLINSAIELGGPSISQHPK
LLALIQDELFRNLMQFGLSMSPLILSMVCSIVLNLYHHLRTELKLQLEAFFSCVVLRLAQSRLGASYQQQEVAME
ALVDFCRQKTFMVEMYANLDCDITCSNVFEDLANLLSKSAFPVNCPLSAMHILSLDGLIAVIQGMAERIGNESHV
SEQGPVDFEEYTPFWTVKCENYADTNHWVPFVRRRKYIKRRLMIGADHFNRDPKKGLEFLQGTHLLPDKLDPQSV
ACFFRYTAGLDKNLVGDFLGNHDEFCVQVLHEFAGTFDFQDMNLDTALRLFLETFRLPGESQKIQRVLEAFSERY
YVQSPHILVNKDAALLLSYSIIMLNTDQHNVQVKKKMTEEDFIRNNRHINGGSDLPREFLSELYHSICKNEIRTI
PEQGVGFPEMTPSRWVDLMRKSKKTSPFIVSDSRAFLDHDMFAIMSGPTIAAISVVFDHAEHEEVFQTCVDGFLA
VAKISACHHLEDVLDDLVVSLCKFTTLLNPSSIEEPVLAFGDDAKARMATITVFTIANRYGNYIRSGWRNILDCI
LRLHKLGLLPARVASDVADDSELSSEPGHGKPVTSSLFASQMPSMGTPRRSSGLMGRFSQLLSLDTEEPRSQPTE
QQLQAHQRTLQTIQNCHIDSIFTESKFLQSDSLLQLARALIWAAGRPQKGNTSPEDEDTAVFCLELLIAITLNNR
DRIVLLWQGVYEHICSIVQSTVMPCALVEKAVFGLLRICQRLLPYKENLADELLRSLQLVLKLDARVADAYCENI
TQEVMRLVKANATHIRSQMGWRTITALLSITARHPEASEAGFEALTFIMSDGVHLSPANYILSVDAARQFAESRV
GQADRSLRALDLMEGSLACLARWSRETKEAAGEEAATKVSLDIGEMWLRLVQGLRKVCLDQREDIRTHALLSLQR
CLMGVEDLSLPHALWLQCFDLVIFTMLDDLLEIAQGHSPKDYRNMEGTLVLAMKLLSKVFLQLLPDLSQLTAFCK
LWLGVLSRMEKYMKAKVRGKKSEKLQELVPELLKNTLFVMRTRGVLAQRGALGSDSLWELTWLHVNNIAPSLQSE
VFPDQGIEQSHPEQGIFSESPESEGTEGVRT*
**>Aquca_045_00153**
MGGLKLQSGTKVMEEEPEHHNCTSNRGALACTVNSEIGAVLAVMRRNVRWGGRYVSGADQLEHSLIQSFKALRKA
IFSWEQPWKTINPIVYLQPFLDIIISDETGAPITGVALSSIYKIMSLDVLDVDTVNVGDAMHLVVDAVTSCRFEV
TDPASEEVVLMKILQVLLACMKSKVAIMLSNQHVCTIVNTCFRVVHQARAKGELLQRIARHTMHELIRCIFSQLP
NMDGRENLLSNGSLCEQEFDGVCKDDNSGSKQLDNGNNSAEYGGQAPSLGLSAYPNSVGNMTSVNSVGSVNDKER
SDLLLMTKPYGIPCMVEIFHFLCSLLNIIEHIESGSRSNTIAYDEDVPLFALGLINSAIELGGSSISQHPKLLAL
IQDELFRNLMQFGLSMSPLILSMVCSIVLNLYHHLRTELKLQFEAFFSCLILRLAQSKFGALYQQQEVAMEALVD
FCRQKTFMVEMYANLDCDITCSNVFEDLANLLSKSAFPVNSPLSAMHILALDGLIAVIQGMAESLGNGSPILEQA
LVDIEEYTPFWTVKCENPTEPNRWIPFVRRRKYIKRRLMTGVDHFNRDLKKGLEFLQGAHLLPDKLDPRSVACFF
RYTTGLDKNLIGDFLGNHDEFCIQVLHEFAGTFDFQNLSLDTALRLFLETFRLPGESQKIQRVLEAFSERYYEQS
PHILANKDAALLLSYSIILLNTDRHNAQVKKKMTEEDFIRNNRHINGGSDLPRVFLSELYHSVCKNEIRTIPEQS
IDFPEMTPSHWVDLMRKSEKTSPYIVCNSRSFLDHDMFTIMSGPTIAALSVVLDHAEDEDVFQTCIDGFLAVAKI
SACLHLDDVLDDLVVSLCKFTTLLNPSSIEDSVLAYGDDTKAKMATFAIFTIANSYGNYIRSGWKNILDCILSLR
KLGLLPSFITSDVTNDSELSSEPDHGKPSASQVPSVSAPQRSISLMGRFSQFLSLDTEEPKSQPPEEHLQVDEHT
LHTIQKCHIDSIFTESTFLHADSLLELARALVWAAGRAQKGNTSPEDEETAVFCLELLITITLNNRDRILLLWQT
VFEYICSIVESTVMPCALVEKAVFGLLRICQRLLPYKENLADELLRSLRLVLKLDARVADAYCENIAHEVMCLVK
ANATHIRSLMGWRTITSLLSITARHPDASEAGFEALMFIMSDGVHLSPANYVLCVDAARQFAESRVGEADRSLCS
LDLMAGSIACLAQWSRENKEAVGDEGATKVSQDIQEMWLRLVQGLKKVCLDQREEIRNHALLLLQRCLMGVDGLP
LSPALWLQCFDLVIFTMLDDLLEISQSLSPKDYRNMEGTLVLAMRLLSKLFLQLMSDLSQLTAFNKLWLGVLSCM
ETYLQAKFRGKKSEKIQELVPELLKNTLLVMRTNGVLEQRSALGGDSLWEVTCLHVNNMAPSLITDVFPDQASDH
SEQKQSIVEGKSEPNVEGVRA*
**>Aquca_007_00074**
MAKAVERNEDDDLLCQQNGKKLSKQKRKQLGISCMLNTEVGAILAVIRRVPDSTSQFISTTEEGFDSSVLNSLKS
LRGLIFSPQQEWHTIDPSIYLAPFLDVVQSDDIPASATAVALSAILKILKLEIFNEKTPGARDAINSIVTGITSC
RLEKTDSTSEDAILIRILQVLIGIMRNRTSYLLTDHAVCTLVNTCFQVVQQSVSRGDSLQWHARHTMHELIQTIY
ARLPEIETHEAENSESETEDADMGINLDSGYGIRSMVDIFHFLCSLLNVVEVVDVDGSTATNSDEDVQLFALTLI
NSAIEISGDGIGKHPKLLRMVQNDLFHHLIHYGMCSNPLLLSMICSTVLNIYHFLRRSIRLQLEAFFTCVLLRVS
NGGVSPQLQEVALEGCISFCRQPTFIIEMYVNYDSDPICRNVFEEIGKLLCKRAFPMPNSPLSSIQVQAFECLLT
MIHNIADSMDEDADSRSDRLHPVVVTEYRPFWIAERNNDDSLESWVESLRMRKAQKRKIMIAADHFNRDEKKGLE
YLKMAHLVPEPRDPKALAYFFRYTPGLEKNKIGDFLGDPDEFNLQVLKEFTHTFAFSGMILDNALRTFLETFRLP
GESQKIQRILEAFSDRFYDQQFSELFVSKDAVFILCYSVIMLNTDQHNPQVKKKMTEEEFIRNNRAINGGKDLPR
EYLSELFQSIATNAITLFGQSGSSIEMNPNRWVELINRAKSMDPFVVCNYHHNLSRDMFACISGPAVATICAVFE
QGDEDEILNECVEGLFSVAKVARFGLEDTLDELLASFCKFTMLLNPYASGEETLYAFSNDLKPRMATLAVFTIAN
KFADVIRGGWRNLVDCLLKLKKLKLLPQSVVEPDTSSPSPSDILTHSKSESGVMFPAFDTGYVITPKTSGLTGRF
SHFLSLDCTEESLTQAGTELEQNLKIIKQCRIGNIFSNSSNFPDDSLQNLGRSLIFAAAGKGQKFSTSVEEEETV
GFCWDFLITITMANLHRFMSFWPSFHENLILVAQLPHFSPCPFSEKAILGLFKVCVQLLSSLQSDKLAEEFIFKS
INLMWKLDKEILDSCSVCITQSVSKLLIEYPANIQTQLGWKSVLHLLSVSGRHPETYEQGVEALISLMSDGTNIS
RQNYAYCIDCAFGFAALKISRLENSWKIIDQMVESVNLLIQWYRTGYTDPGSSVSMSNMSSSSFEESKGHNSSTI
SLFIKLAESLRKTSLVRREEIRNHAVLALQRSFVLTDELGFTSTTCINCFNLVIFAMVDDLHEKMVEYSRRENSI
KEMRSMVATLKLAMELLKEIFLQFLRPLTESSGFRTFWLGLLRRMDTCMKAELGDDSELILQQLVPELLKEMITK
MKEKEILAPKEDNDLWEITYIQIQWIAPSLKEELFPTDY*
**>Aquca_020_00178**
MLNTEVGAVLAVIRRIPDPTSQFISTTEEGFESSVLNSLKSLRGLIFSPQQEWHTIDPSIYLAPFLDVVQSDDIP
AAATAVALSAILKILKLEIFNEKTPGARDAINSIVTGITSCRLEKTDSTSEDAILIRILQVLIGIMRNRTSYLLT

DHAVCILVNTCFQVVQQSVSRGDSLQWNARHTMHDLIQTIYARLPEIEAQEAENSESETEDADMGNNLDSGYGIR
SIVDIFHFLCSLLNVVEVVDVDGSTATNSDEDVQLFALTLINSAIELSGDGIGKHPKLLRMVQNDLFHHLIHYGT
CSSPLLLSMICSTVLNIYHFLRRSIRLQLEAFFIYVLLRVSTEVQPTFIIEMYVNYDSDPICRNVFEKIGKLLCK
KAFPIPNSPLSSMQVQAFEGLLTMIHNIADNMDEDAESRSDRSHPVAITEYRPFWIAERNDDDNLESWVESLRMR
KAQKRKIMIAADHFNRDEKKGLEYLKMAHLVPEPPDPKALAYFFRYTPGLDKNKIGDFLGDPDEFNLQVLKEFTH
TFAFSGMILDNALRTFLETFRLPGESQKIQRILEAFSDRFYDQQFSELFVSKDAVFILCYSVIMLNTDQHNPQVK
KKMTEEEFIRNNRAINGGKDLPREYLSELFQSIATNAITIFGQSGSSIEMNPNRWVELINRAKSMSPFVVCKYHH
SLSRDMFGCISGPAVATICAVFEQGDEDEILNECVEGLFSVARVARFGLEDTLDELLASFCKFTMLLNPYASAEE
TLYAFSNDLKPRMATLAVFTIANKFADVIRGGWRNLVDCLLKLKKLKLLPQSVVEPDTISPSPSDILTRSKSGVM
FPAIDTGYGITRQTSGLTGRFSHFLSLDCTEESLTQAGTEFEQNLKIIKQCRIGNIFNNSSNFPDDSLQNLGRSL
IFAAAGKGQKFSTSVEEEETVGFCWDLLITITMANLHRFMSFWPSFHENLILVAQLPLFSPCPFSEKAILGLFKV
CVQLLSSSQSDKLAEEFIFKSINLMWKLDKEILDSCSVTITQSVSKLLIEYPANIQTQLGWKSVLHLLSVSGRHP
ETYEQGVEALISLMSDGTNISRQNYAYCIDCAFGFAALKISRLENSWKIIEQMVESVNLLIQWYRTGYTDPGSNM
SMSNMSSSSFEESKAHNSSTITLFIKVAEALRKTSLVRREEIRNHAVLSLQRSFALADELGFTSTNCINCFNLVI
FAMVDDLHEKMVEYSRRENSIKEMRSMVATLKLAMELLKEIFLQFLRPLAESSGFRTFWLGVLRRMDTCMKAELG
DDSELILQQLVPELLKEMITKMKEKEILAPKQDNDLWEITYIQIQWIAPSLKEELFPTEY*
**>Kinun_030821**
MGRPKMEPGINAIKEEPEEFDSTSNRGALACMINSEVGAVLAIMRRNVRWGGRFMASDDQLEHALIHSLKALRKQ
LFSLPQQWCTVNPVLYLQPFLDVVRSDETGAPITGVALSSVYKILTLNVIDLNTVNVEEAMHLVVDAVTSCRFEV
TDPASEEVVLMKILQVLLACMKNKAAIVLSNQHVCTIVNTCFRVVHQAGTKGELLQRIARHTMHELVRCIFSRLP
DIGVREVPLANGIASSSKPEVRDNAFGSTELENGNSIAEYNGQPPFVGIASTASMANMMNGNTIGASPNDVQLMI
EPYGVPCMVEIFQFLCSLLNIVEHIGIGPRSNTLTFDEDVPLFALSLINSAIELGGPSICQHPKLLALIQDELFR
NLMQFGLSMSSLILSMVCSIVLNLYHHLRMELKLQLEAFFSCVVLRLAQNRHGASYQQQEVAMEALVDFCRQKTF
MVEMYANLDCDITCINVFEDLANLLSRSAFPVNCPLSAMHILALDGLIAVIQGMAERIGNGSTVSEQVPLDFEEY
IPFWTVKCENYADSSHWVQFVRRRKYIKRRLMIGADHFNRDPKKGLEFLQGTHLLPDKLDPQSVACFFRYTAGLD
KNLVGDFLGNHDDFCVQVLHEFAWTFDFEEMNLDTALRLFLETFRLPGESQKIQRVLEAFSERYYEQSQHILANK
DAALLLSYSLIMLNTDQHNVQVKKKMTEEDFIRNNRHINDGNDLPREFLSELYHNILKNEIRTTPEQGAGFPEMT
PSRWVDLMRKSKKTSPFILSDSRAFLDHDMFAIMSGPTIAAISVVFDHAEQEDVFQTCVDGFLAIAKISACHHLE
DVLDDLVVSLCKFTTLTNPSSIEEPVLAFGDDTKARMATITVFTIANTYADYIRTGWRNILDCILRLHKLGLLPA
RVANDVADDSDISVESGSGKPVANSLSSAAQMPLSMSIPRKSSGLMGRFSQLLSLESEEPRSQPSEQQLAAHQRT
LQTIQKCHIDSIFTESKFLHADSLLQLARALIWAAGRPQRGNNTSPEDEDTAVFFLELLIAITLNNRDRINLLWP
GVYEHISNIVQSTLVPCALVEKAVFGLLRICQRLLPYKENLADELLRSLQLVSKLNDRVAHAYCEHITQEVTHLV
KANAAHIKSQMGWRTIISLLSITARHPDASETGFDAFIFIMSEGAHLSPANYVLCVDASKQFAESRHGQAERSLH
ALDLMAGSISCLARWSHETKGEEAAEKVSRDIGEMWLRLVQGLRKVCLDMREEVRNHALTSLQRCLTGVVDGVNL
GLPHAAWLQCFDMVVFTLLDDLLEISQSHSTKDYRNIEGTLVLALKLLSKVFLLLLHDLSQLTTFCKLWLGVLSR
MEKYMKAKIKGKRSEKLQDLVPELLKNTLLVMKTRGVLIQRSALGGDSLWELTWLHVNNIAPSLQSEVFPDQDSE
NVQQKQNEVVEVNSLV*
**>Kinun_019872**
MGRPKMDFGINAIEEEPEEFDSTSNTGALACMINSEVGAVLAIMRRNVRWGGRFMAGDDQLEHSLIYSLKLLRKQ
IFSWPQQWSTVNPVLYLQPFLDVVRSDETGAPITGVTLSSVYKILTLNVIDLNTVNVEEAMHLVVDAVTSCRFEV
TDPASEEVVLMKILQVLLACMKSKAAIVLSNQHVCTIVNTCFRVVHQAGTKGELLQRIARHTMHELVRCIFSRLP
DIVEGEVPLANGVASSSKPEVRDYNFGSTQLDNGNSFVEYNGQPPSVGIASTASMGNMMNGTIGASPNDLQLMTK
PYGVPCMVEIFHFLCSLLNIVEHIGIGPRSNTLFDEDVPLFALSLINSAIELGGASICQHPKLLALIQDELFRNL
MQFGLSMSSLLLSMVCSIVLNLYHHLRMELKLQLETFFSCVVLRLAQSRYGASYQQQEVAMEALVDFCRQKTFMV
EMYANLDCDITCSNVFEDLSNLLSKSAFPVNCPLSTMHILALDGLIAVIQGMAERIGNGSAVSEQVPFNFEEYIP
FWTVKCEDYADPFHWVPFVRRRKYIKRRLMIGADHFNRDPKKGLEFLQGTHLLPDKLDPQSVACFFRYTAGLDKN
LVGDFLGNHDDFCVQVLHEFAWTFDFEEMNLDTALRLFLETFRLPGESQKIQRVLEAFSERYYEQSQHILANKDA
ALLLSSIILLNTDQHNVQVKKKMTEEDFIRNNRQINDGNDLPREFLSELYHNICKNEIRTTPEQGAGFPEMTPSR
WVDLMRKSKKTSPFILSDSRAFLDHDMFAIMSGPTIAAISVVFDHAEQEDVFQTCVDGFLAIAKISACHHLEDVL
DDLVVSLCKFTTLVNPSSIEEPVLAFGDDSKARMTTITVFTIANTYGDYIRSGWRNILDCILRLHKLGLLPARVA
NDVADNSDISVESGPGKLVTNSLLLAAQMPSSMSTPRKSSGLMGRFSQLLSLESEEPRSQPSEQQLAAHQRTLQT
IQKCHIDSIFTESKFLHADSLLQLARALIGAAGRPQRGNNTSPEDEDTAVFCLELLIAITLNNRDRITILWPGVY
EHISNIVQSTLMPCALVEKAVFGLLRICQRLLPYKENLADELLRSLHLVLKLNDRVAHTYCEHITQEVTRLVKAN
ATHIRSQMGWRTIINLLSTARHPDASETGFDALIFIMSQGVHLSPANYVLCVDASRQFAESRHGQAERSLHALDL
MAGSISCLSRWSHETKGEEAAEKVSRDIGEMWLRLVQGLRKVCLDMREEVRNHALTSLQRCLMGVVDGVNLDLPQ
AAWLQCFDMVVFTLLDDLLEISQSHSTKDYRNIEGTLVLAMKLLSKVFLQLLHDLSQLTTFCKLWLGILTRMVKY
MKAKIKGKRSEKLQDLVPELLKNTLFVMKARGVLIQRSALGGDSLWELTWLHVNNINPSLQCEVFPDLDSENAQQ
KQNEIEGRRQSGVIESIE*
**>Kinun_020572**
MARAIDPNQEEEEEEEEEEEDGRSNSSRQEKKKKTAAIQNKELGVSCMLNTEVGAVLAVIRRVPDPNSQFASSPE
EHYDSHLLNSLKSLRALIFNPLQEWDMIDPSTYLAPFLDVIQSDDIPASATGVALSAILKILKLEIFNLKTPAAR
DTINSIVTGITSCRLEKTDPTSEDAVLMRILQLLTGIMRNPTSVLLTDHAVCTLVNTCFQVVQQSASRGDLLQCN
ARHTMHELIQTIFTRLPDIEVREGESSESDTEDADIMGTSDSGYGARCVVDIFHFLCSLLNVVEVVDMDGGAATS

SDEDVQLFALILINSAIELSGEGIGKHPKLLRMIQDDLFHHLIHYGTCSSPLVLSMICSTVLNIYHCLRRSIRLQ
LEAFFTFVLFRVVNGGGSSQLQEVALEGIISFCRQPTFIIEMYVNYDCDPAFRNVFEDIGKLLCKRAFPAVNSPL
SSLQIQAFEGLLLMIHNIADNIDENKNSGANGHYPVTITEYKPFWTAKCNDEDNETWIEFLRVRRAQKRKMMIAA
NHFNRDEKKGLDYLKISHLVPNPPDPKAFACFYRYIPGLDKNKIGDFLGDPDEFNIQVLKEFTNTFEFSGVILDN
ALRTFLETFRLPGESQKIARVIEAFSERFYDQQSSTVFAGKDAVYILCYSLIMLNTDQHNIQVKKKMTEEEFIRN
NRAINDGKDLPREYLSELFQSISNNEIKVYGETSSSVLLMTPTRWIELINRAKVMEPFIFCNFNISLGRDMFASI
SGPSVAVLSATFEHADEDETLHECIEGLFSVARISRFGLEDTLDELLASFCKFTMLLNPGAQPEDTLHAFSNDLK
ARMAILAIFTIANKFGDSIRGGWRNIMDCLLKLKKLKLLPQSVVEPDTPSTSSSEILKHFKSESGVLFPSADARF
NSRQMSGLIGRFSHFLSLDCTEEMLAQAGSDFEQNLKIIQQCQIGNIFSSSSKLVDDSLHNLGRSLIFAAAGKGQ
KFNTSVEEEETAGFCWNLIMTITSANLHRFLVFWPSFHDYMLVAAQFPFFTPCPFAEKAIISLFKICLKLLGSDQ
SDRLSEEFIFKSINLMWKLDKEILDSCCESITQTVSIILTEYPANIQTQLGWKSVLHLLSVTGRHPETYDQGVQA
LIMFMSEGSHLSRTNYAYCIDCAFGYAALKISPLEKSAKILDLMADSVNLLIQWYRNEYSDPGSNISIVSNTSSS
SVEDGSKILNSSSFTLNLFIKLAESLRKTSLVRREEIRNHAVSALQRSFALADELGFTSTTCINCFNLVIFAMVD
DLHEKMVEYSRRDNSVKEMKSMAGTLKLAMELLNEVFLQFLKPLSQSPGFRTFWLGVLRRMDTCMKADLGTEGEL
KLRELVPELLKKMITKMKEKEILVQKEDNDLWDITYIQIQWIAPTLKEELFPEEQF*

**>AketrGNOMa_chr14**

MGRLKLQPGIKSIEEEPEEFDCTSNRGALACMVNSEVGAVLTVMRRNVRWGGRYVAGDDLLEHSLIQSLKSLRKQ
IFSWQHPWNTINPVVYLQPFLDVIRSDETGAAITGVALSSVYKILTLDVLDLNTVNVEDAMHLVIDAVTNCRFEV
TDPASEVVLMKILQVLLACMKSKVAIVLSNHHVCTIVNTCYRIVHQAESKGELLQRIARHTMHELVRCVFLHLPD
VDNGELPLANDGASSGKQEVGLDKDYTFGSKQLDNGNSGSEYDGQPTSVGFSSMASAGSMASMMNENTMGVGNGK
EPSPNDLQLMTEPYGVPCMVEIFHFLCFRLNSVEHIGMGPSSNAMAFDEDVPLFALGMINSAIELGGCSILQHPK
LLTLIQDELFRILMQFGLSMSPLILSMVCSIVLNLYHHLRMELKLQLEAFFSCVILRLAQSRHGASYQQQEVAME
ALVDFCRQKSFMVEMYANLDCDITCSNVFEDLANVLSKSAFPVNCPLSAMHILALDGLIAVIQGMAERISNGSPV
SEQTPIDFEEYTPFWTVKCENYADPDHWVSFVRRRKYTKRRLMIGADHFNRDPKKGLEFLQGTHLLPDKLDPQSV
ACFFRYTAGLDKNLVGDFLGNHDEFCVKVLHEFAWTFDFQDMNLDTALRLFLETFRLPGESQKIQRVLEAFSERY
YEQSPQILANKDAALLLSYSLIMLNTDQHNVQVKKKMTEEDFIRNNRHINGGNDLPREFLSELYHSICKNEIRTI
PEQGAGFPEMTPSRWIDLMRKSKKTSPFIVCDSRAFLDHDMFAIMSGPTIAAISVVFDHAEHEDVFQTCVDGFLA
VAKISACHHLEDVLDDLVVSLCKFTTLLNPSSVEEPVLAFGDDTKARMATITVFTIANRYGNYIRTGWRNILDCI
LRLHKLGLLPARVASDVADDLELSAEPGHGKPVTSSLSASQMPSMGTPRRSSGLMGRFSQLLSLDSEEPRSQPTE
QQLAAHQRTLQTIQKCHIDSIFTESKFLHADSLLQLARALIWAAGRPQKGNTSPEDEDTAVFCLELLIAITLNNR
DRIILLWQGVYEHISNIVQSTVMPCALVEKAVFGLLRICQRLLPYKENLADELLRSLHLVLKLDARVADAYCEHI
TQEVMRLVKANATHIRSHVGWRTITSLLSITARHPEASEAGFEALTFIMFDGVHLSPANYVLCVDSSRQFAESRV
GQADRSVRALDLMSGSVTCLARWSRETKEAVGEEASAKFSQDIGEMWLRLVQGLRKVCLDQREEVRNHALLSLQR
CLIGVDGVHLPHTSWLQCFDLVIFTMLDDLLEIAQSHSPKDHRNMEGTLVLAMKLLSKVFLQLLHDLSQLTTFCK
LWLGVLSRMEKYMKAKVRGKRSEKLQELVPELLKNTLLVMKARGVLVQRSALGGDSLWELTWLHVNNIAPSLQSE
VFPDQDSGETQPKKSETGGSSETYKDGPVE*

**>AketrGNOMb_chr11**

MNMGKKLLSRTDANEEEPGHSMPSGCALACMINSEIGAMLAVVRRNVRWGGRYMAEDDQLEHSLIQSLKTLRKKL
FSWDHQWHAFDPSIYLLPFLDVIRSDEIGAPITGVALSSVYKILTLGILDPITVNVESAMHLVVDAVTSCRFEVT
DSASEEVVLMKILQVLLACMKSKASFSLDNQHVCTIVNACFRIVHQARTKGELLQRIARHTMHELVRCIFSHLGD
VHHTEYPLYNGGGPSLKPEVSVLEKDRTFGVMQTESRNESPQNASVSAIDRSEEAVNDVDHSLGEPYGVACMVEV
FQFLCSLLNIGEQIGMGSSTNSTEFDEDVPLFALGLINSAIELGGPTINRHPKLLALIQDELFRNLMQFGLSMSP
LILSMVRNIVLNLYLHLRTELKLQLEAFFSCVVLRLAQSKHGDASYQQQEVLMEAIVDFCRQKTFMAEMYANFDC
DISRNNVFEDLGNLLSKSAFPVNSPLSSMHVLSLEGLVVVLQGMAERIGSNSTPVWEGTLLDLEHYLPFWTIKCE
NYSDPNCWVGFVRQRKHIKRKLMTGVDHFNRHPKKGLEFLQKTHLLPDELDAKSVACFFKYTMGLDKNIVGDFLG
SHDEFWVQVLHEFAQTFDFQGMSLDAALRLFLETFRLPGESQKIQRVLEAFSERYYVQSPQVLANKDAALVLSYS
IILLNTDQHNVRVKKKMTVEDFVRNNRHINDGNDLPREFLLELYHSICKNEIRMTPEQGAGFPEVTPSHWINLIH
KSQTASPYITCDPRPFLDRDMFAVLSGPTLAAISVVFDHAEHEDIYQMCVDGFLAMAMISSCSQFVDVLNDLVVS
LCKFTTLLNPSQMDEGILSFADNPKAMMATVTVFTIANKFGDYIHVGWRNIIDCILSLHKLGLLPTIMASDSANT
SSDPVPEKPITNSQPISHVPIRTTSQRSSGLMGRFSQLLSLDTEDPILEPTEEELEAHQRAHQTIQKCQVNSIIT
DSMFLQSDSLLQLVQALILVPSSSLKGKNSSPDEEDTSVFCLELLITITLYNRDRAVLIWPTVYEHIASIVHSTV
TPCTLVEKAVFGLLHICQRLLPYKDNLADELLRSLQLVLKLDARVADAYCEHITQEVADLVKANAVCIKSQMGWR
TITSLLSITARHPDASEAGFEALLFIMSDRTHLSPANYVLFVDASRQFAESWVGSIERSVQALDLMEGSFKCLAG
WRLPPEEVGEIWLRLLRGLSKVCLDQREEVRNRAVLALQRCTMGVEEFCLPPDFWVNCFDLVIFAMLDDLLEIVQ
GQLSRDYRNMEGTFVHVLKLSSKVFLQVVGDLSLSISFSKVWLGILDRMGLYYKVKIRGKKTERLRELVPELLKN
NLLVLKTRGVLVKPSSDTGSSGDTLWELTWPHINNVAPALRAEVFPDQEPELVHAQNEIVSPTMA*

**>AketrGNL2_chr10**

MAKAIDQNRDKEDDGDGAssqqknkkttklknRELSISCMLNTEVGAVLAVIRRVSDPNPQFTPEDNYDSSLINS
LKSLRALIFKPQQEWHTIDPSIYLAPFLDVIQSDDIPASATGVCLSAILKILKLEIFNEKTPGARDAINSVVTGI
TNCRLEKTDPNAEDAVLMRILQLLSGIMRNHTSILLTDHAVCTLVNSCFQVVQQSASKGDLLQCNARHTMHELIQ
TIFARLPDIEDRSGENSESDAEDAETQNIRNSGYGARCAVDIFHFLCSLLNVVEVVDMEGSTATNSDEDVQLFAL
ILINSAIELGGEGIGKYPKLLRTIQDDLFYHLIYYGTCSSPLLFSMICSTVLNIYHFLRRSIRLQLEAFFTYVLL
RVASGGNLSQLQEVALEGIISFCRQPTFIIEMYVNYDCDSVSRNVFEEIGKLLCKRAFPGANNPLSSMQIQAFEA

LLIIIHSIADNMDEDEDRRSSGSYPVVVNEYKPFWTVEDKNNMDMGSWVEFLRVRKTQKRKIMVAADHFNRDEKK
GLDYLKVAHLVPETPDPKALACFYRFTPGLDKNKIGDFLGDPDEFNIQVLKEFTDTFDFSGIILDTALRTFLEAF
RLPGESQKIHRILEAFSERFYDQQSSEIFVTKDAVFILCYSVIMLNTDQHNPQVKKKMTEEEFIRNNRAINGGKD
LPREYLSELFQSISNNAITLFGMSSSFVEMNTNKWVDLINRSKVVEPFIICNFNQSLGRDMFASISGPAIAALSA
TFEHADEDEILHECIEGLFSVARISRFGLEETLDALLASFCKFTLLLNPYATAEETLYAFSNDLKSRMATLAVFT
VANKFGDSIRGGWRNIVDCLLKLKRLKLLPQSVVEPDTASTSSSDILMHSRSESGVIFPSTDTGFGSSRQMSGLI
GRFSHFLSLDTEDSLTHAGSELEQNLKIIQQCRIGNLFSSSSNLLDDSIYNLGRSLIFAAAGKGQKFSTSVEEEE
TVGFCWDLIITITLANLHRFPTFWPCFHDYLLVVAQFPLFSPCPFAEKAIMGLFKVCLKLLGSYRSEKISEDLIF
KSINLMWKLEKEILDTCCNNITQSVSKILTEYPANIQTQLGWKSVLHLLSVAGRHPETYDQGTEALIMLMSDRTH
VSRTNYAYCIDCAFGYAALKISPLEKSWMILDLMADSVKLLIQWYQNDYSDPGSNISNTSSSSFDDSLKALNSSN
STMNLFIKLAESLRKTSLVRREEIRNHAVVALQRSFRLAEDLGFTATNCINCFNLVIFSMVDDLHEKMIEYSRRE
NSVKEMRSMAGTLKLAMELLKEVFLQFLKPLSQNQGFRTFWLGVLRRMDTCMKGELGDECESILLELVPVLLKDM
ITKMKDKEILVHKEDNDLWDITYIQIQWIAPSLKEELFPEEQL*

**>Eca_sc010404_g0150**

MGRLKLQSGINSIEEEPEDCEYVPNIGAFACMLNSEVGAVLAVMRRNVRWGGRYMAGDDQLEHSLIQSLKALRRQ
IFSWKNQWNTINPSLYLQPFLDVVRSDETGAPITGVALSSVYKIMTMDVIDLNTVNVEDAMHSVVDAVTSCRFEV
TDPASEEVVLMKILQVLLACMKSKAAIVLSNQHVCTIVNTCFRVVHQAGTKGELLQRIARHTMHELVRCIFSHLP
DIESRDQSLANGDTPYSNQEMGIDKDYSFGSRQLDNGNGGSEYDGPPSSVGLGSTASTGSMPSMMGGNTFGHNSK
EGGPNDIHLMTEPYGVPCMVEIFNFLCSLLNIIEHGGMGARANTLAFDEDVPLFALGLINSAIELGGSSLRQHPK
LLGLIQDELFRNLMQFGLSMSPLILSMVCSIVLNLYHHLRMELKLQLEAFFSCVILRLAQSRHGATYQQQEVAME
ALVDFCRQKSFMVEMYANLDCDITCTNVFEDLANLLSKSAFPVNCPLSGMHILALDGLIAVIQGMAERIGNGSPI
SEQSPIDFDEYTPFWTVKCENYADPSHWVPFVRRRKYIKRRLMIGADHFNRDPKKGLEFLQGTHLLPDKLDPQSV
ACFFRYTAGLDKNLVGDFLGNHDEFCVQVLHEFAWTFDFQDMNLDTALRLFLETFRLPGESQKIQRVLEAFSDRY
YEQSPQILANKDAALVLSYSLIMLNTDQHNVQVKKKMTEDDFIRNNRHINGGNDLPRDFLVELYHSICKNELRTT
PEQGVGFPEMTPSRWVDLMRKSKKTSPFIMSDSRAFLDHDMFAIMSGPTIAAISVVFDHAEQEDVFQTCVDGFLA
VAKISACHHLEDVLDDLVVSLCKFTTLLNPSSVEEPVLVFGDDTKARMATISVFTIANKYGNYIRTGWRNILDCI
LRLHKLGLLPARVASDVAEDSDLSAESGHGKPVTSSLSSSQMPSMGTPRRSSGLMGRFSQLLSLDTEEPRSQPTE
QQLAAHQRTLQTIQKCHIDSIFTESKFLQADSLLQLARALIWAAGRPQKGNTSPEDEDTAVFCLELLIAITLNNR
DRIVLLWQGVYEHISNIVQSTVMPCALVEKAVFGLLRICQRLLPYKENLADELLRSLQLVLKLDARVADAYCEHI
TQEVMRLVKANTTHIKSQMGWRTITSLLSITARHPEASESGFEALMFIMSDGGIHLSPANYALCVDASRQFAESR
VGHADRSMRAVDLMAGSITCLARWSNEIKDAPGDDSATKVYQDIGEMWLRLVQGLRKVCLDQREEVRNHALLSLQ
KCLTGVEEVRIPHSSWLQCFDLVIFTVLEALLEIAQSHPTKEFRNMEGTLLLAMKLLTKVFLQMLNDLSQLTTFC
KLWLGVLSCMEKYTKAKIRGKKSEKLQELVPELLKNTLVVMKTKGILQQRSALGGDSLWELTWLHVNNIAPSLQS
EVFPDQGPEQKDGGTDVQT*

**>Eca_sc000188_g0230**

MAMESLKLHVGIEANGDAHGHCFPSSSAFACMINSEIGALLAVVRRNVRWGGRYVAEDDQLEHSLIQSLKSLRKK
LFSWDRQWNTVDPSIYLQPFLDVIRSDEIGAPITGVALSSVYKILSLGLLDPTTTNVKDSMHLVVDAVTACRFEV
TDSASEEVVLMKILQVLLACMKSKSSFLLNNQHVCTIVNTCFRIVHQAGTKGELLQRIARHTMHEIVRCIFSHLQ
NIHYTAKLSMLGDGSNDNPKVDLEKDNTFGVKQQDSVNGNPENASVSTGSVGSLVDDTVVGNDNNEQTANDIPNS
MKEPYGVACVIEVFQFLCSLLNAGEQIGMSPRLNWTEFDEDVPLFALGLINSAIELGGPVISGHPALLVLIQEDL
FPNLMQFGLSMSPLILSVVSNIILNLYLHLRVELKLQLEAFFSSIISRLTQSKHGDASYQQQEVLMEALVDLCRQ
KTFMTEMYANFDCDISSSNIFEDICNLLSRSAFPVNSPLSSMHVLALEGLVAVVQGFSGRIENSTTISEQGLLGA
EKYTPFWNEKCENYSDPSSWVGFVRQRKQIKKVLMTGVEHFNRDPKKGLEYLQRTNLLPGELDPESVACFFKYTM
GLDKTIVGDFLGSHEEFWIQVLHEFARTFNFQEMGLDTALRLFLETFRLPGESQKIQRILEVFSERFYEQSPQIL
ANKDAALVLSYSIILLNTDQHNVRVKKKMTVEDFVRNNRRINGGDDLPREYLLELYHSICKNEIRMTPDQGAGFT
EVVVPSHWINLMHKSKTTSPFITCEPKPFLDRDMFSVLSGSTLAAISVVFDHAENEDIYQICVDGFLAMAKLASC
THSVALLDDLVISLCKFTTLLDPSQTDEPIIAFADDSKAMMATETVFTIANNFGDYIRAGWRNILDCILSLHKMG
LVPSSMANNEQSLDPASGKPIIQSTPYVPARTISSQRSSGFMGRFTQLLYLDTEQKNLEPSEEELEAHQRAHQTI
QKCQVSTIFTDSMFLQSESLLQLVQALIWVSNPSQKGKNGSPEDADTAIFCLELLITITLYNRDRIVLLWPVVYE
HIAGIVQSATMPSAFVEKAVFGLLHICQRLLPYKENLADELLRSLQLVLKLDARVADAYCEQITQEIADLVKTNA
ICIKSQMGWRTITSLLSITARHPEASDVGFEALAFVLSDGSHLSPLNYSLCVDACRQFAESRVGSTERSIHALDL
MAGSFTYLARWNLPPQEVGDIWVGLVKGLKKICLDQREEVRNHAILSLQRCTTGIEGFSLPQDSLVQFFDLVIFT
LLDDILEIAQSKTSREYRNMEPTLLHALKLSSKVFLQLIRELSSLSSFPKIWVEILGRIGNYFAVKIRGKKSEKI
RELVPELLKNTLFVMKTNKILVNPSSKSETGSESTLWELTWPLIDNIVPSLKSEVFPDQEAEMTHRPKDTAISEL
PNNGVNTSTASIQS*

**>Eca_sc001331_g0490**

MAKAIEEESDNGNDSQKNKRTTKLQKKELGISCMLNTEVGAVLAVIRRIPDSNSQFLSTPEDNFESSILTSLKSL
RALIFNPQQEWHKIDPSVYIGPFLDVIQSDEIPAAATGVALSAILKILKLEIFNEKTPGSRDAINSIVTGVTSCR
LERTDPTSEEAVLMRVLQVLIGIVRNGASVLLTDHAVCTLINTCFQVVQQSASKGDLLQCNARHTMHELVQTIFA
RLSEIDVSDNSEQDEDDDDEISNNNQESGYGARCMIDIFQFLCSLLNVVEVVDTDGSTATNSDEDVQLFALILIN
SAIELSGEGIRKHPKLLRMIQDDLFHHLIHYGMCSNPLLLSMICSTVLNIYHFLRRYIRLQLEAFFTYVLFRVAA
GGSLQLQEVALEGIISFCRQPTFIIEIFVNYDSDPIFRNVFEEIGKLLCKIAFPIPNNPLSSLQMQAFEGLLIMI
HNIADNIDEDENPASIPSYPVFVTEYKPFWIGETKTNDNIDTWVESIRFRKAQKRKIMIAANHYNRDEKKGLDYL

KIAHLVGDPPDSKAFAWYFRFTPGLDKVKIGDFLGDPDEFHLQVLKDFTDTFNFNGMILDTALRTYLETFRLPGE
SQKIQRILEAFSEKFYDQQASEIFVSKDAVFILCYSLIMLNTDQHNPQVKKKMTEDEFIRNNRAINGGKDLPREY
LLALFQSISNNAITLFGQSGTPVDMNQNRWIDLMNRAKTMEPFILCNFTNCLGRDMFASISGPSVATLSAIFEHV
EEDETLHECIEGLFSVARIARFGLEDTLDELLASFCKFTVLLNPYSSAEETLYAFSNDMKPRMATLGVFTIVNKF
GDSVRGGWRNIMDCLLKLRRLKLLPQSVIEPEVASTSSPDIQSDSKSESGVLFPSFDMGHGSSRQMSGLIGRFSH
FMSLDCTEDSLLQAAGNDFEQNLKIIQQCRIANIFTNSPSLPEESLESLGRSLIFAAAGKGQKFSTPVEEEETVG
FCWELIITVSLANVHRFPSFWPAFHDYLILVAQFPLFSPCPFAEKALLGLFRVCLKLLSSYRSEKVSEELIFKSI
NLMWKLDKEILDTCCEMITQSVSKMLTEYPANIQTQLGWKSVLHLLSISGRHPETYDQGVEALIMLMSDGYHVSR
LNYAYCIDCAFGFAALKISPLEKSWKILDLMADSVNLIIQWYSSGYTDPGSNVSNTSNSSFEEGSKNINSFNFAM
NLFIKLVESLRKTSLVRREEIRNHAVLALQKSFSLADELGFTSTSCMNCFNLVIFAMVDDLHEKLVEYSRRDNST
KEMKSMAGTLKLAMELLTEVFLQFLKPLSQSSGFRTFWLGVLRRMDTCMKTELGDNDESGLQALIPELLKKIITE
MKEKEILVKSEENELWDITHVQIQWIAPSLKGELFPEEEF*

**>Papso_LOC113307666**

MGRLKLQSGINSIEEEPLEECESMTSSGALACMVNSEVGAVLAVMRRNVRWGGRYMAGDDQLEHSLIQSLKSLRR
QIFTWQSQWNTINPALYLQPFLDVIRSDETGAAITGVSLSSVYKILTINMLDLNTVNVEDAMHSVVDAVTSCRFE
VTDPASEEVVLMKILQVLLACMKSKVAVVLSNQHVCTIVNTCFRVVHQAGSKGELLQRIARHTMHELVRCIFLHL
RDIESKEHSSANGSMPYSKKEMGIDKDYAFGTKQLDNGNGNSEYEGVPASIGFASTVTSGSIASMMDANTIGAAN
GKDSPNNVHLMTEPYGVPCMVEIFHFLCSLLNIIEHNGMGPRSNTLAFDEDVPLFALGLINSAIELGGPSLCQHP
KLLGLIQDELFRNLMQFGLSMSPLILSMVCSIVLNLYQHLRMELKLQLEAFFNCVILRLAQSRHGASYQQQEVAM
EALVDFCRQKSFMVEMYANLDCDITCSNAFEDLANLLSKSAFPVNCPLSGMHILALDGLIAVIQGMAERIGNGSP
ISEQSQVDFVEYSPFWTVKCENYADPTHWVPFVRRRKYIKRRLMIGADHFNRDPKKGLEFLQGTHLLPDKLDPQS
VACFFRYTAGLDKNLVGDFLGNHDEFCVQVLHEFAWTFDFEDMNLDTALRLFLETFRLPGESQKIQRVLEAFSER
YYEQSPLILANKDAALLLSYSIIMLNTDQHNVQVKKKMTEEDFIRNNRHINGGGDLPREFLIDLYHSILKNEIRT
TPEQGAGFPEMTPSRWVDLLRKSKKTSPFIMCDSRAFLDHDMFAIMSGPTIAAISVVFDHAEHEDVFQTCVDGFL
AVAKISACHHLEDVLDDLVVSLCKFTTLLNPSSVEEPVLAFGDDTKARMATISVFTIANKYGNYIRTGWRNILDC
ILRLHKLGLLPARVASDVADDSEHSSESGRGQPVTSSLSASQMPSMGTPRRSSGLMGRFSQLLSLDTEEPRSQPT
EQQLAAHQRTLQTIQKCHIDSIFTESKFLQADSLLQLARALIWAAGRPQKGNTSPEDEDTAVFCLELLIAITLNN
RDRIVLLWQGVYEHISNIVQSTVMPCALVEKAVFGLLRICQRLLPYKENLADELLRSLQLVLKLDARVADAYCEH
ITQEVMRLVKANTTHIRSQMGWRTITSLLSITARHPEASESGFEALIFIMSDGGVHLSPANYVLCVDAARQFAES
RVGQSDRSVRAVDLMAGSVTCLARWSKEFKDAPGEDTATKVAQDIGEMWLRLVQGLRKVCLDQREEVRNHALASL
QKCLTGVEGVPLPHASWLSCFDLVIFTALDDLLEIAQGHSPKDYRNMEGTLVLAMKLLAKVFLQLLHDLSQLTTF
CKLWLGVLSRKEKYMRAKIRGKRSEKLQELIPELLKNALLVMKAKGVLQQRSALGGDSLWELTWLHVNNIAPSLQ
SEVFPDQEIGQKHENETNAATYKDVQAVGQ*

**>Papso_LOC113297613**

MTVGSLKMHAGIEENGHGEPGQYIPSSTAFACMINSEIGALLAVVRRNVCWGGRYMSEEDQLEHSLIQSLKSLRK
KLFSWNRQWNAVDPSIYLQPFLDVIRSDEIGAPITGVALSSVYKILSLGILDPTTVNVKDAMHLVVDAVTSCRFE
VTDSASEEVVLMKILQVLLACMKSKASFLLNNQHVCTIVNTCFRIVHQAGTKGELLQRIARHTMNELVRCIFLHL
QHFNHTGKPSTHGNGPPGETEVSVLQKEYTSEDKQLDRSNGSSENASVSVGSLRSLVDDSVVGTGHSEVAVDAVP
NPMNEPYGVACVLEVFQFLCSLLNVGDQIGMSPRLNWTEFDEDVPLFALGLINSSIELGGSFISQHPKLLILIKE
ELFPNLMQFGLSMSPLILSMVCNIILNLYLHLRKELKLQLEAFLSSVIQRLAQSKHGDSSYQQQEVLMEALVDLC
RQKTFMAELYANFDCDISCCNLFEDVSNLLSKSAFPVNSPLSSMHVLALEGLVAVVEGFSERISNSPSFSDKSLL
VLEQYTPFLNEKCKNYADPDSWVGFVRHRKQIKKTLMMGVDHFNRDPKKGLEFLQRTHLLPTELDPQSVALFFRY
TLGLDKNIVGDFLGSHEEFWVQVLQEFARSFNFQDMNLDIALRLFLESFRLPGESQKIQRVLEAFSERFYEQSPQ
ILANKDAALVLSYSIIILLNTDQHNVRVKKKMTVEDFVRNNRRINGGDDLPREFLLELYHSICKNEIKMSPEQGAE
VVTPSHWVNLMHKSKTAPPFITCDSKPFLDRDMFSVLSGPTLAAICVVFDNAENEDIYQTCIDGCLAMAQIASCS
HSVNILDDLVISLCKFTTLLDTSQMDDPILAFADDTKAMMATVTVFTIANKYGDYIRAGWRNILDCILSLHRIGL
LPSVAASNEQSSDPVSGKPIAQSTAYVPVKTLAPQRSSGFMGRFSQLLSLDTEQQSLEPTEEELEAQRRAHQTIE
KCQVSTIFTDSMFLQSDSLLQLVQALIWVSGPSQKGKNGSSEDEDTSIFCLELLTTITLYNRDRIVLLWPGVYEH
IAGIVQSAVMPSAFIEKAVFGLLHICQRLLPYKENLADELLKSLQVILKLDARVADAYCEQITQEIADLVKTNAG
CIKSQMGWRTIASLLSITARHPEASEVGFETLAFIMTDGSHLSAVNYSLCVDACRQFAESRVGSAERSIHALDLM
AGSFTYLTRWKLSPQEASEVWLALVKGLKKVCLDQREEVRNHAILSLQRCTIMIEGFSLLPELWLNFFDLIIFQM
LDDFLEIVQGKSSRDYRNMEITLLHALKLSSKVFLQQIMDLSSLSSFPKLWVGILGRMEIYATIKIRGKKNEKLR
ELVPELLKNSLLVMKTSGILLNPKINPGSDPTLWELTWPIVDNLVPSLKSEVFSDKEPDVTQQQKDSTIPELSNN
GISTSTASIQS*

**>Papso_LOC113316139**

MARAIEEDSDEENGATGVQKNKRFSKLRKKELGISCMLNTEVGAVLAVIRRIPDSNSQFLTEDNYDSTLLTSLKA
LRALIFNPQQEWHKIDPSVYLGPFLDVIQSDEIPASATGVALSAILKILKLDIFNEKTPGVQDAINSIVTGVTSC
RLEKTDSIAEEAVLMKVLQVLIGIMRNGASIYLTDHAVCTLVNTCFQVVQQSASKGDLLQCNARHTMHELVQIIF
ARFPEIDNVRENSESDDDEQLANAHDNRYGARCAIDIFHFLCSLLNVVEVVDTDGSTSTTSDEDVQVFALILINS
AIELSGEEIRAHPRLLKMIQNDLFHHLIHYGSCSSPLLLSMICSTVLNIYHFLRRYVRLQLEAFFSFVLLKVANG
GSSPQLQEVALEGVISFCRQPTFVIEMYVNYDCDPVFRNVFEEIGKLLCKLAFPVPSTPLTSLQMQAFEGLLIVL
NNIADNIDEEANPTSVPSYPVVVSEYKPFWIGESKSNGNTEAWIESLRFRKGQKRKLLIAANHFNRDEKKGLDYL
RITHLTPEPPEAKALALFFRFTPGLDKSKIGDFLGDPDKFYLQVLKEFTDTFSFNGMILDTALRTYLETFRLPGE

SQKIQRILEAFSEKFYDQQSSEIFVSKDAVFILCYSLIMLNTDQHNPQVKKKMTEDEFIRNNRAINGGKDLPRKY
LSELFQSISNNAITIFGQSGAPVLMNPNRWIDIINRSKVTEPFILCSYTNCLGRDMFASIAGPSVAALSAIFEHA
SEDETLHECIEGLFSVSRIARFGLEDTLDELIASFCKFTMLLNPYSTAEETIYAFSNDMKPRMATLGVFSIINKF
GDSVRGGWRNIVDCLLKLKRLKLLPQSLLEPDSTSSADIQRDSKSESGVSFTSFDLGHGSSRQASGLGGTGRFSH
FLSLDCTEDALLQAASGNDFEHNLKIIQQCHIDSIFSNSPTLPEDSLESLGRSLIFAAAGKGQKFSTPVEEEETV
GFCWDLIVTLTLANLHRFSKFWPLFNDYLILVAQFPLFSPCPFAEKAILSLFKICLKLLNSYRTEKMSEENIFKS
INLMWKLDKEILDTCSETITQAVSKMLIEFPANIQTQLGWKSILHLLSISGRHPETYDQGVKALIMLMSDGIHVS
KINYAYCIDCAFGFAALKISPLEKSWKILDLMADSVNLLIQWYRNDYSDPGSNASNNSGSSFEDKNPNSYNSTMN
LFIKLVESLRKTSLVRREDIRNHAVLALQKSFILAEQLGFTSNNCINCFNLVIFAMVDDLHEKLVEYSRRDNAAK
EMKSMAATLKLAMELLKEVFLQFLTPLSQSSGFRTFWLGVLRRMDTCMKTELGDDDETGLQALIPELLKKIITEM
KEKEILVHTEENELWDITHVQIQWIAPSLKEELFPDEF*

**>Macco_1831g8**

MGRLKLQSGINSIEEEPEDCESVSKSGALACMVNSEVGAVLAVMRRNVRWGGRYMAGDDQLEHSLIQSLKALRKQ
IFSWQNQWNTINPALYLQPFLDVIRSDETGAPITGVALSSVYKILTIDILDLNTVNVEDAMHSIVDAVTSCRFEV
TDPASEEVVLMKILQVLLACMKSKAAVVLSNQHVCTIVNTCFRVVHQAGTKGELLQRIARHTMHELVRCIFSHLP
DIDSREHSLANGDTSYSQKEMGLDKDYAFGTKQLDNCNGSSEYDGPPPSVGFASTGSMASMMDGNTIGGGNGKEV
SPNDLHLMTEPYGVPCMVEIFHFLCSLLNIIEHNGMGSRSNTIAFDEDVPLFALGLINSAIELGGPSLCQHPKLL
GLIQDELFRNLMQFGLSMSPLILSMVCSIVLNLYHHLRMELKLQLEAFFSCVILRLAQSRHGASYQQQEVAMEAL
VDFCRQKTFMVEMYANLDCDITCSNVFEDLANLLSKSAFPVNCPLSAMHILALDGLIAVIQGMAERIGNGSPISE
QTPIDFEEYNPFWTVKCENYADPSYWVPFVRRRKYIKRRLMIGADHFNRDPKKGLEFLQGTNLLPDQLDPQSVAC
FFRYTAGLDKNLVGDFLGNHDEFCVQVLHEFAWTFDFQDMNLDTALRLFLETFRLPGESQKIQRVLEAFSERYYE
QSPLILANKDAALLLSYSLIMLNTDQHNVQVKKKMTEEDFIRNNRHINGGNDLPRDFLTELYHSICKNEIRTTPE
QGAGFPEMTPSRWVDLMRKSKKTSPFIMCDSRAFLDHDMFAIMSGPTIAAISVVFDHAEHEDVFQTCVDGFLAVA
KISACHHLEDVLDDLVVSLCKFTTLLNPSSVEEPVLAFGDDTKARMATISVFTIANKYGNYIRTGWRNILDCILR
LHKLGLLPARVASDVADDSELSAESGHGKPVTSSLSASQMPSMGTPRRSSGLMGRFSQLLSLDTEEPRSQPTEQQ
LAAHQRTLQTIQKCHIDSIFTESKFLQADSLLQLARALIWAAGRPQKGNTSPEDEDTAVFCLELLIAITLNNRDR
IVLLWQGVYEHISNIVQSTVMPCALVEKAVFGLLRICQRLLPYKENLADELLRSLQLVLKLDARVADAYCEHITQ
EVMRLVKANTTHIRSQMGWRTITSLLSITARHPEASEAGFEALMFIMSDGGVHLSPANYVLCIDASRQFAESRVG
QADRSVRAVDLMAGSVTCLARWSKETKDAPGEDTAAKVSQDIGEMWLRLVQGLRKVCLDQREEVRNHALLSLQRC
LTGVEGVRLPHASWLQCFDLVLFTVLDDLLEIAQGHSPKDYRNMEGTLVLAMKLLTKVFLQLLHDLSQLTTFCKL
WLGVLSRMEKYMRAKIRGKRNEKLQELIPELLKNTLLVMKTKGVLQQRSALGGDSLWELTWLHVNNIAPSLQSEV
FPDQESEQRQGGETDASGLAASEDVVRT*

**>Macco_9095g21**

MTMGSLKLLAGIEANGQEPRHCIPSSTAFACMINSEIGALLAVVRRNVRWGGRYMTEDDQLEHSLIQSLKSLRKK
LFSWDRQWNAVDPSIYLQPFLDVIRSDEIGAPITGVALSSVYKILTLGILDPTTVNVKDAMHLVVEAVTSCRFEV
TDSASEEVVLMKILQVLLACMKSKAAFLLNNQHVCTIVNTCFRIVHQAGTKGELLQRIARHTMHELVRCIFLHLQ
HIHCTGKPSIQGAEPSVKPEVGVLEKDNTSEVKQPDSVNDSSENASISADSVGSLVNETAVRTGRNEEAVNDVPH
SMREPYGVACVVEVFQFLCSLLNVGDQIGMGPRLNWTEFDEDVPLFALGLINSAIELGGSTVSRHPELLALIKEE
LFPNLMQLGLSMSPLILSMVCNIILNLYLHLRTELKLQLEAFLSSVILRLAQSKHGDTSYQQQEVLMEALVDLCR
QKTFMAEMYANFDCDISCSNVFEDVSNLLSKSAFPVNSPLSSMHVLALEGLVAVVQGFSERIGSSPTFTEQTLLS
VEQYSPFWNEKCENYADPNSWVGFVRHRKQIKKALMMGVDHFNRDPKKGLEFLQRTHLLPNELDPQSVACFFRYT
TGLDKSIVGDFLGNHEDFWVKVLQEFAQTFDFQDMSLDIALRLFLETFRLPGESQKIQRVLEAFSERFYDQSPQI
LANKDAALVLSYSIILLNTDQHNVRVKKKMTEEDFVRNNRRINGGNDLPREFLLDLYNSICKNEIRMTPEQGVGP
EVVMPSHWINLMHKSKTAAPFITCDPKPFLDRDMFSVLSGPTLAAICVVFDHAEKEDIYQTCVDGFLAMAKIASC
SHLVNILDDLVISLCKFTTLLDPSQMDEPILAFADDTKAMMATITVFTIANNFGDYIRTGWRNILDCILSLHKIG
LLPSLVASNEQSSDPVSGKPIIQSASYVPVKTITSQRSSGFMGRFTQLLSLDTEQQNLEPTEEELEARQRAHLTI
EKCQVSTIFTDSMFLQSDSLLQLVQALIWVSGPSQKGKNGSPEDEDTAIFCLELLITITLYNRDRIVLLWPGVYE
HIAGIVQSATMPSAFVEKAVFGLLHICQRLLPYKENLADELLRSLQLILKLDARVADAYCEQITQEIADLVKTNA
MCIKSQMGWRTITSLLSITARHPEASDVGFEALAFVMSDGSHLSPVNYSLCVDACRQFTESRVGSTERSIHALDL
MAGSLTYLTQWKLSPQEAGEIWLGLVKGLRKVSLDQREEVRNHSILSLQRCTAGIEGFSLPQDSWVNFFDLIVFQ
MLDDFLEIVQGKSSRDYRNIENTLLHALKLSSKVFLQQIMDLSPSSSFPKLWMGILGRMEIYTTVKIRGKKSEKL
RELVPELLKNMLLVMKSSRVLVNPSSSSPSSDTTLWEMTWPLIDNLVPNLKSEVFPDREPELTHRQKDTATPTLS
DVNTSTASIQS*

**>Macco_1789g18**

MARAIDEESEEGNDSSSLQKNKRISKLKKKELGISCMLNTEVGAVLAVIRRIPDSNSQFLSTPEENFDSSLLSSL
KSLRALIFNPQQEWHKIDPWIYLGPFLDVIQSDEIPAAATGVALSAILKILKLEIFNKKTPGARDAINSIVTGVT
SCRLEKTDSTSEEAVLMRVLQVLIGIMRNGASILLTDHAVCTLVNTCFQVVQQSASKGDLLQCNARHTMHELVQT
IFAGLPEIDVGENSESENEDDEILPNPDSGYGARCMVDIFHFLCSLLNVVEVVDTDGSTATNSDEDVQLFALILI
NSAIELGGERIRKHPKFLRMIQDDLFHHLIHYGMCSSPLLLSMICSTVLNIYHFLRRCVRLQLEAFFTYVLLRVA
NGGSSSQLQEVALEGIISFCRQPTFIIEMFVNYDCDPVFRNVFEEIGKLLSKIAFPVPNNPLSSLQIHAFEGLLI
TIHNIADNIDEDANPTSTPPYPVLVTEYKPFWIGESKNNENIERWIESLRFRKAQKKKILIAANHFNRDEKKGLD
YLKIAHLVPDPPDPKAFAWFYRFTPGLDKNKIGDFLGDPDELHLQVLKEFTDTFNFNGMILDTALRTYLETFRLP
GESQKIQRILEAFSEKFYDQQSSEIFVSKDAVFILCYSLIMLNTDQHNPQVKKKMTEEEFIRNNRAINGGKDLPR

EYLSELFQSISNNAITLFGQSGAPLEMNPNRWVDLINRAKVMEPFILCNFTNCLGRDMLASISGPTVAALSAIFE
HADEDETLHECVEGLFSVARIARFGLEDTLDELLASFCKFTMLLNPYSTAEETLYAFSNDMKPRMATLGVFTIVN
KFGDSIRGGWRNIVDCLLKLKRLKLLPQSVVEPEIASSSSSDIQRDSKSESGVIFSSFDIGHGSSRQMSGLIGRF
SHFMSLDCTEDLLLQAAGNDFEQNLKIIQQCRIANIITNSPNLPEDSLESLGRSLLFAAAGKGQKFSTPVEEEET
VGFCWDLIITITLANIHRFPTFWPSFNDYLILVAQFPLFSPCPFAEKAILGLFRVCLKLLATYRSEKLSEELIFK
SINLMWKLDKEILDTCSETITQSVSKMLTEYPANIQTQLGWKSVLHLLSISGRHPETYDQGVEALIMLMSDGYHV
SRTNYAYCIDCAFGFAALKISPLEKSWKILDLMADSVNFIIQWYRNGYTDPGSNMSNTSSSSFEDGSKNLNSVNF
TMNLFIKLVESLRKTSLVRREEIRNHAVLALQRSFTLADELDFTSSNCINCFNLVIFAMVDDLHEKLVEYSRREN
STKEMKSMAGTLKLAMELLKEVFLQFLKPLSQSSGFRQFWLGVLRRMDTCMKTELGDDDESGLQGLVPELLKKII
IEMKEKEILVQTEDNELWDITHVQIQWIAPSLKEELFPEEAI*
>**Cerde_20156**
MGRINLQSGIKAIEEEPENGEYQPNGSSLACMINSEIGTVLAVMRRNVRWGGRYISDDDHLEHALIQSLKALRRQ
IFSWKHHWHTISPTVYLQPFLNIIQSEETGAPIAGVALSSVYKILTLDMLDVNTVNIEEAMHSVVDAVTVFHHES
ADPASEEVVFMKKLQVLLACMKSRASSLLSNQHVWKIVHACFTIVHQAGSKGELLQRIARHTMHELIRCIFTHLP
EVDCLEHHRTNGGGNSIKKEVXXDKDYTNKLENGNDNAEYDGAPAAVAVASNSLDSTGSSANLVDGNTLEFGSVK
DTNDILLDLGPYGIPCMVKIFHFLCSLLNIVDNIEMGPSLNPQEFEEDVPLFALALINSAIELGGVSLRRHPKLL
SLIQDELFRNLMQFGLSTSPLILSVVCSIVLNLYNHLRTELKLQIEAFFSCVILRLAQSRNGASYQQQEVVMEAL
VDFCRQKTFMAEIYANLDCDITCNNVFEDLANLLSKSAFPVNRPLSSMHELALDGLIAVIQGMAERISNSSPISE
QPVELDEYTPFWTVKCENYSDPSSWVGFIRRRKYIKRKLMIGADHFNRDPKKGLEYLQGTHLLPDKLDPQSVACF
FRYTAGLDKNLVGDFLGNHDEFCVKVLHEFADTFDFQGMHLDTSLRLFLETFRLPGRVLEAFSERYYEQSPQILA
NKDAAFILSYSLIMLNTDQHNIQVKKKMTEEDFIRNNRRINAGNDLPPEFLSELYHSICKNEIRTIPEQGGTFPE
MTPSRWIDLMHKSRKTSPFILCDSRGFLDHDMFIIMSGPTIAAISVVFDHAEDEDLLRKCIDGFMAVAKISACHH
ledvlddlvVSLCKFTTLMNPSSMDEPVLVFGDDAKARMATVTVFTVANKFGDFIRTAWRNILDCILRLHKIGLL
PARVSSDAADDPELPMDIVHGKPTSNSLSVSHLPAMGTPRRSSGLMGRFSQLLSLDTEEPRSQPTEQQLAAHQRT
LQTIQKCHIDSIFTESKFLHAESLKQLAKALIWAAGRPYKANNTPEDEDTAVFCLELLIAITLNNRDRIILLWRD
VYDHIASIVQSTTMPCALVEKAVFGLLRICQRLLPYKENLADELLRSLQLILKLDARVADAYCENITQEVTRLVK
ANATHIKSQLGWRTITALLSITARHPDASEAGFEALTFVMSEGLHLSPANYLLCVEAARQFAESRVGLTDRSLRS
LELMAGSASCLWKWFPKTEGTEIALKASQDIGDMWLKLLQGLIKICMDQREEVRNRALLSLQSCFVVHESISLPT
ASWLQCldliftllddllDKAQGHSQKEYRNIEGTLSLAMKLVSKLFLQLLPELYHIPSFGKLWLGVLGRMEKYM
KAKIRGKKSEKIHELIPELLKNILFVMKAKGILEQKSTLGGDSLWELTWPHVNNISSTLQTEVFPDQERGITMKQ
SAKGNSSLSDPAILEEEKTGLAGT*
**>Cerde_25991**
MGRINPQSGIKAIEEEHENGEYQPNGSSLACMINSEIGTVLAVMRRNVRWGGRYISDDDQLEQYYHYSADRYISW
KHHWHITTNYLSEEVVFMKKLQVLLACMKTRASSLLSNQHVWKIVHACFTIVHRAGSKGELLQRIARHTMHELVR
CIFTHLPEVDCLEHHKTNGGGNSIKKEVGADKDYTNQLNNGNDNTEYDGAPAAAAFASNSLVSTGPSTNLLDGNT
LEFGTFKNTNDILLDLGPYGIPCMVKIFHFLCSLLNIVDNIEMGPLSNPQEFEEDVPLFALALLNSAIELGGASL
RRHPKLLSLIQDELFRNLMQIGLSTSPLILSVVCSIVLNLYNHLRTELKLQIEAFFSCVILRLAQSRNGASYQQQ
EVVMEALVDFCRQKTFMAEMYANLDCDITCNNVFEDLANLLSKSAFPVNRPLSSMHELALDGLIAVIQGIAERIS
NASPISEQPVELDEYTPFWTVKCENYSDPSSWVGFIRRRKYIKRKLMIGADHFNRDPKKGLEYLQGTHLLPDKLD
PQSVACFFRYTAGLDKNLVGDFLGNHDEFCVKVLHEFAATFDFQGMNLDTSLRLFLETFRLPGEAQKIQRVLEAF
SERYYEQSPQILANKDAAFILSYSLIMLNTDQHNIQVKKKMTEEDFIRNNRRINAGNDLPPEFLSELYHSICKNE
IRTIPEQGGTFLEMTPSRWIDLMHKSRKSSPFILCDSRGFLDRDMFIIMSGPTIAAISVVFDHAEDEDLLRKCVD
GFMAVAKISACHHLEDVLDDLVVSLCKFTTLMNSSSIDEPVLVFGDDAKARMATVTVFTIANKFGDFIRTAWRNI
LDCILRLHKLGLLPARVSSDAADDPELPMDIVHGKPTSSSLTVSHLPAMGTPRRSSGLMGRFSQLLSLDTEEPRS
QPTEQQLAAHQRTLQTIQKCHIDSIFTESKFLHAESLKQLAKALIWAAGRPYKANNTPEDEDTAVFCLELLIAIT
LNNRDRIILLWRDVYDHIASIVQSTTMPCALVEKAVFGLLRICQRLLPYKENLADELLRSLQLILKLDARVADAY
CENITQEVMRLVKANATHIRSQLGWRTITALLSITARHPDASEAGFEALTFVMSEGLHLSPTNYLLCVEAARQFA
ESRVGLTDRSLRSLELMAGSASCLWKWFPKTEGSEIALKTSQDIGDMWLKLLQGLIKICMDQREEVRNRALLSLQ
SCFVVNESVSLPTASWLQCLDLIFTLLDDLLDKAQSHSQKDYRNIEGTLSLAMKLVSKLFLQLLPELYHIPSFGK
LWLGVLGRMEKYMKAKIRGKKSEKIHELIPELLKNILFVMKAKGVLEQKSTVGGDSLWELTWPHVNNISSTLQSE
VFPDQERGAAMKQSAKGNSSLSDPATLEEEKTGLAGT*
**>Cerde_10541**
MVKARDQSHGEKKPKFPNNCKENRKRTNELAISCMLNTEIGAVLAVIRRNSFDSNSSHLPPNEDSYDSPLLNSLK
LLRAHIFNPQREWHKTNPSTFLSPFLDVLQSDDIPSSATGVALASILKILRLNIFDEKTPGAREAINSVVYGITN
CRLEKTDSSLKEAVLSRILQVLVTTMKNPASILLTDHTVCTIVNTCFQVIQQSASRGDLLQRNARHMIHELIQLI
FSRLPGLEVSDEGISEFDLEENEMENGSSSSYGIYCMVDIFHFLCTLLNVVEIVDQDGTAGTNSDEDVQLFALVL
INTAIELGGESIGKHRKLLRLVQDDLFHHVIHYGTCCSPLVLSMICSTVLNLYHYLRRFLKLQLEAFFTFVVLRI
ANGGSSPPLQEVAIEGLVSFCRQPDFIFEMYVNYDCDPLRRNVFEDIGKLLCKSSFLINTPVSGMQAQALEGVVA
IVQKISDNIGNWDSPGPRSTVNHIKTYRPFWEEKCENEEHLETWIEFIRLRKLQKRKIMIAANHFNRDEKKGLQF
LKISHLSSDPPDPKSIAFFYRYTPGLDKNKIGDYLGDPDELHIQVLKEFTETFDFSGIILDTALRTYLETFRLPG
ESQKIQRILEAFSERFYDQQSSEIFVSKDAVFILCYSLIMLNTDQHNPQVKKKMTEEEFIRNNRAINGGKDLPRE
YLSELFQSISNNAITLFGQSGTVMDMNPTRWDELIKRSKIIAPFIERSFEPCITTDLFACISGPSMAALSTIFEH
SDEDEILHECLEGLMSIAISAQYGLQDILDELLASFCKSTALLNPYASAEETLFAFNNDLKPKLATVAVFSICNL

FGTSIRSGWKNMVDCLLKLKRLKLLPQSMIDPYNISTSTNQCLQSKSEQGLMFPSTGNDTHTYGMLGSFSQFLMS
ENKDDSIGNSGCESDENLTIIHQLRIGNIFRESSKLPDESLFCCSRKGQKFVTDVEEEETIGFCWDLLVTISILN
LHRFSIIWPFVHDYLLDAEQFPRFSPCLFSEKAILGLFKICLRLFNSPKYDKLSEELIFKSINYAWKLVMELFEP
RAKEIIDPNRQQMLAKSITQSIIQILVDFPGKVQTQIGWKTLLQLLSITSRNPETYEQGVEAVTSLISDGIYITK
WNYAYFIDSAFGFATLRIIPTDKSTKILDLMANSVHLIVQWYGSAYTDPGSESSNISNSSIDDGLKTGNSNHFAV
NLFIKLGETLRKTSLVRREEIRNHSLYALRKSFAMSDEIGFTSPNCMHFFNHVLFAMVDDLHEKNVEYSMRENSD
REIRSMVGTLKIAMELLVDIFIQFLSILYENPGFRTFWMGMLRRMGTCMKADFGVEENILLRELVPRLLKKMIVK
MKEKNILVPREEDDLWEITYIQIQWIL*

**>Pipni_8.273**

MGRLKLQTGIKAIEEEPSDCESISNKGALACMINSEVGAVLAVMRRNVRWAGRYMAGDDQLEHLLIQNLK
ALRKQIFSWKLQGNMINPVAYLQPFLDVIQSDETGAPITGVALSSVYKILTLDILDLDTVNVVGAMHSVV
DTVTSCRFEVTDPTSEEVVLMKILQVLLACMKSKASVVLYNQHVCTIVNTCFRIVHQAGTKGELLQRIAR
HTMHELIRSIFSHLSDLDHSEGSQGNGGMSPIKPMTGVQGNDAMMERSMENGNAAATYNPPPALLGSTSN
LLVTNFGEETAINNKDFSSDLLASEPYGIPCMVEIFHYLCSLLNYVDQIEMGPRSNPIAYDEDVPLFALG
LINSAVELGGPSVQHHPKLSSLIQDELFRNLMQFGLSMSPLILSMVCSIVLNLYHHLRSELKLQLEAFFS
CVVLRLAESKYGASYQQQEVIMEALVDFCRQKHFMVEMYANLDCDITCANIFEDLANLLSKSAFPVNCPL
SAMHVLALDGLIAVIQGMAERISNGSPVQEKIPVALDEYSPFWTEKCGNYSDPSHWVSFVRRRKYIKRRL
AVGADNFNRDPKKGLEFLQGSHLLPDKLDPQSVACFFRYTSGLDKNLVGDFLGNHDEFCVQVLQEFANTF
DFQDMNLDTALRLFLETFRLPGESQKIQRVLEAFSERYYVQSPHILANKDAAFVLSYSLIMLNTDQHNVQ
VKKKMTEEDFIRNNRRINGGNDLPREFVSELYHSICKNEIRTIPEQGAGFPEMTPSRWTDLMRKSKRTAP
FILCDDGAHLDRDMFAIMSGPTIAAISVVFDHAEQEDVFTTCVEGFLAMAKISACHHLGDVLDDLVVSLC
KFTTLSTPSSVDEPVLAFGDDVKARMATETVFTIANRYGDYVRTGWRNILDCILRLHKLGLLPARVASDA
ADDLDLSSELAQGKPVMGSSITASHIPSIGTPRRSSGLMGRFSQLLSLDTEEPRSQPTEQQLAAHQRTLQ
TIQRCHIDSIFTESKFLQADSLLHLAKALIWIAGRPQKGHSSPEDEDTAVFCLELLIAITLNNRDRIVLL
WQGVYEHIASIVQSTLMPCALVEKAVFGLLRICQRLLPYKENLADELLRSLQLIQKLDARVADSYCENIT
QEVMRLVKANATHIRSQVGWRNITSLLSITARHPEASEAGFEALLFVMSEGAHLSPANFILCVDASRQFA
ESRVGHPHRSMRALDLMAESVASLVRWSREIKEAGEESSKLCHEIGEMWLRLVQGLRKVCLDQREEVRNH
AVSCLQRCLMAEGICVTQALWLQCFDQVIFTMLDDLLEIAQSQKDYKNIEGSLVFALKLLSKVFLHLLTD
LSQATSFCKLWLGVLGRMEKYMKVKVKSKKNEKLPELVPELLKNILLVMKTKGVLVHRTALGGDSLWELT
WYHVKNISPAFQLEVFPDVDQTEARNASPVPIGESYSISLACLMRIDSIESKGRIREG*

**>Pipni_8.277**

MGRLKLQTGIKAIEEEPSDCESISNKGALACMINSEVGAVLAVMRRNVRWAGRYMAGDDQLEHLLIQNLK
ALRKQIFSWKLQGNMINPVAYLQPFLDVIQSDETGAPITGVALSSVYKILTLDILDLDTVNVVGAMHSVV
DTVTSCRFEVTDPTSEEVVLMKILQVLLACMKSKASVVLYNQHVCTIVNTCFRIVHQAGTKGELLQRIAR
HTMHELIRSIFSHLSDLDHSEGSQGNGGMSPIKPMTGVQGNDAMMERSMENGNAAATYNPPPALLGSTSN
LLVTNFGEETAINNKDFSSDLLASEPYGIPCMVEIFHYLCSLLNYVDQIEMGPRSNPIAYDEDVPLFALG
LINSAVELGGPSVQHHPKLSSLIQDELFRNLMQFGLSMSPLILSMVCSIVLNLYHHLRSELKLQLEAFFS
CVVLRLAESKYGASYQQQEVIMEALVDFCRQKHFMVEMYANLDCDITCANIFEDLANLLSKSAFPVNCPL
SAMHVLALDGLIAVIQGMAERISNGSPVQEKIPVALDEYSPFWTEKCGNYSDPSHWVSFVRRRKYIKRRL
AVGADNFNRDPKKGLEFLQGSHLLPDKLDPQSVACFFRYTSGLDKNLVGDFLGNHDEFCVQVLQEFANTF
DFQDMNLDTALRLFLETFRLPGESQKIQRVLEAFSERYYVQSPHILANKDAAFVLSYSLIMLNTDQHNVQ
VKKKMTEEDFIRNNRRINGGNDLPREFVSELYHSICKNEIRTIPEQGAGFPEMTPSRWTDLMRKSKRTAP
FILCDDGAHLDRDMFAIMSGPTIAAISVVFDHAEQEDVFTTCVEGFLAMAKISACHHLGDVLDDLVVSLC
KFTTLSTPSSVDEPVLAFGDDVKARMATETVFTIANRYGDYVRTGWRNILDCILRLHKLGLLPARVASDA
ADDLDLSSELAQGKPVMGSSITASHIPSIGTPRRSSGLMGRFSQLLSLDTEEPRSQPTEQQLAAHQRTLQ
TIQRCHIDSIFTESKFLQADSLLHLAKALIWIAGRPQKGHSSPEDEDTAVFCLELLIAITLNNRDRIVLL
WQGVYEHIASIVQSTLMPCALVEKAVFGLLRICQRLLPYKENLADELLRSLQLIQKLDARVADSYCENIT
QEVMRLVKANATHIRSQVGWRNITSLLSITARHPEASEAGFEALLFVMSEGAHLSPANFILCVDASRQFA
ESRVGHPHRSMRALDLMAESVASLVRWSREIKEAGEESSKLCHEIGEMWLRLVQGLRKVCLDQREEVRNH
AVSCLQRCLMAEGICVTQALWLQCFDQVIFTMLDDLLEIAQSQKDYKNIEGSLVFALKLLSKVFLHLLTD
LSQATSFCKLWLGVLGRMEKYMKVKVKSKKNEKLPELVPELLKNILLVMKTKGVLVHRTALGGDSLWELT
WYHVKNISPAFQLEVFPDVDQTEARNASPVPIGESYSISLACLMRIDSIESKGRIREG*

**>Pipni_6.1394**

MANARLVAAEKEDENDVAGGGDDHANPRQTTTTLSKPGRPLGLACMLNIEISALLVVLRRDQSLPGHFLP
SPEETLDPHLVNSLKSLRATIFNSQHQWQTAHPATYLSPFLDVIQADEVPATATSIALYAVLKILNFDVF
DDGTPGAREAVHSVVTALTNCRLEKTDPEAEDAVLMRVLQVLVAVVRTPAAVLLSDHAICTLVNTCFQVV
HHSSNRADMLKRTARQAMHEMIQVIFARLPEVEEHEEGCLESEGEEGECGANKSGYGATCMVDIFHFLCS
MLNVAQVMDGNGIAAANAEEDKHLFALILINSTIDLGGGEFRNHPKLLGTIQDELFYHLIHYGAVASPIM
LSMVCSTVLNLYQFLRNHLHFQIEAFFTYVLLKTASNGGSTSQQEVAIEAIVNFCRQPRFIIEAYVNYDC
DPLCKNILEETGKLLCKSAFPLSSPLGSAQVQAFEGLVAIVQNIADHVDEDDGDDIYDYKPPRLLEYRPF
WLEKCEKGEDLDEWVDFHRIRKLQKKKIMIAADHFNKDEKKGMDFLKHAHLVPSPPDAKSLALFFRYTPG
LDKNKIGDFLGDPDELNIQVLKEFTATFDFSGVILDTALRTYLETFRLPGESQKIQRILEAFSDVFFEQQ
NSDVFVSKDAVYILCYSLIMLNTDLHNPQVKKKMTEEEFIRNNRAINGGSDLPRAYLSELFQSILTNEIT

IFGERGAGAGMNPNRWADLMKRWRGLEALVVCDFNSRLSRDMFVAFSGPAIATLSAIFEHADNDEVLHEC
MEALFAAARIARHGLPEILDELLACLCKFTTLLNPYAAAEETLFAFSNDLKPRMATLAAFTIANKFGSSI
RGGWRNIVACLLKLKRLKLLPQSVIEPQTYLSSSSSPPMPPADSDVAFPTSQGVDGGPRQSSSIMGRFSQ
FLSMEATEDHAPHGGNDFEQNLKIIQQCGIGNIFNDSGHLPDESLKSLGRALIFAAAGKGQKFSTPVEEE
ETVGFCWELLAAITFINIHRFRIIWPSFHDYLVSVTQFPLFSPTPFSEKAVVSLFNICLKLFTSSLIDKQ
SEDLILKSINMVCKLDKEVLDVCCEHITQSINRILVEFHDQIQTANGWKMLLHFLQFSSRPEKTFDQGVE
ALISLFSSDNRLNETNYLVCIEYAYHFAASKSTPFETRMRILDIMAESANLLVNWAQTRHGDAGNNSSSN
SANEECAKGSMTNLYLNMFIKLAESFKTISLARREELRNHSVVSMGRIFIIAEELGLTSAACSQCFNSVL
FAMVDDLHEKMVQYSMRLNSEEVTSMEGTMMLAMDLLVEVYLQFLKPLSELPEFRQFWLGLLRRMDTCMK
LNLGPSYDNRLRELVPELLKKMIVVMKETGVLVQTNENDLWVTTDFQIQWIAPSVRDELFS*

**>Arifi_013188**

MGRLKLQSGIKAIEEEPSNSESISNSAALSCMINSEIGAVLAVMRRNVRWGGRYMAGDDQLEHSLIQNLKSPRRQ
VFSWQHQWHTINPIIYLQPFLDVIRSDETGAPITGVALSSLHKILTLDILDLDTVNVVDAMHLVVDAVTSCRFEV
TDPASEEVVLMKILQVLLACMKSKASIVLNNQHVCTIVNTCFRIVHQAGTKGELLQRIARHTMHELVRCIFSHLP
DINETKHPAINGSTAPVKQEGSLQVENGNGSSDYDGPTSVGVFGSNNIASKVEDEVASGLASTKELSNDHLATEP
YGIPCMVEIFHFLCSLLNIVEQIEMGPRSNPVAFDEDVPLFALGLINSAIELGGPSICQHPKLLSLIQDELFRNL
MQFGLSMSPLILSMVCSIVLNLYHHLRMELKMQLEAFFSCVILRLAQSRYGASYQQQEVVMEALVDFCRQNQFMP
EMYANLDCDITCSNVFEDLANLLSRSAFPVNSPLSAMHVLALDGLIAVIQGMAERIASGSPAQEHNHMELEEYSP
FWTVKCDNYSDPNHWVGFVRRRKYIKGRLAIGADHFNRDPKKGLEFLQGNHLLPDKLDPQSVACFFRYTAGLDKN
LVGDFLGNHDEFCVQVLHEFAGTFDFQDMNLDTALRLFLETFRLPGESQKIQRVLEAFSERYYEQSPQILANKDA
ALLLSYSLIMLNTDQHNVQVKKKMTEEDFIRNNRRINGGSDLPREFLSELYHSICKNEIRTIPEQGAGFAEMTPS
RWIDLMRKSKNTAPFILCDGGAHLDHDMFAIMSGPTIAAISVVFDHAEQEDVLKTCVEGFLAMAKISACHHLEGV
LDDLVVSLCKFTTLLNPSSVEEPVLAFGDDVKARMATVTVFTIANRYGDYIRTAWRNILDCILRLHKLGLLPARV
ASDAADDSDLSSDPSHGKPVTGTSLSSSHIPSMGTPRRSSGLMGRFSQLLSLDTEEPRSQPTEQQLAAHQRTLQT
IQKCHIDSIFTESKFLQEDSLLHLAKALIWVAGRPQKGNNSPEDEDTAVFCLELLIAITLNNRDRIVLLWQGVYE
HIASIVQSTVMPCALVEKAVFGLLRICQRLLPYKENLADELLRSLQLILKLDARVADAYCEHITQEVLRLVKANA
THIRSQMGWRTITSLLSITARHPEASEAGFEALLFIMSDGAHLTPSNYVLCVDAARQFTESRVGLADRSMRSLDL
MAESAASLAKWSHEIKEAGEEGSKFSQEIGEMWFRLVQGLRKMCLDQREEVRNHAVSSLQRCLTGVDGICITQGL
WLQCFDQVIFTMLDDLLDIAQGHSPKDFRNMEGTLVLAMKLQTKVFLHLLKDIFQATSFCKLWRGVLTRMEKYLK
AKVRGKKSEKLQEQVPELLKNTLLVMKTEGILVQRSALGGDSLWEQTWINVKNIAPSLQSEIFPEEEHTEVGNGG
EVAQRGGS*

**>Arifi_020970**

MTIVRNPKKKEDEPQDKNSSPRPPWETKSDPKQLRDQGLACMLNTEVGAVLAVIRRTQDLSSHIPSDDAYASSLI
HSLKSLRALIFGPQQEWHRIDPTAYLSPFLEIIQSEEIPALATGVALSAILKILKLDIFDEKTSGARDGINSVVL
GLTNCRLEKTNSVSEDAVMMRILQVLMAVMKNPASVLLTDHAVGTVVNTCFNIVQQSAARGDLLQRNARHTMHEL
IQVIFARLPQVEASQEDGLESDAEDGDIASGYGAVCMVNIFHFLCSLLNVVEVVYGEGMTLSSDENVQFFALILI
NTAIELGGQSIEKHPKLLRMIQNDLFHHLIHYGMCSSPLVLSMICSTVLNLYLFLRRHLRLQIEAFFTYVLCKVA
TNYNAPQLQDVAIEGMINFCRQPNFTVEAFVNYDCDPICRNIFEDTMKLLCRTAYPTSSPLSHMQVQAFEGLVAV
IYTLADHMNTEKQMDMGVRRTELTEYTPFWVGKCDNFEDLDTWVEFMRRRKLQKRKIMIAANHFNRDEKKALDFL
KISHILPNPPDAKTLAHFYRYTPGLDKNKIGDFLGDPDEFNVEVLKEHSNTFEFSGMILDTALRTYLETFRLPGE
AQKIQRILEAFSERFYEQQEQELFVSKDAVFILCYSLIMLNTDQHNPQVKKKMTEEEFIKNNREINGGKDLPRDT
LSELFQSISNNAITIFGQNGGTVEMNPNRWTALIKRSNRAEPFMLCNFEPRICREMFAAISGPTIASLSAIFEHG
DEDEMLQECMEGLLAVARIARHGLGDILDELIACLSKFTTLLNPYATEQETLFVFGNDLKPRMATLSVFSIANQA
GDVIRGGWRNILDCMLKLKKLKLLPQLVPTDPDSASSGACNNDTMQHSSSESGIIFPSLNSGFSSGHHLSTATAR
FSQFLAMDNTEDALSVGNDYEHTLRIIQQCRIGSIFTESGTYSCESLQDLGRALIFAAAGKGQKFSTPIEEEETT
GFCWELIIVIACANLHRFPMFWPSFDEHLVVVSQLSLLSSCPFPEKAMKALFRICYKFLSSNQSDALSEELILKS
MDLVYKMDKESIEACCEAITKIFKECPKNFQNPVLGKFLLYMLAITGRHDLTFQCLISLLSDEKKLDKTNYLFIV
DGACAFAMMCSEKSFRILDSVAESVNVLLQWHRTGYSDPGSNASNMSSSSSTEDWSRANMIPYPYPLGLFFKAAQ
MLITVCKDSREDMVKNHAICALQKMFGLGEELVFACTSPISCFTSVLFPLVDELHAKIRENSHLPKSEATEFMTV
SVEDTLLLALRLMAEVFVQFLRPLAEGPGFVNFWQTGILRKMDYYMKADLGYATRLQEVVPEMLKLMITAMMEKN
ILVRKEGDELWDITYLQIQWIAPSLEDDLFSDLKVS*

**>Arifi_003805**

MAEAENSKLPEEDGEEKATEGAEEEEVEEAEEAEAERAGGGGGGGGEEQEEEEEEGEEGEEEEAEEYLLIQPWKL
QSKEKQQKHLALACMLNTEVGAVLAVIRRSPDMENSQFLPSQDENLDPPLLNSLKYLRAFIFSLQQEWRTVDPST
YLSPFLYVIQSNDIPASATGVALSAVLKILKLEIFDQNTPGACNAVNSVVLGLTNCRLERTDPFAEDAVMMRILQ
VLTTIMKNPSSVLLTDDSVKAIVSACFQVVQQAANRTDLLQRHAKHTMHELIQVIFARLPDVEPIEGRADSNGHN
DAGYGARCMVDTFQFLCSLLDGSEETEIDGMTAVIDDEVQLFAVTLINSAIELGGEAIGKHSKLLKMIQDDLFYY
LIQYGVCSNPLVFSMICSTVLNLCNYLRGSLRLQLEAFFTFVLFRVAANSDSPQLQELAMEGLINFCRQPSFLIE
AYVNYDCDPIRRNVFEDIGNLICRNAYATSSPLSSIQVQALEGLVTIIYSILDNLIANNEDLPTGTIEVTEYKPF
WVERCENFDDSEVWVEFVRTRKQQKKKILLAANHFNRDKKKGLEFLKICHLIPDPPDSKSIAHFYRYAPGLDKKK
IGDYLGDPDEFHIQVLKEYTDTFVFTGMILDTALRTYLETFRLPGEAQKIQRILEAFSEKFYDQQPSEMFASKDA
VFILCYSLIMLNTDQHNPKVKKKMTEEEFIRNNRSINGGNDLPREYLSALFYSISNNAITLFAHKGAAMEMNPNL
WVDLINRTNCVEPFFLCSWEPLICREMFVAISGPSVAALSAIFEQTDVDEILHECIEGFMSVAQIASHGLDDILD

ELFTCFCKFTGLFNPYASAEESIMAFSRDLKPKMAILSLFSIAQRYRASIRGGWKTIIDCLLKVKRLKLLPGSVI
DLNTPSSATSNEYLKRSQSETGVILPSFHSGFSSAHSVSSMMSCFSQFLSFDSSEDSSANVGNEFEQNLMIIQQC
RIGNLFCVSSDLPIECLQYLGRALIFAAAGKGQKFSTPTEEEEAVGFCWELLVAITSTYIQQFPKFWPSFHDHLV
AATEFTIIPNTPFTEKAIVGLLKICHKLLGSSQSEILNQDLIFESINLVWKIEKEVLDTCCESITEVISTILIEC
RSNIQSPLVWRMLLHFLRYTACHPEAYDQGVEVLMVLLSEENNLNRFNYWSCVECAFRLAILEDSPLEKSVKLVD
LMADSVKVLVQWHRLVDSDTESTVSSISNISSTSTEGFSKTNSYPLSVFMTVADVLRQICLSPREEIRNQAVLSL
SNIFLLAKELGFNPTNWVSCFNNLFFVTIDEIYEKMVEHSHRADEKEMKRMEGTLMLAFKLLVQAFLKLLLPLSK
SQDFLQLWKGFLRRMDALMKADFGFSCNTELQVLVPELLKKMVTEMKDNGVLVQAEGDELWEATNIHVQWIAPSV
KEELFPDHSKVITSRIEFHHFKDADGQKKMPTPTTGQYLLAIA*
**>LirchGNOMa_sc32**
MGRLKLQPGINSIEEEPGDCDCISNKGALACMVNSEVGAVLAVMRRNVRWGGRYMAGDDQLEHSLIQSLKALRKQ
IFSWEHQWQSINPTIYLQPFLDVIRSDETGAPITGVALTSVYKILTLDMLDLNTVNVEDAMHSVVDAVTSCRFEV
TDPASEEVVLMKILQVLLACMKSKASVVLNNQHVCTIVNTCFRVVHQAGTKGELLQRIARHTMHELVRCIFLHLP
DVHHTEQPSVNGSGASIRSEMGGFDKDHAFGVRQSDNGNGSLEYDAPSASVGFASNASMSTGSTANLADENTLGD
GSSKDTASDMHLTTEPYGIPCMVEIFHFLCSLLNIVEHSGVGPRSNPIVFDEDVPLFALGLINSAIELGGSSIRQ
HPKLLALIQDELFCNLMQFGLSMSPLILSMVCSIVLNLYHHLRMELKLQLEAFFSCVILRLAQSRHGASYQQQEV
AMEALVDFCRQKTFMAEMYANLDCDITCRNVFEDLANLLSKSAFPVNCPLSAMHILALDGLIAVIQGMAERIGNA
SPATEQASVTLEEYTPFWTVKCENYADPNHWVGFVRRRKYIKRILMIGADHFNRDPKKGLEYLQGTHLLPEKLDP
QSVACFFRYTAGLDKNLVGDFLGNHDDFCVRVLEEFAKTFDFQDMNLDTALRLFLETFRLPGESQKIQRVLEAFS
GRYYEQSPQILANKDAALLLSYSLIMLNTDQHNVQVKKKMTEEDFirnnrringgnDLPREFLSELYHSICKNEI
RTIPDQGGSFPEMTPSRWIDLMRKSKKTAPFITCDSRAFLDRDMFAIMSGPTIAAISVVFDHAEHENVFHTCVDG
FLAVAKISACHHledvlddlvvSLCKFTTLLNPSLVEEPVLAFGDDTKARMATITVFTIANRFGDYIRTGWRNIL
DCILRLHKLGLLPARVASDAADDSEPSTDPGHGKPVAGTLSASHIPAMATPRRSSGLMGRFSQLLSLDTEEPRSQ
PTEQQLAAHQRTLQTIQKCHIDSIFTESKFLQADSLLQLSRALVWAAGRPQKGSSSPEDEDTAVFCLELLIAIAL
NNRDRIGLLWQGVYEHIANIVQSTVMPCALVEKAVFGLLRICQRLLPYKENLADELLRSLQLVLKLDARVADAYC
EHITQEVTRLVKANATHIRSPMGWRTITSLLSITARHPEASEAGFEALMFIMSDGVHLSPANYILSVDAARQFAE
SRVGPTERSVRALDLMAESIMCLPRWKEVGEDAPRASSPEIEDMWLRLVQGLRKVCLDQREEVRNHALQSLQRCT
TTNAEAIRLSPASWLQCFDLVIFTMLDDLLEIAHGGHSAKDYRNMEGTLVHAMKLMAKVFLQLLPELSPLTSFCK
LWLGVLGRMEKYTKAKVRAKKCEKLQELVPELLKNTLYVMKNRGVLARTNAIGGDGLWELTWLHVNNIDPSLQSE
VFPDQELEQQARKENEMGSPLLQTAVEGGT*
**>LirchGNOMb_ctg60**
MGRLKLQPGINSIEEEPGDCDCISNKGALACMVNSEVGAVLAVMRRNVRWGGRYMAGDDQLEHSLIQSLKALRKQ
IFSWEHQWQSINPTIYLQPFLDVIRSDETGAPITGVALTSVYKILTLDMLDLNTVNVEDAMHSVVDAVTSCRFEV
TDPASEEVVLMKILQVLLACMKSKASVVLNNQHVCTIVNTCFRVVHQAGTKGELLQRIARHTMHELVRCIFLHLP
DVHHTEQPSVNGSGASIRSEMGGFDKDHAFGVRQSDNGNGSLEYDAPSASVGFASNASMSTGSTANLADENTLGD
GSSKDTASDMHLTTEPYGIPCMVEIFHFLCSLLNIVEHSGVGPRSNPIVFDEDVPLFALGLINSAIELGGSSIRQ
HPKLLALIQDELFCNLMQFGLSMSPLILSMVCSIVLNLYHHLRMELKLQLEAFFSCVILRLAQSRHGASYQQQEV
AMEALVDFCRQKTFMAEMYANLDCDITCRNVFEDLANLLSKSAFPVNCPLSAMHILALDGLIAVIQGMAERIGNA
SPATEQASVTLEEYTPFWTVKCENYADPNHWVGFVRRRKYIKRILMIGADHFNRDPKKGLEYLQGTHLLPEKLDP
QSVACFFRYTAGLDKNLVGDFLGNHDDFCVRVLEEFAKTFDFQDMNLDTALRLFLETFRLPGESQKIQRVLEAFS
GRYYEQSPQILANKDAALLLSYSLIMLNTDQHNVQVKKKMTEEDFIRNNRRINGGNDLPREFLSELYHSICKNEI
RTIPDQGGSFPEMTPSRWIDLMRKSKKTAPFITCDSRAFLDRDMFAIMSGPTIAAISVVFDHAEHENVFHTCVDG
FLAVAKISACHHLEDVLDDLVVSLCKFTTLLNPSLVEEPVLAFGDDTKARMATITVFTIANRFGDYIRTGWRNIL
DCILRLHKLGLLPARVASDAADDSEPSTDPGHGKPVAGTLSASHIPAMATPRRSSGLMGRFSQLLSLDTEEPRSQ
PTEQQLAAHQRTLQTIQKCHIDSIFTESKFLQADSLLQLSRALVWAAGRPQKGSSSPEDEDTAVFCLELLIAIAL
NNRDRIGLLWQGVYEHIANIVQSTVMPCALVEKAVFGLLRICQRLLPYKENLADELLRSLQLVLKLDARVADAYC
EHITQEVTRLVKANATHIRSPMGWRTITSLLSITARHPEASEAGFEALMFIMSDGVHLSPANYILSVDAARQFAE
SRVGPTERSVRALDLMAESIMCLPRWKEVGEDAPRASSPEIEDMWLRLVQGLRKVCLDQREEVRNHALQSLQRCT
TTNAEAIRLSPASWLQCFDLVIFTMLDDLLEIAHGGHSAKDYRNMEGTLVHAMKLMAKVFLQLLPELSPLTSFCK
LWLGVLGRMEKYTKAKVRAKKCEKLQELVPELLKNTLYVMKNRGVLARTNAIGGDGLWELTWLHVNNIDPSLQSE
VFPDQELEQQARKENEMGSPLLQTAVEGGT*
**>LirchGNL2_ctg1255**
MARARDQEEEEEDNGPWRTKGKKNPKPRKELNLACMLNTEVGAVLAAIRRNPDPNSQFLPSHEENYDYALLNSLK
SLRALIFNPQQEWHTIDPSIYLSPFLDVIQSDDIPasatavalsavlKILKLEIFDEKTPGASEAIHAIIAGITN
CRLEKTDPIPEDAVFMRILQLLMMVMKSHASVMLTDHAVCTIVNTCFQIVQQSASRGDLLQRNARHTMHELIQVI
FAHLPEIESSDGGCSESDIEDNEIMSGSRLGYSERCMVDIFHFLCSLLNVVEVVDAEGLGPTNSDEDFQLFALVL
INSAIEMGGEGIGKHRKLLRMIQNDLFHHLIHYGTCSSPIILSMICSTVLNLYHFLRRSLRLQLEAFFTYVLFRA
AAGGSPVQVQEVAIEGIISFCRQPTFIIEAYINYDCDPICGNVFEDIGKLLCKNAFTTSSPLSSMQVQAFEGLVS
IIHKVADSMDSGDSPGPVQPVDVSEYRPFWVEKCENYNDLETWVKFVRLRRFQKKKIMIAANHFNRDEKKGFEFL
KICKLVPDPPDPKSYACFYRYTPGLDKNKIGDFLGDPDELHLQVLKEFTDTFNFSGVILDTALRTFLETFRLPGE
SQKIHRILEAFSEKFYEQQSSEIFVSKDAVFILCYSLIMLNTDQHNPQVKKKMTEEEFIRNNRAINGGKDLPREY
LSELFLSISNNAITLFGQSAASLEMNPNRWVDLIKQSKVMEPFILCNFENRLSRELFVSISGPSVAALSATFEHA
DEDEILQECIEGLLSIARIAQYGLEDILDEILASFCKFTTLLNPYASAEETLFAFSNDLKPRMATLAVFTIANKF

GDSIRGGWRNLVDCllklkrlkllPQSVIEPEMASTSGTDHARHTKSESCVIFPSSHPGLGSSHQNSGLIGRFSQ
FLSLDSSNDDSMANTGSEFEQNLKIIQKCRIGNIFSKSSNLPDEALLNLGRALIFAAAGKGQKFSTPVEEEETVG
FCWELLFTVTSANIYRFLTFWPSFHDYLVLVAQFPLFSPCPFAEKAIIGLFKICLKLLSSPEYDKNHEELIFKSI
NLAWKLDKEILDTCCESITESVCQILNEQAASIQTLLGWKTILHLLSITARHPETFDEGVEAMMSLMSNETLVTK
LNYAYCIDCAFGFAALKMIPLEKSEKIIDLMADSVKLLIQWYQSGYSDPgsntsnfsntssssfeegsKPGNSGN
LPINLFIKLAETLRKTSLVRREEIRNHAISALQQCFLLADDLGFTPPSCLSCFNLVIFAMVDDLHEKMVEYSRRG
NAEREMRSMEATLKMAMELLVEVFLQFLKLLAQNPGFRTFWLGVLRRMDTCMKADLGGSYQSGFQELVPQLLKKM
ITVMKEKEILVQRDGDELWDITNIQIQWIAPSVKEELFPEDF*

**>PeramGNOM_ctg64F**

MGRLKLQSEISSIEEEPGDCEVMSNKGAFACMVNSEVGAVLAVMRRNVRWGGGGRYMAGDDQLEHSLIQSLKTLR
KQIFLWEHQWNTIDPAVYLMPFLDVIRSDETGAPITGVALSSVYKILTVGMLDLNTVNVESALHSVVDAVTSCRF
EVTDPASEEVVLMKILQVLLACMKSKASIVLSNQHVCNIVNTCFRVVHQAGTKGELLQRIARHTMHELVRCIFSH
LPDIDHTKQAITDGGDPSIRTEVGGLDKDYAFGVKQSDNGNGSIEYDAPPSSVGFASNASNFSGSMEGLVDENAV
GYVSNKEATNDMHLTLEPYGVPCMVEIFRFLCSMLNVVEQTGMASRSNPIVFDEDVPLFALGLINSAIELAGPSI
HQHPKLLSLIQDELFCNLMHFGLSMSPLILSTVCSIVLNLFHHLRMDLKLQLEVFFSCVILKLAQSRHGASYQQQ
EVAMEALVDFCRQKSFMAEMYANLDCDITCRNVFEDLANLLSKSAFPVNCPLSVMHILALDGLIAVIQGMAERIE
NSTPVLEQAPVVIEECTPFWTVKCEDYADPNYWVAFVRRRKYIKRMLMIGADHFNRDPKKGLEYLQGTHLLPDKL
DPQSVACFFRYTSGLDKNLVGDFLGNHDEFCVRVLHEFAGTFDFEDMNLDTALRLFLETFRLPGESQKIQRVLEA
FSERYYEQSPHILANKDAALVLSYSLIMLNTDQHNVQVKKKMTEEDFIRNNRRINGGNDLPREYLSELYHSICKN
EIRTIPEQGTGFHEMTPSRWIDLMRKSKKTPPYITCDSRSFLDRDMFAIMSGPTIAAISVVFDHAEHEDVFNTCV
DGFLAVAKISACHHLEDvLDDLVVSLCKFTTLTNPSLVEEPVLAFGDDTKARMATVTVFTIANRFGDYIRTGWRN
ILDCILRLHKLGLLPARVASDAADDSEPSVDPVHGKPVASTVSASHIPTIATPRRSSGLMGRFSQLLSLDTEEPR
SQPTEQQLAAHQRTLQTIQSCHIDNIFTESKFLHADSLLQLARALVWAAGRPQKVSSSPEDEDTAVFCLELLIAI
ALNNRDRISLLWQGVYEHIANIVQSTVMPCALVEKAVFGLLRICQRLLPYKENLADELLRSLQLVLKLDARVADA
YCEHITQEVTSLVKANAMHIRSQMGWRTITSLLSITARHPEASDVGFEALMFIMSDGVHLSPTNYILCLDAARQF
AESRVGSTERSVRALDLMAESITCLSRWKELIEEASKTSQEIDEMWLRLVQGLRKVCLDQREEVRNHALSSLQRC
MTVGAEGIRLSLASWLQCFDQVIFTMLDDFLEITHGGHSTKDYRNMEGTLVHALKLMSKVFLQLLPQLSQLTSFC
KLWLGILGRMEKYTKAKIRGKKSEKLQELIPELLKNTLLVMKSRGILVRTSAIGGDGLWELTWLHVNNVDPSLQS
EVFPDQEQEVARKENENGSSLTPTVVES*

**>PeramGNL2_ctg13F**

MAKVRDQDQEEDGEGEAKEQNPKRLRELGLACMLNTEVGAVLAVIRRTPDPSSQFLPQPEENIDPAILNSLRSLR
ALLFNPLQKWHNIDPSFYLSPFLEVIESDNIPATATGVALSAILKILKLEIFNEKTPGAIDAMHSVVNGITNCRL
ETTDPISEDAVMMRILQVLLATIRNPASVLLSDHAVCTLLNTCFQVVQQSASRGDLLQRSARHTMHEMVQVIFAR
LPLVESSELDVETSTSISYGARCVGDIFHFLCSMLNAVEVVDADGMQTTNTDEDVQLFALVLINSAIELSGERIG
KHPKLMWMIQDDLFHHLIHYGLYSSPLVLSMISSTVLNIYHFLRRSLRLQLEAFFTYVLFRVAAGGSSVQLQEVA
VEGIISFCRQPTFIIEAYINFDCDPIRTNLFEDIGKVLCKIAFPISSPLSSMQIQAFQGLVAMIHNIADKMENDK
RPHPESYALEIPGYQPFWVERCQNYYDLETWvkfvrvrkfqkkkIMIAANHFNRDEKKGLEFLKISHLVSEPPDP
KSYAYFFRYTPGLDKNKIGDFLGDPDEFNLRVLKDFSETFEFSGVILDTALRTFLETFRLPGESQKIQRILEAFS
ERFYEQQSSETFVSKDAVFILCYSLIMLNTDQHNPQVKKKMTEDEFIRNNRAINGGKDLPREYLSELFQSISNNA
ITVFGQSSTSMEMNTSRWVDLMKRSKVVEPYLLCNFQHHLSRELFASISGPSVAALCAIFEHVDEDEILHECIDG
LISIARIAQFGLHDILDEllsslsksttllNPYASAEETLFAFSNDLKPRMATLAVFTIANKFGDSIRGAWRNVV
DCLMKLKRLKLLPQTVVEPNTESTTTHDLSKQSKSESGVTFPSSYTGLGSSRKASGLIGRFSHFLTLDSGDDSMT
SGGSEFEQNLKIIQKCRIGSIFSESSNLPEESLLSLGRALIFAAAGKGQKFSTPIEEEETVGFCWELLITITSAN
LHRFLTIWPSFHDFLLAVAQFPFFSPCPFAEKAIIGLFKICLKLLSSSQTDKIAEELIFKSINLAWKIEKEILDT
CCESITQSISKIVSEYGENLQTFLGWKTILHLLSLTARHPETYDEGVETLILLMSSGALITKLNYAACIDCAFGF
SALKISSLETSTKLIDLMAESVPMLVQWYQLEYSDPESNCSNFSSTTSSSSMEDGMKPNSSLAVNLFVKLADTLR
KTSLVRREEIRNHAVAALKRSFSSADELGFTPSSCINCFNLVIFAMVDDLHEKMVEYSRRGNAEKETKSMEGTLK
MAMELLVEVFLQFLKLLSLNPSFRTFWLGVLRRMDTCMKADLGDNYESQLQFLVPELLKKMITVMKEKEILVQRD
GDELWDITNIQMQWIAPSVKEELFPEDDF*

**>CKAN_00834500**

MGRLKLQSEISSIEEEPGDCEVMSNKGAFACMVNSEVGAVLAVMRRNVRWGGGGRYMAGDDQLEHSLIQSLKTLR
KQIFLWEHQWNTIDPAVYLMPFLDVIRSDETGAPITGVALSSVYKILTVGMLDLNTVNVESALHSVVDAVTSCRF
EVTDPASEEVVLMKILQVLLACMKSKASIVLSNQHVCNIVNTCFRVVHQAGTKGELLQRIARHTMHELVRCIFSH
LPDIDHTKQAITVGGDPSIRTEVGGLDKDYAFGVKQSDNGNGSIEYDAPPSSVGFASNASNFSGSMEGLVDENAV
GYVSNKDAANDMHLTLEPYGVPCMVEIFRFLCSMLNVVEQTGMTSRSNPIVFDEDVPLFALGLINSAIELAGPSI
HQHPKLLSLIQDELFCNLMHFGLSMSPLILSTVCSIVLNLFHHLRMDLKLQLEVFFSCVILKLAQSRHGASYQQQ
EVAMEALVDFCRQKSFMAEMYANLDCDITCRNVFEDLANLLSKSAFPVNCPLSVMHILALDGLIAVIQGMAERIE
NSSPVLEQAQVVIKECTPFWTVKCEDYADPNYWVAFVRRRKYIKRMLMIGADHFNRDPKKGLEYLQGTHLLPEKL
DPQSVACFFRYTSGLDKNLVGDFLGNHDEFCVRVLHEFAGTFDFEDMNLDTALRLFLETFRLPGESQKIQRVLEA
FSERYYEQSPHILANKDAALVLSYSLILLNTDQHNVQVKKKMTEEDFIRNNRLINGGIDFPREYLSELYHSICKN
EIRTIPEQGTGFHEMTPSRWIDLMRKSKKTPPYITCDSRSFLDRDMFAIMSGPTIAAISVVFDHAEHEDVFNTCV
DGFLAVAKISACHHLEDVLDDLVVSLCKFTTLMNPSLVEEPVLAFGDDTKARMATVTVFTIANRFGDYIRTGWRN
ILDCILRLHKLGLLPARVASDAADDSEPSVDPVHGKPVASTVSASHIPTIATPRRSSGLMGRFSQLLSLDTEEPR

SQPTEQQLAAHQRTLQTIQSCHIDNIFTESKFLHADSLLQLARALVWAAGRPQKVSSSPEDEDTAVFCLELLIAI
ALNNRDRISLLWQGVYEHIANIVQSTVMPCALVEKAVFGLLRICQRLLPYKENLADELLRSLQLVLKLDARVADA
YCEHITQEVTSLVKANAMHIRSQMGWRTITSLLSITARHPEASDVGFEALMFIMSDGVHLSPTNYILCLDAARQF
AESRVGSTERSVRALDLMAESITCLSRWKELVEDASKTSQEIDEMWLRLVQGLRKVCLDQREEVRNHALSSLQRC
MTVGAEGIRLSPASWLQCFDQVIFTMLDDFLEITHGGHSTKDYRNMEGTLVHALKLMSKVFLQLLPQLSQLTSFC
KLWLGILGRMEKYTKAKIRGKKSEKLQELIPELLKNTLLVMKSRGILVRTSAIGGDGLWELTWLHVNNVDPSFQS
DVFPDQEQEVARKENENGSSLTPTVVES*
**>CKAN_01177000**
MAKVRDQDQEEDGEGEAKARNPKRLRELGLACMLNTEVGAVLAVIRRTPDLSSQFLPQPEENIDPAILNSLRSLR
ALLFNPLQEWHNIDPSFYLSPFLEVIESDNIPATATGVALSAILKILKLEIFDEKTPGAIDAMHSVVNGITNCHL
ETADPISEDAVMMRILQVLLATIRNPASVLLSDHAVCTLLNTCFQVVQQSASRGDLLQRSARHTMHEMVQVIFAR
LPLVESSELDVETSTSISYGARCMGDIFHFLCSLLNAVEVVDADGMQTTNTDEDVQLFALILINSAIEMSGEGIG
KHPKLMRMIQDDLFHHLIHYGSHSSPLVLSMISSTVLNMYHFLRRSLRLQLEAFFTHVLLRVAAGGSSAQLQEVA
VEGIISFCRQPTFIIEAYVNFDCDPIRTNLFEDVGKVLCKIAFPISSPLSSMQIQAFQGLVAMIHNIADYMENDK
KPRPESYALEIPDYQPFWVERCENYDDLETWVKFVRVRKFQKKKIMIAADHFNRDVKKGLEFLKLSHLVSEPPDP
KSYAYFFRYTPGLDKNMIGDFLGDPDEFNLRVLKDFSETFEFSGVILDTALRTFLETFRLPGESQKIQRILEAFS
ERFYEQQSSEMFVSKDAVFILCYSLIMLNTDQHNAQVKKKMTEDEFIRNNRAINGGKDLPRECLSELFQSISNNA
ITLFGQSSTSMEMNTSRWVDLMKRSNVVEPYILCNFEHHLSRELFASIYGPSVAALCAIFEHVDEDEILHECIDG
LISIARIAQFGLHDILDELLDSLCKSTTLLNPYASAEETLFAFSNDLKPRMATLAVFTIANKFGDSIRGAWRNVV
DCLMKLKRLKLLPQSVVEPNTASTTTHDLSKQSKSESSVTFPSTYTGLGSSRKASGLIGRFSHFLTLDSGDDSMA
SGGSEFEQNLKIIQKCRIGSIFSESSNLPEESLLSLGRALIFAAAGKGQRFSTPIEEEETVGFCWELLITITCAN
LHRFLTIWPSFHDFLLAVAQFPFFSPCPFSEKAIIGLFKICLTLLSSSQTDKIAEELIFKSINLAWKIEKEILDT
CCESITQSISKIVSEYGENLQTFLGWKTSLHLLSLTARHPETYDEGVETLIFLMSSGSLITKLNYAACIDCAFGF
SALKISSLETSTKLINLMADSVPMLVQWYQLEYSDPESHCSNFSSTSSSSMEDGMKPNSSLAVNLFTKLADTLRK
TSLVRREEIRNHAVAALKRSFSSADELGFTPSSCINCFNLVIFAMVDDLHEKMVEYSRKGNTEKETKSMEGTLKM
AMELLVEVFLQFLKLLSLNPSFRTFWLGVLRRMDTCMKADLGDNYESQLQFLVPELLKKMITVMKEKEILVQRDG
DELWDITNIQMQWIAPSVKELFPEDDS*
**>ChisaGNOMa_chr9-59**
MGRLKLQTGIHSIEEEPGDTECMSNKGALACMVNSEVGAVLAVMRRNVRWGGRDDHQLEHPLIRSLKVLRKQIFS
WDHQWHTIDPTIYLQPFLDVIRSDETGAPITGVALSSVYKILTLDMIDLNTVNVETAMHLVVDAVTSCRFEVTDP
ASEEVVLMKILQVLLACMKSKASVVLNNQHVCTIVNTCFRVVHQAGSKGELLQRIARHTMHELVRSIFSHLPEID
QTETPAAHGGEPSIRPEMGGVDKNYASGIKHSDNGNGSLEYDSQQISVGFASNTSLSVACLVDEGAVGDGTHKEA
ASDMHLTTEPYGVPCMVEIFQFLCSLLNIVEQTGMGPRSNPIVFDEDVPLFALGLINSAIELGGPAICRHPKLLS
LIQDELFCNLMHFGLSMSPLILSMVCSIVLNLYHHLRTELKLQLEAFFSCVILRLAQSRYGASYQQQEVAMEALV
DFCRQKTFMAEMYANLDCDITCRNVFEDLANLLSKSAFPVNCPLSSMHILALDGLIAVIQGMAERIGSASPVPEQ
TSVDLEEYTPFWTVKCEDYSDPNHWVRFVRRRKYMKRMLMIGADHFNRDPKKGLEYLQGTHLLPDKLDPQSVACF
FRYTAGLDKNLVGDFLGNHDEFCVRVLNEFAWTFDFQEMNLDTALRLFLETFRLPGESQKIQRVLEAFSERYYEQ
SPHILVNKDAALLLSYSLIMLNTDQHNVQVKKKMTEEDFIRNNRRINDGNDLPRDFLSELYHSICKNEIRTIPDQ
GAGFPEMTPSRWIDLMRKSKKTAPFITCDSKAFLDRDMFAIMSGPTIAAISVVFDHAEHDDVFPTCVDGFLAVAK
ISAYHHLEDVLDDLVVSLCKFTTLLNPSLSEEPVLAFGDNAKARMATVTVFTIANRFGDYIRTGWRNILDCILRL
HKLGLLPARVASDVADDLEPSADPVHGKPIASSLSASHIPAMSTPRRSSGLMGRFSQLLSLDTEEPRSQPTEQQL
AAHQRTLQTIQTCHIVNIFTESKFLQADSLLQLARALVWAAGRPQKGNSSPEDEDTAVFCLELLNAITLNNRDRI
GLLWQLVYEHIANIVQSTVMPCALVEKAVFGLLRICQRLLPYKENLADELLRSLQLVLKLDARVADAYCEHITQE
VTRLVKANATHIRSQMGWRTITALLSITARHPEASDTGFEALTFIMADGVHLSPANYVLCVDASRQFAESRVGPT
ERSVRALDLMADSISCLSRWKELIEASKVSQEIEEMWLRLVQGLRKVCLDQREELRNHALASLQRCMTIGAEVIH
LSAAAWLQCFDLVIFTMLDDLLEIAHGGHSAKDYRNMEGTLVNALKLMSKVFLLLLPELSPLPSFCKLWLGILGR
MEKYAKAKVRGKKSEKLQELVPELLKNTLLVMKSRGILARTSTIGGDGLWELTWLHVNNVDPSLQAEVFPDQEHD
EAGKEKAMANSLLPTAVES*
**>ChisaGNOMb_chr9-13**
MIKLHSMINTIQEESTDCEHMSNRGALACMVNSEVGAVLAVMRRSVQWGGHYMDGDNQLEHSLIQSFKALRKQIF
LCEHQWLEVDPSLYLQPFLDVIRLDETGAPITGVALSSLYKILTFGILDLNTVNIESAMHSVVDAVTSCRFEVTD
PVSEEVVLMKILQVLLACMKSKASIVLDNQHVCSIVNTCFRVVHQAGTKGELLQRIARHTMHELVRCIFSRLPDI
DQTKQPMVNVGGTKVLVYCYYQMSGTEVVCAFEVKQSDNYKGSIEYDAPLESLGFASNASKDAGSVAGFVDESAI
RDVINSDQAKTVLLMTEPYGVPCMVEIFEFLCSMLNDVEQTGIGPRSNPLAFDEDVPLFALGLINSAIELGGPST
CRHPKLLSLIQDELFHNLMHFGLSMSPLILSMVCSIVLNLYHHLRTELKLQLEVFFSCVILRLAEIRHGVSYHQQ
EVAMEALVDFCRQNTFMAEMYANLDCDLTCHNVFEDLANLLSKSAFPVNCPLSTMHVLALDGLIAVIQGMADRIV
NACPILDDALVGLEEYTPFWIAKCDDYTDPSHWVGFVRQRKYIKRMLMIGVDHFNRDPKKGLEYLQGVHLLPDKL
HPHSVACFFRYTAGLDKNLVGDFLGNHDEFCVRVLHEFSGMFDFQDMNLDTALRLFLETFRLPGESQKIQRVLEA
FSERYYEQSPQILVNKDAALVLSYSLILLNTDQHNVQVKRKMTEEDFIRNNRHINGGDDLPREYLSKLYNSICKN
EIRTIPEQGVAFLELTPSRWIDLMCKSKKTAPFIICDSGNFLDRDMFIIMSGPTIAALSVVFDHAEHEDVFQTCV
DGFLSMAKISAhhhledvlddlvvSLCNFTTLVNPSLSDEPVLAFLYDAKARMATVTVFTIANRFGDYIRTGWLN
ILECILRLHRLGLLSTQVASNAVDDLEPSSINVNWKPITSSLSASPLPVFSTPWRSSGFMSRFTQLLSLDTEEPI
SQPTEEQLAAHQCTLQTIQMCHIDDIFTESKFLHADSLLQLARALVWAPGQCQKMSTPLEDEETAVFCLELLIAV

TLNNRDRITLLWQHVYEHIADIIQSTIMPCALVEKAVFGLLHICQRLLPYKENLADELLRSLQLVLKLDARVADT
YCEHITQEIMHLVKANATHIRSHMGWRTIASLLSITARHPEASDVGFEALTFIMSEGVHLSPINYILCLDASRQF
AESCVDSIERSVRALDLMAESVTCLSQWKELVDDASKFSSKEVEEMWLRLLQGLKKVCLDKREEVRNHALSSLQR
CMTAGAEGICVSPGSWLQCFDLVIFPMLDDFLEIVHGGHTAKDYRNMEGTLVHALKLMSKVFLQLLPELSLLPGF
CKLWLEILECMEKYMKAKVRGKKWEKLQELIPELLTNMLLVMKSRGNLVRTSAIGGDRLWELTWLHVNEIDPSLQ
SEIFPVQEQKVAKEENEKGSLLLPTVVEG*
**>ChisaGNL2_chr4**
MAKASDQDQEDEGEDKSPWGQRPKKNSKVKELGLACMLNTEVGTVLAVLRRNPDPNSQFLSPMEDNFDPSLLNSL
RSLRAVIFNPQQEWHSIEPTFYLSPFLEVIQSDDIPAAATGVALSAILKILRLEIFDEKTPGAANAMHLVVGGII
NCRLEKTDPMSEEAVMLRILQVLLATVKNRTSVLLSDHDVCTVVNTCFQVVQQSTNRGDLLQRSARHTMHEMIQH
IFHRLPLIEPTDGGGLESDIEDAEYTSSSNSGYGARCMVDIFHFLCSLLNVVEVVDADGMPTTNTDEDVQLFALV
LINSAIELGGEGIGSHIKLMCLIQDDLFHHLIHYGMYASPLVLSMICSTVLNLYHFLRRXLRLQLEAFFTFVLFR
VATTGCSIQLQEVAVEGIINFCRQPTFIIEAYVNYDCDPIRRNILEDIGRVLCKIAFPISCPLSSMQVQAFEGLV
SVIQNVADNIDNEKSHHPTSYALEVSTYQPFWVERCENYEDQETWVRFVRVRKFQKKKLMIAANHFNRDEKKGLE
FLKITHLVPDPPDPKSYACYFRYTPGLDKNKIGDFLGDPDEFNLQILKEFIDTFDFSCVILDTALRTFLETFRLP
GESQKIQRILEAFSEKFYEQQSSEIFVSKDAVFILCYSLIMLNTDQHNPQVKKKMTEEEFIRNNRAINGGKDLPR
EYLSELFQSISTNAITLVGQSPLSIDMNPNKWVDLIKRSKVVEPYILCDFEHHLSRELFASISGPSIAALSAVFE
QADEDEILNECIEGLISIARIAQFGLEDILDELLASFCKFTTLLNPYASTEETLFAFSNDLKPRMATLAVFSIAN
KYGDSIHGGWKNMVDCLLKLKRLKLLPQSVIEPDTASDNTHDLSKQSKSESSVTFPSTYTGLGSSRKASCFFGRF
AHFLSLDGSDDSMASIGSEFEQNLKIIQQCRIGRIFSESSNLSDESLLNLGRALIFAAAGKGQKFSTPIEEEETV
GFCWELLITATKNNLHRFGTIWPSFHDYLVSVAQFPLFSPCPFAEKAIIGLFKLCLNLLSSSQTDKTSEELIFKS
INLAWKLEKEILDTCCESITESIYKILSEYPEKIETLLGWKTILHLLSVTARHPETYDQGVDAIINLFSDGSHIT
RSNYAYCIDCAFGFAALKISPLEKSAKLIDLMADSVNMLIQWYGSGYSDPGSNISNTSNSSMEDGAKSTNQSNFA
VNLFIKLAETLRKTSLVRREEIRNHAVMALQRCFSLADELGFTPPSCINCFNLVIFAMIDDLHEKMVEYSRRGNS
ERETRSMEGTLKMAMELLVEVFLQFLKLLSQNPGFRTFWLGVLRRMDTCMKADLGDTHESKLQELVPELLKKMIT
AMKDKEILVQQDGDELWEITNIQIQWIAPSVKDELFPEEF*
**>Musac_LOC103986838**
MGRPRLQNSAINPIEEESPDAAEDGSLPPLPPSGGPAAAADLSCVICSEVSAVLAVMRRNVRWGGRYASAADDHH
LEHSLVQSLKSLRRQVFSWGDARRPWCTIEPSAFLRPFVDVVQSDETGAPITGVALSSLYKILTLDLLEPGSGTG
VEAGMHLVVEAVTSCRFEVIDPASEETVLMKILQVMLAIMRSQASIMLSNQHVCTIFNTCFRIVHQAGTKGELLQ
RFSRHTMHELVRCIFSHLPHVEDGGRPPSLKPEIGATDKDQAFGIKQVENGNGSAEPTDTSVRDDNPGSEAENGV
RTMVEPYGISCMVEIFHFLCSLLNVADHIGMSPATNQIAFDEDMPLFALGMINSAIELGGPSISKHPKLLSLIQD
ELFRNLMQFGLSMSPLILSMVCSIVLNLYRHLRTKLKLQLEAFFSCVILRLAQSRYGASYHQQEVAMEALVDFCR
QKTFMAEMYANLDCDITCSNMFEELANLLSRSAFPINCPLSSMHVLALDGLIAVIQGMADRIGNASPSFEQSPLE
LEEYSPFWTVKCANYSDPEHWVKFVRRRKYIKRRLMIGADHFNRDPKKGLEFLQGTHLLPEKLDPQSVACFFRYT
AGLDKNLVGDFLGNHDEFCVQVLHEFAWTFDFQDMNLDTALRLFLETFRLPGESQKIQRVLEAFSERYYEQSPQI
LVDKDAALVLAYSLIMLNTDQHNVQVKKKMTEEDFIRNNRRINGGNDLPREFLSELYHSICRNEIRTAPEQGFGF
TEMSPSRWIDLMQKSKKTSPYIVCDSRPFLDRDMFAIMSGPTIAAISVVFDYAEHEEVFLTCVDGFLAVAKISAY
HHLEDVLDDLVVSLCKFTTLLNSSLVDEPVTAFGDDIKARLATETVFSIANRYGDCIRTGWRNILDCILRLHKLG
LLPARVASDAADDSELPPDSVHGKPVPSSLSTSHIQTMGTPRRSSGLMGRFSQLLSLDTEEPRLQPTEQQLAAHQ
RTMQTIQKCRIDSIFIESKFLHADSLMQIARALIWAAGRPQKVSSSPDDEDTAVFCLELLIAITLNNRDRIGLLW
QGVYEHIANIVQSTVMPCALVEKAVFGLLRICQRLLPYKENLADELLRSLQLVLKLDARVADAYCENITQEVTRL
VKANAAHIKSQMGWRTITSLLSITARHPEASEIGFEALLFIMSEGAHLSPANYVLCIEASRQFAESRVGLTDRSV
RALDLMAESMNCLARWSHETGNAGPETDKISEGIREMWLRLVQALRKICLDQREEVRNHALASLQRCLVGDGVCL
LPSTWLQAFDLVIFTMLDDLLEIAQNHSQKDYRNMEGTLLHAMKLLSKVFLQQWQDLFGLSSFCKLWLGVLSRLE
KYMKVKVRGKKSDKLQELIPELLKNTLLMMKSKGILAKRSTIGGDSLWELTWLHVNNIAPSLQSEVFPGQEMEQL
HSGVQPEGNRSGLLAEPSAS*
**>Musac_LOC103988820**
MGHHRLQNSAIDRNQEESIDAVVDPGGHRNAAAAAVDLACVICSEVSAVLAVMRRNVRWGRRSASADEDDVIEHL
LIRSLKSLRRQVFSWGDARRPWSSIEPSAYLRPFLDVVCSDETSAPITGVALSSVYKILTLDVLDPGAGTGLDTA
MHLIVESVTSCRFEVTDPASEEAVLMKILQVLLAIMRSRASAVLSNQHVCTIVNTCFRVVHQAGTKGELLQRFAR
QTMHELIRCIFSHLPNVKDGGRPPSVKPELGAVDKDQAFGIKQVENGNGSVGPGMTSMADNNMPSPEAPIGAEIM
VEPYGIPCMLEIFHFLCSLLNVADQIGISPAPNQIALDEDVPLFALGMINSVIELGGASICRHPKLLSLIQDELF
RNLMQLGLSVSPLILSMVCSIVLNLYRHLRTELKLQLEAFFSCVILRVTQSRYGASYHQQEVAMEALVDFCRQTT
FMAEMYANFDCDISCSNVFEELANLLSKSAFPVNCPLSSTHVLALDGLIAVIQGMADRIENAPPPSEQTPQELEE
YASFWTMKCEDYSDPDQWVKFVHRMKNIKRKLMIVADHFNRDPKKGLEFLQGTHLLPENLDPNSVACFFRYTTGL
DKNLVGDFLGNHDEFCVQVLHEFARTFDFKDMNMDTALRLFLDTFRLPGESQKIQRVLEAFSERYFEQSPQILVD
KDAALILAYSLIMLNTDQHNVQVKKKMTEEDFIRNNRRINGGNDLPREFLSDLYHSICRNEIRTAPEQGFGFLEM
YPSRWVDLMLKSKTASPYTVCDSRPFLDREMFSVMSGPMIAAISVVFDYAEHEEIFLTCVDGFLAVAKISAHHHL
EEVLDDSVVSLCRFTTLLDSSFIEEPVTIFVDDTKARLALETVFSITNTYGDCIRTGWRNILDCILRLHKLGLLP
APVASDADDDSESSPDSIHGKPVPSPLSTSHVQIMGTRKSSGLIGRFSQLLYFDTEEPSLEPNEQQLAAHQRTLQ
IIQECHIDSIFTESKFLHADSLMQITRALIAAGHPQKITSSPDDEDMAVFCLELLIAITLNNRDRIGLLWQDVYE
HIANIVQSTVMPCALVEKAVFGLLRICQRLLPYKENLADELLRSLQLVLKLDARVADAYCENITQEITRLVKANA

THLKSQMGWRTIASLLSITARHPEASEVGFEALIFIMSEGAHISPANYVLLIEASRHFAESRVGQTDRSVRALDL
MAESINCLARWSRETRDAGEEAEKISEGIREMWLRLVQALRKISLDQREVVRNHALLSLQKCLVAAEGMCLLLST
WLQAFDLVIFTMLDDLLEIVQNHSPKDYRNMEGTFLHAVKLLSKVFLQLLQQLSGQSSFCKLWLGVIGRMEKYMK
IKVRGRKSEKLQELIPELLKNILFVMKSKGILAKGSIVGGDNLWELTWVHVNNIAPYLQSEVFPNQEMEQLHLGA
QTDNSSPRLLAEPSVG*
**>Musac_LOC103990130**
MTDMEKSSVVSRVPRRREIGISCMLNAEVGAVLAVMRRPHDPGVASPDEAANPHLLHSLKALRSLLFHPRQLGEW
RATDPSVYLSPFLDVVQSDDVPAAATGVALSTVLKVLKLDVFDEHTPGARDAIHSIVFAVTNCRLENTDSASEDA
VLMRVLQVLTAVMRTRASVLLTDHAVCTLVNTCFQVVQQSAHRGDLLQRSARHAMHELVQAIFVRLPDVRVPVER
PEAEDEVATTGSYGARCMVDIFHFLCSLLNVGEVIDYADSVGSISSDEDVQLFALVLINSTIELGGEAIGKHPKL
LRIIQDDLFHHLIHYATHTSPLVLSMICSTVLHLYNFLRRCLRLQLEALFTYVMLRIAAGSNGAQLQEVAVEGIT
SFCHQPNFVIETYVNYDCDPLRHNVLEEAARLLCKIAFPVGTPMSPLQIQAFGGIVTIITTIADNIEVDQAPDRE
AYNIDVSDYKPFWVEKCDNSDNSDTWVEFVRMRKLKKKKILIAANHYNRDDKKGLDFLKLSKLVPSPPEPKSLAY
FFRFTPGLDKNKIGDFLGDPDEFNIQVLNEFTGTFDFTGVILDIALRSYLETFRLPGESQKIQRILEAFSDRFYE
QQSSEIFASKDAVFILCYSLIMLNTDQHNPQVKKKMTEEEFIRNNRAINGGMDLPRDYLSELFHSISTNAITLFG
TAGTPTEMSSAHWDDLIKRSRTVEPFITCDYKHKLSREVFVAISGPSVATLAAIFEQTDDEDILHESLDGLISVA
RIARYGLEDILDELLSCFCKLTNLLNPYATAEETLLTFSNELKPRMATLALFTIANKFGESIRGAWKNMIDCLLK
LKRLKLLPQSVIEPDTALAADSKAQTHAKSESGVIFPASHRGAGSNRSVSGLIGRFSQFLSLDNSNDSMISFGSE
LENNLKIIQQCQIGDIFSKSSKLPDESLQNMGRALIFAAAGKGQKFSTPVEEEEIVGFCWDLLVTIASVNLHRFS
AFWPQFHESFAIVSQFPLFSPCPFAEKAIVGLFKIAVKLTASPSRLDKLPEELIFKSINLMWKLDKEILDTCCES
IAESTMQILNVHGKNVQTVFGWKTLLHLLTVAGRHPETFDQGVEALVKLMSEGTHITRYNYPYCIEAAFGFAALK
ISPLDKSSRILSLMADTVNLLIQWHKSGYSDPGSTNSSLSEDGQKAGGSAGNLGANLFMKLVEALRKTSLVRREE
IRNQAVAELRKCFAAAEELDFSPANCLACFNLVIFAMVDDLHEKMLEYSRRENSEREMRSMEGTLKVAMEMLVEV
YLEFLIPLSQSTGFRSFWLGMLRRMDTCMKADLGDHVGVLQELVPVLLKKMIVEMKEKEVLVQEDGNELWDITNI
QIQWIAPSVKDELFPEE*
**>Orysa_LOC9271296**
MGRPRLPGAAGIDPIAEEPPHSAAAAGDGGDAAGLACAISAEASAVLAVMRRSLRHPRAAADDAAADHPLVSSLK
ALRRLVFSPSAAAAPAGAVLRPFLDAVRSEDAGAAVTSASLAALHEVMALMGPSLTGAALREVVDAVASCRFEAG
AEAAAEEAVLMRMLQALLACLRAPAAPALGDQHVCTAVNTCFRVVHQAGAKGELLQRFSWHAMHELIRCVFARLP
QIGSGDGPDGSVKPEMGGMDKNHPFGIGQMENGNGSYASEAVTSDENSADGSGIVVEPYGIPCMVEIFHFLCSLL
NVVEQIGVDEDLPLFALKLINSAIELGGSSIRKHPKLLSLVQDELFRNLMQFGLSMSPLILSMVCSIVLNLYHHL
RTELKLQLEAFFSCIILRLAQPRFGATYHQQEVAMEALVDFCRQKNFMVEMYANLDCDITCRNIFEELANLLSKS
AFPINCPLSSMHILALEGLISVIQGMADRIGNATSRPELLPVELDEYTPFWTVKCENFSDPQHWVKFVRQRKYVK
RRLMIGADHFNRDPKKGLEFLQGTHLLPEKLDPQSVACFFRYTAGLDKNLVGDFLGNHDEFCVQVLHEFAQTFDF
QEMNLDTALRLFLETFRLPGESQKIQRVLEAFSDRYYEQSPQAFANKDTALVLAYSIIMLNTDQHNMQVKKKMTE
EDFIKNNRNINGGSDLPREMLSELYHSICRNEIKTTPEQGMGYFEMSPSRWIDLMRKSKSTSLYIVGDSQPFLDH
DMFAIMSGPTIAAIAVVFDHSEHEEVLLACVDGFLGVAKISAFHHLEDVLDDLVVSLCKFTTLLNTSLVEEPVTA
FGDDLKARLATETLFTIANRYGDYIRTGWRNVLDCILRLHKLGLLPARVASDAADDSEVSAETVQGKPTHSSIST
SHIPVMGTPRKSSGLMGRFSQLLSLDSEEPRSQPTEQQLAAHQRTLQTIQKCRIDSIFTESKFLQPDSLLQLARA
LIWAAGRPQKVASSPDDEDTAVFCLELLIAITLNNRDRIVLLWQGVYEHIANIVQSTVMPCALVEKAIFGLLRIC
QRLLPYKENLADELLRSLQLVLKLDARVADAYCENITQEVARLVKANAGHVKSQMGWRTVVLLLSITARHPDASE
VGFEAIMYIMSEGAHLSLSNYAFCIEASRQFAESRVGLIDRSIRALDLMADSANSLARWSQETKGTGEETDKVLE
AIREMWLKLLQALKKLSLDQREEVRNHALTSLQRCLTATEGVCLQSSTWSHAFDLVIFALLDDLLEISQNHSQKD
YRNMEGSLVLAMKLVAKVYLQLLPDLFGLSSFCKLWLGVLSRMEKYIKIKVRGKRSDKLQELIPELLKNILIAMK
NRGILAKRSTIGGDSLWELTWLHANNISTSLQSDVFPSQEYEQHSSAGSPRGPNGVESRD*
**>Orysa_LOC4329190**
MGGLRAASPSSGAGAGAGAGSDPTPRVAMACVLASEVATVLAIMRRNVRWAGVRYGGDDGADDEHLDHPLIAGLK
SLRRRAASWDTRQWRDVEPLLYLRPFLDVVRSDETGAPITGAALSSLHKILTLDLVGPDAPNVAEAMGAVVEAVT
GCRFEVTDPASEETVLARVLQVLLACVRGRAAPALANRHVCNIVSTCFRVVQQAGTKGELLQRVSRQTMQEVIRC
VFARLPDVDATVVADGQTACSKNQGLSDGEIGNGKSDFVCLNSSGDEVGGGFGVVQDQAMSELFGVPCMVEILQF
LCSLLNIAEDIEVNPRINPIDFDEDVPLFALGLISSAIELSASSINKHPELLAFVQDELFRNLMQFGLSMSPLIL
STVCSIVFTLFYHLRQELKLQLEAFFSCVIIRLGQSRYGASYQQQEVALEALVDFCRQKEFMAEMYANMDCDLQS
SNIFEDLANLLSKSAFPVKSPLSTLNVLALDGLVLVIQAIAERTDNAPQHHEQTVPEISEYFPFWQLKCENTNDP
DQWVRFVHQQKSIKRKLMVGVEHFNRDKKKGFEYLQGAHLLPERLDPRSVALFFRYTPGLDKNLLGDYLGNHDEF
SILVLHEFAKTFDFKEMNLDAALRLFLETFRLPGESQKIQRILEAFSERYYEQSPQMFVNRDAALVLSYSVIMLN
TDQHNIRVKKKMTEEDFIKNNRRINGGNDLPREFLSELYYSICRNEIRTIPEQGAGCSEMSFSRWVDLMWKSKRT
SAYIACDSFPFLDHDMFTIMAGPTVAAISVVFDNVEHEEFLTGCINGFLSVAKLAAFYHLDDVLNDLVVALCKFT
TLLNTSYINDPVTTFGEDTKARMATEAVFTIATTHGDHIRSGWRNIVDCILRLHKISLLPGCLTGDTADDQESSS
DMLPSKLASSRAAPQVVPISTPKKSYGLMGRFSQLLYLDAEESRFQPTEEQLAAQRNASETIKKCQIGTIFTESK
FLQADSLLNLARALTQAAGRPQRITSSLDDESTSVFCLELLITVTLNNRDRIVLLWQGVFEHITHIVQSTVMPCN
LVEKAVFGLLHICQRLLPYKENLVDDLLRSLQLILKLDARVADAYCENITQEVTRLVKGNATHIKSQMGWRTIIS
LLCITARHPDASDVGFEALVFIMSEGAHLSPANFVLSVEASRQFAESRLGSAERSIHALNLMAESVNCLTRWSRE
VKEAGGEADRILEGIAEMWLRLVQALRKVCTDQREEVRNHALLSLHRCLVVDGISVPSSAWLMSFDIIFQLLDEL

LEIAQNYSPKDFRNMEVSLLHAVKLLCKVFLQSLNDISSQSSFSKLWLEVLDMIEKLMKVKVRGRRTEKLQEVIP
ELLKNILLVLKANRVLSKTSTSEENSLWEATWLQVNKIAPSLQPEVFPDSEGDVATQSAKNKSDSPAQSEGVNV*
**>Orysa_LOC4334954**
MARTAAAAVSDDDDDGPPAYTVPRGPRRDPRLKDLGISCMLNTEVAALLAVIRRRPDPYSYLPPAVAAAEEATFA
GLIQSLKNLRGLLFQPRHGAWRCSDPAMYLKPFLDVVQSEESPPAATGVALSSVLKILRIDVFDECSPGARDAIQ
AVLTAVSSCRVERIADPGAEEAVLLRVLQVLAALLRARAAPLLSDHAVCNAVNTCFQVVQHAASGRGSELLQRTA
RHCMHEILQAVFARLPDIPAADAADADDLSVAGFGARCMADIFNFLCTLLLNAPDMVATGDGHGAFITEEDVQLF
SLVLINSAIELGGEAIGKHPKLLRLIQDDLFYHLIHYATESSPLVLSMICSTALNLYHFLRRFLKLQLEAFFMYV
LLRLCGGGGGAQLQEVAIEGLISFCRQPTFVIEMYVNYDCDPMLRNVYEEVGKLLCKAAFPASSPMTTVQLQAFE
GLVNMLTTIADNVEVDKAPDHAAYAVDISDYRLFWVERWDAAAAGGSGNNETWVDFVRKRKLRKKKVAIAANHYN
RDEKKGMEYLRLSQLVPTPPEPRSMAFFLRYSPGLDKNKIGELLGDPEEQSLRVLKEFTETFDFTGVILDTALRT
YLETFRLPGESQKIQRILEAFSERFYEQQTAEVFATKDAAFILCYSLIMLNTDLHNPQVKKKMSEDDFIRNNRAI
NAGKDLPREYLSELFHSISGNAITVFSQASAAAEMTPTRWADLVKRSRAIDPFTPCDFKHKLTREVFVTVSGPAV
ATLAAIFDYTDDEDILNQCVEGLISVARIARYGLEDVLDELLCCLCKFTTLLNPYATTEETLFTFSNELKPRMAT
LALFTITNRFGESVRGAWKNVVDCLLKLKRLKLLPPSLVDQEGGAGAAVSTERLGHRAKSESGVIFPSSHRGAGT
SRHVSGMIGRFSQFLSLDAGGESLLSVGSEFENNLKIIQQCRIGSIFTESGKLPDESVQNLGRALIFAAGGKGQK
FSTPVEEEETVGFCWDLIAVVSSANVHRFTTFWPQLHDCFAAVSQLPLFSPCPFAEKAIVALFRVAVRLLSGGGG
DRMAEELVFKSINLMWKLDKEILDTCCEGISECIVKLLMEYASNVQTPIGWKTVLHLLSVTGRHPETFDQSVAAM
IKLMNDGAQVVTRFNYAACIEAAFGFAALKISPLDISTRILQLMADSVNWLILWHKSGYSDPGNAWSSSSSSSSA
AAMMMMEDASRMGNLATSMFIKLAEALRKTSLVRREEIRNQAVAELGRAFALAAAGELELGPTGSLACFNLVIFA
MVDDLHEKTLEYSRREGAERETRSMEGTLAAATELLADVFVLFLGTLAQGAGFRTFWLGVLRRLDTCIKSDLAAG
GGAGVMQELVPRMLKRMIVEMKDKGVLVAMEGDELWEITHIQIQWIAPAVMEELFPD*
**>Sorbi_LOC8057184**
MGRPRMLGGGAGGIDPIAEEPHHHARSPADGGLGLGPDPAALACAISAEASAVLAVMRRGLRHPRADDAAAEHPL
VASLRALRRLAFSPSPALPAAALRPFLDAVRSEDAGAAVTSASLAALHEVMSLTGAALPGAALREVVDAVASCRF
EAGAEPAAEEAVLMRMLQALLACLRAPAAPALGDQHVCTAVNTCFRVVHQASAKSELLQRFSRHAMHELIRCVFA
RLPQIGSADGVDAAVKPEMGGMDVNHPFGIRQMENGNGSYVSETGASDENSADGSGLIVEPYGVPCMVEIFHFLC
SLLNVVEHIGLDEDLPLFALKLINSAIELGGSSIQKHPKLLSLVQDELFRNLMQFGLSMNPLILSIVCSIALNLY
HHLRTELKLQLEAFFSCIIIRLAQPRFGATYHQQEVAMEALVDFCRQKNFMVEMYANLDCDITCRNVFEELANLL
SKSAFPINCPLSSMHILALEGLIAVIQGMADRIGNATSRPELLPVELDEYTPFWTVKCENFSDPRHWVKFVRQRK
YVKRRLMIGADHFNRDPKKGLEFLQGTHLLPEKLDPQSVACFFRYTAGLDKNLVGDFLGNHDEFCVQVLHEFAQT
FDFHEMNLDTALRLFLETFRLPGESQKIQRVLEAFSDRYYEQSPQSFANKDTALLLSYSIIMLNTDQHNMQVKKK
MTEEDFIKNNRNINGGSDLPREMLSELYHSICRNEIKTTPEQGLGYFEMSPSRWIDLMRKSKSTSPYIVGDSQPF
LDHDMFAVMSGPTIAAIAVVFDHSEHEEVLLTCVDGFLGVAKISAFHHLEDVLDDLVVSLCKFTTLLNTSLVEEP
VTAFGDDLKARLATETLFTIANRYGDYIRTGWRNVLDCILRLHKLGLLPARVASDAADDSELSAEAVQGKAAPSA
VPPSHIPVMGTPRKSSGLMGRFSQLLSLDSEEPRSQPTEQQLAAHQRTLQTIQKCRIDSIFTESKFLQPDSLLQL
ARALIWAAGRPQKVASSPDDEDTAVFCLELLIAITLNNRDRIVLLWQGVYEHIANIVQSTVMPCALVEKAIFGLL
RICKSLLPYKENLADELLRSLQLVLKLDARVADAYCENITQEVARLVKANAAHIKSQMGWRTVILLLSITARHPD
ASEVGFEAIVFIMTEGAHLSLANYGFCIEASRQFAESRVGLADRSVRALDLMSDSVRSLAMWSQEIKATCEEGEK
GLEAIREMWLKLLQALKKLSLDQREEVRNHALASLQRCLTATEEICLQSATWSHAFDLVIFSLLDDLLEISQNHS
QKDYRNMEGSLVLAMKLVAKVYLQLLPDLFGLSSFCKLWLGVLSRMEKYIKIKVRGKRSDKLQEVIPDLLKNILL
VMKNKGILAKRSTIGGDSLWELTWLHANNISTSLLPDVFPSQEYEQQSSAGSPRGPSSVEA*
**>Sorbi_LOC8055663**
MGGLRAASPSGAGSGAVAGAGSDHGAPRLAMACVLASEAATVLAVMRRNVRWAGVRYGGGDGGAGDDEHLDHPLV
AGLKALRRRAAAWGHGRWAGVEPLLYLRPFLDVVRSDETGAPITGAALSSLHKILTLDLVGPGAPAAAEAMAAVV
DAVTACRFEVTDPASEEAVLARVLQVLLACVRSRAAPALANRHVCTIVNTCFRVVQQAGTKGELLQRVSRQTMQE
VIRTVFARLPDIHVTVLSDEQVTGCKNQNLGAEETENGKSDYVCLNSSGDEVGDGSGVVPDKDLMEPFGVPCMVE
ILQFLCSLLNIAEDIKVNPRMSPIDFDEDVPLFALGLINSAIELSASSIHRHQKLLSFVQDELFRNLMHFGLSMS
PLILSTVCSIVFTLFYHLRHELKLQIEAFFSCVILRLAQSRYGASYQQQEVALEALIDFCRQKEFMAEMYANMDC
DLQCSNIFEELANLLSKSAFPVNSPLSALNVLALDGLVAVIQAMAQRTDNAPQHHDQTVPEISEYFPFWQLKCES
SNDPDQWVKFVHQQKSIKRKLMVGVEHFNRDKKKGFEYLQGVHLLPEKLDPHNVALFFRYTPGLDKNLLGDYLGN
HDEFSIRVLHEFARTFDFKDMNLDAALRLFLETFRLPGESQKIQRILEAFSERYYEQSPQMFVNRDAALVLSYSV
IMLNTDQHNVRVKKKMTEEDFIRNNRRINGGNDLPREFLSELYYSICRNEIRTIPEQGAGCSEMSFSRWVDLMWR
SKRTSMYIACDSYPFLDHDMFSVMAGPTVAAVSVVFDNVEHEEVLTGCIDGFLSVAKLAAFYHLDDVLNDLVVAL
SKFTTLLNTSYIDDPVTAFGEDTKARMATEAVFTIATAYGDHIRSGWRNIIDCILRLHKIGLLPGCLTGDTTDDQ
ESSSDSLPGKLASSAPQVLPISTPRKTYGLMGRFSQLLYLDADEPRSQPTEEQLAAQRNASETVKKCQIGTIFTE
SKFLQADSLSNLARALIQAAGRPQKITSSLDDEGISVFCLELLITVTLNNRDRIVLLWQDVFEHITHIVQSTVMP
CNLVEKAVFGLLHICQRLLPYKANLVDDLLRSLQLILKLDARVADAYCENITLEVTRLVKANATHIKSQMGWRTI
ISLLCITARHPDASDAGFEALVFIMSEGAHLSPANFILSVEASRQFAESRLGSAERSIHALNLMADSVNSLIRWS
REVREAGGEADRILEGIAEMWLRLVQALRKVCMDQREEVRNHALLSLHRCLVVDGISVKSSTWLMAFDIIFQLLD
ELLEIAQSYSPKDFRNMEVSLLHAVKLLCKVFLQSLKDLSGQGSFDKLWLEVLDMIEKFMKVKVRGRRTEKLQEA
IPELVKNILMVMKASGILSKTGASENSIWEATWLQVNKIAPSLQSEIFPDNEDGSTTQVEENKLDTQAQSDKNND
Q*

**>Sorbi_LOC8063286**
MARTAGDDDDDGPPPPYAPQRGPRRDPRLKDLGISCMLNTEVGALLAVIRRRPDPYSYLPPAVAAAEEATFAALI
SSLKALRGLLFQPRHGAWRCSDPSTYLTPFLDVVQSEEAPPAATGVALSSVLKILRIDVFDECSPGARDAVHAIL
TALTNCRIERISDAGAEEAVLLRVLQVLGALLRTRAAPLLSDSSVCTAVNTCFQIVQHAASSRGSELLQRTARHC
MHEILQSVFARLPDIRDDADGDDLSLGSGAGFGARCMVDVFNFLCSLLVNASDMVITPDGQGAFTSEEDVMLFSL
VLVNSAVELGGEAIGKHPKLLRLIQDDLFFHLIHYATEYSPLVLSMICSTALNLYHFLRRFLKLQLEAFFVFVLL
RVCGGGNGPQLQEVAVEGLISFCRQPTFVIEMYVNYDCDPLLHNVFEEVGKLLCKAAFPVSGPMSTIQLQAFEGL
VNMVTTIADNVEIDKTPDHDAYAVDVSEFRLFWTERWDSAAAAAAAAAGQRETWVDFVRKRKLRKKKVAVAANHY
NRDQKKGVEFLKLCHLVPTPPDPRSMAYFLRYSPGLDKIKIGEFLGDPDEFNLKVLKEFTETFDFTGAILDTALR
TYLETFRLPGESQKIQRVLEAFSERFFEQQTTGVFATKDAAFILCYSLIMLNTDLHNPQVKKKMSEEDFIRNNRA
INDKKDLPREYLSELFHSISTNAITVFSASAAAVEMTRSRWADLVKRSRALEPFTPCDFKHKLSREVFIAVSGPT
VATLAAIFDGADDEETLNQCVEGLVSVARIARYGLEDVLDELLCCLCKFTTLLNPYATAEETLFTFSNELKPRMA
TLALFTIANRFGESVRGAWKNVVDCLLKLKRLKLLPPSLIDTDGSGGARGGAEQRPGHRHRSSASEAGVIFPATH
RGAGTSRHVSGMIGRFSQFLSLDGGESLLSVGSEFENNLKIIKQCQAGSIFTESAKLPDEALQNLGRALIFAAGG
KGQKFSTPIEEEDTVGFCWDLLALLATANLQRFATFWPPLLECFSAVSQLPLFSPSPFAERAIVALFRVAVRLLS
TPPSQRPSDSRVPEELVFKSINMMWKLDKEILDTCCEGISENIVKLLTEHADGVQTPLGWKTLLHLLTVTGRHPE
TFDQSVAAMIKLMTDGSANITRFNYAAVIEAAFGFAALKISPLDISTKILELMAESVNWLIQWHKSGYSDPGNSG
GFSGSSSSSSSVDEASRMGNLAANMFIKLAEALRKTSLVRREEIRTQAVFELGRAFNLAATGDLDFGPAGCLACF
NLVIFAMVDDLHEKTLEYSRREGAERETRSMEGTLAAAAELLADVFVLLLPTVAQAPGFRTFWLGVLRRMDTCMK
CDLAAGGGDGLMQELVPRMLKRMIMEMKAKEVLVPREGDELWEITHIQIQWIAPAAMDELFP*
**>Bradi_LOC100840992**
MGRPRLLNAGGIDPIAEEPHPPRADADPGGLACAISAEASAVLAVMRRSLRHPRAAADDAAADHPLVSSLKALRR
LVFSPAAAASPSLPAASLRPFLDAVRSEDAGAAVTSASLAALHEVMALTGPSLPGSALREVVDAVASCRFEAGAE
AAAEEAVLMRMLQALLACLRAPAAPALGDQHVCTAVNTCFRVVHQAAAKGELLQRFSRYAMHELVRYIFARLPQI
GSDDGADGTAIPEMGGMDKNHPFGIREMENGNGNYAPEAGTSDENSVDGSGLIVEPYGIPCMVEIFHFLCSLLNV
VEQIGFDEDLPLFALKLINSAIELGGSAIGKHPKLLSLVQDELFRNLMQFGLSISPLILSMVCSIVLNLYHHLRT
ELKMQLEAFFCCIILRLAQPRFGATYHQQEVAMEALVDFCRQKNFMVEMYANLDCDITCRNVFEELANLLSKSAF
PINCPLSSMHILALEGLIAVIQGMADRIGNATSRPELRPVELDEYAPFWTVKCENFLDPQHWVRFVRQRKYVKRR
LMIGADHFNRDPKKGLEFLQGNHLLPEKLDPQSVACFFRYTAGLDKNLVGDFLGNHDEFCVQVLHEFAQTFDFQE
MNLDTALRLFLETFRLPGESQKIQRVLEAFSDRYYEQAPQAFANKDTALLLSYSIIMLNTDQHNMQVKKKMTEED
FIKNNRNINGGSDLPREMLSELYHAICRNEIKTTPEQGMGYLEMSPSRWIDLMRKSKSTSPYIVGDSQPFLDHDM
FAIMSGPTIAAIAVVFDHSEHEEVLLTCVDGFLGIAKISAFHHLEDVLDDLVVSLCKFTTLLNTSLVEEPVTAFG
DDLKARLATETLFTIANRYGDYIRTGWRNVLDCILRLHKLGLLPARVASDAADDSEVYTETVQGKPAPSSISTSH
IPVMGTPRKSSGLMGRFSQLLSLDSEEPRSQPTEQQLAAHQRTLQTIQKCRIDSIFTESKFLQPDSLLQLARALI
WAAGRPQKVASSPDDEDTAVFCLELLIAITLNNRDRIVLLWQGVYEHIANIVQSTVMPCALVEKAIFGLLRICQR
LLPYKENLADELLRSLQLVLKLDARVADAYCENITQEVARLVKANAGHIKSQMGWRTVVLLLSITARHPDASGVG
FEAIMFIMSEGHLSKSNYAICIEASRQFAESRVGLTDRSIRALDLMADSAINLARWSQDTKGSGEEADKGSEAIR
EMWLKLLQALKKLSLDQREEVRNHALISLQRCLTATEGICLQSTTWSHAFDLVIFALLDDLLEIGQNHSQKDYRN
MEGSLVLGMKLVVKVYLQLLPDLFGLSSFCKLWLGVLSRMEKYIKIKVRGKRSDKLQELIPDLLRSILAAMKSQG
ILAKRSTIGGDSLWELTWLHVNNISTGLQSEVFPSQEYEQPNNVGSPRGLNDAETKN*
**>Bradi_LOC100836075**
MGGLRAASPTRPSPPPPRVAMACVVASEIATVLAiMRRNVRWAGVRYGGNDEEHVDHPLIAGLKSLRRAAASWGN
GWPVGGVDPLLYLRPFLDVVRSDETGAPITGAALSSLHKILSLDLVPPGAPNVAEAMGAVVDAVTGCRFEVTDPA
SEEAVLARVLQVLLACVRGRAATALDNRHVCAIVSTCFRVVQQAGTKGELLQRVSRQTMQEVIRCVFSRLPDIDA
ITIVNEQIAGSKNQALGAGEMGNGKSDYVCLNSSGDEVGGGLSVVQDKAMMEPFGVLCMVEILQFLCSLLNIAED
MDVSQRMNPIDFDEDVPLFALGLINSAIELSASSIHRHPKLMSFVQDELFRNLMQFGLSMSPLILSTVCSTVFTL
FYHLRQELKLQVEAFFSCVILRLAQGRYGASYQQQEVALEALVDFCRQKEFMAEMYANMDCDLQCSNVFEDLANL
LSKSAFPVNSPLSALNVLALDGLVAVIQAIAERTDNAHQHHDQAVPEISEYFPFWQLKCESSDDPDQWVRFVHQQ
KGIKRKLMVGVEHFNRDKKKGFEYLQGVHLLPEKLDPCSVALFFRYTPGLDKNLLGDYLGNHDEFSILVLHEFAK
TFDFKEMNLDAALRLFLETFRLPGESQKIQRILEAFSERYYEQSPHMFVNRDAALVLSYSVIMLNTDQHNVRVKK
KMTEEDFIRNNRRINGGNDLPREFLAELYYSICRNEIKTIPEQGAGCSEMSYSRWVDLMWKSKRTSTYIACDSYP
FLDNDIFPIMAGPSVAAISVVFDNVEHEEVLTGCIDGFLSVAKLAAFYHLDDVLNDLVVALCKFTTLLNTSNSDD
PVIAFGEDTKARMATEAVFTIATTYGDHILSGWRNVVDCILRLHKIGLLPGRLTGDTGDDQESSSDSLPSKLAVV
PQLVRINTTKKTYGLMGRFSQLLYLDTDVPGSQPTEEQLAAQRNASETVKKCQIGTIFTESKFLQADSLSNLARA
LVQAAGRPQRITSSLDDEGTSVFCLELLITVTLNNRDRIVLLWQGVYEHITHIVQSTVMPCNLVEKAVFGLLHIC
QRLLPYKENLVDDLLRSLQLILKLDARVADAYCENITLEVTRLVKANATHIKSQMGWRTIISLLCITARHPDASD
AGFEALVFIMSEGAHLSPANFILSVEASRQFAESRLGSAERSIHALNLMSDSVNCLTRWSREVKEAGGEADRILE
GIAEMWLRLVQALRKVCTDQREEVRNHALLSLHRCLVVDGISVSSSAWLMSFDIVFQLLDELLEISQSYSPKDFR
NMEVSLLHAVKLLCKVFLQSLKDLSAQSSFSKLWLEVLDMIEKYAKVRLRGRRTEKLHEAIPELLKNILLVMKAS
GVLSNTSASGENTLWEATWLQVNKIAPSLQSEVFPDNEGEAAAKGEQSMSDSPAQEGQIAVQ*
**>Bradi_LOC100825923**
MARTAASDDDDDGPPTYTMARGNRRDPRLKDLGISCMLNTEVAALLAVIRRRPDPYSYLPPAVAAAEEAVFAGLI
QSLKNLRALLFQPRHGAWRCSDPSMYLTPFLDVVQSEEVPPAATGVALSSVLKILRIDVFDECSPGARDAIQAIL

TAVTNCRIERIADSGAEEAVLLRVLQVLAALLRARAAPLLADSAVCTAVNTCFQVVQHAASSRGSELLQRTARHC
MHEILQAVFSRLPDIRDDAADDDAAVTSGAGFGSRCMVDVFNFLCSLLLNATDMVMTPEGHGAFTSEEDVQLFAL
VLLNSAVELGGEAIGKHPKLLRLIQDDLFYHLIHFATECSPLVLSMICSTVLNLYNFLRRFLKLQLEAFFMFVIL
RVGSGASGLQLQEVAIEGLISFCRQPTFVIEMYVNYDCDPLLRNVYEEVGKLLCKAAYPLSNPMTTVQLQAFEGL
VNMITTIADNVEVEKAPDQAAYNVEISEYRLFWLERWETGEDHGHETWVDFVRKRKLKKKKVAIAANHYNRDEKK
GVEFLKLCYLVPTPPEPKSMAYFLRYSPGLDKVKIGEYLGDPDEFNLQVLKEFTETFDFTGSILDTALRTYLETF
RLPGESQKIQRVLEHFSERFFDQQTAGVFATKDAAFILCYSVIMLNTDLHNPQVKKKMSEDDFIRNNRAINSGKD
LPREYLSELFHSIASNAITMFSQSVTSIEMTTSRWGDLVNRSRSIEPFTPCDFKHKLSREVFIAVSGPAVSTLAA
IFDYTDDEETLNQCVEGLISVARIARYGLDDVLDELLCCLCKFTTLLNPYSTTEETLFTFSNELKPRMSTLALFT
IANRFGESVRGAWKNIVDCLLKLKRLKLLPQSVIEADGSVSSNSDRLSHRPKSELGVIFPSSHRGAGTSRHVSGM
IGRFSQFLSLDNTTESLLSVGSEFENNLKIIQQCRIGSIFTDSGKLPDESLQNLGRALIFAAGGKGQKFSTPIEE
EETVGFCWDLLLVSSANLHRFSSLWPHMHDCFMAVSQLPLFSPCPFAEKAIVALFKIAVKLLPGQPNPDRVAEEL
VCKSINLMWKLDKEILDTCCEGISECIVKLIMDHAGSVQTPIGWKTLLHLLSVTGRHPETFDQSVAALIKLMSDG
AHINRFNYAACIEAAFGFAALKISPLEISTKILELMADSVKWLIQWNKSGYSDPGSTNSSNSSSWAEDASRMGNL
ATSMFIKLAEALRKTSLVRREEIRNQAVADLSRGFAIAAAGDLDLGPAGCLACFNLVIFAMVDDLHEKTLEYSRR
EGAERETRSMEGTLAAATQLLADVFVLFLGTLAQGPGFRTFWLGVLRRMDTCIKSDLAAGGGLGVMQELVPRMLK
RMIVEMKDKEVLVQRDGDELWEITHIQIQWIAPAVKEELFPE*
**>Phala_03g27650**
MGRPRLLGSGGIDPIAEEPPSDGSPSASADLAGLACAISAEASAVLAVMRRSLRYPRSAAAAAADDAAADHPLVS
SLKALRRLVFSPASASAAAVASLPAATLRPFLDAIRSDEAGAAVTSASLAALHEVMALTGAALPGAALREVVDAV
TSCRFEAGAEVAAEEAVLMRILQVLLACLRAPAAAALGDQHVCTAVNTCFRVVHQAGAKGELLQRFSRHAMHELI
RCVFARLPQISGGDGGDRAVKPEMGGTDKNQPFGIRQMENGNGSYTSETGTSDETSVDGSGLVVEPYGVPCMVEI
FHFLCSLLNVVEQIGINPGINAIAFDEDVPLFALRLINSAIELGGSSIRKHPKLLSLVQDELFRNLMQFGLSMSP
LILSMVCSIVLNLYHHLRTELKLQLEAFFSCIIIRLAQPRFGATYHQQEVAMEALVDFCWQKNFMVEMYANLDCD
ITCRNVFEELANLLSKSAFPINCPLSSMHILALDGLIAVIQGMADRIGSETLRPELVPVELDEYTPFWTVKCDNF
SDPQHWVKFVRQRKYVKRRLMIGADHFNRDPKKGLEFLQGTHLLPEKLDPQSVACFFRYTAGLDKNLVGDFLGNH
DEFCVQVLHEFAQTFDFQEMNLDTGLRLFLETFRLPGESQKIQRVLEAFSDRYYEQSPQAFANKDTALLLAYSII
MLNTDQHNMQVKKKMTEEDFIKNNRNINGGSDLPREMLSELYHSICRNEIKTTPEQGMGYFEMSPSRWIDLMRKS
KSTSPYIVGDSQPFLDHDMFAIMSGPTIAAIAVVFDHSEHEEVLLTCVDGFLGVAKISAFHHLEDVLDDLVVSLC
KFTTLLNTSLVEEPVTAFGDDLKARLATETLFTIANRYGHYIRTGWRNVLDCILRLHKLGLLPARVASDAADDSE
LSAETVQGKVTPSSISTSHVPVMGTPRRSSGLMGRFSQLLSLDSEEPRSQPTEQQLAAHQRTLQTIQKCRIDSIF
TESKFLQPDSLLQLARALIWAAGRPQKVASSPDDEDTAVFCLELLIAITLNNRDRIVLLWQGVYEHIANIVQSTV
MPCALVEKAIFGLLRICQRLLPYKENLADELLRSLQLVLKLDARVADAYCENITQEVARLVKANATHIKSPMGWR
TIMLLLSITARHPDASEVGFEAVVFIMSEGAHLSTANYALCIEASRQFAESRVGLIDRSIRALDLMVDSVTCLAQ
WSEETKGTDEAERGLEAIREMWLKLLQALKKLSLDQREEVRNHALTSLQRCLTAAERICLEPATWSHAFDHVIFA
LLDDLLEIGQSHSQKDYRNMEGSLVLAMKLIAKVYLQLLPQLFGLSSFCKLWLGVLSRMEKYIKIKVRGKRSDKL
QELIPELLKSILIVMKNRGDLAKRSTIGGDSLWELTWLHVNNISASLQSEVFPSQEFDQQKDSGCSSGPNGIEAG
ATG*
**>Phala_09g17940**
MGALRAGSPSAPDAGAGVALDPPRRLAMTCVVASEVATVLAVMRRNVRWAGVRYGGDDGGDDEHVDHPLIAGLKS
LRRRAATWDPRRWCAVDPLLYLRPFLDVVRSDETGAPITGAALSSLHKILTLDLVGPDAPNVAEAMDAVVEAVTG
CRFEVTDPASEEAVLARVLQVLLACARGRAAPALANRHVCAIVSTCFRVVQQAGTKGELLQRVSRQTMQEVVRSV
FARLPDIDTTAVADDQIAGSNNRDLGVGEMGNGKSDFVCLNSSGDEVGDGLEVVQDKAMMEPFGVPCMVEILQFL
CSLLNIAEDIEVNPRMNPIDFDEDVPLFALGLINSAIELSASSTHRHPKLLAFVQDELFRNLMQFGLSMSPLILS
MVCSIVFTLFYHLRQELKLQLEAFLSCVILRLAQSRYGASYQQQEVALEALVDFCRQKEFMAEMYANMDCDIQCT
NVFEDLANLLSKSAFPVNSPLSALNVLALDGLVAVVQAIAERTDNAPQHHEQTMPEISEYSPFWQLKCNNSSDPD
QWVRFVRRQKCIKTKLMVGVEHFNRDQKKGFEYLQSVHLLPEKLDPHSVALFFRYTSGLDKNLIGDYLGNHDEFS
IQALHEFAKTFDFKEMNLDTALRLFLETFRLPGESQKIQRILEAFSEWYYEQSPHIFVNRDAALVLSYSVILLNT
DQHNVRVKKKMTEEDFIRNNRRINGGNDLPREFLSELYYSICRNEIRTIPEQGAGCPEMSFSRWVDLMWKSRRTS
PYIVCDPYPFIDHDMFPIMAGPTVAAISVVFDNVEHEDVLAGCINGFLSVAKLAAFYHLDDVLNDLVVSLCKFTN
LLNTSYIDDPVTAFGEDTKARMATEAVFTIATTYGDHIRSGWRNIIDCILRLNKIGLLPGCLTGDTADNHESSSD
AMLSRLASSSASAPQAAQINTPKRTYGLMGRFSQFLYLDAEEPRSQPTEEQLAAQRNASQTVKNCQVGTIFTESK
FLQADSLSNLARALIQAAGRPQKITSSLDDEGTAVFCLELLITVTLNNRDRIVLLWQGVYEHISHIVQSTVMPCN
LVEKAVFGLLHICQRLLPYKENLVDDLLRSLQLILKLDARVADAYCENITQEVTRLVKANATHIKSQMGWRTIIS
LLSITARHPDASDSGFEALVFIMSEGAHLSPANFIISVEASRQFAESRLGPAERSVHALDLMADSVNCLTRWSHE
VKEAGGEADRILEGITEMWLRLVQALRKVCTDQREEVRNHALLSLHRCLVVDGISVSPSAWLMSFDIVFQLLDEL
LEIARNYSPKDFRNMELSLLHTVKLLCKVFLQSLKSLSAQGSFSKLWFEVLDLIEKFMKVKVRGRRTEKLQEAIP
ELLKNILMVMKANGILSKESTSGENSLWEATWLQVNKITPSLQPEVFPDNEGDYGTQDQQNKSDGPAQPEGKSIE
Q*
**>Phala_01g16080**
MARTHSAISDDDDDSPPAHTVHRGPRRDPRLKELGISCMLNTEVAALLAVIRRRPDPYPFLPPAAAAAEEATFTH
LIQSLKNLRAHLFQPRHGAWRCTDPSVYLSPFLDIVQSEEVLPAATGVALSSVLKILRHDVFDECSPGARDATHA
ILAAVTNCRLDRIVDAGAEEAVLLRVLQVLAALLRARAAPLLSDQAVCTAVNTCFQVVQHAGGSRGSELLQRTAR

HCMQEILQAVFARLPDIRAVPETSGAHGSDTEDNMSLTAGAGFGARCMIDIFSFLCSLLLNATDMSMSPEGYGAF
SSEEDVELFALVLINSAIELGGEAIGKHPKLLHLIQDDLFFHLIHYATESSPLVLSMICSTALNLYHFLRRFLKL
QLEAFFMYVLLRVGTGGSGLQLQEVSIEGLISFCQQPTFVIEIYVNYDCDPLSHNVFEEIGRLLCKAVFPTAGPL
STVQVQAFEGLVNLITTVADNIEVEKTPEREAYSIDVLEYRPFWVERWDAADDHNPETWVDFVRKRKLKKKKVAM
AANHYNRDQKKGIEFLKLCHLVPTPPEPRSMGYFFRYTPGLDKNRIGDFLGDPDEFNLRALKEFTETFDFTGDIL
DTALRTYLETFRLPGESQKIQRILEAFSERFFDQQTSDVFATKDAAFILCYSLIMLNTDLHNPQVKKKMTEEDFI
RNNRAINAGKDLPREYLSELFHSISTNAITVFSQTVATVDMTSSRWADLVRRACAVEPYTPCDFKHKLSREVFIA
ISGPTVATLAAIFECTDDEETLNQCIEGLVSVARIARYGLEDVLDELLSCLCKFTTLLNPYATTEETLFTFSNEL
KPRMATLALFTIANRFGESVRGAWKNIVDCLLKLKRLKLLPQSAIEPDDGSSSNSDRSGLRSKSESGVIFPLSHR
GAGTSRHVSGMIGRFSQFLSLEGGGESLLSVGSEFENNLKIIQQCQIGSIFIESSKLPEESLQNLGRALIFAAGG
KGQKFSTPVEEEETVGFGWDLILLVTSANLERFTMFWPQWHECFNAVSQFPLFSPCPFAEKAVVTLFKIAVRMLS
APACLDRLPEELVFKSINLMWKLDKEILDTCCEGISESIVKLLTEHAGSVQTPLGWKTLLHLLSLTGRHPETFDQ
SVAALIKLMADGTHVTLFNYASCIEAAFSFAALKISPLEISTKILELMADSVNWLAQWHRSGYSDPGNSFASNSS
SSSSIEDASRMGNLVTNMFIKLAEALRKTSLVRREEIRNQAVAELRRSFCLAAAGELEFGPAGCLACFNLVIFAM
VDDLHEKTLEYSRREGAERETRSMEGTLAAATELLVDVFVRFLGTLALWPGFRTFWLGVLRRMDTCMKSGLAAGG
GAGVMQELVPQMLKRMIVEMKEKEVLVQREGDELWEITNIQIQWIAPAVKEELFPDKLGTSVGEF*

**>Joasc_18G020200**

MGRPRLLNSGGIDTIAEESPASDASPAAAADDAGLGCAISAEASSVLAVMRRSLRYSRSGGGGGDDGAAEHPLVS
SLKSLRRLVFSPHPQPGAAVLRPFLDAVRSDEAGAAVTGAALAALHEVLSLTGAALPGAALREVVDAVTSCRFEA
AADPAAEEAVLMRILQVLLACLRAPAASLGDQHVCTAVNTCFRVVHQAGAKGELLQRFSRHAMHELVRCIFARL
PQIGAAGDNDGSVKPEMGGTDKNQPFGIKQMENGNGSYAPETGTVDDNSANGGGLVVEPYGVPCMVEIFHFLCSL
LNFVEQIGINPGTNAIAFDEDVPLFALRLINSAIELGGSSIRKHPKLLSLVQDELFRNLMQFGLSMSPLILSMVC
SIALNLYHHLRTELKLQLEAFFACVILRLAQPRFGATYHQQEVAMEALVDFCRQKNFMVEMYANLDCDITCRNVF
EELSNLLSKSAFPVNCPLSSMHILALDGLIAVIQGMADRIGNATSRPELMPVELEEYAPFWTVKSENFSDPQHWV
KFVRQRKYVKRKLMIGADHFNRDPKKGLEFLQGTHLLPEKLDPQSVACFFRYTAGLDKNLVGDFLGNHDEFCVQV
LHEFAQTFDFQEMNLDTALRLFLETFRLPGESQKIQRVLEAFSDRYYEQAPQAFANKDTALLLAYSIIMLNTDQH
NMQVKKKMTEEDFIKNNRNINGGGNLPREMLSELYHSICRNEIRTTPEQGMGYFEMSPSRWIDLMRKSKTTPPYI
VGDSQPFLDHDMFAIMSGPTIAAIAVVFDHSEHEEVLLTCVDGFLAVAKISAFHHLEDVLDDLVVSLCKFTTLLN
TSLVEEPVTAFGDDLKARLATETLFNIANRYGDYIRTGWRNVLDCILRLHKLGLLPARVASDAADDSELSTETVQ
GKPVPSSISTSHIPVMGTPRRSSGLMGRFSQLLSLDSEEPRSQPTEQQLAAHQRTLQTIQKCRIDSIFTESKFLQ
PDSLLQLARALIWAAGRPQKVTSSPDDEDTAVFCLELLIAITLNNRDRIVLLWQGVYEHIANIVQSTVMPCALVE
KAIFGLLRICQRLLPYKENLADELLRSLQLVLKLDARVADAYCENITQEVTRLVKANATHIKSQMGWRTIMLLLS
ITARHPEASEVGFEAVIFIMSEGAHLLTANYALCIEASRQFAESRVGLIDRSIRALDLMADSVKCLARWSQETKG
TDEEAEKMLEGIREMWLKLLQALKKLCLDQREEVRNHALTSLQRCLAGGTEGICLQSATWLQAFDHIIFAMLDDL
LEIGQNHSQKDYRNMEGSLVLAMKLLAKVYLQLLPELFGLSSFCKLWLGVLSRMEKYIKIKVRGKRSEKLQELIP
ELLKNILIVMKNRGILAKRSTIGGDSLWELTWLHVNNISTSLQSEVFPGQESEQQKDSGSPRGGLNGVEAGGPD*

**>Joasc_04G090200**

MGCLKAAVAAEESPPRVAMACVVAAEVATVLAVMRRNVRWGGGGGAAAASAGVRYGGDEGDDEHLDHPLIAGLKT
LRRRAASWDARWWRAVDPLLYLRPFLDVVRSDETGAPITGAALSSLHKILTLDLIGLDTVNVAAAMEVVVEAVTG
CRFEVTDPASEEAVLVRILQVLLACVRGRGAPALTNRHVCAVVSTCFRVVQQAGTKGELLQRVSRQTMQEVVRCV
FARLPEMDGSSSSAATVADGQIGGSRSQDFGVKELENGRSDYVCLNSSGNEVGDGLEVMQDKFMMEPFGIACMVE
ILQFLCSLLNIAEDIEVNPRMNPIDFDEDVPLFALGLINSAIELSASSIHRHPKLLAFVQDELFRNLMQFGLSMS
PLILSMVCSIVLNLFYHLRQELKLQLEAFFSCVILRLAQSRYGASYQQQEVALEALVDFCRQKEFMAEMYANMDC
DIQCTNVFEDLANLLSKSAFPVNSPLSSLHVLALDGLVAVVQAIAERIDNAPQHHEQTVAEISEYSPFWQLKCEN
RNDPDQWVRFVRRQKCIKRKLMVGVEHFNRDQKKGFEYLQGVHLLPEKLDPQSVALFFRYTSGLDKNLIGDFLGN
HDEFSIQVLHEFAKSFDFKEMNLDTALRLFLETFRLPGESQKIQRILEAFSERYYEQSPQIFVNKDAALVLSYSI
ILLNTDQHNVRVKNKMTEEDFVRNNRRINGGNDLPREYLSELYYSICRNEIRTIPEQGAGCPEMSFSHWVDLIWK
SKRSSPYIVCDSYPFLDHDMFAIMAGPTVAAISVVFDHVEHEEVLTGCVDGFLAVAKLAASYHFDDVLNDLVVAL
CKFATLLNSSYIDDPVTAFGEDTRARMATETVFTIATTYGDHIRSGWRNIVDCILRLHKIGLLPGRLTSDAADDP
ESSSDAMPSKPASLSVYPPQIAPISTPKRAYGLMGRFSQLLYLDAEEPRSQPTEEQLAAQRNASQTVKKCQIGTI
FTESKFLQADSLSDLARALIQAAGRPQKITSSLDDEGMAVFCLELLIAVTLNNRDRIVLLWQGVYEHIAHIVQST
VMPCNLVEKAVFGLLHICQRLLPYKENLADDLLRSLQLILKLDARVADAYCENITQEVTRLVKANATHIKSQMGW
RTITSLLSITARHPDASDAGFEALVFIMSEGAHLSPANFVLSVEASRQFAESRLGSAERSVHALDLMAESVNCLA
RWTREVKEVGGEEERILEGITEMWLRLVQALRKVCTDQREEVRNHALLSLHRCLVVEGMYISASAWLQSFDIIFQ
LLDELLEIAQSYSPKDYRNMEVSLLHAVKLLCKVFLQSLKDLSALSSFSKLWLEVLSQIEKFMRAKMRGRRTEKL
QEAIPELLKNILLVMKASGILSKRRSTSGENSLWEPTWLLVNKIAPSLQPEVFPDNEGDHGTQDEQKESDSPAES
QGQSIEQ*

**>Joasc_09G111700**

MARINSAVSDDDDDGGATAQPVHRLARRDPRLKELGVSCMLNTEVAALLAVIRRRQDAPPFLAAAAAAAEETAFP
HLIQSLKTLRSLLFHPRHGAWRCTDPSVYLSPFLDVVQSDQVPAAATGVALSSVLKILRLDVFDECSPGARDAAH
AVMAAVTNCRLERAADAGAEEAVLLRVLQVLAALLRARAAPLLSDHAVCTAVNTCFQVVQQAAGHRGGELLQRTA
RHCMHEILQAVFARLPDVRAPPDAGDNGSDNDNEGPVTAGAGYGARCMIDIFNFLCSLFNVGEMVMSPEGNSAFS
SEEDVQLFALVLINSAIELGGEAIGKHPKLLRMIQDDLFHHLIHCAIQSSPLVLSMICSTVLNLYHFLRRFLKLQ

LEAFFVYVLLRIGAGGSGPQLQEVAIEGTISFCRQPTFVVEMYVNYDCDPLSHNIYEDVGRLLCKAAFPVTNPMT
TIQTQAFEGLVYMITIIADNVEIEKAPDREAYNIDVSEYKPFWVERWDGDDNHETWVEFVRTRKLKKKKVMIAAN
HYNRDEKKGLDFLNLCHLVPSPPEPKSMAYFFRYTPGLNKNKIGDFLGDPDEFNLKVLKEFTETFDFSGAILDTA
LRTYLETFRLPGESQKIQRILEAFSEQFYEQQTSGIFATKDAAFILCYSLIMLNTDLHNPQVKKKMTEEDFIRNN
RAINAGMDLPREYLSELFHSISTNAITVFSQTGASVDMTTSRWADLVKRARTVEPFTPCDFKHKLSREVFIAISG
PTVATLAAIFDYTDDEDILNECVEGLVSVARIARYGLEDVLDELLSCLCKFTTLLNPYATTEETLFTFSNELKPR
MATLALFTIANRFGESVRGAWKNMIDCLLKLKRLKLLPQSVIEPDGGAAANSERAGGRSKSESGIIFPSSHRGVG
TSRHVSGMIGRFSQFLSLEGSAESLLSAGSEFENNLKIIQQCQIGNIFMESAKLPDESLQNLGRALIFAAGGKGQ
KFSTPVEEEETVGFCWDLILLITSTNLQRFTSFWPQLHECFTAVSQLPLFSACPFTEKAIVALFKIALEILSVPA
RLDRLPEELVLKSINLMWKLDKEILDTCCESISEAIVTLLTEHGGNVQTALGWKTLLHLLSVTGRLPETFDQSVG
TLIKMMTDGTHITRFNYAFCIEAAFGFAALKISPLETSTKILELMADSVNWLIQWHKSGYSDPGNSCASSSMSTS
SASMSSVEDAARMGNLAANLFVKLAEALRKTSLVRREEIRNQAVAELRRSFSLAAAGELELGPAGCLACFNLVIF
AMVDDLHEKTLEYSRREGAERETRSMEGTLAASTQLLVEVFVQFLGPLAQGPGFRTFWLGVLRRMDTCMKSDLAT
GGGTGVMQELVPEMLKRMILEMKDKQVLVQREGDELWEITNIQIQWIAPAVKEELFPE*
**>Kobli_KLT10238**
MGRPPRPPNSSIDPIEEEETVDTTTTTSPPSSPSPPLSALQSALSCMILSEIASVVAIMRRHLRHHPRDSDPFQD
PPLLLSFKSLRRLLFSPSASTGLTSSLLLRPFLDTIRSEETGAAVTAAALSAVYNILSFDGPVDVGPALSEVVDA
VTTCRFEVVDQSSEEAVLMKILQVLLECVKGGRDGSVGLDDHHVCTAVNTCFRVVHQAGGKSELLQRFSRNAMQE
LVRRVFGRLPLVGLDPTTDASSTKQEVGTVDKDQPFGVKQMENGTAIGSGTGVPDETIPSTEAPTRTIATPTIEP
YGVPCMVEIFHFLCSLLNVVEQVGVNPGTNTVAFDEDVPLFALRLINIAIELGGPSIKDHPKLLSLVQDELFRNL
MQFGLSMSPLILSMVCSIILNLYQNLRTELKMQLEAFFSCIILRLAQSKFGATYQQQEVAMEALVDFCRQRTFMV
EMYANFDCDITCRNVFEELSNLLSRSAFPLNSPLSSMHVLALEGLIAVVQGMSDRLGSGPLPEQASVQLTEYAPF
WTVKCESYADPGYWVKFVRQRKYIKRRLMIGADHFNRDPKKGLEFLQGTHLLPEKLDPQSVACFFRYTAGLDKNL
VGDFLGNHDEFCVQVLREFAWTFDFEGMNLDTALRLFLETFRLPGESQKIQRVLEAFSERYFEQSPHILANKDAA
LLLSYSLIMLNTDQHNVQVKKKMTEEDFIRNNRHINGGNDLPRDYLTELYHSICRNEIKTIPEQGFGFFEMSPSR
WVDLMRRSKTTPPYIVCGSWPNLDRDMFAVMSGPAIAAVAVVFDHAENAEVLSTCADGFLAIAKISAFHHLEDVL
DDLVVSLCKFTTLLTASAMADEPVTAFGDDLKARLATEALFLIATRYGDFVRTSWRNILDCILRLHKLGLLPARV
ASDAAEDAELAPEVVQGKPLPSSLSTTNMQVMGTPRRSSGLMGRFSQLLSLDTEEPRSQPTEQQLAAHQRTLQTI
QKCRIDSIFTESKFLHADSLLQLARALIWAAGRPQKMANSTDDEDTAVFCLELLIAITLNNRDRIVLLWQGVYEH
ISNIVQSTVLPCALVEKAVFGLLRICQRLLPYKENLADELLRSLQLVAKLDARVADAYCEGIALEVSRLVHANAN
HIRSQAGWRAVMVLLSMTARHQEASEHGFDGLKFIMMDGEHITPANYALCVEASRLFAESRVEVDRSVRALDLLA
GSVTCLTKWSKDAKEAGGDEGDKMAEAVREMWLRLTQSLRKTCLDPRDEVRNHALSCLQKTLIGSDAVCLMSASW
LQAFDLVVFATLDDLLENTSQKDYRNMETSLVIAVRLLSKVYLHLLPDLFGLSSFCKLWLGVLSRMEKCIKAKIR
GKRSEKLQEVITDLLRNILVVMKDRGILAKRSTIGGDSLWDLTWLHVNNISGSLQSEVFPGQEFDPDPRRRDLGA
PAENQE*
**>Kobli_KLT07006**
MVTVAPAVDHEPRLNQLSISCMLNTEVGAVLAVIRRPIDPTIVASAILLDDHLVQSLKSLRAIIFHTDWHTTDPA
VYLSPFLNVIQSDDVPAAATGAAMSAVLKIIRFGVFDDQSPGAFDAVHLVVSAITSCRLERTDPSIEDAILMRVL
QVLLALLHCPAGKLLTDHAICTIVNTCFQVVQQAVNRGDLLQRNARHLMHEVIQIIFLHLPEIKAPEPSSVDLSE
LEVELGSITGFGARCMVDIFNFLCSLLNAGDMVVAPEGYGPADEDVQLFTLVLINSAIELGRESIGRHPKLLHLI
QDDLFHHLIHYATRSGPLVLSMICSTVLNLYHFLRRSLRLQLEAFFVYVLLRIGSGMNGPHLQEVAIESIISFCR
QPTFLIEMFVNYDCDPIRHNVLEEIGKLLCKTAYPQINSQMTPIQIHAFNGLVHIFTTLADSIESTASDQLDGKD
KYKLDMSEYDIALDSDPPEPDSDTWVEFVRKRKLKKNKTMIAAHHYNKDEKKGMEYLKFANLVANPPETKTIARF
FRYTPGLDKHKIGDYLGDPGEFNIQVLKEFTNTFDFTGVILDTALRTFLESFRLPGEAQKIQRIIEVFSEKFFEQ
QSSDIFANKDAVFLLCYSLIMLNTDHHNPQVKKKMTEEEFIKNNRAINEGLDLPREYLSELFHSISTNAIALFSQ
NGSVPKELNANLWGELIRRSLTVDPFIICNFKHELSREVFATISGPSVATLMTIFDYTNDEEIRSECIEGMISVA
RIAQYGLSDVLDELLSCLCKFTTLLNPYNCTEDTQAEFNQSVKMRMAMLALFTIANRFGKSVRGAWKNVIDCIIK
LRRIKLLPPLGIDSSSGPATLVTTGLTANHLKSESGVIFPSSHQGLGTSRYISGMIGRFSQILSLDSGAESMINL
SSESEVNMKLMQQCQIGNIFTDSAKFSEESLHHLGRALIFAASGKGQKFTSVGEEHDTIIFCYDLLLTVTSANLH
RFMAFWQQFHEAFFTLSQYTIMSPYPFPEKAIAPLFKIAGKVISTETAQLDKQAEEAIVKSIHLMVTIENTIDMC
CDIIVASVKALIDEHGGSIQLPVVWRSLLIILSTSGRNPQTFDEAIETLNNLMIGPHLTRSNYLFWVDAAFTLGV
HKTTPLDKSVIILEMLGKSINPILQWHKLSITDPNNPNNNNSEGNMEDLGSRLTPYHLFMKLVEAIGKPSLARRE
EIRNKALIELKRCFQTAAESLDLSPKGCLNCFKNLVLPMVDNLHLKTIDYSRRENMERETISMGGTFREAMEFLA
DFYTIFLTQLWQDNDFMTSWLFVLRRMDTWIKTYYGGEFTMEIHAHIPQVLKRMLEAMKENGMLVPKEGDPLWES
TEVQAKWIAPSVIDELFPK*
**>Anaco_LOC109712899**
MGRPRLQNPSINPIEEEPSDSPDTPAAEAAAAPPPELSALACVVNSEVSAVLAVIRRYLRLGGGRYGSSAAAAAD
EHYLEHPLVLPLKSLRRHVFATAAAAPDPSAYLRPFLDVVCSDEALAPIKSVALSAVHKILTLDAAAAAFAGSPA
AIHGVVEAVASCRLDDADPTAEEAVLMRKLHVLLACMRARPALSDRHVCTIVNTCYGIADQVRAREGLLKELSRH
TLLDLVRCIFGRLPDAARNIPGPPPAVKPQMGGINKDQYFANKQAENGNGTDDNAGNVEASNGGGTTVQQSYGIP
CAVEVFQFLCSRLNVSSEIAPLDEETLVFGLTMINSAIELGGPSFRKHPKLLALIQDELFKNLMSFALLPSPPIL
SMVCSIVLNLYHHLRTELKLQLEAFFSCVILKLSQPRFGATYNQQEAAMEALVDFCRQKTFMAEMYANLDCDITC
SNVFEELANLLSKSAFPINCPLSPMHILALDGLIAVIQGMADRIGNAPSKLEQSSVELEEYTPFWTVKCENYSDP

DYWVKFVRQRKYIKRRLMIGADHFNRDPKKGLEFLQGTHLLPEKLDPQSVACFFRYTAGLDKNLVGDFLGNHDEF
CVQVLHEFARTFDFEGMNLDTALRLFLETFRLPGESQKIQRVLEAFSERYYEQSPHILANKDAALLLSYSLIMLN
TDQHNVQVKKKMTEEDFIRNNRLINGGSDLPREFLSELYHSICKNEIRTIPEQGFGFFEMSYSRWVDLMRKSKST
SPYIVCDSRPFLDHDMFAIMSGPTIAAIAVVFDYAEHEEVLLTCVDGFLAVAKISAFHHLEDVLDDLVVSLCKFT
TLSNSSLVEEPVTAFGDDMKARLATETVFNIANRYGDYIRTGWRNILECILRLHKLGLLPARVASDAADDLELAS
EAVQGKSAPASLSTSHLPIIGTPRRSSGLMGRFSQLLSLDTEEPRSQPTEQQLAAHQRTLQTIQKCRIDSIFTES
KFLQADSLLQLARALIWAAGRPQKVTSSPDDEDTAVFCLELLIAITLNNRDRIALLWQGVYEHIANIVQSTVMPC
ALVEKAVFGLLRICQRLLPYKENLADELLRSLQLVLKLDARVADAYCENITQEVSRLVKANATLIKSHMGWRTIM
SLLSITARHPEASEVGFEALVFIMSEGAHLSPANYVLCVEASRQFAESRVGLTDRSIRALDLMADSVNSLARWSR
ENKEAGEEAEKVLEGIREMWLRLIQAVRKLCLDQREDVRNQALSSLQRCLTNTEGICLLPTSWLQAFDLVIFGML
DDLLEIAQNHSQKDYRNMEGSLLVAMRLLAKVHLQLLPQLFGLSSFCKLWLGVLSRMEKYMKVKVRGKRSEKLQE
LIPELLKGTLQVMKARGILAKRSTIGGDSLWELTWLHVNNISTSLQSEVFPGQESEQELHASKQTDSGSPTQPDN
TSSGASAEVSGDQNGG*
**>Anaco_LOC109718116**
MAGSKHPNSDPGTPVAMPCVVISEVATLLAVMRANVRWAGGAAASSAAVGVRYGDDAGADEYLDHHLVRALKTLR
RRVLSWDPRRWRAVDPILYLKPFLDVIRSDETGAPITGAALSSVRKILALDALDPGAGAAAAAAAAAAAAASAVV
DAVAGCRFEVTDPASEEDVLMKILHVLLACVRGRAGALLGHRQVCTVVNTFFRIVQQAATKGELLQRVSRHTMQE
LVRCIFTRLPDMPVDGAADGPSVANGVGDSEARQIGGLDKDQTFGFESKQYENGNNDNGALTSSANEMAMHVDMK
VVNEPYGIPCMVELFQFLCSLLNSTEDIDMNPRVNPIAFDEDLPLFALGLINSAIELGGSAINRHPKLLNLVKDE
LFHNLMQFGLSRSPLVLSTVCSIVLNLFYHLRDELKVQLETFFSCVILILAQSRYGSSYQQQEVAMEALVDFCRH
ENFMVEMYANMDCDILCSNIFEDLTNLLSKNAFPVNSSLSSLHVLALDGLVAVIQGIAERVDNAPRNSEEISTEI
EEYFPFWLAKCENYADPDQWVRFIRHRKYIKKKLMVGVDHFNRDPKKGFEFLQGVHLLPDKLDPQSVALFFRYSS
GLDKNLIGDFLGSHDEFYIRVLHEFAQTFDFEGMNLDAALRLFLETFRLPGESQKIQRILEAFAERYYEQSPQVL
ANKDAALLLSYSLILLNTDQHNVQVKRKMSEEDFIRNNRGINGGNDLPREFLSEQYYSICRNEIRTIPQQGVGFH
EMSLSRWVDIMQKSKNTSPYILCESCPFLDRDMFAIMAGPTIAAISVVFDYGEYEEVLLKCVDGFLAVAKIAGAH
HLIDVLDDLVVALCKFTTLLNSSLIDEPVIAFGEDTRARMATETVFTIANTYGDYIRNGWKSIIDCILSLHKLGL
ISSRITGDEAADDPESSPDSSFLANPIPSALSTTSQTSSVSTPKRAYGLMGRFSLFLNLDTEESRSQPTEEQLAA
QRRALETIQKCRTGNIFTESKFLQAESLSHLAESLIRAAGRPQRITSGLEEEDTAVFCLELLIAVTLNNRDRIVL
LWQGVYEHIAHIVQCTVMPCNLVEKAVFGLLRICQRLLPYKENLVDDLLKSLQLILKLDARVADAYCENITQEVT
RLVKANATHIKSQTGWRTIISLLSITARHPDASDVGFEALVFIMSEGAHLSPANFILCIEASRQFAESRLGLTER
SVHALDLMGESVSCLARWAREVKEVGGEAERILEGIREMWLRLVQALKKVCVDQREEVRNHALLSLQRCLVGIEG
IYLSPLTWLQSFDIVFHLLDELLEIAQTYSPKDFRNMESSLLHAVKLLSKVFLQSLQDVSGESSFSKLWLGVLEH
LESYMKARMRGRKMEKLQEAVPELLKNTLLVMKANGVLVKRSTSDGSSLWELTWPHVENIAPSLQLEVFPSNELE
PETIDKQKELASWELVGATTASEGSSQAAV*
**>Anaco_LOC109722123**
MAKAHESLSDDETKTDGGAARRRDPRLKELGISCMLNTEVGSILAVIRRPSPSDPIADDPTAFSTASSHLVNSLK
SLRSLIFHPRHSAWRFTDPSVYLSPFLDVIQSDDVPAAATGVALSAVLKILRLDVFDERTPGARDAIHSIVSAIT
NCRLERTDPISEDAVLTRVLQLLSIVLRARASVLLTDHAVCTVINACFQVVQQSAHRTDLLQRSARHTMHDLVQA
VFARLPEVRAAESGDLGSDLEDDDGPPVGYGARCMVDIFHFLCSLLNVGEMVDFIEGYGSPSISSDEDVQLFALV
LINSAVELGGEAIGKHPKLLRMIQDDLFHHLIHYATRSSPLVLSMICSTVLNLYHFLRRSLRLQLEAFFTYALLR
IAAGIGGIQLQEVAIEGAISFLRQPTFAIETYVNYDCDPIRHNVFEEMLKLLCKIAFPVTNPALPMQIQAFEGLV
SVISNVADRIEVEKSPARDSYNLELSDYKPFWIERCEEDESNPDTWVDFVRTRKLRKKKIMIAANHYNRDEKKGF
DYLNLSNLVPSPPDSKSLAYFFRYTPGLDKNKIGDYLGDPSEFNLSVLKEFTETFDFSGVILDIALRTYLETFRL
PGESQKIQRILEAFSERFYEQQSSEIFVSKDAVFILCYSLIMLNTDQHNPQVKKKMTEEEFIRNNRAINGGMDLP
REYLSELFHSISTNAITLFGQTGLPVKLNSNKWADLVKRSRTVEPYITCEFKHKLSRELFAAISGPAVATLAAIF
EHTEDEEILHECVEGLISVARIARYGLEDILDELLCCLCKFTTLLNPYATAEETLFTFSNELKPRMATLALFTIA
NKYGESVRGAWKNLVDCLLKLKRLKVLPQSVIEPGNTSAASNSDVAAHPKTESGVIFPSSHSGFGGTSRYVSGLI
GRFSQFLSLETGADSMLNAGSEFENNLKIIQQCRIGSIFTESSKLPDESLQNLGRALVFAAAGKGQKFSTPVEEE
ETVGFCWDLLAMITTANLHRFTAFWPQLHDSFAVVSQFPLFSPCPFAEKAIVALFKIALKLLSGPAPRLDKLPEE
LIFKSINLMWKLDKEILDTCCESISESTRKILSEHPRNVQTALGWKTLLHLLSVTGRLPETFDQGVEAIIKLMSE
GTYVTRLNYAYCIEAAFGFAALKISPLETSSKILDLMAGSVDWLIQWHKSGFSDPGSGCNSGSSFTSNSSSTSSA
DEPGQRGSPGLAASLFVKLAEALRKTSLVRREEIRNQAVAALRRCFAAAAPAAADAAAEEAAAAAPLLGMGPASM
LACFNQVVFAMVDDLLEKMLEYARRENSEREMRSMEGTLKSAMDLLAEVLVMFLVPISQSPGFRSFWLGVLRRMD
TFMKASLGEAGESALMQETVPALLKRMILEMKEKEVLVQREGDELWEITNIQIQWIAPSVKEELFPEEF*
**>Phoda_LOC103696334**
MGRPRLQNSAINPIEEEPAAESPDAAASSPAWGGGGHHAALACMINSEVSAVLAVMRRNVRWGGRYASAADDHHL
EHSLVQSLKSLRRQVFSWGDARRPWCAVDPSAYLRPFLDVIRSDETGAPITSVALSSIYKILTLDVLDPGAPGAD
AAVHAVVDAVTSCRFEVTDPASEEAVLMKILQVLLACMRSRASSLLSNQHVCTIVNTCFRVVHQAGTKGELLQRF
SRHTMHELIRCIFSHLPDVGDGTEPLSIRSEIGGVDKDQAFGIKQVENGNGSGTVMGSGDDNALGVEVPNGVRIM
VEPYGIACMVEIFHFLCSLLNIGEQIGLSPRSNTIAFDEDVPLFALGLINSAIELGGPSICKHRKLLALIQDELF
RNLMQFGLSMSPLILSMVCSIVLNLYHHLRTELKLQLEAFFSCVILRLAQSRYGASYHQQEVAMEALVDFCRQKT
FMPEMYANFDCDITCTNVFEDLANLLSKSAFPVNCPLSSMHVLALDGLIAVIQGMAERIGNAPPRSEEPVELKEY
TPFWTVKCENCSDPDHLVKFVRRRKYIKRRLMIGADHFNRDPKKGLEFLQGTHLLPEKLDPQSVACFFRYTAGLD

KNLVGDFLGNHDEFCVQVLHEFAGTFDFQDMNLDTALRLFLETFRLPGESQKIQRVLEAFSERYFEQSPQILANK
DAALLLSYSLIMLNTDQHNVQVKKKMTEEDFIRNNRHINDGHDLPREFLSELYHSICRNEIRTIPDQGVGFLEMS
PSRWIDLMRKSKKTSPYIVCDSRPYLDHDMFAIMSGPTIAAISVVFDYAEHEEVFLTCVDGFLAVAKICAFHHLE
DVLDDLVVSLCKFTTLLNTSLVEEPVTAFGDDTKARLAAETVFTIANRYGDHIRTGWRNILDCILRMHKLGLLPA
RVASDASDDSELPLDPIHGKPAPSSLSTSHVPAMGTPRRSSGLMGRFSQLLSLDTEEPRSQPTEQQLAAHQRTLQ
TIQKCHIDSIFTESKFLQADSLLHLARALIWAAGRPQKVTSSPDDEDTAVFCLELLIAITLNNRDRIVLLWQGVY
EHIANIVQSTVMPCALVEKAVFGLLRICQRLLPYKENLADELLRSLQLVLKLDARVADAYCENITQEVTRLVKAN
ATHIKSQMGWRTITSLLSITARHPEASEVGFEALVFIMSDGAQLTPANFILCVEASRQFAESRVGLTDRSVRALD
LMAESLNSLVLWSRETREAVEDAERISEGIKEMWLRLVQALRKVCLDQREEVRNHALLSLQRCLVGAEGISLSSS
SWTQAFDPVIFTMMDDLLEVAQNHSQKDYRNMEGTLLHAMKLLSKVFLHLLQELYGLSTFCKLWLGVLSWMEKYM
KVKVRGKRSEKLQELIPELLKNTLLVMKTRGILARKSTIGGDSLWELTWLHVNNIAPSLQSEVFAGQELEQEVHA
KQRESGTPLQPDGNSSGSPAGASADQSGG*
**>Phoda_LOC103712126**
MGRPRLRNSAINPIEEEPAAESPDAANAASRAAAAAGGRHAALACMINSEVSAVLAVMRRNVRWGSRYASAADDH
HLEHSLVQSLKSLRRQVFSWGDARRPWSAVDPSAYLRPFLDVIRSDETGAPITGVALSSAHKILTLEVLDPGAPG
ADAAVHAVVDAVTSCRFEVTDPASEEAVLMKILQVLLACMRSRASALLSNQHVCTIVNTCFRVVHQAGTKGELLQ
RFSRHTMHELIRCIFSHLPDVGDGAEIDGVGKDRAVGIKQVENGNGSAGSVTGSGDDNFVGGVEVTNGERIMVEP
YGIPCMVEIFHFLCSLLNIVEQIGLSPRSNTIAFDEDVPLFALGLINSAIELGGPSIQKHPKLLALIQDELFWNL
MQFGLSMSPLILSMVCSIVLNLYQHLRRELKLQLEAFFSCVILRLAQSRYGASYHQQEVAMEALVDFCRQKTFMT
EMYANLDCDVTCGNMFEDLANLLSKSAFPVNCPLSSMHVLALDGLIAVIQGMADRRGNAPPHSEEPMELEEYTPF
WTIKCENYSDPDHWVKFVRCRKYIKRRLMIGADHFNRDPKKGLEFLQGTHLLPEKLDPRSVACFFRYTAGLDKNL
VGDFLGNHDEFCVQVLHEFAGTFDFEDMNLDTALRLFLETFLLPGESQKIQRVLEAFSERYFEQSPQILANKDAA
LLLSYSLILLNTDQHNVQVKKKMTEEDFIRNNRHINGGHDLPREFLSELYHSICRNEIRTIPEQGINFIEMSPSR
WIDLMRKSKKTSPYIVCDSRPYLDHDMFTIMSGPTIAAMSVVFDYAEHEEVFLTCVDGFLAVAKISAFHHLEDVL
DDLVVSLCKFTTLLNTSLVEEPVMAFGDDTKARLATETVFNIANRYGDHIRTGWRNILDCILRLHKLGLLPARVA
CDAADDSELPLDPIHGKPPPSSLSTSHVPAMGTPRRSSGLMGRFSQLLSLDTEEPRSQPTEQQLAAHQRTLQTIQ
NCHIDGIFTESKFLQADSLLHLARALIWAAGRPQKVTSSPDDEDTAVFCLELLIAITLNNRDRIVLLWQGVYEHI
ASIVQSTVMPCALLEKAVFGLLRICQRLLPYKENLADELLRSLQLVLKLDARVADAYCENITQEVTRLVKANAIH
IKSQMGWRTITSLLSITARHPEASEVGFEALVFIMSEGAHLSPANYILCVEASRQFADSRVGLTDRSVRALDLMA
ESLNSLARWSRETREVGEEAEKISEGIREMWLRLLQALRKVCLDQREEVRNHALLSLQRSLVGAEGMSLSPSSWT
QAFDPVIVTMLDDLLEISQNHSHKDYRNMEGTLLHAMKLLSKVFLQLLQELCVLSSFCKLWLGVLSRMEKYVKVK
VRGKQSEKLQELIPELLKNMLLVMKARGILAKRSTIGGDSLWELTWLHVSNIAPSLQSQVFAGQELEQVVHAKPK
ESGTPIEPNGSSSGPLAEAPADQSGG*
**>Phoda_LOC103695708**
MARASDDDDADEAAGRGEASGRHRRRDPRLKELGISCMLNTEVGAVLAVIRRPPDPISGFLPGGADEGANPQLLQ
SLKSLRSLIFHPQHSEWRSTEPSVYLXPFLDVIQSDEIPAAATGVALSAVLKILRLNVFEECTPSAREAIHSVVF
AVTNCRLERIEQAAEDAVLMHILQVLVAIVRSRVSILLTDHAVCTIVNTYFQVVQQSVHRGDLLQRSARHAMHEL
IQAIFSRLPEIKVETADRASQAAAADDSDTPISGYGACCMVDIFHFLCSLLNVGEVVDSAEGIGSISSDEDVQLF
VLVLINSAVELGGEAIGSTPKLLRMVQDDLFHHLICCNIIHYAACSSPLVLSMICSAVLNLYHFYRRCLRLQLEA
FFTYVLLRIAAGGSGVQLQEVVIEGILGFCRQPTFAIEMYVNYDCDPIRRNVFEEIGKLLCKTAFPATNPMSPMQ
VQAFEGLVTVITTIADSIEVEQVPDRESYSADVSDYKPFWVERCDSYDDPDTWVEFVRMRKMKKKKVMIAANHYN
RDEKKGLDFLNICHLVPTPPDPKCLAYFFRYTPGLDKNRIGDFLGDPDEFNLRALKEFTETFDFSGAILDTALRP
YLETFWLPGESQKIXRILEAFSERFYEQQSSEIFVSKDAVFILCYSLIMLNTDQHNPQVKKKMTEEEFIRNNRAI
NGGMDLPREYLSELFHSISTNAITLFGPTGLPNELSSNQWADLIKRSRVVEPFIICDFSHKLGRDLFPTISGPSV
ATLAAIFEHTDDEEILQECVEGLISIATIARYGLEDILDELLSCFCKFTTVLNPYATAEETLFAFSNELKPRMAT
VALFTIANKFGDSVRGAWRNIVDCMLKLKRLKLLPQSVIEPENSPSSDPGFTQHAKSESGVIFPSSHLGFGSRRQ
VSGLIGRLSQLLSLDSGGDSMLNCGSEFENNSKIIQQCRIGNIFCESSRLSDESLQNLGRALMFAAAGRGQKFST
PVEEEETVGFCWDILVMISTINIRRFAAFWPQFHDCVTVMSQLPLFSPCPFAEKAIVGLFKIALELFSSPPHVDK
LPEELVFRSINLMWKLDKEILDTCCESIVEGTARILNEHAKNVQTMIGWKTLLHLLSVTGRHPETFDQGVEALMK
LMCEGTHVTRFNYPYCIEAAFGYAALKISPLEKSSKILDLMGDSVNWLIQWHKSGYSDPGSTNSISSMEDGQKMG
AGAGNLEANLFVKQAEALRKTSLVRREEIRNQAVAELRKCFAAAEELDFTPANCLACFNLVIFAMVDDLHEKMLE
YSRRENSEREMKSMEGTLMEAMELLVEVFQRFMVPLSQSPGFRTFWLGVLRRMDTCMKVVIWTLGTRHVRCTSRG
RPHVHPTTPSRFDRT*
**>Phoda_LOC120105401**
MEMQMEMLQRPRWPGARDGDDAYEAAGGGEAGSRHRRRDPRLKELGISCMLNTEVGAVLAVIRRPPDPISGFLPG
GADEGANPQLLQSLKSLRSLIFHPQHSEWRSTEPSIYLSPFLDVIQSDEIPAAATGVALSAVLKILRLNVFEERT
PGAREAIHSVVFAVTNCRLERTEQAAEDAVLMRILQVLVAIVRSRVSVLLTDHAVCTIVNTCFQVVQQSVHRGDL
LQRSARHAMHELIQAIFSRLPEIKVPETADRASQAAAAADSDTPISGYGARCMVDIFHFLCSLLNVGEVVDSAEG
IGSISSDEDVQLFALVLINSAVELGGEAIGKHPKLLRMVQDDLFHHLIHYAACSSPLVLSMICSAVLNLYHFCRR
CLRLQLEAFFTYVLLRIAAGGSGVQLQEVAIEGILGFCRQPTFAIEMYVNYDCDPIRRNVFEEIGKLLCKTAFPA
TNPMSPMQVQAFEGLVTVITTIADSIEVEQVPDRESYSADVSDYKPFWVERCDSYDDPDTWVEFVRMRKMKKKKV
MIAANHYNRDEKKGLDFLNICHLVPTPPDPKCLAYFFRYTPGLDKNRIGDFLGDPDEFNLRALKEFTETFDFSGA
ILDTALRTYLETFRLPGESQKIQRILEAFSERFYEQQSSEIFVSKDAVFILCYSLIMLNTDQHNPQVKKKMTEEE

FIRNNRAINGGMDLPREYLSELFHSISTNAITLFGPTGLPTELSSNQWADLIKRSRVVEPFIICDFSHKLGRDLF
AAISGPSVATLAAIFEHTDDEEILQECVEGLISIATIARYGLEDILDELLSCFCKFTTLLNPYATAEETLFAFSN
ELKPRMATVALFTIANKFGDSVRGAWRNIVDCMLKLKRLKLLPQSVIEPENSPSSDLGFTQHAKSESGVIFPSSH
LGFGSRRQVSGLIGRLSQLLSLDSGGDSMLNCGSEFENNSKIIQQCRIGNIFSESSRLPDESLQNLGRALMFAAA
GRGQKFSTPVEEEETVGFCWDLLVMISTINIQRFAAFWPQFHDCVTVMSQLPLFSPCPFAEKAIVGLFKIALELF
SSPPHVDKLPEELVFRSINLMWKLDKEILDTCCESIVEGTARILNEHAKNVQTMIGWKTLLHLLSVTGRHPETFD
QGVEALMKLMCEGTHVTRFNYPYCIEAAFGYAALKISPLEKSSKILDLMGDSVNWLIQWHKSGYSDPGSTNSSSF
MEDGQKTGAGAGNLGANLFVKLAEALRKTSLVRREEIRNQAVAELRKCFAAAEELDFTPANCLACFNLVIFAMVD
DLHEKMLEYSRRENSEREMKSMEGTLKEAMELLVEVFQRFMVPLSQSPGFRTFWLGVLRRMDTCMKADAGGMGDG
ALQELVPTLLKKMITEMKHKEILVQRDGDELWEITNIQIQWIAPSVKEELFPEEF*
**>Aspof_LOC109850386**
MPSRSSSSAMARLSSMTPSYQRSTKTPRHHRQSPSPDALACVINSEVAAVLAVIAPHVRWGGRYASAATTTSTQL
LSSPLKSVRRQIFSWDTTALGLRGPHSYLXPFLDVIRSTRTGPPRSPASRYHQVYKILTLDALDPRATFNVGEAM
QVVVGCGSSCRFEVTDPASEEAVLMKILQVLLACMKSRASAVLSNQHVCTIVNTCFRVVHQAGTKGELLQRISRQ
TMHELIRCIFSHLAEVNGDGSERSNVKQPEMGGVDMDQGFGIEQAENGNGGDGTVVGAENGLVAAEVPNGMRAMV
EPYGVPCMVEIFHFLCSLLNVVEQIGMSPRSSPIAFDEDVPLFALGLINSAIELGGSSIGKHPKLLALIQDELFR
NLMQFGLSMSPLILSMVCSIVLNLYYHLRTELKLQLEAFFSCVVLRLAQSRYGASYHQQEVAMEALVDFCRQKTF
MPEMYANLDCDITCSNVFEDLANLLSKSAFPVNCPLSAMHVLALDGLIAVIQGMADRIANAPPRMEEASVELEEY
SSFWTVRCENYSDPEHWVRFVRRRKYIKKRLMIGADHFNRDPKKGLEFLQGTHLLPEKLDPQSVACFFRYTAGLD
KNLVGDFLGNHDEFCVQVLHEFAWTFDFEGMNLDTALRLFLETFRLPGESQKIQRVLEAFSERYYEQSPQILANK
DAALLLSYSLIMLNTDQHNVQVKKKMTEEDFIRNNRHINGGNDLPREFLSELYHSILQNEIRTIPDQSIAFPEMT
PSRWIDLMRKSKKTSPYIICDSRPFLDHDMFAIMSGPTIAAISVVFDYAEHEEVFLTCLEGFLAVAKISAYHHLE
DVLDDLVVSLCKFTTLLNASLVDEPVTAFGDDSKARMATEAVFNIANRYGDFIRTGWRNILDCILRLHKLGLLPA
RVASDAADDSELPSDTVHGRPPPSSLSTSLPSMGTPRRSSGLMGRFSQLLSLDTEEPRSQPTEQQLAAHQRTLQT
IQKCHIDSIFTESKFLQADSLLHLARALIWAAGRPQKVTSSPDDEDTAVFCLELLIAITLNNRDRIVLLWQGVYE
HIANIVQSTVMPCALVEKAVFGLLRICQRLLPYKENLADELLRSLQLVLKLDARVADAYCENITQEVTRLVKANA
THIKSHTGWRXTSLLSITARHPEASEVGFEALVFIMSEGAHLSPANYVLCIEAARQFAESRVGLIDRSVRALDLM
AESVVCLARWSTEAKEAGEEAEKIQEGIREMWLRLVQALKKVSIDQREEVRNHAITSLQRCLVGAEGICLSPSSW
LQSFDLVIFAVLDDLLENAQSQSQKEYRNMEGTLLIAMKLLSKVFLQLLQELSGLSSFCKLWLGVLGRMEKYMKV
KVRGKRSEKLQELIPELLKNTLLVMKTKGILAKRSTIGGDSLWDLTWLHVNNIAPSLQTEVFPDQESEQTDRSGS
GVEEGNQGGG*
**>Aspof_LOC109851574**
MARTKDNSDDADDGEARSSPRGGRRRDPRLKDMGISCMLNTEVGSVLAVIRRTPVDLSSSGAGGAPSSPHDIADL
HNSNIINSLKSLRSLLFHPCHPEWRLTNPSVYLAPFLDVVQSDDVPASATGVASAVLKILKLDVFDERTPGARDA
VHSVIFAVTNCRLERTDPASEDAVMMRILQVLTGVMRTRASLLLTDHAVCTIINTCFQXVFSRLPELRSENTSAT
HDEESLSGSDADDDPVVIGYGARCMVDVFHFLCSLLNVGEVVDDGGGVSSDEDVQVFALVLINSAIELGGEGIGK
HGKLLGIIQDDLFHNLIHYATRSSPLVLSMICSTVLNLYQFLRRSLRLQLEAFFTYVLLKIASGSSGLHLQEVAV
EGILGFCRQPTFIIEAYVNYDCDPLHYNVFEELGKLLCKTAFPLSAPMSPMQIQAFEGLVTIIRNIADGVETDKI
PVKDSYDLDISEFKPFWTERCDTYHENTESWVDFVRMRKLKKKKIMVAANHYNRDEKKGLDYLKICQLVPTPPDP
KSLAYFFRYTPGLDKKKIGDYLGDPDELNLKVLAQFTSTFDFSGVTLDTALRTYLESFRLPGESQKIQRILEAFS
DQYYKQQSSDIFASNDAVNILCYSVIMLNTDQHNPQVKKKMTEEEFIRNNRSINGGNDLPREYLSSLFNSIATKA
IAVLVESTGAVLELNANRWADLVKRARASEPFIVCDYKHKLSREVFVTISGPSVATLSAIFEQTDDEDILNECVD
GMISAARIARYGLEDILDELLCCFCKFTTLLNPYATAEETLFTFTNEMKPKMATLAVFTIANKFGDAIRGSWKNL
IDCLLKLKRLRLLPQSLVEPENASKQEPEINSSHIKTESGVIFPSSQFGAGGNNRNASGLMGKFSQFLSMDNGQE
SMLNVGSEFENNLKIIQQCRIGSIFTECSKLPDESLQNLGRSLIFAAGGKGQKFSTSVEEEETVGFCWDLITCIT
SANLHRFSTFWPQFHECLGIVSQFPLFSPCPFAEKAIIAFFKIAVKLLSSSTAHFDKTPEDLIFKTINATWKLDK
DILDTCCCDIADSTTEILNDHVMNVQTMIGWKTLLHMLSITGRHPETFDQGVESLVKLMSNGTHVTRFNYPFCIE
TAFGFAALKIIPLETSFKILDLMAGSVSWLIQWHKSGYSLAANLFLKLAEALRKTSLVRREEIRNQAVQALRRCF
QGTEELEFTPANCLACFNLVIFALVDDLHEKMLEYSRRENSVKEMRSMEVTLIEAMGLLVEVYLQFMVSLSQSPG
FRTFWLGVLRRMDTCMKANMDEAGYGPLQEAVPRLLKKMLVQMKEKEILVQRDGDELWDITNIQIQWIAPQVKEE
LFPDVF*
**>Denca_LOC110105664**
MGRLRLQNPGINPIDEESLADIDTDQSPNPPPDNSCCSSSGHALACMINSKVSAVLAVMRRNVRWSGRYASTADE
YLDHSLIQSLKSLRRQLFSWDTSRWSSIDPTLYLRPFLDVIRSDETGAPITGVALSSVHTILALEVFDAGTAHVE
DAMHAVVNAVTSCRFEVTDPASEEAVLMKILQVLLACMKSRASMVLSNEHVCTIVNTCFRVVHQAGTKGELLQRI
SRNTMHELVRCIFVRLPDVSKSNGKLSSLVEMAGIDKNQAFGMSQMENGNGNYGSLTIPVDDSTVRLEMPNGMKI
MVESYGIPCMVEIFHFLCSLLNIVEQIGISPRTNPMALDEDVPLFALRMINSAIELGGSSISRHPKLLALIQDEL
FRNLMQFSLSMSPLILSTVCSIVLNLYCHLRIQLKLQIEAFFSCVILRLAQGKYGASFHQQEVAMEALVDFCRQK
TFMAEMYANLDCDITCSNVFEDLANLLSRSAFPVNCPLSTMHILSLEGLIAVIQRMADRISNAPPRSRQALLELD
EYTPFWTVKCENYSDPEQWLKFVRCKKYIKRRLMIGADHFNRDPKKGLEFLQGNHLLPEKLDPQSVACFFRYTAG
LDKNLVGDFLGNHDDFCVQVLHEFAGTFDFQDMNLDTALRLFLETFRLPGESQKIQRVLEAFSERYYEQSPHILA
NKDAALLLSYSLIMLNTDQHNVQVKKKMTEEDFIRNNRHINGGNDLPREYLSELYHSICKNEIRTTPDPGIGFLE
MSPSRWIDLLRKSRKTSPYIVCDSQPYLDHDMFAIMSGPTIAAISVVFDYAEHEDVFLTCVDGFLALAKISAYHH

LEDVLDDLVVSLCKFTTLLNTSMAEEPVTAFGDDTKARLATETVFIIANTYGDHIRTGWRNILDCILRLHKLGLL
PARVASDAADDTELLADAAYGKPAVNSLSASHIPSMGTPRRSSGLMGRFSQLLSLDTEEPRTQPTEQQLAAHQRT
LQTIQKCHIDSIFTESKFLHADSLLHLARALIWAAGRPQKVTSSPDDEDTAVFCLELLIAITLNNRDRIFLLWQA
VYDHIANIVQSTVMPSALVEKAVFGILRICQRLLPYKESLADELLRSLQLVLKLDARVADAYCENITQEVSRLVK
ANATHIKSHMGWKTITSLLSITARHTEASEIGFEALVFIMSEGSYLSPANYLLCIEALRQFAESRVGLTDRSTRA
LDLMAESITCLIRWSQEIKDAGEESEKLSEGIRDMWLKLVQALKKLTLDQREEVRNHAMSSLQKCLLEVDGICLL
PSSWLHSFDIVIFMMLDDLIEIAQSQSQKDYRNMEGTLMHAVKLLSKAFLHLLQELSGLSSFCKLWLGVLSRMEK
YMKVKVRGKRSEKIQELVPELLKNILLVMKSKGILAKRSTIGGDSLWELTWLHLNNFAPSLQSEVFTGELELDSS
NHTQSDGSHSAVVEEGSPSG*

**>Denca_LOC110102584**

MGRLGLQNLGINPILEESLADDGGNRLFPLPELAESSSSSCGHAMACMLNSEVSAVLAVMRRNVRWGGRYASAGD
DHLDHSLIQSFKFLRRQLFNWDPNRWSSIEPTLYLRPFLDVIRSDETGAPITGVALSSIHKILSFEMFDAESAHV
EEAMNAIVDAVTSCRFEVTDPVSEEAVLVKILQSLLACMRCRACKVLSNQHVCTIVNTCFRVVNQAGTKGELLQR
ISRHTMHELVRCIFSHLPEVSERTFKASSNAEMVAFEKDESFGLQQLGKSNGNLGLMTSSVNDNVVGLETPDGTR
AMTEPYGIPCMVEIFHFLCSLLNIADEIGISPWTNHMALDEDVPLFALGMINSVIELGRSSIRRHPKLIALIQDE
LFRNLMQFGLSISPLLLSTVCSIVLNLYYHLRTELKLQLEAFFSCVILRLAQGKNGVSYHQQEVALESLVDFCRQ
KTFMAEMYSNLDCDITCCNMFEEITNLLSRCAFPVNSSLSMTHILALEGLIAVIQGMVQRIGNLPPNSQQPLLEL
HEYTPFWCVKCENYSDPEHWLEFVRRRKFIKRRLMNGADHFNRDPKKGLEFLQGIHLLPEKLDPKSVACVFRYTV
GLDKNLVGDFLGNHDEFCVQVLHEFAGTFDFQDMNLDTALRIFLETFRLPGESQKIQRVLEAFSERYYEQSPHIL
VNKDAALLLSYSLIMLNTDQHNMQVKKKMTEEDFIRNNRLINGGNDLPREFLSELYHSICKNEIRITPDPGFGVP
EMSPSRWTDLMRKSKVTSPYVVYDSRPYLDRDMLAIMSGPSIAALSVVFDHAQNEDVLLTCVDGFLAMGKISAYY
HLGDVLDDLVVSLCKFTNLLNSSMFDEPITAFGDDTKARLTTETVFNIANNYGDHIRTGWRNILDCILRLHKVGL
LSAQLATDAAADVEPTEDAEPGKPPMNSVSASNIPSVGITRRSSGLMGRFSQLLYFDTEELKIEPTEQQLAAHQR
ALQTIQKCHINNIFTESKFLHADSLLHLARALILEADRPPNFSSSPEDEDTAVFCLELLIAITLNNRDRIALLWQ
DVYKHIANIVQSIAMPCALIEKAVFGLLRICQRLLPYKEDLTDELLKSLQLVLKLDARVADVYCENITQEVTRLV
KANSTHIKSQMGWKSIISLLSIAARHPDASEFGFEGLVFIMSDGAHLSPANYILCIEASRQFAESRVGLIERSTR
ALDLMAESTICLIRWSQETKKAGGEAEKITEGIQELWLQLIQALRKVTLDQREEVRNHAVALLQKCLAGVEGISF
SPSVRLQSFDIVIFKMLDNLLEIAQGPSQKDYRNMEVTLVHAIKLVSKVFLHLLHELCGLSSFCKLWLGILSRME
RFMKIKFRGKRSEKLQELVPELLKNILLVMKSTGVLVQNSTVGGDSLWELTWFHLNKFAPSLQSEVFPEGELEQK
DHSNQPKTDGSG*

**>Denca_LOC110092941**

MAGVTKDGDTGGHQPPPCSGHRRLCEQGISCTLNTEVGMMLAVIRQPPDPTISPTLHGAPETDSYNSHHVQSLLS
LRSIIFHPLRGSWHSLDPMVYLSPFLDIIQSEDVTAASTSAALYAVLKIMRLDIFSEITPGAREAIHAVVLAITN
CRLELTDPASEDSILMHIHQVLTAVMHSNASVLLTDHAICTIVSSCFQIVQQSADRSDLLQRSARHAMHDLIEGI
YSRLSNMRYSDGGPVDSGSEAEKNALFLHVSYTAQCMIDIFSFLCSLVNVEEPVDPETADASSDEEFQLFVLLLI
NTVVELGGDSINHHPGLLHIIQDDLFHHLIHYSTRSGQRILSITCNIVFSLYHFLRGSLRLQLEAFFIYVLLKIA
KGSYGSQLQEVAVEGITSFCRQPNFIMEIYVNYDCDPIRHNVLEEIVKLLCKTAYPMSSPISSMQLQAFEALISI
IHNIADGIKIHHALSRDAYAIDDSTFKPFWLERCDMHKDPDSWVEFVRMRKLKKKKIMMAANHYNRNEKKGMAFL
NICRLVPSPPDAKSIAYFFRYTPRLDKNKIGDYLGESNDFNIKVLNEFAQTFEFDGVILDTALRTFLETFRLPGE
SQKIHRILEAFSERFYDQQTSGIFASKDAVFILCYSLIMLNTDQHNPQVKKKMTEDEFIRNNRAINGGVDLPREY
LSELFLSISTNAITLFDSSSAVISEMNSHLWANLMKQTRTVEPYILCDYKHKLCREVFNTISGLSVATICVIFEQ
TDDENVLQECIEGLFSIARIARYSLEDILDELLFSLCKYTTLLNPYATAEEIVFSFSNEVKPRMATLAIFTIANK
FGESIRGAWRNLVDCLLKLNRLKLLPPTLLETCRSATEADIDNDGGRHSKSESGSGVIYPPSYLGSDRRNISGLV
GRFTELLSLESGADSMLNAASDMENNLKIIQQCRIDSIFKDSAGLPQESLQHLIRALVFAAAGKGQKFSTSVEEE
ETVGFCWDLIFIIIITANLHRLSAFWMQFNECLALVSQVPQFSPCPFVEKAMVALLRVVILILSSEPPRLQDRQSE
EMVFKSINLTWKLDKDILDACNDSVTEATMKILSKYARNVQTTLGWKTLFHLLSVTGRHPENFDRGVETLIGLMT
EADHITRSNYAYCIEAAFGFAALKISPVEKSSQILNLMAGSVKWIIQWQKSSFYETGNSNMHNSSSSTEEGINKA
SSVQGNSFAATNLFMKLAEALRKTSLVRREEIRNQAVEALGWCFLEVADELEWTAGSYASSFNLVIFAMVDDLHE
KMVEYSRREGMEREMRSMEGTLKMALEQMVEVYLCFMVRLAAGPGFRTFWLGLLRRMDTCMKASLGEEGLPSPVM
QELVPKLLKRMILEMKEKQVLVQSDESDLWEVTNIQIQWIAPSVKEELFPDD*

**>XerviGNOMa_sc13**

MSRSRSTINSISEEEDEPSDQAALACVINAEASAVLAVMRRNVRWGGGVGAVNGRYSPAAAVADPLDHPLVGSLK
SLRRCVFSWGPFSSVDPALYLRPFLDVVRSDETGAPITAAALSALHKILSLDIDAAAAAGNAMRAVVDAVTSCRF
EVTDPASEEAVLVKILQVLLATMRCRAAAVLSNQHVCTVVNTCFRIVHQAGSKGELLQRVSRHMMHELIRTIFAR
LPLVTDSSSQIGMHDKGHVFYTKYTENGNGNGTTEECAALGDGVGGYGIPCMIEIFHFLCSLLNIVEQIGMVPRA
NGVAFDEDVPLFALGLINSAIELGGSSIKKHPKLMSLIQDELFRNLMQFGLSMSPLILSTVCSIVLNLYHHLRTE
LKLQLEAFFSCVVLRLAQNRYGATYQQQEVAMEALVDFCRQKSFMPEVYANLDCDITCSNVFEDLANLLSKSVFP
VNCPLSSMHVLALDGLIAVIQGMADRIGNAPTSTSHRPLELEEYTPFWTVKCEDYADPEHWVRFVRRRKYIKRRL
MIGADHFNRDPKKGLEFLQGTHLLPEKLDPQSVACFFRYTAGLDKNLVGDFLGNHDEFCVQVLHEFAKTFDFQDM
HLDTALRLFLETFRLPGESQKIQRVLEAFSERYYEQSPQILVNKDAALLLSYSLIMLNTDQHNVQVKKKMTEEDF
IRNNRLINGGNDLPREFLSELYHSICRNEIRTIPDQGAGFLEMSPSRWVDLMRKSKNTSPYIVCDFRPFLDHDMF
AIMSGPTIAAISVVFDYAEQEEVFLTCIDGFLAVAKISAYHHLEDVLDDLVVSLCKFTTLLSTSLVDEPLTAFGD
DTKARIATEAVFTIANRYGDYVRTGWRNILACILRLHKLGLLPARVASDAADDPDLSPDPGHGKPAPTSISTSHI

PPIGTPRRSSGLMGRFSQLLSLDTEEPRSQPTEQQLAAHQRTLQTIQKCHIESIFTESKFLQAESLLQLARALIW
AAGRPQKVTSSPDDEDTAVFCLELLIAITLNNRDRIALLWQGVYEHIANIVQSSVMPCSLVERAVFGLLRICQRL
LPYKENLADELLRSLQLVLKLDARVADAYCENITQEVTRLVKANATHVKSQMGWRTITSLLSITARHPEASEVGF
EALVFIMSEGAHISPANFILCIEASRQFAESRVGVTDRSLQALDLMAGCVTCLGRWSREAKEVGEESEKMSEGIR
EMWLRLVQALRKVCTDQRDDVRNHALSSLQKCMVGVDVISLTTSLWLQTFDVILFTLLDDLLEITQNHSQKDYKN
MEITLVSAMKLLSKVFLQQLQELAPLNSFCKLWLGVLGRMEKYMKVKVRGKRSEKLQEVIQELLRNTLLMMKTKG
VLAKRSTIGGDGLWELTWLHANNISTSLQAEVFPDQEIDQDAKATTNPGQSEDVGSPASEGVDGAG*

**>XerviGNOMb_sc7**

MAPAGGDTPAVVPSLSCVIAAEAATVLAVMRRNVRWSSVGVRYAPNAVNDDDDSLDHPLISSLKSLRRRVFLSSR
HVVEMGPTEYLKPFLDVVKSDETGAPITGAALSSIHKLISLFALDSDSARAVIDAVATCRFEVTDSASEEAVLMR
ILQVLLQCVRRGAAAVGGKMIGDRHVCAAVNACFRIVHQARSKSEILQRFSRHTMHELVRCVFLRLPAVDGGTEN
GNTNEDAATSGAQKRVLEVEEADGDTGNVEEPFGIPCMVEIFQFLCSLLNIAEHMGMSPRSNPIAFDEDVPLFAL
GLINSALELGGPWISKHPQLLSQVQDELFHNLMEFGLSMSPLIMSMVFNIVLNLFQHLRTQLKLHIEAFFSCVIL
RLAQSRYGATYQQQEVALEALVDFCRQDTFMVEMYANMDCDISLSNVFEDLINLLSKSAFPINSPLSSLQVLALE
GLVAVVQAMADRLDYAPSVSYEPVADIHEYTPFWTDKCENCQDPDFWVKFVRQRKFIKRKLMIGADHFNRDPKKG
LEFLQGIHLLPEKLDPKSVACFFRYTTGLDKNLLGDYLGNHDQFCIQVLHEFARTFDFKDMNLDTALRLFLETFR
LPGESQKIQRVLEAFSERYYEQSPQILANKDAALILSYSLIMLNTDQHNVQVKKKMTEEDFIRNNSRINGGNDLP
REFLSELYYSICENEIKTIPDQGTGFLEMTYSRWTDLLRKSKRTSQYILCDSRPFLDQDMFAIMSGPTIAALSVV
YDYAEHNEVLLTCIDGFMAVAKISASHHVRGllddllllCKFTTLLSTSSVEEHVIAFGDDLKATLATETVFKI
ANRYGDYIYNSWKNVVDCILTLHKLDFLPAHLTGDTADDTEPLSPSHAKPTANPNLPSRVPSINHRRASSGLMGR
FSQLLSLDAEEVRPQPTEEQLAIHQRTLETIQKCDIDSIFTESKFLQAESLVHLVRALIRAAGQPEKVTGADDED
TAVFCLELLVAVTLNNRDRIKLIWQDVYEHISSIVQSTCMPCPLVEKAVFGLLRICQRLLPYKEDLADDLLRSLQ
LILKLDARVADAYCENITQEVTRLVKANASHIKSQMGWRTIASLLSITARHPDAAEVGFEALVFIMSDGAHLSPA
NFILCVEASRQFAESRVGPSERsihslslmsesvsslsrwsHEIKETHEEADNILEGLREMWLRLVLAMKRTCLD
QREDVRNHALLSLQRCLVAVEGVCLSSTSWIQAFDLVIFPCLDDLLEISQSRSQKDYRSMEESLLRAMKLTSKAF
LQSMQDISSLSNFPNIWLAVLERMEKYMKAKVRGRRSEKLQELIPELLKNTLLVMKTTGILVQKKTPEEDSLWEL
TWLHVNNIDPSLQLEVFPAQEKKTEAS*

**>XerviGNOMc_sc11**

MSPYGSPSSPPRDPRQSSSLYPAIPQTPDGSRRGDLPVAMPSLYCVITAEVATVLAVMRRNVRWSSFGIRYVTDA
ANNDDGSLDHPLISSLKSLRHRVSLSFRNVAELGPTEYLKPFLDVVKSDETGAPITGAALSSIHKLISLFTLDSD
SARAVIDAVATCRFEVTDLASEEAVLMRILQVLLQCIRSGAASQKMVGDRHVCAAVNACFRIVHQARSKSEILQR
FSRHTMHELVLCVFSRLPAIENGNTNEDATTSGAQKQVPEVEEADGGAGIVEEPFGIPCMVEIFQFLCSLLNIAE
HMRMSPRSNPVTFDEDVPLFALGLINSAIELGGPWIGKHPQLLSVVQDELFYNLMEFGLSMSPLIMSMVFSIVLN
LFHHLRTELKLQLEAFFLCVMLRLAESRFGASYQQQEVALEALVDFCRQDNFVVEMYANMDCDISSSNVFEDLIN
LLSKSAFPINSPFSSLQVLALEGLVAVVQAMADRLDCAPSIFYEWIADFHEYTPFWTDKCENCSDPDLWVKFVRQ
RKFIKRKLMIAADHFNRDPKKGLDFLQGIHLLPEKLDPKSVACFFRYTKGLDKNLLGDYLGNHDQFCIQVLHEFA
QTFDFQYMNLDTALRLFLETFRLPGESQKIQRVLEAFSERYYEQSPQILANKDAALILSYSLILLNTDHHNVQVK
KKMTEEDFIRNNRRINGGNDLPRNFLSELYYSICENEIKTIPDQGADFLEMTYSRWTDLLRKSKRTSQYILCDSR
PFLDQDMFAIMSGPTIASLSMVYDYAEREEVLRTCINGFMAVTKIAASHNVRDLLDDLLALLCKFTTLSSSSSME
DHVIAFGDNMKATLATETVFKIANRYGDYIHNGWKNIVDCILTLHKLDLLPVRLTSETVDDTEPLSPFHAKPAAN
PNLPSRVPSMNHRKTSSGLMGRFSLLLSLDAEEVRSQPTEEQLAVHQRTLETIQNCHIDNIFTESKFLQAASLVH
LVRALIRAAGQPEKITGADDEDTRVFCLELLIAVTLNNRDRIKLLWQDVYEHISSIVQSISMPCPLVEKAVFDLL
RICQRLLPYKEDLADDLLRSLQLILKLDARVADAFCESITQEVTRLVKANASHIKSQMGWRTIASLISIAARHPD
AAEVGFEALVFIMSDGAHLSPANFILCVEASRQFAESRVGPSERSIQTLSLMSESVSSLYRWSCEVKEAHEEAEQ
ILEGLREMWLRLVLAMKRACLDQREDVRNHALLSLERCLVAVEGLCLPSSSWIQAFDLVIFPCLDDLFDISQSRS
LKDYRSMEESLVRAIKLTYKAFLQSTQEISSLSNFSKIWLGVLEHMEKYMKAKVRGRQCEKLQELIPELLKTMLL
VIKATGILVQKKSPEGDSLWELTWHHVNNIDSSLQLEVFPLQEKKLNEVDSGTILKPVANISGPLTGVAG*

**>Dioro_LOC120267042**

MGRSRLQNSGINPIeeespdspdasassgpsraSLACVINAEVAAVLAVMRRNVRWGGAGSGAARYLPSAADDHL
DHPLVQSLKSLRGQLFSRDPAHFLDPAAYLRPFLDVVRSDETGAPITGVALSAVYKILTLDGALRPGDGDAIRGV
VDAVTSCRFEVTDPASEEAVLMKILQVLLAAMRSPVAISLSNQHVCTVVNTCFRVVHQAGTKGELLQRVSRHTMH
ELIRCIFSHLPDVGDGTEPLSIKSEMGAVDKDQSFGVNHAEngngnvgslgnsgddgvLEVETPNGMRVMMEPYG
IPCMVEIFHFLCSLLNIVEHMEMSPRMNPIALDEDVPLFALGLINSAVELGGPSIRKHPKLLSLIQDELFRNLMQ
FGLSMSPLILSMVCSIVLNLYHHLRTELKLQLEAFFSCVILRLAQSRYGASYHQQEVAMEALVDFCRQKTFMPEM
YANLDCDITCSNVFEDLANLLSKSAFPVNCPLSSMHVLALDGLIAVIQGMADRIGTAPPRTEQAPLELAEYTPFW
TVKCEDYSDPEEWVKFVRRRKYIKRRLMIGADHFNRDPKKGLEFLQGTHLLPEKLDPQSVACFFRYTAGLDKNLV
GDFLGNHDEFCVQVLHEFAGTFDFEDMNLDTALRLFLETFRLPGESQKIQRVLEAFSERYYEQSPQILANKDAAL
LLSYSLIMLNTDQHNVQVKKKMTEEDFIRNNRLINDGSDLPREFLSELYHSICRNEIRTIPDQGIGFPEMSPSRW
IDLMRKSKKTSKYIICDSRPFLDHDMFAIMSGPTIAAISVVFDYAEQEEVFLTCVDGFLAVAKISAYHHledvld
dlvVSLCKFTTLLSTSLVEEPVTAFGDDMKARLATETVFNIANRYGDFIRTGWRNVLDCILRLHKLGLLPARvas
daaddsevasdpVHGKPVPNSLSTSHIPAMGTPRRSSGLMGRFSQLLSLDTEEPRSQPTEQQLAAHQRTLQTIQK
CHIDSIFTESKFLQADSLLQLARALIWAAGRPQKVTNSPDDEDTAVFCLELLIAITLNNRDRIVLLWKGVYEHIA
NIVQSTVMPCALVEKAVFGLLRICQRLLPYKENLADELLRSLQLVLKLDARVADAYCENITQEVTRLVKANATHI

KSQMGWRTITSLLSITARHPEASEVGFEALVFIMSEGAHLSPANYILCVEASRQFAESRVGLTDRSVRALDLMTE
SVTSLARWSRETRDAGEEAEKISEGIREMWLRLVQALRKVCLDQREEVRNHALSSLQRCLVGLEGLCLSTMSWLQ
AFDLVIFPLMDDLLETAQNHSQKDYRNMEGTLLLSMKLLSKVFLQWLQELSALSSFCKLWLGVLSRMEKFMKAKV
RGKRSEKLQELIPELLKNTLLVMKTKGVLAKRSTIGGDSLWELTWLHVNNMAPSLQLEVFPGQESEQESLAKQKD
SAGSVQPDDNTSAATAEKSTVPEEGVGPGA*
**>Dioro_LOC120254505**
MKPFSDEIDLPKRHRRRDPHLNDAALSCMLNTELGAVLAVIRRPSDLLPSPDDSSSHSSPLLHSLKSLRSLIFHS
DWRSLDPSAYLSPFLAVVQSDDVPAAATAVALSAILKILRLDVFDERTPGARDAIHSVVFSLTNCRLERTDAASE
DSVLMRILQLLVTMMKARAAMLLTDHAVCTVVNTCFQVVQQSANRGDVLQRTARHAMHDLIQSIFSRLPFIKTPN
FAPDPDRACSGSDDEDVDDLIISVEYGSRCMVDIFHFLCSMLNVGEVVESADGLGSLSSDEDVQVFALMLINSAV
ELGGEGMGRHTRLLRMIQDDLFHHLIHYATRSSPLVLSMICSIVLNLYHFLRSSLRLQLEAFFTYVLLKIAAGGN
GVHLQEVAIESLVSFCQQPTFVVEMYVNYDCDPIQQNIFEEVGKLLCKTAFPLTSPMSGMQVQAFEGLTAIVRTI
ADGTDMDMHDPPDKNDYNIDITEFKPFWLERHDSQDSPETWVDFVRMRKLKKKKIMIAANHYNRDEKKGLDFLKI
CHLVESPPDPESFAYFFRYTPGLDKTKIGDYLGDPDEFNLQVLKDFTDTFEFSGVILDTALRTYLDAFRLPGESQ
KIQRILEAFSERYYDQQSSEIFVSKDAVFILCYSLIMLNTDQHNPQVKKKMTEEEFIRNNRAINGGTDLPREYLS
ELFHSIVSNAITFFGQTGPTVVEMNSNRWADLIKRSKKVEPFILCDFKHKLSREVFVSISGPSVATLSAIFEQTD
DEETLHECVEALINIAGIARYGLEDILDELLSCFCKFTTLLNPYATAEETLYVFSNEVKPRMATLAVFTIANKFG
DSIRGAWKNMVDCLLKLKRLRLVPQSVIENDEIPSSETELSHLRSESGVIFPSSHQALSSSKQVSGLIGKFSQFL
SLDTESDSMLNVGSQFENNLKIIQQCRIGSIFRESSKLPDDSLQNLGRSLIFAAGGKGQKFSTSVEEEETVGFSW
ELLATITIANIQRLPNFWPQFHDCLTVVSQFPLFSPCPFAEKAIVSLFKIALKHFSSNPNSNQFSPEKQQIEDLI
FKSINLTWKLDKEILDTCCETITESVTKILIDFPDNVHTTMGWKTLFHLLSVAGRHPETFDQAIEAITKLMTEGT
HITRHNYSYCIEAAFGFAALKISPLEISSKMLDLMSDSVKLLIQWHKSGYSDPGSASISSSSSSSSLTDELPKS
GNIAANLFVKLAEALRKTSLVRREEIRNQAVTLLQRCFDAAADELEFTPANCIACFNFVIFAMVDDLHEKMVEYS
RRDNSEKEMKSMEGTLREAMELLVGVFLKLLVPLAQSPGFRTFWLGVLRRMDTCMKAEGGGEEMEGERGPLQELV
PELLRRMIVEMKEKEILVQKEGDELWDVTNIQIQWIAPALKDELFPEDL*
**>Dioal_08G121600**
MGRSRLQNSGINPIEEESPDSPDASASSGPSRASLACVINAEVAAVLAVMRRNVRWGGAGSGAARYLPSAADDHL
DHPLVQSLKSLRGQLFSRDPAHFLDPAAYLRPFLDVVRSDETGAPITGVALSAVYKILTLDGALRPGDGDAIRGV
VDAVTSCRFEVTDPASEEAVLMKILQVLLAAMRSPVAISLSNQHVCTVVNTCFRVVHQAGTKGELLQRVSRHTMH
ELVRCIFSLLPYVAESTDQPALLNGEMGAVDKDQSFGVNHAENGNGNVGSLGNSGDDGVLGVETPNGMRVMMEPY
GIPCMVEIFHFLCSLLNIVEHMEMSPRMNPIALDEDVPLFALGLINSAVELGGPSIRKHPKLLSLIQDELFRNLM
QFGLSMSPLILSMVCSIVLNLYHHLRTELKLQLEAFFSCVILRLAQSRYGASYHQQEVAMEALVDFCRQKTFMPE
MYANLDCDITCSNVFEDLANLLSKSAFPVNCPLSSMHVLALDGLIAVIQGMADRIGTAPPRTEQAPLELAEYTPF
WTVKCEDYSDPEEWVKFVRRRKYIKRRLMIGADHFNRDPKKGLEFLQGTHLLPEKLDPQSVACFFRYTAGLDKNL
VGDFLGNHDEFCVQVLHEFAGTFDFEDMNLDTALRLFLETFRLPGESQKIQRVLEAFSERYYEQSPQILANKDAA
LLLSYSLIMLNTDQHNVQVKKKMTEEDFIRNNRLINDGSDLPREFLSELYHSICRNEIRTIPDQGIGFPEMSPSR
WIDLMRKSKKTSKYIICDSRPFLDHDMFAIMSGPTIAAISVVFDYAEQEEVFLTCVDGFLAVAKISAYHHLEDVL
DDLVVSLCKFTTLLSTSLVEEPVTAFGDDMKARLATETVFNIANRYGDFIRTGWRNVLDCILRLHKLGLLPARVA
SDAADDSEVASDPVHGKPVPNSLSTSHIPAMGTPRRSSGLMGRFSQLLSLDTEEPRSQPTEQQLAAHQRTLQTIQ
KCHIDSIFTESKFLQADSLLQLARALIWAAGRPQKVTNSPDDEDTAVFCLELLIAITLNNRDRIVLLWKGVYEHI
ANIVQSTVMPCALVEKAVFGLLRICQRLLPYKENLADELLRSLQLVLKLDARVADAYCENITQEVTRLVKANATH
IKSQMGWRTITSLLSITARHPEASEVGFEALVFIMSEGAHLSPANYILCVEASRQFAESRVGLTDRSVRALDLMT
ESVTSLARWSRETRDAGEEAEKISEGIREMWLRLVQALRKVCLDQREEVRNHALSSLQRCLVGLEGLCLSTMSWL
QAFDLVIFPLMDDLLETAQNHSQKDYKNMEGTLLLSMKLLSKVFLQWLQELSALSSFCKLWLGVLSRMEKFMKAK
VRGKRSEKLQELIPELLKNTLLVMKTKGVLAKRSTIGGDSLWELTWLHVNNIAPSLQSEVFPGPGIRAGVTC*
**>Dioal_04G114500**
MLNTELGAVLAVIRRPSDLLPSPDDSSSHSSPLLHSLKSLRSLIFHSDWRSLDPSAYLSPFLAVVQSDDVPAAAT
AVALSAILKILRLDVFDERTPGARDAIHSIIFSLTNCRLERTDTASEDSVLMRILQLLVTMMKARAAVLLTDHAV
CTVVNTCFQVVQQSANRGDVLQRTARHAMHDLIQSIFSSLPSIKTPNFAPDPDPDRACSGSDDDDVDDPIISVEY
GSRCMVDIFHFLCSLLNVGEVVESADGLGSLSSDEDVQVFALMLINSAVELGGECMGRHPRLLRMIQDDLFHHLI
HYATRSSPLVLSMICSIVLNLYHFLRSSLRLQLEAFFTYVLLKIAAGGSGVHLQEVAIESLVSFCQQPTFVVEMY
VNYDCDPIQQNIFEEVGKLLCKTAFPLASPMSGMQVQAFEGLTAIVRTIADGTDIDMDDPPDKNDYNIDITEFKP
FWLERHDSQDSPETWVDFVRMRKLKKKKVMIAANHYNRDEKKGLDFLKICHLVESPPDPESFAYFFRYTPGLDKT
KIGDYLGDPDEFNLQVLKDFTDTFEFSGVILDTALRTYLDAFRLPGESQKIQRILEAFSERYYDQQSSEIFVSKD
AVFILCYSLIMLNTDQHNPQVKKKMTEEEFIRNNRAINGGTDLPREYLSELFHSIVSHAITFFGQTGPTVVEMNS
NRWADLIKRSRKVEPFILCDFKHKLSREVFVSISGPSVATLSAIFEQTDDEETLHECVEALINIAGIARYGLEDI
LDELLSCFCKFTTLLNPYATAEETLYVFSNEVKPRMATLAVFTIANKFGDSIRGAWKNMVDCLLKLKRLRLLPQS
VIENDEIPSSETELSYLRSESGVIFPSSHQALSSSKQVSGLIGKFSQFLSLDTESDSMLNVGSQFENNLKIIQQC
RIGSIFRESYKLPDDSLQNLGRSLIFAAGGKGQKFSTSVEEEETVGFSWELLATITIANIQRLPNFWPQFHDCLT
VVSQFPLFSPCPFAEKAIVSLFKIALKHFSSNPNSNQFSPEKQQTEDLIFKSINLTWKLDKEILDTCCETITESV
TKILIDFPDNVHTTMGWKTLFHLLSVAGRHPETFDQAIEAITKLMSEGTHITRHNYSYCIDAAFGFAALKISPLE
MSSKMLDLMSDSVKLLIQWHKSGYSDPGNASNASISSSSSSSTDEQPKSGNIAANLFVKLAEALRKTSLVRREEI
RNQAVTQLRRCFDAAADELEFTPANCIACFNFVIFAMVDDLHEKMVEYSRRENSEKEMKSMEGTLREAMELLVGV

FLKLLVPLAQSPGFRTFWLGVLRRMDTCMKAVATATAEGGGEEMEGERGPLQELVPELLRRMIVEMKEKEILVQK
EGDELWDVTNIQIQWIAPALKDELFPEDL*
**>SpipoGNOM_chr1**
MGRLRLQSGIKAIEEEEVDDCEGTSSRVALACMINSEVGAVLSVMRRNVRWGGRYVGGDEHLEHSLIQSLKTLRR
QIFSLEHKWHAVNPTLYLKPFLDVIRSDETGAPITGVALSSVYKILTLDILDLNTVNVDNAMYLIVDAVTSCRFE
VTDPASEEVVLMKILQVLLACMKSKASVLLNNQHVCTIVNTCFRVVHQAGTKGELLQRFSHHTMHELVKCIFSHL
KDVEESQLSNGGSPLLTPEVGGLDKGSDFHAKQVENGDGSVGVEDTRLPVGSPVANLVDQNPVAESTNGAQVMSE
PYGIPCMVEIFHFLCSLLNITEQIGVGPNSNPIAFEEDVPLFALRLINSAIELGGSSIQKHPKLLALIQDELFRN
LMQFGLSMSPLIVSMVCSIVLNLYHHLRTELKLQFEAFFSCVILRLAKSRHGSLYQQQEVAMEALVDFCRQNTFM
PEMYANLDCDITCSNVFEDLANLLSKSAFPVNCPLSAMHILALDGLIAIIQGMANRITDACPVPEHAPVVFEEYS
PFWTVKCKNYSDPDIWVKFVRHQKYIKRRLMIGADHFNRDPKKGLEFLQGTHLLPEKLDPQSVACFFRYTAGLDK
NLVGDFLGNHDDFCVQVLHEFAWTFDFHEMNLDTALRLFLETFRLPGESQKIQRVLEAFSERYYEQSPQILANKD
AALLLSYSLIMLNTDQHNVQVKKKMTEEDFIRNNRLINDGSDLPREFLSELYHSICKNEIKTIPDYSVGFPEMTP
SRWIDVMRKSKNTSQYIVYDSRPLLDRDMFAIMSGPTIAAISVVFDHAEQEEVFYTCIDGFMAVAKISACHRLDD
VLDDLVVSLCKFTTLLNASSVEEPVIAFGDDIKARIATETVFSIANRYGDYIHTGWRNILDCILRLHKLGLLPAR
VASDAADDSEYAPDSAHGKLPPSSQNASHLPSMGTPRRSSGLMGRFSQLLSLDTEEPRSQPTEQQLAAHQRTLQT
IQKCHIDTIFTESKFLQAESLLQLAKALTWAAGRPQKVNSSPDDEDTAVFCLELLIAITLNNRDRIVLLWQGVYD
HIANIVQSTVIPCTLVEKAVFGLLRICQRLLPYKENLADELLRSLQLILKLDARVADAYCENITLEVTRLVKANA
SHIKSQMGWRTITSLLSITARHPEASEVGFEALVFIMSEGAHLSPANYILCLEASRQFVESRVGLTDRSVHALDL
MAESMTSLVRWSHEAKRAGEGSEKISEGIRDMWLRLMQALRKICVDQREEIRNHAVTLLQRCLAGSEGIPLSASS
CLQLFDLIIFVVLDDLLEIAQGHLPKDYRNMEGTLVLSVKLLSKVFLQLLQELSPLSSFCKIWLGVLSRMEKYMK
LKVRGKRYEKLQELIPELLKNTLLVMKAKGILVRRSTIGGDSLWELTWLHVNNISPSLQSKVFPGQELQQDSESP
PMGSSLSESAPLPSGDISQAVG*
**>SpipoGNL2_chr8**
MARPRGGDGDGDDSGGGGGRGGDYGEPERRKPVGGRRSNGGRSRRKELGVSCMLNTEVGAVLAVIRRSSSSDGGS
FLPSPDDGSSAGGGGDSHLLHSLKSLRSLIFQPDAGEWRSVEPSVYLGPFIEVVQSGDVPAAATGVALSAILKIL
KLDIFDEKTPGAGEAVHSLVLAATNCRLERTDPASEDAVLMRILQVLTAVLRHPASLLLSDHAVCTVVNTCFQVV
QQSACRGHLLQRNARYAMHELVQTIFSRLPEIQAAEHSDSDTDGDGAGEAGYGARCMVDIFQFLCSLMNVVEVAD
SGEGVGPSCSDEEIQLFALILISSAVELGGESIGHHPRLQRIIQDDLFHHLIHYGTRASPLVLSMICSTVLNLYH
FLRGMAAGSAGAHLQEVAVEGILGFCRQPSFVIEMYVNYDCDPIRRDILEETVRLLCRAAFPTSAPLSPMQIQAF
EGLTTVVDALAENMDLDRSPDGETYQMAISEFRPFWLERSDSHDDDPEAWVEFVRMRKLKKKRIMIAGSHFNRDE
KKGLEFLKISRLAPDYPDAKSIAFFFRYTPGLDKNKMGDFLGDPDEFNIQVLREFTGTFDFSGAILDTALRTFLE
TFRLPGESQKIQRILEAFSERFYEQQSADIFAGMDAVFILCYSLIMLNTDQHNPQVKKKMTEEEFIRNNRAINGG
KDLPREYLSDLFQSISNNAITLFGQSGSAAEMTSSRWFDLVKRSKLDEPFILCDYEHRLCRDAFAVIAGPSVATL
SAIFEQTDDEEILHECIQGFFSVARISQYGLQDIIDELIACFCKFTALLNPYATAEETLFTFSNDVRSRMATLAV
FTIANKFGVSVRGGWRNVLDCLLKLKKLKLLPQLVVESDKSTDSASDFSSHVKAESGLLFSSSSTSRFGGDGGGG
GRRQTSGLMGKFSQFLSLDSGADSLANICGEFENNLKLIQQCQIGSIFAESSKLPDESLQNLGRALIFAAAGKGQ
KFSTPVEEEETVGLCWELLVAITLANIHRFLAFWPQFHDYFTVVAQFPLFSPCPFSEKAILGLFTICLKLFSSPS
AHDKPSEELIFKSVNLMWKLDKEIFDTCCEGIAKAAARLLTEHRGAIQTPLGWKTVLQLLAITGRHQETYDRSVE
ALISLMADGKDITRFNYGYFMEAALGFAASKTSPLEDSLRIIDLMGGSVEWLIQWHKSGHAESGGLLSNASYSSV
EDGSKVVGSSGNVEVSLFVKLAEALRKTSMVRREEIRNHAVATLRRCFAAAAAELEFPPASCIGCFNHIIFAMVD
DLHEKMVEYSRREGSEREMRSMEGTLKGAMELLVDVYLRFLTSLHQSPSFRTFWLGLLRRMDTCMKGDAGVLHDL
VPKLLKKVIIEMKEKEILVQRDGDELWEITNIQIQWIAPDVKDELFPEYF*
**>Zosma_108g00240**
MGCARLQSDIKTIKEEVQEGCDDTETGPVLSCMINSEVGTLLAVMRRNVRWGSRYLAGDNQLDHYLIHSLKILRK
QMFSWGKYWQTIDPVIYLQPFLDVIRSNEIGAPITGVALSSVNKILNLGILDLNTLNIDNAMHLIVDSVTLCRFE
VTDAASEEVILMRILQVLLGCLKCKASSVLTNQHICTVVNTCFRVVHQAENKGELLQRISRHTMHELVRCIFSHL
QNVEVQLSPNDAKPSVKSVDGVLDKDHAFRIKPEETSIIGNFAQDDMVSSVCIDKNTVGPTDMNGIHSITGPYGF
PCAVEIFQFLCSLLNVSEHIEMGSTINPMAFEEDIPLFALGLINSAIELGGPSIQRHPKLLSLVQDELFQNLMQF
GLSMSPLVISMVCSIVLNLYHHLHAELKLQLEAFFSCVILRLTRNRHPASYQQYEVAMEALVDFCRQKNFMAEMY
ANLDCDITCSNVFESLANLLSKSAFPINCPLSPMHLLAFDGLLAVIQGMADRIVDNSPLSEQSSVELEEYTPFWT
LKCENYSDPDHWVRYVRRRKYIKRRLMIGADHFNRDPKKGLEFLQGTHLLPEKLDPQSVACFFRYTSGLDKNLVG
DFLGNHDEFCVQVLHEFARTFDFQNMRLDTALRLFLETFRLPGESQKIHRVLEAFSERYYEQSPHILANKDAALV
LSYSLIMLNTDQHNVQVKKKMSEEDFVRNNRSINGGNDLPREYLSELYHAIYKNELRTIPEQGLSFPEMTPSRWI
DLMQKSKNSSPYISYNSHPFLDRDMFAIMSGPTIAAISVIFDNSEQEEVFHACVDGFLSLAKISACHHLEDVLDD
LLASLCKFTTPTNTSSVEEPVTAFGDDLKARLATETIFNIATRYGDFIRTGWRNLLDCILMLHKLGILPARVASD
AADYTETSVDTTIQEKPILNSQSLSQISTEAPYRSSGLMGRFSQLLSLDTEEPRSQPTEQQLAAHQRTVQTIQKC
SIDSIFTESKFLHTDSLLHLARALILAAGRPQKNISSHDDEDTAVFCLELLIAVTLNNRDRIVLLWQFVYDHIAS
IVQSTVMPCPLVEKAVFGLLKICQRLLPYREELVDDLLKSLQLILKLDARVADAYCENITQEVTRLVKTNGTHIK
SQIGWKTITDLLSITARHPEASEVGFEALVFIMSEGTHLSPANYILCVDAARQFAESRVGLTDRSVQALDMMAES
VACIVRWSLNKVEKSQNGIAEMWLKVIQGLKRVCLDQREDVRNHALLSLQRCTLTVDKNCLSPSLWVQSFDLVIF
LLLNDLLEIAQAQSPKEYKNMEASLVLAIKLLSKLFLHLLKDVSSQSSFPKIWLGVLNLMEKYTKVKLRGKRSEK
LQELIPELLKNILLMMKSNEILVKRNPIVGGDNSLWDLTWLHVIKITPSLQSEIFPGEPHLQA*

**>Zosma_56g00900**
MGYVKLHSGAKTDEETEVVHRGCDDMVRGTALSCIINSEVSAVLSVMRRNIRWAGRYMAGDEQLEHSLIQSLKTL
KKQMFSWGDQWHTIDPVIYLQPFLDVIQSDETSAPIVSIALSSVYKILTLEIFNLNTVNVDKAMHLIVDSVTSCR
FDVTDASEEAEILMILQVLLSCMKNKASVLLSNQHVSIIVKTCFRVVCQADNKGELLQKISRNTIHELIRCIFLH
LQNVEAQLLPNDASSVKSEDDLPNNDISFGIKQINNGNGSFVYDAPETCCSIDKNRIGATKVIGDQLMTNTYGVP
CVMEIFNILFSLLNISENLDMGPRSNTKAFEEDMPLFALGLINSAIESGGSSLKNHPKLLALVQDELFRNLMQFG
LSTSPSVLSMVCSIALNLYHHLRSELKLQLEAFFSCIILRHSKNRHPASFQQQEVAMEALVDFCMQKDFMAEMYA
NLDCDITCSNVFECLSNLLSKGAFPVNCPLSSMHLLALDGLLAVIQGMAGRINNASLISEQTSVELREYTPFWTL
KCENYSDPNQWVGYVRRRKYIKRRLMIGADHFNRDPKKGLEFLQGTHLLPDKLDPQSVACFFRYTSGLDKNLVGD
FLGNHDEFCVQVLHEFAGTFDFQDMSLDTSLRLFLETFRLPGESQKIQRVLEAFSERYYDQSPHILANMDAALVL
SYSLIMLNTDQHNVQVKKKMTEEDFIRNNRSINDGNDLPRDYLSELYHAICKNELKTIPEQGAGFPEMTPSRWID
LMQKSKSTSPYISYGSCPYLDRDMFAMMSGPTIAAISVVFEHAEQEEVFRSCVSGFLALAKISACHHLEDVLDDL
VVSLCRFTTLTNSISIEEPVSAFGDDLKARLATETVFSIVNQYGDFIHTGWRNILDCILRLHKLGLLPARVASDA
SDDTEPSLDAVQGKPVLNSQSLSLTPTGTPRMSSGLMGRFSQLLSLETEEPRSQPSEQQLASHQRTVHTIQKCHI
DSIFTESKLLKPDSLRHLARALVWPYSRSQKNNSSPDDEDTAVFCLELLIAITLNNRDRISLLWQGVYDHTASIV
QSTFMPCPLVEKAVFGLFKICQRLIPYNENLADELLKSLQLILKLDTQVADAYRENITQEVTRLVKANATRIKSH
MGWKTITDLLSITARYLESSEAGFEVVTFIMTDGAFLTPANYILFVDSARQFAESRVSLTDRSIRALGLMADSTA
CLVRWSLDTNPGIEGIDKTYDPIKEMWSHLVKGLRKMCLDPRDEVRNQGLSSLIKCMIGMDGICITSQWQQSFDT
VVFGLLDDLLKITQQGQSPKEYKNIETTLVLAMKFLSKLFLHFLKDLSALSGFCKLWLGVLDRMDKYMNVKFRGK
SEKLHELIPDLLQNILVVMKARGILVQRSAICGDSLWELTWLHVNNIAPSLKSEVFPGDEYAQDL*
**>Zosma_123g00350**
MGRMKFPPPIEEAVEDHGGGWNTRATLPCMINSEIGTVLAVMRRNVRWGGHYMGDENQLEHPLVQSLKTLRNQMF
AWDQQWHTIDPVNYLQPFLDVIQSDETGAPITGVALSSVYKIVTLDILNLNTVNIDKAMPLIADSVTSCRFEVTD
PASEEVVLMKILQVLLACMKSKASVVLSNQHVCNIVSTCFRVVNQAGSKGELLQRISRHTMHELIRCIISRLHNV
ENQSSLTEDGLFLKSELGRLDRDPAFGIMQVENGNGSLDTPGEASSVDENGMVSKHDSVVQLMSGPFGLPCIVEI
FHFLCSLLNISEHIEMGPRSNPVTYEEDMPLFALGLINSAIELAGPSIRIHLQLLSLVKDELFYNLMQLGLSTSP
LIISMVCNIVLNLYHHLHSELKLQIEAFFSCVILRLAKNRHGASYQQQEVAMEALVDFCRQKNFMAEMYANLDCE
ITCDNTFEELVNLLSQSAFPVNSPLSAMHILALDGLIAIVKGMVSRINKSPQILDEVPFKLERYTPFWTAKCDNY
SDPNHWVQYIRHRKYIKKKLMAGVDHFNREPNKGLEFLQGIHLLPEKLDPHSVACFFHYTSGLDKNYVGEFLGNH
DEFCIRVLHEFAKIFDFQDMNLDTALRLFLDTFRLPGESQKIQRVLEAFSERYYEQSPHILANKDAALLLSYSLI
MLNTDQHNVQVKKKMTKEDFIRNNRHINGGNDLPREYLSELYHAICKNELKTIPEQGAGFPEMTPSHWIDIMQKS
KKTSPYIFCNSLPFLDQDMFAIMCGPTIAAISVVFDYAEHDEVFHSCIEGFLVLAKLSASYHLGDVLDDLVTSLC
RFTTLTNSSSVDEPAMEFGDDLKARLATEAVFNLANQYGDFIRIGWRGILDCILRLNKLGFLPARVASDATSDEY
DTSADTEHGKLNSQITSQAPIIVNLRKPSGLMGRFSQLLSLDTEEPRSQPTEQQIAAHNCTIQTIQKCHIDSIFT
ESKFLQADSLLNLVHALISAAAQIQKDNGLLDDEDIAVFCLELLIAITLNNRDRIVILWQSVYDHIASIVQSTSI
PSPLVEKAVFGLLKICQRLLPYKENLADELMRSLQLILKLDARVADAYCENITHEVTRLVKTNATHIKSPVGWRT
ITSLLSITARHPEASECGFEALLFIMSDGAHLSPANYILCIEAARQFAQSRVGLAVRSVRALDLMAESVRCLVRW
SLDTKDAKEGAEKISQGLQEMWLRLVHGLKKVCLDQRDEVRNHALSLLQRCMVGVEDIRLSPESWLQCFDTVIFT
LLDELLDIVQGQSRNEYRNMETSLVLAMKLLSKVFLQSLEDLAPLSSFIELWASFLSQMEKYIKVKIRSRRSEKL
QELIPELLKNTLLVMKAGRILNNKAPISAADNLWELTWVHVNDIVPSLKTEVLSIEELAQDAAAIVVDKDAKEAK
KRTESGHALSLDEGIQAA*
**>Zosma_111g00150**
MVRPKGIGGDCYDDEEGDRRGFGGNSSVQLKEVGVSCMLSTEVGALLAVIRRPVDPNATDVTVADTHVIVALKNL
RSLIFQPHGEWHTIDPVLYISPFINVIQSDEIPAPATALAISAVLKIIRLNIFDEKTPGAGDAIQAVVHGITNCR
LERTDHVTEDAVLMRILQVLLAVMRSRVSAFLTDHAVTTVVNTCFQVVQHSACRGDMLQKNARHTMHELVQIIFI
KLPEIKPNGSGKGSEIEFDEEGGLIGPPVYGTQCMVDIFKFLCSLLNVVVECSDGLGPMSWDEDVQLFALILINS
AIELGGEAIGNHPRLLWTIQDELFYYLIYYGTRSTPLFLSMICSTVLNLYHFLRRSVDVVMLFLLCVRSLRLQLE
AFFRLVLLRVTSSGNSSMAIQEVAVEGIISFCRQPMFVIEMYVNYDSDPICSNVFEEIGKLLCKIAFPTPRPFSP
MQVQAFEGLVSVIGTIAEHKNLDEKPEGESYQVQISDFQPFWLEKCSDITDIRLWTEFVRMRKTMKKRVMIAANH
WNRDEKKGLEFMRVVHLVPDPPDPLDIANFLRYTPGLDKNKIGDYMGDPDEFNLKVLKEFSQTFDFSGVILDTAL
RTFLETFRLPGESQKIQRILETFSERFYEQQTSEIFANKDAVFILCYSLIMLNTDQHNAQVKKKMTEEEFIKNNR
AINNGKDLPRDYLSELFCSISNNAITVFGQNAAAGGVTEMNPNRWVELLKRSRSIETFITPLEFNHKLCRDLFVP
ISGPSVACLSAIFDRADDDEILQDSMKGLLHIARIAQYGLEDILDELLTCFCKCTTLLNPYLTPEETLFSFSHDL
KPRLATLAVFTIANQYGGSVRGEWRNTLDCLLKLNKMRVLPSNIIESSGEKRPPSPSTSSTANLMSQFSQFLTMD
RNLDSMASIGNEFESNLKLIEQCRIGNIFTGSWRLPEESLQNLGRALIFAAAGKNQKFSTPIEEEETVGFCWDLL
MTISAVNINRSSSIWPQFFENFTIVAQFPLFSPCPFAEKGIVGIFSIYVKYLAAGITVGDKERDKVTEEYILKSI
NMMWKFDKVLDTCCDSITDSIVTIFNESGTSIQSILGWKTLLHLLSIAGRHPETYDRAVQALTMVMTVKSATHIT
RYNYAYFIDAAFGFAALRISPLENSLKLLDLMAGSIDCLIQYHTSHTSDPDNATGDPASAMYLSGNLFLKLAEAL
RKTSLVRREEIRNHAVSAIRRIFDRAVDLDFTPTNCIGCFNLVIFAMVDDLHEKTVEYSRRDNSEREMRSMEITL
KAAMEMMVDVYLKFLMVLQLSPAFRTFWLGLLRRMDTCMKANIDGEAGLLQSLVPVLLKKMIREMKAKEVLVERE
GDELYEITSIQIQWIAPSIKDELLM*
**>Taro_018380**

MGRLRLQSGIKVIEEEVVDDCEGASNRAALACMINSEVGAVLAVMRRNVRWSGRYLGGDEHVEHSLIQSLKALRR
QIFSWERQWHAVNPTLYLEPFLDVIRSDETGAPITGVALSSVYKILTIDILDLNTVNVENAMHLIVEAVTSCRFE
VIDPASEEVVLMKILQVLLACIKSKASALLNNQHVCTIVNTCFRVVHQAGTKGELLQRFSRHTMHELVRCIFSHL
PDIEEKPLATGGGALKPEIGGLDKDHALHAKKMENGGGNIGDDDPQHVDSPVATLVDQNTVADVANGSQVMSEPY
GVPCMVEIFHFLCSLLNIAEHIGISPNANPIAFDEDVPLFALKLINSAIELGGPSIQKHPKLFALIQDELFRNLM
QFGLSMSPLIVSMVCSIVLNLYHHLRMELKLQLEAFFSCVILRLAKIKHGFSYQQQEVAMEALVDFCRQQTFMPE
MYANLDCDITCGNVFEDLANLLSKSAFPVNCPLSAMHILALDGLIAVIQGMADRITDASPMSEKTPVEFEEYTPF
WTVKCNNYSDPDSWVKFVRRQKYIKRKLMIGADHFNRDPKKGLEFLQGTHLLPEKLDPQSVACFFRYTSGLDKNL
VGDFLGNHDDFCVQVLHEFARTFDFQDMNLDTALRLFLETFRLPGESQKIQRVLEAFSERYYEQSPQILANKDAA
LLLSYSLIMLNTDQHNVQVKKKMTEEDFIKNNRHINGGNDLPREFLCELYHSICKNEIKTTPDSGAGFPEMTPSR
WIDLMCKSKKTSQYIVYDSRPFLDREMFAIMSGPTIAAISVVFDYAEQEEVFHTCIDGFMAVAKVSACHHLEDVL
DDLVVSLCKFTTLLNASSVEEPVTAFGDDMKARMATETVFSIANRYGDYIRSGWRNILDCILRLHKLGLLPARVA
SDAADDSEQIPDPAQGKLALSSQYASNIPSMGTPRRSSGLMGRFSQLLSLDTEEPRSQPTEQQLAAHQRTLQTIQ
KCHIDSIFTESKFLQADSLLQLAQALIWGAGRLQKVNSSPDDEDTAVFCLELLIAITLNNRDRIVLLWQGVYEHI
ANIVQSTVMPSALVEKAVFGLLRICQRLLPYKENLADELLRSLQLILKLDARVADAYCENITQEVTRLVKANASH
VKSQMGWRTITSLLSITARHPEASEVGFEAMVFIMSEGAHLSPANYVLCLEASRQFVESRVGLTDRSVQALDLMA
QSVTCLAQWSQEAKEMGEGSEKISEGIREMWLRLVQGLRKVSVDQREEVRNHAISLLQRCMGGAEGICLSPASWL
QSFDLVIFAMLDDLLEIAQGHSPKDYRNMEGTLTLAVKLMSKVFLQLLQELSPLSSFCKLWLGVLGCMEKYMKAK
VRGKKSDKLHELIPELVKNTLLVMKARGILVRRSTIGGDSLWELTWLHVNNINPSLQSEIFPEQPEVDPANPVPQ
VSSSAPIAETMPLASEDGIQAGG*
**>Taro_026544**
MGHLRLYSGINSIDEEVADDYQGISKRSALACMINSEVGAVLAVMRRNVRWGGRYVGGDEHFEHSLIQSLKTLRK
QVFSLEQQWHTVNSTLYLQPFLDVIRSDETGAPITGVALSSVNKILTLDILNLNTVNVENAMHLVVDAVTSCRFE
VTDPASEEVVLMKILQVLLACMKTKASVLLTNQQVCTIVNTCFRVVHQAGTKSELLQRISRHTMHELVRCIFSHL
PDIEEQPLAKKMDGLNKDPAFGAKQVESRDGSIEDDAPLPVGSPVTNLVDEKAAAEAANGMQLMTEPYGVPCMVE
IFHFLCSLLNIVEHIGMSPVTNPIAFDEDVPLFALGLINSAVELGGPSIQKHPRLLALVQDELFHNLMQFGLSMS
PLILSMVCSIVLNLYHHLRSELKLQLEAFFSCVILRLAKTKHGVSYQQQEVAMEALVDFCRQRTFMPEMYANFDC
DITCSNVFEDLANLLSRSAFPVNCPLSAMHILALDGLIAVIQGMADRLTDACPVSEQTPVEFHEYTPFWTVKCED
YSDPDTWVKFVRRRKYIKRRLMIGADHFNRDPKKGLEFLQGTHLLPEKLDPQSVACFFRYTSGLDKNLVGDFLGN
HDEFCIQVLHEFAWTFDFKDMNLDTALRLFLETFRLPGESQKIQRVLEAFSERYFEQSPQILVNKDAALLLSYSL
ILLNTDQHNVQVKKKMTEEDFIRNNRHINGGSDLPREFLSELYHSICKNEIRTIPDQVVSFPEMTPSHWIDLMCK
SVKTSQYIVCDTRPFLDHDMFAIMSGPTIAAISVFDYAEQEEVFHTCIDGFMAVAKISACHHLDDVLDDLVVSLC
KFTTLLNASSVEEPVTAFGDDMKARMATETVFNIANRYGDYIRSGWRNILDCILRLHKLGLLPARVASDAADDSD
VSTDSVHGKPVPSSQYSSHLPPMGTPRRSSGLMGRFSQLLSLDTEEPRSQPTEQQLAAHQRTLQTIQKCHVDSIF
TESKFLQADSLLQLARALVWAAGRPQKVSSSPDDEDTAVFCLELLIAITLNNRDRIVLLWQGVYEHIANIVQSTV
MPCALVEKAVFGLLRICQRLLPYKENLADELLRSLQLVLKLDARVADAYCENITQEVTRLVKANASHIKSQMGWR
TIMSLLSITARHPEASEVGFEALVFIMSEGAHLSPANYILCLEAARQFAESRVGLTDRSVHALDLMAESIACLAR
WSHEAKEAGERSEKISEGIREMWLRLVQGLRKVCVDQRDEVRNHALSSLQRCMLGVEGVCLSPHSWLQSFDLVIF
VIIDDLLEIAQGNSPKDYRNMEGTLVLAVKLLSKVFLQLLQELAPLSSFCKLWLGVLSRVEKCMKAKVRGKKSEK
LLELIPELLKNTLLVMKDKGILVRRSTIGGDSLWELTWLHVNNISPSLQSEVFPGHELEQDSGSSPAEALP*
**>Taro_009810**
MGRQRDHDDSDGDGGGGGEPADGARRDRRRSSRSCRGRPPQRQPPSGGRLRELGLSCMLNTEVGAVLAVIRRSSD
AVSGQFLGSPDDAAPPVAADSPLLQSLKSLRSLIFRQGGAGDWRSADPFVYLGPFLDVVQSDDVPASATGVALSA
VLKILKLGIFDEGTPGARDAIHSVVTAITNCRLERTDPASEDAVLMRILQVLTAVLRCPASVHLTDHAVCTVVNT
CFQVVQQSASRGDLLQRNARYTMHELIQTVFSRLPEVDEGVNGGEGYSDSDPEDDGRALSGGSRPQGGYGARCMV
DIFHFLCSLLNVVEVVDSAEGFGSMCFDEDIQLFALILINSAIELGCESVGRHPKLLRMVQDDLFHHLIHYGTRT
SPLVLSMICSTVLNLYHFLRRSLRLQLEAFFTFVLLRTAAGTFGVHLQEVAVEGLISFCRQPTFVIEMYVNYDCD
PVRRDIFEEIVRLLCKTAFPVNSPLGPMQVQAFEGLTAMIAAIVENMDFDRSPEGECYQMQISEFSPFWTERCDD
GGDNHEAWVDFVRTRKFKKKKVMMAANHFNRDEKKGLEYLRISHLVADPPDAKSLACFFRYTPDLDKNKIGDYLG
DPGEFNIQVLHEFAATFAFAGVILDTALRTFLETFRLPGESQKIQRILEAFSERFYDQQSADIFASKDAVFILCY
SLIMLNTDQHNPQVKKKMTEDEFIRNNRAINGGKDLPREYLSELFFSISNNAITLFGQSGSTVEMTTNRWVELVK
SSRTVEPLVLCESEHRLGRDAMAAVAGPSVAALAAVFEHTEEEDVLQECLEALLSVARISQYGLGDIIDQLLSCF
CKFTTFLNPYATADETLFTFTNDVRSRMATLAIFTIANKFGVTVRSGWRNVVDILLKLKRLKLLPPSLMEPDRPT
SDAGGRPQHAKSESSGVVFPLSNPRFSGSRRSSGLMGRFSQFLSLDSAADSLAAVSSEFENNMKIIQQCQIGSIF
SESSKLPDESLQNLGRSLIFAAAGKGQKFSTPVEEEETVAFCWELLVAITVANIQRFPTFWFQFHDYLTAVAQLP
LFSPCPFSEKAIMGLLFICLKLFSSAHDKASEELIFKSINLMWKLDKEILDTCCESIAESTISLLTEHKGAIQTL
LGWKTVFHLLSVTGRHPETYDRSVEALISVMADPRCITRFNYTCCIEAAFGFAALKISTPENSARIIDLMADSVQ
WLIQWHKSGNIESGGSLSSVSTASSSSTEDVLRTAAAASGNVTASLFMKLAEALRKISLVRREEIRNRAVSALRR
CFAAAEELDLPTSNCIGCFNLIIFAMVDDLHEKMLEYSRREGSERELRSMEATLKEAMELLAEVFLQFLGTLYQS
PGFRSFWLGLLRRMDTCIKADLGELGKTSLLHDLVPELLKKMIVEMKEKEILVKRDGDELWDVTNIQIQWIAPAV
KDELFPEDF*
**>Acora_06G147100**
MGRMKLQSGIKAIEEEPPDEMEFNTNRAALACMVNSEVAAVLAIMRRNVRWGGRYMAGDDQLEHPLIQSLKALRR

QIFSWEQHQWHAINPTLYLRPFLDVIRSEETTAPITVVALASVYKILSLDMFDLSTVNVDGAMRSVVETVTSCRF
EVSDPASEEAVLMKILQVLLVCMKSKVSSVLSNQHVLTIVNTCFRIVHQAGTKGELLQRISRYTMQELIQCIFSH
LPNVVEEHQPDQRLVNGGGPSGKTELCGLEKDHVLGVKQLDNGNGLDGFYAKTSDDIPVGTVVSLNGPMASSDSD
LTLDPYGIPCMVEIFHFLCSLLNAAEHIEMGPRMNQITYDEDVPLFALGLINSVIELGGSSISRHPKLLSLVQDE
LFRNLMQFGLSMSPLILSMVSNIVLNLYHHLLAQLKLQFEAFFTCVILRLAQSRYGASYQQQEVAMEALVDFCRQ
KAFMKEMYANFDCDITCSNVFEELTNLLSKSAFPVKCPLSTMHVLALEGLIAVIQGMAERVGNSPTPVYDRAAVR
IEEYSPFWMVKCENYSDPDHWIQFVRRRKYIKRRLMIGADHFNRDPKKGLEFLQGIHLLPEKLDPQSVACFFRYT
AGLDKNLVGDFLGNHDEFCVQVLHEFARTFDFQDMNLDTALRLFLETFRLPGESQKIQRVLEAFSERYYEQSTQI
LANKDAALLLSYSLIMLNTDQHNVQVKKKMTEEDFIRNNRRINDGNDLPREFLSELYHSICQNEIRTIPEQGAGF
AEMTPSRWVDLMRKSRRTTPFLTCGSGAHLDRDMFAIMSGPTIAAISVVFDNAEHEEVFRMCNEGFHAIAKISAC
HHLEDVLDDLVVSLCKFTTLLNPSAVDEPVMAFGDDSKARMATETVFTIANAYGDYIRTGWRNILDCILRLHKLG
LLPARVASDAADDADSPTPPSSNTTTSSSETTHGGRPVLATSALGSVPQHVPAVAMGTPRRSSGLMGRFSQLLSL
DAEEPRSQPTEQQLAAHQRTIQTIQKCHIDSIFTESKFLHADSLLHLAKALINAASRSQKGSMGHHSSPEDEDTA
VFCLELLIAITLNNRDRIRVLWNDVIAHISNIIKTTMMPCTLVERAVFGLLRICQRLLPYKENLAEDLVMSLQLV
LKLDARVADAYCENITVEVARLVKANADLIKSNSGWKMIISLLSATSRHLEASEPGFEALEFIITDGGGRHLTPD
TYKLALEVARQFAISPVVPSERSVRALDLIEGSAACIARWVTELGEGENEEFGNSWVQLVTGLKNFCLDQREDVR
NHAVTCLQRCAVNVHGVTLRPSSWTQIFDPVLFSLLDELLDIAMGHSPKDYRNMEGTLLQAMKLISKAFLHWLKE
MMAMGGGGGVLRKVWLGVLGRMEKYMKAKVRGKRSERLQELVPELLKNMVLVMKANGVLVKRAAAIGGDSLWELT
WLHIRNIAPSLQSEVFPDVEENEGSVEGTAPSVAVALKETAVVASGEGG*

**>Acora_03G000500**

MGASSCSSSAYYMGRMKLQSGIKAIEEEPPDEMECNTNRASLACMINAEVAAVLAIMRRNVRWGGQYMAGDDQLE
HPLIQSLKALRRQIFSWDQHQWYVINPILYLRPFLDVIRSEDTTAPITVVALASVYKILSLDMFDLSTVNVEGAM
WSVVETVTSCRFEVSDPASEEAVLMKILQVLLVCMKSKVSSVLSNQHVLTIVSTCFRIVHQAGTKGELLQRISRY
TMQELIQCIFSHLPNVVEEHQPDQCLVNGGGPFAKSELGGLEKDHALRVKQLDNGNIHDGLSAKTSVDIPVAMVV
SLNGPMASSDTDQTLEPYGIPCMVEIFHFLCSLLNAAELIEMGPRMNQIAYDEDVPLFTLGLINSVIELGGSSIS
RHPKLLSLVQDELFHNLMQFGLSMSPLILSMVCSIVLNLYHHLLAQLKLQFEAFFSCVILRLAQSRYGASYQQQE
VAMEALVDFCRQKAFIKEMYANFDCDITCSNVFEELTNLLSKSAFPVKCPLSTMHVLALEGLISVIQGMVERAGS
SSSLVYDRAAVRVEEYSPFWMVKCENYSDPEHWIQFVRWRKYIKRRLMIGADHFNHDPKKGLEFLQGIHLLPEKL
DPQSVACFFRYTAGLDKNLVGDFLGNHDEFCIQVLQEFARTFDFQDMNLDTALRLFLETFRLPGESQKIQRVLEA
FSERYYEQSPQILANKDAALLLSYSLILLNTDQHNVQVKKKMTEEDFIRNNRLINDGNDLPRDFLSELYHSICQN
EIRTIPEQGAGFAEMTPSRWVNLMRKSRQTTPFLTCGSGAHLDCEMFAIMSGPTIAAISVVFDNAEHEEVFRMCI
EGFHAISKISACHHLNDVLDDLVVSLCKFTTLLNPSSVDEPVMAFGDDVKVRMATETVFTIANGYGDYIRTGWRN
ILDCILRLHKLGLLPARVVSDVADDAESPTPPSSTTTSSYETTHSGRPVLATSALGSVPQHVPAVAMGTPRKSSG
LMGRFSQLLSLDAEEPRAQPTEQQLAAHQRTIQTIQKCHIDSIFTESKFLHADSLLHLAKALINAASRSQKGSMG
HHSSPEDEDTAVFCLELLIAITLNNRDRIRVLWNDVIAHISNIIKTTMMPCTLVERAVFGLLRICQRLLPYKENL
AEDLVMSLQLVLKLDARVADAYCENITVEVARLVKANADLIRSNSGWKIIISLLSVTSRHLEASEPGFEALEFII
TDGGGRHLTPDTYKLALEVARQFAISPVVPLERSMRALDLMEGSATCIARWVTELGEGENKEFGNSWVQLMMGLK
NFCLDQREDVHYHAVTCLQRCVVNVHGVTLRPSSWMQIFDPVLFGLLDELLDIATGHSSKDYRNMEGTLLQAIKL
LSKAFLHWLKEMMVVGGGGVLRKVWLGVLGRMEKYMKAKVRGKRSERLQELVPELLKNMVLVMKANGLLVKRAAA
IGGDSLWELTWLHMRNIAPLLQSEVFPDMEENQSSVEGIAPAVAVALGETTVAASGEVG*

**>Acora_12G089500**

MAKPLDLEDQEKSESEEEDEVADSNTHYISSKPPHINKKRKNHHPHHQPHHLALSCMLNTEIGAVLAVIRRPISD
PHHHYPHPTTPIDPTTPIDPNPTASPFDTPLVASLKSLRARIFDHRRGGLDYRSVDPRAYLSPFLDVVRSDDVPA
AATGAALSAILKLLRLDVFDARTPSAAEAVSSVVAAITNCRLERADPASEDAVLMRVLQVLVAALRCGAAPLLSD
HDVCTVVNTCFQVVQQSAARGDLLQRNARHAMHELVQAIFARLPAIKPPLDARASLSGTDEDESRAYEGYGSRCM
VDIFHFFCSLLNVAEAVDHLDGFPPSITAANEDVQLFALVLINTAVELGGDAIGRHPKLLRMIQDDVFHHLIHYG
SRCGPLVLSMICSAVLNLYNFLRGSLRLQIEAFFVHVLLKMASRGNSIQLQEVAIEGIITFCRQPTFTLEAYINY
DCDPARRDVLEETIKLLCKHAYPTIASPMTPLQVQAFQGLVNVLANIADNVDADDHELAPYKLEFPDNLPVWLDR
PDDPNSWVDFLRARKLQKRRILIAATHFNSDDKKGLEYLRISKIVPENPPDPKSLAWFFRHTSGLDKNKIGDYLG
DPDELNLKVLKEYTEMFHFEGLILDTALRTFLETFRLPGESQKIQRILEAFSDRFYEQQQMSRDVFVSKDVVFIL
CYSLIMLNTDQHNPQVKKKMTEEEFIRNNRAINDGADLAREYLSELFHSISTKAITLSGNSTAAIDMNSNQWGEL
IRRSESVAPFIALPAWSSDHRPIGRDLFASVSGPAVAALAAVFERSDDAETLHECVGALISAARVSRHGLSDVLD
ELLACLCRFTALLNPYATAEETLFAFSHDLKPRMAILAAFTVAARFGAAVRGGGWRSAVDCLLKLKRLKLLPQSV
VEPSAPPPTPPAPPHKKSESASGVIFPPGAAARGGARHASSGGLIGRFSQFLSFESGAESMVLAGGELEHNLKII
QQCKIGSIFSESASLPAESALALMRALIFAAAGKGQKFNTPIEEEETVGFCWELIVAMASANLHRFVELWPYFIE
HLSAVGEQVPLLSTTPFGEKAVVALLEICVRLFQACAPLAPDRDAEELMFRSVALAWKLDKEAVDSSCEAVFECA
RRILRAHGANIQTQNGWKTLLHLITVAGRHQETFDGACEAVVGLLSDSGGANVTGLNYAFVLESAFGLAALRLAP
AERSVEILDALAGSVDRLIEWHRSGNSDPGHNWSVVSMEDGNGAMKGGGSAGAFLRLADVLRKTSLVRREEIRNH
AIVALRRCIRLAAEELELPAQSSVSCFNLVQAMVDDLHSKTVDYARRGEGAWRETRSMESTLRMAMEVMVDVYLL
FLKPISEAPDFRTFWLGLLRRMDTCMKTHLGEEEEEEEETEEEMVMMREMVTVLLKRMIVEMKEKGVLVQREGDD
LWDITNIQIQWISPSIKDELFPEEDF*

**>Nymco_LOC116253329**

MGRPKMQTEINAIAEEPEDNECSSKRAALACMVNSEVGAVLAVMRRNVRWGGRYMTGDDQSEHALVYSLKALRKQ
IFSWQQQWQTINPTTYLKPFLDVIRSDETGAPITGVALSSVYKILTLEIFDLSTVNVEEAMHMVVDAVTSCRFEV
TDPASEEVVLMKILQVLLACMKSKASVMLHNQHVCTIVNTCFRVVHQAGTKGELLQRIARHTMHELVRCIFSHLP
DIGHLESTLASKEGSRHNPDALVDGHEKDYSFGNKSRDNGNGNLEPDVPLVSMSFPSNASVVPASSLVDDNSMEI
ANNLESPNDMHLMNEPYGIPCMVEIFHFMCSLLNIGEHVGVGPQSNPIAFDEDVPLFALGLINSAIELGGLSIRQ
HPKLLALVQDELFRNLMQFGLSMSPLILSMVCSIVLNLYLHLREELKLQLEAFFSCVVLRLTQSRYGATFQQQEV
IMEALVDFCRQPNFMAEMYANLDCDITCSNVFEDLANLLSKSAFPVNCPLSAMHVLALEGLIAVIQGMAERISSA
SPALGHGPVSDLEEYTAFWTVKCESYNDPNYWVKFVRRRKYIKRKLMIGANHFNRDPKKGLEFLQGTHLLPDKLD
PQSVACFFRYTAGLDKSLVGDFLGNHDDFCVQVLHEFARTFDFQEMNLDTALRLFLETFRLPGESQKIQRVLEAF
SERYYEQSPYVLADKDAALVLSYSIIMLNTDQHNVQVKKKMTEEDFIRNNRTINGGRDLPREFLSDLYHSICKNE
IRTTPEQGAGFPEMTPSRWIDLMRKSKTTAPYITCDSRAFLDHDMFAIMSGPTIAAISVVFEHAQHEEVFHMCVE
GFLAVAKISACHHLEDVLDDLVVSLCKSTTLLNPLSSEEPVIAFGDDTKARMATVTVFTIANRFGDYIRSGWRNI
LDCILRLHKLGLLPARVASDAADDSEVSTDSSHGKPAPGAVSVSHMTPMGTPRRSAGLMGRFSQLLSLDTEEPRS
QPTEQQLAAHQRTLQTIQTCHIDSIFTESKFLQADSLLQLARALIWAAGRPQKGNPSPEDEDTAVFCLELLIAIT
LNNRDRIVLLWQGVYEHIASIVQSTVMPCALVEKAVFGLLRICQRLLPYKENLADELLRSLQLILKLDARVADAY
CEHITQEVMRLVKANAAHIRSQPGWRTICSLLSITARHPEASEAGFEALTFIMVEGAHLSPTNYVLCLDASRQFA
ESRLGLTDRSLRALDLMAESLPCLAKWHSVAKEGGDDSSKVSQDIVEMWLKLVQGLRRVCLEQREEVRNHALSLL
QRCLTTADGIGVTPTAWLHCFDHIVFTLIDDLLEIAQGHSPKDYKNMEGSMVLALKLLSKVFLQLLVDLSSLPSF
CKMWLGILTRMDKYGKAKIRGKRCDKLQEQVPEILKNILLVMKNRGVLVQRSALGGDSLWELTWMHVNAFAPFLL
AETFPDQEPHSVDNKTTVVHSSSACQHSDLVAEGVLASGSSNKNGGG*

**>Nymco_LOC116255565**

MVMDGAEEEVKGEGKPMDCSKHQKRMKDRDLGVSCMVNTEVGALLTVMRMNPVDMPTPQFGIHGTEEAVDTSFIN
SLKSLRTFIFNLQQDWHSINPSIYLSPFLDVIQYEEISAAAAGISLSAILKILRLDIFNTGTPRAREAMHSVVEA
IANCRLERANMPADDAILMRILQLLVTCLKCKGSPLLSDNAVCTIVNTCFQIVQQSASRGHVLQRCARHTMHELV
QMIFARLPDLDSSVEILLDNDHISTMNYGIGAIIDIFQFICSLLNLSVEVPFDFDSTVSGPDEDIQLFALILINS
AIELGGEAIARHQKLLGIIQDDLFFHLIHYGSNSSPLVLSMICSTALNLYHHLRSFLKLQLEAFFWFVCLKIAGG
KLYTAQQEEVAIEGLVSFCRQPKFGVEMYMNFDCDPMRENIFEEIGKLLCKKASPGTGPLTSIQSQSFDGMVVMI
QNIADGIDGIDRTNSNLKSPEFYDAPSSSSDCIETTKYRSFWTEKCEDYGDSETVIEFVRMRKCMKKRLMVSSDH
FNRDEKKGLEFLKHSQLVPDPPDAHSIAYFLRYTQGVDKTLVGDYLGDPDEFHIQVLKEFTSTFNFTNMTLDIAL
RNFLETFRLPGEAQKIQRVLEAFAERFYVQSDETFADKDAVMVLCYSLIMLNTDQHNVQVKKKMTEEEFIRNNRN
INGGEDLPREYLSELFQSIANNEITLADQSAASEDMNPSRWIYVINQSNNTDPYISCDTRPWLNRDMFSTLVGPS
IATIAAIFEHTNDDNILKQCMEGFIAVARVARYKFEDVLDELLAMLCKFTTLLNPFSSLEETLFSFAEDVKPKIA
TLTVFTIANKFGDSIRSGWRNIVDCLLRLKKLKLLPPSLSGAEVEAPADRLDLDVPLHSRAESCVVLPLNMNSEG
RFSSLMGKFTAFLSAESNDERMSTGSRSIIDKTLQFINQCCIDSIFSKAKYLDDDSLQHLGRALLMAAAGRTQKF
TSPLDEEDTAGFCWDLIVMITVPNLNRFMSFWPAFHEYLAMTSQLPIFASCPFVEKSLCSLLNISKKLLLQSDSQ
SEELIFKSLQVIWRIDGQVGDACCEEITQLAANVLIEDAKKIQTEFGWQSVLAVLSMGARHPEAFNDGIEALIFL
MSANGAYLTSTNYNMCIETAFSFAASKVSSIEQSKQLIDLMADSMPLLISWHHCGFADPDSSVNSGSSSSFAIQD
SSGSSTSCTTTMFMKLVENLRKMCLIRREEIRNYAVSLLRRCFVKAEELGLTSSAFMLCFNNVAFAMADDLLEKA
VEYSRRENSEKEMHSTEGTLILSMEFLADTYVQFLPLISTSSGFKSFWLGVLRRMDACIMAELGGSDSLKILVQD
LLRRMIIAMKERNILMQKEDNELWDVTVTQILKIAPAVKDELLSEN*

**>Nymco_LOC116252701**

MIGDGGEQRMAEKNSMDNNKRVKRGKEIDLSISCMVNTEVGALLAVMRMSPGDQCASQFIPGADEALDTSLMSGL
KSLRTFIFSIQQDWHSVDPSVYLSPFLEVIQFDEIAAPAVASALSATIKILKLDIFDSTTPHSREAMHLIVDAIT
NCRLERSGHAADDTILMRILQLLISCLRCKASVLLSDSAVCTIVNTCFQVVQQSASRGHLLQRSARLAMHELVQM
IYAHLPEVQRGSRNLADSDNFSSLNYGIGCMVDIFQFICSLLNVAAEVPDFNAMPSSPDEDVESFALVLINSAIE
IAGEAIGEHPKLLKMIQEDLFYNLIRHGSHSTPLVLSLICSVVLNLYNNLRSLLKLQLEAFFRFVCLKIARGKLH
SIQQEEVAIEGLVSFCRQPMFGIEMYTNFDCNPLYSNVFEEIGKLLCKRAFPNNGPLSSIQAQSFEGLVVMIQHI
ADGMEKSRSSKANSNLLSSNDASSFNSNLIRITEYKPFWTEKCERCDDPKAWVEFMRVRKYMKKRLLVSVNHFNL
DEKKGLEFLKHAHLVPDPPEAQSLAYFLRFTPGLDKTMIGDYLGDPDEFHLKVLKEFTSTFDFKNMILDAALRNF
LETFRLPGESQKIQRILEAFAERFYEQSDEMFVDKDAVFVLCYSLIMLNTDQHNAQVKKKMTEEEFIKNNRNING
GKDLPREYLSELFQSISTNEITLFDQSGGSVEMTPSRWINIINQSKTTDAYIFCEGQTQLNHDMFSIIAGPAMAT
LATVFEHADDDQIINQSMEGFLAVARIARYNLEDVLDEILAMLCKFTTLLNPFASLEETFFSLTEDVKPRLATLA
VFTIANKFGDSIRSSWRNVVDCLLRLKRLKLLPPGLAGSDNETPPPRMDLDLPLHSRTESGVIFPSTVNDDRRFS
SLIGKFGQFISTENDEESISTTVGNTNKRNLQFINQCCIEGIFSKSKHLHEESLQNLGKAILLAGAGRGQKFTSP
LDEEETAAFCWDLIVMLTENNIDRFMSFWPAFHDCLGLISQLPLFGSCPFVEKSLFGVLNVTSKLLHKADKTSEE
LIFKSLQLIWKIDSRVVDSCCEALTQISANILKENAKNIQTQFGWKSIIAILSISARSPEAFNDGVEALISLMSD
NVACVTRLNYSICIESAFSFSASKLSSIEKSMQVLDLMADSVNLLVSWLSSGFSDPGSNNSSSTAQDSSRNSISS
IGTMFVRLVESIRKICLVRREEIRNHAVLTLKRCFTKAEGLGFTSTVCLHCFNHVIFAMADDLLEKAIEYSRREN
SEKEMRSMEGTLCLAMEFLTEAFIQFLRTLSSVSGFRSFWMGVLRRMDTCMKTELGDSDKMQILIPTLLKKMILA
MKEHNILVNSEDNELWDITVTQILWIAPAVKDELFPGY*

**>Ambtr_LOC18429558**

MGRPKLQTGIKAIEEERSEDCECTSNRGALACMVNSEVGAVLAVMRRNVRWGGRYMAGDDQLEHTLVQSLKALRR
QIFSWDQNWQSINPAVYLKPFLDVIRSDETGAPITGVALTSVYKILSLEIFDLNTVNVEEAMHSIVDAVTSCRFE

VIDPASEEVVLMKILQVLLACMKSKASVVLSNQHVCTIVNTCFRIVHQAGTKGELLQRIARHTMHELIRCIFAHL
PDIECTEGSSLSNGNAAFIKSDALVGEKDYTFVSKKSENGNGSLDPENPPVSVGFATNASGNSVASLADDNVIGI
GSSNDGASDGHLMTEPYGVPCMVEIFHFLCSLLNFGEHVGMGQRSNTIAFDEDVPLFALGLINSAIELGGLAIER
HAKLLSLIQDELFRNLMQFGLSMSPLILSMVCSVVLNLYHHLRTELKLQLEAFFSCVILRLAQSRYGASYQQQEV
TMEALVDFCRQTSFMSEMYANFDCDITCTNVFEDLANLLSKSAFPVNCPLSAMHILALDGLIAVIQGMAERVGSS
QSLEQGIVGDLEEYNPFWTVKCENYSDASQWVGFVRRRKFIKRRLMIGADHFNRDPKKGLEFLQGTHLLPDKLDP
QSVACFFRYTAGLDKNLVGDFLGNHDDFCVQVLHEFARTFDFEDMNLDTALRLFLETFRLPGESQKIQRVLEAFS
ERYYEQSPHILADKDAALLLSYSLIMLNTDQHNVQVKKKMTEEDFIRNNRHINAGKDLPREFLSDLYQSICKNEI
RTSPEQGAGFPEMTPSHWIDLMKKSKKTPPYIVCDSQAFLDHDMFAIMSGPTIAAISVVFDHAEQEEVFQTCVGG
FLAVAKISASHHLEDVLDDLVVSLCKFTTLLNPVSSVEEPVIAFGDDTKARMATITVFTIANRFGDYIRTGWRNI
LDCILRLHKLGLLPARVASDAADDTELSTDPIHGKPVSSTSLTVSHIPPIGTPRRSSGLMGRFSQLLSLDAEEPR
SQPTEQQLAAHQRTLQTIQKCHIDSIFTESKFLQADSLLQLAKALIWAAGRPQKGGSSSPEDEDTAVFCLELLIA
ITLNNRDRIVLLWQGVYEHIASIVQSTVMPCALVEKAVFGLLRICQRLLPYKENLADELLRSLQLILKLDARVAD
AYCEHITQDVMRLVKANASHIKSQMGWRTISSLLSITARHPEASEPGFEALTFVMAEGAHLTRANYSLCLDASRQ
FAESRVGLTDRSLRALDLMADSVTCLVKWAREAKEAGEDAGQEIGEMWLRLVQGLRKVCLEQREEVRNHALSALQ
RCLTSAEGMGLAPALWLQCFDLVVFTMLDDLLEIAQGHSLKDYRNMEGTLRLAVKLLSKVFLQLLHELSPLPNFC
KLWLGVLGRMDKYMKAKIRGKKTEKLQEEVPELLKNMLLVMKAKGVLVQRSTLGGDSLWELTWLHVNGIAPSLHS
QVFPDQETEQEVKVADTQSPLHRSTDSSAAVSDGSPAAGESNAR*

**>Ambtr_LOC18436789**

MPKEDEDKGCAASQRPARRSERELAISCMINTEVSLVLAVMRRPLDHASPQFLPTEEPPESPLIQSLKALRSFLF
APNDDWAFVDPTLFLSPFLDVIQADDVAAPATGTALSAVLKVLRMEIFDGRTPTAAEAMKAVVEAVANCRLERTD
AGLEDAVLVRIQQVLLACVRCKAGTLLDDRVVCVAFNTCFQVVQQATGRGDLLHRQARHAMHELVQVIFMRLPEL
EGTDDDREDLDSQCEDSSSSSHPSSGYGIACMVDIFYFLCTLLNLVDSSDGEMSGGGFDEDMQLFILILINSAIE
LGGEGIGKHPRLLKIVQDDLLHHLIHYGPRSSPIVLSIICSTVLNLYHHLRCSLKLQLEAFFVHVLLRIIASNVK
FAPSFQKQQQQEVAIEGLISFCRLPQFAMEMYTNFDCDPMCQNVFEQIGNLLCKHAFPVNSPMTSLQVQSLEGLV
VIIQNITDCVLDEQSNEEDNIPVEITSYKTLWTVQCDDYEDSQKWVKFLRERKWFKKKLAIAGDHFNRDQKKGMY
FLRANHLVPEEPDPQSVGHFLRYTMGLDKTLIGDFLGDPDEFNIQVLKELTSTFDFSSVILDIALRTYLETFRLP
GESQKIQRILEAFAEKFYSDQETEMFADKDAVFVLCYSLIMLNTDQHNAQVKKKMTEEEFIRNNRCINAGNDLPR
EYLSELYHSIGNNEITLSSTNDHVSPEKVPSKWTTLTKTSISKMEPYAIWKGGPQLGREMFAAISGPSIAAISAS
FEHADDEEIIRICIEGFSAIAKMANYRLEDALDELLTSLCKFTTLLSPYTSVEDTIYAFADDPKPRVAVITVFAI
ANQYGNSIRSGWRNIVECLLKLNKLKLLPQSIVDNHNDEDEEGENEGTQGNSAANFGKSEAGVIFPSHLHHVLSR
RNSGLMGRFTQFLSLDTCAEDMRTSQVGNEFERNLQVIQQCSIDTIFTRSGSLNIDSLENLAKALVIAAGRAQKY
SSTLSEEEDAVSLCIELLVAISFNCFVEKTLLGLLRICRRIMAQRPEKLAEELIFRSIQFTWKLDAGKILETGCE
PFVRMVSRILIQNAESIHTHVGWRSALSLLSLSARNHEVFEEGADTLVGLMEGGVHITKVNFLMCVDTAFGFAAS
KLCPPEKSMKLLDLMAASVTPLVAWARAGFSDPGSGSGTGSSASGASGDEGMVRGGAIGMFMKLADALRKITLVR
REEIRNHAVSTLCRAIIEADEGLFFSPQACLQCFNTVVFAMVDDLLEKATEKKFMEGTVEMAMEVLVEVFLRLVE
RMGTVSGFRTFWLGVLRRMDTGMKADVGGGSVREMIPGLLRKMIVDMMEKGILVERDGDELWDITNIQIQWIAPS
LKEELFPEDE*

**>SEGI_06272**

MGRPKRKTEIKAIEEEPQECGGTHRRGALACMVNSEVGAVLAVMRRNARWAGRYMAGDDQLEHSLVQALKSLRRQ
IFSWQQPWHMIDPSVYLKPFLDVIRSDETGAQITGVALSAVYKILTLEVFDLNTIHVDDAMHSIVDAVTSCRFEV
TDAASEEVVLMKILQVLLACMKSKASVVLNNQDICTIVNTCFRVVHQAGSKGELLQRVARHTMHELVRAIFSHLP
ELDHAEAATKTNEGERSSVDSKGHVDELGKECRVDGRQFDNGTVDYDSSDMLVSGVLSSHSPFMINSSMEFSDEK
TTSNRVSSKDLANDVQIMIEPYGVPCMVEIFHFLCSLLNIVETMSIGARSNPLAFDEDVPLFALGLINSAVELGG
ESIGRHPKLLALIQDELFRNLMQFGLSSSPLILSMVCSIVLNLYHHLRTKLKLQLEAFFSCVIIRLAQSKYGASY
QQQEVAMEALVDFCRQPTFMSEMYANLDCDITCGNIFEDLGNLLSKSAFPVNCPLSAMHILALDGLIAVIHSMAD
RVTAASPLPEPVPVGEIEEYVPFWEINCESFENPDHWVEFVRRKKYVKRRLMIGADHFNRDPKKGLEFLQGTHLL
PEKLDPQSVACFFRYTAGLDKNLVGDFLGNHDEFCVKVLHEFAKTFDFQEMNLDTALRLFLETFRLPGESQKIQR
VLEAFSERYYEQSSKVLADKDAAFLLSYSLIMLNTDQHNVQVKKKMTEEDFIRNNRHINAGKDLPREYLSELYHS
IVKNEIRTSYEQGIGFPEMTPSRWIDLMRKSEKTPPYIVCDSRPFLDHDMFAIISGPTIAAISVVFDHAEDEDVF
HACVDGFLAVAKMSACHHLEDVLDDLVVSLCKFTTLLNPSASVEEPVLAFGDDTKARMATVTVFTIANEYGDYIR
TGWRNILDCILRLHKLGLLPARVASDAADDPDVANDSIHGKTLSGTPSISHIPVVGTPRRSSGLMGRFSQLLSLD
TDEPRSQPTEQQLAAHQRTVQTIQKCHIDSIFTESKFLQAESLLQLARALIWAAGRPQKGSSSPEDEDTAVFCLE
LLIAITLNNRDRIMLLWQGVYEHIASIVQSTVMPCALVEKAVFGLLRICQRLLPYKESLADELLRSLQLILKLDA
RVADSYCELITQGVMRLVKANAGHIKSSMGWRTISSLLSITARHPEASESGFEALSFIMSDGAHLTQANYVYCLD
ATRAFAESRVGLADRSLRALDLMAESVKCLVRWSKISSGDRKTVGDSREDTLKSSQEIGEMWLRLAHGFRKVCLE
QREEVRNYALLSLQRCLLAAEAISLAPTIWLQCFDQVIFIMLDDLLDIAQGHSPKEYRNMEGTLHHAMKFLSKIF
LQFLDQLLTLPNFRKLWLGVLSHMEKYMKSKLRGKGSEKLHELIPELLKNMLQIMKTRGILVQRSTLGGDSLWEL
TWLHVKGISPSLQTELFPDQEQEGEMEPRSDEYSSAAETVSSEAATVAK*

**SEGI_34273**

MGHLKFPIAMRSIEEEPQQNGCMTRTRGSLACIVNTEVSAVLAVMRRNARWAGHYMAGDDQLDHSLIHSLKELRR
QIFSWKQEWQHINPSIYLKPFLDVIRSDETGAPITGVALSAVYKILTMEVFDLHTVHVDLAMHSVVDAVTSCRFE
VTDPASEEVVLMKILQVLLACMKSKMSLVLNNQDVCTIVNTCFRVVHQAGTKGELLQRTARHTMHELVRSIFAHL

ADLDSTGAHTNSVKKSSLMANSHLENVGKEYISDGNEGSSTGNGNTASETGKSFSASSSVELDDKMADSKIGGKE
LANDEFLREPYGVTGIVEIFQFLCSLLNIVETMGMGTGSSLLALAEDVPLFALGLINSAIELGGPSIGRHPKLLA
LVKDELFHNLIQFGLSSSPLILSTVCSIVLNLYHHLRCELKLQMEAFFTCVIIRLAQGKYGASYQQQEVAMEALV
DFCRQPTFIFEMYANYDCEITCSNVFEDIGNLLSKNAFPVNSPLSAMHILALEGLIAVIHSMADRVNASSSLPEL
TPIRDIEEYEPFWTVTCEKNEDPIHWVPFVQRKKYIKRKLMLGADHFNRDPKKGLEFLQGIHLLPEKLDPLSVAC
FFRYTSGLDKNLVGDFLGNHDEFCVQVLQEFSKSFDFVDMSIDTALRTFLETFRLPGESQKIQRVMEAFAERYYE
QSPQVLADKDAAFVLSYSLIIILNTDRHNVQVKRKMTEEDFIRNNRRINGGKDLPREFLSELYHSILKNEIRTNYE
QSIAIPEMTTSRWIDLIRKSQKTPPYFICDYRPFLDHDMFAIISGPTIAAMSVVFDHAEDEYVFQECVDGFLAVA
QMCACHHLVDVLDDLVVSLCKFTTLLDPSSSVDEDVLAFGDDTKARMVTVTVFTIANRFGDFIRNGWRNILDCIL
RLHKLGLLPARVASEGTDDPDLLNDSVYGKALSEAPPISQLPSITRRRSSGLMGRFSQLLSLESDEPRAQPTEEQ
LAAHQQTMQTIQQCHIDNIFTESKFLQADSLLQLARALIWAAGRPQKSSSPEDEDTPVFCLELLIAISLNNRDRI
MILWQGVHDYIGSIVQTTTMPCALVEKAVFGLLRICQRLLPYKEDLAEELLRSLQLILKLDARVADVYCERITQE
IMQLVKTNAGHIRSSGGWRTVTSLLSITARHPDACEPGFEALSFVMADCAHLTRANYVLCLDAARAFAESRIGMT
DRSLQALDLIAGSVKCLVQFSKADHKNINLLGEDASKSSQETGDMWLRFAHSLRKVCLDQREEVRNHAILLLQQC
FSAVEAICLTPPTWAQLFDQIIFTMLDDLLEIAKGHSPKEYRNMESTLHYAMKLLSKLFLQLLDQLSSLPNFRKL
WLGVLSCMEKYMNSKLRGKGNEKLQELIPELLKNMLLILNDRGVLAKKSALENSSLWEMTWIHVQGISPSLKEEL
FPEKDPEHETELKTKSDEFGMDSEIGFEAAVTGVEQ*

**TaxchGNOMa_chr3**

MGRPRRKIEIKAIEEEPQECGGTRRRGALACMVNSEVGAVLAVMRRNARWAGRYMAGDDQLEHSLVQALKSLRRQ
VFSWQQPWQMIDPSVYLKPFLDVIRSDETGAPITGVALSAVYKILTLEVFDLNTIRVDDAMHSIVDAVTSCRFEV
TDPASEEVVLMKILQVLLACMKSKASVVLNNQDICTIVNTCFRVVHQAGTKGELLQRVARHTMHELVRAIFSHLP
ELDHTEAATTTNGGERPFVDSKGHVDEHGKECRQFDNGTVDYDSADISVSGALASHAPFMINSSMDFSDEKTTGN
KVSSKDLANDVQIMIEPYGVPCMVEIFHFLCSLLNIVETMGIGARSNPLAFDEDVPLFALGLINSAIELGGESIG
QHPKLLALIQDELFRNLMQFGLSSSPLILSMVCSIVLNLYHHLRTKLKLQLEAFFSCVIIRLAQSKYGASYQQQE
VAMEALVDFCRQPTFMSEMYANLDCDITCGNIFEDLGNLLSKSAFPVNCPLSAMHILALDGLIAVIHSMADRVTA
ASPLPEPVPVGEIEECVPFWEINCESFEDPNHWVDFVRRKKYVKRRLMIGADHFNRDPKKGLEFLQGTHLLPEKL
DQQSVACFFRYTAGLDKNLVGDFLGNHDDFCVKVLHEFAKTFDFQEMNLDTALRLFLETFRLPGESQKIQRVLEA
FSERYYEQSSKILADKDAAFLLSYSLIMLNTDQHNVQVKKKMTEDDFIRNNRHINAGKDLPREFLSELYRSIVKN
EIRTSYEQGAGFPEMTPSRWIDLMRKSEKTPPYIVCDSRPFLDHDMFAIISGPTIAAISVVFDHAEDDDVFHACK
DGFHAVAKMSACHHledvlddlvvSLCKFTTLLNPSASVEGPVVAFGDDAKARIATVTVFTIANRYGDYIRTGWR
NILDCILRLHKLGLLparvasdaaddpdvandsIHGKPLSGTPSVSHIPVVGTPRRSSGLMGRFSQLLSLDTDEP
RSQPTEQQLAAHQRTVQTIQKCHIDSIFTESKFLQAESLLQLARALIWAAGRPQKGNSSPEDEDTAVFCLELLIA
ITLNNRDRIMLLWQGVYEHIASIVQSTVMPCALVEKAVFGLLRICQRLLPYKESLADELLRSLQLILKLDARVAD
AYCEHITQEVMRLVKANAGHIKSSMGWRTVSSLLSITARHPDASESGFEALSFIMSDGAHLTQANYVLCLDATRA
FAEARVGLADRSLRALDLMAESVKCLVRWSKMSSGDRKTVGDSREDISKTSQEIGEMWLRLAHGFRKVCLEQREE
VRNHallslqqcllaaeaISLAPTVWLQCFDQVIFIMLDDLLDIAQGHSPKEYRNMEGTLHHAMKFLSKIFLQFL
DQLLTLPNFRKLWLGVLSHMEKYMKSKLRGKGSEKLHELIPELLKNMLHIMKARGILVQRSTLGGDSLWELTWLH
VKGISPSLQAELFPDQEQEGEMEPRSDEYSSAAETMSSEAAAVAK*

**>TaxchGNOMb_chr7**

MGHSKPQLGMRSIEEEPQQNGCVTRGQGSLACIVNAEVSAVLAVMRRNARWAGHYMAGDDELDHSLIHSLKALRR
QIFSWKQEWQSINPSVYLKPFLDVIRSDETGAPITGVALSAVYKILTMEVFDLHTVRVDHAMHSIVDAVTGCRFE
VTDPASEEVVLMKILQVLLACMKSNMSLVLNNQDVCTLVNTCFRVVHQAGTKGELLQRTARHTMHELVRSIFSRL
ADLDSTGTHANDMQRSSLISNSHIDNLGKEYVGdnndgssidnsstdADAGKSFPSTLSYEGSFTVSSSVVLDDN
TDDSRIGGKEVANDKVSRVPYDLTGIVEIFHFLCSLLNVVDTMGVGTGSNHLAIAEDFPLFALGLINAAIELGGA
SIGRHPKLLALVQDELFHNLIQFGLSSSPLILSTVCSIVLNLYHHLRRELKLQMEAFFTCVIIRLAQGKYGASYQ
QQEVAMEALVDFCRQQTFIFEMYANFDCEVTCSNVFEDIGNLLSKNAFPVNCPLSAMHILALEGLIAVIHSMADR
VNASSSLPEPVPVRDIEEYEVFWAVTCESYEDPSHWVPFVRRKKYIKRKLMLGADHFNRDPKKGLEFLQGIHLLP
EKLDPQSVACFFRYTSGLDKNLVGDFLGNHDEFCVWVLKEFAKSFDFVDMSIDTALRTFLETFRLPGESQKIQRV
MEAFAERYYEQSPQVLADKDAAFVLSYSLILLNTDQHNVQVKRKMTEEDFIRNNRRINGGKDLPREFLSELYHSI
LKHEIRTTYEQSMAIPEMTTSRWIDLIRKSQKMPPYIFCDSRPFLDHDMFAIISGPTIAALSVVFDHAEDEYVFQ
ECVDGFLAVAQMCACHHlvdvlddlvvSLCKFTTLLNPSISAEEHVLAFGDDEKARMVTITVFTIANRFGDFIRN
GWRNILDCILRLHKLGLLPARVASDAGDDPDFVIESVHGKASSEAPPLSQLPSINRRRSSGLMGRFSQLLSLESD
EPKTQPTEEQLAAHQRTMQTIQQCHIDSIFTESKFLQADSLLQLARALIWAAGRPRKSSSSEDEDTPVFCLELLI
AIALNNRDRIMILWQGVHEYIGSIVQTTIMPCALVEKAIFGLLRICQRLLPYKEDLAEELLRSLQLILKLDARVA
DVYCERITQEIMQLVKTNAGHIRSSGGWRTVISLLSITARHPDACEPGFEALSFIMADCAHLTRANYVLCLDAAR
AFAESRIGMTDRSLRALDLIAGSVKCLVQFSKIESSDHKTTSLLGEDASKSSQAIGDMWLRLANSLRKVCLDQRE
EVRNHAILSLQRCLSAVEMICLAPSTWSQFFDQVIFSMLDDLLEIAKGHSPKEYRNMEGTLHNAMKLLSKLFLQF
LDQLSALSNFRKLWIGVLSRMEKYMNSKLRGKGNDKLQELIPELLKNMLLILKDRGVLAKKSALETDSLWEMTWI
HVQGISPFLKGELFPEKNPEHESDTQAQASEFVGLDSKVGFQVAVTAVGQETES*

**>Thupl_29380805s0004**

MGRPKRKTEIKAIEEEPQECGGTHRRGALACMVNSEVGSVLAVMRRNARWAGRYMAGDDQLEHSLVQALKSLRRQ
IFIWQQPWHMIDPSVYLKPFLDVIRSDETGGQITGVALSAVYKILTLEVFDLNTIHVDNAMHSIVDAVTSCRFEV
TDAASEEVVLMKILQVLLACMKSKASVVLNNQDICTIVNTCFRVVHQAGSKGELLQRVARHTMHELVRAIFSHLP

ELDYAEAATKTNEGERSSVDSKGHVDELGKECRVDGRQFDNGTADYDSADMLVSGVLTSHSPFMINSSMEFSDEK
TTSNRVSSKDLANDVQIMIEPYGVPCMVEIFHFLCSLLNIVETMSIGARSNPLAFDEDVPLFALGLINSAVELGG
ECIGRHPKLLALIQDELFRNLMQFGLSSSPLILSMVCSIVLNLYHHLRTKLKLQLEAFFSCVIIRLAQSKYGASY
QQQEVAMEALVDFCRQPTFMSEMYANLDCDITCGNIYEDLGNLLSKSAFPVNCPLSAMHILALDGLIAVIHSMAD
RVTAASPLPEPVPVGEIEEYVPFWEINCESFENPDHWVEFVRRKKYVKRRLMIGADHFNRDPKKGLEFLQGTHLL
PEKLDPQSVACFFRYTAGLDKNLVGDFLGNHDEFCVKVLHEFAKTFDFLEMNLDTALRLFLETFRLPGEAQKIQR
VLEAFSERYYEQSSKVIADKDAAFLLSYSLIMLNTDQHNVQVKKKMTEEDFIRNNRHINAGKDLPREYLSELYHS
IVKNEIRTSYEQGIGFPEMTPSRWIDLMRKSEKTPPYIVCDSRPFLDHDMFAIISGPTIAAISVVFDHAEDEDVF
HACVDGFLAVAKMSACHHLEDVLDDLVVSLCKFTTLLNPSASVEEPVLAFGDDTKARMATVTVFTIANEYGDYIR
TGWRNILDCILRLHKLGLLPARVASDAADDPDVANDSIHGKTLSGTPSISHIPVVGTPRRSSGLMGRFSQLLSLD
TDEPRSQPTEQQLAAHQRTVQTIQKCHIDSIFTESKFLQAESLLQLARALIWAAGRPQKGSSSPEDEDTAVFCLE
LLIAITLNNRDRIMLLWQGVYEHIAGIVQSTVMPCALVEKAVFGLLRICQRLLPYKESLADELLRSLQLILKLDA
RVADSYCELITQGVMRLVKANAGHIKSSMGWRTISSLLSITARHPEASESGFEALSFIMSDGAHLTQANYVFCLD
ATRAFAESRVGMADRSLRALDLMAESVKCLVQWSKMSSGDRKTVSDSREDTSKSSQEIGEMWLRLAHGFRKVCLE
QREEVRNYALLSLQRCLLAAEAISLAPTIWLQCFDQVIFIMLDDLLDIAQGHSPKEYRNMEGTLHHAMKFLSKIF
LQFLDQLIALPNFRKLWLGVLSHMEKYMKTKLRGKGSEKLHELIPELLKNMLQIMKTRGVLVQRSTLGGDSLWEL
TWLHVKGISPSLQTELFPDQEQEGEMEPRSDEYSSAAETISSEAVTVAK*

**>Thupl_29378734s0004**

MLADWNYLRGGQAMGHLNFPIAMRSIEEEPQQNGFVTRMGGSLACIVNTEVSAVLAVMRRNARWAGHYMAGDDQL
DHSLIHSLKELRRQIFSWKQEWQSINPSLYLKPFLDVIRSDETGAPITGVALSAVYKILTMEVFDRRTVHVDLAM
HSIVDAVTGCRFEVTDPASEEVVLMKILQVLLACMKSKMCLVLNNQDVCTIVNTCFRVVHQAGTKGELLQRTARH
TMHELVRSIFAHLADLDSTGAHTNSVKRSSLMSNSHSDNVGKEYASDGNEGSSTGNGNADAETGKSFSVSSSVET
DDKMADSKIGGKELANDDVLREPYGITGIVEIFHFLCSLLNIVETMGMGTGSNLLALAEDVPLFALGLINSAIEL
GGTSIGRHPKLLALVQDELFHNLIQFGLSSSPLILSTVCSIVLNLYHHLRCELKLQMEAFFTCVIIRLAQGKYGA
SYQQQEVAMEALVDFCRQPTFIFEMYANYDCEITCSNVFEDIGNLLSKNAFPVNSPLSAMHILALEGLIAVIHSM
ADRVSASSSLPEPTPIRDIEEYEPFWTVTCEEIEDPTQWVPFVRRKKYIKRKLMLGADHFNRDPKKGLEFLQGIH
LLPEKLDPHSVACFFRYTSGLDKNLVGDFLGNHDEFCVQVLLEFSKSFDFVDMSIDTALRTFLETFRLPGEAQKI
QRVMEAFAERYYEQSSQVLADKDAAFVLSYSIILLNTDQHNVQVKKKMTEEDFIRNNRRINGGKDLPREFLSELY
HSILKNEIRTNYDQSIAIPEMTTSRWIDLIRKSRKTPPYIISDYRPFLDHDMFSIISGPTIAAMSVVFDHAEDEY
VFQECVDGFLAVAQMCACHHLVDVLDDLVVSLCKFTTLLDPSSSVDEDVLAFGDDTKAGMVTVTVFTIANRHGDF
IRNGWRNILDCILRLHKLGLLPARVASDGVDDPDISNDSVYGKSLSDTPPISHLPSITRKRSSGLMGRFSQLLSL
ESDEPRTQPTEEQVAAHQRTVQTIQQCNIDNIFTESKFLQADSLLQLARALIWAAGRPQKSGSPEDEDTPVFCLE
LLIAISLNNRDRIMILWQGVHDYIGSIVQTTTMPCALVEKAVFGLLRICQRLLPYKEDLAEELLRSLQLILKLDA
RVADVYCERITQEIMQLVKTNAGHIRSSGGWRTVTSLLSITARHPDACEPGFEALSFVMADCAHLTRANYVLCLD
AARAFAESRIGMTDRSLQALDLIAGSIKCLVQFSKADKNVNLLGEDALKSSQEIGDMWLRFAHSLRKVSLDQREE
VRNHAILSLQQCLSAVDAICLTPSTWAQLFDQIIFTMLDDLLEIAKGHSPKEYRNMESTLHYALKLLSKLFLQLL
DQLSSLPNFRKLWLGVLSCMEKYMNSKLRGKRNEKLQELIPELLKNMLLILNDRGVLAKKSALENSSLWEMTWIQ
VQGISPSLKEELFPEKDPEHETGLKTKSGDSVMDSEIGFEAPVTGVAQ*

**>PicabGNOMa_comp77**

MGRPKLQTGIKAIEEEPEECGGITKRGALACMVNSEVGAVLAVMRRNARWAGRYMAGDDQLEHSLVQSLKNLRRQ
IFSWQQPWHMIDPSIYLKPFMDVIRSDETGAPITGVALSAVYKTLTLEVFDLNTVHVDEAMHSIVDAVTSCRFEV
TDPASEEVVLMKILQVLLACMKSKTSVVLNNQDVCTIVNTCFRVVHQAGTKSELLQRVARHTMHELVRAIFSHLP
ELDHAEAVAAANGSDMPSIKSKVHVDGLPKEHRIDGRQFDNGNADSDSADVSVPGAFASQVDYSDDKTTGTKSST
KELSNDVQILIEPYGVPCMVEIFHFLCSLLNIVETMGIGARSNPLAFDEDVPFFALGLINSAIELGGQSIGQHPK
LLALIQDELFRNLMQFGLSSSPLILSMVCSIVLNLYHHLRMKLKLQLEAFFSCVIIRLAQSKYGASYQQQEVAME
ALVDFCRQPTFMPEMYANLDCDITCANIFEELGNLLSRSAFPVNCPLSAMHILALDGLIAVIHSMADRVTAASPL
PEPVPVGEIEEYAPFWTIKCESYEDSNHWVEFVRRKKYIKRRLLIGADHFNRDAKKGLEFLQGTHLLPEKLDHQS
VACFFRYTAGLDKNLVGDFLGNHDEFCVKVLHEFAKTFDFQDMNLDTALRLFLETFRLPGEAQKIQRVLEAFSER
YYEQSSGILADKDAAFLLSYSLIMLNTDQHNVQVKKKMTEEDFIRNNRLINSGKDLPRDYLSELYHAIVKNEIKT
IYEQSLFPEMNPSRWIDLIRKSQKTPPYIVCDSRPFLDHDMFAIISGPTIAAISVVFDHAEDEDVFHTCVDGFLA
VAKMSACHHLEDVLDDLVVSLCKFTTLLNPAASVEEPLLAFGDDTKARMATVTVFTIANRFGDYIRTGWRNILDC
ILRLHKLGLLPARVASDAAEDPDLANDSMHGKPLSGAPSISHIPVVGTPRRSSGLMGRFSQLLSLDTDEPRSQPT
EQQLAAHQRTVQTIQKCHIDSIFTESKFLQAESLLQLARALIWAAGRPQKGSSSPEDEDTAVFCLELLIAITLNN
RDRIMLLWQGVYEHIAGIVQSTVMPCALVEKAVFGLLRICQRLLPYKENLADELLRSLQLILKLDARVADAYCEH
ITQEVMRLVKANAGHIKSPMGWRTVSSLLSITARHPEASECGFEALSFIMSDGAHLTQANYVLCLDATRAFAESR
VGLADRSLRALDLMTESVKCLVRWSKSVSVDWKTVGDSREDTSKSSQEIGEMWLRLAHGLRKVCLEQREEVRNYA
LLSLQRCLSAADAICLTPSVWLQCFDQVIFTMLDDLLEIAQGHSPKEYRNMEGTLHHAMKFLSKIFLQFLDQLFT
LPNFRKLWLGVLSRMEKYMKAKLRGKGSDKLQELIPELLKNMLLILKARGVVMQRSTLGGDSLWELTWLHVKGIS
PSLQAELFPDLEPEAETEPKSDDYSSAAETVGS*

**>PicabGNOMb_comp95**

MGHSQPQSEIRVNEEQPHRCNNVARRGALACIVNTEVGAVLAVMRRNARWAGHYMSGDDQLEHSLIHSLKALRRK
IFGLQQEWHSINPSVYLKPFLDVIRSDETGAPITGVALSAIYKILTMEVFDLDTVHVEDAMHSIVDAVTSCRFEV
TDPASEEVVLVKILQVLLACMKSKNSIVLSNQDVCTIVNTCFRVVHQAGTKGELLQRTARHTMHELVRSIFARLP

GLDSTTACTNGVESSSIMFNTRVDDSGKESGDDSHKYRNFDNGSADSNSSKKSLSSTSTLEGSFTANSKAELNDK
MVDSRIGEKILADDLGVLKEPYGVAGIVEIFHFLCSLLNIVETIGLGTRSNHLALDEDVPLFALGLINSAIELGG
PAIGKHPKLLALVQDELFYNLMQLGLSSSPLILSMVCSIVLNLYHHLRCQLKLQLEAFFTCVILRLAQGKYGASY
QQQEVAMEALVDFCRQPTFIFEMYANFDCDVTCSNIFEDICNLLSKSAFPVNSPLSAMHVLALEGLIAVIHSMAD
RVNAASSLPEPAPLREIENYEPFWTVTCENYEDPTHWVPFVRRKKYIKRKLMIGADHFNRDTKKGLEFLQGIHLL
PEKLDPQSVACFLRYTWGLEKLLVGDFLGNHDEFCVRVLQEFTKMFDFVGMNIDTALRTFLETFRLPGESQKIQR
VLEAFAERYYEQSPQILVDKDAAFVLAYSLILLNTDQHNAQVKKKMTEEAFIRNNRHINGGKDLPRELLSVLYHS
ILSNEIRTSYDQNLAIPEMTPSRWIDLIQKSQKTPPYIVCDSRPFLDNDMFAIISGPTIAAMSVVFDHAEDEDVF
QECVDGFLAVAKVSACHRLEDVLDDLVASLCKFTTLLNPSASVEEPVLAFGDDTKARMVTVAVFTITNRFGDYIR
NGWRNILDCIIRLHKLGLLPARVASDAADDPELANDSIHGKVSSGGPSISHMPSIGTRRRTSGLMGRFSQLLSLD
SDEPRTQPTEQQLAAQQRTVQTIQQCRIDSIFTESKFLQAESLLQLTRALIWVAGRLQKSSSPEDEDTAVFCLEL
LIAISLNNRDRIMLLWQGIHDYIAGIVQTTIMPCALVEKAVFGLLRICQRLLPYKENLAEELLRSLQLILKLDAR
VADVYCERITQEIMQLVKTNAGHIKSPIGWRTVVSLLSITARHPDASEPGFEALSFIMADGAHLTHANYVLCLDA
ARSFAESRIGTTDRSIRALDLIAGSVKCLVQWSKTDSHASIVVGLSREDTSRSSQEVGDMWLRLVHCFRKVCIDQ
REEVRNHAILSLQRCLSAAEVIYIPPSTWSQCFDQVIFTMLDDLLEIAQGQSRKEYRNMEATLQHAMKFLSKLFL
QLLDQLSSLSNFRKLWLGVLSIMEKYMKAKLRGKGSEKLQELIPELLKNMLLILKARGVLVQKSTLNGDGLWEIT
WLHIQGISPSLKAECFPDQDPEYKPDPKSNASELVEDAAAGYEAVINDGSQVRQEKGEK*
**>AALBA5B993185**
MGRPKLQTGIKAIEEEPQECWGITRRGALACMVNSEVGAVLAVMRRNARWAGRYMAGDDQLEHSLVQSLKNLRRQ
IFSWQQPWHMIDPSIYLKPFMDVIRSDETGAPITGVALSAVYKILTLEVFDLKTIHVDEAMHSIVDAVTSCRFEV
TDPASEEVVLMKILEVLLACMKSKTSVVLNNQDVCTIVNTCFRVVHQAGTKSELLQQVARHTMHELVRAIFAHLP
ELDHTEAVAAVNGSDMHSIKSKVHVDGLPKEHRIDGRQFDNGSADSDSANKSVPGALASQVDYSDDKTMGTKCSS
KDLCNDVQILIEPYGVPCMVEIFHFLCSLLNIVETMGIGARSNPLAFDEDVPFFALGLINSAIELGGEFIGQHPK
LLALIQDELFRNLMQFGLSSSPLILSMVCSIVLNLYHHLRSKLKLQLEAFFSCVIIRLAQSKYGASYQQQEVAME
ALVDFCRQPTFMPEMYANLDCDITCANIFEELGNLLSRSAFPVNCPLSAMHILALDGLIAVINSMADRVTAASPL
PEPVPVGEIEEYAPFWTIKCESYEDANHWVEFVRRKKYIKRRLMIGADHFNRDAKKGLEFLQGTHLLPEKLDHQS
VACFFRYTAGLDKNLVGDFLGNHDEFCVKVLHEFAKTFDFQDMNLDTALRLFLETFRLPGEAQKIQRVLEAFSER
YYEQSSGILADKDAAFLLSYSLIMLNTDQHNVQVKKKMTEEDFIRNNRHINAGKDLPRDYLSELYHAIVKNEIRT
IYEQSVGFPEMNPSRWIDLIRKSEKTPPYIVCDSRPFLDHDMFAIISGPTIAAISVVFDYAEDEDVFHTCVDGFL
AVAKMSACHHLEDVLDDLVVSLCKFTTLLNPAASVEEPLLAFGDDTKARMATVTVFTIANRFGDYIRTGWRNILD
CILRLHKLGLLPARVASDAAEDPDLANDSMHGKPLSGAPSISHIPVVGTPRRSSGLMGRFSQLLSLDTDEPRSQP
TEQQLAAHQRTVQTIQKCHIDSIFTESKFLQAESLLQLARALIWAAGRPQKGSSSPEDEDTAVFCLELLIAITLN
NRDRIMLLWQGVYEHIAGIVQSTVMPCALVEKAVFGLLRICQRLLPYKENLADELLRSLQLILKLDARVADAYCE
HITQEVMRLVKANAGHIKSPMGWRTVSSLLSITARHPEASECGFEALSFIMSDGAHLTQANYVLCLDATRAFAES
RVGLADRSLRALDLMSESVKCLVRWSKSVSGDWKTVGDSREDTSKSSQEIGEMWLRLAHGLRKVCLEQREEVRNY
ALLSLQRCLSAADAICLTPSVWLQCFDQVIFTMLDDLLEIAQGHSPKEYRNMEGTLHQAMKFLSKIFLQFLDQLF
TLPNFRKLWLGVLGRMEKYMKAKLRGKGSDKLQELIPELLKNMLLILKAQGVVMQRSTLGGDSLWELTWLHVKGI
SPSLQAELFPDLEPEAETEPKSDDYSSAAETVGSEVATADSGQNGVNAKKLESEERGGN*
**>AALBA5B1100842**
MGHSQPQNEIRGNEEQTHQGNNVARRGSLACMVNTEIGAVLAVMRRNARWSGHYMSEDDQLEHSLIHSLKVLRRK
IFGLQQEWHSIDPSVYLKPFLDVIRSDETGAPITGVALSAVSKILTMEVFDLNTVHVEDAMHAIVDAVTSCRFEV
TDPASEEVVLVKILHVLLACMKSQNSVVLSNQDVCTIVNTCFRVVHQAGTKGELLQRTARHTMHELVRSIFARLP
DLDSTAACANGVEISSIMSNSHVDDHGKESGDDSHKYGNFDNGSADSDASKKSLSSTLTLDGSFTAISKAELDDK
MVDSRIGEKILADDVGVLREPYGITGIVDIFHFLGSLLNIVEKMGLGTGSNHLALDEDFPLFALGLLNSAIELGG
PAIGRHPKLLALVQDELFENLTQLGLSSSPLILSTVCNIVLNLYHHLRCQVKLQLEAFFTCVILRLAQGKYGASY
QQQEVAMEALVDFCRQPTFISEMYANFDCDITCSNVFEDICNLLSKSAFPVNSPLSAMHVLALEGLIAVIHSMAD
RVNAASSLPDPAPLRDIEEYEPFWTVACENYEDPTHWVPFIRRKKYIKRKLMIGADHFNRDTKKGLEFLQGIHLL
PEKLDPHSVACFFRYTWGLEKLLVGDFLGNHDEFCVQVLQEFTKMFDFVDMNIDTALRTFLETFRLPGESQKIQR
VLEAFAERYYEQSPQILADKDAAFVLAYSLILLNTDQHNAQVKKKMTEEDFIRNNRRINGGKDLPRELLSELYHS
ILSNEIRTSYEQNLAIPEMTPSRSNXXXXXRLEDVLDDLVASMCKFTTLLNPSASVEEPVLAFGDDTKARMVTVA
VFTITNRFGDYIRNGWRNILDCIIRLHKLGLLPARVASDAADDPELANDSIHGKVSSGGLPLSHMPSIRRRTSGL
MGRFSQLLSLDSDEPRTQPTEQQLAAQQRTVQTIQQCRIDSIFTESKFLQAESLLQLTRALIWVAGRLQKSSSAE
DEDTAVFCLELLIAIALNNRDRIMILWQGIHDYIAGIVQTTIMPCALVEKAVFGLLRICQRLLPYKENLAEELLR
SLQLILKLDARVADVYCERITQEIMQLVKTNAGHIKSPIGWRTVVSLLSITARHPDASEPGFEALSFIMADGAHL
TRTNYVLCLDAARSFAESRIGTTDRSMRALDLITGSVKCLVQWSKTDSCDSIAVGISREDTSKSSQEVGXXXSSQ
EVGDMWLRLVHCFRKLSSLSNFRKLWLGVLSVMEKYMKAKLRGKGSEKLQELIPELLKNMLLVLKARGVLVQKST
LNSDSLWEMTWLHIQGISPCLKAECFPDQNPEGKQDPMSNASELVEDAAAGYEAAINDGSQARQEKGEK*
**>Gnemo_TnS000619529**
MHNMRCPNVQNGNNGIEETQECVGFAKRGSLACMVDSEAGAVLAVMRRNGRWASRYMAGDDQLEHLLVQSLKNLR
RQLFSWQQAWKTIEPSIYVKPFLDVIRSDETGAAITEVALSALYKILTIQFFDLKTVNVHEAMHLMVDAVSSCRF
EPTDPASEEVVLMKILQVLLACMKNKASVVLSNQDVCTILNTCFRIVHQAVTKSELLQRVARDTMHELVRVTFAH
LPDLHKSETAVAASGGDTVSLQSEVNVNGLSNGLKVNSRQLDSGAEYEGNTVTVLSKITMQVDGKAYSKEPVTET
QVVVESYGIPCMVEIFHFLCSIMNFGETNDGAARANSLAFDEDIPLFALGLINSAIELGGQSIGEHPKLLSLIHD

ELFRNLMQFGLSPSPLILSKVCSIVLNLYHHLRTKLKLQLEAFFSCVILRLAQSRYGASYQQQEVAMEALVNFCR
QPSFMSEMYANFDSDITCANIFEEIGNLLSKSAFPVNCPLSTMHILSLEGLIAVLHNMADRATVASPVPETLPIG
EIEEYTPFWTIECGNYDDADQWVNFVRRKKYIKRRLMIGADHFNRDPKKGLEFLQGTHLLPEKLDPQSVACFFRY
TAGLDKNLVGDFLGNHDEFCVQVLNEFAKTFDFKDMQLDTALRLFLETFRLPGESQKIQRVLEAFAERYYEQSAG
IFANKDAAFVLSYSLIMLNTDQHNAQVKKKMTEEDFIRNNRRINGGSDFPKDYMSDLYHSIVKNEIRTTLDPGLG
FPEMTACRWTDLMRKSAKASPFIVCDSIPFLDHDMFAIISGPTIAAISVVFDHAEEEDVLHACVDGFLAVAKMSA
SHHLEDVLDDLVVSLCKFTTLLNPVASVEEPVLAFGDDTKARMATVAVFTIANTYGDYIRTGWRNILDCILRLHK
LGLLPARVASDVTDDPDSLNDTVHGKPIAGNPSVSHMPVVGTPRRSSGLMGRFSQLLSLDIDEPRSQPSEQQLAA
HQRTVQTIQKCHIDSIFTESKFLHLESLLQLARALIWAAGRPQKGGSSSEDEDTAVFCLELLIAITLNNRDRIMR
LWQGVYDHIAGIVQSTVMPCALVEKAVFGLLRICQRLLPYKESPADELLRSLQLILKLDARVADAYCEHITQEIM
RLVKANATHINSLMGWRTVSSLLSSTARHPEASEAGFETLSFIMTEGAHLSTLNYALFLDAARAFAESRIGLADR
SLRALDLMAESIKCLVQWSVNNKNNQKPLGNISDDVAKPAQDFGELWLRLAHGLRKVCLEQREEVRNYALMSLQK
CLLGADSIPLSSSTWLQCFDQVIFTMLDDLLEIAQGHSPKEYRNMEGTLYHAVKFISKIFLQLLDQLSVLPNFRK
LWIGVLSRMEKYMKVKIRGKGSEKLQECIPELLRGMLNTLKVKGIIVQRSTLGGDSLWELTWLHLKGISPSLQAE
LFPDKEPEEETEAKSDDYASAAETVSSDAIVVESSQSGIHTKSETKEMDGMMS*

**>Gnemo_TnS000326955**

MGFSTDESGYREKQQHLNNFEEMSKRGALACMVNTEVGAVLAVMRRNARWAGTYMSGDDQLEHSLIHSLKSLRKK
IFSWHDEWQQINPSMYLKPFLDVIRSDETGAPITGVALSAIYKILTMEVFDLNTIHVEEAMHSIVDAVTSCRFEV
TDSASEEVVLMKILQVLLACMKCKNSVVLSNQDVCTIVNTCFRVVHQAGTKGELLQRTARQTMHELVRTIFARLL
DFVPGTNCGNQAESSNSMMTSHAVDCESRCETENNNNPLSNGETNSSELKESLGSNIHSPVHVDSSCHRNDAVTG
SGNGEKKSADDTETLKEPNGITAIIEIFQFLCSLLNVDDTFGVDVGPTHLAFEEDVPLFALGLINSAIELGGPAF
RSHPKLLALIQDDLFSSLMRFGLSPSPLILSTVCSIALNLYHHLRKDIKLQLEAFFTCVIIRLAQGKHGASYQQQ
EVAMEAMVDFCRQPTFVFEMYANFDCEITCSNIFEDVGNALSKSAFPLSLDFPLSSMHILALEGLIALVQNMVER
TGTQLPGPDLKPIKEIVKYEPFWTSTCDNYEDPSSWVTFVCQKKYLKRKLMVGVDHFNREPKKGLEFLKSCHLLP
DHLEPKSVAWFFRYTPGLDKSSLGDFLGNHDEFWVSVLHEFAKTFDFTEMSIDMALRTFLETFRLPGESQKIQRV
LEAFAERYFEESPQILVDKDAAFILAYSLILLNTDRHNAQVKKKMSEEDFIKNNRRINGGNDLPREMLSELYHSI
LTNEIRTQYEHSTAIPEMTPSRWVDLIRKSRSTSPYIMVDSVPLLDQDMFIIISGPTIAAMSVIFDNANDEEILQ
TCIDGFQAVAKLAAFYHLEHVLDDLVVSLCKFTTLLNTPASLEEPILLFGDDMKTRIVTVTVFMIANKYGDHIRT
GWRNIIDCIIRLHKLGLLPSSIGSDGTEDSDGSSDSSNGKASTSIPVTQKLSAVPAKRKSSSLMGRFSQFLYFDS
EETRSQPTEQELAAQQRAVGTIQQCNISSIFKESKFLQAQSLEHLARALIWAAGKPQKGSEDEDTSVFCLQLLIV
VTLNNRDRIMILWQGVHDYIANIVQTIDMPCALVEKAVFGLLRVCQRLLPYKENLAEELLRSLQLILKLDARVAD
VYCERITQEIMQLVKANAGHIKSLIGWRTVASLLSITARHPEASEPGFEALSFIMVEGAHLNQVNYCLCIDAARA
FAESRIGTIERSLKALDLIAGSVKCLAQWSKCVVGEYAAIGVLQENNLISAQDVRDMWQRLLLSIQKICLDQRGD
VRNHAILSLQRCLSAVDASLLQIFSWSECFERLIFKLLDDLFEISQVRSSKEYKSMEVTLQHAIKLMSKFFLQVL
DQLSSMPNFESLWFGVLTRMEKYMKAKLRGRGVERIQELLPELLKNMLLILKARGLLIKTKTNDGKSLWEMTWVH
VQEISPDLMNEIFPDHEVNEKIESTDGTSTIKDSNVNDKVTNADDNSQVTG*

**>Welmi_07742**

MGCPNSHNRLKVVEESQEFVGPAKRGSLACMIDSEVGAVLAVMRRNARWVGRYMASEDQLEHSLVQALKNLRRQL
FSWQQPWNTIDPCAYLKPFLDVVRSDEAGAAITEVALFAVYKILAIEFFNLNTVNVDEAMHSVVDAVTNCRFEIT
DSASEEVVLTKILQVLLVSMKSKSSVVLSNQDICTIVNTCFRIVHQAGSKSELLQRISRDTMHELVRAIFGHLPE
ITQAERAAASAANGGGRPPLHCKANGDVFPNGPKSERRNLDNGTGDFNVINRSAQNAVVPRVDPEKITSGKTGNK
ESVNDAQVLIEPYGVHCMIEIFHFLCSIMNIVETTDIGPRSNSLAFDEDMPLFALGLINSTIELGGQSIGQHPKL
LSLIQDELFRNLMQFGLSPSPLILSKVCSITLNLYHHLCTKLKLQLEAFFSCVIIRLAQSRYGASYQQQEVAMEA
LVNFCRQPTFMSEMYANFDCDITCANIFEEIGNLLSKSAFPVNCPLSAMHILALDGLIAVLHNMADRVTGLSPIP
ETAPIGEIEEYTPFWTIECGNYNDAAQWVNFVRRKKYIKRRLMIGADHFNRDPKKGLEFLQGTHLLPEKLDPQSV
ACFFRYTAGLDKNLVGDFLGNHDEFCVQVLHEFAKTFDFHDMMLDTALRLFLETFRLPGESQKIQRVLEAFSERY
YEQSSGILADKDAAFLLSYSLIMLNTDQHNVQVKKKMTEEDFIRNNRRINAGNDLPREYLSELYQSIVRNEIKTI
YDQGLGFPEMTACRWTDLMRKSAKTSPFIVCDSRPFLDHDMFAIISGPTIAAISVVFDHAEEEDVLHTCVDGFLA
VAKMSASHRLEDVLDDLVVSLCKFTTLLNPIASVEEPVLAFGDDTKARMATVTVFTIANTYGDYIRTGWRNILDC
ILRLHKLGLLPARVASDVTDDPDISNDTNQGKPIPGNPSVSHIPVVGTPRRSSGLMGRFSQLLSLDIDEPRSQPT
EQQLAAHQRTVQTIQKCHIDSIFTESKFLHPESLLQLARALIWAAGRPQKGGSSSEDEDTAVFCLELLIAITLNN
RERVMLLWQGVYDHIAGIIQSTVMPCALVEKAVFGLLRICQRLLPYKENLADELLRSLQLILKLDARVADAYCEH
ITQEIMRLVKANASHINSTMGWRTVSSLLSSTARHPEASEAGFETLSFIMTDGEHLSPANYLFFLDAARAFAESR
VGLADRSLRALDLMAESVKCLVKWSNNFTSHQRTVSSPDDVSKPLQEFGELWLRLAHGLRKVCLEQREEVRNYAL
LSLQSCLLGVDAVPLTPSVWIQCFDQVIFTMLDDLLEIAQGHSPKEYRNMEGTLFQAVKFISKIFLQLLDQLSTL
PNFRKLWIGILSRMEKYMKAKLRGKGSDKLQEYIPELLKSILNAELFPDKETEEETETKSDDYASAAETAELFPD
KETEEETETKSDDYASAAETVSSEVALADSAQNSLTGKLESKEMDTIC*

**>Welmi_23859**

MGHLKVDSVKEKGEKVSKVDAMKRRGALACMVNTEVGAVLAVMRRNARWAGHYTAGDDQLEHSLIHSLKSLRQQI
FSWQHEWQMIDPSVYLKPFLDVIRSDETGAPITGVALSAIYKILVMDVFDLNTIHVDEAMHSIVDAVTSCRFEVT
DSASEEVVLMKILQVLLACMKSKNSVILSNQDVCTIVNTCFRVVHQAGTKGELLQRTARQTMHELVRSIFARLLD
FDSYDSTSVAESFITVIKNSHVAELSKQYRCEINKNYSSNGAPESSEYNESLSTSNVDGVVTVDTSQEVDDQTAD
SINGKWTIKLLIASTGKISVDHGQFLKQPYGITAVVEVFQFLCSLLNVVDTIGAQVGSNHLALEEDVPLFALGLI

NSAIELGGLAFGSHPKLLKLVQDDLFFNLMKFGLSPSPLILSIVCSIVLNLYHYLRQKIKLQLEAFFSCVIIRLA
QGKHGASYQQQEVAMEALVDFCRQPAFISEMYANFDCEITCNNIFEDMGNVLSKNAFPLNVDFPLSSMHILALEG
LIALVGNMTDRANNELSLPDMKPVKEIEKYEPFWTSTCENYEDPLCWVPFIRQRKYLKKKLMVAVDHFNREPKKG
LEFLKANHLLPEHLDPQSVACFFRYTSGLDKSFVGDFLGNHDDFWVFVLHEFARTFDFHKMTIDMALRTFLDTFR
LPGESQKIQRVLEAFAERYYEQSPQILEDKDAAFILAYSLIMLNTDRHNAQVKKKMTEDDFIRNNRRINGGKDLP
REFLSELYHSILTNEIRTTYEQSATVQEMTPSRWFDLIRRSRQAPPYIMSDSLSLLDHDMFAIISGPTIAAMSVV
FDNANNEETLRKCIDGFLAVAKLASRYHLEHVLDDLVGSLCKFTNLINPVASVEEPVLAFGDDTKARMATVAVFT
IANTYGDYIRTGWRNILDCILRLHKLGLLPARVALWSDGADDLDDSANDPNNGKASSGLAAVPKVSGTMTKRRSS
GLMSRFSQLLSFDSEEPRIQPTEQQLAAQQRATETIQRCHIDSIINESKFLQADSLQYLARALVWAAGKPLKSLS
EDEDTSVFCLQLLIAVTLNNRDRILILWQGVHDYIATIVQTVVMPCPLVEKAIFGLLRVCQRLLPYKENLAEELL
RSLQLILKLDAKVADVYCERITQEILQLMKANAGHIKSPVGWRTVASLLSTTARHPEASEPGFEALSFIMVEGAH
LTQINYSLCIDAARAFAESRIGTVERSVKALDLIAGSVKCLLLWSKPHVGEYAAVGLTHEKVLASTQEVRDMWLR
LVFSFQRVCVDQREEVRNHAILSLQRCLSAADVGSFPNFSWPEFFEKLVFKLVDELIDISQCRSPKDYRSMEGTL
QNALKLMSKCFLHLLDQLSSAPDFQSLWLEIFQEQDADSKSELHVKASVIEDVNVAVETAVNNN*

**>Ginbi_01979**

MGRPKLHTGNIKVIEEEPQDCGGLNRRGALACMVNSEVGAVLAVMRRNARWAGRYMAGDDQLEHSLVQSLKSLRR
QIFSWQQPWHMINPAIYLKPFMDVIRSDETGAPITGVALSAVYKILTLEVFDLNTVHVGEAMHSIVDAVTSCRFE
VTDPASEEVVLMKILQVLLACMKSKASVVLNNQDICTIVNTCFRVVHQAGTKGELLQRVARHTMHELVRAIFAHL
PDLDYVDPVPSVDGGDGPCIKSKAHVDGVGKECRADGRQYDNGNGDSVSADISVSSALTSQGPITISSSNDKTTG
SRTSGKELTNDIQIMIEPYGAPCMVEIFHFLCSLLNIVETMGIGARSNPLAFDEDVPLFALGLINSAIELGGQSI
GRHPKLLALIQDELFRNLMQFGLSQSPLILSMVCSIVLNLYHHLRIKLKLQLEAFFSCVIIRLAQSKYGASYQQQ
EVAMEALVDFCRQPTFMSEMYANLDCDITCGNIFEDLGNLLSKSAFPVNCPLSAMHILALDGLIAVIHSMADRVT
AASPLPEPVPVGEIEEYAPFWTIKCESYADSSHWVEFVRRKKYIKRRLMIGADHFNRDPKKGLEFLQGTHLLPEK
LDPQSVACFFRYTSGLDKNLVGDFLGNHDEFCVQVLQEFAKTFDFHDMNLDTALRLFLETFRLPGESQKIQRVLE
AFSERYYEQSSKILADKDAAFLLAYSLIMLNTDQHNVQVKKKMTEDDFIRNNRHINAGKDLPREFLSELYHSIVK
NEIRTSYEQGAGFPEMTPSRWIDLMRKSEKTPPYIVCDSRPFLDHDMFAIISGPTIAAISVVFDHSEDEDVFHTC
VDGFLAVAKMSACHHLEDVLDDLVVSLCKFTTLLNPAASVEEPVLAFGDDTKARMATVTVFTIANRFGDYIRSGW
RNILDCILRLHKLGLLPARVASDAADDPDPANDSMHGKPLSGAPSISHIPAVGTPRRSSGLMGRFSQLLSLDADE
PRSQPTEQQLAAHQRTVQTIQKCHIDSIFTESKFLQAESLLQLARALIWAAGRPQKGSSSPEDEDTAVFCLELLI
AITLNNRDRIMLLWQGVYEHIAGIVQSTVMPCALVEKAVFGLLRICQRLLPYKENLADELLRSLQLILKLDARVA
DAYCEHITQEVMRLVKANAGHIKSPMGWRTVSSLLSITARHPEASESGFEALSFIMADGAHLTQANYVLCLDATR
AFAESRVGLADRSLRALDLMAESVKCLVRWSKAISSDWKTVGDSREDTSKSAQEIGEMWLRLAHGLRKVCLEQRE
EVRNHALLSLQRCLSAAEIICLTPSVWLQCFDQVIFTMVDDLLEIAQGHSPKEYRNMEGTLHHAMKFLSKIFLQF
LDQLFSLPNFRKLWLGVLSRMEKYMKAKLRGKGSDKLQELIPELLKNMLLIMKARGVLVQRSTLGGDSLWELTWL
HVKGISPSLQAELFPDQEQEHDQEPEPEPKSDEYASAAETVSLETATAESGHNLNSGIVHNSRKQESEEGSGK*

**>Ginbi_26378**

MGHAKFRIGIKAIEEGPQQCGSISRRGALACMVNTEVGAVLAVMRRNTRWAGHYMAGDDQLEHSLIHSLKALRRQ
IFSWQQQWHSINPSIYLKPFLDVIRSDETGAPITGVALSTVYKILTMEVFDLNTVHVDEAMHAIVDAVTSCRFEV
TDPASEEVVLMKILQVLLACMKSKTSVVLNNQDVCTIVNTCFRVVHQAGTKGELLQRTARHTMHELVRAIFAHLA
DLDNAITAVHTNGVERPSIMSNSHVDKLEKESQDDGRGGRSLDNGNADSDACEGSLTGAIALEDSLTADTSVGMD
GKLADSRISRKELADDVEVFGERYGILGMVEIFHFLCSLLNIVETMGIGSGSNLLALDEDVPLFALGLINSTIEL
GGQSIGQHPKLFILVQDELFRNLMQFGLSSSPLILSMVCSNVLNLYHHLRCGLKLQLEAFFTCVLIRLAQGKHGA
SYQQQEVAMEALVDFCRQPTFISEMYANFDCDITCSNVFEDLGNLLSKSAFPVNCPLSAMHILALEGLIAVIHSM
ADRVNAASLLHEPAPVKEIEEYVPFWTVTCGNYEDPNQWVPFVRRKKYIKRRLMIGADHFNRDPKKGLEFLQGIH
LLPEKLDPQSVAYFFRYTAGLDKNLVGDFLGNHDEFCVWVLEKFAKTFDFVDMNLDTALRTFLETFRLPGESQKI
QRVLEAFAERYYEQSPHILADKDAAFILAYSLILLNTDQHNTQVKKKMTEDDFIRNNRCINGGKDLPREFLSELY
HSIVKNEIRTSYEQGIAIPEMTPSRWIDLIRKSQKTPPYIVCDSRPFLDHDMFAIISGPTIAALSVVFDHAEDEE
VFQTCVDGFLAVAKMSACHHLEDVLDDLVVSLCKFTTLLNPSASVEEPVLAFGDDTKARMATITVFTIANKFGDY
IRNGWRNILDCILRLHKLGLLPSRVASDAADNPDLAMDSTHGKASSGGPSMSHIPSLGTRRRSSGLMGRFSQLLS
LDSDEPRSQPTEQQLAAHQRTMQTIQQCRIDSIFTESKFLQAESLLQLARALIWAAGRPQKSSSSPEDEDTAVFC
LELLFAITLNNRDRIMILWQGVHEYIAAIVQTTIMPCAFVEKAVFGLLRICQRLLPYKENLAEELLRSLQLILKL
DARVADVYCERITQEVMQLVKANAGHIKSPMGWRTVASLLSITARHPDASEPGFEALSFIMADGAHLTRANYVLC
LDAARAFAESRIGLADRSLRALDLIAESVKCLVQWSKNDLSDHKTICLSQEDTSKSSQEIAEMCLRLAHSLRKVC
LEQREEVRNYAVLSLQRCLSGSEIISLMPSIWLQCFDQVIFTLLDDLLEIAQGRSPKEYRNMDGTLHHAIKFMSK
LFLQFLDQLSSLPSFRKLWIGVLSRMEKYMKAKLRGKGNEKLQELIPELLKNMLLIMKARGVLVQNSTLGGDSLW
ELTRRDVQGISPSLKEEFFPEQVPE*

**>Salcu_s007.g003923**

MRKFSRSNGALACMVNAEIGVVLAVMRRNPRWAGRYMGGDDQLEHSLVYSLKTVRRSIFSWQQPWNKINPSLYLA
PFLDVIRSDETGAPITSVALNSVYKILVFEVFDLQTAHVDEAMHEIVDAVTSCRFEISDIASEEMVLMKILQVLV
AVMKHPTSNLLSDRDVCLVVNTCFRVVHQAVTKGELLQRTARHSMHEIVMSIFAHLPDLDPSLQHLTRSASGSHA
TELVGVDAQNNGTTNGYGLGTTDLPSIRLDKSMDVSCSNVDGSSMKATVELQERRKENGQIVDLVNKVHLVIEPY
GILGIVEIFRYLCSLLNIVDSGIITGQKLLAIDEDVPLFALGLINSAIELGGSSFGCHPELIALVQDDLFRFLMQ
LGLSNMNPLILSMICSIVLNLYHHLRSHLKLQLEAFFSCVIIRLAQGKYGASYQQQEVAMEVLVDFCRQSTYMPE

MYANFDCDITCGNIFEDLGNLLSKSAFPVNCPLSAMHILALEGLIAVIHSMAERAGSLSPATTPEEPFQLNPGQD
YVPFWRLRCESYKDAKSWVKFVQRQKYIKRRLMIGADHFNRDPKKGLEFLQGIRLLPEKLDPQSVACFFRYTTGL
DKNLLGDFLGDPDEFCLQVLAEFAKTFDFQGMNLDAAMRTFLETFRLPGESQKIQRVLEAFAGRYYEQSSQIFAD
KDAAFLLAYSLIMLNTDQHNSQVKKKMTEEDFIKNNRSINGGQDLPREFLSELYHSIVRNEIKISYDNTTGVTEM
TPSRWVGLIHRSHKTSPYIKCDVRPFLDRDMFAIISGPTIAAISVVFDHAEDEDVLRMCVEGFLAVAKISSSHHL
EDVLDDLIVSLCKFTTLLNPNIVEEPVIAFGDDTKARMAAVTVFNIANRYGDYIRTSWRNILDCILRLHKLGLLP
ARVASDSGDSMEILSDNMQGRQIPSGPTLIAVPPAGLRKRSQGLMGRFSQLLSLETDEPRSQPTEQQLAAHQRTM
RAIEACRIENIISQSKFLQAESLLQLVRGLIWAAGRPQKAAGSTEDEDTAVFCLDLLVAVTLNNRDRIMLLWQGV
YEHIANIVQSALIPSTLVEKAVLGLLRICQRLLPYKEDLADELLRSLQLILKLDARVADALCERITQEVMQLVKT
NAKHIRSPMGWRTVSSLLAVTVRHSEASEVGFEALSFIMTEGAHLMRANYVLCLDAARAFAEVSDRGNERAIQAL
DLMADSTKCLIRWSNNSLEHGTNSNGEDATKSIQELGEMWLRLAQGLRKVCGEPREEVRNHAILSLQSSLCAAES
MNLSPSMWRQCFDQVIFVMLDDLLEIALGHSPKDYRNMEYTLQLAMKSVSKIFLKFLEQLVSLPSFRTLWLDLLR
RMNIYRKAKLRGKDNEKIQEVILELLKSILLEMNVRGVLVERNIQDNNKLWELTWHHVQEISPTLQAELLPQLQC
GSEESTDTLSLS*

**>Salcu_s0267.g026961**

MKSKNIDSGAVSCVITAEIGTVLAMMRRNARLAGSYVADDDSWEHSLVHSLKSLRRKLFTWHQPWQEINPLLYLQ
PFLDVVRSDDTSAPITGVALSAIYNILSFEVFDLTTAHIADAMHGLVEGVTSCRFEVIDHGSEEVVLMKILQVLL
AVMKHEASKALSDRDLCLVVSTCFRVVQQATTKGELLQRTARHSMHEIVRVIFAHLPEIEPASIYISRDGSATNS
NINNVGFDAVIRLENVAHAKGCSRNNSSEVVTEASNLEVTDEMPPSESKDVAEPERIPQKPVEPYGIVGIKEILR
DLCCLLDSGVTKGQAVRAIDEDVPLFSLQLIKTAIELGGRSFAFHPELLAVIQDDLFQSIMQLVLANKNPLILSM
VCSVVLNLYYHLRSHLKLQMEAFFSCVIIRLAQGKHGATYQQQEVAMEALVDFCRQPSFMSELYVNFDCDIACSN
LFEDLASLLSKSAFPVNVPLSAMHVLSLEGLVTVIQCMVDRLSHVQKVTPPSVDMPKSEFVPFWTMSCENYKDPN
SWVDFVQRQKLIKQKLMIGADKFNRDQKKGFEFLQGNHLLPKNLSAETVAFFFRHTTGLDKEVIGDFLGDPGEFS
LQVLEEFVNTFDFKDMNLDVAMRVFLSTFRLPGEAQKIHRILEAFSARYFEQSSHVFVDKDAAFVLAYSLILLNT
DQHNSQVKKKMSEEDFIRNNRKINGGQDLPREYLSNLYRSIVKNEIKVSYESGGSAEMSASAWVGLIHKSKKMPP
YLVCDSRAFLDQDMFSIISNQAVAAVSVVFDHAEDEEVLALCEQGFIGVAKLSANYHLLKMLDDLVVSLCNFTTL
LNPGGNTEEIIFSFAEDNRAHTAAQTVFKIATTYGDHIRSGWQSILDCILQIQKIGLLPISILNDSADEMDTSSE
MHGRSVPGNVSVVPTSNRKKSSGFIGRFSQLLYLEPDEPKVQPNEELLAAQQLALKVIEACRVDSIISESKFLQV
DSLLKLANALIQVSGKTQKMGGSFEDEDAAVFCLDLLIAVVLHNRDRILVLWQDVHEHISKIVQRANIPSPLVEK
AVFGLLRICQRLLPYKEELAEELLRSLQVILKLDARVADVLCERITQEVMQLVKANARHIRSPSGWWTVTSLLSI
TARHLEASEAGFETLSFVMLEGAHLSQANYVLFLDAAQAFAEATTGGLARSLNALELMSQSVKCLAKWSDTSLTD
QTSNSNAQELGEMWFKLAQCHKKLIVDPREDVRNHAVLSLQRIMSAAESMHLSASTLEQIFDQIVFVMLDDLLEI
ATKHSPKEFRNMEDTLLFAMKLVSKTFLQFLDPLTTLSVFPTMWSTVLKQMAKYMKTKLRGKGTDKLQESIPELL
KNMLLIMKNRGVLVQESTVDGYTLWDLTWRYAESIAPLLRTELFPEESMNKPEVAIEATPAPEVNIVL*

**>Azfi_s0087.g042331**

MKKLARSSGALACMVNSEIGAVLAVMRRNARWAGRYMGGDADQMEHSLVYSLKALRRSMFNWSQPWPKINPSLYL
TPFLDVIRSDETGAPITGVALSAIYKILVLEVFDPFTAHVAEAIHDIVDAVTSCRFEVTDLASEEVVLMKILQVL
VAIMKHSASDLLSDHDVCLVVNTCFRVVHQAVSKGELLQRTARHSMHEIVRTIFAHLPDLDSSLKLLMSSGCGSH
ESDNVNTTHIEDIDGIRNVSTNGHGSDFTPNSDGASARSSGDLHEGREENGPVSESSLGNKVHSVIEPYGIVGIL
EIFRYLCSLLNIVDSGIISGQSFLAIDEDVPLFALGLINSAVELGGSSFGCHPELIALVQDDLFRFLMQLGLSTI
NPLILSMVCSIVLNLYHHLRSHLKLQFEAFFSCVIIRLAQGKYGATYQQQEVAMEALVDFCRQSTFMPEMYANFD
CDITCGNIFEDLGNLLSKSAFPVNCPLSAMHILALEGLIAVIHSMAERASSTAPIALPLQQIEGQDYAPFWRLKC
ENYKDAKCWVNFVQRQKYIKRRLMIGADHFNRDPKKGLEFLQGIRLLPEKLDPESVACFFRFTTGLDKSLLGDFL
GDPDEFCLQVLAEFAKTFDFEGMALDAAMRTFLETFRLPGEAQKIQRVLEAFAGRYYDQSSHMFADKDAAFLLSY
SLIMLNTDQHNSQVKKKMTEEDFIKNNRSINGGKDMPRDFLSELYHSIIRNEIKITYDNSVGVTEMTPSRWVGLI
HRSQKTSPYIKCDARPFLDRDMFAIISGPTIAAISVVFDHSEDDEDVLRMCVEGFLAVAKISSSHHLEDVLDDLV
VSLCKFMTVLNPASVEEPVIAFGDDTKARTAAVTVFKIANKYGDYIRTSWRNIVDCILRLHKLGLLPTRIANDDG
ETGSAIVQGRPISGGAISIALPPSGAKKRSQGLMGRFSQLLSLETDEPRSQPSEQQLAAHQRTMRAIEACHIDSI
ISESKFLKAESLLQLVRALIWVAGRPQKVGGSSEDEDTANFCLDLLIAVALNNRDRIMLLWPGVYEHISNIVQST
LIPCLLVEKAVFGLLRICQRLLPYKEDLAEELLRSLQLILKLDARVADAFCERITREVMQLVKKNAQHIRSPMGW
RTVSALLSITARHVEASETGFEALSFIMSDGAHLTPANYVLCLDAARAFAEVSVGGNERSIQALNLMSESTKCLI
RWSDASTENGPTNPREKSLQELGEMWLRLAHGLRKVCGEPREEVRNYALLALQRSLSAAENMYLLPSVWGQCFDQ
VVFVLLDDLLEISLRHSPKEYRNMENTLHSATKLVSQIFLQFLDQLTLLPSFRALWVAILSRMKRYREAKLRDND
NQKIEELILDVVKNMLVAMKARGVLVQKSSIGGDSLWELTWHHVYDISPSLRAELLESGSGEKVETQILKE*

**>Azfi_s0094.g043397**

MRKKNVDSGAVACMINAEIGVVLAVMRRNARWTGSYVADDDQWEHSLVHSLKSLRRKIFAWNQPWKDIDPHLYIQ
PFLDVIRSDETSAPITGVALSAIYNILSYEAFDLTTSHIAEAMHGLVEAITSCRFEVIDPASEEVVLMKILQVLL
AIMKHEGSKVLNDKDLCLVVNTCFRVVHQASTKGELLQRTARHSMHEIVRVIFTRLPELQPPLKNLSGNGSSTPS
NNNNIVGFGGAEDPSHSNGDSRNNKLEVSNQVSAPSETLDERIPSSTREDVREPQHRVQISIEPYGIVGIGEIFR
YLCSLLDSGVITGQRIRAIDEDVPLFALGLINTAVELGGNSFSLHTELLAIIQENLFESIMQLALPSKNPLILSM
VCSVVLNLYHHLRSHLKLQLESFFSCIIIRLAQGKHGASYQQQEVAMEALVDFCRQPSFMYEMYVNFDCDISCSN
LFEDLISLLSKSAFPVNSPLSAMHVLSLEGLVAIIQSMVDRLGSIQKPPPPCVDIPISEYVPFWTAICENYEDPK
CWVDFVQRQKFIKHKLMIGADHFSRDQKKGFEFLQGIHLLPEKLDASSVAGFFRYTTGLDKGLLGDYLGDPEEFC

LQVLEEFTKTFDFKGMSLDMAMRVFLATFRLPGEAQKIHRILEAFAARYYEQSSQVLADKDAAFVLAYSLILLNT
DQHNSQVKKKMSEEDFIRNNRKINGGQDLPREYLSDLYHSILKNEIKVSYASGGATNGGTAEMNPSAWVGLVYNS
KKMSPYLICDSRAFLDQDMFSIISGPTVAAVSVIFDHAEDEEVLNLCEQGFFGVAKISASYHLVEVLDDLVVSLC
NFTTLPNSSGSVDELLLCFGEDAKACMAAKTVFKIVNTYGDHIRSSWRNVVDCILQLQKIGLIPSRILNDNADDV
EMSSEILGRTVIGNALSTQTRKRPSGLMGRFSQLLYLEPDEPKTQPSEEQLAAHQEALKVIEACHIDTIISESKF
LQTESLLQLAKALIRVSGRPQKAGGSFEDEDTSVFCLDLLIAVVLQNRDRILLLWHGVHEHISNIVQKASVPSPL
VEKAVFGLLRICQRLLPYKEELAEELLRSLQVILKLDARVADGLCERITHEVMQLVKANARHIKSPSGWKTVASL
LSITARHLEASEPGYETLCFVMFEGAHLTEANYLLFIEAARAFAEASTGGLERSIRSLDLMSESVKSLAKWSNIS
LTNETSNQNQEIGEMWLKLAQSQRRLIVDPREDVRNHAVLSLQRILSGAESMHLSASTLGQVFDHVVFVMLDDLL
EIATKYSPKDYRNMEKTMLFSMKLVSKVFLQFLDQLSSSPSFPATWSTLLKHMEKYMKTKLRGKGSDKLEESIPE
LLKNMLLVMKTRGVMTQECELWELTWRNVQGIAPFLKEELFPQESANKTEVSTDATSAQEANNVVL*
**>Selmo_101122**
MANSRSAARRRRESGALACMISAEVGAVLATMRRNSRWAGRYEEQLDHSLIYSLKLLRRSIFSWTKKPWNSINPC
LYLAPFLDVVRSDETGAPITGTALSAVYKILTSDVFDLRTSHVDEAMHAIVESVTSCRFEVTDPASEEAVLMKIL
QVLLACIGGDMGAVLGHRDVCNVVNTTFRVVHQAGNKSELLQRVARHTMHELVRAIFGHLSSMDPLAGNNGLLVP
WSNVENNVGVVSSSIVGVDYLGNGNAELATGLLVKVSEEGKLRNGEEEQRTIEPYGVPCMVEIFHFLCSLLNMVG
PYGLGESLGGTDEDVPQFALALINSAIEFGGPSFGQHQRLISLVRDELFRNLMQMGLSSNPIILSMVCSIVLNLY
HHLRSYIKLQLEAFFSFIIMRLAQGKFGASYQQQEVAMETLVDFCRQPNFMPEMYANFDCDITCSNTFEELANLL
SKSAFPVNCPLSAMHVLALEGLLAIIHSMADRVDSVPGSPLEPPTFLEIQAYVPFWNMRCDDYKEPSSWVEFVRK
QKYIKRRLMIGADHFNRDPKKGLEFLQGIHLLPEKLDPQSVASFFRYTTGLDKNLLGDFLGDRDDFCLQVLAEFT
RTFDFSSTSLDAALRTFLESFRLPGEAQKIERVMDAFSQRYCEQCVNVFANKDAAFVLAYSVIMLNTDQHKPQVK
KKMSEDDFIRNLRAANGGVDFPREMLSELYQSVAKNEIRISYDLGAGIPEMTHSRWLDLIRRSRKTSPYIVCDSR
PFLDHDMFAIMSGPTIAAISVVFDHAEDEEVLRGCLDGFLAVAKISASHHLEDILDDLVVSLCKFTTLLNPTASM
EEPVLAFGEDSKARMAAVTVFTIANKYGDFIRNGWRNILDCILRLHKLGLLPSRVTSDAADDTDAASESQGKMIA
GGLSVPPLVSAGVRRRSSGLMSRFSQLLSLEADEPRSQPSEQQVAAHQRILQTIEACHIDNIFSESKFLQAESLL
HLAKALIWAAGRPQKGTSSPEDEDTAVFCLELLIAITLNNRDRILLLWQGVYEHMAGIVQTTVVPCLLVEKAVFG
LLRICQRLLPYKENLAEELLRSLQLILKLDARVADAFCERITQEVMQLVRANAGHIKSPMGWRTISSLLSITARH
PDASESGFEALYFIMHDGAHLTPANYVLCLDAARAFAETRVGGVERSLKALDLLSESVNSLIKWSQVATGESDEN
KEDSVRASQELAEMWIRLAQGLRRVCGEQREEVRNCAILSLQRCLTAAESLHLPAMVWIQCFDQVIFVMLDDLLD
IALRNSPKEYRGMEGTLHHAMKFLSKIFLQFLDQLALLPNFRALWLAVLSRMEMYMKARLRGKGSDKLQELVPEL
LRNMLLIMHARGVLVQGSTLGGDSLWELTWHHVQTISPALQPDIFPDEKASSTPSPAAVTPVVSQAVDDPAEQQP
AEQVE*
**>Sphfalx0165s0016**
MGRSSVVSPIGSSGGGGDGSSMEYGLVTRGAVGAYKMRWSGGALACMVNAEVSSVLAVMRRNARWAGRYMAGDDQ
LEHPLIRSLKGLRRRVFGWQREAQWQSINPALYLVPFLDVIRSDETGAQITGVALAAVYKILTLEVFDINTAHVD
VAMHSIVDAVTSCRFEVTDPASEEFVLMKILQVLLACMKSVAGALLSNRDVCTVVNTCFRVVHQAGSKGELLQRT
ARHTMHELVRAIFSHLSSLSPATSTAPFPSLPTTSSELLPSNGSIYATPSLPVKPATGLAIQLTASPELDDNVTG
NGSSRMSEPEGNETATEKGIPISGYTAGEGLTAKRPPQPVTRDGNSEKEGTADYKGAMTTDQPIAIEPYGVPCMV
DIFNFLCSLLNVGDPLGPGQSFLASDEDVPHFALMLINSSIELGGESFGRHPKLLALVQDELFRNLMQMGLSENP
LVLSLVCGIVLNLYHNLRESIKLQLEAFFSFVVIRLAQGRSGASYQQQEVAMEALVDFCRQSTFMPEMYANFDCD
ITLSNTFEDLGNLLTKSAFPVNCPLSAMHVLALEGLLAVIHSMANRVDSAAPGVLPGTPVGVMVAENQEYVPFWT
LKCENYKDPLCWVEFVQHQKYIKRRLMIGADHFNRDPKKGLEFLQGIRLLPEKLDPQSVACFFRYTTGLDKNLLG
DFLGDPDDFCLHVLEEFARTFDFSEMGIDAALRVFLESFRLPGEAQKISRVLEAFAERYYQQSRGILASKDTVYI
LSYAVIMLNTDQHNPQVRKKMTEDDFIKNLRNINNGQDLPRVMLSDLYHAIVRNEIRISYDNGIGVAEMTHSRWV
DLMRRSRSTTPYITCDSRVLLDHDMFAIISGPSIAAISVVFDHAEDEEVLRACIDGFLAVAKISAVHRLEDVLDD
LVVSLCKFTTLLNPTASAEEPVVAFGDDTKARMAAVTVFSIANKFGDFIRTGWRNILDCILRLHKLGLLPARVAS
DATEDADNASDSLQGRLGEGSVTMAPAPAMGTRRRSSGLMSRFSQLLSLESDEPRSAPTEQKLAAHQRTLRTIEA
CHIDQIFNDSKFLHADSLLQLARALVWAAGRPQKGGSSIEDEDTAVFCLELLIAITLNNRDRIMLLWQGVYEHMA
GIVQTTVLPGLLVEKAVFGLLRICQRLLPYKENLAEELLRSLQLILKLDARVADAFCERITQEVMVLVRANAAHI
KSPMGWRTVSSLLSITARHPEASEPGFDALSFIMADGVHLTPANYVLCLDAARAFADARVGGNERSIRALDLLSD
SVGCLTRWSKATSESVEALGSETLEGAPRYSQELAEMWLRLAQGLKRVCLDQREEVRNHAIMCLQRCLAAAESIP
LSATMWAQVFEQVVIALLDELLDIALRHPPKEYRGMEGTLYDAIKFLSKFYLQFLGQLVQLPTFRTLWSQVLSRM
EMYMKAKLRGRGSEKIQELIPELLRNMLQVMHATGILVKTSTVTSDSLWELTWQHVYSISTSLQPNMFSEQTSIL
KGNEQLPVDGNQILHGLVVAGVESGSNPLATGSNSSVGTGDTRHEEKQPLERQ*
**>Cepur_8G135600**
MGRSAHNPSESAMEGVGSGRAMARRSGGALACMVNAEVSAVLAAMRRNVRWQGRYMASDDQMEHPIIRSLKSLRR
RVFSWQQQQSWQSINPSFYLSPFLDVIRSDETGAAITGIALSAVYKIVTLEVFNVKTAHVDVAMHMIVDAVTSCR
FEVTDPASEEVVLMKILQVLLACLKCGAGAVLSNRDVCNVLNTTFRVVHQAGSKGELLQRTARHTMHELVRAVFS
HLSSLEATPALITLPSNSLEAASSNGAVYDKSESSVGNQVDGNGDLSHFEGLETLSGNASEEGSSRLSYSGKREE
ANSRSVVVTDQSTAIEPYGIPCMVEIFNFLCSLLNIADPQSPGQALLASDEDVPHFALLLINSSIELGGESFSRH
PKLLALVQDELFRNLMLMGLSPNALVLSMVCGIVLNLYHHLRSAVKLQLEAFFSYILIRLASGKYGASHQQQEVA
MEALVDFCRQPTFMPEMYANFDCDITLCNTFEDLGNLLSKSAFPVNCPLSAMHVLALEGILAVVHSMADRVDSAA
PALTSSTLSGAMVAENQDYVPFWTLKCENYQDPGSWIEYVKHQKYIKRRLMIGADHFNRDPKKGLEFLQGIRLLP

AKLDPKSVACFFRYTTGLNKDLLGDFLGDPDEFCLQVLEEFARTFDFSDMQIDGALRTFLESFRLPGEAQKIHRV
LEAFSDRFYQQSKGILANKDAAFVLSYSVIMLNTDQHNVQVRKKMTEEDFIKNLRNINDGQDLPRPMLSELYHNI
VRSEIRITYDSGAGVSEMTHSRWVDLMRRSMTTTPYITCDSRPLLDHDMFAIISGPTIAAISVVFDHAEDEEVLR
SCVEGFLAVAKISASHRLQDVLDDLVVSLCKFTTLLNPTASAEEPVVAFGDDTKARMAAITVFSIANKFGDFIRT
AWRNILDCILRLHKLGLLPSRIASDAAEDTDLNGDSAQGKLAGGALGMASMPLSGNRRRSTGLMSRFSQLLSLDA
DEPRSAPTEHQLAAQQRTLRTIESCHIDQIFTDSKFLQSESLLQLARALVWAAGRPQKNGGSSEDEDTAVFCLEL
LFAITLNNRDRIMLLWQGVYEHMAGIVQTTVVPGLLVEKAVFGLLRICQRLLPYKEELAEELLRSLQLILKLDAR
VADAFCERITQEVMVLVRANAGHIKSPMGWRTVSSLLSITARHPEASDPGFEALTFIMQDGAHLTPANYVLCLDA
ARAFAEARVGGSERSIRALDLLSDSVGCLTRWSKAKSASTVVSSEVVEGFPRFSQELAEMWLRLAQGLRRVCLEQ
REEVRNYAIVCLQRCLGAADSITLTPTMWVQSFEQVVLTLLDELLDIAVRYPPKEFRGMEITLMHALKFLSKFYL
QFLDHLAQLPTFRTIWSQVLNRIEMYMKAKLRSRNTEKLQELVPELLQEILQVMYSRKFLVKTDSPTGDSLWELT
WQAVQNISPALKPDILKEPEPLADSKVDPDESPKAEPVAELDTADSLPAGDQGTEEHKPSEEIQ*

**>Cepur_7G143500**

MGRSLEMEGGGGEGGRAGSRRKRSRGALACMVNSEVSAVLAVMRRNARWAGRYSASDDQLEHPLIRSLKGLRRLV
FSWRQPWERIDPCLYLSPFLDVIRSDETGAQITGVALSAVYKMVHLEIFDAETANVDIAMHTIVDSVTSCRFEVT
DPASEEVVLMKILQVLLACMKSGAGAVLSDRDVCIIVNTTFRVVHQAGSKGELLQRTARYTMHELVRVVFSHLPS
LEPTTLATGVTPSSMEISSNSQNAIDSSGAEETDHLGGPVHTVNTPLGDSSSDPAEAERVVINSDEPSENDTVVS
SSIDGRPEGQIAAIETTITAEDSKIEAYGIPCMVEIFSFLCSLLNIEDPQSPGQLVLAGDEDSPQYALMLINSAI
EIGGEALSYHPKLLVLIQDELFRNLMVIGLSQNPLVLSLVFGIVLNLYHHLRRQLKLQLEAFFSFVLIRLASGKH
GATYQQQEVALEALVDLCRQPTFMPEMYSNFDCDISLSNCFEDLGNLLSKSAFPVSGPLSAMHVLALEGLLAVVR
SMADRVDSAAPALASSSSSKNMIIDNQEYVPFWTLKCENYDDPGSWVQFVKHQKYIKARLLLGADHFNRDPKKGL
EFLQGMKLLPPVLDPKSVACFIRYSSGLNKTVIGDYFGDPDDFCLRVLDEFTKTFDFNDMGIDAALRVFLESFRL
PGEAQKIHRVVEAFAERYYEQSKGTLASKDVAFVLSYSVIMLNTDQHNVQVRKKMTEDDFIRNLRKINDGQDLPR
PMLSELFHSIVRNEIRISYNAGAGVADMTHSGWIDLMRRSTSTTPYIICDEQPLLDDEMFPILSGPSIAAISVVF
DHAEDEEVLQLCVDGFLSVAKISASHRLEDVLDDLVVSLCKFTTLLNPFSSDEEPIVAFGDDTRARMATFAVFTI
ANKYGDFIRTGWRNILDCILRLHKLGLLPSRVVNESTEDTDAAGDSAQSKLAGSSSTTAPVPFIKHQRRNTGLMS
RFSQLLSLESEEPPAVPTEEELAAHQRAVRTVESCHIDQIFTDSKFLQAESLLQLARALVWAAGRPHKSGGSTED
EDVAVFCLELLITITLNNRDRIMLLWQGVYEHMAGIIQTAVVPGLLVEKAVFGLLRVCQRLLPYKENLAEELLRS
LQLILKLDARVADTFCERITQEVMVLVRANAAHIKSPMGWRTVTALLSITARHPEASEPGFDALSYIMHDGAHLT
PANYILCVDAARAFAEARVGGPGRSVRALDLLSDSVSCLTRWSKVHSESGDASSSENLEGLSRYSQELAEMWLRL
AQGLRKLCLEQREDVRNHAILCLQRCLSAAEIISLTPAMWTQSFEQVVTLLDELLDLAVRHPKEFKGMDNTLVLI
MKCLSKTYLQFLSQLVKLPAFQTLWFQVLNRMEMYIKAKFRGKNSEKLQELIPELLRNVLQVMHSQGILAQPTTP
SDKSLWDLTWQHVVNIVPTLKPDIFKEPKSEAKDGESAGPQVQASPGSPQTAEVESEYTLVSVPTSRAGDEG*

**>Cepur_9G122800**

MGRSLEMEGGGGEGGRAGSRRKRSRGALACMVNSEVSAVLAVMRRNARWAGRYSASDDQLEHPLIRSLKGLRRLV
FSWRQPWERIDPCLYLSPFLDVIRSDETGAQITGVALSAVYKMVHLEIFDAETANVDIAMHTIVDSVTSCRFEVT
DPASEEVVLMKILQVLLACMKSGAGAVLSDRDVCIIVNTTFRVVHQAGSKGELLQRTARYTMHELVRAVFSHLPS
LEPTTLATGVTPTSMEISSNSENATDSSGAEETDHLGGPVHIVNTPLGDSSSDPAETKRVVINSDEPSENDTVVS
SSIDGRPEGQIAAIETTITAEDSKIEAYGIPCMVEIFSFLCSLLNIEDPQSPGQLVLAGDEDSPQYALMLINSAI
EIGGEALSYHPKLLVLIQDELFRNLMVIGLSQNPLVLSLVFGIVLNLYHHLRRQLKPQLEAFFSFVLIRLASGKH
GATYQQQEVALEALVDLCRQPTFMPEMYSNFDCDISLSNCFEDLGNLLSKSAFPVSGPLSAMHVLALEGLLAVVR
SMADRVDSAAPALASSSSSKNMIIDNQEYVPFWTLKCENYDDPGSWVQFVKHQKYIKARLLLGADHFNRDPKKGL
EFLQGMKLLPPVLDPKSVACFIRYSSGLNKTVIGDYFGDPDDFCLRVLDEFTKTFDFNDMGIDAALRVFLESFRL
PGEAQKIHRVVEAFAERYYEQSKGTLASKDVAFMLSYSVILLNTDQHNVQVRKKMTEDDFIRNLRKINDGQDLPR
PMLSELFHSIVRNEIRISYNAGAGVADMTHSGWIDLMRRSTSTTPYIICDEQPLLDDEMFPILSGPSIAAISVVF
DHAEDEEVLQLCVDGFLSVAKISASHRLEDVLDDLVVSLCKFTTLLNPFSSDEEPIVAFGDDTRARMATFAVFTI
ANKYGDFIRTGWRNILDCILRLHKLGLLPSRVVNESTEDISGDSAQSKLAGSSSTTAPVPFIKHQRRNTGLMSRF
SQLLSLESEEPPAVPTEEELAAHQRAVRTVESCHIDQIFTDSKFLQAESLLQLARALVWAAGRPHKSGGSTEDED
VAVFCLELLITITLNNRDHIMLLWQGVYEHMAGIIQTAVVPGLLVEKAVFGLLRVCQRLLPYKENLAEELLRSLQ
LILKLDARVAETFCERITQEVMVLVRANAAHIKSPMGWRTVTDLLSITARHPEASEPGFDALSYIMHDGAHLTPA
NYILCVDAARAFAEARVGGPGRSVRALDLLSDSVSCLTRWSKVHSESGDASSSENLEGLSRYSQELAEMWLRLAQ
GLRKLCLEQREDVRNHAILCLQRCLSAAEIISLTPAMWIQSFEQVVTLLDELLDLAVRHPKEFKGMDNTLVLIMK
CLSKTYLQFLSQLVKLPAFQTLWFQVLNRMEMYIKAKFRGKHSEKLQELIPELLRNVLQVMHAQGILAQPTTPSD
KSLWDLTWQHVVNIVPTLKPDIFKEPKLEAKDGESAGPQVQASPGSPQTAEVESEYTLVSVPTSRAGDEG*

**>Pp3c22_6150**

MGRSTQNPSTSMEGSAGGEGWGKTRLNGGAVACMMNVEVSAVLAVMRRNARWAGRYMASDDQLEHPLIRSLKSLR
RRIFTWHLQQPWHTIKPALYLSPFLDVIRSDETGAQITGIALSAVYKIITLQVFDVHTMQVDVAMHMIVDAVTSC
RFEVTDPASEEVVLMKILQVLLACMRSGAGAVLSNRDVCNVLNTSFRVVHQAGSKGELLLRTARHTMHELVRAVF
SHLPSLNVTPALIALPSNNPEAASSNGAVYDASETNVDSEVAGNGELADYEGVETFSEDGSRGFPYSGSNGQSRS
VMVTDQISAIEPYGIPCMVEIFSFLCSLLNIADPQSPGQAVLASDEDVPHFALLLINSAIELGGESFSRHPKLLA
LVQDELFRNLMLMGLSPNPLVLSMVCGIVLNLYHHLRTALKLQLEAFFSFILIRLASGNYGATHQQQEVAMEAIV
DFCRQPTFMPEMYANFDCDITLSNTFEDLGNLLSKSAFPVNCPLSAMHVLALEGILAVVHSMADRVDSGASALTS
STLSMVAENQEYVPFWTLKCENYEDPVSWVDHVKHQKYVKRRLMIGADHFNRDPKKGLEFLQGIRLLPAKLDPKS

VACFFRYTTDLNKDLLGDFLGDPDDFCLKVLEEFAGTFNFSEMGIDGALRTFLESFRLPGEAQKIHRVLEAFSER
YYHQSKGIFANKDAAFVLSYSVIMLNTDQHNVQVKKKMTEEDFIKNLRSINDGQDLPRKMLSELYHSIVRSEIKI
SYDGGTGVSELTHSRWVDLMRRSITTTPYITCDSRPLLDHDMFAIISGPTIAAISVVFDHADDEEVLRSCVEGFL
AVAKICASHRLQDVLDDLVVSLCKFTTLLNPLASAEEPVVAFGDDTKARMAAITVFSIANKFGDFIRTGWRNILD
CILRLHKLGLLPSRVPSDPVEDSDLVGDSVQGKLAGSTSGMASMPVTGNRRRSTGLMSRFSQLLSLDADEPRFAP
TEHQLAAQQRTLRTIESCHIDQIFTDSKFLQAESLQQLARALVWAAGRPQKNGGSSEDEDTAVFCLELLFAITLN
NRDRIMLLWQGVYDHMAGIVQTTVVPGLLVEKAVFGLLRICQRLLPYKEDLAEELLRSLQLILKLDARVADAFCE
RITQEVMVLVRENSGHIKSPMGWRTVSSLLSITARHPEASDPGFEALSFIMQDGAHLTPANYVLCLDAARAFAEA
RVGGIERSIRALDLLSDSVGCLKRWSKAKSASTGLSTSEVVEGSSRFSQELAEMWLRLAQGLRRVCLEQREEVRN
YAIICLQRCLAAAGSIALTPTMWIQSFEQVVLTLMDELLDIAVRYPPKEYRGMESTLIHAVKFLSNFYLQFLDQF
AQLPTFRTTWTQVLNRMEMYMNAKLRSRNTEKLQELVPELLRNILKVMHGRKILVQSSTSSGDSFWELTWQLMRS
ISPSLEPDLLNEPISEEGRDAPADLNGSTVAEPGTDAASLPSEDQGTEEHESFEETQ*

**>Pp3c15_11320**

MSRSIGTPGLEMEGEGREQARRSRVSLACMVNYEVSAVLAVMRRNARWAGRFSASDDQLEHPLIRSLKGLRRLVF
TWRQPWPSIDPCLYLSPFLDVIRSDETGAQITGVALSAVYKMVHLNFFDLNTAHVDIAMHTVVDSVTSCRFEVTD
PASEEVVLMKILQVLLACMKSEAGALLSDRDVCNIVNTTFRVVHQAGSKGELLQRTARFTMHELVRAVFSHLPTL
KPTNLTIGVTPLSTEISSNPGNVADQSGSEEGNSFSGPVQTDENPFRDGSLDVADAGRALKISDKSSEEDTIVLP
PLDGKDGGGHAAIGTVGSAEDSNFAAYGVPCMVEIFSFLCSLLNIADPQGLGQLVLASDEDSPQFALMLINSALE
LGGEAFRNHPKLLALIQDELFRNLMEIGLSQNPLVLSLVFGVVLNLYHHLRVLMKLQLEAFFSFVLIRLASGKYG
ATYQQQEVALEALVDFCRQPTFMPEMYANFDCDTSLSNTFEDLVNLLSKNAFPVNCPLSAMHVLALEGLLAVAQS
MADRVDTAVPAFASSTSPSNLAGDNPEYVPFWTLKCENYDDPLSWVQFVKHQKYIKGRLMVGADHFNRDPKKGLE
FLQGMQLLPSEPDPKSLACFIRYCTGLNKSVIGDYLGNPDEFCLRVLDEFAQTFDFSNMGIDAALRVFLEGFRLP
GEAQKIHRIVEAFADRYYQQSKGILASKDAAFVLSYSVIMLNTDQHNKQVRKKMTEDDFIKNLRKINDGQDLPRS
MLSELYHSIVRDEIRISYDSGAGVAEMTHSRWIDLIRRSMITTPYITCDERPLLDYDMFPVLSGPSIAAISVVFD
HAEDEEVMQLCIDGFLAAAKISASHRLEDVVDDLVVSLCKFTTLLNPFSSDEEPVIAFGDDKKARMATVAVFDIA
NKYGDFIRTGWRNILDCILRLQKLGLLPARVANESVEDTDARGDSTSSKLAGSSSTVAPMPDLIRHRRRNTGLMS
RFSQLLSLESDEPPSAPTEEELAAQQRALQCVESCRIDQIFTDSKFLQAESLLQLARAFVWAAGRPHRSGSSTED
EDTAVFCLELLITVTLNNRDRIMLLWQGVYEHMAGIIQTSVFPGLLVEKAVFGLLGVCQRLLPYKEDLAEELLRS
LQLILKLDARVADAFCERITQEVMVLVRANAAHIKSTIGWRTVTSLLSITARHPEASEPGFEALTYVMQDGAHLS
PANYVLCVDAARAFAEARVGGPGRSVRALDLLSDSVGCLTTWSKVHSESADASSGENVEEPSRYTQELTEMWLRL
AQGLRKLCLEQREEVRNHAILCLQGCLSAAEILNLTPVLWAQSFKQVVLNLLDELLELAVRHSKEYKGMEDTLVL
SIKFLSKSYLQFLPQLVNLPSFESSWFQVLNHMEIFIKTKFRGKRSEKLQELIPELLRNILQVMHSQGILVQPTS
PSQDSMWDLTWKHVGSIVPTLKPDILKENRPEVKDGESAVSQEQTPSASIPCDVESEYTIVEKVDSVSTGDQG*

**>Pp3c9_11830**

MSRSVGTPRAEMEGGWRDKARRSKGTLACMVNSEVSAVLAVMRRNARWAGRYSASDDQLEHPLIRSLKGLRRLVF
TWQQPWPSIDPCLYLSPFLDVIRSDETGAQITGVALSAVYKMVQLEIFDVQTANVDIAMHKIVDSVTSCRFEVTD
PASEEVVLMKILQVLLVSLKSGVGAVLSDRDVCNIVNTTFRVVHQAGSKGELLQRTARYTMHELVRAVFSHLPSL
KPTSLAIEIAPHNTESSSSAGKDLNKSGSEEANELSGLVNTSSISSADGSSDLAEAEQPVLVSDEPSEKDAVAHL
LNGSDGSQNAAIETVGSAEDSIVAAYGVPCMVEIFSFLCSLLNIADLLSPGQLVLASDEDSPQFALMLINSALEL
GGDAFKKHPKLLDLIQDTLFRNLMVIGLSQNPIVLSLVFGIVLNLYHHLRGLVKLQLEAFFSFVLIRLASGKHGA
TYQQQEVALEALVDFCRQPTFMPEMYANFDCDTTLSNTFEDLGNLLSKSAFPVNCPLSAMHVLALEGLLAVVRSM
ADRIDTGILVLASSNLGAGNQEYIPFWTLKCEHYDDPSSWVQFVKHQKYIKCRLMIGADHFNRDPKKGLEFLQGM
RLLPSELDPKSVACFIRYSTGLNKTVIGDYLGDPDEFCLRVLDEFAQTFDFSNMRIDSALRLFLESFRLPGEAQK
IHRVVEAFADRYYQQSKGILASKDVAFVLSYSVIMLNTDQHNKQVRNKMTEDDFIRNLRKINDGQDLPRQMLAEL
YHSIVHNEIRISYVSEAGVANMTHSRWIDVMRRSVSTTPYINCDERPLLDHDMFPIISGPSIAALSVVFDHAEDE
EVLQLCIDGFLAVAKLSASHRLEDVLDDLVVSLCKFTTLLNPFPLEEEPVIAFGGDTKARMATVAVFNIANKYGD
FIRTGWRNILDCILRLQKVGLLPAQVANESVEKTNTTGDSAHSKLAGSSSTRIQMPERVRHRRRNTGLMSRFSQL
LSLESDEPPAVPTEEELAAQQRALRTIESCHIHQIFTDSKFLQAESLLQLARALVWSAGRPHKSGGSAEDEDTAV
LCLELLITITLNNRDRIMLLWQGVYEHMAGIIQTSVFPGLLVEKAVFGLLRVCQRLLPYKEDLAEELLRSLQLVL
RLDPRVADAFCERITQEVMVLVRTNAAHIKSPMGWRTVTSLLTVTARHPRASEPGFEALMYIMQAGAHLTPANYV
LCVDAARAFVEARVGGPGRSVRALDLLLDSVGRLTLWSKVHSEGADASGTDSVEGLSRHSQELTEMWVRLAQLLM
KLCLEQRELVRDHAILCLQGCLSAAEMISLTPALLAQSLEQVVLKLLDELLDLAVRHPKEYQGMEDTLFLSTKFL
AKSYLQFLPQLVTLPTFQSLWLQVLKRMEMFINAKFRGKRSEKLQELIPELLRNILQVMHSQGILVQLTKPSQDN
MWDMTWKHVGDIVPTLTPDIIKESSPEVKDGGFAVVGEQMSPASVPSDVESEYTIVEKVDSISLGDQG*

**>AntanGNOMa_sc67**

MACGSLVSDLQPGGGILETSHSNTGVAATGDGGTLACMVNAEIGAVLAVMRRNARWGTRYMAGHDDQHEHSLMHA
LKMLRRRIFSWHQPWQTISPNIYLSPFLEVIRSDETGAPITGVALSAVYKILTLQVFNARTAHVGEAMHSIVDCV
TSCRFEVTDPASEEIVLMKILQVLLACVKTGAGAVLSNGDVLNVVNTCFRVVHQAGSKGELLQRTARHAMHELVR
TIFSRLPLVSVHSGTTPGELQTEGTVSGLQANGANVLGLSSTGSNDGPSLEDGTQKlppqlspsspglalsDGSN
FQDWGAVNKMDGHQGDAASRESYGVMCMVEIFHFLCSLLNVGIPQGPGQSVLASDEDVPLFALGLINSALELGGQ
SFGQHPKLLALVQDELCRNLIQMGLSRQPLVLSMACSIVLNLYHHLRCHVKLQLEAFFNWKLFLLRDNPSGTGRY
GATYQEQEVAMEALVDFCRQPRFMVEMYANFDCDITCSNLFEDLGNLLSKSAFPVNCPLSAMHVLALEGFAVIHS
MADRVDGASVPYSGPAIEEGQEYVPFWTVQCENYEDPQCWVQFVRRCKYIKRRLMIGADHFNRDPKKGLEFLQVI

RFLPEKLDPRSVACFFRYTSGLDKSLLGDFLGDPDEFCLDVLVEFARTFDFTDMSLDSAFLESWPPGEAQKISRV
LDAFAERYYEQSSGILANKDAAYVLAYSVIMLNTDQHNVRVKRKMSEEDFIKNNRSINGGEDLPREMLLDLYHSI
VRDEIKVSYDHGTASGDMTYSRWVDLMRRSRTTSPYIQCDMAQLLDNDMFAMISGPTIAAISVVFDHAEEEEVQQ
ACVDGFLAVAKIAASYRLEDVLDDLAVSLCKFTTLLNPTTSVEEPHCLXGDTKARMAALAVFTIANKFGDYIRGG
WRNILDCILRLHKLGLLASRVASDIADETDGMTSSAHGKVFPGNQALLPAPTGGARRKSSGLMSRFSQLLSLESE
EPRLQPTEQQLAAHQRTLRTIEACHIDNIFTDSRFLQADSLLELAKALIGAAGRSTKGTTPEDEDTAVFCLELLI
AITLNNRDRIMLLWQGVYEHMAGIVQTATTPTPLVEKAVFGLLHICQRLLPYKENLSEELLRSLQLILKLDARVA
DEFCERITQEVMQLVRANAGHIKSMAGWRTVASLLTMTARHPEASDPGFEALVFIMAEGAHMLPANYVLCLDAAR
AFAESRTGGNDSRSALDLLSESVTCLVQWRKVANSDANPAEEVGENGKVIQELADMWLRLAQGLRKVCLDHREEV
RNYALSALQRCIIAADDLNASPAVLAHCFDQIVFPLLDDLLDIVSRQSSKEFRSMEGTLYQAMKFISKSFLHFLD
HLSALPSFGTLWLAVLGHMEKYLKLRLRGKASDRLQEQVPEILKNMLLVMHTRGVLMQRSTLGGDSLLEQSWPQV
QAISPWLQDDMFPEVRAPLAAERSGADKHEVIVEQEVLTTEGAALEASPVAVEAAPEESAHVNSS*
**>AntanGNOMb_AANG383**
MACGSLVSDLQPGGGILETSHSNTGVAATGMVEHWLVW*MQRLGLCWL*CAGMRGGHSHMAGHDDQHEHSLMHAL
KMLRRRIFSWHQPWQTISPNIYLSPFLEVIRSDETGAGAPITGVALSAVYKILTLQVFNARTAHVGEAMHSIVDC
VTSCRFEVTDPASEEIVLMKILQVLLACVKTGAGAVLSNGDVLNVVNTCFRVVHQAGSKGELLQRTARHAMHELV
RTIFSRLPLVSVHSGTTPGELQTEGTVSGLQANGANVLGLSSTGSNDGPSLEDGTQKLPPQLSPSSPGLALSDGS
NFQDWGAVNKMDGHQGDAASRESYGVMCMVEIFHFLCSLLNVGDPMGPGQSVLASDEDVPLFALGLINSALELGG
QSFGQHPKLLALVQDELCRNLIQMGLSRQPLVLSMACSIVLNLYHHLRCHVKLQLEAFFSYVIIRLAQGRYGATY
QEQEVAMEALVDFCRQPRFMVEMYANFDCDITCSNLFEDLGNLLSKSAFPVNCPLSAMHVLALEGLLAVIHSMAD
RVDGASVPYSGPAIEEGQEYVPFWTVQCENYEDPQCWVQFVRRCKYIKRRLMIGADHFNRDPKKGLEFLQVIRFC
QRSLIRVACFFRYTSGLDKSLLGDFLGDPDEFCLDVLVEFARTFDFTDMSLDSALRTFLESFRLPGEAQKISRVL
DAFAERYYEQSSGILANKDAAYVLAYSVIMLNTDQHNVRVKRKMSEEDFIKNNRSINGGEDLPREMLLDLYHSIV
RDEIKVSYDHGTASGDMTYSRWVDLMRRSRTTSPYIQCDMAQLLDNDMFAMISGPTIAAISVVFDHAEEEEVQQA
CVDGFLAVAKIAASYRLEDVLDDLAVSLCKFTTLLNPTTSVEEPAIAFGGDTKARMAALAVFTIANKFGDYIRGG
WRNILDCILRLHKLGLLASRVASDIADETDGMTSSAHGKVFPGNQALLPAPTGGARKSSGLMSRFSQLLSLESEE
PRLQPTEQQLAAHQRTLRTIEACHIDNIFTDSRFLQADSLLELAKALIGAAGRSTKGTTPEDEDTAVFCLELLIA
ITLNNRDRIMLLWQGVYEHMAGIVQTATTPTPLWRRLFWSTSHLPTAASDKENLSEELLRSLQLILKLDAMLLDE
FCERITQEVMQLVRANAGHIKSMAGWRTVASLLTMTARHPEASDPGFEALVFIMAEGAHMLPANYVLCLDAARAF
AESRTGGNDRSIRALDLLSESVTCLVQWRKVANSDANPAEEVGENGKVIQELADMWLRLAQGLRKVCLDHREEVR
NYALSALQRCIIAADDLNASPAVLAHCFDQIVFPLLDDLLDIASRQSSKEFRSMEGTLYQAMKFISKSFLHFLDH
LSALPSFGTLWLAVLGHMEKYLKLRLRGKASDRLQEQVPEILKNMLLVMHTRGVLMQRSTLGGDSLLEQSWPQVQ
AISPWLQDDMFPEVRAPLAAERSGDKHEVIVEQEVLTTEG*
**>Marpo_0100s0023**
MGRSTQSQLGYSPSGEERDEATGRKGRTRTAGALACMVNAEVGAVLSVMRRNARWAGRYISGDEPMEHSLIRNLK
TLRRKIFTWQSPSSPGWQSVNPSFYLSPFLDVIRSDETGAPITGVALSAVYKILTLDIIDVKTVSVEGAIHGIVD
AVTSCRFEVTDPASEEVVLMKILQVLLACMKCGAGLVLSNRDVCVVVNTCFRVVHQAGSKGELLQRTARHTMHEL
VRTIFSHLPDLDPTPGAVNDVEATGNGSIEAAPHRLSTEDSSRSQGTNPSTSNGASGEDGYSEGFTREPERSEPD
SQAEDTLPRKPPRPPSREKMALVVSNKADQDKSHKDGSPALEPYGVPCMVEIFHFLCSLLNITDPTGPGHSLLAS
DEEVPLFALGLINTAIELGGQSFGYHPKLLALVQDELFRNLMQMGLSFTPPVLSMVCSIVLNLYHHLGTFLKLQI
EAFFVYVVIRLAQGKYGAAYQQQEVAMEALVDFCRQPTFMAEMYANFDCDIICSNAFEDIGNLLSKSAFPVNCPL
SAMHVLALEGLLAVIHSMADRIDSLPVTPQLEMVNYTDAPDFVPFWTLKCPSYEDPSCWVEFVRQQKYIKRRLMI
GADHFNRDPKKGLEFLQGIRLLPKNLDPKSVACFFRYTTGLDKRLLGDFLGDHDEFCVQVLQELAKTFDFSDMSL
DAALRTFLESFRLPGEAQKIQRVLEAFAQRYYEQSPVVFANEDAAFTLAYSIIILNTDLHNPQVKRKMTEEDFIR
NNRGINGKQDLPREMLAELYHSIARNEIRISYDNGAGVPEMTHSRWVDLMRRSRSSTPYIKCDSRPLLDHDMFAI
ISGPTIAAISVVFDHAEDEDVLRSCVDGFLAVAKISASHRLEDVLDDLVVSLCKFTTLLNPTASAEEPVVVFGDD
TKARMAAVTVFTIANKWGDYVRTGWRNILDCILRLHKLGLLPSRVASDATDDADAGGDSAQGRVMSGNLSLAVIP
SAGNRRRSTGLMSRFSQLLSLDVDEPRTQPTEQQLAAHQKTLRTIEACHIDNIFSDSKFLQAESLLQLARALIWA
AGRPHKGGSSPEDEDTAVFCLELLIAITLNNRDRIMLLWQMVYEHMAGIVQTSMVPGLLVEKAVFGLLRICQRLL
PYKEDLAEELLRSLQLLLKLDARVAEAFCERITQEMMQLVKANAGHIKSPMGWRTVSSLLASTVRHPEAFGPGFE
TLSFIMSDGAHLTPANYVLCLDAARAFAEARVGGVERSIRALDLLAESVNCLIQWAATCSDSTKGAGGEVFKPET
KDDAIVDTQALGEMWLRLAQVLRRVCGEQREDVRNHAILSLQRCLLSVENIHLPPTIWHPCFEQVIFVMLDDLLD
VALRHPPKEYRGMEGTLLHAMKFLSKIFLQYLSQLSQLPSFRPTWFNILSRMEMYMNARLKGKGSDKLQEVIPEL
LKNMLLIMHNKGVLVQRSTLSGDSLWELTWHHVEAIAPALQSELFPEQEGDPEAESSIPLLGRGDAPATPVGEQD
AEKVKEQILAEKQ*
**>Chabr_g19322**
MMGPRLGEGGGEGRWRDGGGFGSMKLECAGSVACMVNAEIQAVVAVMRQNTRWAATRFSEDDSADDPLLLSFKLL
RKEIFSWQDWNQVSPPRYLIAFLEVIRSPETSAPITSVALSAVYKILSIEIFDANSLDAAGAMHLIVDAVTGCRF
EVTDPASEEVVLIKILQVLLSCLKCRAGALITERDVCNVINTCFRVVHQSVTKGELLQRMSCHIMQEMIRAIFGR
LAYLPEVDGRGISGGSAKGSVSFGRGGERERMARRVDGAAVYRVKKGTRNVESGPGGGGVASERGEEGGGARGEE
NGTVSHGDGGAGGAVGESFGVAGRKEGGAAGGGPAAAAAAAAGGGGGEGGVGGGGFVWSGGDTPVGAEPSAEGGD
GVGGGDGGSERVEEQSSGKMEVPPPYGVSCLVEVFQFLCSLVSLNDGELRTLGPPGSQMAALHDPEDMSLFGLAL
INSALEVGGSNFARHPRLLALVQDELFRNLMQMGLSQNLSLLSLVCGIILSLYRHLRRHLKLQLEAAFEFVLIPL

AQGKLCSYEQQEVAIEALVDLCRQPTFMPDLYANMDCDIMCENTFEDLSNLLSKNAFPVNSPLSAVNVLSLEGLL
AVVHAIARRTAMAAGADSNGGPLDASYLSSIYSTSSSSSVSSLSRVALAGAPECAVSASLSTVSSPTRTGGSKGG
RGGGDGGEVKEYTQFWKDRCADCEDAGVWVEFLRRQKFIKRRLMIGADHFNRDPKKGLEFLQGQRLLPAELDPKS
VACFFRYTPGLDKNLLGEYFGDRDDFNIKVLHAYVSEFDFTDMSLDTALRTFLESFRLPGEAQKIERIMEAFAGR
FFEHCQGVFEHKDAAFVLSYSAIILNTDLHNGQVRRKMTEEEFIRNNKSINNGKDLPRELLVDLYHSIARNELRI
TDSGVVASEMTYSRWVDLIRRSKVVALFVPCEDLPLLDLDMFGIMWGPAIAAISVVFDHAEEEDVLNECRDGFLA
VAKISAYYCMKDVIDNLAVSLCKFTTLLNPGAERPAFAFVEDSKAKMAAETVFFIANNYGDHIRTGWRNLVDCIV
RLHRLRLLPRFFLESEETAGAPGGVSSPPSSSSVQAPPQQAAAGSSPSSPVLRPSSAVVAVLLPDSSGVVGTAPA
PSSESGAAGGGAGGGANHRTRWSMSSGGSSGASSSLGVLPPARGRSGSSGLMGRFSQLWSLETDSPTQDQIAAYQ
FMARAMETCRVEEIFTRSRSLAPESLVQLAKALVAAGPRPQKPAIGGGGGGGGGALALQEEEETAVLCLDLLMQV
TFCNRDRVLLIWSVVRDHLDKIMATSLTPGPLVEKAVFELLRTGLRLMSLSEEVAEELLRSLQLILRLDAKVADS
LCERIASDILQLLKVNFHHIRSPMGWRTVCSLLAKTARHPDAAETGFDALTFIMSDSSHITQANFLACLETGLAF
AEARVGGVDRSMRALTLVGGLVGNLTRWATALGDGGSGGLGGTMGGGGAGYGGGGAGLGADDAAYIQEAGRKELL
DFWTRLVRGVRLVCIEPREDVRDHALLCLQRSLLAADAIVPMPVLCGPAFDQVVFPVLDELLDIAQGQSPKEYRG
MEATLRKAMALLSRIFLHFLHHLAGSPGFQPLWLAVLERMERYMRLATSQGRRGGLLYEQLAENIPELLKNMLLV
MHSRQVLLPVVAGGGGGGGTGIESGDGGSGITQQSQMSMWDLTWQHVSLIAPNFSPETVLGVQVRVQG*

**>ME000774S09573**

MAVENLACALNTEIQAVVAMMRQNATWAMAPHYLEDPEPDDPLLEDFKVLRRNIFKWQDWSRVRPMEYLAPFLEV
IRSEKANTPITGVTLSAVYKMLTLEVFGPGTIDAQHAMHQIVDAVTACRFEVTDQASEEVVLMKILQVLLACLRC
RAGVLLTDRDVCNVVNTCFRVVHQSVTKGELLQRMACHSMQEMVRTIFGKLPALPLSAKGDDPEGTEGTMGVPEG
ADAAAAAAAAAAAAASRQGGATTTPPLQNGNGSAGSSAERDADSASEHAPPGGRQLQQGNGHPTPPAGEEQELKPS
VSTPEGFGEGALGPEPSAGTAGTGDAGSEAAGAGAGAVAMAGGGGGPDAEAKEALRAGEPGGAEASSPASEAPDG
ELETPRGGGGKPHGIACIVEIFHFLCSLLNLDDSLGGQGGAGGGGGGGGVGGEEDMSLLALGLVNSALELGGRHV
ARHPKLLALVQDELFRNLLQLGLSPSALVLSLVCSVVQNLFRSLRAHLKLQLEAFFTHVVVRLAQGRAPHHPQQQ
QQQQLLQEVALETLVDFCRQPGFAPEVYVNFDCDLTCSNTFEDLCSLLSKSAFPVNSPLSATHVLALEGLLALVT
SMAERIDSSSSPSSATSTATGPTPASALGGADGGEAEEEIPHYEPFWERRSESYSDPAKWLPHVTRQKLIKRRLL
VGVDHFNRDAKKGLAYLQAARLLPQPLDPRSVALLFRHAPGLSRSLLGEYLCDHREFNVAVLHEFSATFDFRGLA
LDGALRAYLESFRLQGEAQKIARVLEAFAERYYEQCPHVLANKDAAYILAYSVRRKMTEEEFIRNNRSINDGKDL
PREMLQALYHSIARNEIRVSTNGAHAGAGSGSAGGPPEMTYSRWIDLIRRSQNPDSAVPYILCDPTPAPAPAASI
GAGAGTAPLQPPLPTLPPAAAAALPALLDHEMFAIIAGPTIAAISVVFDHTEQEEVLRRCMDGFLAIARICAHRR
LDGVLNDLVVSLCKFTTLLNPGGGGGGGGGGASFAEEPAVAFGEDAKARMAAVTVFSMANRFGDHITSAWRNILD
CVLRLHKLGLLPAKVGSEASEDLDAAGDAAAAPPVVPPVISRQVRRPSGIMSRFSQFLLLDSEEQQAVPTEAQVA
AHQRTLRTIEACHIDQLFSDSKFLQAEPLQHLTRALVWAAGRPSGPRGGGGGGFASPEDEDTAVFCLELLIAVTL
TNRDRILLLWPGVSEHLAAVVQSASAPGPLVEKAVFGLLRICQRLLPYKEDLADELLRSLQLILRLDARVADTLC
ERITREVLHLVRANAAHVRSPAGWRTVCSLLAITARHPDAAEPGFEALSHIMADGAHLSPVNYVPCLDAARDFAE
ARVGGAERSIRGLDLLAGLVGCLQRWSQAMADSWQGQEGGLQGAAALRGQQELADMWLRLAQAVRRVCTDQREEV
RNHAVACLQRCVLAADGIALPATGAWPPLFDQVVFVMLDDLLDLALREKPKEFRNMEGTLHRALKLVSKVFLHFL
PRLTELPGRGFRALWLAVLSRMEMYLKAKLRGGAGASEQLRENIPELLKNMLLVMHAKGVLAPQSTPGGDSLWEL
TWHHVHSIAPALKDELFPPPPAPAPPPPPPPAPDAAGTAIAPPESAVPNEEAEEVARREVVEEPAGSEQAALQEV
GDGAGVVAEGLASSPQVVAEVEGIENDAEGAERL*

**>SM000017S02750**

MFELGRTTAGGWPAVRPAEYLAPFLEAVRSVETSAPITAAALAAVHRILRLRGALGASTPHAQEAVHAVIDAVTG
CRFEVTDPASEEVVLMKILQVLLACLQCEAGALLTDRDVCNVVNTCFRVVHQAAAKGELLQQMACHTMQEMVRTL
FARLPDLSPSALEDDNEAAAAAFATEGTLSVPEVDSEMRQSSPVSAQVGDNEGGLRPNGPAMPSVNGQLEEDSTL
PSSSSLEEASQAEVTGATDTAGGSDPAVSTATDASSAEGRLPGSLDLVSDAEGSSPGSRGRPPLPPGSRPVWPAP
LDVGASPSEVAAASSPRRLAGSPSSRRSNEAALGQDAGPAGSTVVGRPHGVPCMVEIFHFLCSLLSLDDVAAGLG
PGRPGGGGGEEDMPLFALGLITTALELCGPAVADHPKLLALVQDELFRSLMQLGLSTSPPLLSLVCSAVLSLYQF
LRRHLKLQLEAFFVVNIVRLAQGKYGASYQQQEVAMEALVDFVRQPPFMAEMFANLDCDITCSNAFEDLTSLLSK
SAFPVNSPLTTMHVLALDGLLALVHAIGGRVDALAITPMPSAALAGAGDNGEQTAHLPFWTARCDYGDPARWVGF
VRTRKHVKRRLAIGADHFNRDPKKGLEFLQGAALLPTPADASSVACFFRYTPGLSKSLLGEYFGDPEPFHIAVLR
AFTALFDFTGMLLDAALRTYLESFRLPGEAQKIERVLEAFAAHYYDQCPELLPSENVAHILSYSIIMLNTDLHNL
QVKRKMTEEQFIHNNREIDAGKNLPRETLAELYKSIARNEIRISSDGNAMHGAPEMTYSRWIDLIRRSQTAAPYT
LAGSAAALLDHDMFAIIAGPTIAAISVVFDHTEDEEVLQECMEGFLEVAKICARHRLEGVLDDLVVSLCKFTTLL
NPSSAAFDDPVLVFGEDLKARMAAMTAFTVANRYGDHIGPGWRNILDCILRLHKLGLLPSKVAAMTEAGDEASDG
TDSGAVPQPVAVASLQPMGARKRSTGLMSRFSQLLLLDGDDARAQPTEQQLAAHQRSLRAVEVCHIDQLFTDSRF
LQADSLQQLAKALVWEAGRPQQLTQKSGGGGAGGAAEDEDTAIFCLELLILVALTNRDRIALLWPLVYEYMSSIV
ASATAPGPLVEKAVFGLLHICQRLLPYKEELADELLRSLQLILKLDARVADAFCERITLEVSQLVRTSAGYVRSA
MGWRTVCSLLAITARHPEASEPGFSALEFLMADGAHVTPTNYVPCLDAARQFADARIGGVERSLKALDLLVDLTN
SLQRWSMTVEKDAMASSPAKAAAAEVAEAEQGLPEADEGTSQGALELAELWQRLAHGLRRVSRDHREEVRNHAIL
SLQRCLLAAEHIYVPLSAWGPCFDQVIFVMLDDLLECGVREKPKDFRGMENTLYHAMKLLSRILLQFLEKLSTLS
TFRGLWLAALTRMEMYMKAKVRGGSEELRENIPELLKNLLLVMHARGVLVPAATPSSDSLWELTCHHVEGISASL
IAELFPPPPPAPHAPEGESTQPLRAEEDQSLAVNGAGLLASAVVAKSGHDSSNETNGENGQLVREREGLPTRL*

**>SM000057S18366**

MFELGRTTAGGWPAVRPAEYLAPFLEAVRSVETSALITAAALAAVHRILRLRGALGASTPHAQEAVHAVIDAVTG
CRFEVTDPASEEVVLMKILQVLLACLQCEAGALLTDRDVCNVVNTCFRVVHQAAAKGELLQQMACHTMQEMVRTL
FARLPDLPPSALEDDNEAAAAAFATEGTLSVPEVDPEMRQSSPVSAQVGDNEGGLRPNGPAMPSVNGQLEDDSTL
PSSSSLEEASQAEVTGATDTAGGSDPAVSTATDASLAEGRLPGSLDLASDAEGSSPGSRGRPPLPPGSRPARPAP
LDVGASPSEAAAASSPRRLAGSPSSRRSNEAALGQDAGPAGSTIVGRPHGVPCMVEIFHFLCSLLSLDDAAAGLG
SGRPGGGGGEEDMPLFALGLITTALELCGPAVADHPKLLALVQDELFRSLMQLGLSTSPPLLSLVCSAVLSLYQF
LRRHLKLQLEAFFVVNIVRLAQGKYGASYQQQEVAMEALVDFVRQPPFMAEMFANLDCDITCSNAFEDLTSLLSK
SAFPVNSPLTTMHVLALDGLLALVHAIAGRVDALAITPMTSAALAGAGDNSEQTAHLPFWTARCDYGDPAQWVGF
VRARKHVKRRLAIGADHFNRDPKKGLEFLQGAALLPTPADASSVACFFRYTPGLSKSLLGEYFGDPEPFHIAVLR
AFTALFDFTGMPLDAALRTYLESFRLPGEAQKIERVLEAFAAHYYDQCPELLPSENVAHILSYSIIMLNTDLHNL
QVKRKMTEEQFIHNNREIDAGKNLPRETLAELYKSIARNEIRISSDGNAMHGAPEMTYSRWIDLIRRSQTAAPYI
LAGSAAALLDHDMFAIIAGPTIAAISVVFDHTEDEEVLQECMEGFLEVAKICARHRLEGVLDDLVVSLCKFTTLH
NPSSAAFADPVLVFGEDLKARMAAMTAFTVANRYGDHIGPGWRNILDCILRLHKLGLLPSKVAAMTEAGDEASDG
TDSGAVPQPVVVAPLQPTGARKRSTGLMSRFSQLLLLDGDDARAQPTEQQLAAHQRSLRAVEVCHIDQLFTDSRF
LQADSLQQLAKALVWEAGRPQQVPQKGGGGGAGGAAEDEDTAIFCLELLILVALTNRDRIALLWPLVYEYMSSIV
ASATAPGPLVEKAVFGLLHICQRLLPYKEELADELLRSLQLILKLDARVADAFCERITLEVSQLVRTSAGYVRSA
MGWRTVCSLLAITARHPEASEAGFSALEFLMADGAHVTPTNYVPCLDAARQFADARIGGVERSLKALDLLVGLTN
SLQRWSMTVVKDAMASSPAKAAAAEVAEAEQGVLEADESTSQGALELAELWQRLAHGLRRVSGDHREEVRNHAIL
SLQRCLLAAEHIYVPLSAWGPCFDQVIFVMLDDLLECGVREKPKDFRAMENTLYLAMKLLSRILLQFLEKLSTLS
TFRGLWRAALTRMEMYMKAKVRGGSEELRENIPELLKNLLLVMHARGVLVPAATPSGDSLWELTCHHVEGISASL
IAELFPPPPPAPRSPEAESTQPLLAEQDQSLAVDRAEPLASVVVAKGGHDSSTETNGENGQLVTEQGGLLTRL*
**>SM000082S18366**
MFELGRTTAGGWPAVRPAEYLAPFLEAVRSVETSALITAAALAAVHRILRLRGALGASTPHAQEAVHAVIDAVTG
CRFEVTDPASEEVVLMKILQVLLACLQCEAGALLTDRDVCNVVNTCFRVVHQAAAKGELLQQMACHTMQEMVRTL
FARLPDLPPSALEDDNEAAAAAFATEGTLSVPEVDPEMRQSSPVSAQVGDNEGGLRPNGPAMPSVNGQLEDDSTL
PSSSSLEEASQAEVTGATDTAGGSDPAVSTATDASLAEGRLPGSLDLASDAEGSSPGSRGRPPLPPGSRPARPAP
LDVGASPSEAAAASSPRRLAGSPSSRRSNEAALGQDAGPAGSTIVGRPHGVPCMVEIFHFLCSLLSLDDAAAGLG
SGRPGGGGGEEDMPLFALGLITTALELCGPAVADHPKLLALVQDELFRSLMQLGLSTSPPLLSLVCSAVLSLYQF
LRRHLKLQLEAFFVVNIVRLAQGKYGASYQQQEVAMEALVDFVRQPPFMAEMFANLDCDITCSNAFEDLTSLLSK
SAFPVNSPLTTMHVLALDGLLALVHAIAGRVDALAITPMTSAALAGAGDNSEQTAHLPFWTARCDYGDPAQWVGF
VRARKHVKRRLAIGADHFNRDPKKGLEFLQGAALLPTPADASSVACFFRYTPGLSKSLLGEYFGDPEPFHIAVLR
AFTALFDFTGMPLDAALRTYLESFRLPGEAQKIERVLEAFAAHYYDQCPELLPSENVAHILSYSIIMLNTDLHNL
QVKRKMTEEQFIHNNREIDAGKNLPRETLAELYKSIARNEIRISSDGNAMHGAPEMTYSRWIDLIRRSQTAAPYI
LAGSAAALLDHDMFAIIAGPTIAAISVVFDHTEDEEVLQECMEGFLEVAKICARHRLEGVLDDLVVSLCKFTTLH
NPSSAAFADPVLVFGEDLKARMAAMTAFTVANRYGDHIGPGWRNILDCILRLHKLGLLPSKVAAMTEAGDEASDG
TDSGAVPQPVVVAPLQPTGARKRSTGLMSRFSQLLLLDGDDARAQPTEQQLAAHQRSLRAVEVCHIDQLFTDSRF
LQADSLQQLAKALVWEAGRPQQVPQKGGGGGAGGAAEDEDTAIFCLELLILVALTNRDRIALLWPLVYEYMSSIV
ASATAPGPLVEKAVFGLLHICQRLLPYKEELADELLRSLQLILKLDARVADAFCERITLEVSQLVRTSAGYVRSA
MGWRTVCSLLAITARHPEASEAGFSALEFLMADGAHVTPTNYVPCLDAARQFADARIGGVERSLKALDLLVGLTN
SLQRWSMTVVKDAMASSPAKAAAAEVAEAEQGVLEADESTSQGALELAELWQRLAHGLRRVSGDHREEVRNHAIL
SLQRCLLAAEHIYVPLSAWGPCFDQVIFVMLDDLLECGVREKPKDFRAMENTLYLAMKLLSRILLQFLEKLSTLS
TFRGLWRAALTRMEMYMKAKVRGGSEELRENIPELLKNLLLVMHARGVLVPAATPSGDSLWELTCHHVEGISASL
IAELFPPPPPAPRSPEAESTQPLLAEQDQSLAVDRAEPLASVVVAKGGHDSSTETNGENGQLVTEQGGLLTRL*
**>Kleni_0140**
MAGPQVASLAEGSGWTGPQIACMVNTEINQVMAVMRQNARWAVAPRYLEEELGEDPLLRDFKILRRKLFTWQDWP
KVKPTDYLTPFLEVIRSEETSAPITGVALSAVHKVLSYEIFDAKTAGAAEVMHFIVDAVTGCKFEVTDPASEEVV
LLKILQVLLACLKCGAGPLLSNADVINIINTCLRVAFTKGELLQRTSCQMMQDMVRAIFARLHEFEPAEDGSTQP
GPAVASEQPDGRVLESVPSGAIAEGILSTASTPRSGHSAASTPKSRGAIRSIADIRFGPHPTATEGTPDDGPSLT
PRALKAFEEGKSLVEVVEEETSAVKEDSTLGADSLGPSDSAPVSEALAAAGAGIAAGFDAAVTAVGEAVSTAPVV
AGVPEAGAAPIVPTLSESVPDDTSGSDAVSGAPPLPPPVRVRAEPSDASAGTSGTVVASQPSAAAESLTAPLENG
DSAQKQPAPAASVQASPYGLPCLVEVFRFLCTLIVPEDGADKAQQHDSEDLSIFGLALLSAAIEVAGGSFEKHPR
ILALVTDDLFRNLMQTGQSSNLLVLSLVSGIVLALYQHLKHHLKLQMAAFFEFVVLRAAQGKFYGATYEQQEAAL
EILQDFCRQPSFMADMYANADCDITCSNTFEDLSNLLSKNAFPVNAPLSALHLLSLEGLLAVIQCMADRADNAST
PAGSGSSSPDYNLEEFPQFWNLKCDNWNDPTVWVDFLRRRKYVKRRLMIGADHFNRDPKKGLEFLQGVRLLPEKL
DPESVAFFFRMTPGLNKNLLGEYFGEPDAFNISVLECFTKQFNFTGMALDGALRTYLEAFRLPGEAQKISRVLEA
FASHYYEQCPQNFAHKDAAYVLSYSVIMLNTDQHNGQVKRKMTEEEFIRNNRKINDGKDLPRDMLTHLYYSIAKN
EIRMSSESLAAEPGMTYSRWLDLLRRSQATTPFLSCDPTKPLLDRDMFSVIWGPSIAAISVVFDHADDADTLQQC
MAGFLAVAKISAAHHMEDVFDNLVVSLCKFTTLIGNDKAAVAFGENTKARMATSAVFEIANRHGDTIRSGWRNIL
DCVIRLHKLGLLPTRREQSQNGQGRPSDNQEGSSILANAAAAAAAGTQAGMGRRKSAGIFGTVSRQFQNLLSLDA
GADGERSTIAEHQLAAHQRTLAMVEACRIDDLFTDSKFLHKASLLQLVKALINVSGKPHQKNISTEEEDTALFCL
DLLIAVTLRNRDRILLLWPAVHEHLASIVGSAQSPSPLVEKAVLDLLRVCQRLLPYKEDLGDELLKSLQLVLRLD
ARVADAYAERITREMLQLVKSSVTYIRSPVLWKTVCSLLAATARHPEAAEPGFEALTFIMQDAKHVTPVNFMPCM
DAALAFAEGRVGGVERSVKSLDLLASLSTNLGHWAKTLNAGQADTATSASISGAPASDAAREALDQAVSEAASTS

ATDVPQSPRLTSEKEALIDLWLRLSVGLGRICLDQREDVRNNALECLQRSLLQGEALDVRSGKVWVRVFDQIVFP
TLDELLDLAHKASAKSYRGMEGTLRRAMVLLSRVYLQFLTQLASNPGFANLWVAVLNRAEGYLKSSAPRSAGISM
AENIPELLKNMLLVMNARSILVPEVAPPAEGEAPTLWSLTWQHPIFRSLDPHFREDLFPPPKAAPSQPAPAVEQA
GAEEKQPEGVSTGPVAEAAEAVGEKAADASVAEESKVLVEGSPVEGSKDVAEEGTKPEDVDVAEGYEKVEKEMVE
PIKTNVT*
**>ChlatGNOM_Cat1294**
MISTALRGVEDPTYGMHVPSLLNAEIQGVMAMMRQNAKWAVVPRFTDDEANDDPLLQDFKALRRQIFAWQDWSQV
DPLLYLGPFLEVVRSVETSGPITGMALSAIFKLVTYGVIGVHVVNANEAIRMIADATTHCRFEATDPGSDEVVLM
KILQVLRCPAGALLSDDDVCHIIQTCFRIGHQTGRESELLRRTARQIMQEMVRTLFRRAPPSDHSAPPYSVSCMV
EVFKFMASLITPEETREAEDLCLFGLALVNAALETGGCCFAAHPLLLTVVQNELFYNLTKAGLSSNLSVLSFVCG
ILLNLYVHCREQLKLQLEAFFTHVLLRVAEGKGAPSYEQQEAAIECIVDFCRQASFMSDMYSNLDCDLRFSNVFE
DLFALLSKNAFPVNSPLSGIHLLSLEGLLAIVHSIAERAARDAGAIALASGGSTAVGPVMFPLTDATEYVDYWTA
PCPGPEDPQLEGLLAVIQCMADRADNASTPAGSGSSSPDYNLEEFPQFWNLKCDNWNDPTVLVDCSRKRKYTKRR
LMVGVDHFNRDIKKGFEFLQSICLLPKPLESMAVAQLFRYTPKLNKTAVGEYLGDPGEFNLQVLKEFASMFDFSD
MSLDYALRTFLDSFRLPGEAQKIARIMETFAEQYHRHASNTYANADAAYVLSYSCIMLNTDLHNGQVKKKMTMEE
FIRNNRQINAGQDLPRDMLEELYHSIAKNEIKISSEAMSPSRWVDLLKRAKTGGKFIPCGLSTLYDRDMFVIMWG
PSVAAISVVFDHTEDEDVLRECLDGFISIAKVAAHHRLEDVLDNLVISLCKFTTLLNPTAEKPSVAFGNDTKARM
AAVTVFSIANKCGDYIRAGWRNILDCILRLHKLGLLPSRQRTVRCIEACRIDDIFSESKFLKAESLAQLAKALIW
ASGRSYKDDEDTAVFCLDLLIAITLHNRDRILLLWPLVLAGIIACAQAPGPLVEKAVFDLLKVCQRLLPYKAVIA
DELLRSLQLILKLDAKVADAYAERITAEVLELVRRDVTYIKSPLGWRTVCALLAMTAHHPDAAELGFEALTFLMA
DGAHITPSNFMPCLEAGLAFAEGRLGGADRSLKALDLLGAMTGNLCRWSKAIQSPDRDSTQSLIDLWLPLVGAFC
RLGAEPRAPVRNHSLTMLQRVLLAAESAGIPASVYYAVLLQAVAPMLRDVLELAKKRSGEYGDMGTSLKVAMTML
SKTFLQWLPRLSESPPEFRLLWLHILGCMDQSLRAAPRSEQLLETIPEMVKNMVLVMSTRGFLTRDSIGSTESLY
EASRKVVLSIAPDVAAEAFGSSAAAEASAAPAPQ*

#### Suppl. Figure S1. Duplicated GNOM-related genes in gymnosperms (nextstrain)

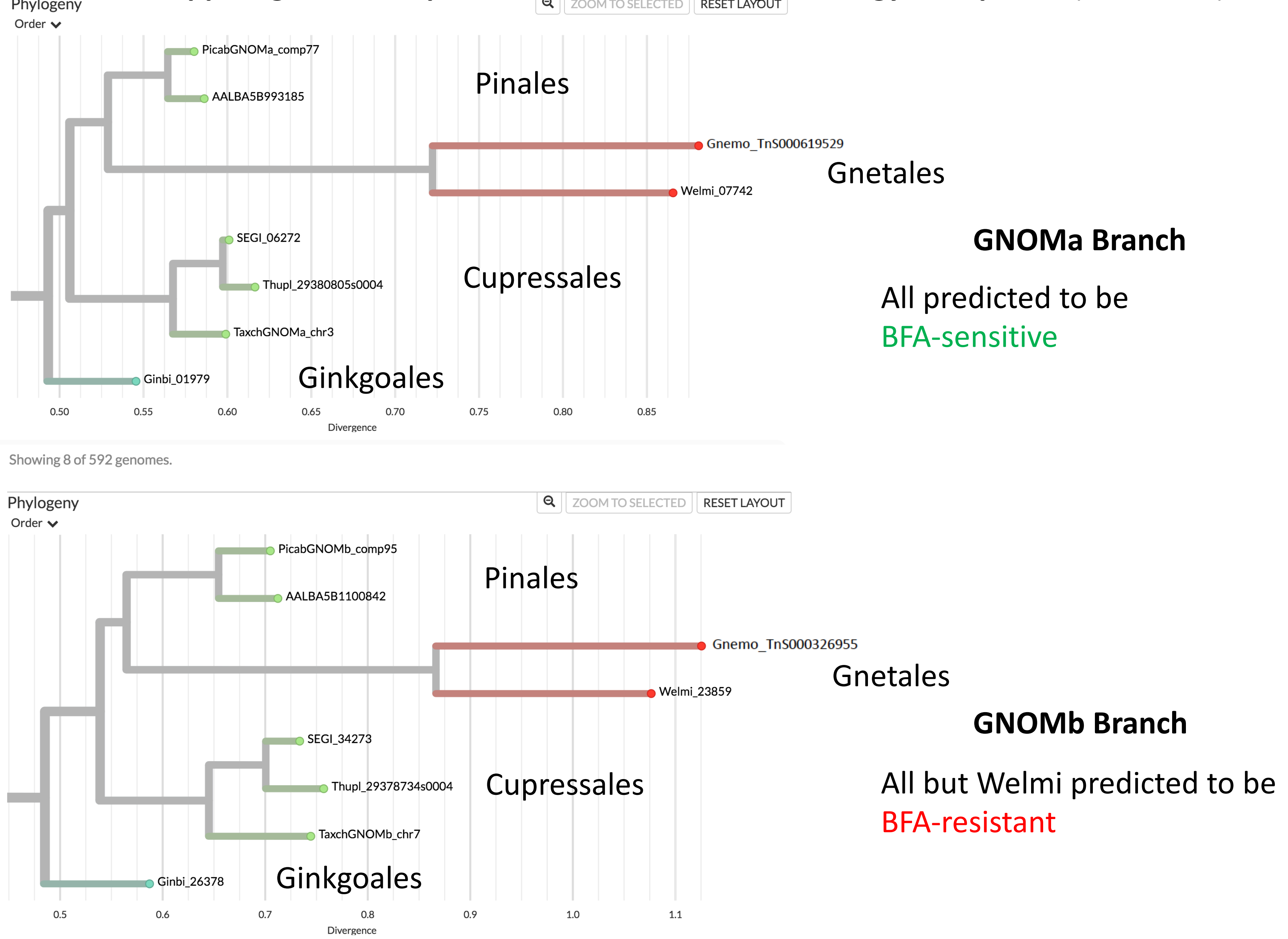

#### Suppl. Figure S2. GNOM-related genes in monocots (nextstrain)

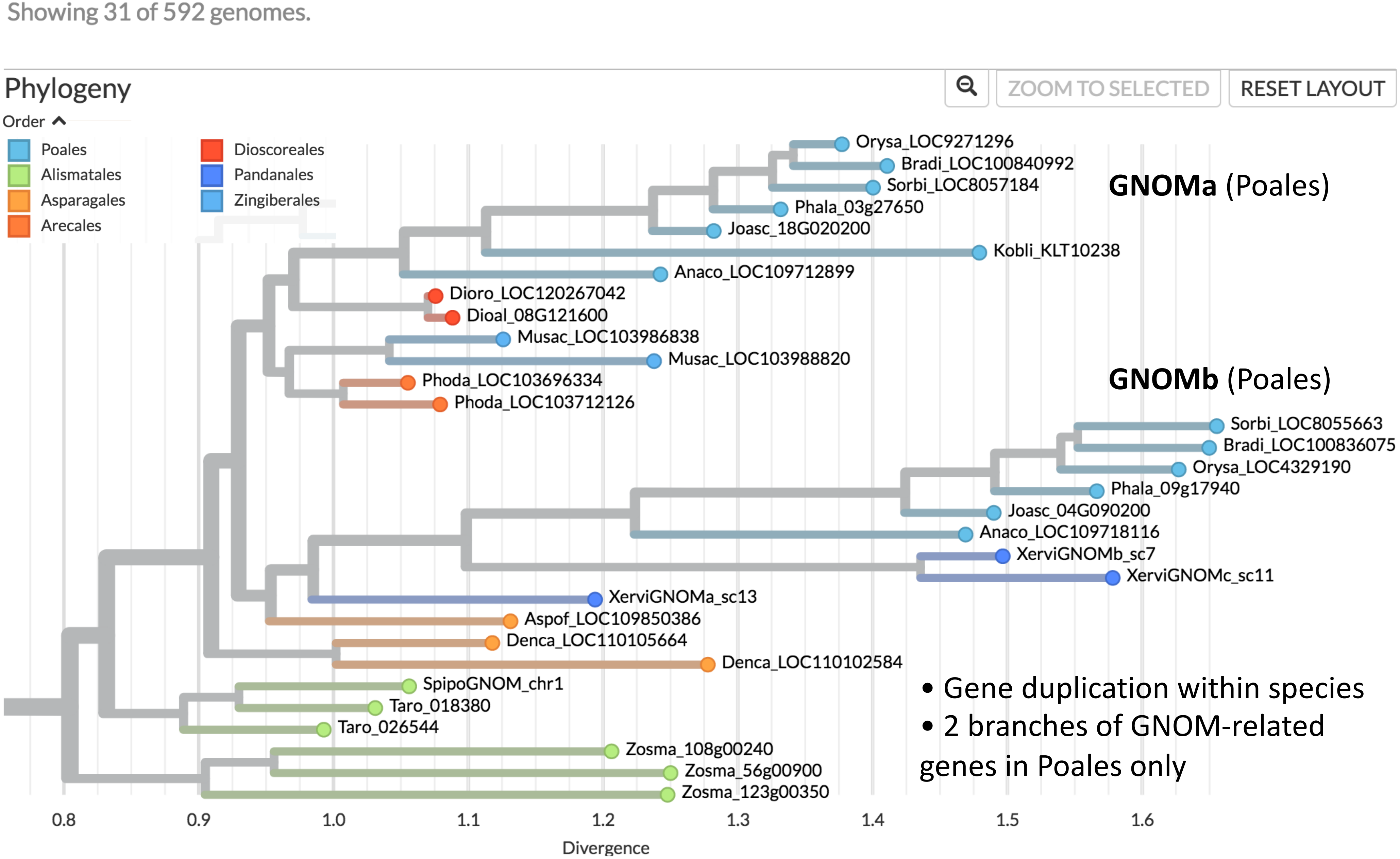

- Gene duplication within species
- 2 branches of GNOM-related genes in Poales only

### Suppl. Figure S3A. Genes flanking GNOM paralogs in OSG1 and OSG2

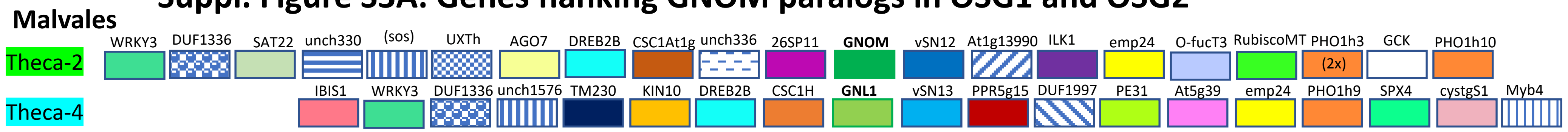

#### GNOM-OSG1

WRKY3 – WRKY family transcription factor; not clearly distinguished from WRKY4 (annotations used interchangeably) (paralog in OSG2)
DUF1336 – domain of unknown function (often annotated as "ENHANCED DISEASE RESISTANCE 2, C-terminal" (Theca_LOC18610328) (paralog in OSG2)
SAT22 – S-acyltransferase 22
unch330 – uncharacterised protein Theca_LOC18610330
(sos) – son of sevenless (misnomer?), Theca_LOC18610331 (paralog in OSG2)
UXTh – UXT homolog (prefoldin-related?)
AGO7 – argonaute 7 (involved in ta-siRNA generation?)
DREB2B – specific member of RAP2 (related to AP2) family (often annotated as RAP2-11) involved in ABA/stress response (paralog in OSG2)
CSC1At1g – CSC1-like family of calcium-sensitive channels member (At1G69450) (paralog in OSG2)
Unch336 – uncharacterised Theca_LOC18610336
26SP11 – 26S proteasome non-ATPase regulatory subunit 11 homolog
vSN12 – vesicle transport v-SNARE 12 (VTI12) (paralog in OSG2)
At1g13990 – uncharacterised Theca_LOC18610340 (DUF3110; next to *AtGNOM* on At-chr1)
ILK1 –serine/threonine-protein kinase containing ankyrin (ANK) repeats (also designated STY46)
emp24 – transmembrane emp24 domain-containing ER protein p24delta9 (paralog in OSG2)
O-fucT3 – O-fucosyltransferase 3 (also annotated as rhamnogalacturonan I rhamnosyltransferase 1 (RTIRT1) or uncharacterised protein At1g04910)
RubiscoMT – Ribulose-1,5 bisphosphate carboxylase/oxygenase large subunit N-methyltransferase (also annotated as fructose-bisphosphate aldolase-lysine N-methyltransferase)
PHO1h3 – phosphate transporter PHO1 homolog 3 (paralog in OSG2)
GCK – uncharacterized protein Theca_LOC18610351 with GCK domain
PHO1h3 – phosphate transporter PHO1 homolog 10 (paralog in OSG2)

#### GNL1-OSG2

IBIS1 – IMPAIRED IN BABA-INDUCED STERILITY 1 (also designated serine/threonine-protein kinase At1g54610 )
unch11576 – uncharacterized protein Theca_LOC18601576 (related to (sos) paralog in OSG1)
TM230 – transmembrane protein 230
KIN10 – SNF1-related protein kinase catalytic subunit alpha KIN10
CSC1H – CSC1-like family of calcium-sensitive channels member HYP1 (paralog in OSG1)
PPR5g15 – pentatricopeptide repeat-containing protein At5g15340, mitochondrial
DUF1997 – uncharacterized protein Theca_LOC18601568
PE31 – pectinesterase 31
At5g39 – uncharacterised protein At5g39570 (misnomer? Theca_LOC18601566)
SPX4 – SPX domain-containing protein 4
cystgS1 – cystathionine gamma-synthase 1, chloroplastic (CGS1)
Myb4 – myb-related protein Myb4 (membeer of MYB transcription factor family, also designated MYB16 or MYB106)

Suppl. Figure 3B. GNOM-related ortho-synteny groups (OSGs) mapped onto *Theobroma cacao* genome

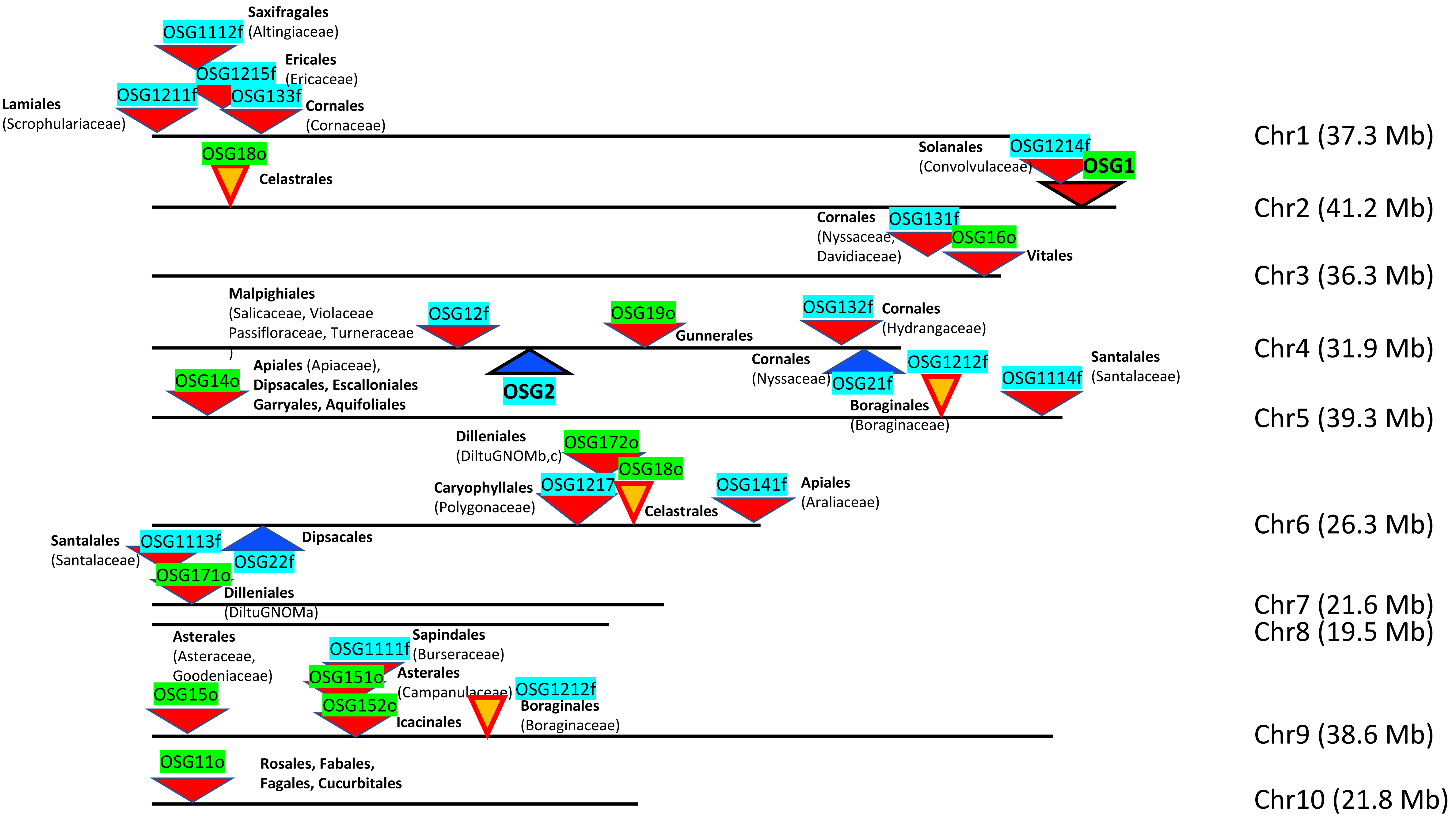

### Suppl. Figure S4. Brassicales cladogram and OSGs

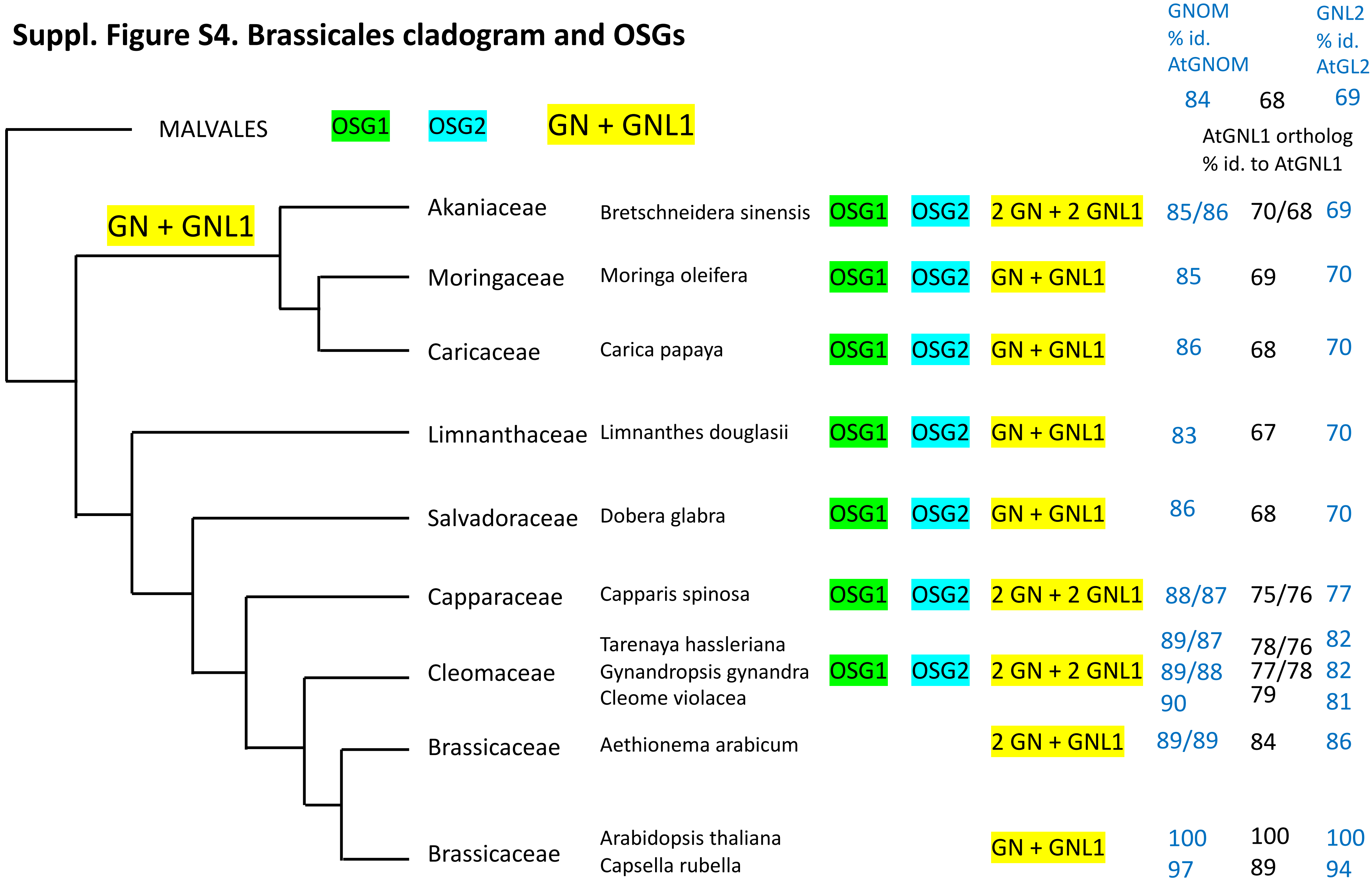

Cladogram based on Zuntini et al. (2024)

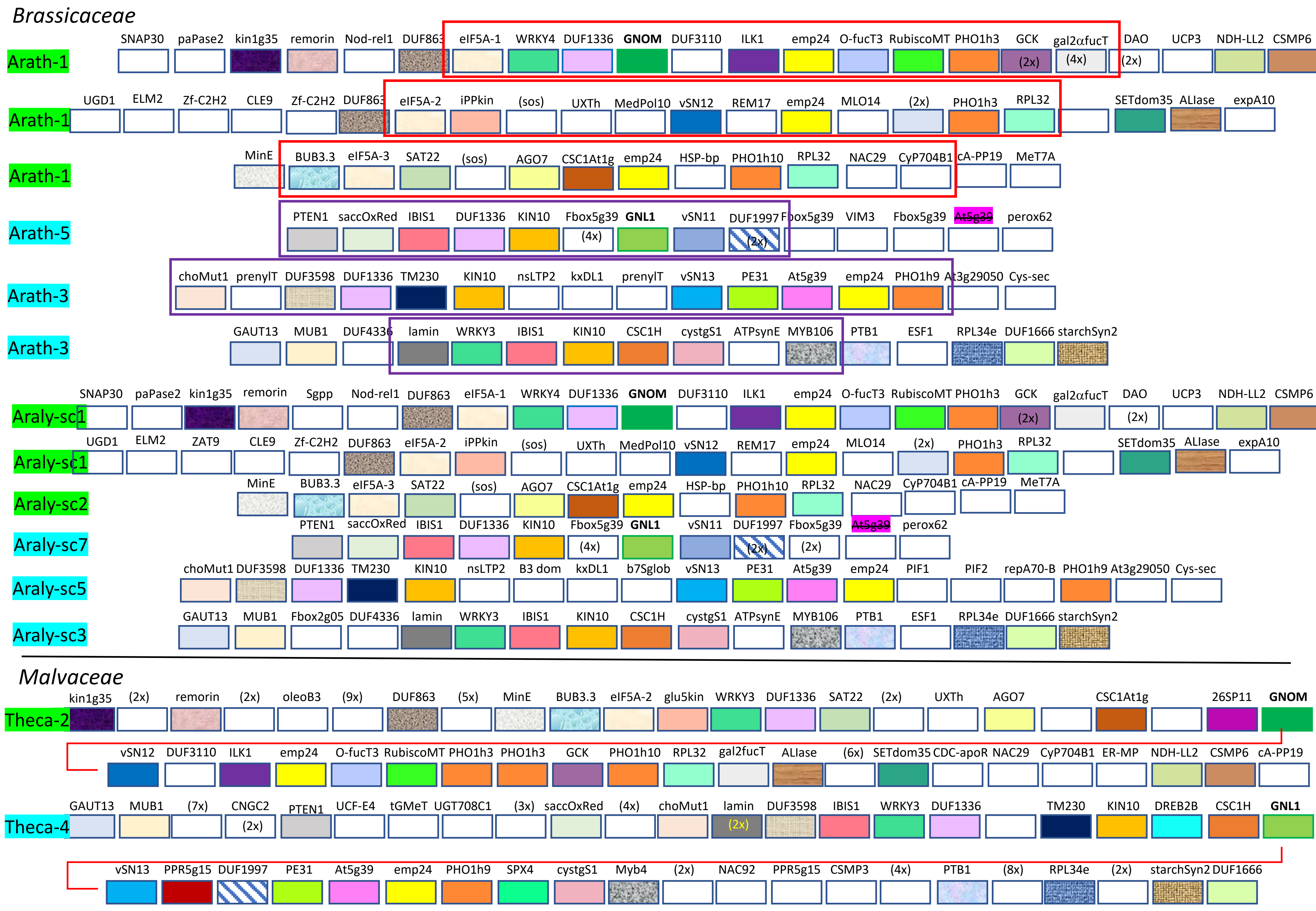

Brassicaceae
Arath-1
SNAP30 paPase2 kin1g35 remorin Nod-rel1 DUF863 eIF5A-1 WRKY4 DUF1336 GNOM DUF3110 ILK1 emp24 O-fucT3 RubiscoMT PHO1h3 GCK gal2αfucT DAO UCP3 NDH-LL2 CSMP6
(2x) (4x) (2x)
Arath-1
UGD1 ELM2 Zf-C2H2 CLE9 Zf-C2H2 DUF863 eIF5A-2 iPPkin (sos) UXTh MedPol10 vSN12 REM17 emp24 MLO14 (2x) PHO1h3 RPL32 SETdom35 ALlase expA10
Arath-1
MinE BUB3.3 eIF5A-3 SAT22 (sos) AGO7 CSC1At1g emp24 HSP-bp PHO1h10 RPL32 NAC29 CyP704B1 cA-PP19 MeT7A
Arath-5
PTEN1 saccOxRed IBIS1 DUF1336 KIN10 Fbox5g39 GNL1 vSN11 DUF1997 Fbox5g39 VIM3 Fbox5g39 At5g39 perox62
(4x) (2x)
Arath-3
choMut1 prenylT DUF3598 DUF1336 TM230 KIN10 nsLTP2 kxDL1 prenylT vSN13 PE31 At5g39 emp24 PHO1h9 At3g29050 Cys-sec
Arath-3
GAUT13 MUB1 DUF4336 lamin WRKY3 IBIS1 KIN10 CSC1H cystgS1 ATPsynE MYB106 PTB1 ESF1 RPL34e DUF1666 starchSyn2
Araly-sc1
SNAP30 paPase2 kin1g35 remorin Sgpp Nod-rel1 DUF863 eIF5A-1 WRKY4 DUF1336 GNOM DUF3110 ILK1 emp24 O-fucT3 RubiscoMT PHO1h3 GCK gal2αfucT DAO UCP3 NDH-LL2 CSMP6
(2x) (2x)
Araly-sc1
UGD1 ELM2 ZAT9 CLE9 Zf-C2H2 DUF863 eIF5A-2 iPPkin (sos) UXTh MedPol10 vSN12 REM17 emp24 MLO14 (2x) PHO1h3 RPL32 SETdom35 ALlase expA10
Araly-sc2
MinE BUB3.3 eIF5A-3 SAT22 (sos) AGO7 CSC1At1g emp24 HSP-bp PHO1h10 RPL32 NAC29 CyP704B1 cA-PP19 MeT7A
Araly-sc7
PTEN1 saccOxRed IBIS1 DUF1336 KIN10 Fbox5g39 GNL1 vSN11 DUF1997 Fbox5g39 At5g39 perox62
(4x) (2x) (2x)
Araly-sc5
choMut1 DUF3598 DUF1336 TM230 KIN10 nsLTP2 B3 dom kxDL1 b7Sglob vSN13 PE31 At5g39 emp24 PIF1 PIF2 repA70-B PHO1h9 At3g29050 Cys-sec
Araly-sc3
GAUT13 MUB1 Fbox2g05 DUF4336 lamin WRKY3 IBIS1 KIN10 CSC1H cystgS1 ATPsynE MYB106 PTB1 ESF1 RPL34e DUF1666 starchSyn2
Malvaceae
Theca-2
kin1g35 (2x) remorin (2x) oleoB3 (9x) DUF863 (5x) MinE BUB3.3 eIF5A-2 glu5kin WRKY3 DUF1336 SAT22 (2x) UXTh AGO7 CSC1At1g 26SP11 GNOM
vSN12 DUF3110 ILK1 emp24 O-fucT3 RubiscoMT PHO1h3 PHO1h3 GCK PHO1h10 RPL32 gal2fucT ALlase (6x) SETdom35 CDC-apoR NAC29 CyP704B1 ER-MP NDH-LL2 CSMP6 cA-PP19
Theca-4
GAUT13 MUB1 (7x) CNGC2 PTEN1 UCF-E4 tGMeT UGT708C1 (3x) saccOxRed (4x) choMut1 lamin DUF3598 IBIS1 WRKY3 DUF1336 TM230 KIN10 DREB2B CSC1H GNL1
(2x) (2x)
vSN13 PPR5g15 DUF1997 PE31 At5g39 emp24 PHO1h9 SPX4 cystgS1 Myb4 (2x) NAC92 PPR5g15 CSMP3 (4x) PTB1 (8x) RPL34e (2x) starchSyn2 DUF1666

*Capparaceae*

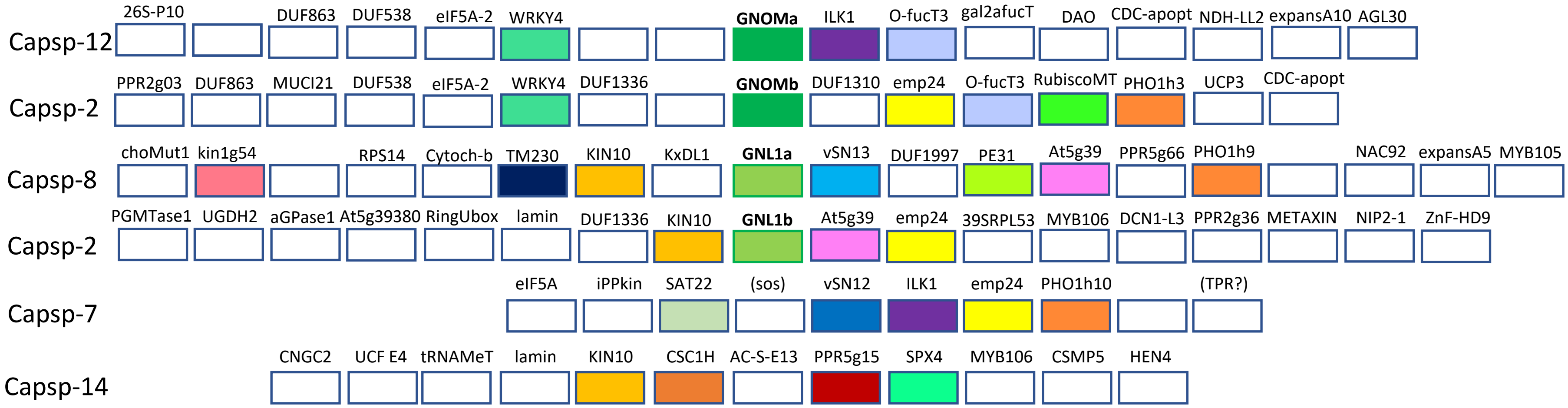

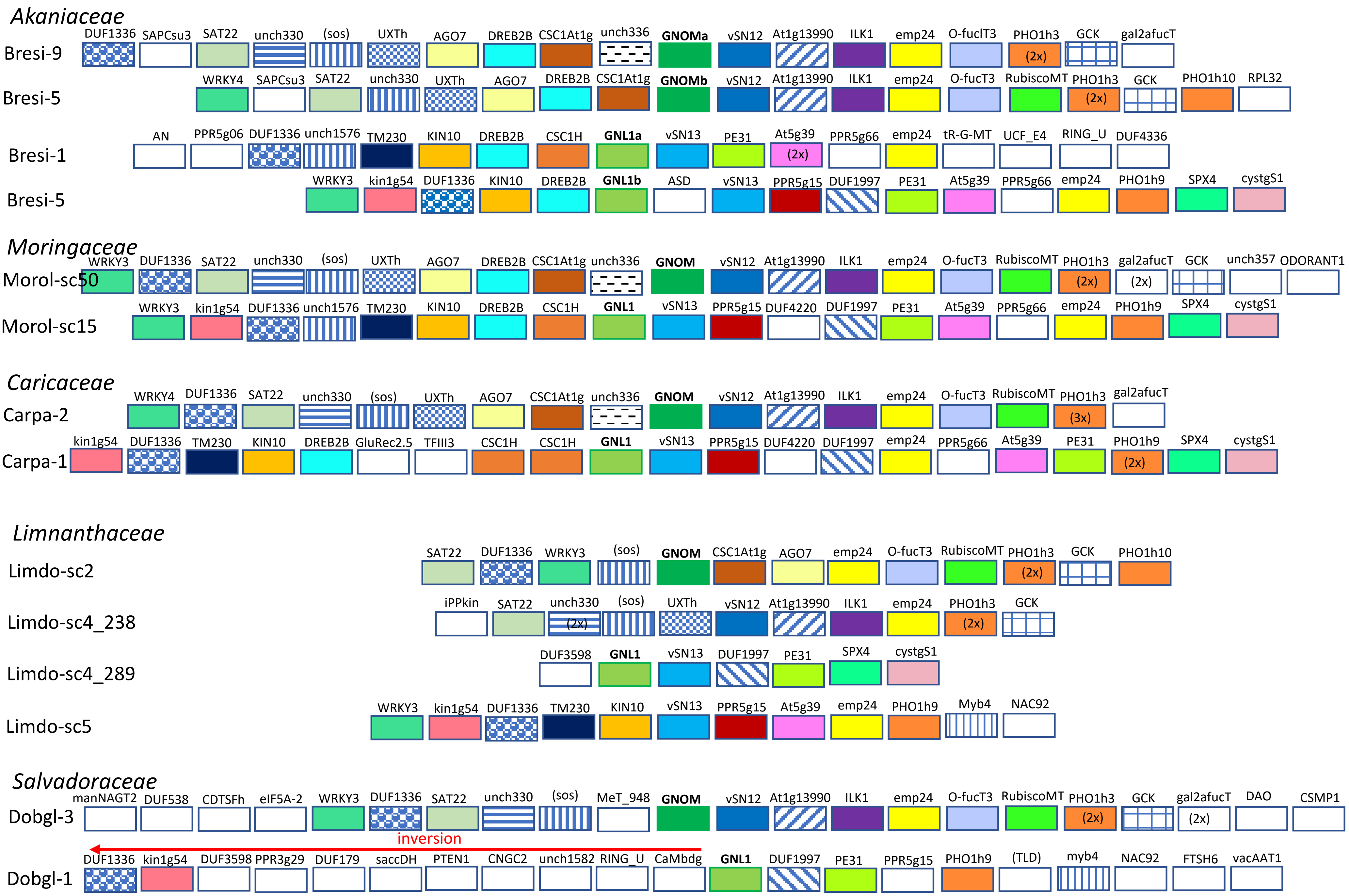
Akaniaceae
Bresi-9
DUF1336 SAPCsu3 SAT22 unch330 (sos) UXTh AGO7 DREB2B CSC1At1g unch336 GNOMa vSN12 At1g13990 ILK1 emp24 O-fuclT3 PHO1h3 (2x) GCK gal2afucT
Bresi-5
WRKY4 SAPCsu3 SAT22 unch330 UXTh AGO7 DREB2B CSC1At1g GNOMb vSN12 At1g13990 ILK1 emp24 O-fucT3 RubiscoMT PHO1h3 (2x) GCK PHO1h10 RPL32
Bresi-1
AN PPR5g06 DUF1336 unch1576 TM230 KIN10 DREB2B CSC1H GNL1a vSN13 PE31 At5g39 (2x) PPR5g66 emp24 tR-G-MT UCF_E4 RING_U DUF4336
Bresi-5
WRKY3 kin1g54 DUF1336 KIN10 DREB2B GNL1b ASD vSN13 PPR5g15 DUF1997 PE31 At5g39 PPR5g66 emp24 PHO1h9 SPX4 cystgS1
Moringaceae
Morol-sc50
WRKY3 DUF1336 SAT22 unch330 (sos) UXTh AGO7 DREB2B CSC1At1g unch336 GNOM vSN12 At1g13990 ILK1 emp24 O-fucT3 RubiscoMT PHO1h3 (2x) gal2afucT (2x) GCK unch357 ODORANT1
Morol-sc15
WRKY3 kin1g54 DUF1336 unch1576 TM230 KIN10 DREB2B CSC1H GNL1 vSN13 PPR5g15 DUF4220 DUF1997 PE31 At5g39 PPR5g66 emp24 PHO1h9 SPX4 cystgS1
Caricaceae
Carpa-2
WRKY4 DUF1336 SAT22 unch330 (sos) UXTh AGO7 CSC1At1g unch336 GNOM vSN12 At1g13990 ILK1 emp24 O-fucT3 RubiscoMT PHO1h3 (3x) gal2afucT
Carpa-1
kin1g54 DUF1336 TM230 KIN10 DREB2B GluRec2.5 TFIII3 CSC1H CSC1H GNL1 vSN13 PPR5g15 DUF4220 DUF1997 emp24 PPR5g66 At5g39 PE31 PHO1h9 (2x) SPX4 cystgS1
Limnanthaceae
Limdo-sc2
SAT22 DUF1336 WRKY3 (sos) GNOM CSC1At1g AGO7 emp24 O-fucT3 RubiscoMT PHO1h3 (2x) GCK PHO1h10
Limdo-sc4_238
iPPkin SAT22 unch330 (2x) (sos) UXTh vSN12 At1g13990 ILK1 emp24 PHO1h3 (2x) GCK
Limdo-sc4_289
DUF3598 GNL1 vSN13 DUF1997 PE31 SPX4 cystgS1
Limdo-sc5
WRKY3 kin1g54 DUF1336 TM230 KIN10 vSN13 PPR5g15 At5g39 emp24 PHO1h9 Myb4 NAC92
Salvadoraceae
Dobgl-3
manNAGT2 DUF538 CDTSFh eIF5A-2 WRKY3 DUF1336 SAT22 unch330 (sos) MeT_948 GNOM vSN12 At1g13990 ILK1 emp24 O-fucT3 RubiscoMT PHO1h3 (2x) GCK gal2afucT (2x) DAO CSMP1
inversion
Dobgl-1
DUF1336 kin1g54 DUF3598 PPR3g29 DUF179 saccDH PTEN1 CNGC2 unch1582 RING_U CaMbdg GNL1 DUF1997 PE31 PPR5g15 PHO1h9 (TLD) myb4 NAC92 FTSH6 vacAAT1

**Suppl. Figure S5. Malvales cladogram and OSGs**

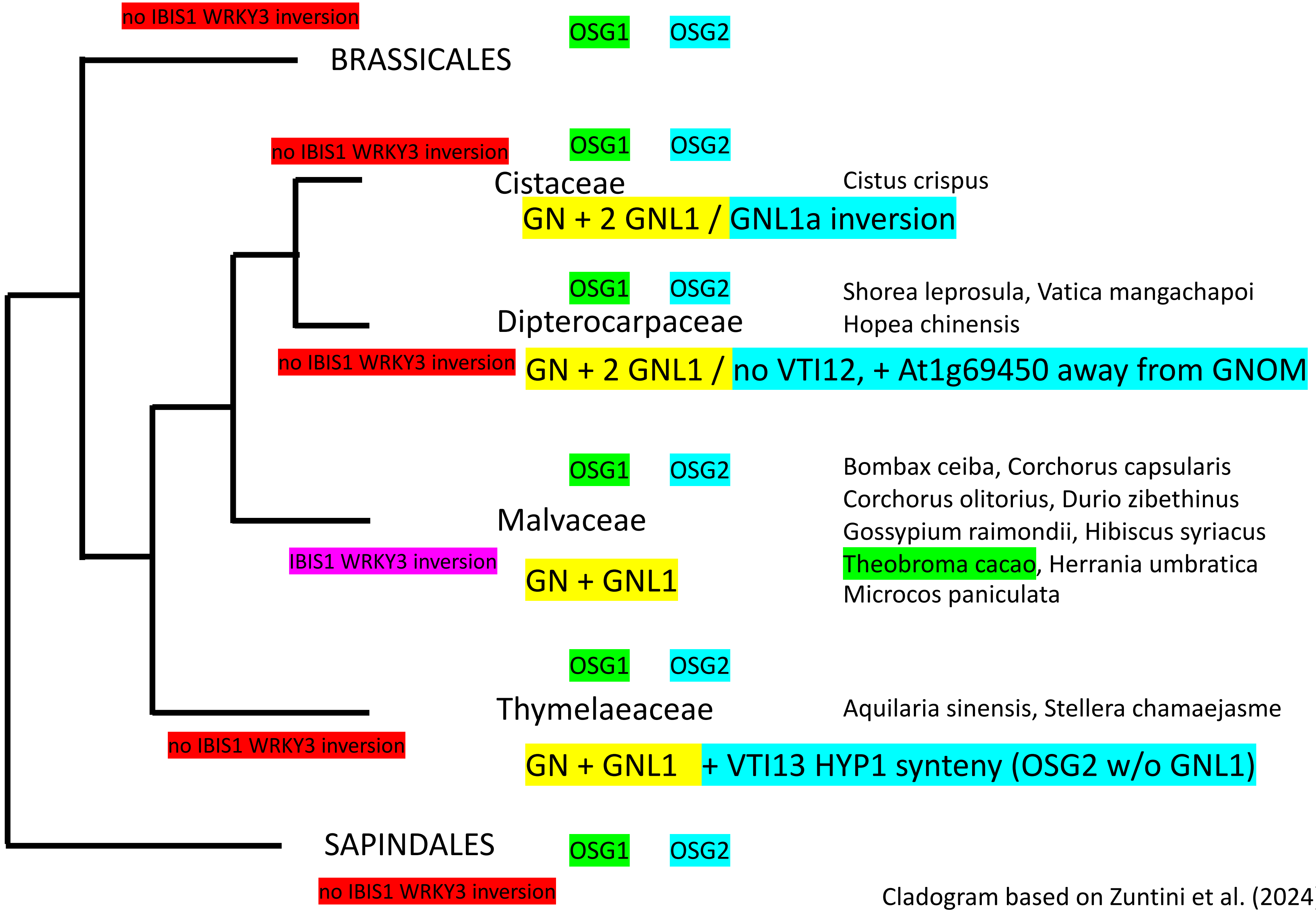

Cladogram based on Zuntini et al. (2024)

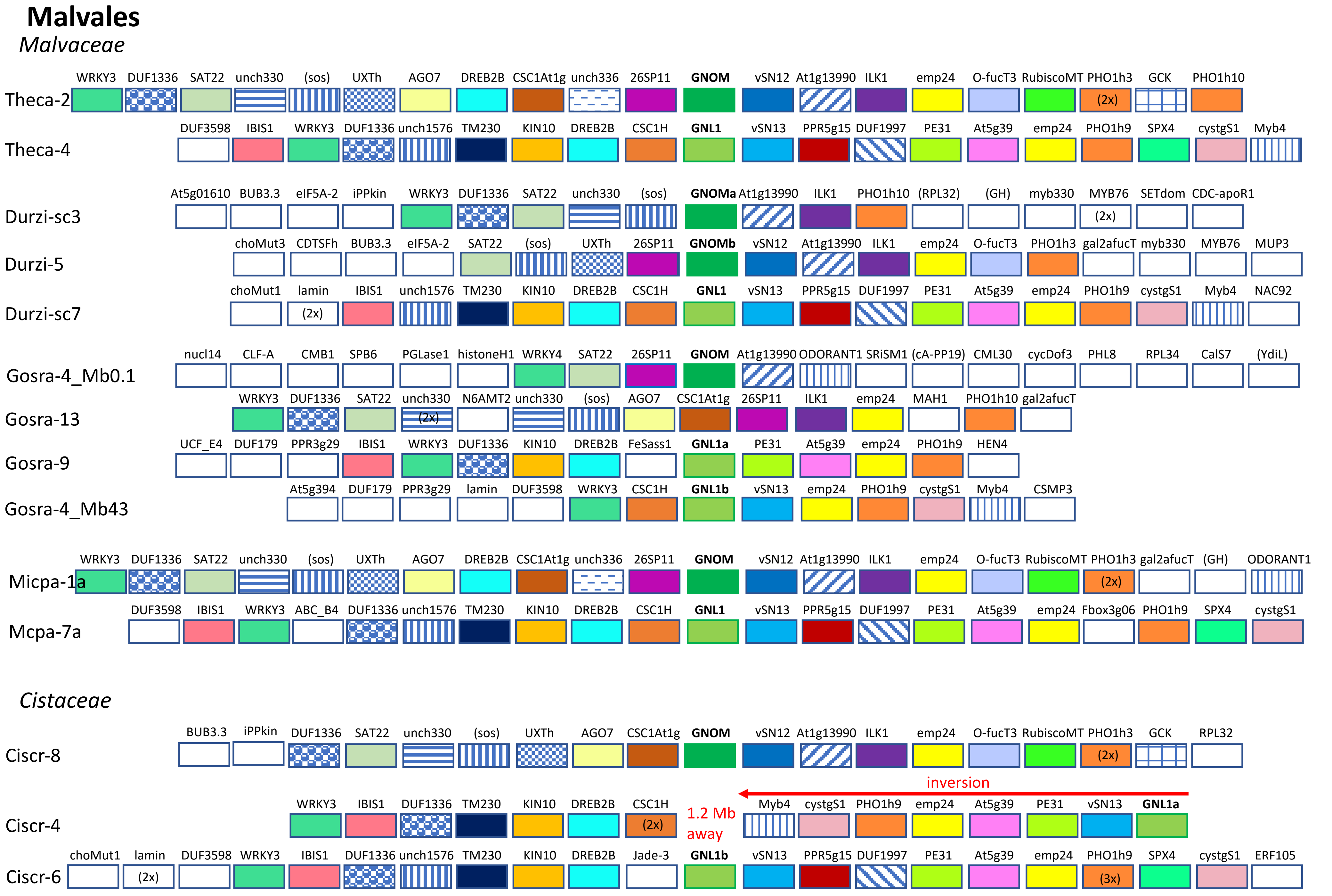
Malvales
Malvaceae
Theca-2
WRKY3 DUF1336 SAT22 unch330 (sos) UXTh AGO7 DREB2B CSC1At1g unch336 26SP11 GNOM vSN12 At1g13990 ILK1 emp24 O-fucT3 RubiscoMT PHO1h3 GCK PHO1h10
(2x)
Theca-4
DUF3598 IBIS1 WRKY3 DUF1336 unch1576 TM230 KIN10 DREB2B CSC1H GNL1 vSN13 PPR5g15 DUF1997 PE31 At5g39 emp24 PHO1h9 SPX4 cystgS1 Myb4
Durzi-sc3
At5g01610 BUB3.3 eIF5A-2 iPPkin WRKY3 DUF1336 SAT22 unch330 (sos) GNOMa At1g13990 ILK1 PHO1h10 (RPL32) (GH) myb330 MYB76 SETdom CDC-apoR1
(2x)
Durzi-5
choMut3 CDTSFh BUB3.3 eIF5A-2 SAT22 (sos) UXTh 26SP11 GNOMb vSN12 At1g13990 ILK1 emp24 O-fucT3 PHO1h3 gal2afucT myb330 MYB76 MUP3
Durzi-sc7
choMut1 lamin IBIS1 unch1576 TM230 KIN10 DREB2B CSC1H GNL1 vSN13 PPR5g15 DUF1997 PE31 At5g39 emp24 PHO1h9 cystgS1 Myb4 NAC92
(2x)
Gosra-4_Mb0.1
nucl14 CLF-A CMB1 SPB6 PGLase1 histoneH1 WRKY4 SAT22 26SP11 GNOM At1g13990 ODORANT1 SRiSM1 (cA-PP19) CML30 cycDof3 PHL8 RPL34 CalS7 (YdiL)
Gosra-13
WRKY3 DUF1336 SAT22 unch330 N6AMT2 unch330 (sos) AGO7 CSC1At1g 26SP11 ILK1 emp24 MAH1 PHO1h10 gal2afucT
(2x)
Gosra-9
UCF_E4 DUF179 PPR3g29 IBIS1 WRKY3 DUF1336 KIN10 DREB2B FeSass1 GNL1a PE31 At5g39 emp24 PHO1h9 HEN4
Gosra-4_Mb43
At5g394 DUF179 PPR3g29 lamin DUF3598 WRKY3 CSC1H GNL1b vSN13 emp24 PHO1h9 cystgS1 Myb4 CSMP3
Micpa-1a
WRKY3 DUF1336 SAT22 unch330 (sos) UXTh AGO7 DREB2B CSC1At1g unch336 26SP11 GNOM vSN12 At1g13990 ILK1 emp24 O-fucT3 RubiscoMT PHO1h3 gal2afucT (GH) ODORANT1
(2x)
Mcpa-7a
DUF3598 IBIS1 WRKY3 ABC_B4 DUF1336 unch1576 TM230 KIN10 DREB2B CSC1H GNL1 vSN13 PPR5g15 DUF1997 PE31 At5g39 emp24 Fbox3g06 PHO1h9 SPX4 cystgS1
Cistaceae
Ciscr-8
BUB3.3 iPPkin DUF1336 SAT22 unch330 (sos) UXTh AGO7 CSC1At1g GNOM vSN12 At1g13990 ILK1 emp24 O-fucT3 RubiscoMT PHO1h3 GCK RPL32
(2x)
inversion
Ciscr-4
WRKY3 IBIS1 DUF1336 TM230 KIN10 DREB2B CSC1H
(2x)
1.2 Mb away
Myb4 cystgS1 PHO1h9 emp24 At5g39 PE31 vSN13 GNL1a
Ciscr-6
choMut1 lamin DUF3598 WRKY3 IBIS1 DUF1336 unch1576 TM230 KIN10 DREB2B Jade-3 GNL1b vSN13 PPR5g15 DUF1997 PE31 At5g39 emp24 PHO1h9 SPX4 cystgS1 ERF105
(2x) (3x)

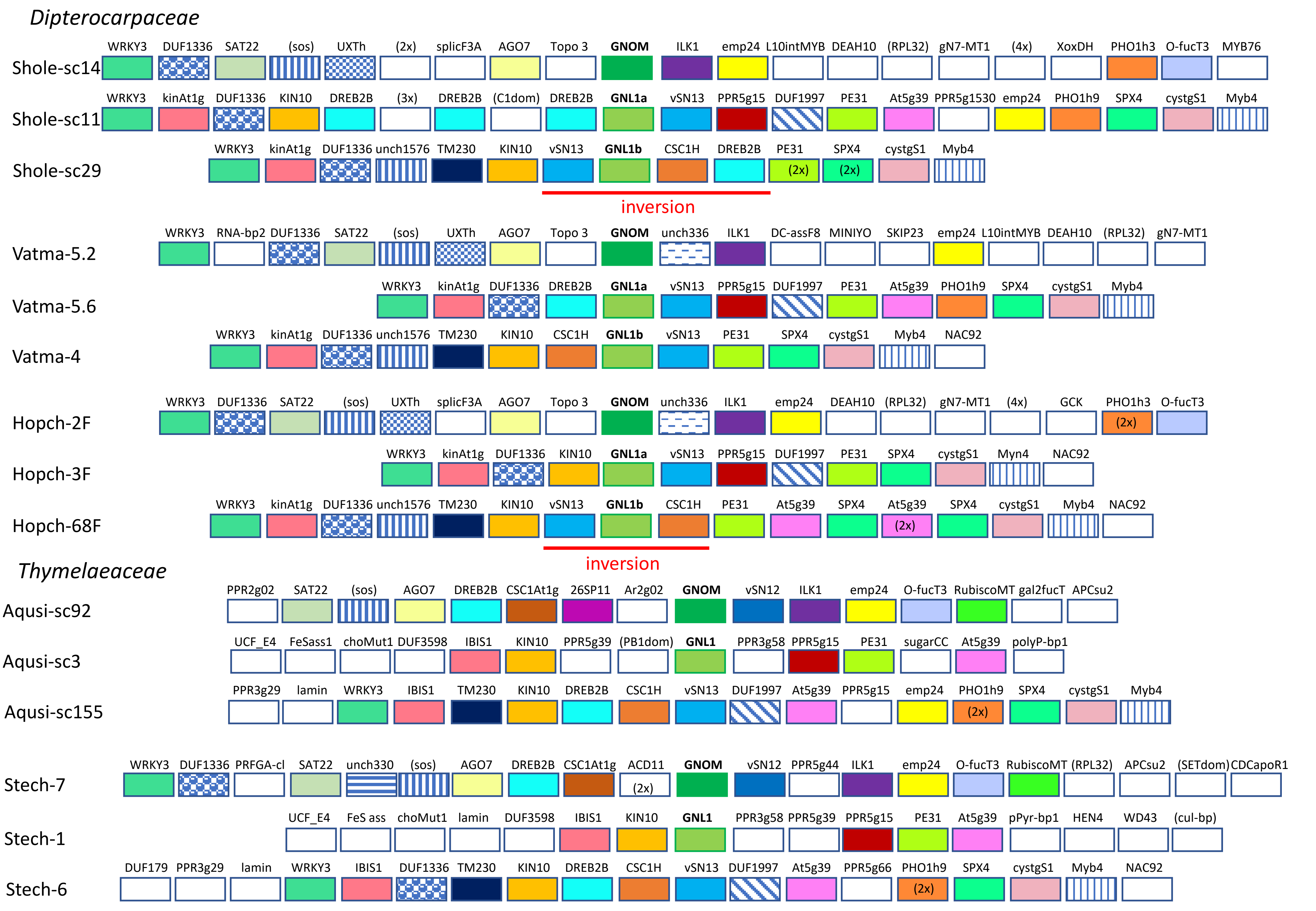

Dipterocarpaceae
Shole-sc14
WRKY3 DUF1336 SAT22 (sos) UXTh (2x) splicF3A AGO7 Topo 3 GNOM ILK1 emp24 L10intMYB DEAH10 (RPL32) gN7-MT1 (4x) XoxDH PHO1h3 O-fucT3 MYB76
Shole-sc11
WRKY3 kinAt1g DUF1336 KIN10 DREB2B (3x) DREB2B (C1dom) DREB2B GNL1a vSN13 PPR5g15 DUF1997 PE31 At5g39 PPR5g1530 emp24 PHO1h9 SPX4 cystgS1 Myb4
Shole-sc29
WRKY3 kinAt1g DUF1336 unch1576 TM230 KIN10 vSN13 GNL1b CSC1H DREB2B PE31 SPX4 cystgS1 Myb4
(2x) (2x)
inversion
Vatma-5.2
WRKY3 RNA-bp2 DUF1336 SAT22 (sos) UXTh AGO7 Topo 3 GNOM unch336 ILK1 DC-assF8 MINIYO SKIP23 emp24 L10intMYB DEAH10 (RPL32) gN7-MT1
Vatma-5.6
WRKY3 kinAt1g DUF1336 DREB2B GNL1a vSN13 PPR5g15 DUF1997 PE31 At5g39 PHO1h9 SPX4 cystgS1 Myb4
Vatma-4
WRKY3 kinAt1g DUF1336 unch1576 TM230 KIN10 CSC1H GNL1b vSN13 PE31 SPX4 cystgS1 Myb4 NAC92
Hopch-2F
WRKY3 DUF1336 SAT22 (sos) UXTh splicF3A AGO7 Topo 3 GNOM unch336 ILK1 emp24 DEAH10 (RPL32) gN7-MT1 (4x) GCK PHO1h3 O-fucT3
(2x)
Hopch-3F
WRKY3 kinAt1g DUF1336 KIN10 GNL1a vSN13 PPR5g15 DUF1997 PE31 SPX4 cystgS1 Myn4 NAC92
Hopch-68F
WRKY3 kinAt1g DUF1336 unch1576 TM230 KIN10 vSN13 GNL1b CSC1H PE31 At5g39 SPX4 At5g39 SPX4 cystgS1 Myb4 NAC92
(2x)
inversion
Thymelaeaceae
Aqusi-sc92
PPR2g02 SAT22 (sos) AGO7 DREB2B CSC1At1g 26SP11 Ar2g02 GNOM vSN12 ILK1 emp24 O-fucT3 RubiscoMT gal2fucT APCsu2
Aqusi-sc3
UCF_E4 FeSass1 choMut1 DUF3598 IBIS1 KIN10 PPR5g39 (PB1dom) GNL1 PPR3g58 PPR5g15 PE31 sugarCC At5g39 polyP-bp1
Aqusi-sc155
PPR3g29 lamin WRKY3 IBIS1 TM230 KIN10 DREB2B CSC1H vSN13 DUF1997 At5g39 PPR5g15 emp24 PHO1h9 SPX4 cystgS1 Myb4
(2x)
Stech-7
WRKY3 DUF1336 PRFGA-cl SAT22 unch330 (sos) AGO7 DREB2B CSC1At1g ACD11 GNOM vSN12 PPR5g44 ILK1 emp24 O-fucT3 RubiscoMT (RPL32) APCsu2 (SETdom) CDCapoR1
(2x)
Stech-1
UCF_E4 FeS ass choMut1 lamin DUF3598 IBIS1 KIN10 GNL1 PPR3g58 PPR5g39 PPR5g15 PE31 At5g39 pPyr-bp1 HEN4 WD43 (cul-bp)
Stech-6
DUF179 PPR3g29 lamin WRKY3 IBIS1 DUF1336 TM230 KIN10 DREB2B CSC1H vSN13 DUF1997 At5g39 PPR5g66 PHO1h9 SPX4 cystgS1 Myb4 NAC92
(2x)

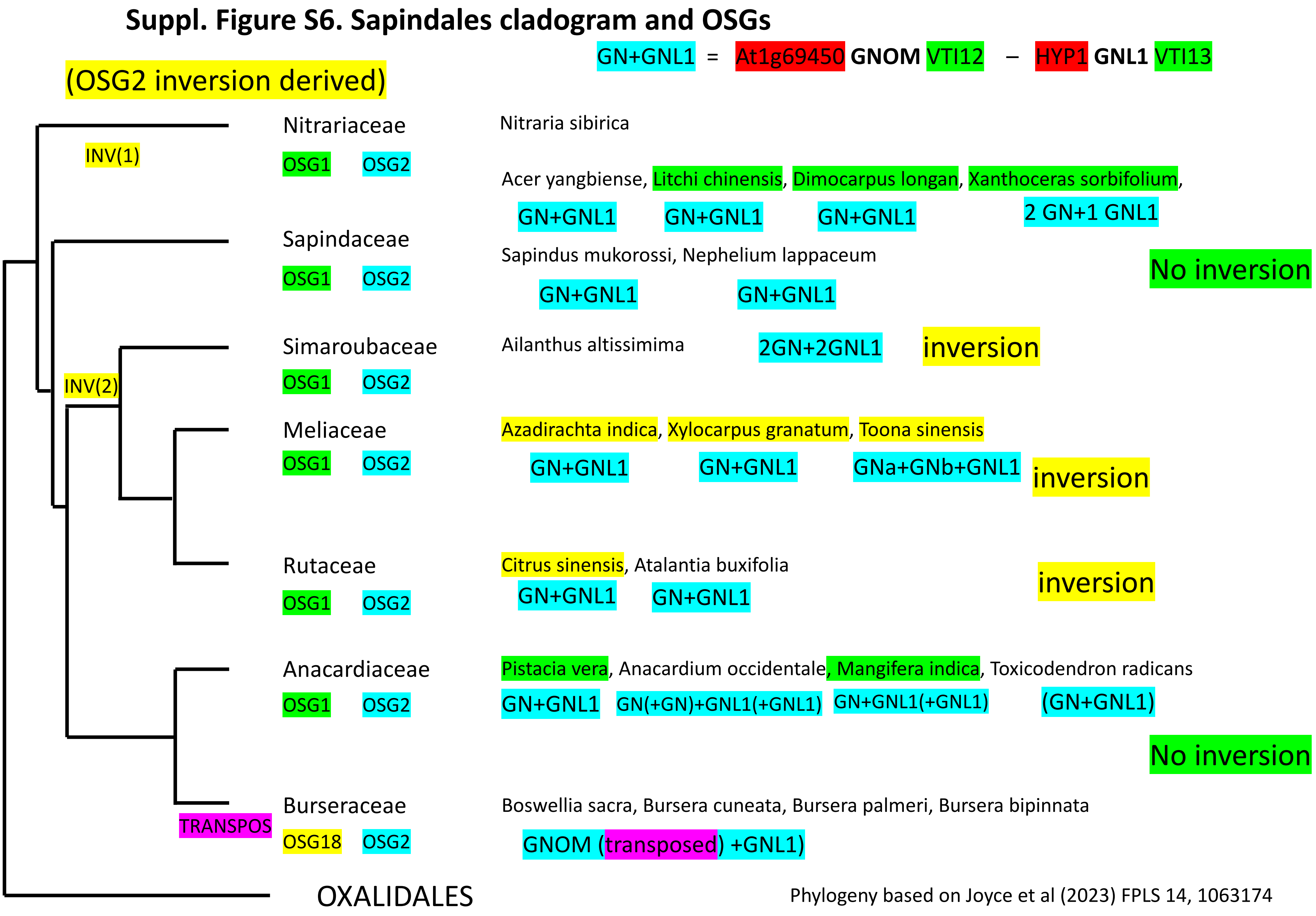

Suppl. Figure S6. Sapindales cladogram and OSGs
GN+GNL1 = At1g69450 GNOM VTI12 – HYP1 GNL1 VTI13
(OSG2 inversion derived)
Nitrariaceae
INV(1)
OSG1 OSG2
Nitraria sibirica
Acer yangbiense, Litchi chinensis, Dimocarpus longan, Xanthoceras sorbifolium,
GN+GNL1 GN+GNL1 GN+GNL1 2 GN+1 GNL1
Sapindaceae
OSG1 OSG2
Sapindus mukorossi, Nephelium lappaceum
GN+GNL1 GN+GNL1
No inversion
Simaroubaceae
INV(2)
OSG1 OSG2
Ailanthus altissimima
2GN+2GNL1
inversion
Meliaceae
OSG1 OSG2
Azadirachta indica, Xylocarpus granatum, Toona sinensis
GN+GNL1 GN+GNL1 GNa+GNb+GNL1
inversion
Rutaceae
OSG1 OSG2
Citrus sinensis, Atalantia buxifolia
GN+GNL1 GN+GNL1
inversion
Anacardiaceae
OSG1 OSG2
Pistacia vera, Anacardium occidentale, Mangifera indica, Toxicodendron radicans
GN+GNL1 GN(+GN)+GNL1(+GNL1) GN+GNL1(+GNL1) (GN+GNL1)
No inversion
Burseraceae
TRANSPOS
OSG18 OSG2
Boswellia sacra, Bursera cuneata, Bursera palmeri, Bursera bipinnata
GNOM (transposed) +GNL1)
OXALIDALES
Phylogeny based on Joyce et al (2023) FPLS 14, 1063174

**Sapindales** Poncirus trifoliata & Murraya koenigii (Rutaceae) GNL1 synteny inversion, too!

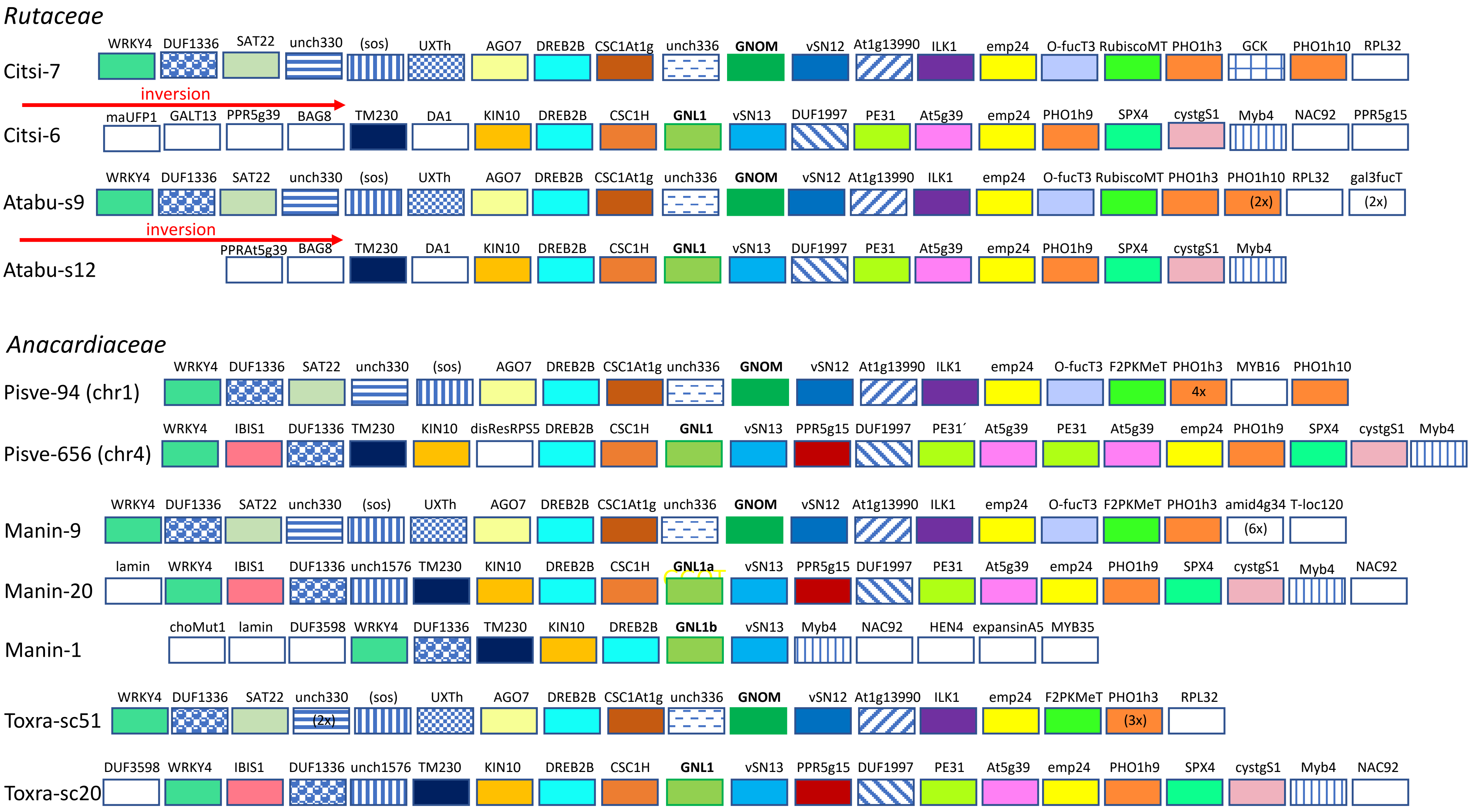

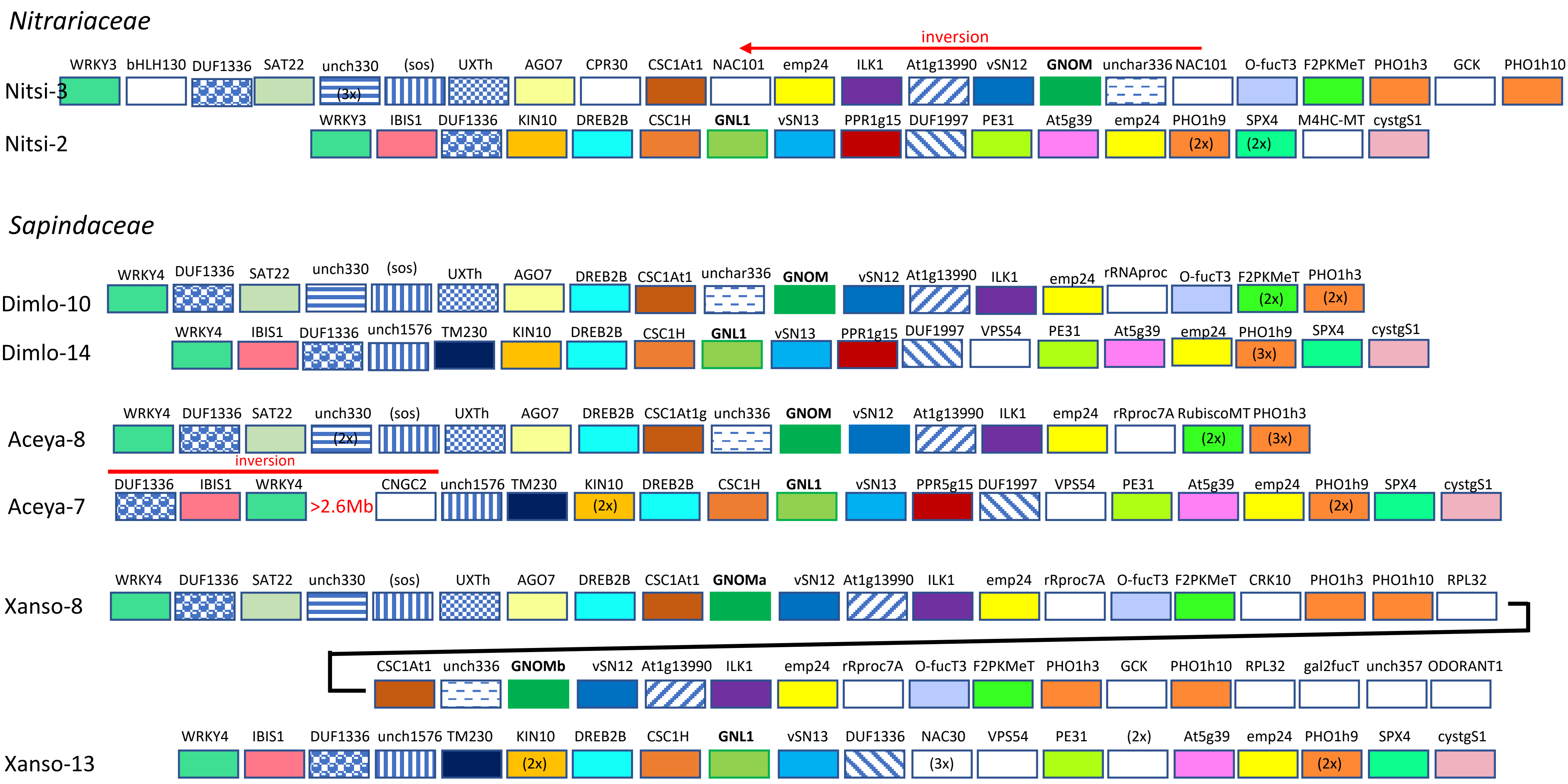
Nitrariaceae
inversion
Nitsi-3
WRKY3 bHLH130 DUF1336 SAT22 unch330 (3x) (sos) UXTh AGO7 CPR30 CSC1At1 NAC101 emp24 ILK1 At1g13990 vSN12 GNOM unchar336 NAC101 O-fucT3 F2PKMeT PHO1h3 GCK PHO1h10
Nitsi-2
WRKY3 IBIS1 DUF1336 KIN10 DREB2B CSC1H GNL1 vSN13 PPR1g15 DUF1997 PE31 At5g39 emp24 PHO1h9 (2x) SPX4 (2x) M4HC-MT cystgS1
Sapindaceae
Dimlo-10
WRKY4 DUF1336 SAT22 unch330 (sos) UXTh AGO7 DREB2B CSC1At1 unchar336 GNOM vSN12 At1g13990 ILK1 emp24 rRNAproc O-fucT3 F2PKMeT (2x) PHO1h3 (2x)
Dimlo-14
WRKY4 IBIS1 DUF1336 unch1576 TM230 KIN10 DREB2B CSC1H GNL1 vSN13 PPR1g15 DUF1997 VPS54 PE31 At5g39 emp24 PHO1h9 (3x) SPX4 cystgS1
Aceya-8
WRKY4 DUF1336 SAT22 unch330 (2x) (sos) UXTh AGO7 DREB2B CSC1At1g unch336 GNOM vSN12 At1g13990 ILK1 emp24 rRproc7A RubiscoMT (2x) PHO1h3 (3x)
inversion
Aceya-7
DUF1336 IBIS1 WRKY4 >2.6Mb CNGC2 unch1576 TM230 KIN10 (2x) DREB2B CSC1H GNL1 vSN13 PPR5g15 DUF1997 VPS54 PE31 At5g39 emp24 PHO1h9 (2x) SPX4 cystgS1
Xanso-8
WRKY4 DUF1336 SAT22 unch330 (sos) UXTh AGO7 DREB2B CSC1At1 GNOMa vSN12 At1g13990 ILK1 emp24 rRproc7A O-fucT3 F2PKMeT CRK10 PHO1h3 PHO1h10 RPL32
CSC1At1 unch336 GNOMb vSN12 At1g13990 ILK1 emp24 rRproc7A O-fucT3 F2PKMeT PHO1h3 GCK PHO1h10 RPL32 gal2fucT unch357 ODORANT1
Xanso-13
WRKY4 IBIS1 DUF1336 unch1576 TM230 KIN10 (2x) DREB2B CSC1H GNL1 vSN13 DUF1336 NAC30 (3x) VPS54 PE31 (2x) At5g39 emp24 PHO1h9 (2x) SPX4 cystgS1

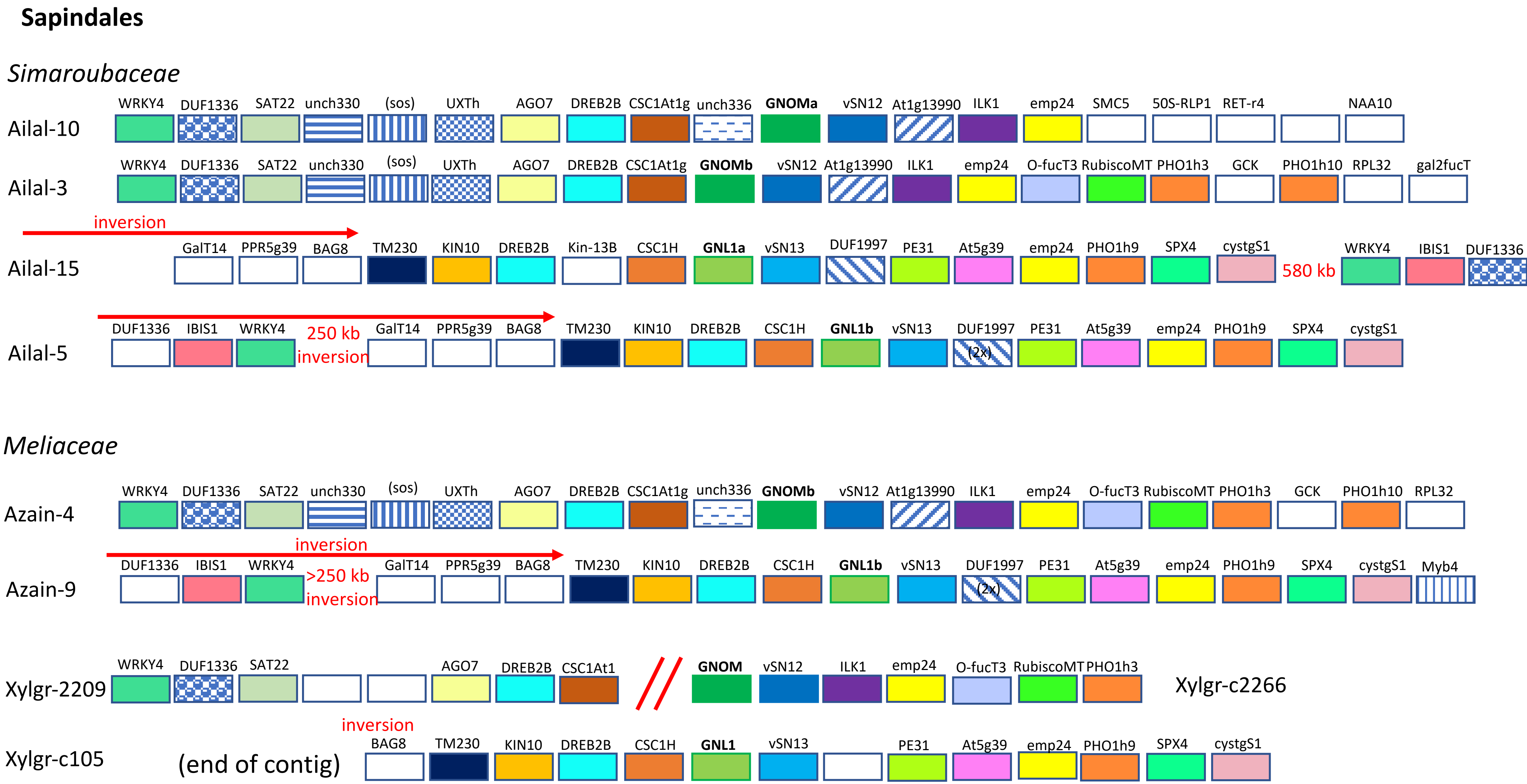
Sapindales
Simaroubaceae
Ailal-10
WRKY4 DUF1336 SAT22 unch330 (sos) UXTh AGO7 DREB2B CSC1At1g unch336 GNOMa vSN12 At1g13990 ILK1 emp24 SMC5 50S-RLP1 RET-r4 NAA10
Ailal-3
WRKY4 DUF1336 SAT22 unch330 (sos) UXTh AGO7 DREB2B CSC1At1g GNOMb vSN12 At1g13990 ILK1 emp24 O-fucT3 RubiscoMT PHO1h3 GCK PHO1h10 RPL32 gal2fucT
inversion
Ailal-15
GalT14 PPR5g39 BAG8 TM230 KIN10 DREB2B Kin-13B CSC1H GNL1a vSN13 DUF1997 PE31 At5g39 emp24 PHO1h9 SPX4 cystgS1 580 kb WRKY4 IBIS1 DUF1336
Ailal-5
DUF1336 IBIS1 WRKY4 250 kb inversion GalT14 PPR5g39 BAG8 TM230 KIN10 DREB2B CSC1H GNL1b vSN13 DUF1997 (2x) PE31 At5g39 emp24 PHO1h9 SPX4 cystgS1
Meliaceae
Azain-4
WRKY4 DUF1336 SAT22 unch330 (sos) UXTh AGO7 DREB2B CSC1At1g unch336 GNOMb vSN12 At1g13990 ILK1 emp24 O-fucT3 RubiscoMT PHO1h3 GCK PHO1h10 RPL32
inversion
Azain-9
DUF1336 IBIS1 WRKY4 >250 kb inversion GalT14 PPR5g39 BAG8 TM230 KIN10 DREB2B CSC1H GNL1b vSN13 DUF1997 (2x) PE31 At5g39 emp24 PHO1h9 SPX4 cystgS1 Myb4
Xylgr-2209
WRKY4 DUF1336 SAT22 AGO7 DREB2B CSC1At1 GNOM vSN12 ILK1 emp24 O-fucT3 RubiscoMT PHO1h3
Xylgr-c2266
inversion
Xylgr-c105
(end of contig)
BAG8 TM230 KIN10 DREB2B CSC1H GNL1 vSN13 PE31 At5g39 emp24 PHO1h9 SPX4 cystgS1

### GNOM transposed > OSG_Theca-chr9, Mb 9.9

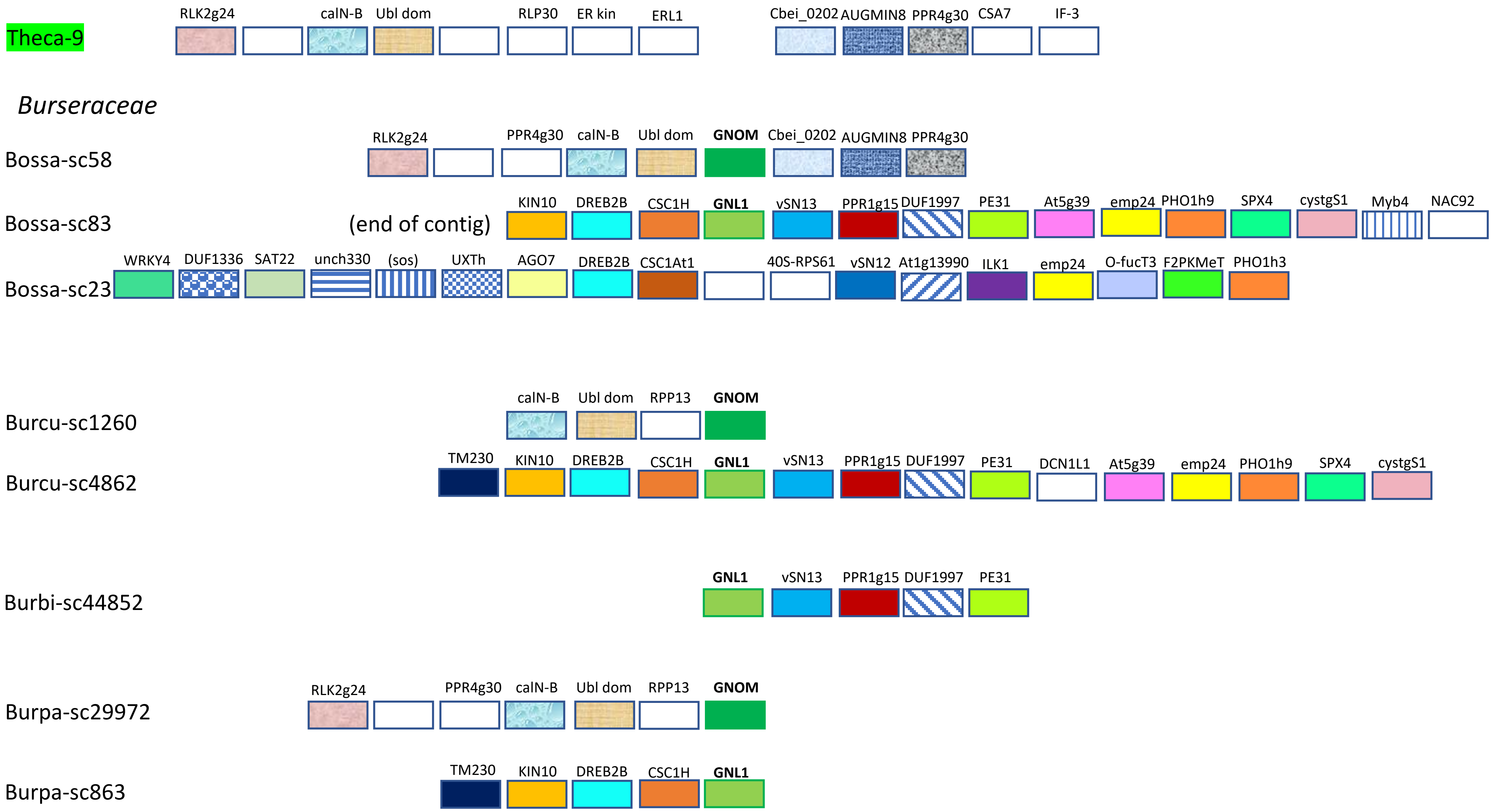

**Suppl.Figure S7. Oxalidales cladogram and OSGs**

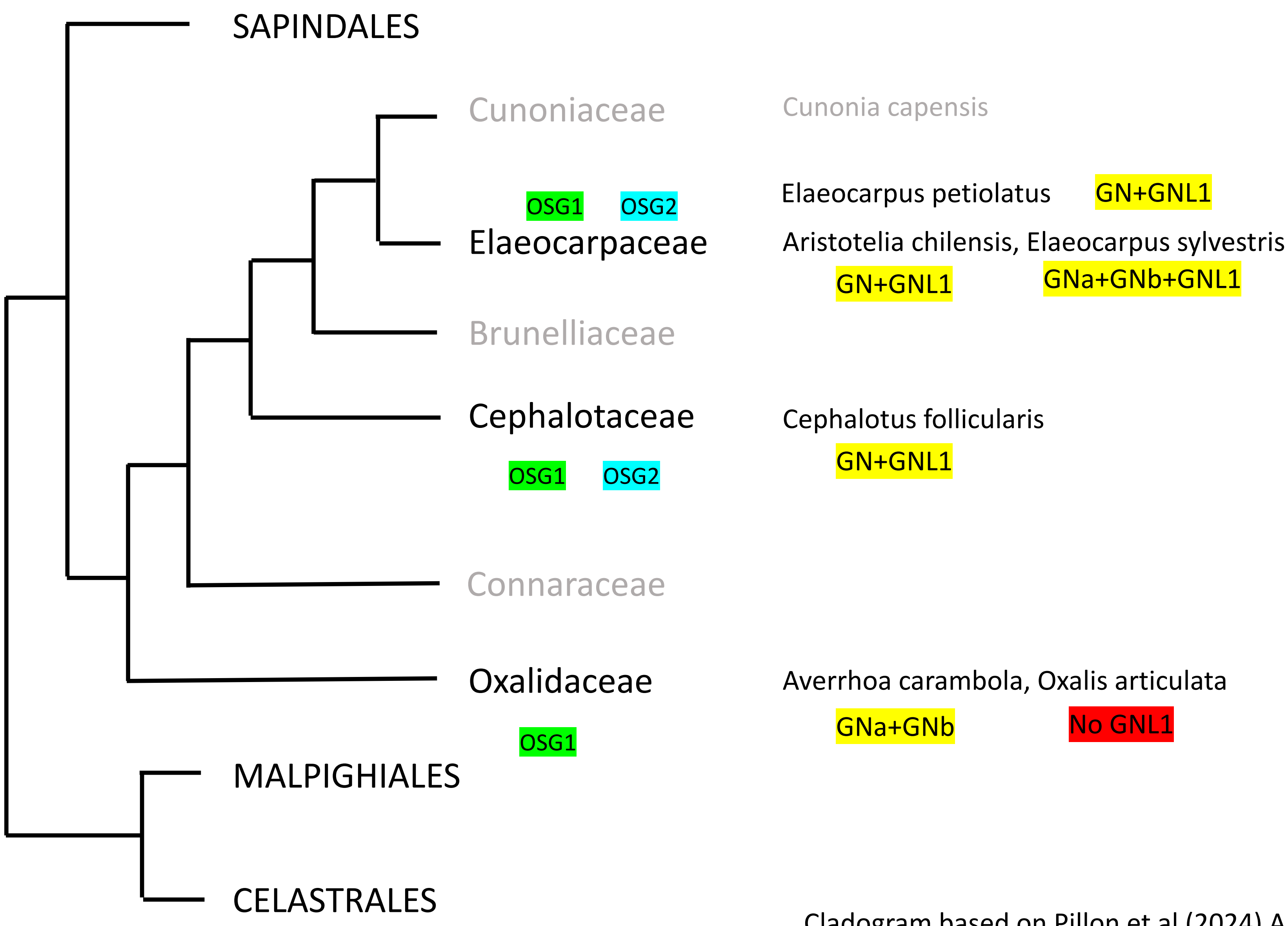

Cladogram based on Pillon et al (2024) Am J Bot 108, 1181

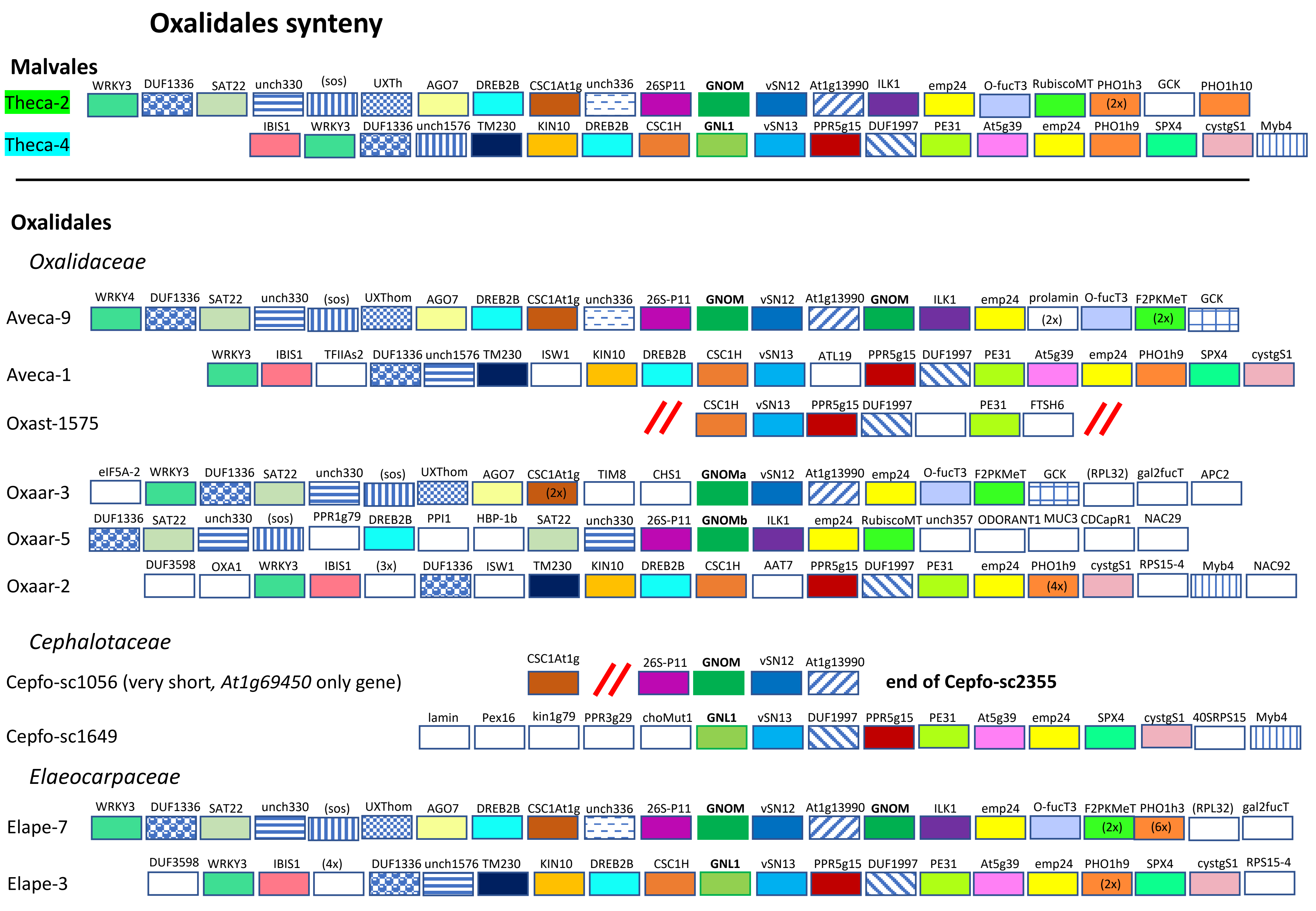
Oxalidales synteny
Malvales
Theca-2
WRKY3 DUF1336 SAT22 unch330 (sos) UXTh AGO7 DREB2B CSC1At1g unch336 26SP11 GNOM vSN12 At1g13990 ILK1 emp24 O-fucT3 RubiscoMT PHO1h3 GCK PHO1h10
(2x)
Theca-4
IBIS1 WRKY3 DUF1336 unch1576 TM230 KIN10 DREB2B CSC1H GNL1 vSN13 PPR5g15 DUF1997 PE31 At5g39 emp24 PHO1h9 SPX4 cystgS1 Myb4
Oxalidales
Oxalidaceae
Aveca-9
WRKY4 DUF1336 SAT22 unch330 (sos) UXThom AGO7 DREB2B CSC1At1g unch336 26S-P11 GNOM vSN12 At1g13990 GNOM ILK1 emp24 prolamin O-fucT3 F2PKMeT GCK
(2x) (2x)
Aveca-1
WRKY3 IBIS1 TFIIAs2 DUF1336 unch1576 TM230 ISW1 KIN10 DREB2B CSC1H vSN13 ATL19 PPR5g15 DUF1997 PE31 At5g39 emp24 PHO1h9 SPX4 cystgS1
Oxast-1575
CSC1H vSN13 PPR5g15 DUF1997 PE31 FTSH6
Oxaar-3
eIF5A-2 WRKY3 DUF1336 SAT22 unch330 (sos) UXThom AGO7 CSC1At1g TIM8 CHS1 GNOMa vSN12 At1g13990 emp24 O-fucT3 F2PKMeT GCK (RPL32) gal2fucT APC2
(2x)
Oxaar-5
DUF1336 SAT22 unch330 (sos) PPR1g79 DREB2B PPI1 HBP-1b SAT22 unch330 26S-P11 GNOMb ILK1 emp24 RubiscoMT unch357 ODORANT1 MUC3 CDCapR1 NAC29
Oxaar-2
DUF3598 OXA1 WRKY3 IBIS1 (3x) DUF1336 ISW1 TM230 KIN10 DREB2B CSC1H AAT7 PPR5g15 DUF1997 PE31 emp24 PHO1h9 cystgS1 RPS15-4 Myb4 NAC92
(4x)
Cephalotaceae
Cepfo-sc1056 (very short, At1g69450 only gene)
CSC1At1g 26S-P11 GNOM vSN12 At1g13990
end of Cepfo-sc2355
Cepfo-sc1649
lamin Pex16 kin1g79 PPR3g29 choMut1 GNL1 vSN13 DUF1997 PPR5g15 PE31 At5g39 emp24 SPX4 cystgS1 40SRPS15 Myb4
Elaeocarpaceae
Elape-7
WRKY3 DUF1336 SAT22 unch330 (sos) UXThom AGO7 DREB2B CSC1At1g unch336 26S-P11 GNOM vSN12 At1g13990 GNOM ILK1 emp24 O-fucT3 F2PKMeT PHO1h3 (RPL32) gal2fucT
(2x) (6x)
Elape-3
DUF3598 WRKY3 IBIS1 (4x) DUF1336 unch1576 TM230 KIN10 DREB2B CSC1H GNL1 vSN13 PPR5g15 DUF1997 PE31 At5g39 emp24 PHO1h9 SPX4 cystgS1 RPS15-4
(2x)

#### Suppl. Figure S8. Malpighiales cladogram and OSGs

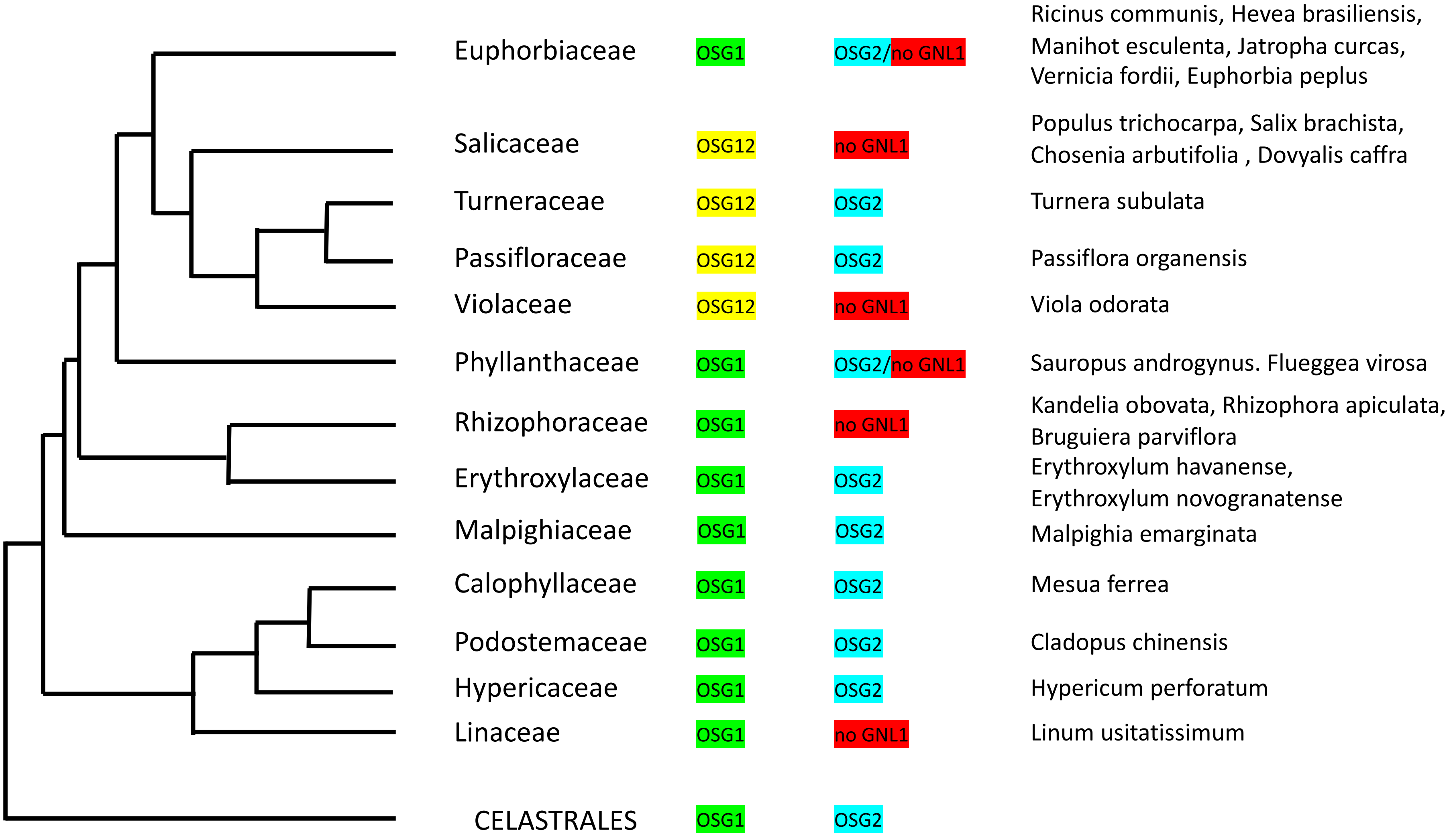

Cladogram based on Cai et al., 2021, Syst Biol 70, 491

### OSG12 Malpighiales (Salicaceae, Passifloraceae, Turneraceae, Violaceae)

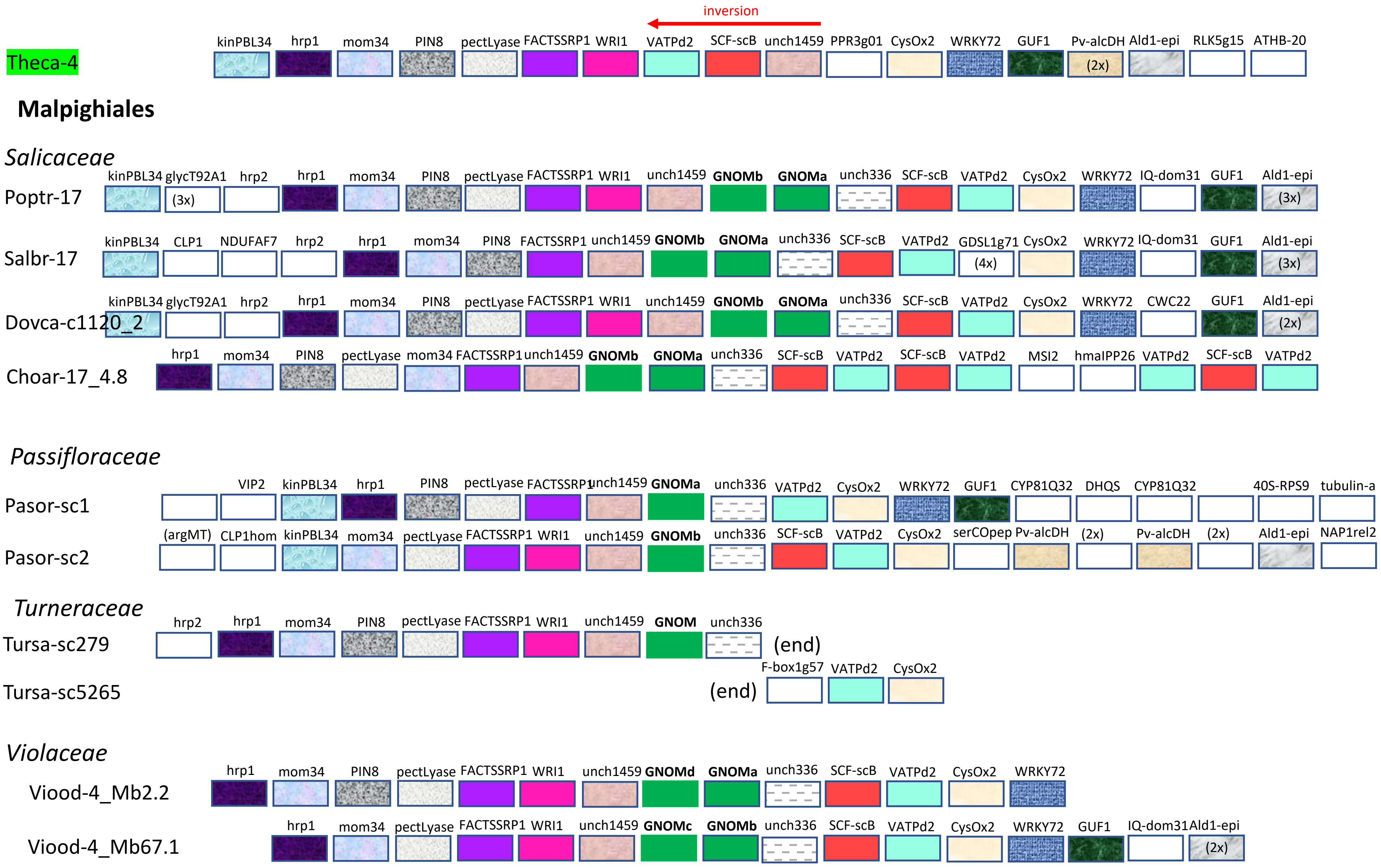

### Loss of GNOM from OSG1 in Malpighiales (Salicaceae, Passifloraceae, Turneraceae, Violaceae)

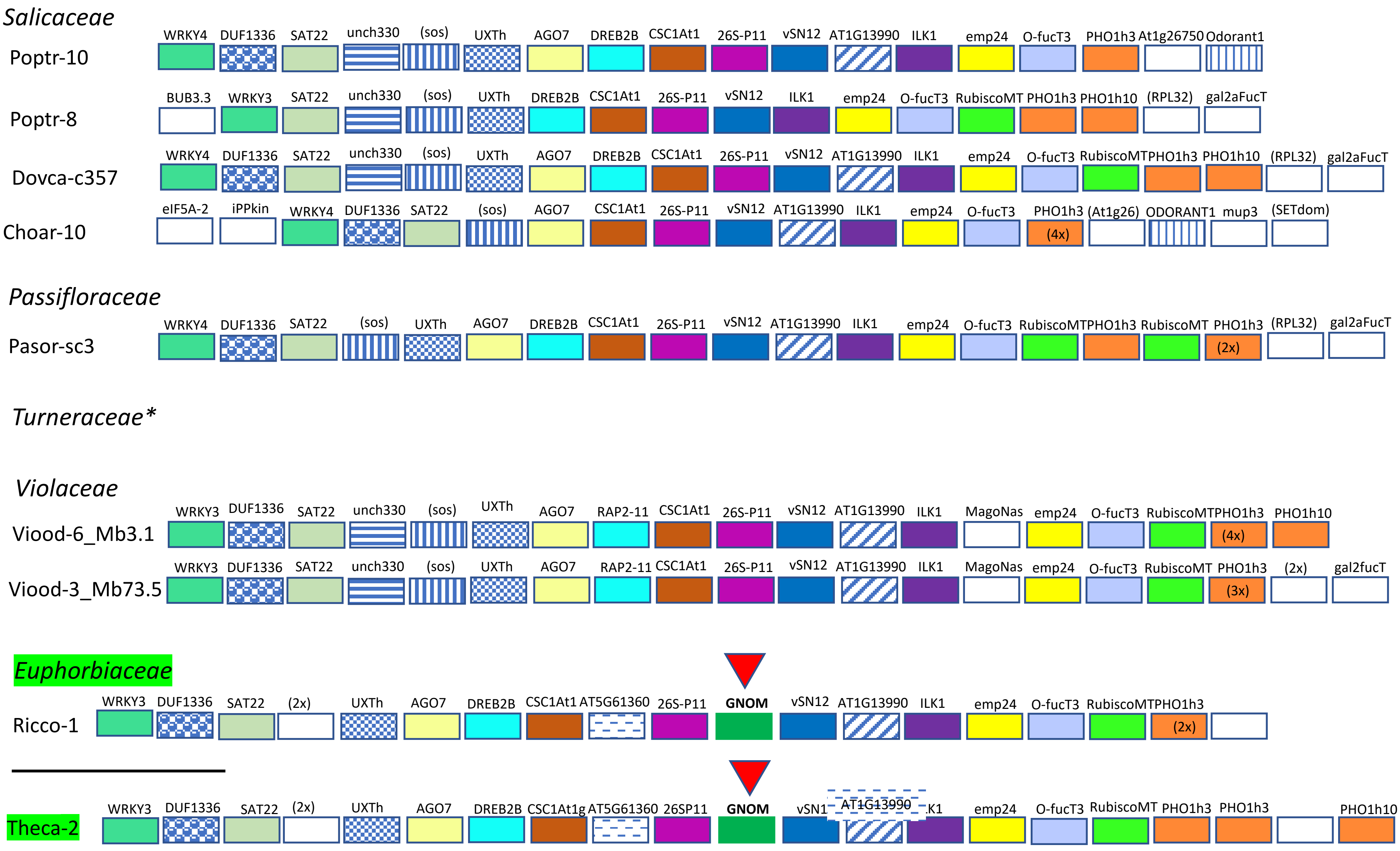

* Only fragments of OSG1 available (short scaffolds)

### OSG2 in Malpighiales

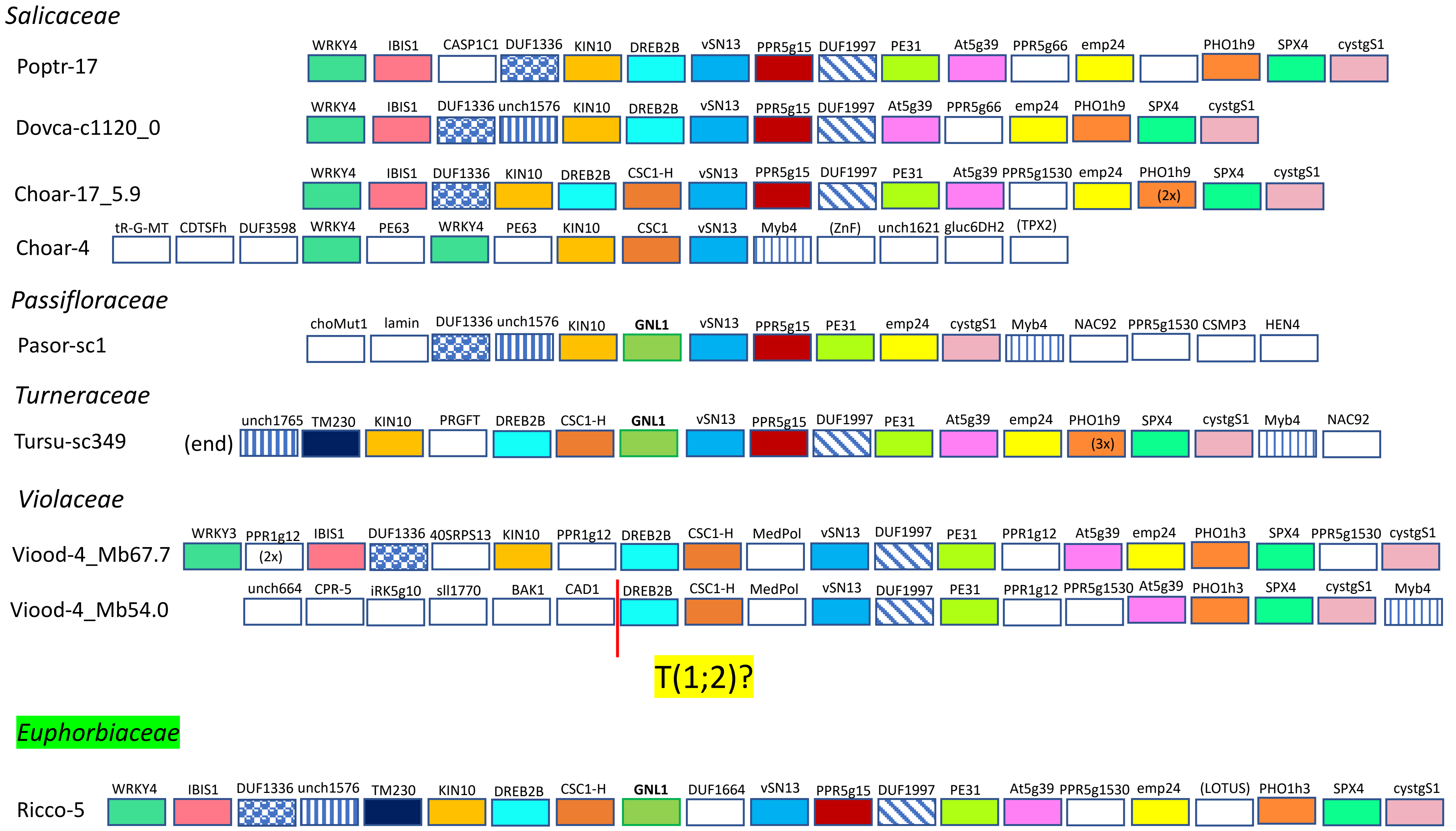

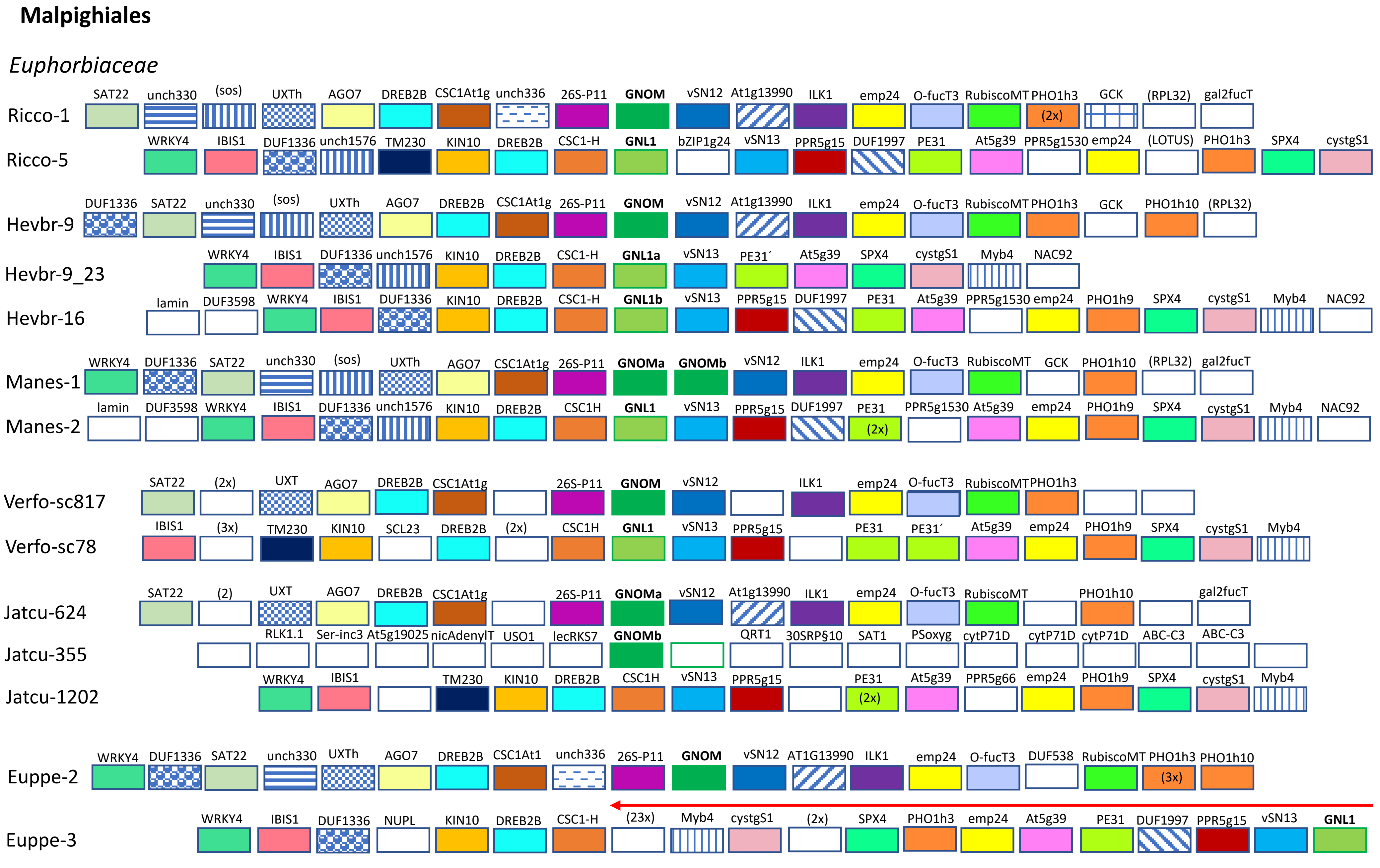

JatcuGNOMb transposed

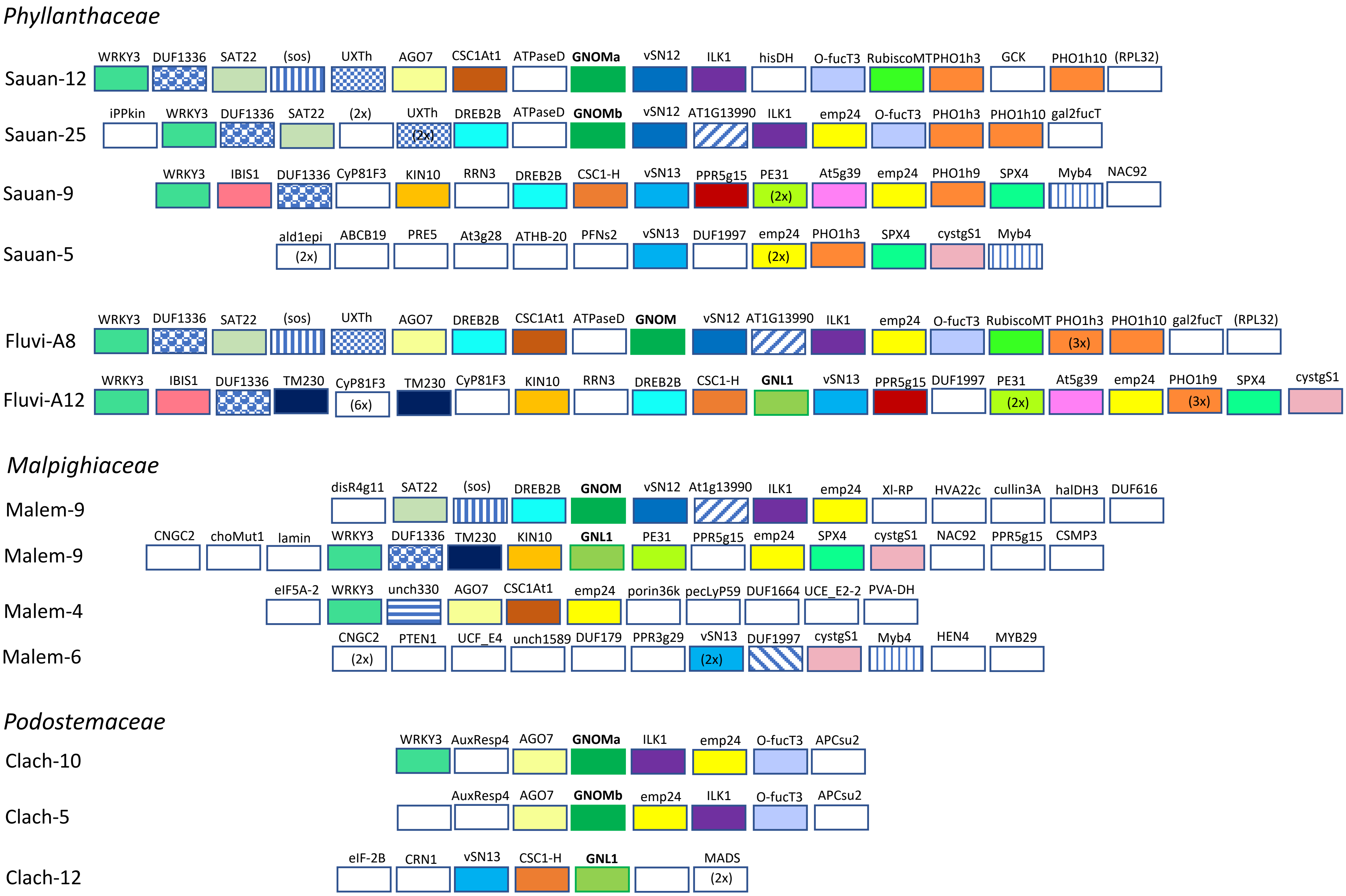
Phyllanthaceae
Sauan-12
WRKY3 DUF1336 SAT22 (sos) UXTh AGO7 CSC1At1 ATPaseD GNOMa vSN12 ILK1 hisDH O-fucT3 RubiscoMT PHO1h3 GCK PHO1h10 (RPL32)
Sauan-25
iPPkin WRKY3 DUF1336 SAT22 (2x) UXTh (2x) DREB2B ATPaseD GNOMb vSN12 AT1G13990 ILK1 emp24 O-fucT3 PHO1h3 PHO1h10 gal2fucT
Sauan-9
WRKY3 IBIS1 DUF1336 CyP81F3 KIN10 RRN3 DREB2B CSC1-H vSN13 PPR5g15 PE31 (2x) At5g39 emp24 PHO1h9 SPX4 Myb4 NAC92
Sauan-5
ald1epi (2x) ABCB19 PRE5 At3g28 ATHB-20 PFNs2 vSN13 DUF1997 emp24 (2x) PHO1h3 SPX4 cystgS1 Myb4
Fluvi-A8
WRKY3 DUF1336 SAT22 (sos) UXTh AGO7 DREB2B CSC1At1 ATPaseD GNOM vSN12 AT1G13990 ILK1 emp24 O-fucT3 RubiscoMT PHO1h3 (3x) PHO1h10 gal2fucT (RPL32)
Fluvi-A12
WRKY3 IBIS1 DUF1336 TM230 CyP81F3 (6x) TM230 CyP81F3 KIN10 RRN3 DREB2B CSC1-H GNL1 vSN13 PPR5g15 DUF1997 PE31 (2x) At5g39 emp24 PHO1h9 (3x) SPX4 cystgS1
Malpighiaceae
Malem-9
disR4g11 SAT22 (sos) DREB2B GNOM vSN12 At1g13990 ILK1 emp24 Xl-RP HVA22c cullin3A halDH3 DUF616
Malem-9
CNGC2 choMut1 lamin WRKY3 DUF1336 TM230 KIN10 GNL1 PE31 PPR5g15 emp24 SPX4 cystgS1 NAC92 PPR5g15 CSMP3
Malem-4
eIF5A-2 WRKY3 unch330 AGO7 CSC1At1 emp24 porin36k pecLyP59 DUF1664 UCE_E2-2 PVA-DH
Malem-6
CNGC2 (2x) PTEN1 UCF_E4 unch1589 DUF179 PPR3g29 vSN13 (2x) DUF1997 cystgS1 Myb4 HEN4 MYB29
Podostemaceae
Clach-10
WRKY3 AuxResp4 AGO7 GNOMa ILK1 emp24 O-fucT3 APCsu2
Clach-5
AuxResp4 AGO7 GNOMb emp24 ILK1 O-fucT3 APCsu2
Clach-12
eIF-2B CRN1 vSN13 CSC1-H GNL1 MADS (2x)

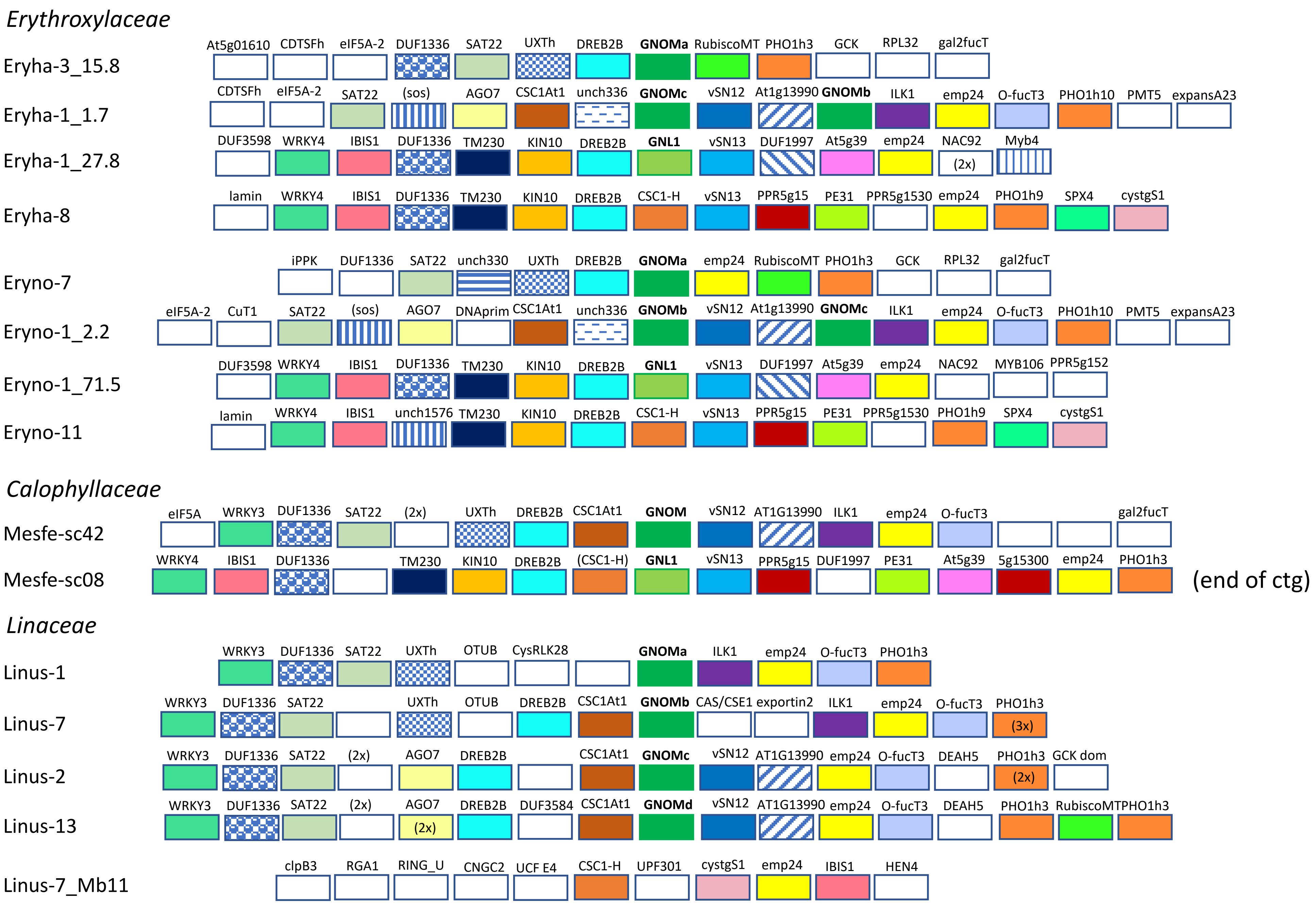
Erythroxylaceae
Eryha-3_15.8
At5g01610 CDTSFh eIF5A-2 DUF1336 SAT22 UXTh DREB2B GNOMa RubiscoMT PHO1h3 GCK RPL32 gal2fucT
Eryha-1_1.7
CDTSFh eIF5A-2 SAT22 (sos) AGO7 CSC1At1 unch336 GNOMc vSN12 At1g13990 GNOMb ILK1 emp24 O-fucT3 PHO1h10 PMT5 expansA23
Eryha-1_27.8
DUF3598 WRKY4 IBIS1 DUF1336 TM230 KIN10 DREB2B GNL1 vSN13 DUF1997 At5g39 emp24 NAC92 (2x) Myb4
Eryha-8
lamin WRKY4 IBIS1 DUF1336 TM230 KIN10 DREB2B CSC1-H vSN13 PPR5g15 PE31 PPR5g1530 emp24 PHO1h9 SPX4 cystgS1
Eryno-7
iPPK DUF1336 SAT22 unch330 UXTh DREB2B GNOMa emp24 RubiscoMT PHO1h3 GCK RPL32 gal2fucT
Eryno-1_2.2
eIF5A-2 CuT1 SAT22 (sos) AGO7 DNAprim CSC1At1 unch336 GNOMb vSN12 At1g13990 GNOMc ILK1 emp24 O-fucT3 PHO1h10 PMT5 expansA23
Eryno-1_71.5
DUF3598 WRKY4 IBIS1 DUF1336 TM230 KIN10 DREB2B GNL1 vSN13 DUF1997 At5g39 emp24 NAC92 MYB106 PPR5g152
Eryno-11
lamin WRKY4 IBIS1 unch1576 TM230 KIN10 DREB2B CSC1-H vSN13 PPR5g15 PE31 PPR5g1530 PHO1h9 SPX4 cystgS1
Calophyllaceae
Mesfe-sc42
eIF5A WRKY3 DUF1336 SAT22 (2x) UXTh DREB2B CSC1At1 GNOM vSN12 AT1G13990 ILK1 emp24 O-fucT3 gal2fucT
Mesfe-sc08
WRKY4 IBIS1 DUF1336 TM230 KIN10 DREB2B (CSC1-H) GNL1 vSN13 PPR5g15 DUF1997 PE31 At5g39 5g15300 emp24 PHO1h3
(end of ctg)
Linaceae
Linus-1
WRKY3 DUF1336 SAT22 UXTh OTUB CysRLK28 GNOMa ILK1 emp24 O-fucT3 PHO1h3
Linus-7
WRKY3 DUF1336 SAT22 UXTh OTUB DREB2B CSC1At1 GNOMb CAS/CSE1 exportin2 ILK1 emp24 O-fucT3 PHO1h3 (3x)
Linus-2
WRKY3 DUF1336 SAT22 (2x) AGO7 DREB2B CSC1At1 GNOMc vSN12 AT1G13990 emp24 O-fucT3 DEAH5 PHO1h3 (2x) GCK dom
Linus-13
WRKY3 DUF1336 SAT22 (2x) AGO7 (2x) DREB2B DUF3584 CSC1At1 GNOMd vSN12 AT1G13990 emp24 O-fucT3 DEAH5 PHO1h3 RubiscoMTPHO1h3
Linus-7_Mb11
clpB3 RGA1 RING_U CNGC2 UCF E4 CSC1-H UPF301 cystgS1 emp24 IBIS1 HEN4

**Malpighiales**

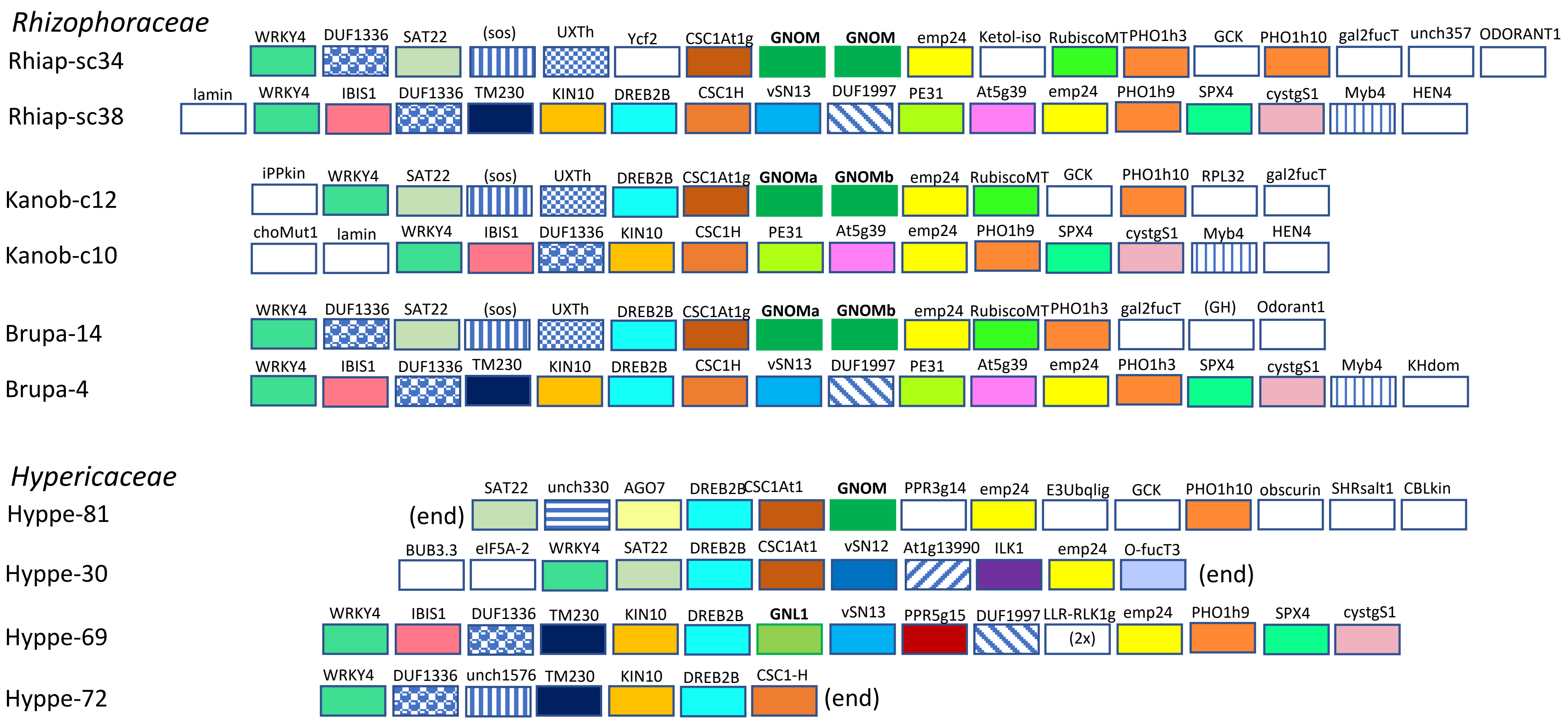

**Suppl. Figure S9. Celastrales cladogram and OSGs**

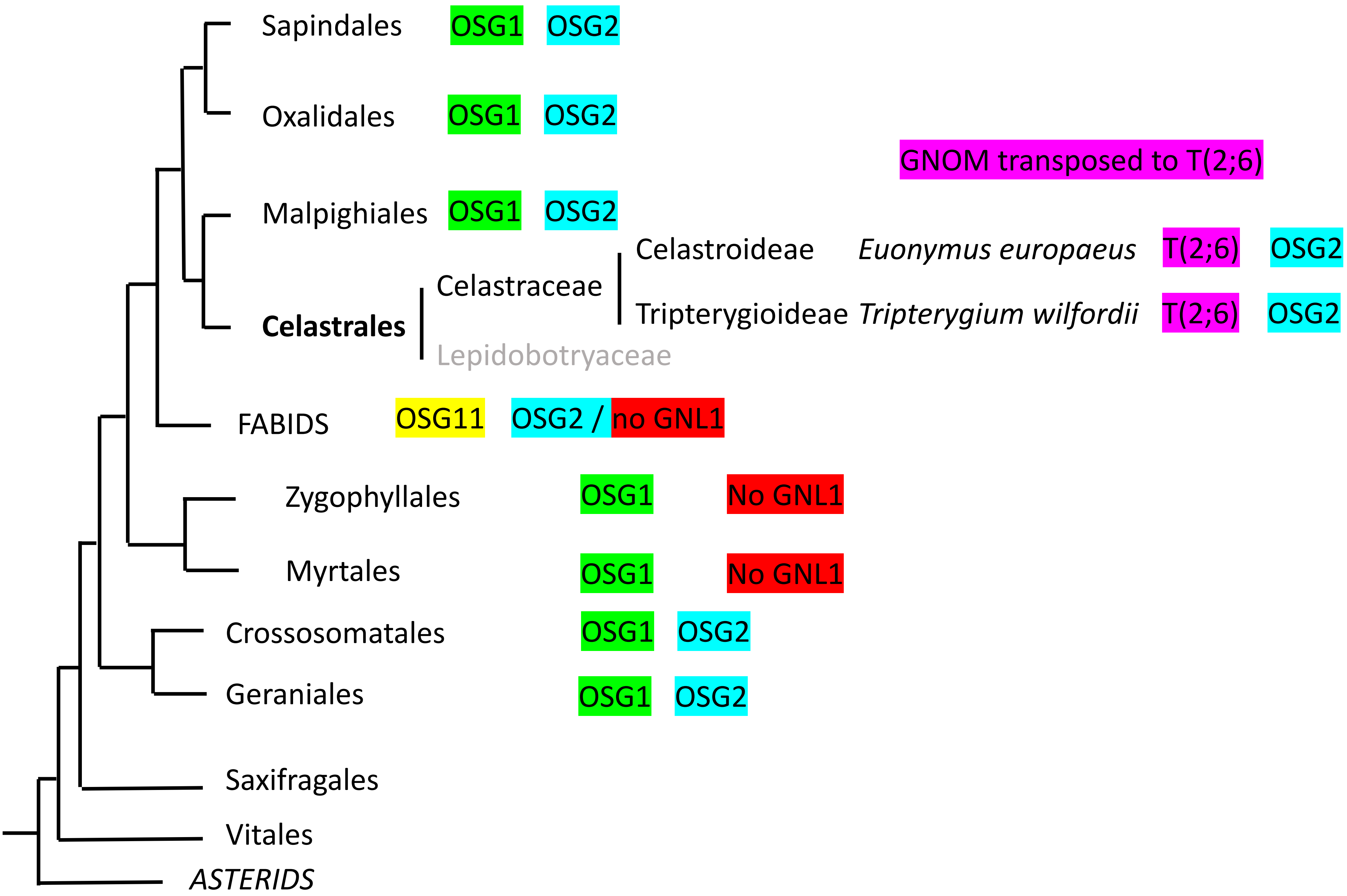

### Celastrales synteny

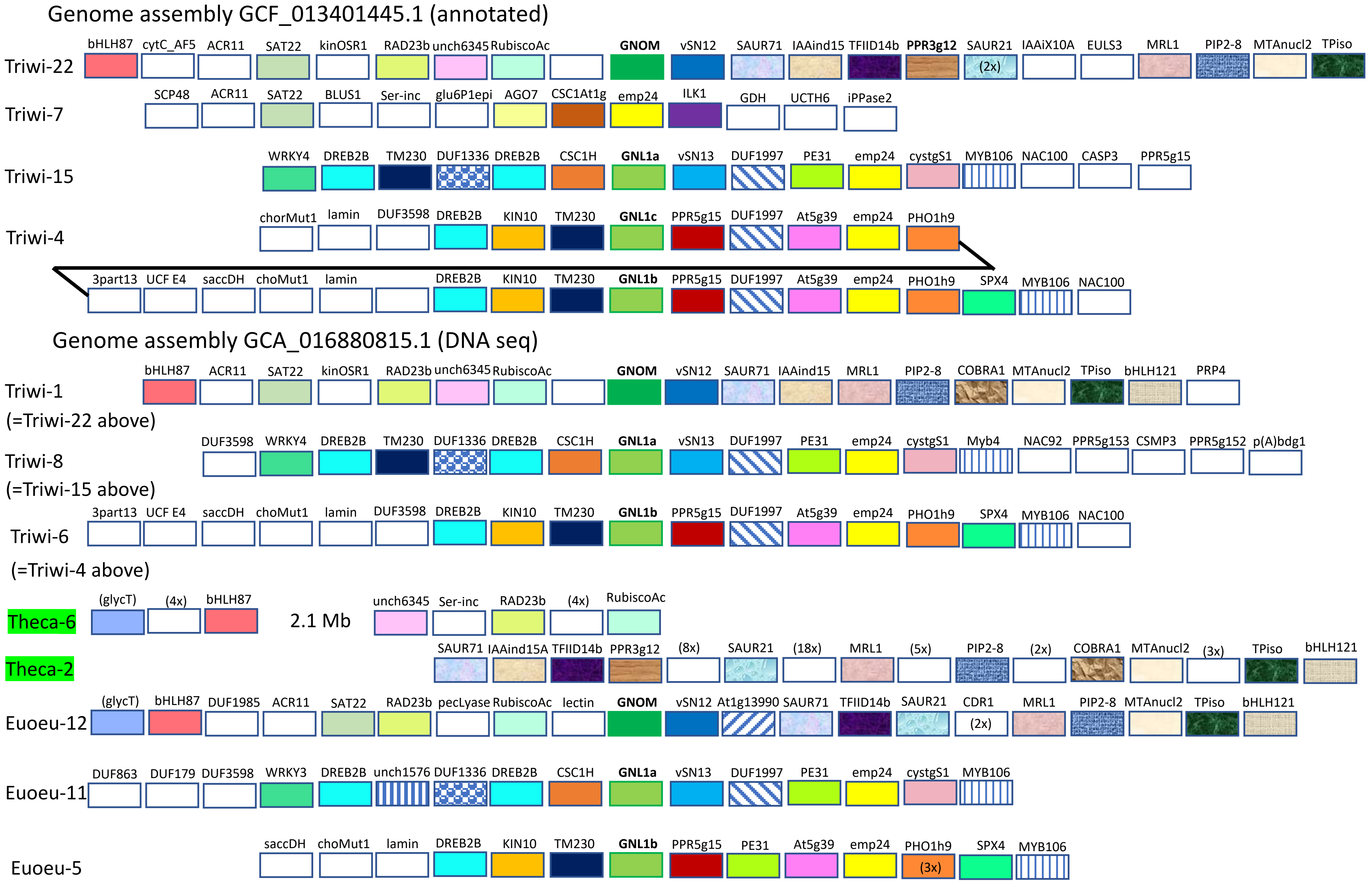

**Suppl. Figure S10. Fagales cladogram and OSGs**

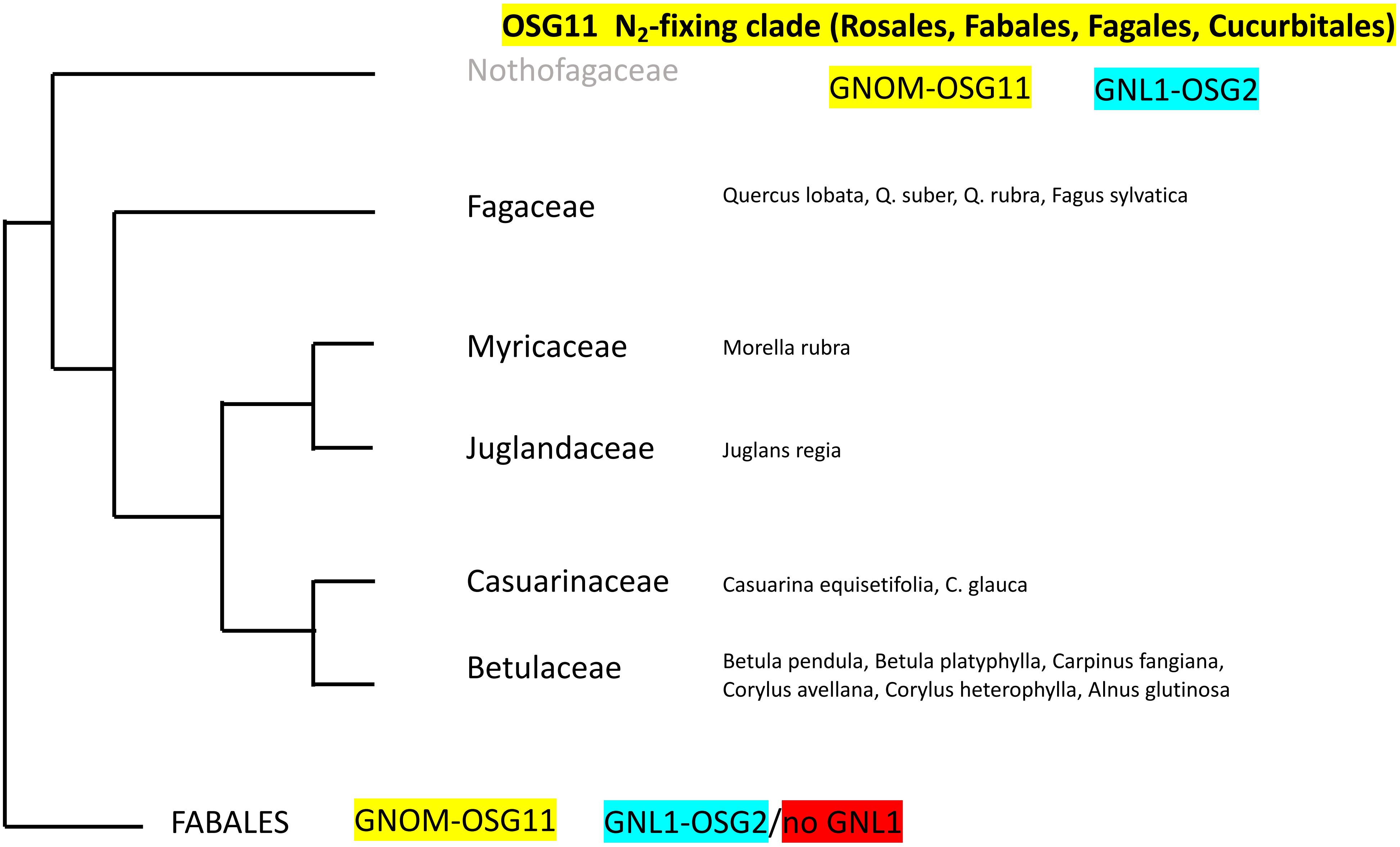

Cladogram based on Zuntini et al. (2024)

**Suppl. Figure S10.** AtGNOM ortholog transposition in **Fagales**

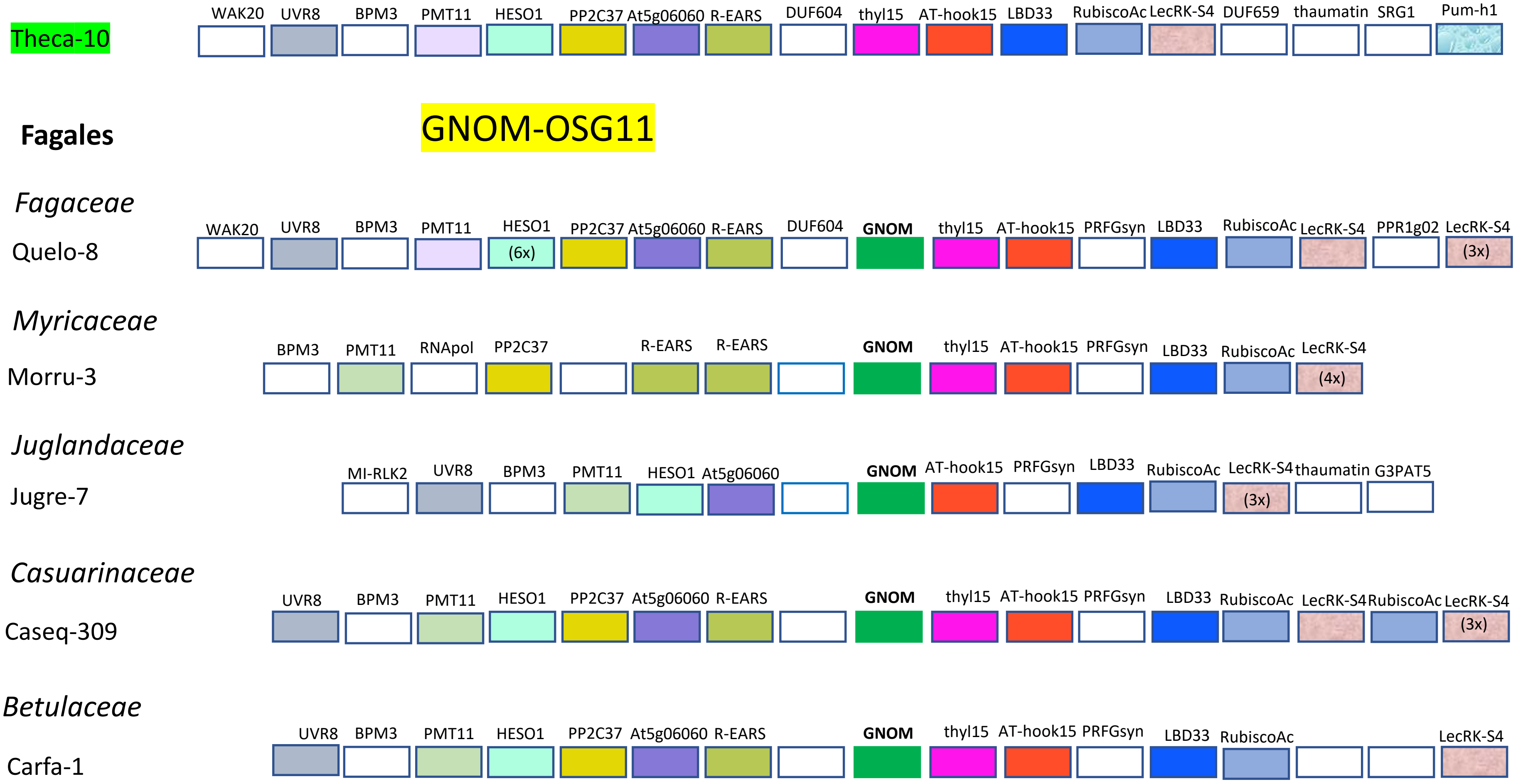

### Loss of GNOM from OSG1 ($N_2$-fixing clade)

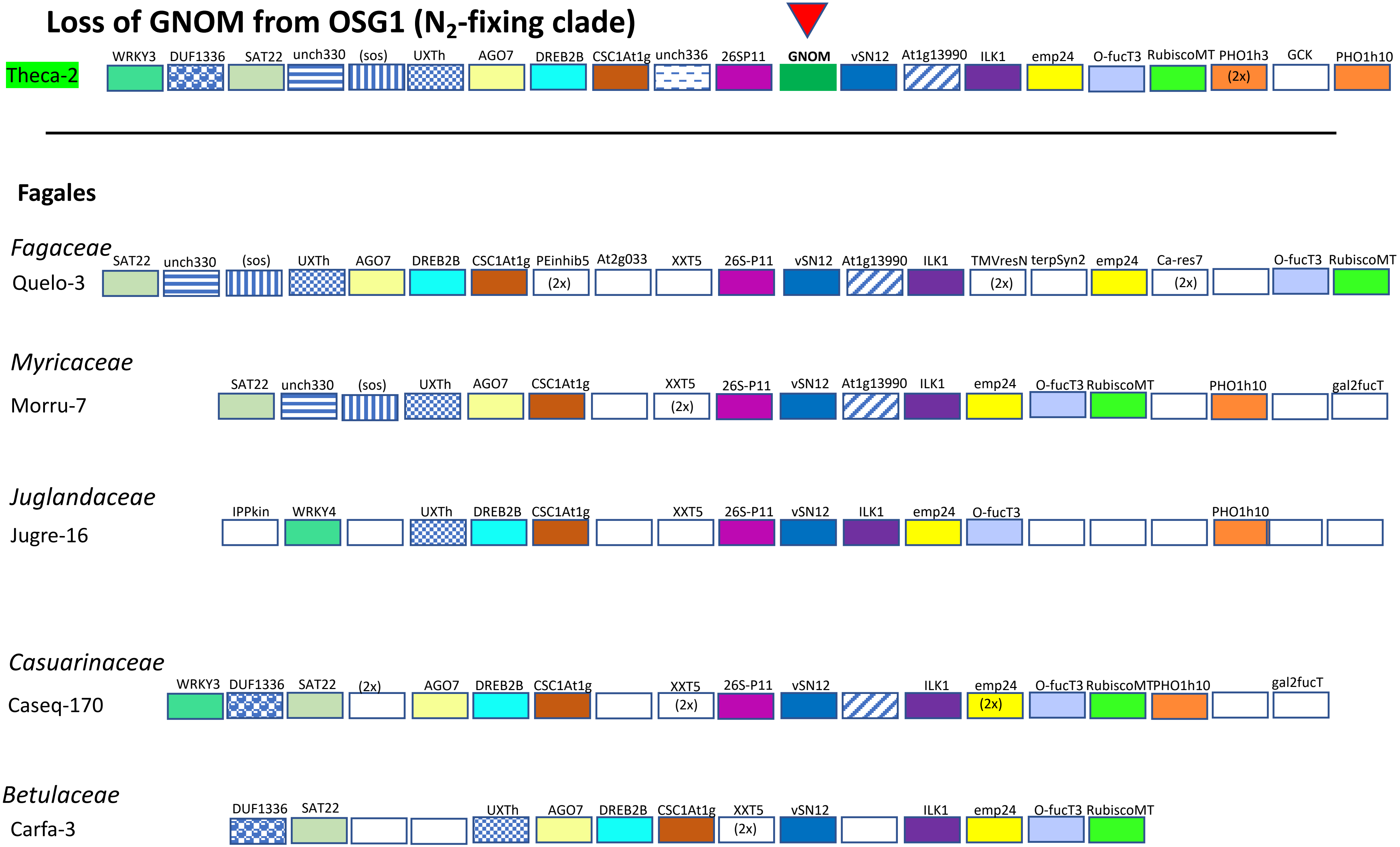

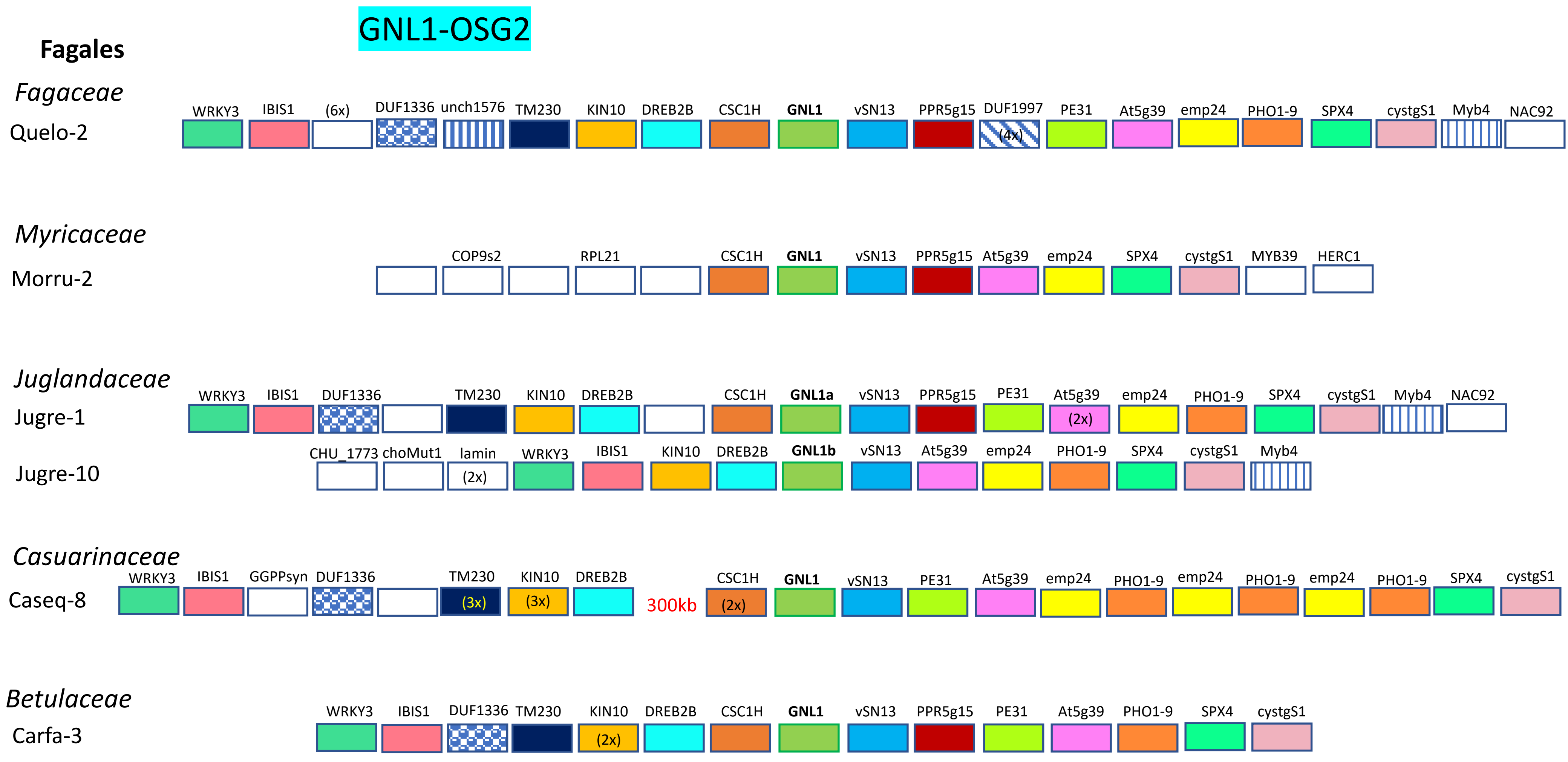
GNL1-OSG2
Fagales
Fagaceae
Quelo-2
WRKY3 IBIS1 (6x) DUF1336 unch1576 TM230 KIN10 DREB2B CSC1H GNL1 vSN13 PPR5g15 DUF1997 (4x) PE31 At5g39 emp24 PHO1-9 SPX4 cystgS1 Myb4 NAC92
Myricaceae
Morru-2
COP9s2 RPL21 CSC1H GNL1 vSN13 PPR5g15 At5g39 emp24 SPX4 cystgS1 MYB39 HERC1
Juglandaceae
Jugre-1
WRKY3 IBIS1 DUF1336 TM230 KIN10 DREB2B CSC1H GNL1a vSN13 PPR5g15 PE31 At5g39 (2x) emp24 PHO1-9 SPX4 cystgS1 Myb4 NAC92
Jugre-10
CHU_1773 choMut1 lamin (2x) WRKY3 IBIS1 KIN10 DREB2B GNL1b vSN13 At5g39 emp24 PHO1-9 SPX4 cystgS1 Myb4
Casuarinaceae
Caseq-8
WRKY3 IBIS1 GGPPsyn DUF1336 TM230 (3x) KIN10 (3x) DREB2B 300kb CSC1H (2x) GNL1 vSN13 PE31 At5g39 emp24 PHO1-9 emp24 PHO1-9 emp24 PHO1-9 SPX4 cystgS1
Betulaceae
Carfa-3
WRKY3 IBIS1 DUF1336 TM230 KIN10 (2x) DREB2B CSC1H GNL1 vSN13 PPR5g15 PE31 At5g39 PHO1-9 SPX4 cystgS1

#### Suppl. Figure S11. Fabales cladogram and OSGs

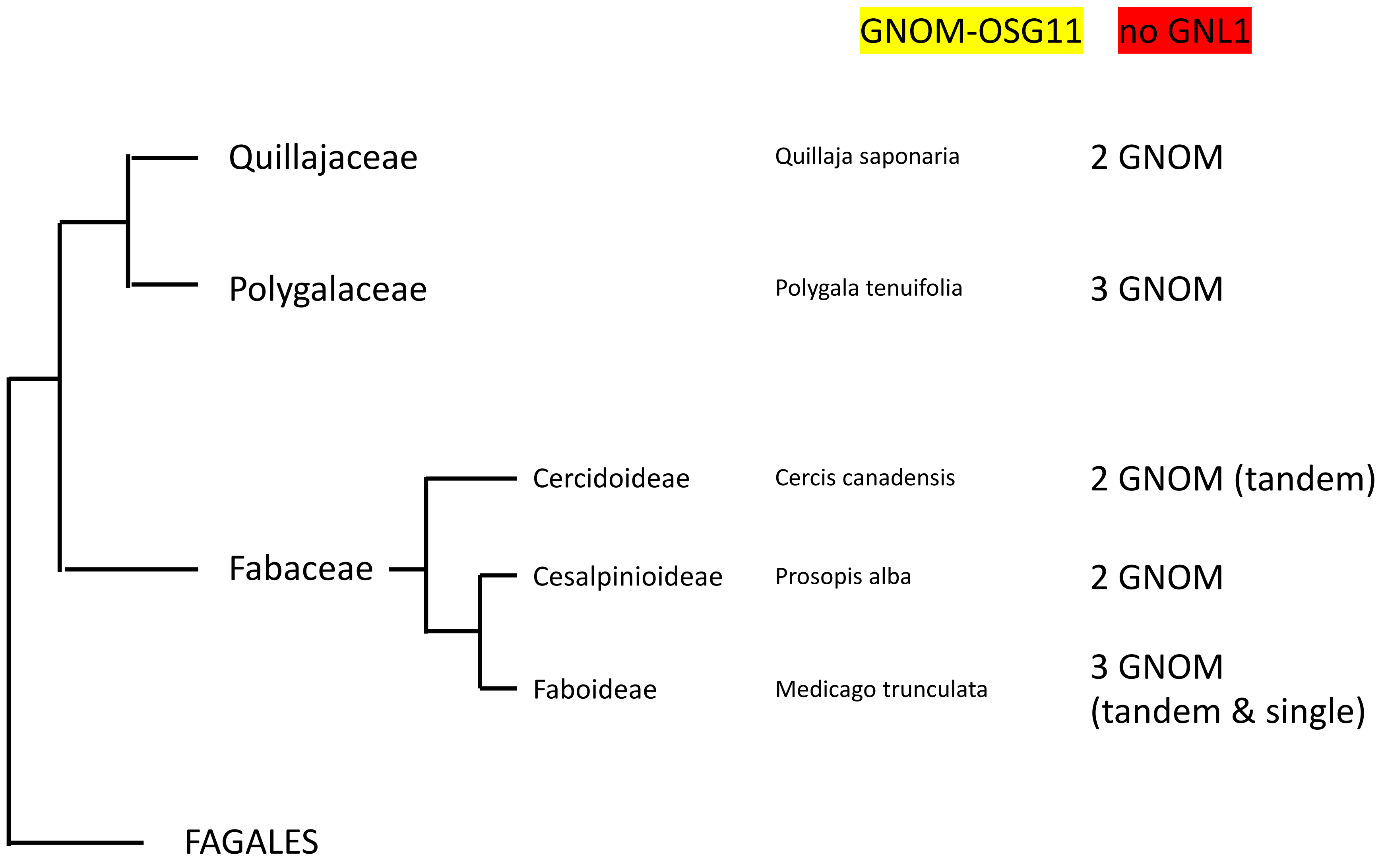

Cladogram based on Zuntini et al. (2024)

### Fabales

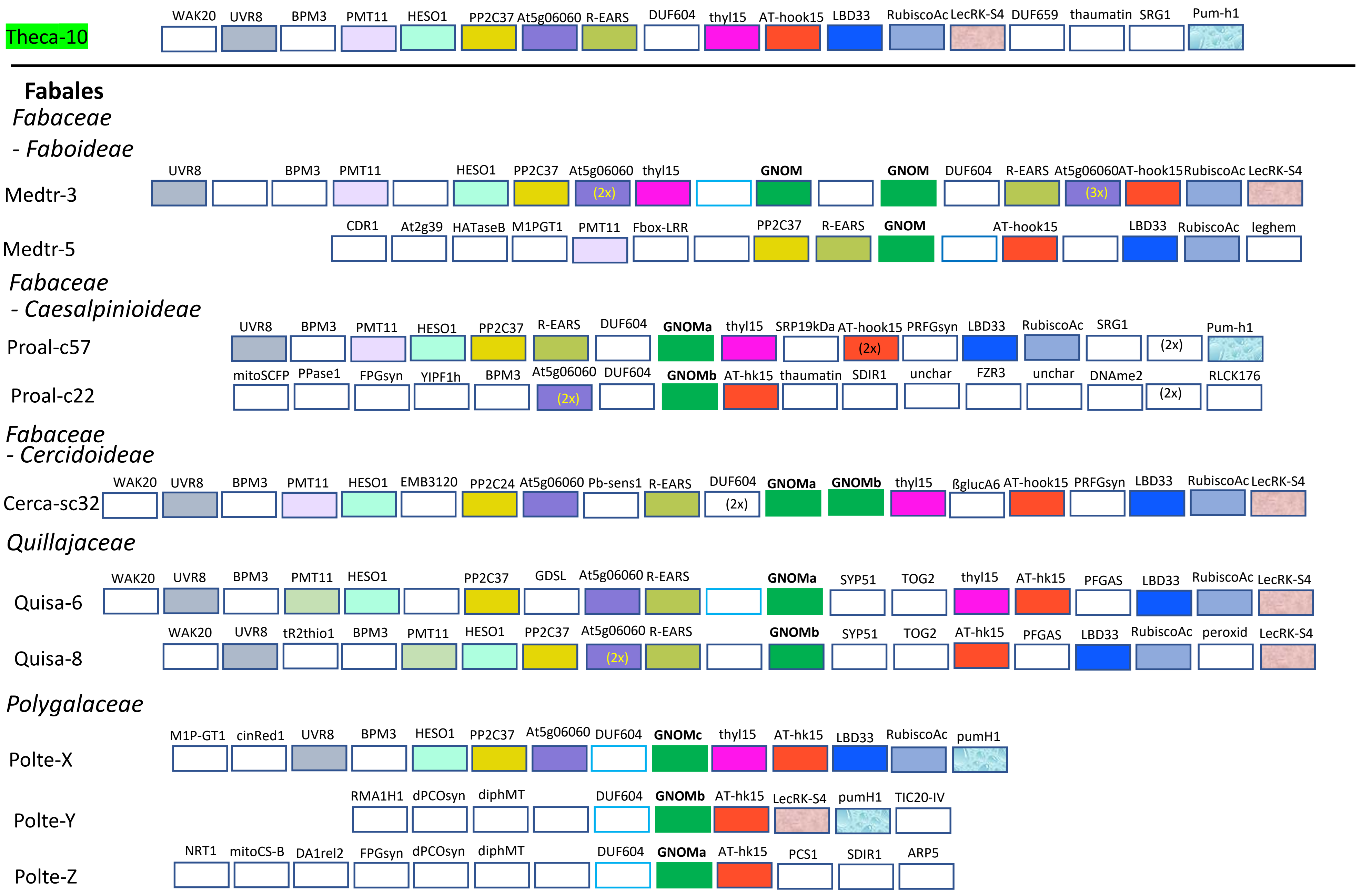

Theca-10
WAK20 UVR8 BPM3 PMT11 HESO1 PP2C37 At5g06060 R-EARS DUF604 thyl15 AT-hook15 LBD33 RubiscoAc LecRK-S4 DUF659 thaumatin SRG1 Pum-h1
Fabales
Fabaceae
- Faboideae
Medtr-3
UVR8 BPM3 PMT11 HESO1 PP2C37 At5g06060 (2x) thyl15 GNOM GNOM DUF604 R-EARS At5g06060 (3x) AT-hook15 RubiscoAc LecRK-S4
Medtr-5
CDR1 At2g39 HATaseB M1PGT1 PMT11 Fbox-LRR PP2C37 R-EARS GNOM AT-hook15 LBD33 RubiscoAc leghem
Fabaceae
- Caesalpinioideae
Proal-c57
UVR8 BPM3 PMT11 HESO1 PP2C37 R-EARS DUF604 GNOMa thyl15 SRP19kDa AT-hook15 (2x) PRFGsyn LBD33 RubiscoAc SRG1 (2x) Pum-h1
Proal-c22
mitoSCFP PPase1 FPGsyn YIPF1h BPM3 At5g06060 (2x) DUF604 GNOMb AT-hk15 thaumatin SDIR1 unchar FZR3 unchar DNAme2 (2x) RLCK176
Fabaceae
- Cercidoideae
Cerca-sc32
WAK20 UVR8 BPM3 PMT11 HESO1 EMB3120 PP2C24 At5g06060 Pb-sens1 R-EARS DUF604 (2x) GNOMa GNOMb thyl15 ßglucA6 AT-hook15 PRFGsyn LBD33 RubiscoAc LecRK-S4
Quillajaceae
Quisa-6
WAK20 UVR8 BPM3 PMT11 HESO1 PP2C37 GDSL At5g06060 R-EARS GNOMa SYP51 TOG2 thyl15 AT-hk15 PFGAS LBD33 RubiscoAc LecRK-S4
Quisa-8
WAK20 UVR8 tR2thio1 BPM3 PMT11 HESO1 PP2C37 At5g06060 (2x) R-EARS GNOMb SYP51 TOG2 AT-hk15 PFGAS LBD33 RubiscoAc peroxid LecRK-S4
Polygalaceae
Polte-X
M1P-GT1 cinRed1 UVR8 BPM3 HESO1 PP2C37 At5g06060 DUF604 GNOMc thyl15 AT-hk15 LBD33 RubiscoAc pumH1
Polte-Y
RMA1H1 dPCOsyn diphMT DUF604 GNOMb AT-hk15 LecRK-S4 pumH1 TIC20-IV
Polte-Z
NRT1 mitoCS-B DA1rel2 FPGsyn dPCOsyn diphMT DUF604 GNOMa AT-hk15 PCS1 SDIR1 ARP5

### Loss of GNOM from OSG1 ($N_2$-fixing clade)

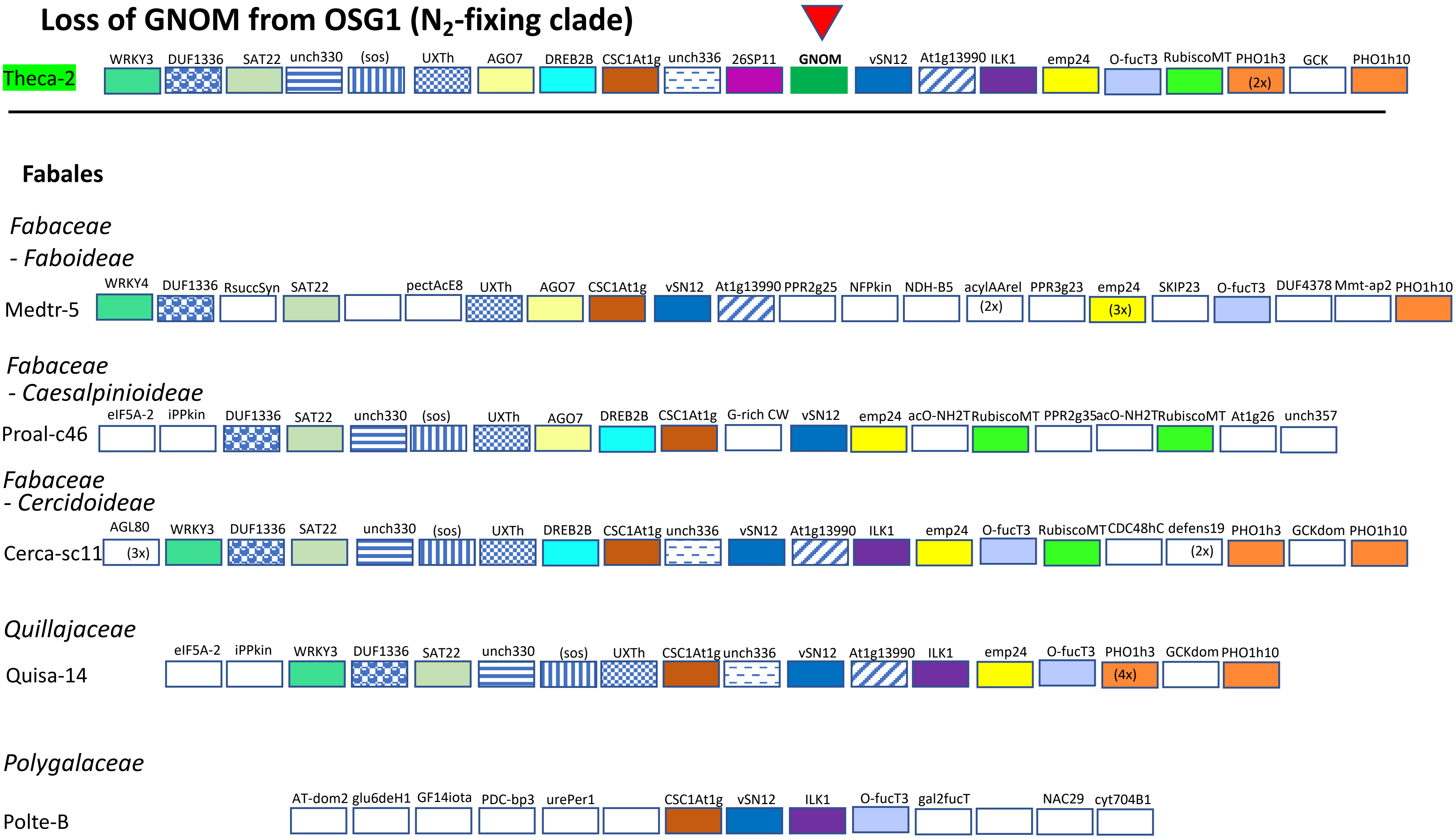

### Loss of GNL1 from OSG2 ($N_2$-fixing clade)

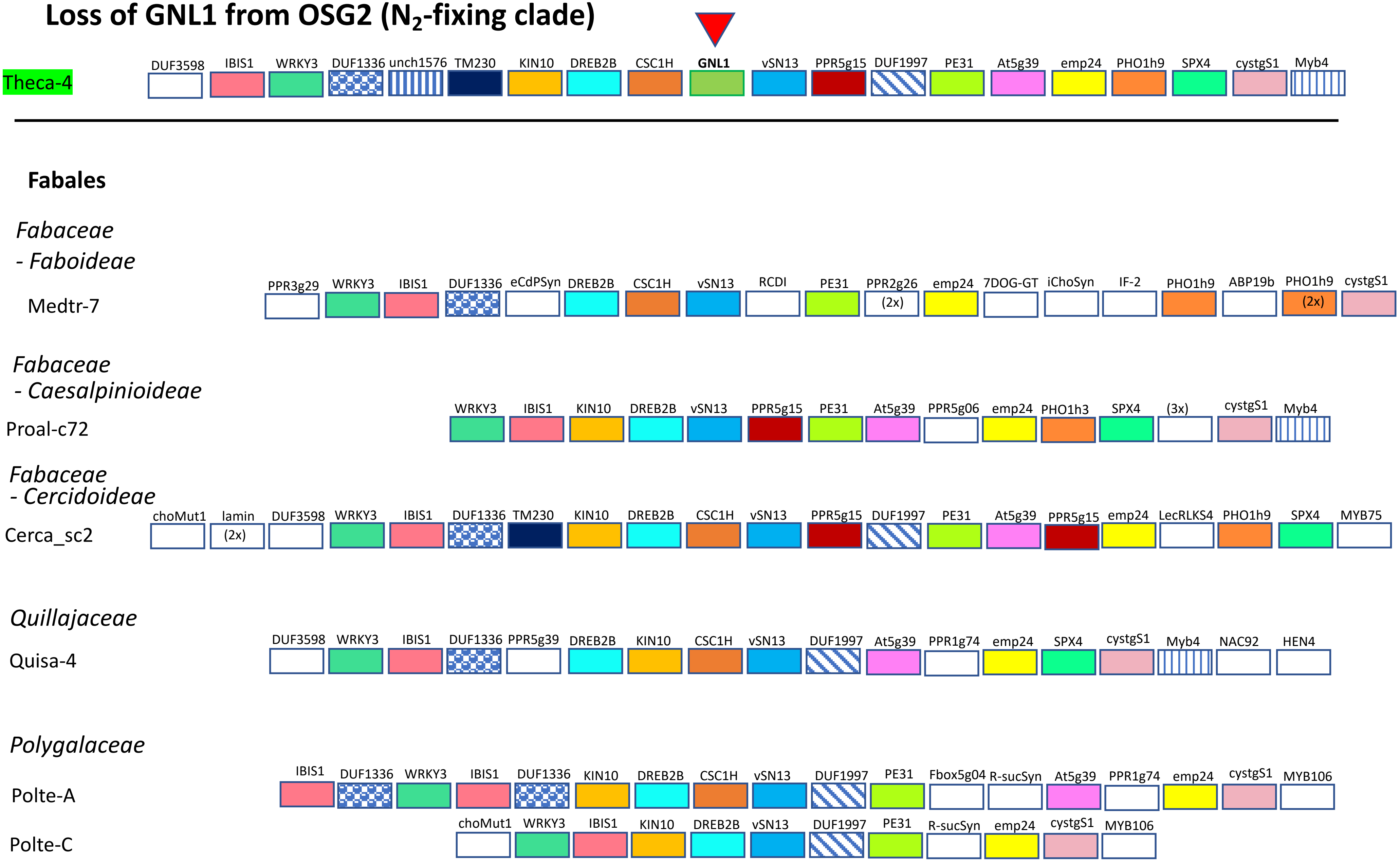

**Suppl. Figur S12. Cucurbitales cladogram and OSG**

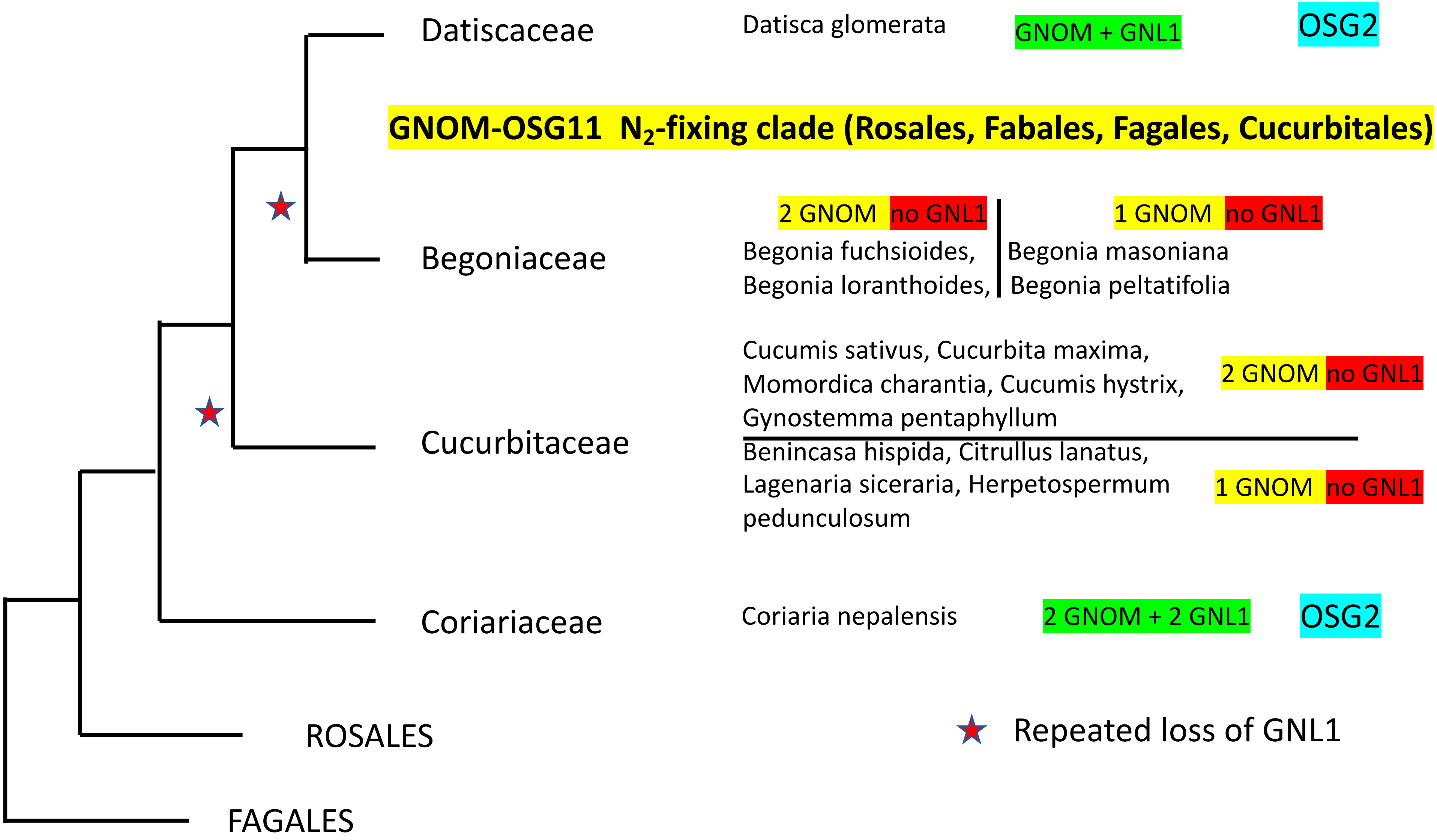

Cladogram based on Zhao et al (2023) Sci Data 10, 259

### Cucurbitales OSG11

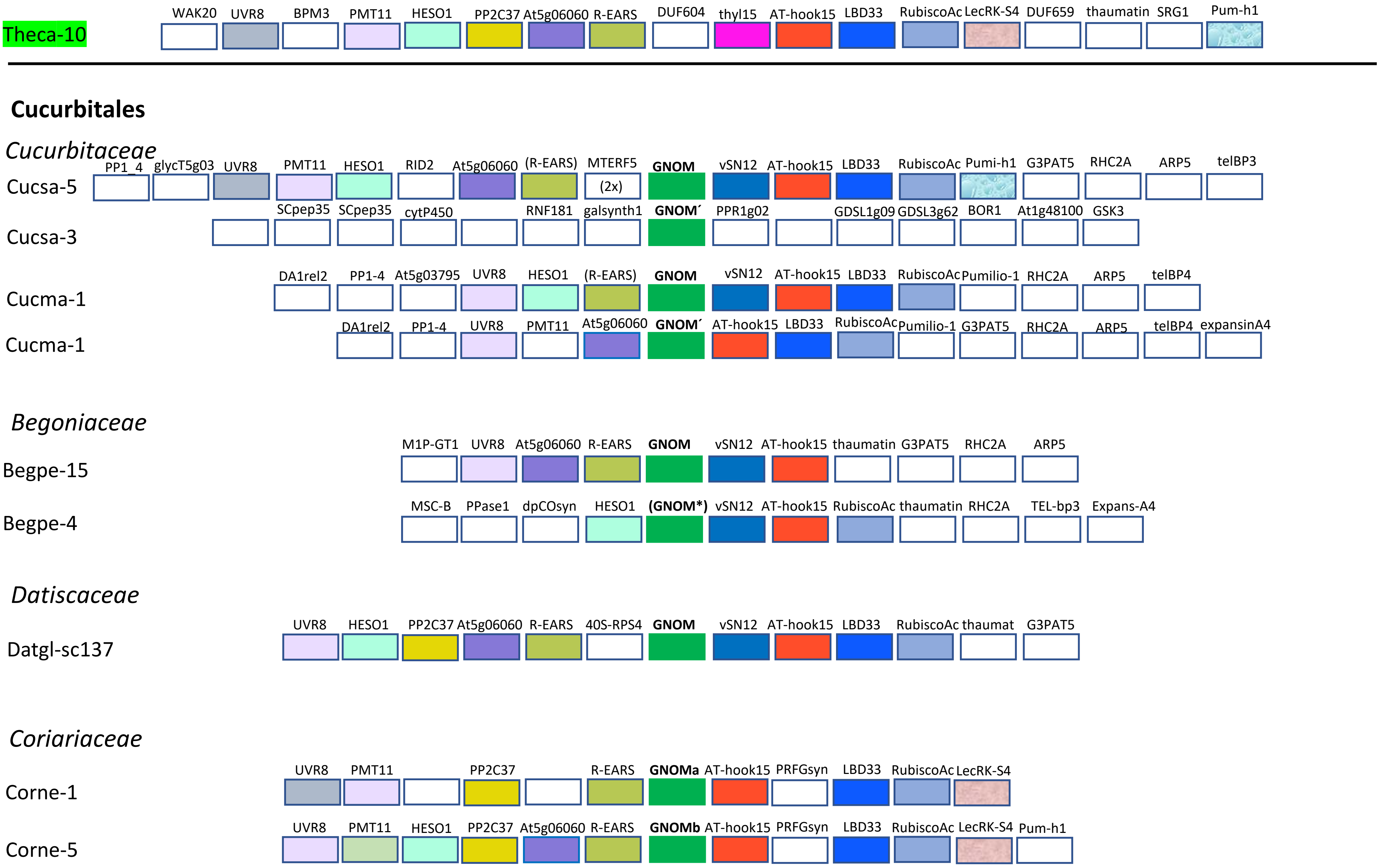

### Loss of GNOM from OSG1 ($N_2$-fixing clade)

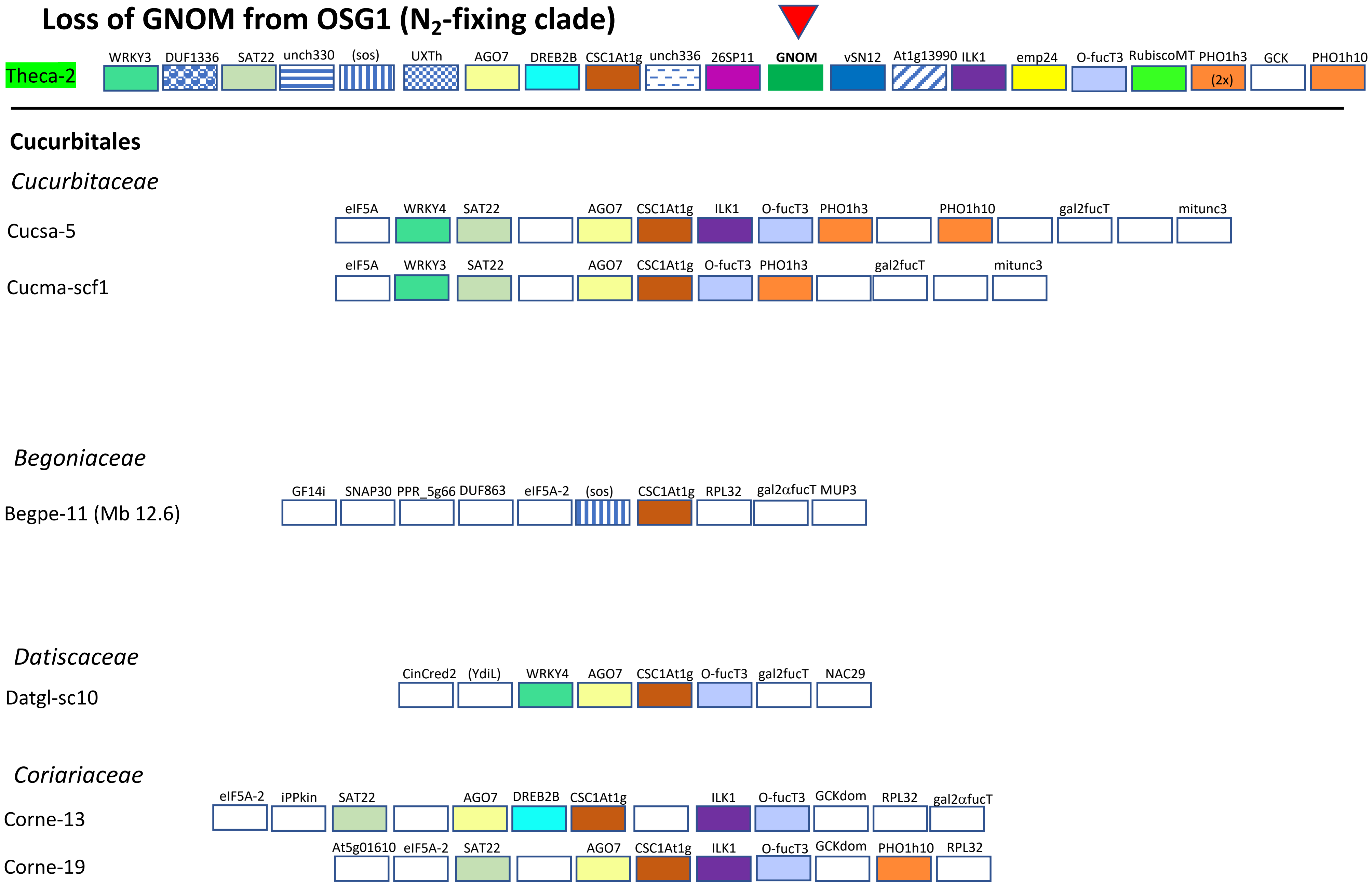

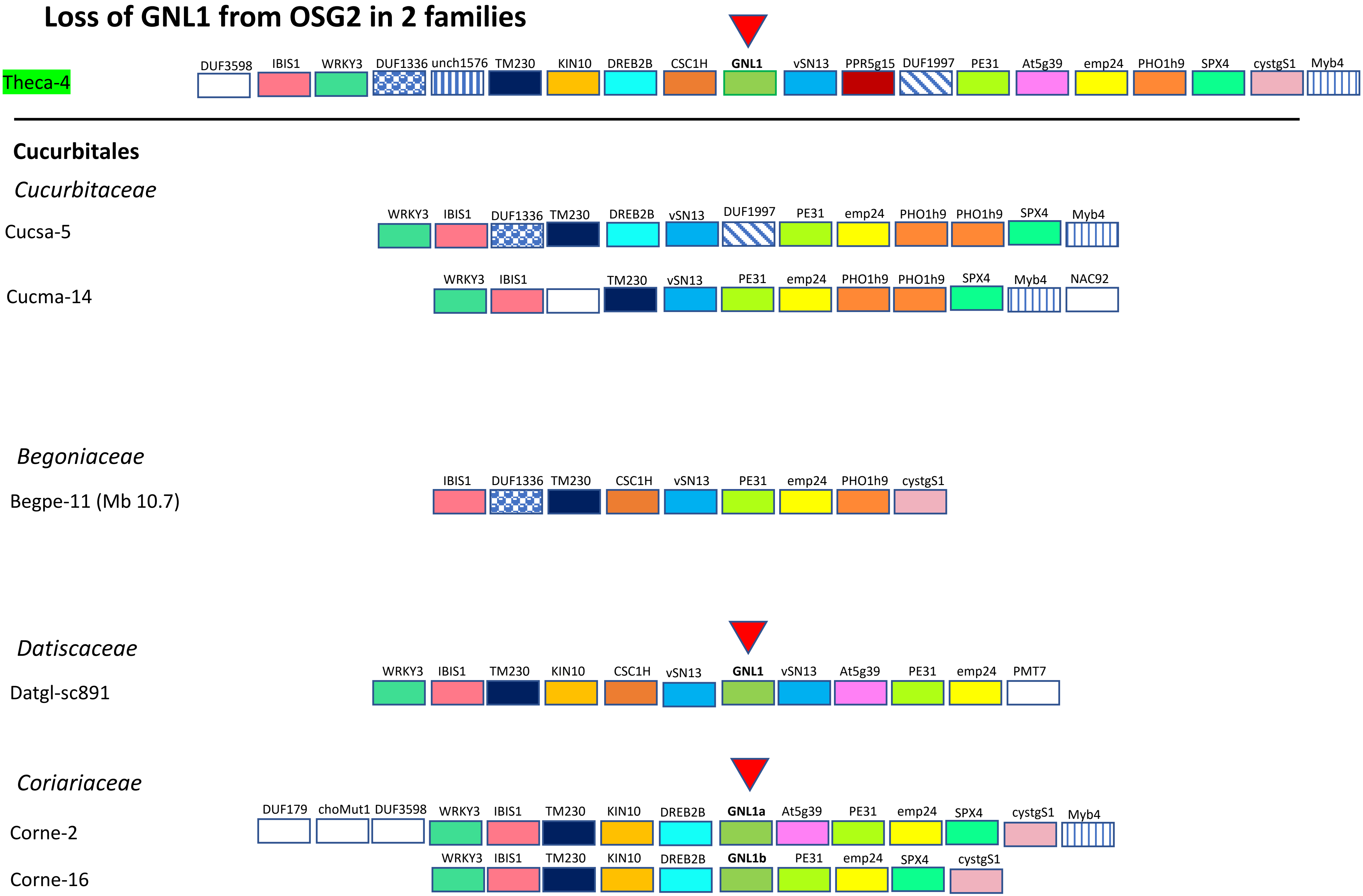
Loss of GNL1 from OSG2 in 2 families
Theca-4
DUF3598 IBIS1 WRKY3 DUF1336 unch1576 TM230 KIN10 DREB2B CSC1H GNL1 vSN13 PPR5g15 DUF1997 PE31 At5g39 emp24 PHO1h9 SPX4 cystgS1 Myb4
Cucurbitales
Cucurbitaceae
Cucsa-5
WRKY3 IBIS1 DUF1336 TM230 DREB2B vSN13 DUF1997 PE31 emp24 PHO1h9 PHO1h9 SPX4 Myb4
Cucma-14
WRKY3 IBIS1 TM230 vSN13 PE31 emp24 PHO1h9 PHO1h9 SPX4 Myb4 NAC92
Begoniaceae
Begpe-11 (Mb 10.7)
IBIS1 DUF1336 TM230 CSC1H vSN13 PE31 emp24 PHO1h9 cystgS1
Datiscaceae
Datgl-sc891
WRKY3 IBIS1 TM230 KIN10 CSC1H vSN13 GNL1 vSN13 At5g39 PE31 emp24 PMT7
Coriariaceae
Corne-2
DUF179 choMut1 DUF3598 WRKY3 IBIS1 TM230 KIN10 DREB2B GNL1a At5g39 PE31 emp24 SPX4 cystgS1 Myb4
Corne-16
WRKY3 IBIS1 TM230 KIN10 DREB2B GNL1b PE31 emp24 SPX4 cystgS1

**Suppl. Figure S13. Rosales cladogram and OSGs**

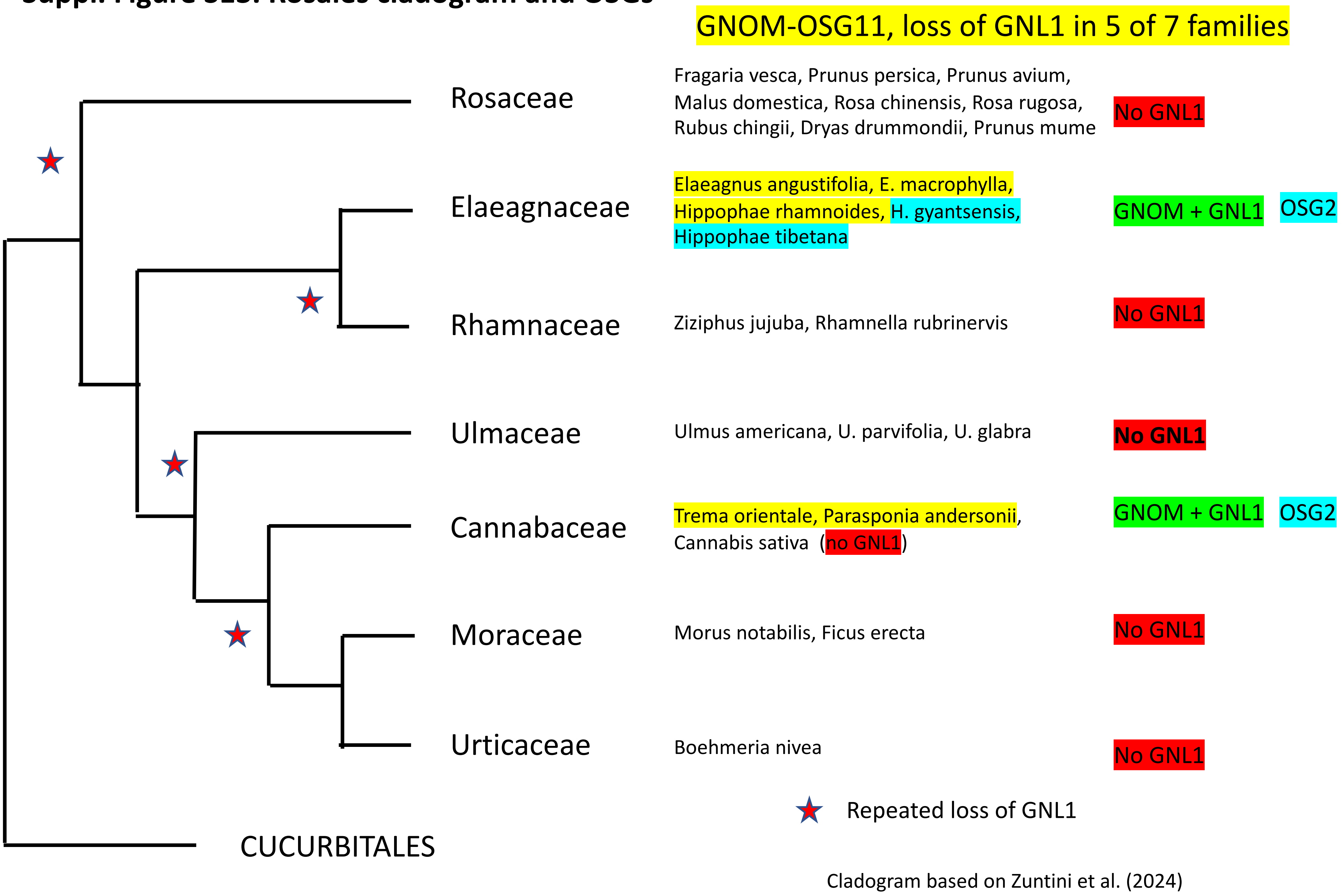

### Rosales OSG11

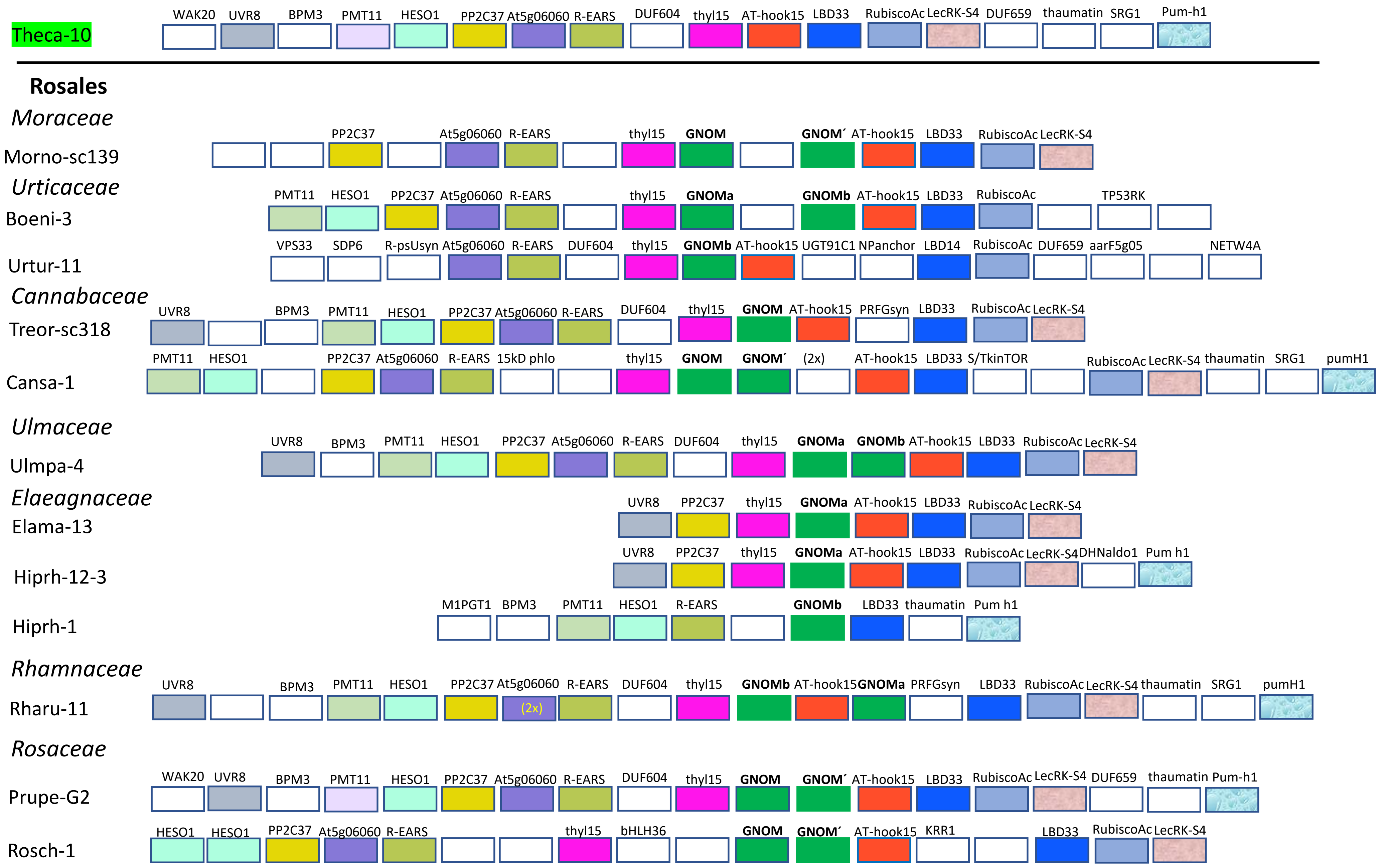

### Loss of GNOM from OSG1 ($N_2$-fixing clade)

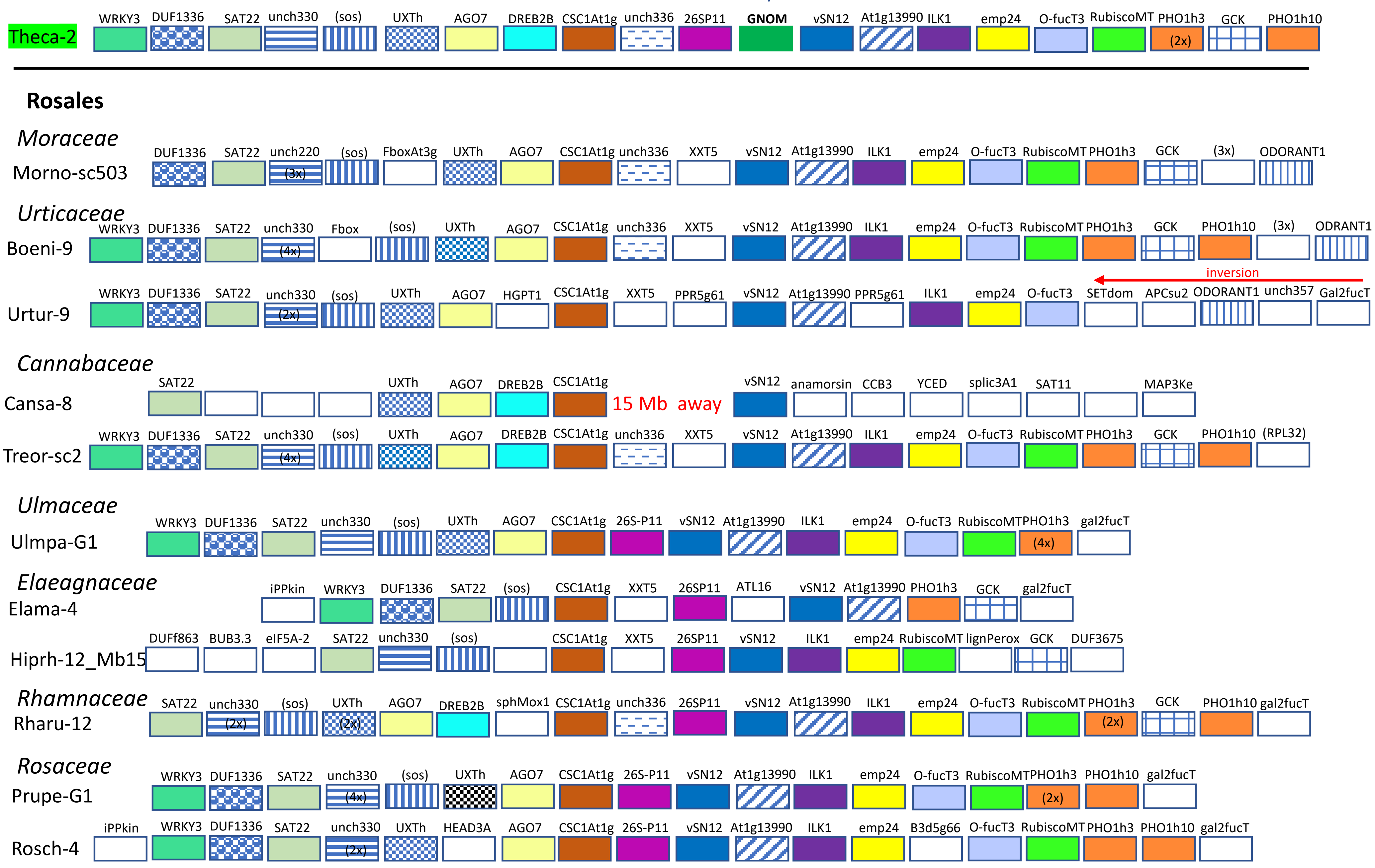

### Loss of GNL1 from OSG2 in 5 families

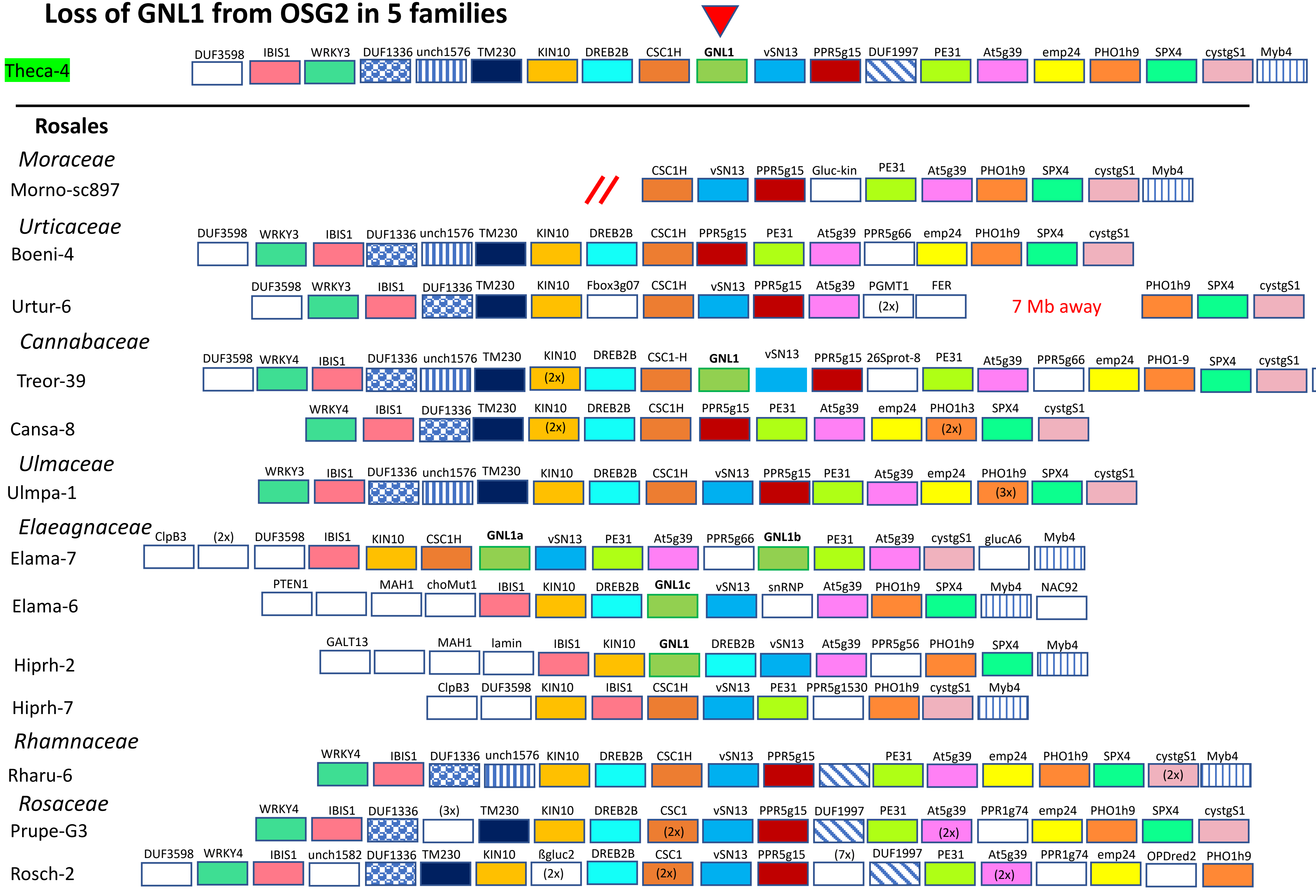

**Suppl. Figure S14. Zygophyllales cladogram and OSGs**

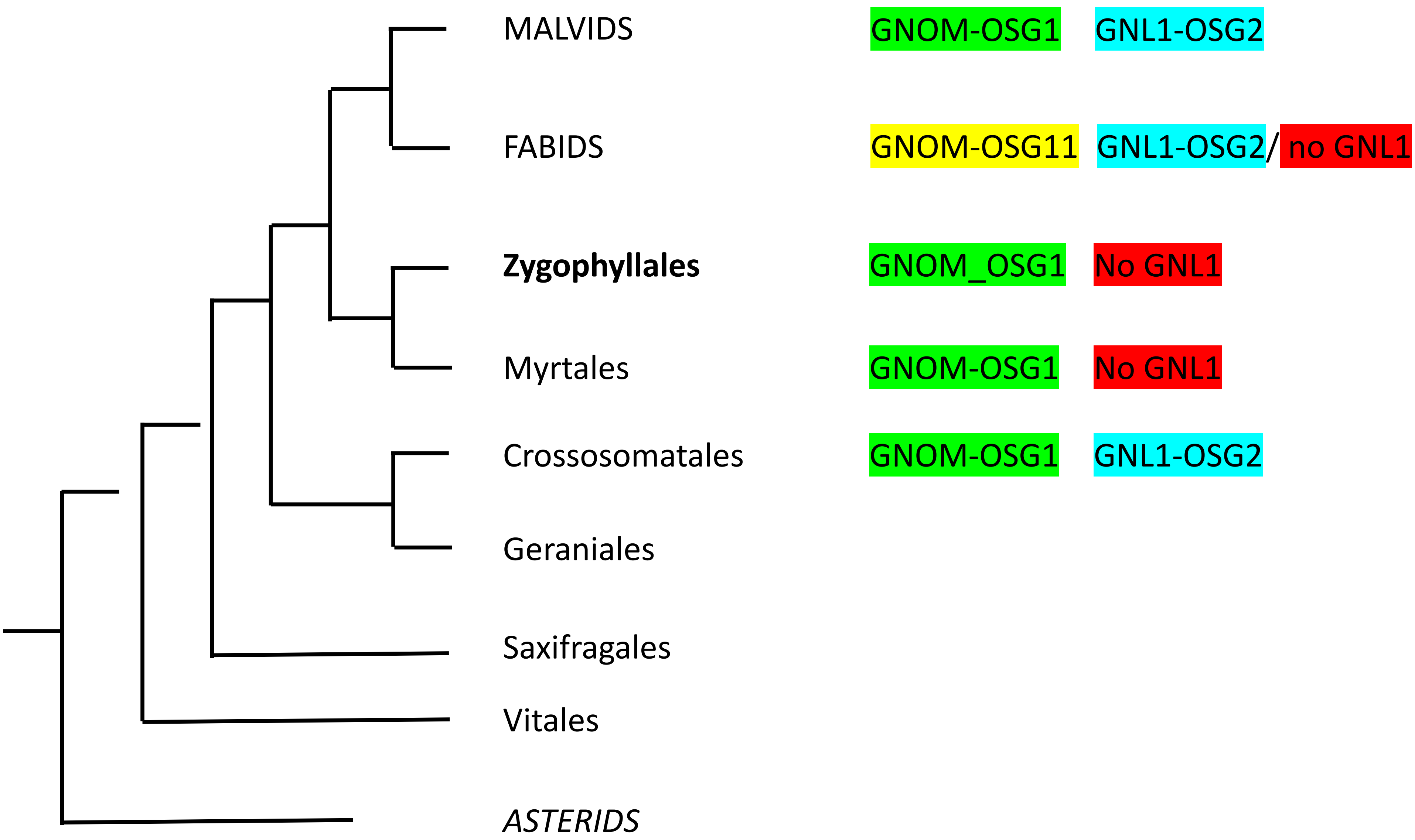

### Zygophyllales synteny

*Zygophyllaceae*

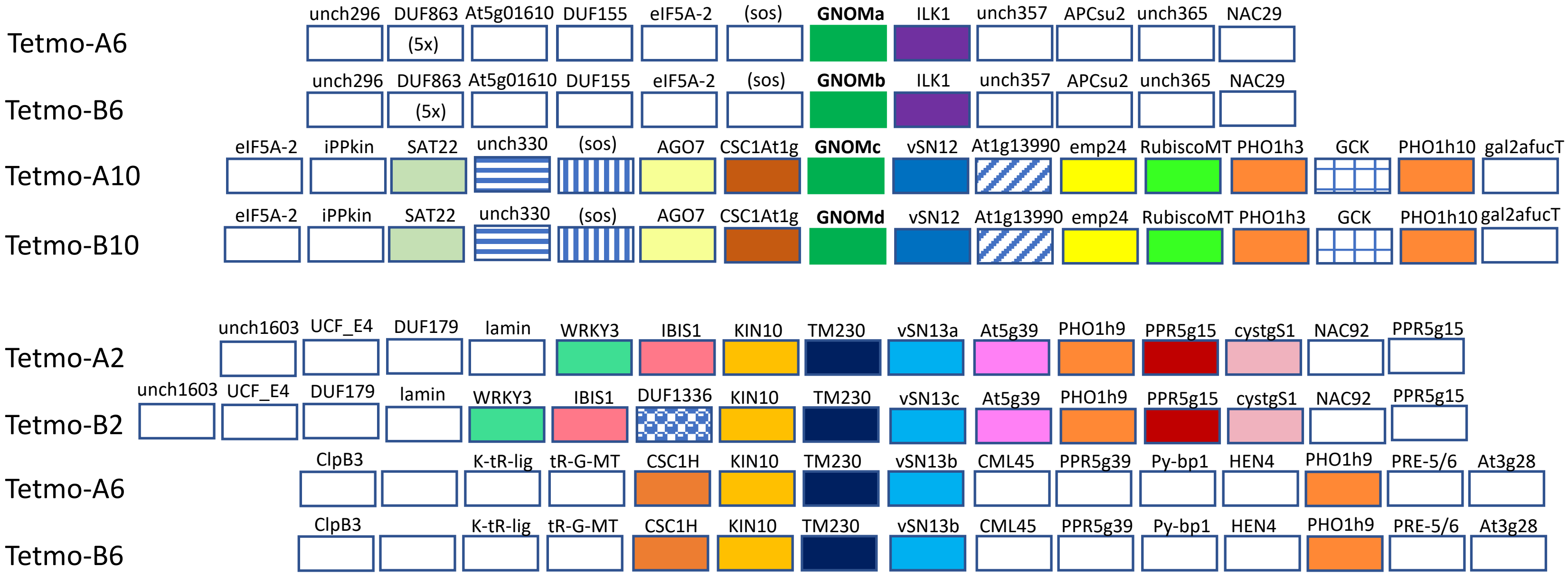

**Suppl. Figure S15. Myrtales cladogram and OSGs**

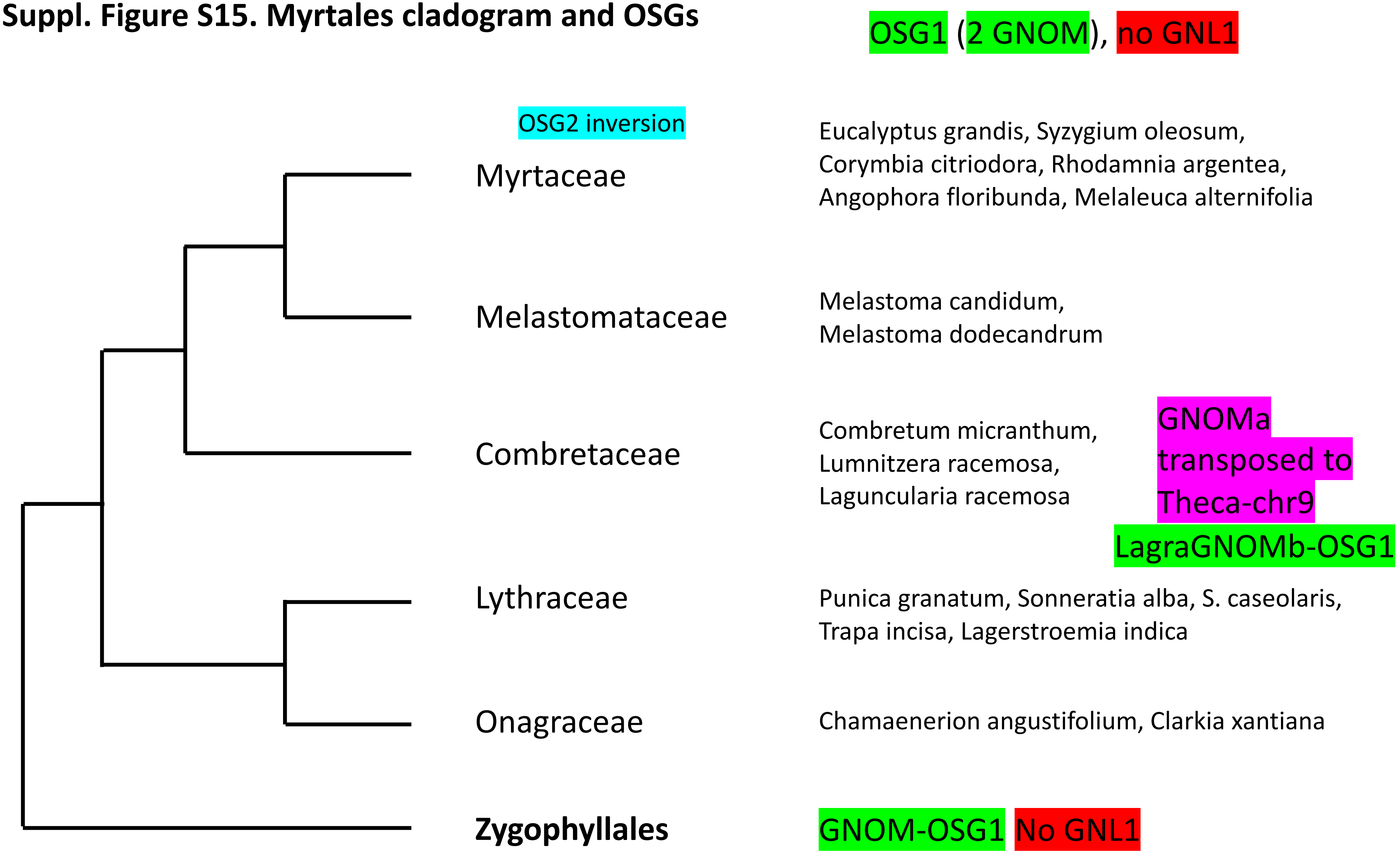

Cladogram based on Maurin et al (2021)

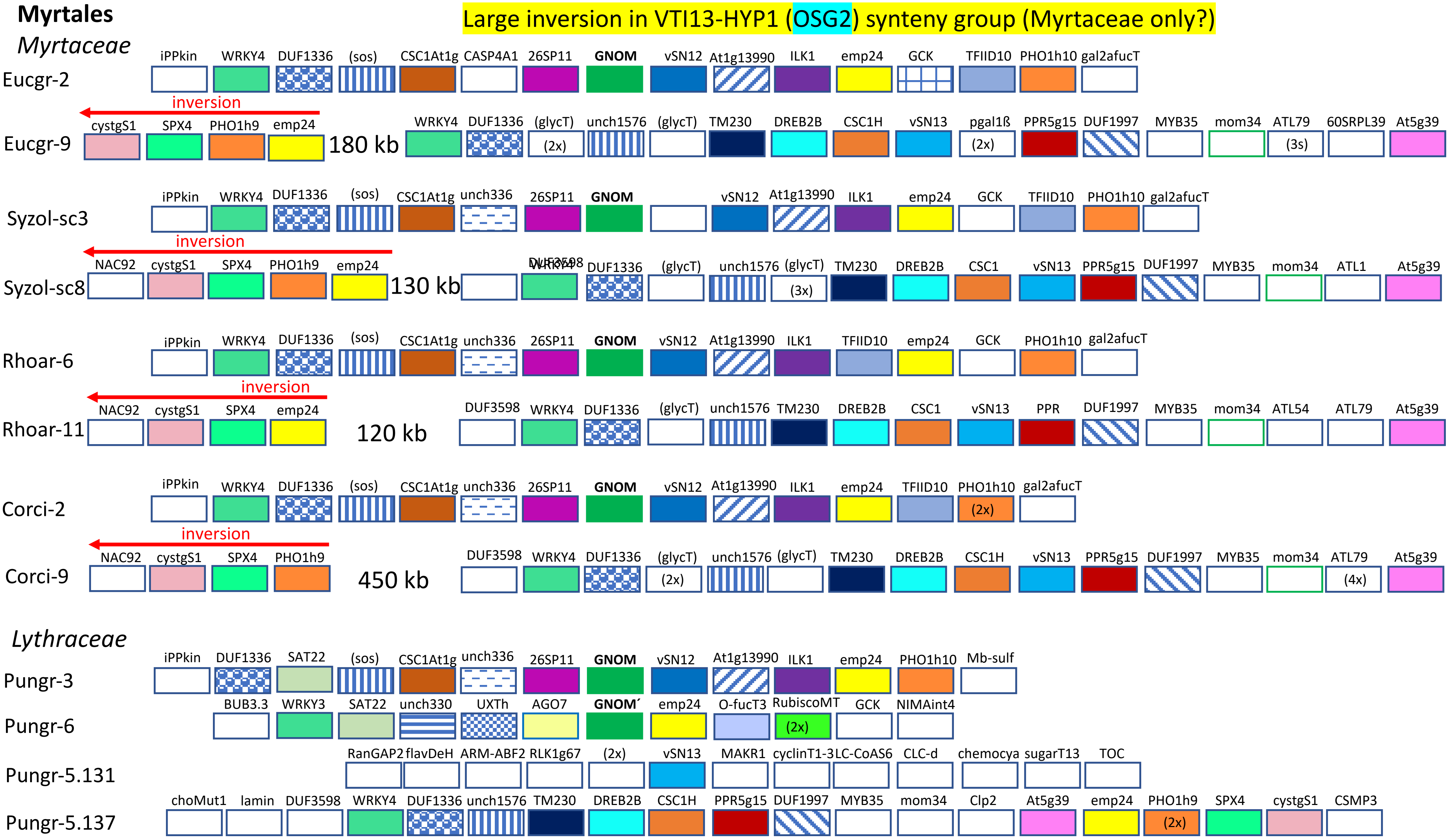

Myrtales
Myrtaceae
Large inversion in VTI13-HYP1 (OSG2) synteny group (Myrtaceae only?)
Eucgr-2
iPPkin WRKY4 DUF1336 (sos) CSC1At1g CASP4A1 26SP11 GNOM vSN12 At1g13990 ILK1 emp24 GCK TFIID10 PHO1h10 gal2afucT
inversion
Eucgr-9
cystgS1 SPX4 PHO1h9 emp24
180 kb
WRKY4 DUF1336 (glycT) (2x) unch1576 (glycT) TM230 DREB2B CSC1H vSN13 pgal1ß (2x) PPR5g15 DUF1997 MYB35 mom34 ATL79 (3s) 60SRPL39 At5g39
Syzol-sc3
iPPkin WRKY4 DUF1336 (sos) CSC1At1g unch336 26SP11 GNOM vSN12 At1g13990 ILK1 emp24 GCK TFIID10 PHO1h10 gal2afucT
inversion
Syzol-sc8
NAC92 cystgS1 SPX4 PHO1h9 emp24
130 kb
DUF3598 WRKY4 DUF1336 (glycT) unch1576 (glycT) (3x) TM230 DREB2B CSC1 vSN13 PPR5g15 DUF1997 MYB35 mom34 ATL1 At5g39
Rhoar-6
iPPkin WRKY4 DUF1336 (sos) CSC1At1g unch336 26SP11 GNOM vSN12 At1g13990 ILK1 TFIID10 emp24 GCK PHO1h10 gal2afucT
inversion
Rhoar-11
NAC92 cystgS1 SPX4 emp24
120 kb
DUF3598 WRKY4 DUF1336 (glycT) unch1576 TM230 DREB2B CSC1 vSN13 PPR DUF1997 MYB35 mom34 ATL54 ATL79 At5g39
Corci-2
iPPkin WRKY4 DUF1336 (sos) CSC1At1g unch336 26SP11 GNOM vSN12 At1g13990 ILK1 emp24 TFIID10 PHO1h10 (2x) gal2afucT
inversion
Corci-9
NAC92 cystgS1 SPX4 PHO1h9
450 kb
DUF3598 WRKY4 DUF1336 (glycT) (2x) unch1576 (glycT) TM230 DREB2B CSC1H vSN13 PPR5g15 DUF1997 MYB35 mom34 ATL79 (4x) At5g39
Lythraceae
Pungr-3
iPPkin DUF1336 SAT22 (sos) CSC1At1g unch336 26SP11 GNOM vSN12 At1g13990 ILK1 emp24 PHO1h10 Mb-sulf
Pungr-6
BUB3.3 WRKY3 SAT22 unch330 UXTh AGO7 GNOM´ emp24 O-fucT3 RubiscoMT (2x) GCK NIMAint4
Pungr-5.131
RanGAP2 flavDeH ARM-ABF2 RLK1g67 (2x) vSN13 MAKR1 cyclinT1-3 LC-CoAS6 CLC-d chemocya sugarT13 TOC
Pungr-5.137
choMut1 lamin DUF3598 WRKY4 DUF1336 unch1576 TM230 DREB2B CSC1H PPR5g15 DUF1997 MYB35 mom34 Clp2 At5g39 emp24 PHO1h9 (2x) SPX4 cystgS1 CSMP3

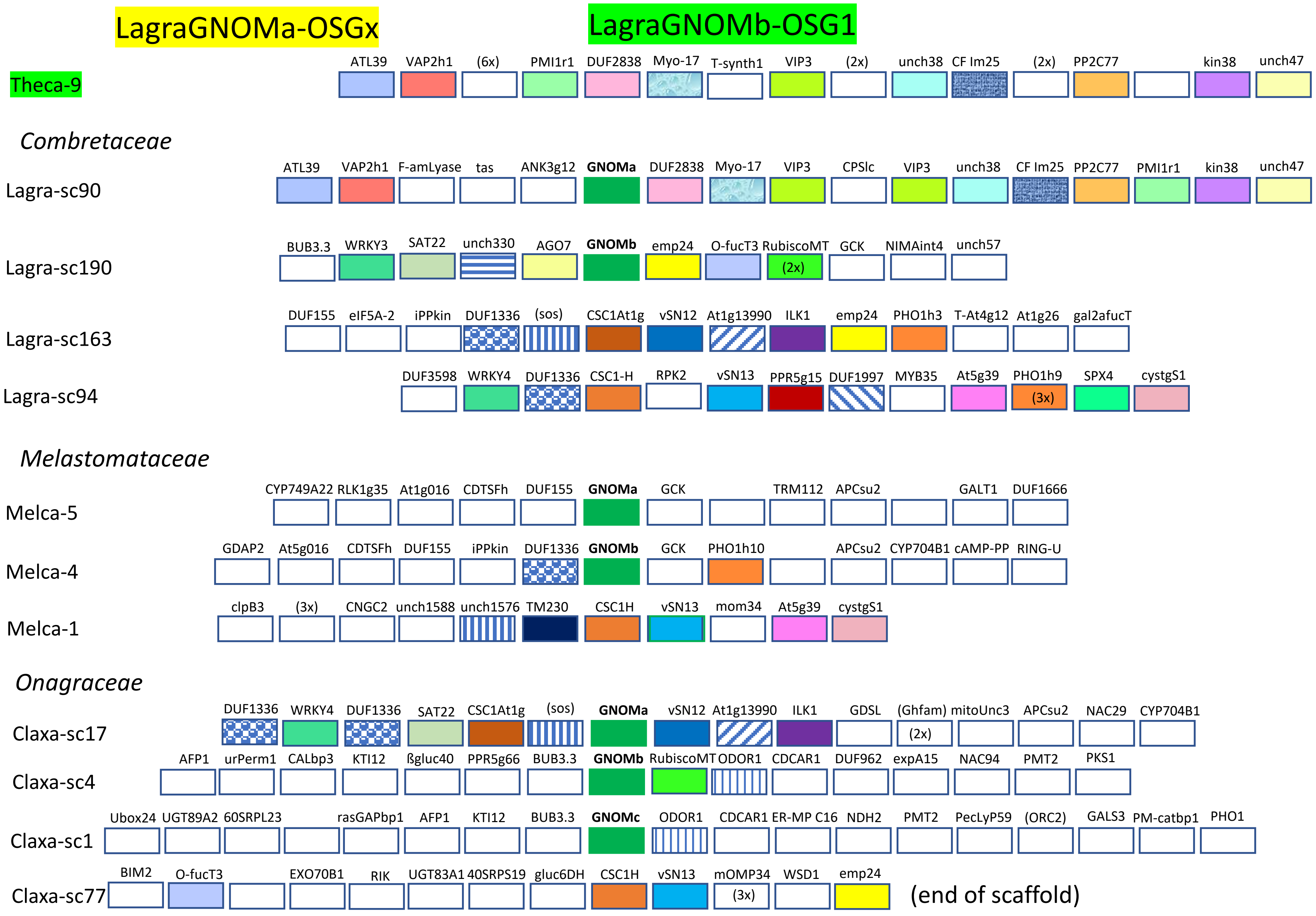

LagraGNOMa-OSGx
LagraGNOMb-OSG1
Theca-9
ATL39 VAP2h1 (6x) PMI1r1 DUF2838 Myo-17 T-synth1 VIP3 (2x) unch38 CF Im25 (2x) PP2C77 kin38 unch47
Combretaceae
Lagra-sc90
ATL39 VAP2h1 F-amLyase tas ANK3g12 GNOMa DUF2838 Myo-17 VIP3 CPSlc VIP3 unch38 CF Im25 PP2C77 PMI1r1 kin38 unch47
Lagra-sc190
BUB3.3 WRKY3 SAT22 unch330 AGO7 GNOMb emp24 O-fucT3 RubiscoMT (2x) GCK NIMAint4 unch57
Lagra-sc163
DUF155 eIF5A-2 iPPkin DUF1336 (sos) CSC1At1g vSN12 At1g13990 ILK1 emp24 PHO1h3 T-At4g12 At1g26 gal2afucT
Lagra-sc94
DUF3598 WRKY4 DUF1336 CSC1-H RPK2 vSN13 PPR5g15 DUF1997 MYB35 At5g39 PHO1h9 (3x) SPX4 cystgS1
Melastomataceae
Melca-5
CYP749A22 RLK1g35 At1g016 CDTSFh DUF155 GNOMa GCK TRM112 APCsu2 GALT1 DUF1666
Melca-4
GDAP2 At5g016 CDTSFh DUF155 iPPkin DUF1336 GNOMb GCK PHO1h10 APCsu2 CYP704B1 cAMP-PP RING-U
Melca-1
clpB3 (3x) CNGC2 unch1588 unch1576 TM230 CSC1H vSN13 mom34 At5g39 cystgS1
Onagraceae
Claxa-sc17
DUF1336 WRKY4 DUF1336 SAT22 CSC1At1g (sos) GNOMa vSN12 At1g13990 ILK1 GDSL (Ghfam) (2x) mitoUnc3 APCsu2 NAC29 CYP704B1
Claxa-sc4
AFP1 urPerm1 CALbp3 KTI12 ßgluc40 PPR5g66 BUB3.3 GNOMb RubiscoMT ODOR1 CDCAR1 DUF962 expA15 NAC94 PMT2 PKS1
Claxa-sc1
Ubox24 UGT89A2 60SRPL23 rasGAPbp1 AFP1 KTI12 BUB3.3 GNOMc ODOR1 CDCAR1 ER-MP C16 NDH2 PMT2 PecLyP59 (ORC2) GALS3 PM-catbp1 PHO1
Claxa-sc77
BIM2 O-fucT3 EXO70B1 RIK UGT83A1 40SRPS19 gluc6DH CSC1H vSN13 mOMP34 (3x) WSD1 emp24
(end of scaffold)

**Suppl. Figure S16. Crossosomatales cladogram and OSG**

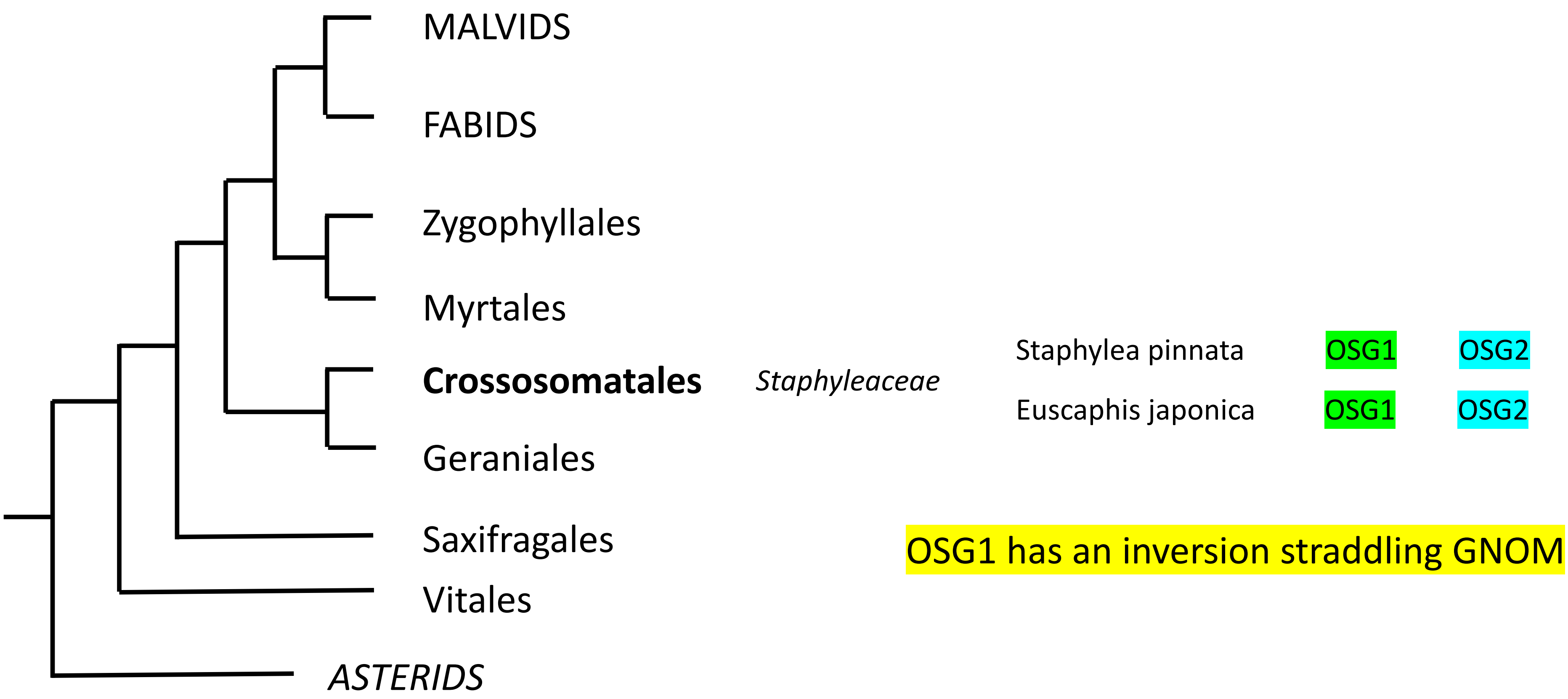

### Crossosomatales

*Staphyleaceae*

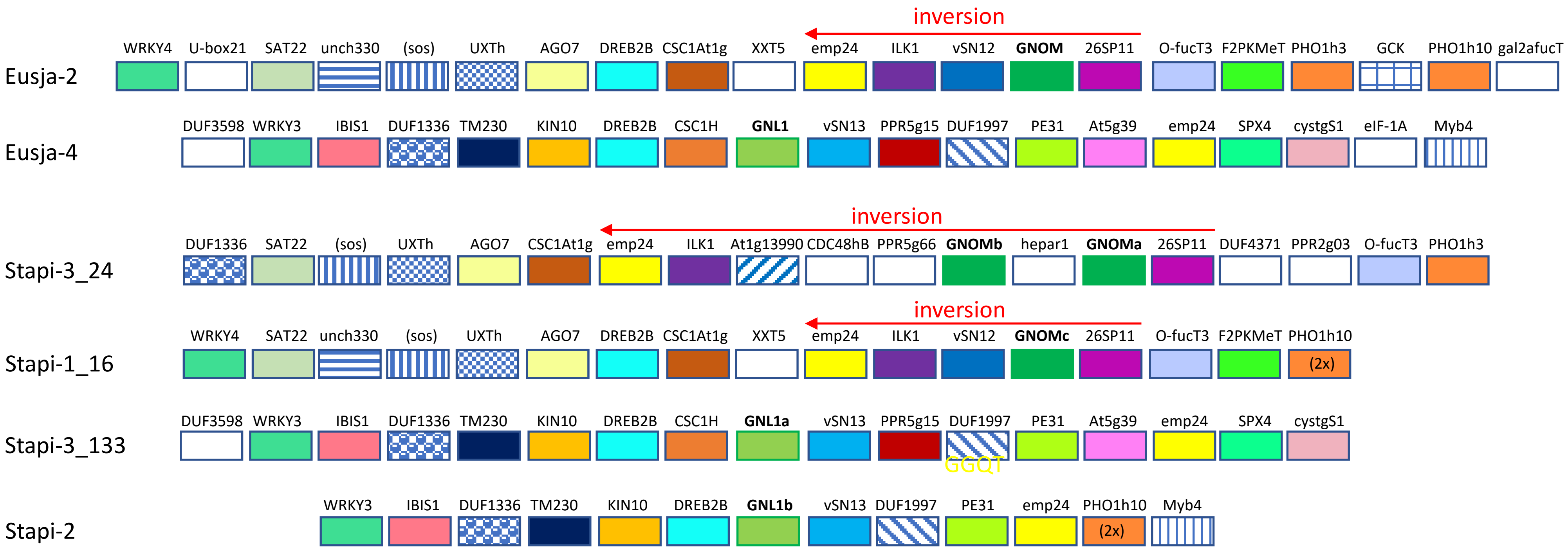

**Suppl. Figure S17. Geraniales cladogram and OSGs**

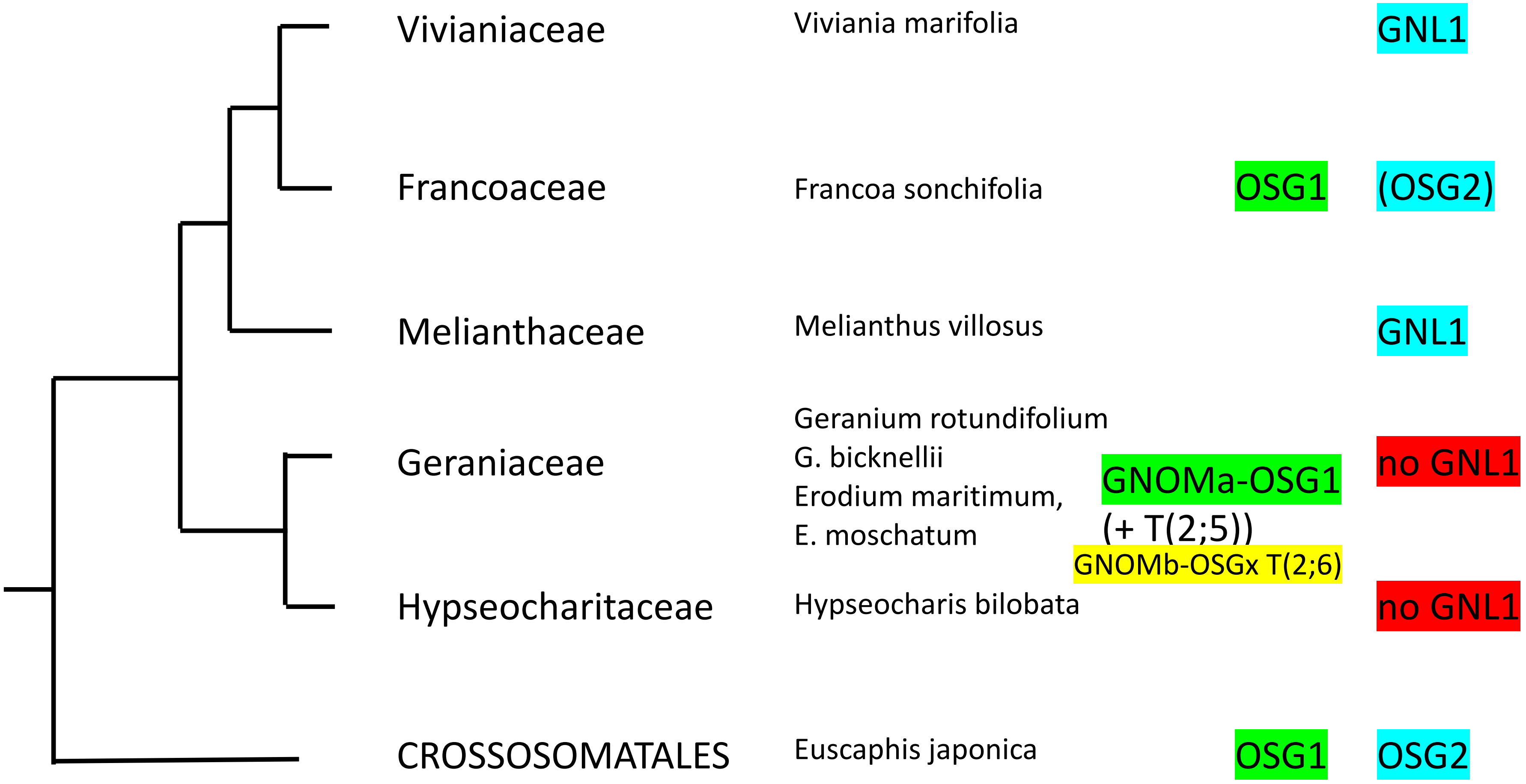

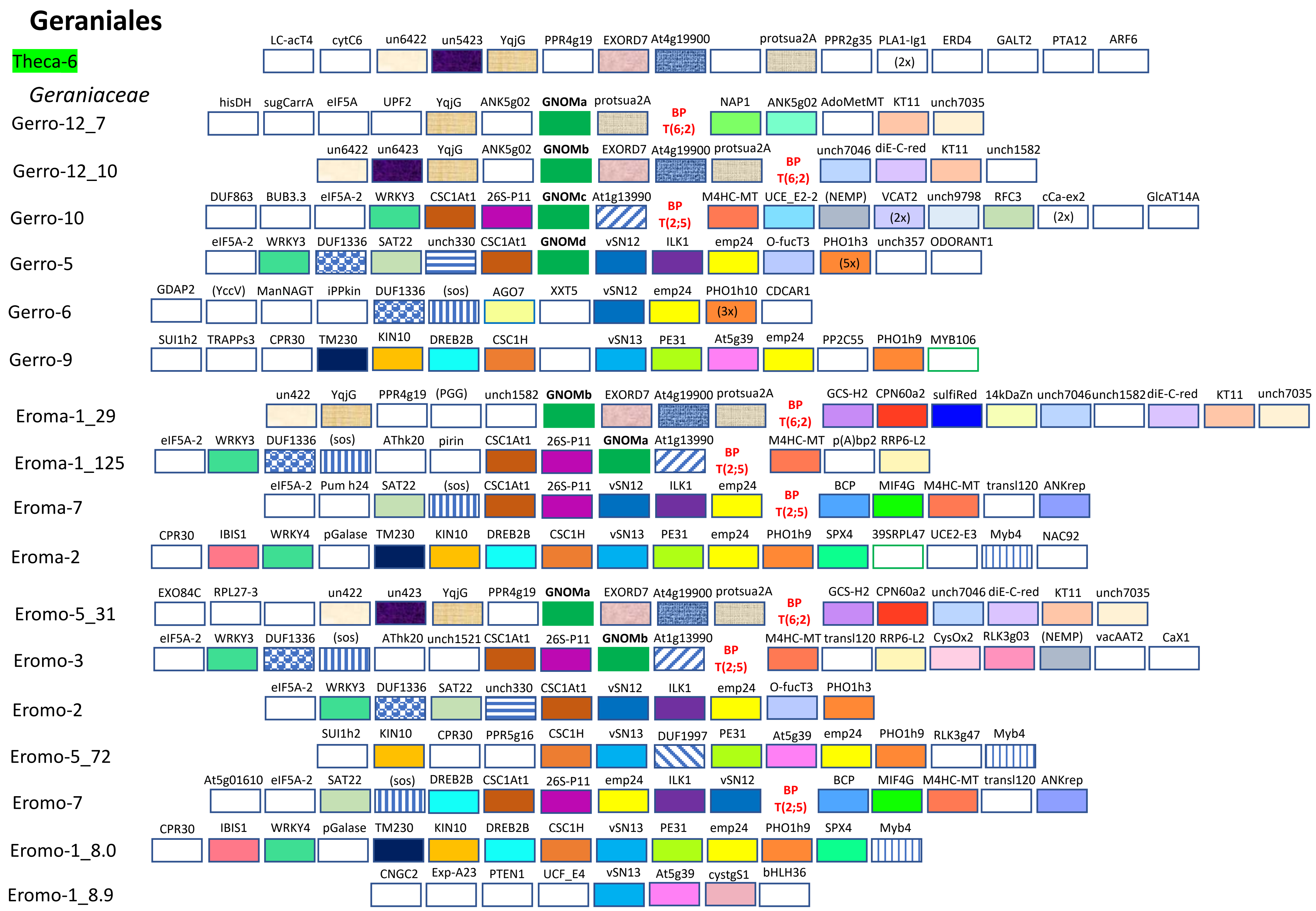

Geraniales
Theca-6
LC-acT4 cytC6 un6422 un5423 YqjG PPR4g19 EXORD7 At4g19900 protsua2A PPR2g35 PLA1-Ig1 (2x) ERD4 GALT2 PTA12 ARF6
Geraniaceae
Gerro-12_7
hisDH sugCarrA eIF5A UPF2 YqjG ANK5g02 GNOMa protsua2A BP T(6;2) NAP1 ANK5g02 AdoMetMT KT11 unch7035
Gerro-12_10
un6422 un6423 YqjG ANK5g02 GNOMb EXORD7 At4g19900 protsua2A BP T(6;2) unch7046 diE-C-red KT11 unch1582
Gerro-10
DUF863 BUB3.3 eIF5A-2 WRKY3 CSC1At1 26S-P11 GNOMc At1g13990 BP T(2;5) M4HC-MT UCE_E2-2 (NEMP) VCAT2 (2x) unch9798 RFC3 cCa-ex2 (2x) GlcAT14A
Gerro-5
eIF5A-2 WRKY3 DUF1336 SAT22 unch330 CSC1At1 GNOMd vSN12 ILK1 emp24 O-fucT3 PHO1h3 (5x) unch357 ODORANT1
Gerro-6
GDAP2 (YccV) ManNAGT iPPkin DUF1336 (sos) AGO7 XXT5 vSN12 emp24 PHO1h10 (3x) CDCAR1
Gerro-9
SUI1h2 TRAPPs3 CPR30 TM230 KIN10 DREB2B CSC1H vSN13 PE31 At5g39 emp24 PP2C55 PHO1h9 MYB106
Eroma-1_29
un422 YqjG PPR4g19 (PGG) unch1582 GNOMb EXORD7 At4g19900 protsua2A BP T(6;2) GCS-H2 CPN60a2 sulfiRed 14kDaZn unch7046 unch1582 diE-C-red KT11 unch7035
Eroma-1_125
eIF5A-2 WRKY3 DUF1336 (sos) AThk20 pirin CSC1At1 26S-P11 GNOMa At1g13990 BP T(2;5) M4HC-MT p(A)bp2 RRP6-L2
Eroma-7
eIF5A-2 Pum h24 SAT22 (sos) CSC1At1 26S-P11 vSN12 ILK1 emp24 BP T(2;5) BCP MIF4G M4HC-MT transl120 ANKrep
Eroma-2
CPR30 IBIS1 WRKY4 pGalase TM230 KIN10 DREB2B CSC1H vSN13 PE31 emp24 PHO1h9 SPX4 39SRPL47 UCE2-E3 Myb4 NAC92
Eromo-5_31
EXO84C RPL27-3 un422 un423 YqjG PPR4g19 GNOMa EXORD7 At4g19900 protsua2A BP T(6;2) GCS-H2 CPN60a2 unch7046 diE-C-red KT11 unch7035
Eromo-3
eIF5A-2 WRKY3 DUF1336 (sos) AThk20 unch1521 CSC1At1 26S-P11 GNOMb At1g13990 BP T(2;5) M4HC-MT transl120 RRP6-L2 CysOx2 RLK3g03 (NEMP) vacAAT2 CaX1
Eromo-2
eIF5A-2 WRKY3 DUF1336 SAT22 unch330 CSC1At1 vSN12 ILK1 emp24 O-fucT3 PHO1h3
Eromo-5_72
SUI1h2 KIN10 CPR30 PPR5g16 CSC1H vSN13 DUF1997 PE31 At5g39 emp24 PHO1h9 RLK3g47 Myb4
Eromo-7
At5g01610 eIF5A-2 SAT22 (sos) DREB2B CSC1At1 26S-P11 emp24 ILK1 vSN12 BP T(2;5) BCP MIF4G M4HC-MT transl120 ANKrep
Eromo-1_8.0
CPR30 IBIS1 WRKY4 pGalase TM230 KIN10 DREB2B CSC1H vSN13 PE31 emp24 PHO1h9 SPX4 Myb4
Eromo-1_8.9
CNGC2 Exp-A23 PTEN1 UCF_E4 vSN13 At5g39 cystgS1 bHLH36

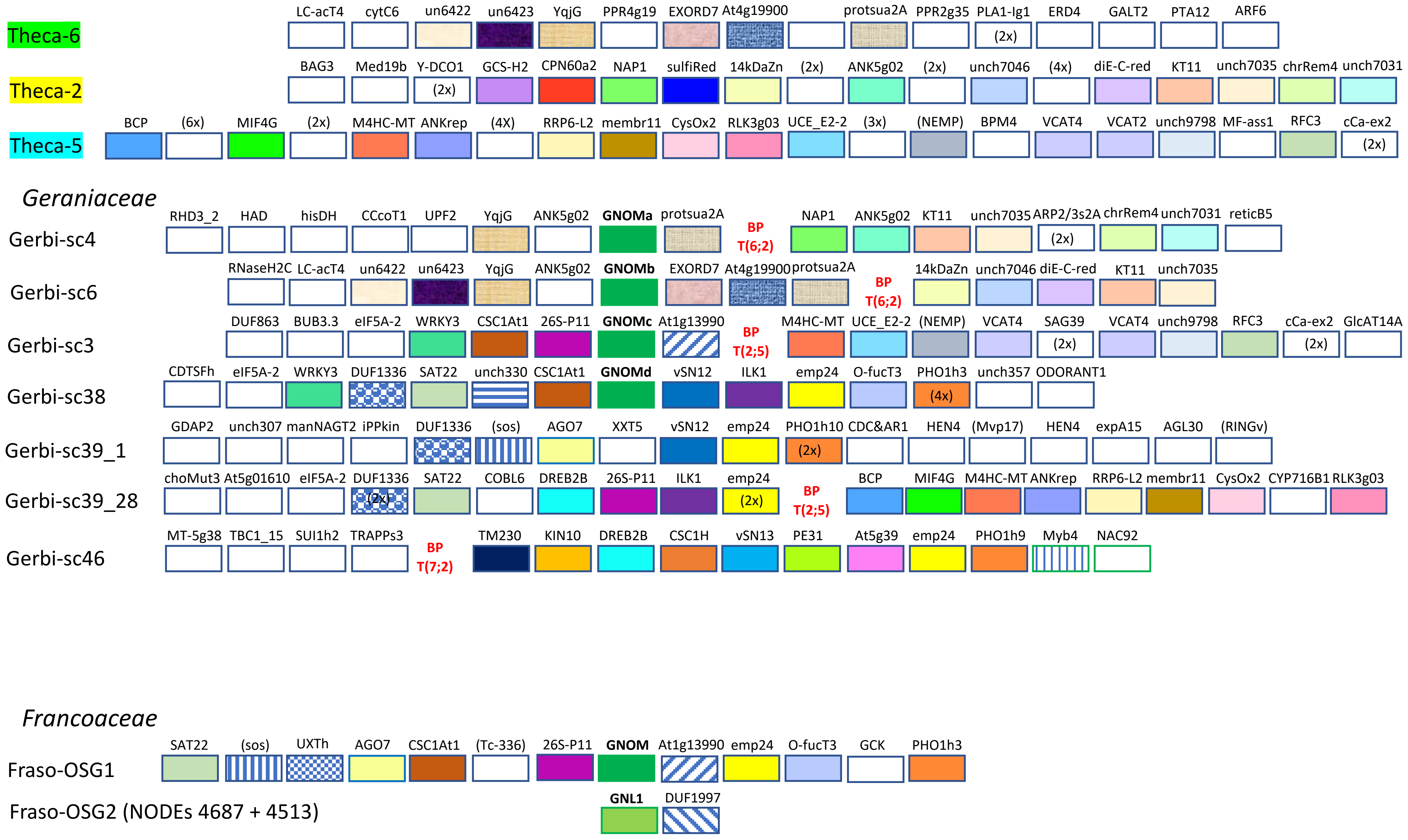

Short contigs in Francoa

#### Suppl. Figure S18. Saxifragales cladogram and OSGs

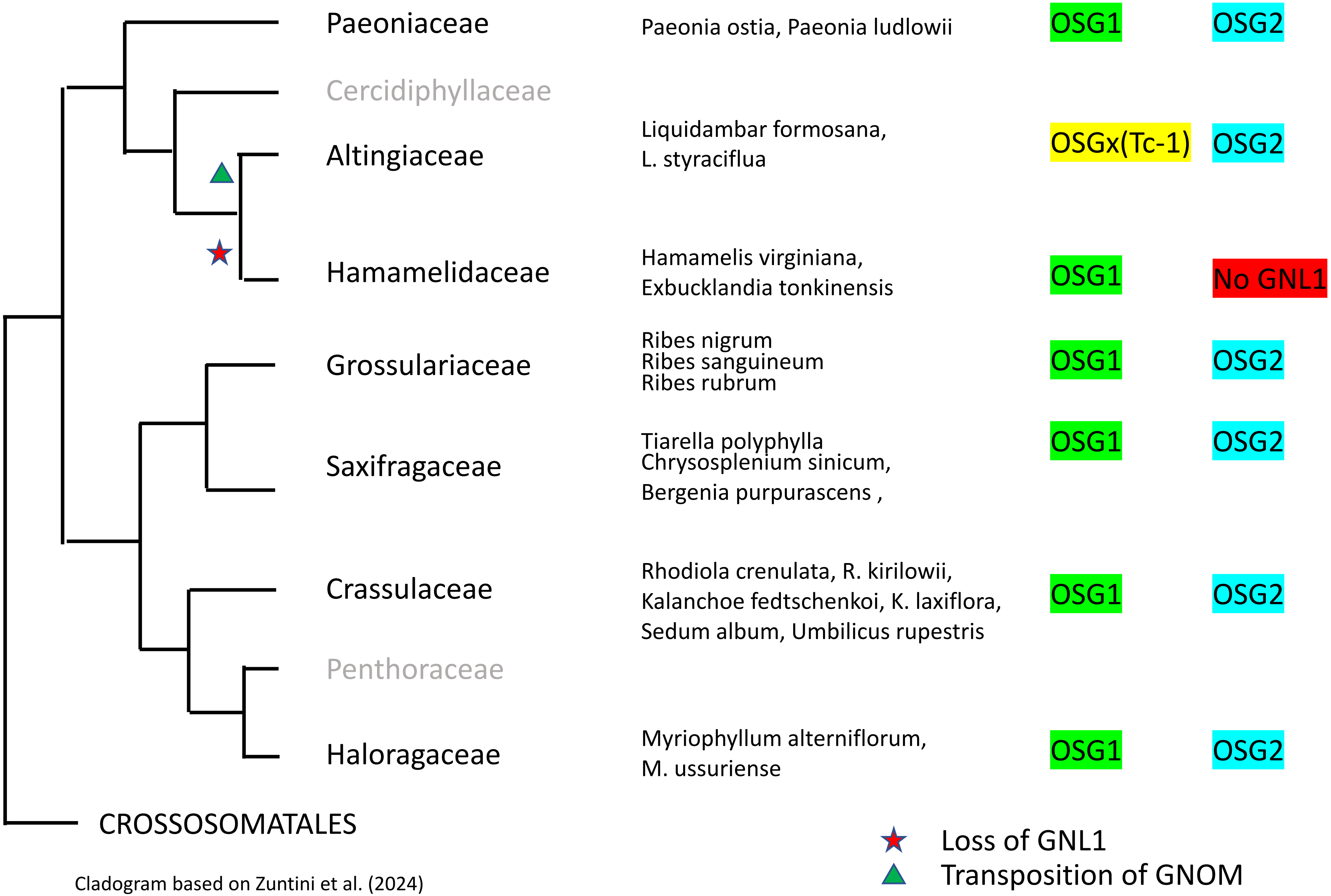

Cladogram based on Zuntini et al. (2024)

### Saxifragales

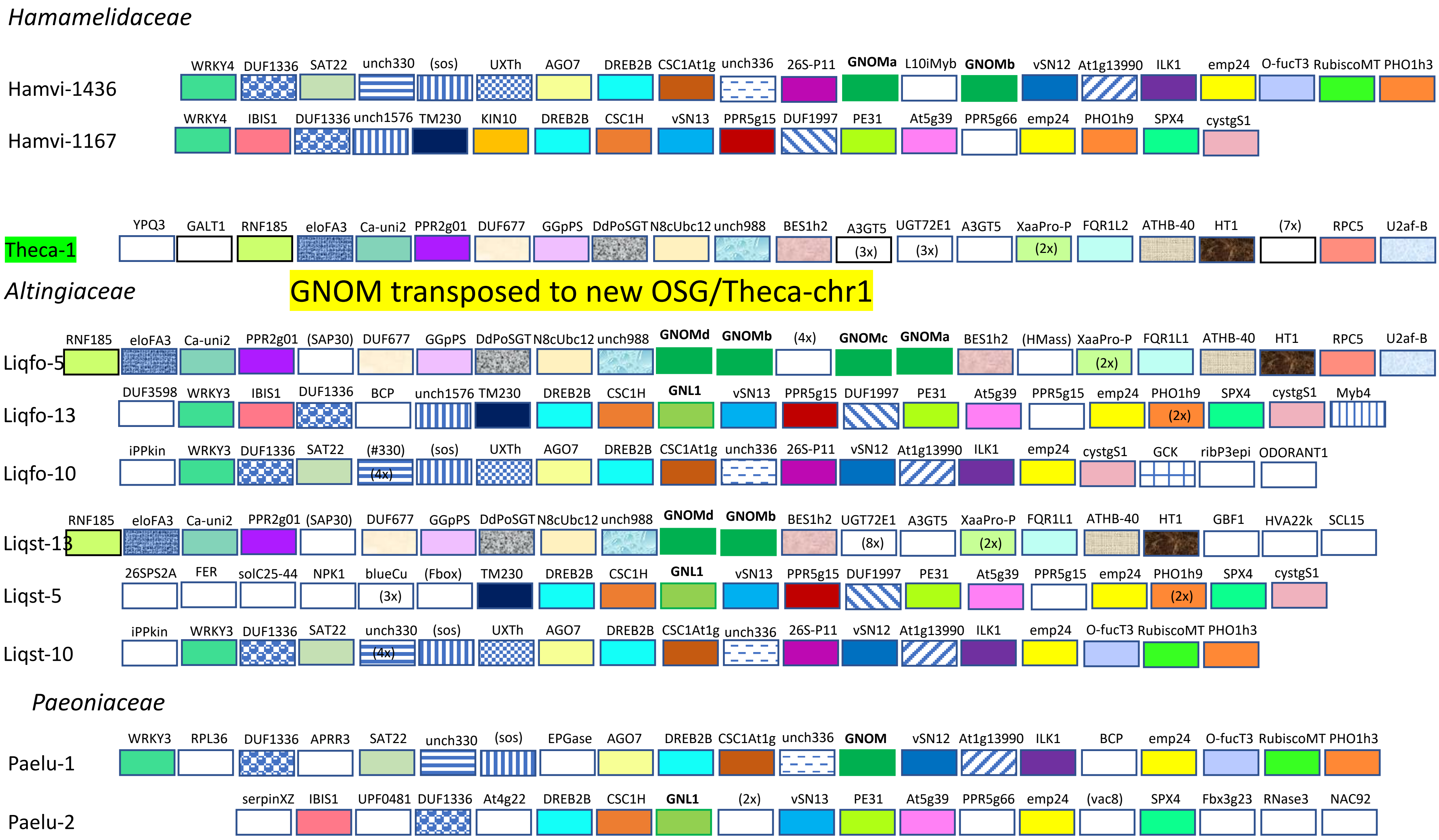

*Crassulaceae*

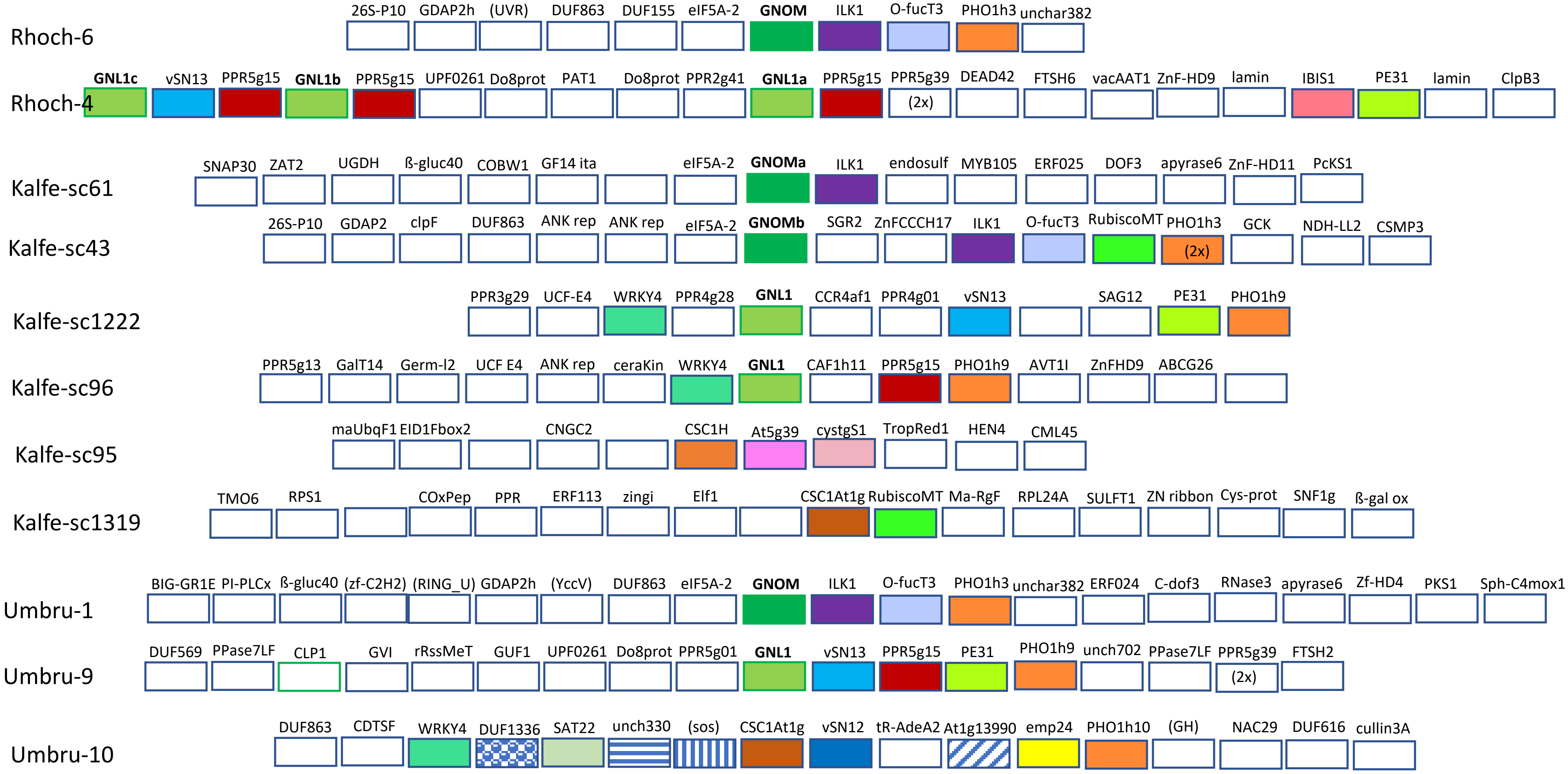

*Saxifragaceae*

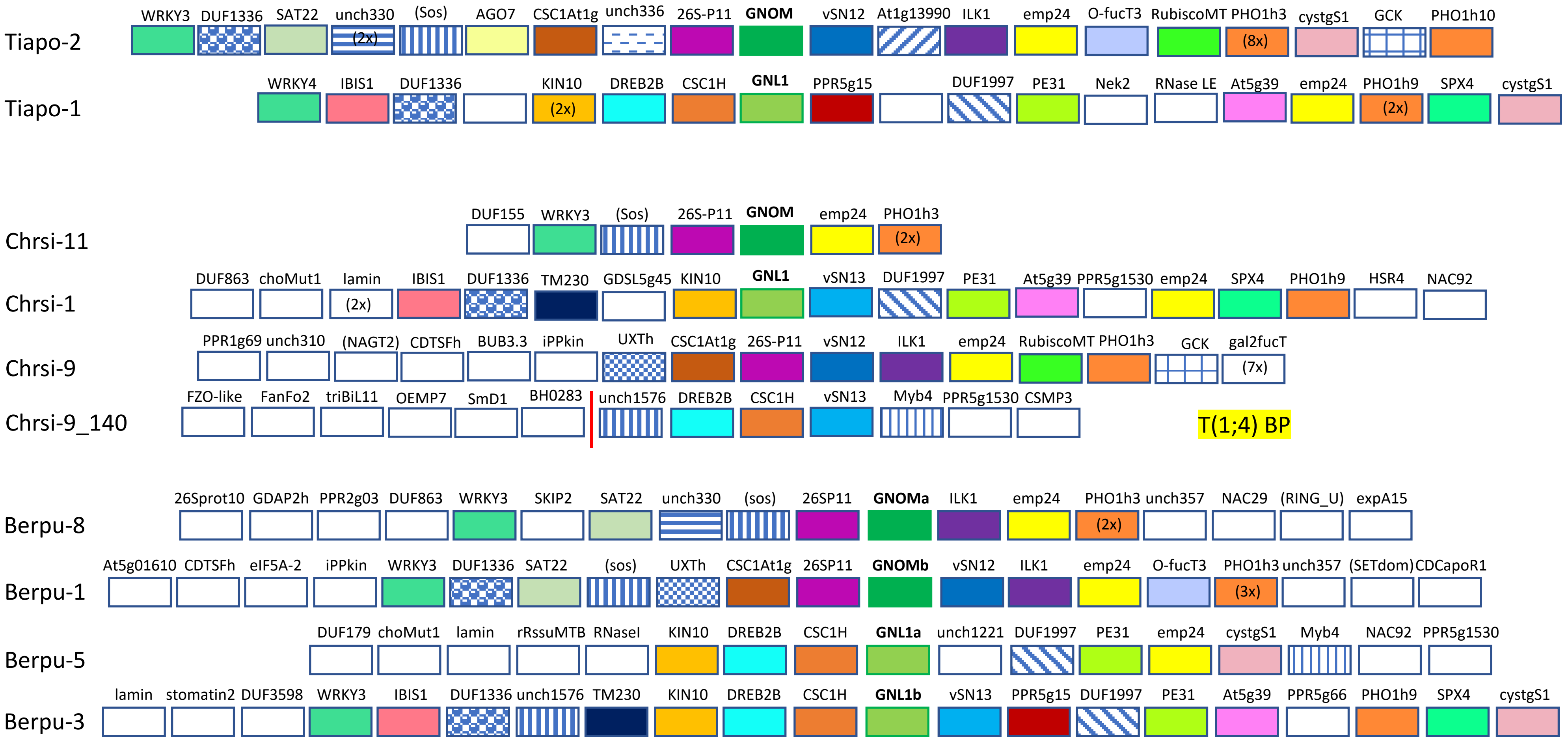

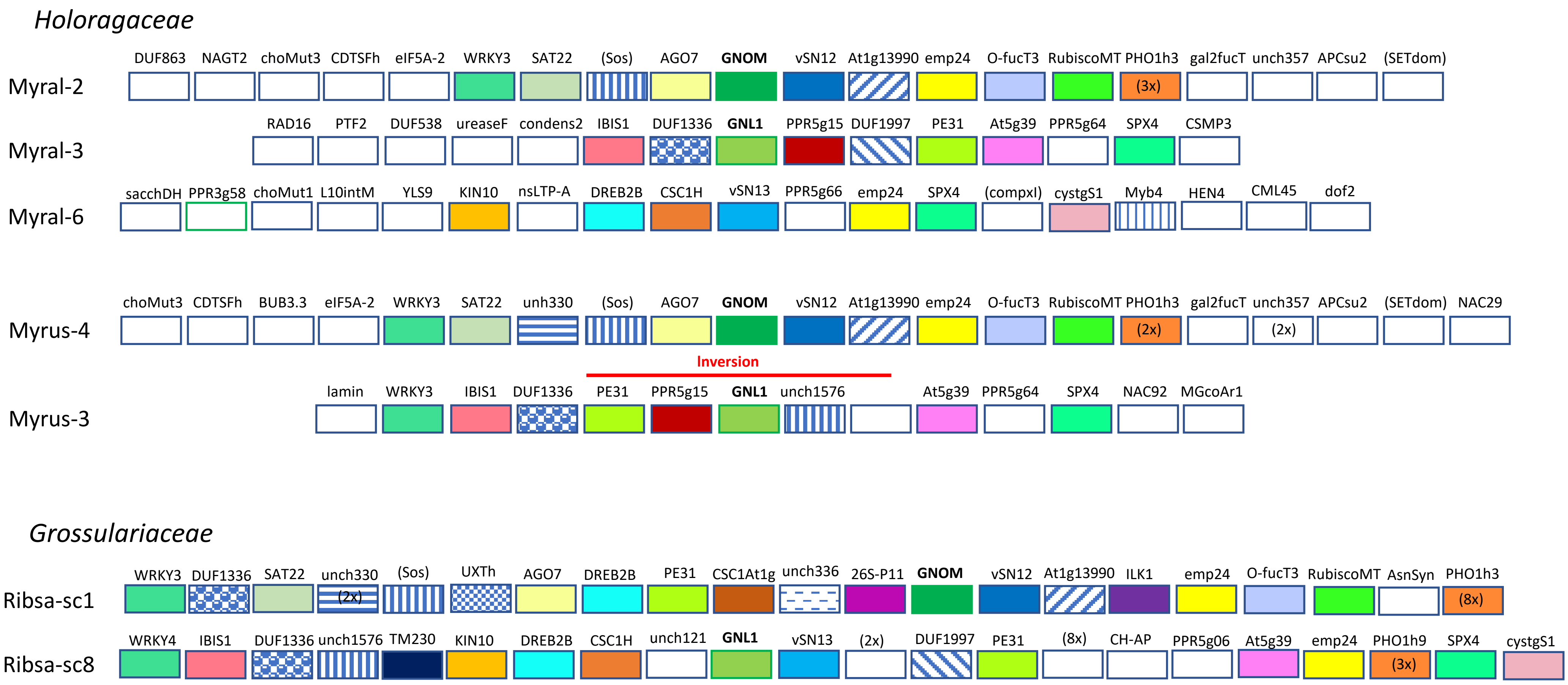
Holoragaceae
DUF863 NAGT2 choMut3 CDTSFh eIF5A-2 WRKY3 SAT22 (Sos) AGO7 GNOM vSN12 At1g13990 emp24 O-fucT3 RubiscoMT PHO1h3 gal2fucT unch357 APCsu2 (SETdom)
Myral-2
(3x)
RAD16 PTF2 DUF538 ureaseF condens2 IBIS1 DUF1336 GNL1 PPR5g15 DUF1997 PE31 At5g39 PPR5g64 SPX4 CSMP3
Myral-3
sacchDH PPR3g58 choMut1 L10intM YLS9 KIN10 nsLTP-A DREB2B CSC1H vSN13 PPR5g66 emp24 SPX4 (compxl) cystgS1 Myb4 HEN4 CML45 dof2
Myral-6
choMut3 CDTSFh BUB3.3 eIF5A-2 WRKY3 SAT22 unh330 (Sos) AGO7 GNOM vSN12 At1g13990 emp24 O-fucT3 RubiscoMT PHO1h3 gal2fucT unch357 APCsu2 (SETdom) NAC29
Myrus-4
(2x)
(2x)
Inversion
lamin WRKY3 IBIS1 DUF1336 PE31 PPR5g15 GNL1 unch1576 At5g39 PPR5g64 SPX4 NAC92 MGcoAr1
Myrus-3
Grossulariaceae
WRKY3 DUF1336 SAT22 unch330 (Sos) UXTh AGO7 DREB2B PE31 CSC1At1g unch336 26S-P11 GNOM vSN12 At1g13990 ILK1 emp24 O-fucT3 RubiscoMT AsnSyn PHO1h3
Ribsa-sc1
(2x)
(8x)
WRKY4 IBIS1 DUF1336 unch1576 TM230 KIN10 DREB2B CSC1H unch121 GNL1 vSN13 (2x) DUF1997 PE31 (8x) CH-AP PPR5g06 At5g39 emp24 PHO1h9 SPX4 cystgS1
Ribsa-sc8
(3x)

**Suppl. Figure S19. Vitales cladogram and OSGs**

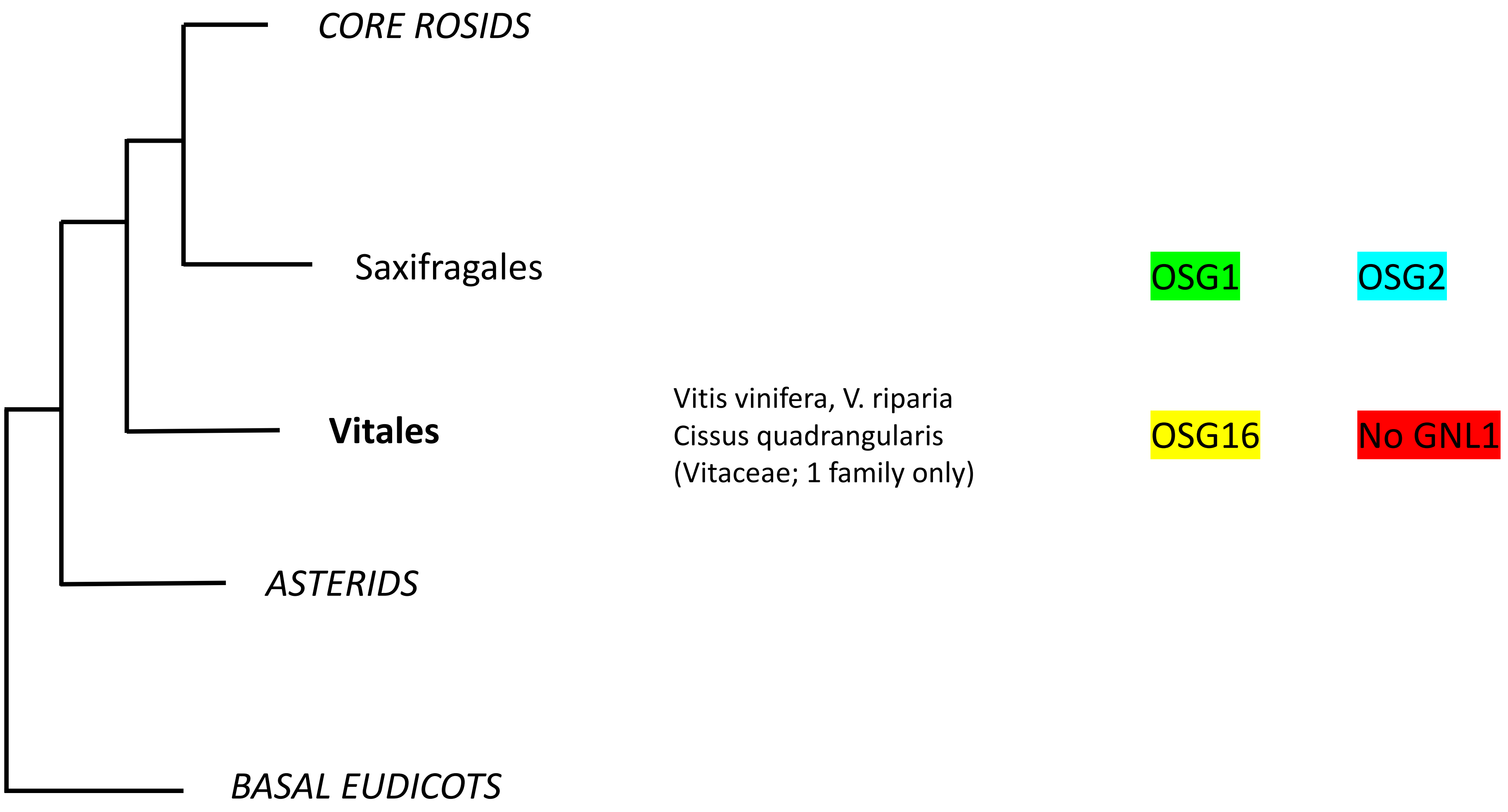

### GNOM-OSG16 (Theca, chr3)

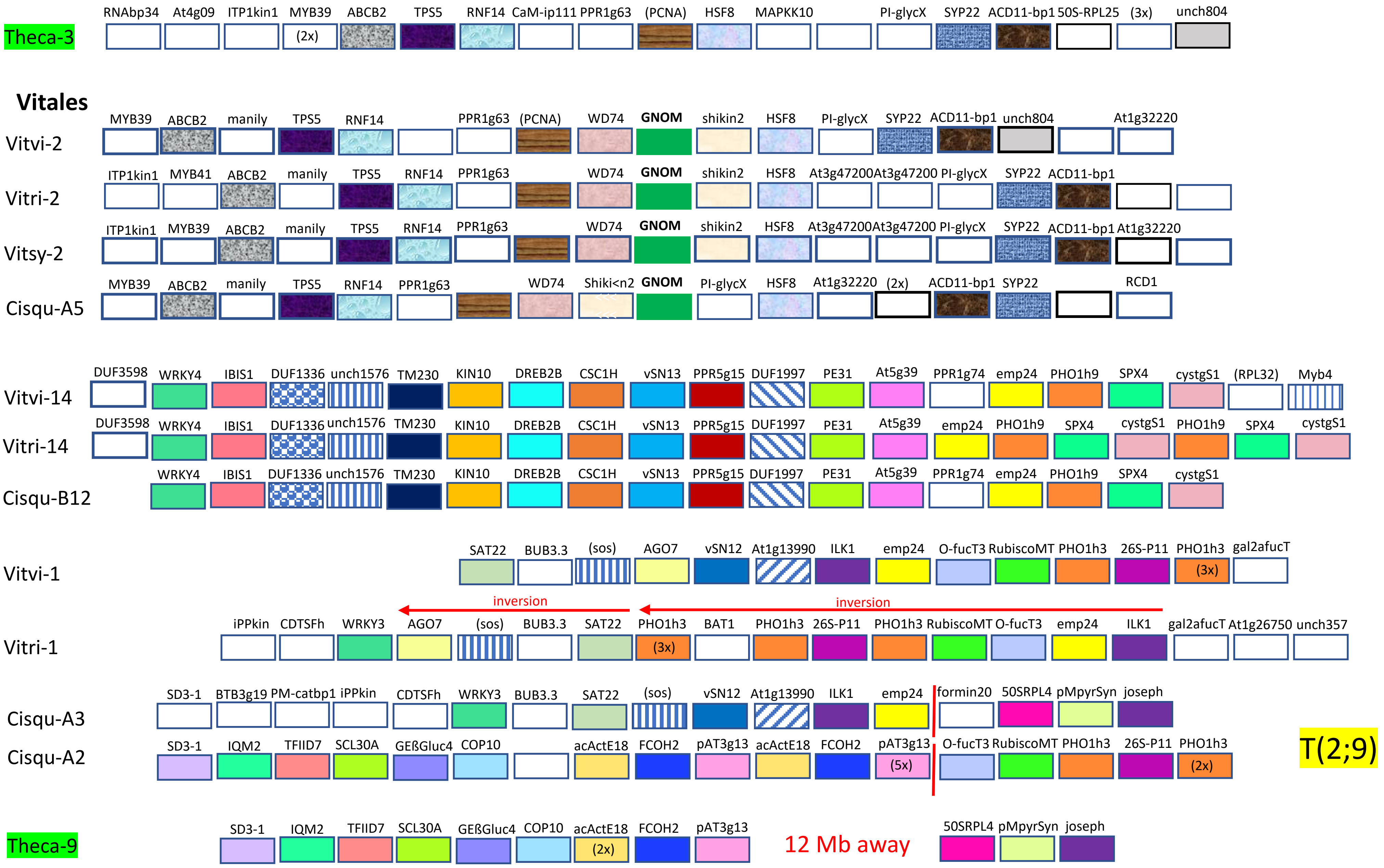

**Suppl. Figure S20. Santalales cladogram and OSGs**

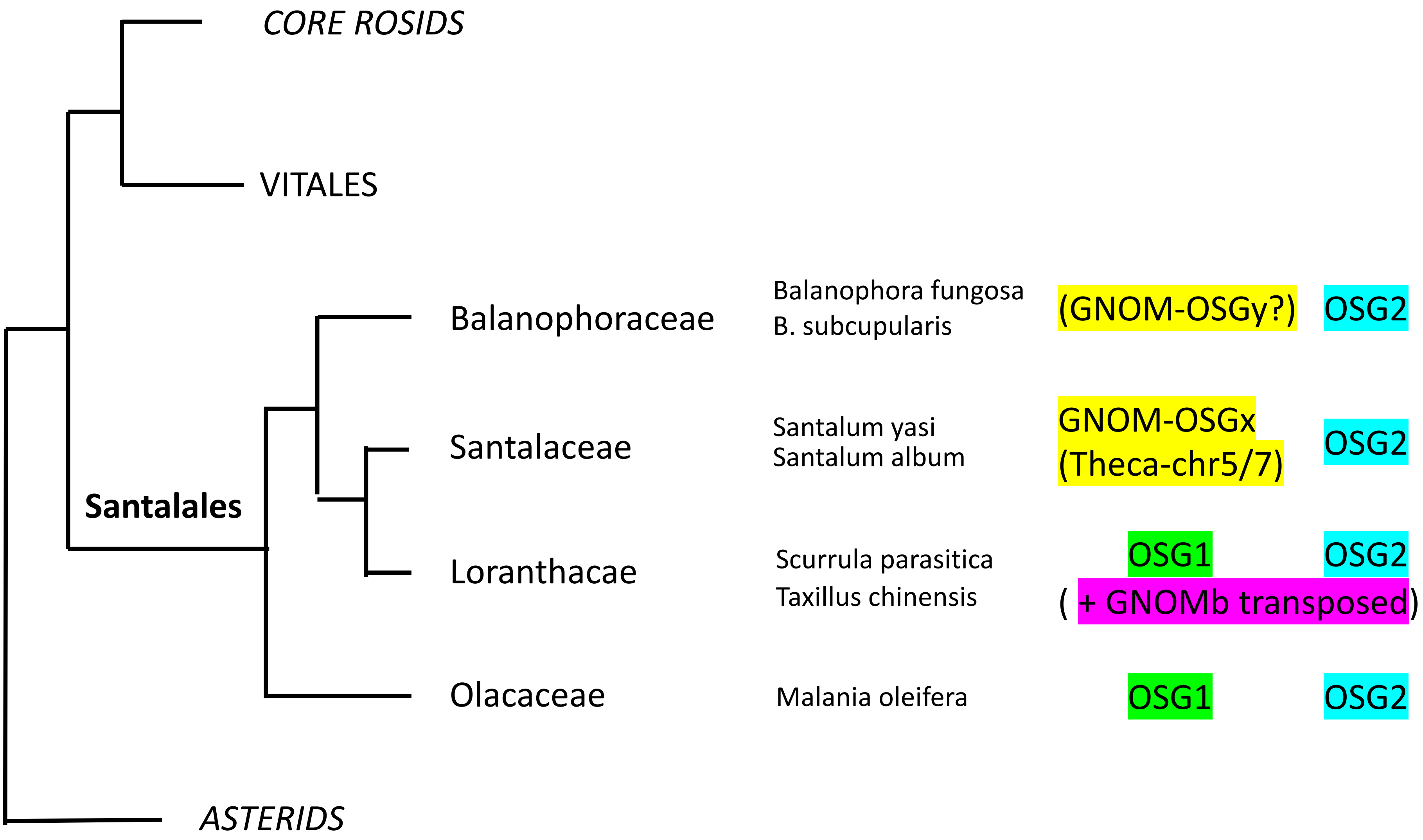

Cladogram based on Zuntini et al. (2024)

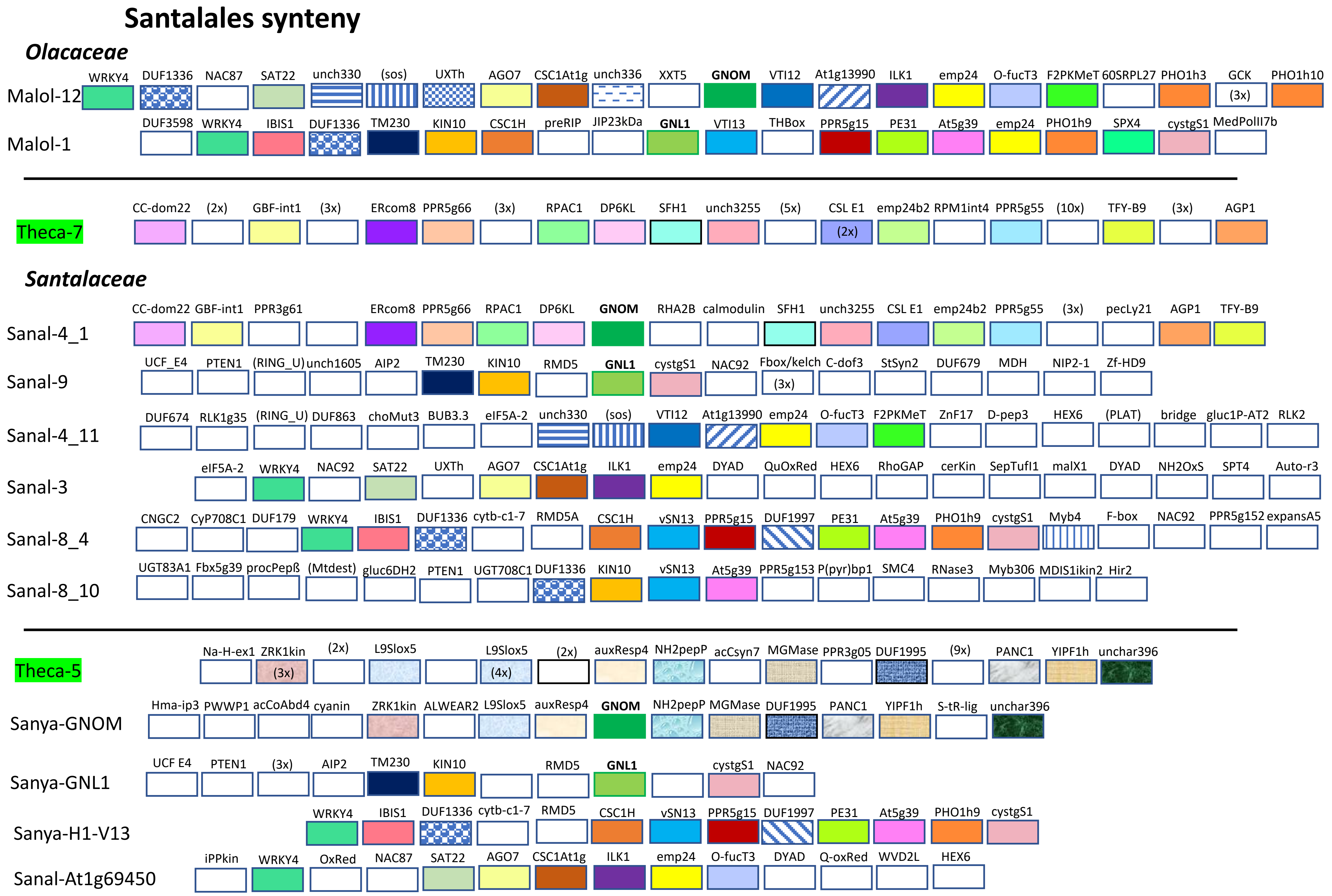
Santalales synteny
Olacaceae
Malol-12 WRKY4 DUF1336 NAC87 SAT22 unch330 (sos) UXTh AGO7 CSC1At1g unch336 XXT5 GNOM VTI12 At1g13990 ILK1 emp24 O-fucT3 F2PKMeT 60SRPL27 PHO1h3 GCK (3x) PHO1h10
Malol-1 DUF3598 WRKY4 IBIS1 DUF1336 TM230 KIN10 CSC1H preRIP JIP23kDa GNL1 VTI13 THBox PPR5g15 PE31 At5g39 emp24 PHO1h9 SPX4 cystgS1 MedPolII7b
Theca-7 CC-dom22 (2x) GBF-int1 (3x) ERcom8 PPR5g66 (3x) RPAC1 DP6KL SFH1 unch3255 (5x) CSL E1 (2x) emp24b2 RPM1int4 PPR5g55 (10x) TFY-B9 (3x) AGP1
Santalaceae
Sanal-4_1 CC-dom22 GBF-int1 PPR3g61 ERcom8 PPR5g66 RPAC1 DP6KL GNOM RHA2B calmodulin SFH1 unch3255 CSL E1 emp24b2 PPR5g55 (3x) pecLy21 AGP1 TFY-B9
Sanal-9 UCF_E4 PTEN1 (RING_U) unch1605 AIP2 TM230 KIN10 RMD5 GNL1 cystgS1 NAC92 Fbox/kelch (3x) C-dof3 StSyn2 DUF679 MDH NIP2-1 Zf-HD9
Sanal-4_11 DUF674 RLK1g35 (RING_U) DUF863 choMut3 BUB3.3 eIF5A-2 unch330 (sos) VTI12 At1g13990 emp24 O-fucT3 F2PKMeT ZnF17 D-pep3 HEX6 (PLAT) bridge gluc1P-AT2 RLK2
Sanal-3 eIF5A-2 WRKY4 NAC92 SAT22 UXTh AGO7 CSC1At1g ILK1 emp24 DYAD QuOxRed HEX6 RhoGAP cerKin SepTufI1 malX1 DYAD NH2OxS SPT4 Auto-r3
Sanal-8_4 CNGC2 CyP708C1 DUF179 WRKY4 IBIS1 DUF1336 cytb-c1-7 RMD5A CSC1H vSN13 PPR5g15 DUF1997 PE31 At5g39 PHO1h9 cystgS1 Myb4 F-box NAC92 PPR5g152 expansA5
Sanal-8_10 UGT83A1 Fbx5g39 procPepß (Mtdest) gluc6DH2 PTEN1 UGT708C1 DUF1336 KIN10 vSN13 At5g39 PPR5g153 P(pyr)bp1 SMC4 RNase3 Myb306 MDIS1ikin2 Hir2
Theca-5 Na-H-ex1 ZRK1kin (3x) (2x) L9Slox5 L9Slox5 (4x) (2x) auxResp4 NH2pepP acCsyn7 MGMase PPR3g05 DUF1995 (9x) PANC1 YIPF1h unchar396
Sanya-GNOM Hma-ip3 PWWP1 acCoAbd4 cyanin ZRK1kin ALWEAR2 L9Slox5 auxResp4 GNOM NH2pepP MGMase DUF1995 PANC1 YIPF1h S-tR-lig unchar396
Sanya-GNL1 UCF E4 PTEN1 (3x) AIP2 TM230 KIN10 RMD5 GNL1 cystgS1 NAC92
Sanya-H1-V13 WRKY4 IBIS1 DUF1336 cytb-c1-7 RMD5 CSC1H vSN13 PPR5g15 DUF1997 PE31 At5g39 PHO1h9 cystgS1
Sanal-At1g69450 iPPkin WRKY4 OxRed NAC87 SAT22 AGO7 CSC1At1g ILK1 emp24 O-fucT3 DYAD Q-oxRed WVD2L HEX6

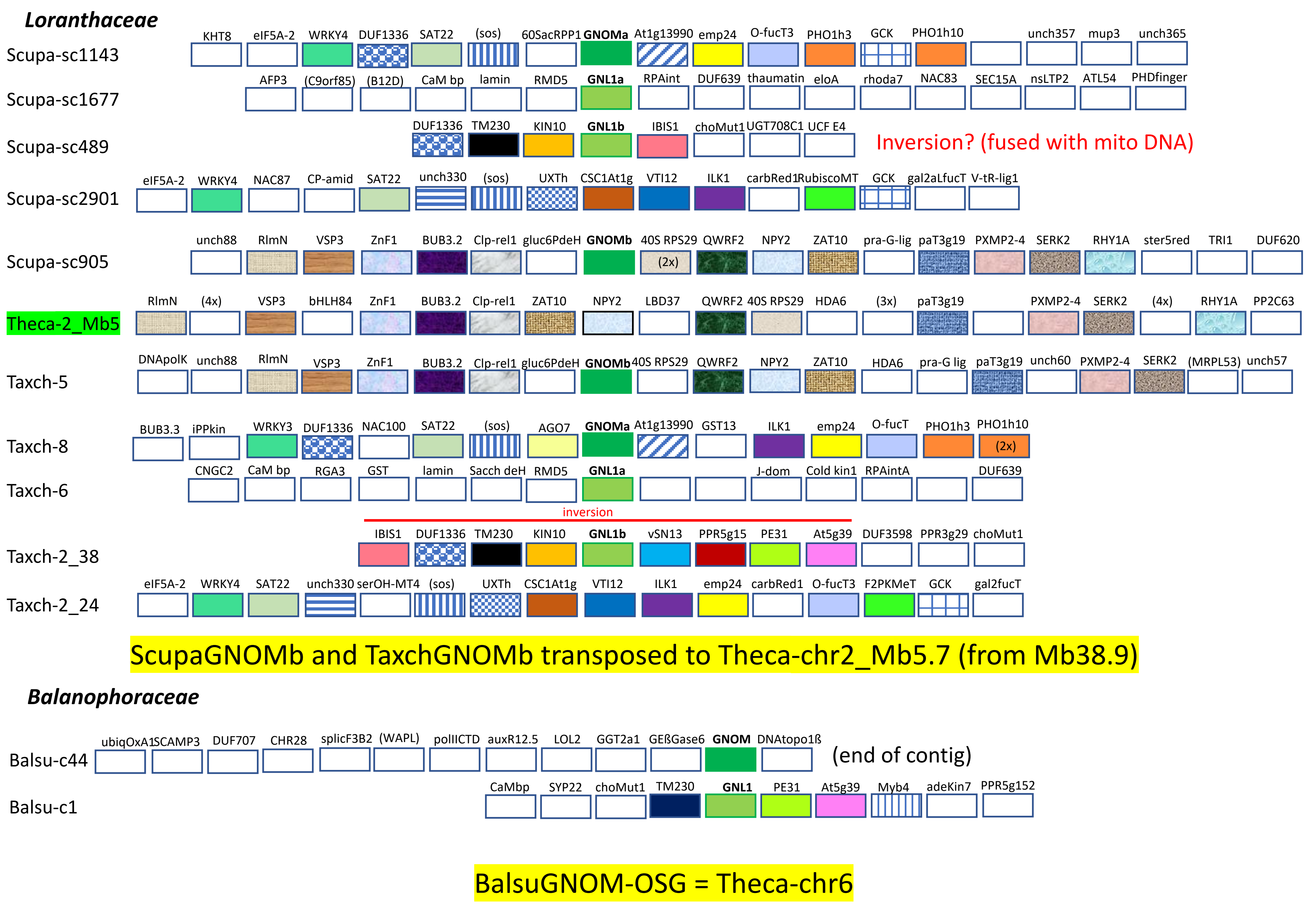

Loranthaceae
Scupa-sc1143
KHT8 eIF5A-2 WRKY4 DUF1336 SAT22 (sos) 60SacRPP1 GNOMa At1g13990 emp24 O-fucT3 PHO1h3 GCK PHO1h10 unch357 mup3 unch365
Scupa-sc1677
AFP3 (C9orf85) (B12D) CaM bp lamin RMD5 GNL1a RPAint DUF639 thaumatin eloA rhoda7 NAC83 SEC15A nsLTP2 ATL54 PHDfinger
Scupa-sc489
DUF1336 TM230 KIN10 GNL1b IBIS1 choMut1 UGT708C1 UCF E4
Inversion? (fused with mito DNA)
Scupa-sc2901
eIF5A-2 WRKY4 NAC87 CP-amid SAT22 unch330 (sos) UXTh CSC1At1g VTI12 ILK1 carbRed1 RubiscoMT GCK gal2aLfucT V-tR-lig1
Scupa-sc905
unch88 RlmN VSP3 ZnF1 BUB3.2 Clp-rel1 gluc6PdeH GNOMb 40S RPS29 (2x) QWRF2 NPY2 ZAT10 pra-G-lig paT3g19 PXMP2-4 SERK2 RHY1A ster5red TRI1 DUF620
Theca-2_Mb5
RlmN (4x) VSP3 bHLH84 ZnF1 BUB3.2 Clp-rel1 ZAT10 NPY2 LBD37 QWRF2 40S RPS29 HDA6 (3x) paT3g19 PXMP2-4 SERK2 (4x) RHY1A PP2C63
Taxch-5
DNApolK unch88 RlmN VSP3 ZnF1 BUB3.2 Clp-rel1 gluc6PdeH GNOMb 40S RPS29 QWRF2 NPY2 ZAT10 HDA6 pra-G lig paT3g19 unch60 PXMP2-4 SERK2 (MRPL53) unch57
Taxch-8
BUB3.3 iPPkin WRKY3 DUF1336 NAC100 SAT22 (sos) AGO7 GNOMa At1g13990 GST13 ILK1 emp24 O-fucT PHO1h3 PHO1h10 (2x)
Taxch-6
CNGC2 CaM bp RGA3 GST lamin Sacch deH RMD5 GNL1a J-dom Cold kin1 RPAintA DUF639
inversion
Taxch-2_38
IBIS1 DUF1336 TM230 KIN10 GNL1b vSN13 PPR5g15 PE31 At5g39 DUF3598 PPR3g29 choMut1
Taxch-2_24
eIF5A-2 WRKY4 SAT22 unch330 serOH-MT4 (sos) UXTh CSC1At1g VTI12 ILK1 emp24 carbRed1 O-fucT3 F2PKMeT GCK gal2fucT
ScupaGNOMb and TaxchGNOMb transposed to Theca-chr2_Mb5.7 (from Mb38.9)
Balanophoraceae
Balsu-c44
ubiqOxA1 SCAMP3 DUF707 CHR28 splicF3B2 (WAPL) polIICTD auxR12.5 LOL2 GGT2a1 GEßGase6 GNOM DNAtopo1ß
(end of contig)
Balsu-c1
CaMbp SYP22 choMut1 TM230 GNL1 PE31 At5g39 Myb4 adeKin7 PPR5g152
BalsuGNOM-OSG = Theca-chr6

### Suppl. Figure S21. Lamiales cladogram and OSGs

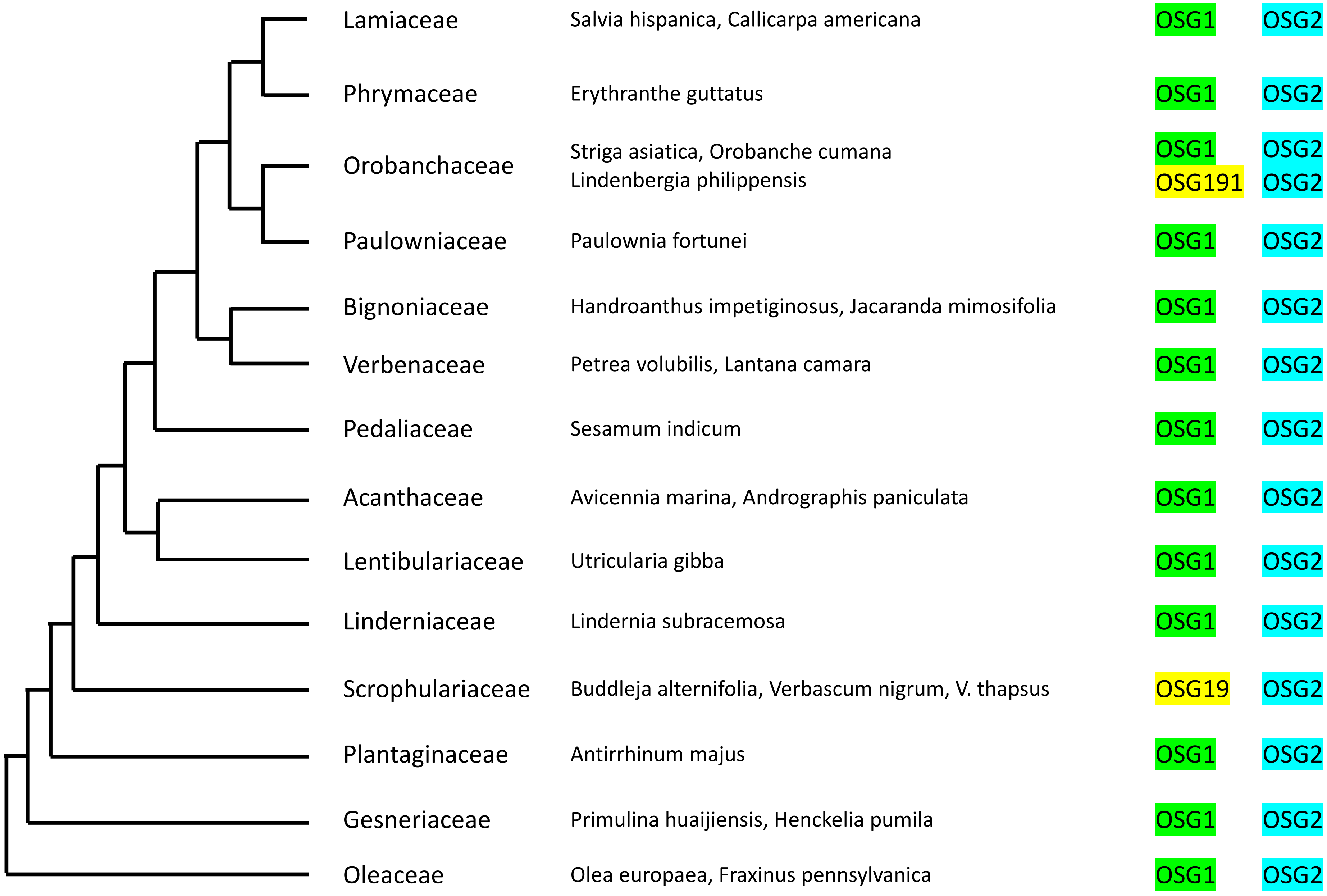

Cladogram based on OTPTI (2019), Fonseca (2021)

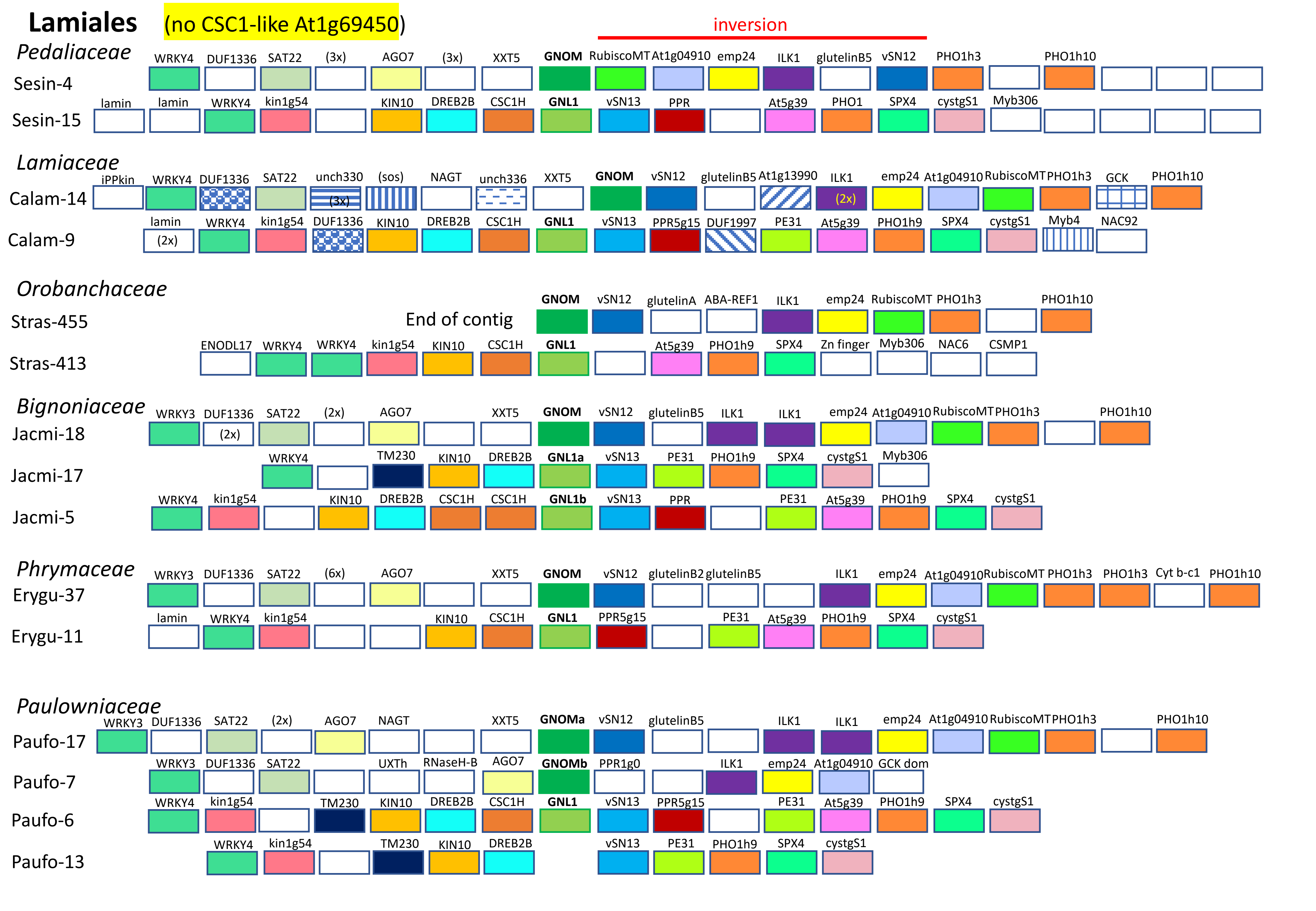
Lamiales
(no CSC1-like At1g69450)
inversion
Pedaliaceae
Sesin-4
WRKY4 DUF1336 SAT22 (3x) AGO7 (3x) XXT5 GNOM RubiscoMT At1g04910 emp24 ILK1 glutelinB5 vSN12 PHO1h3 PHO1h10
Sesin-15
lamin lamin WRKY4 kin1g54 KIN10 DREB2B CSC1H GNL1 vSN13 PPR At5g39 PHO1 SPX4 cystgS1 Myb306
Lamiaceae
Calam-14
iPPkin WRKY4 DUF1336 SAT22 unch330 (3x) (sos) NAGT unch336 XXT5 GNOM vSN12 glutelinB5 At1g13990 ILK1 (2x) emp24 At1g04910 RubiscoMT PHO1h3 GCK PHO1h10
Calam-9
lamin (2x) WRKY4 kin1g54 DUF1336 KIN10 DREB2B CSC1H GNL1 vSN13 PPR5g15 DUF1997 PE31 At5g39 PHO1h9 SPX4 cystgS1 Myb4 NAC92
Orobanchaceae
Stras-455
End of contig
GNOM vSN12 glutelinA ABA-REF1 ILK1 emp24 RubiscoMT PHO1h3 PHO1h10
Stras-413
ENODL17 WRKY4 WRKY4 kin1g54 KIN10 CSC1H GNL1 At5g39 PHO1h9 SPX4 Zn finger Myb306 NAC6 CSMP1
Bignoniaceae
Jacmi-18
WRKY3 DUF1336 (2x) SAT22 (2x) AGO7 XXT5 GNOM vSN12 glutelinB5 ILK1 ILK1 emp24 At1g04910 RubiscoMT PHO1h3 PHO1h10
Jacmi-17
WRKY4 TM230 KIN10 DREB2B GNL1a vSN13 PE31 PHO1h9 SPX4 cystgS1 Myb306
Jacmi-5
WRKY4 kin1g54 KIN10 DREB2B CSC1H CSC1H GNL1b vSN13 PPR PE31 At5g39 PHO1h9 SPX4 cystgS1
Phrymaceae
Erygu-37
WRKY3 DUF1336 SAT22 (6x) AGO7 XXT5 GNOM vSN12 glutelinB2 glutelinB5 ILK1 emp24 At1g04910 RubiscoMT PHO1h3 PHO1h3 Cyt b-c1 PHO1h10
Erygu-11
lamin WRKY4 kin1g54 KIN10 CSC1H GNL1 PPR5g15 PE31 At5g39 PHO1h9 SPX4 cystgS1
Paulowniaceae
Paufo-17
WRKY3 DUF1336 SAT22 (2x) AGO7 NAGT XXT5 GNOMa vSN12 glutelinB5 ILK1 ILK1 emp24 At1g04910 RubiscoMT PHO1h3 PHO1h10
Paufo-7
WRKY3 DUF1336 SAT22 UXTh RNaseH-B AGO7 GNOMb PPR1g0 ILK1 emp24 At1g04910 GCK dom
Paufo-6
WRKY4 kin1g54 TM230 KIN10 DREB2B CSC1H GNL1 vSN13 PPR5g15 PE31 At5g39 PHO1h9 SPX4 cystgS1
Paufo-13
WRKY4 kin1g54 TM230 KIN10 DREB2B vSN13 PE31 PHO1h9 SPX4 cystgS1

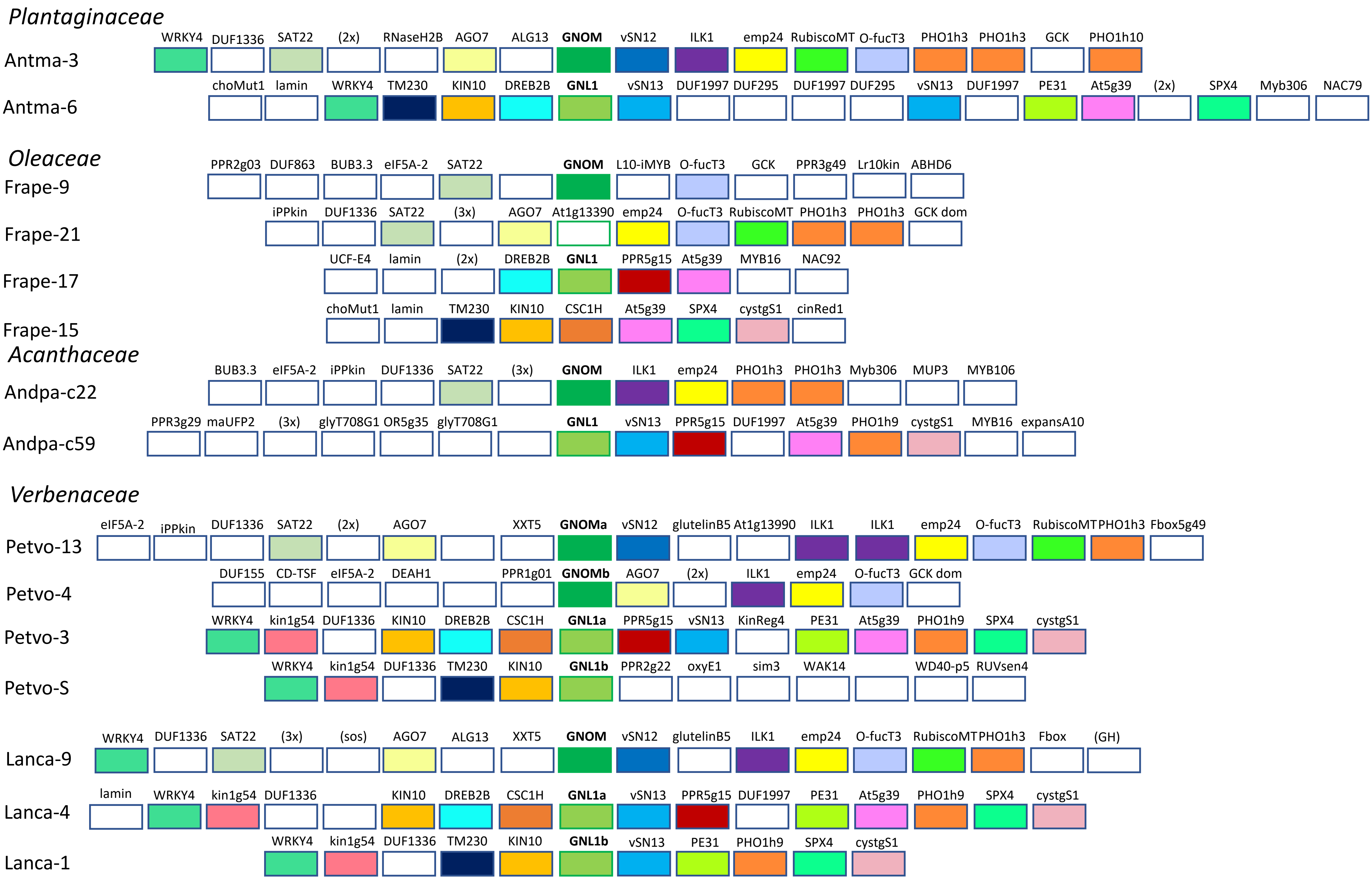
Plantaginaceae
Antma-3
WRKY4 DUF1336 SAT22 (2x) RNaseH2B AGO7 ALG13 GNOM vSN12 ILK1 emp24 RubiscoMT O-fucT3 PHO1h3 PHO1h3 GCK PHO1h10
Antma-6
choMut1 lamin WRKY4 TM230 KIN10 DREB2B GNL1 vSN13 DUF1997 DUF295 DUF1997 DUF295 vSN13 DUF1997 PE31 At5g39 (2x) SPX4 Myb306 NAC79
Oleaceae
Frape-9
PPR2g03 DUF863 BUB3.3 eIF5A-2 SAT22 GNOM L10-iMYB O-fucT3 GCK PPR3g49 Lr10kin ABHD6
Frape-21
iPPkin DUF1336 SAT22 (3x) AGO7 At1g13390 emp24 O-fucT3 RubiscoMT PHO1h3 PHO1h3 GCK dom
Frape-17
UCF-E4 lamin (2x) DREB2B GNL1 PPR5g15 At5g39 MYB16 NAC92
Frape-15
choMut1 lamin TM230 KIN10 CSC1H At5g39 SPX4 cystgS1 cinRed1
Acanthaceae
Andpa-c22
BUB3.3 eIF5A-2 iPPkin DUF1336 SAT22 (3x) GNOM ILK1 emp24 PHO1h3 PHO1h3 Myb306 MUP3 MYB106
Andpa-c59
PPR3g29 maUFP2 (3x) glyT708G1 OR5g35 glyT708G1 GNL1 vSN13 PPR5g15 DUF1997 At5g39 PHO1h9 cystgS1 MYB16 expansA10
Verbenaceae
Petvo-13
eIF5A-2 iPPkin DUF1336 SAT22 (2x) AGO7 XXT5 GNOMa vSN12 glutelinB5 At1g13990 ILK1 ILK1 emp24 O-fucT3 RubiscoMT PHO1h3 Fbox5g49
Petvo-4
DUF155 CD-TSF eIF5A-2 DEAH1 PPR1g01 GNOMb AGO7 (2x) ILK1 emp24 O-fucT3 GCK dom
Petvo-3
WRKY4 kin1g54 DUF1336 KIN10 DREB2B CSC1H GNL1a PPR5g15 vSN13 KinReg4 PE31 At5g39 PHO1h9 SPX4 cystgS1
Petvo-S
WRKY4 kin1g54 DUF1336 TM230 KIN10 GNL1b PPR2g22 oxyE1 sim3 WAK14 WD40-p5 RUVsen4
Lanca-9
WRKY4 DUF1336 SAT22 (3x) (sos) AGO7 ALG13 XXT5 GNOM vSN12 glutelinB5 ILK1 emp24 O-fucT3 RubiscoMT PHO1h3 Fbox (GH)
Lanca-4
lamin WRKY4 kin1g54 DUF1336 KIN10 DREB2B CSC1H GNL1a vSN13 PPR5g15 DUF1997 PE31 At5g39 PHO1h9 SPX4 cystgS1
Lanca-1
WRKY4 kin1g54 DUF1336 TM230 KIN10 GNL1b vSN13 PE31 PHO1h9 SPX4 cystgS1

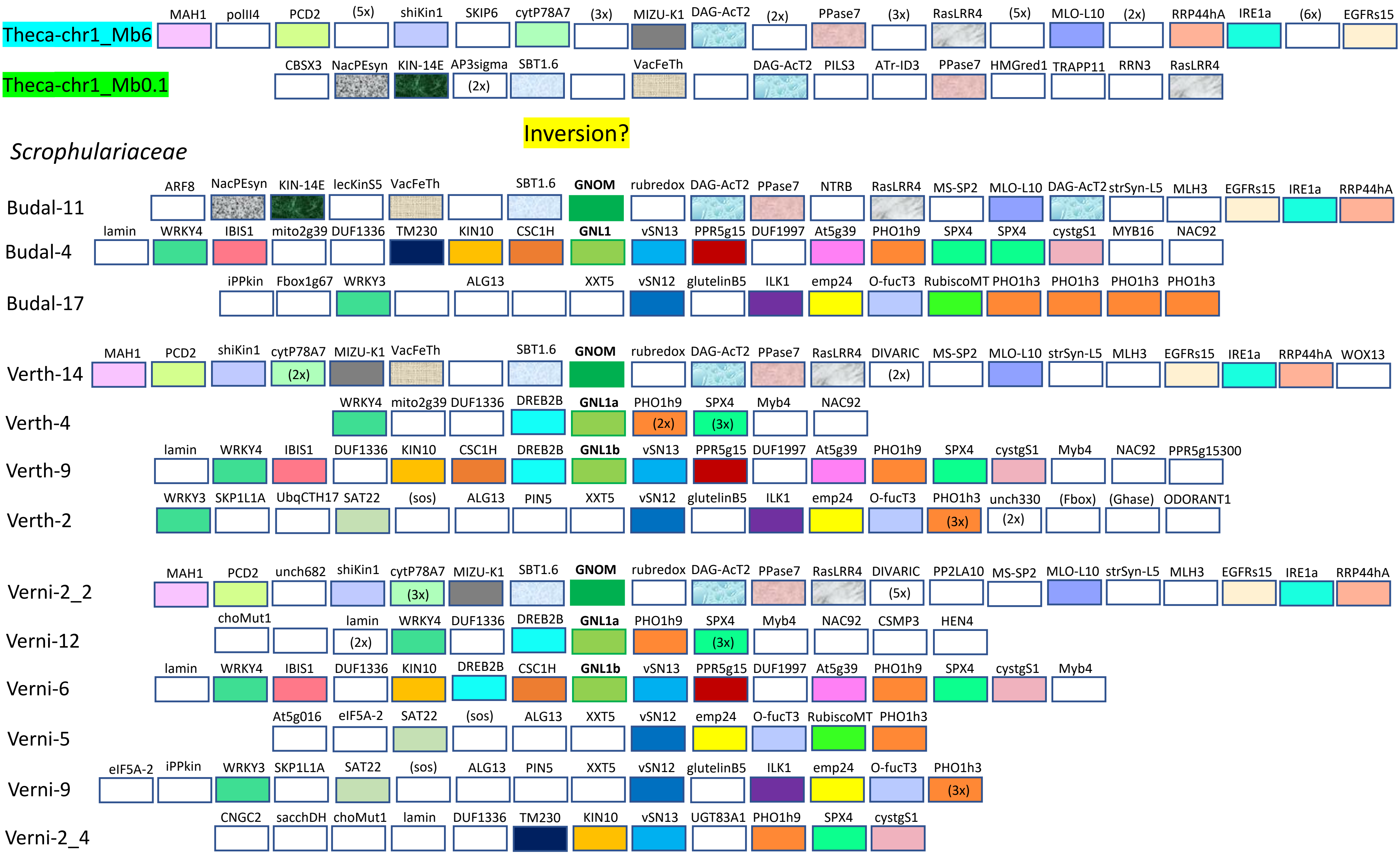

Theca-chr1_Mb6
MAH1 polII4 PCD2 (5x) shiKin1 SKIP6 cytP78A7 (3x) MIZU-K1 DAG-AcT2 (2x) PPase7 (3x) RasLRR4 (5x) MLO-L10 (2x) RRP44hA IRE1a (6x) EGFRs15
Theca-chr1_Mb0.1
CBSX3 NacPEsyn KIN-14E AP3sigma (2x) SBT1.6 VacFeTh DAG-AcT2 PILS3 ATr-ID3 PPase7 HMGred1 TRAPP11 RRN3 RasLRR4
Inversion?
Scrophulariaceae
Budal-11
ARF8 NacPEsyn KIN-14E lecKinS5 VacFeTh SBT1.6 GNOM rubredox DAG-AcT2 PPase7 NTRB RasLRR4 MS-SP2 MLO-L10 DAG-AcT2 strSyn-L5 MLH3 EGFRs15 IRE1a RRP44hA
Budal-4
lamin WRKY4 IBIS1 mito2g39 DUF1336 TM230 KIN10 CSC1H GNL1 vSN13 PPR5g15 DUF1997 At5g39 PHO1h9 SPX4 SPX4 cystgS1 MYB16 NAC92
Budal-17
iPPkin Fbox1g67 WRKY3 ALG13 XXT5 vSN12 glutelinB5 ILK1 emp24 O-fucT3 RubiscoMT PHO1h3 PHO1h3 PHO1h3 PHO1h3
Verth-14
MAH1 PCD2 shiKin1 cytP78A7 (2x) MIZU-K1 VacFeTh SBT1.6 GNOM rubredox DAG-AcT2 PPase7 RasLRR4 DIVARIC (2x) MS-SP2 MLO-L10 strSyn-L5 MLH3 EGFRs15 IRE1a RRP44hA WOX13
Verth-4
WRKY4 mito2g39 DUF1336 DREB2B GNL1a PHO1h9 (2x) SPX4 (3x) Myb4 NAC92
Verth-9
lamin WRKY4 IBIS1 DUF1336 KIN10 CSC1H DREB2B GNL1b vSN13 PPR5g15 DUF1997 At5g39 PHO1h9 SPX4 cystgS1 Myb4 NAC92 PPR5g15300
Verth-2
WRKY3 SKP1L1A UbqCTH17 SAT22 (sos) ALG13 PIN5 XXT5 vSN12 glutelinB5 ILK1 emp24 O-fucT3 PHO1h3 (3x) unch330 (2x) (Fbox) (Ghase) ODORANT1
Verni-2_2
MAH1 PCD2 unch682 shiKin1 cytP78A7 (3x) MIZU-K1 SBT1.6 GNOM rubredox DAG-AcT2 PPase7 RasLRR4 DIVARIC (5x) PP2LA10 MS-SP2 MLO-L10 strSyn-L5 MLH3 EGFRs15 IRE1a RRP44hA
Verni-12
choMut1 lamin (2x) WRKY4 DUF1336 DREB2B GNL1a PHO1h9 SPX4 (3x) Myb4 NAC92 CSMP3 HEN4
Verni-6
lamin WRKY4 IBIS1 DUF1336 KIN10 DREB2B CSC1H GNL1b vSN13 PPR5g15 DUF1997 At5g39 PHO1h9 SPX4 cystgS1 Myb4
Verni-5
At5g016 eIF5A-2 SAT22 (sos) ALG13 XXT5 vSN12 emp24 O-fucT3 RubiscoMT PHO1h3
Verni-9
eIF5A-2 iPPkin WRKY3 SKP1L1A SAT22 (sos) ALG13 PIN5 XXT5 vSN12 glutelinB5 ILK1 emp24 O-fucT3 PHO1h3 (3x)
Verni-2_4
CNGC2 sacchDH choMut1 lamin DUF1336 TM230 KIN10 vSN13 UGT83A1 PHO1h9 SPX4 cystgS1

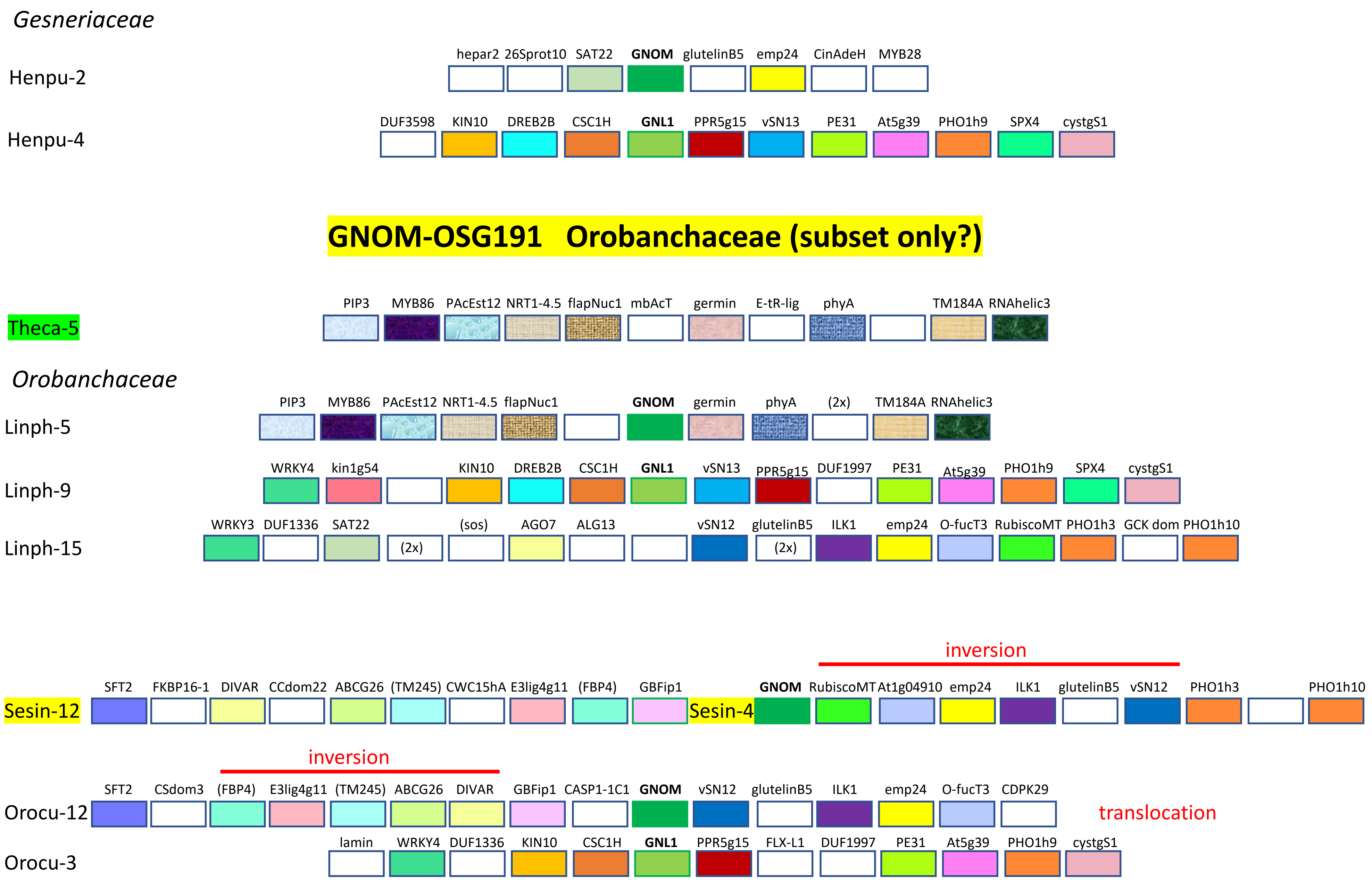
Gesneriaceae
Henpu-2
hepar2 26Sprot10 SAT22 GNOM glutelinB5 emp24 CinAdeH MYB28
Henpu-4
DUF3598 KIN10 DREB2B CSC1H GNL1 PPR5g15 vSN13 PE31 At5g39 PHO1h9 SPX4 cystgS1
GNOM-OSG191 Orobanchaceae (subset only?)
Theca-5
PIP3 MYB86 PAcEst12 NRT1-4.5 flapNuc1 mbAcT germin E-tR-lig phyA TM184A RNAhelic3
Orobanchaceae
Linph-5
PIP3 MYB86 PAcEst12 NRT1-4.5 flapNuc1 GNOM germin phyA (2x) TM184A RNAhelic3
Linph-9
WRKY4 kin1g54 KIN10 DREB2B CSC1H GNL1 vSN13 PPR5g15 DUF1997 PE31 At5g39 PHO1h9 SPX4 cystgS1
Linph-15
WRKY3 DUF1336 SAT22 (2x) (sos) AGO7 ALG13 vSN12 glutelinB5 (2x) ILK1 emp24 O-fucT3 RubiscoMT PHO1h3 GCK dom PHO1h10
inversion
Sesin-12
SFT2 FKBP16-1 DIVAR CCdom22 ABCG26 (TM245) CWC15hA E3lig4g11 (FBP4) GBFip1
Sesin-4
GNOM RubiscoMT At1g04910 emp24 ILK1 glutelinB5 vSN12 PHO1h3 PHO1h10
inversion
Orocu-12
SFT2 CSdom3 (FBP4) E3lig4g11 (TM245) ABCG26 DIVAR GBFip1 CASP1-1C1 GNOM vSN12 glutelinB5 ILK1 emp24 O-fucT3 CDPK29
translocation
Orocu-3
lamin WRKY4 DUF1336 KIN10 CSC1H GNL1 PPR5g15 FLX-L1 DUF1997 PE31 At5g39 PHO1h9 cystgS1

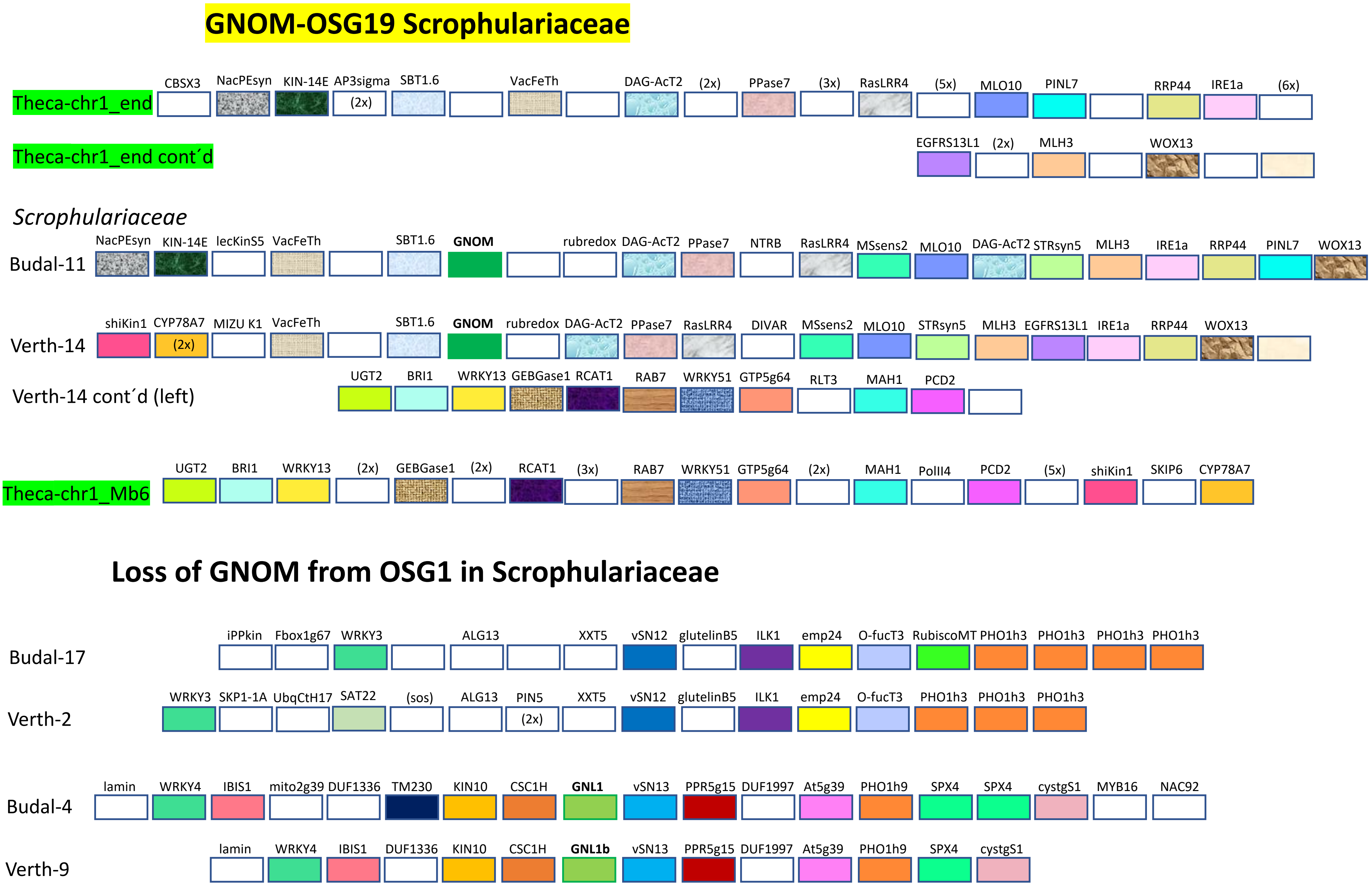
GNOM-OSG19 Scrophulariaceae
Theca-chr1_end
CBSX3 NacPEsyn KIN-14E AP3sigma (2x) SBT1.6 VacFeTh DAG-AcT2 (2x) PPase7 (3x) RasLRR4 (5x) MLO10 PINL7 RRP44 IRE1a (6x)
Theca-chr1_end cont´d
EGFRS13L1 (2x) MLH3 WOX13
Scrophulariaceae
Budal-11
NacPEsyn KIN-14E lecKinS5 VacFeTh SBT1.6 GNOM rubredox DAG-AcT2 PPase7 NTRB RasLRR4 MSsens2 MLO10 DAG-AcT2 STRsyn5 MLH3 IRE1a RRP44 PINL7 WOX13
Verth-14
shiKin1 CYP78A7 (2x) MIZU K1 VacFeTh SBT1.6 GNOM rubredox DAG-AcT2 PPase7 RasLRR4 DIVAR MSsens2 MLO10 STRsyn5 MLH3 EGFRS13L1 IRE1a RRP44 WOX13
Verth-14 cont´d (left)
UGT2 BRI1 WRKY13 GEBGase1 RCAT1 RAB7 WRKY51 GTP5g64 RLT3 MAH1 PCD2
Theca-chr1_Mb6
UGT2 BRI1 WRKY13 (2x) GEBGase1 (2x) RCAT1 (3x) RAB7 WRKY51 GTP5g64 (2x) MAH1 PolII4 PCD2 (5x) shiKin1 SKIP6 CYP78A7
Loss of GNOM from OSG1 in Scrophulariaceae
Budal-17
iPPkin Fbox1g67 WRKY3 ALG13 XXT5 vSN12 glutelinB5 ILK1 emp24 O-fucT3 RubiscoMT PHO1h3 PHO1h3 PHO1h3 PHO1h3
Verth-2
WRKY3 SKP1-1A UbqCtH17 SAT22 (sos) ALG13 PIN5 (2x) XXT5 vSN12 glutelinB5 ILK1 emp24 O-fucT3 PHO1h3 PHO1h3 PHO1h3
Budal-4
lamin WRKY4 IBIS1 mito2g39 DUF1336 TM230 KIN10 CSC1H GNL1 vSN13 PPR5g15 DUF1997 At5g39 PHO1h9 SPX4 SPX4 cystgS1 MYB16 NAC92
Verth-9
lamin WRKY4 IBIS1 DUF1336 KIN10 CSC1H GNL1b vSN13 PPR5g15 DUF1997 At5g39 PHO1h9 SPX4 cystgS1

#### Suppl. Figure S22. Boraginales cladogram and OSGs

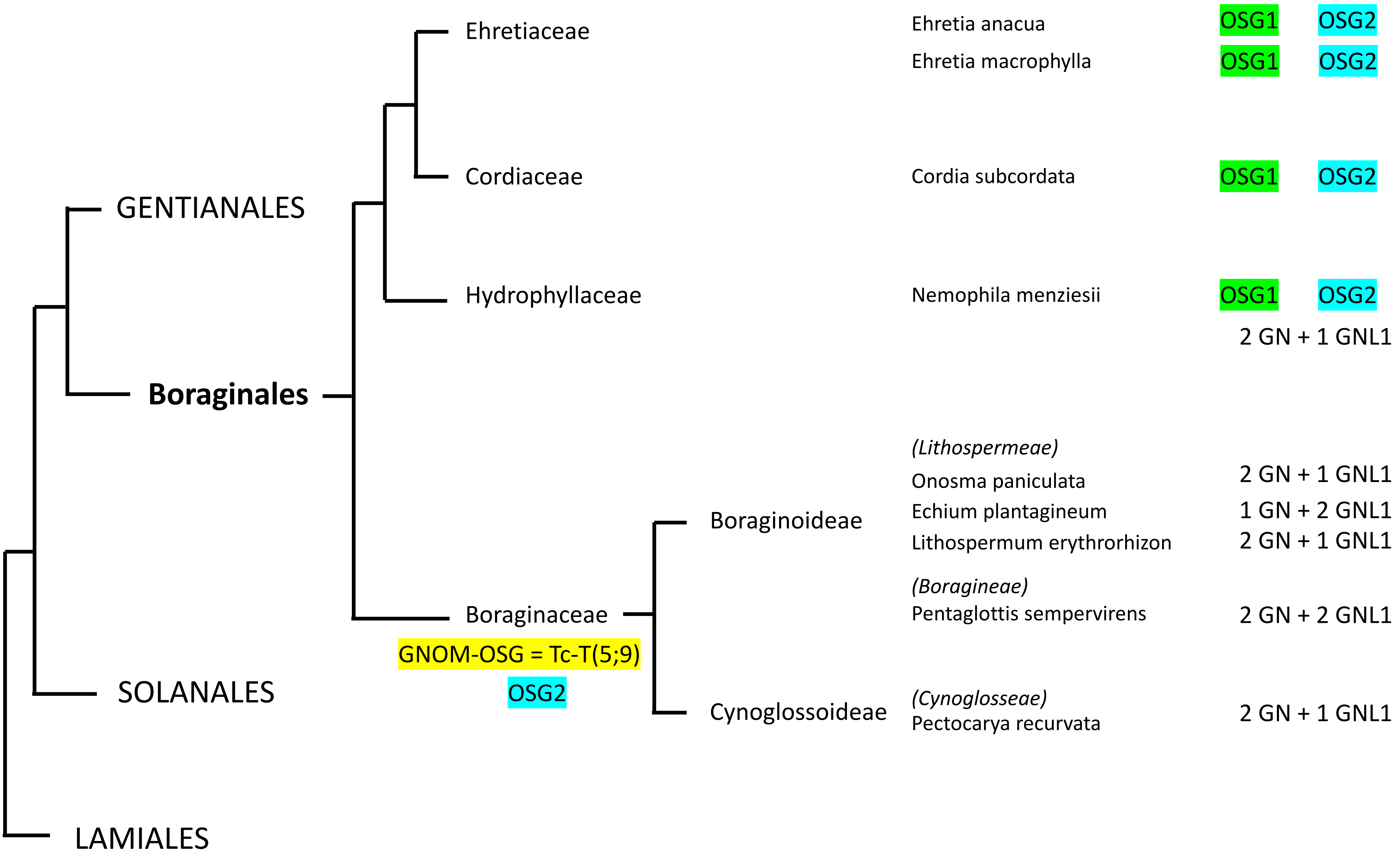

Cladogram based on Weigend et al. (2014); Zhang et al. (2020c)

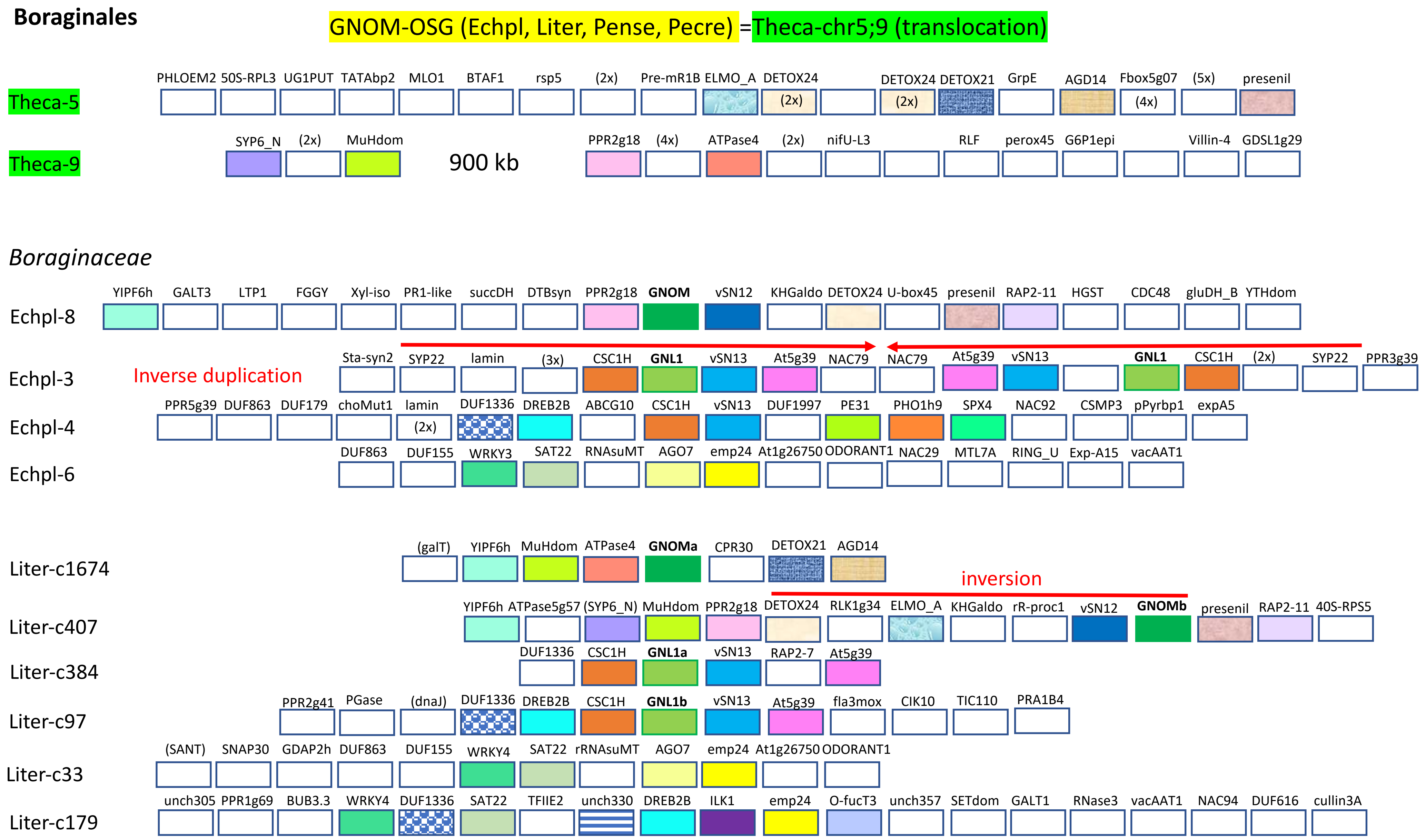
Boraginales
GNOM-OSG (Echpl, Liter, Pense, Pecre) =Theca-chr5;9 (translocation)
Theca-5
PHLOEM2 50S-RPL3 UG1PUT TATAbp2 MLO1 BTAF1 rsp5 (2x) Pre-mR1B ELMO_A DETOX24 (2x) DETOX24 (2x) DETOX21 GrpE AGD14 Fbox5g07 (4x) (5x) presenil
Theca-9
SYP6_N (2x) MuHdom 900 kb PPR2g18 (4x) ATPase4 (2x) nifU-L3 RLF perox45 G6P1epi Villin-4 GDSL1g29
Boraginaceae
Echpl-8
YIPF6h GALT3 LTP1 FGGY Xyl-iso PR1-like succDH DTBsyn PPR2g18 GNOM vSN12 KHGaldo DETOX24 U-box45 presenil RAP2-11 HGST CDC48 gluDH_B YTHdom
Echpl-3
Inverse duplication
Sta-syn2 SYP22 lamin (3x) CSC1H GNL1 vSN13 At5g39 NAC79 NAC79 At5g39 vSN13 GNL1 CSC1H (2x) SYP22 PPR3g39
Echpl-4
PPR5g39 DUF863 DUF179 choMut1 lamin (2x) DUF1336 DREB2B ABCG10 CSC1H vSN13 DUF1997 PE31 PHO1h9 SPX4 NAC92 CSMP3 pPyrbp1 expA5
Echpl-6
DUF863 DUF155 WRKY3 SAT22 RNAsuMT AGO7 emp24 At1g26750 ODORANT1 NAC29 MTL7A RING_U Exp-A15 vacAAT1
Liter-c1674
(galT) YIPF6h MuHdom ATPase4 GNOMa CPR30 DETOX21 AGD14
inversion
Liter-c407
YIPF6h ATPase5g57 (SYP6_N) MuHdom PPR2g18 DETOX24 RLK1g34 ELMO_A KHGaldo rR-proc1 vSN12 GNOMb presenil RAP2-11 40S-RPS5
Liter-c384
DUF1336 CSC1H GNL1a vSN13 RAP2-7 At5g39
Liter-c97
PPR2g41 PGase (dnaJ) DUF1336 DREB2B CSC1H GNL1b vSN13 At5g39 fla3mox CIK10 TIC110 PRA1B4
Liter-c33
(SANT) SNAP30 GDAP2h DUF863 DUF155 WRKY4 SAT22 rRNAsuMT AGO7 emp24 At1g26750 ODORANT1
Liter-c179
unch305 PPR1g69 BUB3.3 WRKY4 DUF1336 SAT22 TFIIE2 unch330 DREB2B ILK1 emp24 O-fucT3 unch357 SETdom GALT1 RNase3 vacAAT1 NAC94 DUF616 cullin3A

### Boraginales

#### *Boraginaceae*

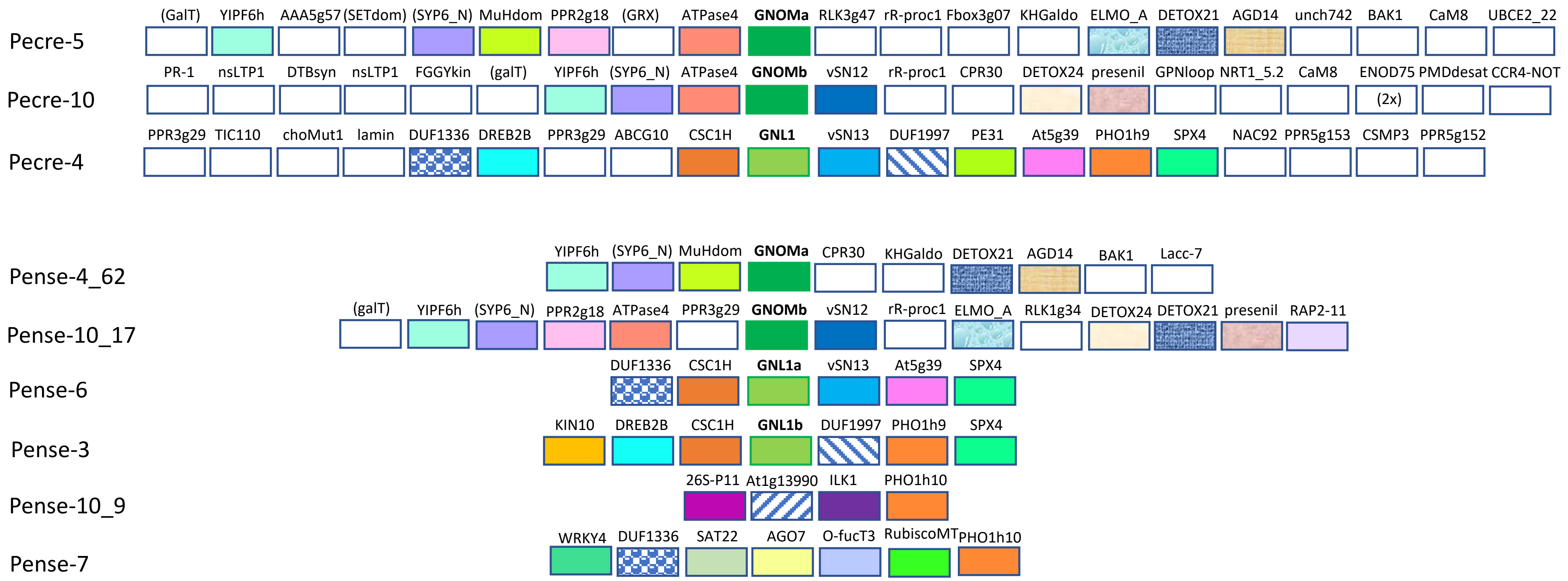

#### Boraginales

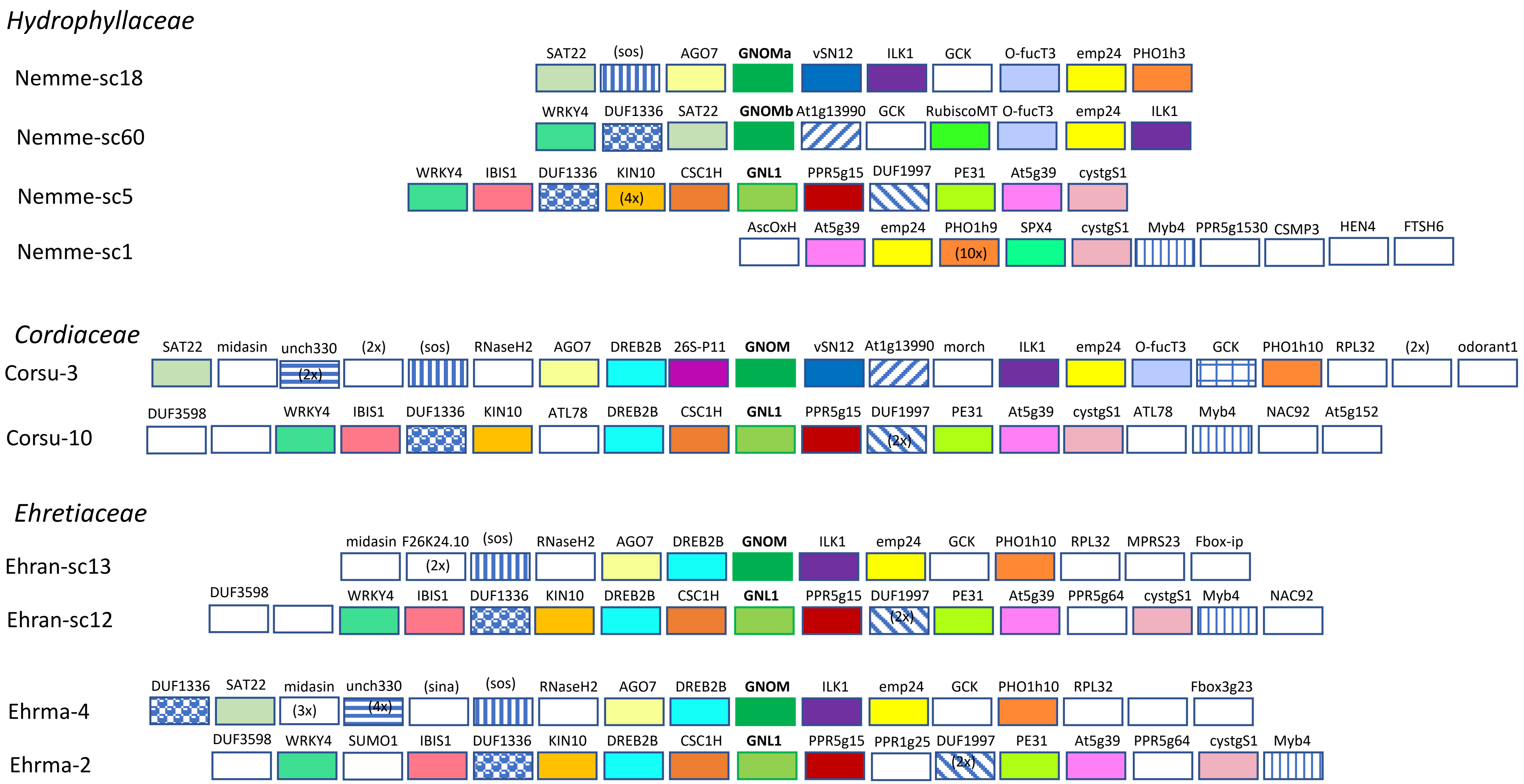

### Suppl. Figure S23. Gentianales cladogram and OSGs

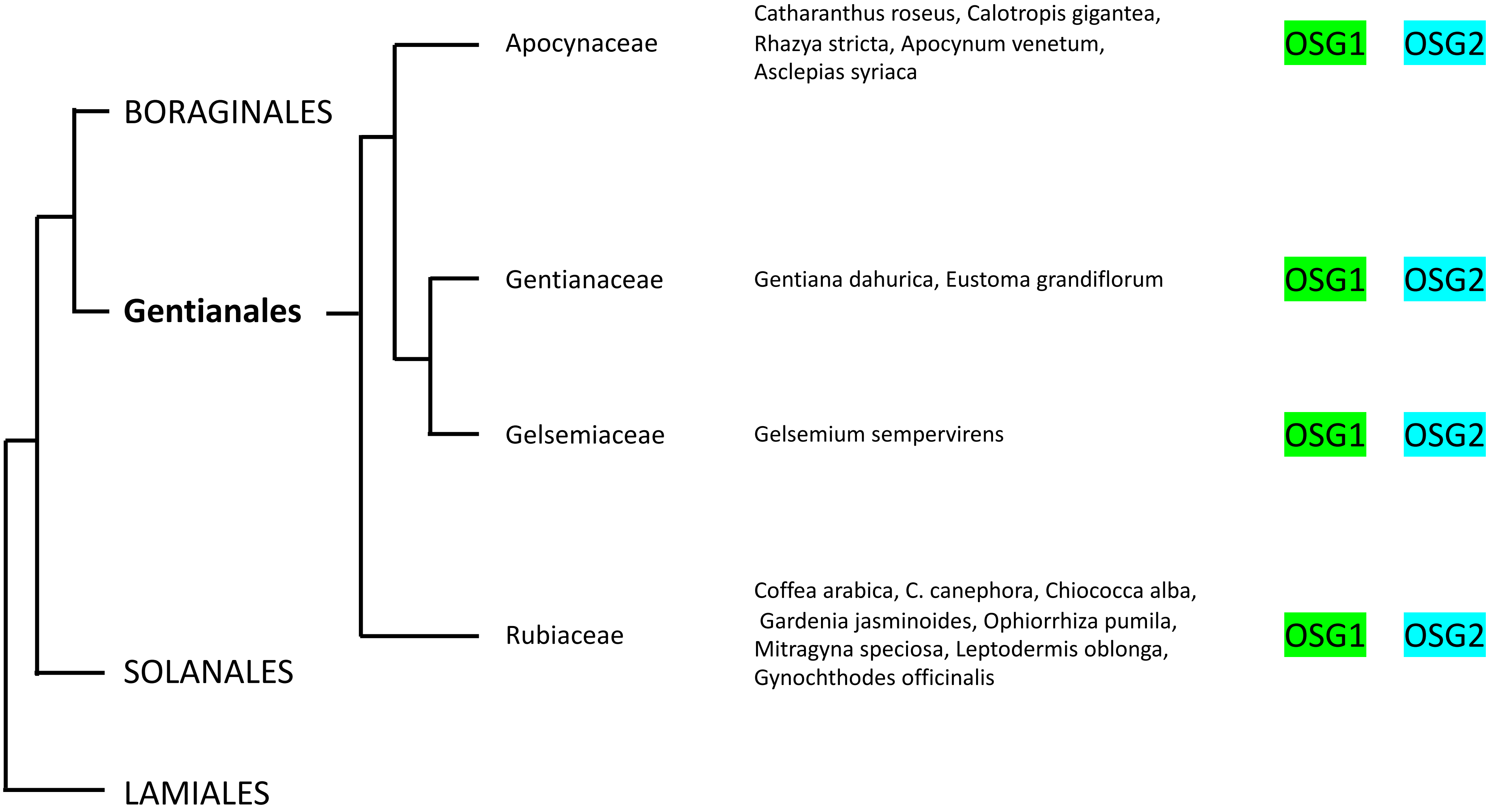

Cladogram based on Zhang et al. (2020c)

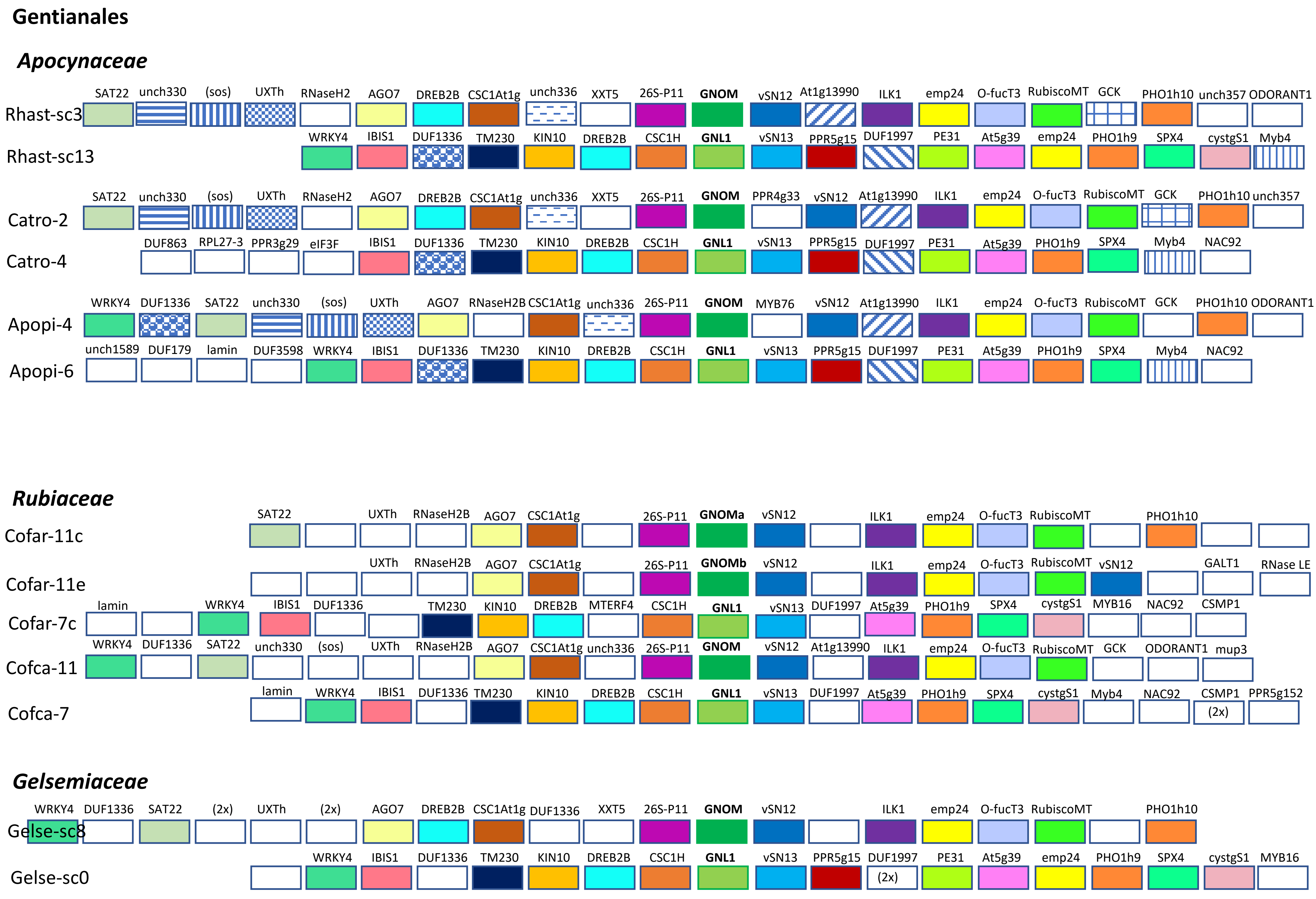
Gentianales
Apocynaceae
Rhast-sc3
SAT22 unch330 (sos) UXTh RNaseH2 AGO7 DREB2B CSC1At1g unch336 XXT5 26S-P11 GNOM vSN12 At1g13990 ILK1 emp24 O-fucT3 RubiscoMT GCK PHO1h10 unch357 ODORANT1
Rhast-sc13
WRKY4 IBIS1 DUF1336 TM230 KIN10 DREB2B CSC1H GNL1 vSN13 PPR5g15 DUF1997 PE31 At5g39 emp24 PHO1h9 SPX4 cystgS1 Myb4
Catro-2
SAT22 unch330 (sos) UXTh RNaseH2 AGO7 DREB2B CSC1At1g unch336 XXT5 26S-P11 GNOM PPR4g33 vSN12 At1g13990 ILK1 emp24 O-fucT3 RubiscoMT GCK PHO1h10 unch357
Catro-4
DUF863 RPL27-3 PPR3g29 eIF3F IBIS1 DUF1336 TM230 KIN10 DREB2B CSC1H GNL1 vSN13 PPR5g15 DUF1997 PE31 At5g39 PHO1h9 SPX4 Myb4 NAC92
Apopi-4
WRKY4 DUF1336 SAT22 unch330 (sos) UXTh AGO7 RNaseH2B CSC1At1g unch336 26S-P11 GNOM MYB76 vSN12 At1g13990 ILK1 emp24 O-fucT3 RubiscoMT GCK PHO1h10 ODORANT1
Apopi-6
unch1589 DUF179 lamin DUF3598 WRKY4 IBIS1 DUF1336 TM230 KIN10 DREB2B CSC1H GNL1 vSN13 PPR5g15 DUF1997 PE31 At5g39 PHO1h9 SPX4 Myb4 NAC92
Rubiaceae
Cofar-11c
SAT22 UXTh RNaseH2B AGO7 CSC1At1g 26S-P11 GNOMa vSN12 ILK1 emp24 O-fucT3 RubiscoMT PHO1h10
Cofar-11e
UXTh RNaseH2B AGO7 CSC1At1g 26S-P11 GNOMb vSN12 ILK1 emp24 O-fucT3 RubiscoMT vSN12 GALT1 RNase LE
Cofar-7c
lamin WRKY4 IBIS1 DUF1336 TM230 KIN10 DREB2B MTERF4 CSC1H GNL1 vSN13 DUF1997 At5g39 PHO1h9 SPX4 cystgS1 MYB16 NAC92 CSMP1
Cofca-11
WRKY4 DUF1336 SAT22 unch330 (sos) UXTh RNaseH2B AGO7 CSC1At1g unch336 26S-P11 GNOM vSN12 At1g13990 ILK1 emp24 O-fucT3 RubiscoMT GCK ODORANT1 mup3
Cofca-7
lamin WRKY4 IBIS1 DUF1336 TM230 KIN10 DREB2B CSC1H GNL1 vSN13 DUF1997 At5g39 PHO1h9 SPX4 cystgS1 Myb4 NAC92 CSMP1 PPR5g152
(2x)
Gelsemiaceae
Gelse-sc8
WRKY4 DUF1336 SAT22 (2x) UXTh (2x) AGO7 DREB2B CSC1At1g DUF1336 XXT5 26S-P11 GNOM vSN12 ILK1 emp24 O-fucT3 RubiscoMT PHO1h10
Gelse-sc0
WRKY4 IBIS1 DUF1336 TM230 KIN10 DREB2B CSC1H GNL1 vSN13 PPR5g15 DUF1997 PE31 At5g39 emp24 PHO1h9 SPX4 cystgS1 MYB16
(2x)

*Gentianaceae*

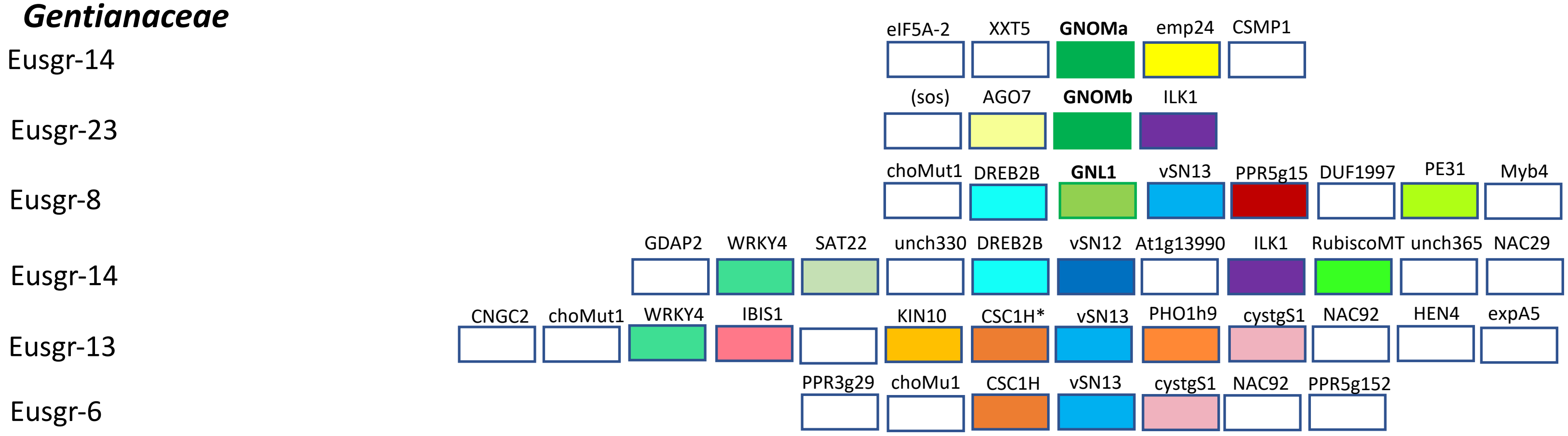

**Suppl. Figure S24. Solanales cladogram and OSGs**

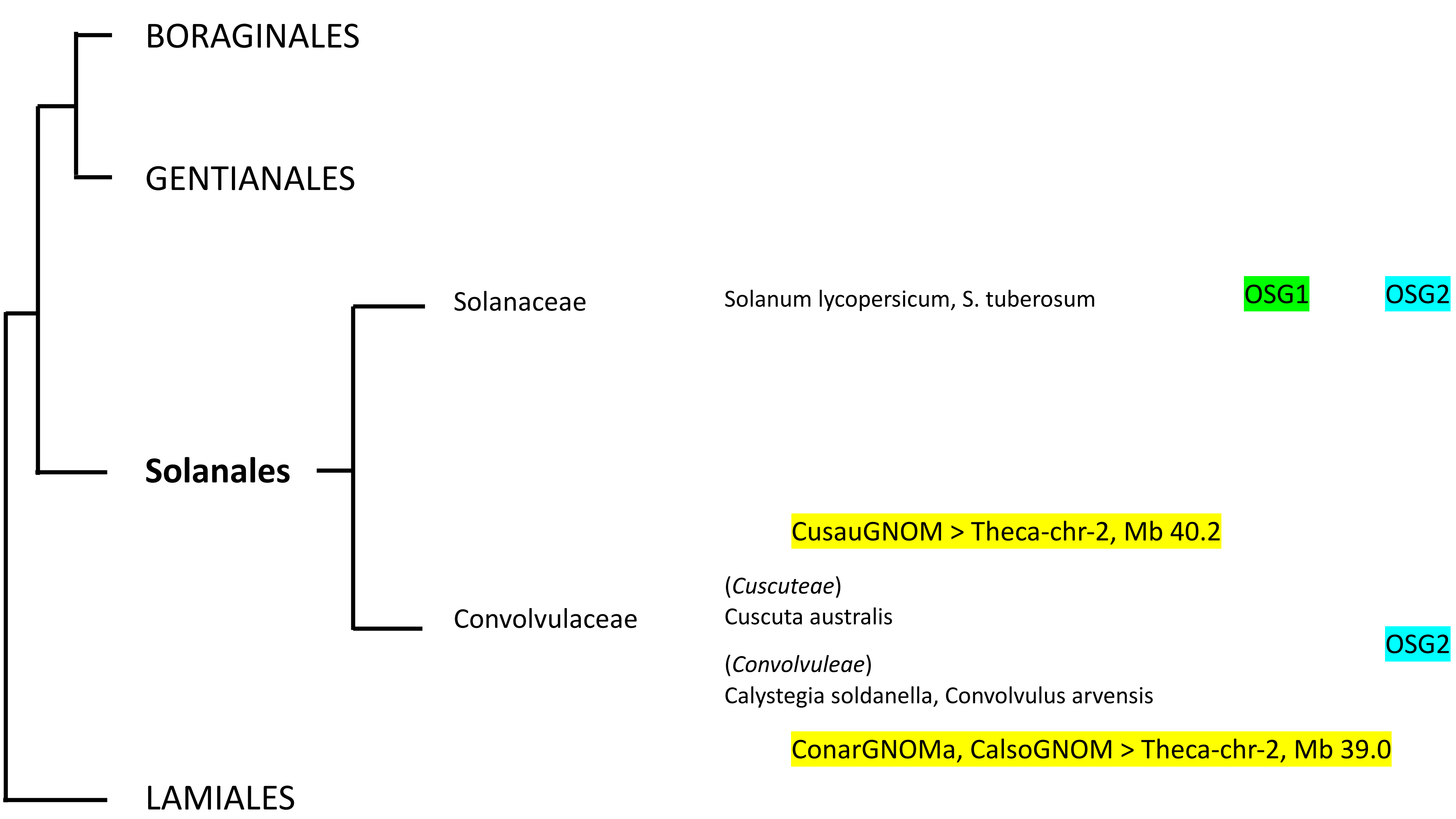

Cladogram based on Zhang et al. (2020c); Zuntini et al. (2024)

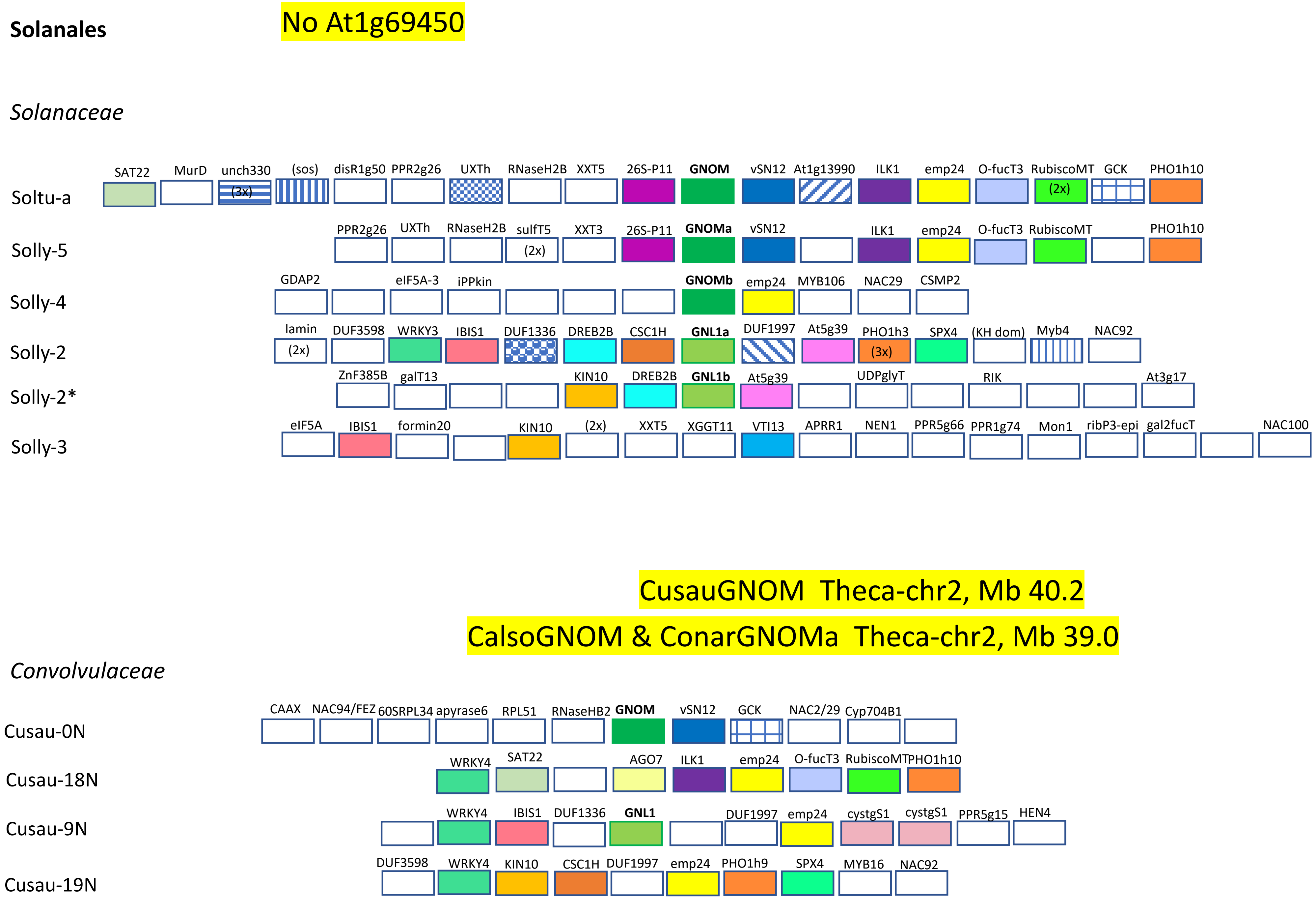

Solanales
No At1g69450
Solanaceae
Soltu-a
SAT22 MurD unch330 (3x) (sos) disR1g50 PPR2g26 UXTh RNaseH2B XXT5 26S-P11 GNOM vSN12 At1g13990 ILK1 emp24 O-fucT3 RubiscoMT (2x) GCK PHO1h10
Solly-5
PPR2g26 UXTh RNaseH2B sulfT5 (2x) XXT3 26S-P11 GNOMa vSN12 ILK1 emp24 O-fucT3 RubiscoMT PHO1h10
Solly-4
GDAP2 eIF5A-3 iPPkin GNOMb emp24 MYB106 NAC29 CSMP2
Solly-2
lamin (2x) DUF3598 WRKY3 IBIS1 DUF1336 DREB2B CSC1H GNL1a DUF1997 At5g39 PHO1h3 (3x) SPX4 (KH dom) Myb4 NAC92
Solly-2*
ZnF385B galT13 KIN10 DREB2B GNL1b At5g39 UDPglyT RIK At3g17
Solly-3
eIF5A IBIS1 formin20 KIN10 (2x) XXT5 XGGT11 VTI13 APRR1 NEN1 PPR5g66 PPR1g74 Mon1 ribP3-epi gal2fucT NAC100
CusauGNOM Theca-chr2, Mb 40.2
CalsoGNOM & ConarGNOMa Theca-chr2, Mb 39.0
Convolvulaceae
Cusau-0N
CAAX NAC94/FEZ 60SRPL34 apyrase6 RPL51 RNaseHB2 GNOM vSN12 GCK NAC2/29 Cyp704B1
Cusau-18N
WRKY4 SAT22 AGO7 ILK1 emp24 O-fucT3 RubiscoMT PHO1h10
Cusau-9N
WRKY4 IBIS1 DUF1336 GNL1 DUF1997 emp24 cystgS1 cystgS1 PPR5g15 HEN4
Cusau-19N
DUF3598 WRKY4 KIN10 CSC1H DUF1997 emp24 PHO1h9 SPX4 MYB16 NAC92

#### Suppl. Figure S25. Aquifoliales cladogram and OSGs

Ilex is the only genus of Aquifoliacae

No GNL1

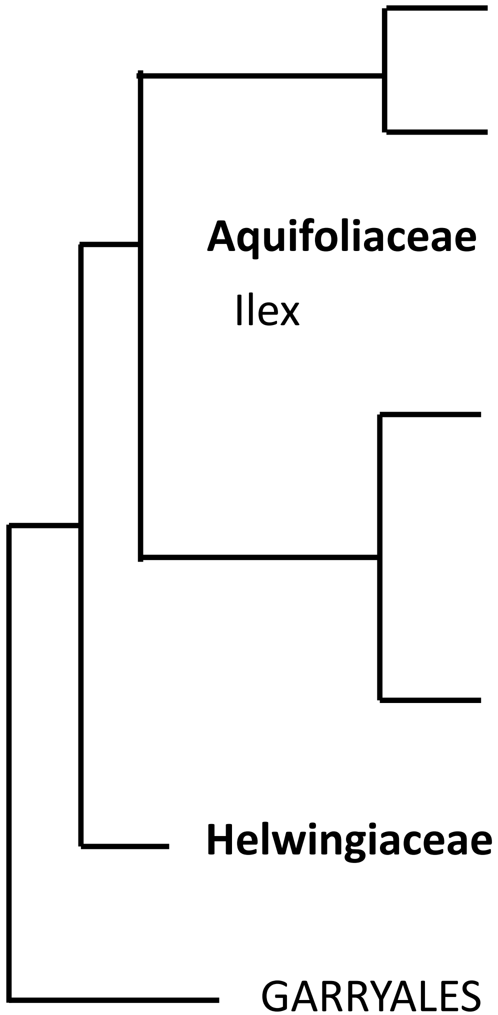

| | | |
|---|---|---|
| Ilex asprella (*clade A*) | GNOMa Theca-chr1 | GNOMb Theca-chr5 (300kb away) |
| Ilex polyneura (*clade B*) | GNOMa Theca-chr1 | GNOMb Theca-chr5 (OSG14) |
| Ilex latifolia (*clade D*) | GNOMa Theca-chr1 | GNOMb Theca-chr5 (300kb away) |
| Ilex aquifolium (*clade D*) | GNOMa Theca-chr1 | GNOMb Theca-chr5 (OSG14) |
| Ilex paraguariensis (*clade E*) | GNOMa Theca-chr1<br>GNOMc (33kb ctg) | GNOMb Theca-chr5 (OSG14)<br>GNOMd Theca-chr5 (300kb away) |
| Helwingia omeiensis | GNOMa, GNOMb, GNOMc all Theca-chr5 (OSG14) | |

Cladogram based on Stull et al (2020); Xu et al. (2022)

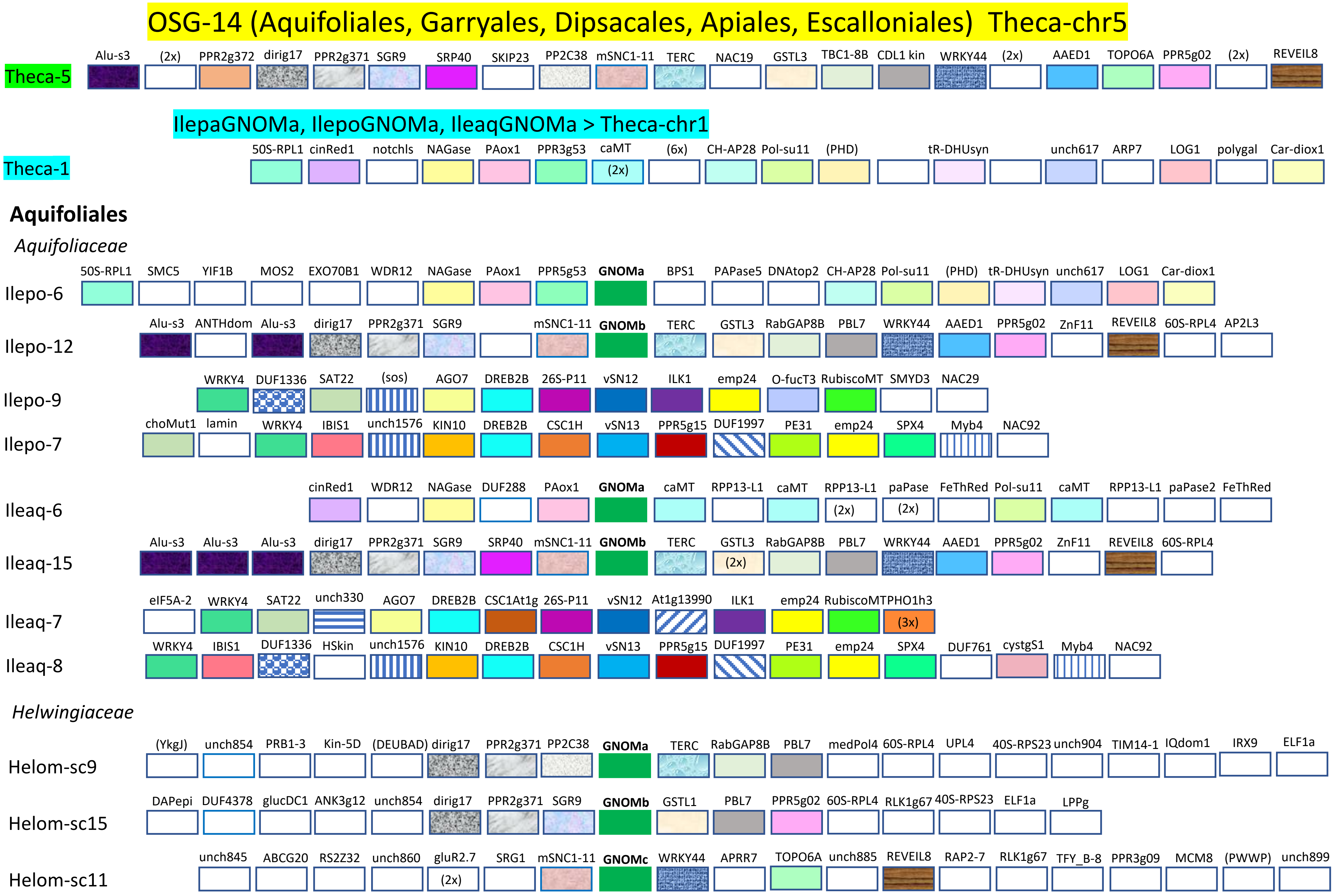

OSG-14 (Aquifoliales, Garryales, Dipsacales, Apiales, Escalloniales) Theca-chr5
Theca-5
Alu-s3 (2x) PPR2g372 dirig17 PPR2g371 SGR9 SRP40 SKIP23 PP2C38 mSNC1-11 TERC NAC19 GSTL3 TBC1-8B CDL1 kin WRKY44 (2x) AAED1 TOPO6A PPR5g02 (2x) REVEIL8
IlepaGNOMa, IlepoGNOMa, IleaqGNOMa > Theca-chr1
Theca-1
50S-RPL1 cinRed1 notchls NAGase PAox1 PPR3g53 caMT (2x) (6x) CH-AP28 Pol-su11 (PHD) tR-DHUsyn unch617 ARP7 LOG1 polygal Car-diox1
Aquifoliales
Aquifoliaceae
Ilepo-6
50S-RPL1 SMC5 YIF1B MOS2 EXO70B1 WDR12 NAGase PAox1 PPR5g53 GNOMa BPS1 PAPase5 DNAtop2 CH-AP28 Pol-su11 (PHD) tR-DHUsyn unch617 LOG1 Car-diox1
Ilepo-12
Alu-s3 ANTHdom Alu-s3 dirig17 PPR2g371 SGR9 mSNC1-11 GNOMb TERC GSTL3 RabGAP8B PBL7 WRKY44 AAED1 PPR5g02 ZnF11 REVEIL8 60S-RPL4 AP2L3
Ilepo-9
WRKY4 DUF1336 SAT22 (sos) AGO7 DREB2B 26S-P11 vSN12 ILK1 emp24 O-fucT3 RubiscoMT SMYD3 NAC29
Ilepo-7
choMut1 lamin WRKY4 IBIS1 unch1576 KIN10 DREB2B CSC1H vSN13 PPR5g15 DUF1997 PE31 emp24 SPX4 Myb4 NAC92
Ileaq-6
cinRed1 WDR12 NAGase DUF288 PAox1 GNOMa caMT RPP13-L1 caMT RPP13-L1 (2x) paPase (2x) FeThRed Pol-su11 caMT RPP13-L1 paPase2 FeThRed
Ileaq-15
Alu-s3 Alu-s3 Alu-s3 dirig17 PPR2g371 SGR9 SRP40 mSNC1-11 GNOMb TERC GSTL3 (2x) RabGAP8B PBL7 WRKY44 AAED1 PPR5g02 ZnF11 REVEIL8 60S-RPL4
Ileaq-7
eIF5A-2 WRKY4 SAT22 unch330 AGO7 DREB2B CSC1At1g 26S-P11 vSN12 At1g13990 ILK1 emp24 RubiscoMT PHO1h3 (3x)
Ileaq-8
WRKY4 IBIS1 DUF1336 HSkin unch1576 KIN10 DREB2B CSC1H vSN13 PPR5g15 DUF1997 PE31 emp24 SPX4 DUF761 cystgS1 Myb4 NAC92
Helwingiaceae
Helom-sc9
(YkgJ) unch854 PRB1-3 Kin-5D (DEUBAD) dirig17 PPR2g371 PP2C38 GNOMa TERC RabGAP8B PBL7 medPol4 60S-RPL4 UPL4 40S-RPS23 unch904 TIM14-1 IQdom1 IRX9 ELF1a
Helom-sc15
DAPepi DUF4378 glucDC1 ANK3g12 unch854 dirig17 PPR2g371 SGR9 GNOMb GSTL1 PBL7 PPR5g02 60S-RPL4 RLK1g67 40S-RPS23 ELF1a LPPg
Helom-sc11
unch845 ABCG20 RS2Z32 unch860 gluR2.7 (2x) SRG1 mSNC1-11 GNOMc WRKY44 APRR7 TOPO6A unch885 REVEIL8 RAP2-7 RLK1g67 TFY_B-8 PPR3g09 MCM8 (PWWP) unch899

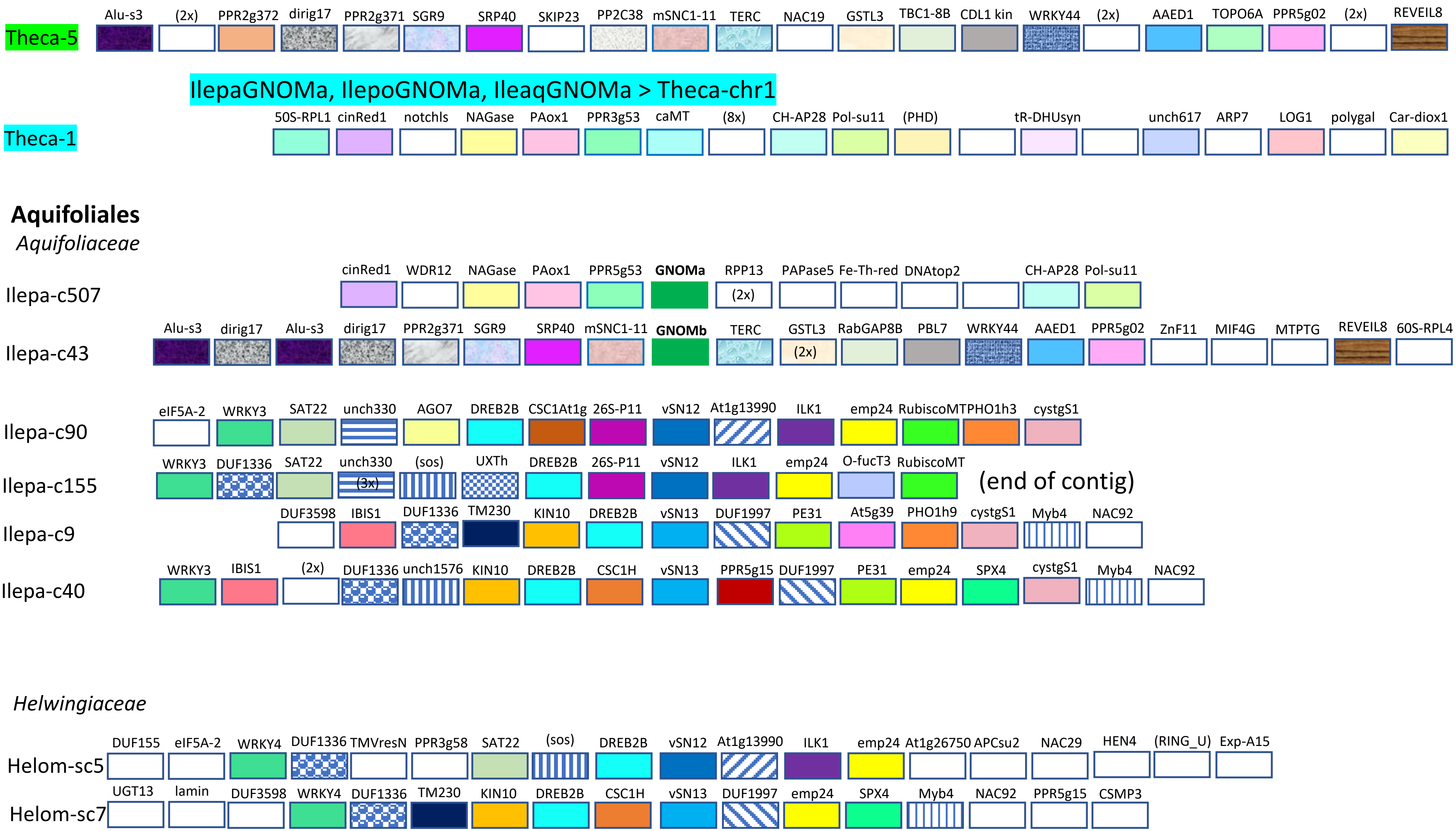

Theca-5
Alu-s3 (2x) PPR2g372 dirig17 PPR2g371 SGR9 SRP40 SKIP23 PP2C38 mSNC1-11 TERC NAC19 GSTL3 TBC1-8B CDL1 kin WRKY44 (2x) AAED1 TOPO6A PPR5g02 (2x) REVEIL8
IlepaGNOMa, IlepoGNOMa, IleaqGNOMa > Theca-chr1
Theca-1
50S-RPL1 cinRed1 notchls NAGase PAox1 PPR3g53 caMT (8x) CH-AP28 Pol-su11 (PHD) tR-DHUsyn unch617 ARP7 LOG1 polygal Car-diox1
Aquifoliales
Aquifoliaceae
Ilepa-c507
cinRed1 WDR12 NAGase PAox1 PPR5g53 GNOMa RPP13 (2x) PAPase5 Fe-Th-red DNAtop2 CH-AP28 Pol-su11
Ilepa-c43
Alu-s3 dirig17 Alu-s3 dirig17 PPR2g371 SGR9 SRP40 mSNC1-11 GNOMb TERC GSTL3 (2x) RabGAP8B PBL7 WRKY44 AAED1 PPR5g02 ZnF11 MIF4G MTPTG REVEIL8 60S-RPL4
Ilepa-c90
eIF5A-2 WRKY3 SAT22 unch330 AGO7 DREB2B CSC1At1g 26S-P11 vSN12 At1g13990 ILK1 emp24 RubiscoMT PHO1h3 cystgS1
Ilepa-c155
WRKY3 DUF1336 SAT22 unch330 (3x) (sos) UXTh DREB2B 26S-P11 vSN12 ILK1 emp24 O-fucT3 RubiscoMT
(end of contig)
Ilepa-c9
DUF3598 IBIS1 DUF1336 TM230 KIN10 DREB2B vSN13 DUF1997 PE31 At5g39 PHO1h9 cystgS1 Myb4 NAC92
Ilepa-c40
WRKY3 IBIS1 (2x) DUF1336 unch1576 KIN10 DREB2B CSC1H vSN13 PPR5g15 DUF1997 PE31 emp24 SPX4 cystgS1 Myb4 NAC92
Helwingiaceae
Helom-sc5
DUF155 eIF5A-2 WRKY4 DUF1336 TMVresN PPR3g58 SAT22 (sos) DREB2B vSN12 At1g13990 ILK1 emp24 At1g26750 APCsu2 NAC29 HEN4 (RING_U) Exp-A15
Helom-sc7
UGT13 lamin DUF3598 WRKY4 DUF1336 TM230 KIN10 DREB2B CSC1H vSN13 DUF1997 emp24 SPX4 Myb4 NAC92 PPR5g15 CSMP3

#### Suppl. Figurre S26. Garryales OSGs

OSG-14 (Aquifoliales, Garryales, Dipsacales, Apiales, Escalloniales) Theca-chr5

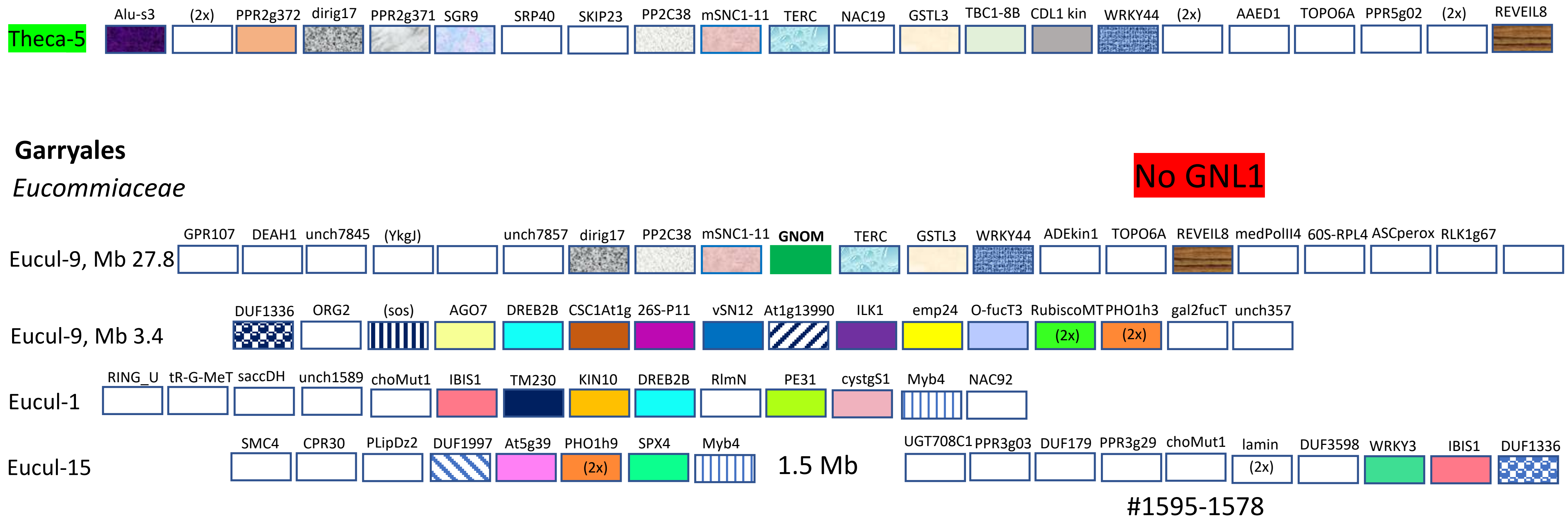

#### 2 families

- Garryaceae
- Eucommiaceae (including *Eucommia ulmoides*)

**Suppl. Figure S27. Apiales cladogram and OSGs**

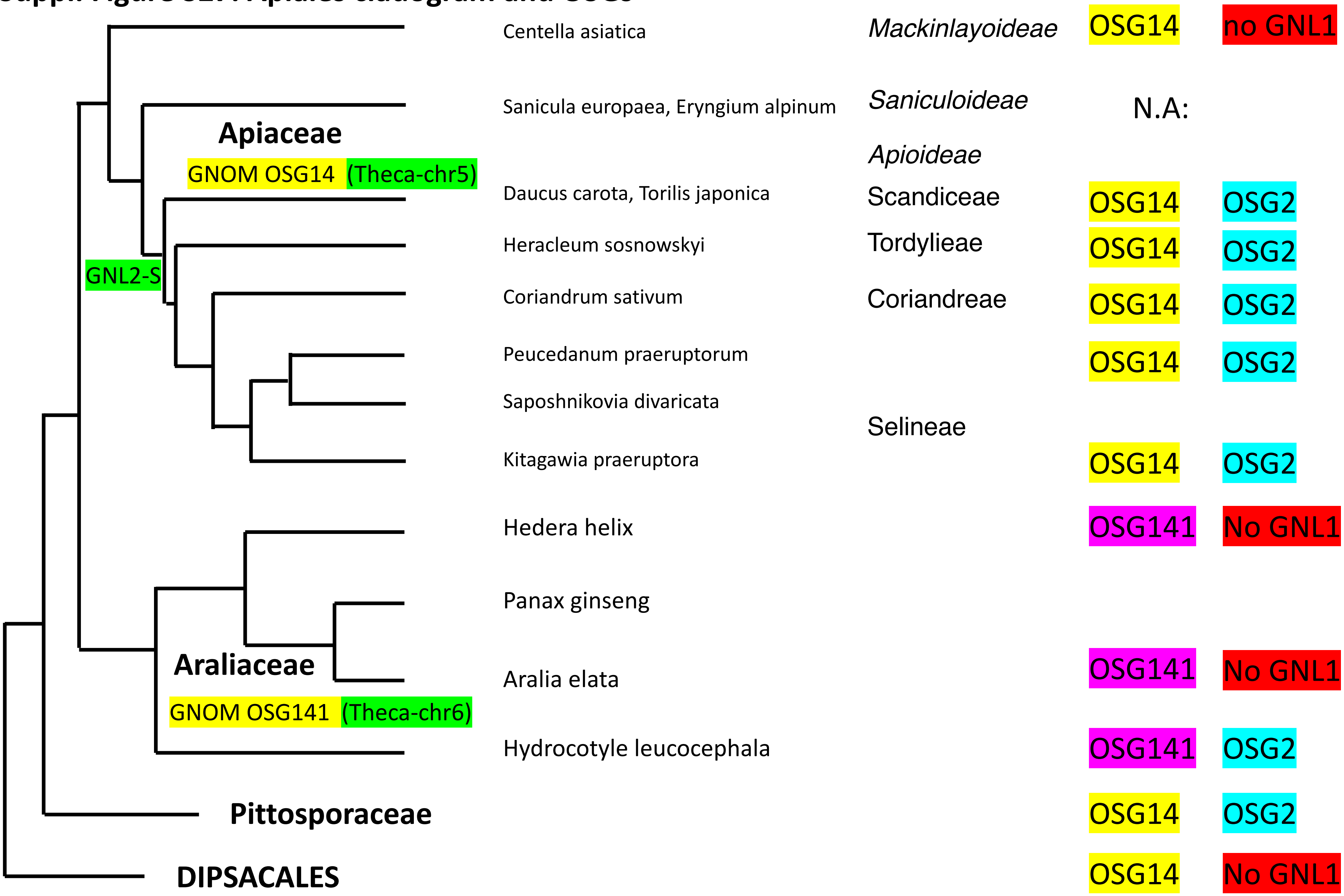

Cladogram based on Clarkson et al. (2021.)

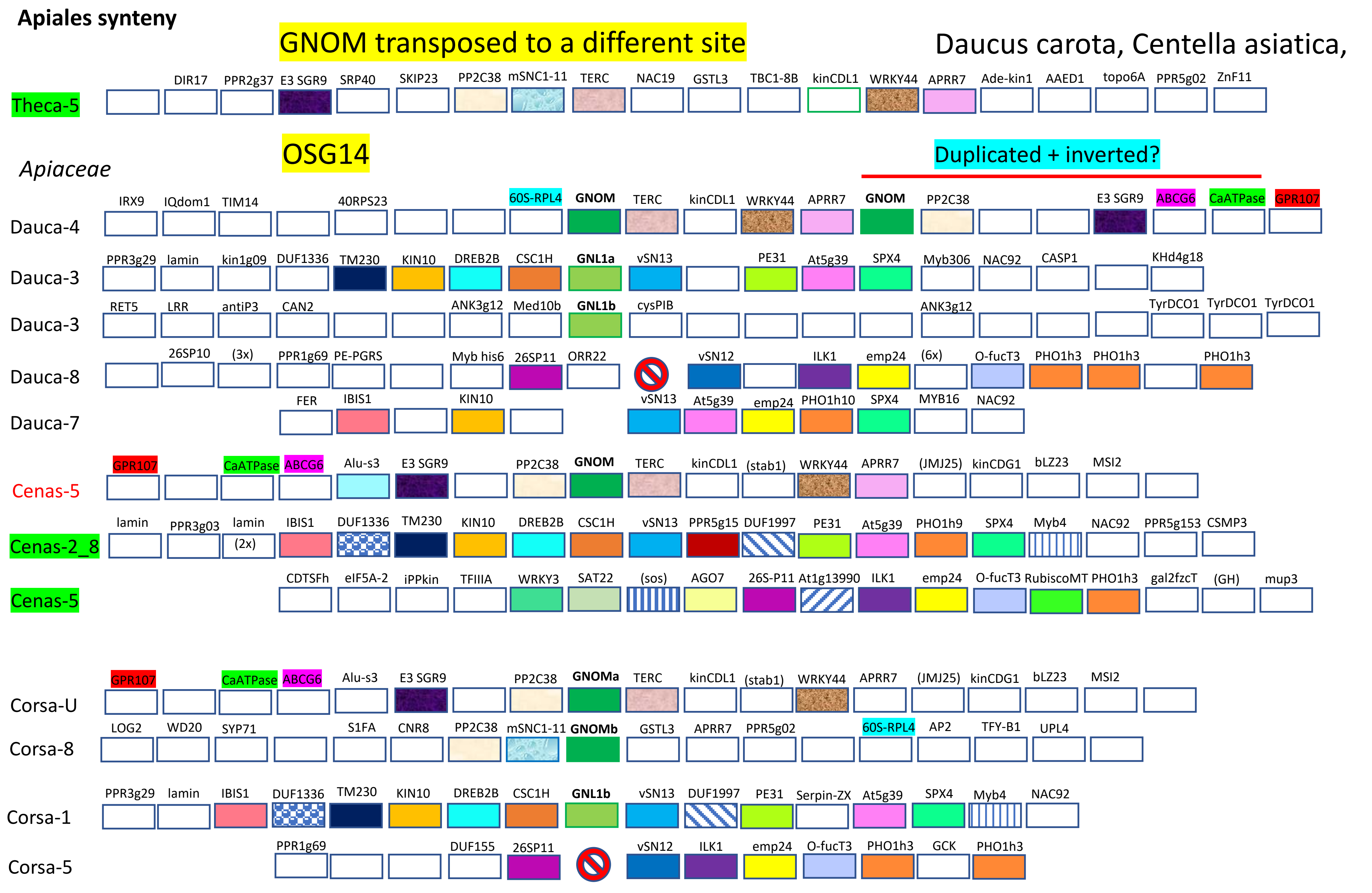
Apiales synteny
GNOM transposed to a different site
Daucus carota, Centella asiatica,
Theca-5
DIR17 PPR2g37 E3 SGR9 SRP40 SKIP23 PP2C38 mSNC1-11 TERC NAC19 GSTL3 TBC1-8B kinCDL1 WRKY44 APRR7 Ade-kin1 AAED1 topo6A PPR5g02 ZnF11
Apiaceae
OSG14
Duplicated + inverted?
Dauca-4
IRX9 IQdom1 TIM14 40RPS23 60S-RPL4 GNOM TERC kinCDL1 WRKY44 APRR7 GNOM PP2C38 E3 SGR9 ABCG6 CaATPase GPR107
Dauca-3
PPR3g29 lamin kin1g09 DUF1336 TM230 KIN10 DREB2B CSC1H GNL1a vSN13 PE31 At5g39 SPX4 Myb306 NAC92 CASP1 KHd4g18
Dauca-3
RET5 LRR antiP3 CAN2 ANK3g12 Med10b GNL1b cysPIB ANK3g12 TyrDCO1 TyrDCO1 TyrDCO1
Dauca-8
26SP10 (3x) PPR1g69 PE-PGRS Myb his6 26SP11 ORR22 vSN12 ILK1 emp24 (6x) O-fucT3 PHO1h3 PHO1h3 PHO1h3
Dauca-7
FER IBIS1 KIN10 vSN13 At5g39 emp24 PHO1h10 SPX4 MYB16 NAC92
Cenas-5
GPR107 CaATPase ABCG6 Alu-s3 E3 SGR9 PP2C38 GNOM TERC kinCDL1 (stab1) WRKY44 APRR7 (JMJ25) kinCDG1 bLZ23 MSI2
Cenas-2_8
lamin PPR3g03 lamin (2x) IBIS1 DUF1336 TM230 KIN10 DREB2B CSC1H vSN13 PPR5g15 DUF1997 PE31 At5g39 PHO1h9 SPX4 Myb4 NAC92 PPR5g153 CSMP3
Cenas-5
CDTSFh eIF5A-2 iPPkin TFIIIA WRKY3 SAT22 (sos) AGO7 26S-P11 At1g13990 ILK1 emp24 O-fucT3 RubiscoMT PHO1h3 gal2fzcT (GH) mup3
Corsa-U
GPR107 CaATPase ABCG6 Alu-s3 E3 SGR9 PP2C38 GNOMa TERC kinCDL1 (stab1) WRKY44 APRR7 (JMJ25) kinCDG1 bLZ23 MSI2
Corsa-8
LOG2 WD20 SYP71 S1FA CNR8 PP2C38 mSNC1-11 GNOMb GSTL3 APRR7 PPR5g02 60S-RPL4 AP2 TFY-B1 UPL4
Corsa-1
PPR3g29 lamin IBIS1 DUF1336 TM230 KIN10 DREB2B CSC1H GNL1b vSN13 DUF1997 PE31 Serpin-ZX At5g39 SPX4 Myb4 NAC92
Corsa-5
PPR1g69 DUF155 26SP11 vSN12 ILK1 emp24 O-fucT3 PHO1h3 GCK PHO1h3

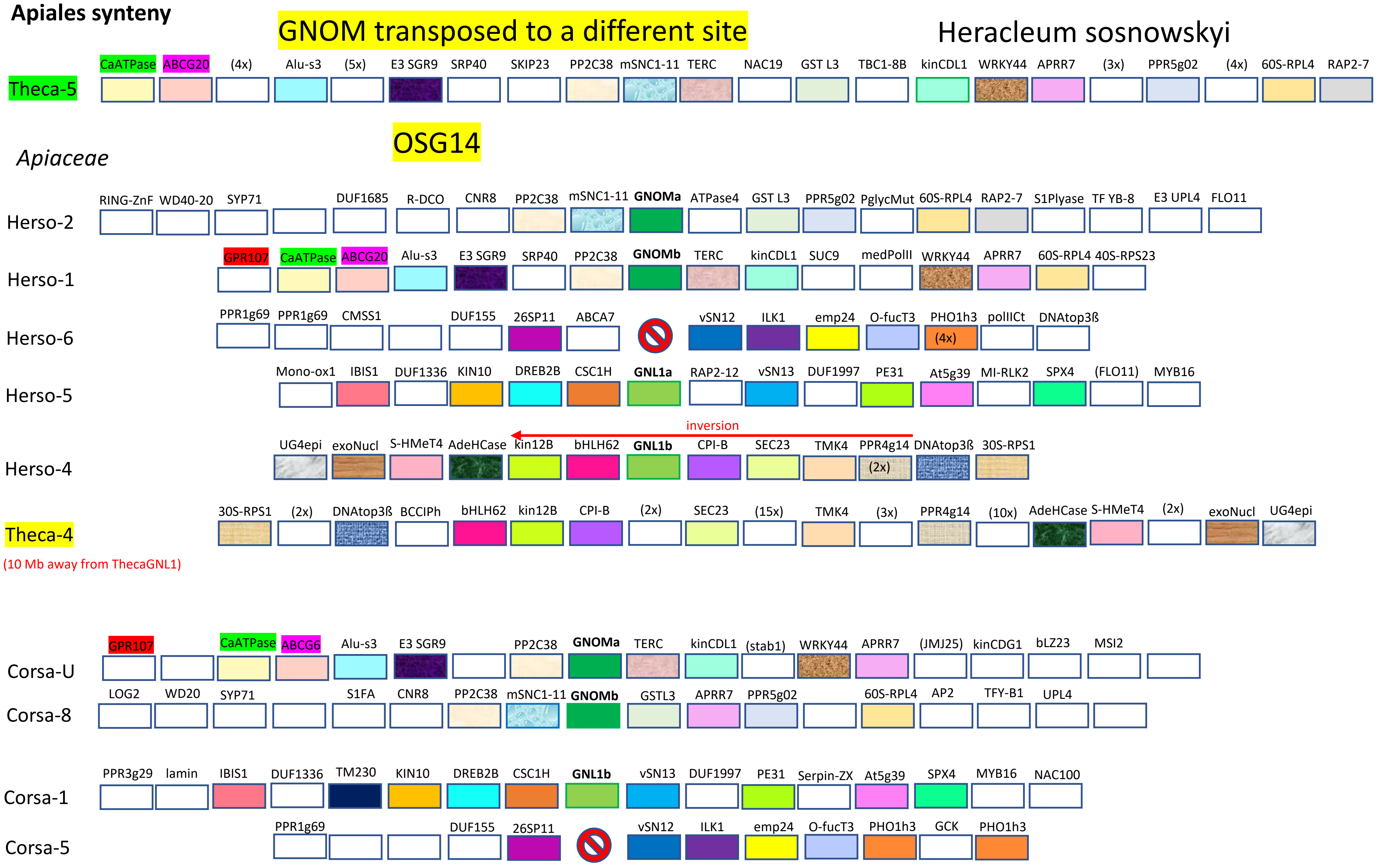
Apiales synteny
GNOM transposed to a different site
Heracleum sosnowskyi
Theca-5
CaATPase ABCG20 (4x) Alu-s3 (5x) E3 SGR9 SRP40 SKIP23 PP2C38 mSNC1-11 TERC NAC19 GST L3 TBC1-8B kinCDL1 WRKY44 APRR7 (3x) PPR5g02 (4x) 60S-RPL4 RAP2-7
OSG14
Apiaceae
Herso-2
RING-ZnF WD40-20 SYP71 DUF1685 R-DCO CNR8 PP2C38 mSNC1-11 GNOMa ATPase4 GST L3 PPR5g02 PglycMut 60S-RPL4 RAP2-7 S1Plyase TF YB-8 E3 UPL4 FLO11
Herso-1
GPR107 CaATPase ABCG20 Alu-s3 E3 SGR9 SRP40 PP2C38 GNOMb TERC kinCDL1 SUC9 medPolII WRKY44 APRR7 60S-RPL4 40S-RPS23
Herso-6
PPR1g69 PPR1g69 CMSS1 DUF155 26SP11 ABCA7 vSN12 ILK1 emp24 O-fucT3 PHO1h3 (4x) polICt DNAtop3ß
Herso-5
Mono-ox1 IBIS1 DUF1336 KIN10 DREB2B CSC1H GNL1a RAP2-12 vSN13 DUF1997 PE31 At5g39 MI-RLK2 SPX4 (FLO11) MYB16
inversion
Herso-4
UG4epi exoNucl S-HMeT4 AdeHCase kin12B bHLH62 GNL1b CPI-B SEC23 TMK4 PPR4g14 (2x) DNAtop3ß 30S-RPS1
Theca-4
30S-RPS1 (2x) DNAtop3ß BCCIPh bHLH62 kin12B CPI-B (2x) SEC23 (15x) TMK4 (3x) PPR4g14 (10x) AdeHCase S-HMeT4 (2x) exoNucl UG4epi
(10 Mb away from ThecaGNL1)
Corsa-U
GPR107 CaATPase ABCG6 Alu-s3 E3 SGR9 PP2C38 GNOMa TERC kinCDL1 (stab1) WRKY44 APRR7 (JMJ25) kinCDG1 bLZ23 MSI2
Corsa-8
LOG2 WD20 SYP71 S1FA CNR8 PP2C38 mSNC1-11 GNOMb GSTL3 APRR7 PPR5g02 60S-RPL4 AP2 TFY-B1 UPL4
Corsa-1
PPR3g29 lamin IBIS1 DUF1336 TM230 KIN10 DREB2B CSC1H GNL1b vSN13 DUF1997 PE31 Serpin-ZX At5g39 SPX4 MYB16 NAC100
Corsa-5
PPR1g69 DUF155 26SP11 vSN12 ILK1 emp24 O-fucT3 PHO1h3 GCK PHO1h3

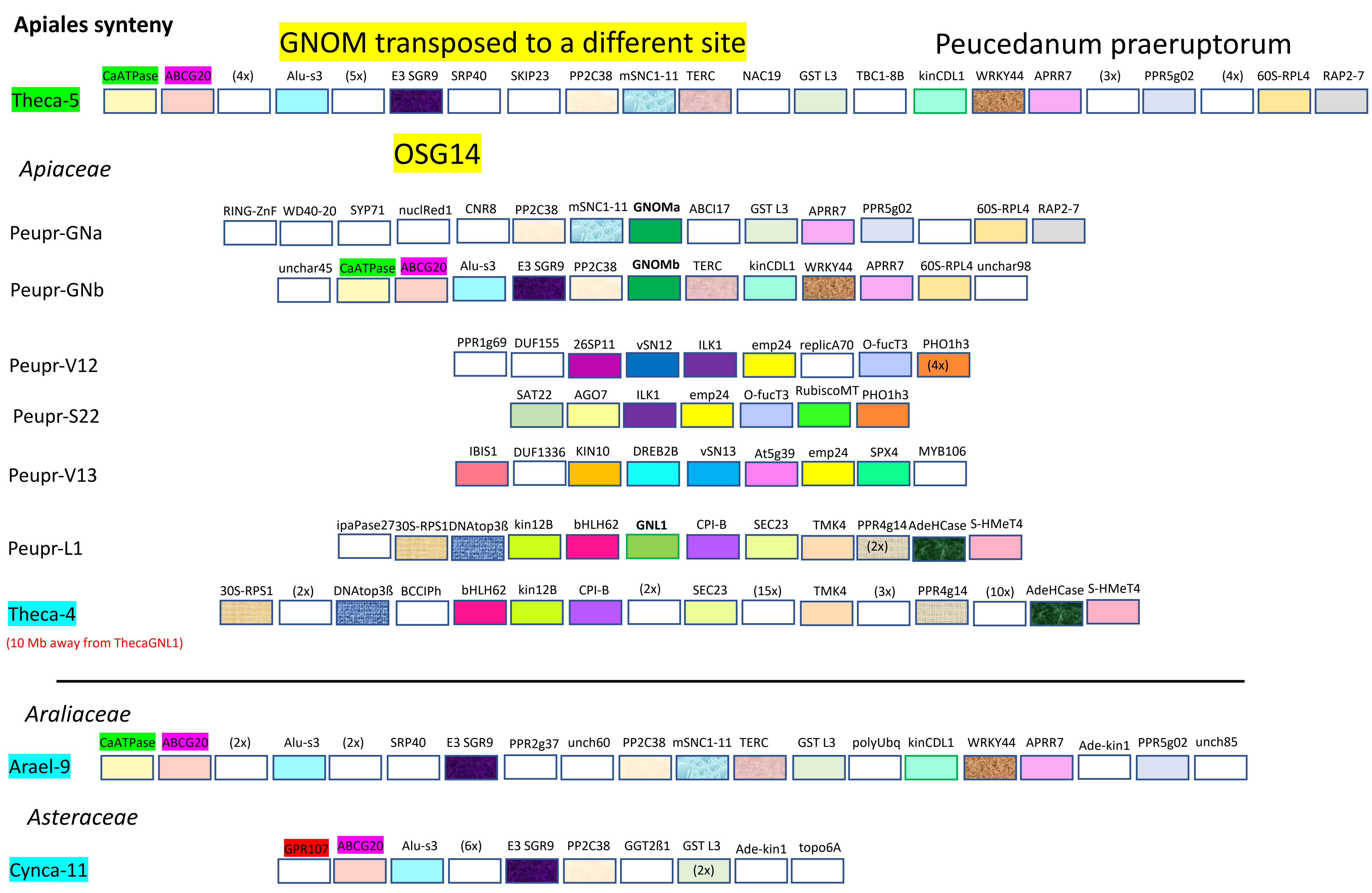
Apiales synteny
GNOM transposed to a different site
Peucedanum praeruptorum
Theca-5
CaATPase ABCG20 (4x) Alu-s3 (5x) E3 SGR9 SRP40 SKIP23 PP2C38 mSNC1-11 TERC NAC19 GST L3 TBC1-8B kinCDL1 WRKY44 APRR7 (3x) PPR5g02 (4x) 60S-RPL4 RAP2-7
OSG14
Apiaceae
Peupr-GNa
RING-ZnF WD40-20 SYP71 nuclRed1 CNR8 PP2C38 mSNC1-11 GNOMa ABCI17 GST L3 APRR7 PPR5g02 60S-RPL4 RAP2-7
Peupr-GNb
unchar45 CaATPase ABCG20 Alu-s3 E3 SGR9 PP2C38 GNOMb TERC kinCDL1 WRKY44 APRR7 60S-RPL4 unchar98
Peupr-V12
PPR1g69 DUF155 26SP11 vSN12 ILK1 emp24 replicA70 O-fucT3 PHO1h3 (4x)
Peupr-S22
SAT22 AGO7 ILK1 emp24 O-fucT3 RubiscoMT PHO1h3
Peupr-V13
IBIS1 DUF1336 KIN10 DREB2B vSN13 At5g39 emp24 SPX4 MYB106
Peupr-L1
ipaPase27 30S-RPS1 DNAtop3ß kin12B bHLH62 GNL1 CPI-B SEC23 TMK4 PPR4g14 (2x) AdeHCase S-HMeT4
Theca-4
(10 Mb away from ThecaGNL1)
30S-RPS1 (2x) DNAtop3ß BCCIPh bHLH62 kin12B CPI-B (2x) SEC23 (15x) TMK4 (3x) PPR4g14 (10x) AdeHCase S-HMeT4
Araliaceae
Arael-9
CaATPase ABCG20 (2x) Alu-s3 (2x) SRP40 E3 SGR9 PPR2g37 unch60 PP2C38 mSNC1-11 TERC GST L3 polyUbq kinCDL1 WRKY44 APRR7 Ade-kin1 PPR5g02 unch85
Asteraceae
Cynca-11
GPR107 ABCG20 Alu-s3 (6x) E3 SGR9 PP2C38 GGT2ß1 GST L3 (2x) Ade-kin1 topo6A

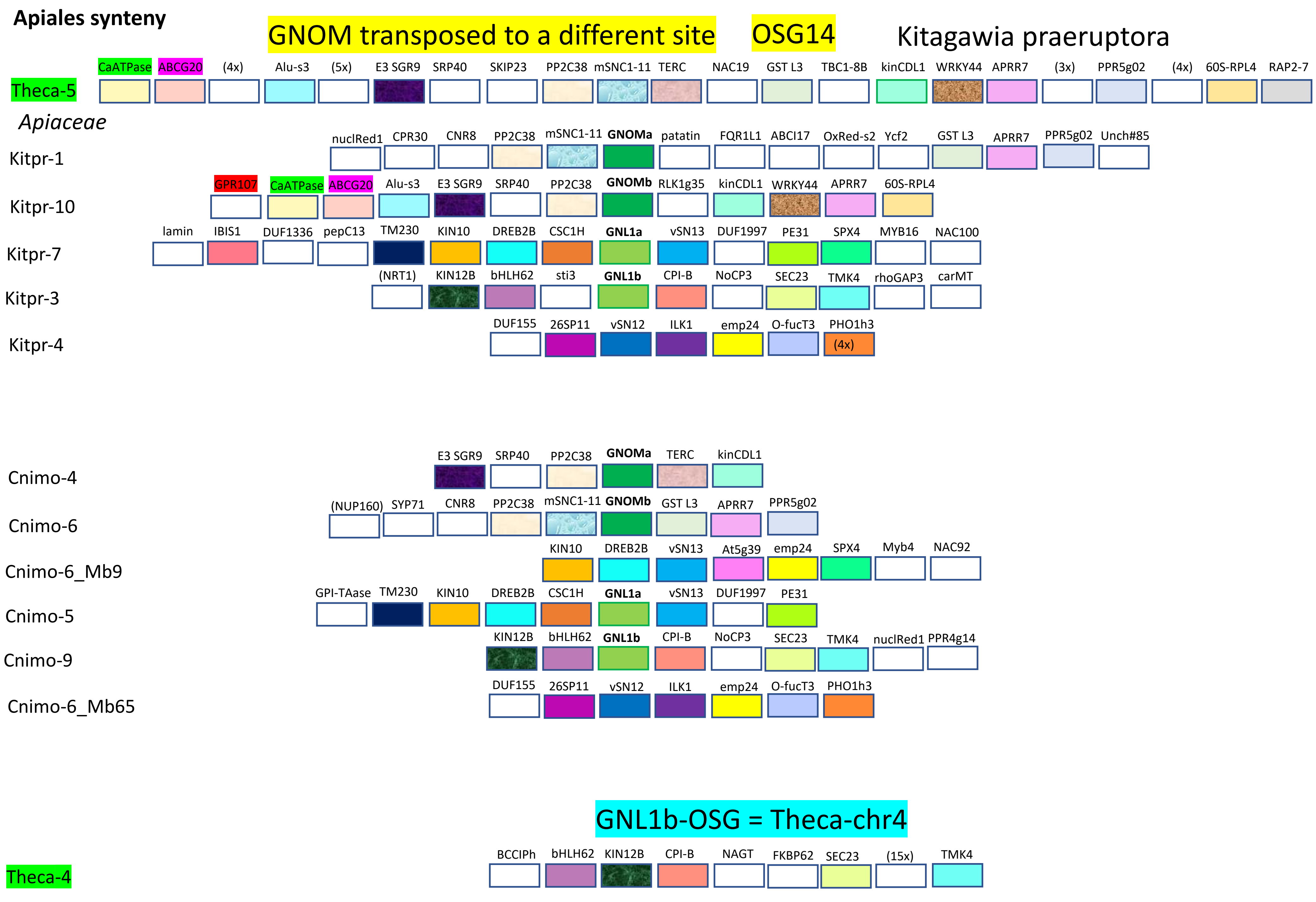
Apiales synteny
GNOM transposed to a different site
OSG14
Kitagawia praeruptora
Theca-5
CaATPase ABCG20 (4x) Alu-s3 (5x) E3 SGR9 SRP40 SKIP23 PP2C38 mSNC1-11 TERC NAC19 GST L3 TBC1-8B kinCDL1 WRKY44 APRR7 (3x) PPR5g02 (4x) 60S-RPL4 RAP2-7
Apiaceae
Kitpr-1
nuclRed1 CPR30 CNR8 PP2C38 mSNC1-11 GNOMa patatin FQR1L1 ABCI17 OxRed-s2 Ycf2 GST L3 APRR7 PPR5g02 Unch#85
Kitpr-10
GPR107 CaATPase ABCG20 Alu-s3 E3 SGR9 SRP40 PP2C38 GNOMb RLK1g35 kinCDL1 WRKY44 APRR7 60S-RPL4
Kitpr-7
lamin IBIS1 DUF1336 pepC13 TM230 KIN10 DREB2B CSC1H GNL1a vSN13 DUF1997 PE31 SPX4 MYB16 NAC100
Kitpr-3
(NRT1) KIN12B bHLH62 sti3 GNL1b CPI-B NoCP3 SEC23 TMK4 rhoGAP3 carMT
Kitpr-4
DUF155 26SP11 vSN12 ILK1 emp24 O-fucT3 PHO1h3 (4x)
Cnimo-4
E3 SGR9 SRP40 PP2C38 GNOMa TERC kinCDL1
Cnimo-6
(NUP160) SYP71 CNR8 PP2C38 mSNC1-11 GNOMb GST L3 APRR7 PPR5g02
Cnimo-6_Mb9
KIN10 DREB2B vSN13 At5g39 emp24 SPX4 Myb4 NAC92
Cnimo-5
GPI-TAase TM230 KIN10 DREB2B CSC1H GNL1a vSN13 DUF1997 PE31
Cnimo-9
KIN12B bHLH62 GNL1b CPI-B NoCP3 SEC23 TMK4 nuclRed1 PPR4g14
Cnimo-6_Mb65
DUF155 26SP11 vSN12 ILK1 emp24 O-fucT3 PHO1h3
GNL1b-OSG = Theca-chr4
Theca-4
BCCIPh bHLH62 KIN12B CPI-B NAGT FKBP62 SEC23 (15x) TMK4

**Apiales synteny**

GNOM transposed to a different site OSG141

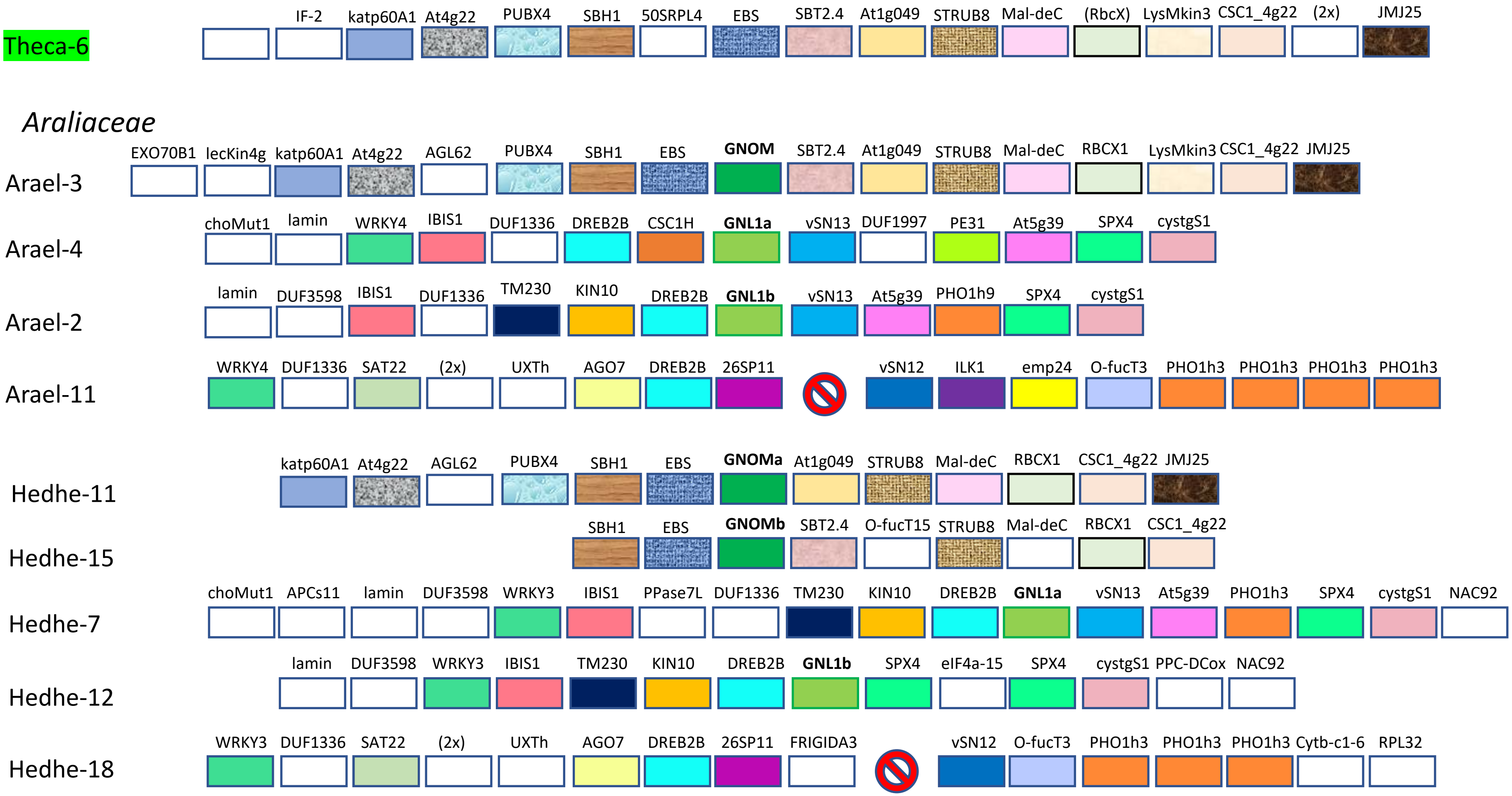

**Apiales synteny**

GNOM transposed to a different site OSG141

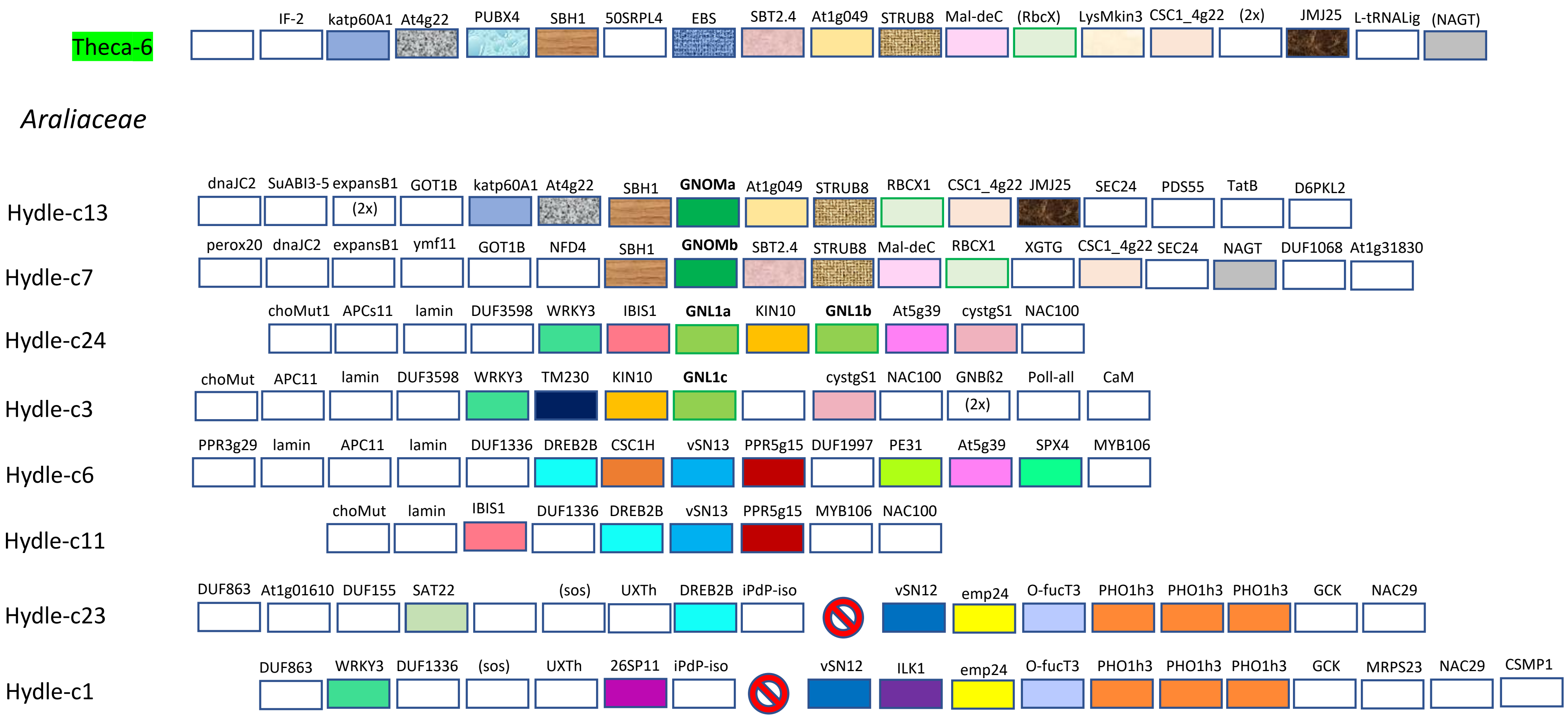

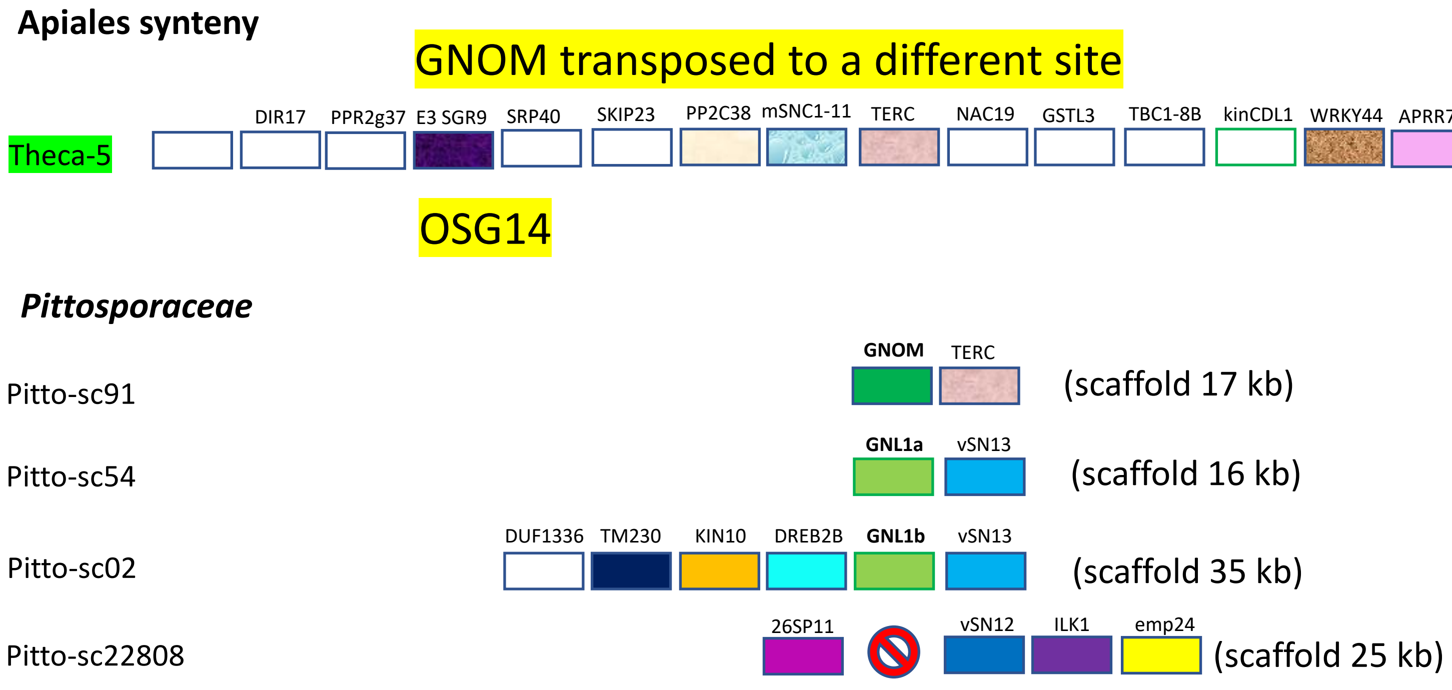
Apiales synteny
GNOM transposed to a different site
Theca-5
DIR17
PPR2g37
E3 SGR9
SRP40
SKIP23
PP2C38
mSNC1-11
TERC
NAC19
GSTL3
TBC1-8B
kinCDL1
WRKY44
APRR7
Ade-kin1
AAED1
topo6A
PPR5g02
ZnF11
OSG14
Pittosporaceae
Pitto-sc91
GNOM
TERC
(scaffold 17 kb)
Pitto-sc54
GNL1a
vSN13
(scaffold 16 kb)
Pitto-sc02
DUF1336
TM230
KIN10
DREB2B
GNL1b
vSN13
(scaffold 35 kb)
Pitto-sc22808
26SP11
vSN12
ILK1
emp24
(scaffold 25 kb)

#### Suppl. Figure S28. Dipsacales cladogram and OSGs

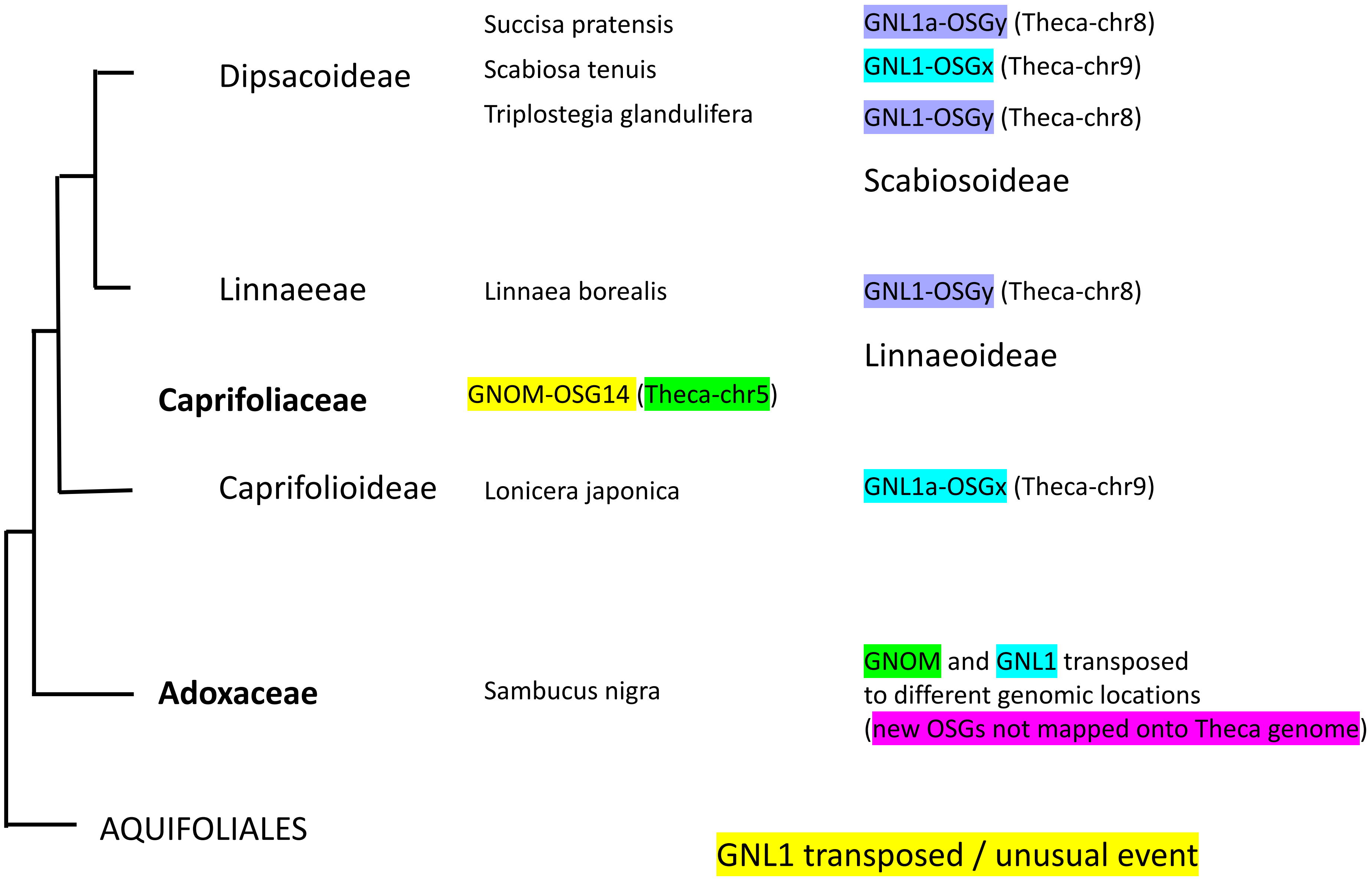

Cladogram based on Lee et al (2021); Zuntini et al. (2024)

### Dipsacales synteny

OSG-14 (Aquifoliales, Garryales, Dipsacales, Apiales, Escalloniales)

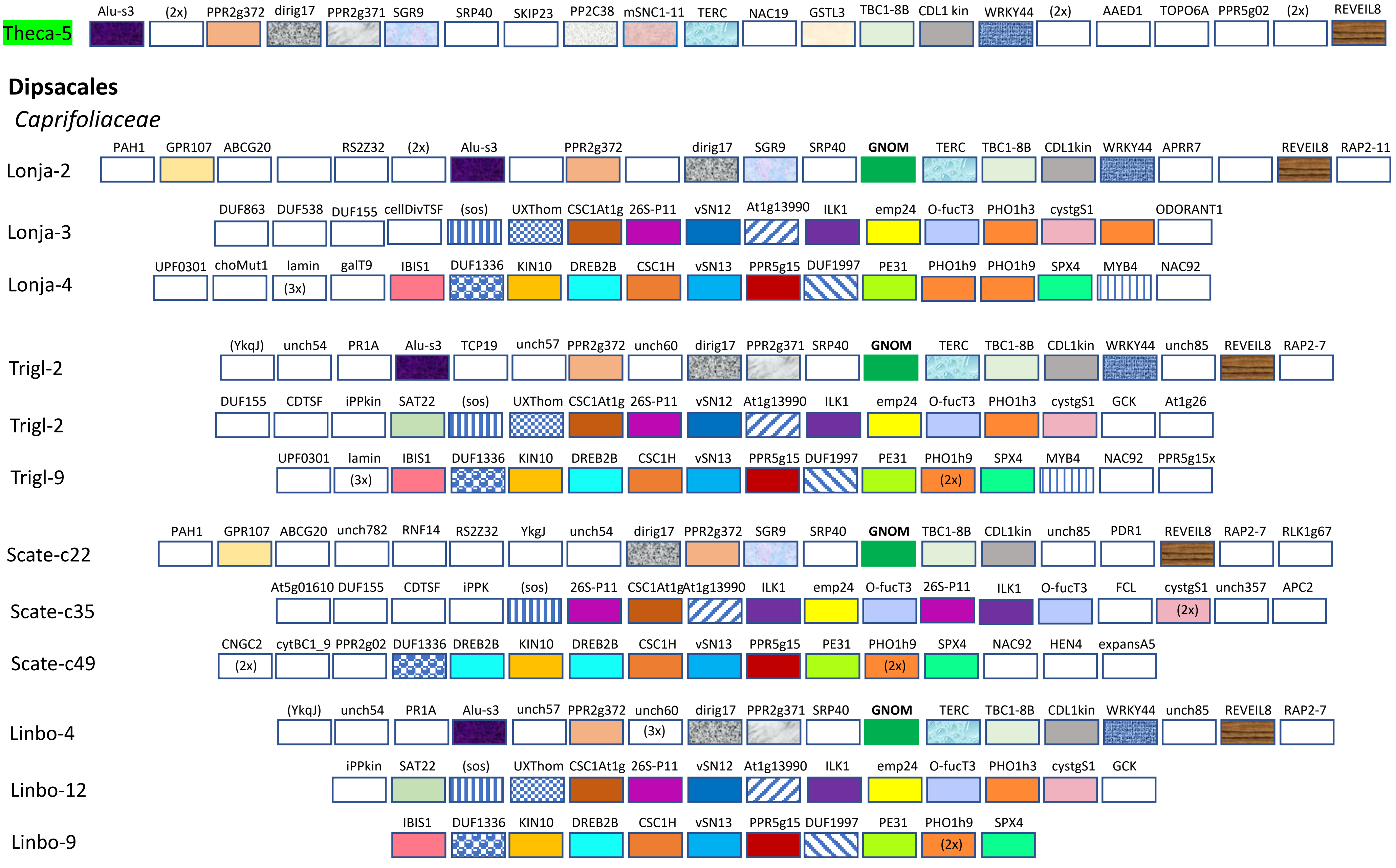

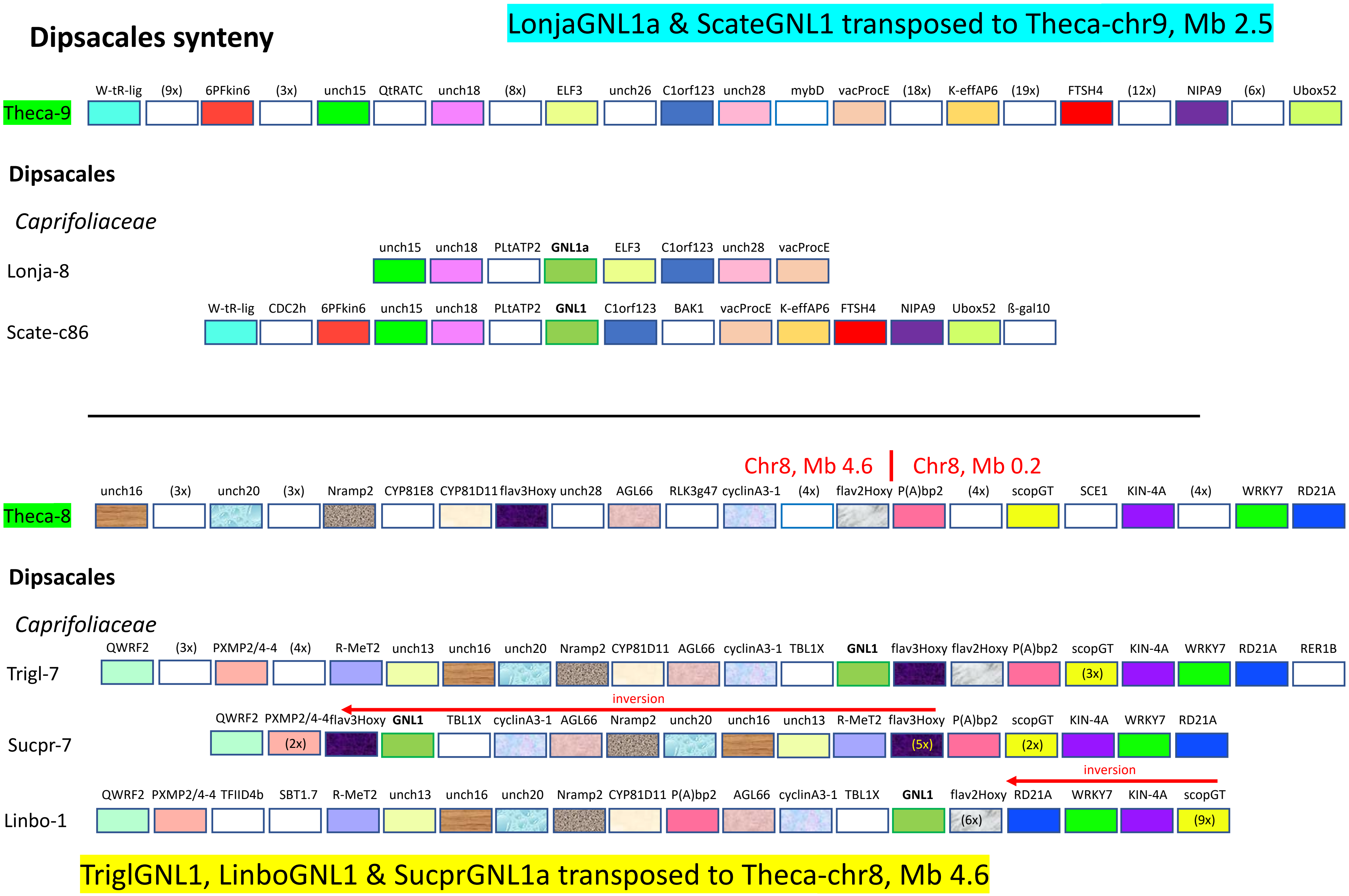
Dipsacales synteny
LonjaGNL1a & ScateGNL1 transposed to Theca-chr9, Mb 2.5
Theca-9
W-tR-lig (9x) 6PFkin6 (3x) unch15 QtRATC unch18 (8x) ELF3 unch26 C1orf123 unch28 mybD vacProcE (18x) K-effAP6 (19x) FTSH4 (12x) NIPA9 (6x) Ubox52
Dipsacales
Caprifoliaceae
Lonja-8
unch15 unch18 PLtATP2 GNL1a ELF3 C1orf123 unch28 vacProcE
Scate-c86
W-tR-lig CDC2h 6PFkin6 unch15 unch18 PLtATP2 GNL1 C1orf123 BAK1 vacProcE K-effAP6 FTSH4 NIPA9 Ubox52 ß-gal10
Chr8, Mb 4.6
Chr8, Mb 0.2
Theca-8
unch16 (3x) unch20 (3x) Nramp2 CYP81E8 CYP81D11 flav3Hoxy unch28 AGL66 RLK3g47 cyclinA3-1 (4x) flav2Hoxy P(A)bp2 (4x) scopGT SCE1 KIN-4A (4x) WRKY7 RD21A
Dipsacales
Caprifoliaceae
Trigl-7
QWRF2 (3x) PXMP2/4-4 (4x) R-MeT2 unch13 unch16 unch20 Nramp2 CYP81D11 AGL66 cyclinA3-1 TBL1X GNL1 flav3Hoxy flav2Hoxy P(A)bp2 scopGT KIN-4A WRKY7 RD21A RER1B
(3x)
inversion
Sucpr-7
QWRF2 PXMP2/4-4 flav3Hoxy GNL1 TBL1X cyclinA3-1 AGL66 Nramp2 unch20 unch16 unch13 R-MeT2 flav3Hoxy P(A)bp2 scopGT KIN-4A WRKY7 RD21A
(2x)
(5x)
(2x)
inversion
Linbo-1
QWRF2 PXMP2/4-4 TFIID4b SBT1.7 R-MeT2 unch13 unch16 unch20 Nramp2 CYP81D11 P(A)bp2 AGL66 cyclinA3-1 TBL1X GNL1 flav2Hoxy RD21A WRKY7 KIN-4A scopGT
(6x)
(9x)
TriglGNL1, LinboGNL1 & SucprGNL1a transposed to Theca-chr8, Mb 4.6

*Adoxaceae*

Sambucus nigra / large genome (11.8 Gb)

GNOM chr15, Mb 230.9

GNL1 chr15, Mb 528.1

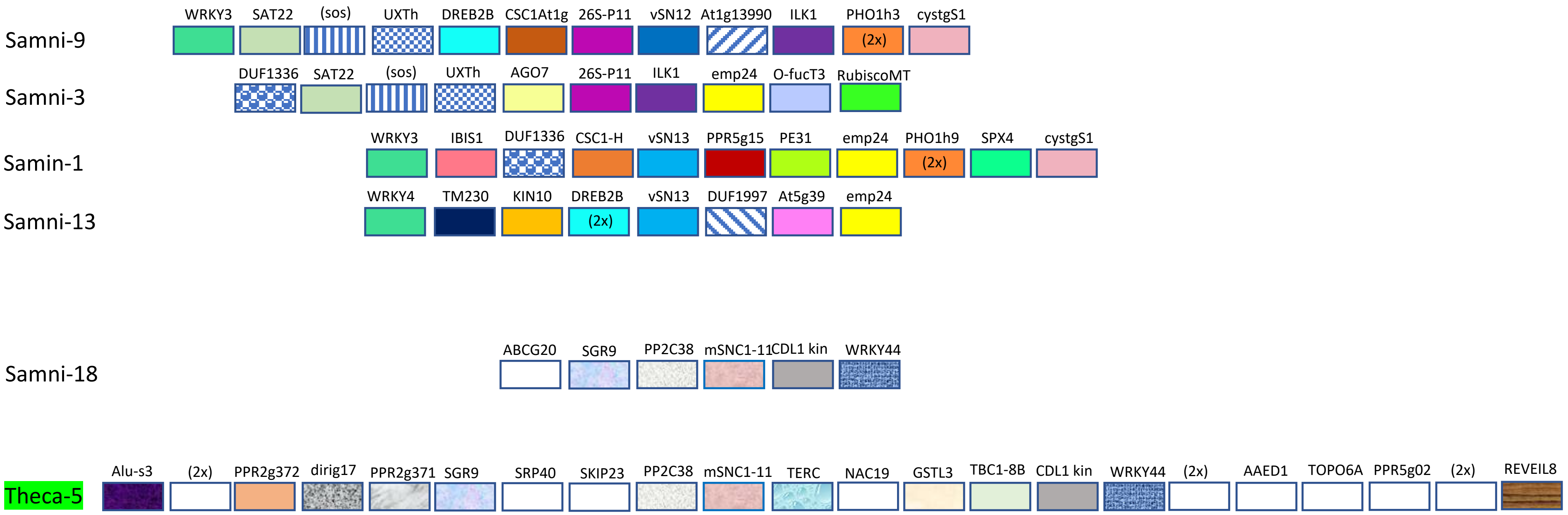

No SamniGNOM in OSG14

OSGs incomplete because of TblastN searches – too many repeated/transposon sequences

#### Suppl. Figure S29. Escalloniales cladogram and OSGs

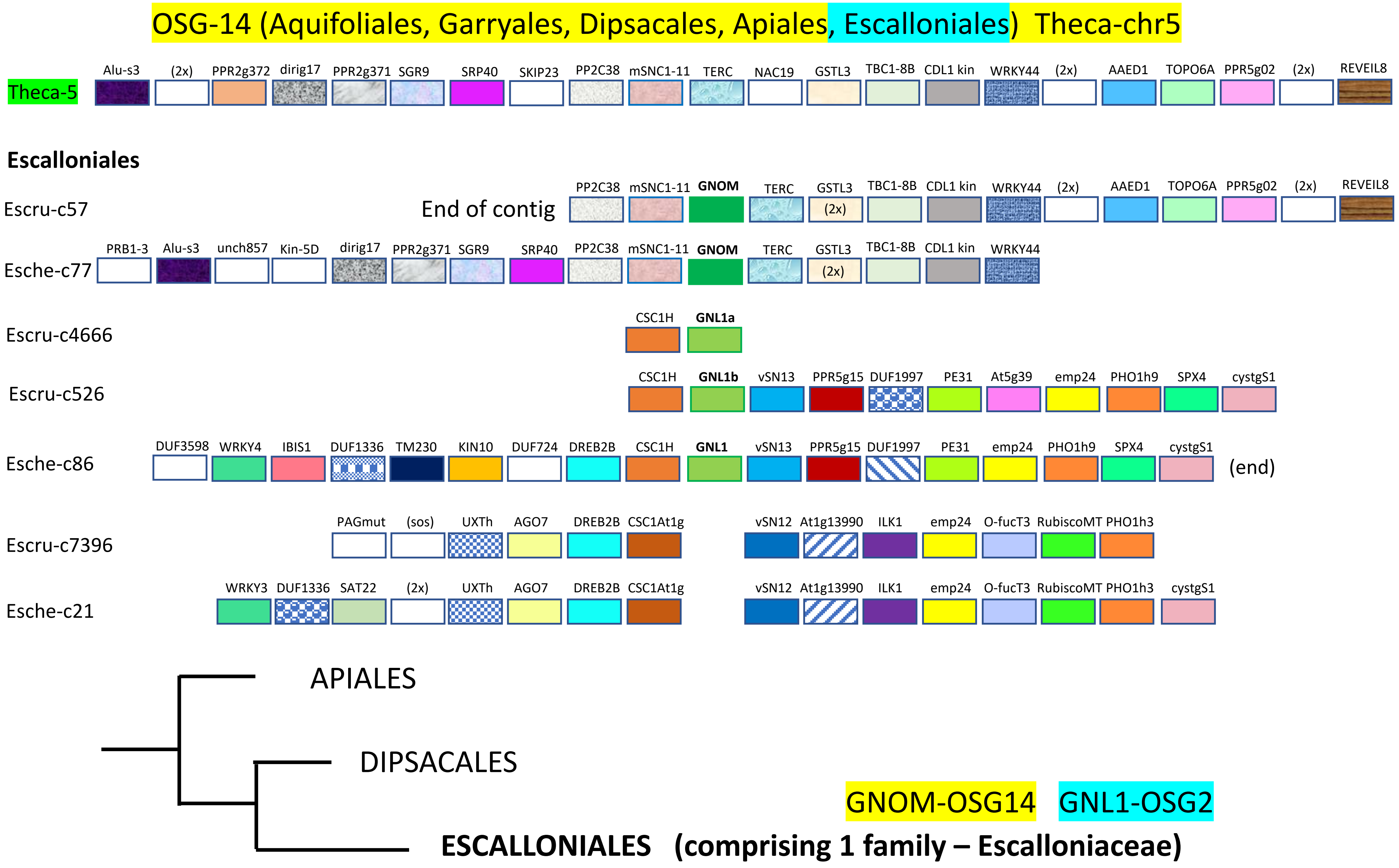

**Suppl. Figure S30. Asterales cladogram and OSGs**

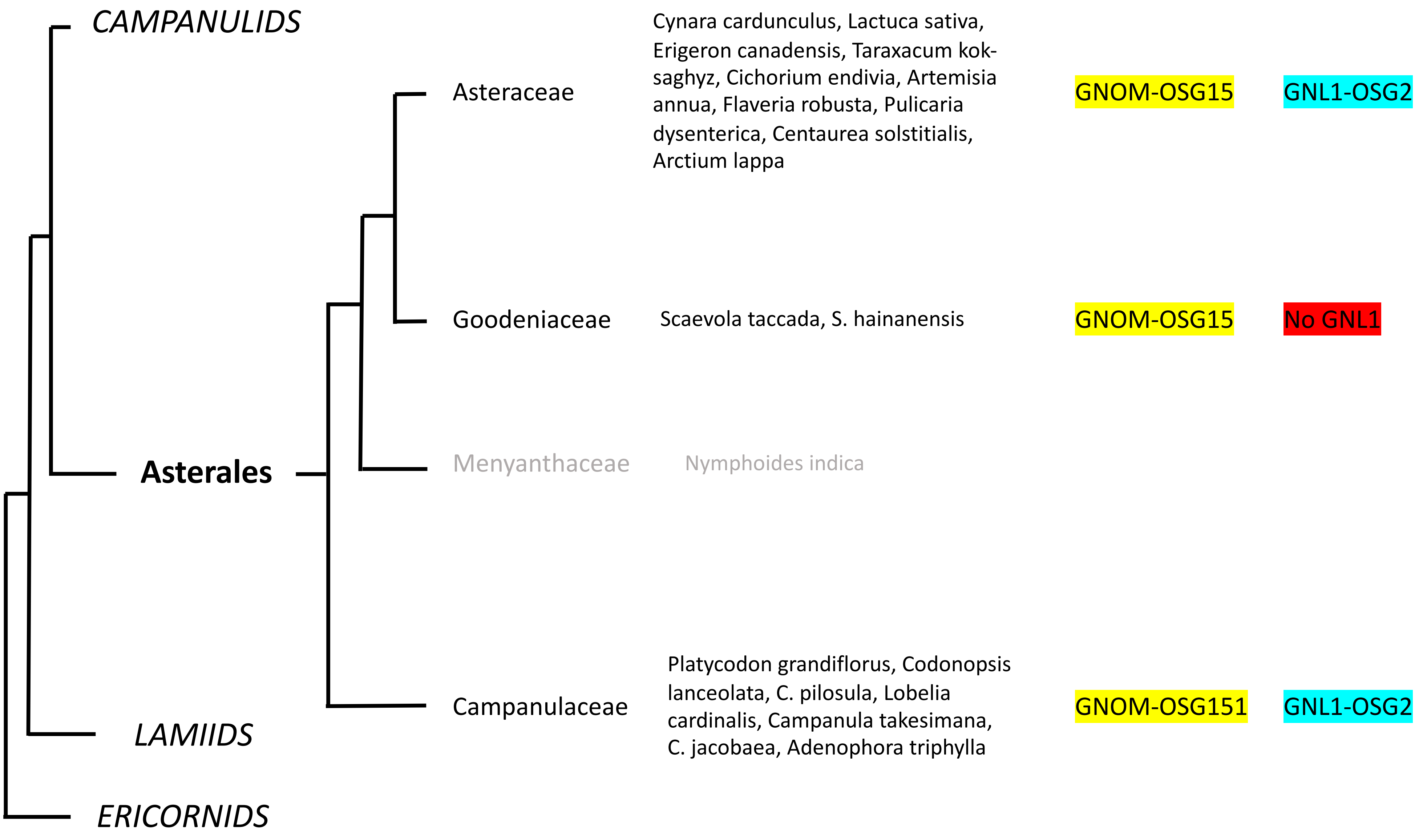

Cladogram based on Zhang et al. (2020c)

**Asterales** (CSC1-like At1g69450 VTI12 synteny) GNOM-OSG15 GNOM transposed

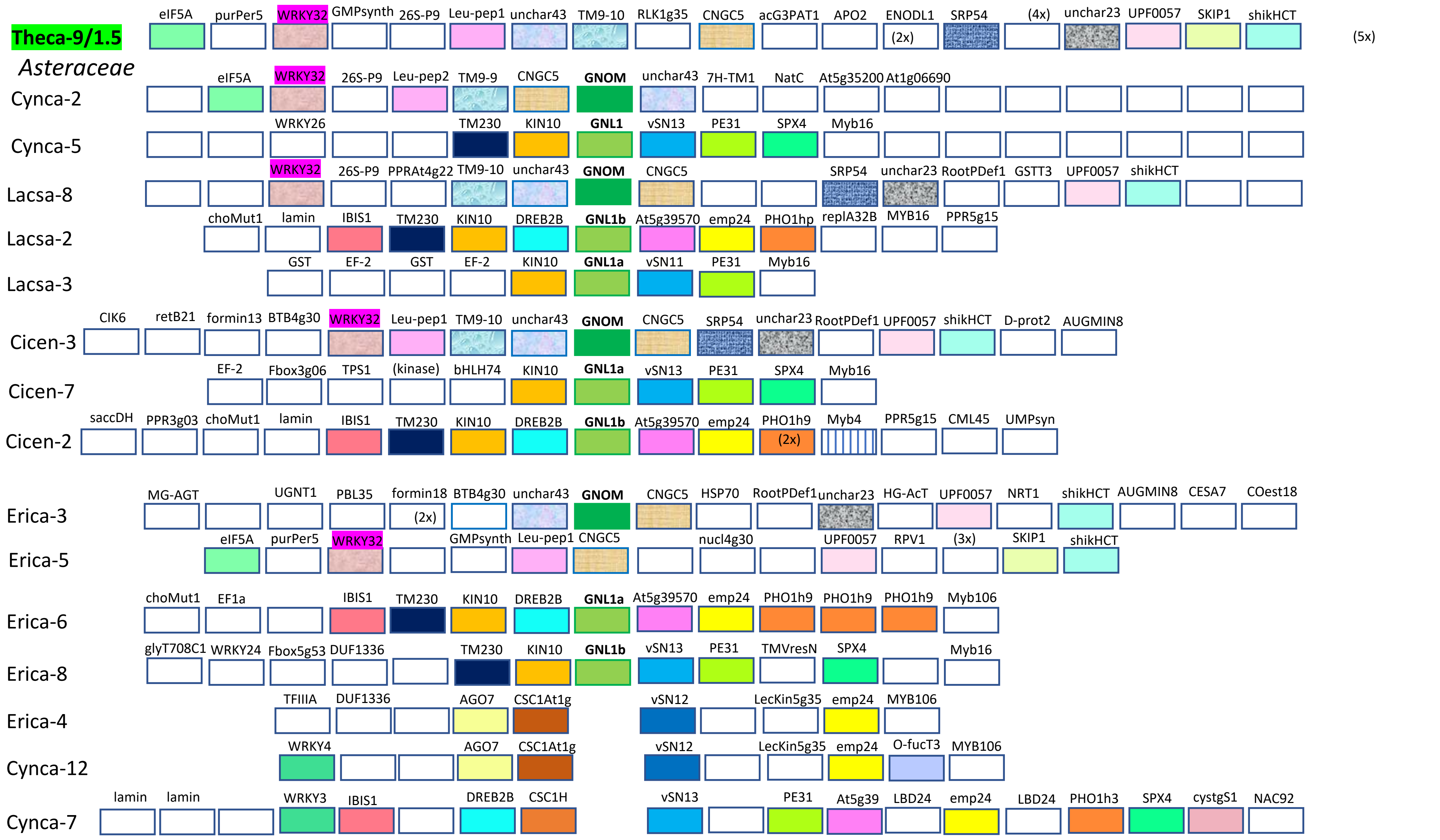

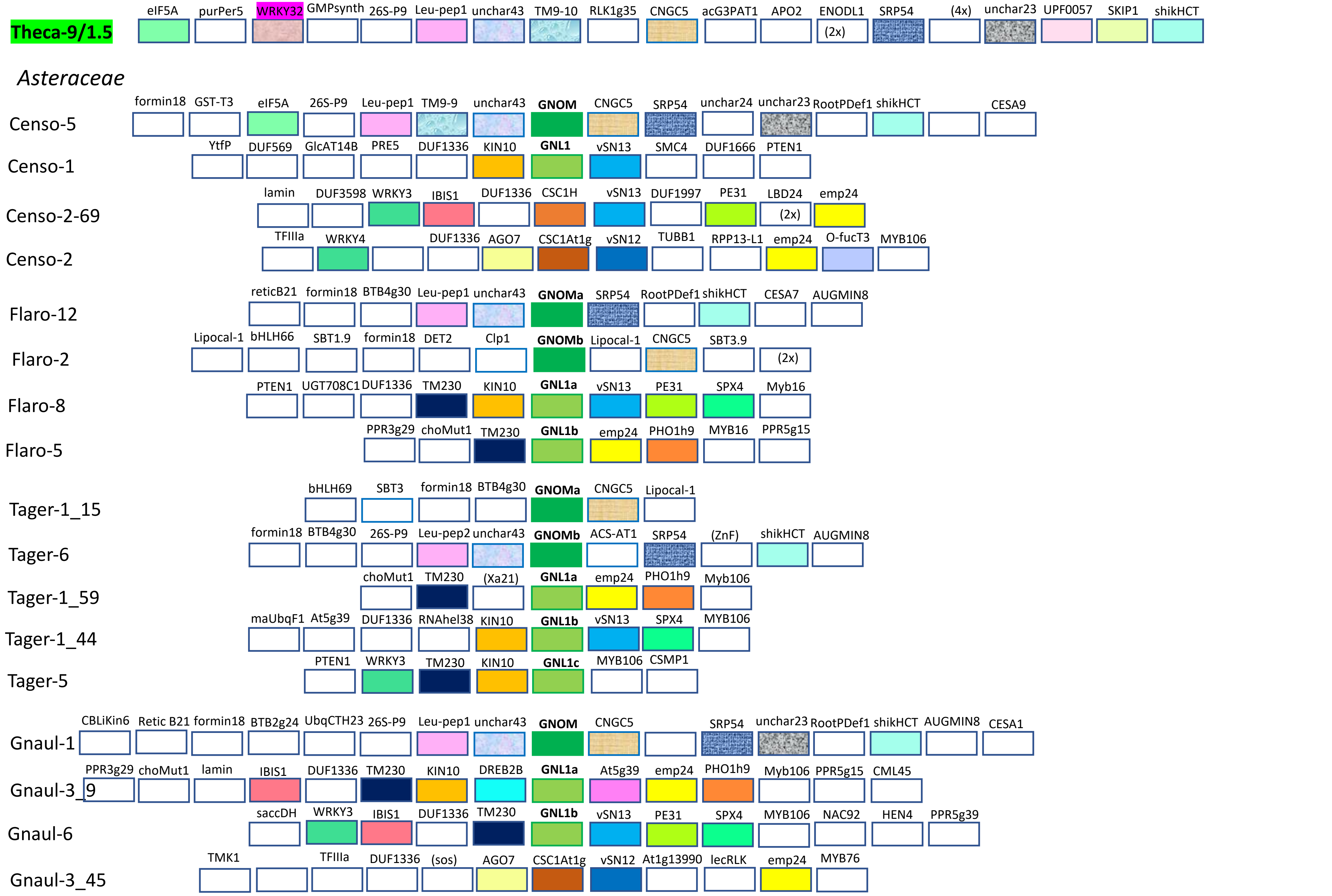
Theca-9/1.5
eIF5A purPer5 WRKY32 GMPsynth 26S-P9 Leu-pep1 unchar43 TM9-10 RLK1g35 CNGC5 acG3PAT1 APO2 ENODL1 (2x) SRP54 (4x) unchar23 UPF0057 SKIP1 shikHCT
Asteraceae
Censo-5
formin18 GST-T3 eIF5A 26S-P9 Leu-pep1 TM9-9 unchar43 GNOM CNGC5 SRP54 unchar24 unchar23 RootPDef1 shikHCT CESA9
Censo-1
YtfP DUF569 GlcAT14B PRE5 DUF1336 KIN10 GNL1 vSN13 SMC4 DUF1666 PTEN1
Censo-2-69
lamin DUF3598 WRKY3 IBIS1 DUF1336 CSC1H vSN13 DUF1997 PE31 LBD24 (2x) emp24
Censo-2
TFIIIa WRKY4 DUF1336 AGO7 CSC1At1g vSN12 TUBB1 RPP13-L1 emp24 O-fucT3 MYB106
Flaro-12
reticB21 formin18 BTB4g30 Leu-pep1 unchar43 GNOMa SRP54 RootPDef1 shikHCT CESA7 AUGMIN8
Flaro-2
Lipocal-1 bHLH66 SBT1.9 formin18 DET2 Clp1 GNOMb Lipocal-1 CNGC5 SBT3.9 (2x)
Flaro-8
PTEN1 UGT708C1 DUF1336 TM230 KIN10 GNL1a vSN13 PE31 SPX4 Myb16
Flaro-5
PPR3g29 choMut1 TM230 GNL1b emp24 PHO1h9 MYB16 PPR5g15
Tager-1_15
bHLH69 SBT3 formin18 BTB4g30 GNOMa CNGC5 Lipocal-1
Tager-6
formin18 BTB4g30 26S-P9 Leu-pep2 unchar43 GNOMb ACS-AT1 SRP54 (ZnF) shikHCT AUGMIN8
Tager-1_59
choMut1 TM230 (Xa21) GNL1a emp24 PHO1h9 Myb106
Tager-1_44
maUbqF1 At5g39 DUF1336 RNAhel38 KIN10 GNL1b vSN13 SPX4 MYB106
Tager-5
PTEN1 WRKY3 TM230 KIN10 GNL1c MYB106 CSMP1
Gnaul-1
CBLiKin6 Retic B21 formin18 BTB2g24 UbqCTH23 26S-P9 Leu-pep1 unchar43 GNOM CNGC5 SRP54 unchar23 RootPDef1 shikHCT AUGMIN8 CESA1
Gnaul-3_9
PPR3g29 choMut1 lamin IBIS1 DUF1336 TM230 KIN10 DREB2B GNL1a At5g39 emp24 PHO1h9 Myb106 PPR5g15 CML45
Gnaul-6
saccDH WRKY3 IBIS1 DUF1336 TM230 GNL1b vSN13 PE31 SPX4 MYB106 NAC92 HEN4 PPR5g39
Gnaul-3_45
TMK1 TFIIIa DUF1336 (sos) AGO7 CSC1At1g vSN12 At1g13990 lecRLK emp24 MYB76

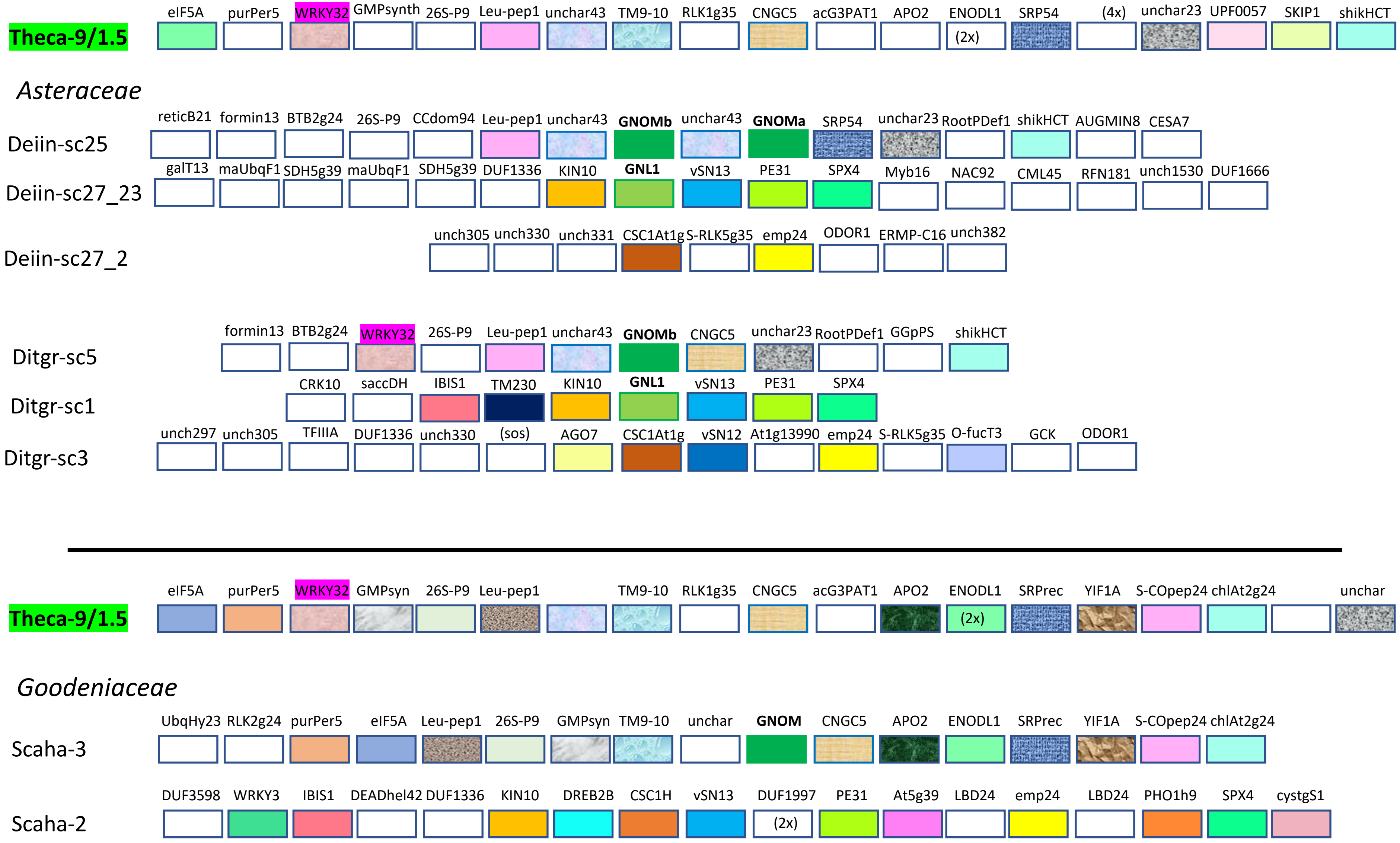
Theca-9/1.5
eIF5A purPer5 WRKY32 GMPsynth 26S-P9 Leu-pep1 unchar43 TM9-10 RLK1g35 CNGC5 acG3PAT1 APO2 ENODL1 (2x) SRP54 (4x) unchar23 UPF0057 SKIP1 shikHCT
Asteraceae
Deiin-sc25
reticB21 formin13 BTB2g24 26S-P9 CCdom94 Leu-pep1 unchar43 GNOMb unchar43 GNOMa SRP54 unchar23 RootPDef1 shikHCT AUGMIN8 CESA7
Deiin-sc27_23
galT13 maUbqF1 SDH5g39 maUbqF1 SDH5g39 DUF1336 KIN10 GNL1 vSN13 PE31 SPX4 Myb16 NAC92 CML45 RFN181 unch1530 DUF1666
Deiin-sc27_2
unch305 unch330 unch331 CSC1At1g S-RLK5g35 emp24 ODOR1 ERMP-C16 unch382
Ditgr-sc5
formin13 BTB2g24 WRKY32 26S-P9 Leu-pep1 unchar43 GNOMb CNGC5 unchar23 RootPDef1 GGpPS shikHCT
Ditgr-sc1
CRK10 saccDH IBIS1 TM230 KIN10 GNL1 vSN13 PE31 SPX4
Ditgr-sc3
unch297 unch305 TFIIIA DUF1336 unch330 (sos) AGO7 CSC1At1g vSN12 At1g13990 emp24 S-RLK5g35 O-fucT3 GCK ODOR1
Theca-9/1.5
eIF5A purPer5 WRKY32 GMPsyn 26S-P9 Leu-pep1 TM9-10 RLK1g35 CNGC5 acG3PAT1 APO2 ENODL1 (2x) SRPrec YIF1A S-COpep24 chlAt2g24 unchar
Goodeniaceae
Scaha-3
UbqHy23 RLK2g24 purPer5 eIF5A Leu-pep1 26S-P9 GMPsyn TM9-10 unchar GNOM CNGC5 APO2 ENODL1 SRPrec YIF1A S-COpep24 chlAt2g24
Scaha-2
DUF3598 WRKY3 IBIS1 DEADhel42 DUF1336 KIN10 DREB2B CSC1H vSN13 DUF1997 (2x) PE31 At5g39 LBD24 emp24 LBD24 PHO1h9 SPX4 cystgS1

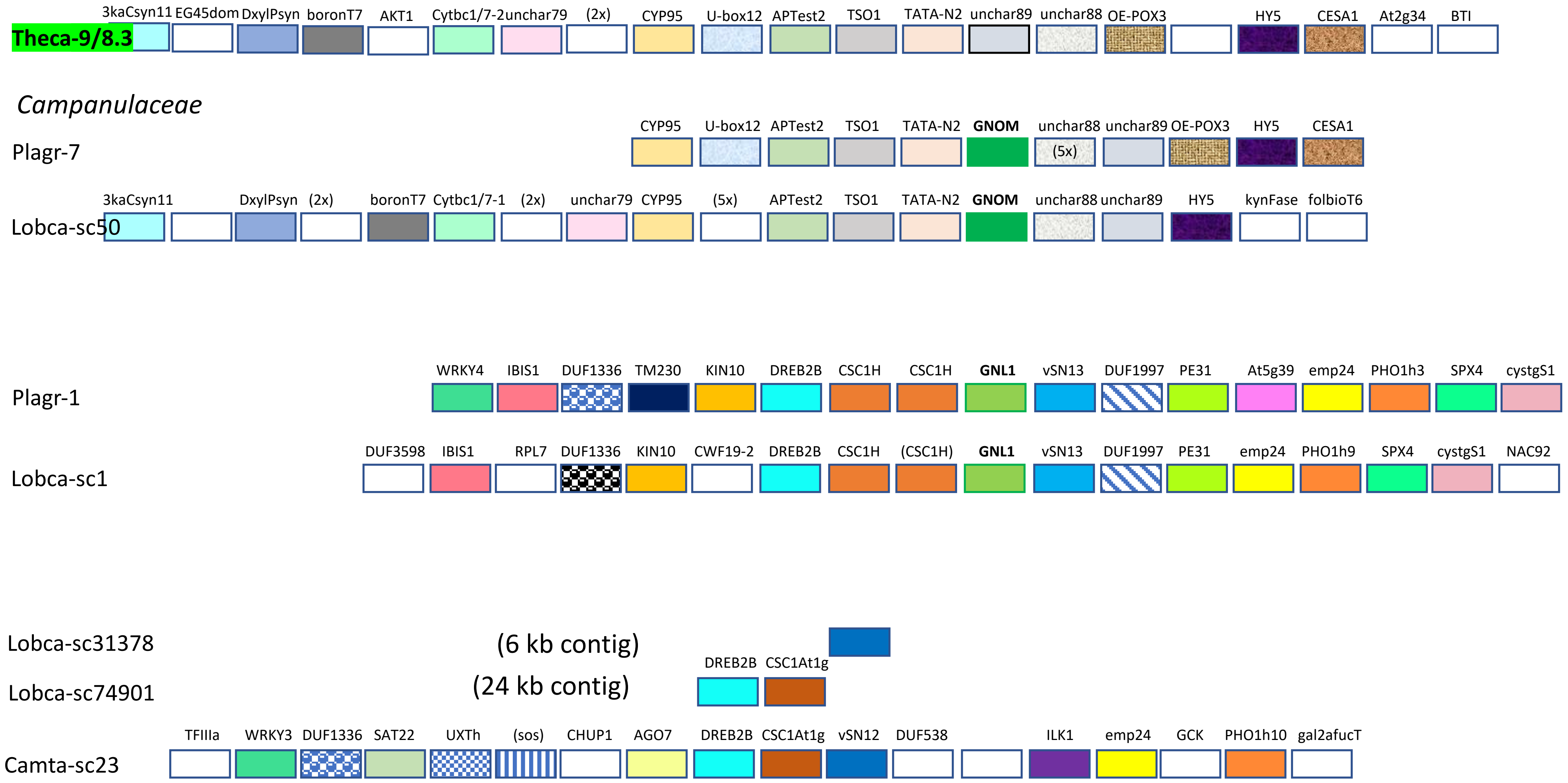
Theca-9/8.3
3kaCsyn11 EG45dom DxylPsyn boronT7 AKT1 Cytbc1/7-2 unchar79 (2x) CYP95 U-box12 APTest2 TSO1 TATA-N2 unchar89 unchar88 OE-POX3 HY5 CESA1 At2g34 BTI
Campanulaceae
Plagr-7
CYP95 U-box12 APTest2 TSO1 TATA-N2 GNOM unchar88 (5x) unchar89 OE-POX3 HY5 CESA1
Lobca-sc50
3kaCsyn11 DxylPsyn (2x) boronT7 Cytbc1/7-1 (2x) unchar79 CYP95 (5x) APTest2 TSO1 TATA-N2 GNOM unchar88 unchar89 HY5 kynFase folbioT6
Plagr-1
WRKY4 IBIS1 DUF1336 TM230 KIN10 DREB2B CSC1H CSC1H GNL1 vSN13 DUF1997 PE31 At5g39 emp24 PHO1h3 SPX4 cystgS1
Lobca-sc1
DUF3598 IBIS1 RPL7 DUF1336 KIN10 CWF19-2 DREB2B CSC1H (CSC1H) GNL1 vSN13 DUF1997 PE31 emp24 PHO1h9 SPX4 cystgS1 NAC92
Lobca-sc31378
(6 kb contig)
Lobca-sc74901
(24 kb contig)
DREB2B CSC1At1g
Camta-sc23
TFIIIa WRKY3 DUF1336 SAT22 UXTh (sos) CHUP1 AGO7 DREB2B CSC1At1g vSN12 DUF538 ILK1 emp24 GCK PHO1h10 gal2afucT

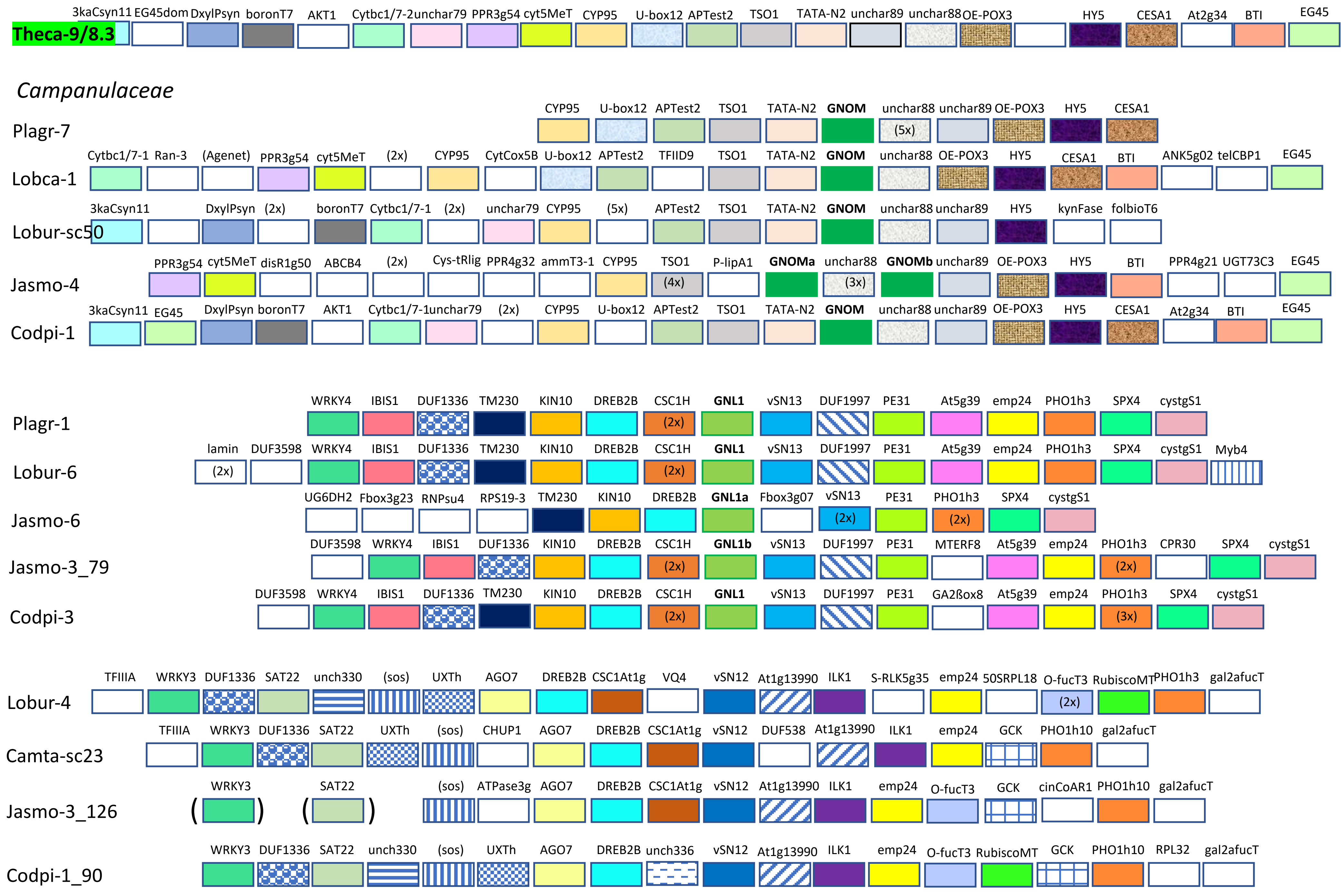

3kaCsyn11 EG45dom DxylPsyn boronT7 AKT1 Cytbc1/7-2 unchar79 PPR3g54 cyt5MeT CYP95 U-box12 APTest2 TSO1 TATA-N2 unchar89 unchar88 OE-POX3 HY5 CESA1 At2g34 BTI EG45
Theca-9/8.3
Campanulaceae
Plagr-7
CYP95 U-box12 APTest2 TSO1 TATA-N2 GNOM unchar88 unchar89 OE-POX3 HY5 CESA1
(5x)
Lobca-1
Cytbc1/7-1 Ran-3 (Agenet) PPR3g54 cyt5MeT (2x) CYP95 CytCox5B U-box12 APTest2 TFIID9 TSO1 TATA-N2 GNOM unchar88 OE-POX3 HY5 CESA1 BTI ANK5g02 telCBP1 EG45
Lobur-sc50
3kaCsyn11 DxylPsyn (2x) boronT7 Cytbc1/7-1 (2x) unchar79 CYP95 (5x) APTest2 TSO1 TATA-N2 GNOM unchar88 unchar89 HY5 kynFase folbioT6
Jasmo-4
PPR3g54 cyt5MeT disR1g50 ABCB4 (2x) Cys-tRlig PPR4g32 ammT3-1 CYP95 TSO1 P-lipA1 GNOMa unchar88 GNOMb unchar89 OE-POX3 HY5 BTI PPR4g21 UGT73C3 EG45
(4x) (3x)
Codpi-1
3kaCsyn11 EG45 DxylPsyn boronT7 AKT1 Cytbc1/7-1 unchar79 (2x) CYP95 U-box12 APTest2 TSO1 TATA-N2 GNOM unchar88 unchar89 OE-POX3 HY5 CESA1 At2g34 BTI EG45
Plagr-1
WRKY4 IBIS1 DUF1336 TM230 KIN10 DREB2B CSC1H GNL1 vSN13 DUF1997 PE31 At5g39 emp24 PHO1h3 SPX4 cystgS1
(2x)
Lobur-6
lamin DUF3598 WRKY4 IBIS1 DUF1336 TM230 KIN10 DREB2B CSC1H GNL1 vSN13 DUF1997 PE31 At5g39 emp24 PHO1h3 SPX4 cystgS1 Myb4
(2x) (2x)
Jasmo-6
UG6DH2 Fbox3g23 RNPsu4 RPS19-3 TM230 KIN10 DREB2B GNL1a Fbox3g07 vSN13 PE31 PHO1h3 SPX4 cystgS1
(2x) (2x)
Jasmo-3_79
DUF3598 WRKY4 IBIS1 DUF1336 KIN10 DREB2B CSC1H GNL1b vSN13 DUF1997 PE31 MTERF8 At5g39 emp24 PHO1h3 CPR30 SPX4 cystgS1
(2x) (2x)
Codpi-3
DUF3598 WRKY4 IBIS1 DUF1336 TM230 KIN10 DREB2B CSC1H GNL1 vSN13 DUF1997 PE31 GA2ßox8 At5g39 emp24 PHO1h3 SPX4 cystgS1
(2x) (3x)
Lobur-4
TFIIIA WRKY3 DUF1336 SAT22 unch330 (sos) UXTh AGO7 DREB2B CSC1At1g VQ4 vSN12 At1g13990 ILK1 S-RLK5g35 emp24 50SRPL18 O-fucT3 RubiscoMT PHO1h3 gal2afucT
(2x)
Camta-sc23
TFIIIA WRKY3 DUF1336 SAT22 UXTh (sos) CHUP1 AGO7 DREB2B CSC1At1g vSN12 DUF538 At1g13990 ILK1 emp24 GCK PHO1h10 gal2afucT
Jasmo-3_126
WRKY3 SAT22 (sos) ATPase3g AGO7 DREB2B CSC1At1g vSN12 At1g13990 ILK1 emp24 O-fucT3 GCK cinCoAR1 PHO1h10 gal2afucT
Codpi-1_90
WRKY3 DUF1336 SAT22 unch330 (sos) UXTh AGO7 DREB2B unch336 vSN12 At1g13990 ILK1 emp24 O-fucT3 RubiscoMT GCK PHO1h10 RPL32 gal2afucT

### Suppl. Figure S31. Ericales cladogram and OSGs

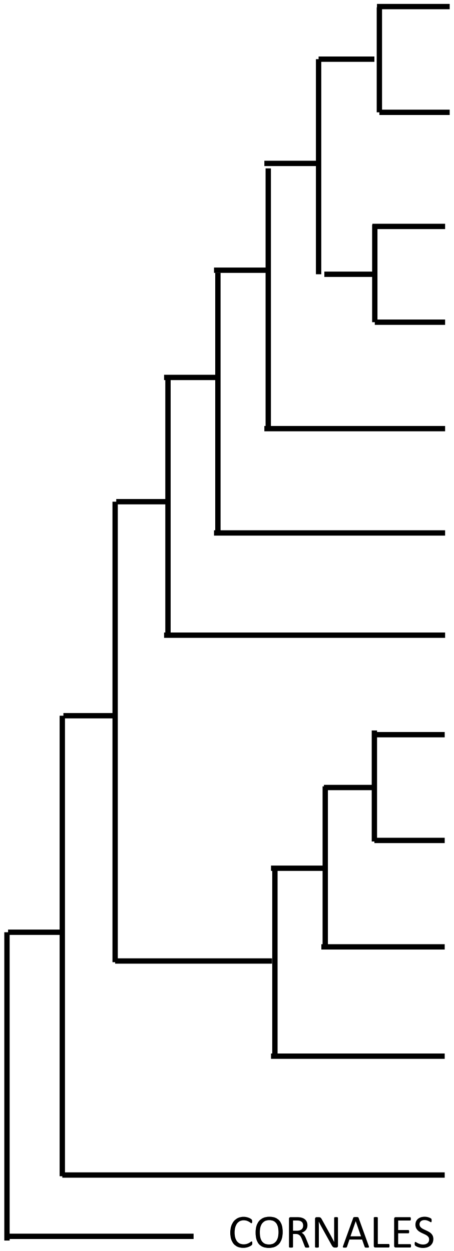

| | | | |
|---|---|---|---|
| Ericaceae | Rhododendron vialii, Gaultheria prostrata, Vaccinium darrowii, Arctostaphylos glauc | OSG1215f | OSG2 |
| Clethraceae | Clethra arborea | OSG1 | OSG2 |
| Actinidiaceae | Actinidia chinensis | OSG1 | OSG2 |
| Roridulaceae | Roridula gorgonias | OSG1 | OSG2 |
| Styracaceae | Sinojackia xylocarpa | | |
| Theaceae | Camellia sinensis | OSG1 | OSG2 |
| Pentaphylacaceae | Euryodendron excelsum | OSG1 | OSG2 |
| Primulacaceae | Aegiceras corniculatum, Androsace saussurei | OSG1 | no GNL1 |
| Ebenaceae | Diospyros oleifera | OSG1 | OSG2 |
| Sapotaceae | Argania spinosa | OSG1 | OSG2 |
| Polemoniaceae | Linanthus parryae | OSG1 | OSG2 |
| Balsaminaceae | Impatiens glandulifera | OSG1 | OSG2 |

Cladogram based on Zhang et al. (2020c)

### Ericales synteny

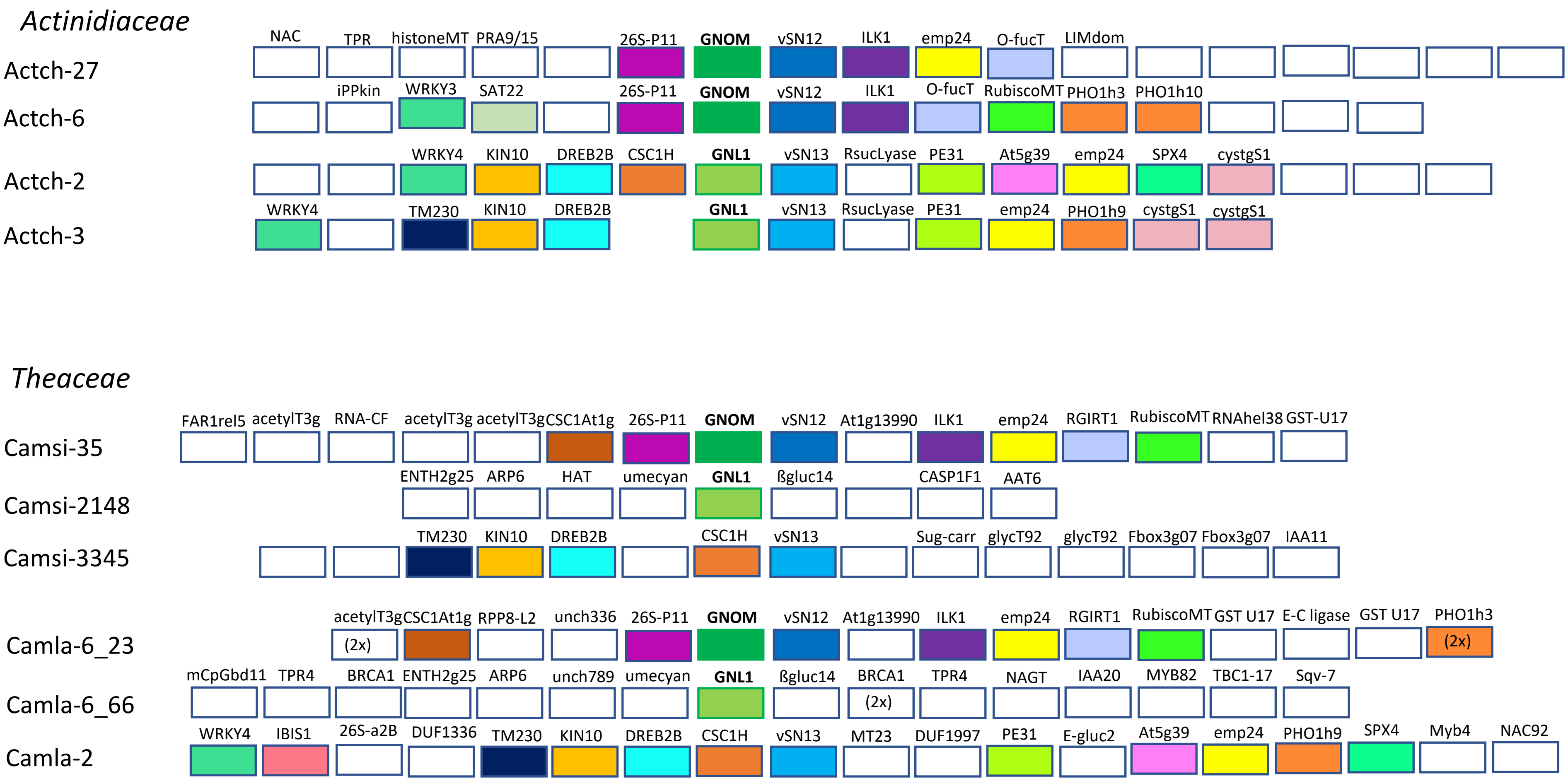

Transposition of GNL1 to Theca-chr9_Mb7.0?

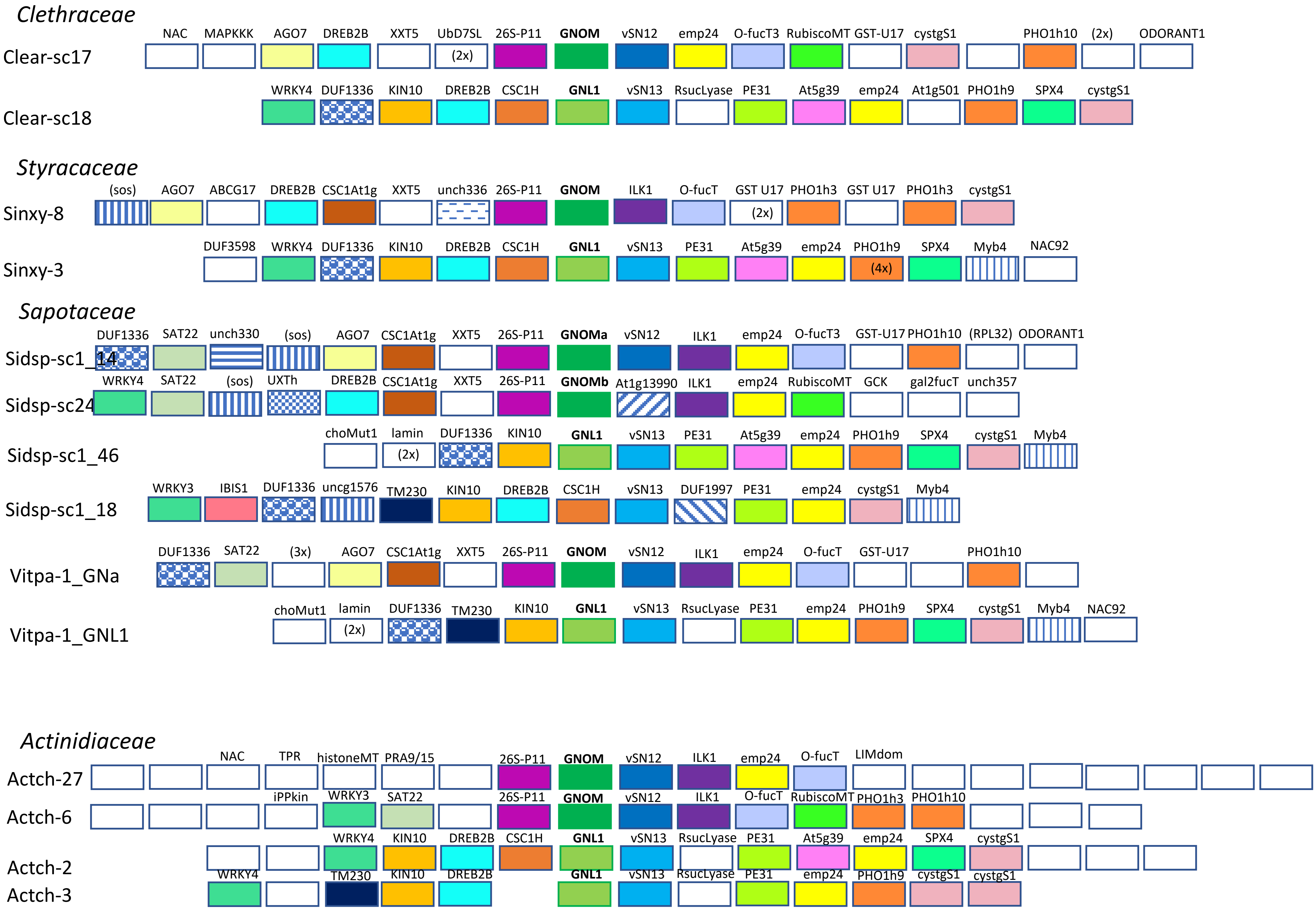

Clethraceae
Clear-sc17
NAC MAPKKK AGO7 DREB2B XXT5 UbD7SL (2x) 26S-P11 GNOM vSN12 emp24 O-fucT3 RubiscoMT GST-U17 cystgS1 PHO1h10 (2x) ODORANT1
Clear-sc18
WRKY4 DUF1336 KIN10 DREB2B CSC1H GNL1 vSN13 RsucLyase PE31 At5g39 emp24 At1g501 PHO1h9 SPX4 cystgS1
Styracaceae
Sinxy-8
(sos) AGO7 ABCG17 DREB2B CSC1At1g XXT5 unch336 26S-P11 GNOM ILK1 O-fucT GST U17 (2x) PHO1h3 GST U17 PHO1h3 cystgS1
Sinxy-3
DUF3598 WRKY4 DUF1336 KIN10 DREB2B CSC1H GNL1 vSN13 PE31 At5g39 emp24 PHO1h9 (4x) SPX4 Myb4 NAC92
Sapotaceae
Sidsp-sc1_14
DUF1336 SAT22 unch330 (sos) AGO7 CSC1At1g XXT5 26S-P11 GNOMa vSN12 ILK1 emp24 O-fucT3 GST-U17 PHO1h10 (RPL32) ODORANT1
Sidsp-sc24
WRKY4 SAT22 (sos) UXTh DREB2B CSC1At1g XXT5 26S-P11 GNOMb At1g13990 ILK1 emp24 RubiscoMT GCK gal2fucT unch357
Sidsp-sc1_46
choMut1 lamin (2x) DUF1336 KIN10 GNL1 vSN13 PE31 At5g39 emp24 PHO1h9 SPX4 cystgS1 Myb4
Sidsp-sc1_18
WRKY3 IBIS1 DUF1336 uncg1576 TM230 KIN10 DREB2B CSC1H vSN13 DUF1997 PE31 emp24 cystgS1 Myb4
Vitpa-1_GNa
DUF1336 SAT22 (3x) AGO7 CSC1At1g XXT5 26S-P11 GNOM vSN12 ILK1 emp24 O-fucT GST-U17 PHO1h10
Vitpa-1_GNL1
choMut1 lamin (2x) DUF1336 TM230 KIN10 GNL1 vSN13 RsucLyase PE31 emp24 PHO1h9 SPX4 cystgS1 Myb4 NAC92
Actinidiaceae
Actch-27
NAC TPR histoneMT PRA9/15 26S-P11 GNOM vSN12 ILK1 emp24 O-fucT LIMdom
Actch-6
iPPkin WRKY3 SAT22 26S-P11 GNOM vSN12 ILK1 O-fucT RubiscoMT PHO1h3 PHO1h10
Actch-2
WRKY4 KIN10 DREB2B CSC1H GNL1 vSN13 RsucLyase PE31 At5g39 emp24 SPX4 cystgS1
Actch-3
WRKY4 TM230 KIN10 DREB2B GNL1 vSN13 RsucLyase PE31 emp24 PHO1h9 cystgS1 cystgS1

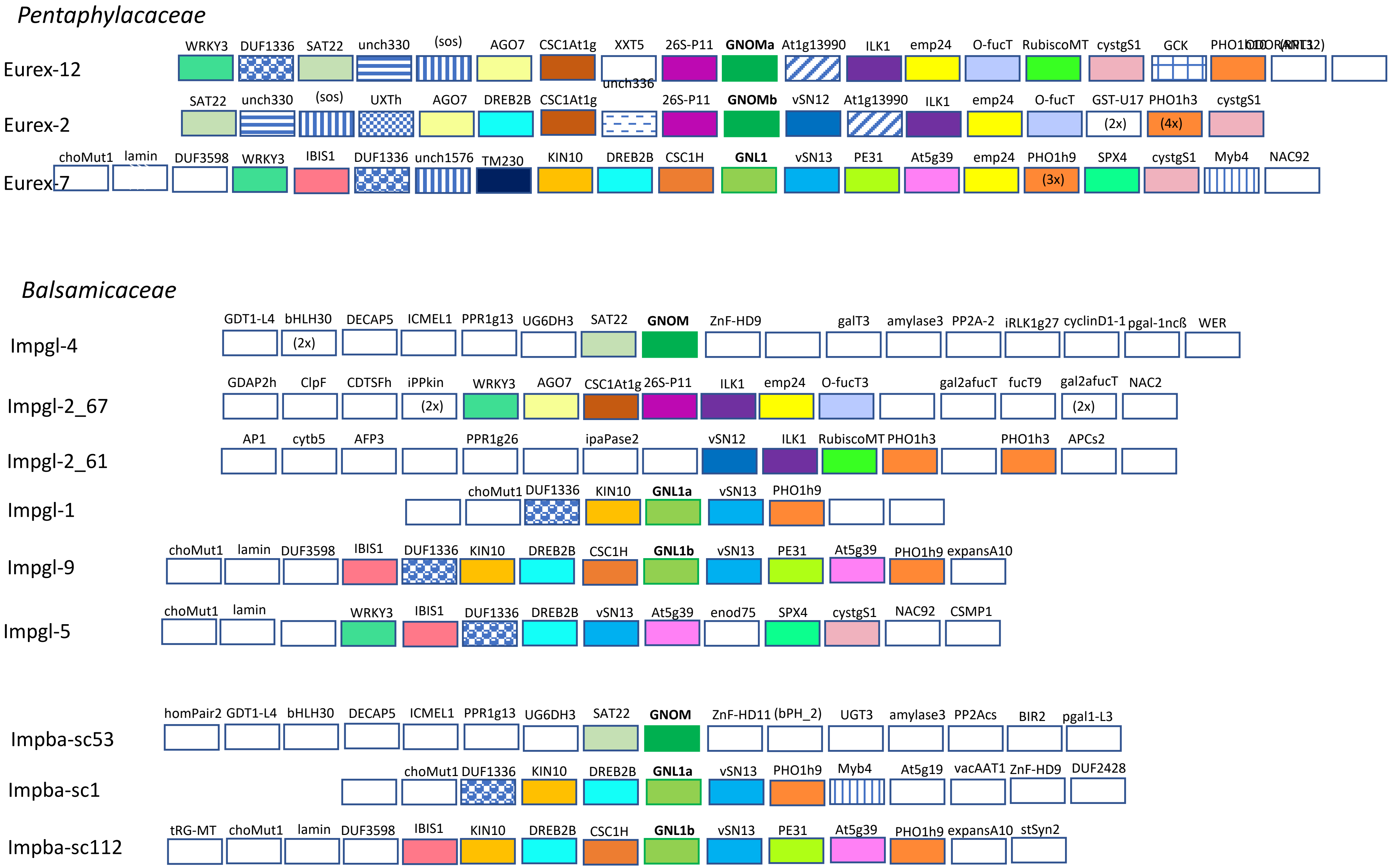
Pentaphylacaceae
Eurex-12
WRKY3 DUF1336 SAT22 unch330 (sos) AGO7 CSC1At1g XXT5 unch336 26S-P11 GNOMa At1g13990 ILK1 emp24 O-fucT RubiscoMT cystgS1 GCK PHO1h10 ODORANT32 (RRT1)
Eurex-2
SAT22 unch330 (sos) UXTh AGO7 DREB2B CSC1At1g 26S-P11 GNOMb vSN12 At1g13990 ILK1 emp24 O-fucT GST-U17 PHO1h3 cystgS1
(2x) (4x)
Eurex-7
choMut1 lamin DUF3598 WRKY3 IBIS1 DUF1336 unch1576 TM230 KIN10 DREB2B CSC1H GNL1 vSN13 PE31 At5g39 emp24 PHO1h9 SPX4 cystgS1 Myb4 NAC92
(3x)
Balsamicaceae
Impgl-4
GDT1-L4 bHLH30 DECAP5 ICMEL1 PPR1g13 UG6DH3 SAT22 GNOM ZnF-HD9 galT3 amylase3 PP2A-2 iRLK1g27 cyclinD1-1 pgal-1ncß WER
(2x)
Impgl-2_67
GDAP2h ClpF CDTSFh iPPkin WRKY3 AGO7 CSC1At1g 26S-P11 ILK1 emp24 O-fucT3 gal2afucT fucT9 gal2afucT NAC2
(2x) (2x)
Impgl-2_61
AP1 cytb5 AFP3 PPR1g26 ipaPase2 vSN12 ILK1 RubiscoMT PHO1h3 PHO1h3 APCs2
Impgl-1
choMut1 DUF1336 KIN10 GNL1a vSN13 PHO1h9
Impgl-9
choMut1 lamin DUF3598 IBIS1 DUF1336 KIN10 DREB2B CSC1H GNL1b vSN13 PE31 At5g39 PHO1h9 expansA10
Impgl-5
choMut1 lamin WRKY3 IBIS1 DUF1336 DREB2B vSN13 At5g39 enod75 SPX4 cystgS1 NAC92 CSMP1
Impba-sc53
homPair2 GDT1-L4 bHLH30 DECAP5 ICMEL1 PPR1g13 UG6DH3 SAT22 GNOM ZnF-HD11 (bPH_2) UGT3 amylase3 PP2Acs BIR2 pgal1-L3
Impba-sc1
choMut1 DUF1336 KIN10 DREB2B GNL1a vSN13 PHO1h9 Myb4 At5g19 vacAAT1 ZnF-HD9 DUF2428
Impba-sc112
tRG-MT choMut1 lamin DUF3598 IBIS1 KIN10 DREB2B CSC1H GNL1b vSN13 PE31 At5g39 PHO1h9 expansA10 stSyn2

### *Primulaceae*

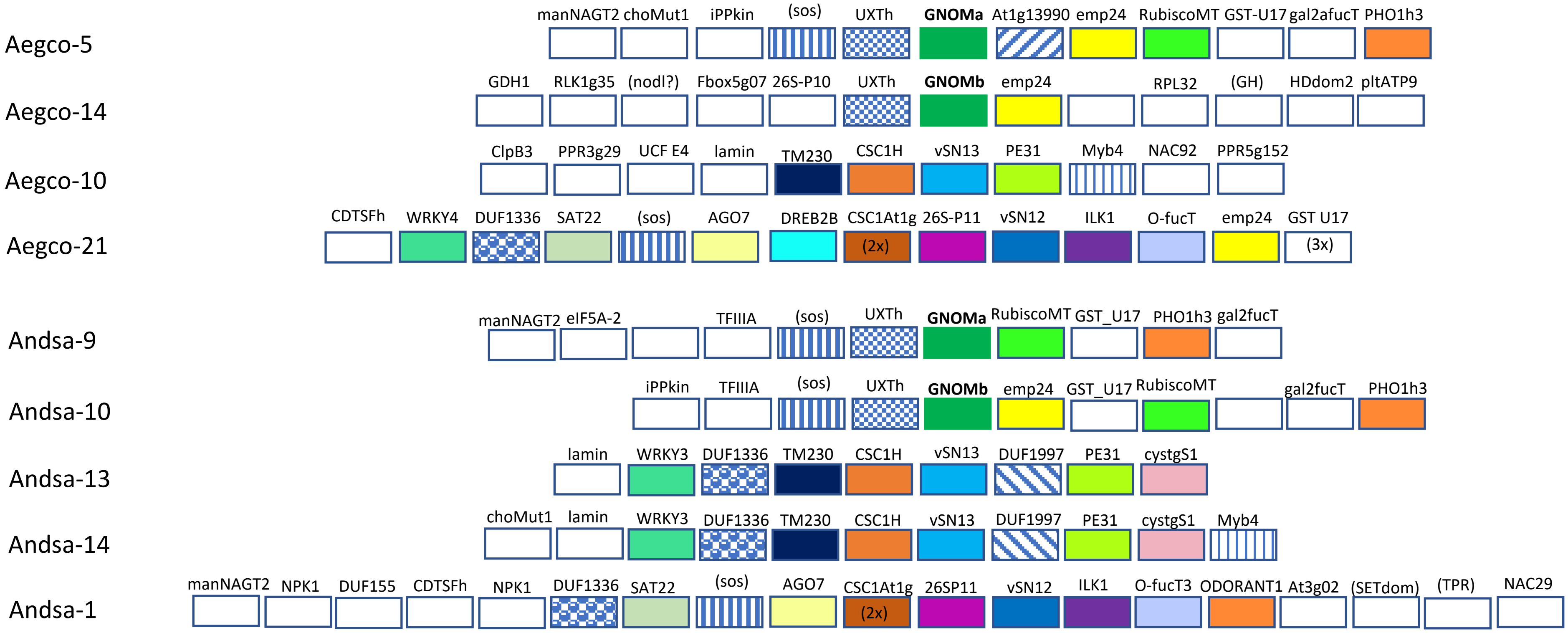

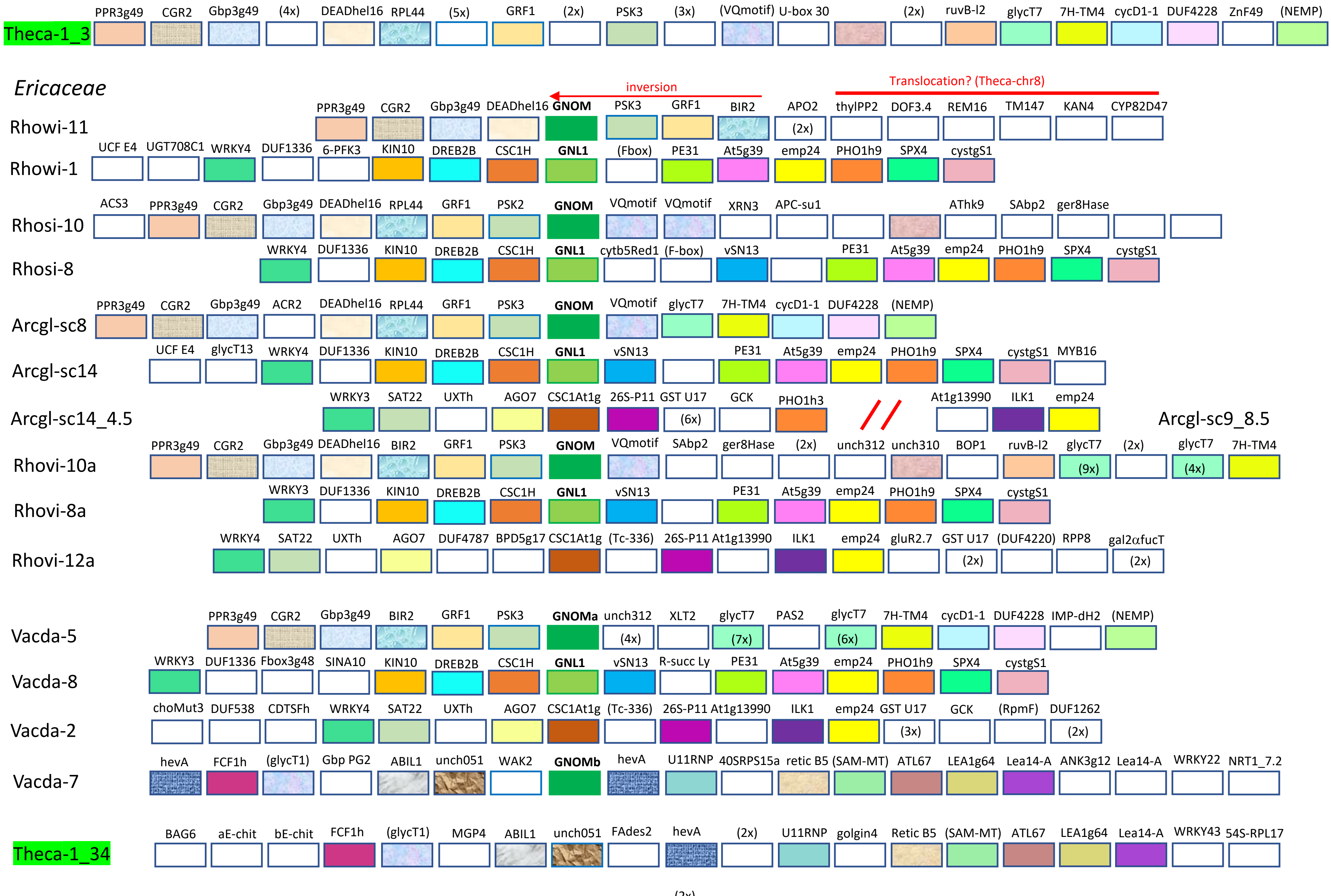

Theca-1_3
PPR3g49 CGR2 Gbp3g49 (4x) DEADhel16 RPL44 (5x) GRF1 (2x) PSK3 (3x) (VQmotif) U-box 30 (2x) ruvB-l2 glycT7 7H-TM4 cycD1-1 DUF4228 ZnF49 (NEMP)
Ericaceae
inversion
Translocation? (Theca-chr8)
Rhowi-11
PPR3g49 CGR2 Gbp3g49 DEADhel16 GNOM PSK3 GRF1 BIR2 APO2 (2x) thylPP2 DOF3.4 REM16 TM147 KAN4 CYP82D47
Rhowi-1
UCF E4 UGT708C1 WRKY4 DUF1336 6-PFK3 KIN10 DREB2B CSC1H GNL1 (Fbox) PE31 At5g39 emp24 PHO1h9 SPX4 cystgS1
Rhosi-10
ACS3 PPR3g49 CGR2 Gbp3g49 DEADhel16 RPL44 GRF1 PSK2 GNOM VQmotif VQmotif XRN3 APC-su1 AThk9 SAbp2 ger8Hase
Rhosi-8
WRKY4 DUF1336 KIN10 DREB2B CSC1H GNL1 cytb5Red1 (F-box) vSN13 PE31 At5g39 emp24 PHO1h9 SPX4 cystgS1
Arcgl-sc8
PPR3g49 CGR2 Gbp3g49 ACR2 DEADhel16 RPL44 GRF1 PSK3 GNOM VQmotif glycT7 7H-TM4 cycD1-1 DUF4228 (NEMP)
Arcgl-sc14
UCF E4 glycT13 WRKY4 DUF1336 KIN10 DREB2B CSC1H GNL1 vSN13 PE31 At5g39 emp24 PHO1h9 SPX4 cystgS1 MYB16
Arcgl-sc14_4.5
WRKY3 SAT22 UXTh AGO7 CSC1At1g 26S-P11 GST U17 (6x) GCK PHO1h3
At1g13990 ILK1 emp24
Arcgl-sc9_8.5
Rhovi-10a
PPR3g49 CGR2 Gbp3g49 DEADhel16 BIR2 GRF1 PSK3 GNOM VQmotif SAbp2 ger8Hase (2x) unch312 unch310 BOP1 ruvB-l2 glycT7 (9x) (2x) glycT7 (4x) 7H-TM4
Rhovi-8a
WRKY3 DUF1336 KIN10 DREB2B CSC1H GNL1 vSN13 PE31 At5g39 emp24 PHO1h9 SPX4 cystgS1
Rhovi-12a
WRKY4 SAT22 UXTh AGO7 DUF4787 BPD5g17 CSC1At1g (Tc-336) 26S-P11 At1g13990 ILK1 emp24 gluR2.7 GST U17 (2x) (DUF4220) RPP8 gal2αfucT (2x)
Vacda-5
PPR3g49 CGR2 Gbp3g49 BIR2 GRF1 PSK3 GNOMa unch312 (4x) XLT2 glycT7 (7x) PAS2 glycT7 (6x) 7H-TM4 cycD1-1 DUF4228 IMP-dH2 (NEMP)
Vacda-8
WRKY3 DUF1336 Fbox3g48 SINA10 KIN10 DREB2B CSC1H GNL1 vSN13 R-succ Ly PE31 At5g39 emp24 PHO1h9 SPX4 cystgS1
Vacda-2
choMut3 DUF538 CDTSFh WRKY4 SAT22 UXTh AGO7 CSC1At1g (Tc-336) 26S-P11 At1g13990 ILK1 emp24 GST U17 (3x) GCK (RpmF) DUF1262 (2x)
Vacda-7
hevA FCF1h (glycT1) Gbp PG2 ABIL1 unch051 WAK2 GNOMb hevA U11RNP 40SRPS15a retic B5 (SAM-MT) ATL67 LEA1g64 Lea14-A ANK3g12 Lea14-A WRKY22 NRT1_7.2
Theca-1_34
BAG6 aE-chit bE-chit FCF1h (glycT1) MGP4 ABIL1 unch051 FAdes2 hevA (2x) U11RNP golgin4 Retic B5 (SAM-MT) ATL67 LEA1g64 Lea14-A WRKY43 54S-RPL17

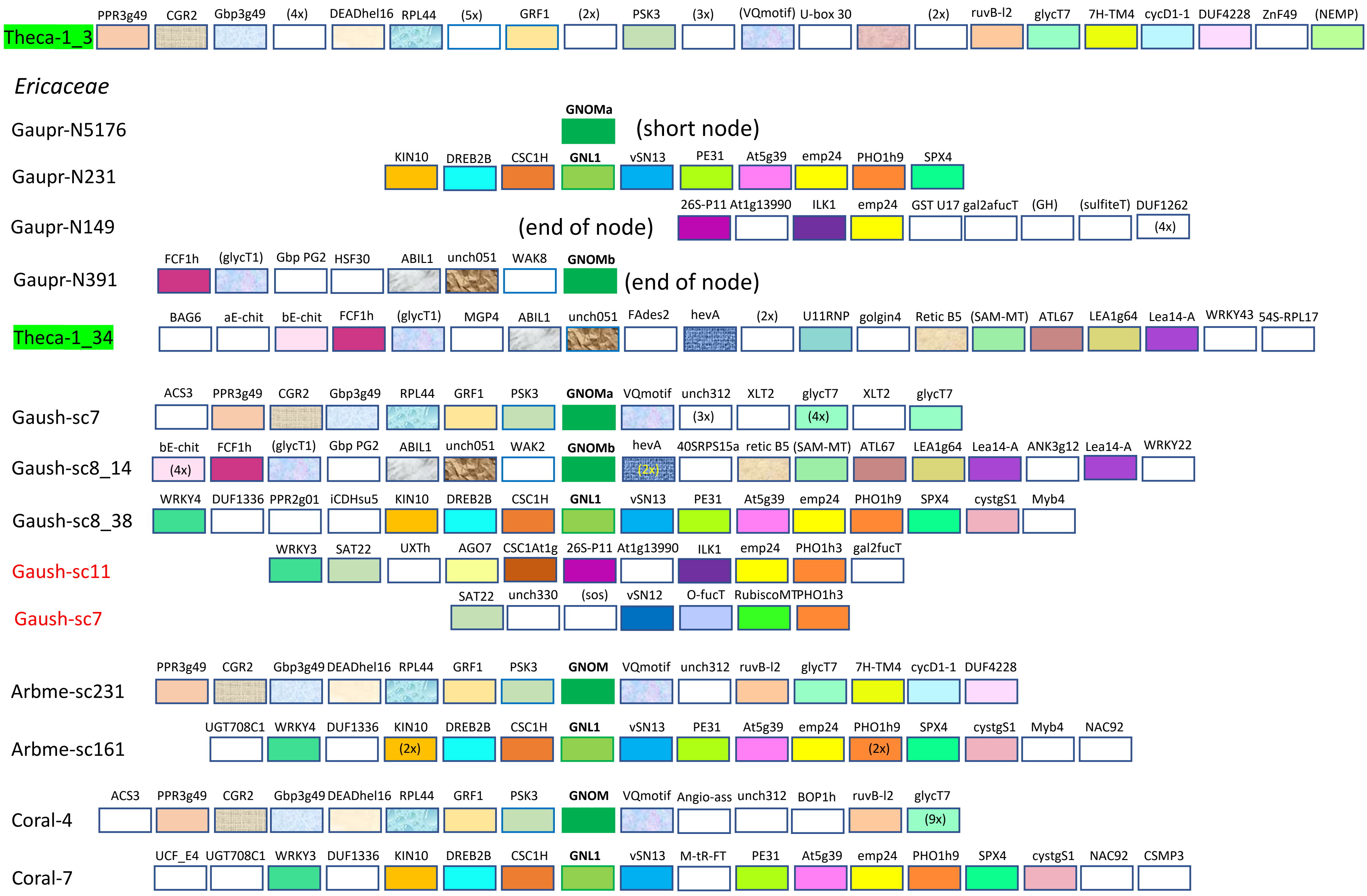
Theca-1_3
PPR3g49 CGR2 Gbp3g49 (4x) DEADhel16 RPL44 (5x) GRF1 (2x) PSK3 (3x) (VQmotif) U-box 30 (2x) ruvB-l2 glycT7 7H-TM4 cycD1-1 DUF4228 ZnF49 (NEMP)
Ericaceae
Gaupr-N5176
GNOMa
(short node)
Gaupr-N231
KIN10 DREB2B CSC1H GNL1 vSN13 PE31 At5g39 emp24 PHO1h9 SPX4
Gaupr-N149
(end of node)
26S-P11 At1g13990 ILK1 emp24 GST U17 gal2afucT (GH) (sulfiteT) DUF1262
(4x)
Gaupr-N391
FCF1h (glycT1) Gbp PG2 HSF30 ABIL1 unch051 WAK8 GNOMb
(end of node)
Theca-1_34
BAG6 aE-chit bE-chit FCF1h (glycT1) MGP4 ABIL1 unch051 FAdes2 hevA (2x) U11RNP golgin4 Retic B5 (SAM-MT) ATL67 LEA1g64 Lea14-A WRKY43 54S-RPL17
Gaush-sc7
ACS3 PPR3g49 CGR2 Gbp3g49 RPL44 GRF1 PSK3 GNOMa VQmotif unch312 XLT2 glycT7 XLT2 glycT7
(3x) (4x)
Gaush-sc8_14
bE-chit FCF1h (glycT1) Gbp PG2 ABIL1 unch051 WAK2 GNOMb hevA 40SRPS15a retic B5 (SAM-MT) ATL67 LEA1g64 Lea14-A ANK3g12 Lea14-A WRKY22
(4x) (2x)
Gaush-sc8_38
WRKY4 DUF1336 PPR2g01 iCDHsu5 KIN10 DREB2B CSC1H GNL1 vSN13 PE31 At5g39 emp24 PHO1h9 SPX4 cystgS1 Myb4
Gaush-sc11
WRKY3 SAT22 UXTh AGO7 CSC1At1g 26S-P11 At1g13990 ILK1 emp24 PHO1h3 gal2fucT
Gaush-sc7
SAT22 unch330 (sos) vSN12 O-fucT RubiscoMT PHO1h3
Arbme-sc231
PPR3g49 CGR2 Gbp3g49 DEADhel16 RPL44 GRF1 PSK3 GNOM VQmotif unch312 ruvB-l2 glycT7 7H-TM4 cycD1-1 DUF4228
Arbme-sc161
UGT708C1 WRKY4 DUF1336 KIN10 DREB2B CSC1H GNL1 vSN13 PE31 At5g39 emp24 PHO1h9 SPX4 cystgS1 Myb4 NAC92
(2x) (2x)
Coral-4
ACS3 PPR3g49 CGR2 Gbp3g49 DEADhel16 RPL44 GRF1 PSK3 GNOM VQmotif Angio-ass unch312 BOP1h ruvB-l2 glycT7
(9x)
Coral-7
UCF_E4 UGT708C1 WRKY3 DUF1336 KIN10 DREB2B CSC1H GNL1 vSN13 M-tR-FT PE31 At5g39 emp24 PHO1h9 SPX4 cystgS1 NAC92 CSMP3

#### Suppl. Figure S32. Cornales cladogram and OSGs

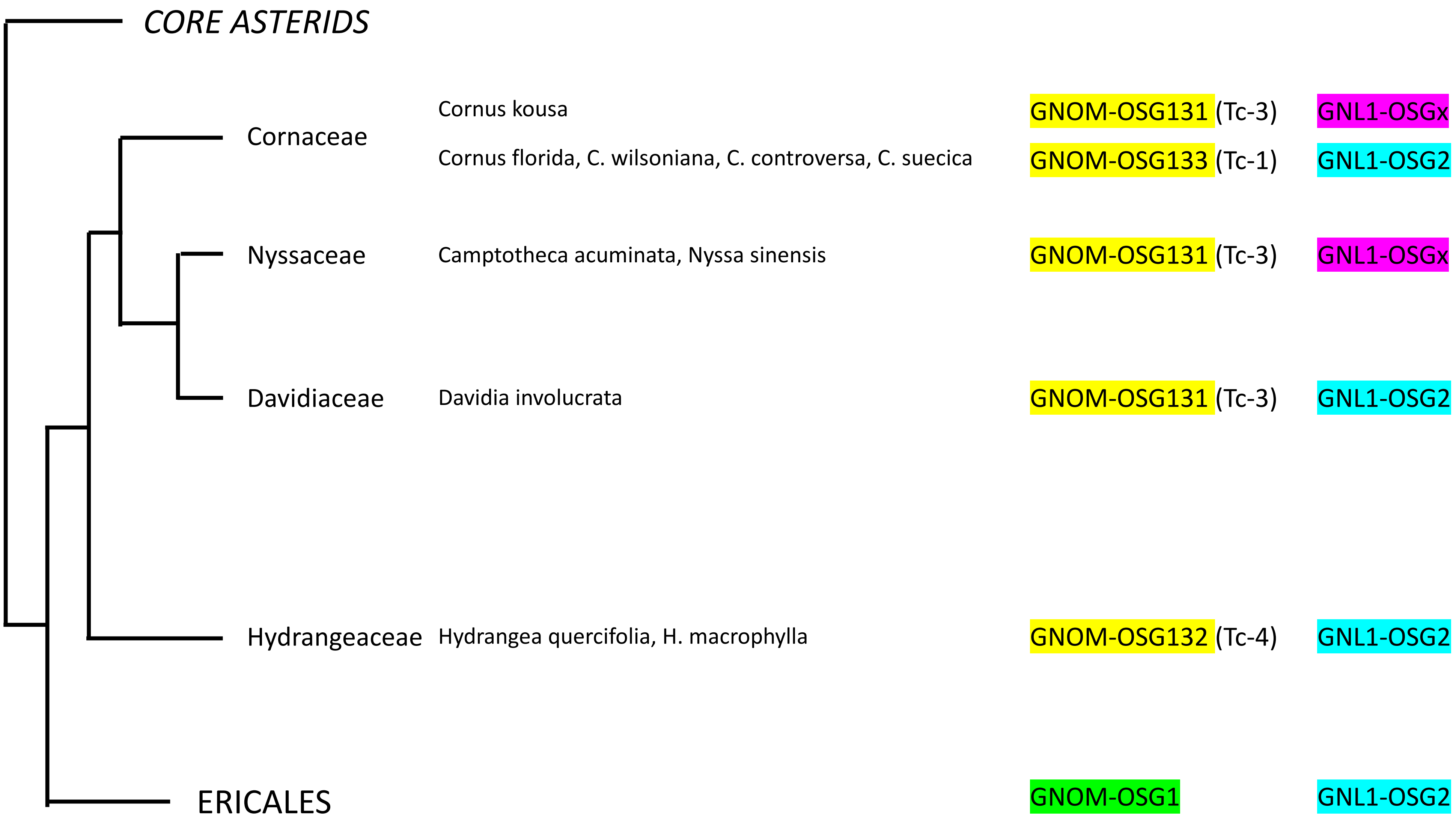

Cladogram based on Zhang et al. (2020c),; Thomas et al. (2021)

**Cornales synteny**

### CornusGNOM-OSG = Theca-chr1

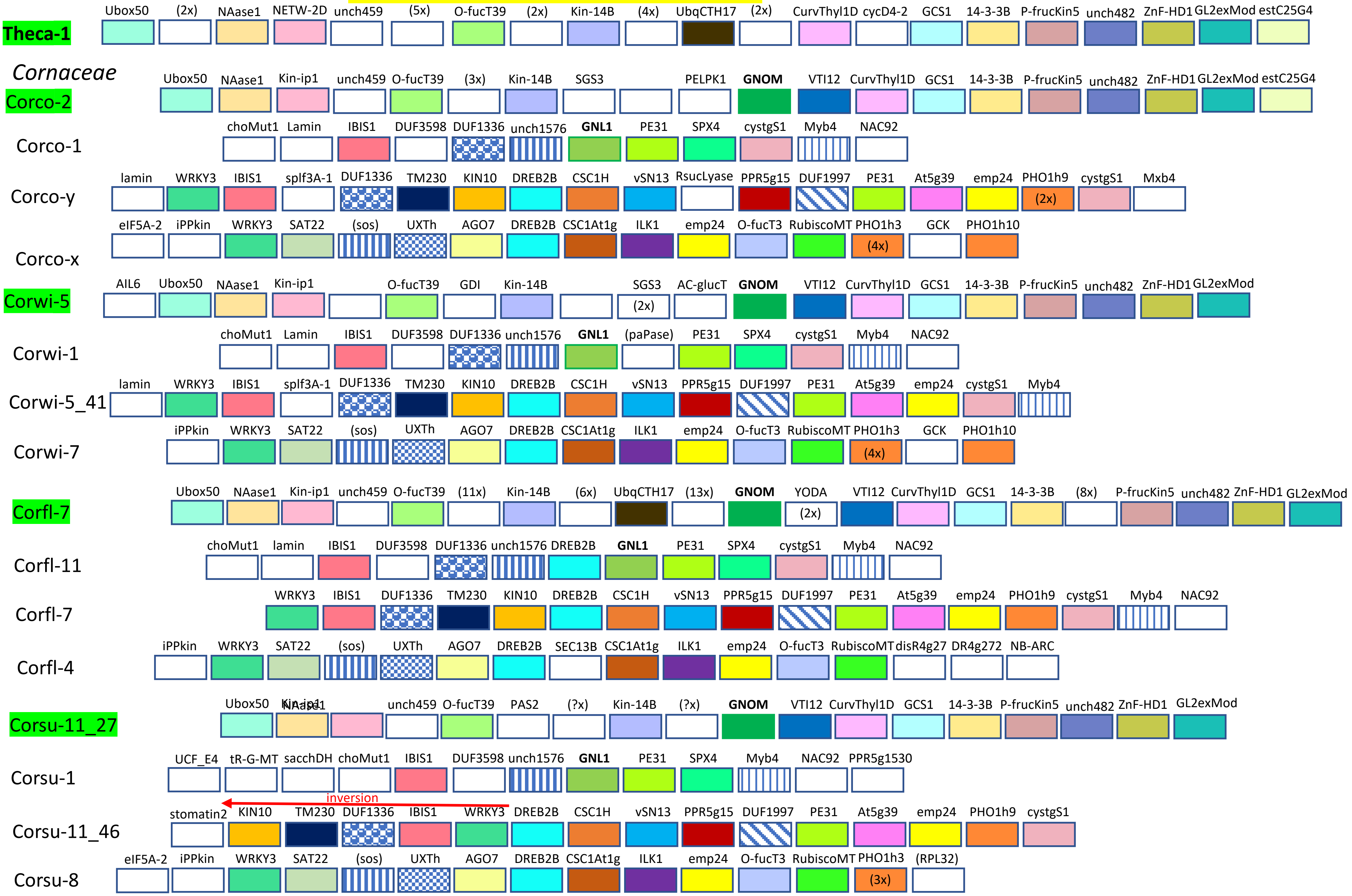

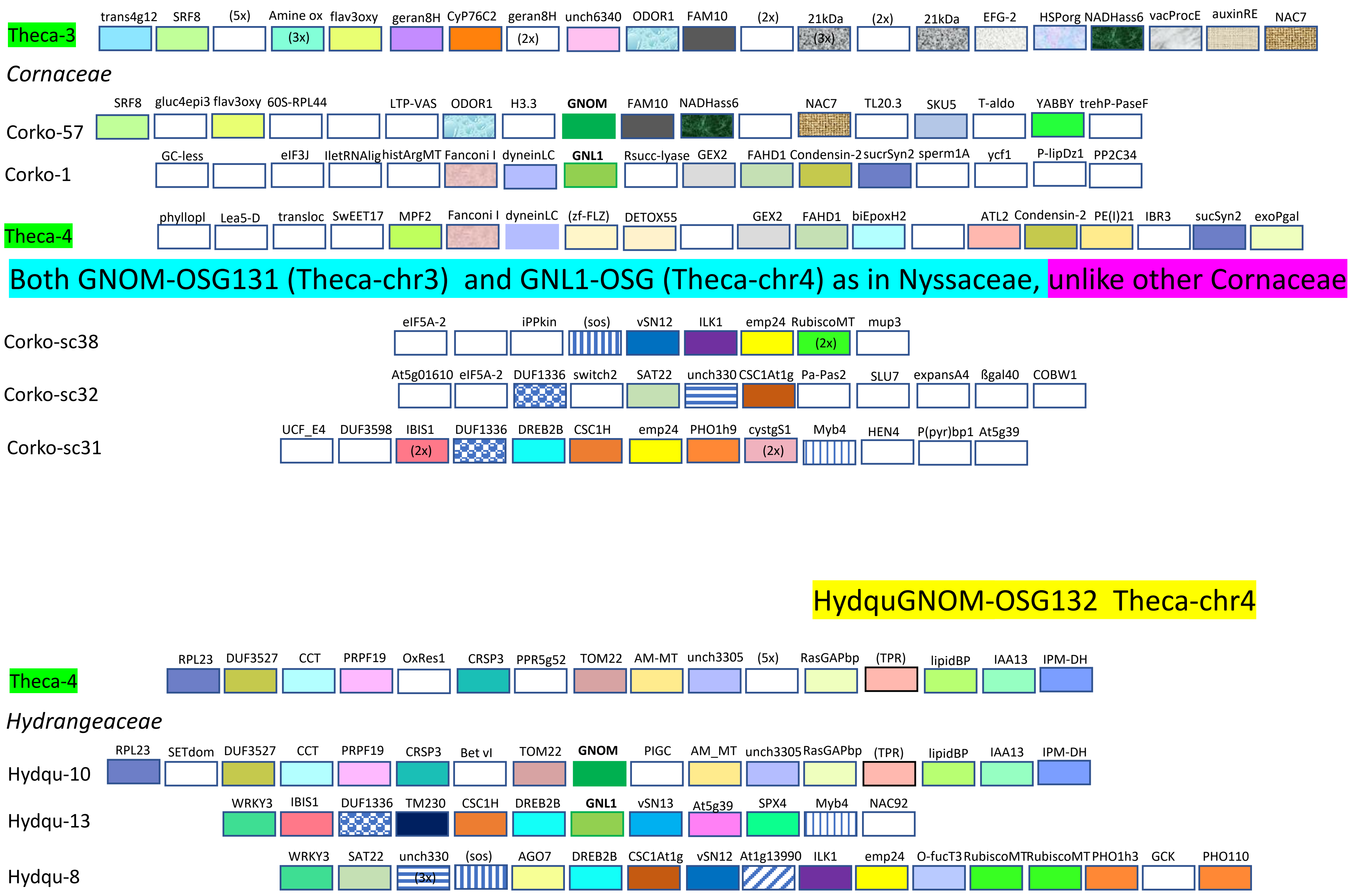
Theca-3
trans4g12 SRF8 (5x) Amine ox (3x) flav3oxy geran8H CyP76C2 geran8H (2x) unch6340 ODOR1 FAM10 (2x) 21kDa (3x) (2x) 21kDa EFG-2 HSPorg NADHass6 vacProcE auxinRE NAC7
Cornaceae
Corko-57
SRF8 gluc4epi3 flav3oxy 60S-RPL44 LTP-VAS ODOR1 H3.3 GNOM FAM10 NADHass6 NAC7 TL20.3 SKU5 T-aldo YABBY trehP-PaseF
Corko-1
GC-less eIF3J IletRNAlig histArgMT Fanconi I dyneinLC GNL1 Rsucc-lyase GEX2 FAHD1 Condensin-2 sucrSyn2 sperm1A ycf1 P-lipDz1 PP2C34
Theca-4
phyllopl Lea5-D transloc SwEET17 MPF2 Fanconi I dyneinLC (zf-FLZ) DETOX55 GEX2 FAHD1 biEpoxH2 ATL2 Condensin-2 PE(I)21 IBR3 sucSyn2 exoPgal
Both GNOM-OSG131 (Theca-chr3) and GNL1-OSG (Theca-chr4) as in Nyssaceae, unlike other Cornaceae
Corko-sc38
eIF5A-2 iPPkin (sos) vSN12 ILK1 emp24 RubiscoMT (2x) mup3
Corko-sc32
At5g01610 eIF5A-2 DUF1336 switch2 SAT22 unch330 CSC1At1g Pa-Pas2 SLU7 expansA4 ßgal40 COBW1
Corko-sc31
UCF_E4 DUF3598 IBIS1 (2x) DUF1336 DREB2B CSC1H emp24 PHO1h9 cystgS1 (2x) Myb4 HEN4 P(pyr)bp1 At5g39
HydquGNOM-OSG132 Theca-chr4
Theca-4
RPL23 DUF3527 CCT PRPF19 OxRes1 CRSP3 PPR5g52 TOM22 AM-MT unch3305 (5x) RasGAPbp (TPR) lipidBP IAA13 IPM-DH
Hydrangeaceae
Hydqu-10
RPL23 SETdom DUF3527 CCT PRPF19 CRSP3 Bet vI TOM22 GNOM PIGC AM_MT unch3305 RasGAPbp (TPR) lipidBP IAA13 IPM-DH
Hydqu-13
WRKY3 IBIS1 DUF1336 TM230 CSC1H DREB2B GNL1 vSN13 At5g39 SPX4 Myb4 NAC92
Hydqu-8
WRKY3 SAT22 unch330 (3x) (sos) AGO7 DREB2B CSC1At1g vSN12 At1g13990 ILK1 emp24 O-fucT3 RubiscoMT RubiscoMT PHO1h3 GCK PHO110

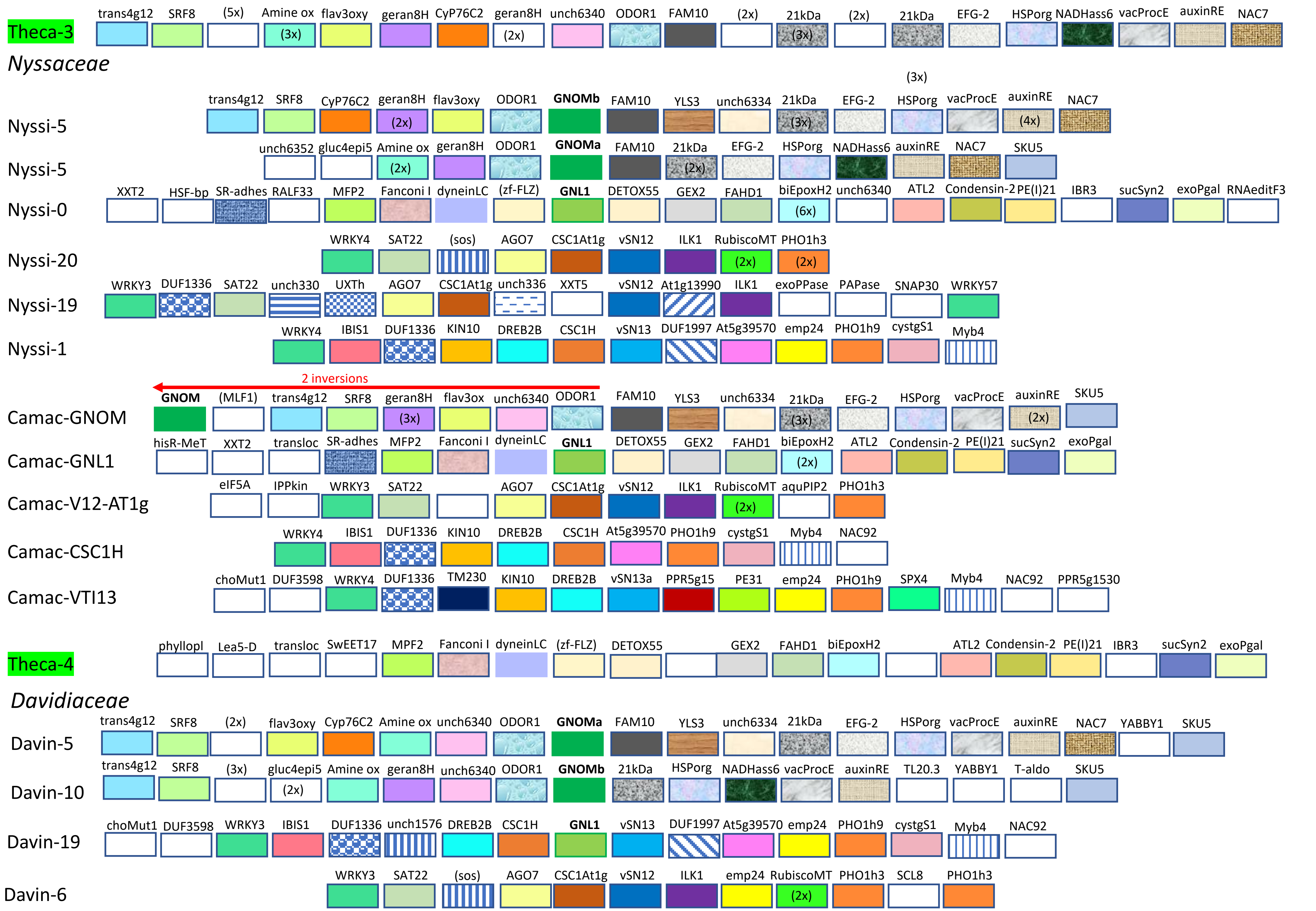

Theca-3
trans4g12 SRF8 (5x) Amine ox (3x) flav3oxy geran8H CyP76C2 geran8H (2x) unch6340 ODOR1 FAM10 (2x) 21kDa (3x) (2x) 21kDa EFG-2 HSPorg NADHass6 vacProcE auxinRE NAC7
Nyssaceae
Nyssi-5
trans4g12 SRF8 CyP76C2 geran8H (2x) flav3oxy ODOR1 GNOMb FAM10 YLS3 unch6334 21kDa (3x) EFG-2 (3x) HSPorg vacProcE auxinRE (4x) NAC7
Nyssi-5
unch6352 gluc4epi5 Amine ox (2x) geran8H ODOR1 GNOMa FAM10 21kDa (2x) EFG-2 HSPorg NADHass6 auxinRE NAC7 SKU5
Nyssi-0
XXT2 HSF-bp SR-adhes RALF33 MFP2 Fanconi I dyneinLC (zf-FLZ) GNL1 DETOX55 GEX2 FAHD1 biEpoxH2 (6x) unch6340 ATL2 Condensin-2 PE(I)21 IBR3 sucSyn2 exoPgal RNAeditF3
Nyssi-20
WRKY4 SAT22 (sos) AGO7 CSC1At1g vSN12 ILK1 RubiscoMT (2x) PHO1h3 (2x)
Nyssi-19
WRKY3 DUF1336 SAT22 unch330 UXTh AGO7 CSC1At1g unch336 XXT5 vSN12 At1g13990 ILK1 exoPPase PAPase SNAP30 WRKY57
Nyssi-1
WRKY4 IBIS1 DUF1336 KIN10 DREB2B CSC1H vSN13 DUF1997 At5g39570 emp24 PHO1h9 cystgS1 Myb4
2 inversions
Camac-GNOM
GNOM (MLF1) trans4g12 SRF8 geran8H (3x) flav3ox unch6340 ODOR1 FAM10 YLS3 unch6334 21kDa (3x) EFG-2 HSPorg vacProcE auxinRE (2x) SKU5
Camac-GNL1
hisR-MeT XXT2 transloc SR-adhes MFP2 Fanconi I dyneinLC GNL1 DETOX55 GEX2 FAHD1 biEpoxH2 (2x) ATL2 Condensin-2 PE(I)21 sucSyn2 exoPgal
Camac-V12-AT1g
eIF5A IPPkin WRKY3 SAT22 AGO7 CSC1At1g vSN12 ILK1 RubiscoMT (2x) aquPIP2 PHO1h3
Camac-CSC1H
WRKY4 IBIS1 DUF1336 KIN10 DREB2B CSC1H At5g39570 PHO1h9 cystgS1 Myb4 NAC92
Camac-VTI13
choMut1 DUF3598 WRKY4 DUF1336 TM230 KIN10 DREB2B vSN13a PPR5g15 PE31 emp24 PHO1h9 SPX4 Myb4 NAC92 PPR5g1530
Theca-4
phyllopl Lea5-D transloc SwEET17 MPF2 Fanconi I dyneinLC (zf-FLZ) DETOX55 GEX2 FAHD1 biEpoxH2 ATL2 Condensin-2 PE(I)21 IBR3 sucSyn2 exoPgal
Davidiaceae
Davin-5
trans4g12 SRF8 (2x) flav3oxy Cyp76C2 Amine ox unch6340 ODOR1 GNOMa FAM10 YLS3 unch6334 21kDa EFG-2 HSPorg vacProcE auxinRE NAC7 YABBY1 SKU5
Davin-10
trans4g12 SRF8 (3x) gluc4epi5 (2x) Amine ox geran8H unch6340 ODOR1 GNOMb 21kDa HSPorg NADHass6 vacProcE auxinRE TL20.3 YABBY1 T-aldo SKU5
Davin-19
choMut1 DUF3598 WRKY3 IBIS1 DUF1336 unch1576 DREB2B CSC1H GNL1 vSN13 DUF1997 At5g39570 emp24 PHO1h9 cystgS1 Myb4 NAC92
Davin-6
WRKY3 SAT22 (sos) AGO7 CSC1At1g vSN12 ILK1 emp24 RubiscoMT (2x) PHO1h3 SCL8 PHO1h3

### Suppl. Figure S33. Caryophyllales cladogram and OSGs

No GNL1

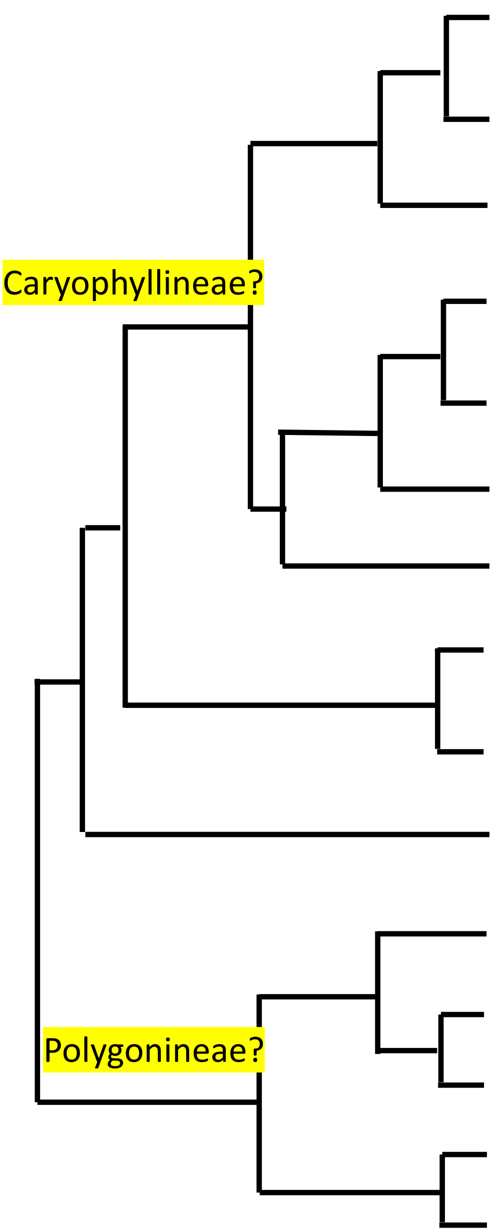

| | | |
|---|---|---|
| Nyctaginaceae | Bougainvillea glabra | OSG1 |
| Phytolaccaceae | Phytolacca americana | OSG1 |
| Aizoaceae | Tetragonia tetragonoides | OSG1 |
| Portulacaceae | Portulaca amilis, P. oleraceae | OSG1 |
| Cactaceae | Carnegiea gigantea, Opuntia basilaris | OSG1 |
| Talinaceae | Talinum fruticosum | OSG1 |
| Molluginaceae | Pharnaceum exiguum | OSG1 |
| Amaranthaceae | Amaranthus hypochondriacus, A. cruentus, Chenopodium quinoa, C. pallidicaule, Spinacia oleracea, Beta vulgaris, Suaeda aralocaspica, Haloxylon ammodendron, Salicornia ramosissima | OSG1 |
| Caryophyllaceae | Dianthus caryophyllus, Corrigiola litoralis, Heliosperma pusillum, Gypsophila paniculata, Saponaria officinalis, Spergularia bocconei | diverse **See below**) |
| Simmondsiaceae | Simmondsia chinensis | OSG1 |
| Tamaricaceae | Tamarix chinensis | OSG1 |
| Polygonaceae | Fagopyrum tartaricum, Fallopia convolvulus Polygonum aviculare | GNOMaOSG-Tc6 |
| Plumbaginaceae | Limonium bicolor | Not analysed |
| Nepenthaceae | Nepenthes mirabilis, N. gracilis | OSG1 |
| Droseraceae | Drosera capensis, D. spatulata | Not analysed (DrospGNOMa-OSG1) |

Cladogram based on OTPTI (2019); Li et al. (2021a)

#### Caryophyllaceae cladogram

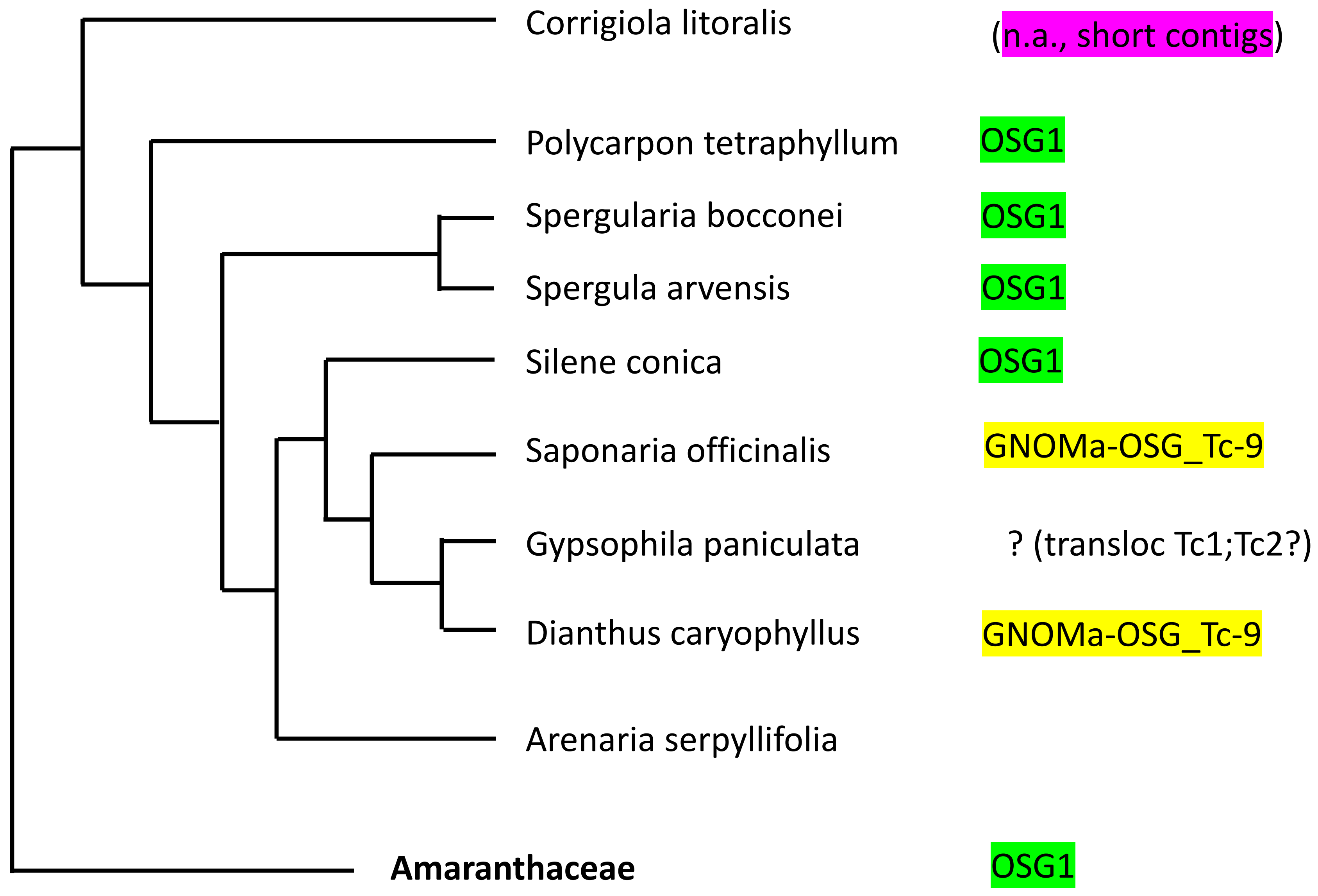

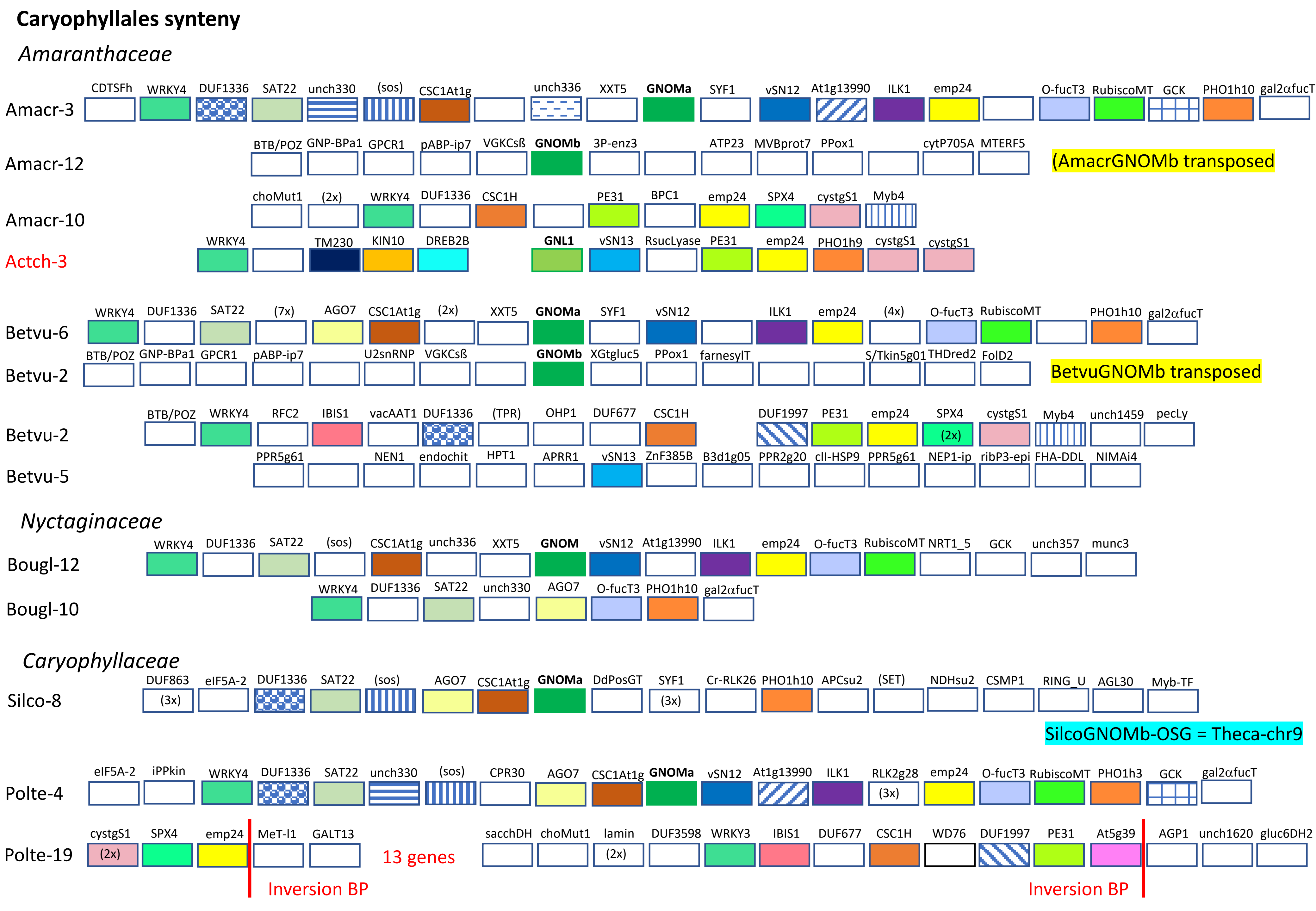
Caryophyllales synteny
Amaranthaceae
Amacr-3
CDTSFh WRKY4 DUF1336 SAT22 unch330 (sos) CSC1At1g unch336 XXT5 GNOMa SYF1 vSN12 At1g13990 ILK1 emp24 O-fucT3 RubiscoMT GCK PHO1h10 gal2αfucT
Amacr-12
BTB/POZ GNP-BPa1 GPCR1 pABP-ip7 VGKCsß GNOMb 3P-enz3 ATP23 MVBprot7 PPox1 cytP705A MTERF5
(AmacrGNOMb transposed
Amacr-10
choMut1 (2x) WRKY4 DUF1336 CSC1H PE31 BPC1 emp24 SPX4 cystgS1 Myb4
Actch-3
WRKY4 TM230 KIN10 DREB2B GNL1 vSN13 RsucLyase PE31 emp24 PHO1h9 cystgS1 cystgS1
Betvu-6
WRKY4 DUF1336 SAT22 (7x) AGO7 CSC1At1g (2x) XXT5 GNOMa SYF1 vSN12 ILK1 emp24 (4x) O-fucT3 RubiscoMT PHO1h10 gal2αfucT
Betvu-2
BTB/POZ GNP-BPa1 GPCR1 pABP-ip7 U2snRNP VGKCsß GNOMb XGtgluc5 PPox1 farnesylT S/Tkin5g01 THDred2 FolD2
BetvuGNOMb transposed
Betvu-2
BTB/POZ WRKY4 RFC2 IBIS1 vacAAT1 DUF1336 (TPR) OHP1 DUF677 CSC1H DUF1997 PE31 emp24 SPX4 (2x) cystgS1 Myb4 unch1459 pecLy
Betvu-5
PPR5g61 NEN1 endochit HPT1 APRR1 vSN13 ZnF385B B3d1g05 PPR2g20 cII-HSP9 PPR5g61 NEP1-ip ribP3-epi FHA-DDL NIMAi4
Nyctaginaceae
Bougl-12
WRKY4 DUF1336 SAT22 (sos) CSC1At1g unch336 XXT5 GNOM vSN12 At1g13990 ILK1 emp24 O-fucT3 RubiscoMT NRT1_5 GCK unch357 munc3
Bougl-10
WRKY4 DUF1336 SAT22 unch330 AGO7 O-fucT3 PHO1h10 gal2αfucT
Caryophyllaceae
Silco-8
DUF863 (3x) eIF5A-2 DUF1336 SAT22 (sos) AGO7 CSC1At1g GNOMa DdPosGT SYF1 (3x) Cr-RLK26 PHO1h10 APCsu2 (SET) NDHsu2 CSMP1 RING_U AGL30 Myb-TF
SilcoGNOMb-OSG = Theca-chr9
Polte-4
eIF5A-2 iPPkin WRKY4 DUF1336 SAT22 unch330 (sos) CPR30 AGO7 CSC1At1g GNOMa vSN12 At1g13990 ILK1 RLK2g28 (3x) emp24 O-fucT3 RubiscoMT PHO1h3 GCK gal2αfucT
Polte-19
cystgS1 (2x) SPX4 emp24 MeT-I1 GALT13 13 genes sacchDH choMut1 lamin (2x) DUF3598 WRKY3 IBIS1 DUF677 CSC1H WD76 DUF1997 PE31 At5g39 AGP1 unch1620 gluc6DH2
Inversion BP
Inversion BP

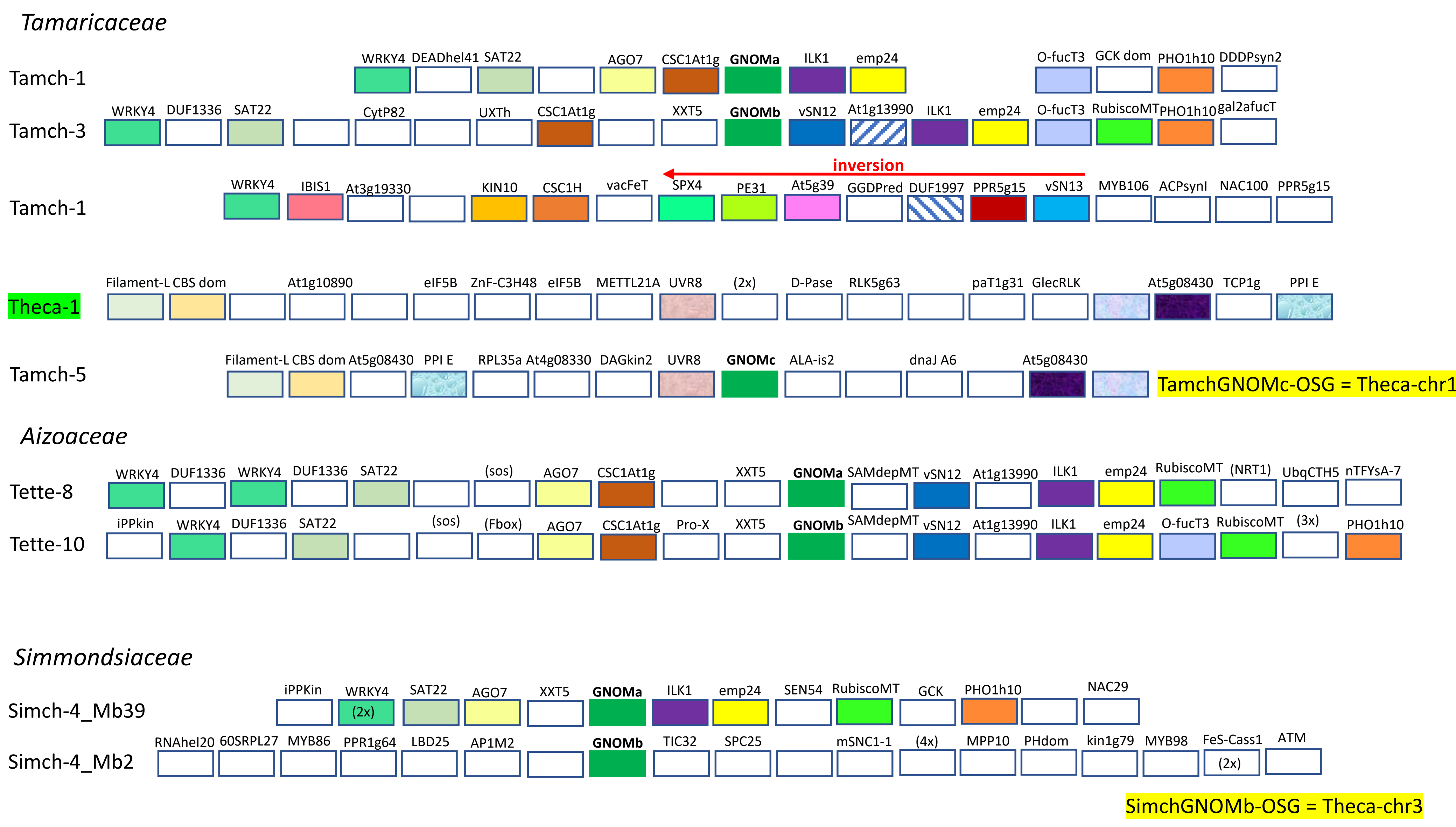
Tamaricaceae
Tamch-1
WRKY4 DEADhel41 SAT22 AGO7 CSC1At1g GNOMa ILK1 emp24 O-fucT3 GCK dom PHO1h10 DDDPsyn2
Tamch-3
WRKY4 DUF1336 SAT22 CytP82 UXTh CSC1At1g XXT5 GNOMb vSN12 At1g13990 ILK1 emp24 O-fucT3 RubiscoMT PHO1h10 gal2afucT
inversion
Tamch-1
WRKY4 IBIS1 At3g19330 KIN10 CSC1H vacFeT SPX4 PE31 At5g39 GGDPred DUF1997 PPR5g15 vSN13 MYB106 ACPsynI NAC100 PPR5g15
Theca-1
Filament-L CBS dom At1g10890 eIF5B ZnF-C3H48 eIF5B METTL21A UVR8 (2x) D-Pase RLK5g63 paT1g31 GlecRLK At5g08430 TCP1g PPI E
Tamch-5
Filament-L CBS dom At5g08430 PPI E RPL35a At4g08330 DAGkin2 UVR8 GNOMc ALA-is2 dnaJ A6 At5g08430
TamchGNOMc-OSG = Theca-chr1
Aizoaceae
Tette-8
WRKY4 DUF1336 WRKY4 DUF1336 SAT22 (sos) AGO7 CSC1At1g XXT5 GNOMa SAMdepMT vSN12 At1g13990 ILK1 emp24 RubiscoMT (NRT1) UbqCTH5 nTFYsA-7
Tette-10
iPPkin WRKY4 DUF1336 SAT22 (sos) (Fbox) AGO7 CSC1At1g Pro-X XXT5 GNOMb SAMdepMT vSN12 At1g13990 ILK1 emp24 O-fucT3 RubiscoMT (3x) PHO1h10
Simmondsiaceae
Simch-4_Mb39
iPPKin WRKY4 (2x) SAT22 AGO7 XXT5 GNOMa ILK1 emp24 SEN54 RubiscoMT GCK PHO1h10 NAC29
Simch-4_Mb2
RNAhel20 60SRPL27 MYB86 PPR1g64 LBD25 AP1M2 GNOMb TIC32 SPC25 mSNC1-1 (4x) MPP10 PHdom kin1g79 MYB98 FeS-Cass1 (2x) ATM
SimchGNOMb-OSG = Theca-chr3

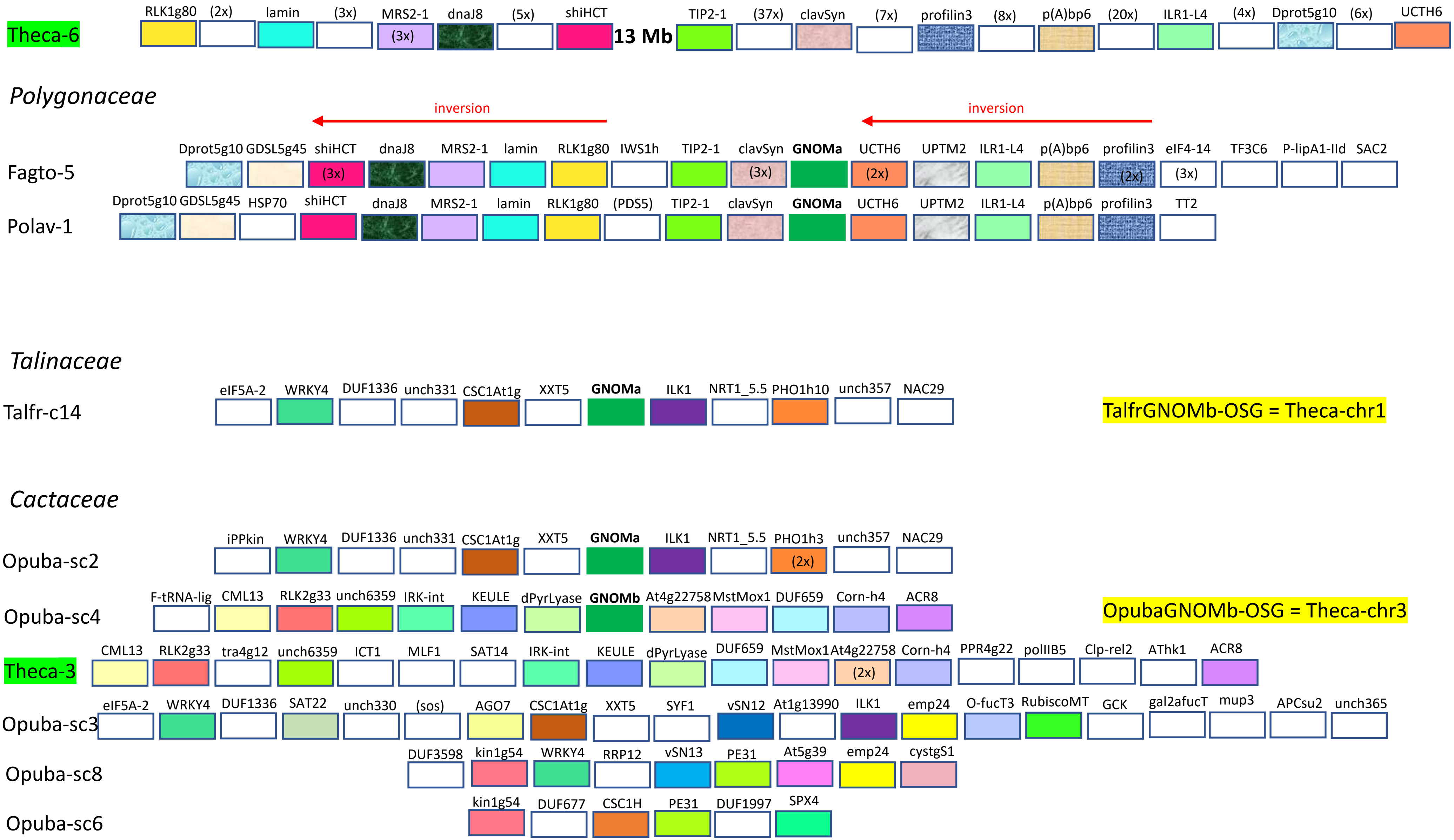

Theca-6
RLK1g80 (2x) lamin (3x) MRS2-1 (3x) dnaJ8 (5x) shiHCT 13 Mb TIP2-1 (37x) clavSyn (7x) profilin3 (8x) p(A)bp6 (20x) ILR1-L4 (4x) Dprot5g10 (6x) UCTH6
Polygonaceae
inversion
inversion
Fagto-5
Dprot5g10 GDSL5g45 shiHCT (3x) dnaJ8 MRS2-1 lamin RLK1g80 IWS1h TIP2-1 clavSyn (3x) GNOMa UCTH6 (2x) UPTM2 ILR1-L4 p(A)bp6 profilin3 (2x) eIF4-14 (3x) TF3C6 P-lipA1-IId SAC2
Polav-1
Dprot5g10 GDSL5g45 HSP70 shiHCT dnaJ8 MRS2-1 lamin RLK1g80 (PDS5) TIP2-1 clavSyn GNOMa UCTH6 UPTM2 ILR1-L4 p(A)bp6 profilin3 TT2
Talinaceae
Talfr-c14
eIF5A-2 WRKY4 DUF1336 unch331 CSC1At1g XXT5 GNOMa ILK1 NRT1_5.5 PHO1h10 unch357 NAC29
TalfrGNOMb-OSG = Theca-chr1
Cactaceae
Opuba-sc2
iPPkin WRKY4 DUF1336 unch331 CSC1At1g XXT5 GNOMa ILK1 NRT1_5.5 PHO1h3 (2x) unch357 NAC29
Opuba-sc4
F-tRNA-lig CML13 RLK2g33 unch6359 IRK-int KEULE dPyrLyase GNOMb At4g22758 MstMox1 DUF659 Corn-h4 ACR8
OpubaGNOMb-OSG = Theca-chr3
Theca-3
CML13 RLK2g33 tra4g12 unch6359 ICT1 MLF1 SAT14 IRK-int KEULE dPyrLyase DUF659 MstMox1 At4g22758 (2x) Corn-h4 PPR4g22 polIIB5 Clp-rel2 AThk1 ACR8
Opuba-sc3
eIF5A-2 WRKY4 DUF1336 SAT22 unch330 (sos) AGO7 CSC1At1g XXT5 SYF1 vSN12 At1g13990 ILK1 emp24 O-fucT3 RubiscoMT GCK gal2afucT mup3 APCsu2 unch365
Opuba-sc8
DUF3598 kin1g54 WRKY4 RRP12 vSN13 PE31 At5g39 emp24 cystgS1
Opuba-sc6
kin1g54 DUF677 CSC1H PE31 DUF1997 SPX4

### Suppl. Figure S34. Dilleniales cladogram and OSGs

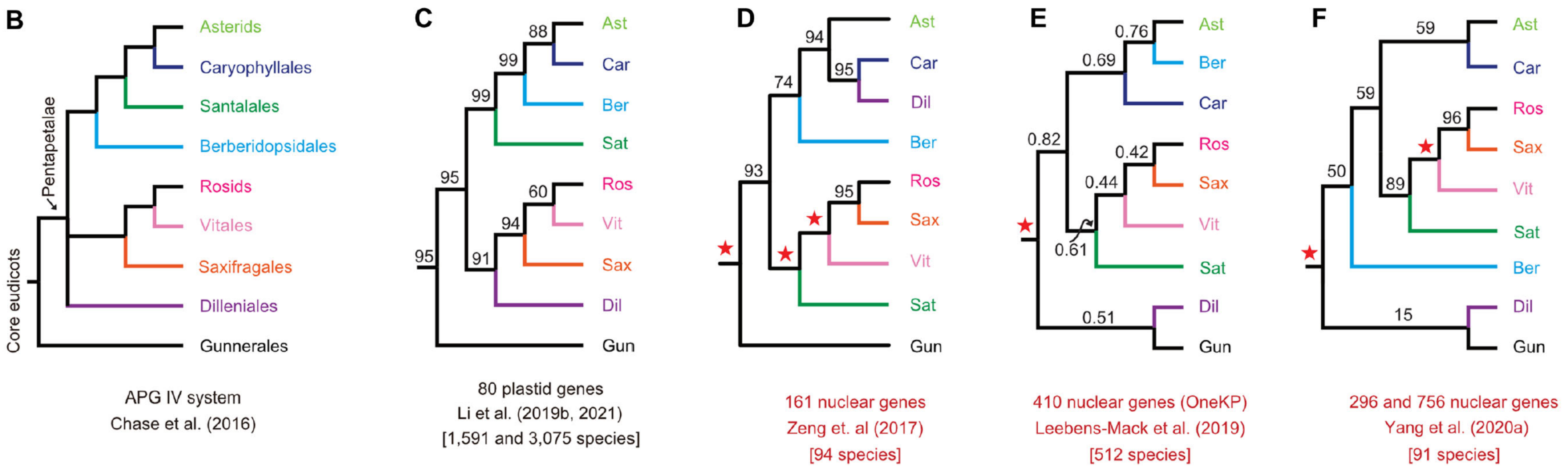

Phylogenetic position of Dilleniales uncertain. Alternative cladograms from Zhang and Ma (2024)

### OSG-17 Dilleniales (GNOMa-OSG171, GNOMb,c-OSG172)

#### Dillenia turbinata (sc8), Dillenia suffruticosa (sc22) GNOMa-OSG171 (Theca-chr7)

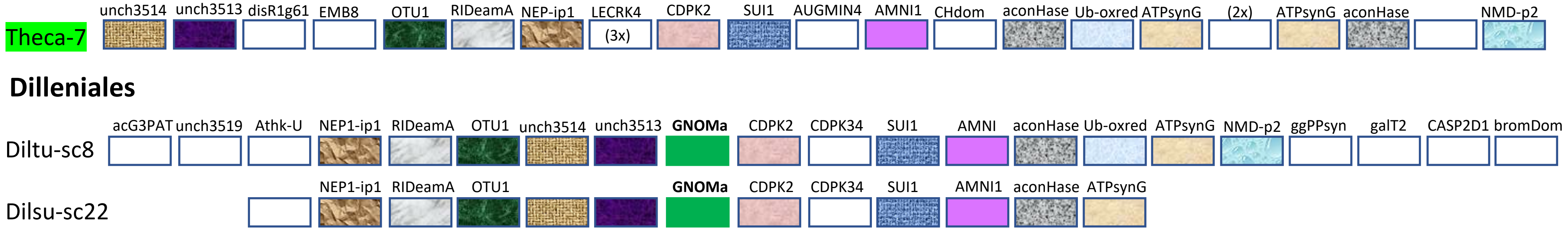

#### Dillenia turbinata (sc28, sc25), Dillenia suffruticosa (sc9, sc12) GNOMb,c-OSG171 (Theca-chr6)

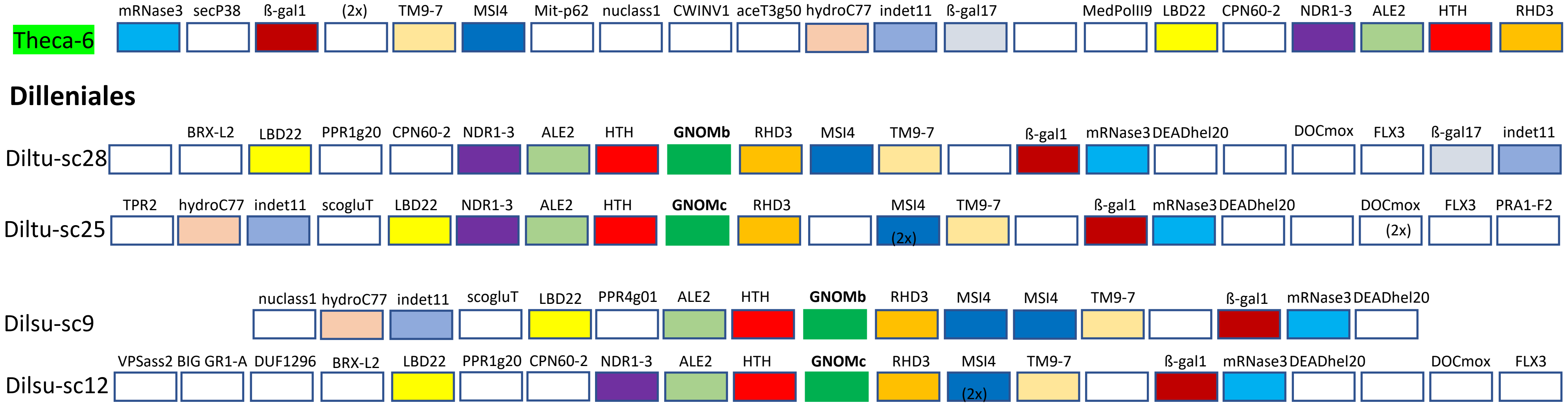

### Loss of GNOM from OSG1 in Dilleniales

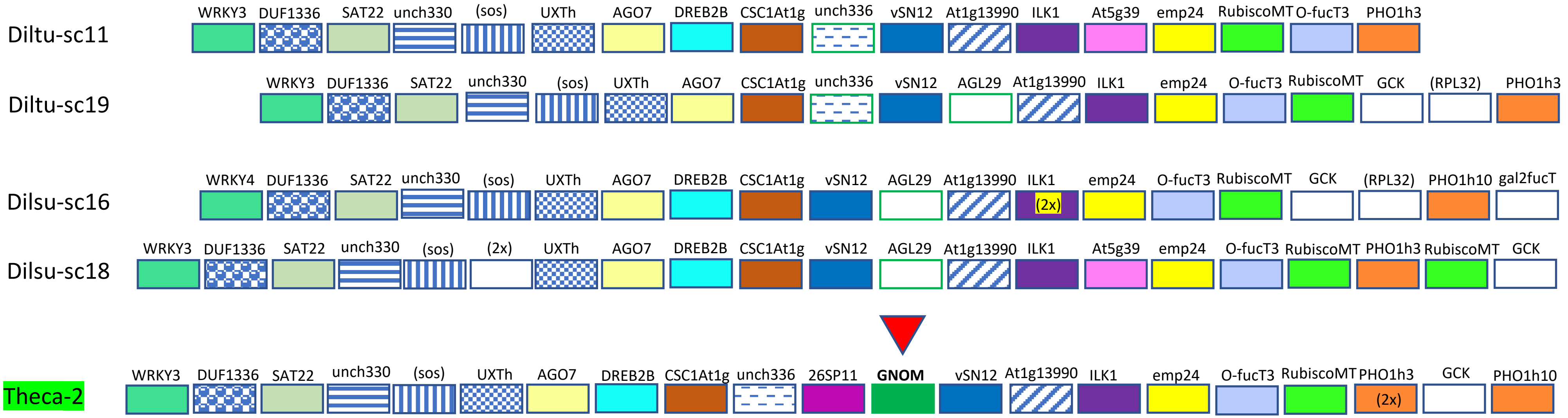

### Loss of GNL1 from OSG2 in Dilleniales

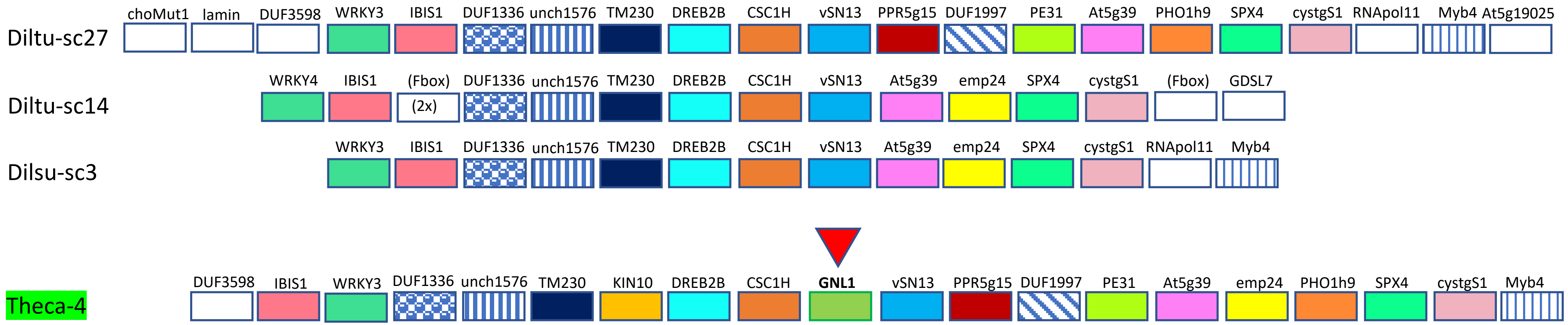

### Suppl. Figure S35. Icacinales cladogram and OSGs

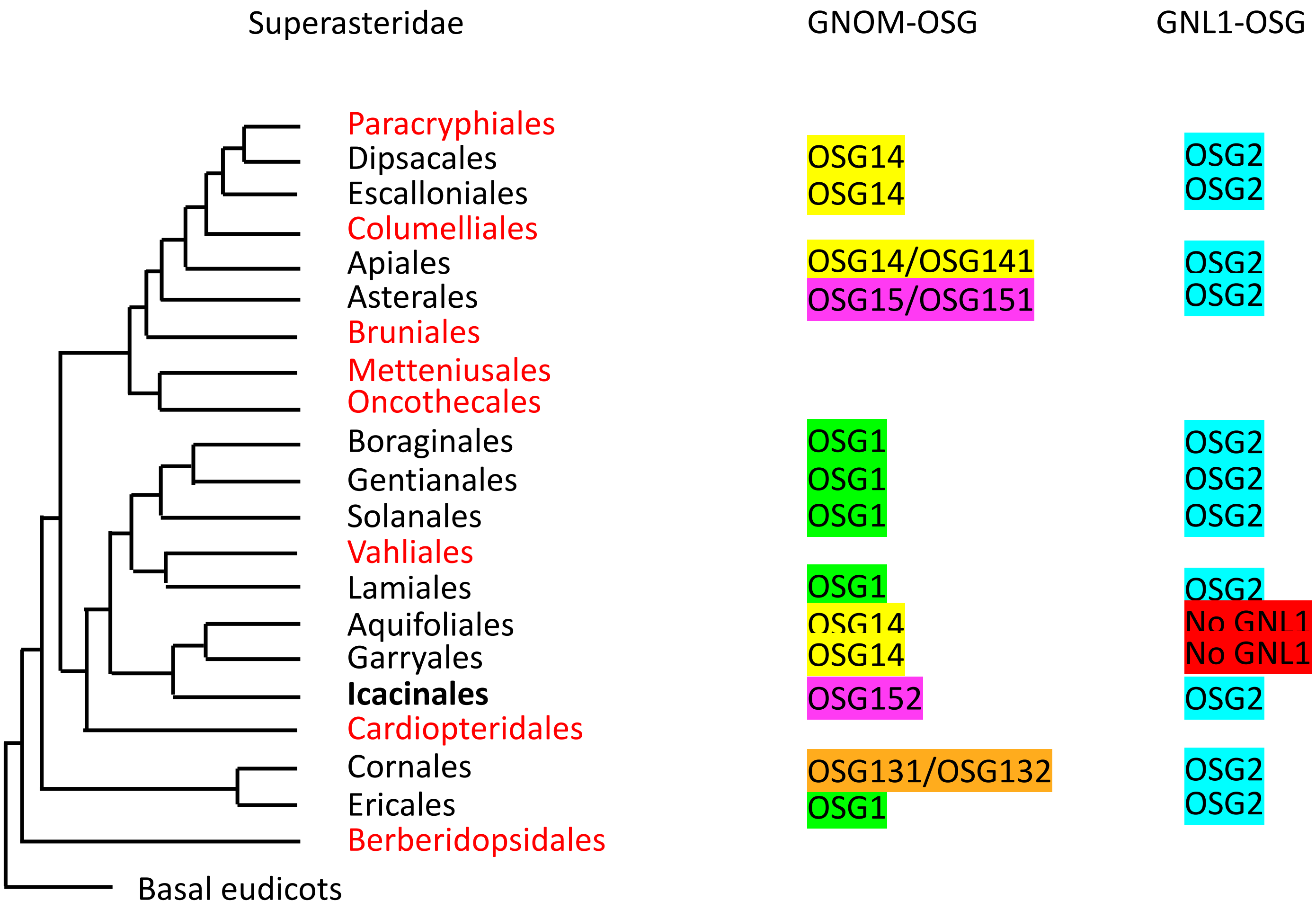

Note discrepancy between cladogram and shared GNOM transposition.
Orders in red not analysed. For genome sequence of *Iodes seguinii* (Icacinales), see Gong et al. (2025).

Cladogram based on Zuntini et al (2024)

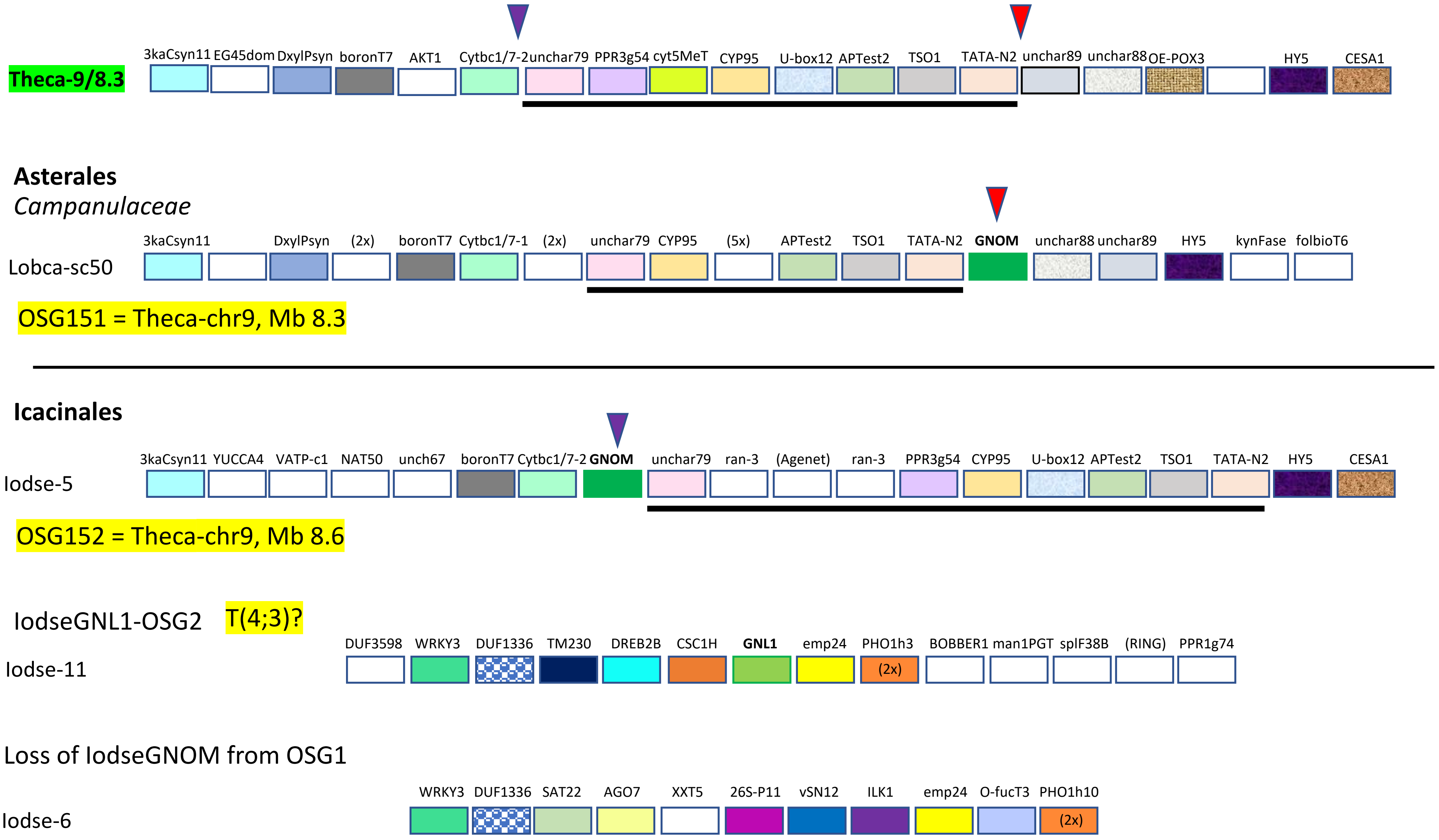

**Suppl. Figure S35.** Transposition of *AtGNOM* ortholog in *Campanulaceae* ( ) and Icacinales ( ) Genes separating the two insertion sites are underlined.

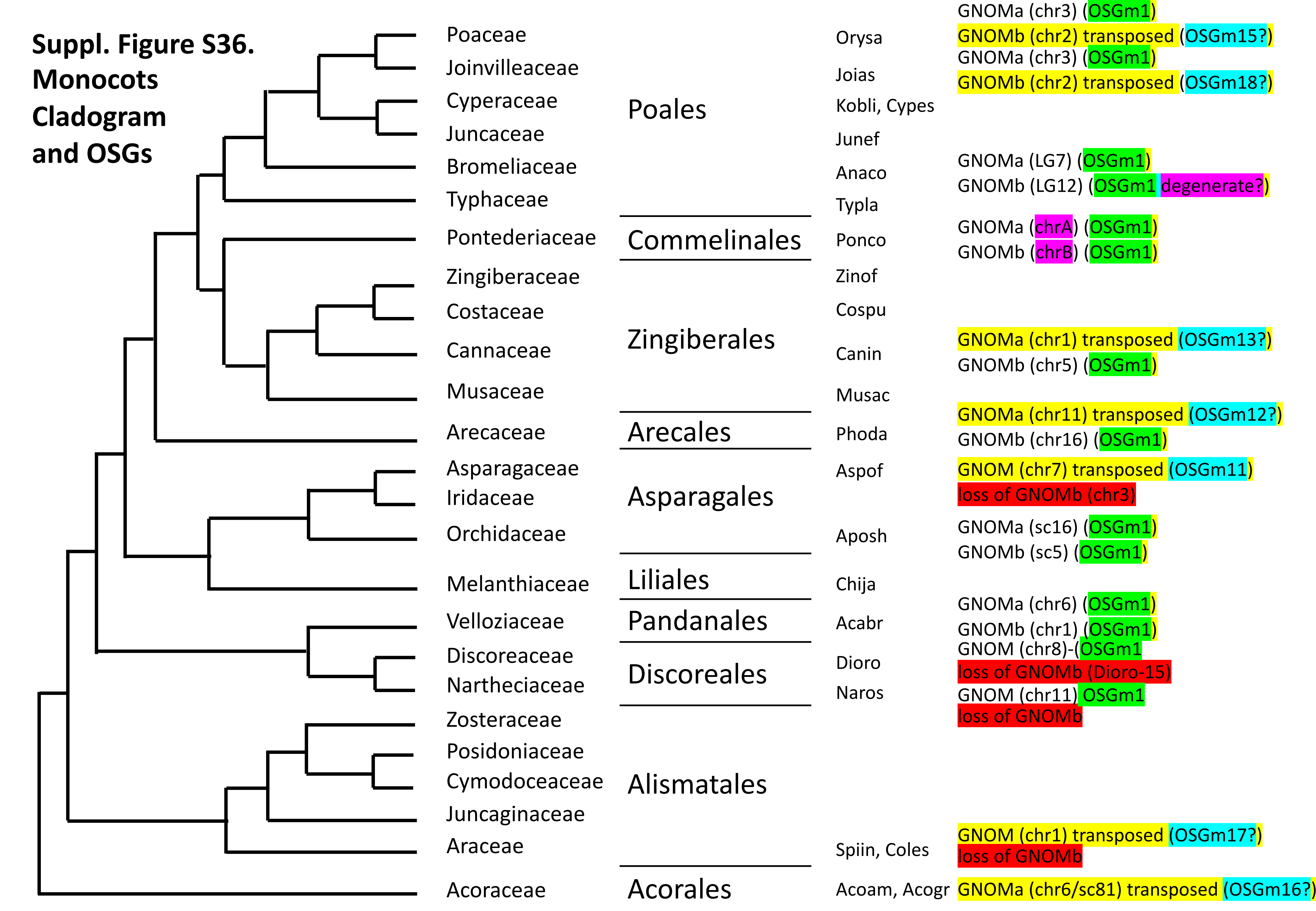

**Suppl. Figure S36. Monocots Cladogram and OSGs**

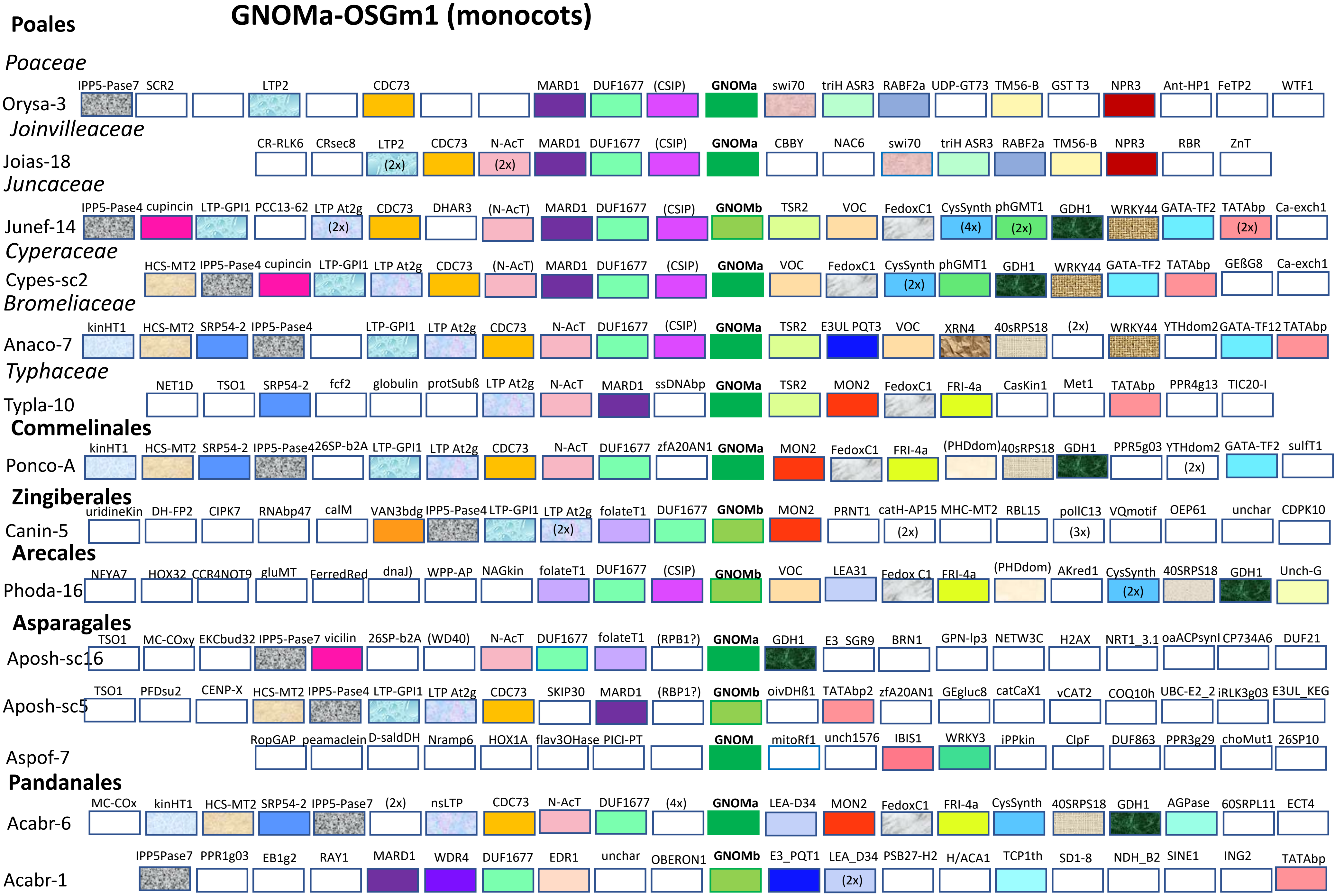

GNOMa-OSGm1 (monocots)
Poales
Poaceae
Orysa-3
IPP5-Pase7 SCR2 LTP2 CDC73 MARD1 DUF1677 (CSIP) GNOMa swi70 triH ASR3 RABF2a UDP-GT73 TM56-B GST T3 NPR3 Ant-HP1 FeTP2 WTF1
Joinvilleaceae
Joias-18
CR-RLK6 CRsec8 LTP2 (2x) CDC73 N-AcT (2x) MARD1 DUF1677 (CSIP) GNOMa CBBY NAC6 swi70 triH ASR3 RABF2a TM56-B NPR3 RBR ZnT
Juncaceae
Junef-14
IPP5-Pase4 cupincin LTP-GPI1 PCC13-62 LTP At2g (2x) CDC73 DHAR3 (N-AcT) MARD1 DUF1677 (CSIP) GNOMb TSR2 VOC FedoxC1 CysSynth (4x) phGMT1 (2x) GDH1 WRKY44 GATA-TF2 TATAbp (2x) Ca-exch1
Cyperaceae
Cypes-sc2
HCS-MT2 IPP5-Pase4 cupincin LTP-GPI1 LTP At2g CDC73 (N-AcT) MARD1 DUF1677 (CSIP) GNOMa VOC FedoxC1 CysSynth (2x) phGMT1 GDH1 WRKY44 GATA-TF2 TATAbp GEßG8 Ca-exch1
Bromeliaceae
Anaco-7
kinHT1 HCS-MT2 SRP54-2 IPP5-Pase4 LTP-GPI1 LTP At2g CDC73 N-AcT DUF1677 (CSIP) GNOMa TSR2 E3UL PQT3 VOC XRN4 40sRPS18 (2x) WRKY44 YTHdom2 GATA-TF12 TATAbp
Typhaceae
Typla-10
NET1D TSO1 SRP54-2 fcf2 globulin protSubß LTP At2g N-AcT MARD1 ssDNAbp GNOMa TSR2 MON2 FedoxC1 FRI-4a CasKin1 Met1 TATAbp PPR4g13 TIC20-I
Commelinales
Ponco-A
kinHT1 HCS-MT2 SRP54-2 IPP5-Pase4 26SP-b2A LTP-GPI1 LTP At2g CDC73 N-AcT DUF1677 zfA20AN1 GNOMa MON2 FedoxC1 FRI-4a (PHDdom) 40sRPS18 GDH1 PPR5g03 YTHdom2 (2x) GATA-TF2 sulfT1
Zingiberales
Canin-5
uridineKin DH-FP2 CIPK7 RNAbp47 calM VAN3bdg IPP5-Pase4 LTP-GPI1 LTP At2g (2x) folateT1 DUF1677 GNOMb MON2 PRNT1 catH-AP15 (2x) MHC-MT2 RBL15 pollC13 (3x) VQmotif OEP61 unchar CDPK10
Arecales
Phoda-16
NFYA7 HOX32 CCR4NOT9 gluMT FerredRed dnaJ) WPP-AP NAGkin folateT1 DUF1677 (CSIP) GNOMb VOC LEA31 Fedox C1 FRI-4a (PHDdom) AKred1 CysSynth (2x) 40SRPS18 GDH1 Unch-G
Asparagales
Aposh-sc16
TSO1 MC-COxy EKCbud32 IPP5-Pase7 vicilin 26SP-b2A (WD40) N-AcT DUF1677 folateT1 (RPB1?) GNOMa GDH1 E3_SGR9 BRN1 GPN-lp3 NETW3C H2AX NRT1_3.1 oaACPsynl CP734A6 DUF21
Aposh-sc5
TSO1 PFDsu2 CENP-X HCS-MT2 IPP5-Pase4 LTP-GPI1 LTP At2g CDC73 SKIP30 MARD1 (RBP1?) GNOMb oivDHß1 TATAbp2 zfA20AN1 GEgluc8 catCaX1 vCAT2 COQ10h UBC-E2_2 iRLK3g03 E3UL_KEG
Aspof-7
RopGAP peamaclein D-saldDH Nramp6 HOX1A flav3OHase PICI-PT GNOM mitoRf1 unch1576 IBIS1 WRKY3 iPPkin ClpF DUF863 PPR3g29 choMut1 26SP10
Pandanales
Acabr-6
MC-COx kinHT1 HCS-MT2 SRP54-2 IPP5-Pase7 (2x) nsLTP CDC73 N-AcT DUF1677 (4x) GNOMa LEA-D34 MON2 FedoxC1 FRI-4a CysSynth 40SRPS18 GDH1 AGPase 60SRPL11 ECT4
Acabr-1
IPP5Pase7 PPR1g03 EB1g2 RAY1 MARD1 WDR4 DUF1677 EDR1 unchar OBERON1 GNOMb E3_PQT1 LEA_D34 (2x) PSB27-H2 H/ACA1 TCP1th SD1-8 NDH_B2 SINE1 ING2 TATAbp

**Poales** GNOMb-OSGs (monocots)

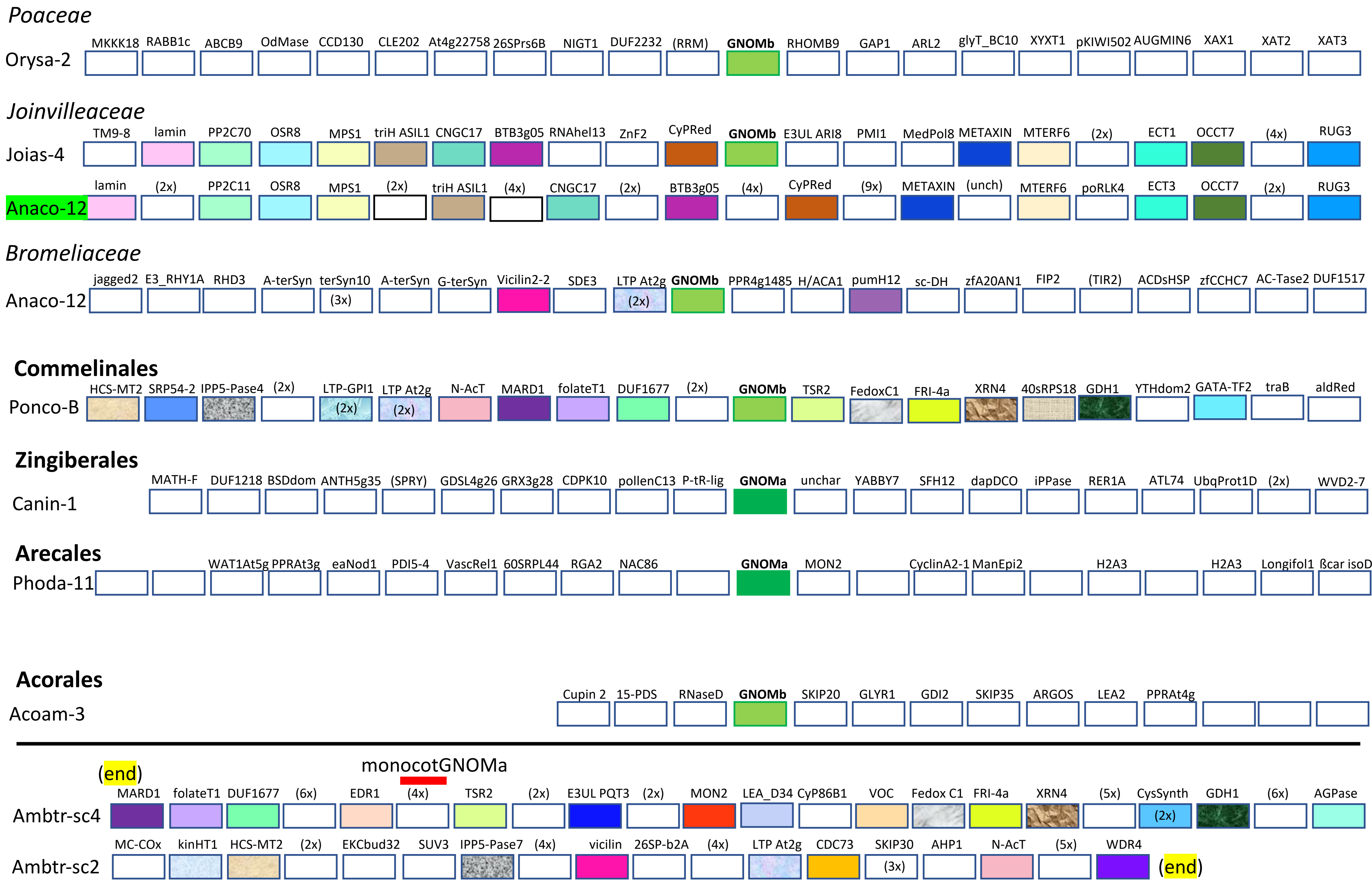

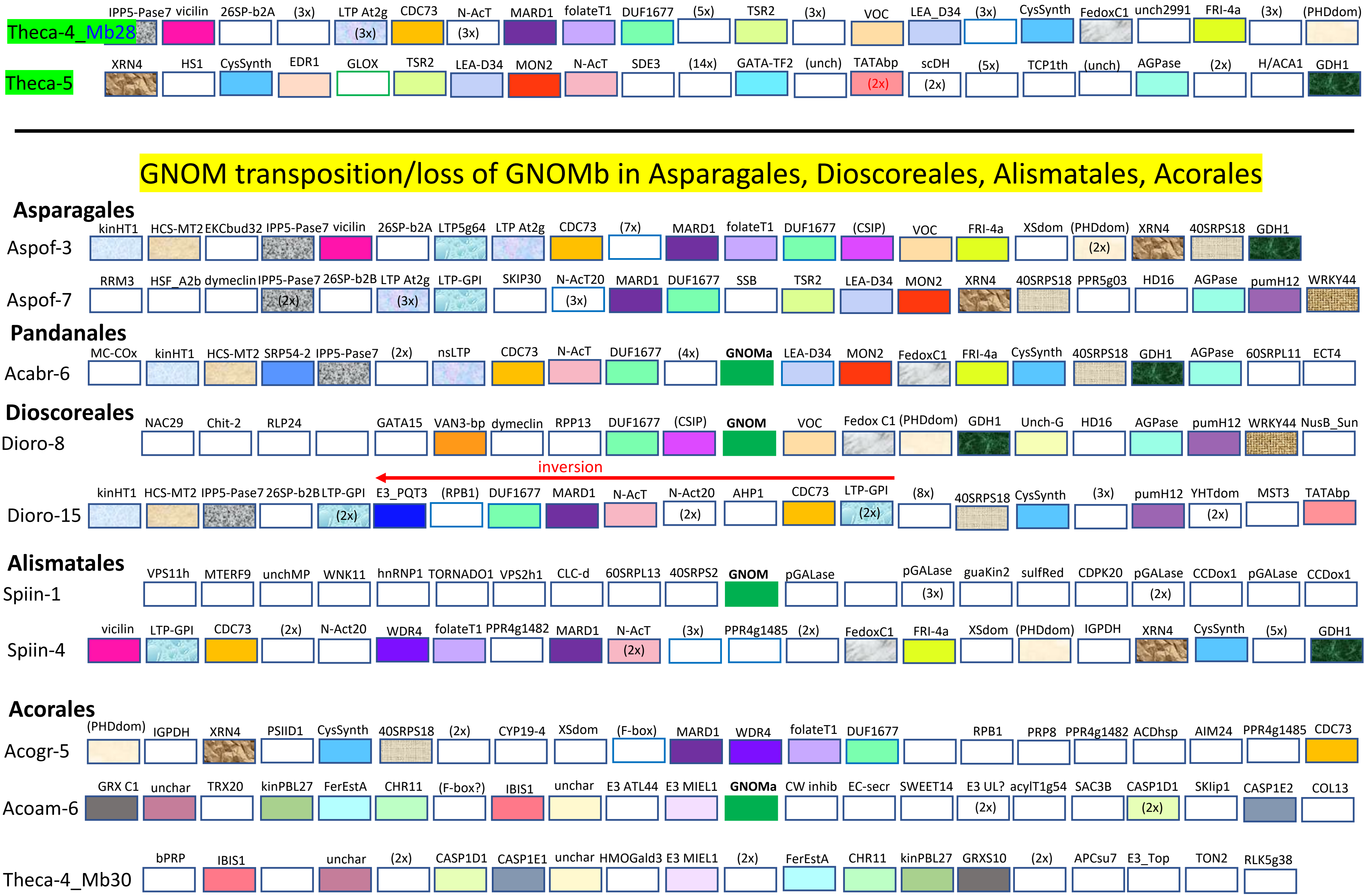

Theca-4_Mb28
IPP5-Pase7 vicilin 26SP-b2A (3x) LTP At2g (3x) CDC73 N-AcT (3x) MARD1 folateT1 DUF1677 (5x) TSR2 (3x) VOC LEA_D34 (3x) CysSynth FedoxC1 unch2991 FRI-4a (3x) (PHDdom)
Theca-5
XRN4 HS1 CysSynth EDR1 GLOX TSR2 LEA-D34 MON2 N-AcT SDE3 (14x) GATA-TF2 (unch) TATAbp (2x) scDH (2x) (5x) TCP1th (unch) AGPase (2x) H/ACA1 GDH1
GNOM transposition/loss of GNOMb in Asparagales, Dioscoreales, Alismatales, Acorales
Asparagales
Aspof-3
kinHT1 HCS-MT2 EKCbud32 IPP5-Pase7 vicilin 26SP-b2A LTP5g64 LTP At2g CDC73 (7x) MARD1 folateT1 DUF1677 (CSIP) VOC FRI-4a XSdom (PHDdom) (2x) XRN4 40SRPS18 GDH1
Aspof-7
RRM3 HSF_A2b dymeclin IPP5-Pase7 (2x) 26SP-b2B LTP At2g (3x) LTP-GPI SKIP30 N-AcT20 (3x) MARD1 DUF1677 SSB TSR2 LEA-D34 MON2 XRN4 40SRPS18 PPR5g03 HD16 AGPase pumH12 WRKY44
Pandanales
Acabr-6
MC-COx kinHT1 HCS-MT2 SRP54-2 IPP5-Pase7 (2x) nsLTP CDC73 N-AcT DUF1677 (4x) GNOMa LEA-D34 MON2 FedoxC1 FRI-4a CysSynth 40SRPS18 GDH1 AGPase 60SRPL11 ECT4
Dioscoreales
Dioro-8
NAC29 Chit-2 RLP24 GATA15 VAN3-bp dymeclin RPP13 DUF1677 (CSIP) GNOM VOC Fedox C1 (PHDdom) GDH1 Unch-G HD16 AGPase pumH12 WRKY44 NusB_Sun
inversion
Dioro-15
kinHT1 HCS-MT2 IPP5-Pase7 26SP-b2B LTP-GPI (2x) E3_PQT3 (RPB1) DUF1677 MARD1 N-AcT N-Act20 (2x) AHP1 CDC73 LTP-GPI (2x) (8x) 40SRPS18 CysSynth (3x) pumH12 YHTdom (2x) MST3 TATAbp
Alismatales
Spiin-1
VPS11h MTERF9 unchMP WNK11 hnRNP1 TORNADO1 VPS2h1 CLC-d 60SRPL13 40SRPS2 GNOM pGALase pGALase (3x) guaKin2 sulfRed CDPK20 pGALase (2x) CCDox1 pGALase CCDox1
Spiin-4
vicilin LTP-GPI CDC73 (2x) N-Act20 WDR4 folateT1 PPR4g1482 MARD1 N-AcT (2x) (3x) PPR4g1485 (2x) FedoxC1 FRI-4a XSdom (PHDdom) IGPDH XRN4 CysSynth (5x) GDH1
Acorales
Acogr-5
(PHDdom) IGPDH XRN4 PSIID1 CysSynth 40SRPS18 (2x) CYP19-4 XSdom (F-box) MARD1 WDR4 folateT1 DUF1677 RPB1 PRP8 PPR4g1482 ACDhsp AIM24 PPR4g1485 CDC73
Acoam-6
GRX C1 unchar TRX20 kinPBL27 FerEstA CHR11 (F-box?) IBIS1 unchar E3 ATL44 E3 MIEL1 GNOMa CW inhib EC-secr SWEET14 E3 UL? (2x) acylT1g54 SAC3B CASP1D1 (2x) SKlip1 CASP1E2 COL13
Theca-4_Mb30
bPRP IBIS1 unchar (2x) CASP1D1 CASP1E1 unchar HMOGald3 E3 MIEL1 (2x) FerEstA CHR11 kinPBL27 GRXS10 (2x) APCsu7 E3_Top TON2 RLK5g38

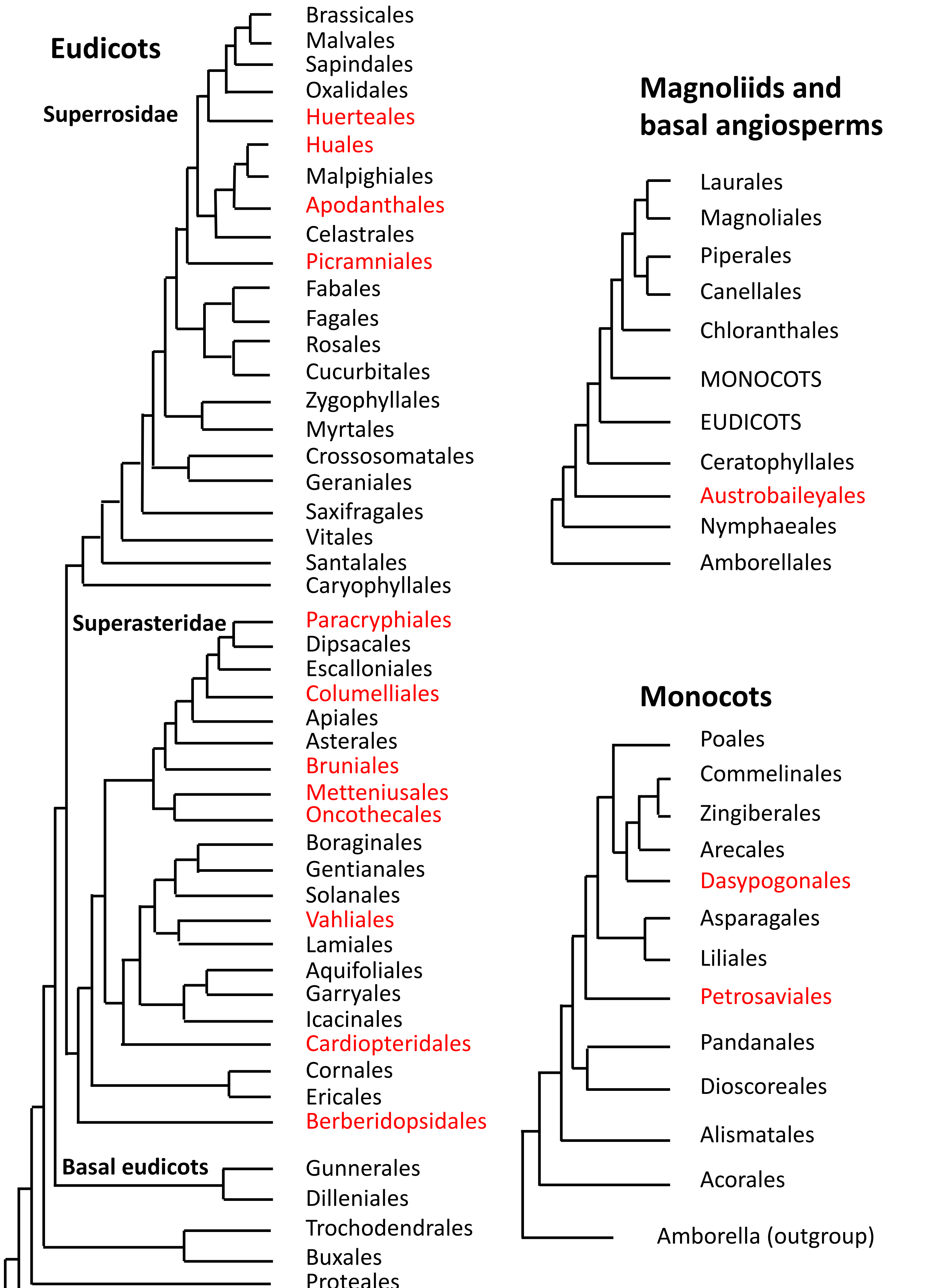

**SupplFig. S37. Cladogram of angiosperm orders**
Orders in red not analysed in this study.
Cladogram based on Zuntini et al. (2024)

#### Suppl. Figure S38. Origin of OSG1 and OSG2 in early-diverging dicots

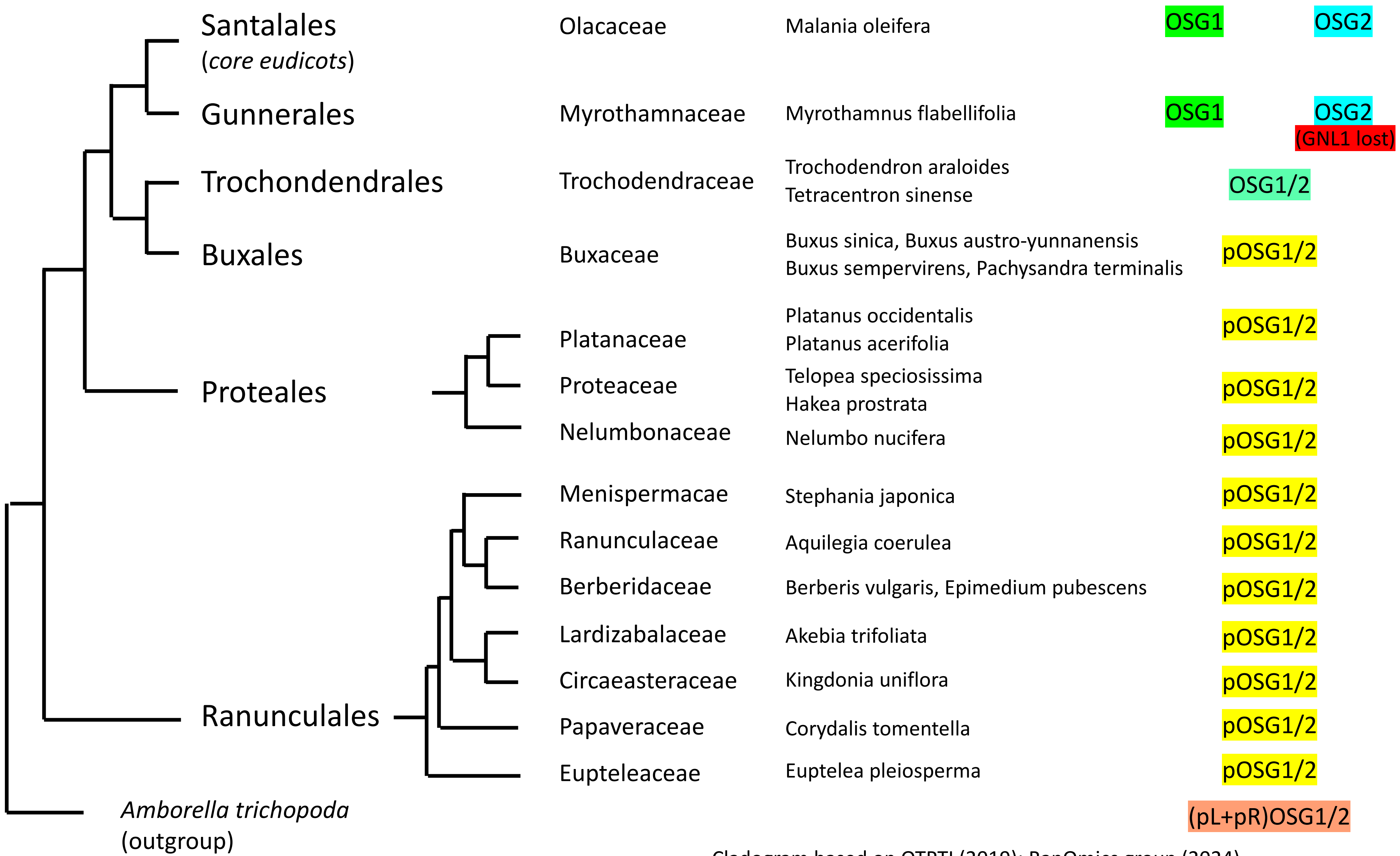

Cladogram based on OTPTI (2019); RanOmics group (2024)

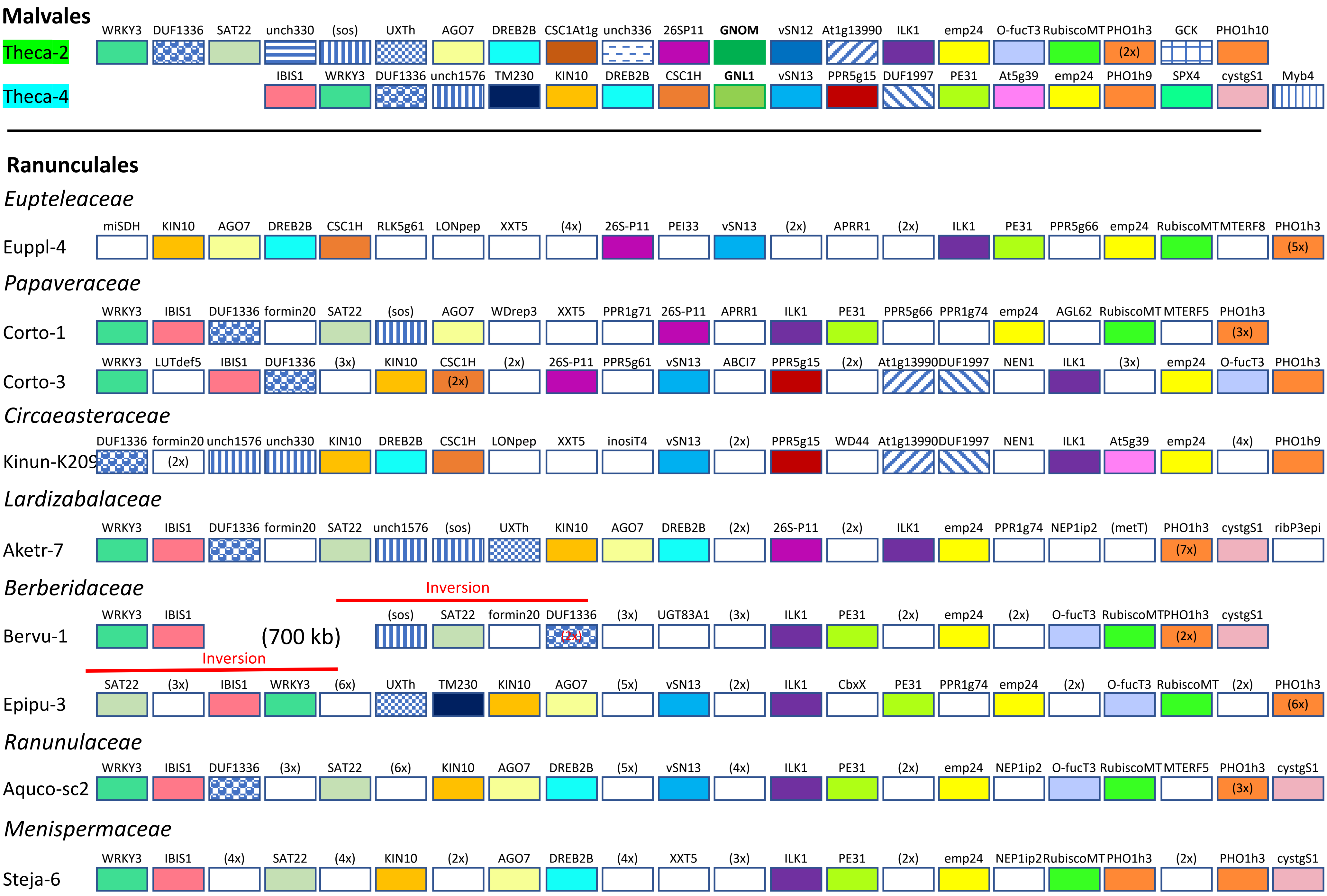
Malvales
Theca-2
WRKY3 DUF1336 SAT22 unch330 (sos) UXTh AGO7 DREB2B CSC1At1g unch336 26SP11 GNOM vSN12 At1g13990 ILK1 emp24 O-fucT3 RubiscoMT PHO1h3 GCK PHO1h10
(2x)
Theca-4
IBIS1 WRKY3 DUF1336 unch1576 TM230 KIN10 DREB2B CSC1H GNL1 vSN13 PPR5g15 DUF1997 PE31 At5g39 emp24 PHO1h9 SPX4 cystgS1 Myb4
Ranunculales
Eupteleaceae
Euppl-4
miSDH KIN10 AGO7 DREB2B CSC1H RLK5g61 LONpep XXT5 (4x) 26S-P11 PEI33 vSN13 (2x) APRR1 (2x) ILK1 PE31 PPR5g66 emp24 RubiscoMT MTERF8 PHO1h3
(5x)
Papaveraceae
Corto-1
WRKY3 IBIS1 DUF1336 formin20 SAT22 (sos) AGO7 WDrep3 XXT5 PPR1g71 26S-P11 APRR1 ILK1 PE31 PPR5g66 PPR1g74 emp24 AGL62 RubiscoMT MTERF5 PHO1h3
(3x)
Corto-3
WRKY3 LUTdef5 IBIS1 DUF1336 (3x) KIN10 CSC1H (2x) 26S-P11 PPR5g61 vSN13 ABCI7 PPR5g15 (2x) At1g13990 DUF1997 NEN1 ILK1 (3x) emp24 O-fucT3 PHO1h3
(2x)
Circaeasteraceae
Kinun-K209
DUF1336 formin20 unch1576 unch330 KIN10 DREB2B CSC1H LONpep XXT5 inosiT4 vSN13 (2x) PPR5g15 WD44 At1g13990 DUF1997 NEN1 ILK1 At5g39 emp24 (4x) PHO1h9
(2x)
Lardizabalaceae
Aketr-7
WRKY3 IBIS1 DUF1336 formin20 SAT22 unch1576 (sos) UXTh KIN10 AGO7 DREB2B (2x) 26S-P11 (2x) ILK1 emp24 PPR1g74 NEP1ip2 (metT) PHO1h3 cystgS1 ribP3epi
(7x)
Berberidaceae
Inversion
Bervu-1
WRKY3 IBIS1 (700 kb) (sos) SAT22 formin20 DUF1336 (3x) UGT83A1 (3x) ILK1 PE31 (2x) emp24 (2x) O-fucT3 RubiscoMT PHO1h3 cystgS1
(2x) (2x)
Inversion
Epipu-3
SAT22 (3x) IBIS1 WRKY3 (6x) UXTh TM230 KIN10 AGO7 (5x) vSN13 (2x) ILK1 CbxX PE31 PPR1g74 emp24 (2x) O-fucT3 RubiscoMT (2x) PHO1h3
(6x)
Ranunulaceae
Aquco-sc2
WRKY3 IBIS1 DUF1336 (3x) SAT22 (6x) KIN10 AGO7 DREB2B (5x) vSN13 (4x) ILK1 PE31 (2x) emp24 NEP1ip2 O-fucT3 RubiscoMT MTERF5 PHO1h3 cystgS1
(3x)
Menispermaceae
Steja-6
WRKY3 IBIS1 (4x) SAT22 (4x) KIN10 (2x) AGO7 DREB2B (4x) XXT5 (3x) ILK1 PE31 (2x) emp24 NEP1ip2 RubiscoMT PHO1h3 (2x) PHO1h3 cystgS1

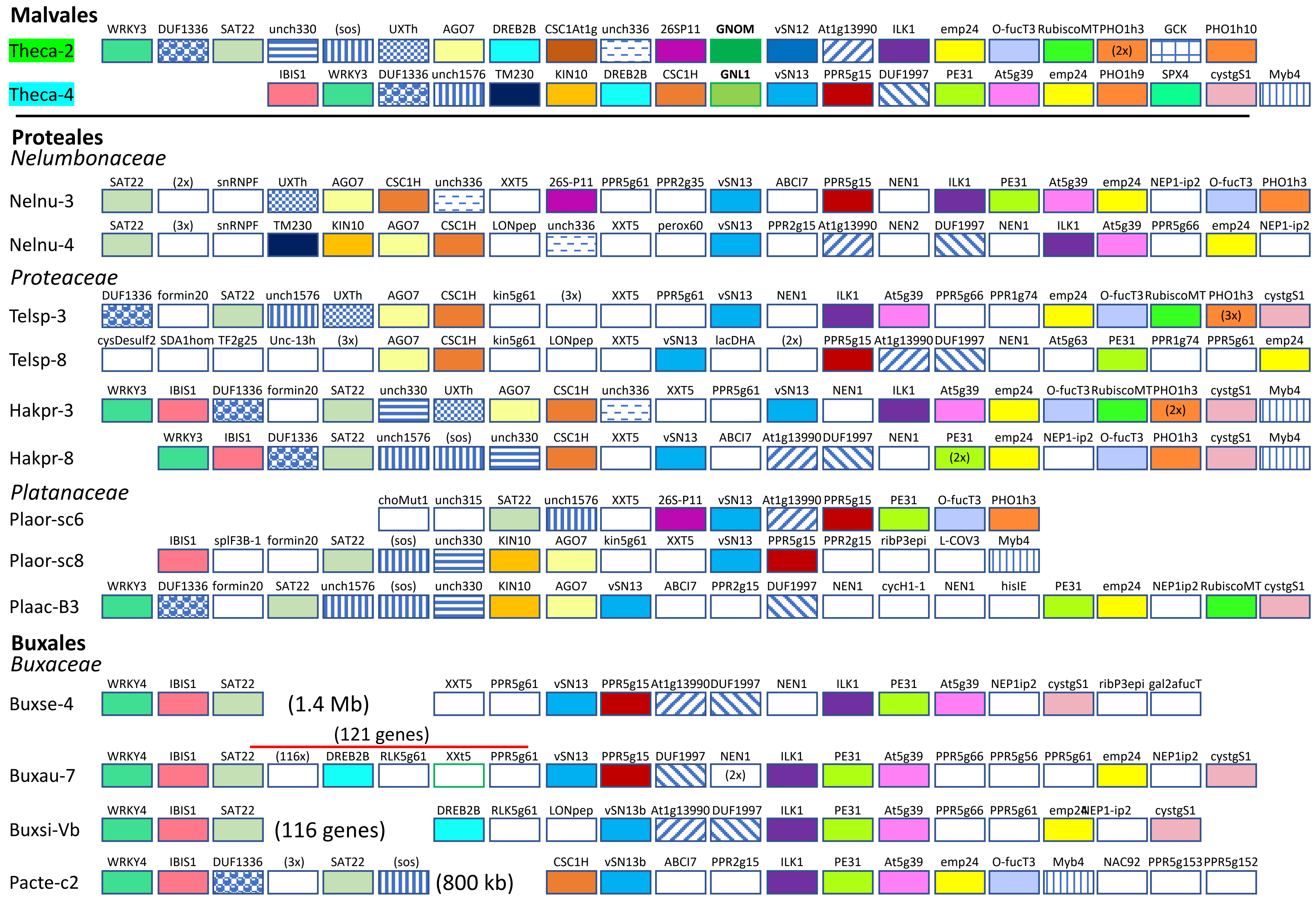
Malvales
Theca-2
WRKY3 DUF1336 SAT22 unch330 (sos) UXTh AGO7 DREB2B CSC1At1g unch336 26SP11 GNOM vSN12 At1g13990 ILK1 emp24 O-fucT3 RubiscoMT PHO1h3 GCK PHO1h10
(2x)
Theca-4
IBIS1 WRKY3 DUF1336 unch1576 TM230 KIN10 DREB2B CSC1H GNL1 vSN13 PPR5g15 DUF1997 PE31 At5g39 emp24 PHO1h9 SPX4 cystgS1 Myb4
Proteales
Nelumbonaceae
Nelnu-3
SAT22 (2x) snRNPF UXTh AGO7 CSC1H unch336 XXT5 26S-P11 PPR5g61 PPR2g35 vSN13 ABCI7 PPR5g15 NEN1 ILK1 PE31 At5g39 emp24 NEP1-ip2 O-fucT3 PHO1h3
Nelnu-4
SAT22 (3x) snRNPF TM230 KIN10 AGO7 CSC1H LONpep unch336 XXT5 perox60 vSN13 PPR2g15 At1g13990 NEN2 DUF1997 NEN1 ILK1 At5g39 PPR5g66 emp24 NEP1-ip2
Proteaceae
Telsp-3
DUF1336 formin20 SAT22 unch1576 UXTh AGO7 CSC1H kin5g61 (3x) XXT5 PPR5g61 vSN13 NEN1 ILK1 At5g39 PPR5g66 PPR1g74 emp24 O-fucT3 RubiscoMT PHO1h3 cystgS1
(3x)
Telsp-8
cysDesulf2 SDA1hom TF2g25 Unc-13h (3x) AGO7 CSC1H kin5g61 LONpep XXT5 vSN13 lacDHA (2x) PPR5g15 At1g13990 DUF1997 NEN1 At5g63 PE31 PPR1g74 PPR5g61 emp24
Hakpr-3
WRKY3 IBIS1 DUF1336 formin20 SAT22 unch330 UXTh AGO7 CSC1H unch336 XXT5 PPR5g61 vSN13 NEN1 ILK1 At5g39 emp24 O-fucT3 RubiscoMT PHO1h3 cystgS1 Myb4
(2x)
Hakpr-8
WRKY3 IBIS1 DUF1336 SAT22 unch1576 (sos) unch330 CSC1H XXT5 vSN13 ABCI7 At1g13990 DUF1997 NEN1 PE31 emp24 NEP1-ip2 O-fucT3 PHO1h3 cystgS1 Myb4
(2x)
Platanaceae
Plaor-sc6
choMut1 unch315 SAT22 unch1576 XXT5 26S-P11 vSN13 At1g13990 PPR5g15 PE31 O-fucT3 PHO1h3
Plaor-sc8
IBIS1 splF3B-1 formin20 SAT22 (sos) unch330 KIN10 AGO7 kin5g61 XXT5 vSN13 PPR5g15 PPR2g15 ribP3epi L-COV3 Myb4
Plaac-B3
WRKY3 DUF1336 formin20 SAT22 unch1576 (sos) unch330 KIN10 AGO7 vSN13 ABCI7 PPR2g15 DUF1997 NEN1 cycH1-1 NEN1 hisIE PE31 emp24 NEP1ip2 RubiscoMT cystgS1
Buxales
Buxaceae
Buxse-4
WRKY4 IBIS1 SAT22 XXT5 PPR5g61 vSN13 PPR5g15 At1g13990 DUF1997 NEN1 ILK1 PE31 At5g39 NEP1ip2 cystgS1 ribP3epi gal2afucT
(1.4 Mb)
(121 genes)
Buxau-7
WRKY4 IBIS1 SAT22 (116x) DREB2B RLK5g61 XXt5 PPR5g61 vSN13 PPR5g15 DUF1997 NEN1 ILK1 PE31 At5g39 PPR5g66 PPR5g56 PPR5g61 emp24 NEP1ip2 cystgS1
(2x)
Buxsi-Vb
WRKY4 IBIS1 SAT22 DREB2B RLK5g61 LONpep vSN13b At1g13990 DUF1997 ILK1 PE31 At5g39 PPR5g66 PPR5g61 emp24 NEP1-ip2 cystgS1
(116 genes)
Pacte-c2
WRKY4 IBIS1 DUF1336 (3x) SAT22 (sos) CSC1H vSN13b ABCI7 PPR2g15 ILK1 PE31 At5g39 emp24 O-fucT3 Myb4 NAC92 PPR5g153 PPR5g152
(800 kb)

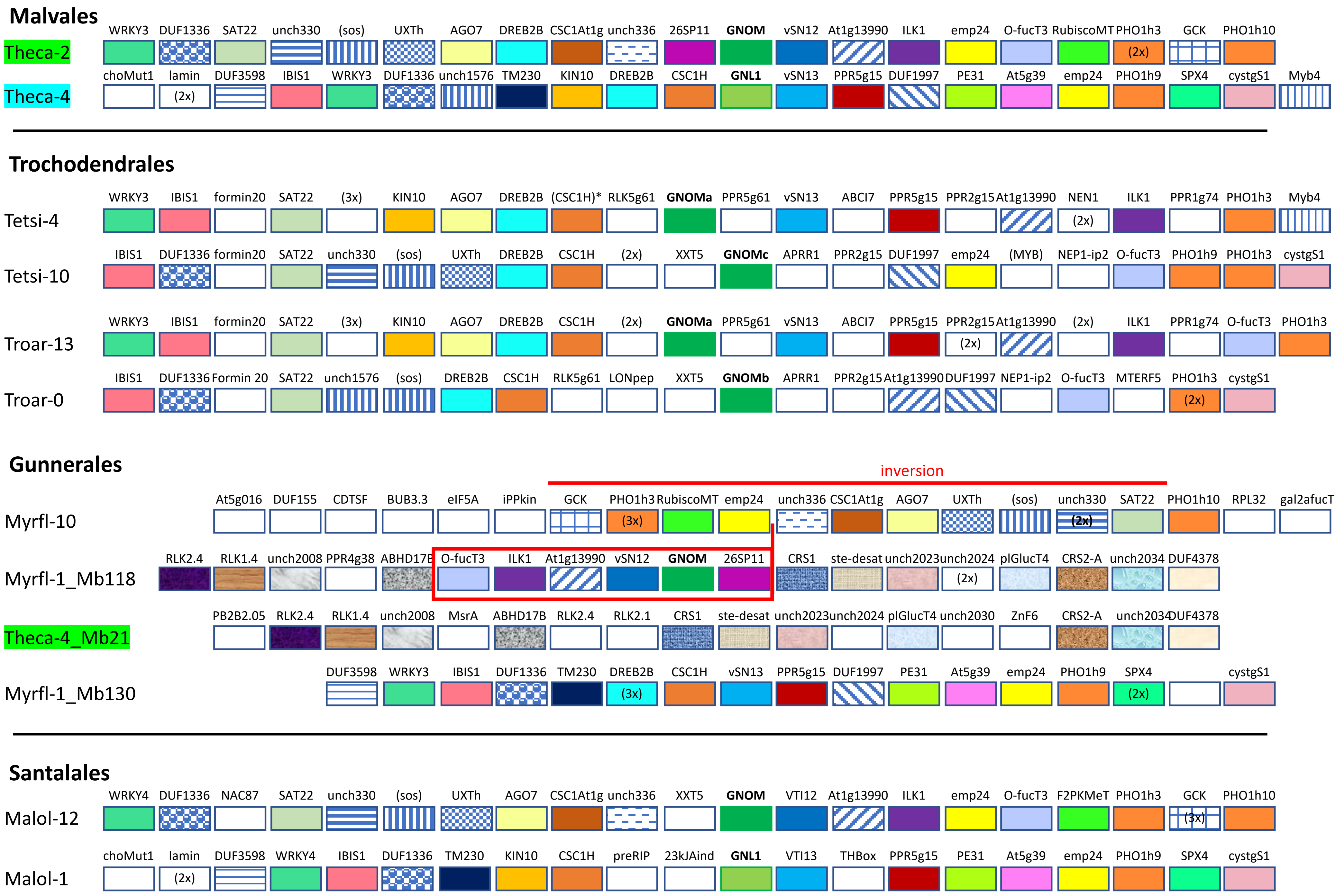
Malvales
Theca-2
WRKY3 DUF1336 SAT22 unch330 (sos) UXTh AGO7 DREB2B CSC1At1g unch336 26SP11 GNOM vSN12 At1g13990 ILK1 emp24 O-fucT3 RubiscoMT PHO1h3 GCK PHO1h10
(2x)
Theca-4
choMut1 lamin DUF3598 IBIS1 WRKY3 DUF1336 unch1576 TM230 KIN10 DREB2B CSC1H GNL1 vSN13 PPR5g15 DUF1997 PE31 At5g39 emp24 PHO1h9 SPX4 cystgS1 Myb4
(2x)
Trochodendrales
Tetsi-4
WRKY3 IBIS1 formin20 SAT22 (3x) KIN10 AGO7 DREB2B (CSC1H)* RLK5g61 GNOMa PPR5g61 vSN13 ABCI7 PPR5g15 PPR2g15 At1g13990 NEN1 ILK1 PPR1g74 PHO1h3 Myb4
(2x)
Tetsi-10
IBIS1 DUF1336 formin20 SAT22 unch330 (sos) UXTh DREB2B CSC1H (2x) XXT5 GNOMc APRR1 PPR2g15 DUF1997 emp24 (MYB) NEP1-ip2 O-fucT3 PHO1h9 PHO1h3 cystgS1
Troar-13
WRKY3 IBIS1 formin20 SAT22 (3x) KIN10 AGO7 DREB2B CSC1H (2x) GNOMa PPR5g61 vSN13 ABCI7 PPR5g15 PPR2g15 At1g13990 (2x) ILK1 PPR1g74 O-fucT3 PHO1h3
(2x)
Troar-0
IBIS1 DUF1336 Formin 20 SAT22 unch1576 (sos) DREB2B CSC1H RLK5g61 LONpep XXT5 GNOMb APRR1 PPR2g15 At1g13990 DUF1997 NEP1-ip2 O-fucT3 MTERF5 PHO1h3 cystgS1
(2x)
Gunnerales
inversion
Myrfl-10
At5g016 DUF155 CDTSF BUB3.3 eIF5A iPPkin GCK PHO1h3 RubiscoMT emp24 unch336 CSC1At1g AGO7 UXTh (sos) unch330 SAT22 PHO1h10 RPL32 gal2afucT
(3x) (2x)
Myrfl-1_Mb118
RLK2.4 RLK1.4 unch2008 PPR4g38 ABHD17B O-fucT3 ILK1 At1g13990 vSN12 GNOM 26SP11 CRS1 ste-desat unch2023 unch2024 plGlucT4 CRS2-A unch2034 DUF4378
(2x)
Theca-4_Mb21
PB2B2.05 RLK2.4 RLK1.4 unch2008 MsrA ABHD17B RLK2.4 RLK2.1 CRS1 ste-desat unch2023 unch2024 plGlucT4 unch2030 ZnF6 CRS2-A unch2034 DUF4378
Myrfl-1_Mb130
DUF3598 WRKY3 IBIS1 DUF1336 TM230 DREB2B CSC1H vSN13 PPR5g15 DUF1997 PE31 At5g39 emp24 PHO1h9 SPX4 cystgS1
(3x) (2x)
Santalales
Malol-12
WRKY4 DUF1336 NAC87 SAT22 unch330 (sos) UXTh AGO7 CSC1At1g unch336 XXT5 GNOM VTI12 At1g13990 ILK1 emp24 O-fucT3 F2PKMeT PHO1h3 GCK PHO1h10
(3x)
Malol-1
choMut1 lamin DUF3598 WRKY4 IBIS1 DUF1336 TM230 KIN10 CSC1H preRIP 23kJAind GNL1 VTI13 THBox PPR5g15 PE31 At5g39 emp24 PHO1h9 SPX4 cystgS1
(2x)

**Suppl. Table S1A. GBF1-type ARF-GEFs in Charophyta (7 orders, 8 families, 9 species)**

| Order | Species | Gene name (chr/ctg/sc #) | Protein Size (# aa) | BFA res / sens | % id. AtGNOM | % id. AtGNL1 | database | ref |
|---|---|---|---|---|---|---|---|---|
| (Mesostigmatophyceae) Mesostigmatales | Mesostigma viride | MesviGNOM (sc64, Mb 0.5) | (1211) | sens | (46) | (44) | NCBI | GCA_009 746045.1 |
| (Chlorokybophyceae) Chlorokybales | Chlorokybus atmophyticus | ChlatGNOM (ctg_Cat1294, Mb 0.2) | 1309 | sens | 52 | 44 | NCBI | GCA_009 103225.1 |
| (Klebsormidiophyceae) Klebsormidiales | Klebsormidium fl. (nitens) | KleniGNOM Kfl00216_0140 (ctg kfl00216, Mb 0.1) | 1732 | sens | 50 | 46 | NCBI | GCA_000 708835.1 |
| (Charophyceae) Charales | Chara braunii | ChabrGNOM Chabr_g19322 (sc230, Mb 0.9) | 1718 | res | 61 | 49 | NCBI | GCA_003 427395.1 |
| (Zygnematophyceae) Desmidiales | Penium margaritaceum | PenmaGNOMa pm007614g0010 (ctg PM002557, kb 30) | 1681 | sens | 57 | 48 | NCBI | GCA_023 375785.1 |
| | | PenmaGNOMb pm002557g0040 (ctg PM002557, Mb 0.1) | 1827 | sens | 50 | 43 | NCBI | GCA_023 375785.1 |
| (Zygnematophyceae) Spirogloeales | Spirogloea muscicola | SpimuGNOMa (ctg 0214 s82, kb 345) | 1573 | sens | 59 | 50 | NCBI | GCA_009 602725.1 |
| | | SpimuGNOMb (ctg 0214s17, kb 10) | 1573 | sens | 59 | 49 | NCBI | GCA_009 602725.1 |
| | | SpimuGNOMc (ctg 0214s57, kb 270) | 1574 | sens | 58 | 49 | NCBI | GCA_009 602725.1 |
| (Zygnematophyceae) Zygnematales (Zygnemataceae) | Zygnema cf. cylindricum SAG698-1a | ZygcyGNOM Zygcyl6981a (sc 3410, Mb 0.4) | 1592 | sens | 59 | 50 | NCBI | GCA_037 447655.1 |
| (Zygnematophyceae) Zygnematales (Zygnemataceae) | Zygnema circumcarinatum UTEX 1560 | ZygciGNOM Zygcir1560 (chr 10, Mb 1.4) | 1521 | sens | 53 | 48 | NCBI | GCA_037 447605.1 |
| (Zygnematophyceae) Zygnematales (Mesotaeniaceae) | Ancylonema nordenskioldii | AncnoGNOM ctg12227 | 1533 | sens | 52 | 47 | NCBI | GCA_050 230475.1 |

**Suppl. Table S1B. GBF1-type ARF-GEFs in Bryophyta, Lycophyta, ferns (12 orders, 12 families, 15 species)**

| Order | Species | Gene name (chr/ctg/sc #) | Protein Size (# aa) | BFA res / sens | % id. AtGN | % id. AtGNL1 | database | ref |
|---|---|---|---|---|---|---|---|---|
| **Bryophyta s.l.** | | | | | | | | |
| LIVERWORTS (Marchantiophyta) Marchantiales | Marchantia polymorpha subsp. ruderalis | MarpoGNOM MPTK2_3g25100 (chr3, Mb29.5) | 1513 | res | 65 | 55 | NCBI | GCA_0378339 65.1 |
| HORNWORTS (Anthocerotophyta) Anthocerotales | Anthoceros angustus | AntanGNOMa_ (scAANG162, Mb 0.5) | 1490 | sens | 62 | 55 | NCBI | GCA_0109091 65.1 |
| | | AntanGNOMb (scAANG383, kb20) | 1465 | sens | 61 | 53 | NCBI | GCA_0109091 65.1 |
| HORNWORTS (Anthocerotophyta) Anthocerotales | Anthoceros agrestis OXF | AntagGNOM (utg58l.310) (utg58l, Mb 1.5)) | 1500 | sens | 64 | 56 | UZH | pep file |
| MOSSES/ BRYOPHYTA s.s. Funariales | Physcomitrium patens | PhypaGNOMa LOC112274759 (chr22, Mb 3.5) | 1482 | sens | 62 | 54 | NCBI | GCF_0000024 25.5 |
| | | PhypaGNOMb LOC112292439 (chr15, Mb 7.5) | 1498 | sens | 59 | 53 | NCBI | GCF_0000024 25.5 |
| | | PhypaGNOMc LOC11228663 (chr9, Mb 8.0) | 1493 | sens | 59 | 52 | NCBI | GCF_0000024 25.5 |
| MOSSES Hypnales (Thuidiaceae) | Thuidium tamariscinum | ThutaGNOM (chr 1, Mb 32.3) | 1487 | sens | 63 | 54 | NCBI | GCA_9470348 85.1 |
| MOSSES Sphagnales | Sphagnum fallax | SphfaGNOM Sphfalx16G018200 | 1553 | sens | 64 | 55 | Phytozome 13 | S.fallax v1.1 |
| MOSSES Dicranales | Ceratodon purpureus | CerpuGNOMa CepurR40.8G135600 | 1489 | sens | 64 | 54 | Phytozome 13 | C. purpureus v1.1 |
| | | CrepuGNOMb CepurR40.7G143500 | 1497 | sens | 61 | 53 | Phytozome 13 | C. purpureus v1.1 |
| | | CerpuGNOMc CepurR40.9G122800 | 1495 | res | 60 | 53 | Phytozome 13 | C. purpureus v1.1 |
| MOSSES Bryales | Pohlia nutans | PohnuGNOMa (chr 5, Mb 6.7) | 1485 | sens | 64 | 55 | NCBI | GCA 022496805.1 |
| | | PohnuGNOMb chr 6, Mb 6.3) | 1486 | sens | 64 | 55 | NCBI | GCA 022496805.1 |

| | | | | | | | | |
|---|---|---|---|---|---|---|---|---|
| **Lycophyta** | | | | | | | | |
| LYCOPHYTES Selaginellales (Selaginellaceae) | Selaginella moellendorffii | SelmoGNOMa Selmo_LOC9653513 (sc69, Mb 0.6) | 1436 | sens | 66 | 56 | NCBI | GCF_0001434 15.4 |
| | | SelmoGNOMb Selmo_LOC9657677 (sc24, Mb 1.1) | 1436 | sens | 66 | 56 | NCBI | GCF_0001434 15.4 |
| LYCOPHYTES Lycopodiales (Lycopodiaceae) | Diaphasiastrum complanatum | DiacoGNOMa Dicom.09G064300 (chr 9, Mb 35.8) | 1519 | sens | 64 | 54 | Phytozome 13 | Diaphasiastrum complanatum v3.1 |
| | | DiacoGNOMb Dicom.01G064200 (chr 1, Mb 35.4) | 1498 | res | 58 | 53 | Phytozome 13 | Diaphasiastrum complanatum v3.1 |
| **ferns** | | | | | | | | |
| Salviniales (Salviniaceae) | Salvinia cucullata | SalcuGNOMa Sacu_s007.g003923 | 1437 | sens | 64 | 56 | fernbase | Salvinia cucullata_v1.1 |
| | | SalcuGNOMb Sacu_s0267.g026961 | 1418 | res | 56 | 52 | fernbase | Salvinia cucullata_v1.1 |
| Salviniales (Salviniaceae) | Azolla filiculoides | AzofiGNOMa Azfi_s0087.g042331 | 1421 | sens | 66 | 57 | fernbase | Azolla filiculoides |
| | | AzofiGNOMb Azfi_s0094.g043397 | 1416 | res | 60 | 52 | fernbase | Azolla filiculoides |
| Cyatheales (Cyatheaceae) | Alsophila spinulosa | AlsspGNOMa Aspi01Gene41929 | 1460 | sens | 67 | 58 | Figshare | Pep file |
| | | AlsspGNOMb Aspi01Gene65565 | 1443 | sens | 62 | 55 | Figshare | Pep file |
| | | AlsspGNOMc Aspi01Gene60238 | 1445 | res | 62 | 55 | Figshare | Pep file |
| Polypodiales (Pteridaceae) | Ceratopteris richardii | CerriGNOMa KP509_02G006600 (chr 2, Mb 16.9) | 1440 | sens | 67 | 58 | NCBI | GCA_0203108 75.1 |
| | | CerriGNOMb KP509_15G051500 (chr 15, Mb 135.3) | 1399 | res | 54 | 50 | NCBI | GCA_0203108 75.1 |
| Polypodiales (Pteridaceae) | Adiantum capillus-veneris | AdicaGNOMa Adc10687 (chr22, Mb 53.1) | 1441 | sens | 67 | 58 | NCBI | GCA_0145293 85.2 |
| | | AdicaGNOMb (chr19, Mb 70.9) | 1425 | res | 55 | 50 | NCBI | GCA_0145293 85.2 |

**Suppl. Table S1C. GBF1-type ARF-GEFs in gymnosperms (5 orders, 7 families, 13 species)**

| Order | Species | Gene name (chr/ctg/sc #) | Protein Size (# aa) | BFA res / sens | % id. AtGN | % id. AtGNL1 | database | ref |
|---|---|---|---|---|---|---|---|---|
| Ginkgoales (Ginkgoaceae) | Ginkgo biloba | GinbiGNOMa Gb_01979 | 1498 | sens | 74 | 61 | Giga Science | Ginkgo biloba |
| | | GinbiGNOMb Gb_26378 | 1450 | res | 71 | 59 | Giga Science | Ginkgo biloba |
| Cycadales (Cycadaceae) | Cycas panzhihuaensis | CycpaGNOMa CYCAS_003811 (chr1, Mb 1234.2) | 1498 | sens | 74 | 61 | NCBI | GCA_0232 13395.1 |
| | | CycasGNOMb CYCAS_005782 (chr10, Mb 537.1) | 1490 | res | 68 | 58 | NCBI | GCA_0232 13395.1 |
| Gnetales (Welwitschiaceae) | Welwitschia mirabilis | WelmiGNOMa W.mirabilis.07742 (chr 3, Mb 119.7) | 1473 | sens | 68 | 58 | Dryad | Pep file |
| | | WelmiGNOMb W.mirabilis.23859 (chr 6, Mb 80.9) | 1414 | sens | 61 | 57 | Dryad | Pep file |
| Gnetales (Gnetaceae) | Gnetum montanum | GnemoGNOMa TnS000619529 (sc619529, kb 270) | 1478 | sens | 69 | 58 | NCBI | GCA_0156 80685.1 |
| | | GnemoGNOMb TnS000326955 (sc326955, kb 60) | 1476 | res | 61 | 56 | NCBI | GCA_0156 80685.1 |
| Pinales (Pinaceae) | Picea abies | PicabGNOMa (ctg PA09770556.1) | 1483 | sens | 73 | 61 | NCBI | GCA_9000 67695.1 |
| | | PicabGNOMb (comp95351) (4 ctgs) | 1487 | res | 68 | 58 | NCBI | GCA_9000 67695.1 |
| Pinales (Pinaceae) | Pinus taeda | PintaGNOMa (sc110652, kb 75) | 1483 | sens | 71 | 63 | NCBI | GCA_0004 04065.3 |
| | | PintaGNOMb (sc21345, kb 210) | 1482 | res | 67 | 57 | NCBI | GCA_0004 04065.3 |
| Pinales (Pinaceae) | Larix sibirica | LarsiGNOMa (sc4207121) | 1475 | sens | 72 | 60 | NCBI | GCA_0041 51065.1 |
| | | LarsiGNOMb (sc5111451) | 1488 | sens | 67 | 58 | NCBI | GCA_0041 51065.1 |

| | | | | | | | | |
|---|---|---|---|---|---|---|---|---|
| Cupressales (Taxaceae) | Taxus chinensis | TaxchGNOMa (chr 3, Mb 169.7) | 1470 | sens | 77 | 60 | NCBI | GCA_0197 76745.2 |
| | | TaxchGNOMb (chr 7, Mb 230.1) | 1479 | res | 70 | 60 | NCBI | GCA_0197 76745.2 |
| Cupressales (Taxaceae) | Torreya grandis | TorgrGNOMa (chr 7) TG7g00959 | 1474 | sens | 72 | 60 | figshare | Pep file |
| | | TorgrGNOMb (chr 8) TG8g02666 | 1475 | res | 66 | 58 | figshare | Pep fil |
| Cupressales (Cupressaceae) | Sequoiadendron giganteum | SeqgiGNOMa SEGI_06272 (chr 5, Mb 134.0) | 1474 | sens | 73 | 61 | NCBI | GCA_0071 15665.2 |
| | | SeggiGNOMb SEGI_34273 (chr 3, Mb 673.3) | 1461 | res | 67 | 57 | NCBI | GCA_0071 15665.2 |
| Cupressales (Cupressaceae) | Thuja plicata | ThuplGNOMa (sc29380805, Mb 0.5) | 1474 | sens | 72 | 60 | NCBI | GCA_0185 84345.1 |
| | | ThuplGNOMb (sc29378734, Mb 0.5) | 1473 | res | 67 | 58 | NCBI | GCA_0185 84345.1 |
| Cupressales (Cupressaceae) | Cupressus sempervirens | CupseGNOMa (chr5, Mb 914) | 1474 | sens | 73 | 61 | NCBI | GCA_0287 49045.1 |
| | | CupseGNOMb (chr3, Mb 354) | 1461 | res | 67 | 58 | NCBI | GCA_0287 49045.1 |
| Cupressales (Cupressaceae) | Cryptomeria japonica | CryjaGNOMa LOC131050749 (chr5, Mb 780.5) | 1474 | sens | 73 | 61 | NCBI | GCF_0302 72615.1 |
| | | CryjaGNOMb LOC131035957 (chr3, Mb 323.3) | 1472 | res | 67 | 58 | NCBI | GCF_0302 72615.1 |

**Suppl. Table S1D. GBF1-type ARF-GEFs in basal angiosperms, magnoliids, chloranthales, ceratophyllales (8 orders, 12 families, 21 species)**

| Order | Species | Gene name (chr/ctg/sc #) | Protein Size (# aa) | BFA res / sens | % id. AtGN | % id. AtGNL1 | % id. AtGNL2 | database | ref |
|---|---|---|---|---|---|---|---|---|---|
| **Basal angiosperms** | | | | | | | | | |
| Amborellales (Amborellaceae) | Amborella trichopoda | AmbtrGNOM LOC18429558 (chr3, Mb 57.2) | 1469 | sens | 78 | 63 | | NCBI | GCA_046862475.1 |
| | | AmbtrGNL2 LOC18436789 (chr11, Mb 37.4) | 1389 | sens | | | 49 | NCBI | GCA_046862475.1 |
| Nymphaeales (Nymphaeaceae) | Nymphaea colorata | NymcoGNOM LOC116253329 (chr1, Mb 12.7)) | 1472 | sens | 76 | 63 | | NCBI | GCF_008831285.2 |
| | | NymcoGNL2a LOC116255565 (chr6, Mb 6.1) | 1396 | sens | | | 50 | NCBI | GCF_008831285.2 |
| | | NymcoGNL2b LOC116252701 (chr 1, Mb 27.4) | 1388 | sens | | | 50 | NCBI | GCF_008831285.2 |
| Nymphaeales (Nymphaeaceae) | Nymphaea thermarum | NymthGNOM_ EJ110_NYTH52200 (sc1144, kb 25) | 1472 | sens | 76 | 63 | | NCBI | GCA_011799765.1 |
| | | NymthGNL2a EJ110_NYTH30733 (sc328, kb 235) | 1396 | sens | | | 55 | NCBI | GCA_011799765.1 |
| | | NymthGNL2b EJ110_NYTH30720 (sc328, kb 180) | 1394 | sens | | | 50 | NCBI | GCA_011799765.1 |
| | | NymthGNL2c EJ110_NYTH06403 (sc23, kb 395) | 1391 | sens | | | 51 | NCBI | GCA_011799765.1 |
| Nymphaeales (Nymphaeaceae) | Euryale ferox | EurfeGNOMa g30206 (chr. 4, Mb 1.3) | 1467 | sens | 77 | 63 | | CoGe | Euryale ferox |
| | | EurfeGNOMb g23146 (chr. 2, Mb 1.5) | 1465 | sens | 76 | 63 | | CoGe | Euryale ferox |
| | | EurfeGNOMc g04013 (chr. 3, Mb 6.1) | 1467 | sens | 75 | 63 | | CoGe | Euryale ferox |

| | | | | | | | | | |
|---|---|---|---|---|---|---|---|---|---|
| | | EurfeGNL2a g32575 (chr. 14, Mb 5.8) | 1396 | sens | | | 49 | CoGe | Euryale ferox |
| | | EurfeGNL2b g16136 (chr. 12, Mb 0.2) | 1388 | sens | | | 51 | CoGe | Euryale ferox |
| **Magnoliids** | | | | | | | | | |
| Piperales (Saururaceae) | Saururus chinensis | SauchGNOM Sc010_0661 (sc10, Mb 6.6) | 1448 | res | 77 | 62 | | GWH | Pep file |
| | | SauchGNL2 Sc008_1622 (sc8, Mb 13.9) | 1392 | sens | | | 59 | GWH | Pep file |
| Piperales (Piperaceae) | Piper nigrum | PipniGNOMa Pn8.273 | 1458 | sens | 76 | 62 | | cotton.hzau.edu.cn | Pep file |
| | | PipniGNOMb Pn8.277 | 1458 | sens | 76 | 62 | | cotton.hzau.edu.cn | Pep file |
| | | PipniGNL2 Pn6.1394 | 1391 | sens | | | 53 | cotton.hzau.edu.cn | Pep file |
| Piperales (Aristolochiaceae) | Aristolochia fimbriata | ArifiGNOM H6P81_013188 (chr 4, 20.1) | 1433 | sens | 78 | 63 | | NCBI | GCA_019845555.1 |
| | | ArifiGNL2a H6P81_020970 (chr 7, Mb 21.1) | 1386 | sens | | | 54 | NCBI | GCA_019845555.1 |
| | | ArifiGNL2b H6P81_003805 (chr 1, Mb 40.1) | 1362 | sens | | | 52 | NCBI | GCA_019845555.1 |
| Piperales (Aristolochiaceae) | Aristolochia contorta | AricoGNOM EVM0013862 (LG3, Mb 25.1) | 1433 | sens | 78 | 62 | | CoGe | Aristolochia contorta |
| | | AricoGNL2a EVM0013422 (LG3, Mb 7.0) | (1373) | res | | | 51 | CoGe | Aristolochia contorta |
| | | AricoGNL2b EVM0004522 (LG1, Mb 30.7) | (1497) | sens | | | 50 | CoGe | Aristolochia contorta |
| Magnoliales (Magnoliaceae) | Liriodendron chinense | LirchGNOMa (Sc32, Mb 8.6) | 1455 | sens | 78 | 63 | | NCBI | GCA_003013855.2 |

| | | | | | | | | | |
|---|---|---|---|---|---|---|---|---|---|
| | | LirchGNOMb (ctg60, Mb 5.8) | 1455 | sens | 78 | 63 | | NCBI | GCA_0030 13855.2 |
| | | LirchGNL2 (ctig1255, Mb 0.4) | 1392 | sens | | | 61 | NCBI | GCA_0030 13855.2 |
| Magnoliales (Magnoliaceae) | Magnolia biondii | MagbiGNOM MBI03446 (chr 05, Mb 106.1) | 1455 | sens | 78 | 62 | | Dryad | Pep file |
| | | MagbiGNL2 MBI11792 (chr 01, Mb 53.9) | 1391 | sens | | | 61 | Dryad | Pep file |
| Magnoliales (Magnoliaceae) | Magnolia officinalis | MagofGNOM (group 5, Mb 93.6) | 1455 | sens | 78 | 62 | | NCBI | GCA_0210 15155.1 |
| | | MagofGNL2 (group 1, Mb 92.6) | 1391 | sens | | | 61 | NCBI | GCA_0210 15155.1 |
| Magnoliales (Magnoliaceae) | Magnolia sinica | MagsiGNOM LOC131243939 (chr 4, Mb 113.9) | 1455 | sens | 78 | 62 | | NCBI | GCF_0299 62835.1 |
| | | MagsiGNL2 LOC131254124 (chr 1, Mb 111.5) | 1391 | sens | | | 62 | NCBI | GCF_0299 62835.1 |
| Magnoliales (Annonaceae) | Annona cherimola | AnnchGNOM Ach24280 (chr5, kb 101) | 1452 | sens | 77 | 61 | | NCBI | GCA_0304 90835.1 |
| | | AnnchGNL2 Ach26572 (chr5, Mb 28.1) | 1387 | sens | | | 59 | NCBI | GCA_0304 90835.1 |
| Canellales (Canellaceae) | Warburgia ugandensis | WarugGNOMa wug10312 (chr4, Mb 50.7) | 1468 | sens | 79 | 63 | | NCBI | GCA_0352 36585.1 |
| | | WarugGNOMb wug4143 (chr10, Mb 5.9) | 1437 | res | 78 | 62 | | NCBI | GCA_0352 36585.1 |
| | | WarugGNL2 wug9485 (chr14, Mb 45.2) | 1398 | sens | | | 60 | NCBI | GCA_0352 36585.1 |
| Laurales (Lauraceae) | Persea americana | PeramGNOM MRB53_001684 (chr1, Mb 27.4) | 1441 | sens | 77 | 62 | | NCBI | GCA_0298 52735.1 |
| | | PeramGNL2 MRB53_017315 (chr5, Mb 57.5) | 1378 | sens | | | 61 | NCBI | GCA_0298 52735.1 |

| | | | | | | | | | |
|---|---|---|---|---|---|---|---|---|---|
| Laurales (Lauraceae) | Cinnamomum micranthum | CinmiGNOM CKAN_00834500 (sc0003, Mb 25.0) | 1453 | res | 77 | 62 | | NCBI | GCA_0035 46025.1 |
| | | CinmiGNL2 CKAN_01177000 (sc0004, Mb 24.4) | 1377 | sens | | | 61 | NCBI | GCA_0035 46025.1 |
| Laurales (Calycanthaceae) | Chimonanthus salicifolius | ChisaGNOMa (chr 9, Mb 59.9) | 1444 | sens | 77 | 61 | | NCBI | GCA_0133 50335.1 |
| | | ChisaGNOMb (chr 9, Mb 13.9) | 1455 | res | 70 | 59 | | NCBI | GCA_0133 50335.1 |
| | | ChisaGNL2 (chr 4, Mb 71.8) | 1390 | sens | | | 61 | NCBI | GCA_0133 50335.1 |
| Laurales (Calycanthaceae) | Chimonanthus praecox | ChiprGNOMa (chr8, Mb 28.1) | 1444 | sens | 77 | 61 | | NCBI | GCA_0221 13865.1 |
| | | ChiprGNOMb (chr8, Mb 19.0) | 1452 | res | 71 | 59 | | NCBI | GCA_0221 13865.1 |
| | | ChiprGNL2 (chr4, Mb 64.1) | 1375 | sens | | | 61 | NCBI | GCA_0221 13865.1 |
| **Chloranthales (sister to magnoliids)** | | | | | | | | | |
| Chloranthales (Chloranthaceae) | Chloranthus spicata | ChlspGNOMa Cspi18331 (sc 7, Mb 8.0) | 1469 | res | 76 | 63 | | CNGB | GWHBFSJ 00000000 |
| | (GWH Nov 21) | ChlspGNOMb Cspi14874 (sc 7, Mb 7.8) | 1405 | res | 72 | 59 | | CNGB | GWHBFSJ 00000000 |
| | | (no GNL2) | | | | | | | |
| Chloranthales (Chloranthaceae) | Chloranthus sessilifolius | ChlseGNOMa (ctg 15.725) (LG13, Mb 43.1) | 1470 | sens | 78 | 64 | | NCBI | GCA_0210 18995.1 |
| | | ChlseGNOMb (ctg 22.381) (LG9, Mb 109.3) | 1474 | sens | 76 | 62 | | NCBI | GCA_0210 18995.1 |
| | | ChlseGNL2a (ctg 19.311) (LG12, Mb 68.5) | 1391 | sens | | | 59 | NCBI | GCA_0210 18995.1 |
| | | ChlseGNL2b (ctg 9.1129+9.1128) | 1295 | sens | | | 57 | NCBI | GCA_0210 18995.1 |

| | | | | | | | | | |
|---|---|---|---|---|---|---|---|---|---|
| | | (LG6, Mb 80.4) | | | | | | | |
| **Ceratophyllales (sister to eudicots)** | | | | | | | | | |
| Ceratophyllales (Ceratophyllaceae) | Ceratophyllum demersum | CerdeGNOMa prefix20156 (chr. 3, Mb 23.1) | 1449 | sens | 75 | 62 | | CoGe | Ceratophyll um demersum |
| | | CerdeGNOMb prefix25991 (chr. 9, Mb 31.8) | 1387 | sens | 73 | 59 | | CoGe | Ceratophyll um demersum |
| | | CerdeGNL2 prefix10541 (chr. 3, Mb 84.4) | 1377 | sens | | | 56 | CoGe | Ceratophyll um demersum |

**Suppl. Table S1E. GBF1-type ARF-GEFs in monocots (10 orders, 26 families, 52 species)**

| Order | Species | Gene name (chr/ctg/sc #) | Protein Size (# aa) | BFA res / sens | % id. AtGN | % id. AtGNL1 | % id. AtGNL2 | database | ref |
|---|---|---|---|---|---|---|---|---|---|
| Acorales (Acoraceae) | Acorus americanus | AcoamGNOMa Acora.06G147100 (chr6, Mb 18.0) | 1474 | sens | 70 | 57 | | Phytozome 13 | A.americanus v1.1 |
| | | AcoamGNOMb Acora.03G000500 (chr3, Mb 0.1) | 1484 | res | 69 | 56 | | Phytozome 13 | A.americanus v1.1 |
| | | AcoamGNL2 Acora.12G089500 (chr12, Mb 18.2) | 1451 | sens | | | 51 | Phytozome 13 | A.americanus v1.1 |
| Acorales (Acoraceae) | Acorus gramineus | AcogrGNOMa DACA020116 (chr81, kb 130) | 1474 | sens | 69 | 56 | | NCBI | GCA_030737 835.1 |
| | | AcogrGNOMb DACA006115 (chr5, Mb 26.7) | 1476 | res | 68 | 56 | | NCBI | GCA_030737 835.1 |
| | | AcogrGNL2 DACA005488 (chr12, Mb 7.7) | 1454 | sens | | | 51 | NCBI | GCA_030737 835.1 |
| Acorales (Acoraceae) | Acorus calamus | AcocaGNOMa B17g01428 (chr17, Mb 19.7) | 1470 | sens | 70 | 57 | | CNGB | GWH pep file |
| | | AcocaGNOMb A05g00509 (chr5, Mb 4.6) | 1474 | sens | 70 | 56 | | CNGB | GWH pep file |
| | | AcocaGNOMc B04g02020 (chr4, Mb 29.7) | 1470 | res | 69 | 56 | | CNGB | GWH pep file |
| | | AcocaGNL2a B22g00940 (chr22, Mb 17.7) | 1432 | sens | | | 50 | CNGB | GWH pep file |
| | | AcocaGNL2b A10g00534 (chr 10, Mb 5.5) | 1074 | sens | | | 52 | CNGB | GWH pep file |
| Alismatales (Araceae) | Spirodela polyrhiza | SpipoGNOM Spipo3G0098500 (chr1, Mb 1.2) | 1448 | sens | 75 | 64 | | NCBI | GCA_900492 545.1 |

| | | | | | | | | | |
|---|---|---|---|---|---|---|---|---|---|
| | | SpipoGNL2<br>(chr 8, Mb 7.6) | 1400 | sens | | | 54 | NCBI | GCA_900492<br>545.1 |
| Alismatales<br>(Araceae) | Spirodela<br>intermedia | SpiinGNOM<br>SI8410_01000257<br>(chr 1, Mb 1.0) | 1447 | sens | 74 | 62 | | NCBI | GCA_902729<br>315.2 |
| | | SpiinGNL2<br>SI8410_09013719<br>(chr 9, Mb 12.4) | 1415 | | | | 57 | NCBI | GCA_902729<br>315.2 |
| Alismatales<br>(Araceae) | Colocasia<br>esculenta | ColesGNOMa<br>(chr1, Mb 23.3) | (1460) | sens | 77 | 62 | | NCBI | GCA_014218<br>235.1 |
| | | ColesGNOMb<br>(chr8, Mb 170.1) | (1445) | res | 77 | 63 | | NCBI | GCA_014218<br>235.1 |
| | | ColesGNL2<br>(chr2, Mb 15.8) | (1373) | sens | | | 55 | NCBI | GCA_014218<br>235.1 |
| Alismatales<br>(Araceae) | Lemna minuta | LemmuGNOM<br>(chr11, Mb 15.7) | 1445 | sens | 71 | 61 | | NCBI | GCA_024174<br>645.1 |
| | | LemmuGNL2<br>(chr2, Mb 22.6) | 1375 | sens | | | 54 | NCBI | GCA_024174<br>645.1 |
| Alismatales<br>(Araceae) | Wolffia<br>australiana | WolauGNOM (chr8,<br>Mb 15.1) | 1429 | sens | 70 | 61 | | NCBI | GCA_029677<br>425.1 |
| | | WolauGNL2 (chr5,<br>Mb 21.1) | | sens | | | | NCBI | GCA_029677<br>425.1 |
| Alismatales<br>(Juncaginacae) | Triglochin<br>maritima | TrimaGNOMa<br>(chr21, Mb 23.8) | 1409 | sens | 76 | 63 | | NCBI | GCA_963682<br>175.1 |
| | | TrimaGNOMb<br>(chr23, Mb 24.2) | 1409 | sens | 75 | 63 | | NCBI | GCA_963682<br>175.1 |
| | | TrimaGNOMc<br>(chr10, Mb 0.5) | 1420 | sens | 70 | 60 | | NCBI | GCA_963682<br>175.1 |
| | | TrimaGNL2 (chr11,<br>Mb 8.2) | | | | | n.a. | NCBI | GCA_963682<br>175.1 |
| Alismatales<br>(Cymodoceaceae) | Amphibolis<br>antarctica | AmpanGNOMa<br>(scf7180000526928) | 1437 | sens | 74 | 62 | | NCBI | GCA_024699<br>675.1 |
| | | AmpanGNOMb<br>(scf7180000526361) | 1438 | res | 72 | 60 | | NCBI | GCA_024699<br>675.1 |
| | | AmpanGNL2<br>(scf7180000530710) | 1376 | sens | | | 55 | NCBI | GCA_024699<br>675.1 |

| | | | | | | | | | |
|---|---|---|---|---|---|---|---|---|---|
| Alismatales (Cymodoceaceae) | Cymodocea nodosa | CymnoGNOMa (chr5, Mb 17.5) | 1437 | sens | 74 | 62 | | NCBI | GCA_036874 025.1 |
| | | CymnoGNOMb (chr2, Mb 2.3) | 1441 | res | 71 | 58 | | NCBI | GCA_036874 025.1 |
| | | CymnoGNL2 (chr11, Mb 0.7) | | | | | n.a. | NCBI | GCA_036874 025.1 |
| Alismatales (Posidoniaceae) | Posidonia australis | PosauGNOMa (scf7180004758438) | 1441 | sens | 76 | 62 | | NCBI | GCA_024699 735.1 |
| | | PosauGNOMb (scf7180003738803) | 1436 | res | 73 | 61 | | NCBI | GCA_024699 735.1 |
| | | PosauGNL2 (scf7180003767994) | 1409 | sens | | | 56 | NCBI | GCA_024699 735.1 |
| Alismatales (Zosteraceae) | Zostera marina | Zosma_108g00240 (sc108) | 1413 | sens | 70 | 59 | | NCBI | GCA_001185 155 |
| | | Zosma_56g00900 (sc56) | 1415 | sens | 69 | 57 | | NCBI | GCA_001185 155 |
| | | Zosma_123g00350.1 (sc123) | 1443 | sens | 67 | 59 | | NCBI | GCA_001185 155 |
| | | Zosma_111g00160-150 (sc111) Zosma_111g00160+ 00150 | 1357 | sens | | | 54 | NCBI | GCA_001185 155 |
| Dioscoreales (Dioscoreaceae) | Dioscorea rotundata | Dioro_LOC12026704 2 (chr8) (Dioro_chr8_525232 9) | 1455 | sens | 75 | 62 | | NCBI | GCA_009730 915.2 |
| | | Dioro_LOC12025450 5 (Dioro_chr4_593013 5) | 1399 | sens | | | 58 | NCBI | GCA_009730 915.2 |
| Dioscoreales (Dioscoreaceae) | Dioscorea alata | Dioal.08G121600 | 1422 | sens | 75 | 62 | | NCBI | GCA_020875 875.1 |
| | | Dioal.04G114500 | 1379 | sens | | | 58 | NCBI | GCA_020875 875.1 |
| Dioscoreales (Dioscoreaceae) | Dioscorea dumetorum | DioduGNOM | (1470) | sens | 73 | 65 | | NCBI | GCA_902712 375.1 |

| | | | | | | | | | |
|---|---|---|---|---|---|---|---|---|---|
| | | DioduGNL2 | (1380) | sens | | | 55 | NCBI | GCA_902712 375.1 |
| Dioscoreales (Nartheciaceae) | Narthecium ossifragum | NarosGNOM (chr11, Mb 3.8) | 1456 | sens | 75 | 61 | | NCBI | GCA_963920 575.1 |
| | | NarosGNL2 (chr8, Mb 0.8) | 1413 | | | | 56 | NCBI | GCA_963920 575.1 |
| Pandanales (Velloziaceae) | Xerophyta viscosa | XerviGNOMa (sc0013, Mb 2.5) | 1422 | sens | 72 | 60 | | NCBI | GCA_002076 135.1 |
| | | XerviGNOMb (sc0007, Mb 0.3) (aa243-266 missing) | 1377 | sens | 65 | 57 | | NCBI | GCA_002076 135.1 |
| | | XerviGNOMc_sc0011 (aa239-266 missing) | 1420 | res | 63 | 55 | | NCBI | GCA_002076 135.1 |
| | | XerviGNL2_sc0029 (aa223-1276 wrong sequence) | ? | ? | | | 61 | NCBI | GCA_002076 135.1 |
| Pandanales (Velloziaceae) | Acanthochlamys bracteata | AcabrGNOMa (chr6, Mb 8.1) | 1434 | sens | 73 | 61 | | NCBI | GCA_019914 995.1 |
| | | AcabrGNOMb (chr1, Mb 9.7) | 1419 | res | 64 | 57 | | NCBI | GCA_019914 995.1 |
| | | AcabrGNL2 (chr13, Mb 1.8) | 1383 | sens | | | 57 | NCBI | GCA_019914 995.1 |
| Liliales (Melanthiaceae) | Chionographis japonica | ChijaGNOMa (chr2) | 1432 | sens | 77 | 62 | | NCBI | GCA_947650 365.1 |
| | | ChijaGNOMb (chr9) | 1437 | sens | 68 | 58 | | NCBI | GCA_947650 365.1 |
| | | ChijaGNL2 (chr 11) | 1395 | sens | | | 57 | NCBI | GCA_947650 365.1 |
| Asparagales (Orchidaceae) | Dendrobium catenatum | Denca_LOC1101056 64 (sc260) DencaGNOMa | 1445 | sens | 74 | 61 | | NCBI | GCA_001605 985.2 |
| | | Denca_LOC1101025 84 (sc637) DencaGNOMb | 1437 | sens | 68 | 58 | | NCBI | GCA_001605 985.2 |
| | | Denca_LOC1100929 41 (sc840) | 1402 | sens | | | 51 | NCBI | GCA_001605 985.2 |

| | | | | | | | | | |
|---|---|---|---|---|---|---|---|---|---|
| Asparagales (Orchidaceae) | Apostasia shenzhenica | AXF42_Ash010487 (sc5, Mb 3.0) AposhGNOMb | 1427 | res | 67 | 59 | | NCBI | GCA_002786 265.1 |
| | | AXF42_Ash011579 (sc16, Mb 2.2) reannotated AposhGNOMa | (1441) | sens | 73 | 61 | | NCBI | GCA_002786 265.1 |
| | | AXF42_Ash002502 (sc86) | 1396 | | | | 54 | NCBI | GCA_002786 265.1 |
| Asparagales (Orchidaceae) | Apostasia ramifera | AporaGNOMa (sc105) | (1460) | sens | 78 | 62 | | NCBI | GCA_019297 695.1 |
| | | AporaGNOMb (sc989) | (1445) | res | 68 | 57 | | NCBI | GCA_019297 695.1 |
| | | AporaGNL2 (sc10) | (1393) | sens | | | 54 | NCBI | GCA_019297 695.1 |
| Asparagales (Orchidaceae) | Vanilla planifolia | HPP92_015831 (chr7) | (1450) | sens | 76 | 65 | | NCBI | GCA_016413 895.1 |
| | | (chr 7) (fragments, pseudo?) | (1440) | sens | 45 | 37 | | NCBI | GCA_016413 895.1 |
| | | HPP92_015832 (chr7) | (1450) | sens | 85 | 65 | | NCBI | GCA_016413 895.1 |
| | | HPP92_002719 (chr1) | (1370) | sens | | | 55 | NCBI | GCA_016413 895.1 |
| Asparagales (Iridaceae) | Iris pallida | IripaGNOMa M6B38_209310 ssc1000940 | 1420 | sens | 74 | 62 | | NCBI | GCA_029216 955.1 |
| | | IripaGNOMb M6B38_370335 ssc42672 | 1420 | sens | 74 | 62 | | NCBI | GCA_029216 955.1 |
| | | IripaGNOMc M6B38_399180 ssc39381 | 1416 | sens | 73 | 61 | | NCBI | GCA_029216 955.1 |
| | | IripaGNOMd M6B38_124610 ssc44928 | 1415 | sens | 73 | 61 | | NCBI | GCA_029216 955.1 |
| | | IripaGNL2a (not annotated) ssc39659 | 1379 | sens | | | 56 | NCBI | GCA_029216 955.1 |

| | | | | | | | | | |
|---|---|---|---|---|---|---|---|---|---|
| | | IripaGNL2b (not annotated) ssc41173 | 1353 | sens | | | 55 | NCBI | GCA_029216955.1 |
| Asparagales (Asparagaceae) | Asparagus officinalis | Aspof_LOC109850386 (chr 7) | 1435 | sens | 75 | 62 | | NCBI | GCF_001876935.1 |
| | | Aspof_LOC109851574 (chr 8) | 1356 | sens | | | 56 | NCBI | GCF_001876935.1 |
| Asparagales (Asparagaceae) | Asparagus setaceus | AspseGNOM_OF25554 | 1425 | sens | 76 | 64 | | NCBI | GCA_012295165.1 |
| | | AspseGNL2_OF18517 | 1407 | sens | | | 57 | NCBI | GCA_012295165.1 |
| Arecales (Arecacae) | Phoenix dactylifera | Phoda_LOC103696334 (chr 11, GNOMa) chr11T0156900 | 1454 | sens | 76 | 62 | | NCBI | GCF_009389715.1 |
| | | Phoda_LOC103712126 (chr 16, GNOMb) chr16T0076100 | 1451 | res | 74 | 62 | | NCBI | GCF_009389715.1 |
| | | Phoda_LOC103695708 (Phoda_GNL2a) 000276FT0001900 | 1365 | sens | | | 56 | NCBI | GCF_009389715.1 |
| | | Phoda_LOC120105401 (Phoda_GNL2b) | 1405 | sens | | | 58 | NCBI | GCF_009389715.1 |
| Arecales (Arecacae) | Calamus simplicifolius | CalsiGNOMa (sc2304) | (1454) | sens | 78 | 62 | | NCBI | GCA_900491605.1 |
| | | CalsiGNOMb (sc2554) | (1454) | res | 78 | 63 | | NCBI | GCA_900491605.1 |
| | | CalsiGNL2a (sc2274) | (1393) | sens | | | 57 | NCBI | GCA_900491605.1 |
| | | CalsiGNL2b (sc176) | (1420) | sens | | | 61 | NCBI | GCA_900491605.1 |
| Arecales (Arecacae) | Metroxylon sagu | MetsaGNOMa (ctg 8250) | 1437 | sens | 76 | 62 | | NCBI | GCA_017589505.3 |
| | | MetsaGNOMb (ctg 4670) | 1454 | res | 75 | 62 | | NCBI | GCA_017589505.3 |
| | | MetsaGNL2 (ctg 4500) | 1390 | sens | | | 59 | NCBI | GCA_017589505.3 |
| Arecales (Arecacae) | Elaeis guineensis | Elagu_LOC105034005 | 1446 | res | 75 | 62 | | NCBI | GCF_000442705.1 |

| | | | | | | | | | | |
|---|---|---|---|---|---|---|---|---|---|---|
| | | GNOMa (sc00415) | | | | | | | | |
| | | Elagu_LOC105036111<br>GNOMb (sc02793) | 1454 | sens | 76 | 62 | | NCBI | | GCF_000442705.1 |
| | | Elagu_LOC105046398<br>GNL2 (chr 1) | 1396 | sens | | | 59 | NCBI | | GCF_000442705.1 |
| Zingiberales (Musaceae) | Musa acuminata | Musac_LOC103988820 (chr 6) | 1441 | sens | 71 | 59 | | NCBI | | GCF_000313855.2 |
| | | Musac_LOC103986838 (chr 6) | 1445 | sens | 75 | 61 | | NCBI | | GCF_000313855.2 |
| | | Musac_LOC103990130 (chr 1) | 1367 | sens | | | 56 | NCBI | | GCF_000313855.2 |
| Zingiberales (Musaceae) | Ensete glaucum | EnsglGNOMa<br>(chr eg08, Mb 41.8) | 1442 | sens | 75 | 61 | | NCBI | | GCA_021527575.1 |
| | | EnsglGNOMb<br>(chr eg08, Mb 54.7) | 1442 | sens | 70 | 60 | | NCBI | | GCA_021527575.1 |
| | | EnsglGNL2 (chr eg01) | 1368 | sens | | | 57 | NCBI | | GCA_021527575.1 |
| Zingiberales (Zingiberaceae) | Zingiber officinale | LOC121984128 (chr1B)<br>ZinofGNOMaB | 1432 | sens | 73 | 60 | | NCBI | | GCA_018446385.1 |
| | | LOC122038315 (chr1A)<br>ZinofGNOMaA | 1431 | sens | 73 | 60 | | NCBI | | GCA_018446385.1 |
| | | LOC122014938 (chr8B)<br>ZinofGNOMcB | 1430 | sens | 71 | 60 | | NCBI | | GCA_018446385.1 |
| | | LOC122005619 (chr7B)<br>ZinofGNOMbB | 1424 | sens | 69 | 59 | | NCBI | | GCA_018446385.1 |
| | | LOC122001327 (chr7A)<br>ZinofGNOMbA | 1424 | sens | 69 | 59 | | NCBI | | GCA_018446385.1 |
| | | LOC122012704 (chr8A)<br>ZinfGNOMcA | 1428 | sens | 71 | 60 | | NCBI | | GCA_018446385.1 |

| | | | | | | | | | |
|---|---|---|---|---|---|---|---|---|---|
| | | LOC121988887 (chr6B) ZinofGNOMdB | 1423 | res | 67 | 58 | | NCBI | GCA_018446 385.1 |
| | | LOC122048561 (chr2B) ZinofGNL2b | 1358 | sens | | | 55 | NCBI | GCA_018446 385.1 |
| | | LOC122043976 (chr2A) ZinofGNL2a | 1358 | sens | | | 55 | NCBI | GCA_018446 385.1 |
| Zingiberales (Zingiberaceae) | Wurfbainia villosa | WurviGNOMa (chr4, Mb 14.5) | 1431 | sens | 73 | 60 | | NCBI | GCA_025175 895.1 |
| | | WurviGNOMb (chr19, Mb 15.5) | 1429 | res | 73 | 60 | | NCBI | GCA_025175 895.1 |
| | | WurviGNOMc (chr20, Mb 6.0) | 1423 | sens | 70 | 60 | | NCBI | GCA_025175 895.1 |
| | | WurviGNOMd (chr23, Mb 29.4) | 1423 | sens | 69 | 59 | | NCBI | GCA_025175 895.1 |
| | | WurvoGNL2 (chr11, Mb 58.8) | 1366 | sens | | | 55 | NCBI | GCA_025175 895.1 |
| Zingiberales (Costaceae) | Costus lasius | CoslaGNOMa (chr9, Mb 98.9) | 1443 | sens | 75 | 61 | | NCBI | GCA_027563 935.2 |
| | | CoslaGNOMb (chr7, Mb 1.8) | 1450 | sens | 68 | 59 | | NCBI | GCA_027563 935.2 |
| | | CoslaGNOMc (chr4, Mb 6.9) | 1450 | res | 66 | 57 | | NCBI | GCA_027563 935.2 |
| | | CoslaGNL2 (chr5, Mb 8.5) | 1372 | sens | | | 55 | NCBI | GCA_027563 935.2 |
| Zingiberales (Costaceae) | Costus pulverulentus | CospuGNOMa (chr9, Mb 40.1) | 1443 | sens | 75 | 61 | | NCBI | GCA_027562 315.1 |
| | | CospuGNOMb (chr7, Mb 1.7) | 1450 | sens | 68 | 59 | | NCBI | GCA_027562 315.1 |
| | | CospuGNOMc (chr4, Mb 6.2) | 1450 | res | 66 | 57 | | NCBI | GCA_027562 315.1 |
| | | CospuGNL2 (chr5, Mb 8.3) | 1373 | sens | | | 55 | NCBI | GCA_027562 315.1 |
| Zingiberales (Marantaceae) | Thalia dealbata | ThadeGNOMa (chr6, Mb 25.6) | 1443 | sens | 75 | 61 | | NCBI | GCA_030544 415.1 |

| | | | | | | | | | |
|---|---|---|---|---|---|---|---|---|---|
| | | ThadeGNOMb (chr6, Mb 22.0) | 1422 | sens | 70 | 59 | | NCBI | GCA_030544 415.1 |
| | | ThadeGNL2 (chr6, Mb 17.5) | 1368 | sens | | | 54 | NCBI | GCA_030544 415.1 |
| Zingiberales (Cannaceae) | Canna indica | CaninGNOMa (chr1, Mb 78.8) Cni_G02893 | 1443 | sens | 75 | 62 | | NCBI | GCA_034359 265.1 |
| | | CaninGNOMb (chr5, Mb 78.6) Cni_G17668 | 1435 | sens | 70 | 61 | | NCBI | GCA_034359 265.1 |
| | | CaninGNL2 (chr1, Mb 23.8) Cni_G01726 | 1370 | sens | | | 55 | NCBI | GCA_034359 265.1 |
| Commelinales (Pontederiaceae | Pontederia paniculata | PonpaGNOMa (sc1340) | (1407) | sens | 72 | 61 | | NCBI | GCA_001647 135.1 |
| | | PonpaGNOMb (sc304) | 1444 | sens | 74 | 62 | | NCBI | GCA_001647 135.1 |
| | | PonpaGNL2 (sc168) | 1390 | sens | | | 56 | NCBI | GCA_001647 135.1 |
| Commelinales (Pontederiaceae) | Pontederia cordata | PoncoGNOMa 022494 | 1444 | sens | 74 | 62 | | Figshare | Pep file |
| | | PoncoGNOMb 018924 | 1453 | sens | 73 | 62 | | Figshare | Pep file |
| | | PoncoGNL2 005405 | 1396 | sens | | | 55 | Figshare | Pep file |
| Poales (Typhaceae) | Typha latifolia | TyplaGNOMa tig00027959 (chr 10) | 1440 | sens | 74 | 61 | | NCBI | GCA_019914 945.1 |
| | | TyplaGNOMb tig00027902 (chr 4) | 1438 | sens | 69 | 59 | | NCBI | GCA_019914 945.1 |
| | | TyplaGNL2 tig00000870 (chr 11) | 1391 | sens | | | 58 | NCBI | GCA_019914 945.1 |
| Poales (Bromeliaceae) | Puya raimondii | PuyraGNOMa (sc4) | (1460) | sens | 74 | 59 | | NCBI | GCA_016920 855.1 |
| | | PuyraGNOMb (sc4) | (1450) | res | 67 | 60 | | NCBI | GCA_016920 855.1 |
| | | PuyraGNL2 (sc2) | (1390) | sens | | | 54 | NCBI | GCA_016920 855.1 |

| | | | | | | | | | |
|---|---|---|---|---|---|---|---|---|---|
| Poales (Bromeliaceae) | Ananas comosus | AnacoGNOMa LOC109712899 (chr7) | 1441 | sens | 68 | 56 | | NCBI | GCF_001540 865.1 |
| | | AnacoGNOMb LOC109718116, (chr12) | 1464 | res | 64 | 57 | | NCBI | GCF_001540 865.1 |
| | | AnacoGNL2 LOC109722123, (chr16) | 1419 | sens | | | 56 | NCBI | GCF_001540 865.1 |
| Poales (Juncaceae) | Juncus effusus | JunefGNOMa LUZ60_004684 (chr 13, Mb 6.8) | 1396 | sens | 59 | 52 | | NCBI | GCA_027726 005.1 |
| | | JunefGNOMb LUZ60_005372 (chr 14, Mb 8.9) | 1424 | sens | 69 | 56 | | NCBI | GCA_027726 005.1 |
| | | JunefGNL2 LUZ60_000419 (chr 1, Mb 7.5) | | | | | 48 | NCBI | GCA_027726 005.1 |
| Poales (Cyperaceae) | Kobresia (Carex) littledalei | KobliGNOM FCM35_KLT10238 (chr 20, Mb 0.9) | 1431 | sens | 68 | 57 | | NCBI | GCA_011114 355.1 |
| | | KobliGNL2 FCM35_KLT07006 (chr 16, Mb 10.6) | 1369 | sens | | | 49 | NCBI | GCA_011114 355.1 |
| Poales (Cyperaceae) | Cyperus esculentus | CypesGNOMb (sc19, Mb 0.6) | 1421 | sens | 60 | 53 | | NCBI | GCA_030179 965.1 |
| | | CypesGNOMa (sc2, Mb 5.9) | 1414 | sens | 68 | 57 | | NCBI | GCA_030179 965.1 |
| | | CypesGNL2 (sc5, Mb 7.3) | 1369 | sens | | | 48 | NCBI | GCA_030179 965.1 |
| Poales (Joinvilleaceae) | Joinvillea ascendens | Joasc.18G020200 (chr18) | 1424 | sens | 71 | 58 | | Phytozome 13 | V1.1 |
| | | Joasc.04G090200 (chr4) | 1432 | res | 65 | 56 | | Phytozome 13 | V1.1 |
| | | Joasc.09G111700 (chr9) | 1408 | sens | | | 54 | Phytozome 13 | V1.1 |
| Poales (Poaceae) | Pharus latifolius | PhalaGNOMa Pl03g27650 (chr3, Mb 81.9) | 1428 | sens | 70 | 57 | | NCBI | GCA_019359 835.1 |

| | | | | | | | | | |
|---|---|---|---|---|---|---|---|---|---|
| | | PhalaGNOMb Pl09g17940 (chr9, Mb 58.6) | 1426 | res | 63 | 56 | | NCBI | GCA_019359 835.1 |
| | | PhalaGNL2 Pl01g16080 (chr1, Mb 23.5) | 1415 | sens | | | 54 | NCBI | GCA_019359 835.1 |
| Poales (Poaceae) | Oryza sativa | OrysaGNOMa LOC9271296 (chr 3, 27.0) | 1410 | sens | 70 | 56 | | NCBI | GCF_034140 825.1 |
| | | OrysaGNOMb LOC4329190 (chr 2, Mb 13.1) | 1424 | sens | 62 | 55 | | NCBI | GCF_034140 825.1 |
| | | OrysaGNL2 LOC4334954 (chr 4, Mb 1.1) | 1407 | sens | | | 54 | NCBI | GCF_034140 825.1 |
| Poales (Poaceae) | Brachypodium distachyon | BradiGNOMa LOC100840992 (chr 1, Mb9.7) | 1407 | sens | 69 | 56 | | NCBI | GCF_000005 505.3 |
| | | BradiGNOMb LOC100836075 (chr 1, 71.7) | 1412 | sens | 63 | 55 | | NCBI | GCF_000005 505.3 |
| | | BradiGNL2 LOC100825923 (chr 2, 48.3) | 1392 | sens | | | 54 | NCBI | GCF_000005 505.3 |
| Poales (Poaceae) | Sorghum bicolor | SorbiGNOMa LOC8057184 (chr 1, Mb 11.5) | 1411 | sens | 69 | 56 | | NCBI | GCF_000003 195.3 |
| | | SorbiGNOMb LOC8055663 (chr 1, Mb 58.2) | 1426 | sens | 63 | 54 | | NCBI | GCF_000003 195.3 |
| | | SorbiGNL2 LOC8063286 (chr 8, Mb 47.6) | 1412 | sens | | | 52 | NCBI | GCF_000003 195.3 |

**Suppl. Table S1F. GBF1-type ARF-GEFs in basal eudicots (5 orders, 13 families, 26 species)**

| Order (family) | Species | Gene name (chr/ctg/sc #) | Protein Size (# aa) | BFA res / sens | % id. AtGN | % id. AtGNL1 | % id. AtGNL2 | database | ref |
|---|---|---|---|---|---|---|---|---|---|
| Ranunculales (Eupteleaceae) | Euptelea pleiosperma | EupplGNOMa (chr5, Mb 29.5) | 1461 | sens | 81 | 63 | | NCBI | GCA_0350484 95.1 |
| | | EupplGNOMb (chr1, Mb 23.8) | 1467 | res | 80 | 65 | | NCBI | GCA_0350484 95.1 |
| | | EupplGNL2 (chr10, Mb 31.2) | 1402 | sens | | | 64 | NCBI | GCA_0350484 95.1 |
| Ranunculales (Papaveraceae) | Eschscholzia californica | EsccaGNOM EscalH1.4G086700 | 1444 | sens | 80 | 62 | | Phytozome 13 | Eschscholzia californ HAP1 |
| | | EsccaGNL1 EscalH1.5G076100 | 1439 | res | 64 | 56 | | Phytozome 13 | Eschscholzia californ HAP1 |
| | | EsccaGNL2 EscalH1.5G227500 | 1390 | sens | | | 63 | Phytozome 13 | Eschscholzia californ HAP1 |
| Ranunculales (Papaveraceae) | Papaver somniferum | PapsoGNOMa LOC113307666 (chr9, Mb 168.7) | 1455 | sens | 80 | 63 | | NCBI | GCF_0035736 95.1 |
| | | PapsoGNOMb LOC113297613 (chr7, Mb 73.1) | 1436 | res | 64 | 56 | | NCBI | GCF_0035736 95.1 |
| | | PapsoGNL2 LOC113316139 (chr1, Mb 218.1) | 1388 | sens | | | 61 | NCBI | GCF_0035736 95.1 |
| Ranunculales (Papaveraceae) | Macleaya cordata | MaccoGNOMa BVC80_1831g8 (sc1831, kb 35) | 1453 | sens | 81 | 65 | | NCBI | GCA_0021747 75.1 |
| | | MaccoGNOMb BVC80_9095g21 (sc9095, kb 155) | 1436 | res | 66 | 57 | | NCBI | GCA_0021747 75.1 |
| | | MaccoGNL2 BVC80_1789g18 (sc1789, kb 245) | 1392 | sens | | | 63 | NCBI | GCA_0021747 75.1 |
| Ranunculales (Papaveraceae) | Corydalis tomentella | CortoGNOMa ctg2A40T160 (chr 2, Mb 8.0) | 1444 | sens | 80 | 63 | | CNGB | GWHAORS00 000000 |
| | (GWH May 22) | CortoGNOMb ctg79R5T20 (chr 2, Mb 13.9) | 1531 | sens | 66 | 55 | | CNGB | GWHAORS00 000000 |

| | | | | | | | | | |
|---|---|---|---|---|---|---|---|---|---|
| | | CortoGNL2a ctg39A16T100 (chr 1, Mb 23.1) | 1399 | sens | | | 62 | CNGB | GWHAORS00 000000 |
| | | CortoGNL2b ctg18A14T148 (chr 5, Mb 2.0) | 1388 | sens | | | 63 | CNGB | GWHAORS00 000000 |
| Ranunculales (Lardizabalaceae) | Akebia trifoliata | AketrGNOMa (chr 14, Mb 1.3) | 1455 | sens | 80 | 63 | | NCBI | GCA_0179794 45.1 |
| | | AketrGNOMb (chr 11, Mb 40.3) | 1415 | res | 67 | 57 | | NCBI | GCA_0179794 45.1 |
| | | AketrGNL2 (chr 10, Mb 0.4) | 1393 | sens | | | 62 | NCBI | GCA_0179794 45.1 |
| Ranunculales (Circaeasteraceae) | Kingdonia uniflora | KinunGNOMa GIB67_030821 (sc_K1406, Mb 3.3) | 1441 | sens | 78 | 63 | | NCBI | GCA_0140581 05.1 |
| | | KinunGNOMb GIB67_019872 (sc_K629, Mb 4.1) | 1446 | res | 78 | 63 | | NCBI | GCA_0140581 05.1 |
| | | KinunGNL2 GIB67_020572 (sc_K1706, Mb 1.1) | 1406 | sens | | | 64 | NCBI | GCA_0140581 05.1 |
| Ranunculales (Berberidaceae) | Berberis thunbergii | BerthGNOMa (sc28F, kb 150) | (1460) | sens | 78 | 61 | | NCBI | GCA_0032901 65.1 |
| | | BerthGNOMb (sc333T, kb 190) | (1460) | res | 82 | 61 | | NCBI | GCA_0032901 65.1 |
| | | BerthGNOMc (sc211F, kb 235) | (1430) | res | 65 | 57 | | NCBI | GCA_0032901 65.1 |
| | | BerthGNL2 (sc1831F, kb 80) | (1390) | sens | | | 60 | NCBI | GCA_0032901 65.1 |
| Ranunculales (Berberidaceae) | Epimedium pubescens | EpipuGNOMa Ebr01G003690 (chr 1, Mb 29.8) | 1072 | sens | 80 | 62 | | CNGB | GWH pep file |
| | | EpipuGNOMb Ebr04G051320 (chr 4, Mb 402.3) | 1354 | sens | 64 | 57 | | CNGB | GWH pep file |
| | | No EpipuGNL2 | | | | | | | |
| Ranunculales (Ranunculaceae) | Aquilegia coerulea | AqucoGNOMa | 1456 | sens | 80 | 63 | | NCBI | GCA_0027385 05.1 |

| | | | | | | | | | |
|---|---|---|---|---|---|---|---|---|---|
| | | AQUOO_014_00263 (sc14, Mb 1.4) | | | | | | | |
| | | AqucaGNOMb AQUCO_045_00153 (sc45, Mb 1.1) | 1446 | res | 73 | 61 | | NCBI | GCA_002738505.1 |
| | | AqucaGNL2a AQUCO_007_00074 (sc7, Mb 0.6) | 1390 | sens | | | 63 | NCBI | GCA_002738505.1 |
| | | AqucaGNL2b AQUCO_020_00178 (sc20, Mb 1.3) | 1389 | sens | | | 63 | NCBI | GCA_002738505.1 |
| Ranunculales (Ranunculaceae | Aquilegia kansuensis | AqukaGNOMa (chr 6, Mb 1.4) | 1456 | sens | 80 | 63 | | NCBI | GCA_020826895.1 |
| | | AqukaGNOMb (chr 4, Mb 33.6) | 1446 | res | 73 | 61 | | NCBI | GCA_020826895.1 |
| | | AqukaGNL2a (chr5, Mb 11.9) | 1393 | sens | | | 63 | NCBI | GCA_020826895.1 |
| | | AqukaGNL2b (chr 7, Mb 37.6) | 1392 | sens | | | 63 | NCBI | GCA_020826895.1 |
| Ranunculales (Menispermaceae) | Tinospora cordifolia | TincoGNOMa (sc13802) | 1466 | sens | 81 | 64 | | NCBI | GCA_030144865.1 |
| | NCBI Jun23 | TincoGNOMb (sc14092) | 1441 | res | 63 | 56 | | NCBI | GCA_030144865.1 |
| | | TincoGNL2 (sc252, kb 50) | 1383 | sens | | | 59 | NCBI | GCA_030144865.1 |
| Ranunculales (Menispermaceae) | Stephania japonica | StejaGNOMa Sjap_007090 (chr2, Mb 74.5) | 1461 | sens | 80 | 63 | | NCBI | GCA_039657345.1 |
| | NCBI May 24 | StejaGNOMb Sjap_004118 (chr2, Mb 2.6) | 1439 | res | 63 | 56 | | NCBI | GCA_039657345.1 |
| | annotated | StejaGNL2 Sjap_011312 (chr4, Mb 37.1) | 1385 | sens | | | 57 | NCBI | GCA_039657345.1 |
| Proteales (Nelumbonaceae) | Nelumbo nucifera | NelnuGNOMa LOC104605370 (sc53, Mb 1.8) | 1469 | sens | 79 | 64 | | NCBI | GCF_000365185.1 |
| | (NCBI Aug 18) | NelnuGNOMb LOC104612314 | 1473 | sens | 78 | 62 | | NCBI | GCF_000365185.1 |

| | | | | | | | | | |
|---|---|---|---|---|---|---|---|---|---|
| | | (sc123, Mb 0.3) | | | | | | | |
| | | NelnuGNL2 LOC104585650 (sc142, Mb 0.6) | 1385 | sens | | | 61 | NCBI | GCF_000365185.1 |
| Proteales (Proteaceae) | Macadamia integrifolia | MacinGNOMa LOC122062215 (chr 2, Mb 17.6) | 1472 | sens | 81 | 63 | | NCBI | GCF_013358625.1 |
| | | MacinGNOMb LOC122067401 (sc289, kb 120) | 1436 | sens | 66 | 58 | | NCBI | GCF_013358625.1 |
| | | MacinGNL2 LOC122072250 (chr 2, Mb 34.4) | 1404 | sens | | | 62 | NCBI | GCF_013358625.1 |
| Proteales (Proteaceae) | Hakea prostrata | HakprGNOMa (chr7, Mb 6.7) | 1470 | sens | 81 | 63 | | NCBI | GCA_041222555.1 |
| | | HakprGNOMb (chr1, Mb 70.5) | 1432 | res | 65 | 58 | | NCBI | GCA_041222555.1 |
| | | HakprGNOMc (chr1, Mb 76.7) | 1434 | sens | 64 | 57 | | NCBI | GCA_041222555.1 |
| | | HakprGNL2 (chr11, Mb 0.6) | 1366 | sens | | | 56 | NCBI | GCA_041222555.1 |
| Proteales (Proteaceae) | Telopea speciosissima | TelspGNOMa LOC122667501 (chr 7, Mb 9.4) | 1472 | sens | 80 | 63 | | NCBI | GCF_018873765.1 |
| | | TelspGNOMb LOC122669738 (chr 7, Mb 50.3) | 1471 | sens | 77 | 61 | | NCBI | GCF_018873765.1 |
| | | TelspGNOMc LOC122662494 (chr 1, Mb 81.3) | 1441 | sens | 66 | 58 | | NCBI | GCF_018873765.1 |
| | | TelspGNL2a LOC122672147 (chr 8, Mb 21.0) | 1399 | sens | | | 62 | NCBI | GCF_018873765.1 |
| | | TelspGNL2b LOC122668736 (chr 7, Mb 58.8) | (1400) | sens | | | 56 | NCBI | GCF_018873765.1 |
| Proteales (Platanaceae) | Platanus orientalis | PlaorGNOMa (no DCB) (sc146347) | 1228 | sens | 80 | 63 | | NCBI | GCA_025762735.1 |

| | | | | | | | | | |
|---|---|---|---|---|---|---|---|---|---|
| | | PlaorGNOMb (sc288847) aa1441-1471 missing | 1440 | res | 81 | 63 | | NCBI | GCA_025762735.1 |
| | | PlaorGNOMc (no DCB) (sc3, Mb 7.9) | 1140 | sens | 82 | 64 | | NCBI | GCA_025762735.1 |
| | | PlaorGNL2 (sc1, Mb 1.1) | 1398 | sens | | | 63 | NCBI | GCA_025762735.1 |
| Proteales (Platanaceae) | Platanus acerifolia | PlaacGNOMa (ctgA6) PaA6G274240 | 1472 | sens | 81 | 64 | | Dryad | Pep file |
| | | PlaacGNOMb (ctgB6) PaB6G307410 | 1503 | sens | 78 | 63 | | Dryad | Pep file |
| | | PlaacGNOMc (ctgA6) PaA6G283070 | 1476 | res | 81 | 63 | | Dryad | Pep file |
| | | PlaacGNL2 (ctgA6) PaA6G283520 | 1398 | sens | | | 63 | Dryad | Pep file |
| Buxales (Buxaceae) | Buxus sinica | BuxsiGNOMa BUSI_023311 | 1486 | sens | 79 | 62 | | Dryad | Pep file |
| | | BuxsiGNOMb BUSI_016854 | 1440 | res | 64 | 56 | | Dryad | Pep file |
| | | BuxsiGNOMc BUSI_016855 (internal deletions) | 1255 | res | 58 | 47 | | Dryad | Pep file |
| | | BuxsiGNL2 BUSI_014142 | 1405 | res | | | 66 | Dryad | Pep file |
| Buxales (Buxaceae) | Buxus austro-yunnanensis | BuxauGNOMa GWHGBHNJ021904 (LG12, Mb 7.4) | 1467 | sens | 81 | 63 | | CNGB | GWH pep file |
| | | BuxauGNOMb GWHGBHNJ015817 (LG08, Mb 14.5) | 1440 | res | 64 | 56 | | CNGB | GWH pep file |
| | | BuxauGNL2 GWHGBHNJ019994 (LG11, Mb 21.4) | 1402 | res | | | 66 | CNGB | GWH pep file |
| Buxales (Buxaceae) | Buxus sempervirens | BuxseGNOMa (chr 12, Mb 7.9) | 1467 | sens | 81 | 63 | | NCBI | GCA_947561395.1 |
| | | BuxseGNOMb (chr 6, Mb 28.0) | 1440 | res | 64 | 56 | | NCBI | GCA_947561395.1 |

| | | | | | | | | | |
|---|---|---|---|---|---|---|---|---|---|
| | | BuxseGNL2 (chr 8, Mb 40.0) | 1403 | res | | | 66 | NCBI | GCA_947561395.1 |
| Buxales (Buxaceae) | Pachysandra terminalis | PacteGNOMa (sc9, Mb 6.9) | 1279 | ? (gap) | 79 | 68 | | NCBI | GCA_031762675.1 |
| | | PacteGNOMb (sc20, Mb 3.1) | 1437 | res | 66 | 57 | | NCBI | GCA_031762675.1 |
| | | PacteGNL2 (sc3, Mb 11.5) | 1387 | sens | | | 64 | NCBI | GCA_031762675.1 |
| Trochondendrales (Trochodendraceae) | Trochodendron aralioides | TroarGNOMa EVMgroup0.675 | 1468 | sens | 82 | 64 | | Giga Science | Aral chr pep file |
| | | TroarGNOMc EVMgroup12.117 | 1440 | sens | 67 | 58 | | Giga Science | Aral chr pep file |
| | | TroarGNOMb EVMgroup13.1500 | 1465 | res | 81 | 65 | | Giga Science | Aral chr pep file |
| | | TroarGNL2a EVMgroup17.673 | 1401 | sens | | | 67 | Giga Science | Aral chr pep file |
| | | TroarGNL2b EVMgroup17.668 | 996 | sens | | | 63 | Giga Science | Aral chr pep file |
| Trochondendrales (Trochodendraceae) | Tetracentron sinense | TetsiGNOMa HHK36_007283 (chr 4, Mb 50.5) | 1465 | sens | 80 | 64 | | NCBI | GCA_015143295.1 |
| | | TetsiGNOMb HHK36_000870 (chr 1, Mb 18.9) | 1439 | res | 66 | 58 | | NCBI | GCA_015143295.1 |
| | | TetsiGNOMc HHK36_015395 (chr 10, Mb 35.3) | 1469 | sens | 81 | 64 | | NCBI | GCA_015143295.1 |
| | | TetsiGNL2 HHK36_012649 (chr 8, Mb 9.6) | 1399 | sens | | | 67 | NCBI | GCA_015143295.1 |
| Gunnerales (Myrothamnaceae) | Myrothamnus flabellifolia | MyrflGNOM MyflaH2.01G225100 (chr1, Mb 117.2) | 1461 | sens | 82 | 63 | | Phytozome 13 | M.flabellifolia HAP2 |
| | | MyrflGNL2 MyflaH2.05G046800 (chr5, Mb 27.2) | ~~1400~~ | sens | | | 66 | Phytozome 13 | M.flabellifolia HAP2 |

**Suppl. Table S1G. GBF1-type ARF-GEFs in superasterids (15 orders, 65 families, 156 species)**

| Order | Species | Gene name (chr/ctg/sc #) | Protein Size (# aa) | BFA res / sens | % id. AtGN | % id. AtGNL1 | % id. AtGNL2 | database | ref |
|---|---|---|---|---|---|---|---|---|---|
| Dilleniales (Dilleniaceae) | Dillenia turbinata | DiltuGNOMa RJ641_034610 (sc8, Mb 5.0) | 1472 | sens | 83 | 62 | | NCBI | GCA_037126225.1 |
| | | DiltuGNOMb RJ641_023984 (sc28, Mb 13.4) | 1455 | sens | 70 | 59 | | NCBI | GCA_037126225.1 |
| | | DiltuGNOMc RJ641_019877 (sc25, Mb 1.8) | 1455 | res | 70 | 60 | | NCBI | GCA_037126225.1 |
| | | DiltuGNL2a RJ641_026917 (chr4, Mb 3.9) | | | | | n.a. | NCBI | GCA_037126225.1 |
| | | DiltuGNL2b RJ641_026094 /sc3, Mb 26.2) | | | | | n.a. | NCBI | GCA_037126225.1 |
| Dilleniales (Dilleniaceae) | Dillenia suffruticosa | DilsuGNOMa (sc22, 2.7) | 1472 | sens | 83 | 62 | | NCBI | GCA_033439265.1 |
| | | DilsuGNOMb (sc9, Mb 23.2) | 1459 | sens | 70 | 60 | | NCBI | GCA_033439265.1 |
| | | DilsuGNOMc (sc12, Mb 22.4) | 1459 | res | 70 | 61 | | NCBI | GCA_033439265.1 |
| | | DilsuGNL2a (sc7, Mb 22.0) | 1397 | sens | | | 65 | NCBI | GCA_033439265.1 |
| | | DilsuGNL2b (sc6, Mb 3.7) | 1392 | sens | | | 64 | NCBI | GCA_033439265.1 |
| Caryophyllales (Tamaricaceae) | Tamarix chinensis | TC01G3028 / TamchGNOMa | 1456 | sens | 81 | 62 | | GigaSci database | Pep file |
| | | TC03G0796 / TamchGNOMb | 1446 | res | 78 | 60 | | GigaSci database | Pep file |
| | | TC05G2206 / TamchGNOMc | 1465 | sens | 68 | 60 | | GigaSci database | Pep file |
| Caryophyllales (Polygonaceae) | Fagopyrum tartaricum | FagtaGNOMa (chr 5, Mb 18.6) | 1443 | sens | 81 | 62 | | NCBI | GCA_002319775.1 |

| | | | | | | | | | |
|---|---|---|---|---|---|---|---|---|---|
| | | FagtaGNOMb (chr 5, Mb 43.3) | 1442 | sens | 68 | 59 | | NCBI | GCA_002319775.1 |
| | | FagtaGNOMc (chr 6, Mb 36.2) | 1450 | res | 66 | 57 | | NCBI | GCA_002319775.1 |
| | | FagtaGNL2a (chr 5, Mb 51.6) | 1360 | sens | | | 56 | NCBI | GCA_002319775.1 |
| | | FagtaGNL2b (chr 6, Mb 47.9) | 1368 | sens | | | 59 | NCBI | GCA_002319775.1 |
| Caryophyllales (Polygonaceae) | Polygonum aviculare | PolavGNOMa (chr1, Mb 28.0) | 1446 | sens | 81 | 62 | | NCBI | GCA_934048045.1 |
| | | PolavGNOMb (chr10, Mb 2.5) | 1446 | sens | 81 | 62 | | NCBI | GCA_934048045.1 |
| | | PolavGNL2 (chr 2, Mb 38.5) | 1369 | sens | | | 59 | NCBI | GCA_934048045.1 |
| Caryophyllales (Plumbaginaceae) | Limonium bicolor | LimbiGNOMa (chr 2, Mb 22.2) | 1458 | sens | 80 | 62 | | NCBI | GCA_023374045.1 |
| | | LimbiGNOMb (chr 1; Mb 387.4) | 1460 | res | 67 | 58 | | NCBI | GCA_023374045.1 |
| | | LimbiGNOMc (chr 1; Mb 387.1) | 1389 | res | 59 | 56 | | NCBI | GCA_023374045.1 |
| | | LimbiGNL2 (chr 7) | 1426 | sens | | | 54 | NCBI | GCA_023374045.1 |
| Caryophyllales (Nepenthaceae) | Nepenthes gracilis | NepgrGNOMa (sc15) Nepgr_017127 | 1450 | sens | 80 | 61 | | NCBI | GCA_033239525.1 |
| | | NepgrGNOMb (sc8) Nepgr_009522 | 1452 | sens | 69 | 58 | | NCBI | GCA_033239525.1 |
| | | NepgrGNOMc (sc11) Nepgr_012222 | (1429) | res | 62 | 54 | | NCBI | GCA_033239525.1 |
| | | NepgrGNOMd (sc4) Nepgr_003594 | (1086) | sens | 71 | 53 | | NCBI | GCA_033239525.1 |
| | | NepgrGNL2 (sc20) Nepgr_020765 | 1363 | sens | | | 63 | NCBI | GCA_033239525.1 |

| | | | | | | | | | |
|---|---|---|---|---|---|---|---|---|---|
| Caryophyllales (Droseraceae) | Drosera capensis | DrocaGNOM_LIEC01011435 jcf7180002683404 | (1450) | sens | 78 | 62 | | NCBI | GCA_001925005.1 |
| | | DrocaGNL1a_LIEC01010116 jcf7180002682079 | (1465) | sens | 68 | 56 | | NCBI | GCA_001925005.1 |
| | | DrocaGNL1b_LIEC01011850 jcf7180002683820 | (1470) | res | 67 | 56 | | NCBI | GCA_001925005.1 |
| | | DrocaGNL1c_LIEC01006883 jcf7180002678744 | (1460) | sens | 64 | 56 | | NCBI | GCA_001925005.1 |
| | | DrocaGNL2_LIEC01010600 | (1375) | sens | | | 59 | NCBI | GCA_001925005.1 |
| | | DrocaGNOM_LIEC01008622 C-term fragm (end of contig) | (450) | | 73 | 60 | | NCBI | GCA_001925005.1 |
| | | DrocaGNL1_LIEC01011346 N-term fragm (middle of contig) | (516) | | 69 | 68 | | NCBI | GCA_001925005.1 |
| Caryophyllales (Simondsiaceae) | Simmondsia chinensis | SimchGNOMa Sc04g0007880 (sc10, Mb 36.9) | 1449 | sens | 81 | 62 | | NCBI | GCA_018398585.1 |
| | | SimchGNOMb Sc04g0001650 (sc10, Mb 2.0) | 1443 | res | 70 | 60 | | NCBI | GCA_018398585.1 |
| | | SimchGNL2 Sc18g0001130 (sc18_1) | 1379 | sens | | | 61 | NCBI | GCA_018398585.1 |
| Caryophyllales (Caryophyllaceae) | Dianthus caryophyllus | DiacaGNOMa (LG15) | 1448 | sens | 79 | 61 | | NCBI | GCA_023091065.1 |
| | | DiacaGNOMb (LG11) | 1450 | sens | 66 | 58 | | NCBI | GCA_023091065.1 |
| | | DiacaGNL2 (LG08) | 1351 | sens | | | 61 | NCBI | GCA_023091065.1 |
| Caryophyllales (Caryophyllaceae) | Heliosperma pusillum | HelpuGNOMa KSS87_017689 jcf7180000347315 | 1450 | sens | 81 | 62 | | NCBI | GCA_022085475.1 |

| | | | | | | | | | |
|---|---|---|---|---|---|---|---|---|---|
| | | HelpuGNOMb KSS87_008652 jcf7180000417999 | 1441 | sens | 65 | 58 | | NCBI | GCA_022085475.1 |
| | | HelpuGNL2 jcf7180000349207 | 1347 | sens | | | 60 | NCBI | GCA_022085475.1 |
| Caryophyllales (Caryophyllaceae) | Gypsophila paniculata | Gpan04g00496 (chr4) GyppaGNOMa (sc12, Mb 2.8) | 1448 | sens | 80 | 62 | | NCBI | GCA_032274805.1 |
| | (CNGB pep file Oct 22) | Gpan12g00130 (chr12) GyppaGNOMb (sc180566) | 1451 | sens | 66 | 57 | | NCBI | GCA_032274805.1 |
| | NCBI Oct 23 | Gpan05g01547 (chr5) GyppaGNL2 (sc183188) | 1347 | sens | | | 61 | NCBI | GCA_032274805.1 |
| Caryophyllales (Caryophyllaceae) | Saponaria officinalis | SapofGNOMa Sapof.08G052500 (chr8, Mb 19.3) | 1448 | sens | 81 | 62 | | NCBI | GCA_040167595.1 |
| | | SapofGNOMb Sapof.03G130900 (chr3, Mb 154.4) | 1448 | sens | 64 | 57 | | NCBI | GCA_040167595.1 |
| | | SapofGNOMc Sapof.09G070700 (chr9, Mb 51.9) | 1082 | sens | 67 | 52 | | NCBI | GCA_040167595.1 |
| | | SapofGNL2 (chr4) Sapof.04G028000 | 1350 | sens | | | 61 | NCBI | GCA_040167595.1 |
| Caryophyllales (Amaranthaceae) | Amaranthus hypochondriacus | AmahyGNOM (chr 3) | (1450) | sens | 80 | 69 | | NCBI | GCA_000753965.2 |
| | | AmahyGNL1 (chr 12) | (1460) | res | 62 | 54 | | NCBI | GCA_000753965.2 |
| | | AmahyGNL2 (chr 14) | (1371) | | | | 58 | NCBI | GCA_000753965.2 |
| Caryophyllales (Amaranthaceae) | Amaranthus cruentus | Amacr03g15130 (chr3) | 1460 | sens | 79 | 61 | | NCBI | GCA_048987545.1 |
| | | Amacr12g11020 (chr 12) | 1447 | res | 65 | 57 | | NCBI | GCA_048987545.1 |
| | | Amacr14g03290 (chr 14) | 1351 | sens | | | 59 | NCBI | GCA_048987545.1 |

| | | | | | | | | | |
|---|---|---|---|---|---|---|---|---|---|
| Caryophyllales (Amaranthaceae) | Chenopodium quinoa | LOC110726932 ChequGNOMd | 1439 | res | 67 | 59 | | NCBI | GCA_001683475.1 |
| | | LOC110713128 ChequGNOMa | 1444 | sens | 82 | 61 | | NCBI | GCA_001683475.1 |
| | | LOC110696589 ChequGNOMb | 1446 | sens | 80 | 61 | | NCBI | GCA_001683475.1 |
| | | LOC110732048 (2 isoforms) ChequGNOMc | 1443 | sens | 80 | 61 | | NCBI | GCA_001683475.1 |
| | | LOC110707811 | 1347 | sens | | | 59 | NCBI | GCA_001683475.1 |
| Caryophyllales (Amaranthaceae) | Spinacia oleracea | LOC110804592 (chr6) | 1438 | sens | 79 | 61 | | NCBI | GCA_020520425.1 |
| | | LOC110785052 (chr2) | 1446 | sens | 81 | 61 | | NCBI | GCA_020520425.1 |
| | | LOC110776164 (chr4) | 1434 | res | 67 | 59 | | NCBI | GCA_020520425.1 |
| | | LOC110777245 (ctg30) | 1352 | sens | | | 59 | NCBI | GCA_020520425.1 |
| Caryophyllales (Amaranthaceae) | Beta vulgaris | Betvu_LOC104897851 (chr6) | 1443 | sens | 81 | 62 | | NCBI | GCA_026745355.1 |
| | | Betvu_LOC104906035 (chr2) | 1437 | res | 66 | 57 | | NCBI | GCA_026745355.1 |
| | | Betvu_LOC104902176 (chr8) | 1369 | | | | 60 | NCBI | GCA_026745355.1 |
| Caryophyllales (Molluginaceae) | Pharnaceum exiguum | PhaexGNOM (sc2487) | (1460) | sens | 83 | 62 | | NCBI | GCA_900322385.1 |
| | | PhaexGNL2 (sc3276) | (1350) | sens | | | 60 | NCBI | GCA_900322385.1 |
| Caryophyllales (Talinaceae) | Talinum fruticosum | TalfrGNOMa (Tf_contig_014) | 1440 | sens | 81 | 62 | | NCBI | GCA_021018815.1 |
| | | TalfrGNOMb (Tf_contig_100) | 1426 | sens | 70 | 55 | | NCBI | GCA_021018815.1 |

| | | | | | | | | | |
|---|---|---|---|---|---|---|---|---|---|
| | | TalfrGNL2 (Tf_contig_099) | 1359 | sens | | | 62 | NCBI | GCA_021018815.1 |
| Caryophyllales (Cactaceae) | Carnegiea gigantea | CargiGNOMa (sc8504) | 1439 | sens | 80 | 63 | | NCBI | GCA_029747015.1 |
| | | CargiGNOMb (sc1597) | 1441 | sens | 75 | 60 | | NCBI | GCA_029747015.1 |
| | | CargiGNL2 (sc1566) | 1355 | sens | | | 62 | NCBI | GCA_029747015.1 |
| Caryophyllales (Portulacaceae) | Portulaca amilis | FUN_023444 (sc4) PoramGNOMa | 1440 | sens | 81 | 63 | | Phytozome 13 | V1.0 |
| | | FUN_037122 (sc7) PoramGNOMb | 1451 | res | 66 | 53 | | Phytozome 13 | V1.0 |
| | | FUN_026381 (sc5) PoramGNL2 | 1369 | sens | | | 61 | Phytozome 13 | V1.0 |
| Caryophyllales (Aizoaceae) | Mesembryanthemum crystallinum | MescrGNOM (sc3010) | 1447 | sens | 81 | 62 | | NCBI | GCA_030267885.1 |
| | | MescrGNL2 (sc2180) | 1360 | sens | | | 61 | NCBI | GCA_030267885.1 |
| Caryophyllales (Aizoaceae) | Tetragonia tetragonoides | TetteGNOMa (chr8, Mb 30.5) | 1439 | sens | 82 | 62 | | NCBI | GCA_038025085.1 |
| | | TetteGNOMb (chr10, Mb 38.7) | 1439 | sens | 82 | 62 | | NCBI | GCA_038025085.1 |
| | | TetteGNL2a (chr4, Mb 39.4) | 1355 | sens | | | 61 | NCBI | GCA_038025085.1 |
| | | TetteGNL2b (chr6, Mb 42.6) | 1249 | sens | | | 60 | NCBI | GCA_038025085.1 |
| Cornales (Hydrangeaceae) | Hydrangea quercifolia | Hyque.10G111800 (chr 10) | 1458 | sens | 83 | 63 | | Phytozome 13 | V1.1 |
| | | Hyque.13G093400 (chr 13) | 1436 | res | 72 | 65 | | Phytozome 13 | V1.1 |
| | | Hyque.18G092600 (chr 18) | 1368 | sens | | | 68 | Phytozome 13 | V1.1 |
| Cornales (Hydrangeaceae) | Hydrangea macrophylla | HydmaGNOM (sc 0112F.1) | 1463 | sens | 83 | 63 | | NCBI | GCA_033977485.1 |

| | | | | | | | | | |
|---|---|---|---|---|---|---|---|---|---|
| | | HydmaGNL1 (sc1607F.1) | 1448 | res | 71 | 64 | | NCBI | GCA_033977485.1 |
| | | HydmaGNL2 (sc 0518F.1) | 1363 | sens | | | 68 | NCBI | GCA_033977485.1 |
| Cornales (Cornaceae) | Cornus controversa | CorcoGNOM (chr 2, Mb 26.0) Cco\|G0010105 | 1470 | sens | 83 | 64 | | NCBI | GCA_026545225.1 |
| | | CorcoGNL1 (chr 1, Mb 124.2) Cco\|G0008165 | 1463 | res | 70 | 64 | | NCBI | GCA_026545225.1 |
| | Figshare May23 annotation | CorcoGNL2 (chr 7) Cco\|G0031546 | 1389 | sens | | | 65 | NCBI | GCA_026545225.1 |
| Cornales (Cornaceae) | Cornus florida | CorflGNOM (chr7, Mb 44.8) LOC132267122 | 1466 | sens | 81 | 63 | | NCBI | GCA_030987335.1 |
| | | CorflGNL1a (chr11; Mb 134.966) LOC132286102 | 1463 | sens | 68 | 64 | | NCBI | GCA_030987335.1 |
| | | CorflGNL1b (chr11; Mb 134.957) LOC132282205 | 1358 | sens | 62 | 64 | | NCBI | GCA_030987335.1 |
| | | CorflGNL2 (chr10, Mb 90.2) LOC132280074 | 1383 | sens | | | 65 | NCBI | GCA_030987335.1 |
| Cornales (Cornaceae) | Cornus wilsoniana | CorwiGNOM (chr5) CW05G04750 | 1470 | sens | 83 | 64 | | CNGB | GWH pep file |
| | | CorwiGNL1 (chr1) CW01G46140 | 1463 | res | 70 | 64 | | CNGB | GWH pep file |
| | | CorwiGNL2 (chr8) CW08G13890 | 1007 | sens | | | 69 | CNGB | GWH pep file |
| Cornales (Cornaceae) | Cornus suecica | CorsuGNOM (chr11, Mb 27.3) | | sens | 82 | 63 | | NCBI | GCA_963922005.1 |
| | | CorsuGNL1 (chr1, Mb 3.7) | | res | 67 | 63 | | NCBI | GCA_963922005.1 |
| | | CorsuGNL2 (chr3, Mb 73.4) | | sens | | | 65 | NCBI | GCA_963922005.1 |

| | | | | | | | | | |
|---|---|---|---|---|---|---|---|---|---|
| Cornales (Nyssaceae) | Camptotheca acuminata | Cac_g011358 | 1465 | sens | 81 | 63 | | Dryad | Pep file |
| | | Cac_g006307 | 1465 | res | 71 | 66 | | Dryad | Pep file |
| | | Cac_g011770 | 1380 | sens | | | 68 | Dryad | Pep file |
| Cornales (Nyssaceae) | Nyssa sinensis | Nyssi_F0562_000634 (LG0) | 1463 | res | 72 | 66 | | NCBI | GCA_008638375.1 |
| | | Nyssi_F0562_012948 (LG5) | 1443 | sens | 83 | 64 | | NCBI | GCA_008638375.1 |
| | | Nyssi_F0562_012781 (LG5) | 1436 | sens | 79 | 61 | | NCBI | GCA_008638375.1 |
| | | Nyssi_F0562_009521 + 009520 (LG4) NyssiGNL2 | 1372 | sens | | | 68 | NCBI | GCA_008638375.1 |
| Cornales (Davidiaceae) | Davidia involucrata | GWHGABJS030942+ GWHGABJS030943 (chr 5) DavinGNOMa | 1461 | sens | 83 | 63 | | CNGB | GWH Pep file |
| | | GWHGABJS005498+ GWHGABJS005497 (chr 10) DavinGNOMb | 1349 | sens | 84 | 66 | | CNGB | GWH Pep file |
| | | GWHGABJS020601 (chr 19) DavinGNL1 | 14633 | res | 72 | 66 | | CNGB | GWH Pep file |
| | | GWHGABJS021366 (chr 2) DavinGNL2 | 1384 | sens | | | 68 | CNGB | GWH Pep file |
| Ericales (Balsaminaceae) | Impatiens glandulifera | ImpglGNOM (chr 4, Mb 57.3) | 1421 | sens | 78 | 60 | | NCBI | GCA_907164915.1 |
| | | ImpglGNL1a (chr 1, Mb 65.2) | 1441 | res | 65 | 62 | | NCBI | GCA_907164915.1 |
| | | ImpglGNL1b (chr 9, Mb 28.6) LOC124914662 | 1454 | res | 63 | 59 | | NCBI | GCA_907164915.1 |
| | | ImpglGNL2 (chr 2, Mb 11.2) LOC124924335 | 1375 | sens | | | 63 | NCBI | GCA_907164915.1 |
| Ericales (Balsaminaceae) | Impatiens balsamina | ImpbaGNOM (sc53, Mb 3.8) | 1416 | sens | 79 | 61 | | NCBI | GCA_040721895.1 |

| | | | | | | | | | |
|---|---|---|---|---|---|---|---|---|---|
| | | ImpbaGNL1a (sc1, Mb 15.3) | 1445 | res | 65 | 62 | | NCBI | GCA_0407 21895.1 |
| | | ImpbaGNL1b (sc112, Mb 6.6) | 1455 | res | 64 | 59 | | NCBI | GCA_0407 21895.1 |
| | | ImpbaGNL2 (sc21, Mb 3.3) | 1367 | sens | | | 63 | NCBI | GCA_0407 21895.1 |
| Ericales (Polemoniaceae) | Linanthus parryae | LinpaGNOMa (SCAF_2; Mb 153.65) | 1457 | sens | 80 | 62 | | NCBI | GCA_0230 55425.1 |
| | | LinpaGNOMb (SCAF_11, Mb12.3) | 1460 | res | 71 | 59 | | NCBI | GCA_0230 55425.1 |
| | | LinpaGNL2 (SCAF_14) | 1359 | sens | | | 65 | NCBI | GCA_0230 55425.1 |
| Ericales (Sapotaceae) | Argania spinosa (Sideroxylum spinosum) | ArgspGNOMa jcf7180001103702 | (1430) | sens | 84 | 63 | | NCBI | GCA_0480 23285.2 |
| | | ArgspGNOM/GNOMb jcf7180001164485 | (1200) | sens | 73 | 56 | | NCBI | GCA_0480 23285.2 |
| | | ArgspGNL1 jcf7180001094971 | (1450) | res | 69 | 62 | | NCBI | GCA_0480 23285.2 |
| | | ArgspGNL2 jcf7180001163719 | (1370) | sens | | | 65 | NCBI | GCA_0480 23285.2 |
| Ericales (Sapotaceae) | Vitellaria paradoxa | Vitpa01g15290 (chr 1) (VitpaGNOMa) | 1463 | sens | 82 | 64 | | NCBI | GCA_0199 16065.1 |
| | ) | Vitpa05g17920 (chr 5) (VitpaGNOMb) | 1471 | sens | 74 | 59 | | NCBI | GCA_0199 16065.1 |
| | | Vitpa01g38230 (chr 1)(+VTI13) (VitpaGNL1) | 1465 | res | 69 | 64 | | NCBI | GCA_0199 16065.1 |
| | | Vitpa09g12830 (chr 9) (VitpaGNL2) | 1374 | sens | | | 66 | NCBI | GCA_0199 16065.1 |
| Ericales (Ebenaceae) | Diospyros lotus | DioloGNOMa (chr 7) LOC127806012 | 1458 | sens | 84 | 63 | | NCBI | GCA_0146 33365.1 |
| | | DioloGNOMb (chr 10) LOC127811807 | 1460 | res | 70 | 63 | | NCBI | GCA_0146 33365.1 |

| | | | | | | | | | |
|---|---|---|---|---|---|---|---|---|---|
| | | DioloGNL2a (chr 1) | | sens | | | | NCBI | GCA_0146 33365.1 |
| | | DioloGNL2b (chr 2) | | sens | | | | NCBI | GCA_0146 33365.1 |
| Ericales (Primulaceae) | Aegiceras corniculatum | THS.05G0197980 (chr 5) Aegco_05G0197980 | 1457 | sens | 79 | 61 | | CNGB | CNA00177 38 |
| | | THS.14G0264380 (chr 14) Aegco_14G0264380 | 1457 | sens | 79 | 61 | | CNGB | CNA00177 38 |
| | | THS.02G0321210 (chr 2) Aegco_02G0321210 | 1359 | sens | | | 63 | CNGB | CNA00177 38 |
| Ericales (Primulaceae) | Primula veris | BG8809307 (ctg239) (PriveGNOM) | 1448 | sens | 79 | 61 | | NCBI | GCA_9636 82055.1 |
| | | BG8816056 (ctg1350) (PriveGNL2) | 1360 | sens | | | 65 | NCBI | GCA_9636 82055.1 |
| Ericales (Primulaceae) | Primula vulgaris | PvLHv1_174150 (PrivuGNOM) PrivuGNOMa (chr6, Mb 15.8) | 1448 | sens | 79 | 61 | | NCBI | GCA_9639 20805.1 |
| | | PrivuGNOMb (chr2, Mb 0.1) | 1450 | sens | 71 | 55 | | NCBI | GCA_9639 20805.1 |
| | | PvLHv1_059820 (PrivuGNL2) PrivuGNL2 (chr7, Mb 14.1) | 1360 | sens | | | 65 | NCBI | GCA_9639 20805.1 |
| Ericales (Pentaphylacaceae) | Euryodendron excelsum | EurexGNOMa (chr12, Mb 6.0) | 1464 | res | 82 | 63 | | NCBI | GCA_0330 60915.1 |
| | | EurexGNOMb (chr2, Mb 10.0) | 1469 | sens | 84 | 63 | | NCBI | GCA_0330 60915.1 |
| | | EurexGNL1 (chr7, Mb 4.9) | 1461 | sens | 72 | 65 | | NCBI | GCA_0330 60915.1 |
| | | EurexGNL2 (chr5, Mb 9.1) | 1395 | sens | | | 68 | NCBI | GCA_0330 60915.1 |
| Ericales (Theaceae) | Camellia sinensis | Camsi_LOC1143063 71 (sc35) HYC85_021014 (chr 10) | 1465 | sens | 81 | 62 | | NCBI | GCA_0041 53795.1 |

| | | | | | | | | | |
|---|---|---|---|---|---|---|---|---|---|
| | | Camsi_LOC114271324 (sc2148) HYC85_016072 (chr 7) | 1441 | res | 70 | 63 | | NCBI | GCA_004153795.1 |
| | | Camsi_LOC114282320 (sc3154) HYC85_015997 (chr 7) | 1378 | sens | | | 67 | NCBI | GCA_004153795.1 |
| Ericales (Styracaceae) | Sinojackia xylocarpa | SinxyGNOM (chr8, Mb 69.7) Ssp08G023500 | 1162 | sens | 85 | 65 | | NGDC | GHW Pep file |
| | | SinxyGNL1 (chr3, Mb 66.6) Ssp03G011960 | 1460 | res | 69 | 64 | | NGDC | GHW Pep file |
| | | SinxyGNL2 (chr3, Mb 71.3) Ssp03G014540 | 1345 | sens | | | 67 | NGDC | GHW Pep file |
| Ericales (Roridulaceae) | Roridula gorgonias | rgor_13293 Rorgo_13293 | 1462 | sens | 83 | 63 | | Dryad | Pep file |
| | | rgor_03512 Rorgo_03512 | 1455 | sens | 68 | 63 | | Dryad | Pep file |
| | | rgor_08858 Rorgo_08858 | 1386 | sens | | | 65 | Dryad | Pep file |
| Ericales (Actinidiaceae) | Actinidia chinensis | RUSV2a.006361 (chr 6) ActchGNOMa | 1462 | sens | 83 | 63 | | figshare | Pep file |
| | | RUSV2a.031499 (chr 27)ActchGNOMb | 1462 | sens | 82 | 62 | | figshare | Pep file |
| | | RUSV2a.043104 (chr 2) ActchGNL1a | 1461 | res | 69 | 63 | | figshare | Pep file |
| | | RUSV2a.004073 (chr 3) ActchGNL1b | 1460 | sens | 70 | 63 | | figshare | Pep file |
| | | RUSV2a.009920 (chr 9) ActchGNL2 | 1386 | | | | 66 | figshare | Pep file |
| Ericales (Clethraceae) | Clethra arborea | carb_17208 ClearGNOM | (1462) | sens | 83 | 62 | | Dryad | Pep file |
| | | carb_18756 ClearGNL1 | 1456 | res | 67 | 63 | | Dryad | Pep file |

| | | | | | | | | | |
|---|---|---|---|---|---|---|---|---|---|
| | | carb_06455 ClearGNL2 | 1374 | sens | | | 65 | Dryad | Pep file |
| Ericales (Ericaceae) | Rhododendron williamsianum | Rhowi_C3L33_18534 (Rhowi_C3L33_18533 RhowiGNOM (LG11) | 14xx | sens | 81 | 65 | | NCBI | GCA_0097 46105.1 |
| | | Rhowi_C3L33_01496 RhowiGNL1 (LG01) | 1371 | res | 65 | 61 | | NCBI | GCA_0097 46105.1 |
| | | Rhowi_C3L33_03657 RhowiGNL2 (LG02?) | 1352 | | | | 65 | NCBI | GCA_0097 46105.1 |
| Ericales (Ericaceae) | Rhododendron vialii | RhoviGNOM (chr 10a) | 1469 | sens | 81 | 62 | | NCBI | GCA_0302 53575.1 |
| | | RhoviGNL1 (chr 8a) LOC131335108 | 1451 | res | 68 | 62 | | NCBI | GCA_0302 53575.1 |
| | | RhoviGNL2 (chr 3a) | 1368 | sens | | | 66 | NCBI | GCA_0302 53575.1 |
| Ericales (Ericaceae) | Vaccinium myrtillus | VacmyGNOMa (chr 5) | (1472) | sens | 82 | 64 | | NCBI | GCA_0169 20895.1 |
| | | VacmyGNOMb (chr 7) | (1470) | sens | 83 | 63 | | NCBI | GCA_0169 20895.1 |
| | | VacmyGNOMc (chr 8) | (1452) | res | 67 | 62 | | NCBI | GCA_0169 20895.1 |
| | | VacmyGNL2 (chr 11) | (1375) | | | | 66 | NCBI | GCA_0169 20895.1 |
| Ericales (Ericaceae) | Vaccinium darrowii | VacdaGNOMb (chr 7)Vadar_006062 | 1471 | sens | 80 | 63 | | NCBI | GCA_0209 21065.1 |
| | | VacdaGNOMa (chr 5) Vadar_003698+Vadar _003622 | 1471 | sens | 81 | 63 | | NCBI | GCA_0209 21065.1 |
| | | VacdaGNL1 (chr 8) Vadar_021760 | 1449 | res | 67 | 62 | | NCBI | GCA_0209 21065.1 |
| | | VacdaGNL2 (chr 11) Vadar_027043 | 1371 | sens | | | 66 | NCBI | GCA_0209 21065.1 |
| Ericales (Ericaceae) | Arctostaphylos glauca | ArcglGNOM (SCAF_8, Mb 22.1) | 1472 | sens | 82 | 63 | | NCBI | GCA_0199 85065.2 |
| | NCBI Feb 22 | ArcglGNL1 (SCAF_14, Mb 14.7) | 1452 | sens | 67 | 61 | | NCBI | GCA_0199 85065.2 |

| | | | | | | | | | |
|---|---|---|---|---|---|---|---|---|---|
| | | ArcglGNL2a (SCAF_2, Mb 4.8) | 1369 | sens | | | 65 | NCBI | GCA_019985065.2 |
| | | ArcglGNL2b (SCAF_10, Mb 18.1) | 1370 | sens | | | 66 | NCBI | GCA_019985065.2 |
| Ericales (Ericaceae) | Arbutus unedo | ArbunGNOM (ctg_14799) | 1473 | sens | 81 | 63 | | NCBI | GCA_014822125.1 |
| | | ArbunGNL1 (ctg_17973) | 1452 | sens | 67 | 61 | | NCBI | GCA_014822125.1 |
| | | ArbunGNL2 (ctg_22936) | 1369 | sens | | | 66 | NCBI | GCA_014822125.1 |
| Ericales (Ericaceae) | Calluna vulgaris | CalvuGNOM (chr4. Mb 38.7) | 1470 | sens | 81 | 62 | | NCBI | GCA_964145215.1 |
| | | CalvuGNL1 (chr6, Mb 3.8) | 1451 | res | 65 | 61 | | NCBI | GCA_964145215.1 |
| | | CalvuGNL2 (chr7, Mb4.7) | 1365 | sens | | | 65 | NCBI | GCA_964145215.1 |
| Ericales (Ericaceae) | Corema album | CoralGNOM (chr4, Mb 32.5) | 1470 | sens | 81 | 62 | | NCBI | GCA_964187925.1 |
| | | CoralGNL1 (chr7 Mb 34.0) | 1450 | res | 67 | 62 | | NCBI | GCA_964187925.1 |
| | | CoralGNL2 (chr9, Mb 36.5) | 1367 | sens | | | 65 | NCBI | GCA_964187925.1 |
| Asterales (Asteraceae) | Cynara cardunculus | Cynca_LOC112507087 (LG2) | 1449 | sens | 82 | 63 | | NCBI | GCA_001531365.2 |
| | | Cynca_LOC112512756 (LG5) | 1444 | sens | 65 | 60 | | NCBI | GCA_001531365.2 |
| | | Cynca_LOC112516397 (LG8) | 1396 | sens | | | 62 | NCBI | GCA_001531365.2 |
| Asterales (Asteraceae) | Lactuca sativa | LacsaGNOM (chr 8)LOC111878063 | 1434 | sens | 81 | 63 | | NCBI | GCA_002870075.4 |
| | | LacsaGNL1a (chr 3)LOC111879745 | 1443 | sens | 63 | 59 | | NCBI | GCA_002870075.4 |
| | | LacsaGNL1b (chr 2) LOC111920050 | 1380 | res | 59 | 55 | | NCBI | GCA_002870075.4 |
| | | LacsaGNL2 (chr 5) LOC111897921 | 1393 | sens | | | 62 | NCBI | GCA_002870075.4 |

| | | | | | | | | | |
|---|---|---|---|---|---|---|---|---|---|
| Asterales (Asteraceae) | Erigeron (= Conyza) canadensis | LOC122591235 EricaGNOM (chr 3) | 1435 | sens | 81 | 62 | | NCBI | GCA_010389155.1 |
| | | LOC122605937 EricaGNL1a (chr 6) | 1409 | res | 63 | 58 | | NCBI | GCA_010389155.1 |
| | | LOC122580400 EricaGNL1b (chr 8) | 1427 | sens | 62 | 59 | | NCBI | GCA_010389155.1 |
| | | LOC122597551 EricaGNL2 (chr 4) | 1388 | sens | | | 61 | NCBI | GCA_010389155.1 |
| Asterales (Asteraceae) | Taraxacum kok-saghyz | TarkoGNOM (chr4, Mb 6.9) LXL04_018119 | 1470 | sens | 80 | 61 | | NCBI | GCA_047496595.1 |
| | | TarkoGNL1 (chr5, Mb 81.8) LXL04_024132 | 1427 | sens | 62 | 59 | | NCBI | GCA_047496595.1 |
| | | TarkoGNL2 (chr1, Mb 51.7) LXL04_002565 | 1396 | sens | | | 62 | NCBI | GCA_047496595.1 |
| Asterales (Asteraceae) | Arctium lappa | ArclaGNOM (LG01, Mb 143.9) L6452_03048 | 1449 | sens | 82 | 63 | | NCBI | GCA_023525745.1 |
| | | ArclaGNL1 (LG15, Mb 1.0) L6452_38471 | 1438 | sens | 64 | 59 | | NCBI | GCA_023525745.1 |
| | | ArclaGNL2 ( LG7, Mb 3.3) L6452_21561 | 1397 | sens | | | 62 | NCBI | GCA_023525745.1 |
| Asterales (Asteraceae) | Centaurea solstitialis | CensoGNOM (chr 5, Mb 55.0) OSB04_021089 | 1446 | sens | 81 | 63 | | NCBI | GCA_030169165.1 |
| | | CensoGNL1 (chr 1, Mb 1.6) | 1435 | sens | 63 | 59 | | NCBI | GCA_030169165.1 |
| | | CensoGNL2 (chr 8, Mb 33.4) | 1394 | sens | | | 62 | NCBI | GCA_030169165.1 |
| Asterales (Asteraceae) | Carthamus tinctorius | CartiGNOM (chr4, Mb 52.9) Ct_G20439 | 1446 | sens | 81 | 63 | | CNGB | GWH Pep file |
| | | CartiGNL1 (sc279840) | 1436 | sens | 64 | 60 | | CNGB | GWH Pep file |

| | | | | | | | | | |
|---|---|---|---|---|---|---|---|---|---|
| | | CartiGNL2 | | | | | n.a. | CNGB | GWH Pep file |
| Asterales (Asteraceae) | Pulicaria dysenterica | PuldyGNOM (chr4, Mb 73.3) | 1441 | sens | 81 | 62 | | NCBI | GCA_947179395.2 |
| | | PuldyGNL1a (chr3, Mb 85.9) | 1425 | sens | 62 | 58 | | NCBI | GCA_947179395.2 |
| | | PuldyGNL1b (chr8, Mb 57.4) | 1439 | res | 62 | 58 | | NCBI | GCA_947179395.2 |
| | | PuldyGNL2 (chr7, Mb 69.0) | 1400 | sens | | | 62 | NCBI | GCA_947179395.2 |
| Asterales (Asteraceae) | Gnaphalium uliginosum | GnaulGNOM (chr1, Mb 59.6) | 1431 | sens | 81 | 63 | | NCBI | GCA_963930675.1 |
| | | GnaulGNL1a (chr3, Mb 9.4) | 1422 | res | 63 | 60 | | NCBI | GCA_963930675.1 |
| | | GnaulGNL1b (chr6. Mb 10.6) | 1425 | sens | 61 | 59 | | NCBI | GCA_963930675.1 |
| | | GnaulGNL2 (chr2, Mb 1.5) | 1399 | sens | | | 61 | NCBI | GCA_963930675.1 |
| Asterales (Asteraceae) | Logfia minima | LogmiGNOMa (chr2, Mb 42.8) | 1431 | sens | 82 | 64 | | NCBI | GCA_963921325.1 |
| | | LogmiGNOMb (chr1, Mb 8.9) | 1431 | sens | 81 | 63 | | NCBI | GCA_963921325.1 |
| | | LogmiGNL1a (chr10, Mb 8.1) | 1424 | res | 63 | 60 | | NCBI | GCA_963921325.1 |
| | | LogmiGNL1b (chr8, Mb 8.0) | 1424 | res | 63 | 59 | | NCBI | GCA_963921325.1 |
| | | LogmiGNL1c (chr7, Mb 9.2) | 1425 | sens | 61 | 59 | | NCBI | GCA_963921325.1 |
| | | LogmiGNL1d (chr6, Mb 8.6) | 1425 | sens | 61 | 58 | | NCBI | GCA_963921325.1 |
| | | LogmiGNL2 (chr3, Mb 1.1) | 1395 | sens | | | 61 | NCBI | GCA_963921325.1 |
| Asterales (Asteraceae) | Silybum marianum | SilmaGNOM (chr1, Mb 50.4) Smar01g036550 | 1450 | sens | 81 | 63 | | NCBI | GCA_036898165.1 |

| | | | | | | | | | |
|---|---|---|---|---|---|---|---|---|---|
| | | SilmaGNL1 (chr3, Mb 51.8) Smar03g036650 | 1446 | sens | 64 | 60 | | NCBI | GCA_036898165.1 |
| | | SilmaGNL2 (chr7, Mb 1.4) Smar07g001320 | 1397 | sens | | | 62 | NCBI | GCA_036898165.1 |
| Asterales (Asteraceae) | Dittrichia graveolens | DitgrGNOM (sc5, Mb 25.0) | 1442 | sens | 81 | 63 | | NCBI | GCA_038502395.1 |
| | | DitgrGNL1 (sc1, Mb 87.4) | 1424 | sens | 62 | 59 | | NCBI | GCA_038502395.1 |
| | | DitgrGNL2 (sc9, Mb 5.0) | | | | | n.a. | NCBI | GCA_038502395.1 |
| Asterales (Campanulaceae) | Platycodon grandiflorus | PlagrGNOM (chr 7, Mb 41.86) | 1462 | sens | 81 | 64 | | NCBI | GCA_016624345.1 |
| | | PlagrGNL1 (chr 1) PGJG377720 | 1447 | res | 68 | 64 | | NCBI | GCA_016624345.1 |
| | | PlagrGNL2 (chr 9) PGJG245830 | 1372 | sens | | | 66 | NCBI | GCA_016624345.1 |
| Asterales (Campanulaceae) | Codonopsis lanceolata | CodlaGNOM (chr 1) Cl_43450 | 1462 | sens | 82 | 63 | | NCBI | GCA_013146195.2 |
| | | CodlaGNL1 (chr 3) Cl_35770 | 1443 | res | 68 | 64 | | NCBI | GCA_013146195.2 |
| | | CodlaGNL2 (chr 8) | 1360 | sens | | | 66 | NCBI | GCA_013146195.2 |
| Asterales (Campanulaceae) | Campanula takesimana | CamtaGNOMa (sc285) | 1459 | sens | 83 | 63 | | NCBI | GCA_026119855.1 |
| | | CamtaGNOMb (sc929) | 1445 | sens | 81 | 63 | | NCBI | GCA_026119855.1 |
| | | CamptaGNL1 (sc1281) | 1428 | sens | 68 | 62 | | NCBI | GCA_026119855.1 |
| | | CamtaGNL2 (sc104) | 1365 | sens | | | 65 | NCBI | GCA_026119855.1 |
| Asterales (Campanulaceae) | Lobelia cardinalis | LobcaGNOM (sc50, Mb 9.2) | 1462 | sens | 83 | 64 | | NCBI | GCA_028566635.1 |
| | | LobcaGNL1 (sc1, Mb 9.1) | 1446 | res | 67 | 63 | | NCBI | GCA_028566635.1 |

| | | | | | | | | | |
|---|---|---|---|---|---|---|---|---|---|
| | | LobcaGNL2 (sc54, Mb 11.3) | 1374 | sens | | | 65 | NCBI | GCA_0285 66635.1 |
| Asterales (Campanulaceae) | Adenophora triphylla var. japonica | AdetrGNOMa (C0083) | 1458 | sens | 82 | 63 | | NCBI | GCA_0224 95995.1 |
| | | AdetrGNOMb (C4151) | 1458 | sens | 81 | 63 | | NCBI | GCA_0224 95995.1 |
| | | AdetrGNL1a (C0733) | 1445 | sens | 67 | 62 | | NCBI | GCA_0224 95995.1 |
| | | AdetrGNL1b (C3039) | 1440 | sens | 67 | 61 | | NCBI | GCA_0224 95995.1 |
| | | AdetrGNL2 (C2139) | 1364 | sens | | | 66 | NCBI | GCA_0224 95995.1 |
| Asterales (Campanulaceae) | Jasione montana | JasmoGNOMa (chr4, Mb 5.82) | 1454 | sens | 81 | 63 | | NCBI | GCA_9640 59625.1 |
| | | JasmoGNOMb (chr4, Mb 5.87) | 1434 | sens | 77 | 61 | | NCBI | GCA_9640 59625.1 |
| | | JasmoGNL2 | 1368 | sens | | | 66 | NCBI | GCA_9640 59625.1 |
| Escalloniales (Escalloniaceae) | Escallonia rubra | EscruGNOM (tig00007257) RJ640_007258 | ~~1565~~ 1459 | sens | 83 | 63 | | NCBI | GCA_0330 65855.1 |
| | | EscruGNL1a (tig00004666) RJ640_002103 | 1460 | sens | 69 | 63 | | NCBI | GCA_0330 65855.1 |
| | | EscrGNL1b (tig00010526) RJ640_025651 | 1460 | sens | 68 | 63 | | NCBI | GCA_0330 65855.1 |
| | | EscruGNL2 (tig00010172) RJ640_024279 | 1378 | sens | | | 65 | NCBI | GCA_0330 65855.1 |
| Escalloniales (Escalloniaceae) | Escallonia herrerae | EscheGNOM (tig00013077) RJ639_008991 | 1459 | sens | 84 | 64 | | NCBI | GCA_0330 70095.1 |
| | | EscheGNL1 (tig00014886) RJ639_040520 | 1460 | sens | 69 | 64 | | NCBI | GCA_0330 70095.1 |
| | | EscheGNL2a (tig00010405) | 1390 | sens | | | 66 | NCBI | GCA_0330 70095.1 |

| | | | | | | | | | |
|---|---|---|---|---|---|---|---|---|---|
| | | RJ639_032265 | | | | | | | |
| | | EscheGNL2b (tig00009607) RJ639_001463 | | sens | | | n.a. | NCBI | GCA_033070095.1 |
| Dipsacales (Caprifoliaceae) | Lonicera japonica | LonjaGNOM (chr 2) Lj2C104G13 + Lj2C104T12 (GWHGAAZE005595 + GWHGAAZE005594 | 1402 | sens | 79 | 61 | | NCBI | GCA_021464415.1 |
| | | Lj7A647T56 (chr 7) LonjaGNL2, GWHPAAZE025641 | 1382 | sens | | | 66 | NCBI | GCA_021464415.1 |
| | | GWHGAAZE028431 (chr 8) LonjaGNL1a | 1392 | sens | 66 | 63 | | NCBI | GCA_021464415.1 |
| | | GWHGAAZE011704 (chr 3) LonjaGNL1b | 1433 | res | 68 | 64 | | NCBI | GCA_021464415.1 |
| | | GWHGAAZE011896 (chr 3) LonjaGNL1e | 1392 | res | 65 | 59 | | NCBI | GCA_021464415.1 |
| Dipsacales (Caprifoliaceae) | Triplostegia glandulifera | TriglGNOM (chr2) TgChr02G09021 | 1450 | sens | 83 | 63 | | Figshare | Pep file |
| | | TriglGNL1 (chr7) TgChr07G24888 | 1416 | sens | 68 | 64 | | Figshare | Pep file |
| | | TriglGNL2 (chr5) TgChr05G17241 | 1172 | sens | | | 73 | Figshare | Pep file |
| Dipsacales (Adoxaceae) | Sambucus nigra | SamniGNOM (chr 15, Mb 230) | 1461 | sens | 83 | 63 | | NCBI | GCA_949130495.1 |
| | | SamniGNL1 (chr 15, Mb 528) | 1461 | res | 67 | 63 | | NCBI | GCA_949130495.1 |
| | | SamniGNL2a (chr 4) | | sens | | | n.a. | NCBI | GCA_949130495.1 |
| | | SamniGNL2b (chr 8) | | sens | | | n.a. | NCBI | GCA_949130495.1 |
| | | SamniGNL2c (chr 3, Mb 498) | | | | | n.a. | NCBI | GCA_949130495.1 |
| | | SamniGNL2d (chr 3, Mb 510) | | | | | n.a. | NCBI | GCA_949130495.1 |

| | | | | | | | | | |
|---|---|---|---|---|---|---|---|---|---|
| Apiales (Apiaceae) | Daucus carota (NEW/NCBI) | LOC108218573 (chr 4) DCAR_014959 / GNOMa | 1464 | sens | 80 | 62 | | NCBI | GCF_0016 25215.2 |
| | | LOC108218683 (chr 4) DCAR_014955 / GNOMb | 1443 | sens | 80 | 63 | | NCBI | GCF_0016 25215.2 |
| | | LOC108214229 (chr 3) DCAR_011871/GNL1b | 1446 | res | 66 | 63 | | NCBI | GCF_0016 25215.2 |
| | | LOC108212563 (chr 3) DCAR_011077/GNL1a | 1427 | res | 64 | 62 | | NCBI | GCF_0016 25215.2 |
| | | LOC108199672 (chr 8) DCAR_028721 | 1347 | sens | | | 62 | NCBI | GCF_0016 25215.2 |
| Apiales (Apiaceae) | Coriandrum sativum | CsUnG03098 / CorsaGNOMa | 1465 | sens | 82 | 62 | | Coriander Genomics Database | Pep file |
| | | Cs08G02544 + Cs08G02545 / CorsaGNOMb | 1369 | sens | 82 | 62 | | Coriander Genomics Database | Pep file |
| | | Cs07G02353 / CorsaGNL1a | 1446 | sens | 66 | 63 | | Coriander Genomics Database | Pep file |
| | | Cs01G02092 / CorsaGNL1b | 1447 | res | 67 | 63 | | Coriander Genomics Database | Pep file |
| | | Cs05G00077 (aa544-1373) CorsaGNL2 | 822 | sens | | | 61 | Coriander Genomics Database | Pep file |
| Apiales (Apiaceae) | Peucedanum praeruptorum | PeuprGNOMa BHQH00008554 | 763 | sens | 80 | 64 | | Figshare | Pep file |
| | | PeuprGNOMb BHQH00041507 | 1483 | sens | 81 | 61 | | Figshare | Pep file |
| | | PeuprGNL1 BHQH00020051 | 1446 | sens | 67 | 63 | | Figshare | Pep file |
| | | PeoprGNL2 BHQH00020799 | 1367 | sens | | | 62 | Figshare | Pep file |
| Apiales (Apiaceae) | Kitagawia praeruptora | KitprGNOMa (chr1, Mb 160.7) | 1444 | sens | 81 | 62 | | NCBI | GCA_0376 93485.1 |

| | | | | | | | | | |
|---|---|---|---|---|---|---|---|---|---|
| | | KitprGNOMb (chr10, Mb 80.3) | 1465 | sens | 82 | 61 | | NCBI | GCA_037693485.1 |
| | | KitprGNL1a (chr7, Mb 52.0) | 1446 | res | 66 | 63 | | NCBI | GCA_037693485.1 |
| | | KitprGNL1b (chr3, Mb 38.9) | 1446 | sens | 67 | 63 | | NCBI | GCA_037693485.1 |
| | | KitprGNL2 (chr11, Mb 149.3) | 1367 | sens | | | 62 | NCBI | GCA_037693485.1 |
| Apiales (Apiaceae) | Cnidium monnieri | CnimoGNOMa (chr4, Mb 86.9) | 1466 | sens | 82 | 61 | | NCBI | GCA_040143745.1 |
| | NCBI Jun 24 | CnimoGNOMb (chr6, Mb 3.0) | 1444 | sens | 81 | 62 | | NCBI | GCA_040143745.1 |
| | | CnimoGNL1a (chr5, Mb 27.3) | 1446 | res | 66 | 63 | | NCBI | GCA_040143745.1 |
| | | CnimoGNL1b (chr9, Mb 61.0) | 1446 | sens | 66 | 63 | | NCBI | GCA_040143745.1 |
| | | CnimoGNL2 (chr6, Mb 97.6) | 1367 | sens | | | 62 | NCBI | GCA_040143745.1 |
| Apiales (Araliaceae) | Hedera helix | HedheGNOMa (chr 11, Mb 21.3) | 1468 | sens | 82 | 64 | | NCBI | GCA_947179155.2 |
| | | HedheGNOMb (chr 15, Mb 20.8) | 1463 | sens | 82 | 64 | | NCBI | GCA_947179155.2 |
| | | HedheGNOMc/GNL1a (chr 7, Mb 40.1) | 1448 | sens | 68 | 64 | | NCBI | GCA_947179155.2 |
| | | HedheGNOMd/GNL1b (chr 12, Mb 37.6) | 1448 | res | 67 | 64 | | NCBI | GCA_947179155.2 |
| | | HedheGNL2 (chr 11, Mb 46.8) | 1381 | sens | | | 66 | NCBI | GCA_947179155.2 |
| Apiales (Araliaceae) | Hydrocotyle leucocephala | HydleGNOMa (ctg13) HyleuH1.13G017500 | 1461 | sens | 83 | 63 | | NCBI | GCA_964106935.1 |
| | | HydleGNOMb (ctg 7) HyleuH1.07G112200 | 1462 | sens | 83 | 63 | | NCBI | GCA_964106935.1 |
| | | HydleGNL1a (ctg 24) HyleuH2.24G030900 | 1436 | sens | 68 | 64 | | NCBI | GCA_964106935.1 |
| | | HydleGNL1b (ctg 24) HyleuH1.24G031100 | 1436 | sens | 67 | 63 | | NCBI | GCA_964106935.1 |

| | | | | | | | | | |
|---|---|---|---|---|---|---|---|---|---|
| | | HydleGNL1c (ctg 3) HyleuH1.03G062800 aa1-140missing | 1294 | sens | 64 | 60 | | NCBI | GCA_964106935.1 |
| | | HydleGNL2 (ctg 8) HyleuH1.08G088300 | 1370 | sens | | | 65 | NCBI | GCA_964106935.1 |
| Apiales (Pittosporaceae) | Pittosporum topira | PittoGNOM (sc 48391) | 1445 | sens | 82 | 62 | | NCBI | GCA_032359685.1 |
| | | PittoGNL1a (sc50254) | 1447 | sens | 69 | 63 | | NCBI | GCA_032359685.1 |
| | | PittoGNL1b (sc 57402) | 1393 | res | 65 | 60 | | NCBI | GCA_032359685.1 |
| | | PittoGNL2 (sc55040) | 1359 | sens | | | 66 | NCBI | GCA_032359685.1 |
| Aquifoliales (Aquifoliaceae) | Ilex asprella | Ilex_043370 IleasGNOMa (chr 8) | 1460 | sens | 84 | 64 | | NCBI | GCA_023539305.1 |
| | | Ilex_030306-030303 (1175aa) IleasGNOMb (chr 13) | 1441 | res | 71 | 57 | | NCBI | GCA_023539305.1 |
| | | Ilex_041692 IleasGNL2 (chr 5) | 1370 | sens | | | 68 | NCBI | GCA_023539305.1 |
| Aquifoliales (Aquifoliaceae) | Ilex polyneura | IlepoGNOMa (chr 6) GWHGBDNW027520 | 1460 | sens | 83 | 64 | | HortRes | Pep file |
| | | IlepoGNOMb (chr 12) GWHGBDNW006086 (deletion aa944-1146) | 1218 | | 72 | 57 | | HortRes | Pep file |
| | | IlepoGNL2a (chr 3) GWHGBDNW020750 | 1383 | sens | | | 67 | HortRes | Pep file |
| | | IlepoGNL2b (chr 4) GWHGBDNW022502 (only aa644-1391) | 606 | sens | | | 57 | HortRes | Pep file |
| Aquifoliales (Aquifoliaceae) | Ilex aquifolium | IleaqGNOMa (chr 6, Mb 35.5) | 1460 | sens | 83 | 64 | | NCBI | GCA_951799425.1 |
| | | IleaqGNOMb (chr 15, Mb 26.7) | 1440 | sens | 80 | 61 | | NCBI | GCA_951799425.1 |
| | | IleaqGNL2 (chr 3, Mb 47.4) | 1383 | sens | | | 67 | NCBI | GCA_951799425.1 |
| Aquifoliales (Helwingiacae) | Helwingia omeiensis | HOGene026623 (sc9) HelomGNOMa | 1458 | sens | 82 | 63 | | Fiigshare | Pep file |

| | | | | | | | | | |
|---|---|---|---|---|---|---|---|---|---|
| | | HOGene040796 (sc15) HelomGNOMb | 1438 | sens | 82 | 63 | | Fiigshare | Pep file |
| | | HOGene029268 (sc11) HelomGNOMc | 1460 | sens | 80 | 63 | | Fiigshare | Pep file |
| | | HOGene036203 (sc14) HelomGNL2 | 1321 | sens | | | 64 | Fiigshare | Pep file |
| Garryales (Eucommiaceae) | Eucommia ulmoides | Eucul_013014 (chr 9) GWHPAAAL013014 (sc78) | 1467 | sens | 81 | 63 | | NCBI | GCA_0166 47705.1 |
| | | Eucul_026230 (chr 6) GWHPAAAL026230 (sc949) | 1387 | sens | | | 64 | NCBI | GCA_0166 47705.1 |
| Icacinales (Icacinaceae) | Iodes seguinii | IodseGNOM (chr5) | 1463 | sens | 81 | 64 | | CNGB | GWHERDR 00000000 |
| | | IodseGNL1 (chr11) | 1457 | res | 68 | 63 | | CNGB | GWHERDR 00000000 |
| | | IodseGNL2 (chr10) | 1380 | sens | | | 63 | CNGB | GWHERDR 00000000 |
| Solanales (Solanaceae) | Solanum lycopersicum | Solyc05g013040 | 1449 | sens | 80 | 62 | | NCBI | GCA_0365 12215.2 |
| | | Solyc04g005560 | 1443 | sens | 79 | 62 | | NCBI | GCA_0365 12215.2 |
| | | Solyc02g067010 | 1448 | sens | 64 | 60 | | NCBI | GCA_0365 12215.2 |
| | | Solyc02g088290 | 1410 | sens | 64 | 61 | | NCBI | GCA_0365 12215.2 |
| | | Solyc03g070470 | 1368 | sens | | | 64 | NCBI | GCA_0365 12215.2 |
| Solanales (Solanaceae) | Solanum pennellii | SolpeGNOMa LOC107020592 | 1449 | sens | 81 | 62 | | NCBI | GCA_0014 06875.2 |
| | | SolpeGNOMb LOC107018127 | 1444 | sens | 79 | 61 | | NCBI | GCA_0014 06875.2 |
| | | SolpeGNL1a LOC107011435 | 1416 | sens | 65 | 61 | | NCBI | GCA_0014 06875.2 |
| | | SolpeGNL1b LOC107009047 | 1448 | sens | 64 | 61 | | NCBI | GCA_0014 06875.2 |

| | | | | | | | | | |
|---|---|---|---|---|---|---|---|---|---|
| | | SolpeGNL2 LOC107013389 | 1381 | sens | | | 64 | NCBI | GCA_001406875.2 |
| Solanales (Solanaceae) | Nicotiana sylvestris | Nicsy_LOC104237751 NicsyGNOMa | 1448 | sens | 81 | 62 | | NCBI | GCA_000393655.2 |
| | | Nicsy_LOC104213471 DCB domain missing | 1199 | sens | 80 | 62 | | NCBI | GCA_000393655.2 |
| | | Nicsy_LOC104236878 NicsyGNL1a | 1442 | sens | 66 | 62 | | NCBI | GCA_000393655.2 |
| | | Nicsy_LOC104236979 NicsyGNL1b | 1448 | sens | 65 | 61 | | NCBI | GCA_000393655.2 |
| | | Nicsy_ LOC104236920 NicsyGNL2 | 1379 | sens | | | 65 | NCBI | GCA_000393655.2 |
| Solanales (Solanaceae) | Petunia secreta# | PetseGNOMa (Scf2416325) | (1449) | sens | 81 | 63 | | NCBI | GCA_029990175.1 |
| | | PetseGNOMb (Scf2408355) | (1443) | sens | 80 | 60 | | NCBI | GCA_029990175.1 |
| | | PetseGNL1a (Scf2410847) | (1445) | res | 68 | 60 | | NCBI | GCA_029990175.1 |
| | | PetseGNL1b (Scf2415079) | (1430) | sens | 69 | 62 | | NCBI | GCA_029990175.1 |
| | | PetseGNL2 (Scf2257687) | (1381) | sens | | | 62 | NCBI | GCA_029990175.1 |
| Solanales (Solanaceae) | Datura stramonium | DatstGNOM (sc433) HAX54_033742 | 1447 | sens | 81 | 62 | | NCBI | GCA_018107945.1 |
| | | DatstGNL1a (sc33) HAX54_027185 | 1442 | sens | 65 | 62 | | NCBI | GCA_018107945.1 |
| | | DatstGNL1b (sc8778) HAX54_050343 | 1450 | sens | 64 | 61 | | NCBI | GCA_018107945.1 |
| | | DatstGNL2 (sc4092) HAX54_032160 | 1379 | | | | 66 | NCBI | GCA_018107945.1 |
| Solanales (Convolvulaceae) | Cuscuta australis | CusauGNOM DM860_000320 | 1440 | sens | 79 | 61 | | NCBI | GCA_003260385.1 |
| | | CusauGNL1 DM860_005101 | 1453 | sens | 63 | 58 | | NCBI | GCA_003260385.1 |
| | | CusauGNL2 DM860_001637 | 1354 | sens | | | 60 | NCBI | GCA_003260385.1 |

| | | | | | | | | | |
|---|---|---|---|---|---|---|---|---|---|
| Solanales (Convolvulaceae) | Ipomoea nil | Iponi_LOC109166281 (IponiGNOMa / sc 3087) | 1458 | sens | 83 | 62 | | NCBI | GCA_001879475.1 |
| | | Iponi_LOC109153697 (IponiGNOMb / sc 2516) | 1460 | res | 65 | 62 | | NCBI | GCA_001879475.1 |
| | | Iponi_LOC109180388 (IponiGNOMc / sc 1026) | 1464 | sens | 64 | 60 | | NCBI | GCA_001879475.1 |
| | | Iponi_LOC109147747 (IponiGNL2 / sc 2036) | 1354 | sens | | | 65 | NCBI | GCA_001879475.1 |
| Gentianales (Rubiaceae) | Coffea arabica | Cofar_LOC113716666 (chr 11c) | 1448 | sens | 82 | 63 | | NCBI | GCA_036785885.1 |
| | | Cofar_LOC113718983 (chr 11e) | 1454 | sens | 82 | 63 | 5200bp | NCBI | GCA_036785885.1 |
| | | Cofar_LOC113697893 (chr 7c) | 1458 | res | 66 | 61 | | NCBI | GCA_036785885.1 |
| | | Cofar_LOC113692407 (chr 6c) GNL2a | 1377 | sens | | | 64 | NCBI | GCA_036785885.1 |
| | | Cofar_LOC113695788 (chr 6e) GNL2b | 1377 | sens | | | 64 | NCBI | GCA_036785885.1 |
| Gentianales (Rubiaceae) | Coffea canephora | CofcaGNOM Cc11_g12600 + Cofca_T00032293001 | 1448 | sens | 82 | 63 | | NCBI | GCA_036785865.1 |
| | | CofcaGNL1 Cc07_g03650 Cofca_T00039609001 | 1458 | res(?) | 66 | 61 | | NCBI | GCA_036785865.1 |
| | | CofcaGNL2 Cc06_g09520 Cofca_T00041903001 | 1377 | sens | | | 64 | NCBI | GCA_036785865.1 |
| Gentianales (Rubiaceae) | Gardenia jasminoides | GarjaGNOM chr 11 | (1450) | sens | 82 | 62 | | NCBI | GCA_013103745.1 |
| | | GarjaGNL1 chr 9 | (1460) | res | 68 | 62 | | NCBI | GCA_013103745.1 |
| | | GarjaGNL2 chr 8 (DNA seq) | (1370) | sens | | | 63 | NCBI | GCA_013103745.1 |
| Gentianales (Rubiaceae) | Chiococca alba | ChialGNOM / g3955 | 1447 | sens | 83 | 62 | | Dryad | Pep file |

| | | | | | | | | | |
|---|---|---|---|---|---|---|---|---|---|
| | | ChialGNL1 / g16800 | 1458 | res | 66 | 61 | | Dryad | Pep file |
| | | ChialGNL2 / g8037 | 1378 | sens | | | 64 | Dryad | Pep file |
| Gentianales (Rubiaceae) | Ophiorrhiza pumila | OphpuGNOM (chr 11) | 1450 | sens | 81 | 63 | | NCBI | GCA_016586305.1 |
| | | OphpuGNL1 (chr 7) | (1450) | sens | 69 | 62 | | NCBI | GCA_016586305.1 |
| | | OphpuGNL2 (chr 6) Opuchr06_g0006290 | 1373 | sens | | | 65 | NCBI | GCA_016586305.1 |
| Gentianales (Rubiaceae) | Mitragyna speciosa | g16541 (GNOM) | 1452 | sens | 83 | 63 | | NCBI | GCA_024721245.1 |
| | | g28202 (GNL1) | 1458 | res | 67 | 62 | | NCBI | GCA_024721245.1 |
| | | g5965 (GNL2a) | 1378 | sens | | | 65 | NCBI | GCA_024721245.1 |
| Gentianales (Rubiaceae) | Leptodermis oblonga | LepobGNOM (LG2) | (1354) | sens | 81 | 62 | | NCBI | GCA_016801395.1 |
| | | LepobGNL1a (LG1) | (1450) | res | 67 | 61 | | NCBI | GCA_016801395.1 |
| | | LepobGNL1b (LG4) | (1467) | res | 67 | 59 | | NCBI | GCA_016801395.1 |
| | | LepobGNL2 (LG1) | (1370) | sens | | | 60 | NCBI | GCA_016801395.1 |
| Gentianales (Rubiaceae) | Gynochthodes officinalis | GynofGNOM (LG10) | | sens | 82 | 64 | | NCBI | GCA_046128155.1 |
| | | GynofGNL1 (LG7) | | sens | 70 | 60 | | NCBI | GCA_046128155.1 |
| | | GynofGNL2 (LG1) | | sens | | | | NCBI | GCA_046128155.1 |
| Gentianales (Apocynaceae) | Catharanthus roseus | CRO_018704 | 1451 | sens | 82 | 64 | | NCBI | GCA_028651295.1 |

| | | | | | | | | | |
|---|---|---|---|---|---|---|---|---|---|
| | | CRO_021518 | 1458 | sens | 65 | 61 | | NCBI | GCA_028651295.1 |
| | | CRO_013169 | 1368 | sens | | | 64 | NCBI | GCA_028651295.1 |
| Gentianales (Apocynaceae) | Calotropis gigantea | CalgiGNOM / cal_g003986 | 1457 | sens | 82 | 63 | | Dryad | Pep file |
| | | CalgiGNL1 / cal_g002145 | 1459 | sens | 67 | 62 | | Dryad | Pep file |
| | | CalgiGNL2 / cal_g000739 | 1393 | sens | | | 57 | Dryad | Pep file |
| Gentianales (Apocynaceae) | Rhazya stricta | RhastGNOM / sc3_609 | (1450) | sens | 84 | 64 | | NCBI | GCA_001752375.1 |
| | | RhastGNL1 / sc13 | (1460) | sens | 71 | 64 | | NCBI | GCA_001752375.1 |
| | | RhastGNL2 / sc10 | (1375) | sens | | | 61 | NCBI | GCA_001752375.1 |
| | | RhastGNL2 / sc1 | (1380) | sens | | | 59 | NCBI | GCA_001752375.1 |
| Gentianales (Apocynaceae) | Apocynum venetum | ApoveGNOM (ctg00132) | (1450) | sens | 82 | 64 | | NCBI | GCA_049998395.1 |
| | | ApoveGNL1a (ctg00123) | (1460) | sens | 71 | 63 | | NCBI | GCA_049998395.1 |
| | | ApoveGNL1b (ctg00256) | (1460) | sens | 74 | 64 | | NCBI | GCA_049998395.1 |
| | | ApoveGNL2 (ctg00128) | (1374) | sens | | | 57 | NCBI | GCA_049998395.1 |
| Gentianales (Apocynaceae) | Cynanchum aphyllum | CynwiGNOM (ctg04473) | 1457 | sens | 82 | 63 | | NCBI | GCA_025388415.1 |
| | | CynwiGNL1 (ctg02870) | 1459 | sens | 67 | 62 | | NCBI | GCA_025388415.1 |
| | | CynwiGNL2 (ctg05191) | 1413 | sens | | | 57 | NCBI | GCA_025388415.1 |

| | | | | | | | | | |
|---|---|---|---|---|---|---|---|---|---|
| Gentianales (Gentianaceae) | Gentiana dahurica | GendaGNOMa (chr 10, Mb 30.4) | 1452 | sens | 79 | 61 | | NCBI | GCA_024500145.1 |
| | | GendaGNOMb (chr 13, Mb 73.9) | 1453 | sens | 62 | 58 | | NCBI | GCA_024500145.1 |
| | | GendaGNOMc (chr 1, Mb 27.2) | (1370) | res | 58 | 55 | | NCBI | GCA_024500145.1 |
| | | GendaGNL2 (chr 10, Mb 8.3) | 1385 | sens | | | 61 | NCBI | GCA_024500145.1 |
| Gentianales (Gelsemiaceae) | Gelsemium sempervirens | GelseGNOM scaffold_8 | (1442) | sens | 82 | 62 | | Dryad | Pep file |
| | | GelseGNL1 scaffold_0 | 1461 | res | 67 | 62 | | Dryad | Pep file |
| | | GelseGNL2 scaffold_147 | 1375 | sens | | | 64 | Dryad | Pep file |
| Boraginales (Boraginaceae) | Echium plantagineum | EchplGNOMa, LG8 (genomic DNA) QFAX02000008 | 1426 | sens | 78 | 61 | | NCBI | GCA_003412495.2 |
| | | EchplGNOMb, LG3 (26.11 Mb; 2 genes 100 kb apart) QFAX02000003 | 1429 | sens | 64 | 60 | | NCBI | GCA_003412495.2 |
| | | EchplGNOMc, LG3 (26.21 Mb; gene 2) QFAX02000003 | 1429 | sens | 64 | 61 | | NCBI | GCA_003412495.2 |
| | | EchplGNL2, LG4 (genomic DNA) QFAX02000004 | 1372 | sens | | | 63 | NCBI | GCA_003412495.2 |
| Boraginales (Boraginaceae) | Lithospermum erythrorhizon | Leryth_024862 (contig_1784) LiterGNOMa | 1443 | sens | 79 | 62 | | NCBI | GCA_019722365.1 |
| | | Leryth_011110 (contig_258) (LiterGNOMb) | 1423 | sens | 79 | 62 | | NCBI | GCA_019722365.1 |
| | | Leryth_025748 (contig_1791) LiterGNL1 | 1456 | sens | 65 | 61 | | NCBI | GCA_019722365.1 |
| | | Leryth_026338 (contig_1512) (LiterGNL2) | 1376 | sens | | | 64 | NCBI | GCA_019722365.1 |

| | | | | | | | | | |
|---|---|---|---|---|---|---|---|---|---|
| Boraginales (Cordiaceae) | Cordia subcordata | CorsuGNOM (ctg 20) | 1454 | sens | 82 | 64 | | NCBI | GCA_0285 54755.1 |
| | | CorsuGNL1 (ctg 19) | 1463 | res | 69 | 63 | | NCBI | GCA_0285 54755.1 |
| | | CorsuGNL2 (ctg 8) | 1380 | sens | | | 64 | NCBI | GCA_0285 54755.1 |
| Boraginales (Ehretiaceae) | Ehretia anacua | EhranGNOM (sc 13) EhanaH1.13G059800 | 1394 | sens | 83 | 64 | | Phytozome 1 | HAP1 v1.1 |
| | | EhranGNL1 (sc 12) EhanaH1.12G126500 | 1463 | sens | 67 | 64 | | Phytozome 1 | HAP1 v1.1 |
| | | EhranGNL2 (sc 9) EhanaH1.09G100900 | 1379 | sens | | | 65 | Phytozome 1 | HAP1 v1.1 |
| Boraginales (Ehretiaceae) | Ehretia macrophylla | EhrmaGNOM (chr4, Mb 65.0) | 1454 | sens | 83 | 64 | | NCBI | GCA_0379 74665.1 |
| | | EhrmaGNL1 (chr2, Mb 49.1) | 1463 | sens | 68 | 63 | | NCBI | GCA_0379 74665.1 |
| | | EhrmaGNL2 (chr16, Mb 11.6) | 1366 | sens | | | 65 | NCBI | GCA_0379 74665.1 |
| Lamiales (Oleaceae) | Olea europaea | ~~OleeuGNOM_ctg_Oe6 _s00085~~ OLEA9_A035984 (chr11)GNOM | 1450 | sens | 80 | 64 | | NCBI | GCA_0027 42605.1 |
| | | ~~OleeuGNL1_ctg_Oe6 _s05964~~ OLEA9_A009008 (chr 10) GNL1a | 1460 | sens | 65 | 61 | | NCBI | GCA_0027 42605.1 |
| | | ~~OleeuGNL1_ctg_Oe6 _s08155~~ OLEA9_A112886 (chr 14) GNL1b | 1446 | sens | 64 | 61 | | NCBI | GCA_0027 42605.1 |
| | | OleeuGNL2_ctg_Oe6 _s03921 | (1375) | sens | | | 63 | NCBI | GCA_0027 42605.1 |
| Lamiales (Oleaceae) | Forsythia suspensa | ForsuGNOM Contig01028 (chr 4) | (1450) | sens | 82 | 70 | | NCBI | GCA_0236 38005.1 |
| | | ForsuGNL1 Contig01032 (chr 11) 2 GNL1 tandem | (1450) | res | 71 | 63 | | NCBI | GCA_0236 38005.1 |

| | | | | | | | | | |
|---|---|---|---|---|---|---|---|---|---|
| | | ForsuGNL1 defective Contig01032 (40 kb away) | (1050) | res | 63 | 54 | | NCBI | GCA_0236 38005.1 |
| | | ForsuGNL2 Contig00008 (chr 3) | (1375) | sens | | | 62 | NCBI | GCA_0236 38005.1 |
| Lamiales (Oleaceae) | Fraxinus excelsior | FraexGNOMa (chr 7) | 1461 | res | 79 | 63 | | NCBI | GCA_9652 26085.1 |
| | | FraexGNOMb (chr 15) | 1461 | res | 80 | 63 | | NCBI | GCA_9652 26085.1 |
| | | FraexGNOMc (chr 8) | 1456 | sens | 66 | 62 | | NCBI | GCA_9652 26085.1 |
| | | FraexGNL2 (chr 1) | 1365 | sens | | | 65 | NCBI | GCA_9652 26085.1 |
| Lamiales (Oleaceae) | Fraxinus pennsylvanica | FrapeGNOMa (chr 9) | 1448 | sens | 79 | 62 | | NCBI | GCA_9121 72775.2 |
| | | FrapeGNOMb (chr 17) | 1460 | res | 65 | 62 | | NCBI | GCA_9121 72775.2 |
| | | FrapeGNL2 (chr 1) | 1366 | sens | | | 65 | NCBI | GCA_9121 72775.2 |
| Lamiales (Oleaceae) | Osmanthus fragrans | OsmfrGNOMa (chr 13) | (1450) | sens | 80 | 62 | | NCB) | GCA_0193 95295.1 |
| | | OsmfrGNOMb (chr 7) | (1440) | sens | 72 | 61 | | NCB) | GCA_0193 95295.1 |
| | | OsmfrGNOMc (chr 14) | (1430) | res | 69 | 60 | | NCB) | GCA_0193 95295.1 |
| | | OsmfrGNL2 (chr 1) | (1360) | sens | | | 63 | NCB) | GCA_0193 95295.1 |
| Lamiales (Gesneriaceae) | Primulina huaijiensis | PrihuGNOMa (chr 14) | 1445 | sens | 79 | 61 | | NCBI | GCA_0122 95235.2 |
| | | PrihuGNOMc (chr 3, Mb 28.8) | 1420 | sens | 64 | 61 | | NCBI | GCA_0122 95235.2 |
| | | PrihuGNOMb (chr 3, Mb 29.7) | 1443 | sens | 79 | 62 | | NCBI | GCA_0122 95235.2 |
| | | PrihuGNOMd (sc40239) | 1458 | sens | 64 | 61 | | NCBI | GCA_0122 95235.2 |

| | | | | | | | | | |
|---|---|---|---|---|---|---|---|---|---|
| | | PrihuGNL2 (chr 3, Mb 24.7) | 1362 | sens | | | 64 | NCBI | GCA_0122 95235.2 |
| Lamiales (Gesneriaceae) | Henckelia pumila | HenpuGNOM (chr2, Mb 117.3) | 1445 | sens | 79 | 62 | | NCBI | GCA_0335 68475.2 |
| | | HenpuGNL1 (chr4, Mb 29.2) | 1458 | sens | 65 | 61 | | NCBI | GCA_0335 68475.2 |
| | | HenpuGNL2 (chr4, Mb 43.7) | | | | | n.a. | NCBI | GCA_0335 68475.2 |
| Lamiales (Plantaginaceae) | Antirrhinum majus | Am03g32310 AntmaGNOM (chr 3) | 1450 | sens | 80 | 61 | | CNGB | Pep file |
| | | Am06g34280 AntmaGNL1 (chr 6) | 1435 | sens | 65 | 61 | | CNGB | Pep file |
| | | Am06g18620 AntmaGNL2 (chr 6) | 1377 | sens | | | 64 | CNGB | Pep file |
| Lamiales (Plantaginaceae) | Plantago ovata | PlaovGNOMa (chr1, Mb 28.9) 100% id. Pov_00004964 | 1442 | sens | 79 | 62 | | NCBI | GCA_0282 74465.1 |
| | | PlaovGNOMb (chr1, Mb 135.5) 100% id. Pov_00012506 | 1441 | sens | 65 | 62 | | NCBI | GCA_0282 74465.1 |
| | | PlaovGNL2 (chr4, Mb 0.5) | 1358 | sens | | | 61 | NCBI | GCA_0282 74465.1 |
| Lamiales (Scrophulariaceae) | Buddleja alternifolia | Bualt11G0038600 BudalGNOM (chr 11) | 1446 | sens | 81 | 63 | | NCBI | GCA_0194 26215.1 |
| | | Bualt04G0142100 BudalGNL1 (chr 4) | 1438 | sens | 67 | 63 | | NCBI | GCA_0194 26215.1 |
| | | Bualt07G0105600 BudalGNL2 (chr 7) | 1366 | sens | | | 65 | NCBI | GCA_0194 26215.1 |
| Lamiales (Scrophulariaceae) | Verbascum thapsus | VerthGNOM (chr14, Mb2.5) | 1445 | sens | 80 | 62 | | NCBI | GCA_9636 69365.1 |
| | | VerthGNL1a (chr4, Mb 21.4) | 1432 | sens | 67 | 62 | | NCBI | GCA_9636 69365.1 |
| | | VerthGNL1b (chr9, Mb 4.1) | 1431 | sens | 66 | 63 | | NCBI | GCA_9636 69365.1 |

| | | | | | | | | | |
|---|---|---|---|---|---|---|---|---|---|
| | | VerthGNL2 (chr5, Mb 6.2) | 1359 | sens | | | 65 | NCBI | GCA_963669365.1 |
| Lamiales (Linderniaceae) | Lindernia brevidens | LinbrGNOM SWDC01000156 Sc7 | (1450) | sens | 78 | 62 | | NCBI | GCA_004919715.1 |
| | | LinbrGNL1a SWDC01000148 Sc12 | (1450) | sens | 64 | 62 | | NCBI | GCA_004919715.1 |
| | | LinbrGNL1b SWDC01000151 Sc2 | (1450) | sens | 64 | 58 | | NCBI | GCA_004919715.1 |
| | | LinbrGNL2 SWDC01000155 Sc6 | (1375) | sens | | | 60 | NCBI | GCA_004919715.1 |
| Lamiales (Lentibulariaceae) | Genlisea aurea | EPS60000 (sc7838) | 1435 | sens | 74 | 59 | | NCBI | GCA_000441915.1 |
| | | EPS65079 | 1424 | sens | 63 | 61 | | NCBI | GCA_000441915.1 |
| | | EPS71757 | 1382 | sens | 51 | 51 | | NCBI | GCA_000441915.1 |
| | | EPS59939 | 1366 | sens | | | 61 | NCBI | GCA_000441915.1 |
| Lamiales (Lentibulariaceae) | Utricularia gibba | UtrgiGNOM ctg749. | 1450 | sens | 76 | 61 | | NCBI | GCA_002189035.1 |
| | | UtrgiGNL1 ctg736. | 1443 | sens | 60 | 60 | | NCBI | GCA_002189035.1 |
| | | UtrgiGNL2 (chr 1) | 1357 | sens | | | 60 | NCBI | GCA_002189035.1 |
| Lamiales (Lentibulariaceae) | Utricularia reniformis | UtrreGNOM (sc233, Mb 0.2) | 1450 | sens | 77 | 60 | | NCBI | GCA_009725065.1 |
| | | UtrreGNL1 (sc174, Mb 0.4) | 1445 | sens | 60 | 59 | | NCBI | GCA_009725065.1 |
| | | UtrreGNL2 (sc5, Mb 1.0) | 1361 | sens | | | 61 | NCBI | GCA_009725065.1 |
| Lamiales (Lentibulariaceae) | Pinguicula primuliflora | PinprGNOM (sc20, Mb 0.01) | 1438 | sens | 78 | 61 | | NCBI | GCA_025802335.1 |
| | | PinprGNL1 (sc15, Mb 0.01) | 1441 | sens | 62 | 60 | | NCBI | GCA_025802335.1 |

| | | | | | | | | | |
|---|---|---|---|---|---|---|---|---|---|
| | | PinprGNL2 (sc1, Mb 0.9) | 1369 | sens | | | 61 | NCBI | GCA_025802335.1 |
| Lamiales (Acanthaceae) | Andrographis paniculata | AndpaGNOM (chr 19) | 1486 | sens | 80 | 62 | | NCBI | GCA_009805555.1 |
| | | AndpaGNL1 (chr 2) | 1451 | sens | 64 | 61 | | NCBI | GCA_009805555.1 |
| | | AndpaGNL2 (chr 10) | 1359 | | | | 62 | NCBI | GCA_009805555.1 |
| Lamiales (Acanthaceae) | Avicennia marina | MSTRG.7779/ AvimaGNOM (chr 9) | 1457 | sens | 79 | 61 | | NCBI | GCA_019155195.1 |
| | | MSTRG.3029/ AvimaGNL1 (chr 16) | 1439 | res | 65 | 61 | | NCBI | GCA_019155195.1 |
| | | AvimaGNL2 (chr 27) | (1373) | sens | | | 62 | NCBI | GCA_019155195.1 |
| Lamiales (Pedaliaceae) | Sesamum indicum | Sesin_LOC105159998 (LG4) | 1453 | sens | 81 | 62 | | NCBI | GCA_000512975.1 |
| | | Sesin_LOC105177279 (LG15) | 1439 | res | 67 | 62 | | NCBI | GCA_000512975.1 |
| | | Sesin_LOC105158259 (LG3) | 1356 | sens | | | 65 | NCBI | GCA_000512975.1 |
| Lamiales (Verbenaceae) | Petrea volubilis | PetvoGNOMa (chr 13) Petvo.13G010580 | 1457 | sens | 82 | 63 | | NCBI | GCA_026212405.1 |
| | | PetvoGNOMb (chr 4) Petvo.4G002870 | 1445 | sens | 77 | 60 | | NCBI | GCA_026212405.1 |
| | | PetvoGNL1a (chr 3) Petvo.3G004050 | 1437 | sens | 67 | 62 | | NCBI | GCA_026212405.1 |
| | | PetvoGNL1b (unplaced) Petvo.S008240 | 1297 | sens | 58 | 55 | | NCBI | GCA_026212405.1 |
| | | PetvoGNL2 (chr 2) Petvo.2G006980 | 1338 | sens | | | 62 | NCBI | GCA_026212405.1 |
| Lamiales (Verbenaceae) | Verbena officinalis | VerofGNOM (chr4, Mb 40.0) | 1448 | sens | 80 | 62 | | NCBI | GCA_958496215.1 |
| | | VerofGNL1a (chr1, Mb 22.7) | 1440 | sens | 66 | 62 | | NCBI | GCA_958496215.1 |

| | | | | | | | | | |
|---|---|---|---|---|---|---|---|---|---|
| | | VerofGNL1b (chr3, Mb 34.0) | 1411 | sens | 61 | 59 | | NCBI | GCA_958496215.1 |
| | | VrofGNL2 (chr7, Mb 14.0) | 1357 | sens | | | 63 | NCBI | GCA_958496215.1 |
| Lamiales (Verbenaceae) | Lantana camara | LancaGNOM (chr9, Mb 9.4) | 1450 | sens | 80 | 62 | | NCBI | GCA_037501455.1 |
| | | LancaGNL1a (chr4, Mb 4.1) | 1434 | res | 66 | 63 | | NCBI | GCA_037501455.1 |
| | | LancaGNL1b (chr1, Mb 1.6) | 1413 | sens | 61 | 59 | | NCBI | GCA_037501455.1 |
| | | LancaGNL2 (chr5, Mb 78.1) | 1348 | sens | | | 62 | NCBI | GCA_037501455.1 |
| Lamiales (Bignoniaceae) | Handroanthus impetiginosus | HanimGNOM (sc4829) CDL12_22191 | 1445 | sens | 82 | 63 | | NCBI | GCA_002762385.1 |
| | | HanimGNL1 (sc5545) CDL12_24118 | 1440 | res | 76 | 64 | | NCBI | GCA_002762385.1 |
| | | HanimGNL2 (sc3298) CDL12_17220 | 1364 | sens | | | 65 | NCBI | GCA_002762385.1 |
| Lamiales (Bignoniaceae) | Jacaranda mimosifolia | Jmimo29902 / JacmiGNOM (chr 18) | 1450 | sens | 81 | 63 | | NCBI | GCA_018894105.1 |
| | | Jmimo29048 / JacmiGNL1a (chr 17) | (1420) | sens | 64 | 60 | | NCBI | GCA_018894105.1 |
| | | Jmimo10406 / JacmiGNL1b (chr 5) | 1440 | sens | 66 | 63 | | NCBI | GCA_018894105.1 |
| | | Jmimo15631 / JacmiGNL2 (chr 8) | 1372 | sens | | | 65 | NCBI | GCA_018894105.1 |
| Lamiales (Paulowniaceae) | Paulownia fortunei | PaufoGNOMa (chr17) Pfo_023964 | 1456 | sens | 81 | 63 | | NCBI | GCA_040144305.1 |
| | | PaufoGNL1 (chr 6) Pfo_008936 | 1442 | sens | 67 | 62 | | NCBI | GCA_040144305.1 |
| | | PaufoGNOMb (chr 7) pseudo? sequencing errors? | (1453) | sens | 84 | 69 | | NCBI | GCA_040144305.1 |
| | | PaufoGNL2 (chr 8) Paufo_011839 | 1365 | sens | | | 65 | NCBI | GCA_040144305.1 |

| | | | | | | | | | |
|---|---|---|---|---|---|---|---|---|---|
| Lamiales (Orobanchaceae) | Striga asiatica | STAS_24345 (sc455) | 1453 | sens | 80 | 61 | | NCBI | GCA_0086 36005.1 |
| | | STAS_22812 (sc413) | 1443 | sens | 65 | 61 | | NCBI | GCA_0086 36005.1 |
| | | STAS_20940 + aa182-334 fragment (synthetic) (sc37) | (1174) | | | | 63 | NCBI | GCA_0086 36005.1 |
| Lamiales (Orobanchaceae) | Lindenbergia philippensis | Liphi.05G047100 (chr 5) | 1445 | sens | 80 | 61 | | Phytozome 13 | V1.0 |
| | | Liphi.09G029200 (chr 9) | 1431 | sens | 67 | 63 | | Phytozome 13 | V1.0 |
| | | Liphi.08G097200 (chr 8) | 1373 | sens | | | 65 | Phytozome 13 | V1.0 |
| Lamiales (Orobanchaceae) | Orobanche cumana | 12G01204/ OrocuGNOM | 1453 | sens | 79 | 60 | | CNGB | GWH Pep file |
| | | 3G02103 / OrocuGNL1 | 1442 | sens | 66 | 61 | | CNGB | GWH Pep file |
| | | 2G00339 / OrocuGNL2 | 1369 | sens | | | 64 | CNGB | GWH Pep file |
| Lamiales (Phrymaceae) | Mimulus (Erythranthe) guttatus | Migut.L01245 (sc37) Erygu_LOC10596756 3 | 1451 | sens | 80 | 63 | | NCBI | GCA_0005 04015.1 |
| | | Migut.H00675 (sc11) Erygu_LOC10597090 4 | 1442 | res | 66 | 62 | | NCBI | GCA_0005 04015.1 |
| | | Migut.I00641 (sc7) Erygu_LOC10597603 5 | 1372 | sens | | | 64 | NCBI | GCA_0005 04015.1 |
| Lamiales (Lamiaceae) | Salvia hispanica | SalhiGNOMa / SHI_00000013 Salhi_LOC12521128 6 (chr 3) | 1444 | sens | 80 | 61 | | NCBI | GCA_0231 19035.1 |
| | | SalhiGNOMb / SHI_00015079 Salhi_LOC12520590 5 (chr 2) | 1449 | sens | 80 | 61 | | NCBI | GCA_0231 19035.1 |
| | | SalhiGNL1 / SHI_00016683 Not in NCBI genome | 1440 | res | 66 | 62 | | NCBI | GCA_0231 19035.1 |

| | | | | | | | | | |
|---|---|---|---|---|---|---|---|---|---|
| | | SalhiGNL2 / SHI_00029737 Salhi_LOC125189731 (chr 5) | 1385 | sens | | | 62 | NCBI | GCA_023119035.1 |
| Lamiales (Lamiaceae) | Scutellaria baicalensis | ScubaGNOM (chr 2) | 1452 | sens | 80 | 63 | | NCBI | GCA_005771605.1 |
| | | ScubaGNL1 (chr 1) | 1437 | res | 66 | 63 | | NCBI | GCA_005771605.1 |
| | | ScubaGNL2 (chr 3) | 1342 | sens | | | 61 | NCBI | GCA_005771605.1 |
| Lamiales (Lamiaceae) | Perilla citriodora | C2S52_015980 (contig_PC02_8) | 1455 | sens | 81 | 62 | | NCBI | GCA_026008535.1 |
| | | C2S52_002474 (contig_PC02_3) | 1441 | res | 68 | 62 | | NCBI | GCA_026008535.1 |
| | | C2S52_018783 DNA missing (contig_PC02_9) | 665 | sens | | | | NCBI | GCA_026008535.1 |
| | | C2S52_001248 (contig_PC02_1) | 1366 | sens | | | 62 | NCBI | GCA_026008535.1 |
| Lamiales (Lamiaceae) | Tectona grandis | TecgrGNOM / Tg15g11250 | 1457 | sens | 82 | 63 | | GigaSci | Pep file |
| | | TecgrGNL1 / Tg09g16010 | 1440 | res | 66 | 62 | | GigaSci | Pep file |
| | | TecgrGNL2 / Tg10g09750 | 1376 | sens | | | 64 | GigaSci | Pep file |
| Lamiales (Lamiaceae) | Ajuga decumbens | AjudeGNOM / g7586 | 1446 | sens | 81 | 63 | | Figshare | Pep file |
| | | AjudeGNL1 / g24258 | 1438 | sens | 67 | 62 | | Figshare | Pep file |
| | | AjudeGNL2 / g29748 | 1366 | sens | | | 60 | Figshare | Pep file |
| Lamiales (Lamiaceae) | Mentha longifolia | MenloGNOMa (chr 12) | (1445) | sens | 81 | 63 | | NCBI | GCA_001642375.2 |
| | | MenloGNOMb (chr 2, 19.5 Mb) | (1450) | sens | 81 | 60 | | NCBI | GCA_001642375.2 |
| | | MenloGNL1 (chr 2, 19.5 Mb) | (1440) | res | 68 | 61 | | NCBI | GCA_001642375.2 |

| | | | | | | | | | |
|---|---|---|---|---|---|---|---|---|---|
| | | MenloGNL2 (chr 2, 8.2 Mb) | (1370) | sens | | | 61 | NCBI | GCA_0016 42375.2 |
| Lamiales (Lamiaceae) | Agastache rugosa | AgaruGNOM (sc1) 37190 | 1450 | sens | 81 | 62 | | NCBI | GCA_0314 70985.1 |
| | | AgaruGNL1a (sc3) 02070 | 1444 | res | 66 | 62 | | NCBI | GCA_0314 70985.1 |
| | | AgaruGNL1b (sc6) 03340 (95% id. AgaruGNL1a) | 1442 | res | 66 | 62 | | NCBI | GCA_0314 70985.1 |
| | | AgaruGNL2 (sc8) 18380 | 1363 | sens | | | 62 | NCBI | GCA_0314 70985.1 |

**Suppl. Table S1H. GBF1-type ARF-GEFs in superrosids (17 orders, 70 families, 164 species)**

| Order | Species | Gene name (chr/ctg/sc #) | Protein Size (# aa) | BFA res / sens | % id. AtGN | % id. AtGNL1 | % id. AtGNL2 | database | ref |
|---|---|---|---|---|---|---|---|---|---|
| Santalales (Olacaceae) | Malania oleifera | Maole12G0095000 (chr12) 100% id. Maole_005233 MalolGNOM | 1456 | sens | 84 | 63 | | NCBI | GCF_029873635.1 |
| | | Maole01G0176400 (chr1) 100% id. Maole_012409 MalolGNL1 | 1465 | res | 69 | 64 | | NCBI | GCF_029873635.1 |
| | | Maole04G0021300 (chr4) 100% id. Maole_013225 MalolGNL2a | 1410 | sens | | | 62 | NCBI | GCF_029873635.1 |
| | | Maole09G0160400 (chr9) 97% id. Maole_014154 MalolGNL2b | 1411 | sens | | | 61 | NCBI | GCF_029873635.1 |
| Santalales (Santalaceae) | Santalum album | SanalGNOM (chr2, Mb 12.0) | 1450 | sens | 81 | 63 | | NCBI | GCA_034195605.1 |
| | | SanalGNL1 (chr8, Mb 12.3) | 1464 | res | 68 | 64 | | NCBI | GCA_034195605.1 |
| | | SanalGNL2 (chr9, Mb 3.2) | | | | | 63 | NCBI | GCA_034195605.1 |
| Santalales (Santalaceae) | Santalum yasi | gene10972_Y1 SanyaGNOM | 1450 | sens | 81 | 62 | | Figshare | Pep file |
| | | gene21469_Y1 SanyaGNL1 | 1465 | res | 68 | 64 | | | |
| | | gene13746_Y1 SanyaGNL2 | 1405 | sens | | | 63 | | |
| Santalales (Loranthaceae) | Taxillus chinensis | TaxchGNOMb (chr5, Mb54.18) | 1409 | sens | 77 | 59 | | NCBI | GCA_023512835.1 |
| | | TaxchGNOMa (chr 8, Mb10.75) | 1441 | sens | 78 | 59 | | NCBI | GCA_023512835.1 |
| | | TaxchGNL1b (chr 2, Mb 38.05) | 1454 | sens | 68 | 62 | | NCBI | GCA_023512835.1 |

| | | | | | | | | | |
|---|---|---|---|---|---|---|---|---|---|
| | | TaxchGNL1a (chr 6, Mb 24.59) | 1468 | res | 66 | 61 | | NCBI | GCA_0235 12835.1 |
| | | TaxchGNL2 (chr 3, Mb 13.7) | (1380) | sens | | | (59) | NCBI | GCA_0235 12835.1 |
| Santalales (Balanophoraceae) | Balanophora fungosa | BalfuGNOM (ctg35) BALA12295 | 1393 | sens | 60 | 51 | | CNGB | GWH Pep file |
| | | BalfuGNL1 (ctg7706) BALA05290 | 1386 | res | 51 | 51 | | CNGB | GWH Pep file |
| | | BalfuGNL2 (ctg8) BALA14597 | 952 | sens | | | n.a. | CNGB | GWH Pep file |
| Vitales (Vitaceae) | Vitis vinifera | Viv_NC_012008 Vitvi_LOC100247152 (chr2) | 1470 | sens | 83 | 63 | | NCBI | GCA_0307 04535.1 |
| | | Viv_NC_012017 Vitvi_LOC100255904 (chr11) | 1390 | sens | | | 66 | NCBI | GCA_0307 04535.1 |
| Vitales (Vitaceae) | Vitis riparia | LOC117929271 (chr2) VitriGNOM | 1470 | sens | 83 | 63 | | NCBI | GCF_0043 53265.1 |
| | | LOC117925203 (chr11) VitriGNL2 | 1390 | sens | | | 66 | NCBI | GCF_0043 53265.1 |
| Saxifragales (Saxifragaceae) | Tiarella polyphylla | TiapoGNOM (LG2) | 1469 | sens | 84 | 63 | | CNGB | GWH Üeü file |
| | | TiapoGNL1 (LG1) | 1460 | sens | 70 | 66 | | CNGB | GWH Üeü file |
| | | TiapoGNL2 (LG7) | 1371 | sens | | | 67 | CNGB | GWH Üeü file |
| Saxifragales (Saxifragaceae) | Chrysosplenium sinicum | ChrsiGNOM (chr11) Cs11T0015900 | 1438 | sens | 81 | 62 | | Figshare | Pep file |
| | | ChrsiGNL1 (chr1) Cs1T0139900 | 1462 | sens | 66 | 63 | | Figshare | Pep file |
| | | ChrsiGNL2 (chr12) Cs12T0061100 (aa322-364 deleted) | 1316 | sens | | | 62 | Figshare | Pep file |
| Saxifragales (Crassulaceae) | Kalanchoe fedtschenkoi | Kaladp0061s0170 Kalfe_61s0170 | 1457 | sens | 82 | 63 | | NCBI | GCA_0023 12845.1 |
| | | Kaladp0043s0155 | 1449 | sens | 80 | 62 | | NCBI | GCA_0023 12845.1 |

| | | | | | | | | | |
|---|---|---|---|---|---|---|---|---|---|
| | | Kaladp1222s0075 | 1449 | sens | 64 | 61 | | NCBI | GCA_0023 12845.1 |
| | | Kaladp0096s0081 Kalfe_96s0081 | 1477 | res | 63 | 60 | | NCBI | GCA_0023 12845.1 |
| | | Kaladp0031s0117 Kalfe_31s0117 | 1391 | sens | | | 55 | NCBI | GCA_0023 12845.1 |
| Saxifragales (Crassulaceae) | Kalanchoe laxiflora | Kalaxd.06G122200 (chr 6) KallaGNOMa | 1457 | sens | 83 | 61 | | Phytozome 13 | V1.1 |
| | | Kalaxd.05G133200 (chr 5) KallaGNOMb | 1446 | sens | 80 | 61 | | Phytozome 13 | V1.1 |
| | | Kalaxd.07G141300 (chr 7) KallaGNL1a | 1452 | res | 65 | 60 | | Phytozome 13 | V1.1 |
| | | Kalaxd.04G035400 (chr 4) KallaGNL1b | 1444 | sens | 66 | 62 | | Phytozome 13 | V1.1 |
| | | Kalaxd.11G129200 (chr 11) KallaGNL2 | 1387 | sens | | | 55 | Phytozome 13 | V1.1 |
| Saxifragales (Crassulaceae) | Sedum album | SedalGNOM (ctg 13) | 1455 | sens | 80 | 61 | | NCBI | GCA_0064 09495.1 |
| | | SedalGNL1a (ctg 317 | 1432 | sens | 64 | 60 | | NCBI | GCA_0064 09495.1 |
| | | SedalGNL1b (ctg 25) | 1449 | res | 58 | 56 | | NCBI | GCA_0064 09495.1 |
| | | SedalGNL2 (ctg 425) | 1389 | sens | | | 55 | NCBI | GCA_0064 09495.1 |
| Saxifragales (Crassulaceae) | Rhodiola kirilowii | RhokiGNOMa (chr10, Mb 4.3) | 1453 | sens | 81 | 62 | | NCBI | GCA_0408 69135.1 |
| | (NCBI Nov23) | RhokiGNOMb (chr1, Mb 51.7) | 1433 | sens | 63 | 60 | | NCBI | GCA_0408 69135.1 |
| | | RhokiGNOMc (chr1, Mb 51.5) | (1298) | | | | | NCBI | GCA_0408 69135.1 |
| | | RhokiGNL2 (chr6, Mb 28.4) | | | | | | NCBI | GCA_0408 69135.1 |
| Saxifragales (Crassulaceae) | Umbilicus rupestris | UmbruGNOM (chr1, Mb 1.1) | 1456 | sens | 81 | 62 | | NCBI | GCA_9636 78935.1 |

| | | | | | | | | | |
|---|---|---|---|---|---|---|---|---|---|
| | (NCBI Dec23) | UmbruGNL1 (chr9, Mb 12.8) | 1436 | sens | 64 | 61 | | NCBI | GCA_963678935.1 |
| | | UmbruGNL2 (chr20, Mb 1.0) | 1393 | sens | | | 55 | NCBI | GCA_963678935.1 |
| Saxifragales (Paeoniaceae) | Paeonia ludlowii | PaeluGNOM (chr 1) PL-1G204610 | 1460 | sens | 83 | 64 | | Figshare | Pep file |
| | | PaeluGNL1 (chr 2) PL-2G114280 | 1422 | res | 68 | 64 | | Figshare | Pep file |
| | | PaeluGNL2 (chr 3) PL-3G169270 | 1177 | sens | | | 66 | Figshare | Pep file |
| Saxifragales (Altingiaceae) | Liquidambar formosana | LiqfoGNOMa (chr5, Mb 39.08) | 1470 | res | 82 | 64 | | NCBI | GCA_039720395.1 |
| | | LiqfoGNOMb (chr5, Mb 38.97) | 1470 | res | 82 | 64 | | NCBI | GCA_039720395.1 |
| | | LiqfoGNOMc (chr5, Mb 39.07) | 1470 | sens | 84 | 64 | | NCBI | GCA_039720395.1 |
| | | LiqfoGNOMd (chr5, Mb 38.95) | 1470 | sens | 84 | 64 | | NCBI | GCA_039720395.1 |
| | | LiqfoGNL1 (chr13, Mb 7.4) | 1452 | sens | 71 | 66 | | NCBI | GCA_039720395.1 |
| | | LiqfoGNL2 (chr15, Mb 30.6) L1049_000212 | 1402 | sens | | | 67 | NCBI | GCA_039720395.1 |
| Saxifragales (Altingiaceae) | Liquidambar styraciflua | LiqstGNOMa (chr13, Mb 23.58) | 1470 | sens | 84 | 64 | | NCBI | GCA_041721615.1 |
| | | LiqstGNOMb (chr13, Mb 23.59) | 1470 | res | 82 | 63 | | NCBI | GCA_041721615.1 |
| | | LiqstGNL1 (chr5, Mb 40.2) | 1452 | sens | 71 | 66 | | NCBI | GCA_041721615.1 |
| | | LiqstGNL2 (chr15, Mb 15.2) | 1402 | sens | | | 67 | NCBI | GCA_041721615.1 |
| Saxifragales (Hamamelidaceae) | Hamamelis virginiana | HamviGNOMa (ctg1436) | 1469 | sens | 84 | 63 | | NCBI | GCA_017140935.1 |
| | (same contig) | HamviGNOMb (ctg1436) | 1467 | res | 89 | 63 | | NCBI | GCA_017140935.1 |

| | | | | | | | | | |
|---|---|---|---|---|---|---|---|---|---|
| | | HamviGNL2 (ctg1273) | 1398 | sens | | | 67 | NCBI | GCA_017140935.1 |
| Geraniales (Geraniaceae) | Geranium maculatum | GermaGNOMa (sc5) | 1468 (1455) | res | 81 | 63 | | NCBI | GCA_028566255.1 |
| | | GermaGNOMb (sc315179) | 1444 | sens | 74 | 60 | | NCBI | GCA_028566255.1 |
| | | GermaGNL2 | (1127) | sens | | | 57 | NCBI | GCA_028566255.1 |
| Geraniales (Geraniaceae) | Geranium rotundifolium | GerroGNOMa (chr11, Mb 7.0) | 1456 | sens | 83 | 62 | | NCBI | GCA_963920655.1 |
| | | GerroGNOMb (chr12, Mb 10.1) | 1457 | res | 82 | 53 | | NCBI | GCA_963920655.1 |
| | | GerroGNOMc (chr10, Mb 26.4) | 1448 (1467) | sens | 73 | 60 | | NCBI | GCA_963920655.1 |
| | | GerroGNOMd (chr5, Mb 34.5) | 1442 | sens | 73 | 60 | | NCBI | GCA_963920655.1 |
| | | GerroGNL2 (chr12, Mb 3.4) | 1351 | sens | | | 59 | NCBI | GCA_963920655.1 |
| Geraniales (Geraniaceae) | Erodium maritimum | EromaGNOMa (chr1, Mb 125.4) | 1455 | sens | 74 | 59 | | NCBI | GCA_963921825.1 |
| | | EromaGNOMb (chr1, Mb 29.1) | 1454 | res | 82 | 62 | | NCBI | GCA_963921825.1 |
| | | EromaGNL2 (chr1, Mb 2.2) | 1365 | sens | | | 59 | NCBI | GCA_963921825.1 |
| Geraniales (Francoaceae) | Francoa sonchifolia | FrasoGNOMa (NODE_10629 + NODE_849) | ~~1236~~ 1454 | sens | 82 | 63 | | NCBI | GCA_036897585.1 |
| | | FrasoGNOMb (NODE_6300) | 1458 | sens | 81 | 63 | | NCBI | GCA_036897585.1 |
| | | FrasoGNL1 (NODES_4687_4513) | 1461 | sens | 66 | 64 | | NCBI | GCA_036897585.1 |
| | | FrasoGNL2 (NODE_9140) | 1366 | sens | | | 62 | NCBI | GCA_036897585.1 |

| | | | | | | | | | |
|---|---|---|---|---|---|---|---|---|---|
| Crossosomatales (Staphyleaceae) | Euscaphis japonica | EusjaGNOM (LG 02) (GWHGBCHS007551 + GWHGBCHS007550) | (1311) | sens | 76 | 58 | | NGDC | GWH Pep file |
| | ( | EusjaGNL1 (LG 04) (GWHGBCHS011854) | 1464 | res | 67 | 65 | | NGDC | GWH Pep file |
| | | EusjaGNL2 (LG 03) (GWHGBCHS008788) | 1383 | sens | | | 68 | NGDC | GWH Pep file |
| Myrtales (Myrtaceae) | Eucalyptus grandis | Eucgr.B03196 Eucgr_LOC104433426 (chr 2) | 1460 | sens | 83 | 63 | | NCBI | GCA_016545825.1 |
| | | Eucgr.C03862 Eucgr_LOC104440351 (chr 2) | 1387 | sens | | | 67 | NCBI | GCA_016545825.1 |
| Myrtales (Myrtaceae) | Syzygium oleosum | Syzol_LOC115671905 | 1460 | sens | 84 | 63 | | NCBI | GCA_021117445.2 |
| | | Syzol_LOC115690464 | 1386 | sens | | | 67 | NCBI | GCA_021117445.2 |
| Myrtales (Myrtaceae) | Corymbia citriodora | Cocit.B2628 (chr 2) | 1460 | sens | 84 | 63 | | NCBI | GCA_014858505.1 |
| | | Cocit.C2224 | 1386 | sens | | | 66 | NCBI | GCA_014858505.1 |
| Myrtales (Myrtaceae) | Rhodamnia argentea | Rhoar_LOC115755598 | 1460 | sens | 84 | 63 | | NCBI | GCA_020921035.1 |
| | | Rhoar_LOC115729910 | 1386 | sens | | | 66 | NCBI | GCA_020921035.1 |
| | | Rhoar_LOC115729907 | 1386 | sens | | | 66 | NCBI | GCA_020921035.1 |
| Myrtales (Myrtaceae) | Angophora floribunda | AngflGNOM (chr 2) | (1460) | sens | 84 | 65 | | NCBI | GCA_014182895.1 |
| | | AngflGNL2 (chr 3) | (1382) | sens | | | 64 | NCBI | GCA_014182895.1 |
| Myrtales (Melastomataceae) | Melastoma dodecandrum | GWHGBCLA017923 (LG05) MeldoGNOMa | 1455 | sens | 83 | 63 | | CNGB | GWH Pep file |
| | | GWHGBCLA006626 (LG01) MeldoGNOMb | 1453 | res | 81 | 62 | | CNGB | GWH Pep file |
| | | GWHGBCLA003796 (LG01) MeldoGNL2 | 1400 | sens | | | 61 | CNGB | GWH Pep file |

| | | | | | | | | | |
|---|---|---|---|---|---|---|---|---|---|
| Myrtales (Melastomataceae) | Melastoma candidum | MelcaGNOMb (chr 4, Mb 15.5) MLD38_011920 | 1453 | res | 81 | 62 | | NCBI | GCA_0236 53495.1 |
| | | MelcaGNOMa (chr 5) MLD38_017003 | 1455 | sens | 82 | 63 | | NCBI | GCA_0236 53495.1 |
| | | MelcaGNL2 (chr 4, Mb 29.9) MLD38_015326 | 1383 | sens | | | 63 | NCBI | GCA_0236 53495.1 |
| Myrtales (Combretaceae) | Combretum micranthium | CommiGNOMa Cmmi469 | 1462 | sens | 84 | 63 | | NCBI | GCA_0241 38025.1 |
| | | CommiGNOMb Cmmi489 | 1452 | res | 63 | 53 | | NCBI | GCA_0241 38025.1 |
| | | CommiGNOMc Cmmi419 | 1373 | res | 65 | 57 | | NCBI | GCA_0241 38025.1 |
| | | CommiGNL2 Cmmi97 | 1392 | sens | | | 63 | NCBI | GCA_0241 38025.1 |
| Myrtales (Combretaceae) | Lumnitzera racemosa | LumraGNOMa (Lura7, Mb 87.4) | 1463 | sens | 84 | 63 | | NCBI | GCA_0241 38005.1 |
| | | LumraGNOMb (Lura6) | 1471 | res | 65 | 56 | | NCBI | GCA_0241 38005.1 |
| | | LumraGNL2 (Lura7, Mb 78.3) | 1390 | sens | | | 63 | NCBI | GCA_0241 38005.1 |
| Myrtales (Combretaceae) | Laguncularia racemosa | LagraGNOMa (sc90,Mb 2.9) | 1458 | sens | 85 | 63 | | NCBI | GCA_0379 54665.1 |
| | | LagraGNOMb (sc190, Mb 0.5) | 1471 | res | 65 | 56 | | NCBI | GCA_0379 54665.1 |
| | | LagraGNL2 (sc12, Mb 0.4) | 1391 | sens | | | 63 | NCBI | GCA_0379 54665.1 |
| Myrtales (Lythraceae) | Punica granatum | PungrGNOM LOC116199793 (chr 3) | 1460 | sens | 84 | 64 | | NCBI | GCA_0076 55135.2 |
| | | PungrGNL1 LOC116209960 (chr 6) | 1457 | res | 67 | 58 | | NCBI | GCA_0076 55135.2 |
| | | PungrGNL2 LOC116200818 (chr 3) | 1371 | sens | | | 64 | NCBI | GCA_0076 55135.2 |
| Myrtales (Lythraceae) | Sonneratia alba | SonalGNOMa (sc30) | (1460) | sens | 82 | 61 | | NCBI | GCA_9001 74645.1 |

| | | | | | | | | | |
|---|---|---|---|---|---|---|---|---|---|
| | | SonalGNOMb (sc4) | (1469) | res | 68 | 58 | | NCBI | GCA_900174645.1 |
| | | SonalGNL2 (sc17) | 1381 | sens | | | 62 | NCBI | GCA_900174645.1 |
| Myrtales (Onagraceae) | Chamaenerion angustifolium | ChaanGNOMa (ctg291) (chr3) | 1455 | sens | 82 | 61 | | NCBI | GCA_946814005.1 |
| | | ChaanGNOMb (ctg1951) (chr9) | 1456 | sens | 81 | 61 | | NCBI | GCA_946814005.1 |
| | | ChaanGNOMc (ctg251) (chr13) | (1450) | sens | 62 | 55 | | NCBI | GCA_946814005.1 |
| | | ChaanGNOMd (ctg321) (chr12) | | sens | 63 | 54 | | NCBI | GCA_946814005.1 |
| | | ChaanGNL2 (ctg81) (chr11) | 1385 | sens | | | 63 | NCBI | GCA_946814005.1 |
| Myrtales (Onagraceae) | Clarkia xantiana | ClaxaGNOMa (sc100017) | 1455 | sens | 81 | 61 | | NCBI | GCA_036873135.1 |
| | | ClaxaGNOMb (sc100004) | 1382 | sens | 61 | 53 | | NCBI | GCA_036873135.1 |
| | | ClaxaGNOMc (sc100001, Mb 5.2) | 1381 | sens | 61 | 54 | | NCBI | GCA_036873135.1 |
| | | ClaxaGNL2a (sc100001, Mb 37.9) | 1416 | sens | | | 61 | NCBI | GCA_036873135.1 |
| | | ClaxaGNL2b (sc100010) | 1387 | sens | | | 62 | NCBI | GCA_036873135.1 |
| Zygophyllales (Zygophyllaceae) | Tetraena mongolica | TetmoGNOMa (chr A6) TmoA06G007330 | 1460 | sens | 81 | 62 | | figshare | Pep file |
| | | TetmoGNOMb (chr B6) TmoB06G007140 | 1460 | sens | 81 | 61 | | figshare | Pep file |
| | | TetmoGNOMc (chr A10) TmoA10G007160 | 1465 | sens | 69 | 60 | | figshare | Pep file |
| | | TetmoGNOMd (chr B10) TmoB10G007680 | 1465 | sens | 69 | 60 | | figshare | Pep file |

| | | | | | | | | | |
|---|---|---|---|---|---|---|---|---|---|
| | | TetmoGNL2a (ctg 581) Tmoptg000058lG000 680 | 1377 | sens | | | 65 | figshare | Pep file |
| | | TetmoGNL2b (chr A4) TmoA04G023750 | 1377 | sens | | | 65 | figshare | Pep file |
| | | TetmoGNL2c (chr B4) TmoB04G024260 | 1377 | sens | | | 65 | figshare | Pep file |
| Fabales (Fabaceae) | Medicago trunculata | Medtr_LOC25489342 (chr 3) (Medtr3g068130) | 1474 | sens | 80 | 61 | | NCBI | GCA_0484 18095.1 |
| | | Medtr_LOC25489344 (chr 3) (Medtr3g068140) | 1473 | sens | 71 | 61 | | NCBI | GCA_0484 18095.1 |
| | | Medtr_LOC11438427 (chr 5) (Medtr5g080650) | 1465 | sens | 71 | 61 | | NCBI | GCA_0484 18095.1 |
| | | Medtr_LOC11437178 (chr 6) (Medtr6g086260) | 1384 | sens | | | 62 | NCBI | GCA_0484 18095.1 |
| Fabales (Fabaceae) | Phaseolus vulgaris | Phvul.001G235300 | 1473 | sens | 83 | 63 | | Phytozome 13 | V2.1 |
| | | Phvul.008G231400 | 1473 | sens | 72 | 61 | | Phytozome 13 | V2.1 |
| | | Phvul.001G235400 | 1464 | res | 70 | 59 | | Phytozome 13 | V2.1 |
| | | Phvul.004G161900 | 1377 | sens | | | 63 | Phytozome 13 | V2.1 |
| Fabales (Fabaceae) | Vigna angularis | Vigan_LOC108329051 (ViganGNOMa, chr 3) | 1473 | sens | 83 | 64 | | NCBI | GCA_0168 08095.1 |
| | | Vigan_LOC108329296 (ViganGNOMb, chr 3) | 1465 | res | 69 | 59 | | NCBI | GCA_0168 08095.1 |
| | | Vigan_108338084 (ViganGNOMc, chr 7) | 1493 | sens | 72 | 62 | | NCBI | GCA_0168 08095.1 |
| | | LOC108345258 (ViganGNL2, chr 10) | 1362 | sens | | | 61 | NCBI | GCA_0168 08095.1 |
| Fabales (Fabaceae) | Abrus precatorius | Abrpr_LOC113846820 AbrprGNOMb | 1471 | sens | 81 | 62 | | NCBI | GCA_0039 35025.1 |

| | | | | | | | | | |
|---|---|---|---|---|---|---|---|---|---|
| | | Abrpr_LOC113849046 AbrprGNOMa | 1465 | sens | 83 | 64 | | NCBI | GCA_0039 35025.1 |
| | | Abrpr_LOC113846103 AbrprGNOMc | 1473 | sens | 74 | 62 | | NCBI | GCA_0039 35025.1 |
| | | Abrpr_LOC11386679 5 | 1361 | sens | | | 63 | NCBI | GCA_0039 35025.1 |
| Fabales (Fabaceae) | Prosopis alba | LOC114732932 GNOMa (ctg 12957) | 1465 | res | 82 | 63 | | NCBI | GCA_0047 99145.2 |
| | | LOC114748431 GNOMb (ctg 5722) | 1449 | sens | 76 | 61 | | NCBI | GCA_0047 99145.2 |
| | | LOC114759906 GNL2a | 1365 | sens | | | 64 | NCBI | GCA_0047 99145.2 |
| Fabales (Fabaceae) | Cercis canadensis | CercaGNOMa (sc32) | 1469 | sens | 83 | 63 | | NCBI | GCA_0032 55065.1 |
| | | CercaGNOMb (sc32) | 1475 | res | 76 | 63 | | NCBI | GCA_0032 55065.1 |
| | | CercaGNL2 (sc515) | 1390 | sens | | | 65 | NCBI | GCA_0032 55065.1 |
| Fabales (Fabaceae) | Acacia acuminata | AcaacGNOMa tig00000237 | 1465 | res | 82 | 62 | | NCBI | GCA_0190 22655.1 |
| | | AcaacGNOMb tig00063063 | 1468 | sens | 75 | 61 | | NCBI | GCA_0190 22655.1 |
| | | AcaacGNL2 tig00000281 | 1362 | sens | | | 62 | NCBI | GCA_0190 22655.1 |
| | | EpefaGNL2 (ctg 3765000007) | 1354 | sens | | | 66 | NCBI | GCA_0190 22655.1 |
| Fabales (Fabaceae) | Inga leiocalycina | IngleGNOMa (chr6, Mb 16.3) | 1462 | res | 82 | 63 | | NCBI | GCA_9632 42795.1 |
| | | IngleGNOMb (chr7, Mb 0.2) | 1469 | sens | 75 | 61 | | NCBI | GCA_9632 42795.1 |
| | | IngleGNL2 (chr3, Mb 22.4) | 1367 | sens | | | 62 | NCBI | GCA_9632 42795.1 |
| Fabales (Quillajaceae) | Quillaja saponaria | QuisaGNOMa (chr 6) | 1466 | sens | 83 | 63 | | NCBI | GCA_0293 79385.1 |
| | (NCBI Mar 23) NEW | QuisaGNOMb (chr 8) | 1454 | sens | 83 | 63 | | NCBI | GCA_0293 79385.1 |

| | | | | | | | | | |
|---|---|---|---|---|---|---|---|---|---|
| | | 93% id. QuisaGNOMa | | | | | | | |
| | | QuisaGNL2 (chr 4) | 1388 | sens | | | 67 | NCBI | GCA_029379385.1 |
| Fabales (Polygalaceae) | Polygala tenuifolia | PolteGNOMa | 1462 | sens | 82 | 63 | | TCMPG | Pep file |
| | | PolteGNOMb | 1462 | res | 82 | 63 | | TCMPG | Pep file |
| | | PolteGNOMc | 1452 | sens | 80 | 61 | | TCMPG | Pep file |
| | | PolteGNL2 | 1385 | sens | | | 62 | TCMPG | Pep file |
| Rosales (Rosaceae) | Fragaria vesca | Frave_LOC101300393 (LG7, Mb 16.21) (mrna26324A) | 1471 | sens | 82 | 62 | | NCBI | GCA_000184155.1 |
| | | Frave_LOC101300670 (LG7, Mb 16.22) mrna26324B | 1467 | res | 75 | 62 | | NCBI | GCA_000184155.1 |
| | | Frave_LOC101305794 (LG7, Mb 11.98) (mrna07469) | 1444 | sens | 78 | 60 | | NCBI | GCA_000184155.1 |
| | | Frave_LOC101307629 (LG1) (mrna02092) | 1376 | sens | | | 65 | NCBI | GCA_000184155.1 |
| Rosales (Rosaceae) | Prunus persica | Prupe.2G239200 (= LOC18785443) | 1467 | sens | 83 | 63 | | NCBI | GCA_000346465.2 |
| | | Prupe.2G239100 (= LOC18784935) | 1473 | sens | 78 | 62 | | NCBI | GCA_000346465.2 |
| | | Prupe.7G228800 Prupe_LOC18770144 (chr G7) | 1384 | sens | | | 67 | NCBI | GCA_000346465.2 |
| Rosales (Rosaceae) | Rosa chinensis | Rosch_LOC112182803 | 1471 | sens | 83 | 62 | | NCBI | GCA_002994745.2 |
| | | Rosch_LOC112182802 | 1465 | sens | 77 | 62 | | NCBI | GCA_002994745.2 |

| | | | | | | | | | |
|---|---|---|---|---|---|---|---|---|---|
| | | Rosch_LOC1121908 48 | 1361 | sens | | | 67 | NCBI | GCA_00 2994745. 2 |
| Rosales (Rhamnaceae) | Ziziphus jujuba | Zizju_LOC107413409 (chr 3) | 1465 | sens | 84 | 63 | | NCBI | GCA_0317 55915.1 |
| | (almost tandem) | Zizju_LOC107413408 (chr 3) | 1469 | res | 83 | 63 | | NCBI | GCA_0317 55915.1 |
| | | Zizju_LOC107431830 (chr 12) | 1378 | sens | | | 66 | NCBI | GCA_0317 55915.1 |
| Rosales (Rhamnaceae) | Rhamnella rubrinervis | FNV43_RR24766 (LG11) | 1471 | res | 83 | 63 | | NCBI | GCA_0078 44105.2 |
| | | FNV43_RR24768 (LG11) | 1460 | sens | 83 | 63 | | NCBI | GCA_0078 44105.2 |
| | | RharuGNL2 (LG7) | 1380 | sens | | | 68 | NCBI | GCA_0078 44105.2 |
| Rosales (Elaeagnaceae | Elaeagnus angustifolia | EVM0003851 / ElaanGI Contig 86422 | 1453 | sens | 83 | 61 | | Figshare | Pep file |
| | | EVM0015105 / ElaanGI Contig 19186 | 1454 | sens | 67 | 65 | | Figshare | Pep file |
| | | EVM0028040 / ElaanGI Contig 150397 | 1453 | res | 64 | 63 | | Figshare | Pep file |
| | | EVM0021459 / ElaanGI Contig 94613 | 1453 | sens | 67 | 65 | | Figshare | Pep file |
| | | EVM0016313 / ElaanGNL2 | 1375 | sens | | | 66 | Figshare | Pep file |
| Rosales (Elaeagnaceae | Elaeagnus macrophylla | ElamaGNOMa (chr 13, Mb 2.3) | 1453 (1471) | sens | 83 | 62 | | NCBI | GCA_0293 78325.1 |
| | | ElamaGNOMb (chr 7, Mb 24.0) ElamaGNL1a | 1454 | sens | 66 | 65 | | NCBI | GCA_0293 78325.1 |
| | | ElamaGNOMc (chr 7, Mb 23.9) ElamaGNL1b | 1456 | sens | 67 | 65 | | NCBI | GCA_0293 78325.1 |
| | | ElamaGNOMd (chr 6, Mb 26.3) ElamaGNL1c | 1452 | res | 66 | 65 | | NCBI | GCA_0293 78325.1 |

| | | | | | | | | | |
|---|---|---|---|---|---|---|---|---|---|
| | | ElamaGNL2 (chr 2, Mb 1.3) ElamaGNL1c | (1374) | sens | | | 66 | NCBI | GCA_029378325.1 |
| Rosales (Elaeagnaceae | Hippophae gyantsensis | HipgyGNOMa (chr12, Mb 4.1) | 1454 | sens | 82 | 62 | | NCBI | GCA_030763125.1 |
| | | HipgyGNOMb (chr1, Mb 11.8) | 1452 | sens | 79 | 60 | | NCBI | GCA_030763125.1 |
| | | HipgyGNL1a (chr2, Mb 58.9) | 1455 | res | 66 | 66 | | NCBI | GCA_030763125.1 |
| | | HipgyGNL1b (chr1, Mb 81.4) aa432-1149 missing | | --- | | | | NCBI | GCA_030763125.1 |
| | | HipgyGNL2 (chr10, Mb 1.1) | 1376 | sens | | | 66 | NCBI | GCA_030763125.1 |
| Rosales (Elaeagnaceae | Hippophae tibetana | HiptiGNOMa (chr5, Mb 38.0) | 1454 | sens | 82 | 62 | | NCBI | GCA_037013495.1 |
| | | HiptiGNOMb (chr1, Mb 138.3) | 1452 | sens | 78 | 60 | | NCBI | GCA_037013495.1 |
| | | HiptiGNL1a (chr3, Mb 76.8) | 1459 (1454) | res | 66 | 65 | | NCBI | GCA_037013495.1 |
| | | HiptiGNL1b (chr1, Mb 4.9) | gaps | | | | | NCBI | GCA_037013495.1 |
| | | HiptiGNL2 (chr9, Mb 1.1) | 1376 | sens | | | 66 | NCBI | GCA_037013495.1 |
| Rosales (Ulmaceae) | Ulmus americana | UlmamGNOMa (sc14739) | 1457 | sens | 83 | 63 | | NCBI | GCA_010015005.3 |
| | | UlmamGNOMb (sc13178) | 1465 | res | 76 | 63 | | NCBI | GCA_010015005.3 |
| | | UlmamGNL2 (sc1203) | 1373 | sens | | | 67 | NCBI | GCA_010015005.3 |
| Rosales (Ulmaceae) | Ulmus parvifolia | UlmpaGNOMa (chr4, Mb 83.2) | 1466 | sens | 84 | 64 | | NCBI | GCA_041381145.1 |
| | | UlmpaGNOMb (chr4, Mb 83.2) | 1468 | res | 75 | 63 | | NCBI | GCA_041381145.1 |
| | | UlmpaGNL2 (chr7, Mb 31.9) | 1373 | sens | | | 67 | NCBI | GCA_041381145.1 |

| | | | | | | | | | |
|---|---|---|---|---|---|---|---|---|---|
| Rosales (Ulmaceae) | Ulmus glabra | UlmglGNOMa (chr2, Mb 52.5) | 1466 | sens | 85 | 64 | | NCBI | GCA_964106905.1 |
| | | UlmglGNOMb (chr2, Mb 52.6) | 1468 | res | 76 | 63 | | NCBI | GCA_964106905.1 |
| | | UlmglGNL2 (chr6, Mb 32.1) | 1373 | sens | | | 67 | NCBI | GCA_964106905.1 |
| Rosales (Cannabaceae) | Trema orientale | TorRG33x02_270070 (scf00318) | 1466 | sens | 84 | 63 | | NCBI | GCA_002914845.1 |
| | | TorRG33x02_079340 (scf00039) | 1456 | sens | 69 | 67 | | NCBI | GCA_002914845.1 |
| | | TorRG33x02_095630 (scf00050) | 1380 | sens | | | 66 | NCBI | GCA_002914845.1 |
| Rosales (Cannabaceae) | Parasponia andersonii | PanWU01x14_256170 | 1466 | sens | 84 | 63 | | NCBI | GCA_002914805.1 |
| | | PanWU01x14_321370 | 1456 | sens | 70 | 66 | | NCBI | GCA_002914805.1 |
| | | PanWU01x14_058240 | 1380 | sens | | | 66 | NCBI | GCA_002914805.1 |
| Rosales (Cannabaceae) | Cannabis sativa | LOC115707648 (chr 1) | 1464 | sens | 83 | 63 | | NCBI | GCA_029168945.1 |
| | | LOC115704658 (chr 1) | 1467 | res | 77 | 61 | | NCBI | GCA_029168945.1 |
| | | LOC115716528 (chr 5) | 1389 | | | | 65 | NCBI | GCA_029168945.1 |
| Rosales (Cannabaceae) | Humulus lupulus | HumluGNOMa (chr8, Mb 120.5) LOC133798386 | 1467 | sens | 84 | | | NCBI | GCA_963169125.1 |
| | | HumluGNOMb (chr8, Mb 120.5) LOC133798387 | 1454 | res | 78 | | | NCBI | GCA_963169125.1 |
| | | HumluGNL2 (chr 5) LOC133780431 | 1376 | | | | | NCBI | GCA_963169125.1 |
| Rosales (Moraceae) | Morus notabilis | Morno_LOC21408061 (sc139) | 1479 | sens | 83 | 63 | | NCBI | GCA_000414095.2 |
| | tandem | Morno_LOC21408060 (sc139) | 1465 | res | 78 | 62 | | NCBI | GCA_000414095.2 |

| | | | | | | | | | |
|---|---|---|---|---|---|---|---|---|---|
| | | Morno_LOC21403706 | 1373 | sens | | | 66 | NCBI | GCA_0004 14095.2 |
| Rosales (Moraceae) | Ficus erecta | FicerGNOMa hctg0032F_007.1 | (1465) | sens | 84 | 63 | | NCBI | GCA_0086 35985.1 |
| | | FicerGNOMb hctg0032F_007.1 | (1465) | res | 80 | 62 | | NCBI | GCA_0086 35985.1 |
| | | FicerGNL2 hctg0023F_012.1 | (1377) | sens | | | 60% | NCBI | GCA_0086 35985.1 |
| Rosales (Urticaceae) | Boehmeria nivea | BoeniGNOMa (chr 3, 6.5 Mb) | (1465) | sens | 85 | 63 | | NCBI | GCA_0181 32145.1 |
| | | BoeniGNOMb (chr 3, 6.5 Mb) | (1440) | res | 78 | 63 | | NCBI | GCA_0181 32145.1 |
| | | BoeniGNL2 (chr 8) | | | | | 64 | NCBI | GCA_0181 32145.1 |
| Rosales (Urticaceae) | Urtica urens | UrturGNOMa (chr9, Mb 21.7) | 1463 | sens | 81 | 62 | | NCBI | GCA_9582 96335.1 |
| | | UrturGNOMb (chr11, Mb 4.8) | 1444 | res | 70 | 60 | | NCBI | GCA_9582 96335.1 |
| | | UrturGNL2 (chr3, Mb 11.5) | 1356 | sens | | | 58 | NCBI | GCA_9582 96335.1 |
| Cucurbitales (Coriariaceae) | Coriaria nepalensis | CorneGNOMa (chr 1, Mb 56.1) | 1457 | sens | 82 | 61 | | NCBI | GCA_0271 86245.1 |
| | | CorneGNOMb (chr 5, Mb 1.0) | 1457 | sens | 81 | 61 | | NCBI | GCA_0271 86245.1 |
| | | CorneGNL1a (chr 2, Mb 2.9) | 1430 | sens | 67 | 65 | | NCBI | GCA_0271 86245.1 |
| | | CorneGNL1b (chr 16, Mb 6.9) | 1430 | res | 68 | 66 | | NCBI | GCA_0271 86245.1 |
| | | CorneGNL2 (chr 8, Mb 2.6) | 1380 | sens | | | 64 | NCBI | GCA_0271 86245.1 |
| Cucurbitales (Cucurbitaceae) | Cucumis sativus | Cucsa_LOC1012046 38 (chr 5) (Cucsa.165100) | 1469 | sens | 81 | 63 | | NCBI | GCA_0000 04075.3 |
| | | Cucsa_LOC1012079 38 (chr 3) (Cucsa.192630) | 1471 | res | 80 | 63 | | NCBI | GCA_0000 04075.3 |

| | | | | | | | | | |
|---|---|---|---|---|---|---|---|---|---|
| | | Cucsa_LOC101210509 (chr 2) (Cucsa.116300) | 1388 | sens | | | 65 | NCBI | GCA_000004075.3 |
| Cucurbitales (Cucurbitaceae) | Cucurbita maxima | Cucma_LOC111490452 | 1468 | sens | 81 | 63 | | NCBI | GCA_002738345.1 |
| | | Cucma_LOC111476959 | 1469 | res | 79 | 62 | | NCBI | GCA_002738345.1 |
| | | Cucma_LOC111465412 | 1381 | sens | | | 66 | NCBI | GCA_002738345.1 |
| Cucurbitales (Cucurbitaceae) | Momordica charantia | LOC111010039 MomchGNOMa (chr 9) | 1470 | sens | 81 | 63 | | NCBI | GCA_001995035.1 |
| | | LOC111025550 / MomchGNOMb (chr 2) | 1469 | sens | 80 | 62 | | NCBI | GCA_001995035.1 |
| | | LOC111024228 / MomchGNL2 (chr 8) | 1384 | sens | | | 66 | NCBI | GCA_001995035.1 |
| Cucurbitales (Cucurbitaceae) | Benincasa hispida | BenhiGNOM LOC120080467 (chr 6) | 1468 | sens | 81 | 63 | | NCBI | GCA_009727055.1 |
| | | BenhiGNL2 LOC120079606 (chr 6) | 1391 | sens | | | 65 | NCBI | GCA_009727055.1 |
| Cucurbitales (Datiscaceae) | Datisca glomerata | Datgl137S20405 (sc137) | 1470 | sens | 84 | 62 | | NCBI | GCA_003255025.1 |
| | | Datgl891S35907 (sc891) | 1450 | res | 69 | 66 | | NCBI | GCA_003255025.1 |
| | | Datgl574S11525 (sc574) | 1380 | sens | | | 65 | NCBI | GCA_003255025.1 |
| Cucurbitales (Begoniaceae) | Begonia fuchsioides | Begfu1501S08048 (sc1501) | 1469 | sens | 82 | 61 | | NCBI | GCA_003255005.1 |
| | | Begfu4464S35236 (sc4464) | 1469 | res | 81 | 61 | | NCBI | GCA_003255005.1 |
| | | Begfu1692S08615 (sc1692) | 1389 | sens | | | 66 | NCBI | GCA_003255005.1 |
| Cucurbitales (Begoniaceae) | Begonia peltatifolia | BegpeGNOM (chr 15) | 1469 | sens | 82 | 61 | | NCBI | GCA_022433055.1 |
| | | BegpeGNL2 (chr 2) | 1389 | sens | | | 66 | NCBI | GCA_022433055.1 |

| | | | | | | | | | |
|---|---|---|---|---|---|---|---|---|---|
| Fagales (Fagaceae) | Quercus lobata | Quelo_LOC115954940 (chr 8) | 1469 | sens | 81 | 63 | | NCBI | GCA_001633185.5 |
| | | Quelo_LOC115976920 (chr 2) | 1463 | res | 70 | 68 | | NCBI | GCA_001633185.5 |
| | | Quelo_LOC115981000 (chr 3) | 1424 | sens | | | 65 | NCBI | GCA_001633185.5 |
| Fagales (Fagaceae) | Quercus glauca | QueglGNOM (chr 8) | 1469 | sens | 82 | 63 | | NCBI | GCA_023736055.1 |
| | | QueglGNL1 (chr 2) | 1464 | res | 70 | 68 | | NCBI | GCA_023736055.1 |
| | | QueglGNL2 (chr 3) | 1394 | sens | | | 65 | NCBI | GCA_023736055.1 |
| Fagales (Fagaceae) | Fagus sylvatica | FagsyGNOM (chr 9) | 1469 | sens | 84 | 63 | | NCBI | GCA_964211995.1 |
| | | FagsyGNL1 (chr 8) | 1461 | res | 70 | 68 | | NCBI | GCA_964211995.1 |
| | | FagsyGNL2 (chr 12) | 1385 | sens | | | 68 | NCBI | GCA_964211995.1 |
| Fagales (Myricaceae) | Morella rubra | CJ030_MR3G014762 (chr 3) | 1468 | sens | 83 | 63 | | NCBI | GCA_003952965.2 |
| | | CJ030_MR2G025548 (chr 2) | 1461 | res | 68 | 67 | | NCBI | GCA_003952965.2 |
| | | CJ030_MR0G007998 (sc418) | 1375 | sens | | | 66 | NCBI | GCA_003952965.2 |
| Fagales (Juglandaceae) | Juglans regia | Jugre_LOC108991852 JugreGNOM (chr 7) | 1467 | sens | 82 | 62 | | NCBI | GCA_001411555.2 |
| | | Jugre_LOC109006351 JugreGNL1a (chr 1) | 1464 | sens | 69 | 68 | | NCBI | GCA_001411555.2 |
| | | Jugre_LOC108985951 JugreGNL1b (chr 10) | 1460 | res | 68 | 67 | | NCBI | GCA_001411555.2 |
| | | Jugre_LOC108992663 | 1394 | sens | | | 66 | NCBI | GCA_001411555.2 |
| Fagales (Casuarinaceae) | Casuarina equisetifolia | CaseqGNOM (sc309) | 1471 | sens | 83 | 63 | | NCBI | GCA_028551475.1 |
| | | CaseqGNL1 (sc190) | 1462 | res | 69 | 67 | | NCBI | GCA_028551475.1 |

| | | | | | | | | | |
|---|---|---|---|---|---|---|---|---|---|
| | | CaseqGNL2 (sc144) | 1418 | sens | | | 63 | NCBI | GCA_0285 51475.1 |
| Fagales (Betulaceae) | Betula pendula | Bpev01.c0365.g0008 (chr 5) (Bp_FXXK01000366) | 1471 | sens | 83 | 63 | | NCBI | GCA_9001 84695.1 |
| | | Bpev01.c0281.g0095 (chr 2) (Bp_FXXK01000282) | 1467 | res | 69 | 68 | | NCBI | GCA_9001 84695.1 |
| | | Bpev01.c1159.g0023 (chr 10) (Bp_FXXK01001160) | 1392 | sens | | | 68 | NCBI | GCA_9001 84695.1 |
| Fagales (Betulaceae) | Corylus heterophylla | CorheGNOM (chr 6) | 1471 | sens | 84 | 62 | | NCBI | GCA_0164 03345.1 |
| | | CorheGNL1 (chr 5) | 1462 | sens | 70 | 68 | | NCBI | GCA_0164 03345.1 |
| | | CorheGNL2 (chr 11) | 1369 | sens | | | 66 | NCBI | GCA_0164 03345.1 |
| Fagales (Betulaceae) | Alnus glutinosa | AlnglGNOM (sc62290) | 1469 | sens | 83 | 63 | | NCBI | GCA_9589 79055.1 |
| | | AlnglGNL1 (sc33409) | 1465 | res | 70 | 68 | | NCBI | GCA_9589 79055.1 |
| | | AlnglGNL2 (sc17582) | | | | | | NCBI | GCA_9589 79055.1 |
| Fagales (Betulaceae) | Carpinus fangiana | Carfa_FH972_001430 (chr 1) | 1471 | sens | 83 | 62 | | NCBI | GCA_0069 37295.1 |
| | | Carfa_FH972_007592 (chr 3) | 1461 | sens | 70 | 68 | | NCBI | GCA_0069 37295.1 |
| | | Carfa_FH972_003276 (chr 1) | 1392 | sens | | | 68 | NCBI | GCA_0069 37295.1 |
| Fagales (Betulaceae) | Corylus avellana | CoravGNOM (chr ca7) | (1470) | sens | 84 | 61 | | NCBI | GCA_9010 00735.2 |
| | | CoravGNL1 (chr ca5) | (1460) | sens | 70 | 67 | | NCBI | GCA_9010 00735.2 |
| | | CoravGNL2 (chr ca11) | (1390) | sens | | | 65 | NCBI | GCA_9010 00735.2 |
| Celastrales (Celastraceae) | Tripterygium wilfordii | TriwiGNOM (chr 1, kb 138) | (1462) | sens | 84 | 63 | | NCBI | GCA_0134 01445.1 |

| | | | | | | | | | |
|---|---|---|---|---|---|---|---|---|---|
| | | LOC119991987, chr22 | | | | | | | |
| | (new genome NCBI Feb 21) | TriwiGNL1a (chr 8, Mb 12.5) LOC120016528, chr15 | (1463) | sens | 68 | 66 | | NCBI | GCA_013401445.1 |
| | (2 TriwiGNL1) | TriwiGNL1b (chr 6, Mb 5.6) LOC119996657, chr 4; LOC119996659, chr 4 | (1463) | res | 68 | 65 | | NCBI | GCA_013401445.1 |
| | | TriwiGNL2 (chr 6, Mb 3.5) | (1380) | sens | | | 67 | NCBI | GCA_013401445.1 |
| Celastrales (Celastraceae) | Euonymus europaeus | EuoeuGNOM (chr12, Mb 3.8) | 1463 | sens | 82 | 63 | | NCBI | GCA_963580455.1 |
| | NCBI Nov 2023 | EuoeuGNL1a (chr11, Mb 4.8) | 1454 | sens | 68 | 66 | | NCBI | GCA_963580455.1 |
| | | EuoeuGNL1b (chr5, Mb 27.2) | 1460 | res | 67 | 66 | | NCBI | GCA_963580455.1 |
| | | EuoeuGNL2 (chr4, Mb 22.1) | 1376 | sens | | | 68 | NCBI | GCA_963580455.1 |
| Malpighiales (Malpighiaceae) | Malpighia emarginata | MalemGNOM NRA309ch09g29826 (chr9) | 1464 | sens | 83 | 63 | | plantgarden | Pep file |
| | | MalemGNL1 NRA309ch09g31492 (chr9) | 1442 | sens | 67 | 66 | | plantgarden | Pep file |
| | | MalemGNL2 NRA309ch08g27243 (chr8) | 1372 | sens | | | 65 | plantgarden | Pep file |
| Malpighiales (Linaceae) | Linum usitatissimum | Lus10026627 / LinusGNOMa (chr1, Mb 0.7) | 1446 | sens | 80 | 61 | | NCBI | GCA_048169125.1 |
| | | Lus10030432 / LinusGNOMb (chr7, Mb 9.6) | 1449 | sens | 80 | 60 | | NCBI | GCA_048169125.1 |
| | | Lus10037140 / LinusGNOMd (chr13, Mb 9.1) | 1407 | sens | 77 | 62 | | NCBI | GCA_048169125.1 |
| | | Lus10036789 / LinusGNOMc (chr2, Mb 19.7) | 1403 | sens | 79 | 64 | | NCBI | GCA_048169125.1 |

| | | | | | | | | | |
|---|---|---|---|---|---|---|---|---|---|
| | | Lus10021771 (LinusGNL2a, chr1, Mb 17.8?) | 1372 | sens | | | 61 | NCBI | GCA_0481 69125.1 |
| | | Lus10034615 (LinusGNL2b, chr7, Mb 13.9?) | 1387 | sens | | | 62 | NCBI | GCA_0481 69125.1 |
| Malpighiales (Linaceae) | Linum bienne | LinbiGNOMa (sc 960) | 1446 | sens | 80 | 61 | | NCBI | GCA_0106 65285.1 |
| | | LinbiGNOMb (sc 62) | 1446 | sens | 80 | 61 | | NCBI | GCA_0106 65285.1 |
| | | LinbiGNOMc (sc 614) | 1375 | sens | 79 | 63 | | NCBI | GCA_0106 65285.1 |
| | | LinbiGNL2a (sc 1) | 1387 | sens | | | 62 | NCBI | GCA_0106 65285.1 |
| | | LinbiGNL2b (sc 82) | 1387 | sens | | | 62 | NCBI | GCA_0106 65285.1 |
| Malpighiales (Linaceae) | Linum tenue | LinteGNOMa (LG10, Mb 16.0) LITE_LOCUS6964 | 1459 | sens | 79 | 61 | | NCBI | GCA_9461 22785.1 |
| | | LinteGNOMb (LG10, Mb 35.2) LITE_LOCUS8312 | 1459 | sens | 80 | 62 | | NCBI | GCA_9461 22785.1 |
| | | LinteGNOMc (LG6) LITE_LOCUS35573 | 1459 | sens | 80 | 62 | | NCBI | GCA_9461 22785.1 |
| | | LinteGNL1a (LG10, Mb 39.0) LITE_LOCUS8722 | 1453 | sens | 65 | 64 | | NCBI | GCA_9461 22785.1 |
| | | LinteGNL1b (LG9, Mb 34.6) LITE_LOCUS50160 | 1459 | sens | 64 | 63 | | NCBI | GCA_9461 22785.1 |
| | | LinteGNL2a (LG 10, Mb 8.0) LITE_LOCUS6516 | 1379 | sens | | | 62 | NCBI | GCA_9461 22785.1 |
| | | LinteGNL2b (LG9, Mb 55.9) LITE_LOCUS51829 | 1381 | sens | | | 63 | NCBI | GCA_9461 22785.1 |
| | | LinteGNL2c (LG7) LITE_LOCUS42764 | 1381 | sens | | | 63 | NCBI | GCA_9461 22785.1 |

| | | | | | | | | | |
|---|---|---|---|---|---|---|---|---|---|
| Malpighiales (Hypericacae) | Hypericum perforatum | HyppeGNOM (ctg 60781) | (1465) | sens | 83 | 60 | | NCBI | GCA_041475425.1 |
| | | HyppeGNL1 (ctg 62069) | (1440) | sens | 66 | 65 | | NCBI | GCA_041475425.1 |
| | | (no GNL2?) | | | | | | NCBI | GCA_041475425.1 |
| Malpighiales (Hypericacae) | Hypericum androsaemum | HypanGNOM (chr16, Mb 6.7) | 1471 | sens | 82 | 63 | | NCBI | GCA_963995505.1 |
| | | HypanGNL1a (chr4, Mb 3.4) | 1434 | sens | 65 | 64 | | NCBI | GCA_963995505.1 |
| | | HypanGNL1b (chr12, Mb 14.7) | 1434 | res | 65 | 64 | | NCBI | GCA_963995505.1 |
| | | HypanGNL2 (chr11, Mb 13.1) | 1400 | sens | | | 62 | NCBI | GCA_963995505.1 |
| Malpighiales (Calophyllaceae) | Mesua ferrea | MesfeGNOMa sc MesFer2642 | (1463) | sens | 84 | 64 | | NCBI | GCA_904814375.1 |
| | | MesfeGNOMb sc MesFer2750 | (1450) | sens | 86 | 62 | | NCBI | GCA_904814375.1 |
| | | MesfeGNL1a sc MesFer2408 | (1444) | sens | 72 | 66 | | NCBI | GCA_904814375.1 |
| | | MesfeGNL1b sc MesFer6600 ( | (1431) | sens | 69 | 65 | | NCBI | GCA_904814375.1 |
| | | MesfeGNL2 sc MesFer1296 | (1400) | sens | | | 64 | NCBI | GCA_904814375.1 |
| Malpighiales (Podostemaceae) | Cladopus chinensis | ClachGNOMa GWHPACFD001936 / chr 10 | 1415 | sens | 75 | 58 | | CNGB | GWH Pep file |
| | | ClachGNOMb GWHPACFD019647 / chr 5 | 1415 | sens | 75 | 58 | | CNGB | GWH Pep file |
| | | ClachGNL1 GWHPACFD003699 / chr 12 | 1375 | sens | 59 | 59 | | CNGB | GWH Pep file |
| | | ClachGNL2 GWHPACFD025206 (tig00001399) | 1320 | sens | | | 56 | CNGB | GWH Pep file |

| | | | | | | | | | |
|---|---|---|---|---|---|---|---|---|---|
| Malpighiales (Rhizophoraceae) | Kandelia obovata | KanobGNOMa (chr15) GWHPACBH002465 | (1455) | sens | 82 | 63 | | NCBI | GCA_0214 64305.1 |
| | | KanobGNOMb (chr15) GWHPACBH002465 | (1455) | res | 75 | 62 | | NCBI | GCA_0214 64305.1 |
| | | KanobGNL2 (chr 3) | (1376) | sens | | | 65 | NCBI | GCA_0214 64305.1 |
| Malpighiales (Rhizophoraceae) | Rhizophora apiculata | RhiapGNOMa (sc34) | (1465) | sens | 83 | 62 | | NCBI | GCA_0378 32385.1 |
| | | RhiapGNOMb (sc34) | (1460) | res | 75 | 61 | | NCBI | GCA_0378 32385.1 |
| | | RhiapGNL2 (sc6, 1.06Mb) | (1385) | sens | | | 65 | NCBI | GCA_0378 32385.1 |
| Malpighiales (Rhizophoraceae) | Bruguiera parviflora | BrupaGNOMa (chr 14) | (1470) | sens | 83 | 62 | | NCBI | GCA_0198 04595.1 |
| | | BrupaGNOMb (chr 14) | (1470) | sens | 76 | 60 | | NCBI | GCA_0198 04595.1 |
| | | BrupaGNL2 (chr 2) | (1390) | sens | | | 65 | NCBI | GCA_0198 04595.1 |
| Malpighiales (Erythroxylaceae) | Erythroxylum havanense | EryhaGNOMa (F:2324357 3) | 1470 | sens | 82 | 63 | | NCBI | GCA_0247 52395.1 |
| | | EryhaGNOMb (F:2324357 1) (Mb 1.68) | 1471 | sens | 82 | 63 | | NCBI | GCA_0247 52395.1 |
| | | EryhaGNOMc (F:2324357 1) (Mb 1.69) | 1443 | sens | 69 | 56 | | NCBI | GCA_0247 52395.1 |
| | | EryhaGNL1 (F:2324357 1) (Mb 27.8) | 1435 | sens | 66 | 67 | | NCBI | GCA_0247 52395.1 |
| | | EryhaGNL2 (F:2324357 8) | 1379 | sens | | | 67 | NCBI | GCA_0247 52395.1 |
| Malpighiales (Erythroxylaceae) | Erythroxylum novogranatense | ErynoGNOMa (LG07) | 1469 | sens | 81 | 63 | | NCBI | GCA_0298 91385.1 |
| | | ErynoGNOMb (LG01, Mb 2.3) | 1470 | sens | 82 | 63 | | NCBI | GCA_0298 91385.1 |
| | | ErynoGNOMc (LG01, Mb 2.2) | 1434 | sens | 69 | 57 | | NCBI | GCA_0298 91385.1 |

| | | | | | | | | | |
|---|---|---|---|---|---|---|---|---|---|
| | | ErynoGNL1 (LG01, Mb 71.5) | 1436 | sens | 66 | 66 | | NCBI | GCA_0298 91385.1 |
| | | ErynoGNL2 (LG11) | 1379 | sens | | | 67 | NCBI | GCA_0298 91385.1 |
| Malpighiales (Phyllanthaceae) | Flueggea virosa | FluviGNOM (chrA8, Mb 3.9) | 1467 | sens | 84 | 63 | | NCBI | GCA_0387 89575.1 |
| | | FluviGNL1 (chrA12, Mb 11.1) | 1458 | res | 66 | 66 | | NCBI | GCA_0387 89575.1 |
| | | FluviGNL2 (chrA5, Mb 11.6) | 1376 | sens | | | 68 | NCBI | GCA_0387 89575.1 |
| Malpighiales (Phyllanthaceae) | Sauropus androgynus | SauanGNOMa (LG12) San12G007300 | 1469 | sens | 82 | 63 | | CNGB | GWH Pep file |
| | | SauanGNOMb (LG25) San25G003480 | 1459 | sens | 80 | 61 | | CNGB | GWH Pep file |
| | | SauanGNL2a (LG18) San18G004080 | 1364 | sens | | | 67 | CNGB | GWH Pep file |
| | | SauanGNL2b (LG19) San19G000820 | 1364 | sens | | | 67 | CNGB | GWH Pep file |
| Malpighiales (Euphorbiaceae) | Ricinus communis | Ricco_LOC8281853 (chr 1) (scf_1106159297436) (Rc_29813.m001539) | 1470 | sens | 84 | 64 | | NCBI | GCA_0195 78655.1 |
| | | Ricco_LOC8283443 (chr 5) (scf_1106159291216) (Rc_29635.m000459) | 1450 | res | 68 | 68 | | NCBI | GCA_0195 78655.1 |
| | | Ricco_LOC107261358 (chr 1) (scf_1106159298012) (Rc_29751.m001855) | 1377 | sens | | | 67 | NCBI | GCA_0195 78655.1 |
| Malpighiales (Euphorbiaceae) | Manihot esculenta | Manes.01G032800 Manes_LOC1106252 74 (LG1) | 1469 | sens | 85 | 62 | | NCBI | GCA_0016 59605.2 |
| | | Manes.01G032600 Manes_LOC1106252 43 (LG1) | 1469 | sens | 83 | 62 | | NCBI | GCA_0016 59605.2 |
| | | Manes.02G049000 | 1435 | res | 66 | 66 | | NCBI | GCA_0016 59605.2 |

| | | | | | | | | | |
|---|---|---|---|---|---|---|---|---|---|
| | | Manes_LOC110609669 (chr 2) | | | | | | | |
| | | Manes.03G019000 Manes_LOC110610972 (chr 3) | 1381 | sens | | | 67 | NCBI | GCA_001659605.2 |
| Malpighiales (Euphorbiaceae) | Jatropha curcas | Jatcu_LOC105630489 (sc624) JatcuGNOMa | 1466 | sens | 85 | 63 | | NCBI | GCA_014843425.1 |
| | | Jatcu_LOC105633077 (sc355) JatcuGNOMb | 1466 | sens | 78 | 62 | | NCBI | GCA_014843425.1 |
| | | Jatcu_LOC105638327 (sc202) | 1377 | sens | | | 67 | NCBI | GCA_014843425.1 |
| Malpighiales (Euphorbiaceae) | Hevea brasiliensis | HevbrGNOM (chr9, Mb 95.5) LOC110645704 | 1470 | sens | | | | NCBI | GCA_030052815.1 |
| | | HevbrGNL1a (chr9, Mb 23.5) LOC110663786 | 1460 | res | | | | NCBI | GCA_030052815.1 |
| | | HevbrGNL1b (chr16) LOC110640282 (pseudo?) | 1461 | res | | | | NCBI | GCA_030052815.1 |
| | | HevbrGNL2a (chr4) LOC110653973 | 1379 | sens | | | | NCBI | GCA_030052815.1 |
| | | HevbrGNL2b (chr11) LOC110637295 (pseudo?) | (1360) | sens | | | | NCBI | GCA_030052815.1 |
| Malpighiales (Euphorbiaceae) | Vernicia fordii | VerfoGNOM GWHPAAEU054407 | 1470 | sens | 85 | 63 | | CNGB | GWH Pep file |
| | | VerfoGNL1 GWHPAAEU053260 | 1461 | res | 67 | 67 | | CNGB | GWH Pep file |
| | | VerfoGNL2 GWHPAAEU035762 | 1377 | sens | | | 67 | CNGB | GWH Pep file |
| Malpighiales (Euphorbiaceae) | Euphorbia peplus | EuppeGNOM (chr 2) | 1468 | sens | 82 | 62 | | NCBI | GCA_028411795.1 |
| | | EuppeGNL1 (chr 3) | 1460 | sens | 67 | 66 | | NCBI | GCA_028411795.1 |
| | | EuppeGNL2 (chr 8) | 1404 | sens | | | 63 | NCBI | GCA_028411795.1 |

| | | | | | | | | | |
|---|---|---|---|---|---|---|---|---|---|
| Malpighiales (Salicaceae) | Populus trichocarpa | Potri.017G078900.1 (POPTR_0017s11500) (Poptr_LOC18107297) (chr17) | 1470 | sens | 84 | 63 | | NCBI | GCA_000002775.4 |
| | | Potri.017G078800.1 (POPTR_0017s11490) (Poptr_LOC18107296) (chr17) | 1464 | res | 77 | 60 | | NCBI | GCA_000002775.4 |
| | | Potri.018G075300.1 (POPTR_0018s06100) (Poptr_LOC7458753) (chr18) | 1375 | sens | | | 64 | NCBI | GCA_000002775.4 |
| Malpighiales (Salicaceae) | Salix brachista | DKX38_026992 (chr 17 | 1470 | sens | 84 | 63 | | NCBI | GCA_049639585.1 |
| | | DKX38_026991 | 1458 | res | 77 | 61 | | NCBI | GCA_049639585.1 |
| | | DKX38_028175 | 1374 | sens | | | 63 | NCBI | GCA_049639585.1 |
| Malpighiales (Salicaceae) | Dovyalis caffra | DovcaGNOMa (ctg1120, Mb 2.3) DCAF_LOCUS667 | 1468 | sens | 84 | 63 | | NCBI | GCA_963924115.1 |
| | | DovcaGNOMb (ctg1120, Mb 2.4) DCAF_LOCUS668 | 1444 | res | 78 | 59 | | NCBI | GCA_963924115.1 |
| | | DovcaGNL2 (ctg_602, Mb 2.7) DCAF_LOCUS10142 | 1380 | sens | | | 65 | NCBI | GCA_963924115.1 |
| Malpighiales (Violaceae) | Viola odorata | ViodGNOMa (chr4, Mb 2.2) | 1454 | sens | 82 | 63 | | NCBI | GCA_963691705.1 |
| | | ViodGNOMb (chr4, Mb 67.1) | 1454 | res | 82 | 63 | | NCBI | GCA_963691705.1 |
| | | ViodGNOMc (chr4, Mb 67.1) | 1446 | sens | 72 | 58 | | NCBI | GCA_963691705.1 |
| | | ViodGNOMd (chr4, Mb 2.2) | 1427 | sens | 72 | 57 | | NCBI | GCA_963691705.1 |
| | | ViodGNL2a (chr8, Mb 11.2) | 1375 | sens | | | 65 | NCBI | GCA_963691705.1 |
| | | ViodGNL2b (chr4, Mb 10.8) | 1376 | sens | | | 65 | NCBI | GCA_963691705.1 |

| | | | | | | | | | |
|---|---|---|---|---|---|---|---|---|---|
| Malpighiales (Passifloraceae) | Passiflora edulis | PasedGNOMa ZX.08G0021780 | 1467 | sens | 82 | 62 | | NCBI | GCA_0021 56105.1 |
| | | PasedGNOMb ZX.08G0021710 | 1109 | sens | 70 | 55 | | NCBI | GCA_0021 56105.1 |
| | | PasedGNL1 ZX.01G0007300 | 1459 | res | 65 | 66 | | NCBI | GCA_0021 56105.1 |
| | | PasedGNL2 ZX.04G0001150 | 1372 | sens | | | 62 | NCBI | GCA_0021 56105.1 |
| Malpighiales (Passifloraceae) | Passiflora organensis | PasorGNOMa (sc1, Mb 1.8) | 1467 | sens | 82 | 63 | | NCBI | GCA_0197 39055.1 |
| | | PasorGNOMb (sc2, Mb 16.6) | 1468 | sens | 81 | 61 | | NCBI | GCA_0197 39055.1 |
| | | PasorGNL1 (sc1, Mb 2.1) | 1460 | res | 66 | 65 | | NCBI | GCA_0197 39055.1 |
| | | PasorGNL2 (sc1, Mb 9.4) | 1373 | sens | | | 62 | NCBI | GCA_0197 39055.1 |
| Malpighiales (Turneraceae) | Turnera subulata | Tsubulata_008874 TursuGNOM | 1463 | sens | 85 | 63 | | NCBI | GCA_0283 86065.3 |
| | | Tsubulata_010547 TursuGNL1 | 1456 | sens | 67 | 66 | | NCBI | GCA_0283 86065.3 |
| | | Tsubulata_006009 TursuGNL2 | 1370 | sens | | | n.a. | NCBI | GCA_0283 86065.3 |
| Oxalidales (Oxalidaceae) | Averrhoa carambola | GWHPABKE019952 (chr 9) | 1453 | sens | 83 | 62 | | CNGB | GWH Pep file |
| | | GWHPABKE019955 (chr 9) | 1443 | sens | 77 | 61 | | CNGB | GWH Pep file |
| | | GWHPABKE014651 (chr 7) | 1379 | sens | | | 69 | CNGB | GWH Pep file |
| Oxalidales (Oxalidaceae) | Oxalis articulata | OxaarGNOMa (chr3, Mb 39.1) | 1442 | sens | 84 | 63 | | NCBI | GCA_0491 90865.1 |
| | | OxaarGNOMb (chr5, Mb 45.6) | 1428 | sens | 72 | 61 | | NCBI | GCA_0491 90865.1 |
| | | OxaarGNL2 (chr1, Mb 51.3) | 1373 | sens | | | 67 | NCBI | GCA_0491 90865.1 |
| Oxalidales (Cephalotaceae) | Cephalotus follicularis | CFOL_v3_24675 (sc2355) | 1454 | sens | 81 | 63 | | NCBI | GCA_0019 72305.1 |

| | | | | | | | | | |
|---|---|---|---|---|---|---|---|---|---|
| | | CFOL_v3_21055 (sc1649) | 1411 | res | 66 | 65 | | NCBI | GCA_0019 72305.1 |
| | | CFOL_v3_22974 (sc1985) | 1385 | sens | | | 67 | NCBI | GCA_0019 72305.1 |
| Oxalidales (Elaeocarpaceae) | Aristotelia chilensis | ArichGNOM | 1452 | sens | 83 | 63 | | NCBI | GCA_0089 21755.1 |
| | | ArichGNL1 | (1440) | res | 70 | 70 | | NCBI | GCA_0089 21755.1 |
| | | ArichGNL2 | (1376) | sens | | | 69 | NCBI | GCA_0089 21755.1 |
| Sapindales (Nitrariaceae) | Nitraria sibirica | NitsiGNOM (LG 03) LG03.303 | 1467 | sens | 82 | 63 | | Figshare | Pep file |
| | | NitsiGNL1 (LG 02) LG02.231 | 1457 | sens | 68 | 67 | | Figshare | Pep file |
| | | NitsiGNL2 (LG 07) LG07.1058 | 1378 | sens | | | 67 | Figshare | Pep file |
| Sapindales (Sapindaceae) | Acer yangbiense | EZV62_017872 (chr 8) | 1467 | sens | 84 | 64 | | NCBI | GCA_0080 09225.1 |
| | | EZV62_015425 (chr 7) | 1465 | res | 69 | 68 | | NCBI | GCA_0080 09225.1 |
| | | EZV62_012732 | 1387 | sens | | | 69 | NCBI | GCA_0080 09225.1 |
| Sapindales (Sapindaceae) | Xanthoceras sorbifolium | JRO89_XS05G00374 00 XansoGNOM (chr 5) | 1472 | sens | 84 | 64 | | NCBI | GCA_0493 50635.1 |
| | | JRO89_XS12G02339 00 XansoGNL1 (chr 12) | 1463 | res | 69 | 68 | | NCBI | GCA_0493 50635.1 |
| | | JRO89_XS04G02362 00 XansoGNL2 (chr4) | 1381 | sens | | | 69 | NCBI | GCA_0493 50635.1 |
| Sapindales (Sapindaceae) | Dimocarpus longan | DimloGNOM (chr 10) | (1471) | sens | 84 | 63 | | NCBI | GCA_0229 84855.1 |
| | | DimloGNL1 (chr 14) | (1461) | res | 69 | 66 | | NCBI | GCA_0229 84855.1 |
| | | DimloGNL2 (chr 1) | (1386) | sens | | | 68 | NCBI | GCA_0229 84855.1 |

| | | | | | | | | | |
|---|---|---|---|---|---|---|---|---|---|
| Sapindales (Sapindaceae) | Sapindus mukorossi | GWHGBECP015250 (chr 8) SapmuGNOM | 1470 | sens | 84 | 64 | | CNGB | GWH Pep file |
| | | GWHGBECP023790 (chr 13) SapmuGNL1 | 1460 | res | 66 | 66 | | CNGB | GWH Pep file |
| | | GWHGBECP024656 (chr 14) SapmuGNL2 | 1387 | sens | | | 69 | CNGB | GWH Pep file |
| Sapindales (Anacardiaceae) | Anacardium occidentale | Anaoc.0003s0363 / AnaocGNOM (sc14) | 1464 | sens | 84 | 64 | | NCBI | GCA_026122585.1 |
| | | Anaoc.0002s0742 ( | (1365) | sens | 82 | 63 | | NCBI | GCA_026122585.1 |
| | | Anaoc.0018s0976 AnaocGNL1a (sc206) | 1456 | sens | 70 | 66 | | NCBI | GCA_026122585.1 |
| | | Anaoc.0010s1629 AnaocGNL1b (sc218) | 1461 | res | 69 | 67 | | NCBI | GCA_026122585.1 |
| | | Anaoc.0019s0415 | 1380 | sens | | | 66 | NCBI | GCA_026122585.1 |
| Sapindales (Anacardiaceae) | Pistacia vera | Pisve_LOC116120515 (chr 1)/ctg94, Mb 1.4 | 1464 | sens | 84 | 65 | | NCBI | GCA_008641045.1 |
| | | Pisve_LOC116112061 (chr 4, Mb 28.6) / ctg656, Mb 0.1 | 1457 | res | 71 | 69 | | NCBI | GCA_008641045.1 |
| | | PisveGNL2 Pisve_LOC116127888 Ctg196, Mb 0.8 | 1382 | sens | | | 68 | NCBI | GCA_008641045.1 |
| Sapindales (Anacardiaceae) | Mangifera indica | ManinGNOM (chr 9) / (chr 4) | (1452) | sens | 84 | 62 | | NCBI | GCA_011075055.1 |
| | | ManinGNL1a (chr 20) / (chr15) | (1460) | sens | 69 | 67 | | NCBI | GCA_011075055.1 |
| | | ManinGNL1b (chr 1) / (chr 9) | (1460) | res | 69 | 67 | | NCBI | GCA_011075055.1 |
| | | ManinGNL2 (chr 7) / (chr 17) | (1370) | sens | | | 64 | NCBI | GCA_011075055.1 |
| Sapindales (Anacardiaceae) | Toxicodendron radicans | ToxraGNOM (ctg 370) | 1464 | sens | 84 | 65 | | NCBI | GCA_009867345.2 |
| | | ToxraGNL1 (ctg 723) | 1456 | res | 71 | 69 | | NCBI | GCA_009867345.2 |

| | | | | | | | | | |
|---|---|---|---|---|---|---|---|---|---|
| | | ToxraGNL2 (ctg 597) | 1386 | sens | | | 68 | NCBI | GCA_009867345.2 |
| Sapindales (Burseraceae) | Boswellia sacra | BossaGNOM (scf7180000069358) | 1466 | sens | 84 | 64 | | NCBI | GCA_013180625.2 |
| | | BossaGNL1 (scf7180000069383) | 1441 | res | 70 | 69 | | NCBI | GCA_013180625.2 |
| | | BossaGNL2 (scf7180000049982) | 1380 | sens | | | 69 | NCBI | GCA_013180625.2 |
| Sapindales (Burseraceae) | Bursera cuneata | BurcuGNOM (sc1260) | 1466 | sens | 85 | 64 | | NCBI | GCA_026212285.1 |
| | | BurcuGNL1 (sc4862) | 1443 | res | 69 | 69 | | NCBI | GCA_026212285.1 |
| | | BurcuGNL2 (sc31078) | 1380 | sens | | | 69 | NCBI | GCA_026212285.1 |
| Sapindales (Burseraceae) | Bursera bipinnata | BurbiGNOM (sc41007) | 1466 | sens | 84 | 64 | | NCBI | GCA_026225755.1 |
| | | BurbiGNL1 (sc44852) | 1444 | res | 69 | 69 | | NCBI | GCA_026225755.1 |
| | | BurbiGNL2 (sc874) | | | | | 69 | NCBI | GCA_026225755.1 |
| Sapindales (Burseraceae) | Bursera palmeri | BurpaGNOM (sc29972) | 1465 | sens | 85 | 64 | | NCBI | GCA_026225775.1 |
| | | BurpaGNL1 (sc853) | 1444 | res | 69 | 69 | | NCBI | GCA_026225775.1 |
| | | BurpaGNL2 (sc25316) | 1380 | sens | | | 69 | NCBI | GCA_026225775.1 |
| Sapindales (Simaroubaceae) | Ailanthus altissimus | AilalGNOMa (chr 10) | 1468 | sens | 84 | 63 | | NCBI | GCA_046736235.1 |
| | | AilalGNOMb (chr 3) | 1469 | sens | 84 | 63 | | NCBI | GCA_046736235.1 |
| | | AilalGNL1a (chr 15) | 1467 | sens | 70 | 68 | | NCBI | GCA_046736235.1 |
| | | AilalGNL1b (chr 5) | 1463 | res | 69 | 67 | | NCBI | GCA_046736235.1 |
| | | AilalGNL2 (chr 4) | 1382 | sens | | | 69 | NCBI | GCA_046736235.1 |

| | | | | | | | | | |
|---|---|---|---|---|---|---|---|---|---|
| Sapindales (Meliaceae) | Azadirachta indica NEW | AzainGNOM (chr 4) | 1469 | sens | 85 | 64 | | NCBI | GCA_022749755.1 |
| | | AzainGNL1 (chr 9) | 1462 | sens | 69 | 68 | | NCBI | GCA_022749755.1 |
| | | AzainGNL2 (chr 7) | 1383 | sens | | | 70 | NCBI | GCA_022749755.1 |
| Sapindales (Meliaceae) | Xylocarpus granatum | XylgrGNOM (ctg 2266) | (1470) | sens | 91 | 62 | | NCBI | GCA_019650275.1 |
| | | XylgrGNL1 (ctg 105) | (1440) | res | 69 | 66 | | NCBI | GCA_019650275.1 |
| | | XylgrGNL2 (ctg 3646) | (1381) | sens | | | 69 | NCBI | GCA_019650275.1 |
| Sapindales (Rutaceae) | Citrus sinensis | Citsi_LOC102627469 (chr 7) | 1469 | sens | 84 | 63 | | NCBI | GCA_022201045.1 |
| | | Citsi_LOC102624619 (chr 6) | 1453 | res | 70 | 68 | | NCBI | GCA_022201045.1 |
| | | Citsi_LOC102625158 (chr 4) | 1380 | sens | | | 69 | NCBI | GCA_022201045.1 |
| Sapindales (Rutaceae) | Atalantia buxifolia | AtabuGNOM (sc9317) | 1469 | sens | 84 | 63 | | NCBI | GCA_030444775.1 |
| | | AtabuGNL1 (sc12963) | 1451 | res | 70 | 68 | | NCBI | GCA_030444775.1 |
| | | AtabuGNL2 (sc8059) | (1350) | sens | | | 67 | NCBI | GCA_030444775.1 |
| Sapindales (Rutaceae) | Murraya koenigii | MurkoGNOM (chr 4) | 1469 | sens | 84 | 63 | | NCBI | GCA_025617385.1 |
| | | MurkoGNL1 (chr 9) | 1468 | sens | 70 | 68 | | NCBI | GCA_025617385.1 |
| | | MurkoGNL2 (chr 7) | 1380 | sens | | | 69 | NCBI | GCA_025617385.1 |
| Malvales (Thymelaeaceae) | Aquilaria sinensis | AqusiGNOM (sc92.109) | 1468 | sens | 85 | 64 | | NCBI | GCA_005392925.1 |
| | | AqusiGNL1 (sc3.680) | 1473 | res | 65 | 66 | | NCBI | GCA_005392925.1 |
| | | AqusiGNL2 (sc277.24) | 1374 | sens | | | 66 | NCBI | GCA_005392925.1 |

| | | | | | | | | | |
|---|---|---|---|---|---|---|---|---|---|
| Malvales (Thymelaeaceae) | Stellera chamaejasme | StechGNOM SchG030265 (chr 7, Mb 2.4) | 1464 | sens | 83 | 64 | | NCBI | GCA_0245 86325.1 |
| | | StechGNL1 SchG027613 (chr 1, Mb 41.2) | 1454 | res | 65 | 65 | | NCBI | GCA_0245 86325.1 |
| | | StechGNL2 SchG015129 (chr 4, Mb 48.9) | 1374 | sens | | | 64 | NCBI | GCA_0245 86325.1 |
| Malvales (Malvaceae) | Theobroma cacao | ThecaGNOM (chr 2) LOC18610338 | 1468 | sens | 84 | 62 | | NCBI | GCA_0002 08745.2 |
| | | ThecaGNL1 (chr4) LOC18601571 | 1456 | res | 69 | 68 | | NCBI | GCA_0002 08745.2 |
| | | ThecaGNL2 (chr9) LOC18588627, | 1378 | sens | | | 69 | NCBI | GCA_0002 08745.2 |
| Malvales (Malvaceae) | Gossypium raimondii | Gosra_LOC10577959 6 (chr 12) (Gorai_12G186300) | 1464 | sens | 84 | 63 | | NCBI | GCA_0256 98545.1 |
| | | Gosra_LOC10579815 7 (chr 6) (Gorai_6G013900) | 1459 | res | 66 | 67 | | NCBI | GCA_0256 98545.1 |
| | | Gosra_LOC10578054 3 (chr 12) (Gorai_12G051700) | 1430 | sens | 63 | 64 | | NCBI | GCA_0256 98545.1 |
| | | Gosra_LOC10576918 8 (chr 9) (Gorai_9G023900) | 1379 | sens | | | 69 | NCBI | GCA_0256 98545.1 |
| Malvales (Malvaceae) | Durio zibethinus | Durzi_LOC111281800 (DurziGNOMa, sc3) | 1468 | sens | 84 | 63 | | NCBI | GCA_0023 03985.1 |
| | | Durzi_LOC111286997 DurziGNOMb sc5) | 1467 | sens | 82 | 62 | | NCBI | GCA_0023 03985.1 |
| | | Durzi_LOC111291690 (DurziGNL1a, sc7) | 1456 | res | 68 | 68 | | NCBI | GCA_0023 03985.1 |
| | | Durzi_LOC111300567 (DurziGNL1b, sc13) | (1456) | sens | 75 | 72 | | NCBI | GCA_0023 03985.1 |
| | | Durzi_LOC111315992 (DurziGNL1c, sc22) | (1440) | res | 65 | 66 | | NCBI | GCA_0023 03985.1 |

| | | | | | | | | | |
|---|---|---|---|---|---|---|---|---|---|
| | | Durzi_LOC111284518 (DurziGNL2a, sc 4) | 1379 | sens | | | 69 | NCBI | GCA_002303985.1 |
| | | Durzi_LOC111303223 (DurziGNL2b, sc 15) | 1372 | sens | | | 68 | NCBI | GCA_002303985.1 |
| Malvales (Malvaceae) | Corchorus capsularis | CCACVL1_06544 (ctg 08109) | 1448 | sens | 83 | 63 | | NCBI | GCA_001974805.1 |
| | | CCACVL1_13722 (ctg 10430) | 1447 | sens | 69 | 68 | | NCBI | GCA_001974805.1 |
| | | CCACVL1_09433 | 1359 | sens | | | 68 | NCBI | GCA_001974805.1 |
| Malvales (Malvaceae) | Corchorus olitorius | COLO4_27976 (CorolGNOM, contig 19753) | 1448 | sens | 84 | 63 | | NCBI | GCA_001974825.2 |
| | | COLO4_12228 (CorolGNL1, contig 17904) | 1447 | sens | 69 | 68 | | NCBI | GCA_001974825.2 |
| | | COLO4_34896 | 1373 | sens | | | 68 | NCBI | GCA_001974825.2 |
| Malvales (Dipterocarpaceae) | Shorea leprosula | SholeGNOM (sc00014) | 1467 | sens | 85 | 63 | | NCBI | GCA_020521535.1 |
| | | SholeGNL1a (sc00011) | 1459 | sens | 71 | 68 | | NCBI | GCA_020521535.1 |
| | | SholeGNL1b (sc00029) | 1461 | res | 69 | 68 | | NCBI | GCA_020521535.1 |
| | | SholeGNL2 (sc00035) | 1383 | sens | | | 68 | NCBI | GCA_020521535.1 |
| Brassicales (Akaniaceae) | Bretschneidera sinensis | BresiGNOMa (chr 9) | (1455) | sens | 85 | 61 | | NCBI | GCA_023935145.1 |
| | | BresiGNOMb (chr 5, Mb 32.3) | (1462) | sens | 86 | 61 | | NCBI | GCA_023935145.1 |
| | | BresiGNL1a (chr 1) | (1465) | sens | 70 | 70 | | NCBI | GCA_023935145.1 |
| | | BresiGNL1b (chr 5, Mb 92.3) | (1458) | sens | 69 | 68 | | NCBI | GCA_023935145.1 |
| | | BresiGNL2a (chr 2) | (1384) | sens | | | 69 | NCBI | GCA_023935145.1 |

| | | | | | | | | | |
|---|---|---|---|---|---|---|---|---|---|
| | | BresiGNL2b (chr 5, Mb 114.3) | (1374) | Sens? | | | 50 | NCBI | GCA_023935145.1 |
| Brassicales (Moringaceae) | Moringa oleifera | Morol09g08130 (chr 9) MorolGNOM (sc 11123+ sc 29959) | 1471 | sens | 85 | 64 | | NCBI | GCA_021397835.1 |
| | | Morol03g07330 (chr 3) MorolGNL1 (sc 6476) | 1461 | res | 70 | 69 | | NCBI | GCA_021397835.1 |
| | | Morol06g05910 (chr6) | 1387 | sens | | | 70 | NCBI | GCA_021397835.1 |
| Brassicales (Caricaceae) | Carica papaya | CarpaGNOM (chr 2) | 1468 | sens | 86 | 64 | | NCBI | GCA_021527605.1 |
| | | CarpaGNL1 (chr 1) | 1463 | res | 68 | 68 | | NCBI | GCA_021527605.1 |
| | | CarpaGNL2 (chr 7) | 1392 | sens | | | 70 | NCBI | GCA_021527605.1 |
| Brassicales (Capparaceae) | Capparis spinosa | CapspGNOMa (chr12) Cs12G009480 | 1461 | sens | 88 | 63 | | Figshare | Pep file |
| | | CapspGNOMb (chr 2) Cs02G006420 (aa459-1431) | 990 | sens | 87 | 60 | | Figshare | Pep file |
| | | CapspGNL1a (chr 8) Cs08G004760 | 1477 | res | 63 | 75 | | Figshare | Pep file |
| | | CapspGNL1b (chr 2) Cs02G004040 | 1459 | res | 64 | 76 | | Figshare | Pep file |
| | | CapspGNL2 (chr 10) Cs10G005990 | 1354 | sens | | | 77 | Figshare | Pep file |
| Brassicales (Cleomaceae) | Tarenaya hassleriana | Tarha_LOC104824072 | 1449 | sens | 89 | 63 | | NCBI | GCA_000463585.1 |
| | | Tarha_LOC104805178 | 1459 | sens | 87 | 61 | | NCBI | GCA_000463585.1 |
| | | Tarha_LOC104805485 | 1444 | sens | 64 | 78 | | NCBI | GCA_000463585.1 |
| | | Tarha_LOC104823043 | 1445 | ? (V) | 63 | 76 | | NCBI | GCA_000463585.1 |
| | | Tarha_LOC104799071 | 1376 | sens | | | 82 | NCBI | GCA_000463585.1 |

| | | | | | | | | | |
|---|---|---|---|---|---|---|---|---|---|
| Brassicales (Cleomaceae) | Cleome violacea | Clevi.0004s1596 | 1456 | sens | 90 | 62 | | Phytozome 13 | V2.1 |
| | | Clevi.0004s2072 | 1459 | res | 65 | 79 | | Phytozome 13 | V2.1 |
| | | Clevi.0068s0067 | 1376 | | | | 81 | Phytozome 13 | V2.1 |
| Brassicales (Brassicaceae) | Aethionema arabicum | AetarGNOMa (KE151134) (LG2) | (1450) | sens | 89 | 63 | | NCBI | GCA_000411095.1 |
| | | AetarGNOMb (KE151808) | (1450) | sens | 89 | 63 | | NCBI | GCA_000411095.1 |
| | | AetarGNL1 (KE152569) | (1442) | res | 68 | 84 | | NCBI | GCA_000411095.1 |
| | | AetarGNL2 (KE152478) (LG1) | (1375) | | | | 86 | NCBI | GCA_000411095.1 |
| Brassicales (Brassicaceae) | Microthlaspi erraticum | MERR_LOCUS32759 GNOM | 1453 | sens | 96 | 62 | | NCBI | GCA_902728155.2 |
| | | MERR_LOCUS42656 GNL1 | 1445 | res | 64 | 88 | | NCBI | GCA_902728155.2 |
| | | MERR_LOCUS31978 GNL2 | 1376 | sens | | | 93 | NCBI | GCA_902728155.2 |
| Brassicales (Brassicaceae) | Capsella rubella | CapruGNOM (sc1) LOC17897608 | 1454 | sens | 97 | 63 | | NCBI | GCA_000375325.1 |
| | | CapruGNL1 (sc7) LOC17879376 | 1445 | res | 61 | 89 | | NCBI | GCA_000375325.1 |
| | | CapruGNL2 (sc6) LOC17881284 | 1377 | sens | | | 94 | NCBI | GCA_000375325.1 |
| Brassicales (Brassicaceae) | Eutrema salsugineum | Thhal_v10009607m (LOC18993478) (chr1) | 1454 | sens | 97 | 62 | | NCBI | GCA_016617915.1 |
| | | Thhal_v10027619m (chr 4) | 1447 | res | 65 | 90 | | NCBI | GCA_016617915.1 |
| | | Thhal_v10012445m (sc 8) | 1376 | sens | | | 93 | NCBI | GCA_016617915.1 |
| Brassicales (Brassicaceae) | Brassica napus | LOC106346538 (chr A6) | 1448 | sens | 95 | 62 | | NCBI | GCA_020379485.1 |

| | | | | | | | | | |
|---|---|---|---|---|---|---|---|---|---|
| | | LOC106377450 (chr C5) | 1448 | sens | 95 | 62 | | NCBI | GCA_020379485.1 |
| | | LOC106446048 (chr A4) | 1433 | res | 64 | 87 | | NCBI | GCA_020379485.1 |
| | | LOC106405785 (chr C6) | 1447 | res | 64 | 87 | | NCBI | GCA_020379485.1 |
| | | LOC106382937 (sc2177) | 1447 | res | 64 | 87 | | NCBI | GCA_020379485.1 |
| | | LOC106432323 (chr C1) | 1437 | res | 64 | 87 | | NCBI | GCA_020379485.1 |
| | | LOC106352670(chr A2) | 1444 | res | 64 | 87 | | NCBI | GCA_020379485.1 |
| | | LOC106431314 (chr C9) | 1373 | sens | | | 92 | NCBI | GCA_020379485.1 |
| | | LOC106370806 (chr A10) | 1373 | sens | | | 93 | NCBI | GCA_020379485.1 |
| Brassicales (Brassicaceae) | Brassica rapa NEW (2020) | LOC103872089 (chr A6) | 1448 | sens | 95 | 62 | | NCBI | GCA_000309985.3 |
| | | LOC103849503 (chr A4) | 1433 | res | 64 | 87 | | NCBI | GCA_000309985.3 |
| | | LOC103850143 (chr A7) | 1446 | res | 63 | 87 | | NCBI | GCA_000309985.3 |
| | | LOC103845738 (chr A10) | 1373 | sens | | | 93 | NCBI | GCA_000309985.3 |
| Brassicales (Brassicaceae) | Camelina neglecta | CamneGNOM (chr 1) | 1452 | sens | 98 | 62 | | NCBI | GCA_023864065.1 |
| | | CamneGNL1 (chr 5) | 1443 | res | 62 | 90 | | NCBI | GCA_023864065.1 |
| | | CamneGNL2 (chr 6) | 1383 | sens | | | 94 | NCBI | GCA_023864065.1 |
| Brassicales (Brassicaceae) | Boechera stricta | Bostr_13671s0057 | 1454 | sens | 98 | 63 | | NCBI | GCA_018361395.1 |
| | | Bostr_10058s0047 | 1445 | res | 63 | 91 | | NCBI | GCA_018361395.1 |
| | | Bostr_2570s0012 | 1151 | sens | 61 | 81 | | NCBI | GCA_018361395.1 |

| | | | | | | | | | |
|---|---|---|---|---|---|---|---|---|---|
| | | Bostr.26527s0239 | 1383 | sens | | | 95 | NCBI | GCA_018361395.1 |
| Brassicales (Brassicaceae) | Arabidopsis lyrata | AL1G25820.t1 | 1454 | sens | 99 | 63 | | NCBI | GCA_000004255.1 |
| | | AL7G50810.t1 | 1445 | res | 62 | 94 | | NCBI | GCA_000004255.1 |
| | | AL6G31120.t1 (LOC9310023) | 1376 | sens | | | 98 | NCBI | GCA_000004255.1 |
| Brassicales (Brassicaceae) | Arabidopsis thaliana | GNOM / At1g13980 | 1451 | sens | 100 | 62 | | | |
| | | GNL1 / AT5G39500 | 1443 | res | 62 | 100 | | | |
| | | GNL2 / AT5G19610 | 1375 | sens | | | 100 | | |

#### Suppl. Table S2. Rescue of *gnom*$^{sgt}$ by different *GNOM*-related paralogs

##### Genotype of mother plant: *BranaGNOM-YFP/- gnom/GNOM*

| Line | *gnom* ($PPT^R$) | *gnom* ($PPT^s$) | WT-like seedlings ($PPT^R$) | WT-like seedlings ($PPT^s$) | n |
|---|---|---|---|---|---|
| #1 | 0 | 22 (5.9%) | 277 (74.5%) | 73 (19.6%) | 372 |
| #2 | 0 | 24 (5.5%) | 333 (76.0%) | 81 (18.5%) | 438 |
| Expected (rescue) | 0% | 6.25% | 75% | 18.75% | |
| Expected (No rescue) | 18.75% | 6.25% | 56.25% | 18.75% | |

##### Genotype of mother plant: *BranaGNL1-YFP/- gnom/GNOM*

| Line | *gnom* ($PPT^R$) | *gnom* ($PPT^s$) | WT-like seedlings ($PPT^R$) | WT-like seedlings ($PPT^s$) | n |
|---|---|---|---|---|---|
| #1 | 110 (18.6%) | 35 (5.9%) | 329 (55.5%) | 119 (20.0%) | 593 |
| #2 | 87 (17.1%) | 27 (5.3%) | 290 (57.1%) | 104 (20.5%) | 508 |
| Expected (rescue) | 0% | 6.25% | 75% | 18.75% | |
| Expected (No rescue) | 18.75% | 6.25% | 56.25% | 18.75% | |

##### Genotype of mother plant: *PoptrGNOMa-YFP/- gnom/GNOM*

| | *gnom* ($PPT^R$) | *gnom* ($PPT^s$) | WT-like seedlings ($PPT^R$) | WT-like seedlings ($PPT^s$) | n |
|---|---|---|---|---|---|
| #1 | 0 | 11 (4.8%) | 180 (79.3%) | 36 (15.9%) | 227 |
| #2 | 0 | 14 (6.4%) | 173 (78.6%) | 33 (15.0%) | 220 |
| Expected (rescue) | 0% | 6.25% | 75% | 18.75% | |
| Expected (No rescue) | 18.75% | 6.25% | 56.25% | 18.75% | |

##### Genotype of mother plant: *PoptrGNOMb-YFP/PtGNOMb gnom/GNOM*

| Line | *gnom* ($PPT^R$) | WT-like seedlings ($PPT^R$) | Partially rescued seedlings ($PPT^R$) | n |
|---|---|---|---|---|
| #1 | 0 | 297 (76.5%) | 91 (23.5%) | 388 |
| #2 | 16(3.4%) | 358 (76.2%) | 96 (20.4%) | 470 |
| Expected (rescue) | 0% | 100% | | |
| Expected (No rescue) | 25% | 75% | | |

**Genotype of mother plant: *OrysaGNOMa-YFP/- gnom/GNOM***

| Line | *gnom* (PPT$^R$) | *gnom* (PPT$^s$) | WT-like seedlings (plus partial rescue) (PPT$^R$) | WT-like seedlings (PPT$^s$) | n |
|---|---|---|---|---|---|
| #1 | 0 | 12(5.2%) | 181 (78.7%) | 37 (16.1%) | 230 |
| #2 | 0 | 14(5.5%) | 193 (76.3%) | 46 (18.2%) | 253 |
| Expected (rescue) | 0% | 6.25% | 75% | 18.75% | |
| Expected (No rescue) | 18.75% | 6.25% | 56.25% | 18.75% | |

**Genotype of mother plant: *OrysaGNOMb-YFP/OrysaGNOMb-YFP gnom/GNOM***

| Line | *gnom* (PPT$^R$) | WT-like seedlings (PPT$^R$) | Partially rescued seedlings (PPT$^R$) | n |
|---|---|---|---|---|
| #1 (plant 1) | 14 (4.9%) | 237 (83.7%) | 32 (11.3%) | 283 |
| #1/(plant 2) | 38 (7.1%) | 428 (79.7%) | 71 (13.2%) | 537 |
| Expected (rescue) | 0% | 100% | | |
| Expected (No rescue) | 25% | 75% | | |

**Genotype of mother plant: *OrysaGNOMb/- gnom/GNOM***

| Line | *gnom* (PPT$^R$) | *gnom* (PPT$^s$) | WT-like seedlings (PPT$^R$) | WT-like seedlings (PPT$^s$) | n |
|---|---|---|---|---|---|
| #2 | 119 (16.3%) | 38 (5.2%) | 438 (60%) | 135 (18.5%) | 730 |
| Expected (rescue) | 0% | 6.25% | 75% | 18.75% | |
| Expected (No rescue) | 18.75% | 6.25% | 56.25% | 18.75% | |

#### Note

OsGNOMb had low activity such that (partial) rescue of *gnom-sgt* was only detected when 2 copies of the transgene were present

***Marchantia polymorpha***

*gnom$^{sgt}$* homozygosity of 20 wild-type looking seedlings was confirmed by PCR, from two independent lines (T2) growing on PPT selection plate.

| | Primer | Nucleotide sequence |
|---|---|---|
| | GNovertag_S | GAAAGTGAAAGTAAGAGGC |
| | GNovertag_AS | CGTAGAGAGGTGTTACATAAG |
| | YFP-AS | CATGTGATCGCGCTTCTC |
| | mCherry_F | CCTTCCCGGGATGGTGAGCAAGGGCGAGGAGGA |
| | mCherry_R | CCTTTCTAGATTACTTGTACAGCTCGTCCATGC |
| | GNOM-Prom-KpnI-S | TTTTTGGTACCGGTGTGTATGATAATGAATATTG |
| | GNOM-Prom-ApaI-AS | TTTTTGGGCCCTTAATCTGCTCAAATCTTCAGCCAG |
| | GNOM-YFP-SacI-AvrII-S | TTTTTGAGCTCCCTAGGATGGTGAGCAAGGGCGAGG |
| | GN-UTR-SacI-AS | TTTTTGAGCTCAATCGAAATCCGTCTCCCGGAGC |
| | GNOM-Promotor-Seq | AGGTTCCATTAATTGGGCCT |
| | BranaGNOM-Seq1 | GCCAACAGATCTGAAATC |
| | BranaGNOM-Seq2 | CGCTGGATGGCTTAATTG |
| | BranaGNOM-Seq3 | GCATAAATCCAAGAAAAC |
| | BranaGNOM-Seq4 | CTGCCTGGAATTGCTAAT |
| | BranaGNOM-Seq1a | GGTGTATCTTCTCCCATCT |
| | BranaGNL1_ApaI-S | TTTTGGGCCCATGGGGTATCAGAATCACCATCCAT |
| | BranaGNL1_NotI-AS | TTTTGCGGCCGCGTTTTCCGGTACCACCGGAGTATCC |
| | BranaGNL1-Seq1 | ATCGTATTATGATTTATCTG |
| | BranaGNL1-Seq2 | GTGATATAACATGCAGTAATG |
| | BranaGNL1-Seq3 | GATTCAGATGATCCCAGATC |
| | BranaGNL1-Seq4 | AGATAAAGCTAGTGCTGTC |
| | BranaGNL1-Seq5 | GCTAGACGAACTGCTCGCTG |
| | BranaGNL1-Ins-R | ATGCTATAGGGTTAGATCTG |
| | PoptrGNOMa-Seq1 | GCGCTGAACAGACATTGG |
| | PoptrtGNOMa-Seq2 | GGATGGCTGAAAGGATAG |
| | PoptrGNOMa-Seq3 | TGATCTAATGCACAAGTC |
| | PoptrGNOMa-Seq4 | CTGCCTTGAATTGCTCAT |
| | PoptrGNOMa-Seq5 | GGAGTCTGCAAAATTATCACA |
| | PoptrGNOMb-CDS-ApaI-S | TTTTGGGCCCATGGGGAGAATAAAGCTACAGTCTG |
| | PoptrGNOMb-CDS-NotI-AS | TTTTGCGGCCGCACCTCCAGTGCCAGCAGCACATTCAG |
| | PoptrGNOMb-Seq1 | GTAACATCTCTGCTCTCAATG |
| | PoptrGNOMb-Seq2 | GATTTGCACCTCATGACATTG |
| | PoptrGNOMb-Seq3 | CGGAGCTGATCACTTCAATC |
| | PoptrGNOMb-Seq4 | ATTGGATGCTGTAGTTGTG |
| | PoptrGNOMb-Seq5 | TTGAGGTCACTGCAACTTG |
| | PoptrGR2-Seq6 | GAAGGGACACTTGTCATTGC |
| | OrysaGNOMb-ApaI-F | TTTTGGGCCCATGGGCGGCCTGAGGGCAGCGTCGC |
| | OrysaGNOMb-NotI-R | TTTTGCGGCCGCAACATTCACGCCTTCAGATTGTGCT |
| | OrysaGNOMb-Seq1 | CCTTCCTGTAAAGATTTCG |
| | OrysaGNOMb-Seq2 | TGTGTTATCATAAGACTTGG |
| | OrysaGNOMb-Seq3 | AGTAATCATGCTCAACACAG |
| | OrysaGNOMb-Seq4 | ATTGTATTTAGATGCTGAAG |
| | OrysaGNOMb-Seq5 | GTGAATTGTCTTACACGCTG |
| | $\Delta DCB^{OsGNOMa}$-ApaI-F | TTTTGGGCCCATGCTACTTGTGCAGATGGGTGGC |
| | $\Delta DCB^{OsGNOMa}$-NotI-R | TTTTGCGGCCGCGTCCCGGGACTCAAC |
| | OrysaGNOMa-Seq1 | TCTTGCTCTGGAAGGTCTG |
| | OrysaGNOMa-Seq1n | GAATTCTGCAGATGGCAGTG |
| | OrysaGNOMa-Seq2 | GTCGAAGTCAACATCTCTG |
| | OrysaGNOMa-Seq3 | TGTCTGGAACTGCTAATCGC |
| | OrysaGNOMa-Seq4 | ACTTACTAGAGATCAGCCAG |
| | OrysaGNOMaseq_R1 | CAACTCCAATTTGCTCGACA |
| | OrysaGNOMaseq_R2 | CGAGCAGCGCCTGCAGCATC |
